# Genome-wide association studies of global *Mycobacterium tuberculosis* resistance to thirteen antimicrobials in 10,228 genomes

**DOI:** 10.1101/2021.09.14.460272

**Authors:** The CRyPTIC Consortium, Sarah G Earle, Daniel J Wilson

**Affiliations:** University of Oxford

## Abstract

The emergence of drug resistant tuberculosis is a major global public health concern that threatens the ability to control the disease. Whole genome sequencing as a tool to rapidly diagnose resistant infections can transform patient treatment and clinical practice. While resistance mechanisms are well understood for some drugs, there are likely many mechanisms yet to be uncovered, particularly for new and repurposed drugs. We sequenced 10,228 *Mycobacterium tuberculosis* (MTB) isolates worldwide and determined the minimum inhibitory concentration (MIC) on a grid of twofold concentration dilutions for 13 antimicrobials using quantitative microtiter plate assays. We performed oligopeptide- and oligonucleotide-based genome-wide association studies using linear mixed models to discover resistance-conferring mechanisms not currently catalogued. Use of MIC over binary resistance phenotypes increased heritability for the new and repurposed drugs by 26-37%, increasing our ability to detect novel associations. For all drugs, we discovered uncatalogued variants associated with MIC, including in the *Rv1218c* promoter binding site of the transcriptional repressor *Rv1219c* (isoniazid), upstream of the *vapBC20* operon that cleaves 23S rRNA (linezolid) and in the region encoding an α-helix lining the active site of Cyp142 (clofazimine, all *p*<10^-7.7^). We observed that artefactual signals of cross resistance could be unravelled based on the relative effect size on MIC. Our study demonstrates the ability of very large-scale studies to substantially improve our knowledge of genetic variants associated with antimicrobial resistance in *M. tuberculosis*.

## Introduction

Tuberculosis (TB) continues to represent a major threat to global public health, with the World Health Organization (WHO) estimating 10 million cases and 1.4 million deaths in 2019 alone [1]. Multidrug resistance (MDR) poses a major challenge to tackling TB; it is estimated that there were 465,000 cases of rifampicin resistant TB in 2019, of which 78% were resistant to the first-line drugs rifampicin and isoniazid – called MDR-TB [1]. While treatment is 85% successful overall, that drops to 57% for rifampicin-resistant and MDR-TB [1]; underdiagnosis and treatment failures then amplify the problem by encouraging onward transmission of MDR-TB [2]. New treatment regimens for MDR-TB are therefore an important focus, introducing new and repurposed drugs such as bedaquiline, clofazimine, delamanid and linezolid [3, 4]; however resistance is already emerging [5,6,7].

Understanding mechanisms of resistance in TB is important for developing rapid susceptibility tests that improve individual patient treatment, recommending drug regimens that reduce the development of MDR and developing new and improved drugs that expand treatment options [8, 9]. Genomics can accelerate drug susceptibility testing, replacing slower culture-based methods by predicting resistance from the sequenced genome rather than directly phenotyping the bacteria [10]. Genome sequencing-based susceptibility testing for first-line drugs has achieved sensitivities of 91.3-97.5% and specificities of 93.6-99.0% [11], surpassing the thresholds for clinical accreditation, motivating its adoption by multiple public health authorities [12]. In low-resource settings, molecular tests such as Cepheid GeneXpertⓇ and other line probe assays offer rapid and more economical susceptibility testing by genotyping a panel of known resistance-conferring genetic variants [13], with performance close to that achieved by whole genome sequencing [14, 15]. However, the limited number of resistance-conferring mutations that can be included in such tests can lead to missed MDR diagnoses and incorrect treatment [11, 16]. Both approaches rely on the development and maintenance of resistance catalogues of genetic variants [17, 11].

In the discovery of resistance-conferring variants, traditional molecular approaches have been replaced by high-throughput, large-scale whole genome sequencing studies of hundreds to thousands of resistant and susceptible clinical isolates [18,19,20,21,22,23]. Despite the strong performance of genome-based resistance prediction for first-line drugs, knowledge gaps remain, especially for second-line drugs [24,25,17]. There are numerous challenges in the pursuit of previously uncatalogued resistance mechanisms. Very large sample sizes are needed to identify rarer resistance mechanisms with confidence. The lack of recombination in *Mycobacterium tuberculosis* makes it difficult to pinpoint resistance variants unless they arise on multiple genetic backgrounds, reiterating the need for large sample sizes. Sophisticated analyses are required that attempt to disentangle genetic causation from correlation [26]. A reliance on a binary resistance/sensitivity classification paradigm has hindered reproducibility for some drugs, by failing to mirror the continuous nature of resistance [27,28,29].

The aim of *Comprehensive Resistance Prediction for Tuberculosis: an International Consortium* (*CRyPTIC*) was to address these challenges by assembling a global collection of over 10,000 *M. tuberculosis* isolates from 27 countries followed by whole-genome sequencing and semi-quantitative determination of minimum inhibitory concentration (MIC) to 13 first- and second-line drugs using a bespoke 96-well broth micodilution plate assay. The development of novel, inexpensive, high-throughput drug susceptibility testing assays allowed us to conduct the project at scale, while investigating MIC on a grid of twofold concentration dilutions [30, 31]. Here we report the identification of previously uncatalogued resistance-conferring variants through 13 genome-wide association studies (GWAS) investigating MIC values in 10,228 *M. tuberculosis* isolates. We employed a linear mixed model (LMM) to identify putative causal variants while controlling for confounding and genome-wide linkage disequilibrium (LD). We developed a novel approach to testing associations at both 10,510,261 oligopeptides (11-mers) and 5,530,210 oligonucleotides (31-mers) to detect relevant genetic variation in both coding and non-coding sequences, and to avoid a reference-based mapping approach that can inadvertently miss significant variation. We report previously uncatalogued variants associated with MIC for all 13 drugs, focusing on variants in the 20 most significant genes per drug. We highlight notable discoveries for each drug, and demonstrate the ability of large-scale studies to improve our knowledge of genetic variants associated with antimicrobial resistance in *M. tuberculosis*.

## Results

CRyPTIC collected isolates from 27 countries worldwide, oversampling for drug resistance [31]. 10,228 genomes were included in total across the GWAS analyses; 533 were lineage 1, 3581 lineage 2, 805 lineage 3, and 5309 lineage 4. Due to rigorous quality control, we dropped samples for each drug as detailed in the methods, resulting in a range of 6,388-9,418 genomes used in each GWAS (**Figure 1**). Minimum inhibitory concentrations (MICs) were determined on a grid of twofold concentration dilutions for 13 antimicrobials using quantitative microtiter plate assays: first-line drugs ethambutol, isoniazid and rifampicin; second-line drugs amikacin, ethionamide, kanamycin, levofloxacin, moxifloxacin and rifabutin and the new and repurposed drugs bedaquiline, clofazimine, delamanid and linezolid. The phenotype distributions differed between the drugs, with low numbers of sampled resistant isolates for the new and repurposed drugs which have not yet been widely used in tuberculosis treatment (**Figure 1**, **Supplementary Figure 1**). Assuming log_2_ MIC epidemiological cut-offs (ECOFFs) of 0.25 (bedaquiline, clofazimine), 0.12 (delamanid) and 1 mg/L (linezolid) [31], the GWAS featured 66 isolates resistant to bedaquiline, 97 resistant to clofazimine, 77 resistant to delamanid and 67 resistant to linezolid. We performed oligopeptide- and oligonucleotide-based GWAS analyses, controlling for population structure using linear mixed models (LMMs). We focused initially on oligopeptides, interpreting oligonucleotides only where necessary for clarifying results.

**Figure 1.**
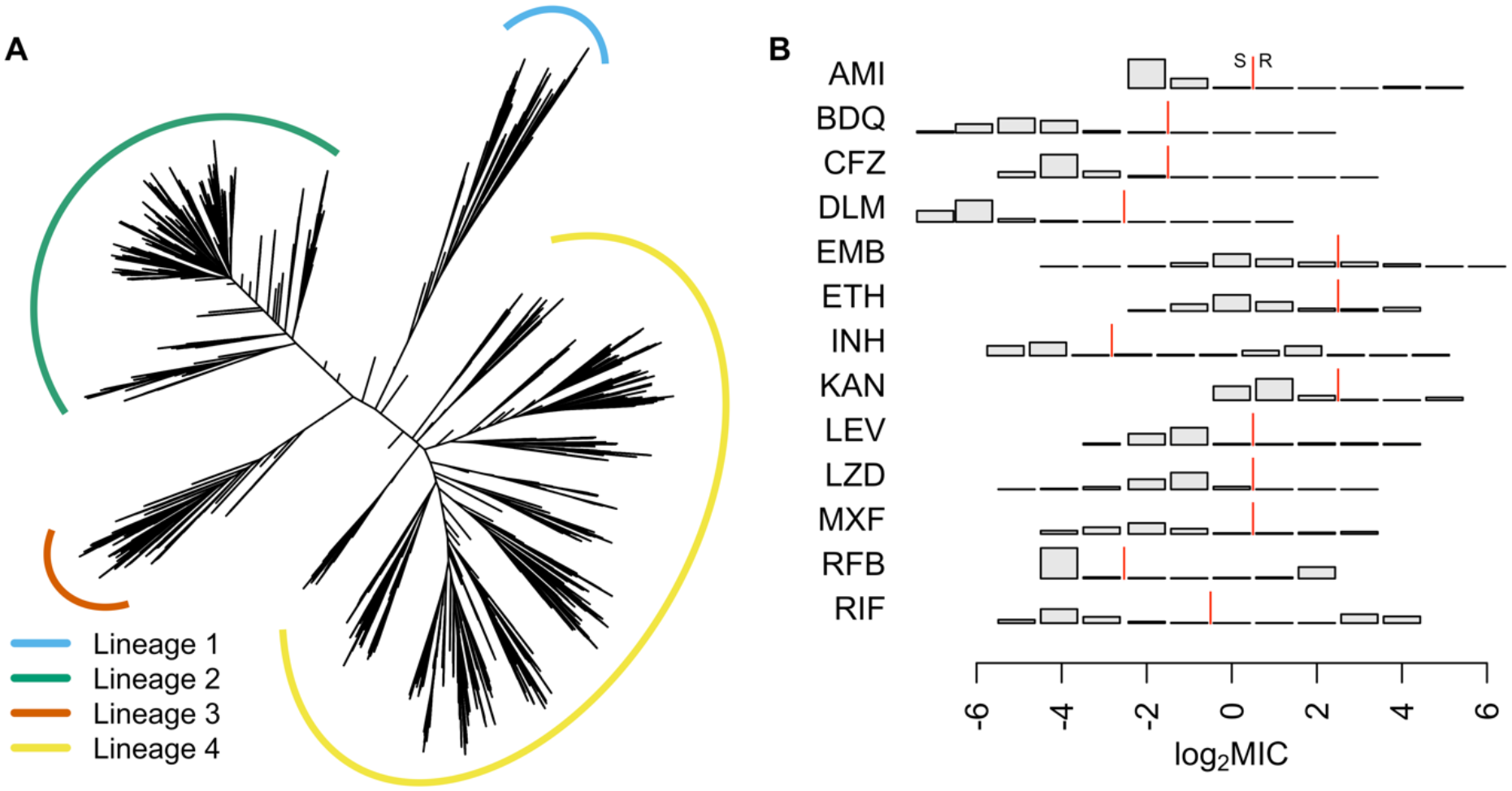
**A** Phylogeny of 10,228 isolates sampled globally by CRyPTIC used in the GWAS analyses. Lineages are coloured yellow (lineage 1), green (2), blue (3) and orange (4). Branch lengths have been square root transformed to visualise the detail at the tips. **B** Distributions of the log_2_ MIC measurements for all 13 drugs in the GWAS analyses, amikacin (AM), bedaquiline (BDQ), clofazimine (CFZ), delamanid (DLM), ethambutol (EMB), ethionamide (ETH), isoniazid (INH), kanamycin (KAN), levofloxacin (LEV), linezolid (LZD), moxifloxacin (MXF), rifabutin (RFB) and rifampicin (RIF). The red line indicates the ECOFF breakpoint for binary resistance versus sensitivity calls [31].

Estimates of sample heritability (variance in the phenotype explained by additive genetic effects) were higher for MIC compared to binary resistant vs. sensitive phenotypes for the new and repurposed drugs bedaquiline, clofazimine, delamanid and linezolid by at least 26%. Across drugs, binary heritability ranged from 0-94.7% and MIC heritability from 36.0-95.6%, focusing on oligopeptides (**Figure 2, Supplementary Figure 2 and Supplementary Table 1**). For delamanid, binary heritability was not significantly different from zero (2.99×10^-6^; 95% confidence interval (CI) 0.0-0.5%), while MIC heritability was 36.0% (95% CI 28.9-43.1%). Heritability estimates were more similar between binary and MIC phenotypes for the remaining drugs, differing by −3.6 to +5.2%.

**Figure 2.**
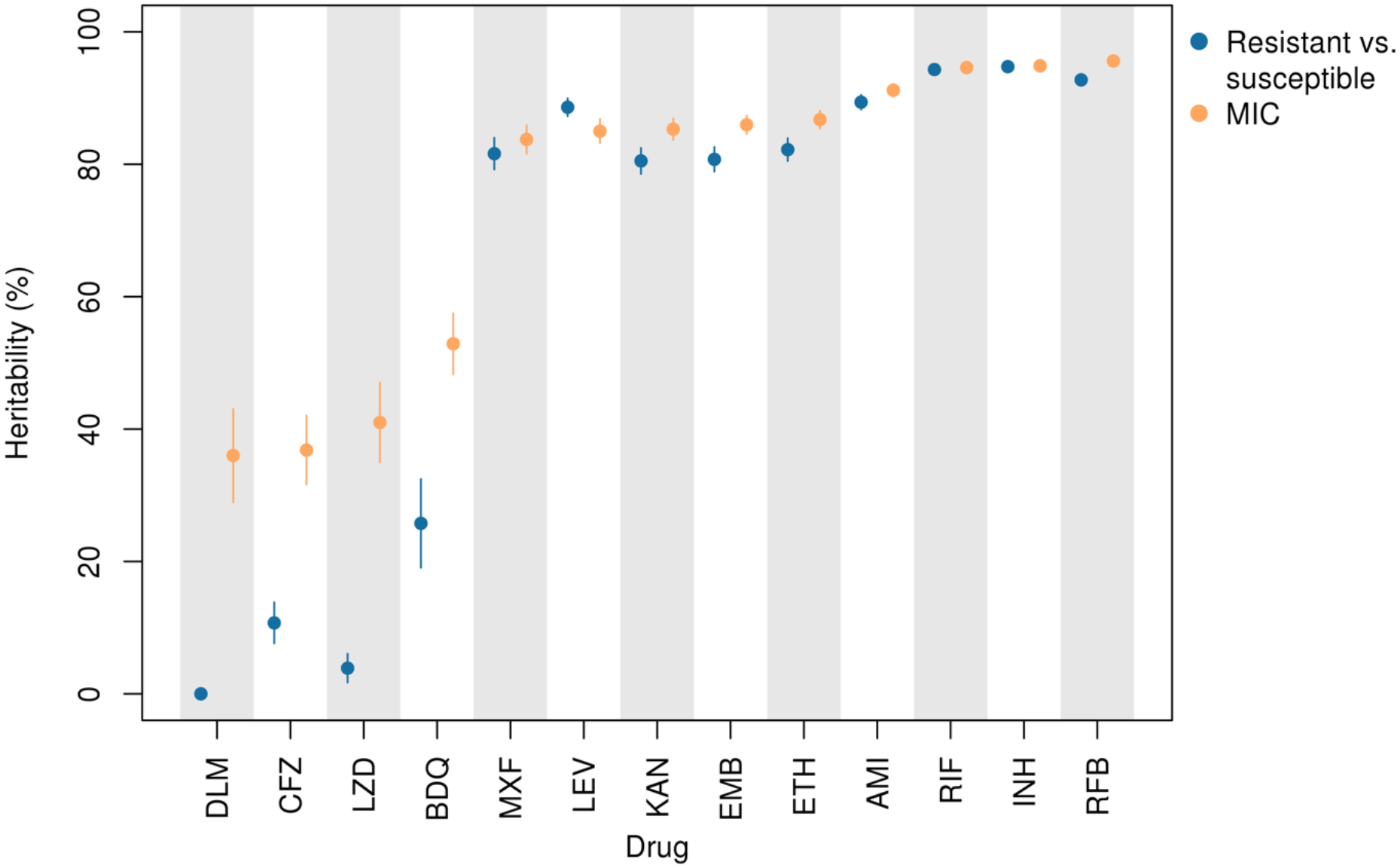
MIC heritability (orange) versus binary (resistant/sensitive) heritability (blue) assuming additive genetic variation in oligopeptide presence/absence across 13 drugs, DLM (delamanid), clofazimine (CFZ), linezolid (LZD), bedaquiline (BDQ), moxifloxacin (MXF), levofloxacin (LEV), kanamycin (KAN), ethambutol (EMB), ethionamide (ETH), amikacin (AM), rifampicin (RIF), isoniazid (INH), rifabutin (RFB). Lines depict 95% confidence intervals. MIC heritability was at least 26% higher than binary heritability for the new and repurposed drugs bedaquiline, clofazimine, delamanid and linezolid.

GWAS identified oligopeptide variants associated with changes in MIC for all 13 drugs after controlling for population structure (**Figure 3**, **Table 1**, **Supplementary Figure 3-4**). In total, across the drugs, we tested for associations at 10,510,261 variably present oligopeptides and 5,530,210 oligonucleotides; these captured substitutions, insertions and deletions. The drugs differed in the number of genes or intergenic regions that were significant, the drugs with fewest significant genes being isoniazid (12), levofloxacin (13) and moxifloxacin (6). We defined the significance of a gene or intergenic region by the most significant oligopeptide within it, and assessed all significant variants above a 0.1% minor allele frequency (MAF) threshold for the top 20 significant genes. The top 20 genes for each drug are detailed in **Table 1**. Some variants were identified in novel genes, some were novel variants in known genes, and some were known variants. We highlight examples of these (in reverse order) in the following sections. Highlighted examples have been chosen to exclude genes or variants in LD with other regions where possible; some are in LD with other less significant variants.

**Figure 3.**
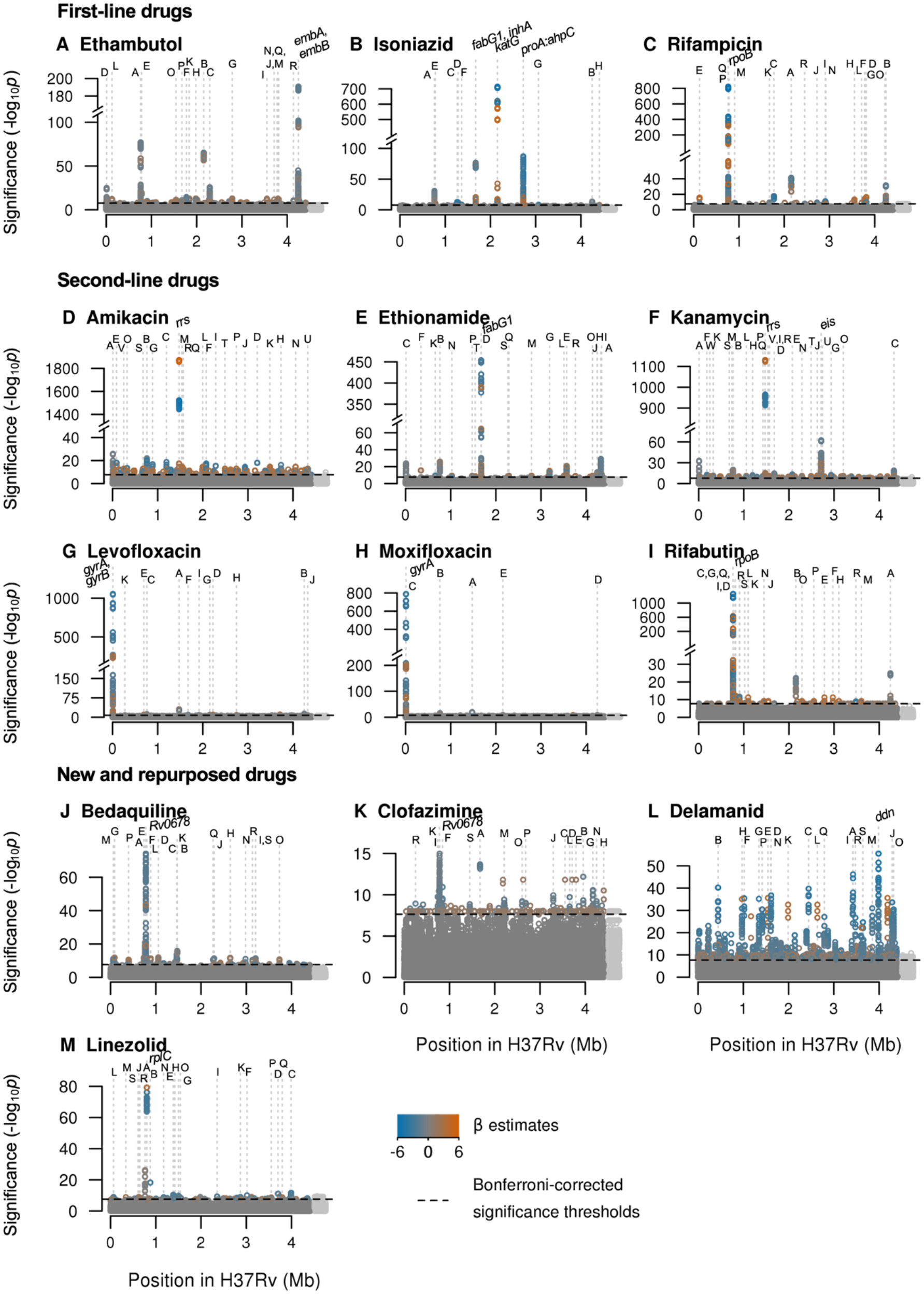
Manhattan plots of regions containing oligopeptide variants associated with MIC across 13 drugs. Significant oligopeptides are coloured by the direction (orange=increase, blue=decrease) and magnitude of their effect size on MIC, estimated by LMM [32]. Bonferroni-corrected significance thresholds are shown by the black dashed lines. The top 20 genes ranked by their most significant oligopeptides are annotated alphabetically. Gene names separated by colons indicate intergenic regions. Gene names for those annotated with letters can be found in Table 1. Oligopeptides were aligned to the H37Rv reference; unaligned oligopeptides are plotted to the right in light grey.

**Figure 4.**
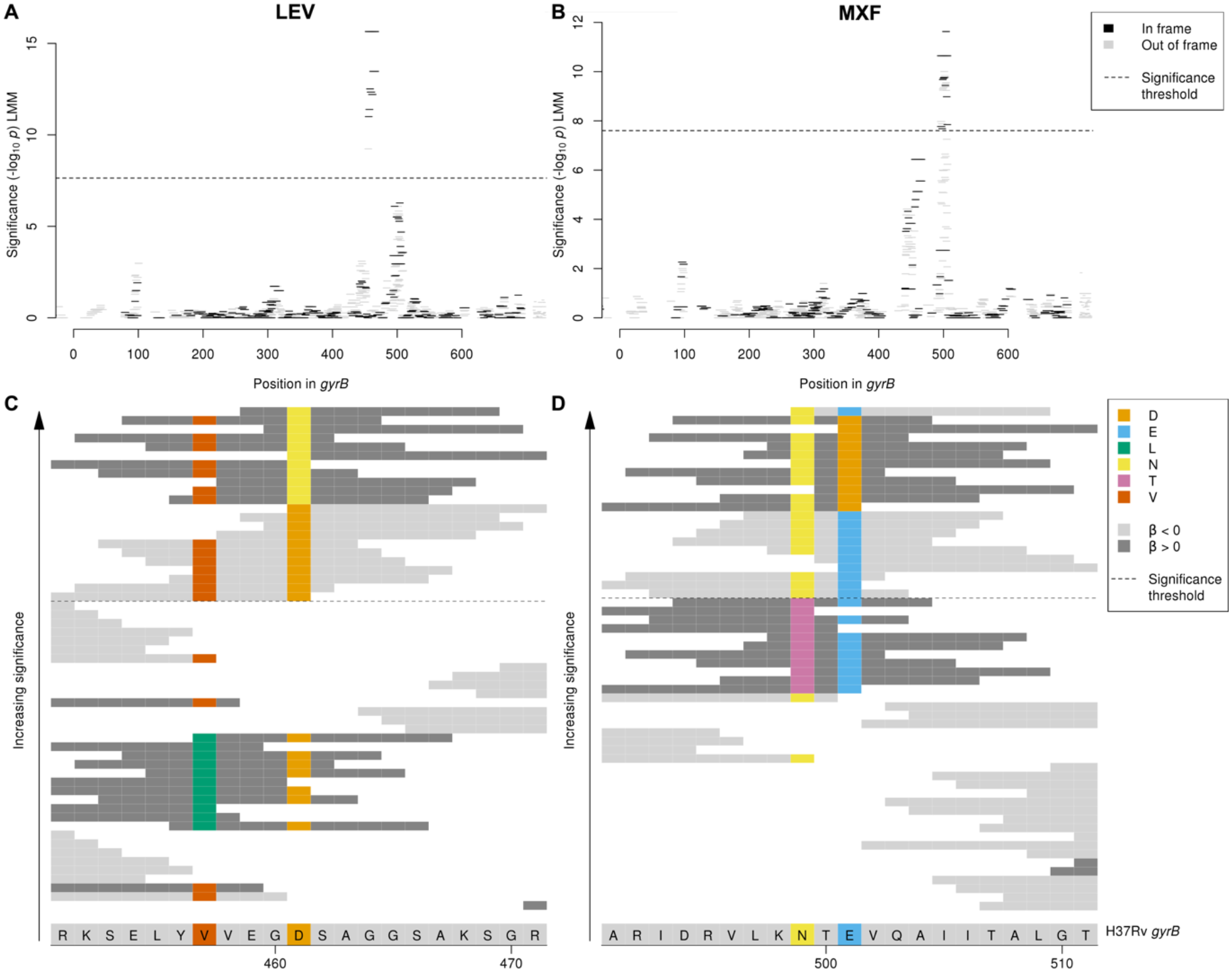
Interpreting significant oligopeptide variants for levofloxacin and moxifloxacin MIC in *gyrB*. Oligopeptide Manhattan plots are shown for **A** levofloxacin **B** moxifloxacin. Oligopeptides are coloured by the reading frame that they align to, black for in frame and grey for out of frame in *gyrB*. Oligopeptides aligned to the region by nucmer but not realigned by BLAST are shown in grey on the right hand side of the plots. The black dashed lines indicate the Bonferroni-corrected significance thresholds – all oligopeptides above the line are genome-wide significant. Alignment is shown of oligopeptides significantly associated with **C** levofloxacin and **D** moxifloxacin. The H37Rv reference codons are shown at the bottom of the figure, grey for an invariant site, coloured at variant site positions. The background colour of the oligopeptides represents the direction of the β estimate, light grey when β < 0 (associated with lower MIC), dark grey when β> 0 (associated with higher MIC). Oligopeptides are coloured by their amino acid residue at variant positions only.

**Table 1.**
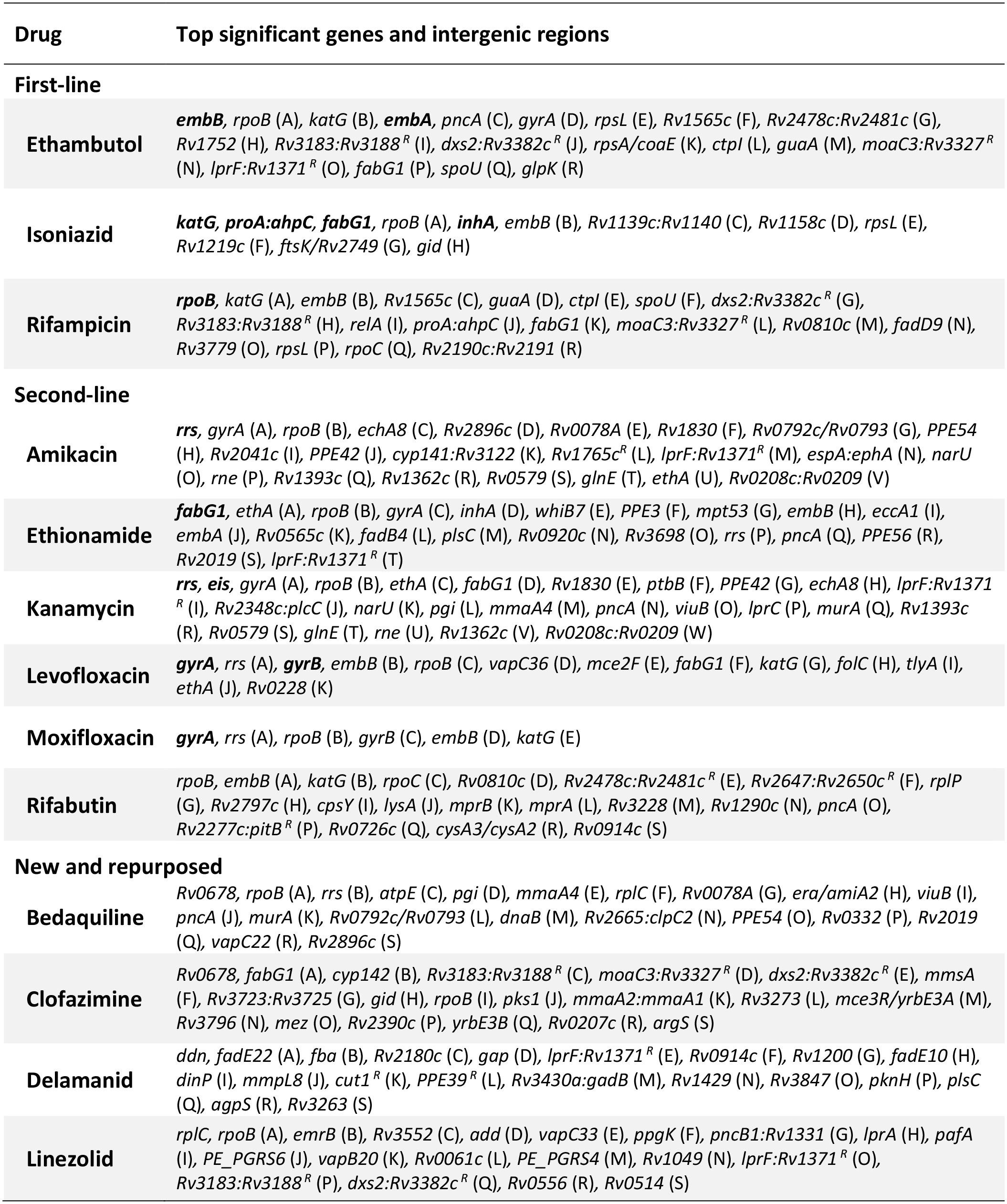
The top genes or intergenic regions ranked by their most significant oligopeptides per drug, up to a maximum of 20 (more only when the 20^th^ was tied). Genes are highlighted in bold if they were catalogued for that drug by [17, 11]. Gene names separated by colons indicate intergenic regions. Genes or intergenic regions capturing repeat regions are highlighted with the superscript *^R^*. Alphabetic characters following gene names are used to cross-reference with the corresponding Manhattan plots in Figure 3.

We assessed whether the top genes for each drug were in either of two previously described resistance catalogues [17, 11]; we describe variants not in these catalogues as uncatalogued (**Table 1**). The interpretation of oligopeptides and oligonucleotides required manual curation to determine the underlying variants they tagged, and the most significant oligopeptide or oligonucleotide for each allele captured by the significant signals are described in **Supplementary table 2** and the **Supplementary text**. For 8/13 drugs with previously catalogued resistance determinants, the most significant GWAS signal in CRyPTIC was a previously catalogued variant, consistent with previous GWAS [18,19,20,21,22,23]. The most significant catalogued variants for each drug were (lowercase for nucleotides, uppercase for amino acids): *rrs* a1401g (amikacin, kanamycin), *embB* M306V (ethambutol), *fabG1* c−15t (ethionamide), *katG* S315T (isoniazid), *gyrA* D94G (levofloxacin, moxifloxacin), and *rpoB* S450L (rifampicin) [17, 11]. For the remaining drugs with no previously catalogued resistance determinants, the genes identified by the top signals were: *Rv0678* (bedaquiline, clofazimine), *ddn* (delamanid), *fabG1* (ethionamide), *katG* (isoniazid), *rplC* (linezolid) and *rpoB* (rifabutin). The top variants identified for each drug were all significant at *p*<1.04×10^-15^.

For many drugs, the direction of effect of the most significant oligopeptide variants was to decrease MIC (**Supplementary Figure 5**), implying that low-MIC oligopeptides and oligonucleotides are more likely to be genetically identical across strains than high-MIC haplotypes. This would be consistent with the independent evolution of increased MIC from a shared, low-MIC TB ancestor. Uncatalogued variants significantly associated with MIC are important because they could improve resistance prediction and shed light on underlying resistance mechanisms; they may be novel or previously implicated in resistance but not to a standard of evidence sufficient to be catalogued. We discuss the choice of catalogues in the Discussion [17, 11].

We next looked at uncatalogued variants in known resistance-conferring genes. We identified uncatalogued variants in *gyrB* associated with levofloxacin and moxifloxacin MIC (minimum *p*-value levofloxacin: *p*<10^-15.6^, moxifloxacin: *p*<10^-11.6^. The primary mechanisms of resistance to the fluoroquinolones levofloxacin and moxifloxacin are mutations in *gyrA* or *gyrB*, the subunits of DNA gyrase. The *gyrB* Manhattan plots for levofloxacin and moxifloxacin both contained two adjacent peaks within the gene, but for each drug just one of the two peaks was significant, and these differed between the drugs (**Figure 4**). Interpretation of oligopeptides and oligonucleotides requires an understanding of the variants that they capture, which we visualised by aligning them to H37Rv and interpreting the variable sites (e.g. **Figure 4C-D**). For levofloxacin the peak centred around amino acid 461. Significant oligopeptides captured amino acids 461 and 457, which are both uncatalogued [17, 11] with 457 falling just outside of the *gyrB* quinolone resistance-determining region (QRDR-B) [33]. Oligopeptides capturing 461N were associated with increased MIC (e.g. **N**SAGGSAKSGR, -log_10_*p* = 15.65, effect size β = 2.46, present in 15/7300 genomes). Oligopeptides capturing the reference alleles at codons 461 and 457 were significantly associated with lower MIC (e.g. 461D: **D**SAGGSAKSGR, -log_10_*p* = 13.47, β = −2.14, present in 7278/7300 genomes; 457V/461D: SELY**V**VEG**D**SA, -log_10_*p* = 12.51, β = −1.96, present in 7272/7300 genomes). For moxifloxacin, the peak centred around amino acid 501. Significant oligopeptides captured amino acids 499 and 501. Oligopeptides capturing 501D were associated with increased MIC (e.g. NT**D**VQAIITAL, -log_10_*p* = 10.64, β = 1.86, present in 23/6388 genomes). Oligopeptides capturing the reference allele at codons 499 and 501 were associated with lower MIC (e.g. **N**T**E**VQAIITAL, -log_10_*p* = 11.63, β = −1.33, present in 6332/6388 genomes). Amino acids 461 and 501 are at the interface between *gyrB* and the bound fluroquinolone [34]. *gyrB* is included in the reference catalogues for predicting levofloxacin but not moxifloxacin resistance, therefore our results support inclusion in future moxifloxacin catalogues [17, 11].

Next we looked at specific examples of significant associations identified by GWAS in genes not catalogued by [17, 11] for each of the drugs. A well-recognized challenge in GWAS for antimicrobial resistance is the presence of artefactual cross resistance. To mitigate this risk, we preferentially highlight variants significantly associated with a single drug. However, many catalogued resistance variants demonstrated artefactual cross resistance. For example, variants in the rifampicin resistance determining region were in the top 20 significant associations for all drugs except for delamanid (**Table 1**). Interestingly, we observed that the magnitude of effect sizes was often larger on MIC of the drug to which catalogued variants truly confer resistance (**Supplementary Figure 6**). For example, the effect sizes for significant oligopeptides in *rpoB* were greater for rifampicin and rifabutin than for all other drugs. This suggests that the β estimates could help to prioritise drugs for follow up when genes are significantly associated with multiple drugs.

### First-line drugs

#### Ethambutol and rifampicin

Oligonucleotides downstream of *spoU* (*Rv3366*) were significantly associated with ethambutol and rifampicin MIC (minimum *p*-value *p*<10^-10.0^, **Supplementary Figure 7**). SpoU is a tRNA/rRNA methylase, shown to have DNA methylation activity [35]. As the association was outside of the coding region, we interpreted oligonucleotides for this association. Oligonucleotides associated with increased MIC captured the relatively common adenine 20 nucleotides downstream of the stop codon (e.g. C**A**AACCAGCCGGTATGCGCACAACGAAGCTC, RIF: -log_10_*p* = 12.82, β = 3.19, present in 159/8394 genomes; EMB: -log_10_*p* = 10.86, β = 1.36, present in 163/7081 genomes). This mutation has been identified in previous association studies as associated with rifampicin and ethambutol resistance [36, 37] but has not been catalogued. The new evidence provided by CRyPTIC supports re-evaluation of this putative resistance-conferring variant. The simultaneous association of *spoU* with rifampicin and ethambutol may be an example of artefactual cross resistance. The effect sizes on MIC for rifampicin (β = 3.19) were larger than for ethambutol (β= 1.36), suggesting prioritisation of the rifampicin association over the ethambutol association reported here.

#### Isoniazid

Oligopeptides in *Rv1219c* were significantly associated with isoniazid MIC (minimum *p*-value *p*<10^-8.5^, **Supplementary Figure 8**). Rv1219c represses transcription of the Rv1217c-Rv1218c multidrug efflux transport system [38]. It binds two motifs, a high-affinity intergenic sequence in the operon’s promoter, and a low-affinity intergenic sequence immediately upstream of *Rv1218c* [38]. The peak signal of association coincides with the C-terminal amino acids 188-189 in the low-affinity binding domain of Rv1219c. Multiple extremely low frequency oligopeptides were associated with increased MIC, present in just one or two genomes. In contrast, oligopeptides containing the reference alleles at codons 188-189 were present in 8919/8929 genomes and strongly associated with decreased MIC (e.g. EVYT**EG**LLADR, -log_10_*p* = 8.46, β = −3.63, present in 8919/8929 genomes). Substitutions at these positions may therefore derepress the multidrug efflux transport system. Indeed, overexpression of Rv1218c has been observed to correlate with higher isoniazid MIC in vitro [39].

### Second-line drugs

#### Amikacin and kanamycin

Oligopeptides in *PPE42* (*Rv2608*) were significantly associated with aminoglycoside MIC, for both amikacin and kanamycin (minimum *p*-value *p*<10^-12.8^, **Supplementary Figure 9**). PPE42 is an outer membrane-associated PPE-motif family protein and potential B cell antigen. It elicits a high humoral and low T cell response [40] and is one of four antigens in the vaccine candidate ID93 [41]. The C-terminal major polymorphic tandem repeats (MPTRs) contain a region of high antigenicity [40]. The peak association with MIC occurred halfway along the coding sequence. The oligopeptides most associated with higher MIC captured a premature stop codon at position 290 (e.g. PLLE**\***AARFIT, amikacin -log_10_*p* = 11.25, β = 3.12, present in 38/8430 genomes; kanamycin -log_10_*p* = 10.25, β = 2.33, present in 40/8748 genomes). A nearby premature stop codon at amino acid 484 was previously identified in a multi-drug resistant strain [42], supporting the proposition that truncation of PPE42 enhances aminoglycoside resistance.

#### Ethionamide

Oligopeptides and oligonucleotides upstream and within the transcriptional regulator *whiB7* (*Rv3197A*) were significantly associated with ethionamide MIC (minimum *p*-value *p*<10^-18.2^, **Supplementary Figure 10**). Oligonucleotides associated with higher MIC captured a single-base guanine deletion 177 bases upstream of *whiB7*, within the 5’ untranslated region [43] (e.g. AACCGTGTCGCCGCCGCGACTGACGAGTCCT, -log_10_*p* = 18.18, β = 2.16, present in 46/8287 genomes), while oligopeptides associated with higher MIC captured multiple substitutions within the AT-hook motif known to bind AT-rich sequences [44, 45] (e.g. DQGSIVSQQHP, -log_10_*p* = 10.85, β = 1.96, present in 22/8287 genomes). Substitutions in the AT-hook motif may disrupt the binding with the *whiB7* promoter sequence, while deletions upstream of *whiB7* have been shown to result in overexpression of WhiB7 [46]. WhiB7 is induced by antibiotic treatment and other stress conditions and activates its own expression along with other drug resistance genes, for example *tap* and *erm* [45]. Variants in and upstream of another *whiB*-like transcriptional regulator, *whiB6*, were previously found to be associated with resistance to ethionamide [19, 47], capreomycin, amikacin, kanamycin and ethambutol [22, 23]. WhiB7 has been implicated in cross-resistance to multiple drugs, including macrolides, tetracyclines and aminoglycosides [45, 46], however activation of WhiB7 is not induced by all antibiotics, for instance isoniazid [43]. Interestingly, oligopeptides and oligonucleotides in or upstream of *whiB7* were not found to be significantly associated with any of the other 12 antimicrobials. This could indicate yet another mechanism by which *whiB7* is involved in resistance to anti-tuberculosis drugs.

#### Levofloxacin

Oligopeptides in *tlyA* (*Rv1694*) were significantly associated with MIC of the fluoroquinolone levofloxacin (minimum *p*-value *p*<10^-7.8^, **Supplementary Figure 11**). *tlyA* encodes a methyltransferase which methylates ribosomal RNA. Variants in *tlyA*, including loss-of-function mutations, confer resistance to the aminoglycosides viomycin and capreomycin [48] by knocking out its methyltransferase activity [49]. An extremely low frequency oligopeptide was associated with increased MIC, and captured a one-nucleotide adenosine insertion between positions 590 and 591 in codon 198 in a conserved region [50]. In contrast, oligopeptides containing the reference alleles in this region were associated with decreased MIC (e.g. GKGQVGPGGV**V**, -log_10_*p* = 7.83, β = −1.86, present in 7281/7300 genomes). The resulting frameshift likely mimics the knockout effect of deleting the 27 C-terminal residues of TlyA, which ablates methyltransferase activity [51]. While loss-of-function mutations conferring antimicrobial resistance were previously reported to specifically increase aminoglycoside MIC, fluoroquinolones were not investigated [52]. The signal in *tlyA* may therefore reveal genuine, previously unidentified cross-resistance.

#### Rifabutin

Oligonucleotides in *cysA2* (*Rv0815c*) and *cysA3* (*Rv3117*) were significantly associated with rifabutin MIC (minimum *p*-value *p*<10^-7.7^, **Supplementary Figure 12**). They encode identical proteins, which are putative uncharacterised thiosulfate:cyanide sulfurtransferases, known as rhodaneses, belonging to the essential sulfur assimilation pathway, secreted during infection [53]. No genome-wide significant signals associated specific oligopeptides or oligonucleotides with higher MIC. Significant oligonucleotides that aligned to *cysA2* and *cysA3* were associated with lower MIC. They captured two variants: a synonymous nucleotide substitution, a thymine at position 117 in codon 39, and a non-synonymous nucleotide substitution, a guanine at position 103 inducing amino acid substitution 35D (e.g. CATAT**G**ACCGTGACCATAT**T**GCCGGCGCGAT, -log_10_*p* = 7.74, β = −2.65, present in 9396/9418 genomes). These positions coincide with the rhodanese characteristic signature in the N-terminal region, important for rhodanese stability [54]. However, the mechanism of resistance against rifabutin remains to be elucidated.

### New and repurposed drugs

#### Bedaquiline

Oligonucleotides situated in the region of overlap at the 3’ ends of *amiA2* (*Rv2363*) and *era* (*Rv2364c*) were significantly associated with bedaquiline MIC (minimum *p*-value *p*<10^-10.5^, **Supplementary Figure 13**). These genes encode an amidase and a GTPase, respectively, on opposite strands. Of the two top oligonucleotides associated with higher MIC, the first captures two substitutions that are synonymous in *era*, 7-19 nucleotides upstream of the stop codon, and 3’ non-coding in *amiA2*, 4-16 nucleotides downstream of the stop codon (e.g. CCCCAAACAGCT**T**GGCCGACTGGG**G**TTTTAG, -log_10_*p* = 10.47, β = 1.26, present in 7919/8009 genomes). The second additionally captures a variant that induces a non-synonymous guanine substitution at position 1451 in *amiA2*, and is 3’ intergenic in *era*, one nucleotide downstream of the stop codon (e.g. CAAACAGCT**T**GGCCGACTGGG**G**TTTTAG**C**TC, -log_10_*p* = 7.87, β = 0.88, present in 7898/8009 genomes). Interestingly, AmiA2 has previously been identified at lower abundance in MDR compared to sensitive isolates [55], and Era (but not AmiA2) has been shown to be required for optimal growth of H37Rv [56]. These variants may therefore enhance tolerance to bedaquiline.

#### Clofazimine

Oligopeptides in *cyp142* (*Rv3518c*), which encodes a cytochrome P450 enzyme with substrates of cholesterol/cholest-4-en-3-one, were significantly associated with clofazimine MIC (minimum *p*-value *p*<10^-12.2^, **Supplementary Figure 14**). Oligopeptides associated with higher MIC captured the amino acid residue 176I (e.g. EDFQIT**I**DAFA, -log_10_*p* = 7.99, β = 1.14, present in 100/7297 genomes). The association signal falls within the F *α*-helix of CYP142, which lines the entrance to the active site with largely hydrophobic residues, forming part of the substrate binding pocket [57, 58]. Homology with CYP125 suggests that residue 176 captured by the GWAS is within 5 Å of the binding substrate [58]. The potential for cytochrome P450 enzymes as targets for anti-tuberculosis drugs has been highlighted [59]; CYP142 is inhibited by azole drugs [59] and has been found to form a tight complex with nitric oxide (NO) [60]. The anti-mycobacterial activity of clofazimine has been shown to produce reactive oxygen species [61], therefore the substitution identified by the GWAS may disrupt the binding of NO to CYP142. Methionine and isoleucine are both hydrophobic residues, so the mechanism for how this would disrupt binding is unknown.

#### Delamanid

Oligonucleotides in *pknH* (*Rv1266c*), which encodes a serine/threonine-protein kinase, were significantly associated with delamanid MIC (minimum *p*-value *p*<10^-30.2^, **Supplementary Figure 15**). Delamanid is a prodrug activated by deazaflavin-dependent nitroreductase which inhibits cell wall synthesis. PknH phosphorylates the adjacent gene product EmbR [62], enhancing its binding of the promoter regions of the *embCAB* operon [63]. Mutations in *embAB* are responsible for ethambutol resistance [64]. The peak GWAS signal localized to the C-terminal periplasmic domain of PknH [62]. Oligonucleotides below our MAF threshold captured extremely low frequency triplet deletions of either ACG at nucleotides 1645-7 or GAC at nucleotides 1644-6. In contrast, oligonucleotides containing the reference alleles in this region were associated with decreased MIC (e.g. CAAGACGGTCACCGTCACGAATAAGGCCAAG, -log_10_*p* = 30.21, β = −3.29, present in 7555/7564 genomes). These variants likely disrupt intramolecular disulphide binding linking the two highly conserved alpha helices that form the V-shaped cleft of the C-terminal sensor domain [65]. Since NO is released upon activation of DLM, and deletion of PknH alters sensitivity to nitrosative and oxidative stresses [66], these rare variants may alter tolerance to delamanid mediated by NO.

#### Linezolid

Oligonucleotides in *vapB20* (*Rv2550c*) were significantly associated with linezolid MIC (minimum *p*-value *p*<10^-8.6^, **Supplementary Figure 16**). VapB20 is an antitoxin cotranscribed with its complementary toxin VapC20 [67]. The latter modifies 23S rRNA [68], the target of linezolid which inhibits protein synthesis by competitively binding 23S rRNA. The peak signal in *vapB20* occurred just upstream of the promotor and VapB20 binding sites, 21 nucleotides upstream of the −35 region [68]. Oligonucleotides below our MAF threshold associated with increased MIC shared a cytosine 33 nucleotides upstream of *vapB20*, replacing the reference nucleotide thymine which was associated with decreased MIC (e.g. GAATCGG**A**TGCTTGCCGCTGGCTGCCGAGTT, -log_10_*p* = 8.60, β = −2.02, present in 6724/6732 genomes). This substitution may derepress the toxin, which could interrupt linezolid binding by cleaving the Sarcin-Ricin loop of 23S rRNA.

## Discussion

In this study we tested oligopeptides and oligonucleotides for association with quantitative MIC measurements for 13 antimicrobials to identify novel resistance determinants. Analysing MIC rather than binary resistance phenotypes enabled identification of variants that cause subtle changes in MIC. This is important, on the one hand, because higher rifampicin and isoniazid MIC in sensitive isolates are associated with increased risk of relapse after treatment [69]. Conversely, low-level resistance among isolates resistant to RIF and isoniazid mediated by particular mutations may sometimes be overcome by increasing the drug dose, or replacing rifampicin with rifabutin, rather than changing to less desirable drugs with worse side effects [70,71,72,73,74]. The investigation of MIC was particularly effective at increasing heritability for the new and repurposed drugs.

The MICs were positively correlated between many drugs, particularly amongst first-line drugs. Consequently, many of the 10,228 isolates we studied were MDR and XDR. In GWAS, this generates artefactual cross resistance, in which variants that cause resistance to one drug appear associated with other drugs to which they do not confer resistance. In practice, it is difficult to distinguish between associations that are causal versus artefactual without experimental evidence. Nevertheless, we found frequent evidence of artefactual cross resistance: several genes and intergenic regions featured among the top 20 strongest signals of association to multiple drugs, including *rpoB* (12 drugs), *embB* (7), *fabG1* (7), *rrs* (6), *gyrA* (6), *katG* (6), *lprF:Rv1371* (6), *pncA* (5), *ethA* (4), *Rv3183:Rv3188* (4) *dxs2:Rv3382c* (4), *rpsL* (3) and *moaC3:Rv3327* (3). Among previously catalogued variants, we observed that the estimated effect sizes were usually larger in magnitude for significant true associations than significant artefactual associations (**Supplementary figure 4**). In future GWAS, this relationship could help tease apart true versus artefactual associations when a uncatalogued variant is associated with multiple drugs.

We focused on variants in the top twenty most significant genes identified by GWAS for each of the 13 drugs, classifying significant oligopeptides and oligonucleotides according to whether the variants they tagged were previously catalogued among known resistance determinants, or not. While the interpretation of oligopeptides and oligonucleotides required manual curation to determine the underlying variants they tagged, the approach had the advantage of avoiding reference-based variant calling which can miss important signals, particularly at difficult-to-map regions. For 8/13 drugs with previously catalogued resistance determinants, the most significant GWAS signal in CRyPTIC was a previously catalogued variant. Among the uncatalogued variants there are promising signals of association, including in the *Rv1218c* promoter binding site of the transcriptional repressor *Rv1219c* (associated with MIC for isoniazid) upstream of the *vapBC20* operon that cleaves 23S rRNA (linezolid) and in the region encoding a helix lining the active site of *cyp142* (clofazimine). These variants would benefit from further investigation via replication studies in independent populations, experimental exploration of proposed resistance mechanisms, or both.

We elected to classify significant variants as catalogued versus uncatalogued, rather than known versus novel, for several reasons. The catalogues represent a concrete, pre-existing knowledgebase collated by expert groups for use in a clinical context [17, 11]. We chose [17, 11] as they are the most recent and up to date catalogues available for the drugs we investigated. The inclusion criteria for variants to be considered catalogued are therefore stringent; it follows that a class of variants exist that have been reported in the literature but not assimilated into the catalogues [17, 11]. The literature is vast and heterogenous, with evidence originating from molecular, clinical and genome-wide association studies. Inevitably, some uncatalogued variants in the literature will be false positives, while others will be real but did not meet the standard of evidence or clinical relevance for cataloguing. Evidence from CRyPTIC that supports uncatalogued variants in the latter group is of equal or greater value than the discovery of completely novel variants, because it contributes to a body of independent data supporting their involvement. For instance, *gyrB* did not appear in the catalogues we used for moxifloxacin [17, 11]. Yet our rediscovery of *gyrB* 501D complements published reports associating the substitution with moxifloxacin resistance [75,76,77], strongly enhancing the evidence in favour of inclusion in future catalogues. Indeed, the recent WHO prediction catalogue, published after the completion of this study and which draws on the CRyPTIC data analysed here includes the E501D resistance-associated variant [78]. Moreover, of the five new genes added to the forthcoming WHO catalogue [78] but not featuring in the catalogues [17, 11] used here – *eis* (amikacin), *ethA* (ethionamide), *inhA* (ethionamide), *rplC* (linezolid), *gyrB* (moxifloxacin) – we identify all as containing significant variants by GWAS except one, *eis* (amikacin).

The combination of a very large dataset exceeding 10,000 isolates and quantification of resistance via MIC enabled the CRyPTIC study to attribute a large proportion of fine-grained variability in antimicrobial resistance in *M. tuberculosis* to genetic variation. Compared to a parallel analysis of binary resistance phenotypes in the same samples, we observed an increase in heritability of 26.1-37.1% for the new and repurposed drugs bedaquiline, clofazimine, delamanid and linezolid. The improvement was most striking for delamanid, whose heritability was not significantly different to zero for the binary resistance phenotype. In contrast, the scope for improvement was marginal for the better-studied drugs isoniazid and rifampicin, where MIC heritabilities of 94.6-94.9% were achieved. This demonstrates the ability of additive genetic variation to explain almost all the phenotypic variability in MIC for these drugs. Nevertheless, we were still able to find uncatalogued hits for these drugs. The very large sample size also contributed to increased heritability compared to previous pioneering studies. Compared to Farhat et al 2019 [22] who estimated the heritability of MIC phenotypes in 1452 isolates, we observed increases in heritability of 2.0% (kanamycin), 3.3% (amikacin), 14.0% (isoniazid), 10.8% (rifampicin), 11.2% (ethambutol) and 19.4% (moxifloxacin). Furthermore, many of the uncatalogued signals we report here as significant detected rare variants at below 1% minor allele frequency, underlining the ability of very large-scale studies to improve our understanding of antimicrobial resistance not only quantitatively, but to tap otherwise unseen rare variants that reveal new candidate resistance mechanisms.

## Materials and Methods

### Sampling frames

CRyPTIC collected isolates from 27 countries worldwide, oversampling for drug resistance, as described in detail in [31]. Clinical isolates were subcultured for 14 days before inoculation onto one of two CRyPTIC designed 96-well microtiter plates manufactured by ThermoFisher. The first plate used (termed UKMYC5) contained doubling-dilution ranges for 14 different antibiotics, the second (UKMYC6) removed para-aminosalicylic acid due to poor results on the plate [30] and changed the concentration of some drugs. Para-aminosalicylic acid was therefore not included in the GWAS analyses. Phenotype measurements were determined to be high quality, and included in the GWAS analyses, if three independent methods (Vizion, AMyGDA and BashTheBug) agreed on the value [31]. Sequencing pipelines differed slightly between the CRyPTIC sites, but all sequencing was performed using Illumina, providing an input of matched pair FASTQ files containing the short reads.

15,211 isolates were included in the initial CRyPTIC dataset with both genomes and phenotype measurements after passing genome quality control filters [31, 79], however some plates were later removed due to problems identified at some laboratories with inoculating the plates [31]. Genomes were also excluded if they met any of the following criteria, determined by removing samples at the outliers of the distributions: (i) no high quality phenotypes for any drugs; (ii) total number of contigs > 3000; (iii) total bases in contigs < 3.5×10^6^ or > 5×10^6^; (iv) number of unique oligonucleotides < 3.5×10^6^ or > 5×10^6^; (v) sequencing read length not 150/151 bases long. This gave a GWAS dataset of 10,422 genomes used to create the variant presence/absence matrices. We used Mykrobe [80,79,81] to identify *Mycobacterium* genomes not belonging to lineages 1-4 or representing mixtures of lineages. This led to the exclusion of 193 genomes, which were removed from GWAS by setting the phenotypes to NA. The number of genomes with a high quality phenotype for at least one of the 13 drugs was therefore 10,228. Of these 533 were lineage 1, 3581 lineage 2, 805 lineage 3, and 5309 lineage 4. Due to rigorous quality control described above, only samples with high quality phenotypes were tested for each drug, resulting in a range of 6,388-9,418 genomes used in each GWAS.

### Phylogenetic inference

A pairwise distance matrix was constructed for the full CRyPTIC dataset based on variant calls [79]. For visualisation of the dataset, a neighbour joining tree was built from the distance matrix using the ape package in R and subset to the GWAS dataset. Negative branch lengths were set to zero, and the length was added to the adjacent branch. The branch lengths were square rooted and the tree annotated by lineages assigned by Mykrobe [80].

### Oligonucleotide/oligopeptide counting

To capture SNP-based variation, indels, and combinations of SNPs and indels, we pursued oligonucleotide and oligopeptide-based approaches, focusing primarily on oligopeptides. Where helpful for clarifying results, we interpreted significant associations using oligonucleotides. Sequence reads were assembled *de novo* using Velvet Optimiser [82] with a starting lower hash value of half the read length, and a higher hash value of the read length minus one; if these were even numbers they were lowered by one. If the total sequence length of the reads in the FASTQ file was greater than 1×10^9^, then the reads were randomly subsampled prior to assembly down to a sequence length of 1×10^9^ which is around 227x mean coverage. For the oligopeptide analysis, each assembly contig was translated into the six possible reading frames in order to be agnostic to the correct reading frame. 11 amino acid long oligopeptides were counted in a one amino acid sliding window from these translated contigs. 31bp nucleotide oligonucleotides were also counted from the assembled contigs using dsk [83]. For both oligonucleotide and oligopeptide analyses, a unique set of variants across the dataset was created, with the presence or absence of each unique variant determined per genome. An oligonucleotide/oligopeptide was counted as present within a genome if it was present at least once. This resulted in 60,103,864 oligopeptides and 34,669,796 oligonucleotides. Of these, 10,510,261 oligopeptides and 5,530,210 oligonucleotides were variably present in the GWAS dataset of 10,228 genomes.

### Oligonucleotide/oligopeptide alignment

We used the surrounding context of the contigs that the oligopeptides/oligonucleotides were identified in to assist with their alignment. First, we aligned the contigs of each genome to the H37Rv reference genome [84] using nucmer [85], keeping alignments above 90% identity, assigning a H37Rv position to each base in the contig. Version 3 of the H37Rv strain (NC_000962.3) was used as the reference genome throughout the analysis. All numbering refers to the start positions in the H37Rv version 3 GenBank file. This gave a position for each oligonucleotide identified in the contigs, and after translating the six possible reading frames of the contig, each oligopeptide too. Each oligonucleotide/ oligopeptide was assigned a gene or intergenic region (IR) or both in each genome. These variant/gene combinations were then merged across all genomes into unique variant/gene combinations, where a variant could be assigned to multiple genes or intergenic regions. Variant/gene combinations were then kept if seen in five or more genomes. In some specific regions where significant oligonucleotides or oligopeptides appeared to be capturing an invariant region, a threshold of just one genome was used to visualise low frequency variants in the region. This was used only for interpretation of the signal in the region, and not for the main analyses. To improve alignment for the most significant genes and intergenic regions, all oligonucleotides/oligopeptides in the gene/IR plus those that aligned to a gene/IR within 1kb were re-aligned to the region using BLAST. Alignments were kept if above 70% identity, recalculated along the whole length of the oligonucleotide/oligopeptide assuming the whole oligonucleotide/oligopeptide aligned. Oligopeptides were aligned to all six possible reading frames and only the correct reading frame was interpreted. An oligonucleotide/oligopeptide was interpreted as unaligned if it did not align to any of the six possible reading frames. A region was determined to be significant if it contained significant oligopeptides above a minor allele frequency (MAF) of 0.1% that were assigned to the region that also aligned using BLAST. If no significant oligopeptides aligned to the correct reading frame of a protein, or if the significant region was intergenic, then oligonucleotides were assessed.

### Covariates

Isolates were sampled from 9 sites and minimum inhibitory concentrations (MIC) were measured on two versions of the quantitative microtiter plate assays, UKMYC5 and UKMYC6 [31]. UKMYC6 contained adjusted concentrations for some drugs. Therefore in order to account for possible batch effects, we controlled for site plus plate type in the LMM by coding them as binary variables. These plus an intercept were included as covariates in the GWAS analyses.

### Testing for locus effects

We performed association testing using linear mixed model (LMM) analyses implemented in the software GEMMA to control for population structure [32]. Significance was calculated using likelihood ratio tests. We computed the relatedness matrix from the presence/absence matrix using Java code which calculates the centred relatedness matrix. GEMMA was run using no minor allele frequency cut-off to include all variants. When assessing the most significant regions for each drug, we excluded oligopeptides below 0.1% MAF. To understand the full signal at these regions, oligo-peptides and nucleotides were visualised in alignment figures to interpret the variants captured. When assessing the gene highlighted for each drug, we assessed the LD (*r*^2^) of the most significant oligo-peptide or nucleotide in the gene with all other top oligo-peptides or nucleotides for the top 20 genes for the drug. The top variants in the genes noted were not in high LD with known causal variants, in some cases they were in LD with other top 20 gene hits that were less significant.

### Correcting for multiple testing

Multiple testing was accounted for by applying a Bonferroni correction calculated for each drug. The unit of correction for all studies was the number of unique “phylopatterns”, i.e. the number of unique partitions of individuals according to variant presence/absence for the phenotype tested. An oligopeptide/oligonucleotide was considered to be significant if its *p*-value was smaller than *α*/*n*_p_, where we took *α* =0.05 to be the genome-wide false positive rate (i.e. family-wide error rate, FWER) and *n*_p_ to be the number of unique phylopatterns above 0.1% MAF in the genomes tested for the particular drug. The -log_10_*p* significance thresholds for the oligopeptide analyses were: 7.69 (amikacin, kanamycin), 7.65 (bedaquiline), 7.64 (clofazimine, levofloxacin), 7.67 (delamanid, ethionamide), 7.62 (ethambutol, linezolid), 7.70 (isoniazid), 7.60 (moxifloxacin), 7.71 (rifabutin) and 7.68 (rifampicin). The -log_10_*p* significance thresholds for the oligonucleotide analyses were: 7.38 (amikacin, kanamycin), 7.34 (bedaquiline, clofazimine, levofloxacin), 7.36 (delamanid, ethionamide), 7.32 (ethambutol), 7.39 (isoniazid, rifabutin), 7.33 (linezolid), 7.31 (moxifloxacin) and 7.37 (rifampicin).

### Estimating sample heritability

Sample heritability is the proportion of the phenotypic variation that can be explained by the bacterial genotype assuming additive effects. This was estimated using the LMM null model in GEMMA [32] from the presence vs. absence matrices for both oligopeptides and oligonucleotides separately. Sample heritability was estimated for the MIC phenotype as well as for the binary sensitive vs. resistant phenotype. The binary phenotypes were determined using the epidemiological cutoff (ECOFF), defined as the MIC that encompasses 99% of wild type isolates [31], all those below the ECOFF were considered susceptible, and those above the ECOFF were considered to be resistant.

## Acknowledgements

### Acknowledgements – funders

This work was supported by Wellcome Trust/Newton Fund-MRC Collaborative Award (200205/Z/15/Z); and Bill & Melinda Gates Foundation Trust (OPP1133541). Oxford CRyPTIC consortium members are funded/supported by the National Institute for Health Research (NIHR) Oxford Biomedical Research Centre (BRC), the views expressed are those of the authors and not necessarily those of the NHS, the NIHR or the Department of Health, and the National Institute for Health Research (NIHR) Health Protection Research Unit in Healthcare Associated Infections and Antimicrobial Resistance, a partnership between Public Health England and the University of Oxford, the views expressed are those of the authors and not necessarily those of the NIHR, Public Health England or the Department of Health and Social Care. J.M. is supported by the Wellcome Trust (203919/Z/16/Z). Z.Y. is supported by the National Science and Technology Major Project, China Grant No. 2018ZX10103001. K.M.M. is supported by EMBL’s EIPOD3 programme funded by the European Union’s Horizon 2020 research and innovation programme under Marie Skłodowska Curie Actions. T.C.R. is funded in part by funding from Unitaid Grant No. 2019-32-FIND MDR. R.S.O. is supported by FAPESP Grant No. 17/16082-7. L.F. received financial support from FAPESP Grant No. 2012/51756-5. B.Z. is supported by the National Natural Science Foundation of China (81991534) and the Beijing Municipal Science & Technology Commission (Z201100005520041). N.T.T.T. is supported by the Wellcome Trust International Intermediate Fellowship (206724/Z/17/Z). G.T. is funded by the Wellcome Trust. R.W. is supported by the South African Medical Research Council. J.C. is supported by the Rhodes Trust and Stanford Medical Scientist Training Program (T32 GM007365). A.L. is supported by the National Institute for Health Research (NIHR) Health Protection Research Unit in Respiratory Infections at Imperial College London. S.G.L. is supported by the Fonds de Recherche en Santé du Québec. C.N. is funded by Wellcome Trust Grant No. 203583/Z/16/Z. A.V.R. is supported by Research Foundation Flanders (FWO) under Grant No. G0F8316N (FWO Odysseus). G.M. was supported by the Wellcome Trust (098316, 214321/Z/18/Z, and 203135/Z/16/Z), and the South African Research Chairs Initiative of the Department of Science and Technology and National Research Foundation (NRF) of South Africa (Grant No. 64787). The funders had no role in the study design, data collection, data analysis, data interpretation, or writing of this report. The opinions, findings and conclusions expressed in this manuscript reflect those of the authors alone. L.G. was supported by the Wellcome Trust (201470/Z/16/Z), the National Institute of Allergy and Infectious Diseases of the National Institutes of Health under award number 1R01AI146338, the GOSH Charity (VC0921) and the GOSH/ICH Biomedical Research Centre (www.nihr.ac.uk). A.B. is funded by the NDM Prize Studentship from the Oxford Medical Research Council Doctoral Training Partnership and the Nuffield Department of Clinical Medicine. D.J.W. is supported by a Sir Henry Dale Fellowship jointly funded by the Wellcome Trust and the Royal Society (Grant No. 101237/Z/13/B) and by the Robertson Foundation. A.S.W. is an NIHR Senior Investigator. T.M.W. is a Wellcome Trust Clinical Career Development Fellow (214560/Z/18/Z). A.S.L. is supported by the Rhodes Trust. R.J.W. receives funding from the Francis Crick Institute which is supported by Wellcome Trust, (FC0010218), UKRI (FC0010218), and CRUK (FC0010218). T.C. has received grant funding and salary support from US NIH, CDC, USAID and Bill and Melinda Gates Foundation. The computational aspects of this research were supported by the Wellcome Trust Core Award Grant Number 203141/Z/16/Z and the NIHR Oxford BRC. Parts of the work were funded by the German Center of Infection Research (DZIF). The Scottish Mycobacteria Reference Laboratory is funded through National Services Scotland. The Wadsworth Center contributions were supported in part by Cooperative Agreement No. U60OE000103 funded by the Centers for Disease Control and Prevention through the Association of Public Health Laboratories and NIH/NIAID grant AI-117312. Additional support for sequencing and analysis was contributed by the Wadsworth Center Applied Genomic Technologies Core Facility and the Wadsworth Center Bioinformatics Core. SYNLAB Holding Germany GmbH for its direct and indirect support of research activities in the Institute of Microbiology and Laboratory Medicine Gauting. N.R. thanks the Programme National de Lutte contre la Tuberculose de Madagascar.

Computation used the Oxford Biomedical Research Computing (BMRC) facility, a joint development between the Wellcome Centre for Human Genetics and the Big Data Institute supported by Health Data Research UK and the NIHR Oxford Biomedical Research Centre.

### Wellcome Trust Open Access

This research was funded in part, by the Wellcome Trust/Newton Fund-MRC Collaborative Award [200205/Z/15/Z]. For the purpose of Open Access, the author has applied a CC BY public copyright licence to any Author Accepted Manuscript version arising from this submission. This research was funded, in part, by the Wellcome Trust [214321/Z/18/Z, and 203135/Z/16/Z]. For the purpose of open access, the author has applied a CC BY public copyright licence to any Author Accepted Manuscript version arising from this submission.

### Acknowledgements – people

We thank Faisal Masood Khanzada and Alamdar Hussain Rizvi (NTRL, Islamabad, Pakistan), Angela Starks and James Posey (Centers for Disease Control and Prevention, Atlanta, USA), and Juan Carlos Toro and Solomon Ghebremichael (Public Health Agency of Sweden, Solna, Sweden), Iñaki Comas and Álvaro Chiner-Oms (Instituto de Biología Integrativa de Sistemas, Valencia, Spain; CIBER en Epidemiología y Salud Pública, Valencia, Spain; Instituto de Biomedicina de Valencia, IBV-CSIC, Valencia, Spain).

### Ethics Statement

Approval for CRyPTIC study was obtained by Taiwan Centers for Disease Control IRB No. 106209, University of KwaZulu Natal Biomedical Research Ethics Committee (UKZN BREC) (reference BE022/13) and University of Liverpool Central University Research Ethics Committees (reference 2286), Institutional Research Ethics Committee (IREC) of The Foundation for Medical Research, Mumbai (Ref nos. FMR/IEC/TB/01a/2015 and FMR/IEC/TB/01b/2015), Institutional Review Board of P.D. Hinduja Hospital and Medical Research Centre, Mumbai (Ref no. 915-15-CR [MRC]), scientific committee of the Adolfo Lutz Institute (CTC-IAL 47-J / 2017) and in the Ethics Committee (CAAE: 81452517.1.0000.0059) and Ethics Committee review by Universidad Peruana Cayetano Heredia (Lima, Peru) and LSHTM (London, UK).

### Members of the CRyPTIC consortium (in alphabetical order)

Correspondence to: Daniel J Wilson

Sarah G Earle^4^, Daniel J Wilson^4^, Ivan Barilar^29^, Simone Battaglia^1^, Emanuele Borroni^1^, Angela Pires Brandao^2,3^, Alice Brankin^4^, Andrea Maurizio Cabibbe^1^, Joshua Carter^5^, Daniela Maria Cirillo^1^, Pauline Claxton^6^, David A Clifton^4^, Ted Cohen^7^, Jorge Coronel^8^, Derrick W Crook^4^, Viola Dreyer^29^, Vincent Escuyer^9^, Lucilaine Ferrazoli^3^, Philip W Fowler^4^, George Fu Gao^10^, Jennifer Gardy^11^, Saheer Gharbia^12^, Kelen Teixeira Ghisi^3^, Arash Ghodousi^1,13^, Ana Luíza Gibertoni Cruz^4^, Louis Grandjean^33^, Clara Grazian^14^, Ramona Groenheit^44^, Jennifer L Guthrie^15,16^, Wencong He^10^, Harald Hoffmann^17,18^, Sarah J Hoosdally^4^, Martin Hunt^19,4^, Zamin Iqbal^19^, Nazir Ahmed Ismail^20^, Lisa Jarrett^21^, Lavania Joseph^20^, Ruwen Jou^22^, Priti Kambli^23^, Rukhsar Khot^23^, Jeff Knaggs^19,4^, Anastasia Koch^24^, Donna Kohlerschmidt^9^, Samaneh Kouchaki^4,25^, Alexander S Lachapelle^4^, Ajit Lalvani^26^, Simon Grandjean Lapierre^27^, Ian F Laurenson^6^, Brice Letcher^19^, Wan-Hsuan Lin^22^, Chunfa Liu^10^, Dongxin Liu^10^, Kerri M Malone^19^, Ayan Mandal^28^, Mikael Mansjö^44^, Daniela Matias^21^, Graeme Meintjes^24^, Flávia de Freitas Mendes^3^, Matthias Merker^29^, Marina Mihalic^18^, James Millard^30^, Paolo Miotto^1^, Nerges Mistry^28^, David Moore^31,8^, Kimberlee A Musser^9^, Dumisani Ngcamu^20^, Hoang Ngoc Nhung^32^, Stefan Niemann^29, 48^, Kayzad Soli Nilgiriwala^28^, Camus Nimmo^33^, Nana Okozi^20^, Rosangela Siqueira Oliveira^3^, Shaheed Vally Omar^20^, Nicholas Paton^34^, Timothy EA Peto^4^, Juliana Maira Watanabe Pinhata^3^, Sara Plesnik^18^, Zully M Puyen^35^, Marie Sylvianne Rabodoarivelo^36^, Niaina Rakotosamimanana^36^, Paola MV Rancoita^13^, Priti Rathod^21^, Esther Robinson^21^, Gillian Rodger^4^, Camilla Rodrigues^23^, Timothy C Rodwell^37,38^, Aysha Roohi^4^, David Santos-Lazaro^35^, Sanchi Shah^28^, Thomas Andreas Kohl^29^, Grace Smith^21,12^, Walter Solano^8^, Andrea Spitaleri^1,13^, Philip Supply^39^, Utkarsha Surve^23^, Sabira Tahseen^40^, Nguyen Thuy Thuong Thuong^32^, Guy Thwaites^32,4^, Katharina Todt^18^, Alberto Trovato^1^, Christian Utpatel^29^, Annelies Van Rie^41^, Srinivasan Vijay^42^, Timothy M Walker^4,32^, A Sarah Walker^4^, Robin Warren^43^, Jim Werngren^44^, Maria Wijkander^44^, Robert J Wilkinson^45,46,26^, Penelope Wintringer^19^, Yu-Xin Xiao^22^, Yang Yang^4^, Zhao Yanlin^10^, Shen-Yuan Yao^20^, Baoli Zhu^47^

### Institutions

1 IRCCS San Raffaele Scientific Institute, Milan, Italy

2 Oswaldo Cruz Foundation, Rio de Janeiro, Brazil

3 Institute Adolfo Lutz, São Paulo, Brazil

4 University of Oxford, Oxford, UK

5 Stanford University School of Medicine, Stanford, USA

6 Scottish Mycobacteria Reference Laboratory, Edinburgh, UK

7 Yale School of Public Health, Yale, USA

8 Universidad Peruana Cayetano Heredia, Lima, Perú

9 Wadsworth Center, New York State Department of Health, Albany, USA

10 Chinese Center for Disease Control and Prevention, Beijing, China

11 Bill & Melinda Gates Foundation, Seattle, USA

12 UK Health Security Agency, London, UK

13 Vita-Salute San Raffaele University, Milan, Italy

14 University of New South Wales, Sydney, Australia

15 The University of British Columbia, Vancouver, Canada

16 Public Health Ontario, Toronto, Canada

17 SYNLAB Gauting, Munich, Germany

18 Institute of Microbiology and Laboratory Medicine, IMLred, WHO-SRL Gauting, Germany

19 EMBL-EBI, Hinxton, UK

20 National Institute for Communicable Diseases, Johannesburg, South Africa

21 Public Health England, Birmingham, UK

22 Taiwan Centers for Disease Control, Taipei, Taiwan

23 Hinduja Hospital, Mumbai, India

24 University of Cape Town, Cape Town, South Africa 25 University of Surrey, Guildford, UK

26 Imperial College, London, UK 27 Université de Montréal, Canada

28 The Foundation for Medical Research, Mumbai, India

29 Research Center Borstel, Borstel, Germany

30 Africa Health Research Institute, Durban, South Africa

31 London School of Hygiene and Tropical Medicine, London, UK

32 Oxford University Clinical Research Unit, Ho Chi Minh City, Viet Nam

33 University College London, London, UK

34 National University of Singapore, Singapore 35 Instituto Nacional de Salud, Lima, Perú

36 Institut Pasteur de Madagascar, Antananarivo, Madagascar 37 FIND, Geneva, Switzerland

38 University of California, San Diego, USA

39 Univ. Lille, CNRS, Inserm, CHU Lille, Institut Pasteur de Lille, U1019 - UMR 9017 - CIIL - Center for Infection and Immunity of Lille, F-59000 Lille, France

40 National TB Reference Laboratory, National TB Control Program, Islamabad, Pakistan

41 University of Antwerp, Antwerp, Belgium

42 University of Edinburgh, Edinburgh, UK

43 Stellenbosch University, Cape Town, South Africa 44 Public Health Agency of Sweden, Solna, Sweden

45 Wellcome Centre for Infectious Diseases Research in Africa, Cape Town, South Africa

46 Francis Crick Institute, London, UK

47 Institute of Microbiology, Chinese Academy of Sciences, Beijing, China

48 German Center for Infection Research (DZIF), Hamburg-Lübeck-Borstel-Riems, Germany

