## Supplementary Figures and Tables for "Genome-wide association studies of global *Mycobacterium tuberculosis* resistance to thirteen antimicrobials in 10,228 genomes"

### Supplementary Information

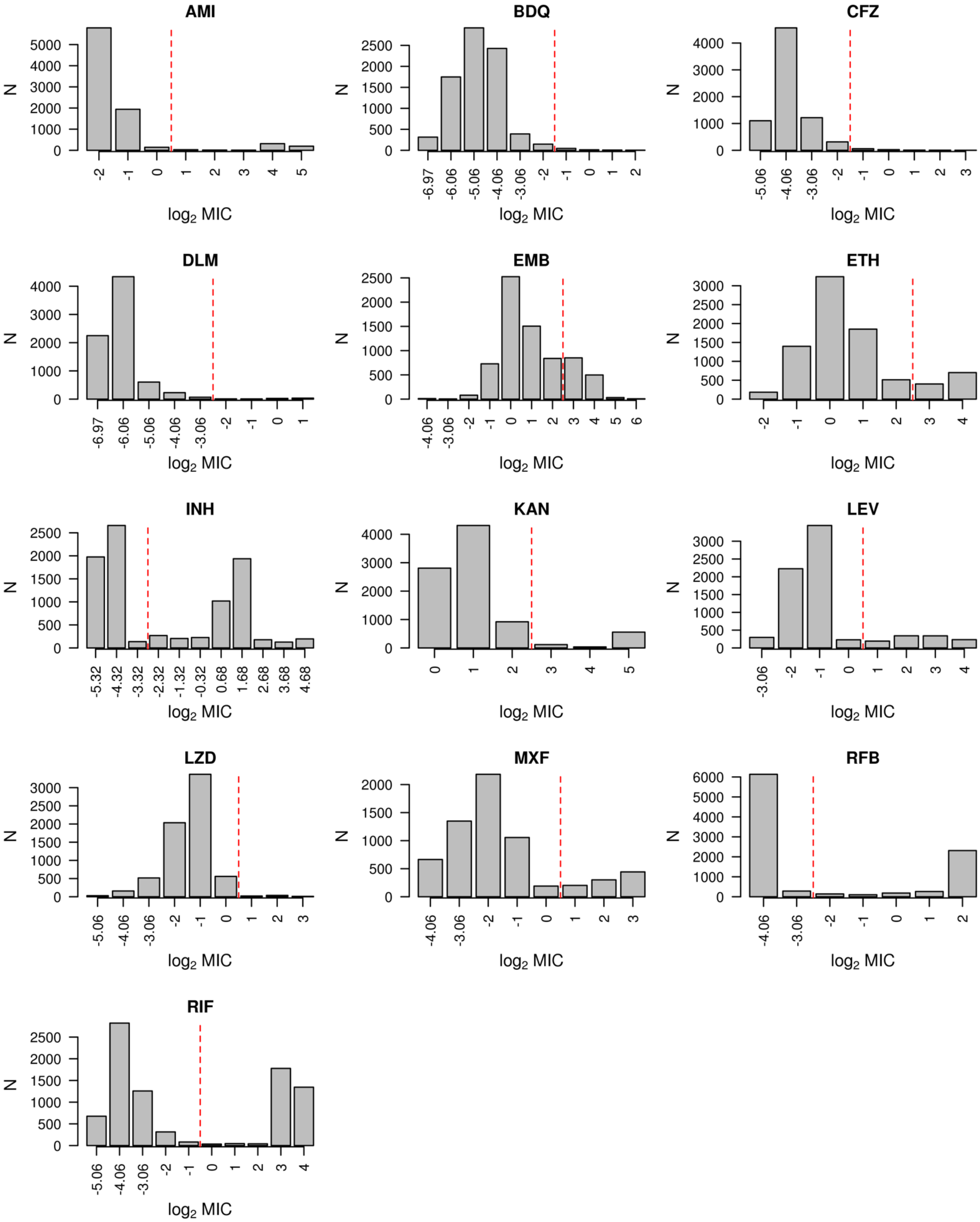


**Supplementary Figure 1** Distributions of the $\log_2$ MIC measurements for all 13 drugs in the GWAS analyses, amikacin (AMI), bedaquiline (BDQ), clofazimine (CFZ), delamanid (DLM), ethambutol (EMB), ethionamide (ETH), isoniazid (INH), kanamycin (KAN), levofloxacin (LEV), linezolid (LZD), moxifloxacin (MXF), rifabutin (RFB) and rifampicin (RIF). The red dashed line indicates the ECOFF, measurements to the left of the ECOFF are considered sensitive, those to the right are considered resistant.

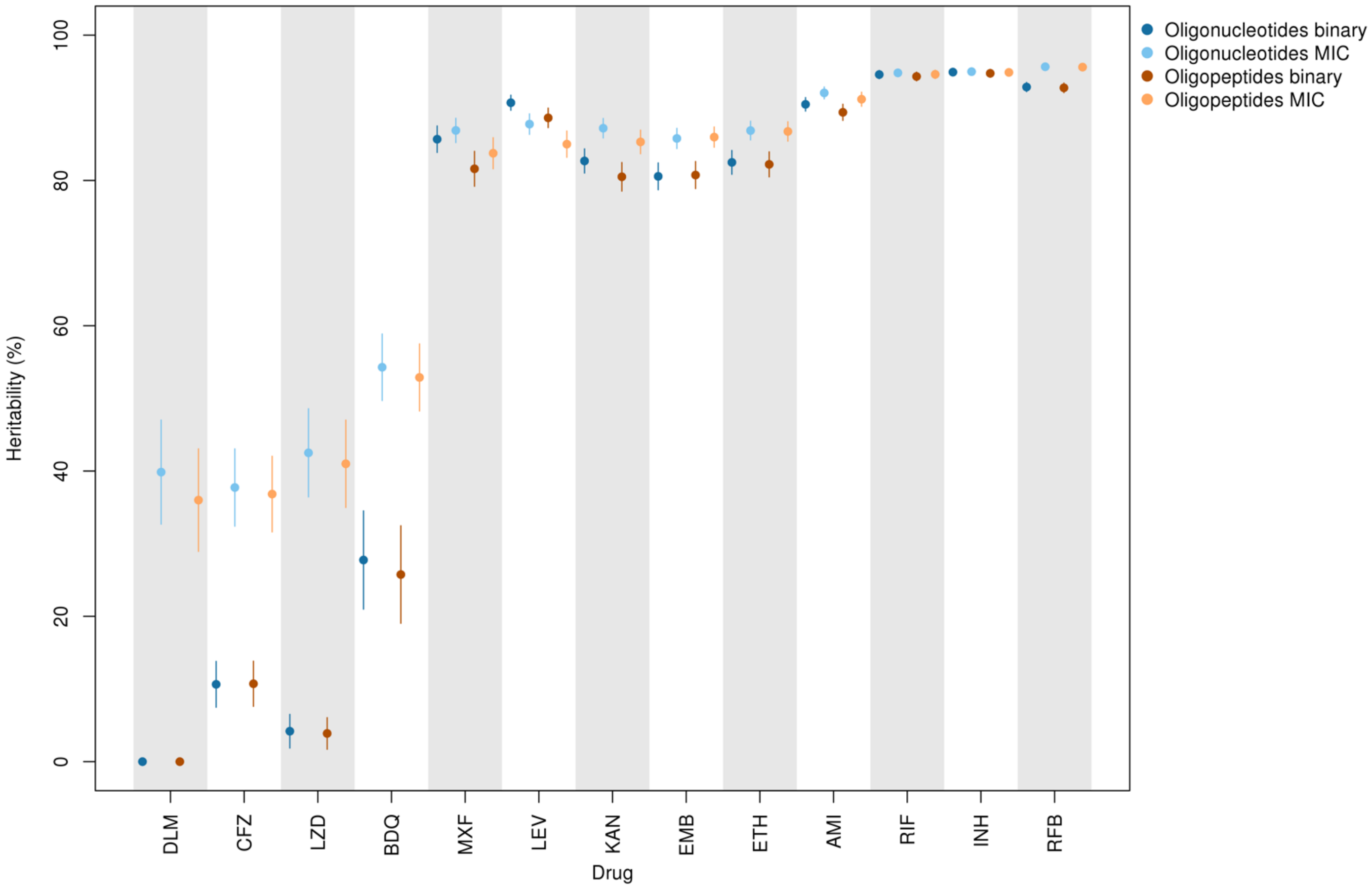


**Supplementary Figure 2** Oligopeptide and oligonucleotide sample heritability estimates for binary resistant vs. sensitive phenotypes compared to semi-quantitiative MIC phenotypes. Sample heritability estimates and 95% confidence intervals are shown for the 13 drugs, DLM (delamanid), clofazimine (CFZ), linezolid (LZD), bedaquiline (BDQ), moxifloxacin (MXF), levofloxacin (LEV), kanamycin (KAN), ethambutol (EMB), ethionamide (ETH), amikacin (AM), rifampicin (RIF), isoniazid (INH), rifabutin (RFB). When estimating heritability of the same phenotype, the oligopeptide and oligonucleotide estimates are very similar.

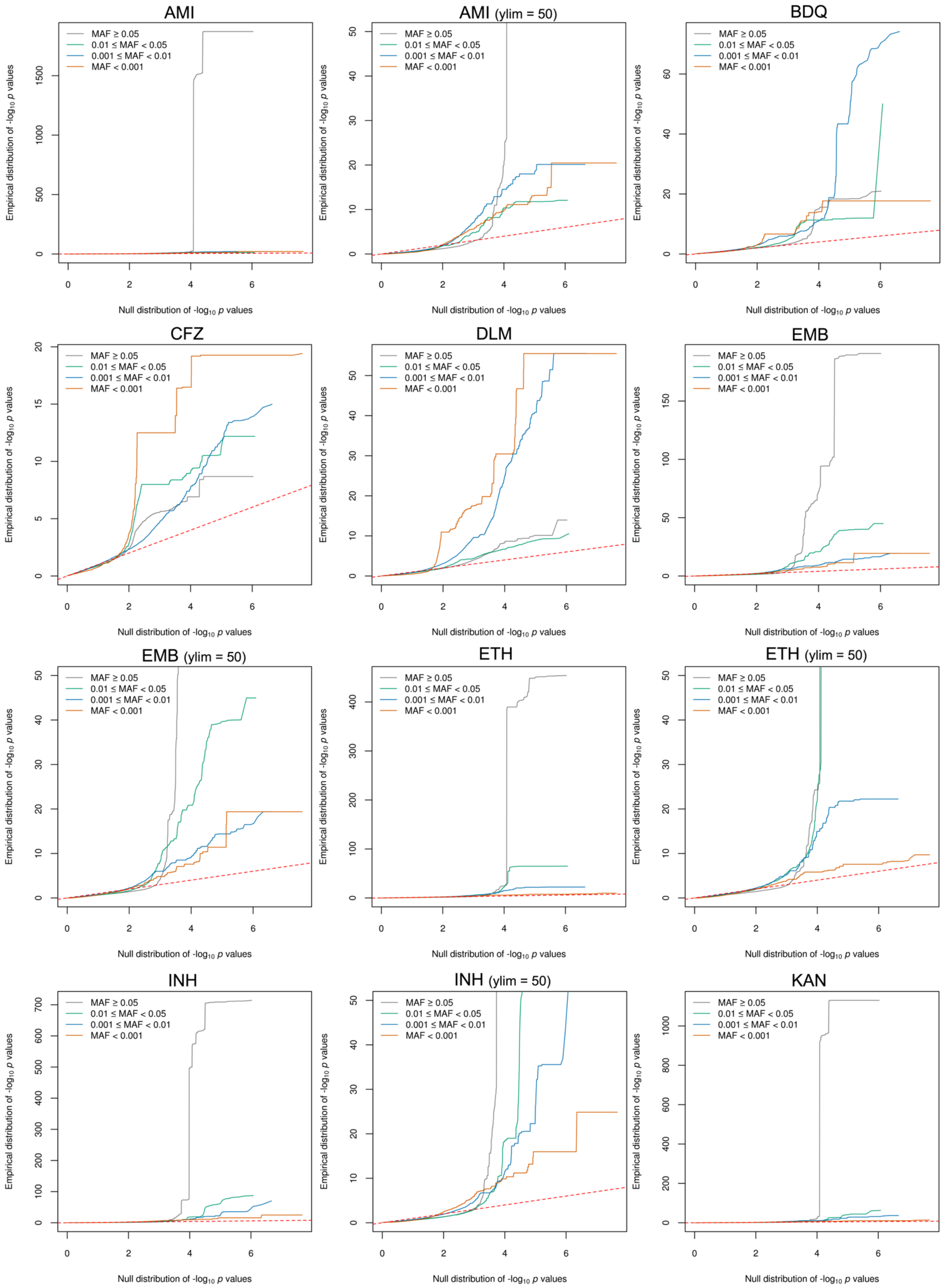


**Supplementary Figure 3 QQ plots for the oligopeptide analyses, part A.** Comparing the empirical distribution of p-values to the expected distribution under the null hypothesis for the drugs amikacin (AMI), bedaquiline (BDQ), clofazimine (CFZ), delamanid (DLM), ethambutol (EMB), ethionamide (ETH), isoniazid (INH), kanamycin (KAN). Oligopeptides in the orange (MAF < 0.1%) were not initially analysed, only used for signal interpretation.

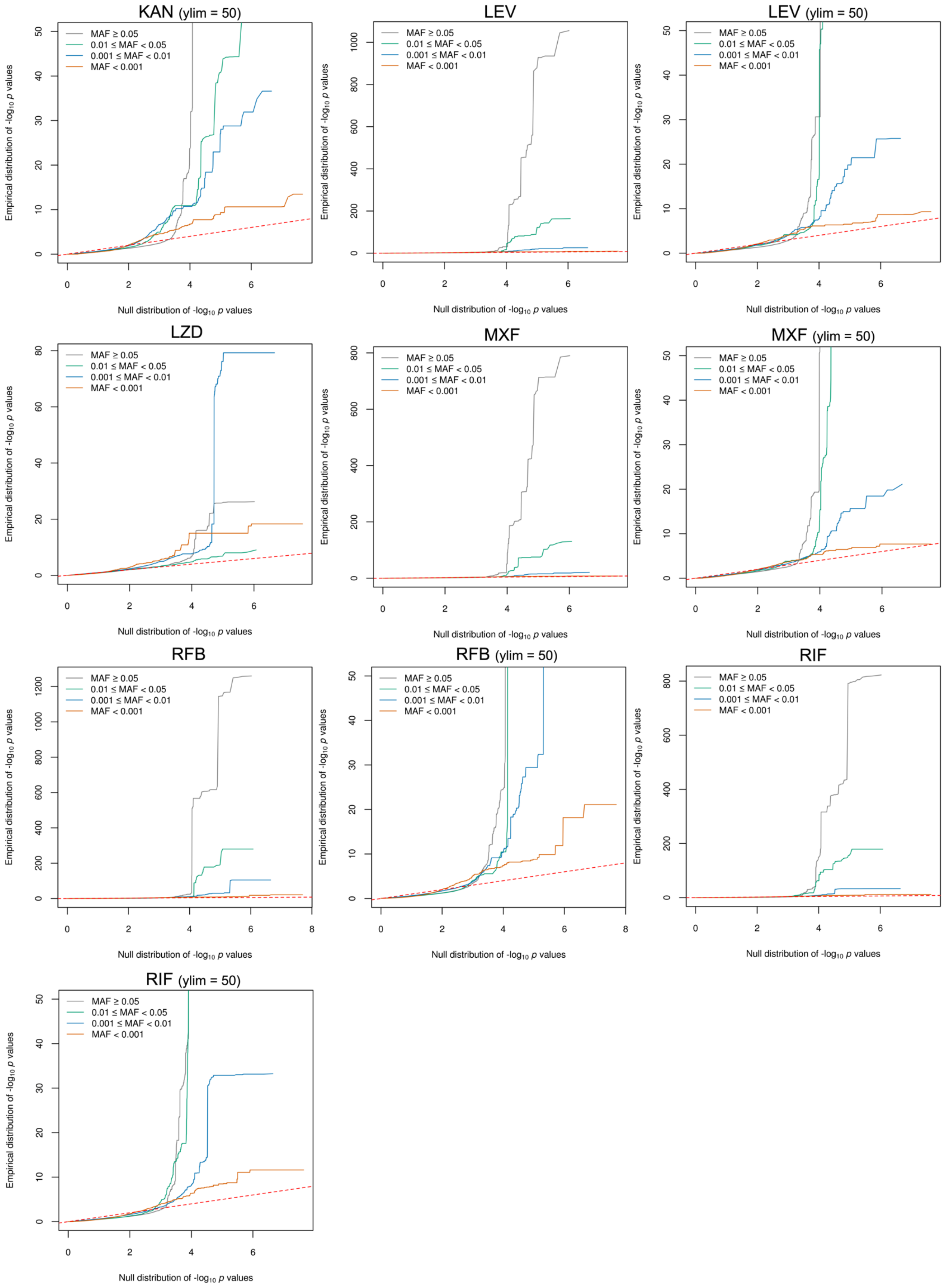


**Supplementary Figure 4 QQ plots for the oligopeptide analyses, part B.** Comparing the empirical distribution of p-values to the expected distribution under the null hypothesis for the kanamycin (KAN), levofloxacin (LEV), linezolid (LZD), moxifloxacin (MXF), rifabutin (RFB) and rifampicin (RIF). Oligopeptides in the orange (MAF < 0.1%) were not initially analysed, only used for signal interpretation.

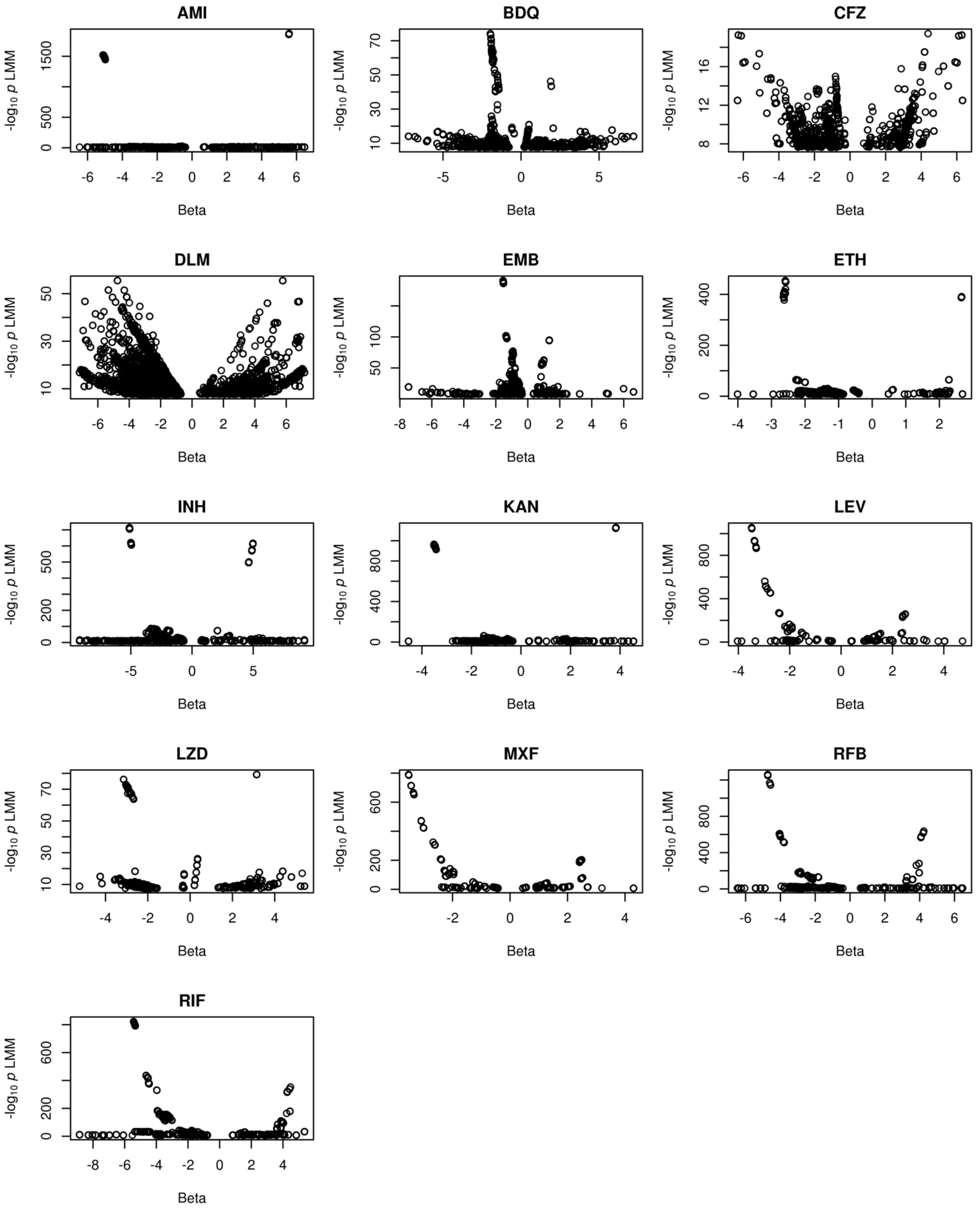


**Supplementary Figure 5** Effect size (beta) estimates and -log$_{10}$ *p*-values for all significant oligopeptide variants for each drug, amikacin (AMI), bedaquiline (BDQ), clofazimine (CFZ), delamanid (DLM), ethambutol (EMB), ethionamide (ETH), isoniazid (INH), kanamycin (KAN), levofloxacin (LEV), linezolid (LZD), moxifloxacin (MXF), rifabutin (RFB) and rifampicin (RIF). For many of the drugs, the most significant oligopeptides were associated with lower MIC.

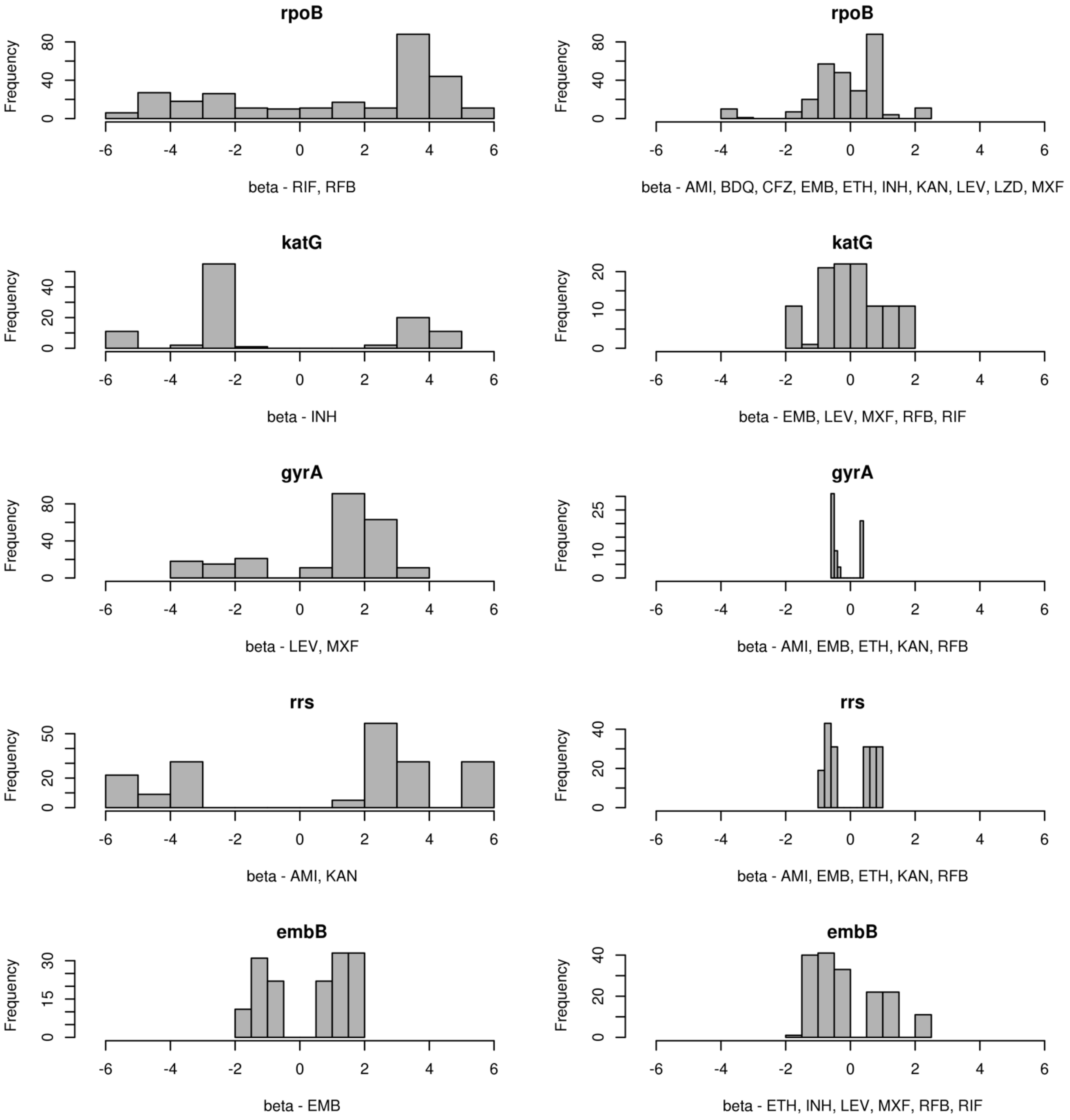


**Supplementary Figure 6** Significant oligopeptide (*rpoB, katG, gyrA, embB*) and oligonucleotide (*rrs*) effect size (beta) estimates for known resistance genes plus the flanking 33 amino acids (oligopeptides) or 100 bases (oligonucelotides). On the left the beta estimates are shown for all significant oligopeptides for the drugs the gene is causal for, on the right the beta estimates are shown for the same gene, but for the drugs they are artefactually associated to. For many drugs, the beta estimate is lower when the gene is significant due to artefactual cross resistance. Drug name abbreviations are as follows: amikacin (AMI), bedaquiline (BDQ), clofazimine (CFZ), delamanid (DLM), ethambutol (EMB), ethionamide (ETH), isoniazid (INH), kanamycin (KAN), levofloxacin (LEV), linezolid (LZD), moxifloxacin (MXF), rifabutin (RFB) and rifampicin (RIF).

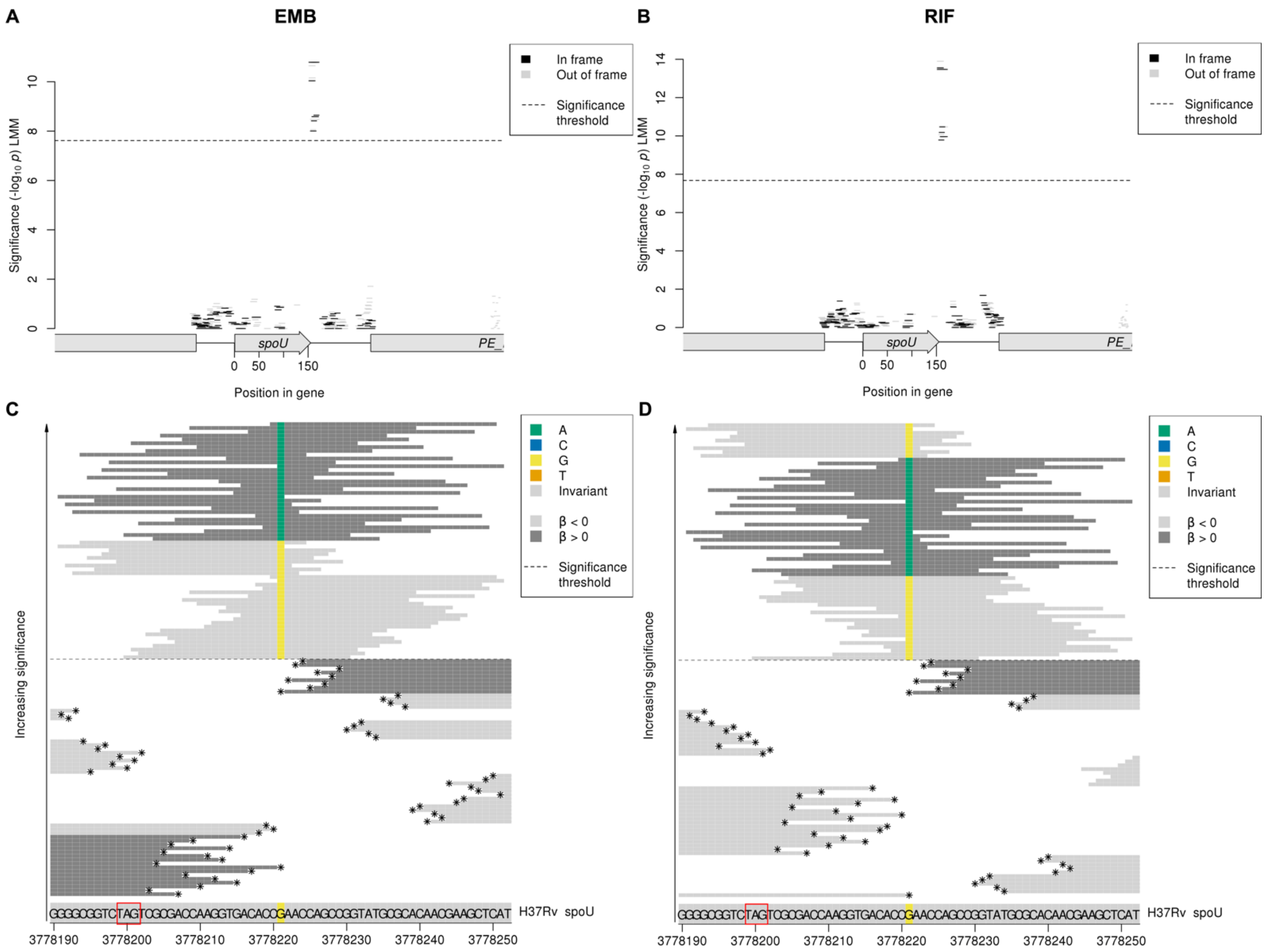


**Supplementary Figure 7** Variants in *spoU* associated with ethambutol (EMB) and rifampicin (RIF) MIC. Manhattan plots showing the oligopeptide association results for the *spoU* coding region **A** ethambutol and **B** rifampicin, and oligonucleotide alignment plots showing close ups of the significant region just downstream of *spoU* for **C** ethambutol and **D** rifampicin. The black dashed lines indicate the Bonferroni-corrected significance thresholds. In the Manhattan plots, oligopeptides are coloured by the reading frame that they align to, black for the correct reading frame for *spoU*. Oligopeptides assigned to the region but did not align using BLAST are shown in grey on the right hand side of the plots. In the oligonucleotide alignment plots, the H37Rv reference codons are shown at the bottom of the figure, grey for an invariant site, coloured at variant site positions. The oligonucleotides that aligned to the region are plotted from least significant at the bottom to most significant at the top. The background colour of the oligonucleotides represents the direction of the β estimate, light grey when β < 0 (associated with lower MIC), dark grey when β > 0 (associated with higher MIC). Oligonucleotides are coloured by their amino acid residue at all variant positions. Oligonucleotides below the MAF threshold and not included in the analysis, but visualised here for signal interpretation, are marked by *s. The *spoU* stop codon is highlighted in red in the alignment plots.

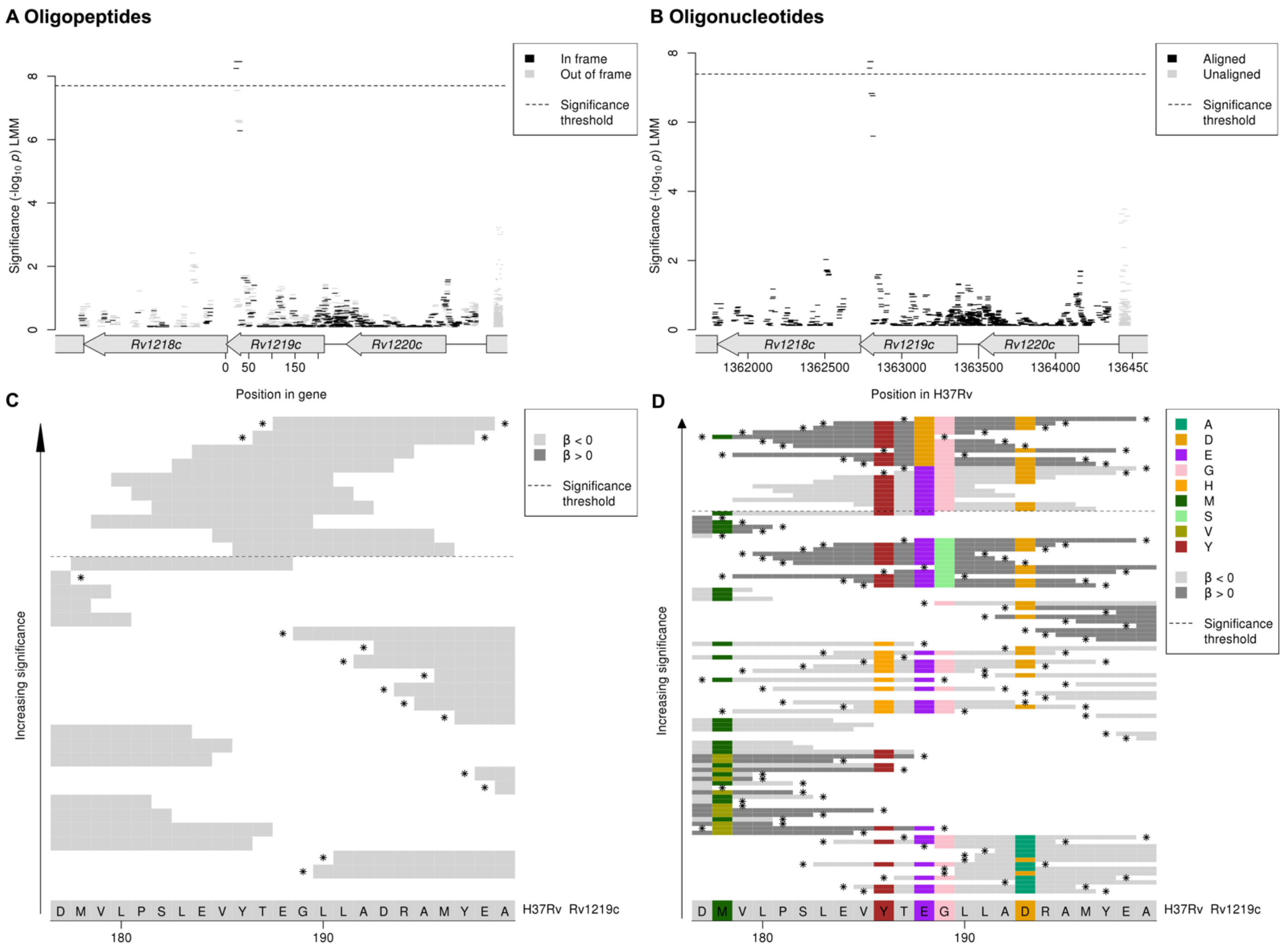


**Supplementary Figure 8** Variants in *Rv1219c* associated with isoniazid MIC. Manhattan plots showing the association results for the *Rv1219c* coding region for the **A** oligopeptides and **B** oligonucleotides, and oligopeptide alignment plots showing close ups of the significant region in *Rv1219c* for **C** oligopeptides present in five or more genomes in the full GWAS dataset and **D** oligopeptides present in at least one genome in the full GWAS dataset. The black dashed lines indicate the Bonferroni-corrected significance thresholds. In the Manhattan plots, oligopeptides are coloured by the reading frame that they align to, black for the correct reading frame for *Rv1219c*. Oligo-peptides and nucleotides assigned to the region but did not align using BLAST are shown in grey on the right hand side of the plots. In the oligopeptide alignment plots, the H37Rv reference codons are shown at the bottom of the figure, grey for an invariant site, coloured at variant site positions. The oligopeptides that aligned to the region are plotted from least significant at the bottom to most significant at the top. The background colour of the oligopeptides represents the direction of the β estimate, light grey when β < 0 (associated with lower MIC), dark grey when β > 0 (associated with higher MIC). Oligopeptides are coloured by their amino acid residue at all variant positions. Oligo-peptides and nucleotides below the MAF threshold and not included in the analysis, but visualised here for signal interpretation, are marked by *s.

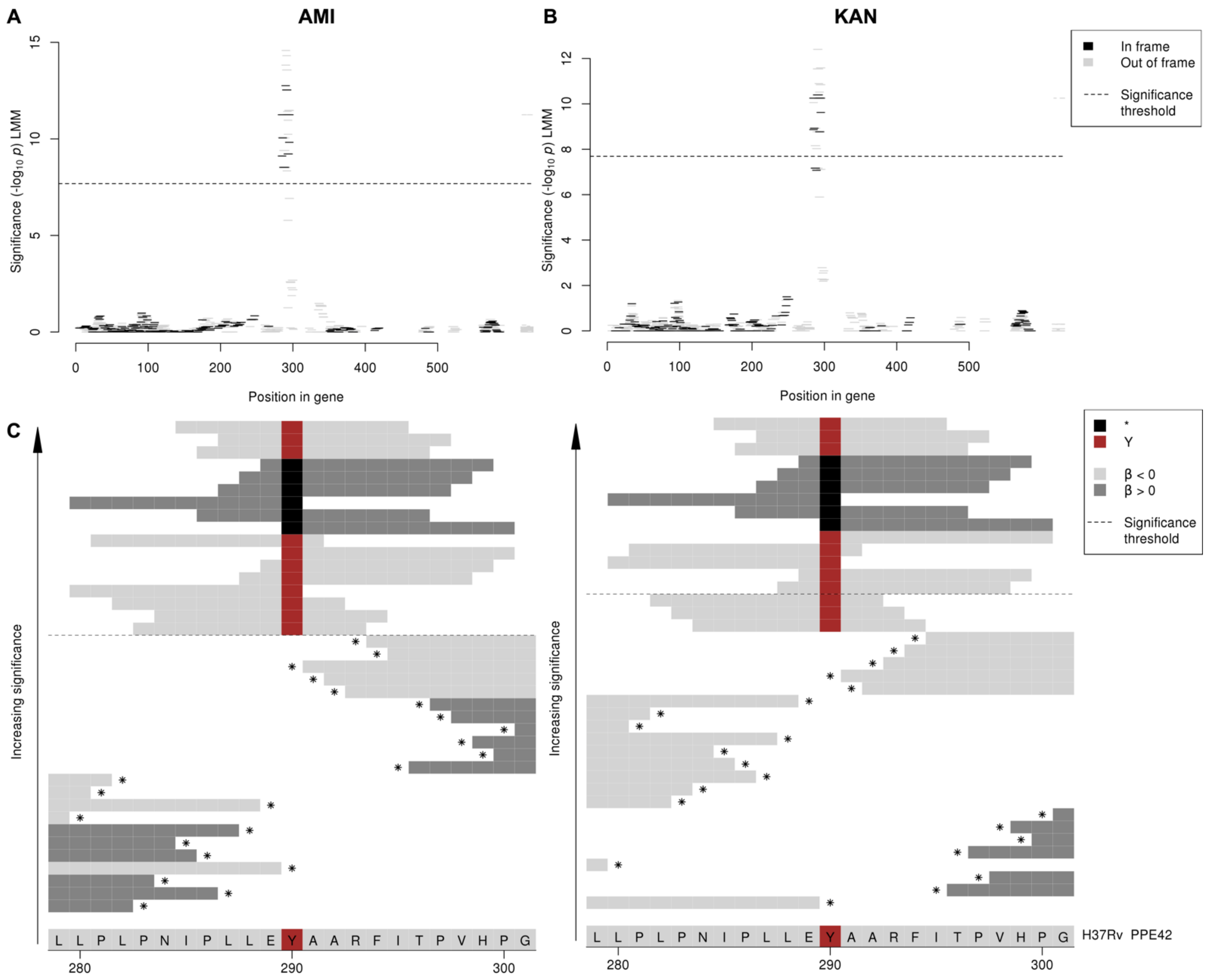


**Supplementary Figure 9** Variants in *PPE42* associated with amikacin (AMI) and kanamycin (KAN) MIC. Manhattan plots showing the oligopeptide association results for the *PPE42* coding region **A** amikacin and **B** kanamycin, and oligopeptide alignment plots showing close ups of the significant region in *PPE42* for **C** amikacin and **D** kanamycin. Black dashed lines indicate the Bonferroni-corrected significance thresholds. In the Manhattan plots, oligopeptides are coloured by the reading frame that they align to, black for the correct reading frame for *PPE42*. Oligopeptides assigned to the region but did not align using BLAST are shown in grey on the right hand side of the plots. In the oligopeptide alignment plots, the H37Rv reference codons are shown at the bottom of the figure, grey for an invariant site, coloured at variant site positions. The oligopeptides that aligned to the region are plotted from least significant at the bottom to most significant at the top. The background colour of the oligopeptides represents the direction of the β estimate, light grey when β < 0 (associated with lower MIC), dark grey when β > 0 (associated with higher MIC). Oligopeptides are coloured by their amino acid residue at all variant positions. Oligopeptides below the MAF threshold and not included in the analysis, but visualised here for signal interpretation, are marked by *s.

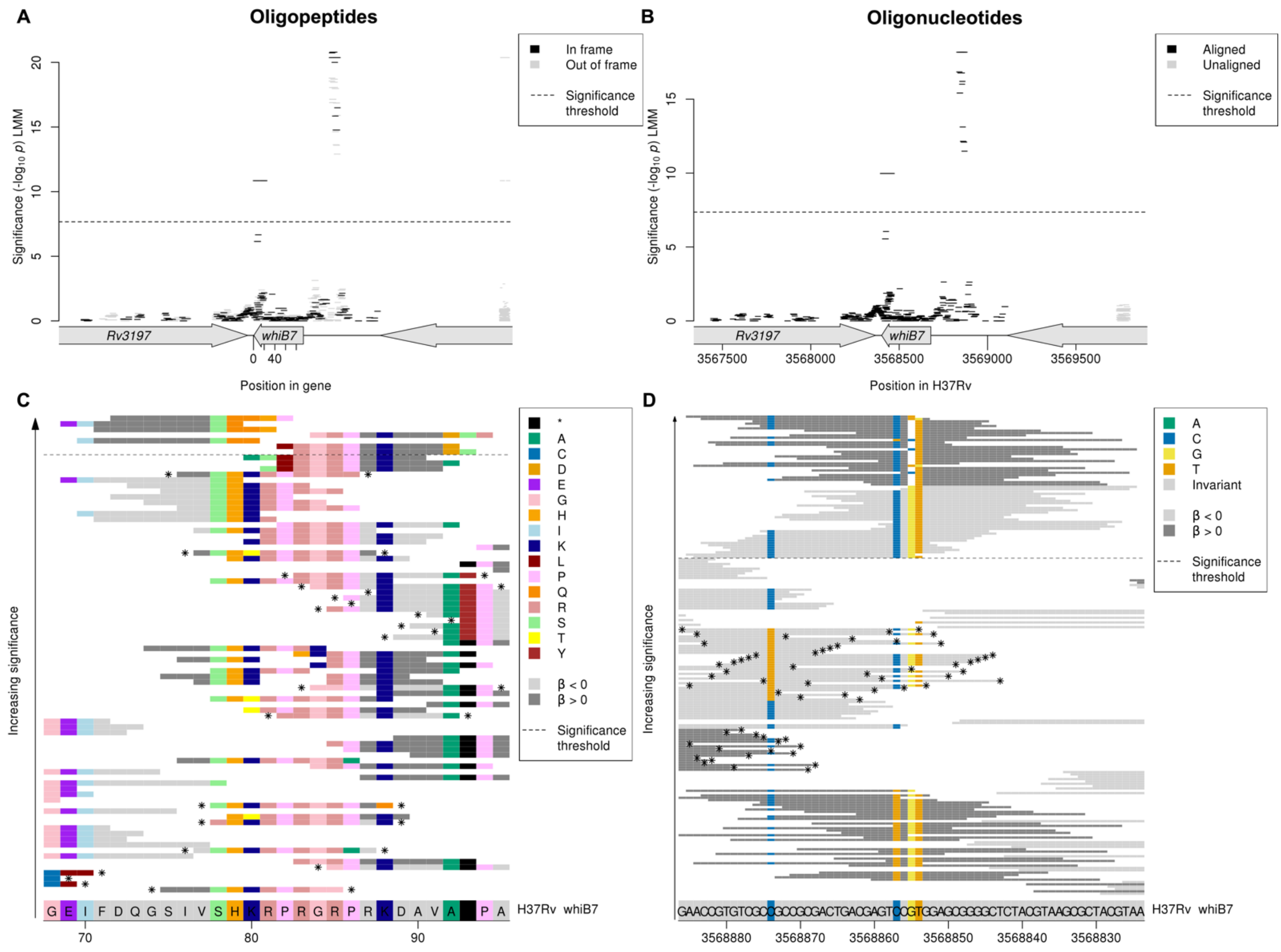


**Supplementary Figure 10** Variants in and upstream of *whiB7* associated with ethionamide MIC. Manhattan plots showing the association results for *whiB7* for the **A** oligopeptides and **B** oligonucleotides, and alignment plots showing close ups of significant regions in *whib7* for **C** oligopeptides in the C-terminal end of the coding region and **D** oligonucleotides in the upstream intergenic region. The black dashed lines indicate the Bonferroni-corrected significance thresholds. In the Manhattan plots, oligopeptides are coloured by the reading frame that they align to, black for the correct reading frame for *whiB7*. Oligo-peptides and nucleotides assigned to the region but did not align using BLAST are shown in grey on the right hand side of the plots. In the alignment plots, the H37Rv reference codons are shown at the bottom of the figure, grey for an invariant site, coloured at variant site positions. The oligo-peptides and nucleotides that aligned to the region are plotted from least significant at the bottom to most significant at the top. The background colour of the oligo-peptides and nucleotides represents the direction of the β estimate, light grey when $\beta < 0$ (associated with lower MIC), dark grey when $\beta > 0$ (associated with higher MIC). Oligo-peptides and nucleotides are coloured by their allele at all variant positions. Oligo-peptides and nucleotides below the MAF threshold and not included in the analysis, but visualised here for signal interpretation, are marked by *s.

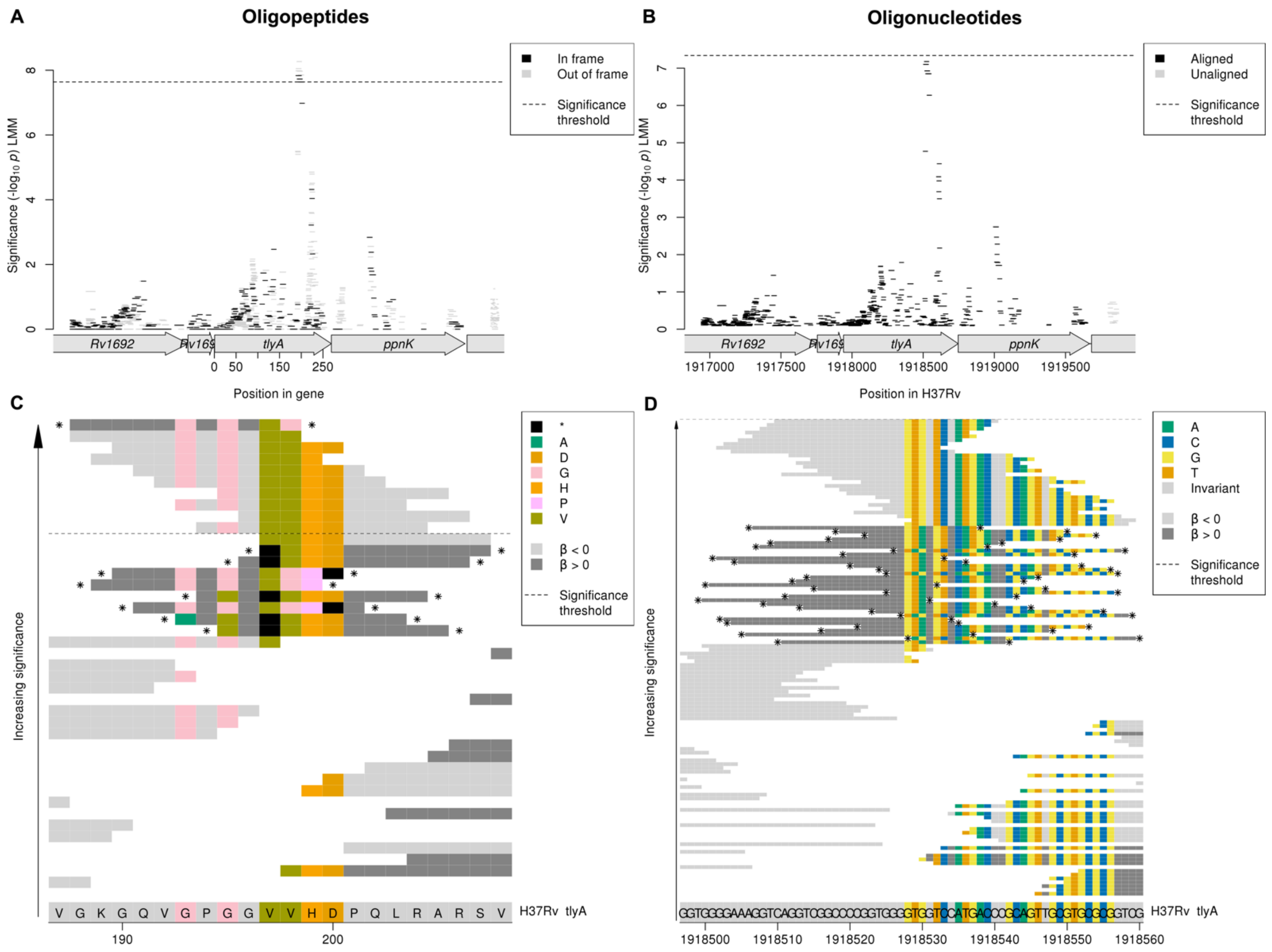


**Supplementary Figure 11** Variants in *tlyA* associated with levofloxacin MIC. Manhattan plots showing the association results for the *tlyA* coding region for the **A** oligopeptides and **B** oligonucleotides, and alignment plots showing close ups of the significant region in *tlyA* for the **C** oligopeptides **D** oligonucleotides. The black dashed lines indicate the Bonferroni-corrected significance thresholds, no oligonucleotides were significant in this region. In the Manhattan plots, oligopeptides are coloured by the reading frame that they align to, black for the correct reading frame for *tlyA*. Oligo-peptides and nucleotides assigned to the region but did not align using BLAST are shown in grey on the right hand side of the plots. In the alignment plots, the H37Rv reference alleles are shown at the bottom of the figure, grey for an invariant site, coloured at variant site positions. The oligo-peptides and nucleotides that aligned to the region are plotted from least significant at the bottom to most significant at the top. The background colour of the oligo-peptides and nucleotides represents the direction of the β estimate, light grey when β < 0 (associated with lower MIC), dark grey when β> 0 (associated with higher MIC). Oligo-peptides and nucleotides are coloured by their allele at all variant positions. Oligo-peptides and nucleotides below the MAF threshold and not included in the analysis, but visualised here for signal interpretation, are marked by *s.

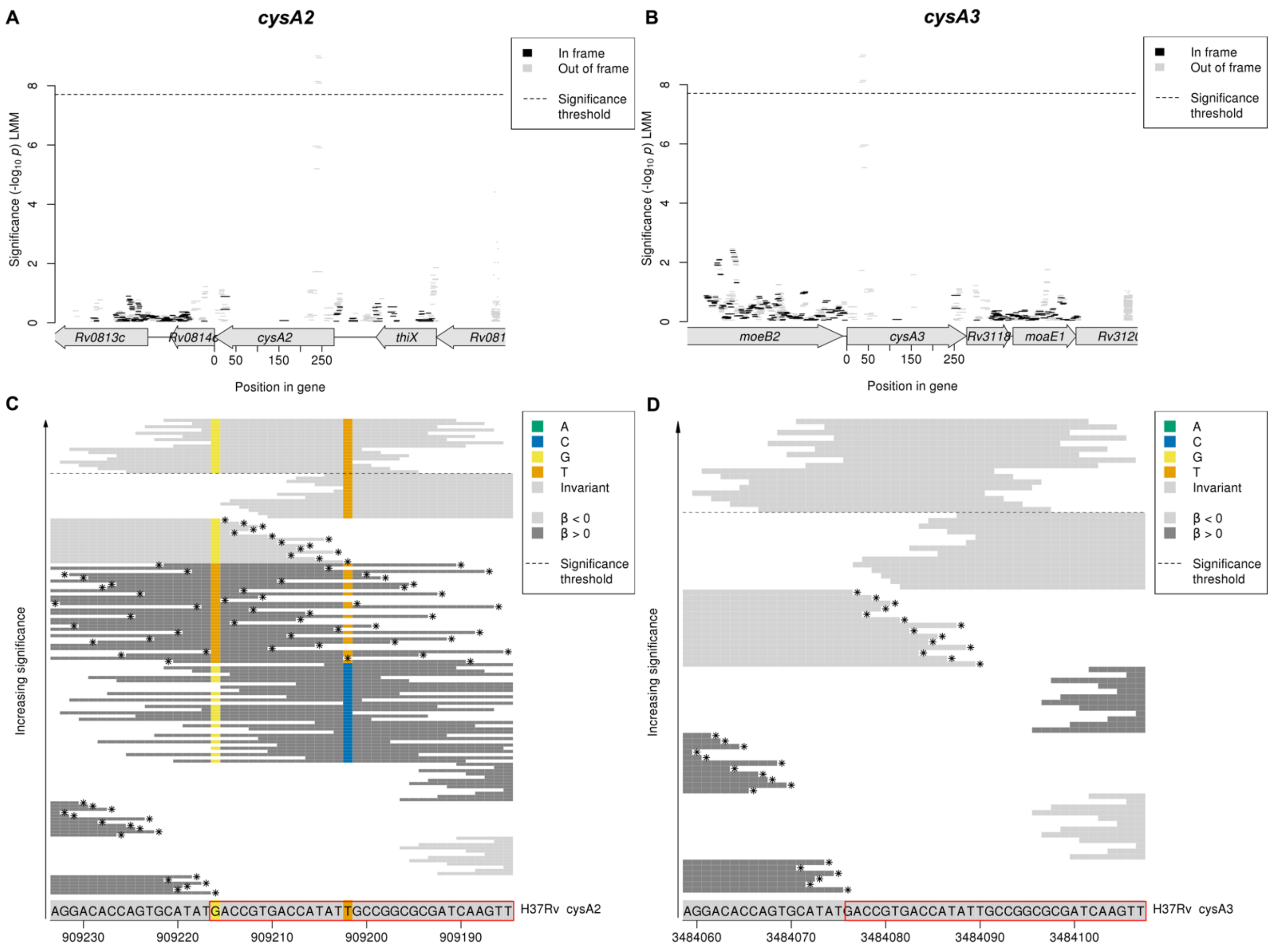


**Supplementary Figure 12** in *cysA2* and *cysA3* associated with rifabutin MIC. Manhattan plots showing the association results for the coding region for **A** *cysA2* and **B** cysA3, and oligonucleotide alignment plots showing close ups of the significant region for **C** *cysA2* **D** *cysA3*. The black dashed lines indicate the Bonferroni-corrected significance thresholds. The significant oligonucleotides that align to *cysA2* and *cysA3* are the same. In the Manhattan plots, oligopeptides are coloured by the reading frame that they align to, black for the correct reading frame for *cysA2* or *cysA3*. Oligopeptides assigned to the region but did not align using BLAST are shown in grey on the right hand side of the plot. In the oligonucleotide alignment plots, the H37Rv reference alleles are shown at the bottom of the figure, grey for an invariant site, coloured at variant site positions. The oligonucleotides that aligned to the region are plotted from least significant at the bottom to most significant at the top. The background colour of the oligonucleotides represents the direction of the β estimate, light grey when β < 0 (associated with lower MIC), dark grey when β> 0 (associated with higher MIC). Oligonucleotides are coloured by their allele at all variant positions. Oligonucleotides below the MAF threshold and not included in the analysis, but visualised here for signal interpretation, are marked by *s. The region that encodes the rhodanese characteristic signature in the N-terminal region is highlighted in red in the alignment plots.

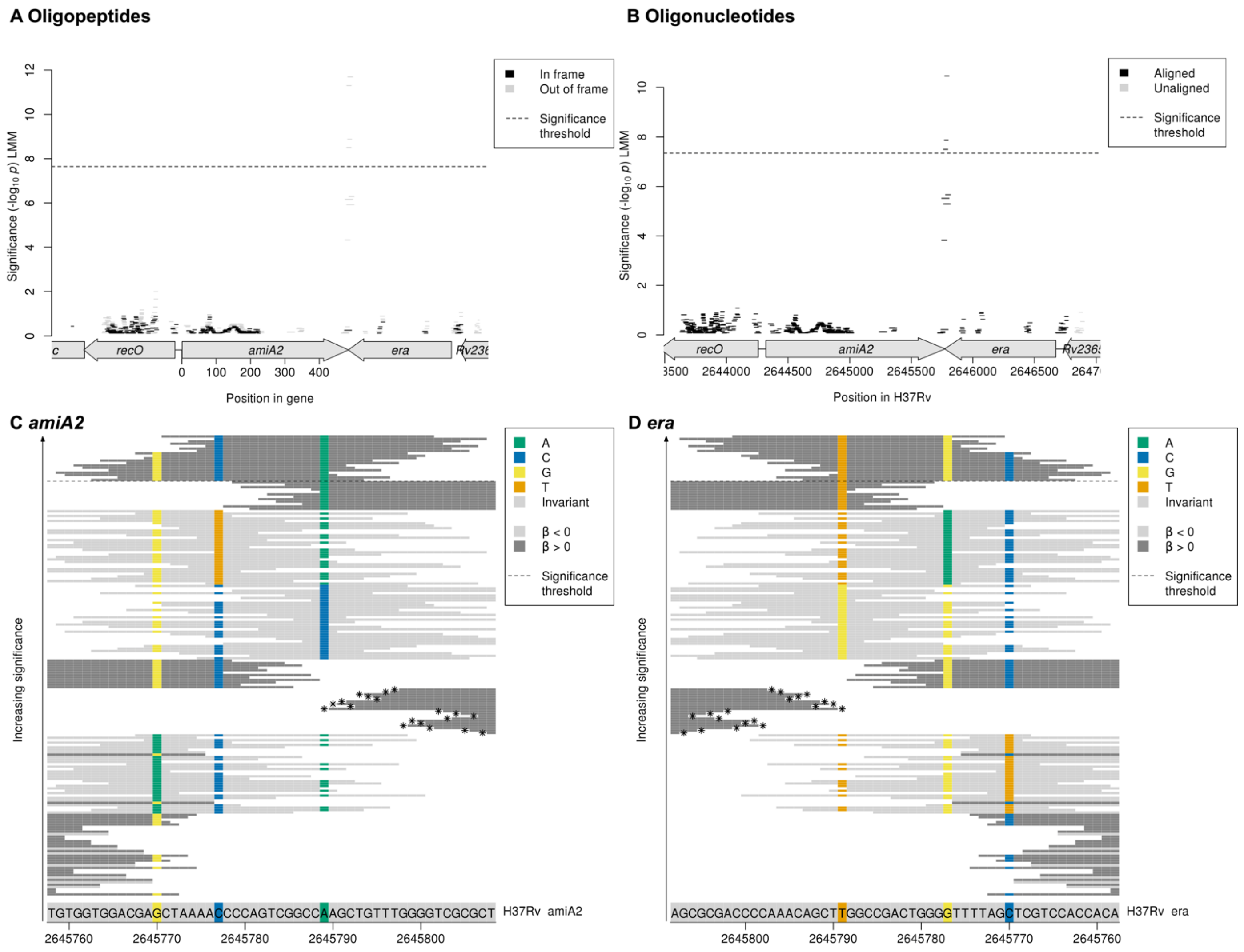


**Supplementary Figure 13** Variants in *amiA2* and *era* associated with bedaquiline MIC. Manhattan plots showing the association results for the *amiA2/era* coding region for the **A** oligopeptides and **B** oligonucleotides, and oligonucleotide alignment plots showing close ups of the significant region in *amiA2/era* in the correct reading frame for **C** *amiA2* **D** *era*. The black dashed lines indicate the Bonferroni-corrected significance thresholds. In the Manhattan plots, oligopeptides are coloured by the reading frame that they align to, black for the correct reading frame for *amiA2*. Oligo-peptides and nucleotides assigned to the region but did not align using BLAST are shown in grey on the right hand side of the plots. In the oligonucleotide alignment plots, the H37Rv reference alleles are shown at the bottom of the figure, grey for an invariant site, coloured at variant site positions. The oligonucleotides that aligned to the region are plotted from least significant at the bottom to most significant at the top. The background colour of the oligonucleotides represents the direction of the $\beta$ estimate, light grey when $\beta < 0$ (associated with lower MIC), dark grey when $\beta > 0$ (associated with higher MIC). Oligonucleotides are coloured by their allele at all variant positions. Oligonucleotides below the MAF threshold and not included in the analysis, but visualised here for signal interpretation, are marked by *s.

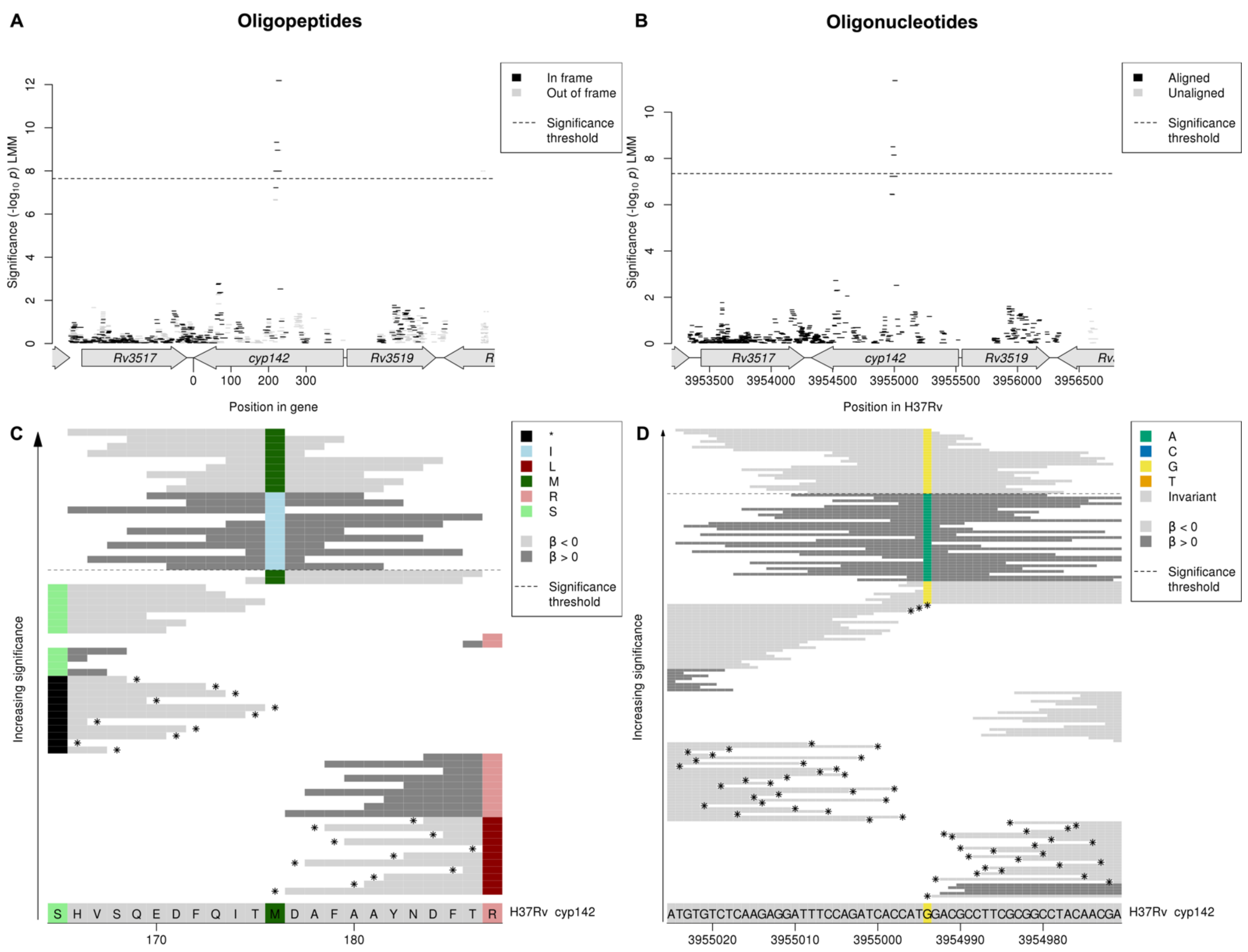


**Supplementary Figure 14** Variants in *cyp142* associated with clofazimine MIC. Manhattan plots showing the association results for the *cyp142* coding region for the **A** oligopeptides and **B** oligonucleotides, and alignment plots showing close ups of the significant region in *cyp142* for the **C** oligopeptides **D** oligonucleotides. The black dashed lines indicate the Bonferroni-corrected significance thresholds. In the Manhattan plots, oligopeptides are coloured by the reading frame that they align to, red for the correct reading frame for *cyp142*. Oligo-peptides and nucleotides assigned to the region but did not align using BLAST are shown in grey on the right hand side of the plots. In the alignment plots, the H37Rv reference alleles are shown at the bottom of the figure, grey for an invariant site, coloured at variant site positions. The oligo-peptides and nucleotides that aligned to the region are plotted from least significant at the bottom to most significant at the top. The background colour of the oligo-peptides and nucleotides represents the direction of the β estimate, light grey when $\beta < 0$ (associated with lower MIC), dark grey when $\beta > 0$ (associated with higher MIC). Oligo-peptides and nucleotides are coloured by their allele at all variant positions. Oligo-peptides and nucleotides below the MAF threshold and not included in the analysis, but visualised here for signal interpretation, are marked by *s.

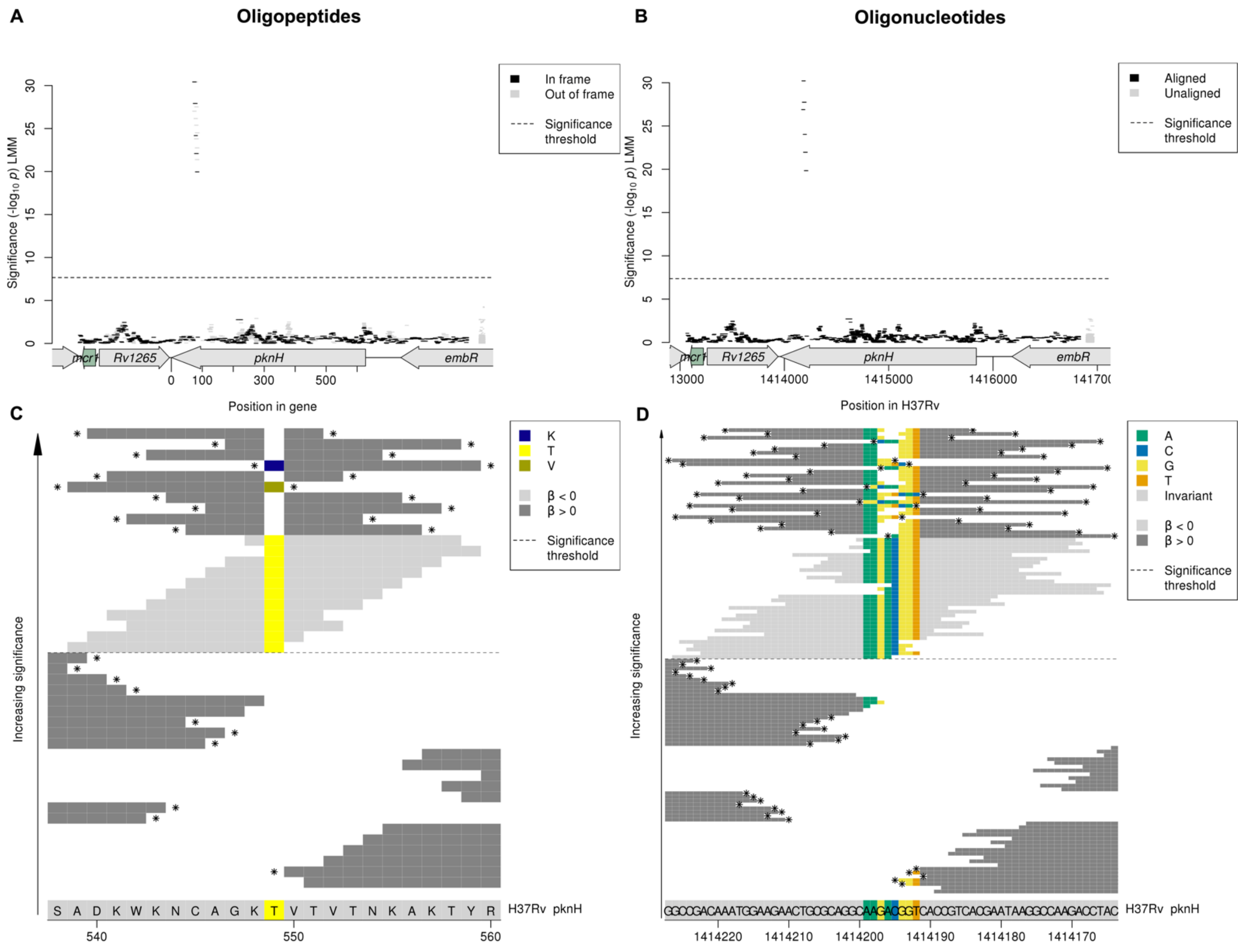


**Supplementary Figure 15** Variants in *pknH* associated with delamanid MIC. Manhattan plots showing the association results for the *pknH* coding region for the **A** oligopeptides and **B** oligonucleotides, and alignment plots showing close ups of the significant region in *pknH* for the **C** oligopeptides **D** oligonucleotides. The black dashed lines indicate the Bonferroni-corrected significance thresholds. In the Manhattan plots, oligopeptides are coloured by the reading frame that they align to, black for the correct reading frame for *pknH*. Oligo-peptides and nucleotides assigned to the region but did not align using BLAST are shown in grey on the right hand side of the plots. In the alignment plots, the H37Rv reference alleles are shown at the bottom of the figure, grey for an invariant site, coloured at variant site positions. The oligo-peptides and nucleotides that aligned to the region are plotted from least significant at the bottom to most significant at the top. The background colour of the oligo-peptides and nucleotides represents the direction of the β estimate, light grey when β < 0 (associated with lower MIC), dark grey when β> 0 (associated with higher MIC). Oligo-peptides and nucleotides are coloured by their allele at all variant positions. Oligo-peptides and nucleotides below the MAF threshold and not included in the analysis, but visualised here for signal interpretation, are marked by *s.

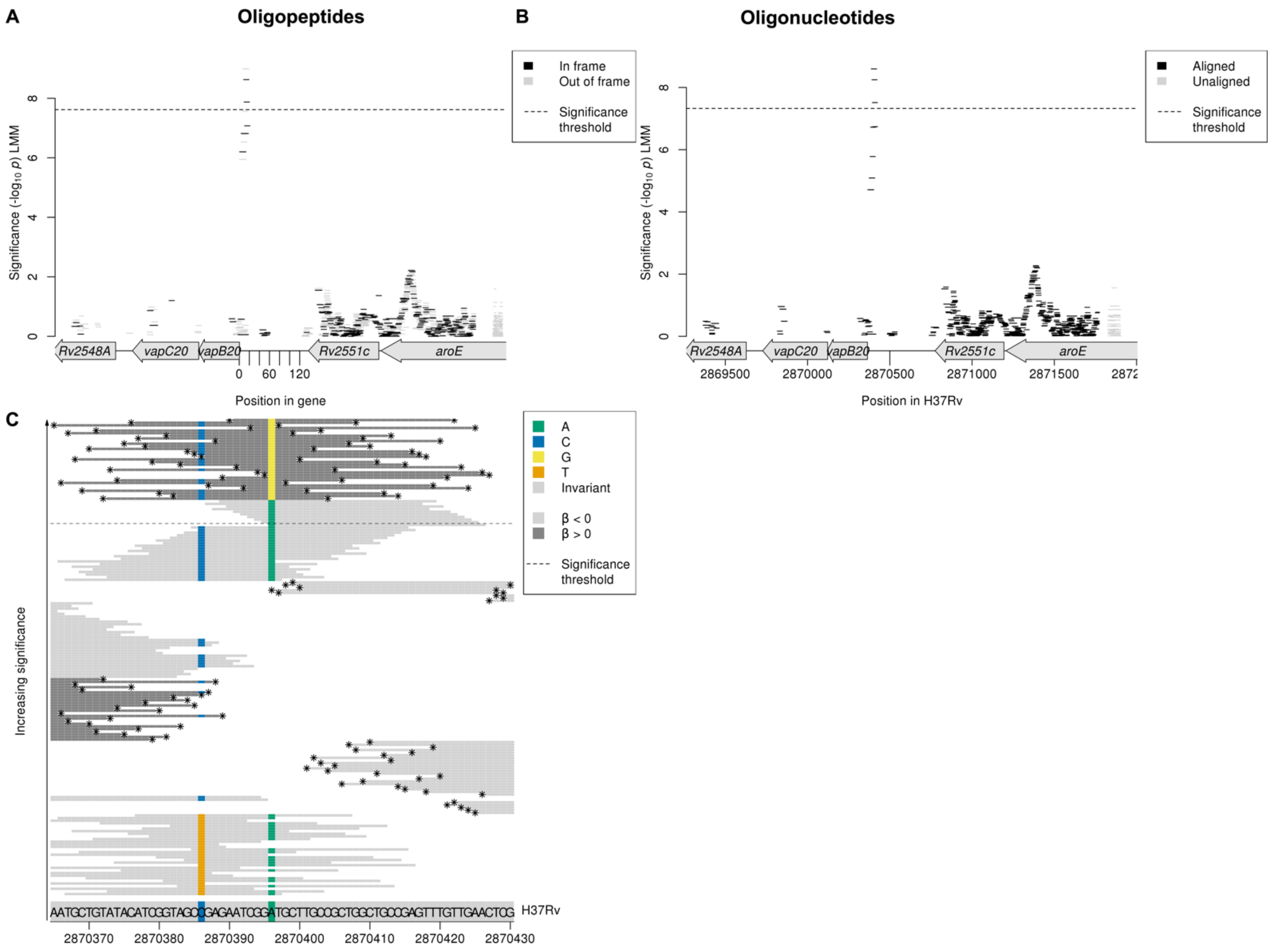


**Supplementary Figure 16** Variants in *vapB20* associated with linezolid MIC. Manhattan plots showing the association results for *vapB20* for the **A** oligopeptides and **B** oligonucleotides, and **C** oligonucleotide alignment plot showing a close up of the significant region just upstream of *vapB20*. The black dashed lines indicates the Bonferroni-corrected significance thresholds. In the Manhattan plots, oligopeptides are coloured by the reading frame that they align to, black for the correct reading frame for *amiA2*. Oligo-peptides and nucleotides assigned to the region but did not align using BLAST are shown in grey on the right hand side of the plots. In the oligonucleotide alignment plot, the H37Rv reference alleles are shown at the bottom of the figure, grey for an invariant site, coloured at variant site positions. The oligonucleotides that aligned to the region are plotted from least significant at the bottom to most significant at the top. The background colour of the oligonucleotides represents the direction of the β estimate, light grey when β < 0 (associated with lower MIC), dark grey when β> 0 (associated with higher MIC). Oligonucleotides are coloured by their allele at all variant positions. Oligo-peptides and nucleotides below the MAF threshold and not included in the analysis, but visualised here for signal interpretation, are marked by *s.

| | Heritability estimates and 95% confidence intervals | | | |
|---|---|---|---|---|
| | **Oligopeptides MIC** | **Oligopeptides binary** | **Oligonucleotides MIC** | **Oligonucleotides binary** |
| **Delamanid** | 36.0 (28.94-43.05) | 0 (-0.49-0.49) | 39.86 (32.69-47.02) | 0 (-0.45-0.45) |
| **Clofazimine** | 36.83 (31.62-42.04) | 10.72 (7.61-13.83) | 37.73 (32.41-43.05) | 10.64 (7.49-13.8) |
| **Linezolid** | 41 (34.98-47.01) | 3.88 (1.7-6.06) | 42.51 (36.44-48.58) | 4.19 (1.87-6.51) |
| **Bedaquiline** | 52.88 (48.26-57.5) | 25.76 (19.04-32.47) | 54.29 (49.71-58.86) | 27.75 (20.99-34.52) |
| **Moxifloxacin** | 83.74 (81.6-85.88) | 81.6 (79.2-84.01) | 86.88 (85.21-88.56) | 85.67 (83.85-87.5) |
| **Levofloxacin** | 85 (83.18-86.81) | 88.61 (87.27-89.95) | 87.75 (86.34-89.17) | 90.69 (89.66-91.73) |
| **Kanamycin** | 85.3 (83.68-86.92) | 80.5 (78.53-82.47) | 87.19 (85.85-88.53) | 82.68 (81.01-84.35) |
| **Ethambutol** | 85.97 (84.57-87.36) | 80.74 (78.88-82.61) | 85.78 (84.39-87.17) | 80.56 (78.71-82.41) |
| **Ethionamide** | 86.74 (85.41-88.08) | 82.21 (80.49-83.94) | 86.86 (85.57-88.15) | 82.49 (80.85-84.13) |
| **Amikacin** | 91.18 (90.21-92.15) | 89.38 (88.26-90.49) | 92.05 (91.23-92.86) | 90.47 (89.54-91.41) |
| **Rifampicin** | 94.6 (94.03-95.17) | 94.31 (93.71-94.91) | 94.8 (94.27-95.34) | 94.57 (94.01-95.14) |
| **Isoniazid** | 94.87 (94.37-95.38) | 94.74 (94.2-95.29) | 94.97 (94.47-95.48) | 94.91 (94.37-95.45) |
| **Rifabutin** | 95.6 (95.13-96.07) | 92.75 (92.09-93.41) | 95.65 (95.19-96.1) | 92.86 (92.22-93.5) |

**Supplementary Table 1** Oligopeptide and oligonucleotide sample heritability estimates for binary resistant vs. sensitive phenotypes compared to semi-quantitiative MIC phenotypes. Sample heritability estimates and 95% confidence intervals are shown for the 13 drugs.
