## Supplementary Text for "Genome-wide association studies of global *Mycobacterium tuberculosis* resistance to thirteen antimicrobials in 10,228 genomes"

Oligopeptide (protein kmer) and oligonucleotide (nucleotide kmer) variant interpretation

*Rv2180c* ... 90
*gap* ... 91
*lprF:Rv1371* [R] ... 92
*Rv0914c* ... 92
*Rv1200* ... 94
*fadE10* ... 95
*dinP* ... 95
*mmpL8* ... 96
*cut1* ... 97
*PPE39* ... 98
*Rv3430a:gadB* ... 99
*Rv1429* ... 100
*Rv3847* ... 101
*pknH* ... 102
*plsC* ... 104
*agpS* ... 105
*Rv3263* ... 105
Ethambutol (EMB) ... 107
*embB* ... 107
*rpoB* ... 111
*katG* ... 113
*embA* ... 113
*pncA* ... 115
*gyrA* ... 118
*rpsL* ... 119
*Rv1565c* ... 119
*Rv2478c:Rv2481c* [R] ... 120
*Rv1752* ... 121
*Rv3183:Rv3188* [R] ... 123
*dxs2:Rv3382c* [R] ... 124
*rpsA/coaE* ... 125
*ctpI* ... 127
*guaA* ... 128
*moaC3:Rv3327* [R] ... 129
*lprF:Rv1371* [R] ... 129
*fabG1* ... 131
*spoU* ... 132
*glpK* ... 133

Ethionamide (ETH) .... 135
*fabG1* .... 135
*ethA* .... 137
*rpoB* .... 144
*gyrA* .... 145
*inhA* .... 146
*whiB7* .... 148
*PPE3* .... 150
*mpt53* .... 151
*embB* .... 152
*eccA1* .... 153
*embA* .... 154
*Rv0565c* .... 154
*fadB4* .... 156
*plsC* .... 157
*Rv0920c* .... 158
*Rv3698* .... 159
*rrs* .... 160
*pncA* .... 161
*PPE56* .... 162
*Rv2019* .... 163
*lprF:Rv1371* [R] .... 164
Isoniazid (INH) .... 165
*katG* .... 165
*proA:ahpC* .... 170
*fabG1* .... 172
*rpoB* .... 174
*inhA* .... 175
*embB* .... 177
*Rv1139c:Rv1140* .... 179
*Rv1158c* .... 180
*rpsL* .... 182
*Rv1219c* .... 183
*ftsK/Rv2749* .... 185
*gid* .... 187
Kanamycin (KAN) .... 188
*rrs* .... 188
*eis* .... 189

*gyrA* ... 191
*rpoB* ... 192
*ethA* ... 193
*fabG1* ... 195
*Rv1830* ... 196
*ptbB* ... 196
*PPE42* ... 197
*echA8* ... 198
*lprF:Rv1371* [R] ... 199
*Rv2348c:plcC* ... 200
*narU* ... 201
*pgi* ... 202
*mmaA4* ... 203
*pncA* ... 204
*viuB* ... 206
*lprC* ... 207
*murA* ... 208
*Rv1393c* ... 209
*Rv0579* ... 210
*glnE* ... 211
*rne* ... 213
*Rv1362c* ... 214
*Rv0208c:Rv0209* ... 215
Levofloxacin (LEV) ... 217
*gyrA* ... 217
*rrs* ... 219
*gyrB* ... 220
*embB* ... 221
*rpoB* ... 222
*vapC36* ... 223
*mce2F* ... 224
*fabG1* ... 225
*katG* ... 227
*folC* ... 228
*tlyA* ... 228
*ethA* ... 230
*Rv0228* ... 231
Linezolid (LZD) ... 232

*rplC* ... 232
*rpoB* ... 233
*emrB* ... 234
*Rv3552* ... 235
*add* ... 237
*vapC33* ... 238
*ppgK* ... 239
*pncB1:Rv1331* ... 241
*lprA* ... 242
*pafA* ... 243
*PE_PGRS6* ... 244
*vapB20* ... 245
*Rv0061c* ... 246
*PE_PGRS4* ... 248
*Rv1049* ... 249
*lprF:Rv1371* [R] ... 250
*Rv3183:Rv3188* [R] ... 250
*dxs2:Rv3382c* [R] ... 251
*Rv0556* ... 252
*Rv0514* ... 253
Moxifloxacin (MXF) ... 255
*gyrA* ... 255
*rrs* ... 257
*rpoB* ... 258
*gyrB* ... 259
*embB* ... 260
*katG* ... 261
Rifabutin (RFB) ... 262
*rpoB* ... 262
*embB* ... 265
*katG* ... 268
*rpoC* ... 269
*Rv0810c* ... 269
*Rv2478c:Rv2481c* [R] ... 270
*Rv2647:Rv2650c* [R] ... 271
*rplP* ... 272
*Rv2797c* ... 273
*cpsY* ... 274

### Amikacin (AMI)

#### *rrs*

Capturing the rRNA *rrs* between the two coding genes *murA* and *ogt*.

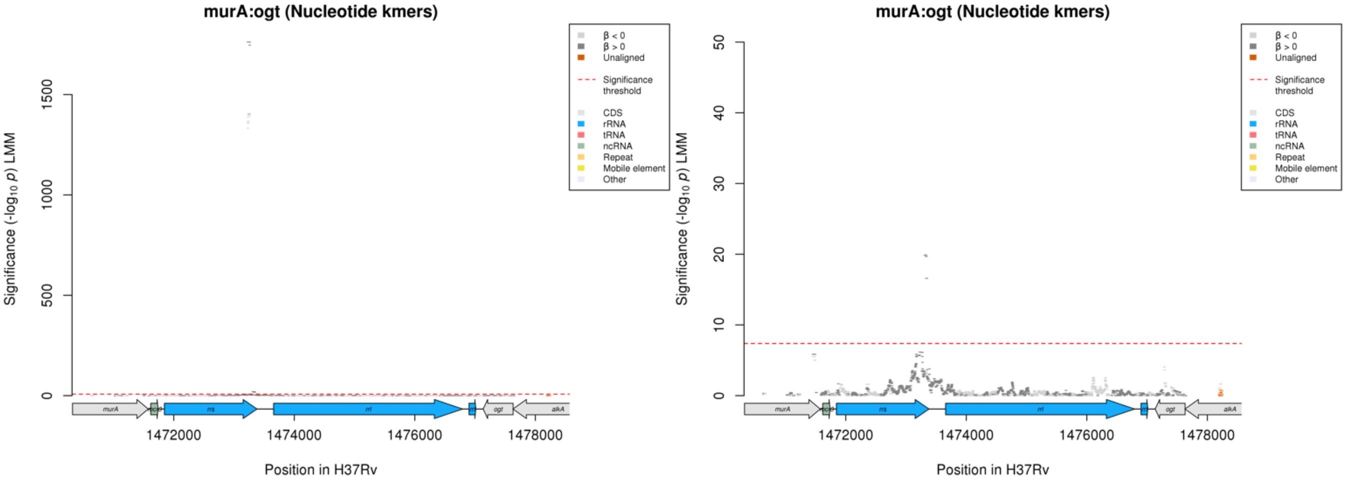

Nucleotide kmers:

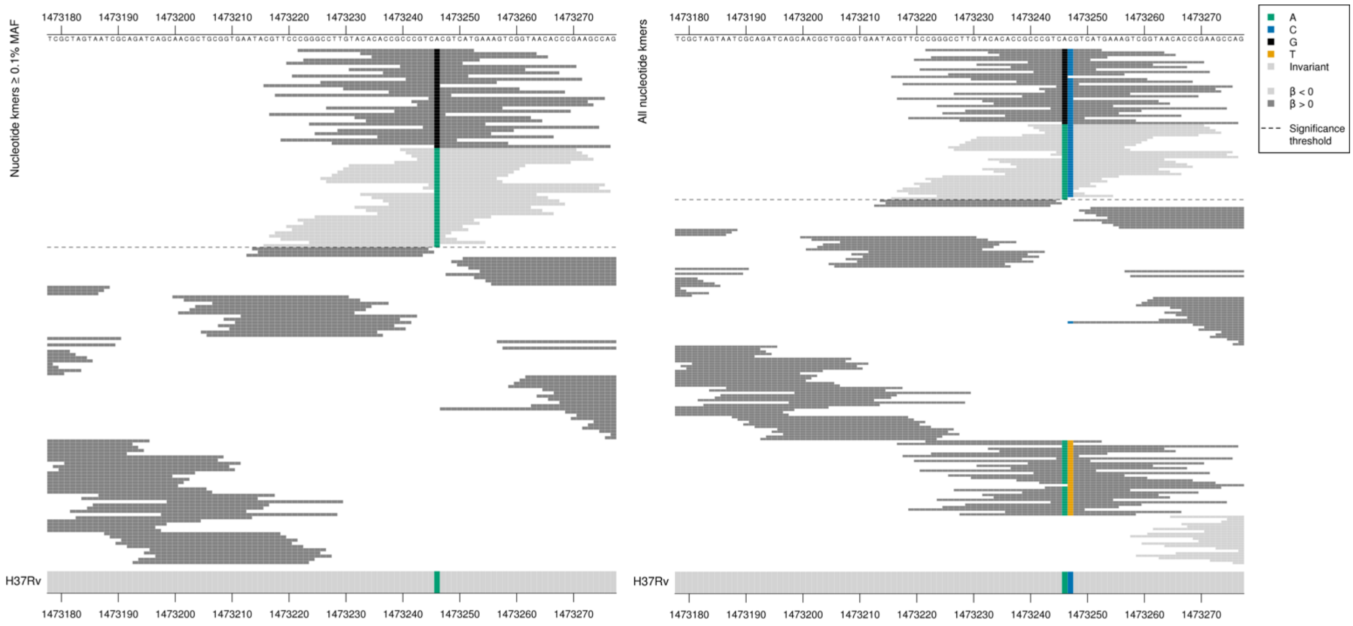

*rrs* = 16S ribosomal RNA

| Kmer | $-\log_{10} p$ | β | MAC | MAF | Ps | Variants |
|---|---|---|---|---|---|---|
| CGGGCCTTGTACACACCGCCCGTCGCGTCAT | 1761.62 | 5.57 | 507 | 6.01 | 1473222 | 1473246/1401G, 1473247/1402C |
| CGTTCCCGGGCCTTGTACACACCGCCCGTCG | 1761.62 | 5.57 | 507 | 6.01 | 1473216 | 1473246/1401G |
| CCCGTCACGTCATGAAAGTCGGTAACACCCG | 1405.86 | -5.06 | 554 | 6.57 | 1473240 | 1473246/1401A, 1473247/1402C |
| CGTTCCCGGGCCTTGTACACACCGCCCGTCA | 1332.59 | -4.95 | 558 | 6.62 | 1473216 | 1473246/1401A |

Amikacin (AMI)

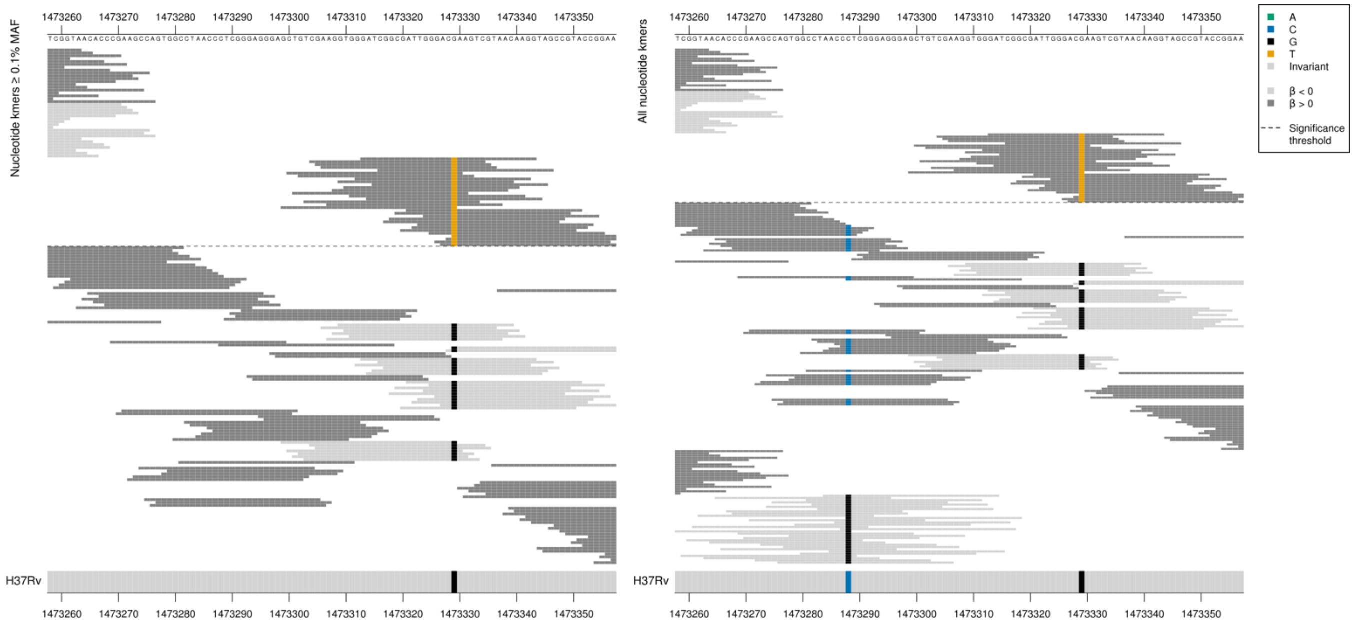

*rrs* ends position 1473382.

| Kmer | $-\log_{10} p$ | β | MAC | MAF | Ps | Variants |
|---|---|---|---|---|---|---|
| GATCGGCGATTGGGACTAAGTCGTAACAAGG | 19.88 | 2.76 | 11 | 0.13 | 1473313 | 1473329/1484T |

#### *gyrA*

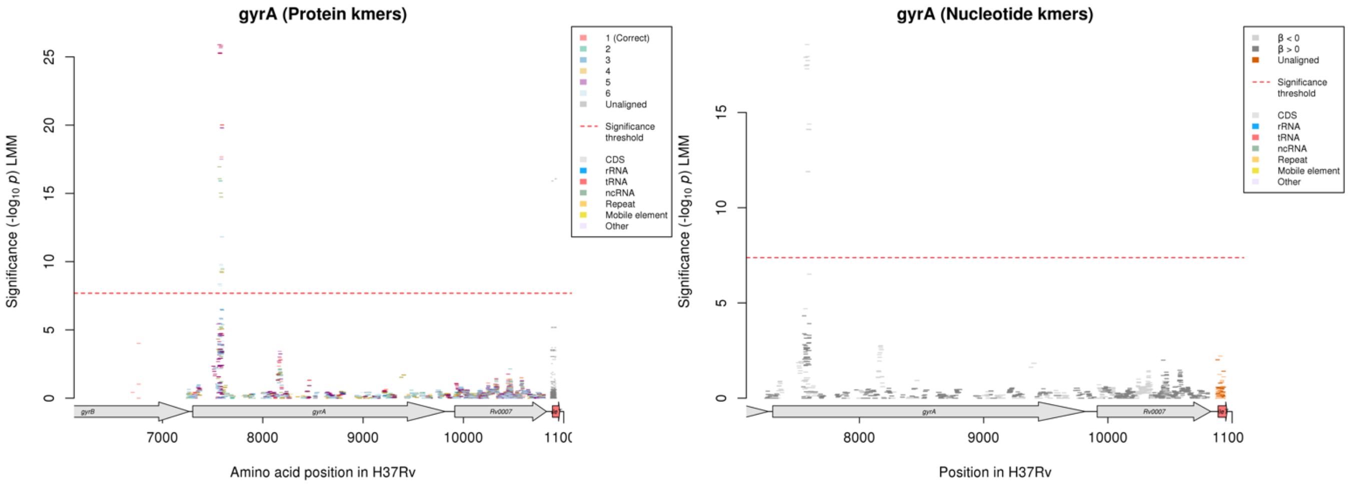

Protein kmers:

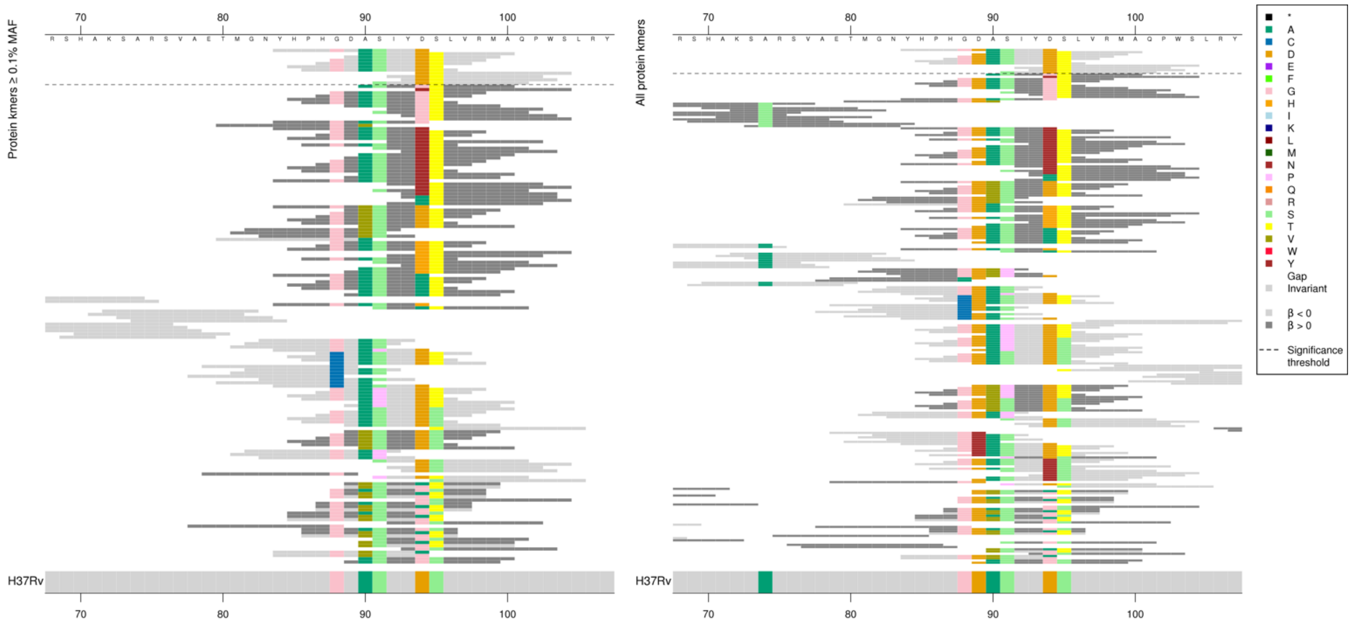

*gyrA* = DNA gyrase subunit A

| Kmer | $-\log_{10} p$ | β | MAC | MAF | Ps | Variants |
|---|---|---|---|---|---|---|
| YHPHGDASIYD | 25.88 | -0.59 | 1296 | 15.37 | 84 | 88G, 89D, 90A, 91S, 94D |
| ASIYDTLVRMA | 25.87 | -0.60 | 1733 | 20.56 | 90 | 90A, 91S, 94D, 95T |
| DASIYDTLVRM | 25.73 | -0.60 | 1740 | 20.64 | 89 | 89D, 90A, 91S, 94D, 95T |
| HPHGDASIYDT | 25.26 | -0.59 | 1756 | 20.83 | 85 | 88G, 89D, 90A, 91S, 94D, 95T |
| DTLVRMAQPWS | 20.01 | -0.55 | 1344 | 15.94 | 94 | 94D, 95T |
| SIYDTLVRMAQ | 17.67 | -0.50 | 1409 | 16.71 | 91 | 91S, 94D, 95T |

*rpoB*

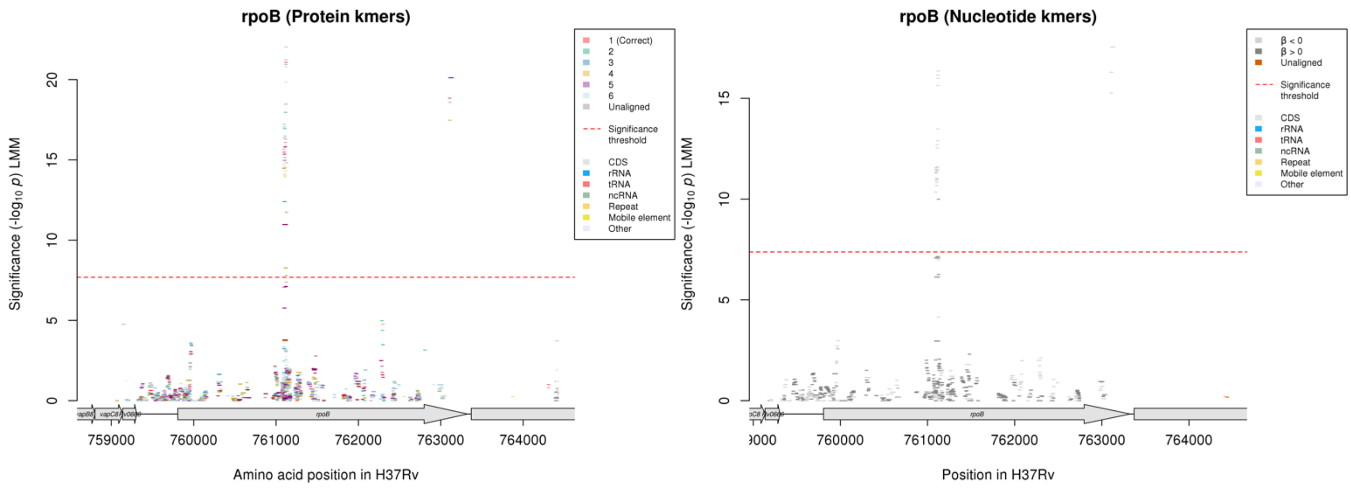

#### Protein kmers:

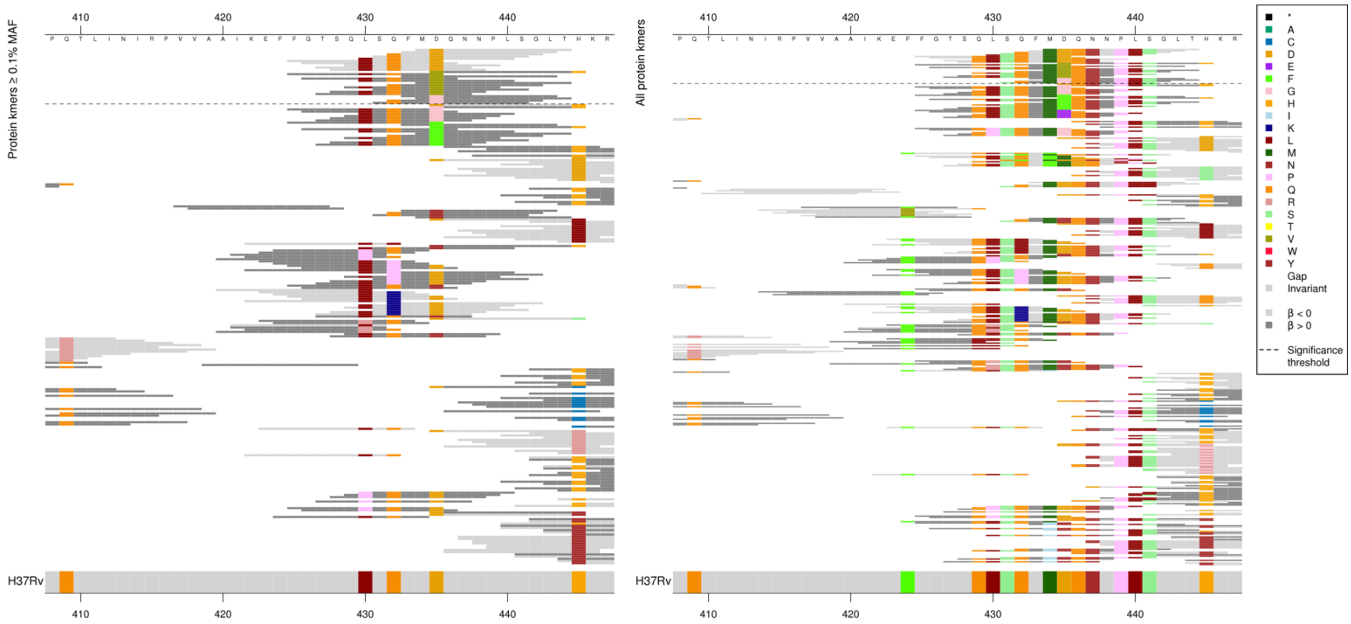

| Kmer | -log₁₀ *p* | β | MAC | MAF | Ps | Variants |
|---|---|---|---|---|---|---|
| MDQNNPLSGLT | 21.08 | -0.73 | 621 | 7.37 | 434 | 434M, 435D, 436Q, 437N, 439P, 440L, 441S |
| QFMDQNNPLSG | 16.29 | -0.61 | 663 | 7.86 | 432 | 432Q, 434M, 435D, 436Q, 437N, 439P, 440L, 441S |
| SQFMDQNNPLS | 15.84 | -0.60 | 666 | 7.90 | 431 | 431S, 432Q, 434M, 435D, 436Q, 437N, 439P, 440L, 441S |
| LSQFMDQNNPL | 15.79 | -0.56 | 734 | 8.71 | 430 | 430L, 431S, 432Q, 434M, 435D, 436Q, 437N, 439P, 440L |
| FGTSQLSQFMD | 15.66 | -0.55 | 737 | 8.74 | 425 | 429Q, 430L, 431S, 432Q, 434M, 435D |
| GTSQLSQFMDQ | 15.50 | -0.55 | 739 | 8.77 | 426 | 429Q, 430L, 431S, 432Q, 434M, 435D, 436Q |
| QLSQFMDQNNP | 15.36 | -0.55 | 740 | 8.78 | 429 | 429Q, 430L, 431S, 432Q, 434M, 435D, 436Q, 437N, 439P |
| TSQLSQFMDQN | 15.36 | -0.55 | 742 | 8.80 | 427 | 429Q, 430L, 431S, 432Q, 434M, 435D, 436Q, 437N |
| VQNNPLSGLTH | 10.98 | 0.66 | 446 | 5.29 | 435 | 435V, 436Q, 437N, 439P, 440L, 441S, 445H |
| FGTSQLSQFMV | 10.97 | 0.66 | 447 | 5.30 | 425 | 435V (+others) |
| FMGQNNPLSGL | 8.27 | 1.26 | 34 | 0.40 | 433 | 434M, 435G, 436Q, 437N, 439P, 440L, 441S (+ others) |

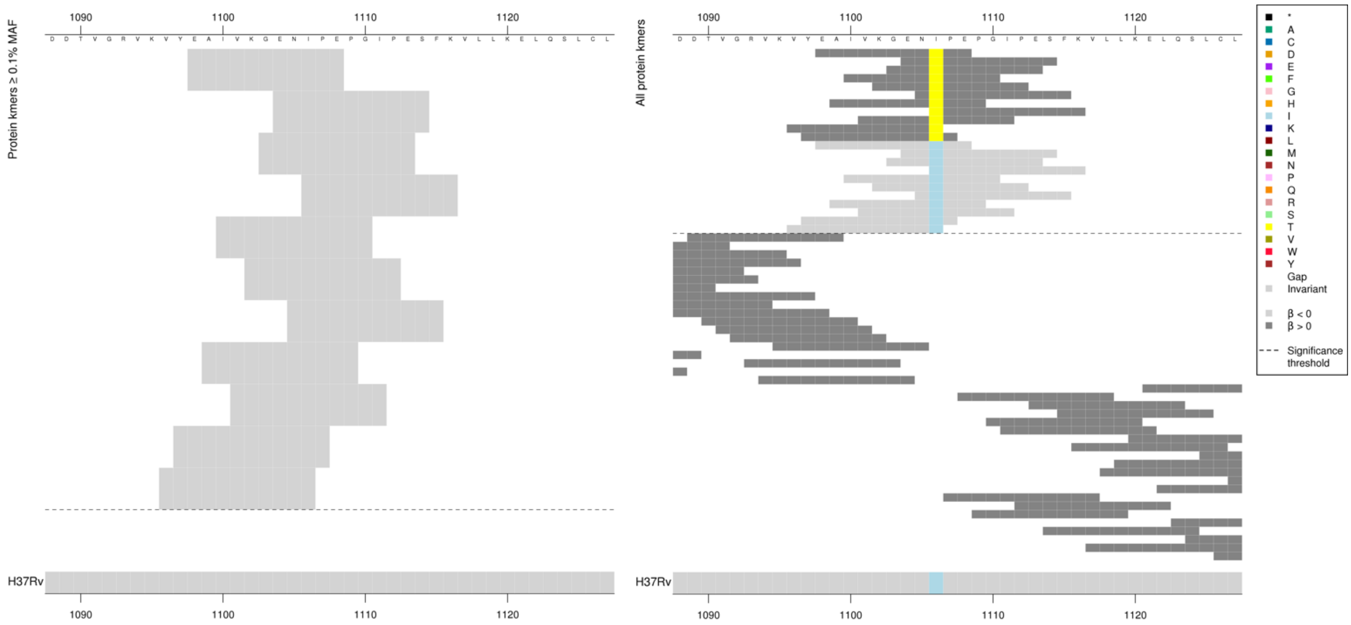

Low MAF kmers = 1106T.

| Kmer | -log₁₀ *p* | β | MAC | MAF | Ps | Variants |
|---|---|---|---|---|---|---|
| EAIVKGENIPE | 20.12 | -3.59 | 9 | 0.11 | 1098 | 1106I |

### *echA8*

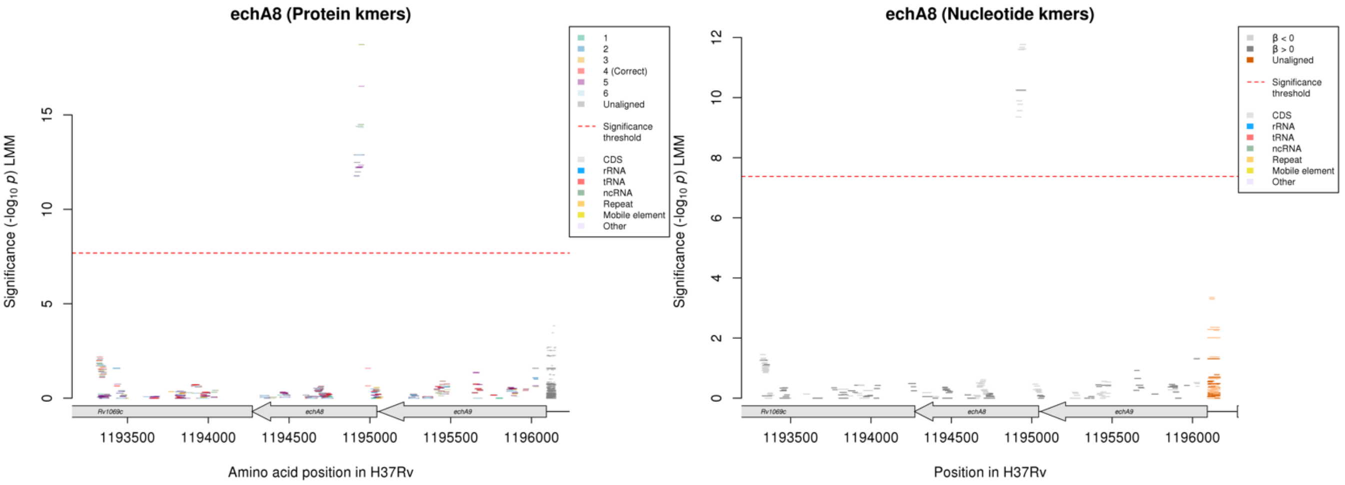

No significant kmers aligned to the correct reading frame of *echA8* in the protein kmer analysis but there were significant kmers in the nucleotide kmer analysis.

Nucleotide kmers:

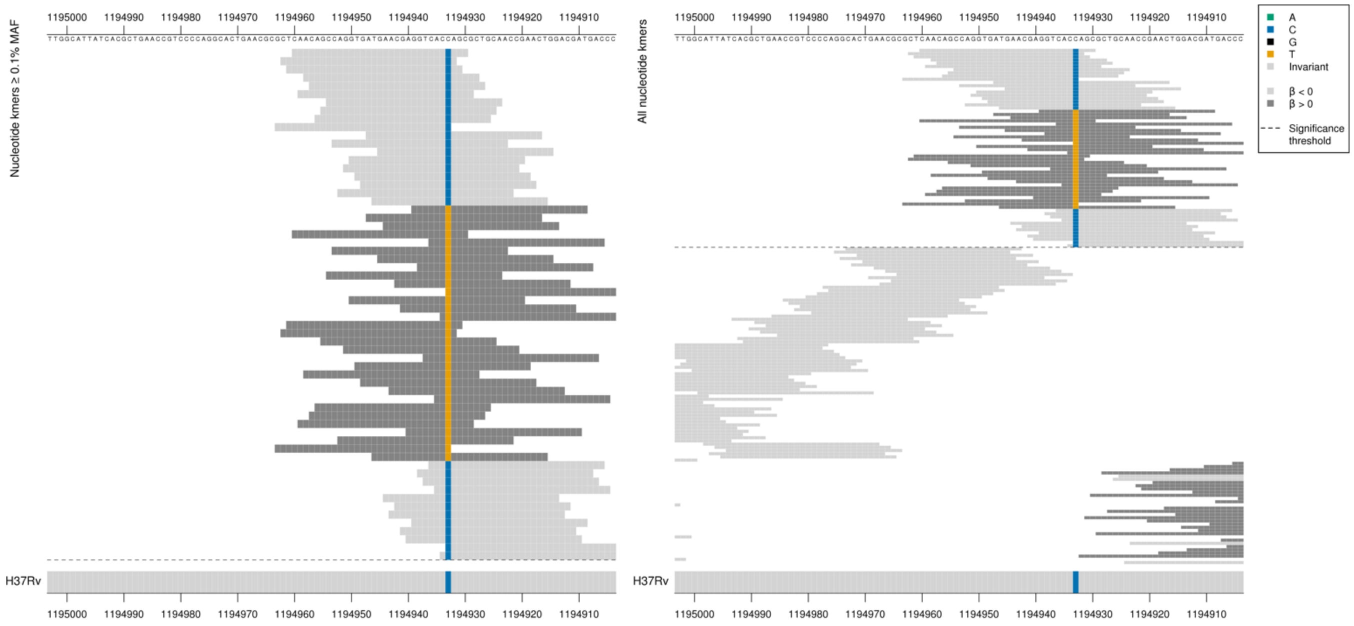

*echA8* = enoyl-CoA hydratase EchA8

Low MIC-associated = ACC = Thr / T
High MIC-associated = ACT = Thr / T
Synonymous mutation 1194923/121

| Kmer | $-\log_{10} p$ | β | MAC | MAF | Ps | Variants |
|---|---|---|---|---|---|---|
| CAACAGCCAGGTGATGAACGAGGTCACCAGC | 11.76 | -2.32 | 17 | 0.20 | 1194960 | 1194923/121C |
| GGTCACTAGCGCTGCAACCGAACTGGACGAT | 10.24 | 2.51 | 14 | 0.17 | 1194939 | 1194923/121T |

# *Rv2896c*

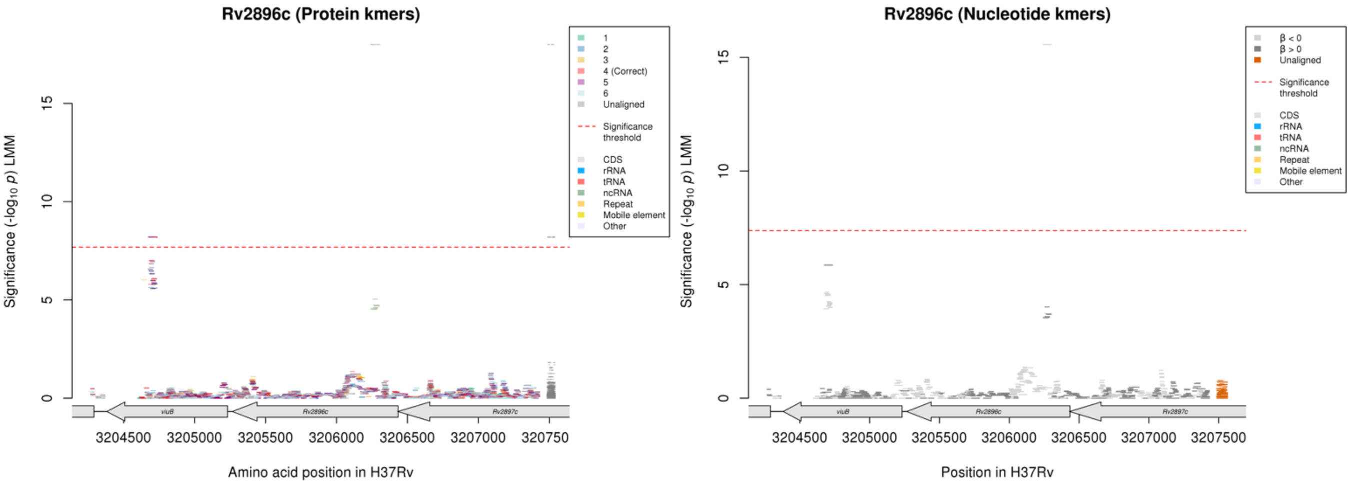

No significant kmers aligned to the correct reading frame of *Rv2896c* in the protein kmer analysis, but there were significant kmers in the nucleotide analysis.

Nucleotide kmers:

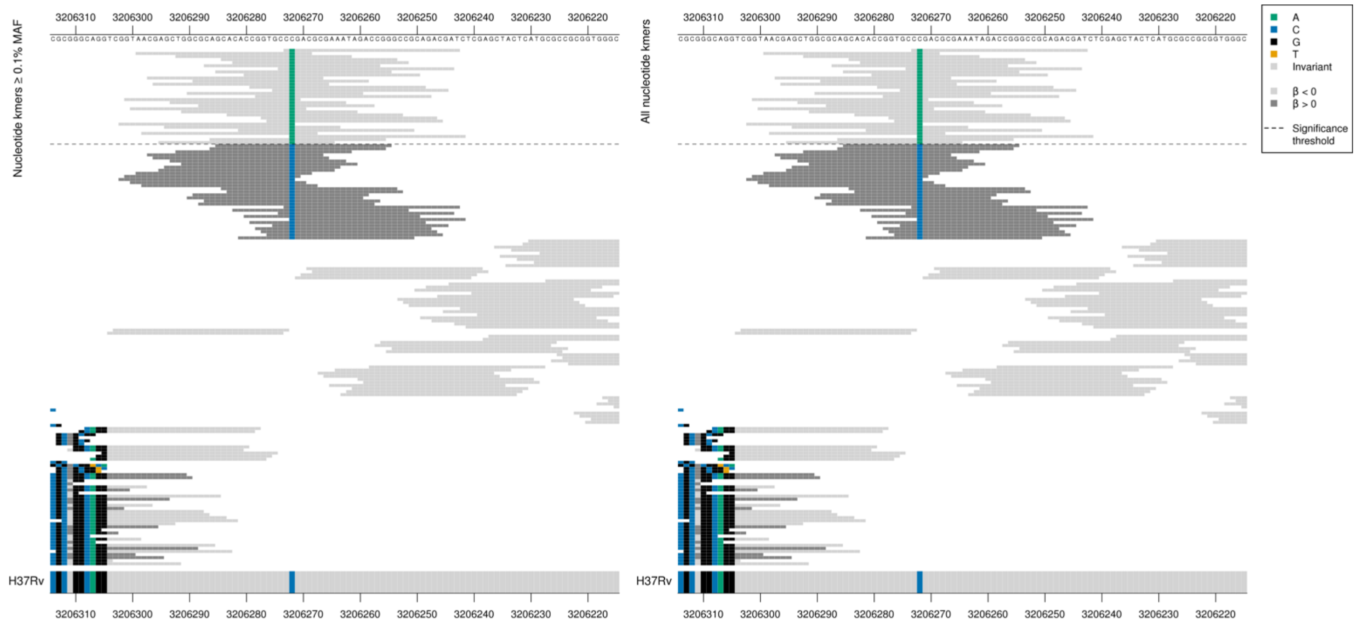

Low MIC-associated = AGA = Arg / R

Kmers below significance threshold = CGA = Arg / R

Synonymous mutation 3206272/163 – possible LD with *Rv0078A*

| Kmer | $-\log_{10} p$ | β | MAC | MAF | Ps | Variants |
|---|---|---|---|---|---|---|
| CAGACGCGAAATAGACCGGGCCGCAGACGAT | 15.57 | -2.53 | 12 | 0.14 | 3206273 | 3206272/163A |

# Rv0078A

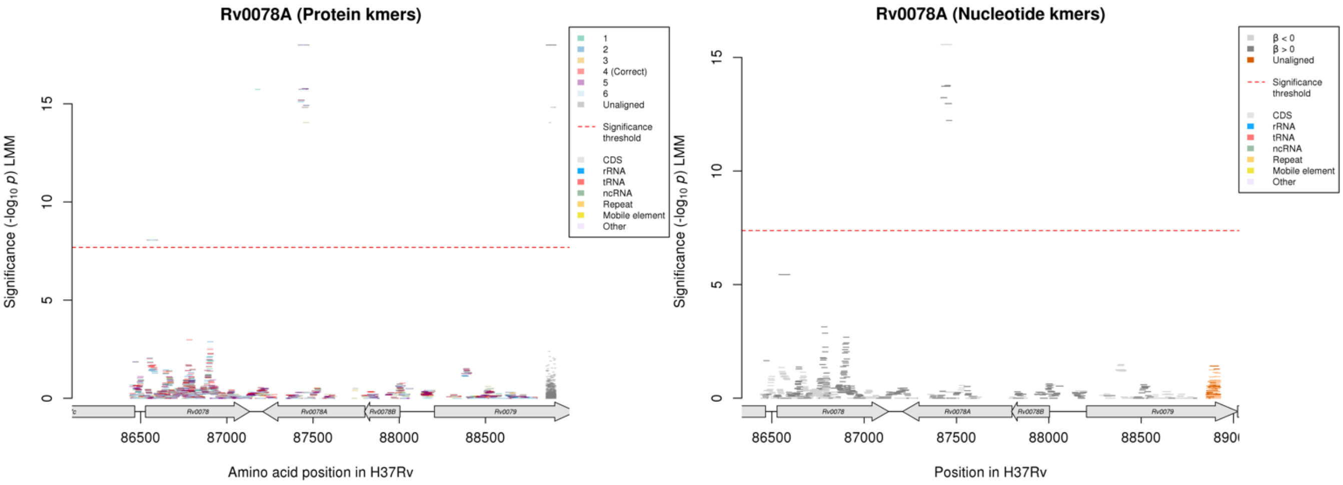

This region is also significant for BDQ, but with an opposite direction of effect.

Protein kmers:

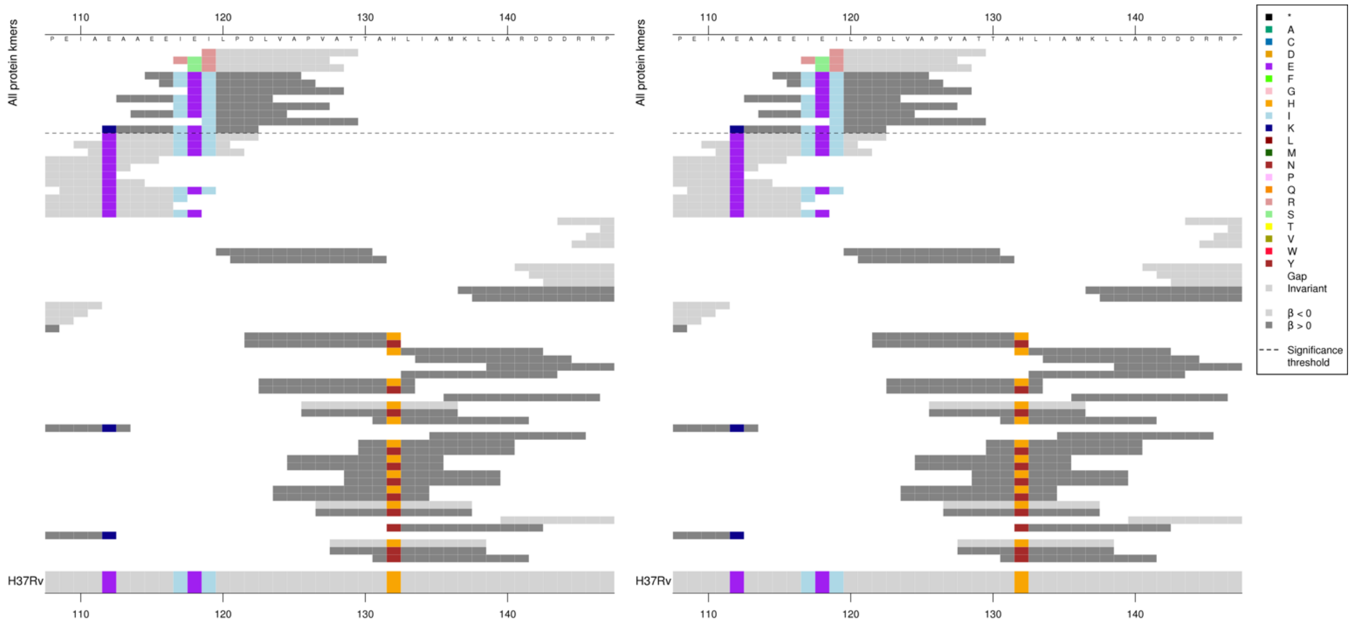

Nucleotide kmers:

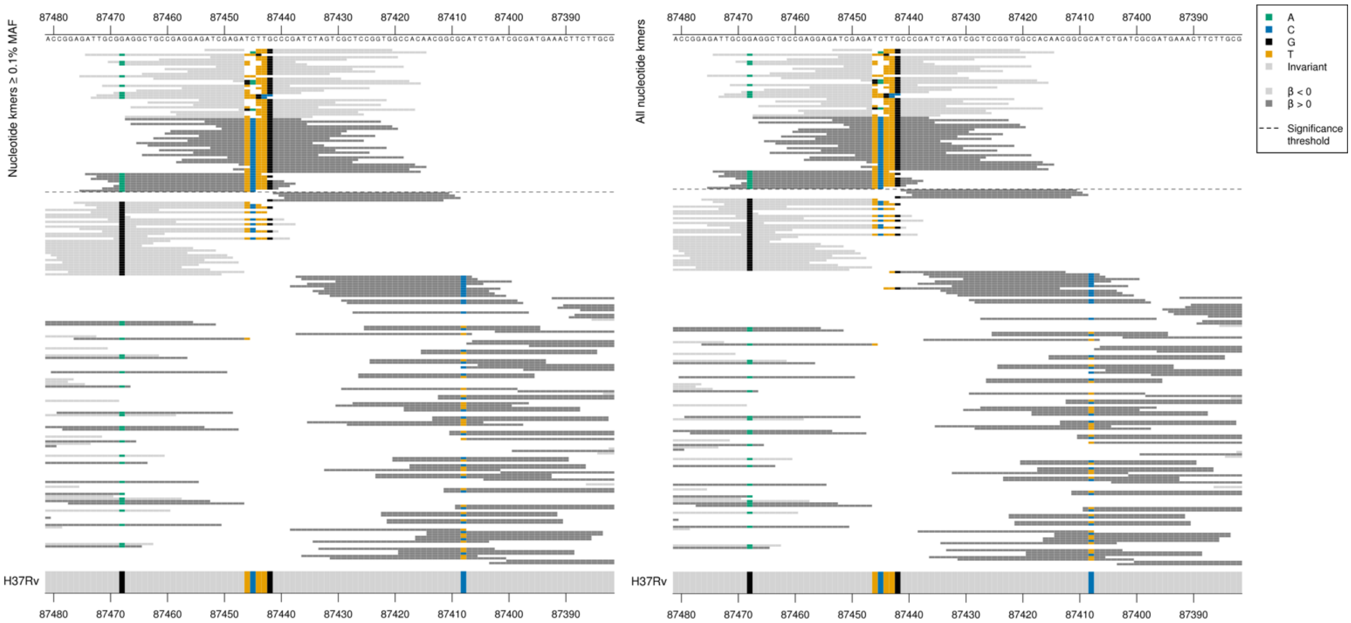

The nucleotide kmer alignments show that there were deletions in the low MIC-associated kmers relative to the reference. Deletion of CT at positions 356-357 (87446-87445) in low MIC-associated kmers.
All high MIC-associated kmers share the base 87445C.
All high MIC-associated kmers in the first group that don't cover 87468/334A contain 87446-87445CT.

| Kmer | $-\log_{10} p$ | β | MAC | MAF | Ps | Variants |
|---|---|---|---|---|---|---|
| ATCGAGATTGCCCGATCTAGTCGCTCCGGTG | 15.57 | -2.53 | 12 | 0.14 | 87453 | 87446-87445/356-357 TC deletion |
| AGGCTGCCGAGGAGATCGAGATCTTGCCCGA | 13.77 | 2.19 | 18 | 0.21 | 87467 | 87446-87445/356-357 TC |
| ATTGCGAAGGCTGCCGAGGAGATCGAGATCT | 12.97 | 2.03 | 2947 | 34.96 | 87474 | 87468/334A, 87446-87445/356-357 TC |

## *Rv1830*

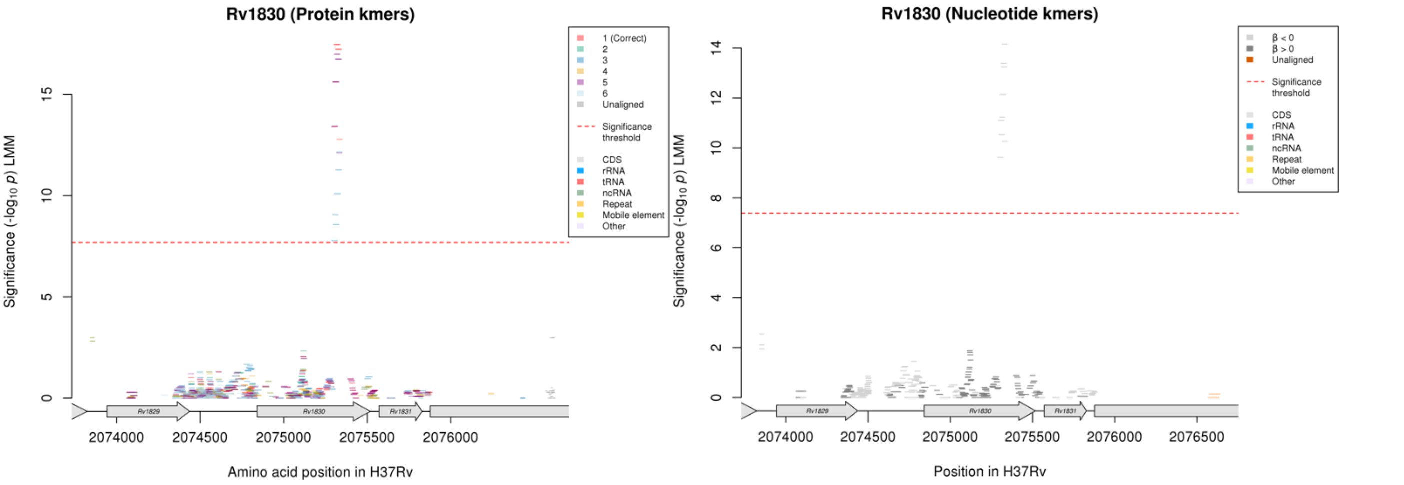

Protein kmers:

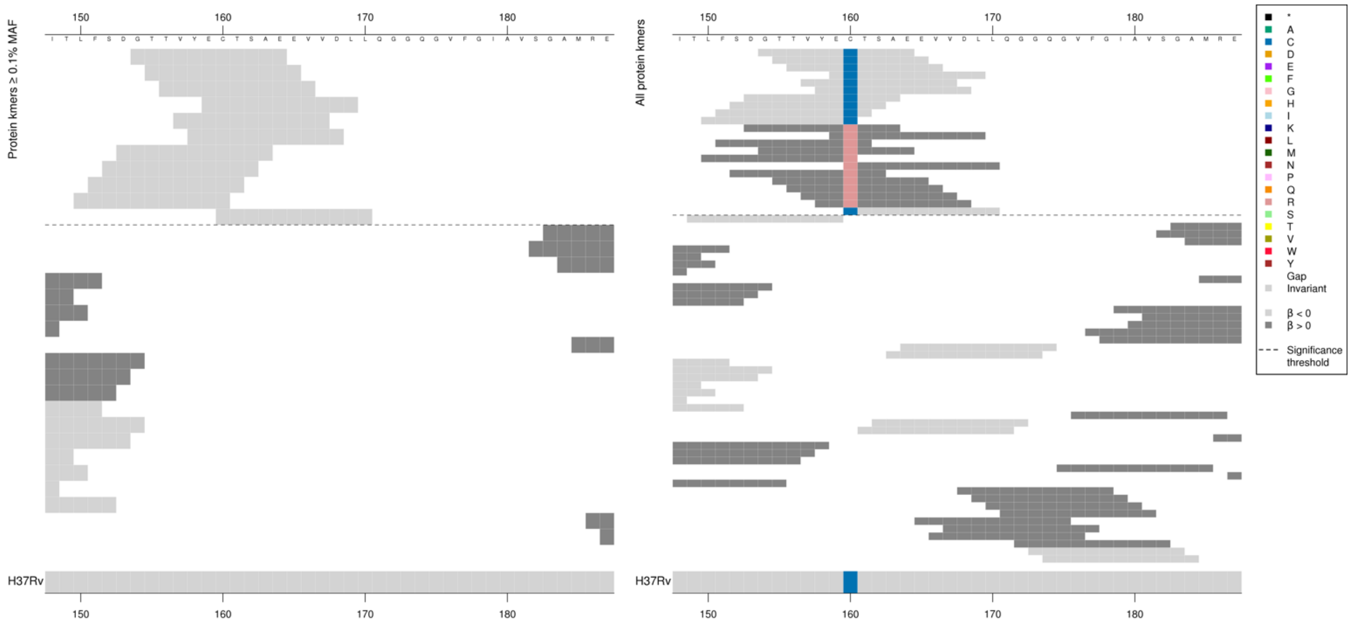

*Rv1830* = HTH-type transcriptional regulator

Low MAF kmers = 160R.

| Kmer | $-\log_{10} p$ | β | MAC | MAF | Ps | Variants |
|---|---|---|---|---|---|---|
| GTTVYECTSAE | 17.45 | -3.23 | 12 | 0.14 | 154 | 160C |

# Rv0792c/Rv0793

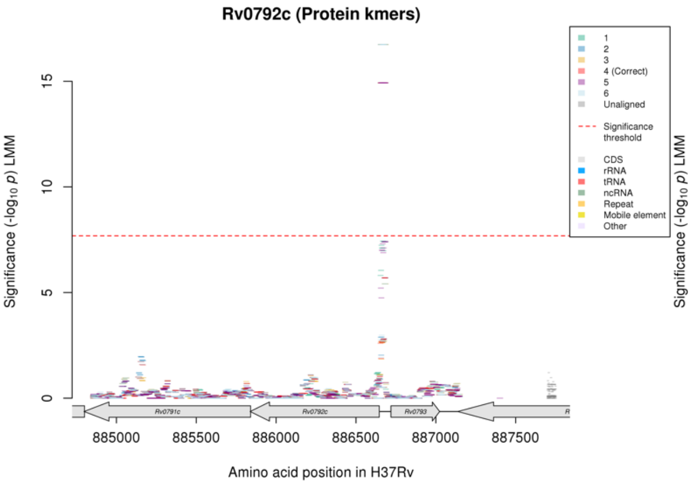

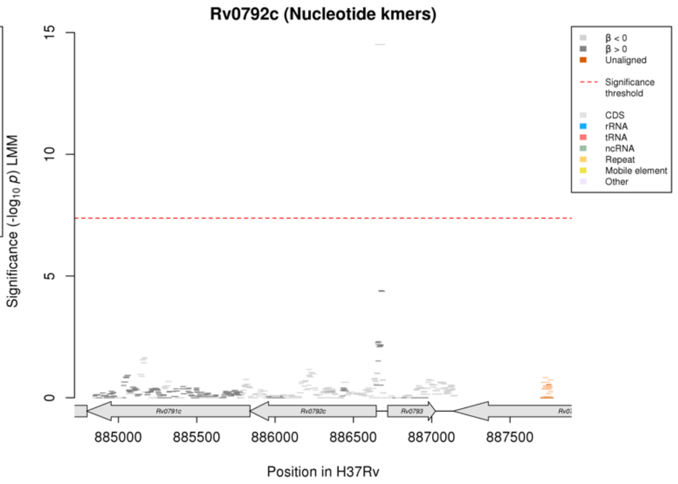

Nucleotide kmers:

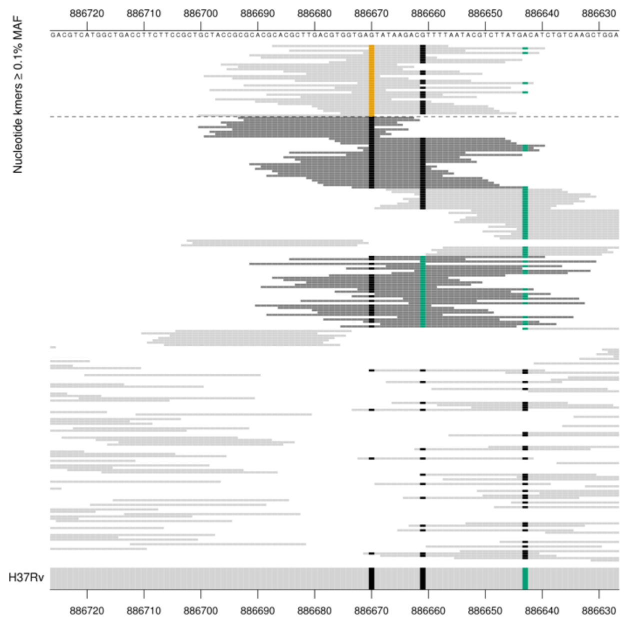

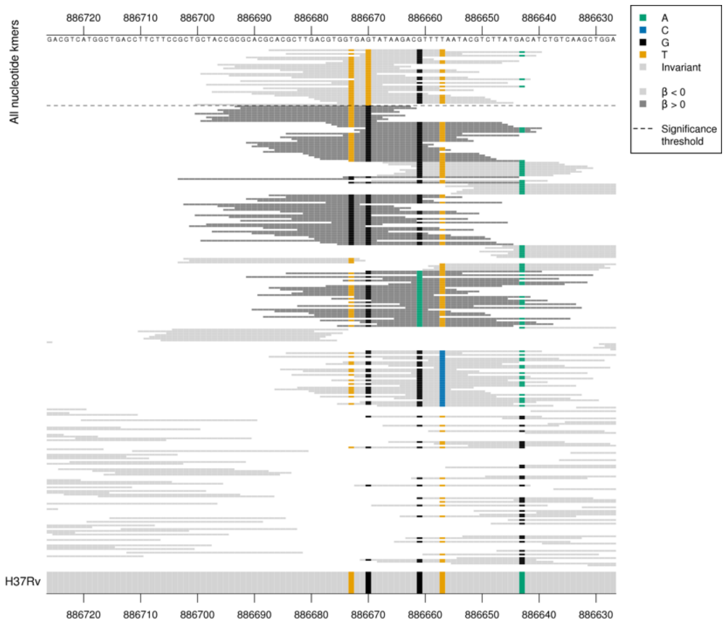

*Rv0792c* = transcriptional regulator

*Rv0792c* start position is 886646 on the reverse strand.
Mutation in possible promoter region, position 886670/-24.
Rv0793 start position is 886719 on the forward strand.
Mutation in possible promoter region, position 886670/-49.

| Kmer | $-\log_{10} p$ | β | MAC | MAF | Ps | Variants |
|---|---|---|---|---|---|---|
| CACGCTTGACGTGGTGATTATAAGACGTTTT | 14.51 | -2.34 | 13 | 0.15 | 886687 | 886670/-24 T Rv0792c/ 886670/-49 A Rv0793 |

### *PPE54*

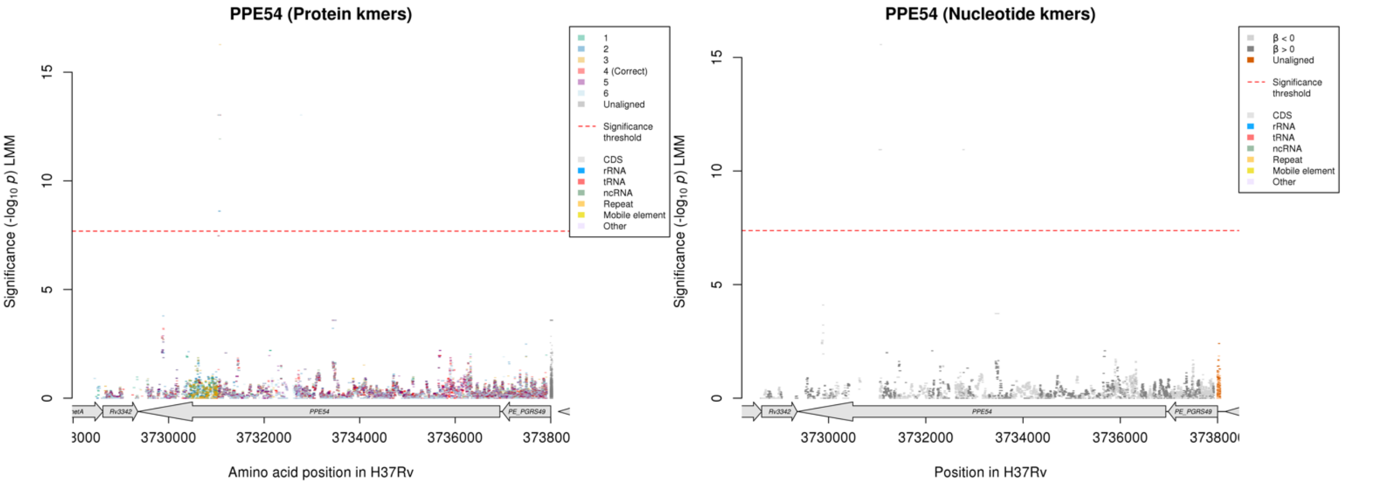

For the correct reading frame, the significant protein kmers aligned to just one region of the gene *PPE54*. For the nucleotide kmer analysis, there were significant kmers in two regions of the gene.

Protein kmers:

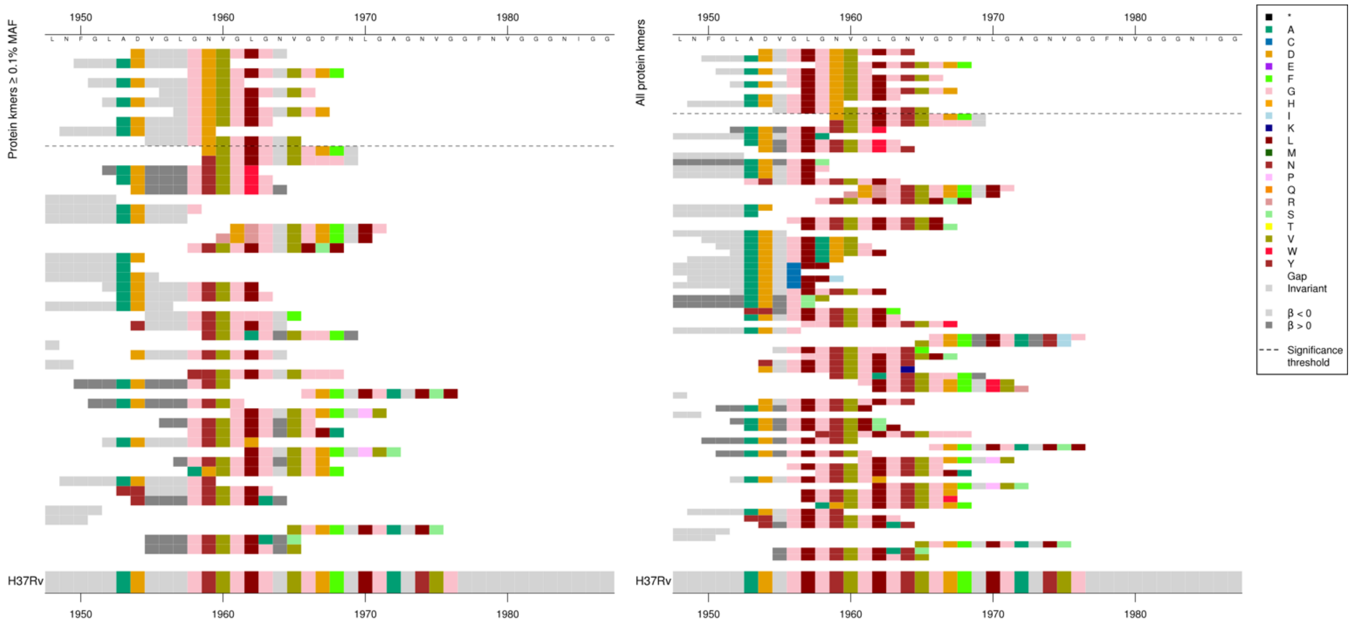

Nucleotide kmers:

The same region for the nucleotide kmer analysis that the significant protein kmers aligned to:

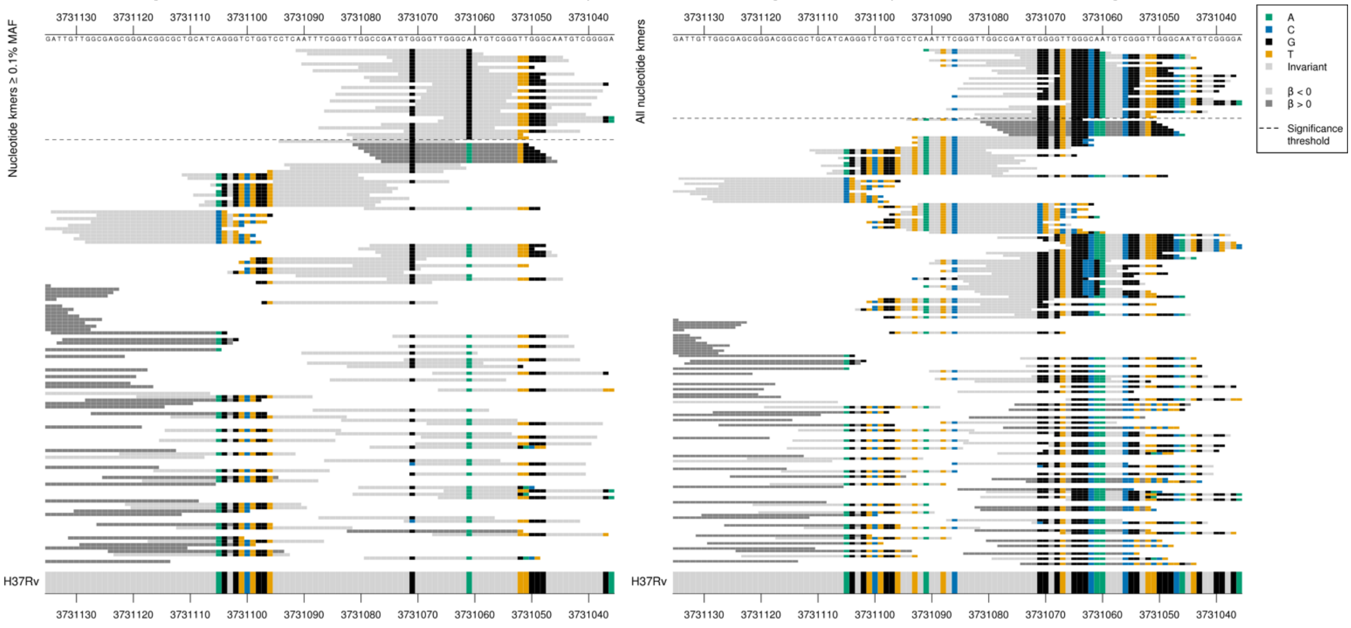

The second region containing significant kmers in *PPE54* in the nucleotide kmer analysis:

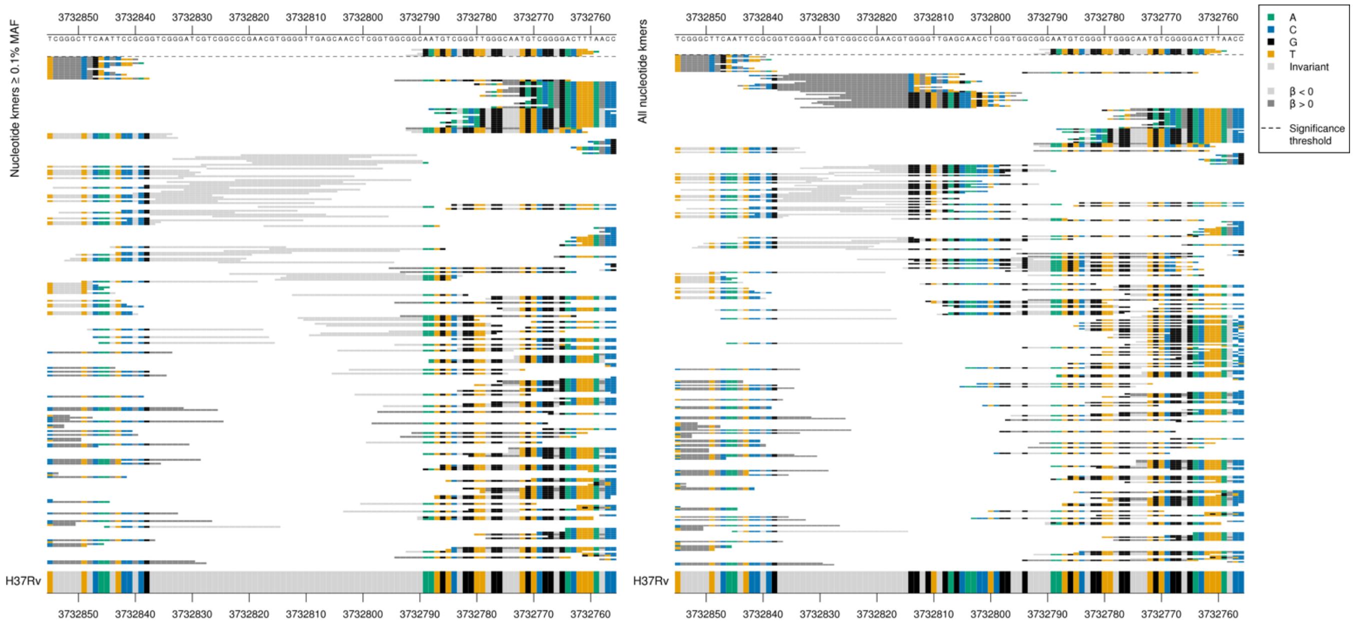

These four kmers aligned equally well with the region in first alignment above. Capturing 31 kmers and therefore likely a single mutation.

*PPE54* begins at position 3736935.
Mutation at position 3731061, first position in codon 1959, GAT = Asp / D

| Kmer | $-\log_{10} p$ | β | MAC | MAF | Ps | Variants |
|---|---|---|---|---|---|---|
| DVGLGDVGLGN | 13.03 | -2.37 | 11 | 0.13 | 1954 | 1959D |

# *Rv2041c*

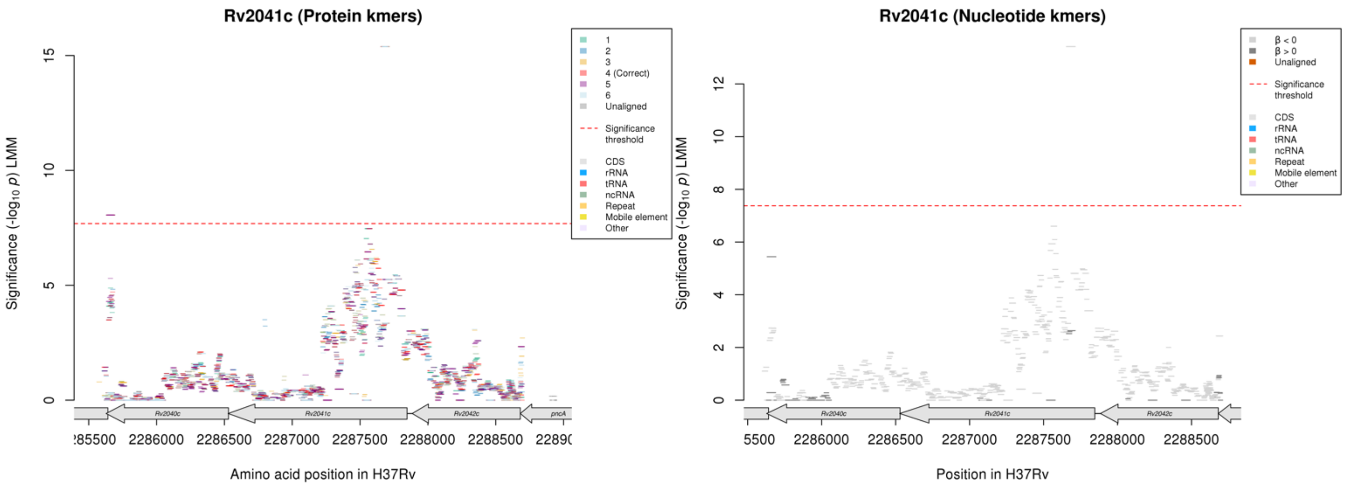

*Rv2041c* product = sugar ABC transporter substrate-binding lipoprotein

#### Amikacin (AMI)

##### Protein kmers:

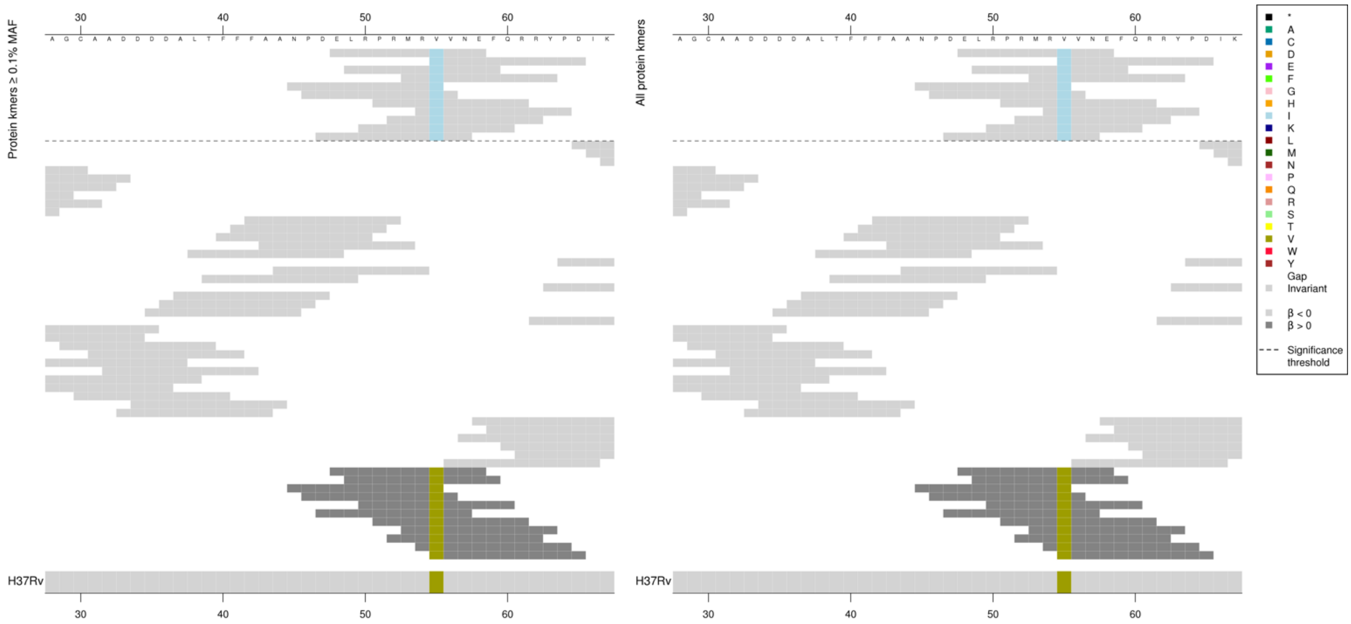

| Kmer | $-\log_{10} p$ | β | MAC | MAF | Ps | Variants |
|---|---|---|---|---|---|---|
| ELRPRMRIVNE | 15.39 | -2.37 | 15 | 0.18 | 48 | 55I |

#### *PPE42*

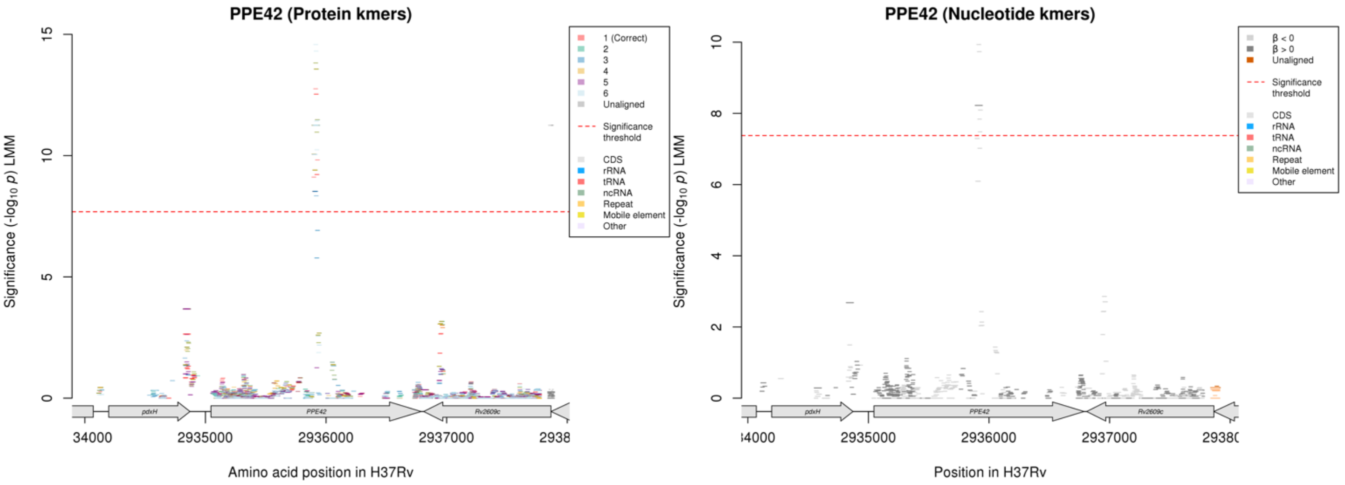

##### Protein kmers:

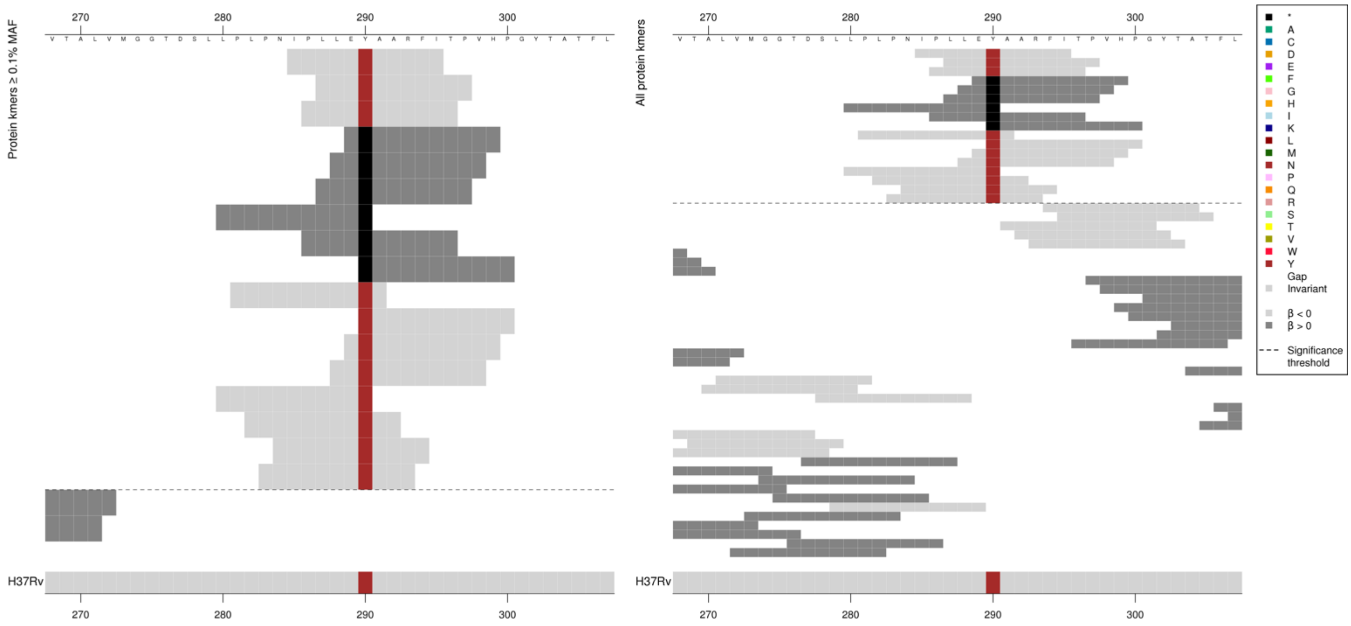

Premature stop codon is found in kmers associated with high MICs.

| Kmer | $-\log_{10} p$ | β | MAC | MAF | Ps | Variants |
|---|---|---|---|---|---|---|
| IPLLEYAARFI | 12.76 | -2.67 | 42 | 0.50 | 285 | 290Y |
| E*AARFITPVH | 11.25 | 3.12 | 38 | 0.45 | 289 | 290* |

#### *cyp141:Rv3122*

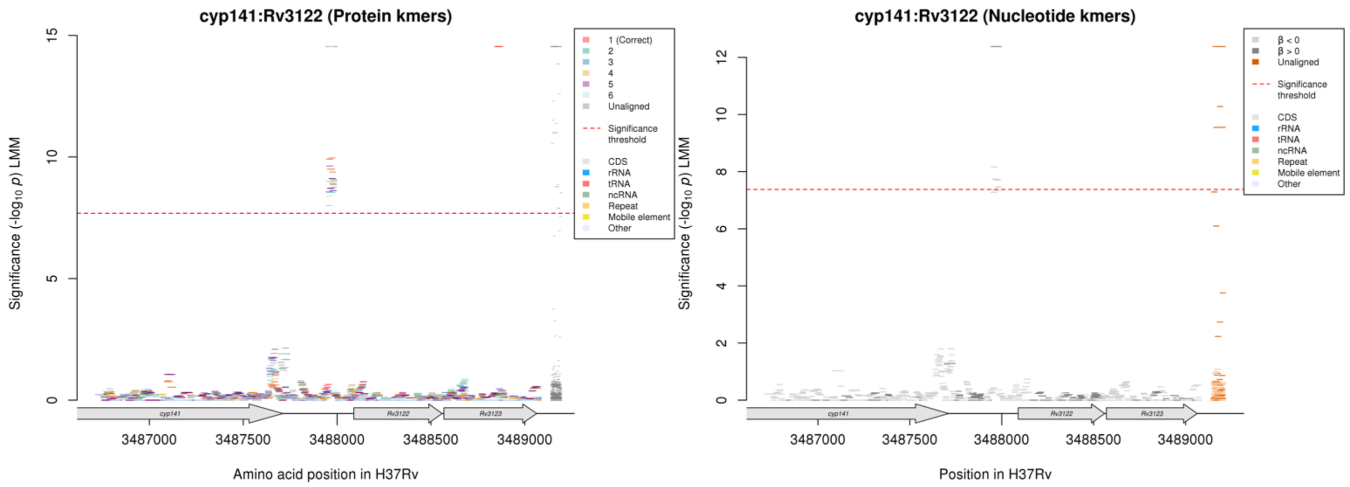

Nucleotide kmers:

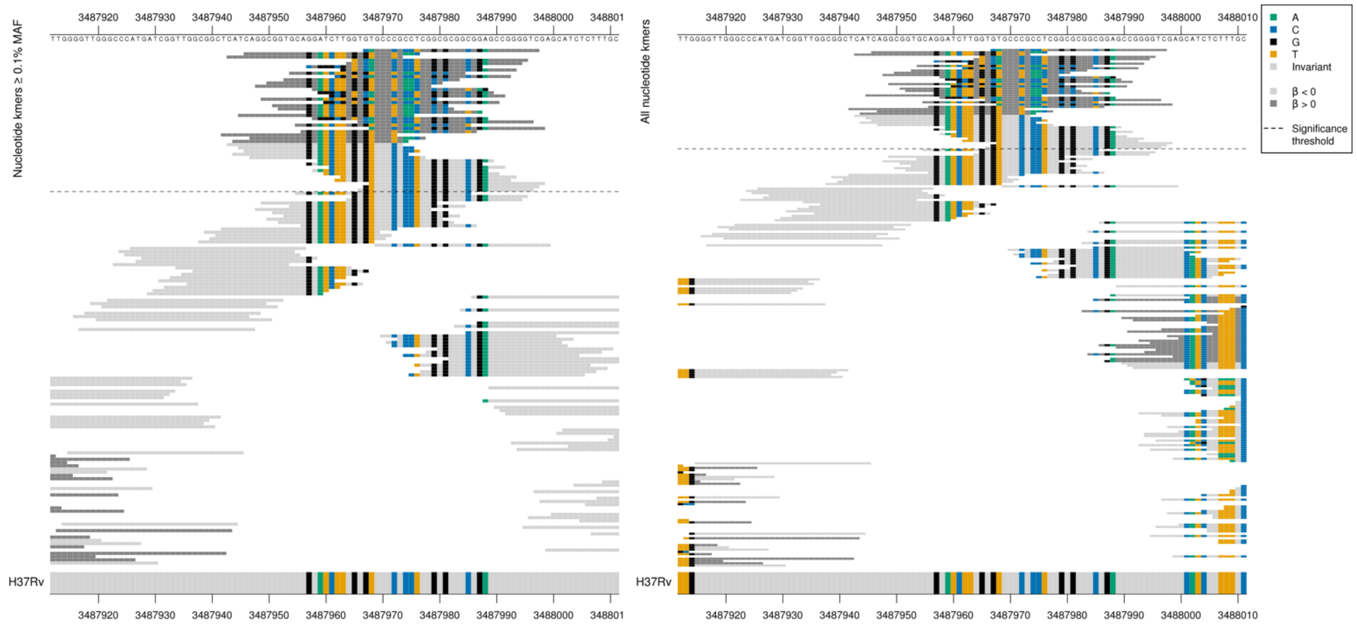

Repetitive region, kmers align to all three regions, alignments were not unique. Kmers in LD with *espA:ephA* and likely tagging that association. The *espA:ephA* kmer alignments were unique alignments.

| Kmer | $-\log_{10} p$ | β | MAC | MAF | Ps | Variants |
|---|---|---|---|---|---|---|
| CAGCCCGCCTCGGCGCGGCGGAGCCGGGGTC | 12.38 | 2.66 | 9 | 0.11 | 3487967 | Multiple differences to reference |
| CATCAGGCGGTGCAGGATCTTGGTGTGCCCG | 8.16 | -1.43 | 40 | 0.47 | 3487943 | WT |

## *Rv1765c* R

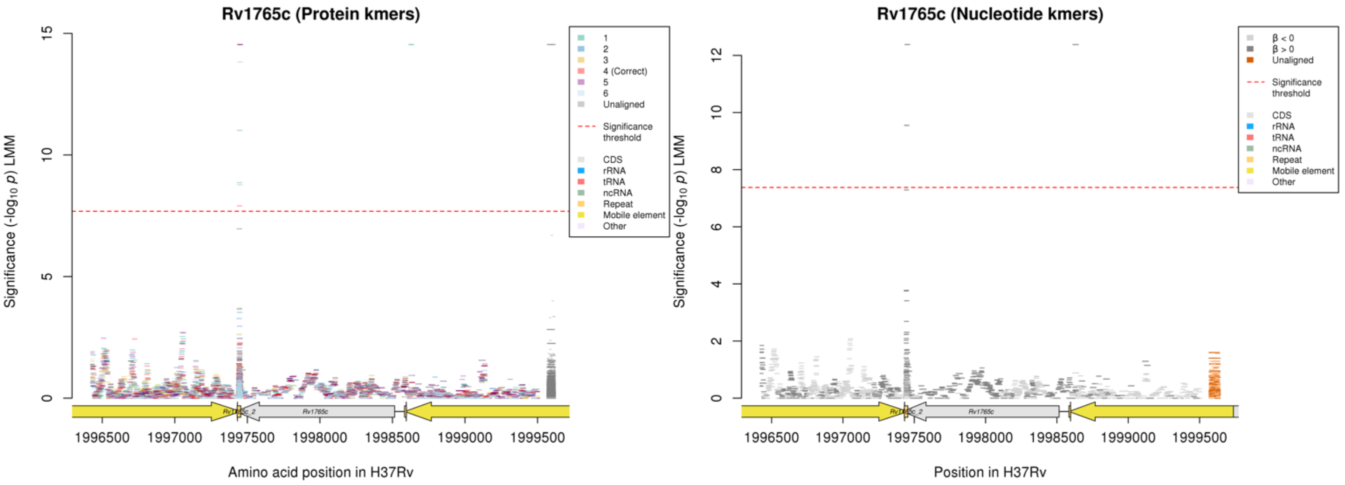

Non-unique alignments: Number of significant kmers:
*Rv1765c* = 19 (9 match *lprF:Rv1371*, 11 match *cyp141:3122*)
*lprF:Rv1371* = 24 (4 match *cyp141:Rv3122*, 9 match *Rv1765c*)
*cyp141:Rv3122* = 44 (4 match *lprF:Rv1371*, 11 match *Rv1765c*)

Nucleotide kmers:

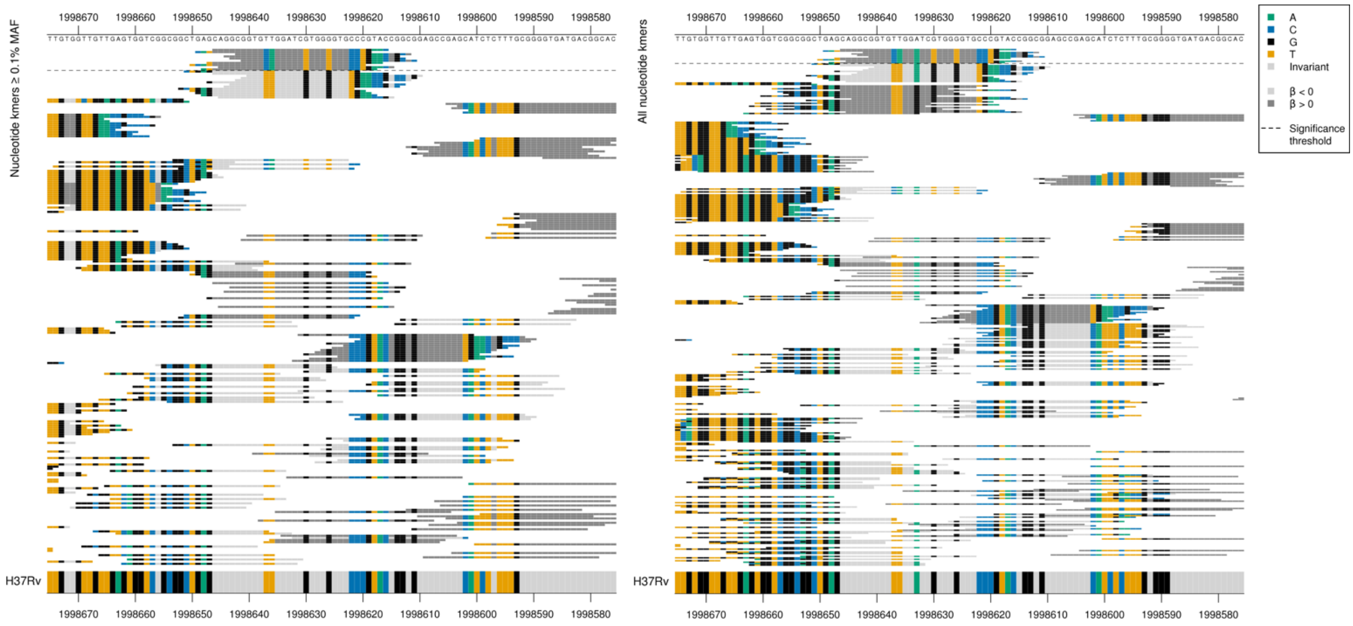

This region is upstream to Rv1765c, and falls within the mobile element.

| Kmer | $-\log_{10} p$ | β | MAC | MAF | Ps | Variants |
|---|---|---|---|---|---|---|
| CAGGCGGTGCAGGATCTTGGTGTGCCTGAAC | 12.38 | 2.66 | 9 | 0.11 | 1998646 | Multiple differences to reference |

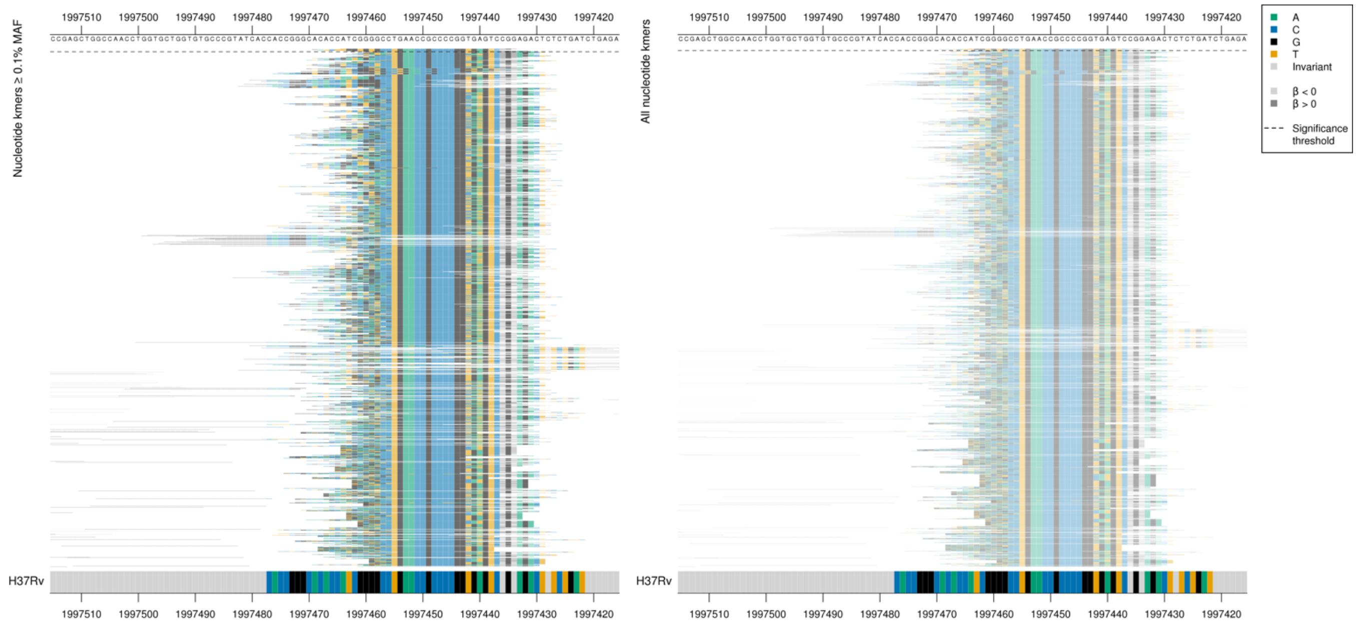

This region falls within Rv176c, the repeat region. The coding sequence is on the reverse strand positions 1998515-1997418, the repeat region is at positions 1997455-1997428 in H37Rv. This top kmer also aligned to the intergenic region lprF:Rv1371.

| Kmer | $-\log_{10} p$ | β | MAC | MAF | Ps | Variants |
|---|---|---|---|---|---|---|
| TGGTGTGCCTGAACCGCCCCGGTGAGTCCGG | 12.38 | 2.66 | 9 | 0.11 | 1997464 | Multiple differences to reference |

#### *lprF:Rv1371* $^{R}$

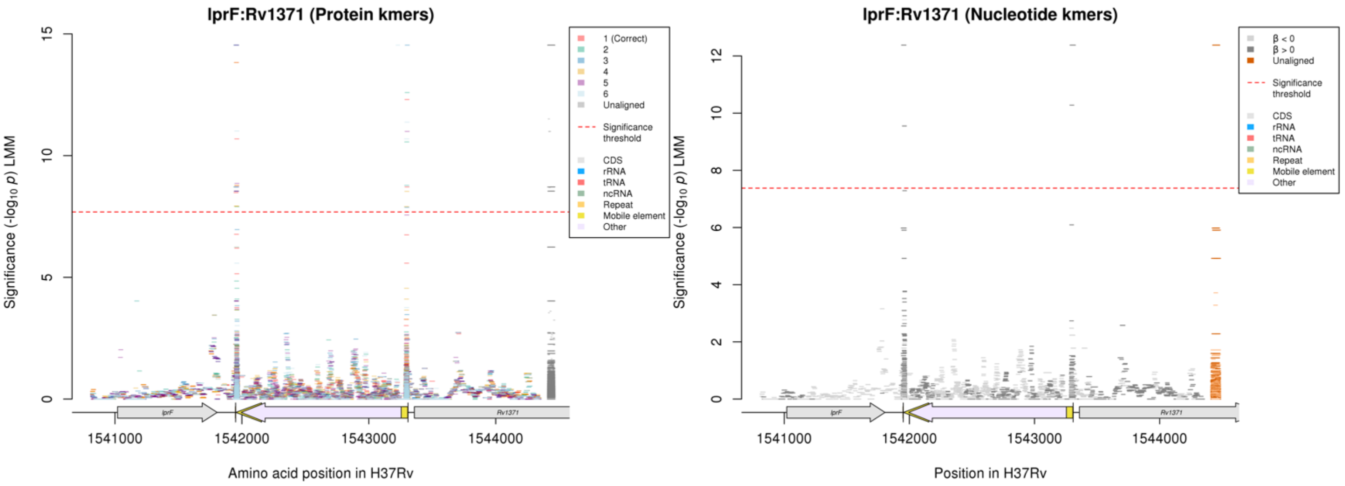

#### Nucleotide kmers:

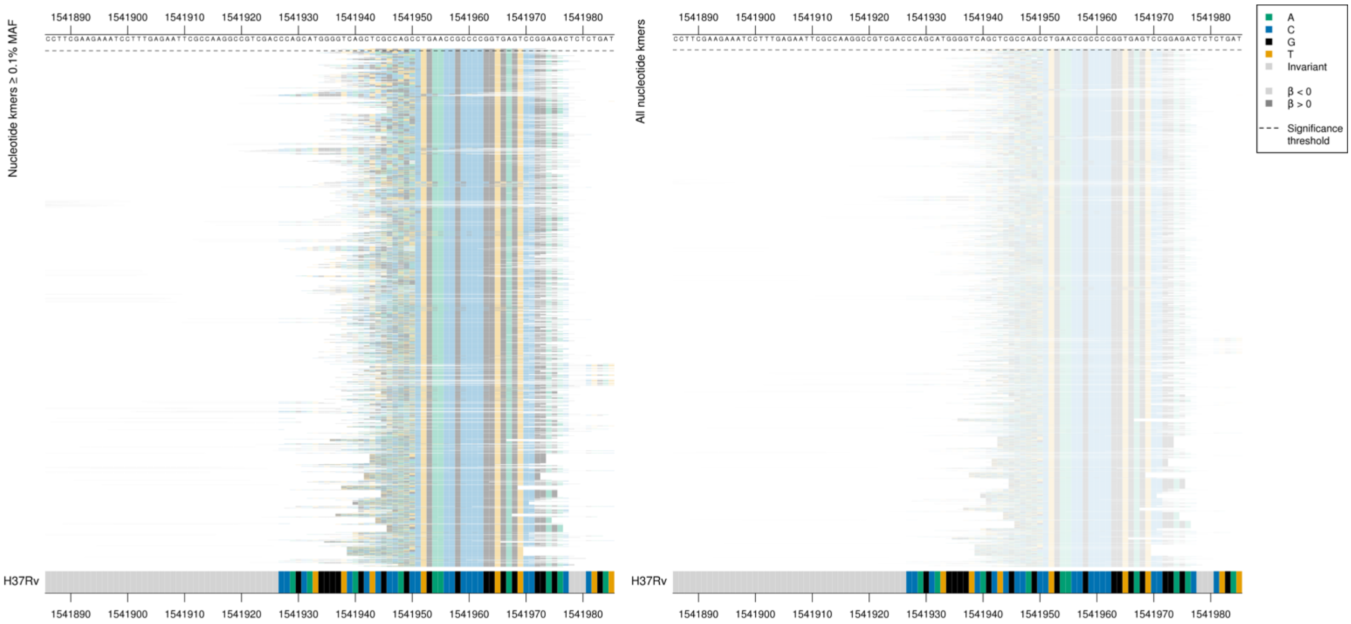

| Kmer | $-\log_{10} p$ | β | MAC | MAF | Ps | Variants |
|---|---|---|---|---|---|---|
| TGGTGTGCCTGAACCGCCCCGGTGAGTCCGG | 12.38 | 2.66 | 9 | 0.11 | 1541943 | Multiple differences to reference |

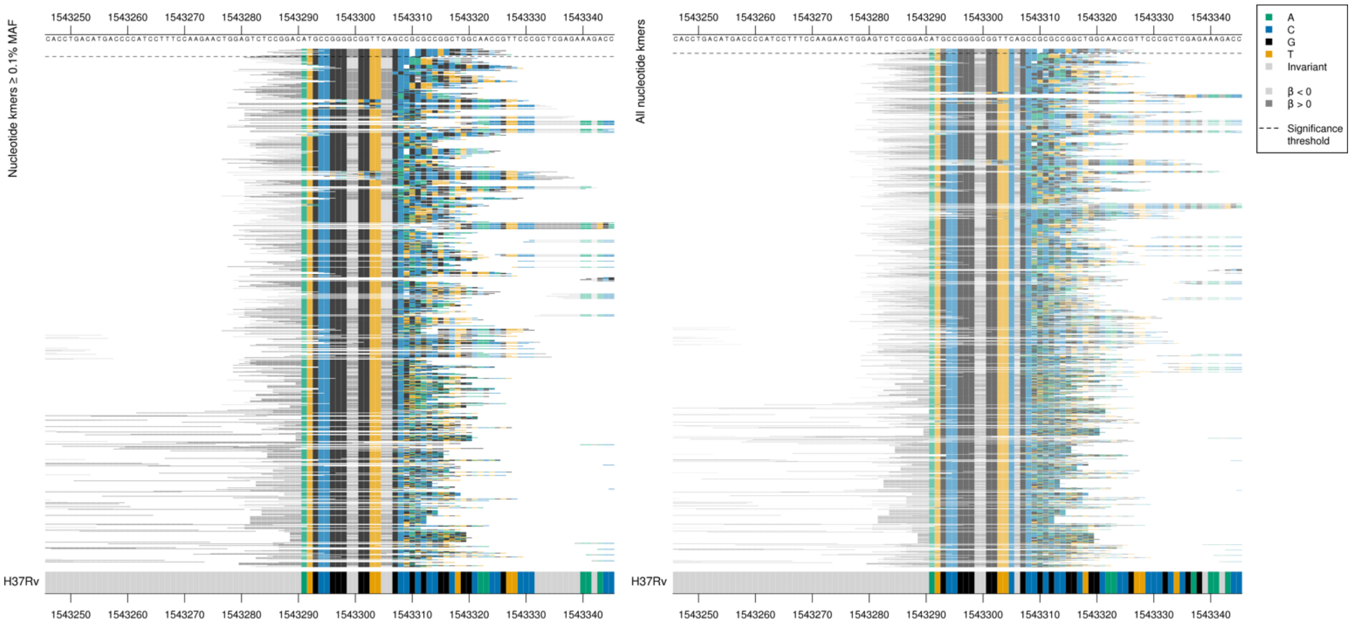

| Kmer | $-\log_{10} p$ | β | MAC | MAF | Ps | Variants |
|---|---|---|---|---|---|---|
| ATGCCGGGGCGGTTCAGCCCGCCTCGGCGCG | 12.38 | 2.66 | 9 | 0.11 | 1543291 | Multiple differences to reference |

### *espA:ephA*

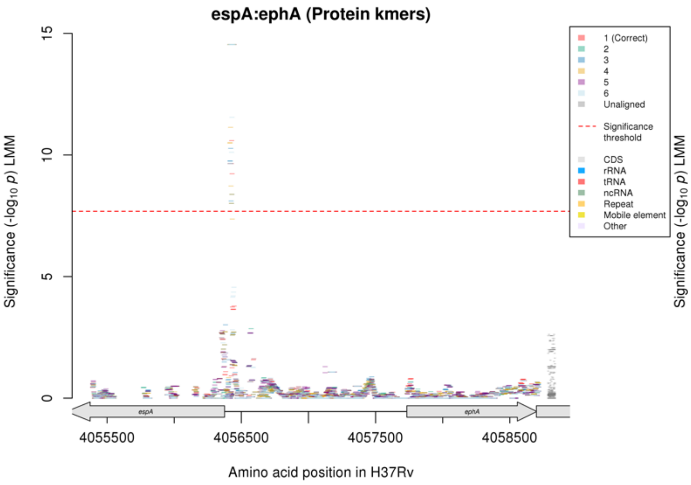

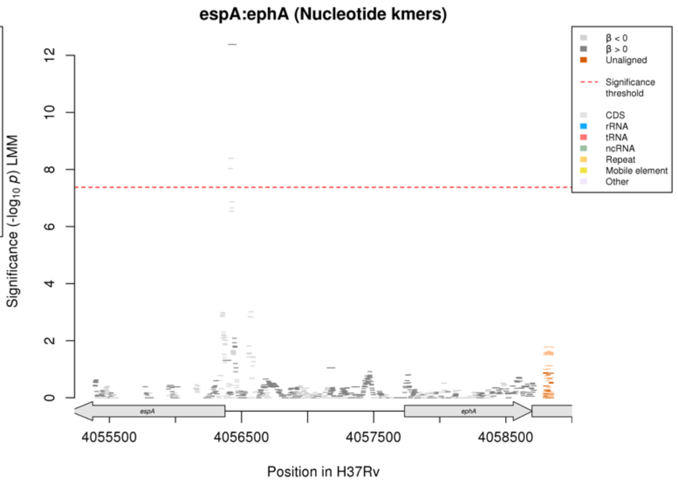

Nucleotide kmers:

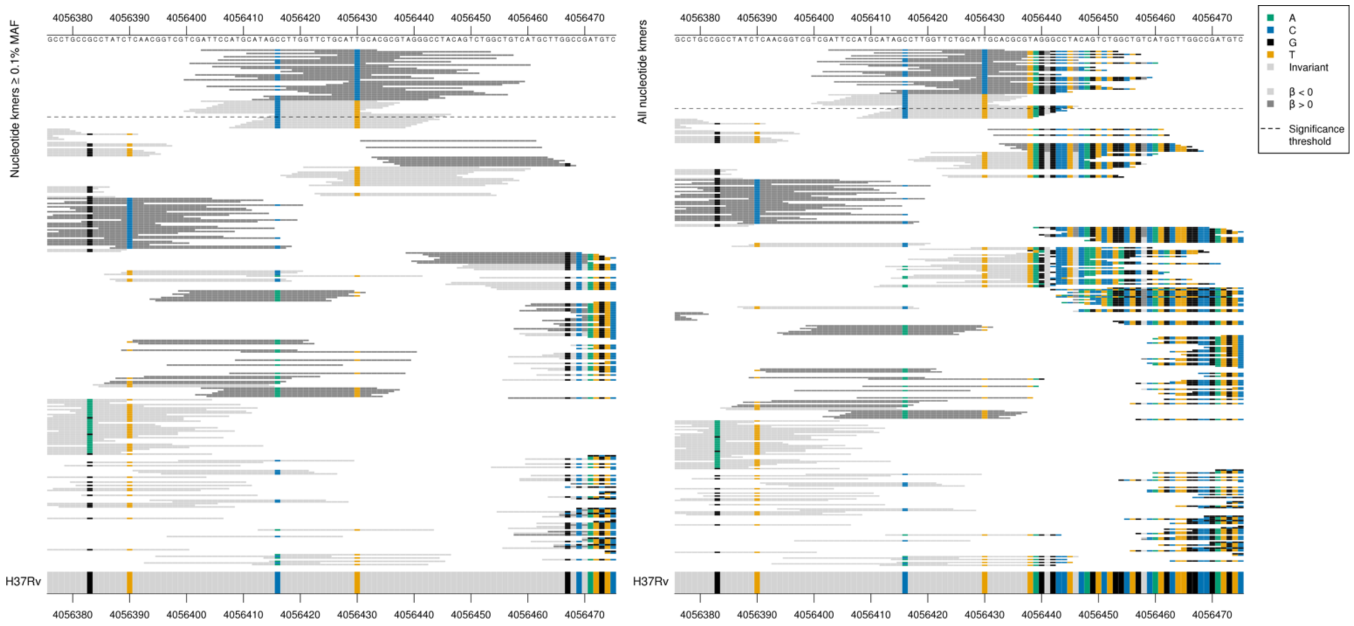

Alignments shown on the forward strand, *espA* encoded on the reverse strand.
*espA* product = ESX-1 secretion-associated protein EspA
*espA* begins at position 4056375, just off from the alignment.
Mutations in possible promoter region, positions 4056430/-55 and 4056416/-41.

| Kmer | $-\log_{10} p$ | β | MAC | MAF | Ps | Variants |
|---|---|---|---|---|---|---|
| ATTCCATGCATAGCCTTGGTTCTGCATCGCA | 12.38 | 2.66 | 9 | 0.11 | 4056403 | 4056430/-55G espA |
| ATTCCATGCATAGCCTTGGTTCTGCATTGCA | 8.39 | -1.72 | 4147 | 49.19 | 4056403 | 4056430/-55A, 4056416/-41G espA |

#### *narU*

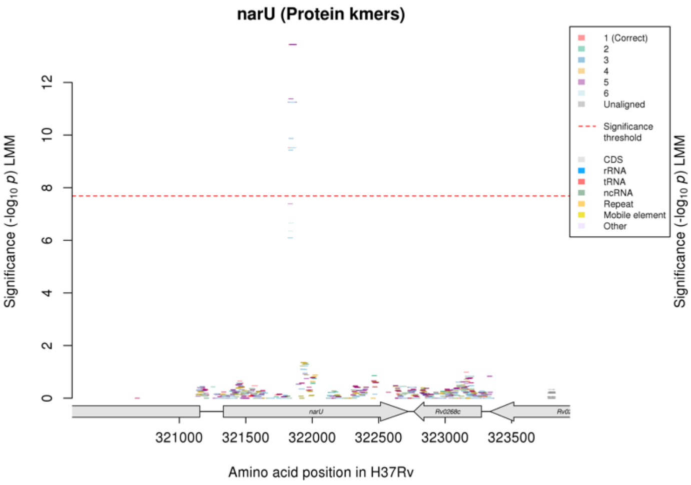

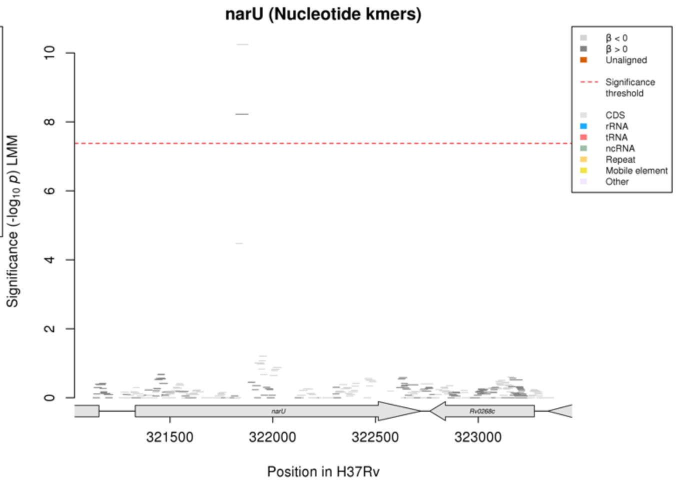

Protein kmers:

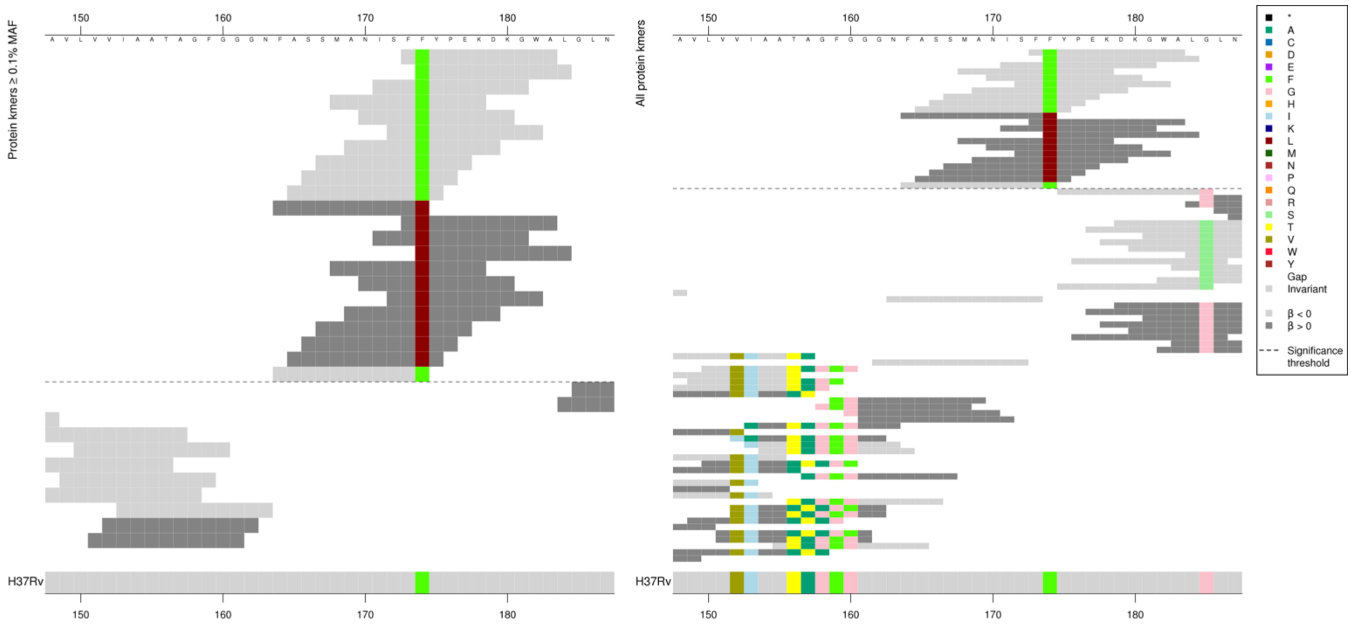

| Kmer | -log10 *p* | β | MAC | MAF | Ps | Variants |
|---|---|---|---|---|---|---|
| FFYPEKDKGWA | 13.44 | -2.85 | 42 | 0.50 | 173 | 174F |
| FASSMANISFL | 11.25 | 3.12 | 38 | 0.45 | 164 | 174L |

#### *rne*

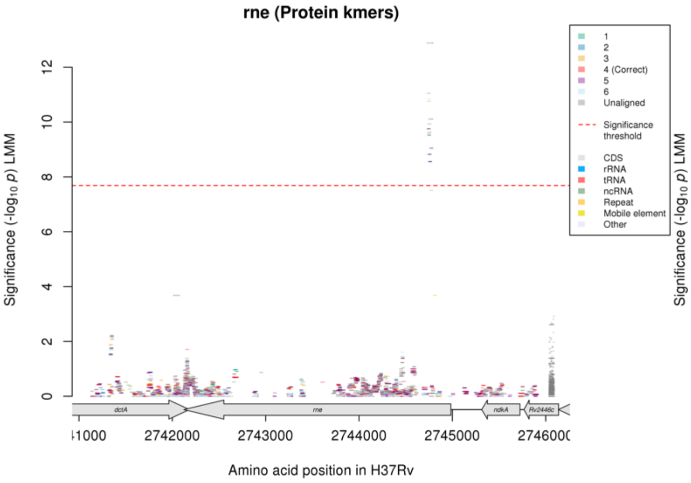

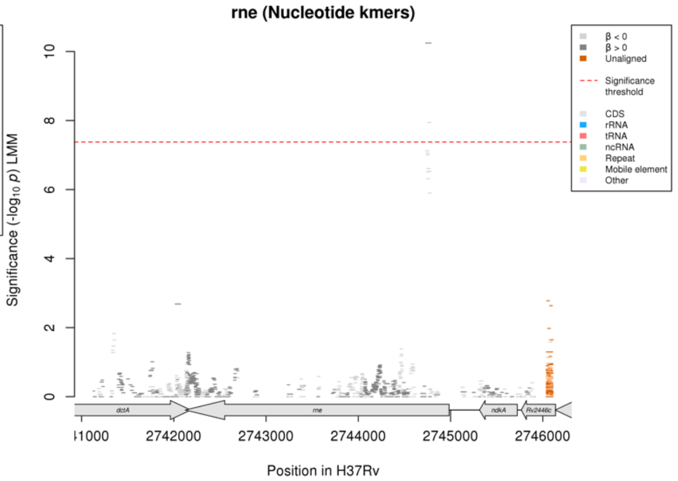

Nucleotide kmers:

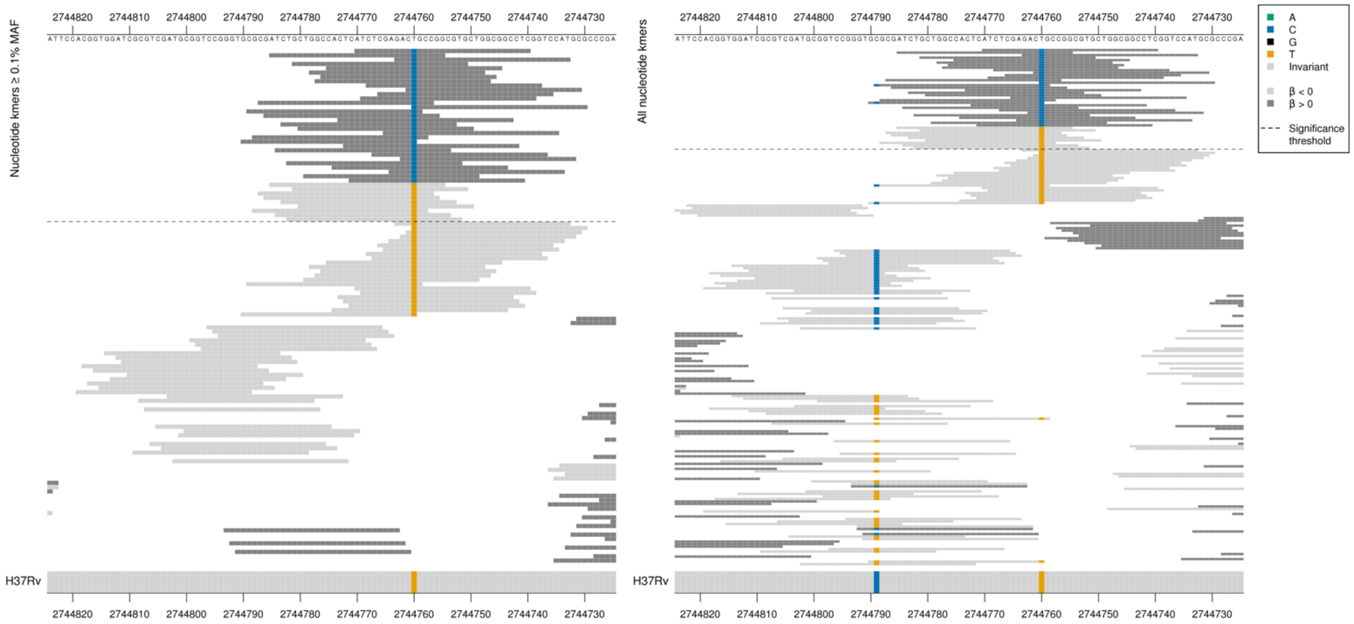

*rne* product = ribonuclease E

High MIC associated kmers = ACC = Thr / T
Low MIC associated kmers = ACT = Thr / T
Synonymous mutation at position 2744760/225

| Kmer | -log$_{10}$ $p$ | β | MAC | MAF | Ps | Variants |
|---|---|---|---|---|---|---|
| ATCTCGAGACCGCCGGCGTGCTGGCGGCCTC | 10.24 | 2.51 | 14 | 0.17 | 2744770 | 2744760/225C |
| ATCTGCTGGCCACTCATCTCGAGACTGCCGG | 7.94 | -1.85 | 22 | 0.26 | 2744785 | 2744760/225T |

## *Rv1393c*

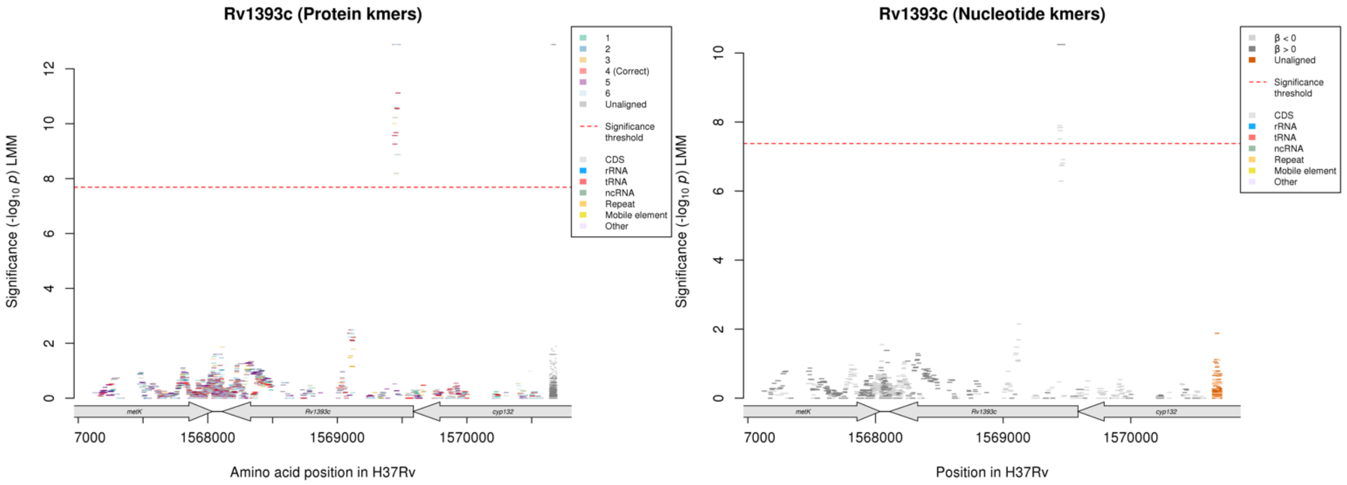

Protein kmers:

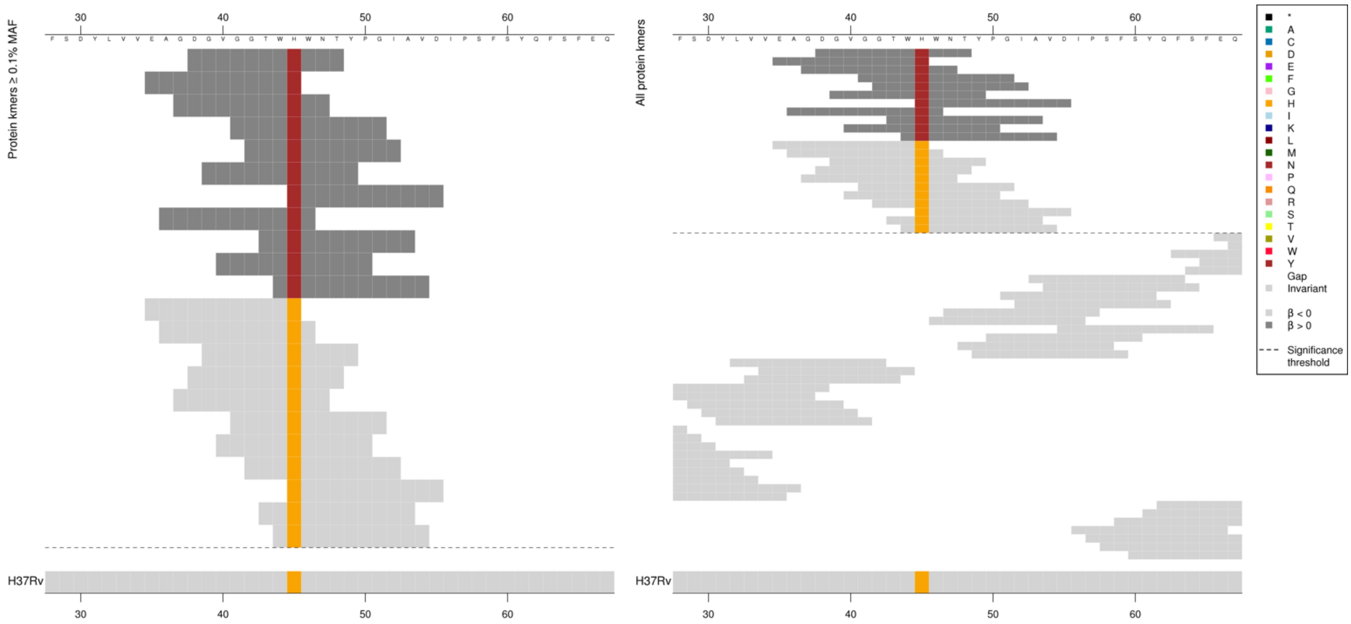

*Rv1393c* product = monoxygenase

The kmers captured a substitution at codon 45. Both the low MIC-associated H and the high MIC-associated N containing kmers were significantly associated.

| Kmer | $-\log_{10} p$ | β | MAC | MAF | Ps | Variants |
|---|---|---|---|---|---|---|
| DGVGGTWNWNT | 12.88 | 2.81 | 14 | 0.17 | 38 | 45N |
| EAGDGVGGTWH | 11.12 | -2.30 | 20 | 0.24 | 35 | 45H |

# *Rv1362c*

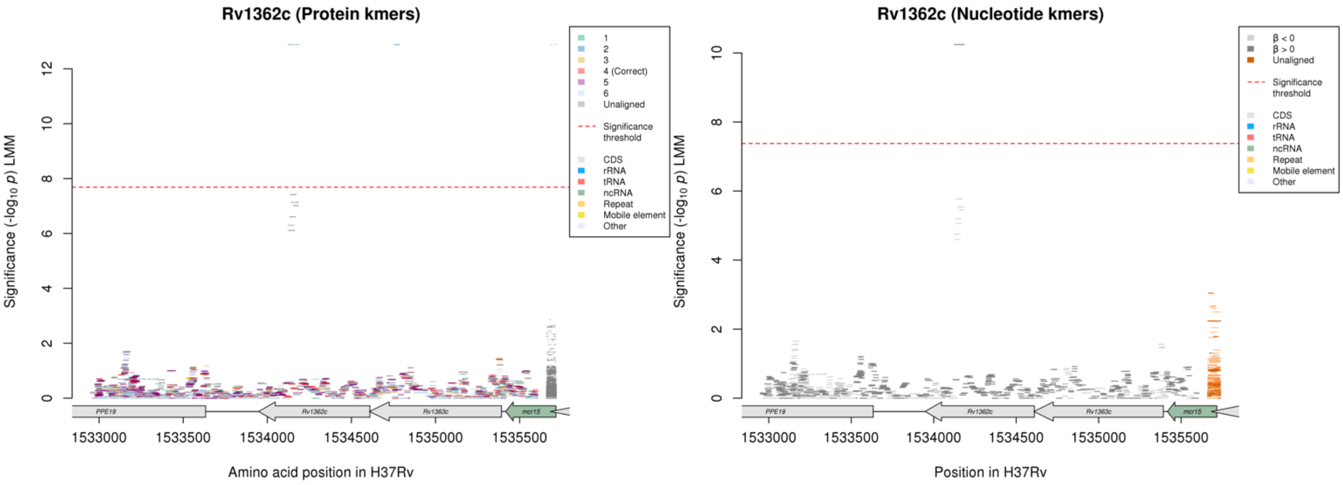

### Amikacin (AMI)

Nucleotide kmers:

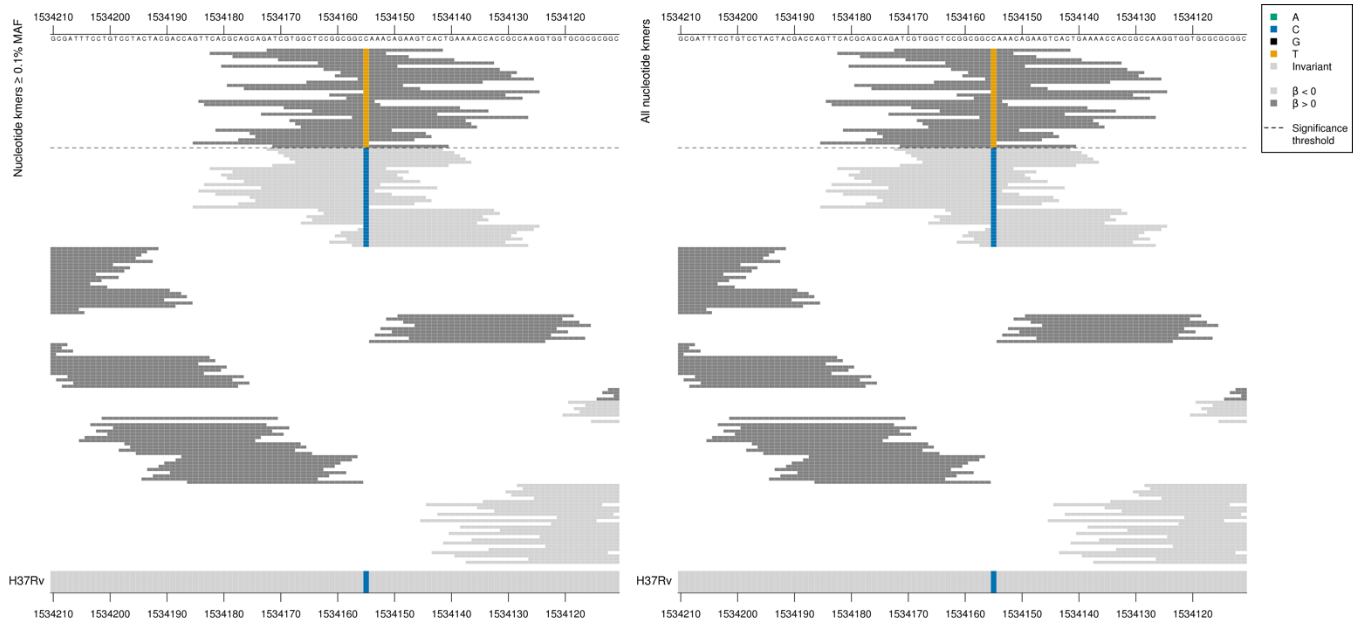

*Rv1362c* product = membrane protein

High MIC-associated kmers = GCT = Ala / A
Kmers below significance threshold = GCC = Ala / A
Synonymous mutation at position 1534155/456

| Kmer | $-\log_{10} p$ | β | MAC | MAF | Ps | Variants |
|---|---|---|---|---|---|---|
| ATCGTGGCTCCGGCGGCTAAACAGAAGTCAC | 10.24 | 2.51 | 14 | 0.17 | 1534172 | 1534155/456T |

## *Rv0579*

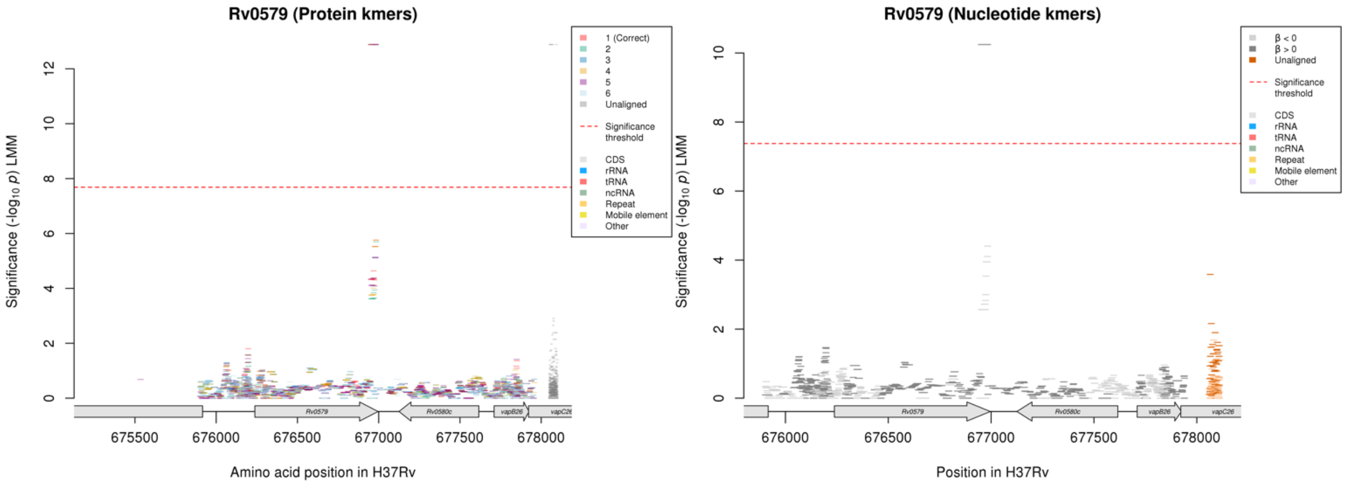

Protein kmers:

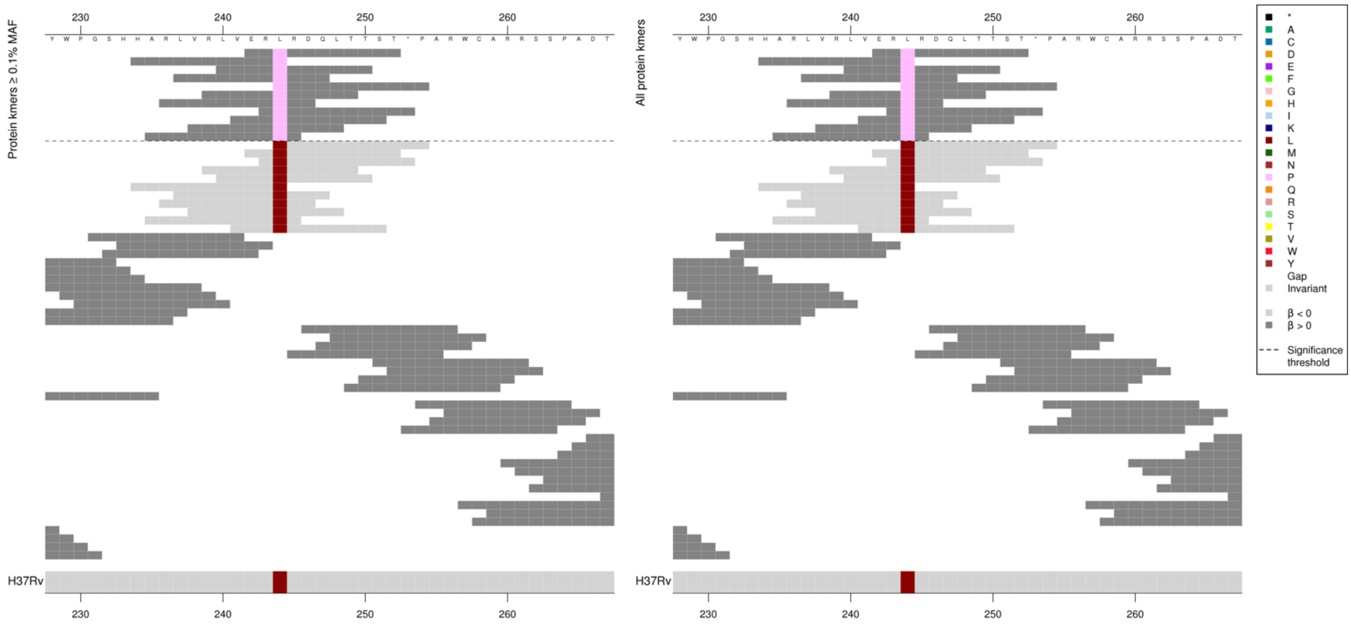

Stop codon is codon 253. The high MIC-associated 244P containing kmers were significantly associated, however the kmers containing 244L were not significant.

| Kmer | -log₁₀ *p* | β | MAC | MAF | Ps | Variants |
|---|---|---|---|---|---|---|
| ERPRDQLTTST | 12.88 | 2.81 | 14 | 0.17 | 242 | 244P |

#### *glnE*

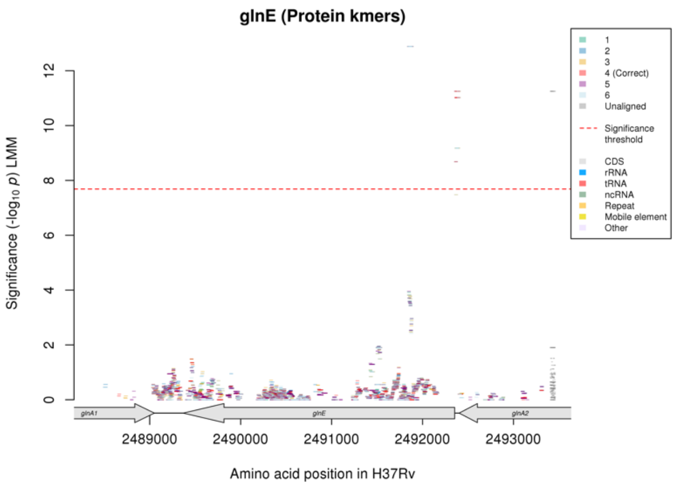

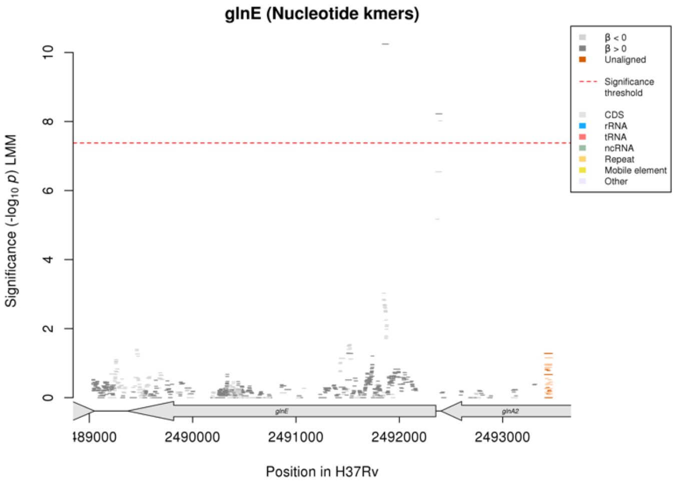

Nucleotide kmers:

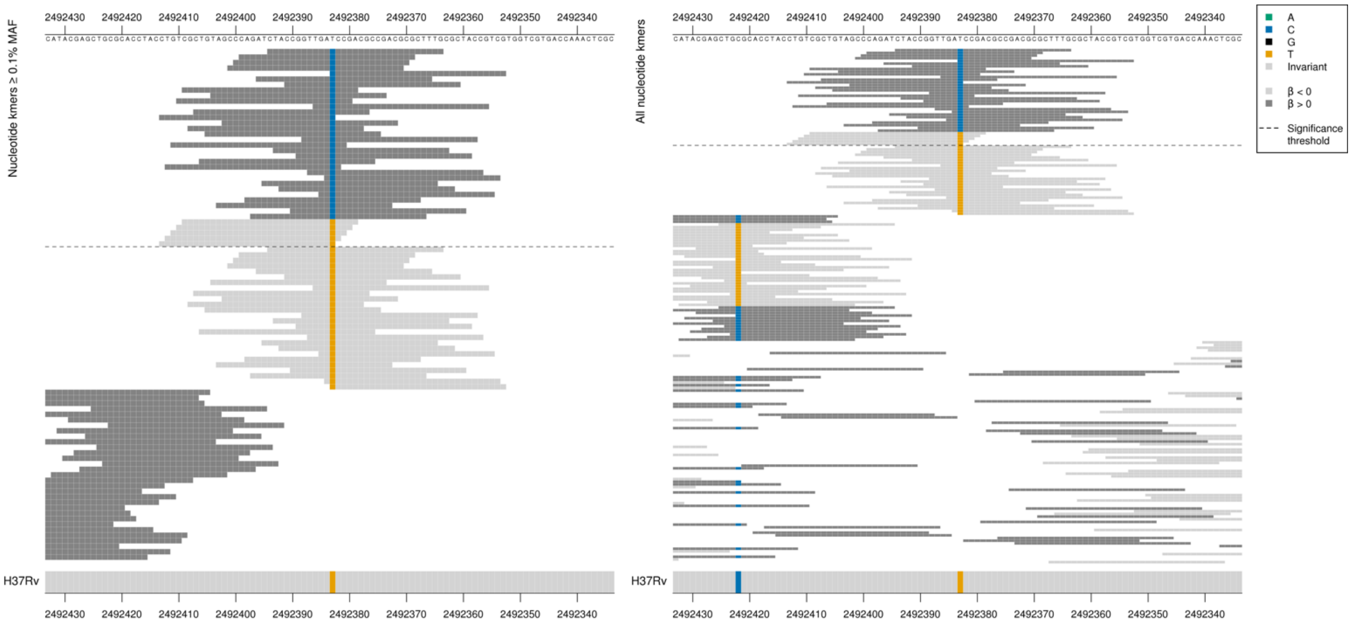

*glnE* is coded on the reverse strand, starting at position 2492353, *glnA2* ends position 2492402. Kmers just upstream of the start codon at position 2492383/-30 in the -35 region, changing the -35 from from the low MIC-associated sequence TTGA**T**C, to the high MIC-associated sequence TTGA**C**C.

| Kmer | $-\log_{10} p$ | β | MAC | MAF | Ps | Variants |
|---|---|---|---|---|---|---|
| CTACCGGTTGACCCGACGCCGACGCGCTTTG | 8.22 | 2.75 | 38 | 0.45 | 2492394 | 2492383/-30 C |
| CGCTGTAGCCCAGATCTACCGGTTGATCCGA | 8.02 | -2.67 | 39 | 0.46 | 2492409 | 2492383/-30 T |

Nucleotide kmers:

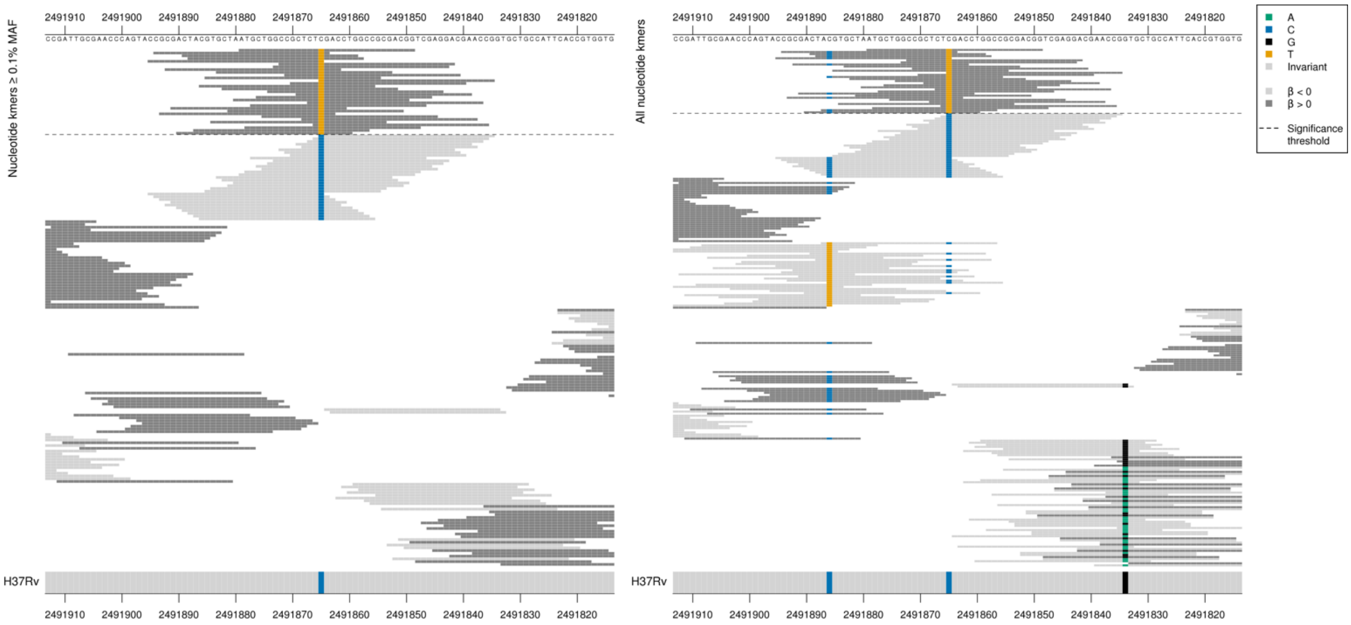

High MIC associated = CTT = Leu / L
Kmers below significance threshold = CTC = Leu / L
Synonymous mutation at position 2491865/489
The top kmer does not align to the other linked genes.

| Kmer | $-\log_{10} p$ | β | MAC | MAF | Ps | Variants |
|---|---|---|---|---|---|---|
| ATGCTGGCCGCTCTTGACCTGGCCGCGACGG | 10.24 | 2.51 | 14 | 0.17 | 2491879 | 2491865/489 T |

### *ethA*

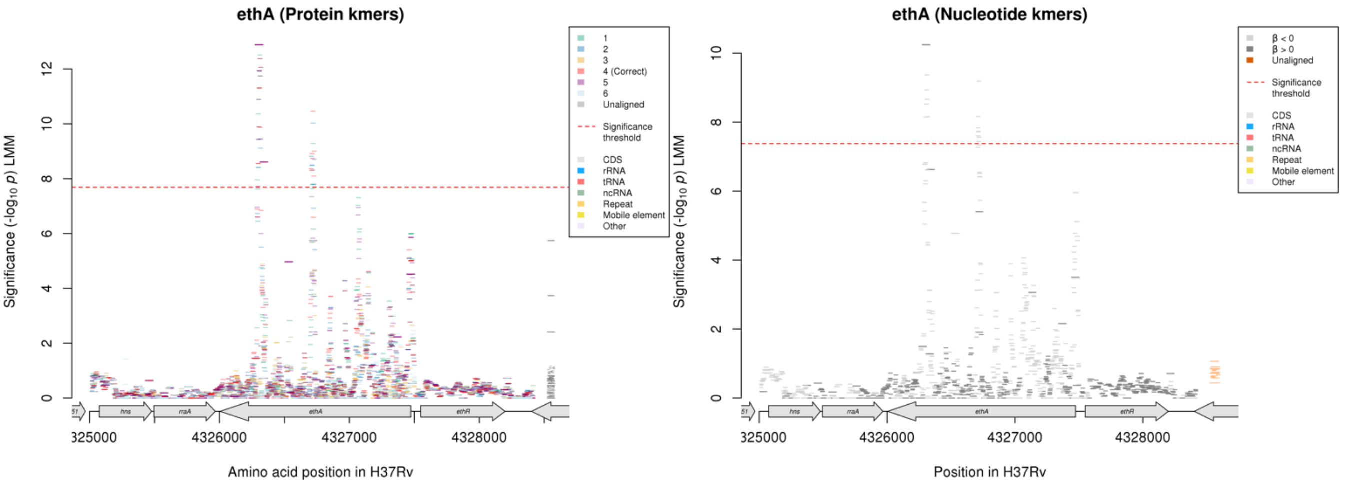

Protein kmers:

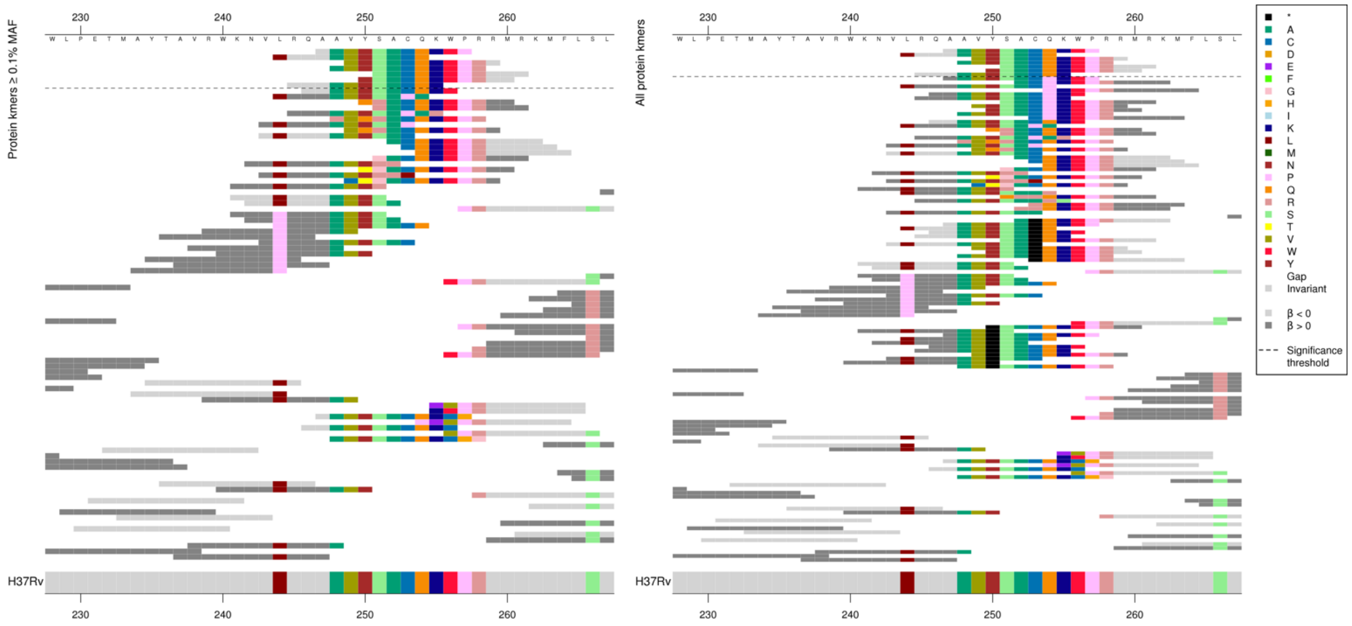

First alignment shows low MIC-associated kmers which were identical to the reference. The top high MIC (although not significant) kmers captured multiple amino acid changes.

The same region for the nucleotide kmer analysis to see the variation in the kmers that were not significant:

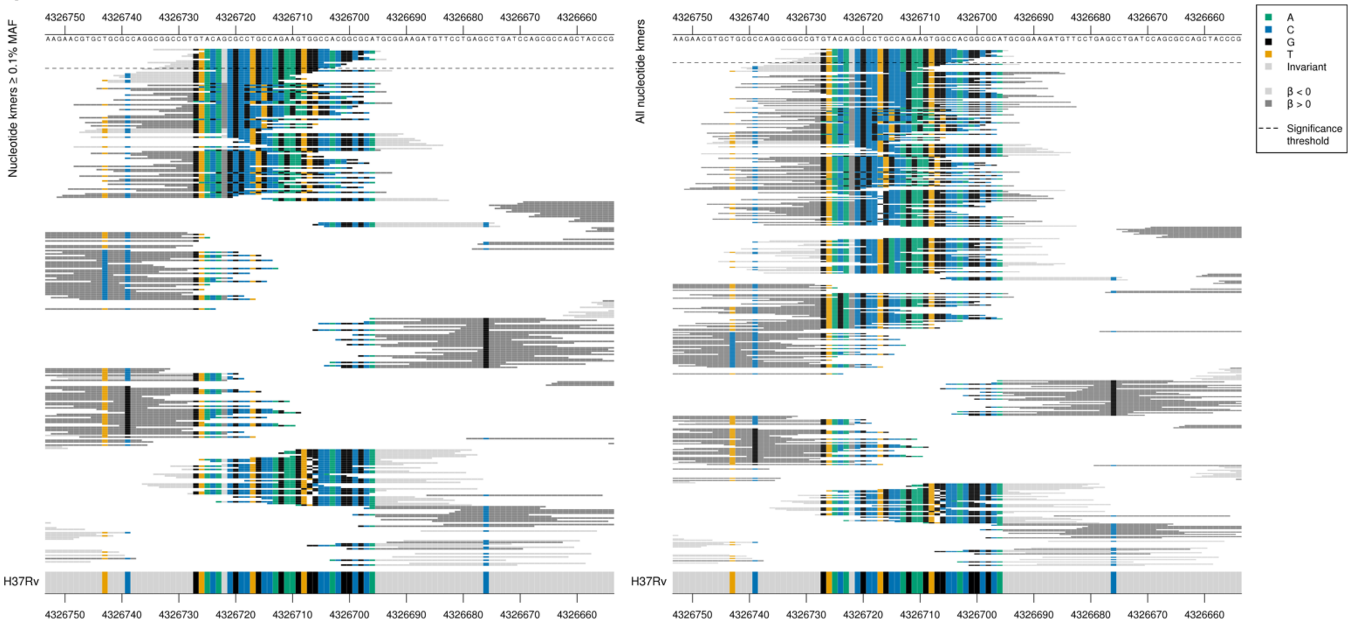

Mutation and a deletion in kmers that were not significant.

| Kmer | $-\log_{10} p$ | $\beta$ | MAC | MAF | Ps | Variants |
|---|---|---|---|---|---|---|
| AAVYSACQKWP | 10.46 | -0.78 | 108 | 1.28 | 247 | WT |

Protein kmers:

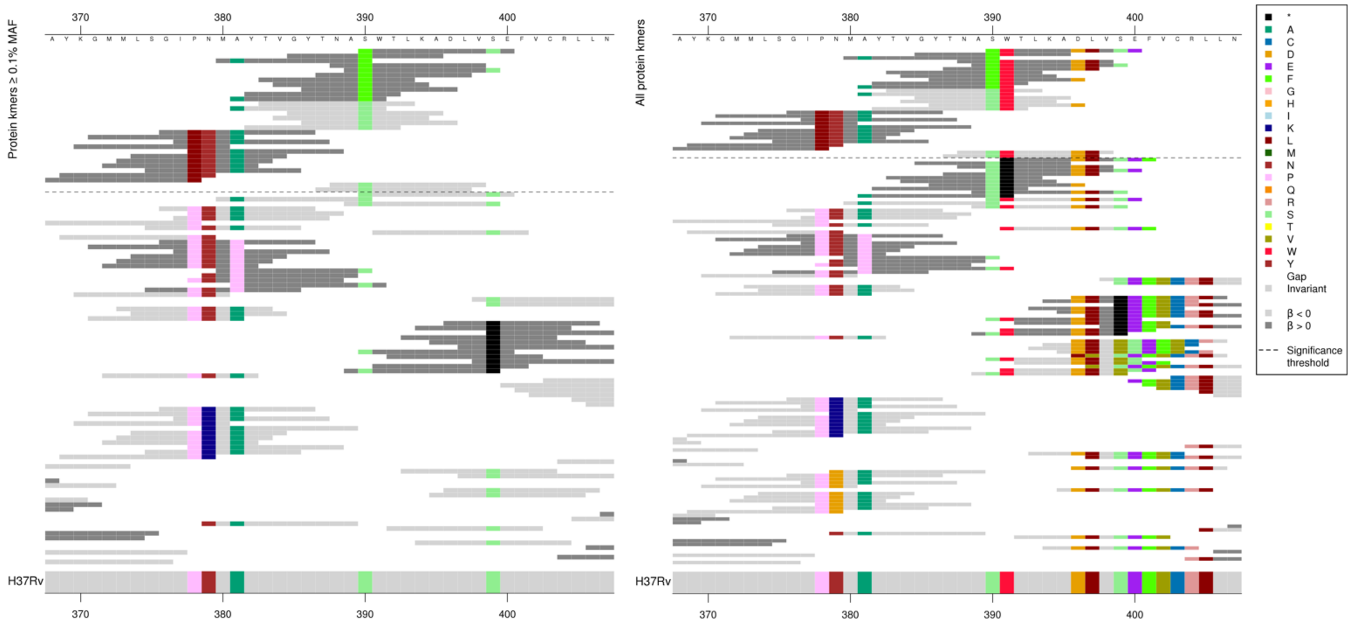

The second alignment mainly captured two significant substitutions. The first is at codon 378, where the L containing high MIC-associated kmers were significantly associated. The second substitution is at codon 390, where kmers containing the low MIC-associated S and high MIC-associated F were both significantly associated. For each substitution the *p*-values varied, so it is likely that the surrounding substitutions influence the significance of the kmers.

| Kmer | $-\log_{10} p$ | $\beta$ | MAC | MAF | Ps | Variants |
|---|---|---|---|---|---|---|
| FWTLKADLVSE | 12.88 | 2.81 | 14 | 0.17 | 390 | 390F |
| TVGYTNASWTL | 12.38 | -1.31 | 56 | 0.66 | 383 | 390S |
| AYTVGYTNASW | 12.06 | -1.42 | 216 | 2.56 | 381 | 381A, 390S, 391W |
| GYTNASWTLKA | 11.31 | -1.22 | 58 | 0.69 | 385 | 390S, 391W |
| YTNASWTLKAD | 9.89 | -1.10 | 60 | 0.71 | 386 | 390S, 391W, 396D |
| GILNMAYTVGY | 8.61 | 1.85 | 44 | 0.52 | 376 | 378L |
| NASWTLKADLV | 8.55 | -1.02 | 64 | 0.76 | 388 | 390S, 391W, 396D, 397L |

# *Rv0208c:Rv0209*

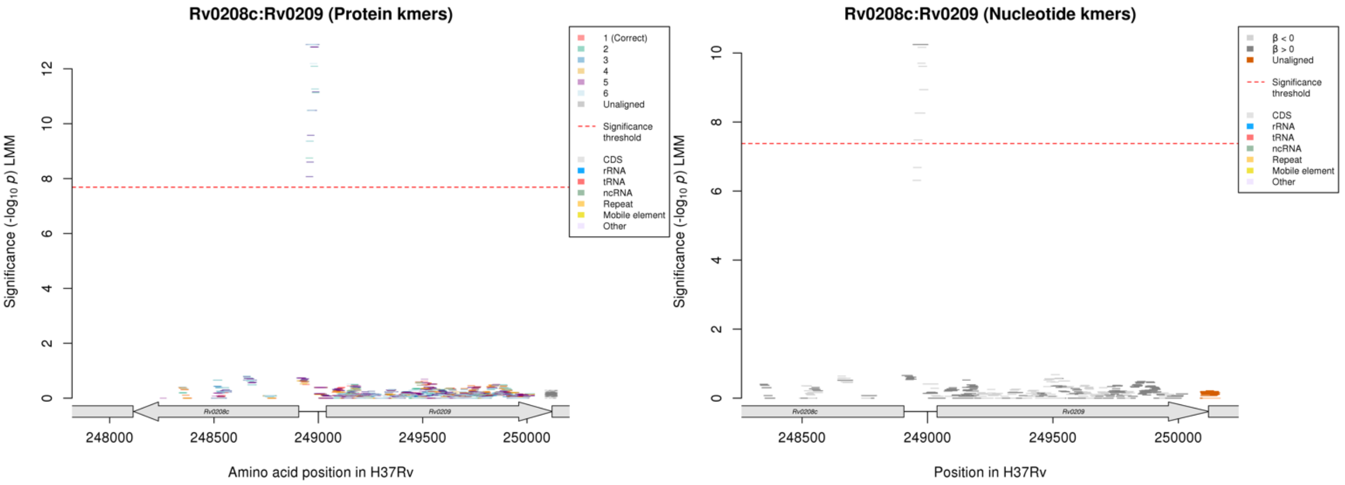

Nucleotide kmers:

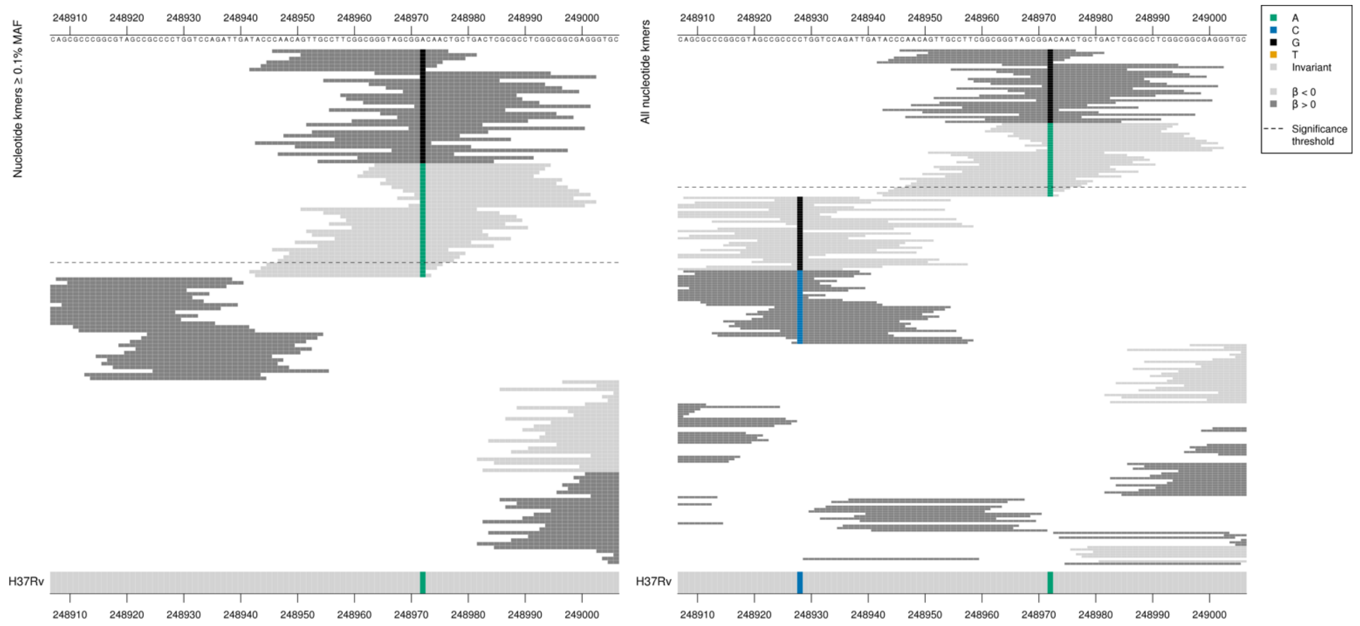

*Rv0208c* product = tRNA (guanine-N(7)-)-methyltransferase
*Rv0209* product = HP

Gene *Rv0208c* starts at position 248906 on the reverse strand and *Rv0209* starts at position 249038 on the forward strand. The kmers captured a mutation in the intergenic region, position 248972, 67bp from both genes.
Significant kmers on the forward strand:
High MIC-associated kmers = G
Low MIC-associated kmers = A

| Kmer | -log₁₀ *p* | β | MAC | MAF | Ps | Variants |
|---|---|---|---|---|---|---|
| CAACAGTTGCCTTCGGCGGGTAGCGGGCAAC | 10.24 | 2.51 | 14 | 0.17 | 248946 | 248972 (67bp from both genes) G (forward strand) |
| GGTAGCGGACAACTGCTGACTCGCGCCTCGG | 10.16 | -2.49 | 15 | 0.18 | 248964 | 248972 (67bp from both genes) A (forward strand) |

### Bedaquiline (BDQ)

## *Rv0678*

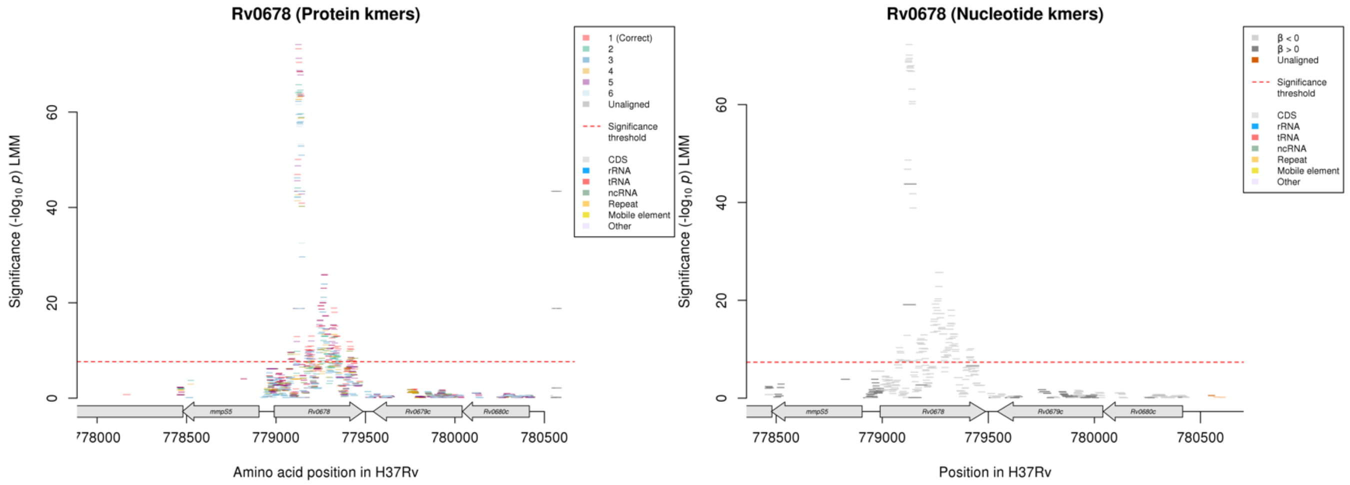

Protein kmers:

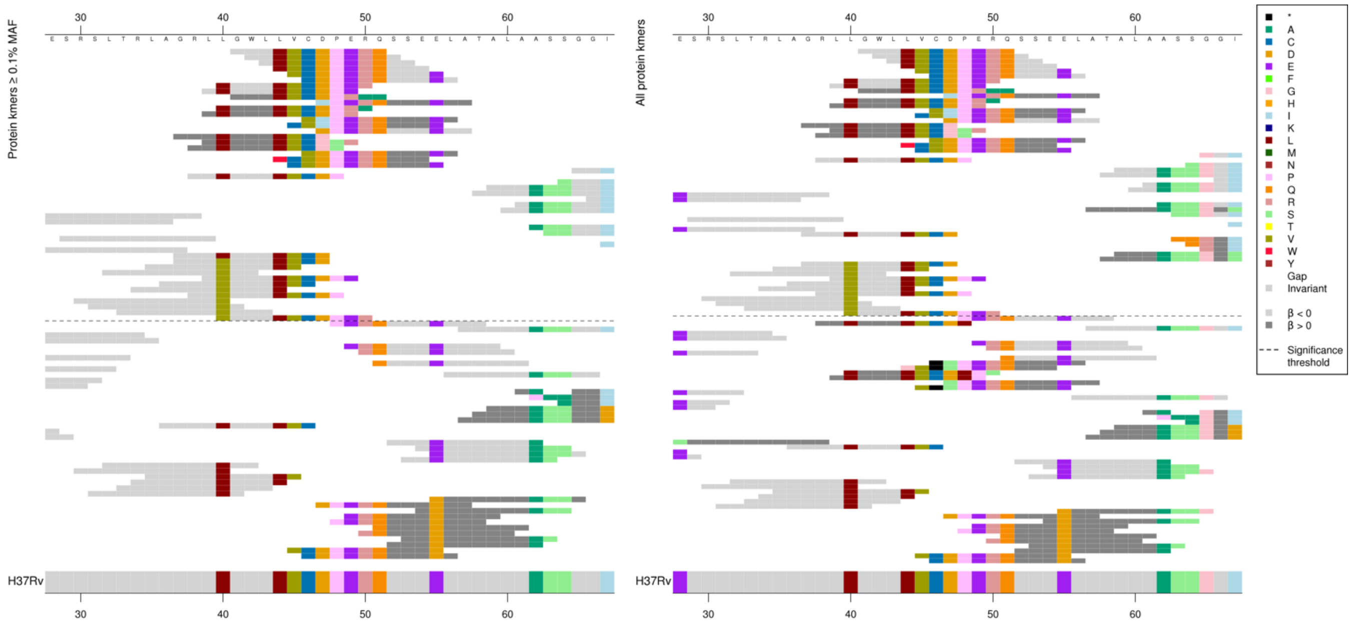

The following nucleotide alignments are showing the same region as the amino acid alignments above:

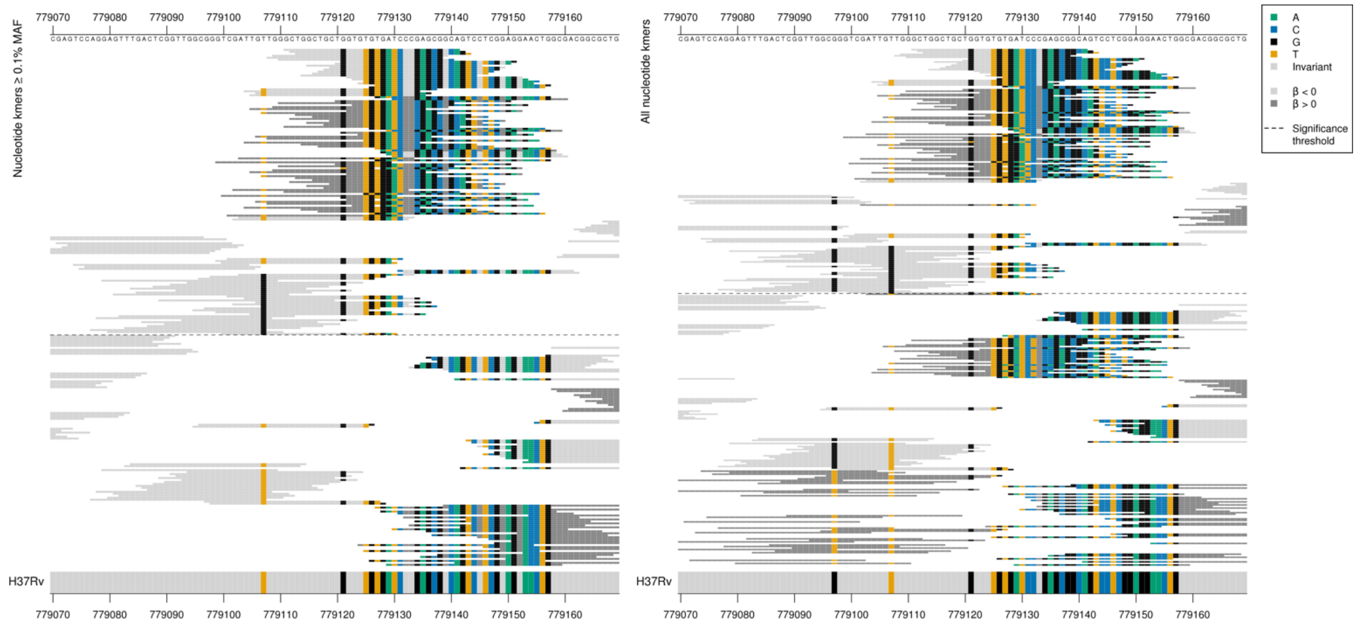

*Rv0678* begins at position 778990. These show that the first group of high MIC-associated kmers contained a C insertion around positions 779131-779133. Other high MIC-associated kmers contained a G insertion following the G at position 779128. Low MIC-associated kmers were a range of *p*-values depending on the positions they covered, so including the top kmer per group of low MIC-associated kmers in the table. All high MIC-associated kmers that captured the C insertion were the same *p*-value, as were the high MIC-associated kmers capturing the G insertion.
Low MAF kmers = 779132/143T.

| Kmer | $-\log_{10} p$ | β | MAC | MAF | Ps | Variants |
|---|---|---|---|---|---|---|
| GCTGGCTGCTGGTGTGTGATCCCGAGCGGCA | 72.27 | -1.92 | 72 | 0.90 | 779111 | WT between positions 779104-779157/115-168 |
| GATCCCCGAGCGGCAGTCCTCGGAGGAACTG | 43.77 | 1.94 | 40 | 0.50 | 779127 | C insertion in 'CCC' positions 779131-779133/142-144 |
| GATCCCGAGCGGCAGTCCTCGGAGGAACTGG | 41.79 | -1.64 | 60 | 0.75 | 779128 | WT between positions 779128-779160/139-171 |
| ATTGTTGGGCTGGCTGCTGGTGTGTGGATCC | 19.11 | 2.07 | 14 | 0.17 | 779103 | G insertion between 779127-779129/138-140 |
| ATTGTTGGGCTGGCTGCTGGTGTGTGATCCC | 12.83 | -1.02 | 44 | 0.55 | 779103 | WT between positions 779102-779133/113-144 |
| ATTTCGAGTCCAGGAGTTTGACTCGGTTGGC | 10.31 | -1.63 | 13 | 0.16 | 779066 | WT between positions 779066-779103/77-114 |
| TCGAGTCCAGGAGTTTGACTCGGTTGGCGGG | 7.85 | -1.35 | 15 | 0.19 | 779069 | WT between positions 779067-779162/78-173 |
| ATTGGTGGGCTGGCTGCTGGTGTGTGATCCC | 7.52 | -1.61 | 11 | 0.14 | 779103 | 779107/118G |

Protein kmers:

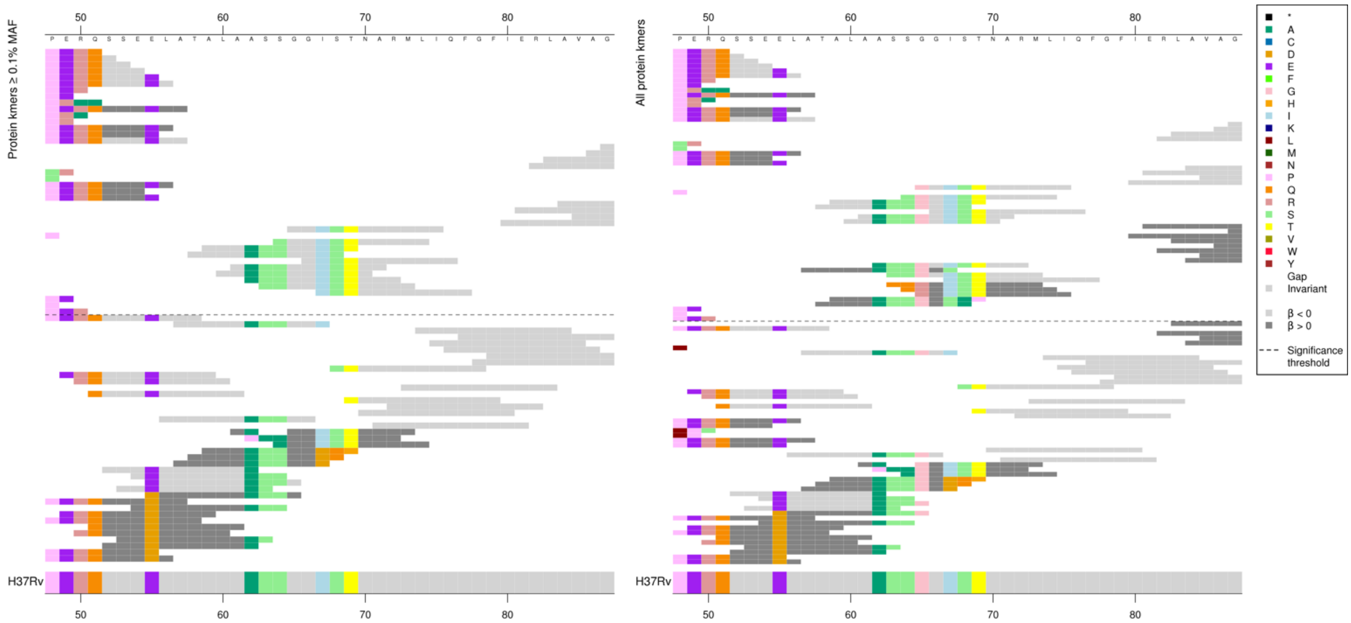

The same region for the nucleotide kmer analysis:

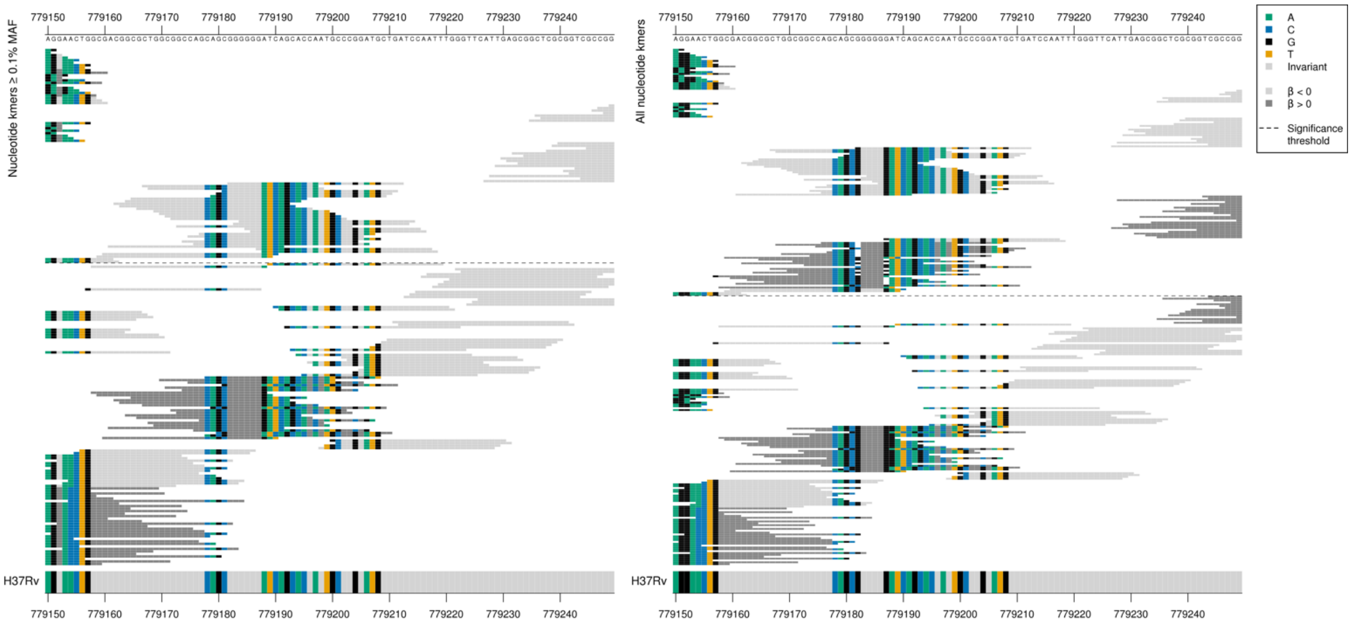

These kmers in *Rv0678* were not significant for the CFZ analysis.

Low MAF kmers = G deletion between positions 779182-779187/193-198, below the 0.1% MAF cutoff.

| Kmer | -log10 *p* | β | MAC | MAF | Ps | Variants |
|---|---|---|---|---|---|---|
| GGGGGGATCAGCACCAATGCCCGGATGCTGA | 12.93 | -1.37 | 25 | 0.31 | 779182 | WT between positions 779161-779218/172-229 |
| GACGGCGCTGGCGGCCAGCAGCGGGGGGATC | 7.81 | -0.97 | 49 | 0.61 | 779160 | WT between 779159-779190/170-201 |

Protein kmers:

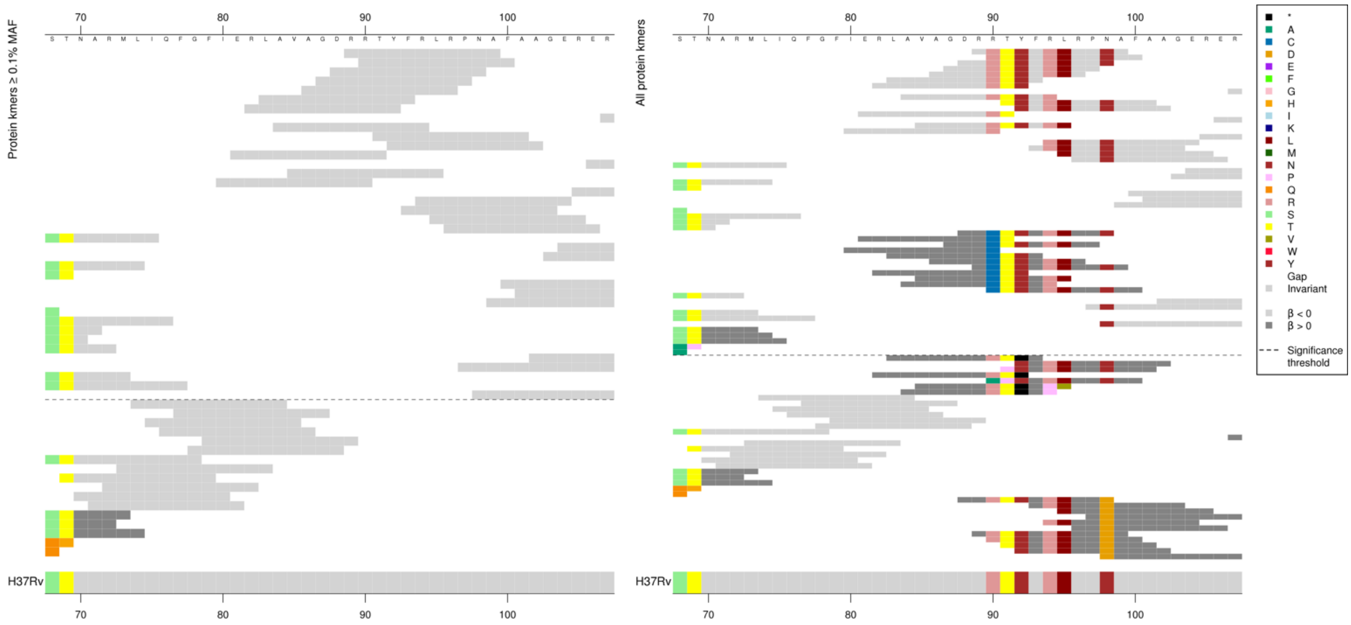

The same region for the nucleotide kmer analysis:

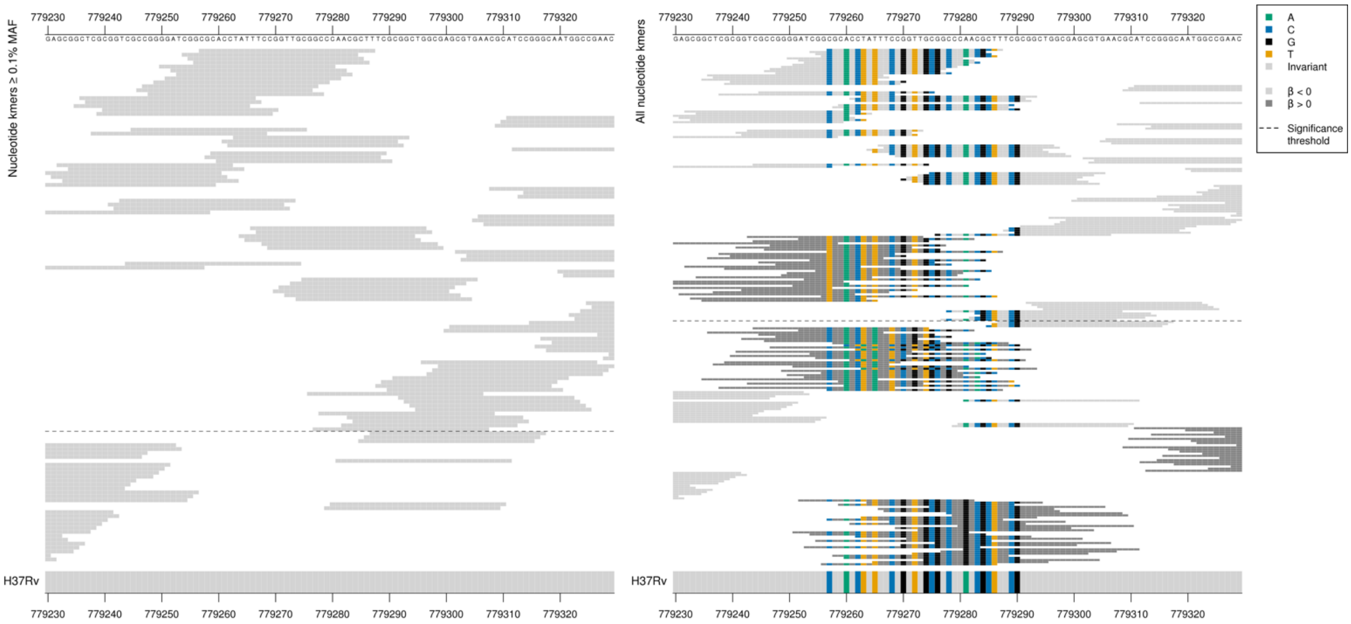

The top low MIC-associated kmers captured the reference sequence, including codons 90R and 98N which were the same as the reference, the other reference sequence kmers covered the same region over a range of *p*-values but not all covered both codons. Just reporting the top kmer in the table.
Low MAF kmers = 90C.

Kmers below significance threshold = 98D
Kmers below significance threshold = Indels

| Kmer | -log$_{10}$ *p* | β | MAC | MAF | Ps | Variants |
|---|---|---|---|---|---|---|
| RRTYFRLRPNA | 25.87 | -1.89 | 28 | 0.35 | 89 | WT / 90R, 98N |

Protein kmers:

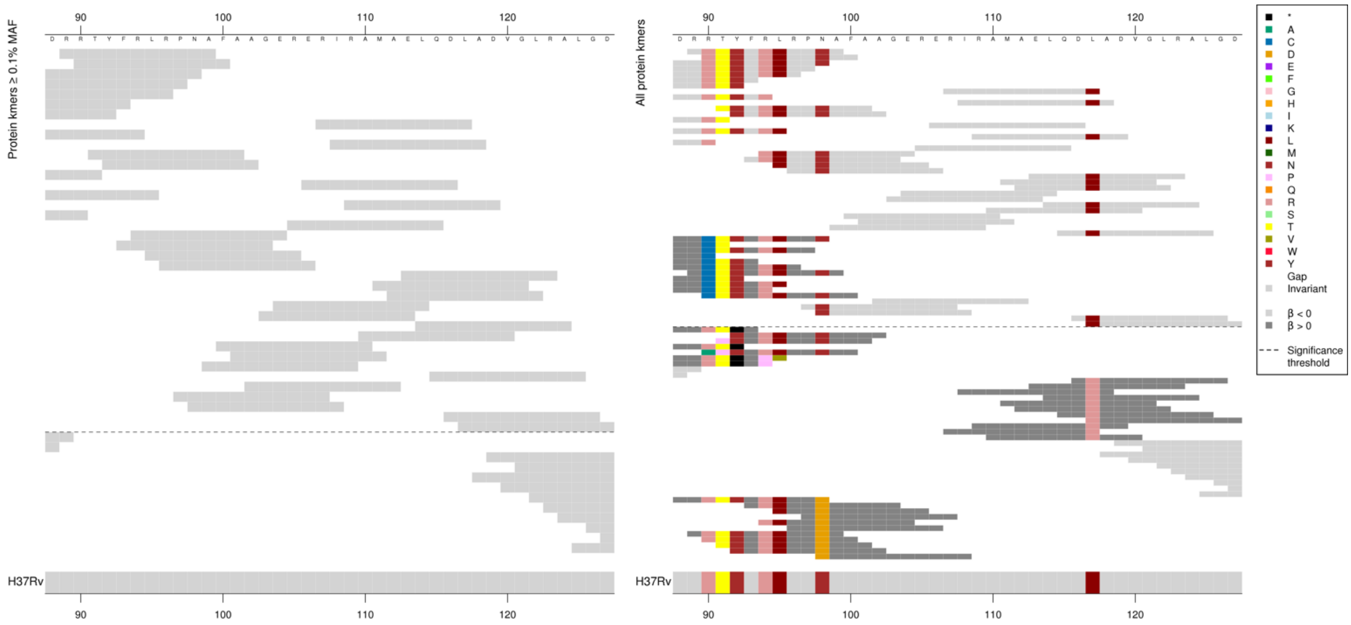

Kmers below significance threshold = 117R
There were also some significant low MIC-associated kmers in the region which were WT and do not cover codon 117, these were possibly in LD or capturing a region with very rare variation not analysed here.

| Kmer | -log$_{10}$ *p* | β | MAC | MAF | Ps | Variants |
|---|---|---|---|---|---|---|
| RLAVAGDRRTY | 19.37 | -1.91 | 22 | 0.27 | 82 | WT / 117L |

Protein kmers:

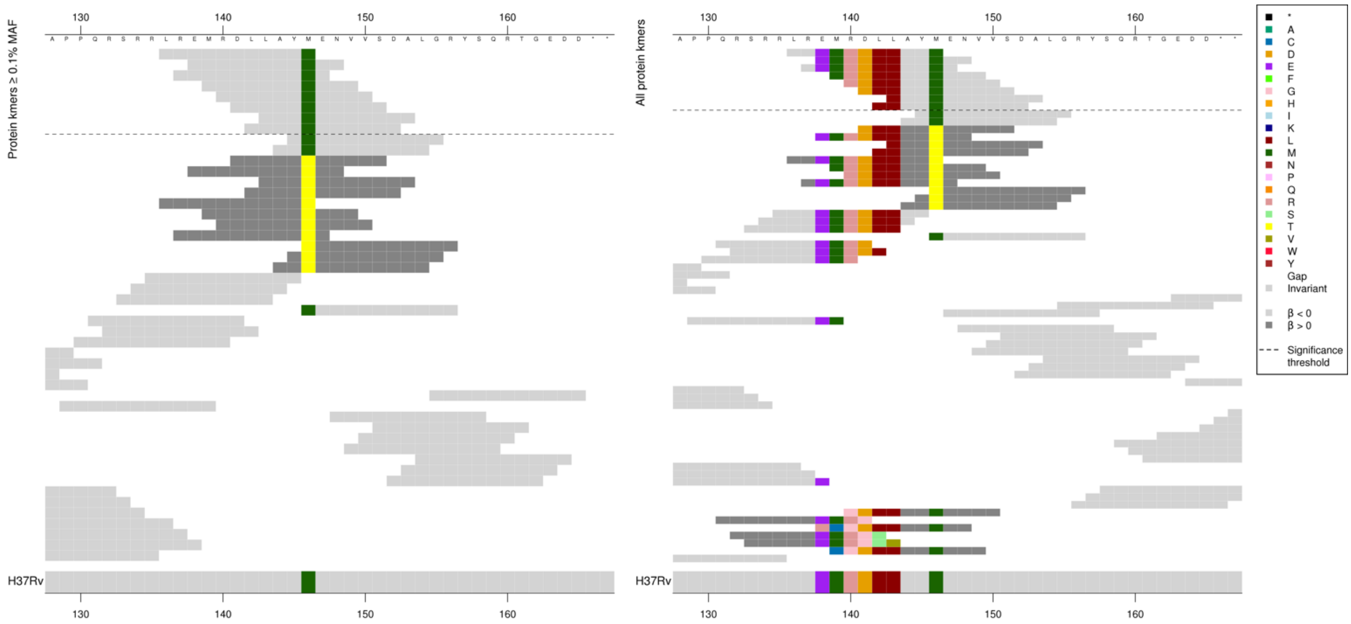

The same region for the nucleotide kmer analysis:

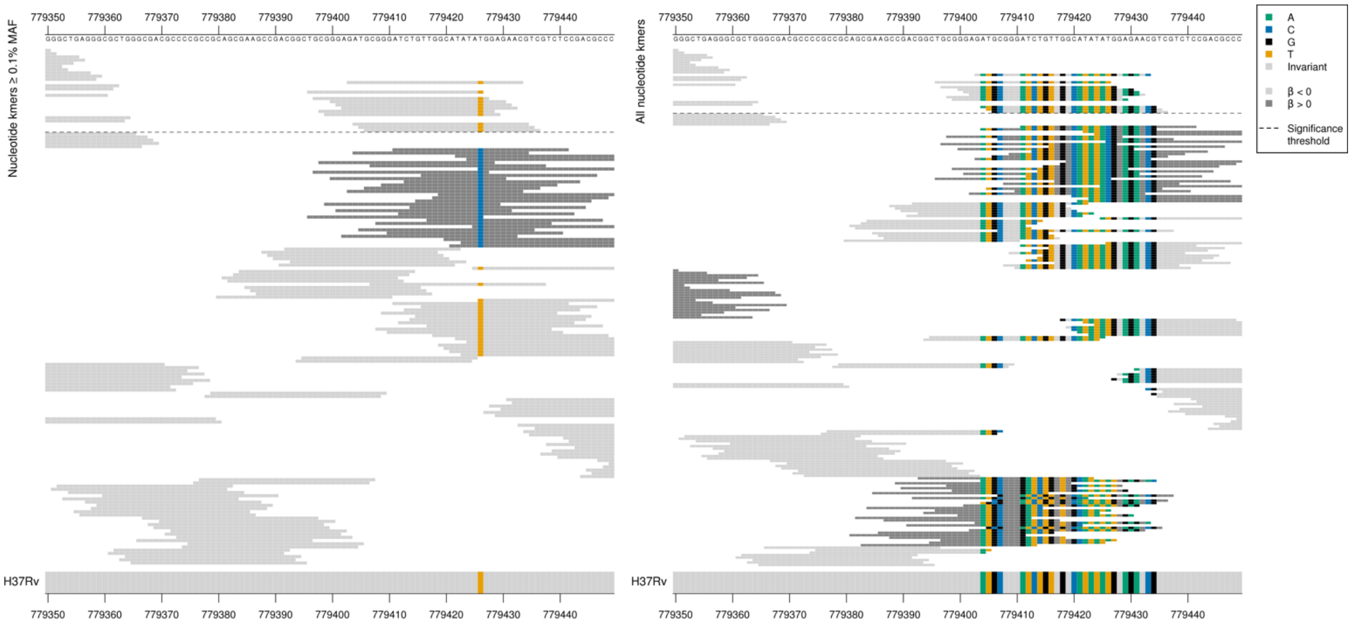

The kmers in this region were not significant for the CFZ analysis.
All five low MIC-associated kmers were different *p*-values, just showing the top kmer to represent capturing codon 1476M. It is likely that the low MIC-associated kmers were also capturing the reference codons across the region where low MAF not significant kmers contained indels.

Kmers below significance threshold = codon 146T
Kmers below significance threshold (below 0.1% MAF) = indels

| Kmer | $-\log_{10} p$ | β | MAC | MAF | Ps | Variants |
|---|---|---|---|---|---|---|
| LREMRDLLAYM | 11.84 | -1.45 | 26 | 0.32 | 136 | WT / 146M |

#### *rpoB*

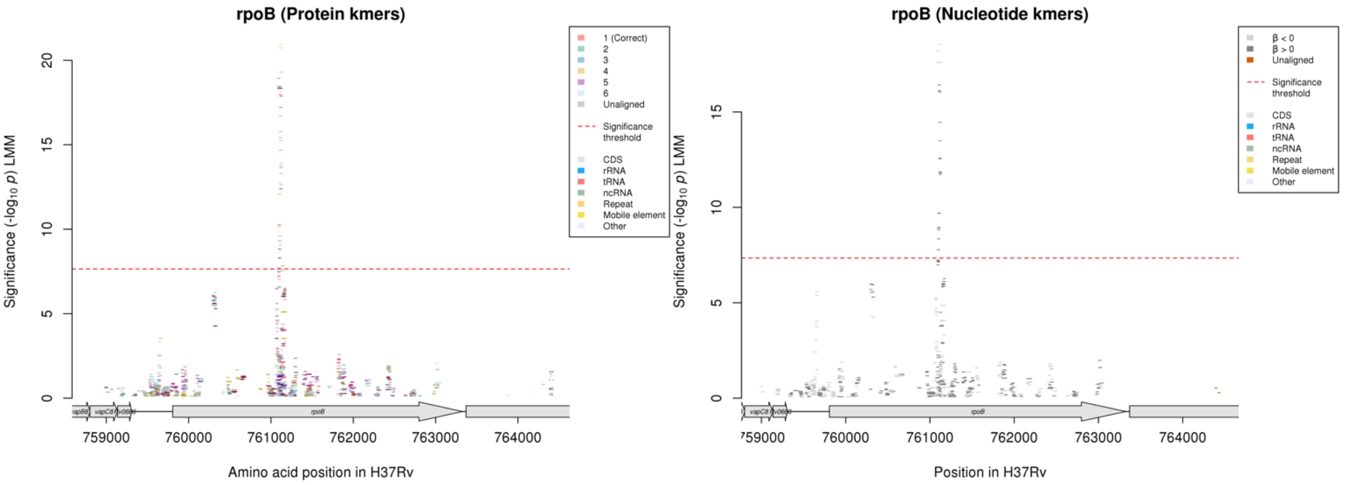

Protein kmers:

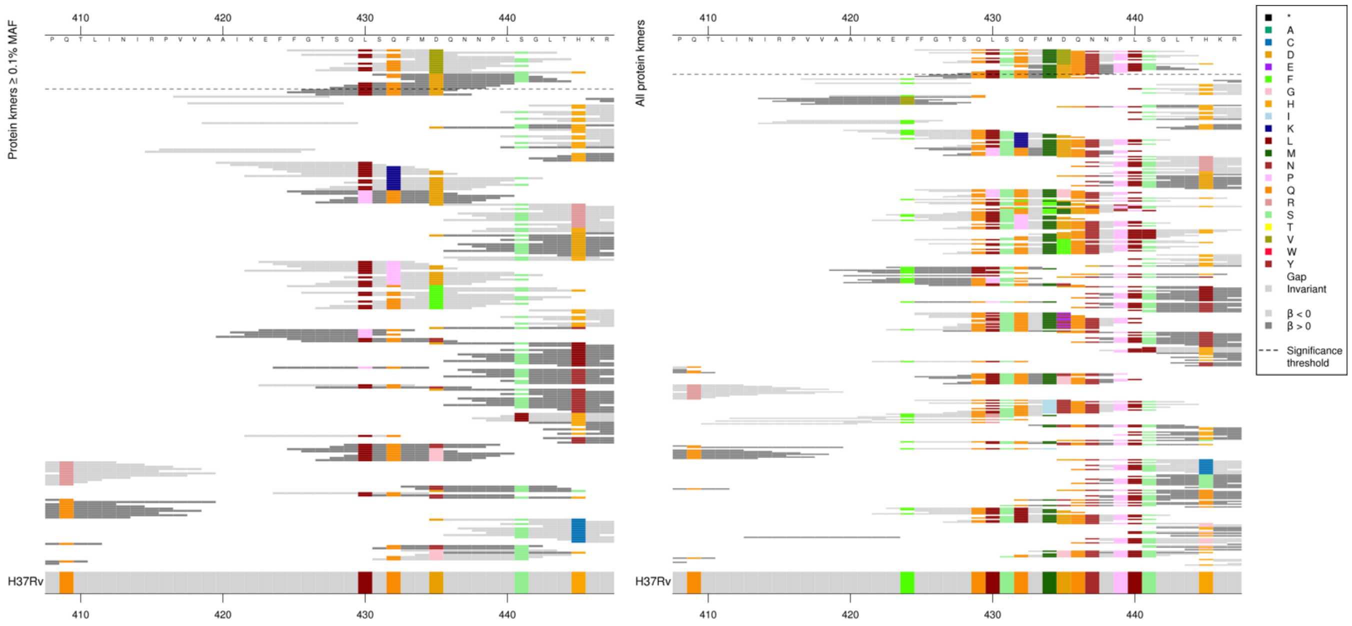

This substitution was significant for RIF and RFB, however for BDQ the substitutions had an opposite direction of effect.

The low and high MIC-associated kmer *p*-values varied depending on which other codons were covered, but for this table just reporting the most significant kmer capturing 435 for each.

| Kmer | $-\log_{10} p$ | β | MAC | MAF | Ps | Variants |
|---|---|---|---|---|---|---|
| FGTSQLSQFMV | 18.34 | -0.59 | 410 | 5.12 | 425 | 435V |
| SQFMDQNNPLS | 16.69 | 0.42 | 624 | 7.79 | 431 | 435D |

*rrs*

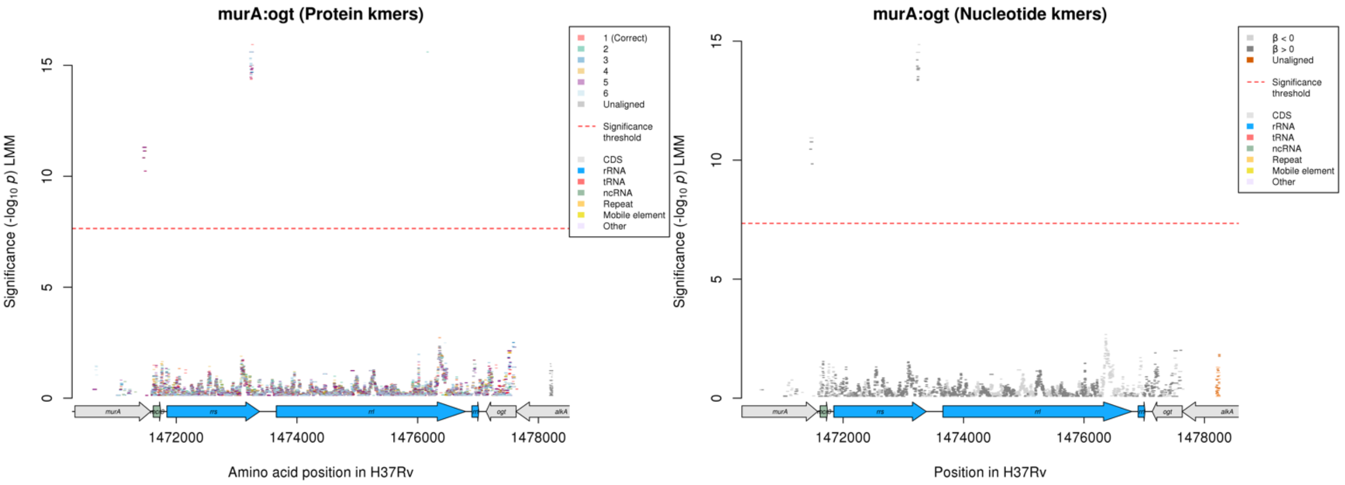

Nucleotide kmers:

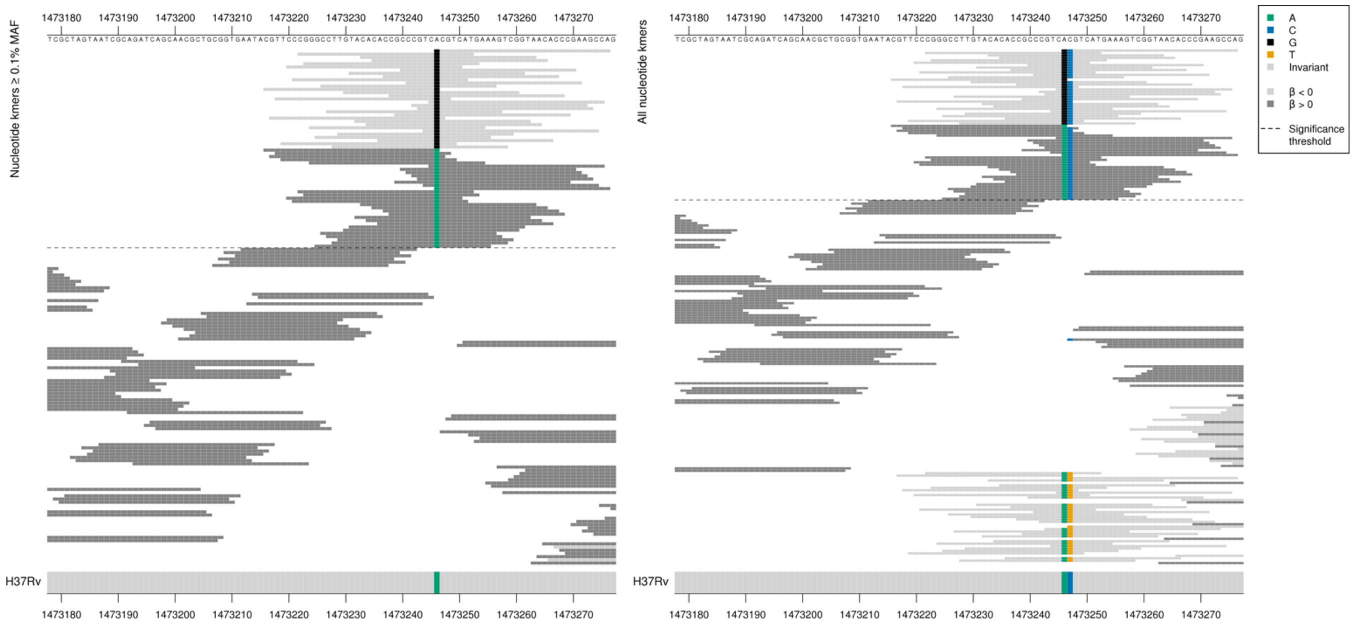

This mutation was significant for AMI and KAN, however for BDQ there was an opposite direction of effect for the significant kmers at position 1401. For AMI and KAN, the G was associated with high MIC. Here for BDQ, the A was associated with high MICs.

| Kmer | $-\log_{10} p$ | β | MAC | MAF | Ps | Variants |
|---|---|---|---|---|---|---|
| GCGTCATGAAAGTCGGTAACACCCGAAGCCA | 14.86 | -0.44 | 484 | 6.04 | 1473246 | 1473246/1401G |
| CGTTCCCGGGCCTTGTACACACCGCCCGTCA | 14.21 | 0.42 | 519 | 6.48 | 1473216 | 1473246/1401A |

#### *atpE*

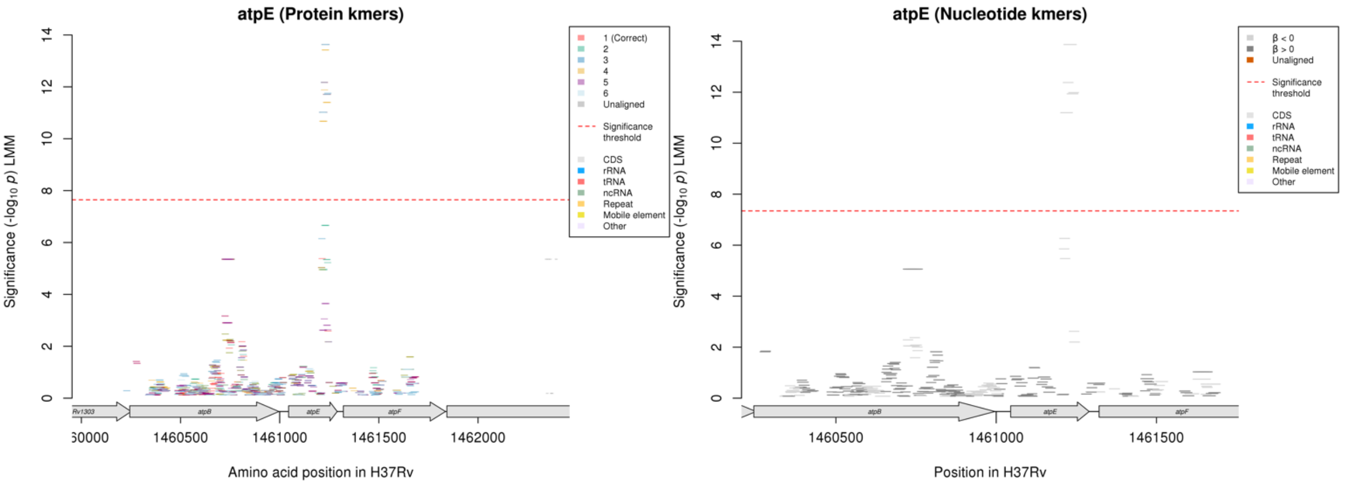

Protein kmers:

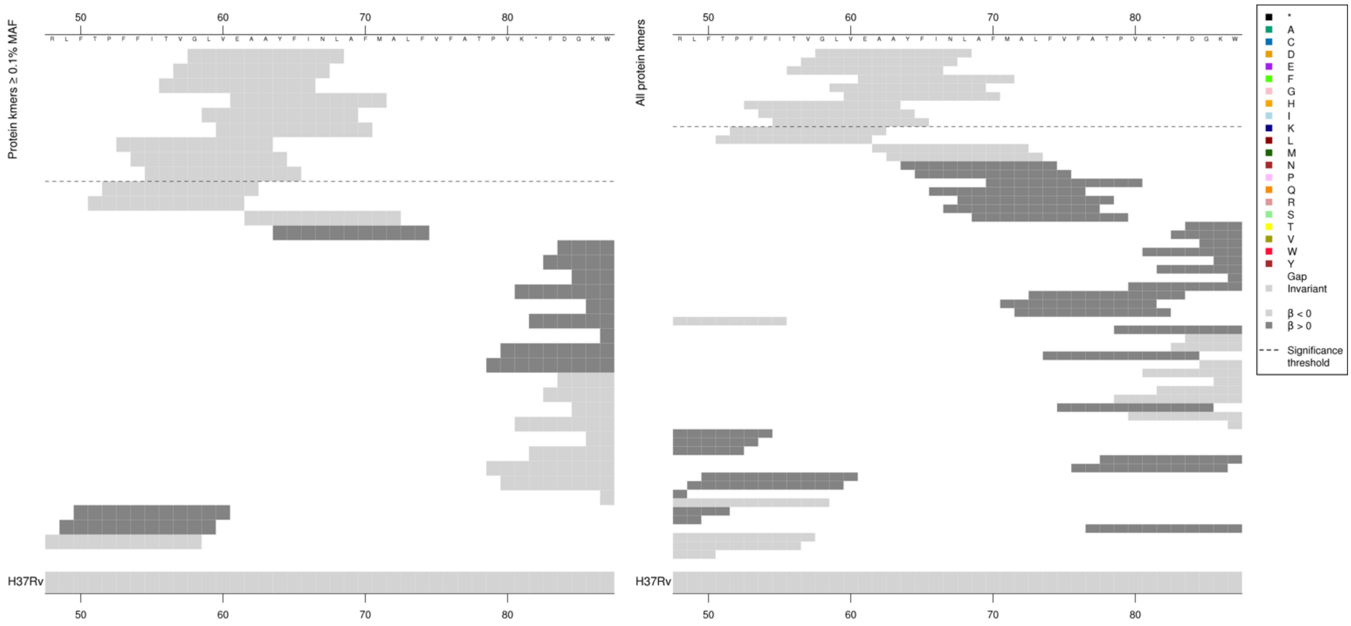

The same region for the nucleotide kmers, to check if the protein kmers were capturing a synonymous mutation:

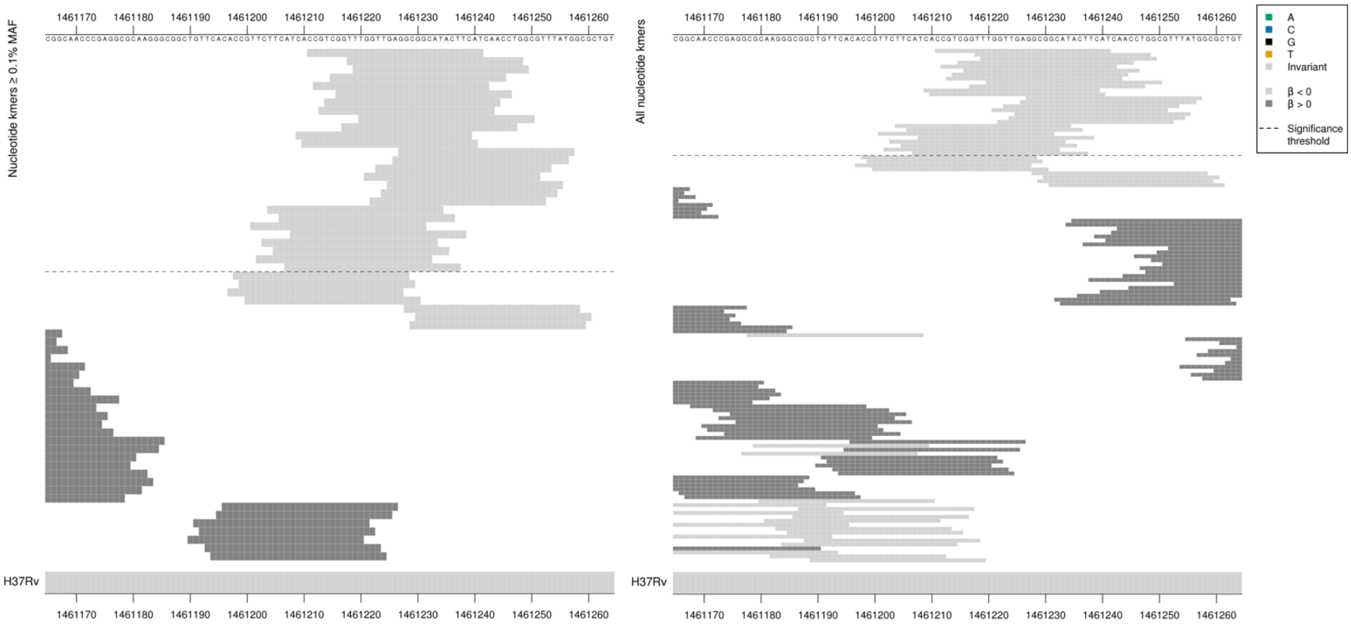

Removing the threshold requiring a kmer to be aligned to a gene/intergenic region in at least five genomes across the full dataset. Showing the alignment after lowering the threshold to just one genome:

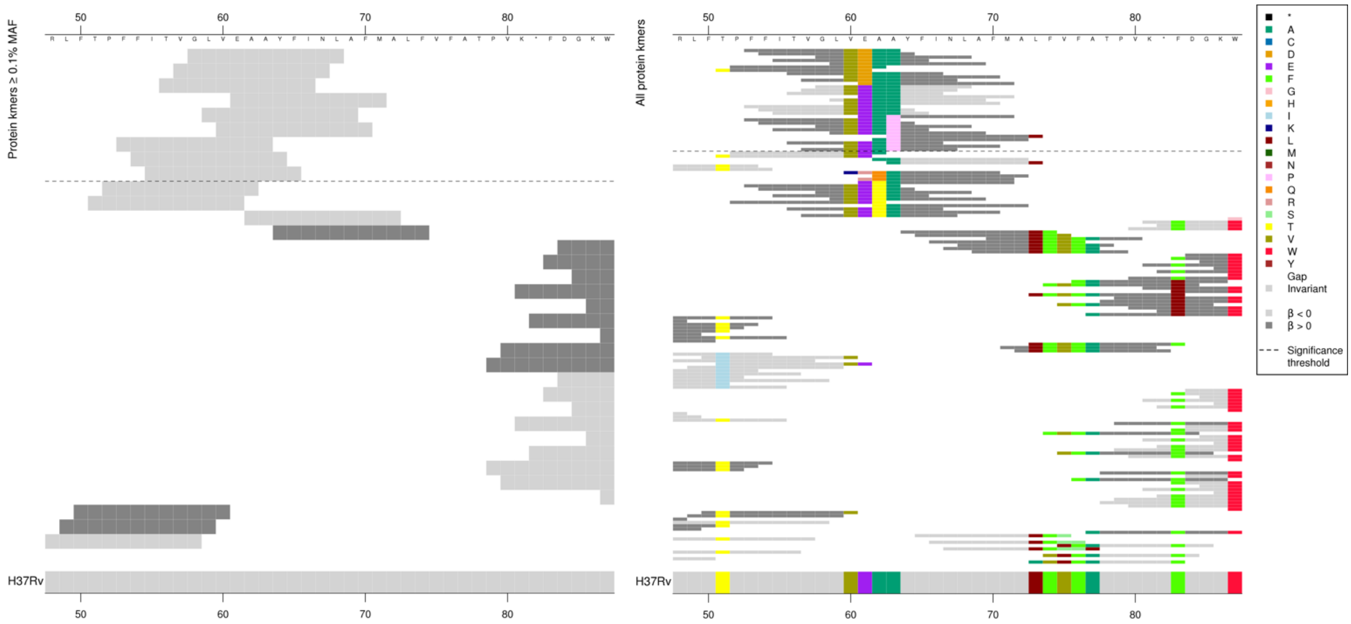

Protein kmers present in five or fewer genomes across the full dataset capturing:

61D

$-\log_{10}p$ = 16.64, beta = 4.10, MAC = 3, Ps = 53

63P

$-\log_{10}p$ = 10.63, beta = 4.15, MAC = 2, Ps = 61

*atpE* product = ATP synthase subunit C

Capturing a region identical to the reference as significantly associated with low MIC.

| Kmer | $-\log_{10} p$ | β | MAC | MAF | Ps | Variants |
|---|---|---|---|---|---|---|
| GLVEAAYFINL | 13.63 | -1.97 | 14 | 0.17 | 58 | WT / 61E, 63A |

#### *pgi*

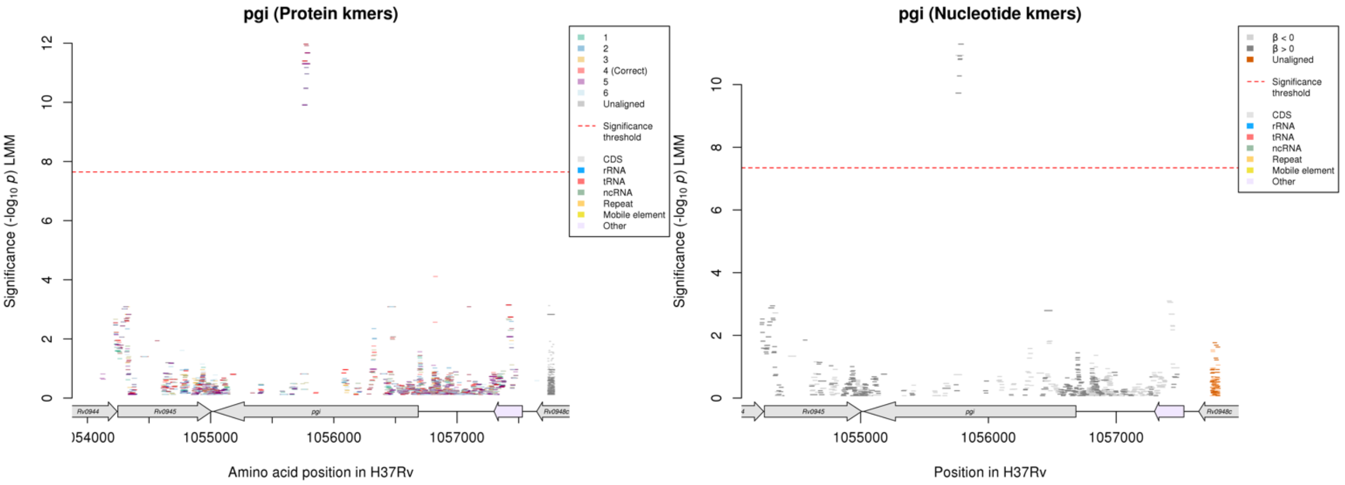

Protein kmers:

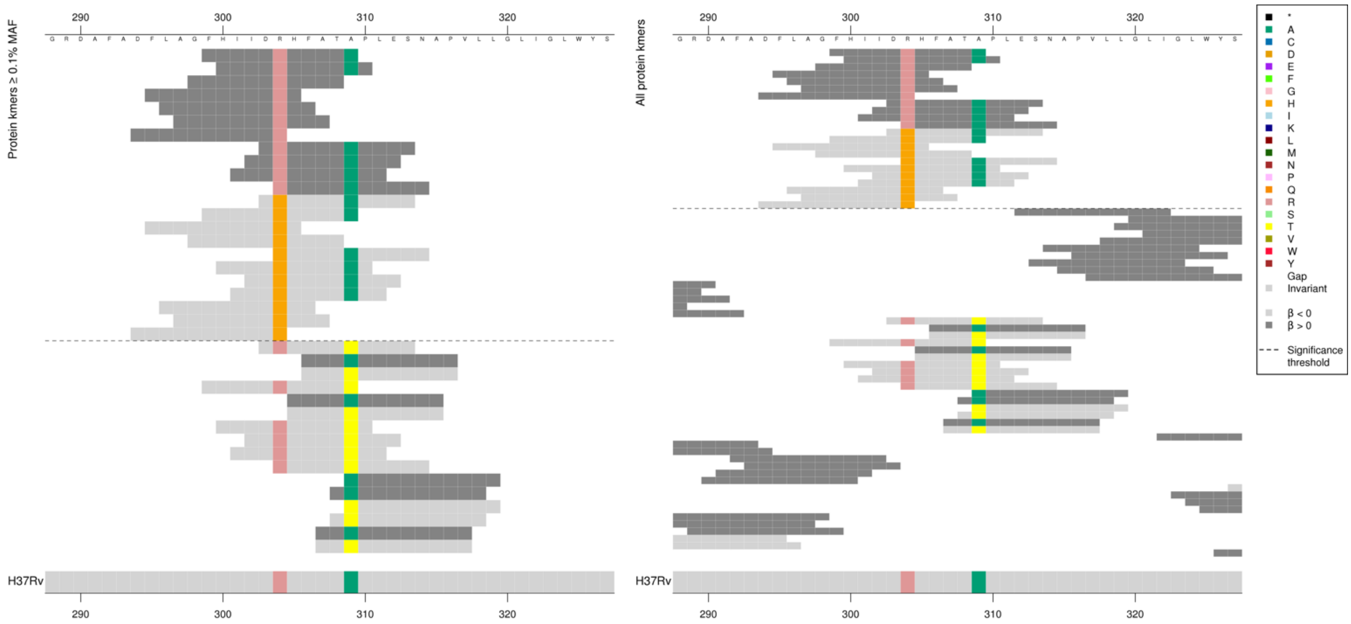

*pgi* produce = glucose-6-phosphate isomerase

| Kmer | -log₁₀ *p* | β | MAC | MAF | Ps | Variants |
|---|---|---|---|---|---|---|
| FHIIDRHFATA | 11.97 | 1.01 | 123 | 1.54 | 299 | 304R |
| DHHFATAPLES | 11.30 | -1.06 | 111 | 1.39 | 303 | 304H |

#### *mmaA4*

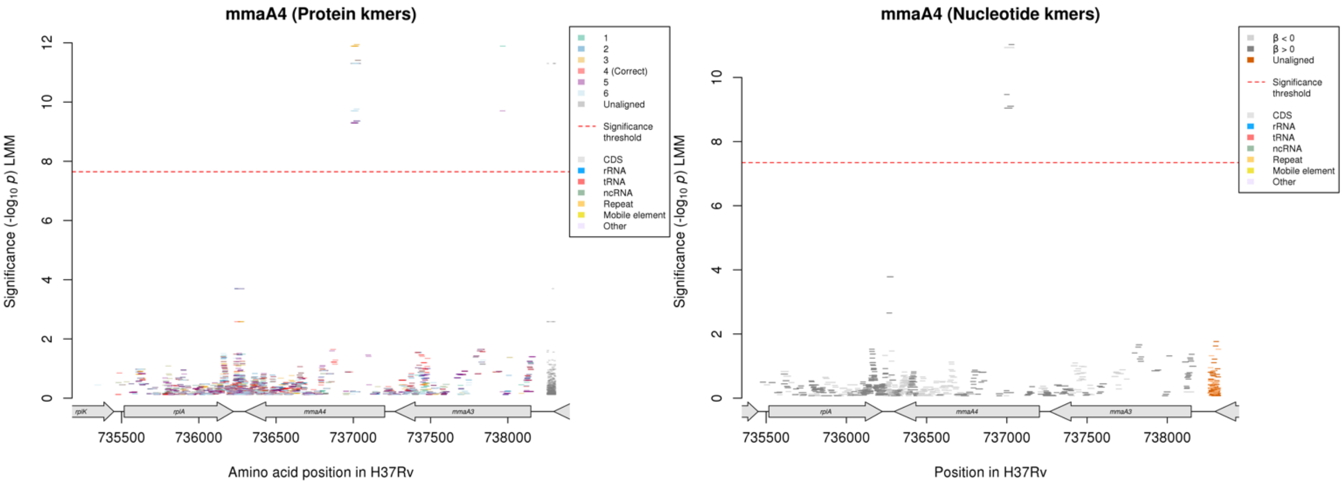

No significant protein kmers aligned to the correct reading frame.

Nucleotide kmers:

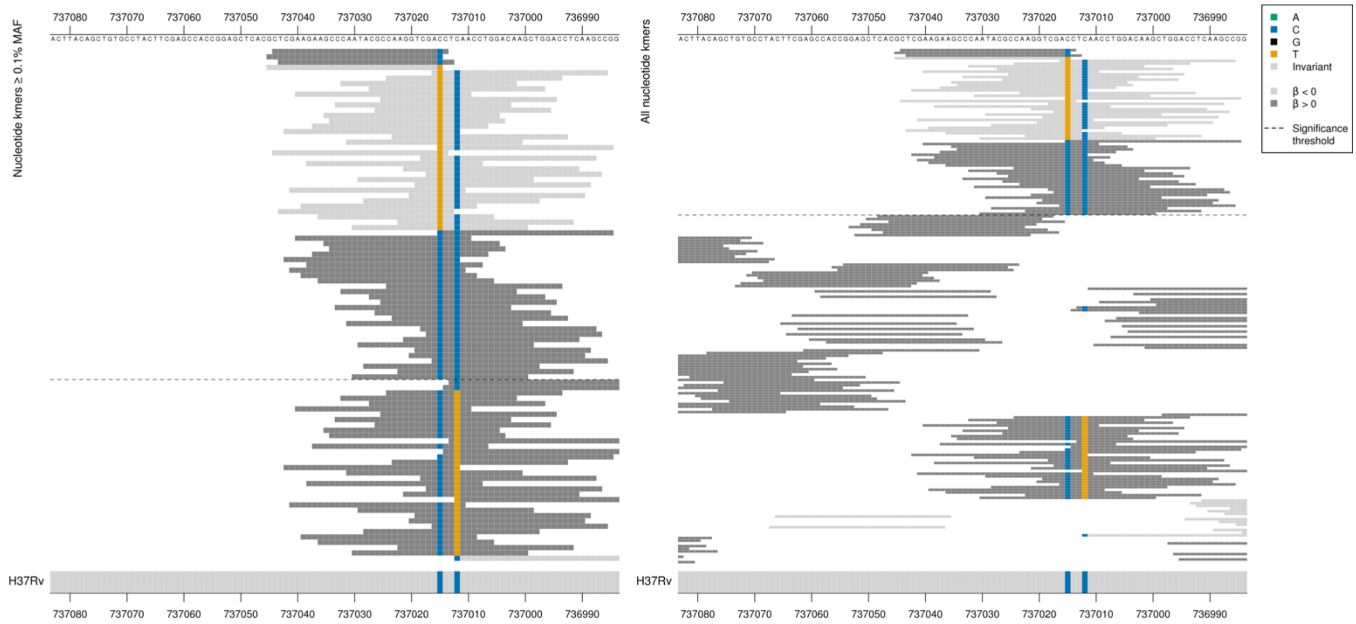

Synonymus mutation at position 737015/189.
High MIC-associated kmers = GAC = Asp / D
Low MIC-associated kmers = GAT = Asp / D

The high MIC-associated kmers were less significant when covering the mutation at position 737012/192.
Synonymous mutation at position 737012/192
High MIC-associated kmers = CTC = Leu / L
Low MIC-associated kmers = CTT = Leu / L

| Kmer | $-\log_{10} p$ | β | MAC | MAF | Ps | Variants |
|---|---|---|---|---|---|---|
| CTCGAAGAAGCCCAATACGCCAAGGTCGACC | 11.02 | 1.03 | 117 | 1.46 | 737044 | 737015/189C |
| GCTCGAAGAAGCCCAATACGCCAAGGTCGAT | 10.93 | -1.07 | 111 | 1.39 | 737045 | 737015/189T |
| CCTCAACCTGGACAAGCTGGACCTCAAGCCG | 9.47 | 0.89 | 125 | 1.56 | 737015 | 737015/189C, 737012/192C |

#### *rplC*

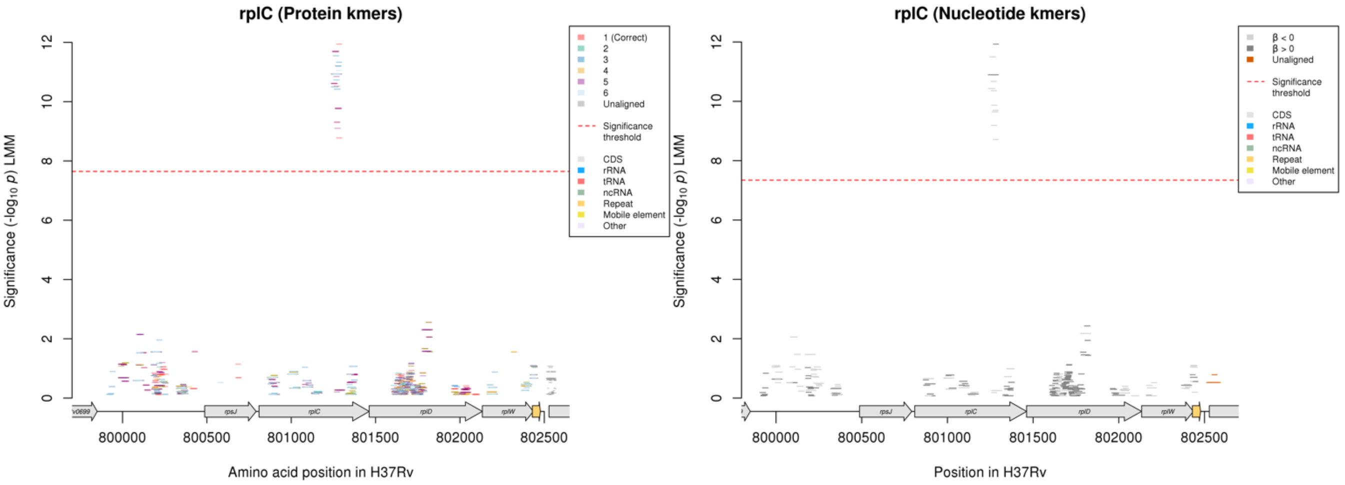

Protein kmers:

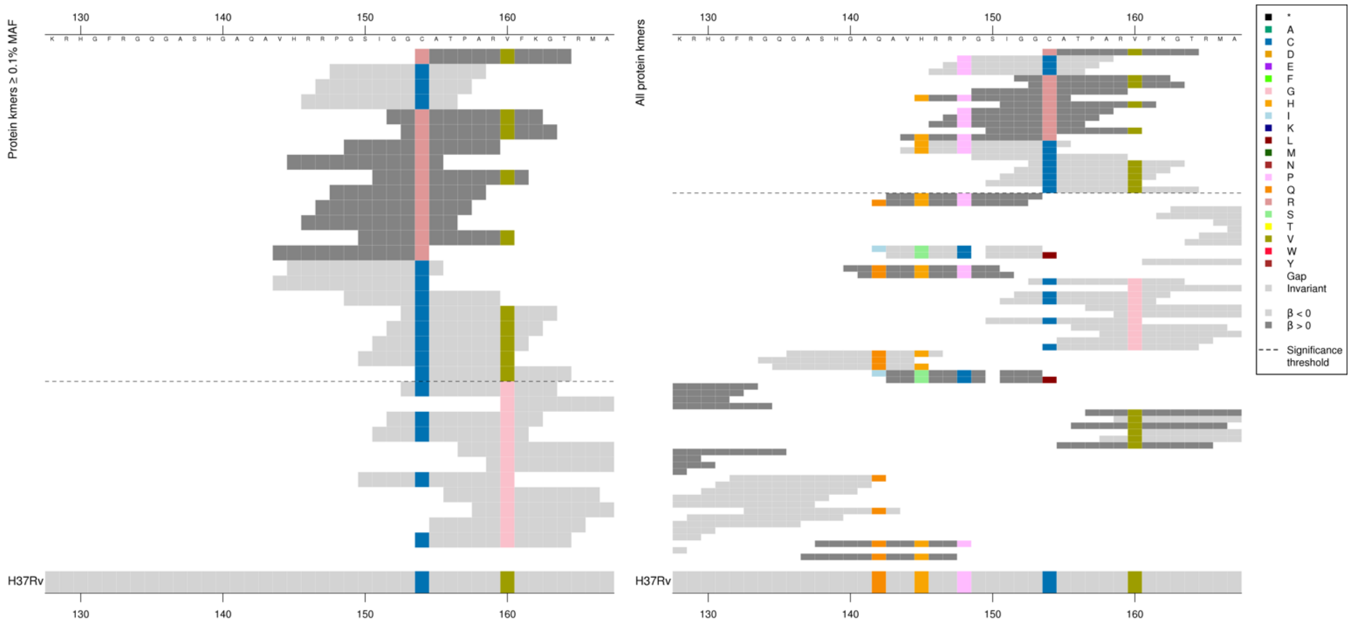

*rplC* product = 50S ribosomal protein L3

Mix of *p*-values for the significant kmers, largely capturing the variant at amino acid 154, but other variants possibly influenced the significance slightly.

| Kmer | $-\log_{10} p$ | β | MAC | MAF | Ps | Variants |
|---|---|---|---|---|---|---|
| RATPARVFKGT | 11.94 | 1.23 | 27 | 0.34 | 154 | 154R |
| PGSIGGCATPA | 11.69 | -1.29 | 24 | 0.30 | 148 | 154C |

## *Rv0078A*

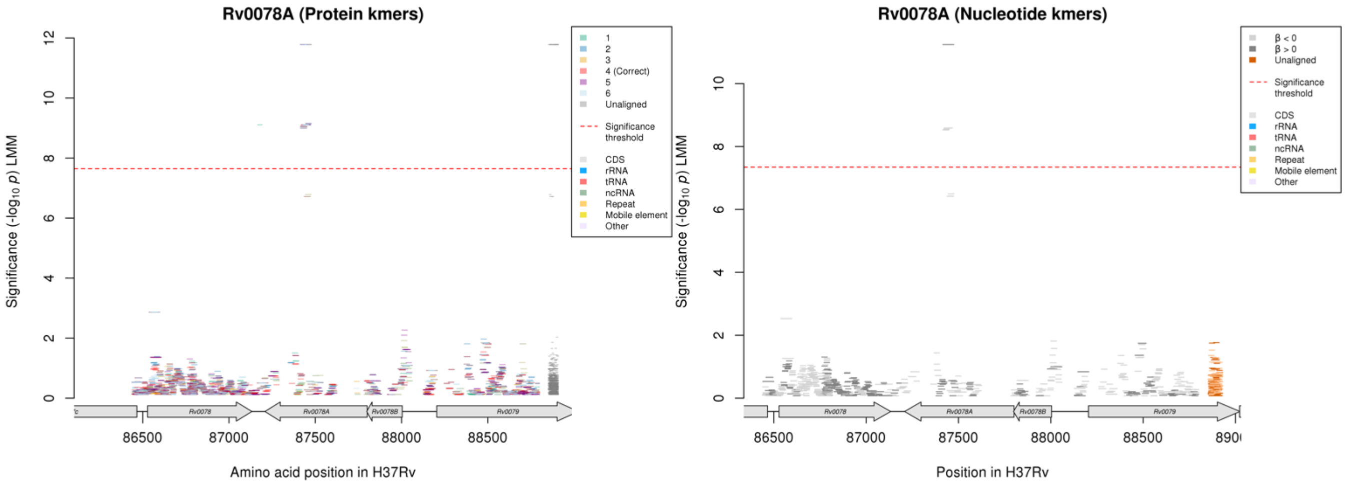

Protein kmers:

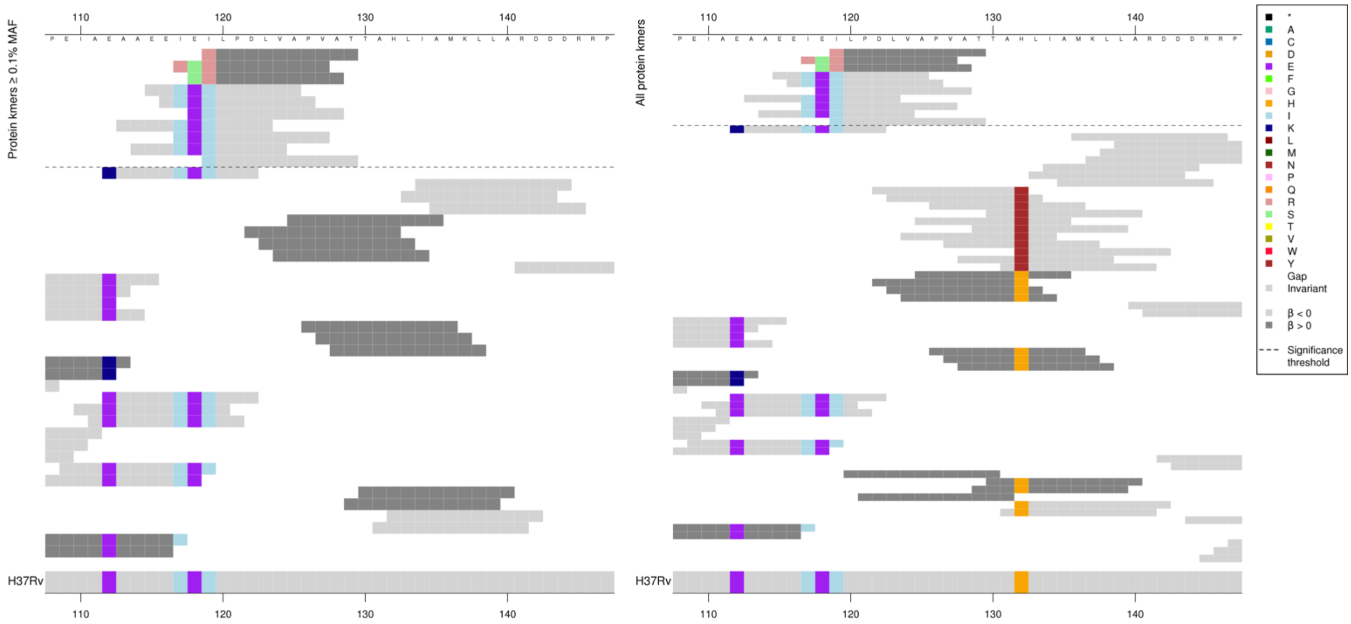

The same region for the nucleotide kmer analysis:

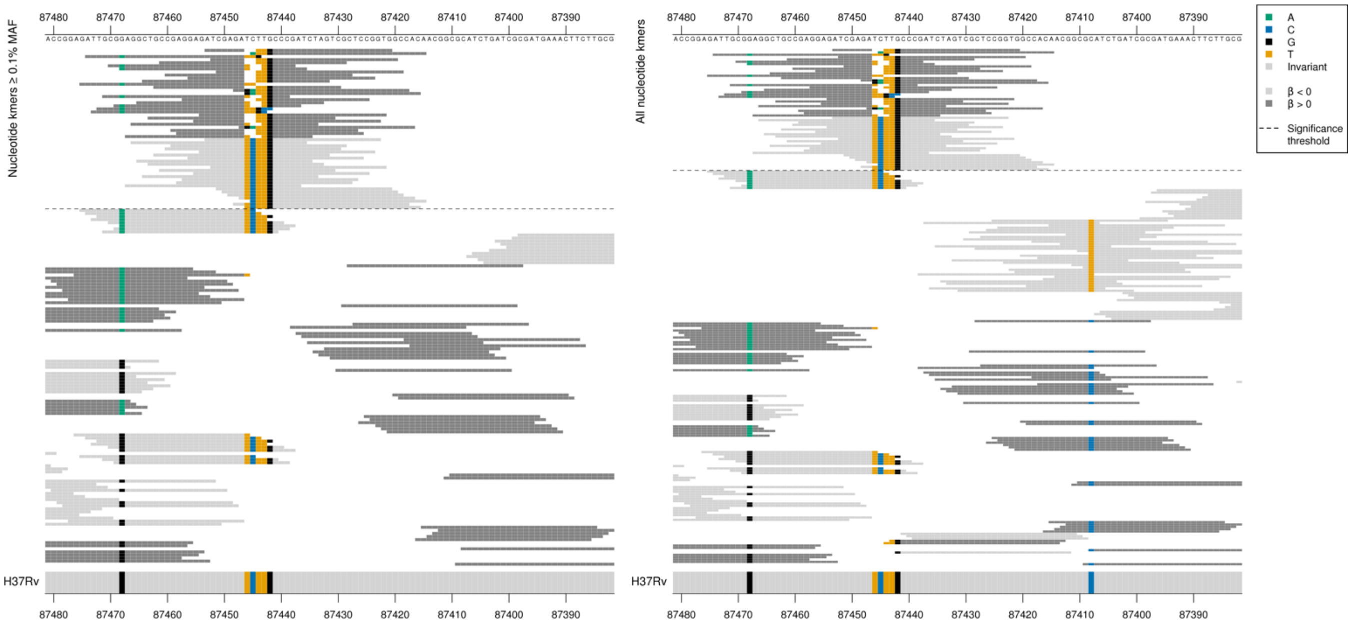

These kmers in the BDQ analysis had the opposite direction of effect to the AMI analysis.

| Kmer | $-\log_{10} p$ | β | MAC | MAF | Ps | Variants |
|---|---|---|---|---|---|---|
| ATCGAGATTGCCCGATCTAGTCGCTCCGGTG | 11.24 | 1.52 | 19 | 0.24 | 87453 | 87446-87445/356-357 CT deletion |
| ATCGAGATCTTGCCCGATCTAGTCGCTCCGG | 8.59 | -1.19 | 24 | 0.30 | 87453 | WT |

### *amiA2 and era*

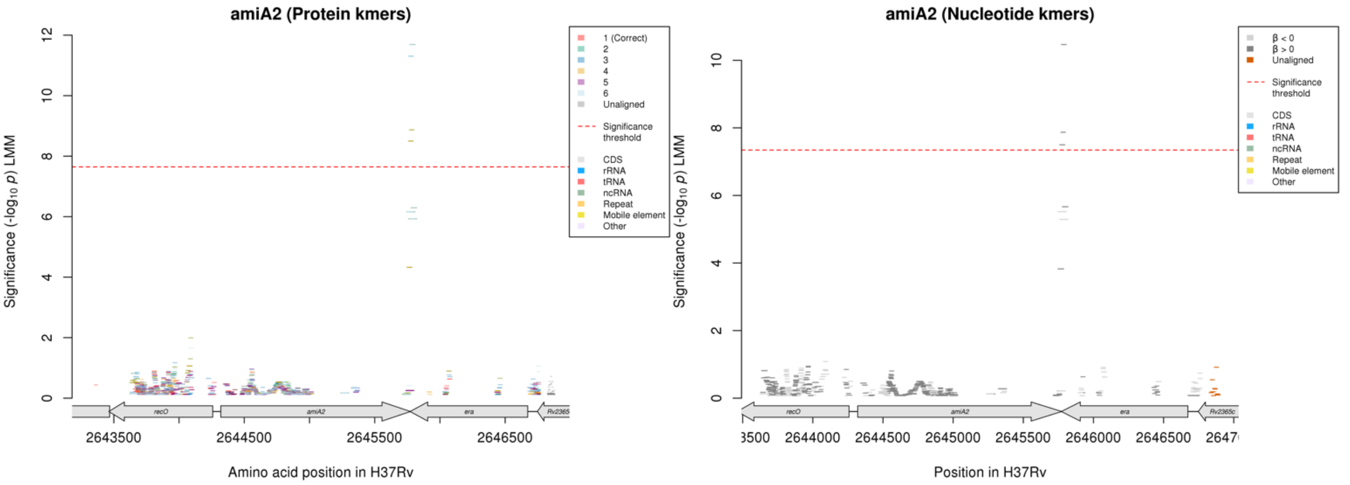

*amiA2* ends at position 2645774 *era* ends at position 2645771, therefore the genes overlap by a few bp.
*amiA2* coding strand nucleotide kmers:

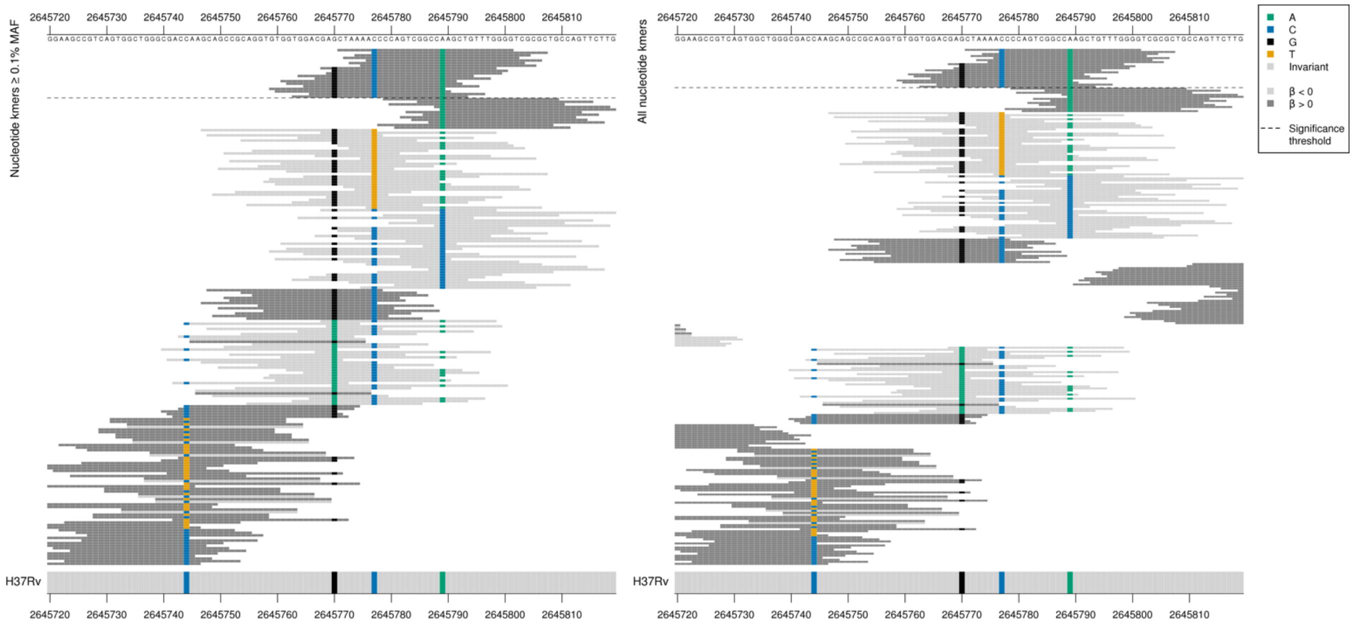

*era* coding strand (same kmers, opposite strand) nucleotide kmers:

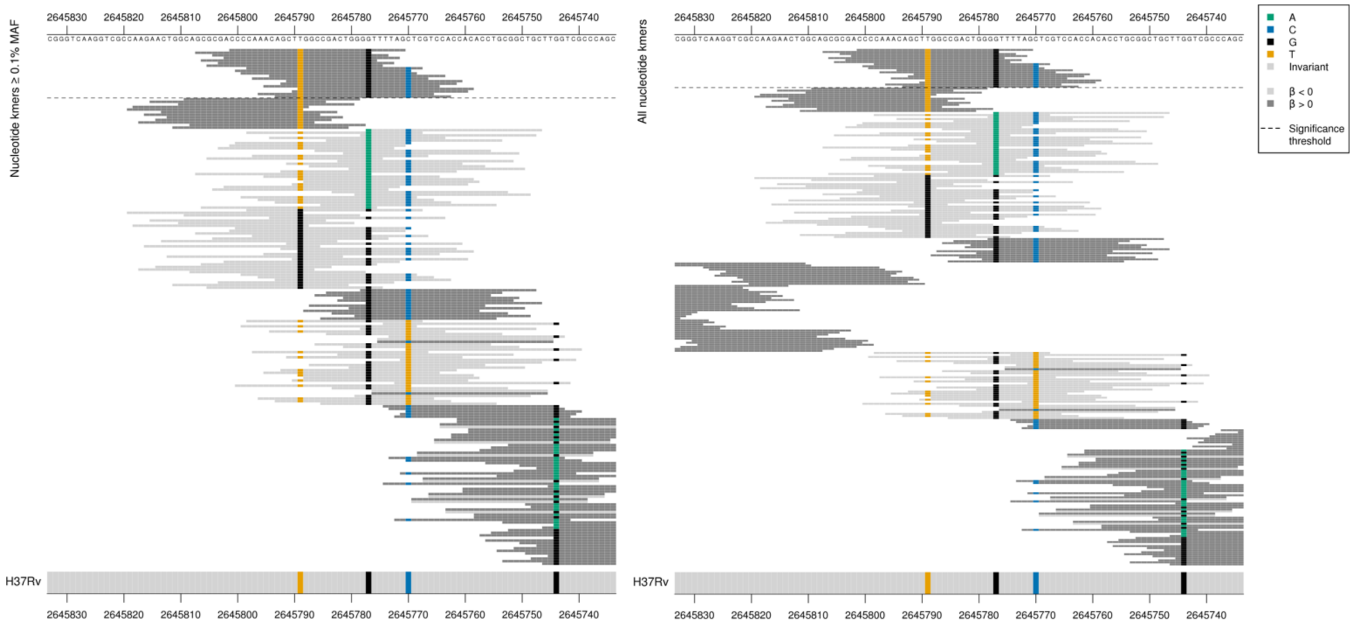

Appears that the significant, high MIC-associated kmers were capturing a combination of alleles at positions 2645777 and 2645789 and 2645770.

Most significant kmers covered variants at positions 2645777 and 2645789. These mutations fall just after the stop codon for *amiA2*.
Mutations in *era* in the correct reading frame:
2645777:
High MIC-associated = GGG = Gly / G
Kmers below significance threshold = GGA = Gly / G
2645789:
High MIC-associated = CTT = Leu / L
Kmers below significance threshold = CTG = Leu / L
Both synonymous.

The kmers were slightly less significant when also covering the variant at position 2645770, which is 1bp after the stop codon for *era* but before the stop codon of *amiA2*.
Mutations in *amiA2* in the correct reading frame:
High MIC-associated = AGC = Ser / S
Kmers below significance threshold = AAC = Asn / N

*amiA2* product = amidase
*era* product = GTPase Era

| Kmer | $-\log_{10} p$ | β | MAC | MAF | Ps | Variants |
|---|---|---|---|---|---|---|
| CCCCAAACAGCTTGGCCGACTGGGGTTTTAG | 10.47 | 1.26 | 90 | 1.12 | 2645801 | 2645777/897G and 2645789/885T in era |
| CAAACAGCTTGGCCGACTGGGGTTTTAGCTC | 7.87 | 0.88 | 111 | 1.39 | 2645798 | 2645777/897G and 2645789/885T in era, 2645770/1451G (484S aa) in amiA2 |

#### *viuB*

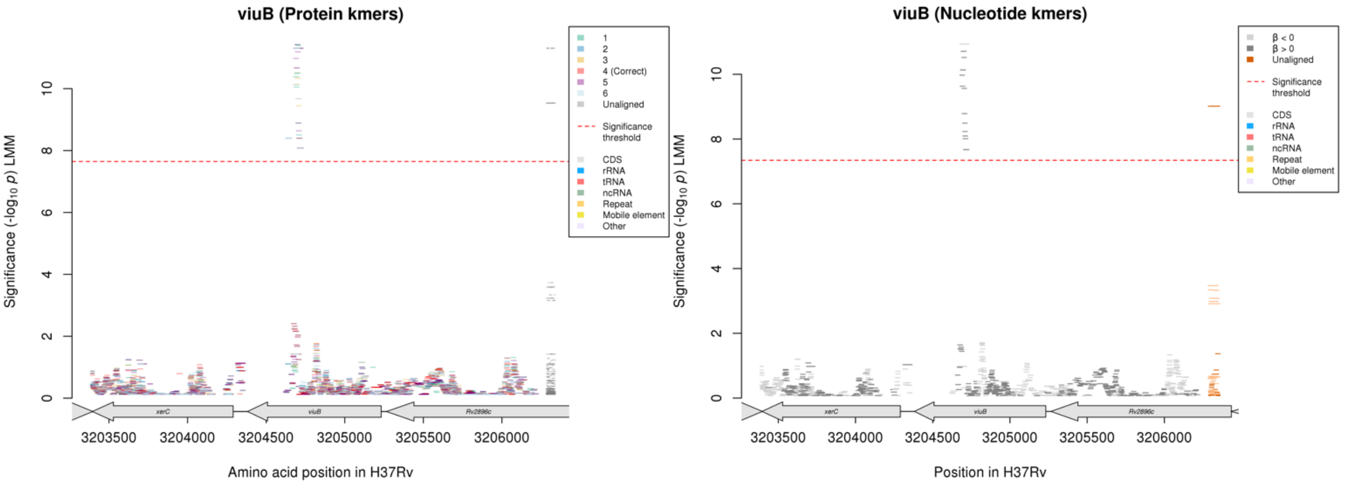

No significant protein kmers aligned to the correct reading frame.

Nucleotide kmers:

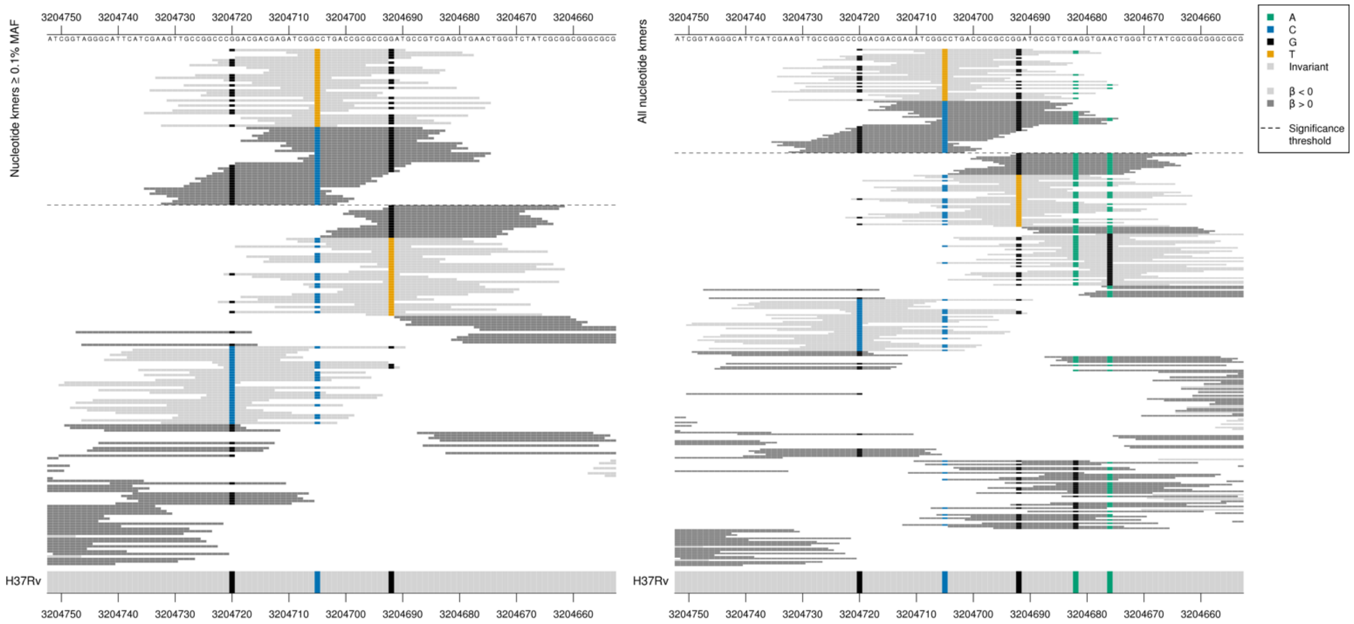

*viuB* product = mycobactin utilization protein ViuB
Gene begins at position 3205232.
Captured a combination of mutations, including synonymous and non-synonymous mutations.

3204705 (codon 176):
High MIC-associated = GGC = Gly / G
Low MIC-associated = GGT = Gly / G

3204720 (codon 171):
High MIC-associated = CCG = Pro / P
Kmers below significance threshold = CCC = Pro / P

3204692 (codon 181):
High MIC-associated = GAT = Asp / D
Kmers below significance threshold = TAT = Tyr / Y

3204676 (codon 186):
High MIC-associated = AAC = Asn / N
Kmers below significance threshold = AGC = Ser / S

3204682 (codon 184):
Low and high MIC-associated = GAG = Glu / E
Kmers below significance threshold= GGG = Gly / G

All low MIC-associated kmers were the same *p*-value, capturing 3204705/528T. The high MIC-associated kmers varied in *p*-value depending the positions covered.

Kmers below significance threshold = 3204682G
Kmers below significance threshold = 3204720C, 3204692T, 3204676G

| Kmer | $-\log_{10} p$ | β | MAC | MAF | Ps | Variants |
|---|---|---|---|---|---|---|
| GGACGACGAGATCGGTCTGACCGCGCCGGAT | 10.93 | -1.07 | 111 | 1.39 | 3204720 | 3204705/528T |
| CGACGAGATCGGCCTGACCGCGCCGGATGCC | 10.71 | 0.84 | 177 | 2.21 | 3204717 | 3204705/528C, 3204692/541G |
| ATCGGCCTGACCGCGCCGGATGCCGTCGAGG | 10.13 | 0.80 | 180 | 2.25 | 3204710 | 3204705/528C, 3204692/541G, 3204682/551A |
| CCTGACCGCGCCGGATGCCGTCGAGGTGAAC | 9.97 | 0.75 | 187 | 2.33 | 3204705 | 3204705/528C, 3204692/541G, 3204682/551A, 3204676/557A |
| GGACGACGAGATCGGCCTGACCGCGCCGGAT | 8.79 | 0.69 | 218 | 2.72 | 3204720 | 3204720/513G, 3204705/528C, 3204692/541G |
| CCCGGACGACGAGATCGGCCTGACCGCGCCG | 8.49 | 0.76 | 164 | 2.05 | 3204723 | 3204720/513G, 3204705/528C |

### *pncA*

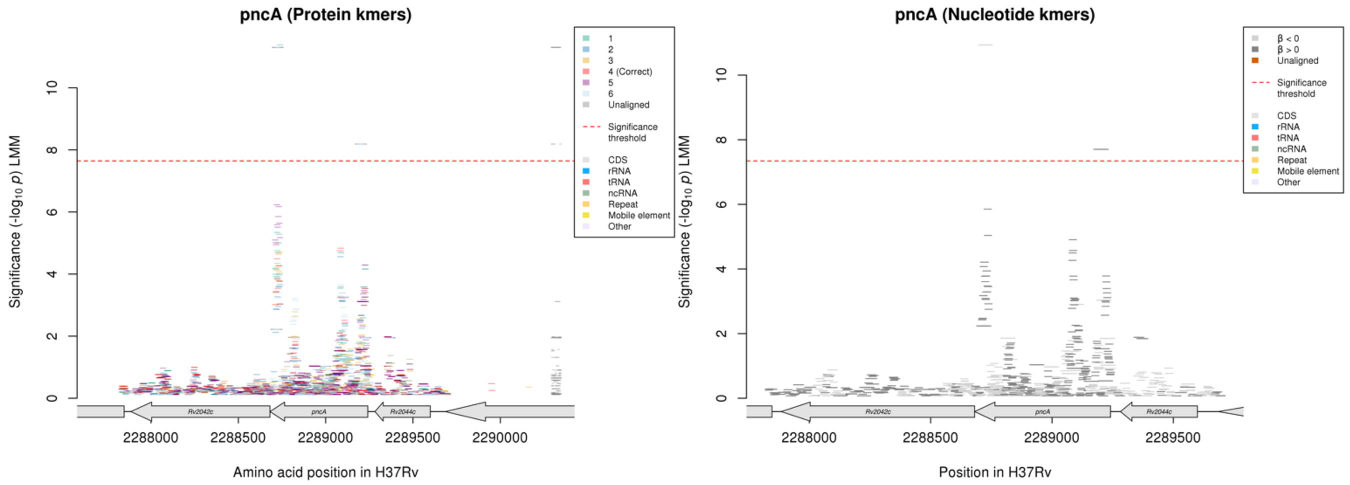

Protein kmers:

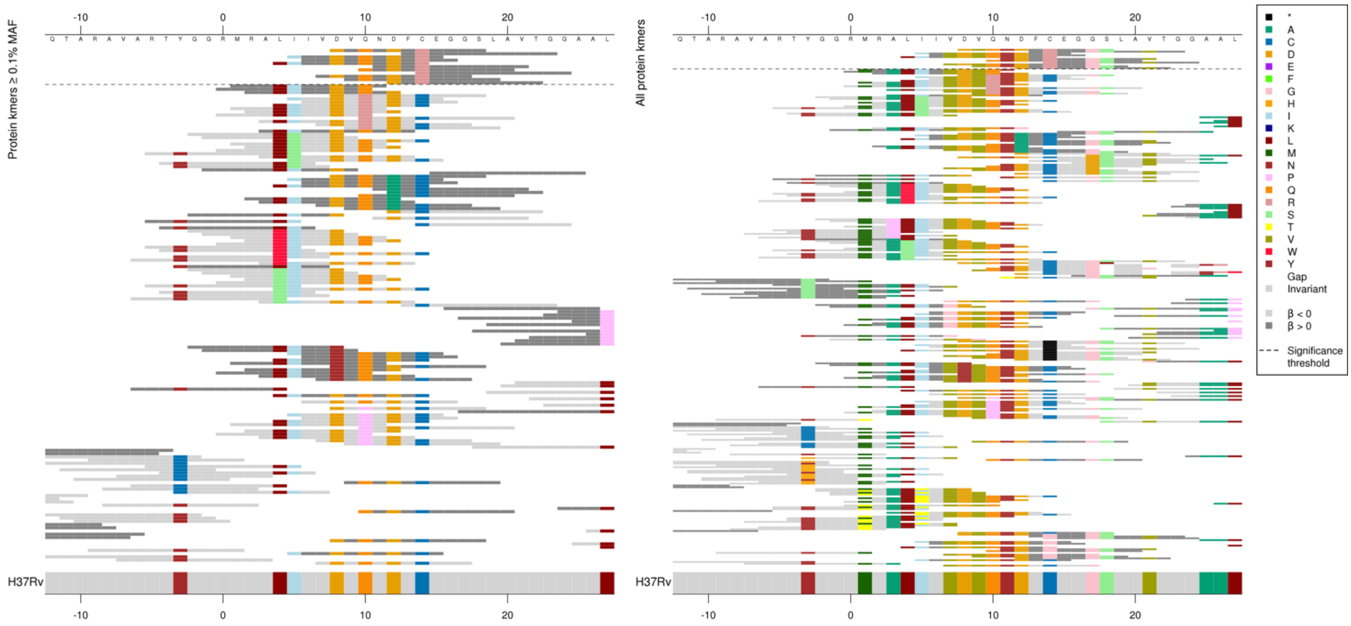

The substitution at codon 14 in *pncA* was not significant for any other drugs where *pncA* is a top gene.

| Kmer | $-\log_{10} p$ | β | MAC | MAF | Ps | Variants |
|---|---|---|---|---|---|---|
| DVQNDFREGGS | 8.19 | 0.91 | 42 | 0.52 | 8 | 14R |

Protein kmers:

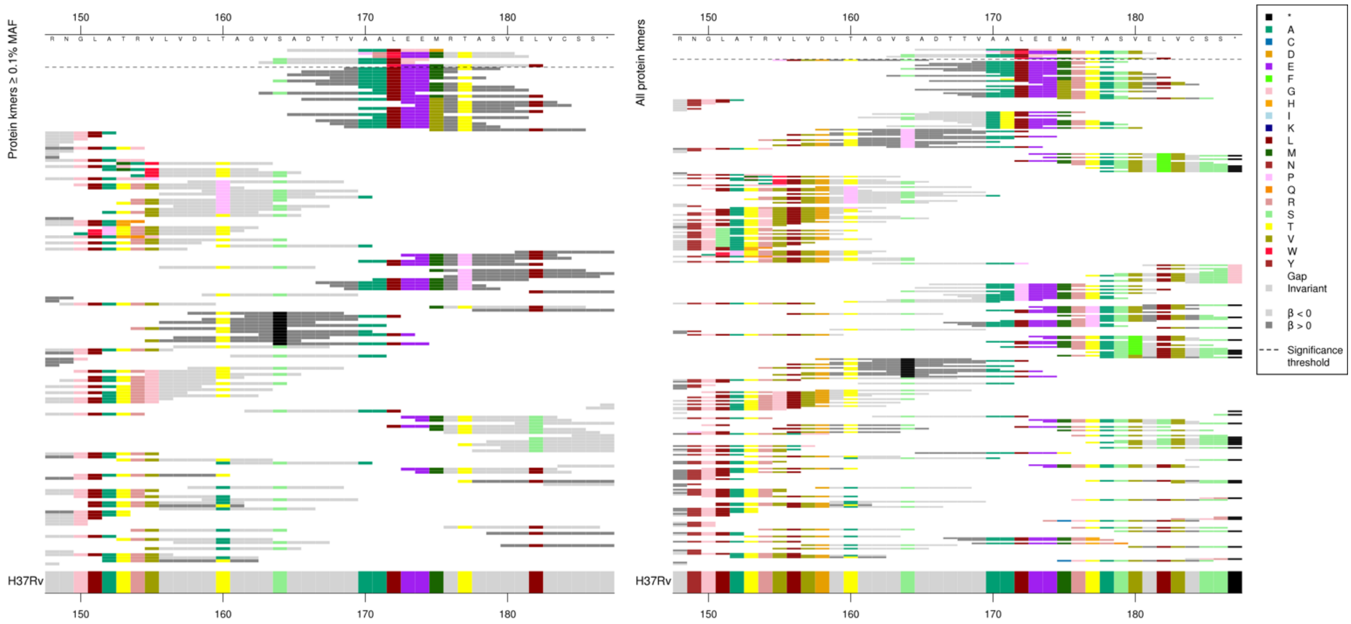

Nucleotide kmers in the same region:

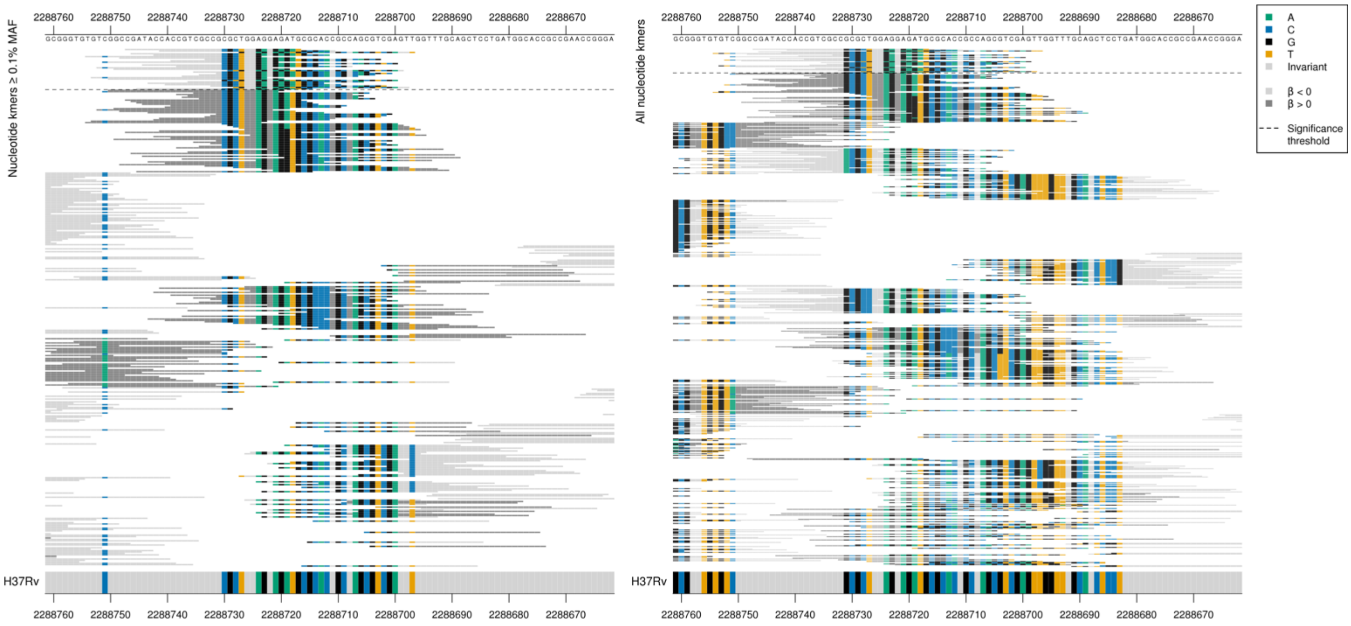

This is also a different region of *pncA* to that seen for other drugs which have *pncA* as a top gene.

The kmers captured an insertion of a G between positions 2288727-2288724/515-518, changing GG to GGG.

| Kmer | $-\log_{10} p$ | β | MAC | MAF | Ps | Variants |
|---|---|---|---|---|---|---|
| CGATACCACCGTCGCCGCGCTGGGAGGAGAT | 10.93 | -1.07 | 111 | 1.39 | 2288747 | G insertion between 2288727-2288724/515-518 |

### *murA*

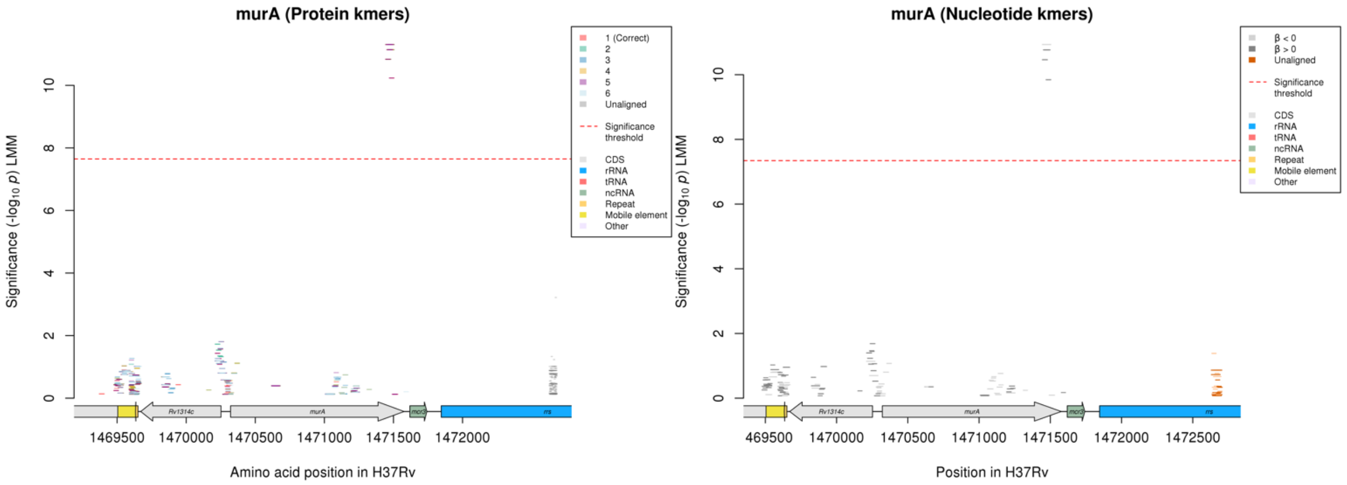

Protein kmers:

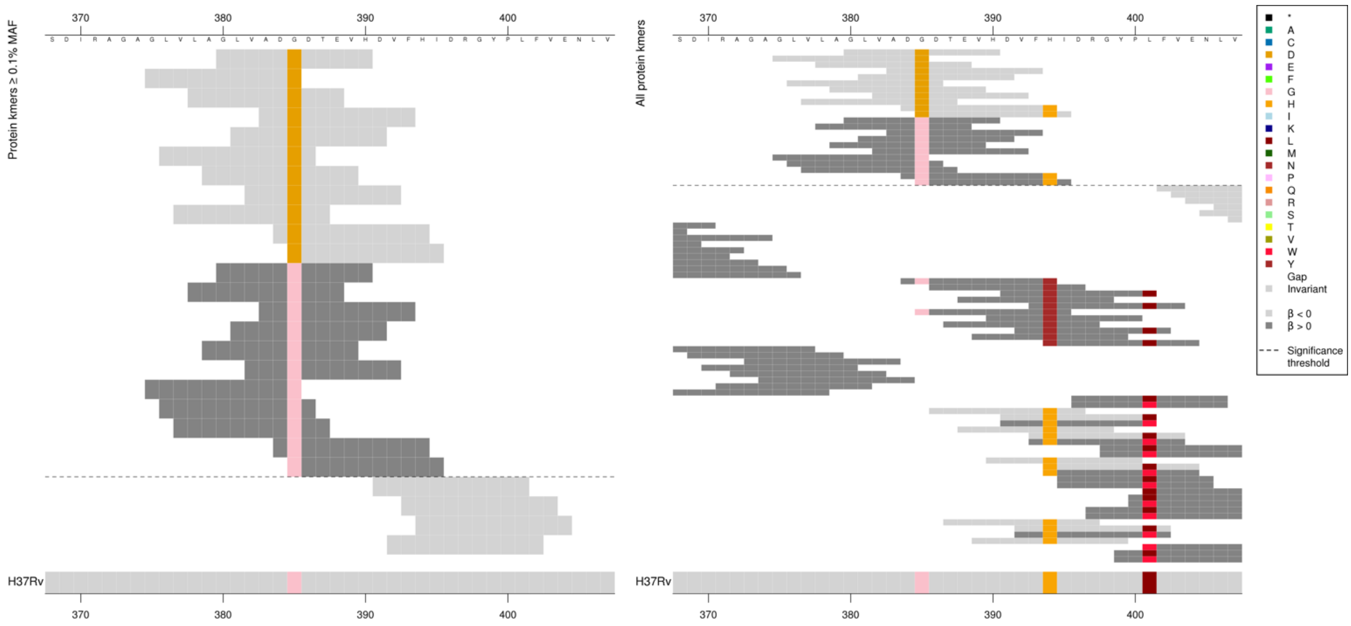

*murA* product = UDP-N-acetylglucosamine 1-carboxyvinyltransferase

| Kmer | -log₁₀ *p* | β | MAC | MAF | Ps | Variants |
|---|---|---|---|---|---|---|
| GLVADDDTEVH | 11.30 | -1.06 | 111 | 1.39 | 380 | 385D |
| GLVADGDTEVH | 11.14 | 1.04 | 113 | 1.41 | 380 | 385G |

# *Rv0792c/Rv0793*

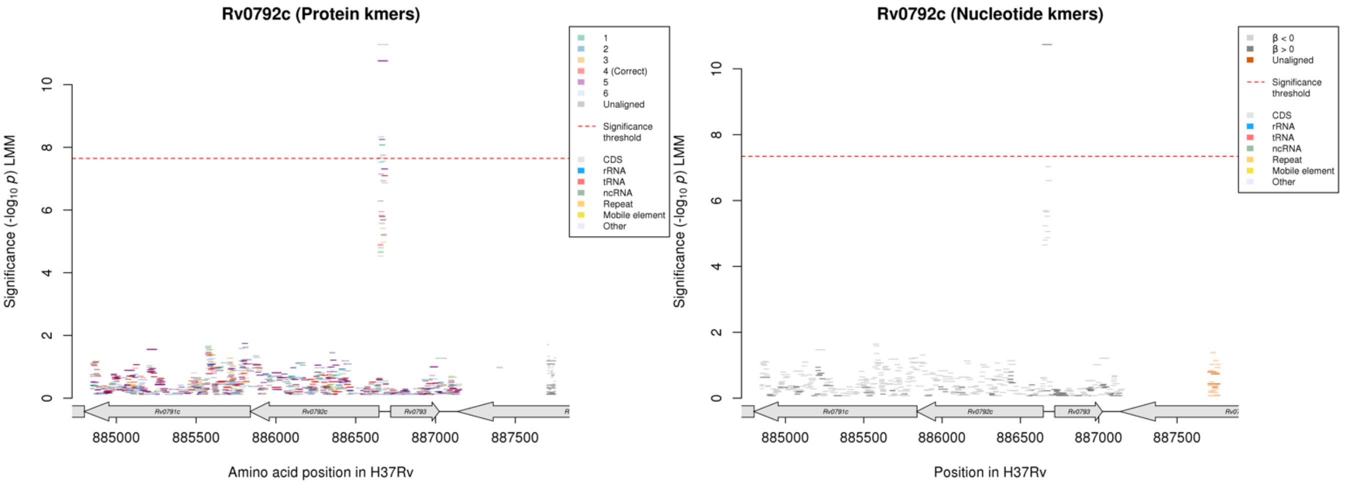

Nucleotide kmers:

Coding strand for *Rv0792c*:

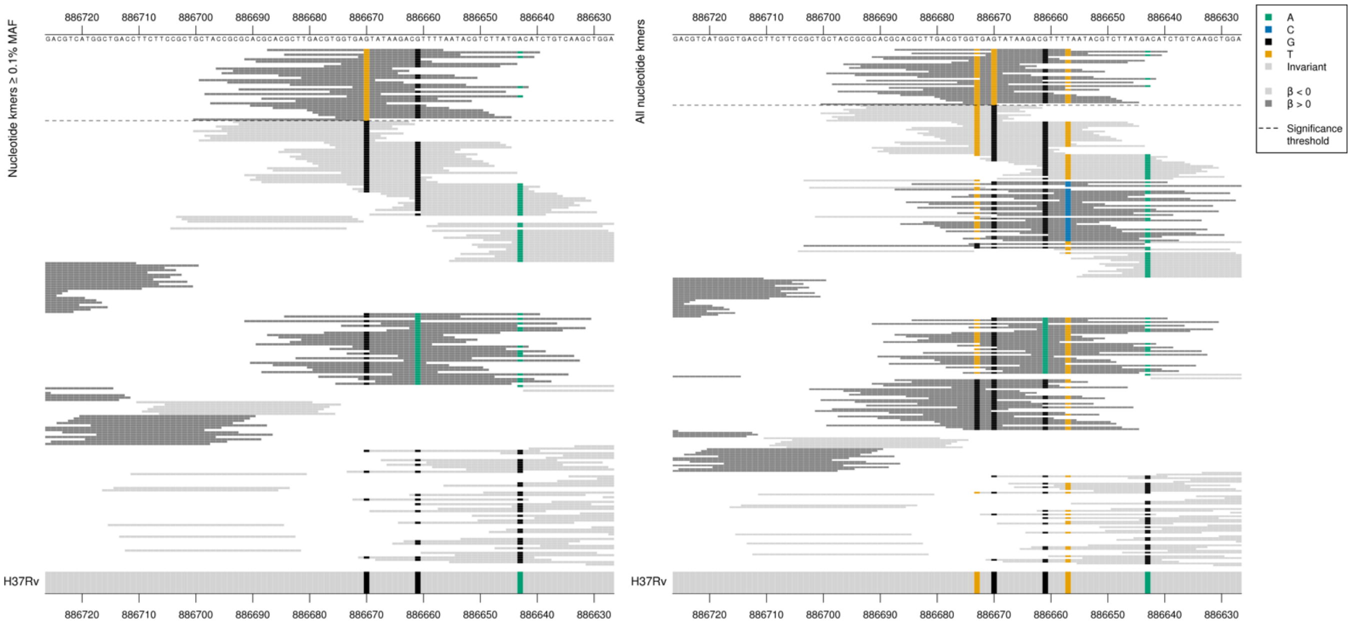

*Rv0792c* product = transcriptional regulator

*Rv0793* product = monooxygenase

*Rv0792c* starts at position 886646, *Rv0793* starts at position 886719. Possible promoter region mutation. Captured a mutation at position 886670, as found in the AMI GWAS but with an opposite direction of effect for BDQ.

| Kmer | -log10 *p* | β | MAC | MAF | Ps | Variants |
|---|---|---|---|---|---|---|
| CACGCTTGACGTGGTGATTATAAGACGTTTT | 10.74 | 1.44 | 20 | 0.25 | 886687 | 886670/-24T Rv0792c, 886670/-49A Rv0793 |

### *dnaB*

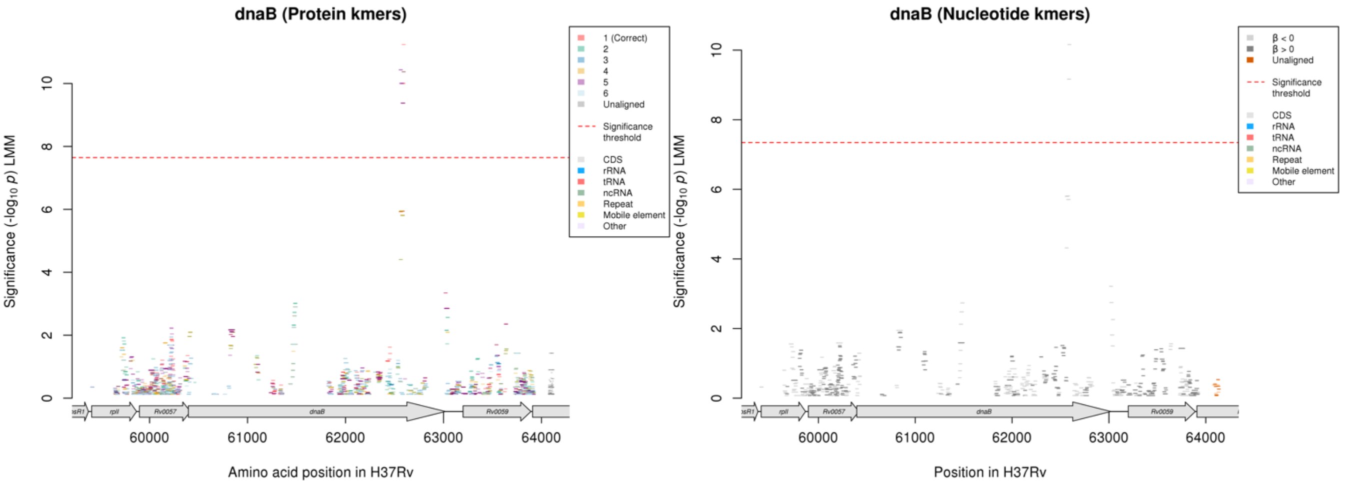

Protein kmers:

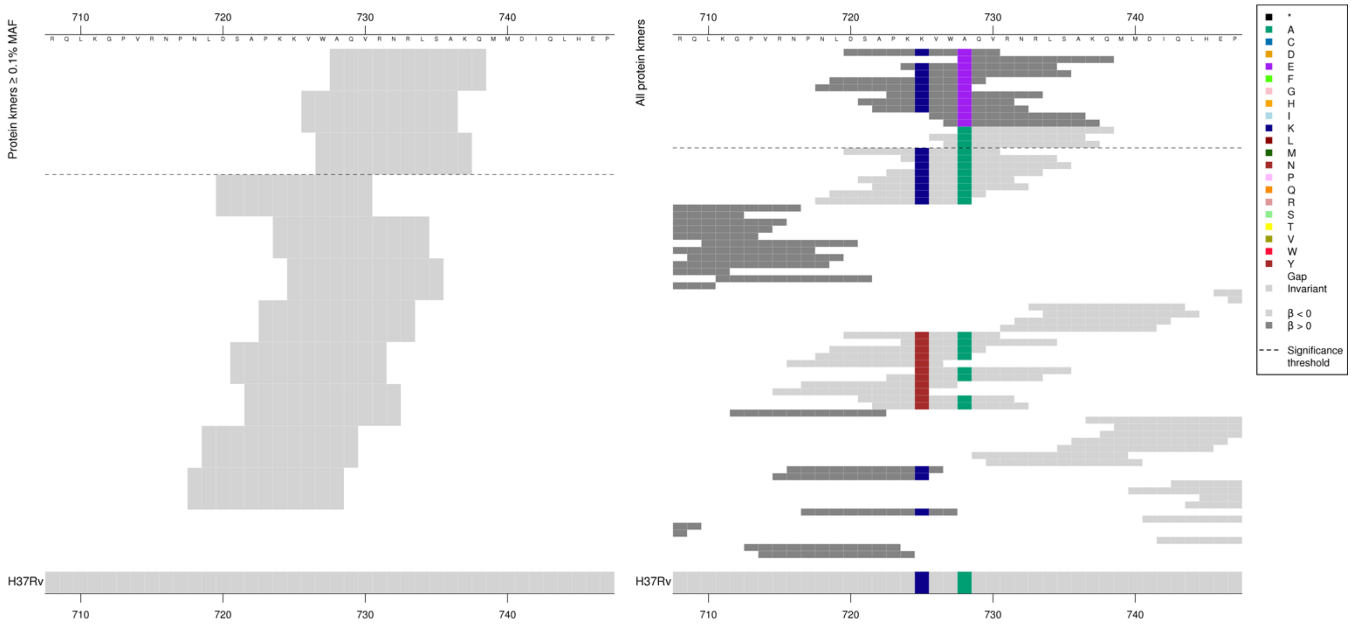

*dnaB* product = replicative DNA helicase

Only the 728A containing low MIC-associated kmers were above the frequency cut off to have been included in the analysis.
Low MAF kmers = 728E.

| Kmer | $-\log_{10} p$ | β | MAC | MAF | Ps | Variants |
|---|---|---|---|---|---|---|
| AQVRNRLSAKQ | 11.24 | -2.36 | 9 | 0.11 | 728 | 728A |

### *Rv2665:clpC2*

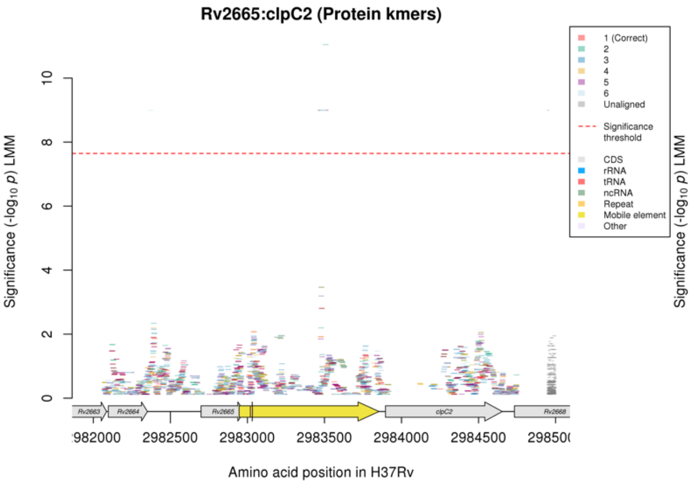

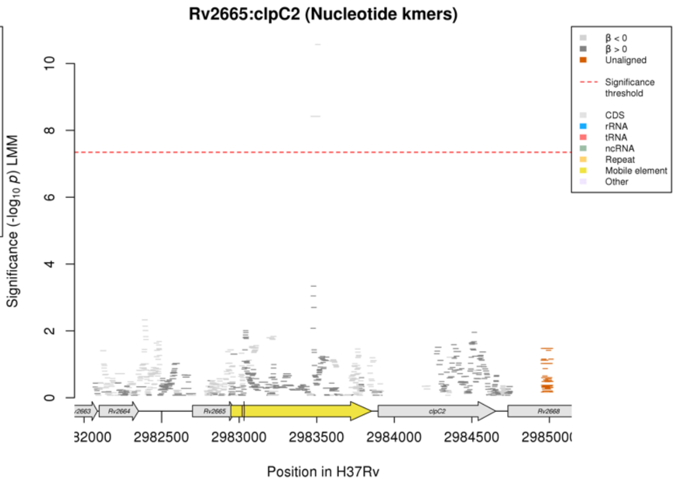

Nucleotide kmers:

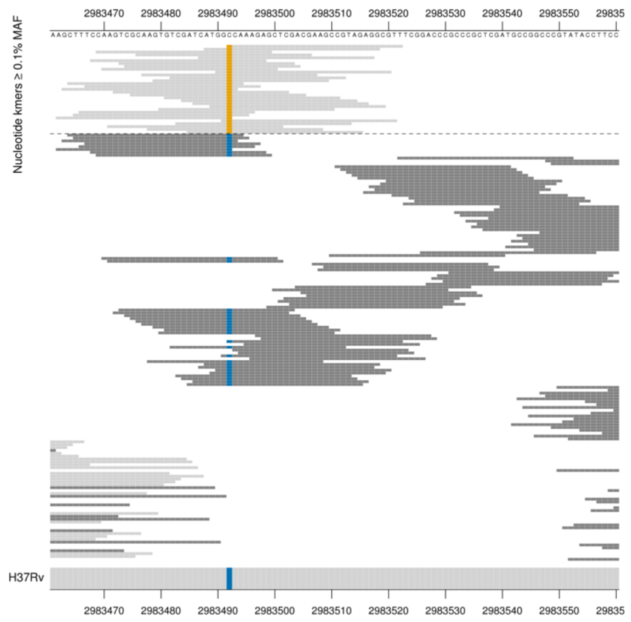

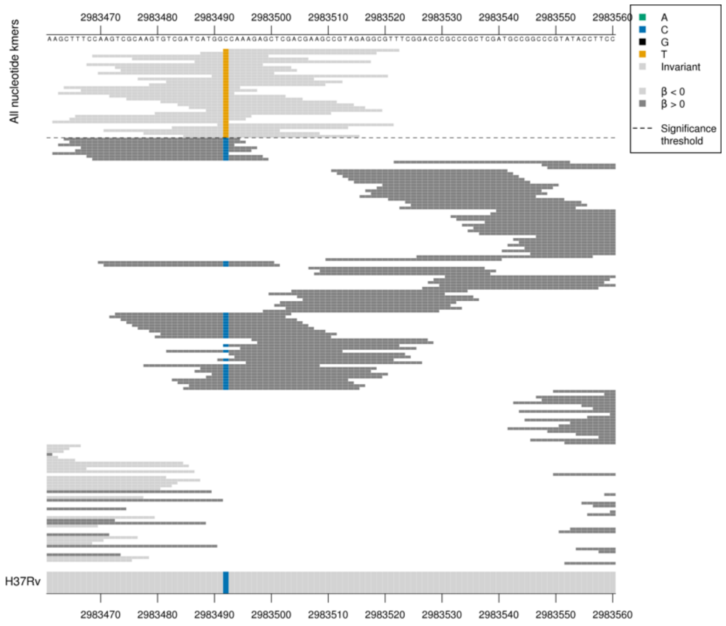

*clpC2* product = ATP-dependent protease ATP-binding subunit ClpC

Capturing a mutation in a mobile element at position 2983492. Only the T containing low MIC-associated kmers were significantly associated.

| Kmer | $-\log_{10} p$ | β | MAC | MAF | Ps | Variants |
|---|---|---|---|---|---|---|
| TCAAAGAGCTCGACGAAGCCGTAGAGGCGTT | 10.57 | -1.58 | 55 | 0.69 | 2983492 | 2983492T |

### *PPE54*

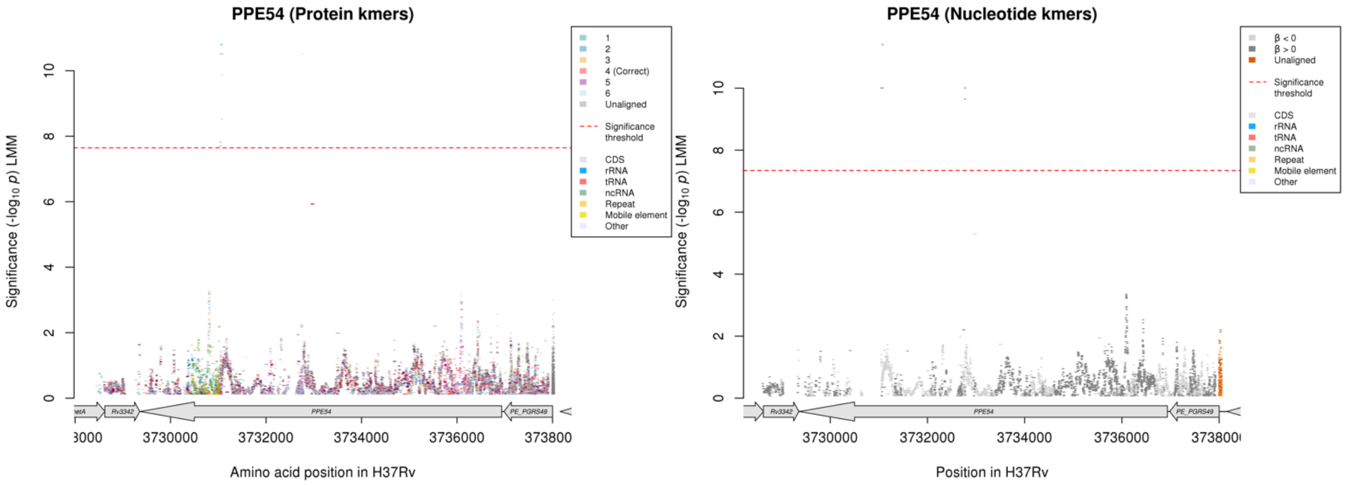

As seen for AMI, for the correct protein kmer reading frame there were significant kmers in just one region of the gene, for the nucleotide kmer analysis there were significant kmers in two regions of the gene.

Protein kmers:

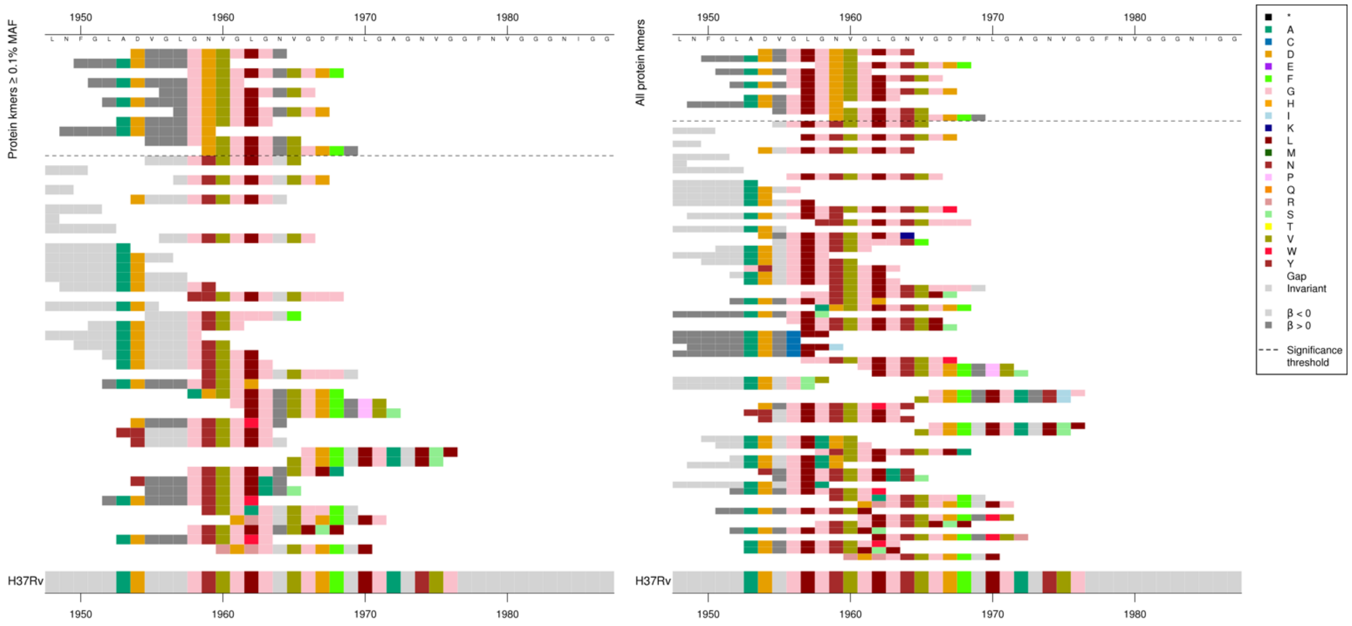

One of the regions for the nucleotide kmer analysis:

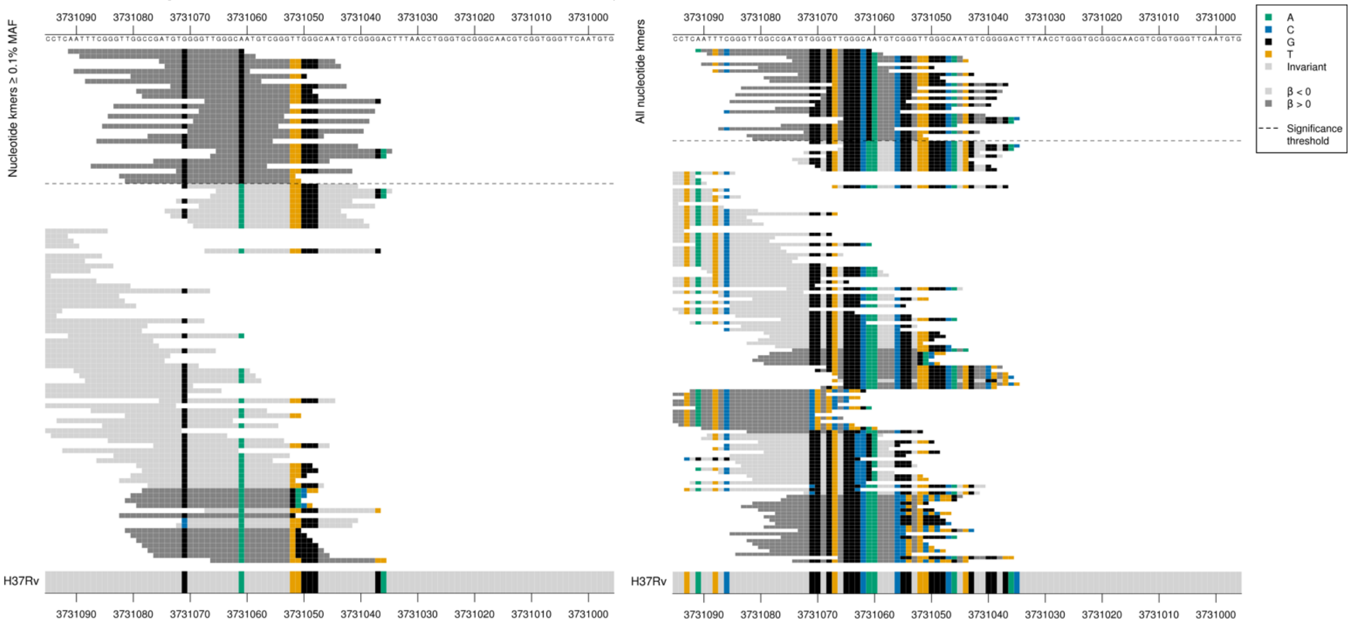

The second significant region in the nucleotide analysis was the same as was seen for AMI. The second region contains kmers that align to both regions.

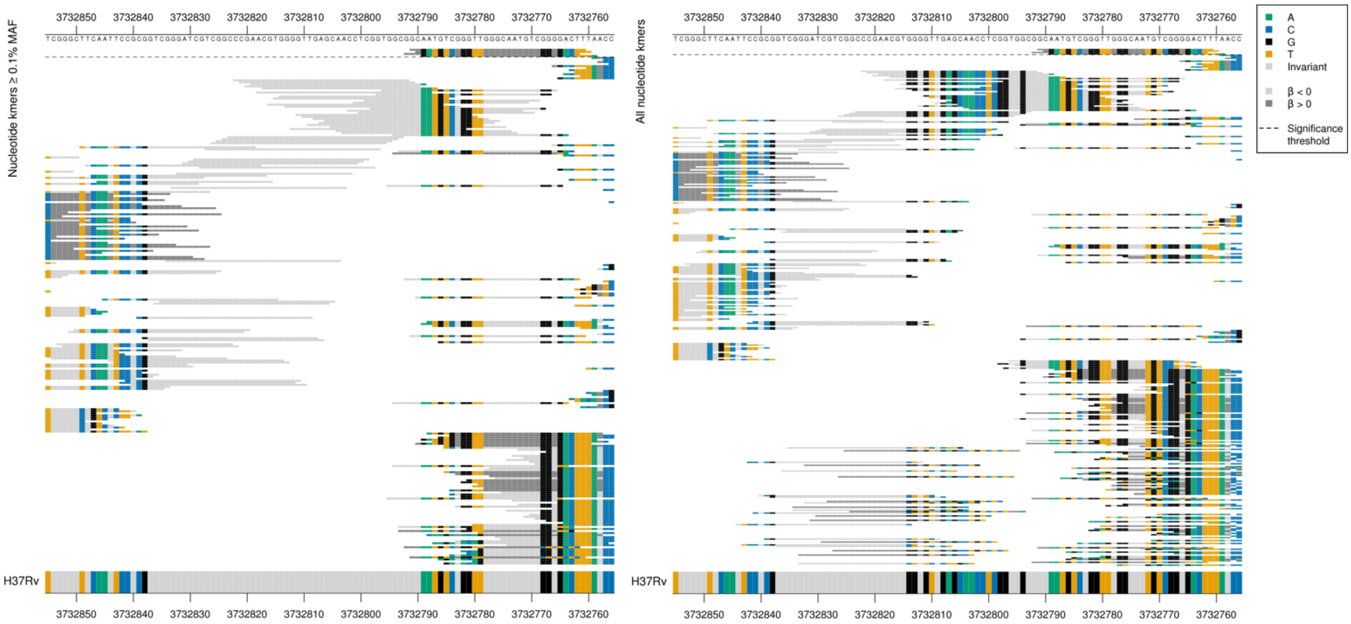

These kmers were significant for the AMI analysis but with an opposite direction of effect.
All significant kmers covered amino acids 1958-1959.

High MIC-associated kmers = 1959D
Kmers below significance threshold = 1959N + others

| Kmer | $-\log_{10} p$ | β | MAC | MAF | Ps | Variants |
|---|---|---|---|---|---|---|
| DVGLGDVGLGN | 10.51 | 1.48 | 18 | 0.22 | 1954 | 1959D |

## Rv0332

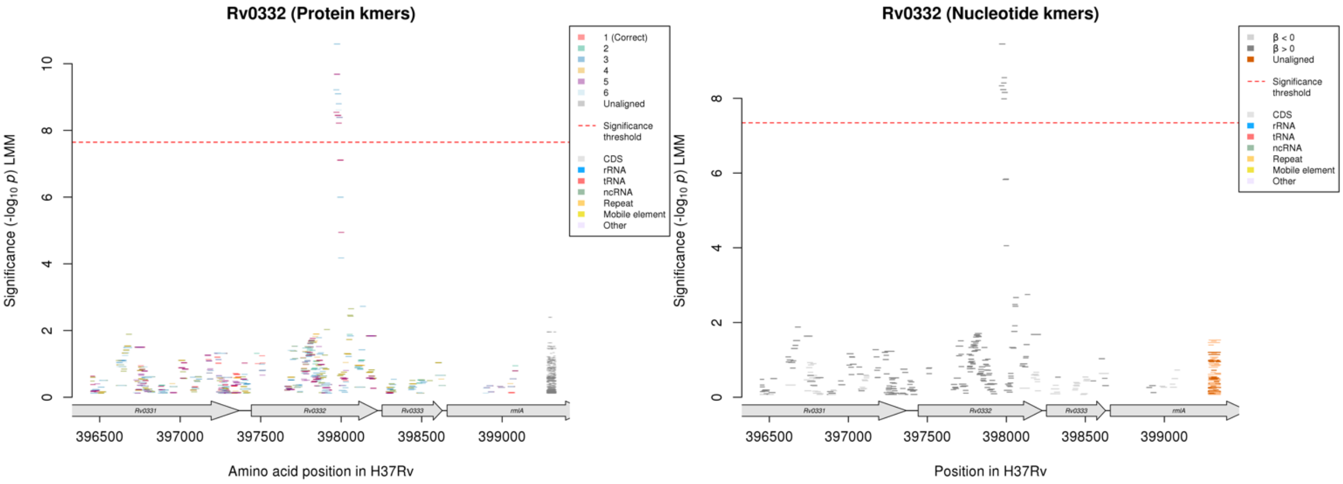

Protein kmers:

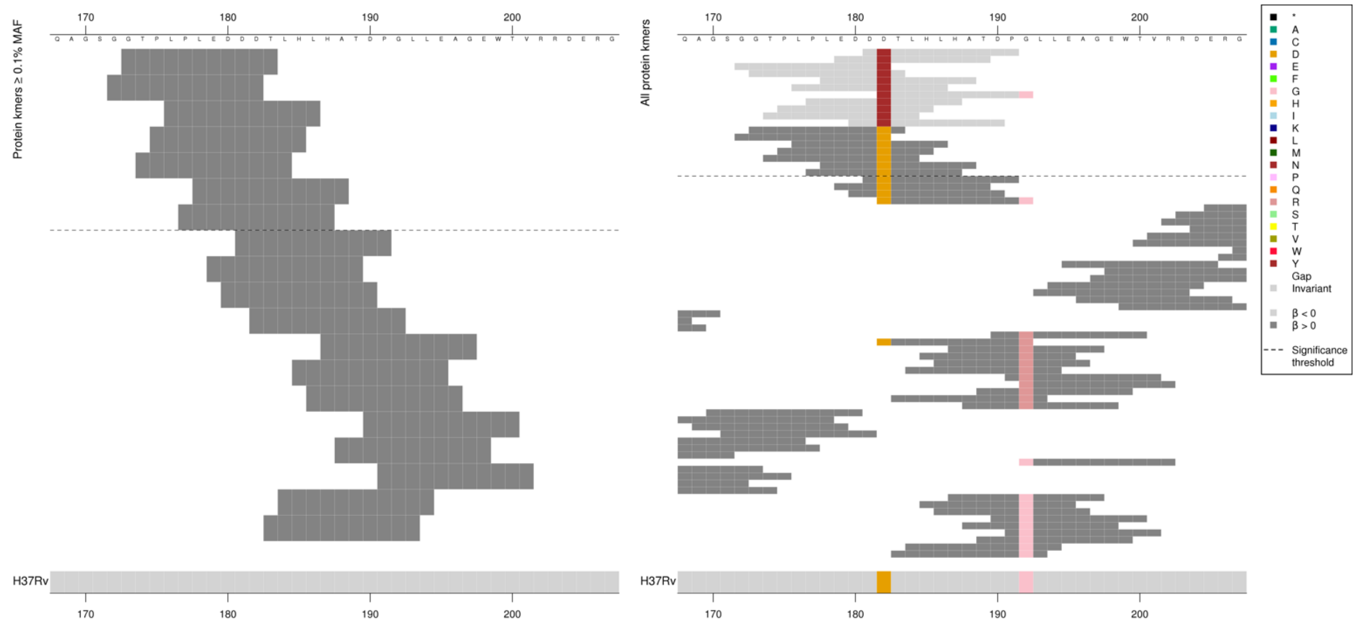

*Rv0332* product = HP

Low MAF kmers = 182N.

| Kmer | -log₁₀ *p* | β | MAC | MAF | Ps | Variants |
|---|---|---|---|---|---|---|
| GTPLPLEDDDT | 9.68 | 1.89 | 12 | 0.15 | 173 | 182D |

# *Rv2019*

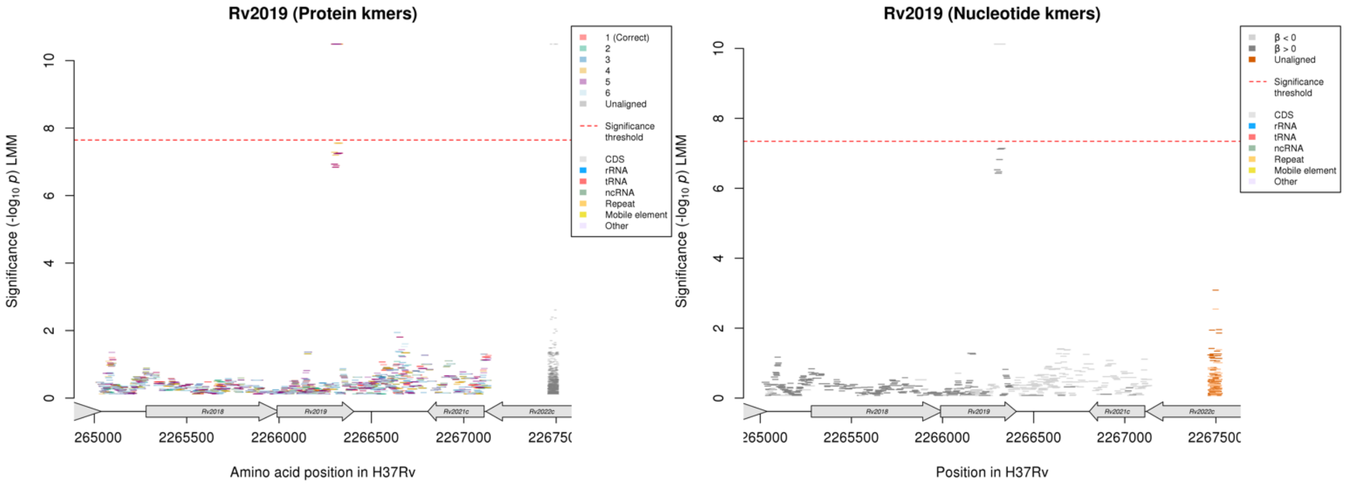

Protein kmers:

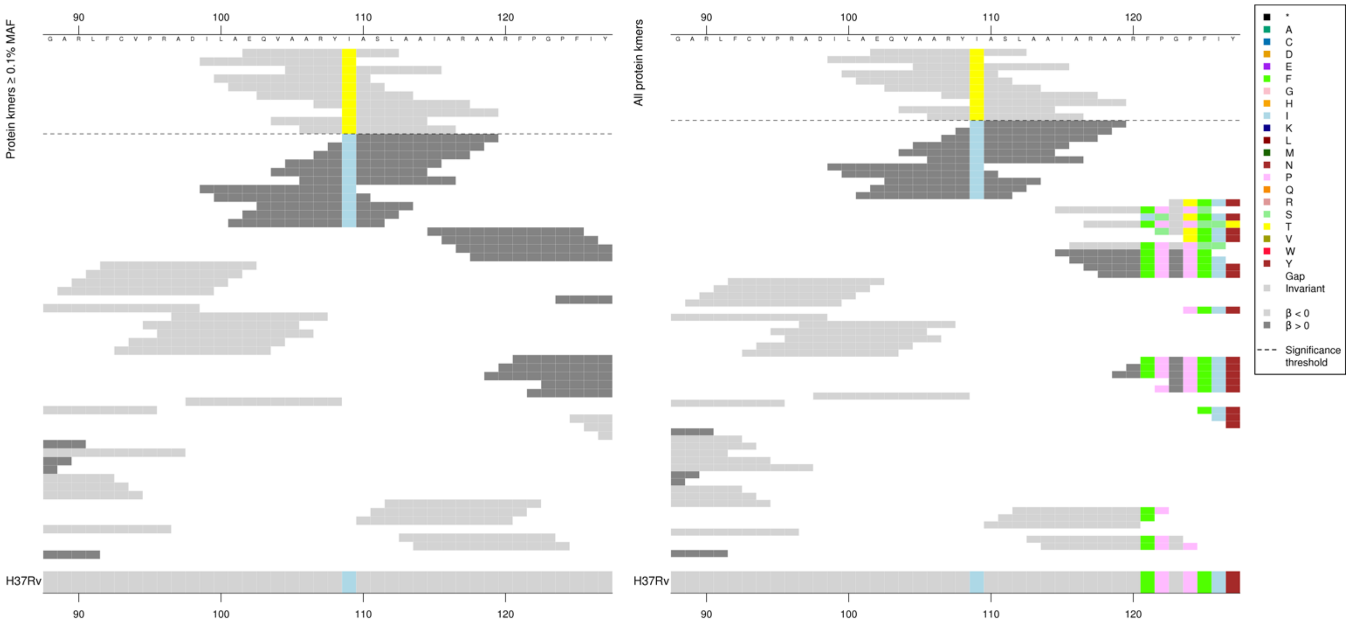

*Rv2019* product = HP.
Low MIC-associated = 109T, Kmers below significance threshold = 109I

| Kmer | -log$_{10}$ *p* | β | MAC | MAF | Ps | Variants |
|---|---|---|---|---|---|---|
| EQVAARYTASL | 10.49 | -1.02 | 114 | 1.42 | 102 | 109T |

#### *vapC22*

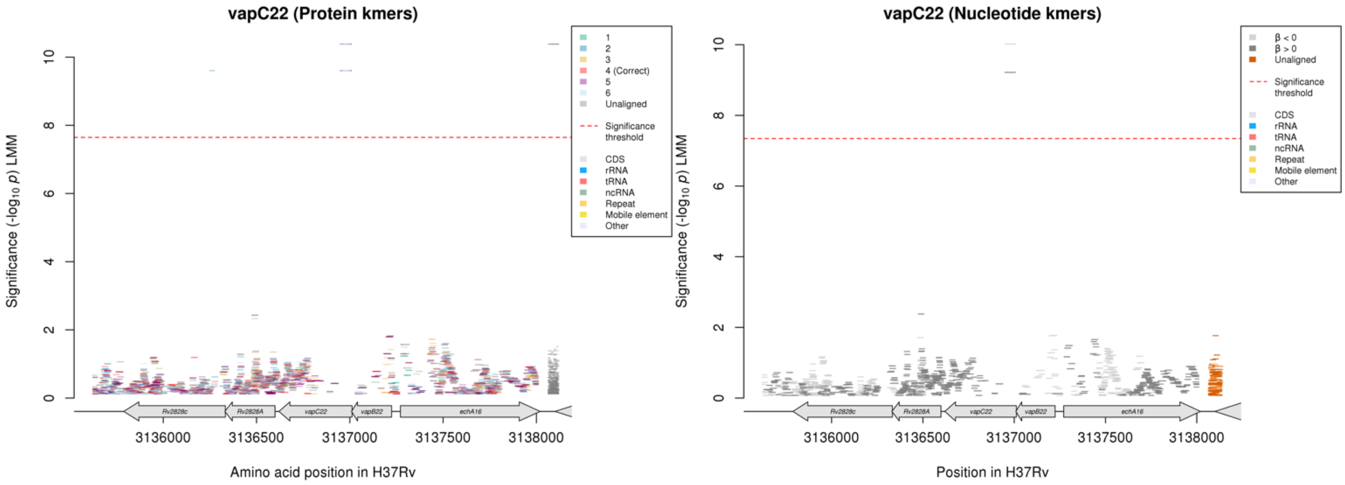

Protein kmers:

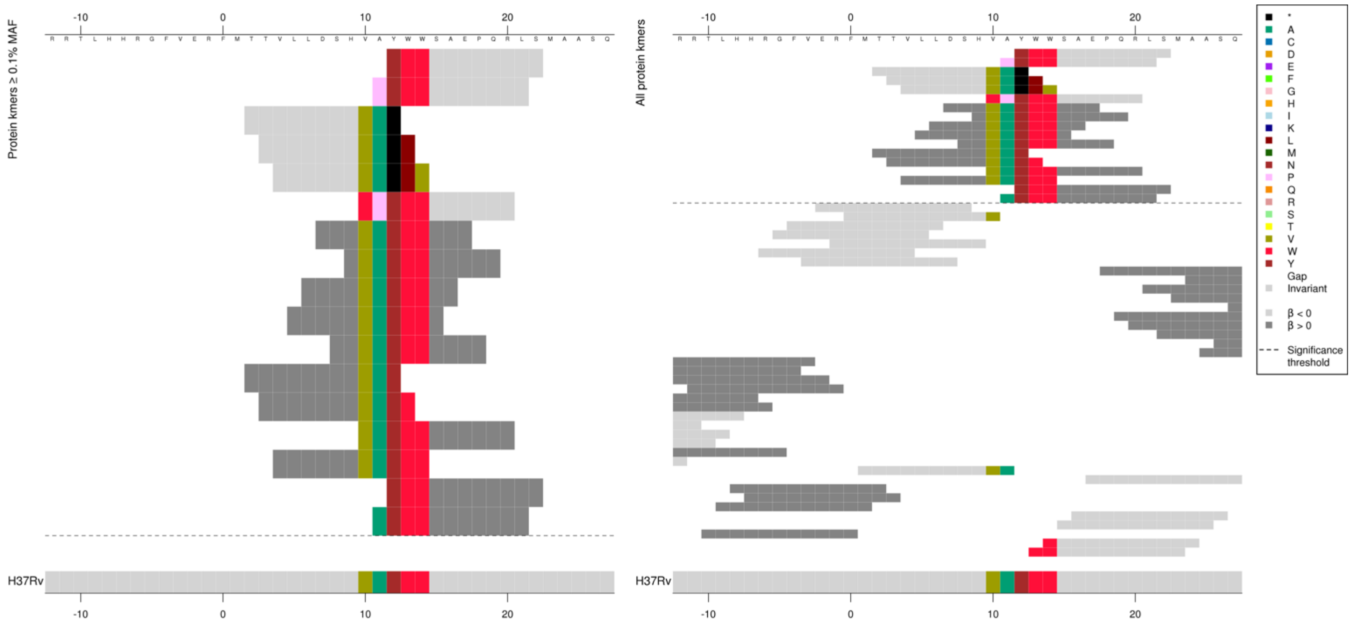

The same region for the nucleotide kmer analysis:

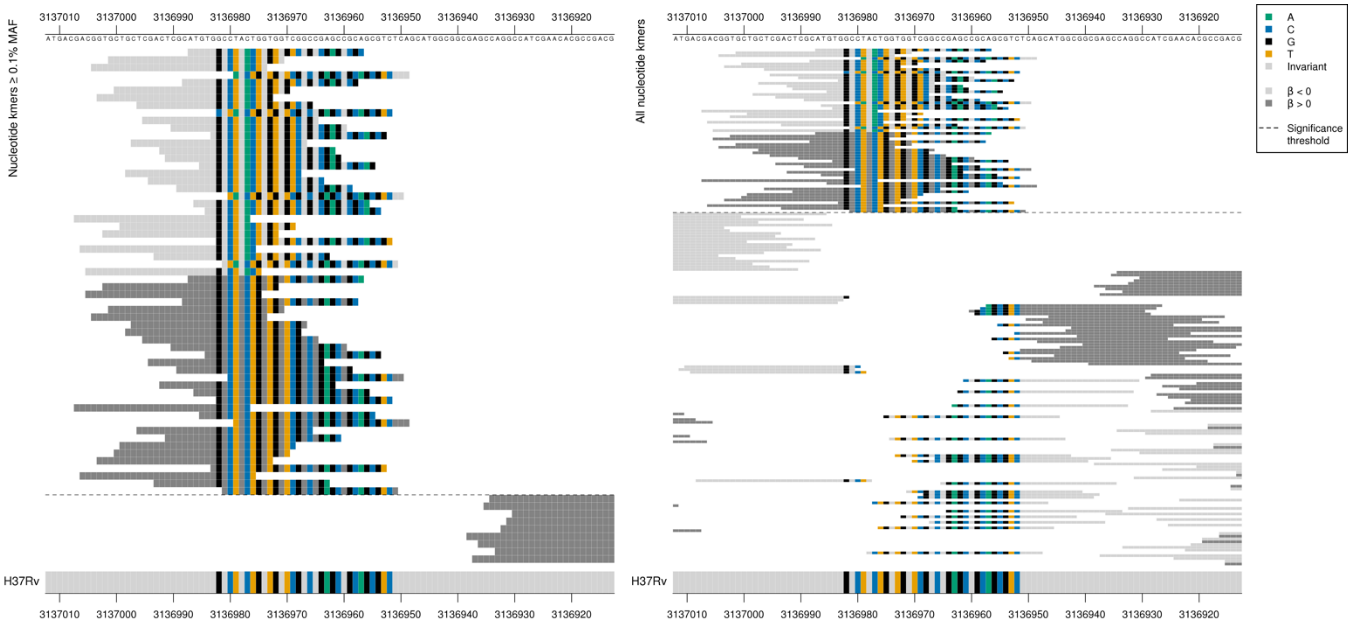

*vapC22* product = ribonuclease VapC22

Low MIC-associated kmers capturing an insertion of an A between positions 3136979-3136977/34-36.

| Kmer | $-\log_{10} p$ | β | MAC | MAF | Ps | Variants |
|---|---|---|---|---|---|---|
| ATGTGGCCTAACTGGTGGTCGGCCGAGCCGC | 10.02 | -1.03 | 113 | 1.41 | 3136987 | A insertion between 3136979-3136977/34-36 |
| ATGTGGCCTACTGGTGGTCGGCCGAGCCGCA | 9.22 | 0.94 | 116 | 1.45 | 3136987 | WT |

# *Rv2896c*

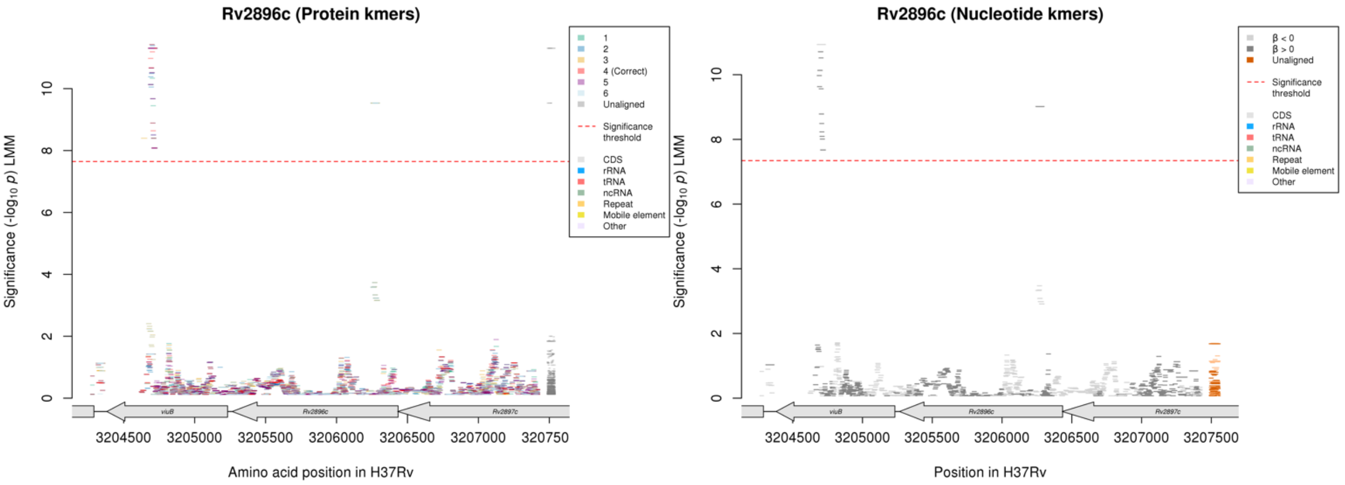

No significant kmers aligned to the correct reading frame of *Rv2896c* in the protein kmer analysis, but there were significant kmers in the nucleotide analysis.
These kmers in *Rv2896c* were also significant for AMI where they have the opposite direction of effect, but they were not significant for KAN.

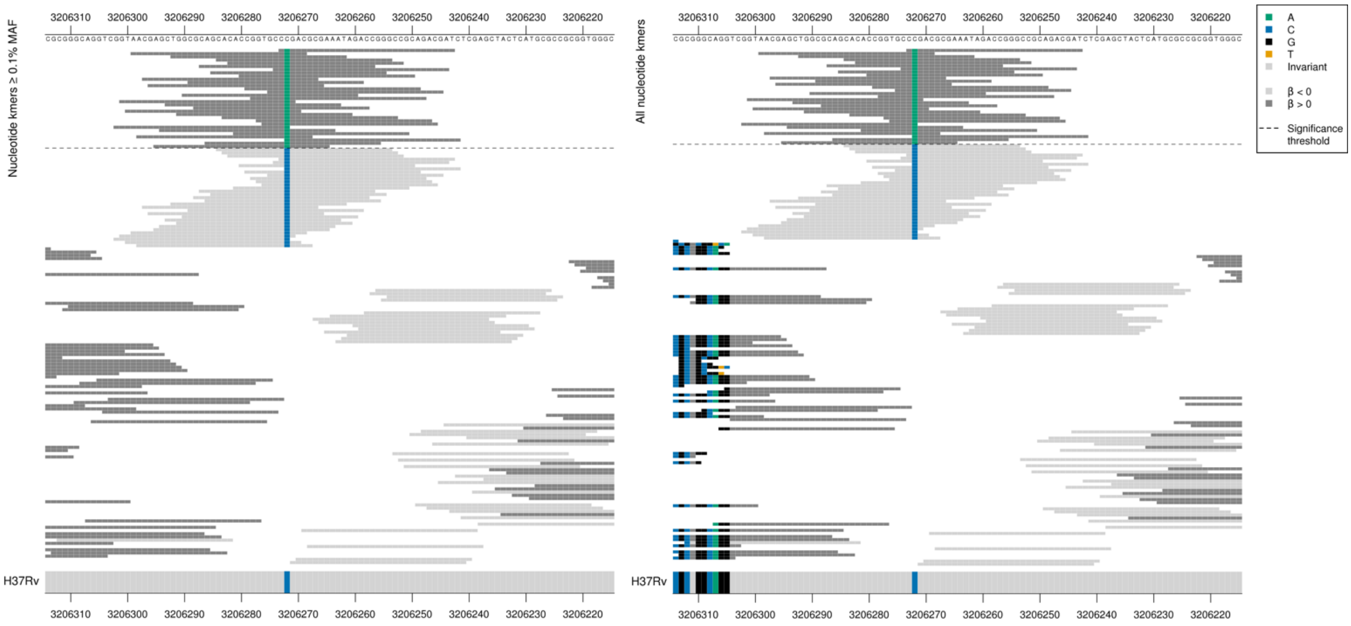

| Kmer | $-\log_{10} p$ | β | MAC | MAF | Ps | Variants |
|---|---|---|---|---|---|---|
| CAGACGCGAAATAGACCGGGCCGCAGACGAT | 9.02 | 1.37 | 18 | 0.22 | 3206273 | 3206272/163A |

### Clofazimine (CFZ)

## *Rv0678*

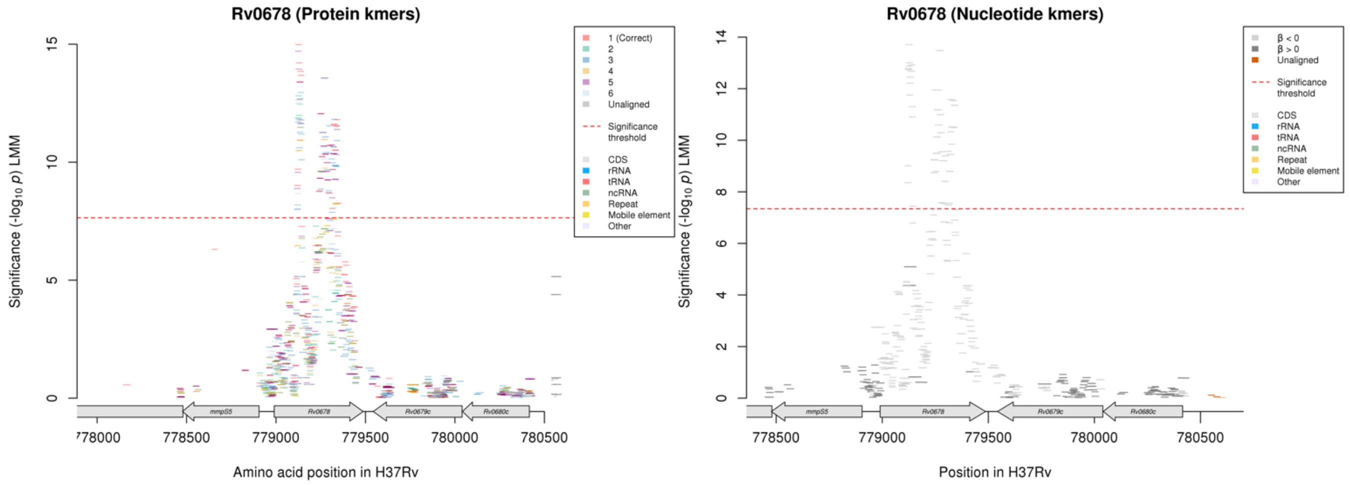

Protein kmers:

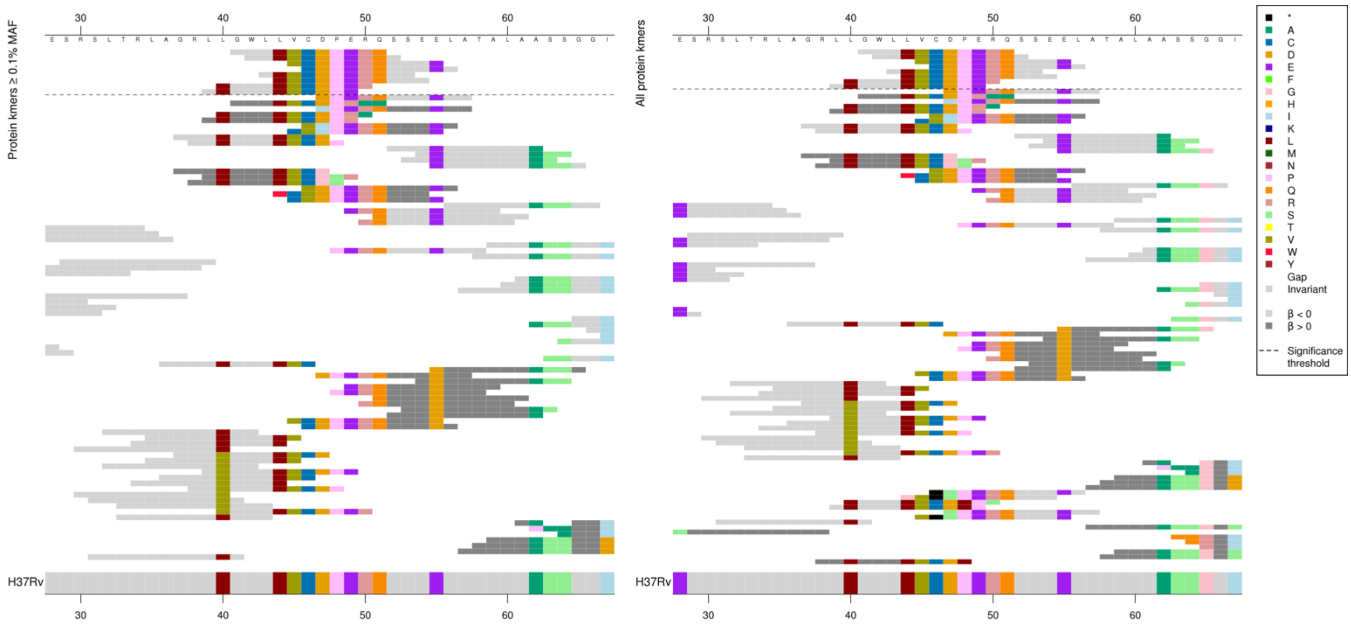

The same region for the nucleotide kmer analysis:

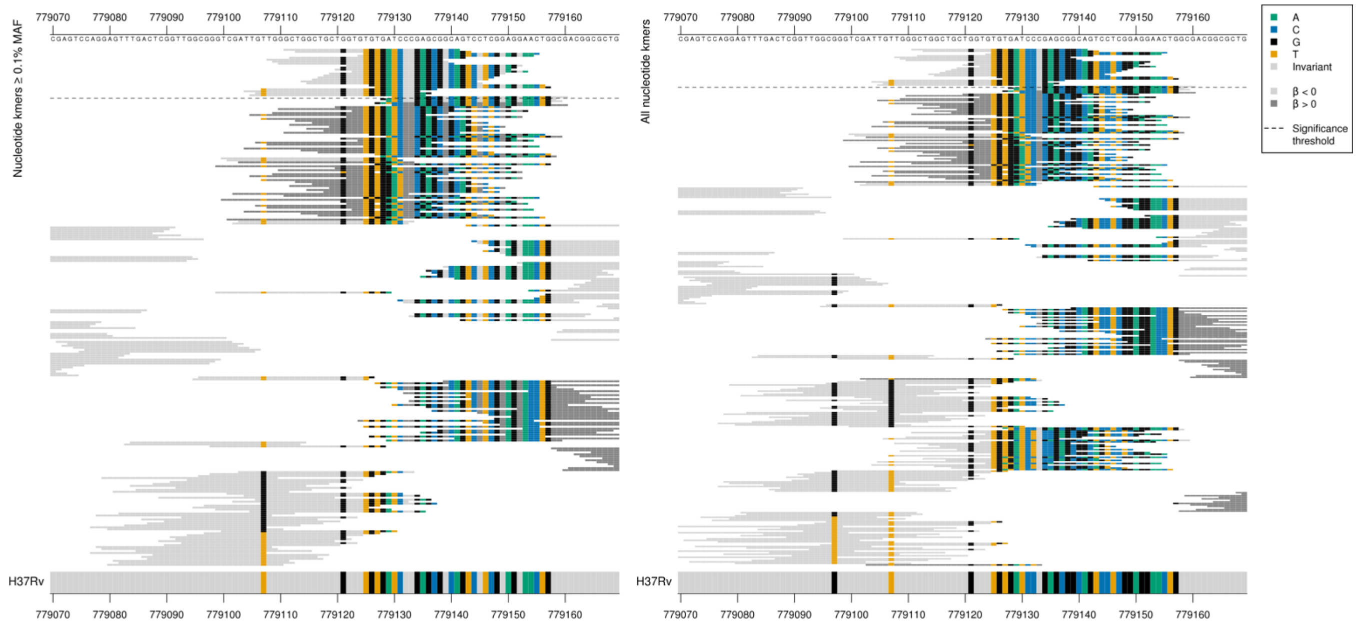

For CFZ, only the low MIC kmers were significantly associated. For BDQ, high MIC kmers were also significantly associated.

| Kmer | $-log_{10}\,p$ | β | MAC | MAF | Ps | Variants |
|---|---|---|---|---|---|---|
| GWLLVCDPERQ | 14.98 | -0.82 | 48 | 0.66 | 41 | WT between amino acids 39-56 |

Protein kmers:

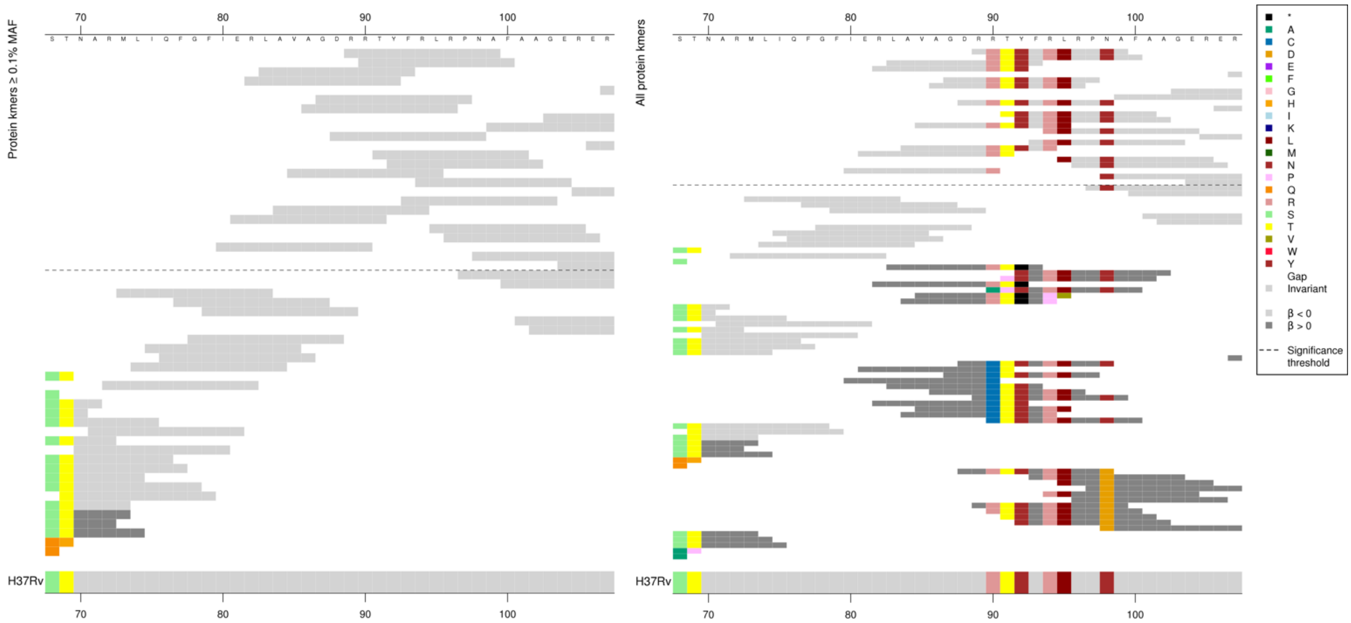

The top low MIC-associated kmers captured the reference sequence, including codons 90R and 98N which were the same as the reference. The other reference sequence kmers covered the same region over a range of *p*-values but not all covered both codons. For BDQ, the high MIC-associated kmers containing 90C were significantly associated.

Kmers below significance threshold = 90C, 98D, indels

| Kmer | $-log_{10}\,p$ | β | MAC | MAF | Ps | Variants |
|---|---|---|---|---|---|---|
| RRTYFRLRPNA | 13.56 | -1.08 | 27 | 0.37 | 89 | WT / 90R, 98N |

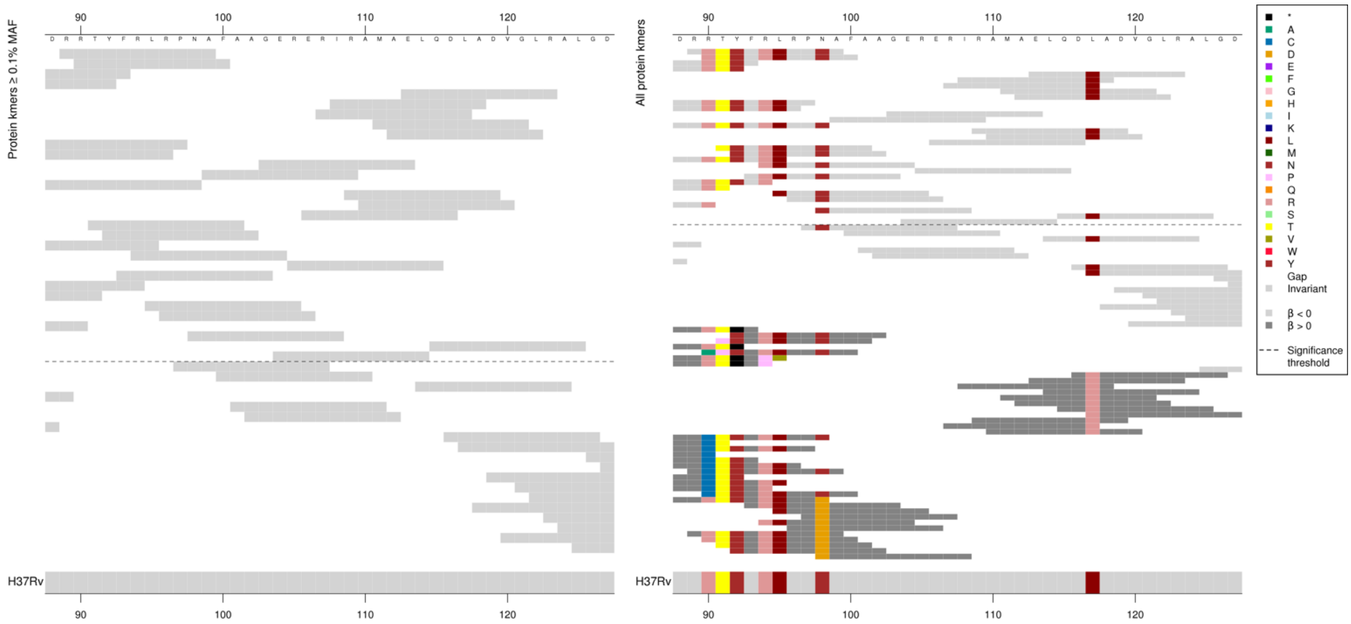

Kmers below significance threshold = 117R

There were also some significant low MIC-associated kmers in the region which were the same as the reference and did not cover codon 117, these were possibly in LD with other variants or were capturing regions containing rare variation not analysed here.

| Kmer | $-\log_{10} p$ | β | MAC | MAF | Ps | Variants |
|---|---|---|---|---|---|---|
| ELQDLADVGLR | 11.80 | -1.29 | 16 | 0.22 | 113 | WT / 117L |

#### *fabG1*

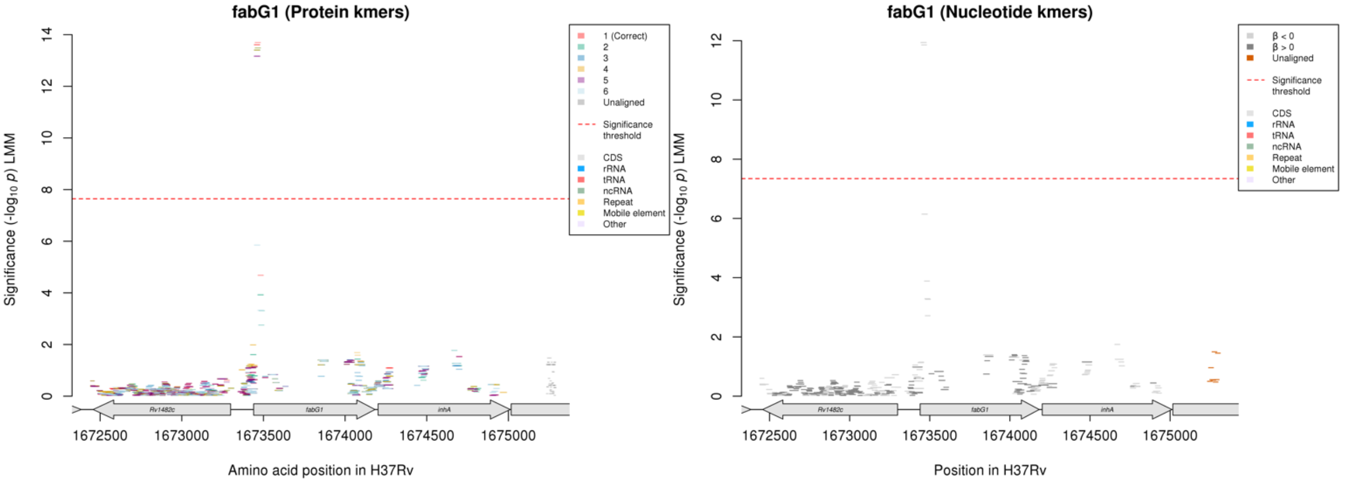

Protein kmers:

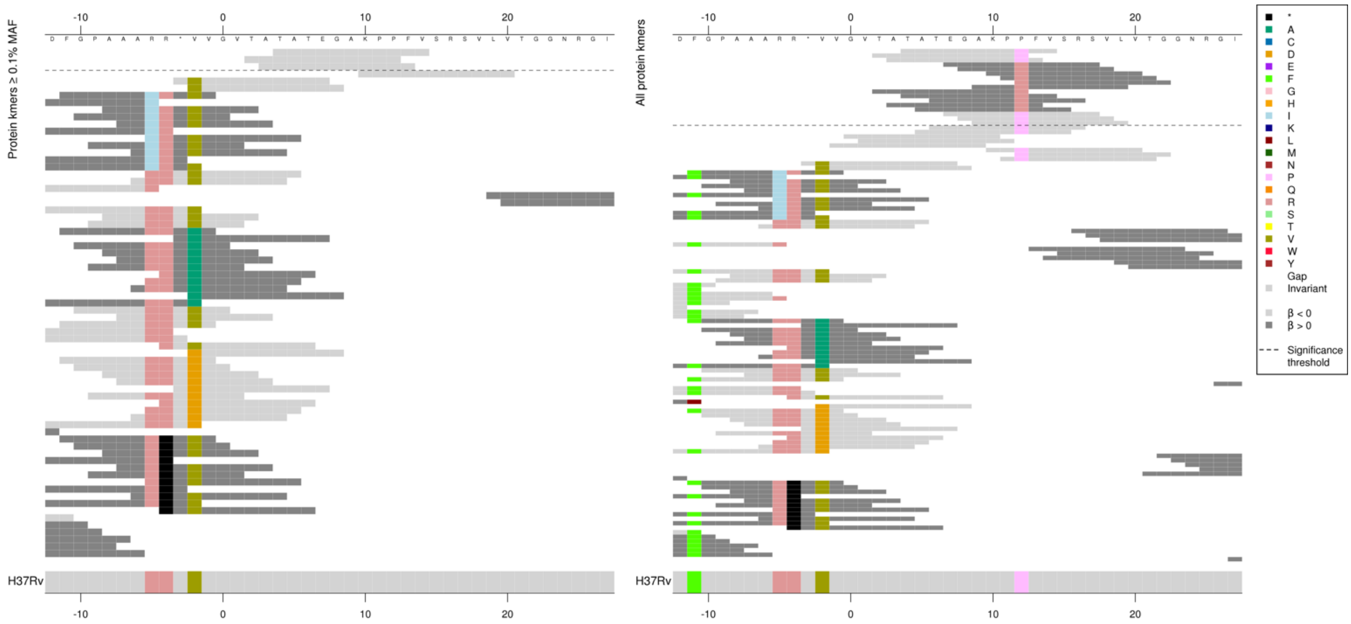

*fabG1* product = 3-oxoacyl-ACP reductase FabG

Capturing a substitution at codon 12, all kmers were low frequency. Codon 12P kmers were present in six-ten genomes.
Low MAF kmers = 12R.

| Kmer | $-\log_{10} p$ | β | MAC | MAF | Ps | Variants |
|---|---|---|---|---|---|---|
| TATEGAKPPFV | 13.69 | -1.88 | 9 | 0.12 | 4 | 12P |

### *cyp142*

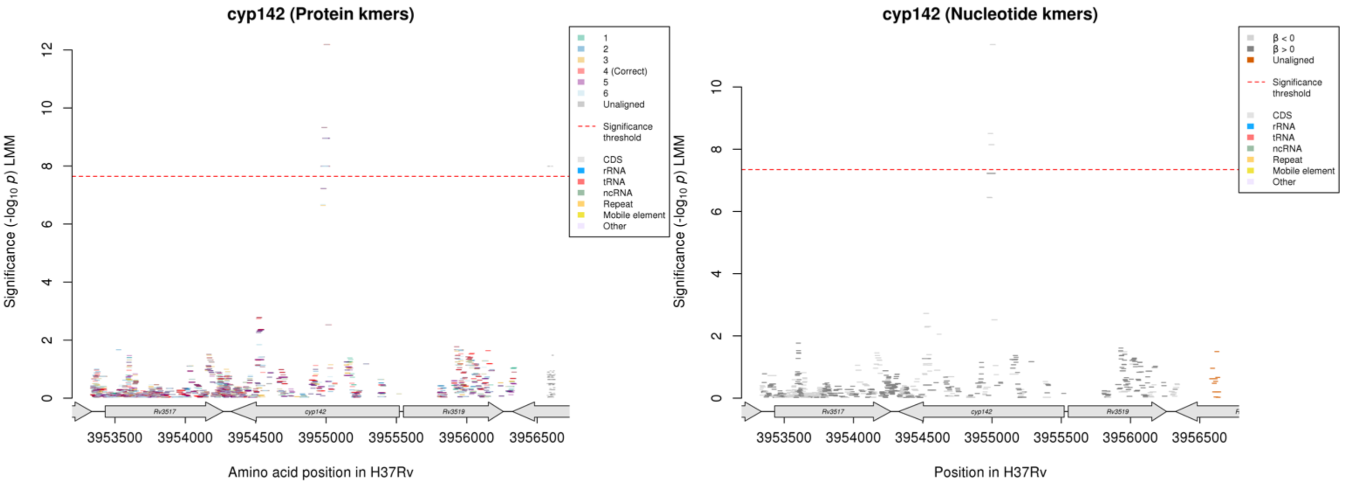

Protein kmers:

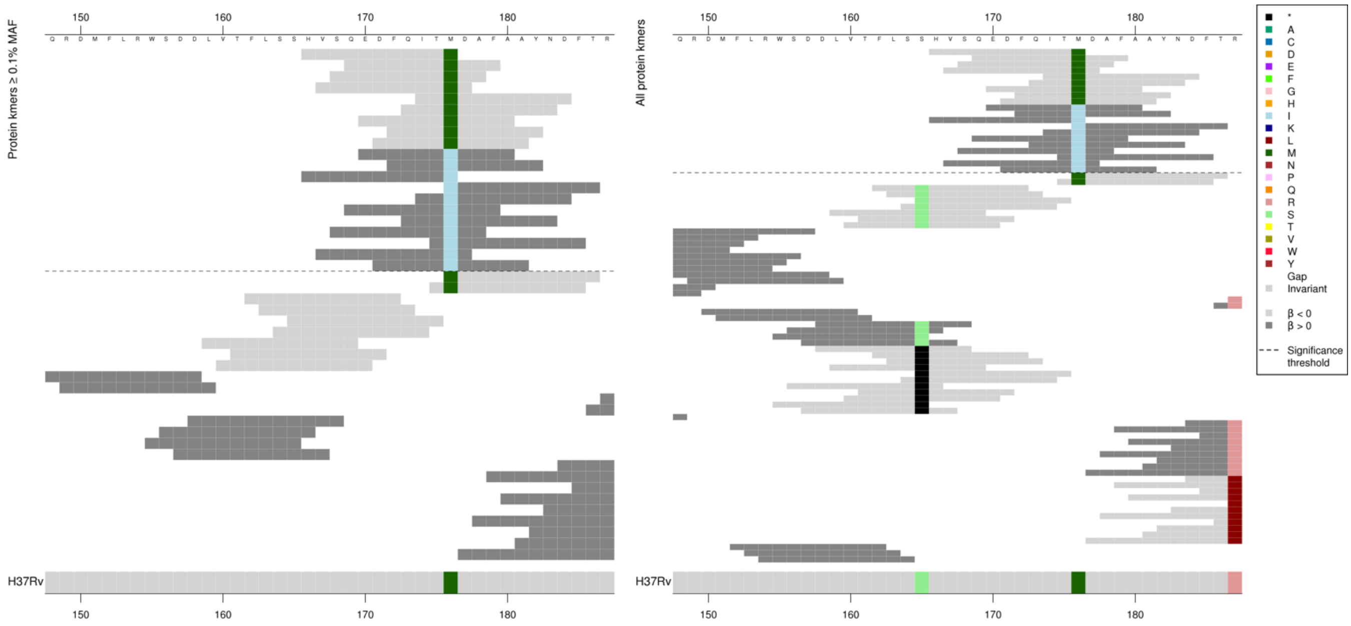

Removing the threshold requiring a kmer to be aligned to a gene/intergenic region in at least five genomes across the full dataset. Showing the protein kmer alignment after lowering the threshold to just one genome:

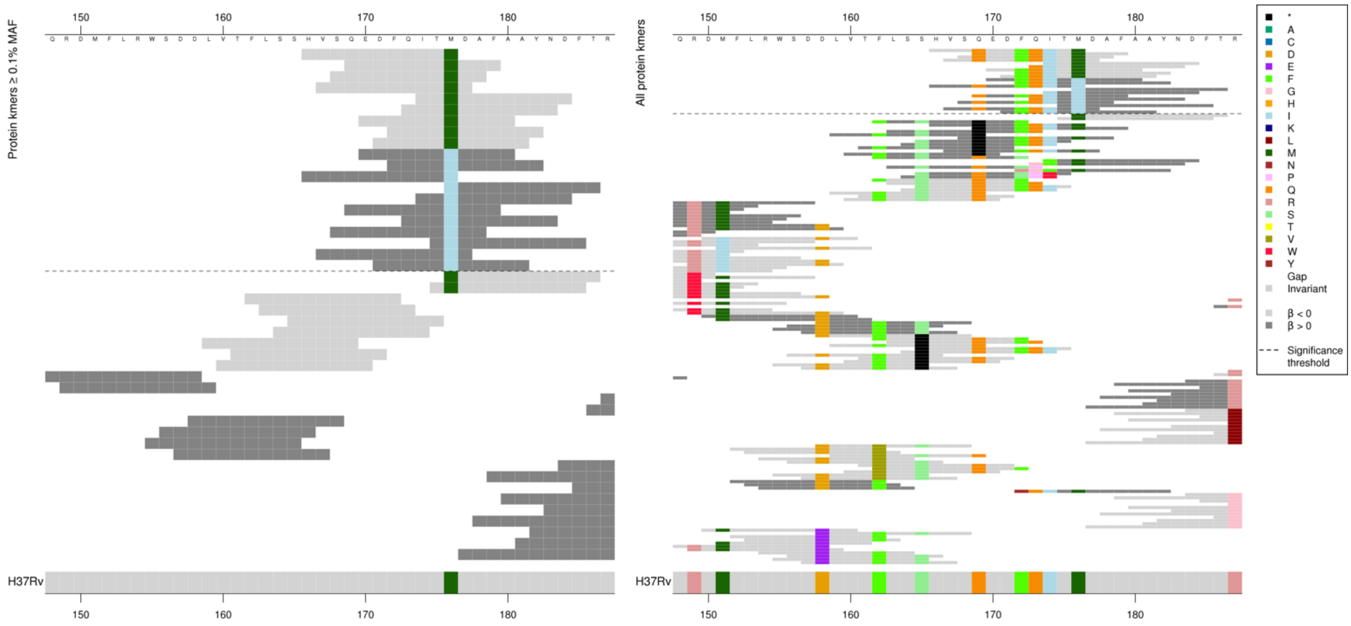

Nucleotide kmers:

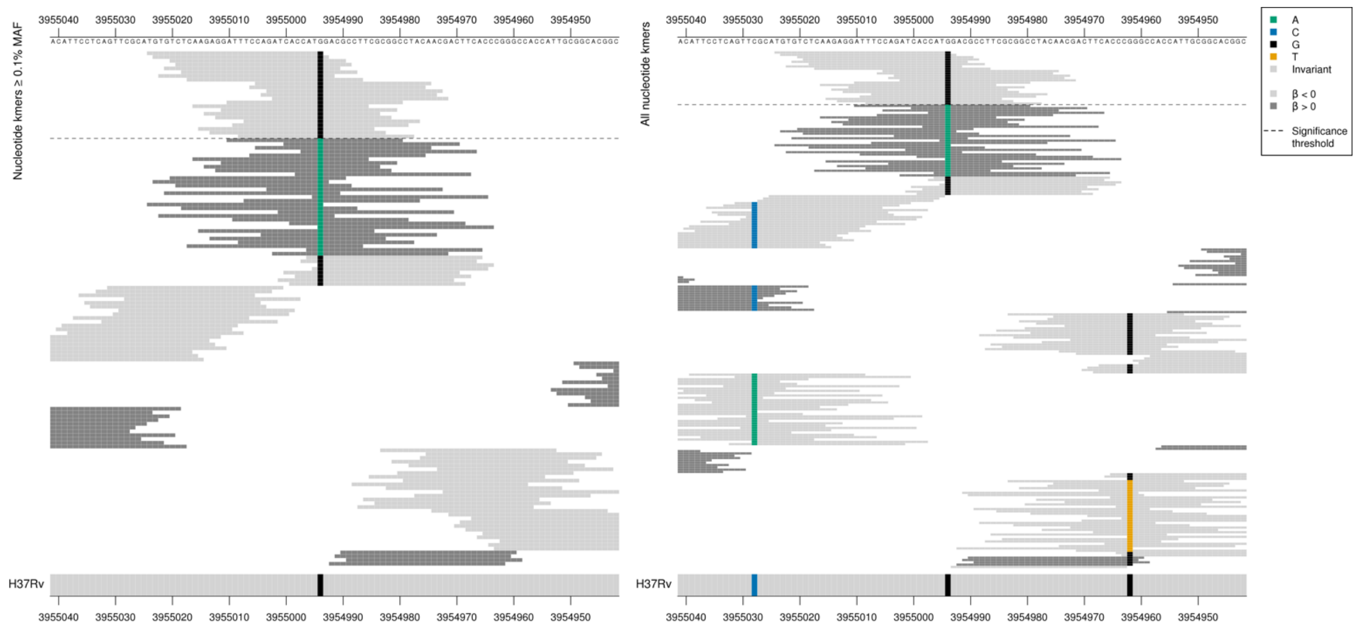

Removing the threshold requiring a kmer to be aligned to a gene/intergenic region in at least five genomes across the full dataset. Showing the nucleotide kmer alignment after lowering the threshold to just one genome:

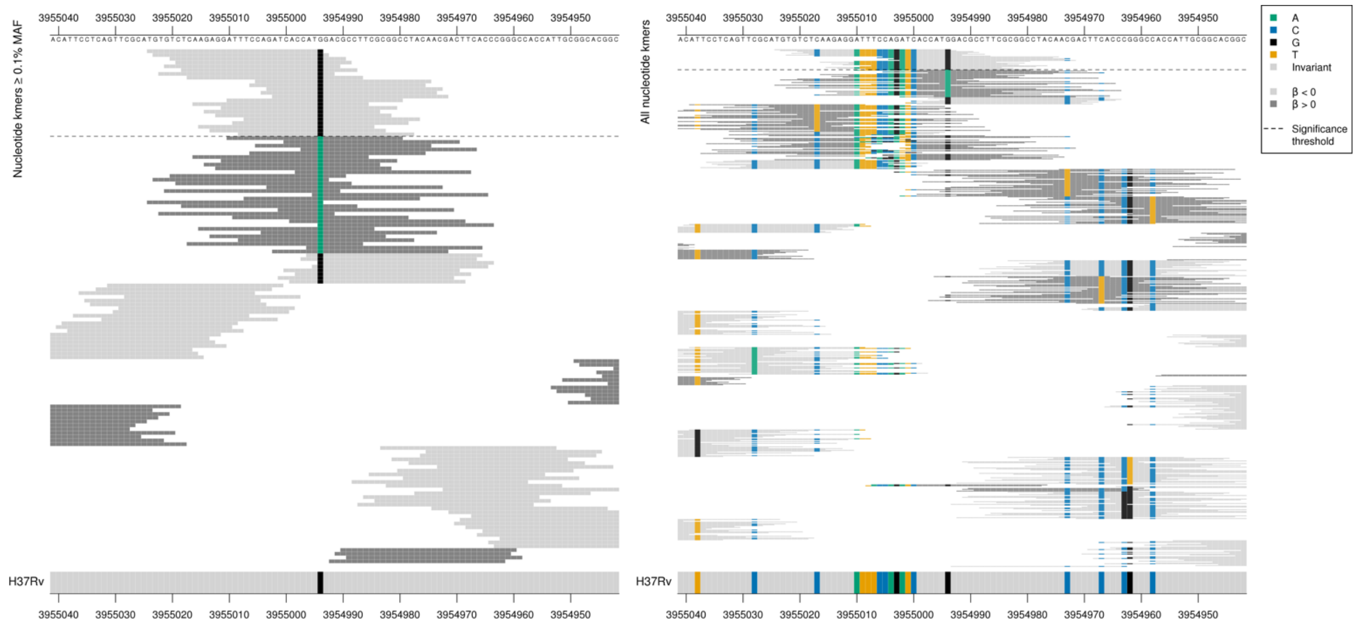

The kmers below the significance threshold capturing the stop codon were present in just two isolates (e.g. FLSSHVS*EDF, $-\log_{10}p$ = 4.89, beta = 2.24, MAC = 2)
Kmers capturing the indels shown by the nucleotide kmers (e.g. kmer ATGTGTCTCAAGAGGATTCACCATGGACGCC) were also present in just two isolates ($-\log_{10}p$ = 2.90, beta = 1.68, MAC = 2)

| Kmer | $-\log_{10} p$ | β | MAC | MAF | Ps | Variants |
|---|---|---|---|---|---|---|
| HVSQEDFQITM | 12.18 | -1.19 | 106 | 1.45 | 166 | 176M |
| EDFQITIDAFA | 7.99 | 1.14 | 100 | 1.37 | 170 | 176I |

## *Rv3183:Rv3188*$^{R}$

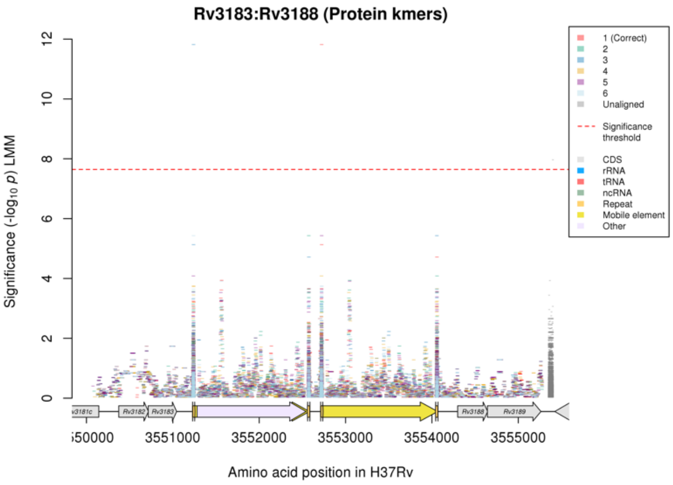

No significant kmers in the nucleotide kmer analysis.
Repeat regions at the ends of a mobile element. One significant kmer above the significant threshold and MAF threshold.

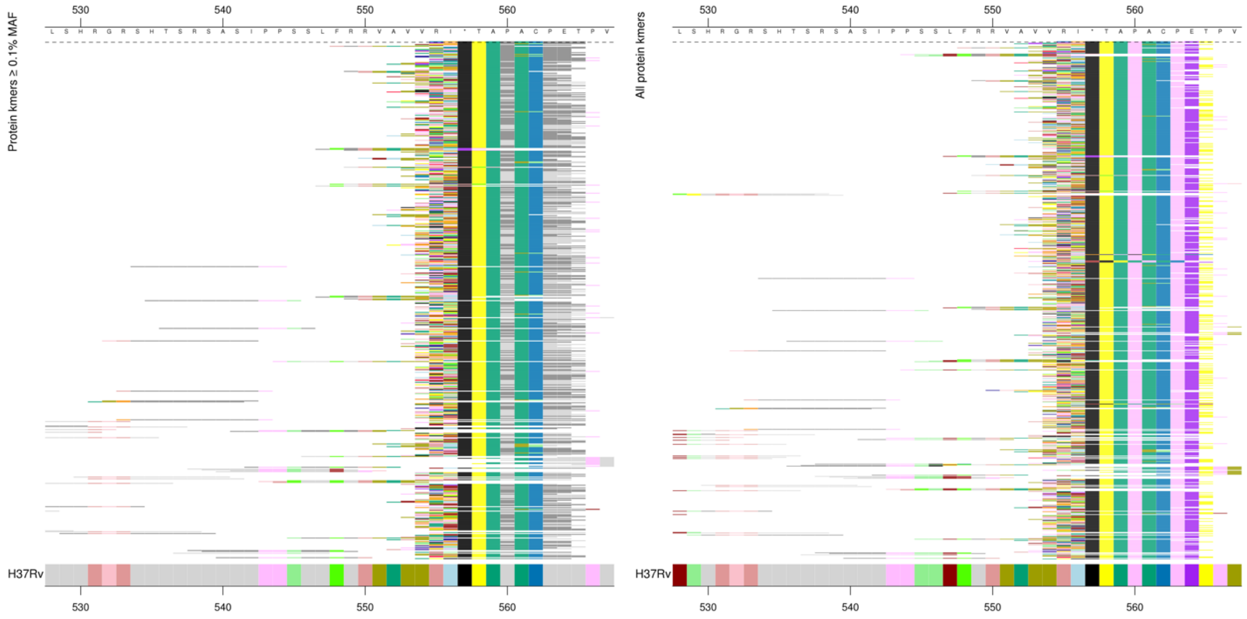

| Kmer | -log$_{10}$ *p* | β | MAC | MAF | Ps | Variants |
|---|---|---|---|---|---|---|
| CH*TAPACPET | 11.82 | 1.23 | 104 | 1.43 | 555 | Repeat region – kmer out of frame as in intergenic region |

### *moaC3:Rv3327*[R]

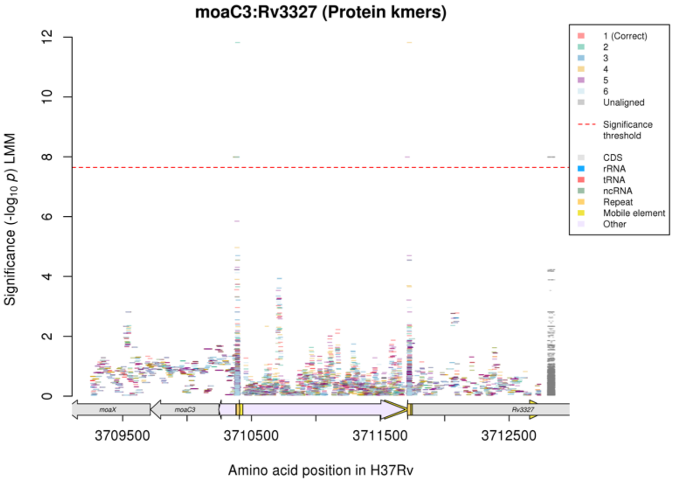

No significant kmers in the nucleotide kmer analyses.
Repeat regions at the ends of a mobile element.
Two kmers above the significance threshold.

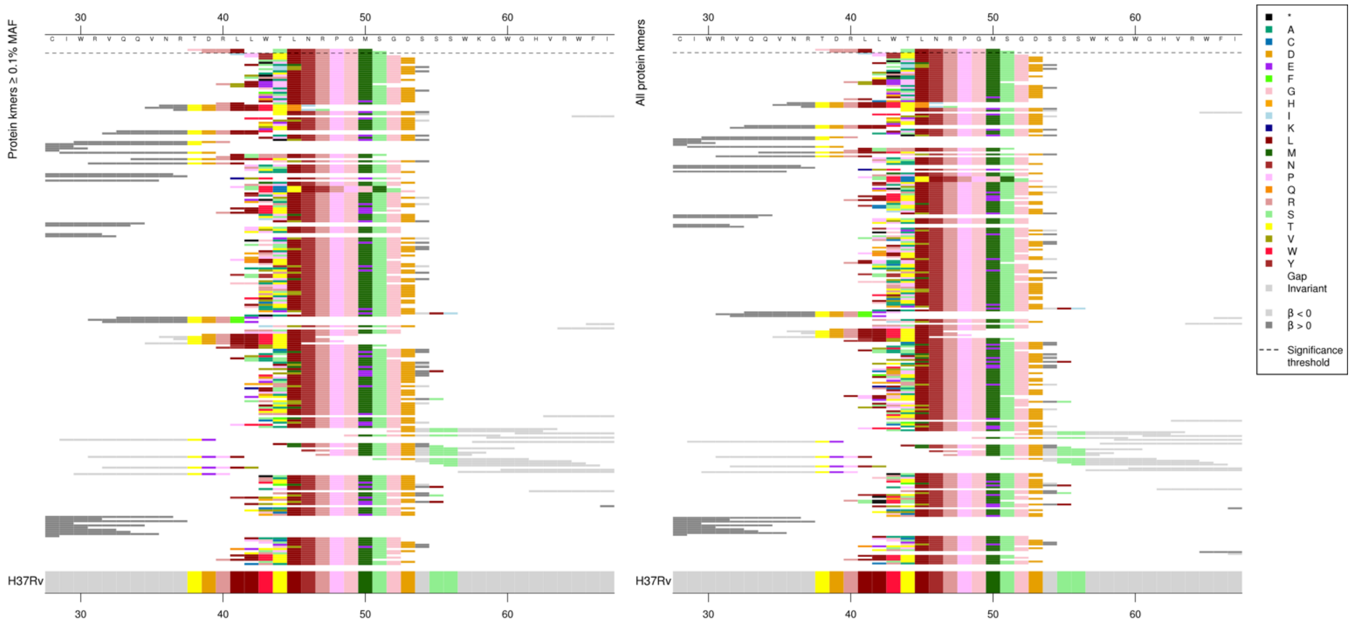

| Kmer | $-\log_{10} p$ | β | MAC | MAF | Ps | Variants |
|---|---|---|---|---|---|---|
| GRRLSLNRPGM | 7.99 | 1.14 | 100 | 1.37 | 38 | Repeat region – kmer out of frame as in intergenic region |

### *dxs2:Rv3382c* [R]

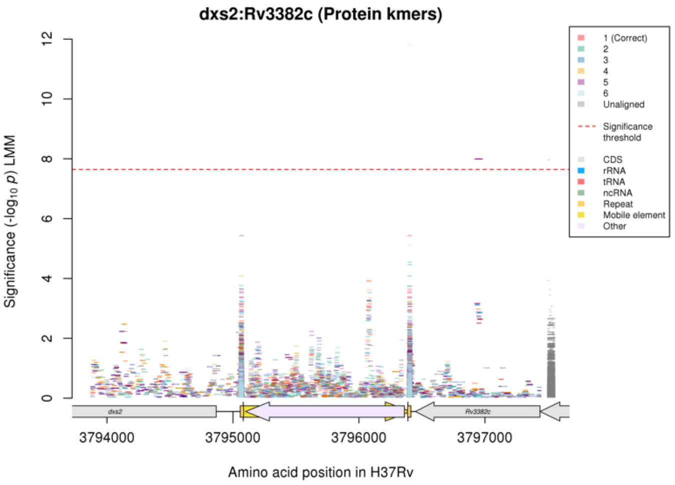

No significant kmers in the nucleotide kmer analysis.
Repeat regions at the ends of a mobile element.

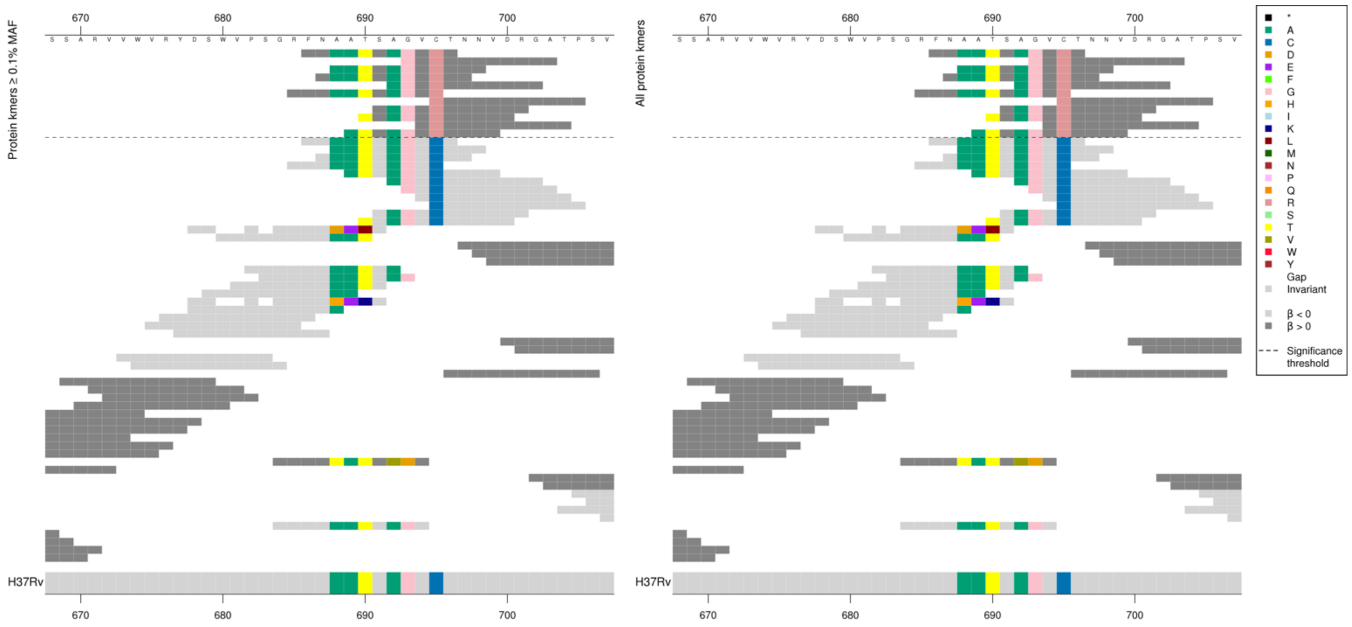

| Kmer | $-\log_{10} p$ | β | MAC | MAF | Ps | Variants |
|---|---|---|---|---|---|---|
| FNAATSAGVRT | 7.99 | 1.14 | 100 | 1.37 | 686 | Kmer out of frame as in intergenic region |

### *mmsA*

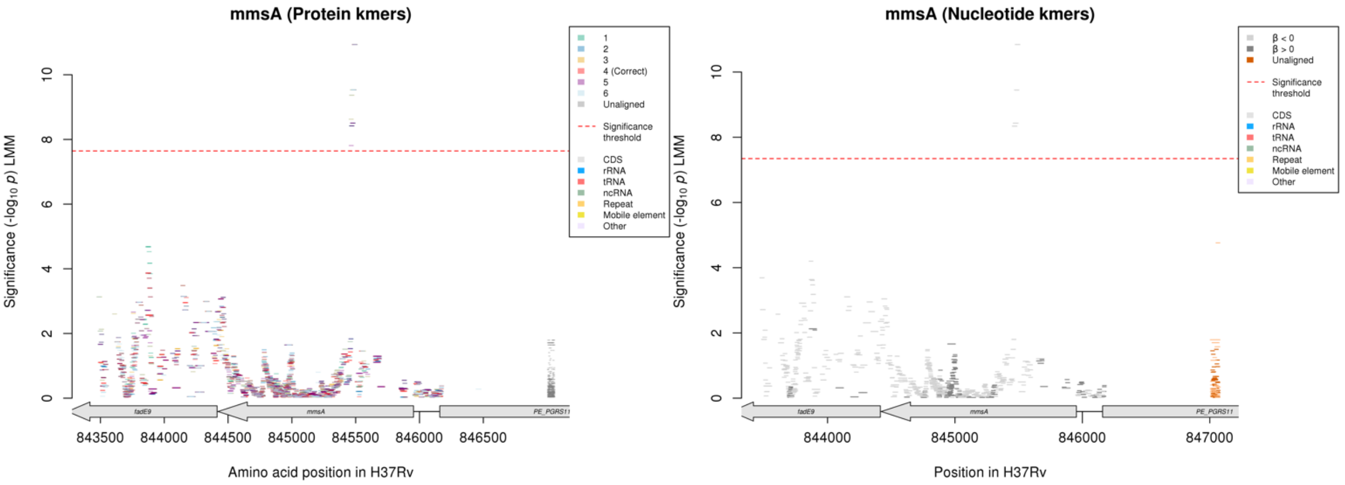

No significant protein kmers align to the correct reading frame.

Nucleotide kmers:

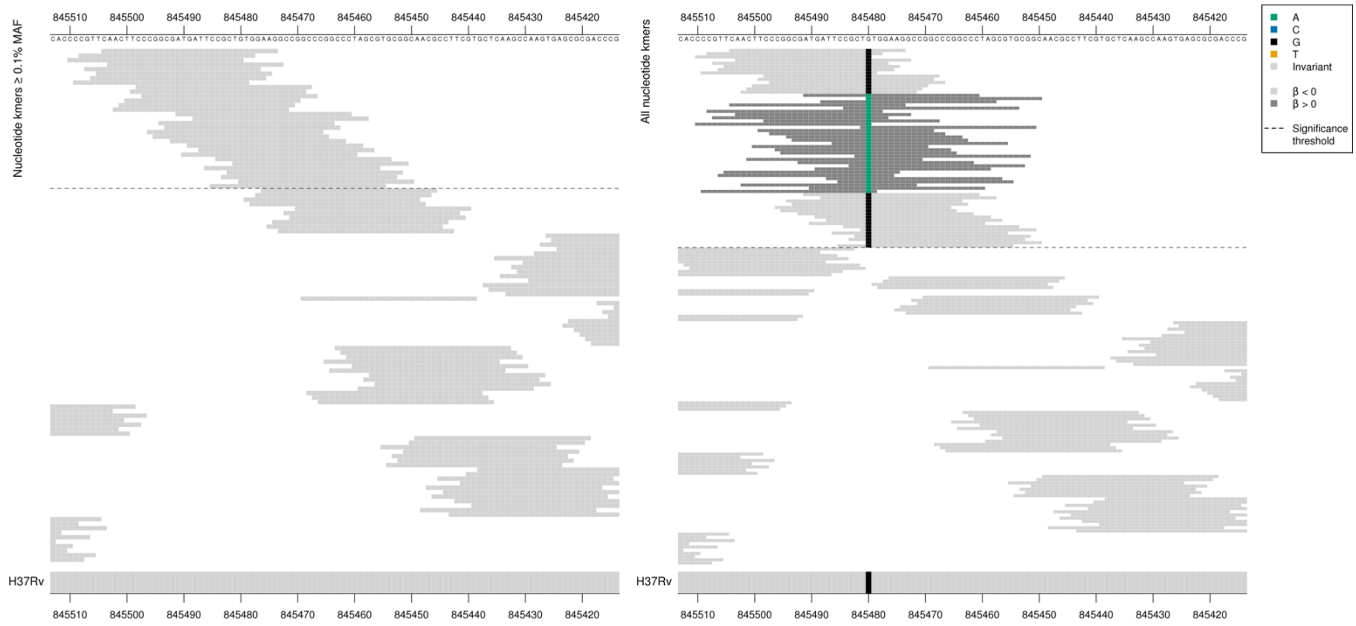

Low MIC-associated kmers = CTG = Leu / L

High MIC-associated (low frequency) kmers = CTA = Leu / L

Synonymous mutation at position 845480/474.

Low MAF kmers = 845480/474A.

| Kmer | $-\log_{10} p$ | β | MAC | MAF | Ps | Variants |
|---|---|---|---|---|---|---|
| CAACTTCCCGGCGATGATTCCGCTGTGGAAG | 10.84 | -1.74 | 8 | 0.11 | 845504 | 845480/474G |

# *Rv3723:Rv3725*

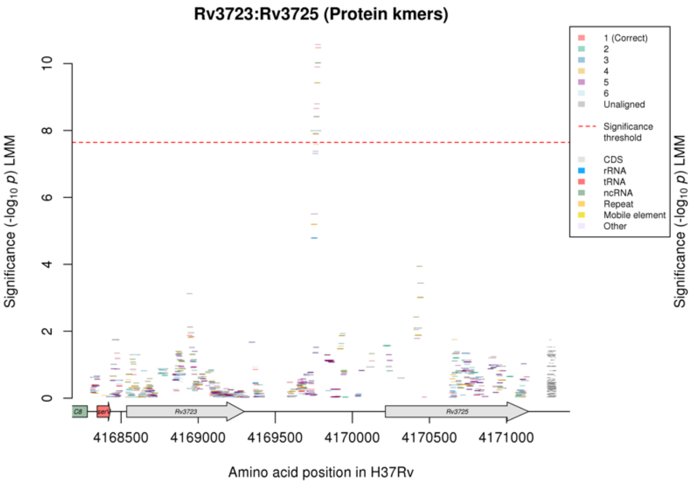

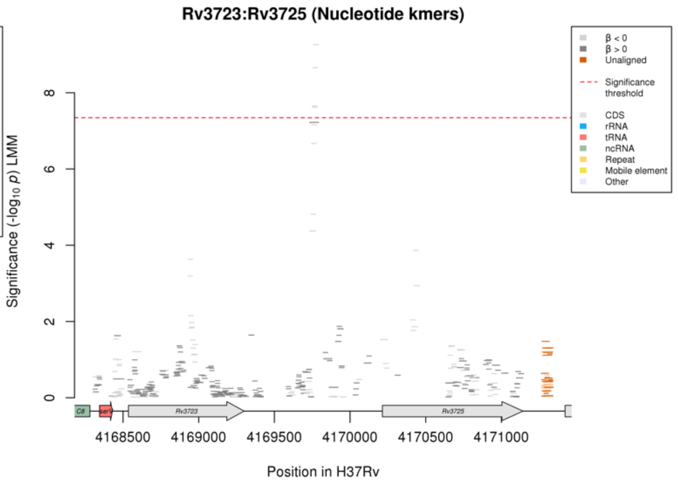

#### Nucleotide kmers:

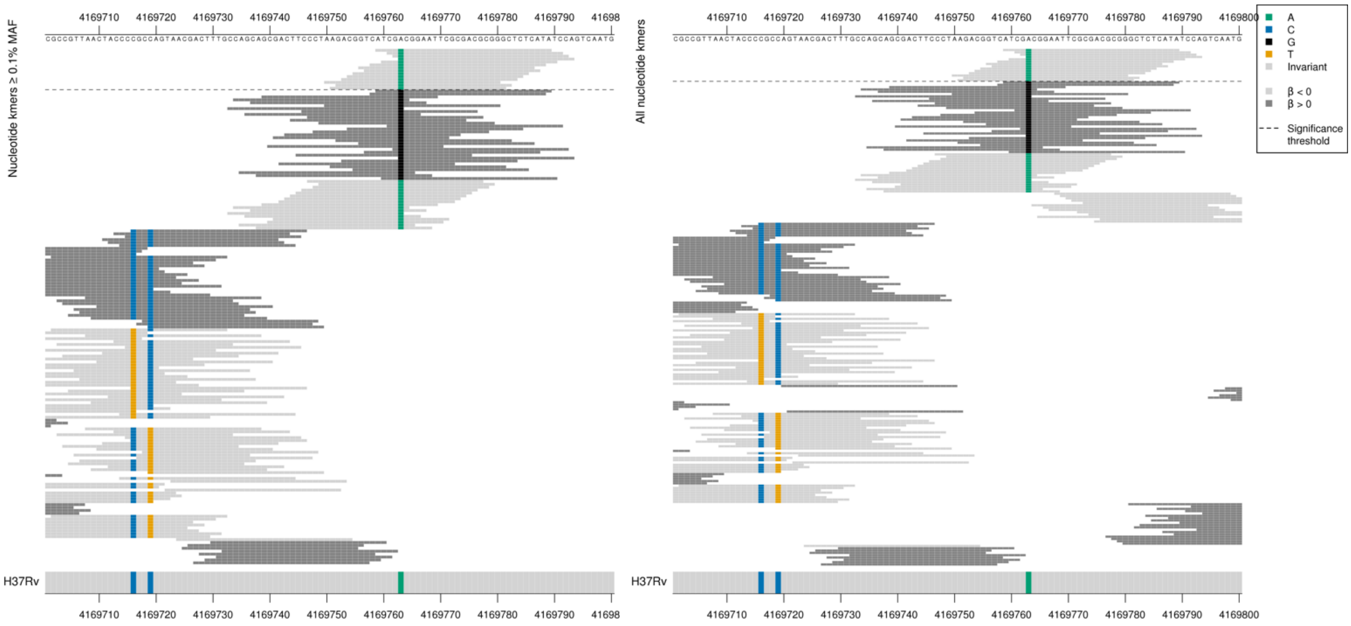

| Kmer | -log$_{10}$ *p* | β | MAC | MAF | Ps | Variants |
|---|---|---|---|---|---|---|
| ATCGACGGAATTCGCGACGCGGGCTCTCATA | 9.27 | -1.21 | 102 | 1.40 | 4169759 | 4169763A |

### *gid*

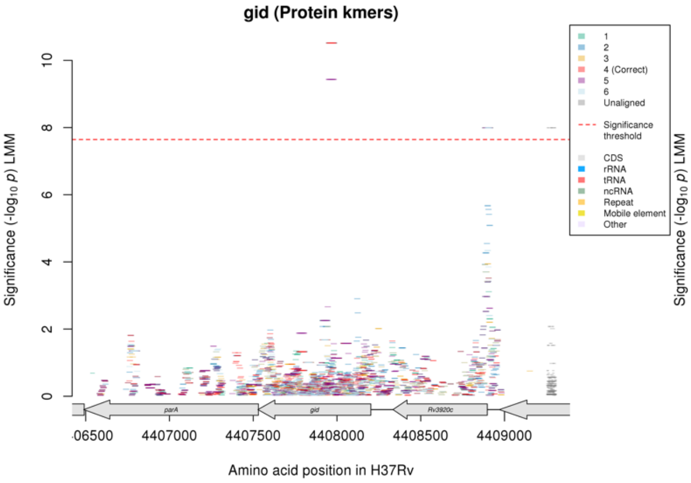

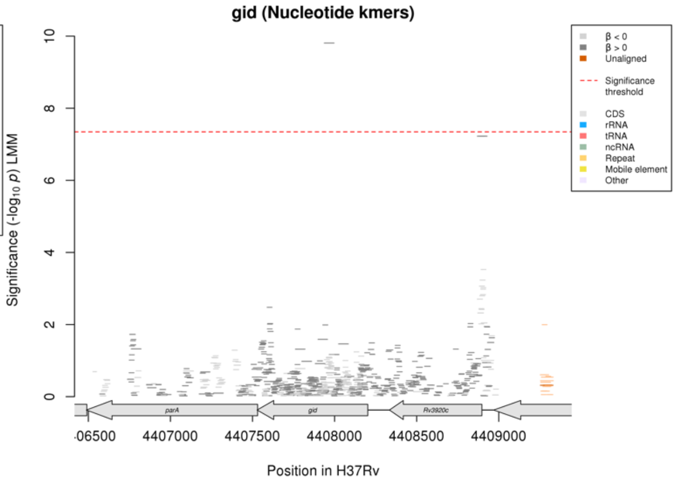

### Clofazimine (CFZ)

Protein kmers:

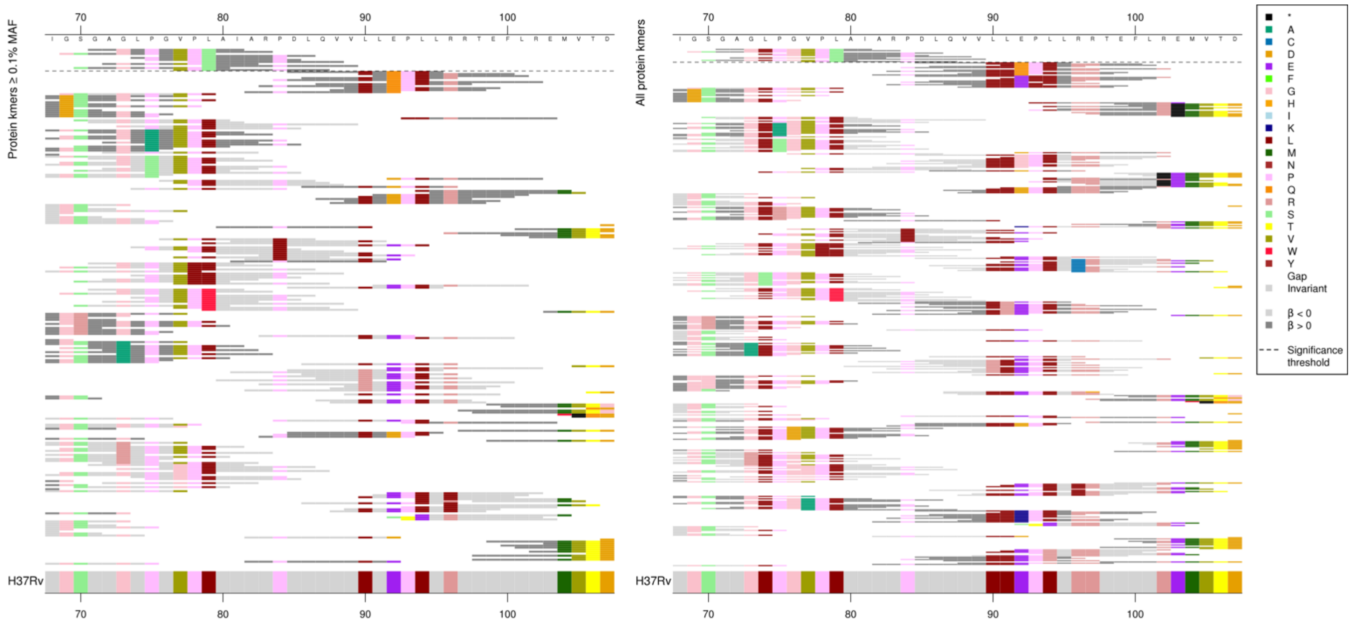

High MIC-associated kmers = 79S

Kmers below significance threshold = 79L/79W

| Kmer | $-\log_{10} p$ | β | MAC | MAF | Ps | Variants |
|---|---|---|---|---|---|---|
| GAGLPGVPSAI | 10.52 | 1.11 | 104 | 1.43 | 71 | 79S |

#### *rpoB*

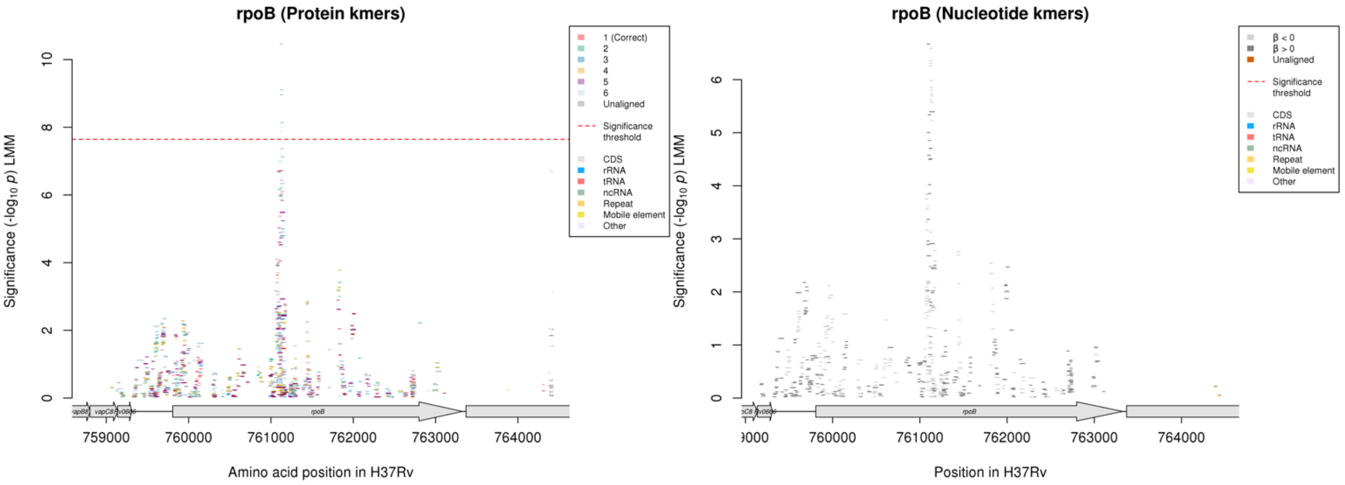

None of the significant protein kmers align to the correct reading frame.

Nucleotide kmers fall just below the significance threshold with a peak in the same region:

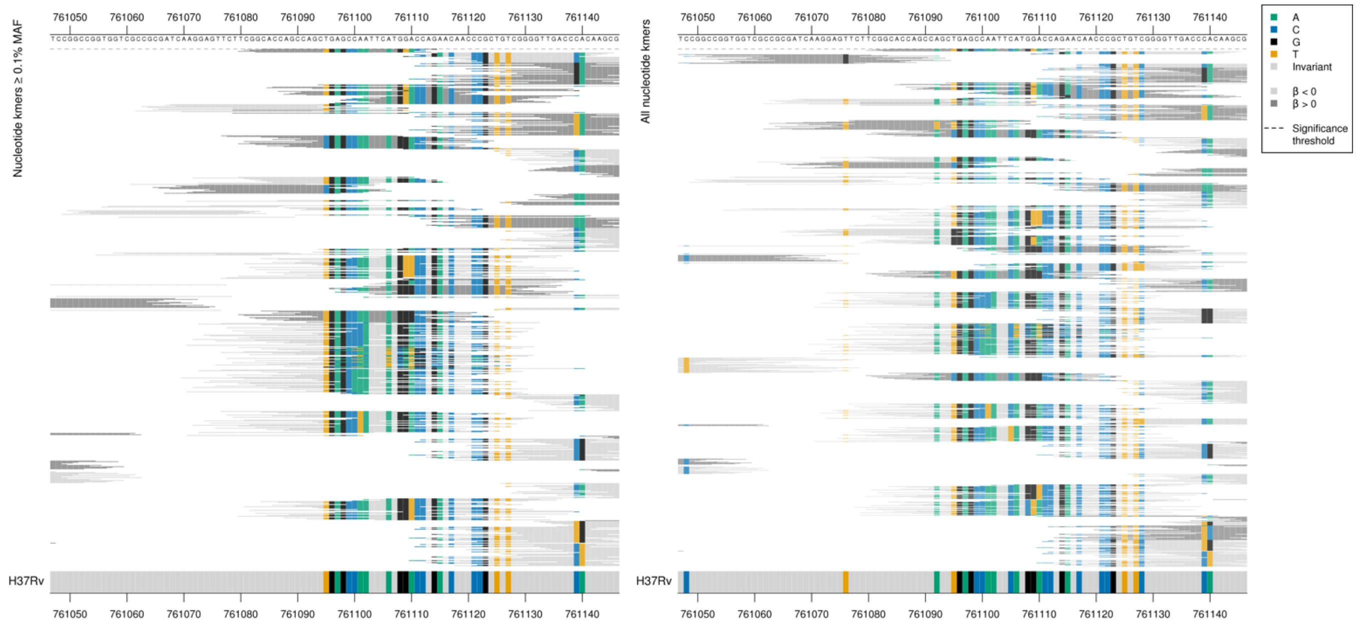

| Kmer | $-\log_{10} p$ | β | MAC | MAF | Ps | Variants |
|---|---|---|---|---|---|---|
| CACCAGCCAGCTGAGCCAATTCATGTACCAG | 6.67 | 0.48 | 61 | 0.84 | 761084 | WT covering positions 761109-761139/1303-1333 (codons 435-445) |

#### *pks1*

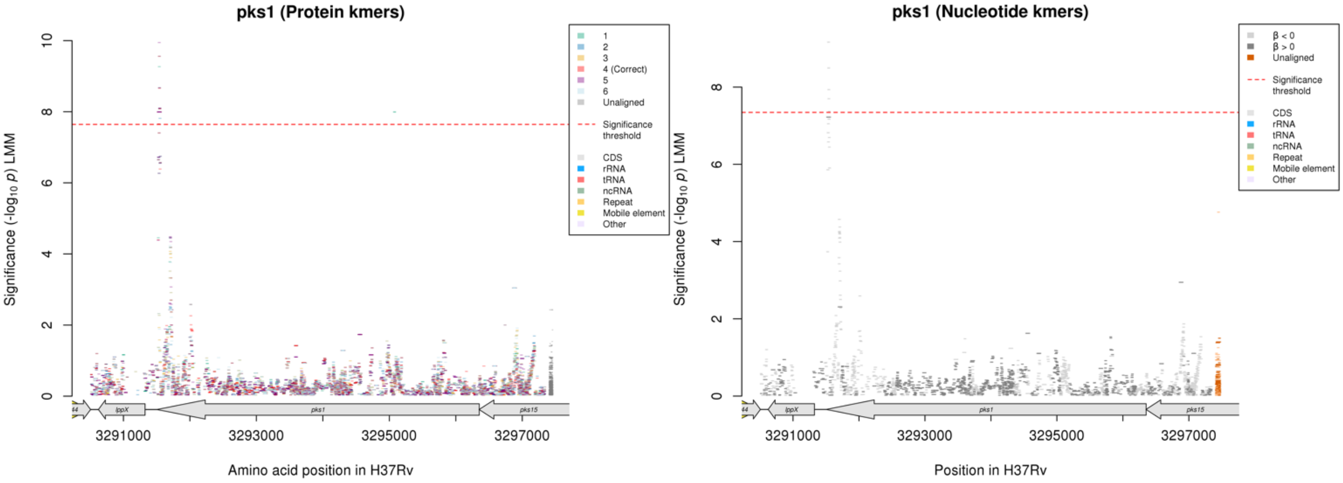

Protein kmers:

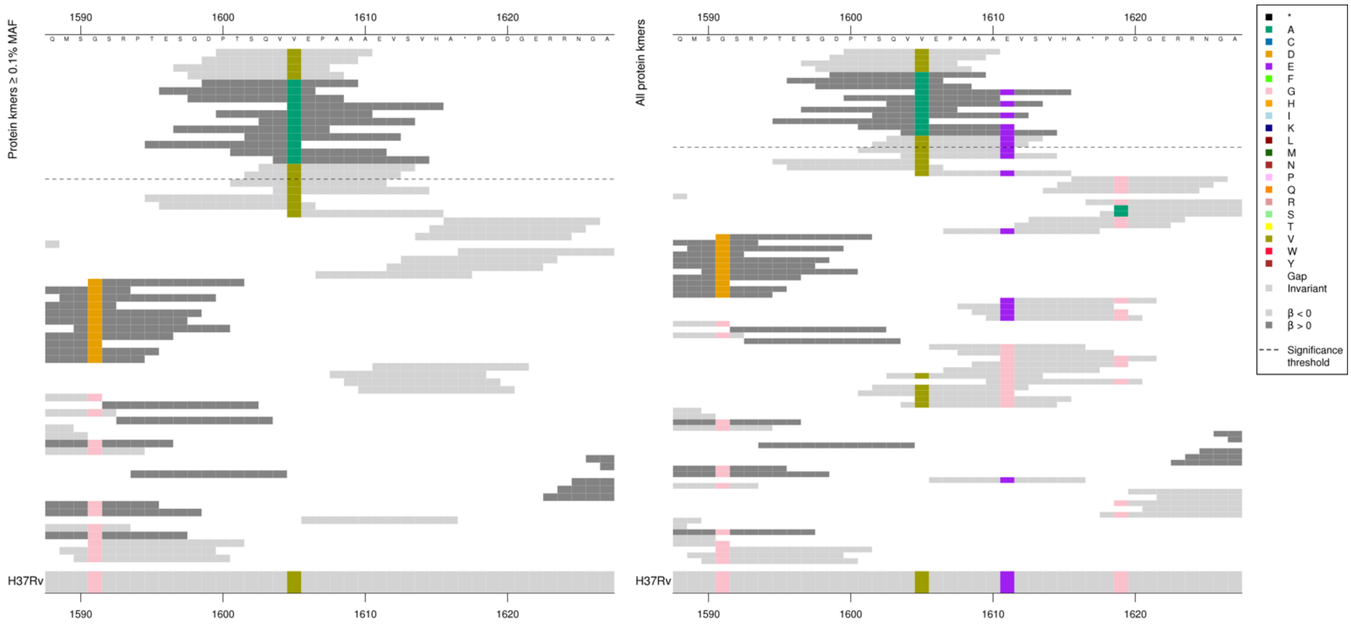

| Kmer | -log10 *p* | β | MAC | MAF | Ps | Variants |
|---|---|---|---|---|---|---|
| PTSQVVEPAAA | 9.56 | -0.96 | 112 | 1.53 | 1600 | 1605V |
| DPTSQVAEPAA | 7.99 | 1.14 | 100 | 1.37 | 1599 | 1605A |
| QVVEPAAAEVS | 7.65 | -0.77 | 117 | 1.60 | 1603 | 1605V, 1611E |

#### *mmaA2:mmaA1*

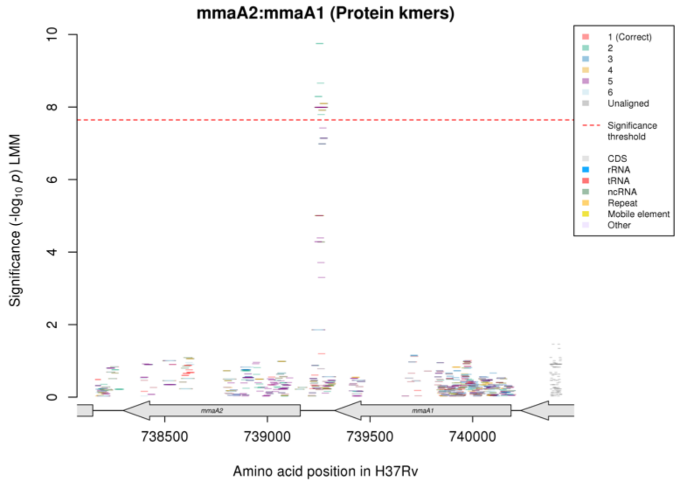

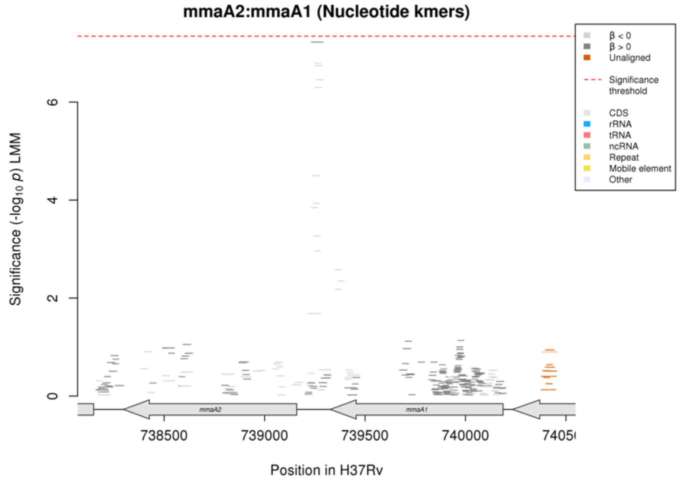

No significant kmers in the nucleotide kmer analysis, but the peak is in the same region and the alignments are below:

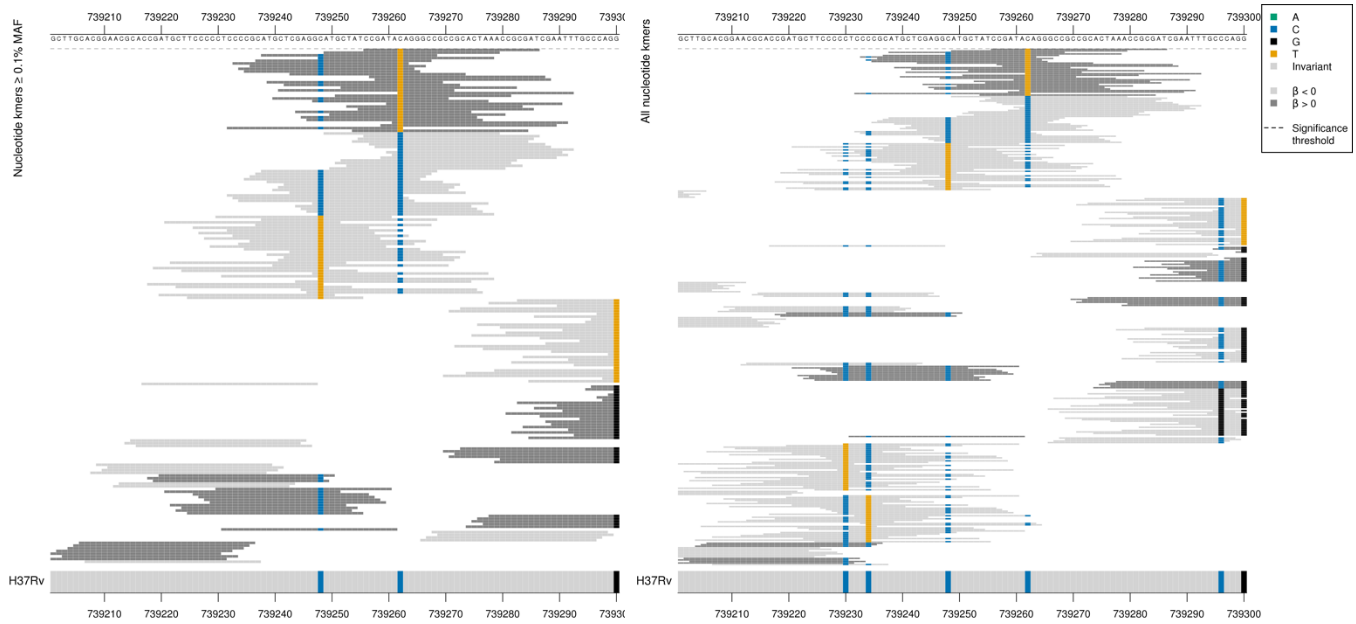

Capturing a mutation at position 739262, 102bp upstream of *mmaA2*, and 65bp downstream of *mmaA1*.

| Kmer | $-\log_{10} p$ | β | MAC | MAF | Ps | Variants |
|---|---|---|---|---|---|---|
| CCGATATAGGGCCGCCGCACTAAACCGCGAT | 7.22 | 1.14 | 101 | 1.36 | 739256 | 739262T (forward strand) |

## *Rv3273*

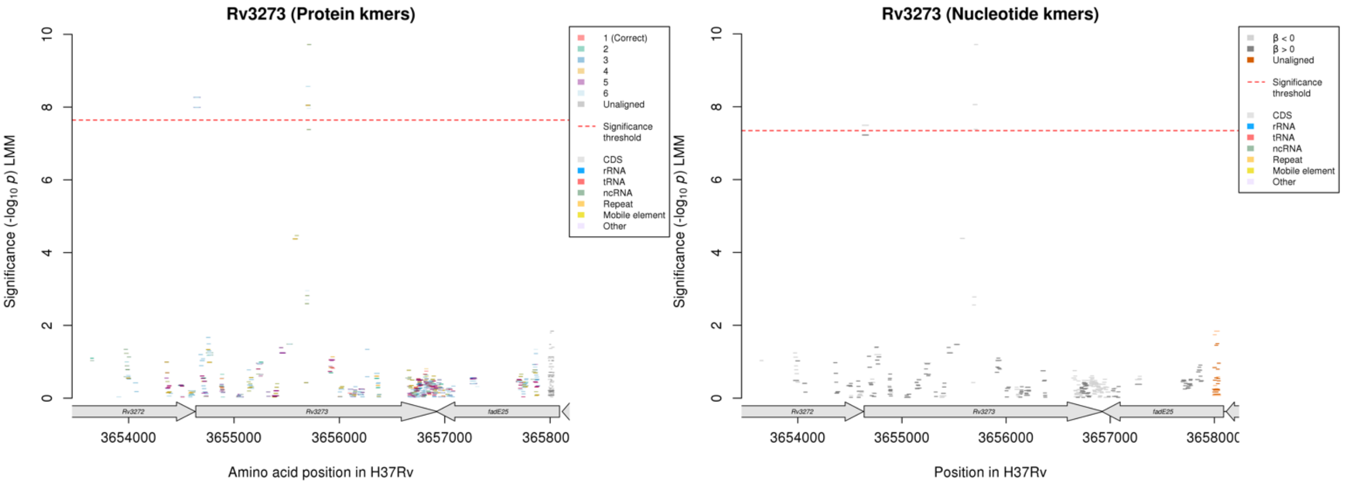

Only low frequency kmers aligned to the correct reading frame for the protein kmer analysis.
Rv3273 starts at position 3654637
Rv3272 ends at position 3654632

Nucleotide kmers:

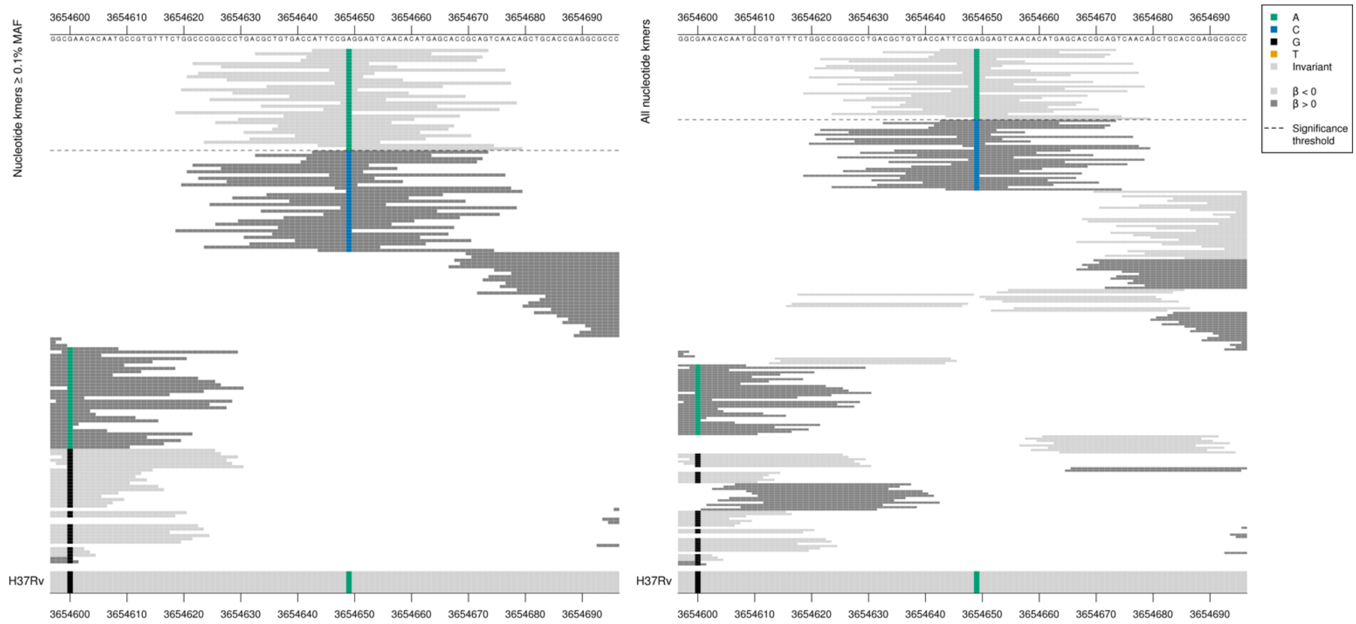

Capturing synonymous mutation at position 3654649, position 13 in Rv3273.
AGG = Arg/R
CGG = Arg/R

| Kmer | -log$_{10}$ *p* | β | MAC | MAF | Ps | Variants |
|---|---|---|---|---|---|---|
| ATTCCGAGGAGTCAACACATGAGCACCGCAG | 7.49 | -1.12 | 101 | 1.38 | 3654643 | 3654649/13A |

Nucleotide kmers:

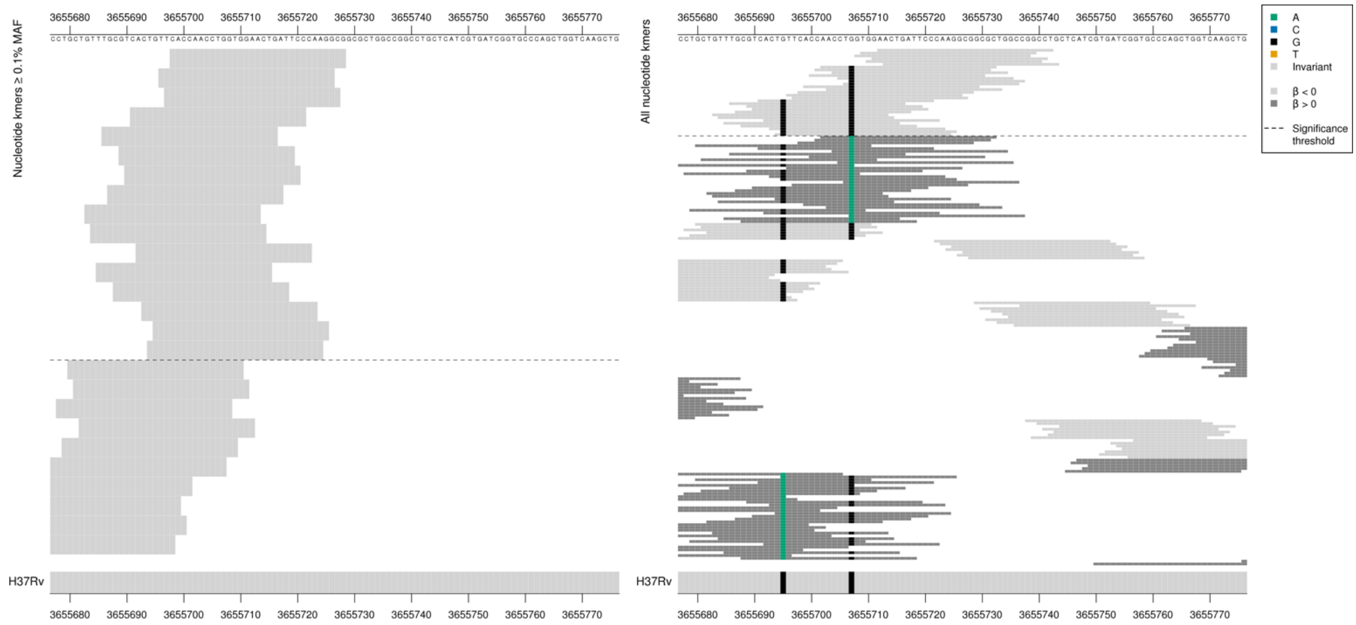

WT kmers and synonymous mutations at positions 3655695 (1059) and 3655707 (1071)
3655695 (1059)
CTG = Leu / L
CTA = Leu / L
3655707 (1071)
CTG = Leu / L
CTA = Leu / L

Low MAF kmers = WT positions 3655708-3655743, WT / 3655707/1071 G

| Kmer | $-\log_{10} p$ | β | MAC | MAF | Ps | Variants |
|---|---|---|---|---|---|---|
| CACTGTTCACCAACCTGGTGGAACTGATTCC | 8.06 | -1.19 | 13 | 0.18 | 3655691 | WT / 3655707/1071 G, 3655695/1059G |

#### *mce3R:yrbE3A*

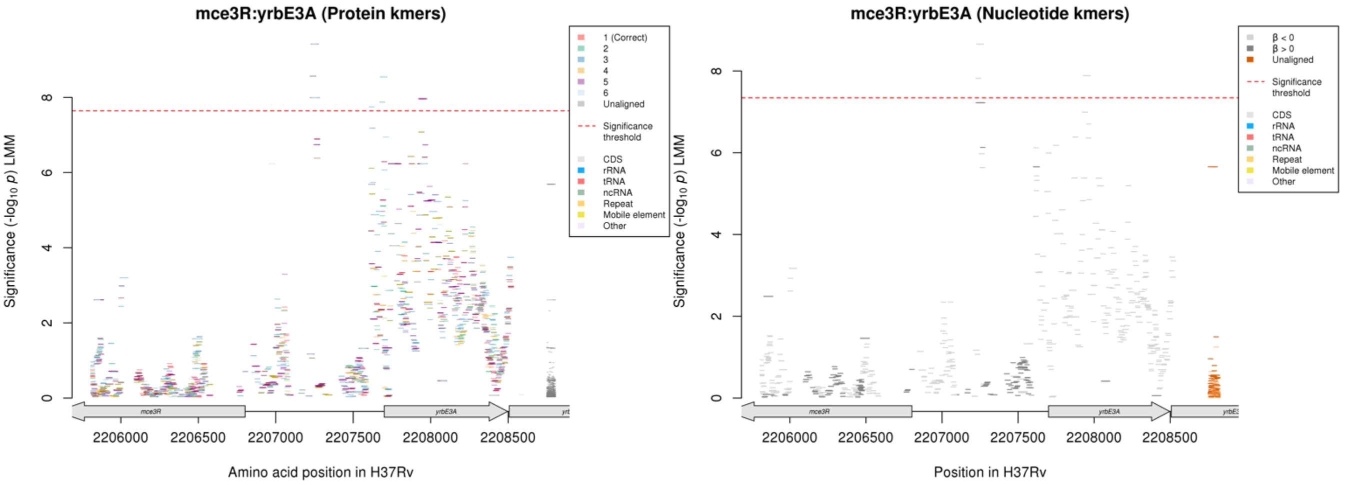

Some kmers aligned to the wrong reading frame at the start of the gene, but no significant kmers in this region in the nucleotide kmer analysis.

Nucleotide kmers:

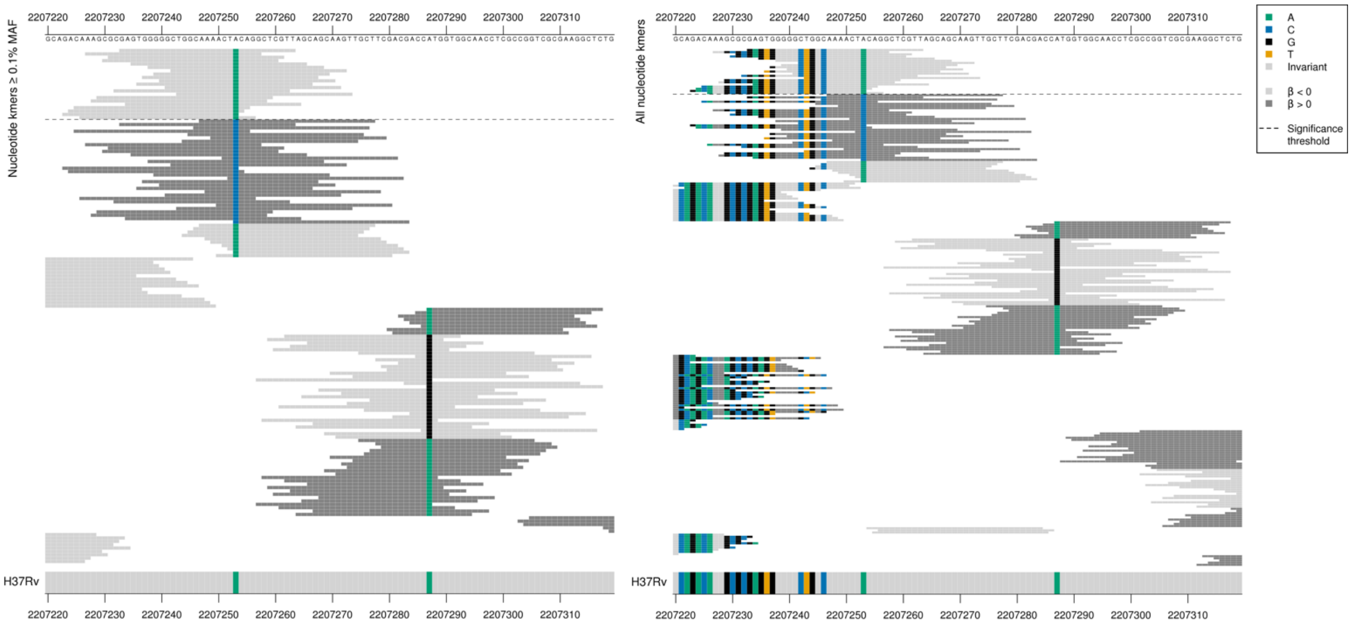

Whole region is inflated in significance. *mce3R* begins at position 2206802 on the reverse strand. *yrbE3A* begins at position 2207700 on the forward strand.

| Kmer | $-\log_{10} p$ | β | MAC | MAF | Ps | Variants |
|---|---|---|---|---|---|---|
| GAGTGGGGGCTGGCAAAACTACAGGCTCGTT | 8.66 | -1.14 | 103 | 1.41 | 2207233 | 2207253/-447A yrbE3A or 2207253/-451T mce3R |

# *Rv3796*

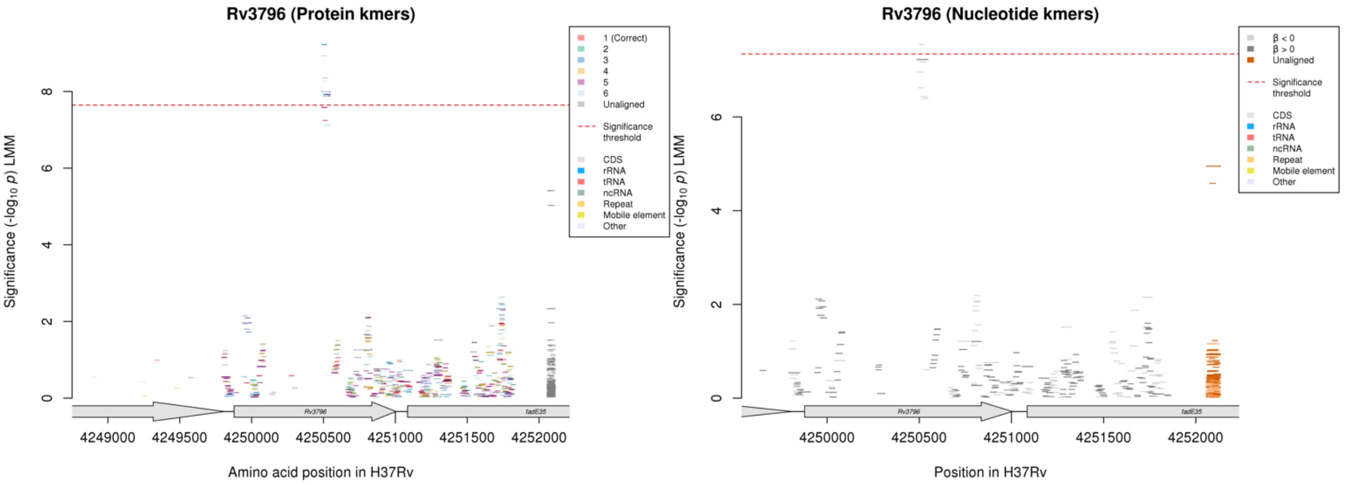

#### Protein kmers:

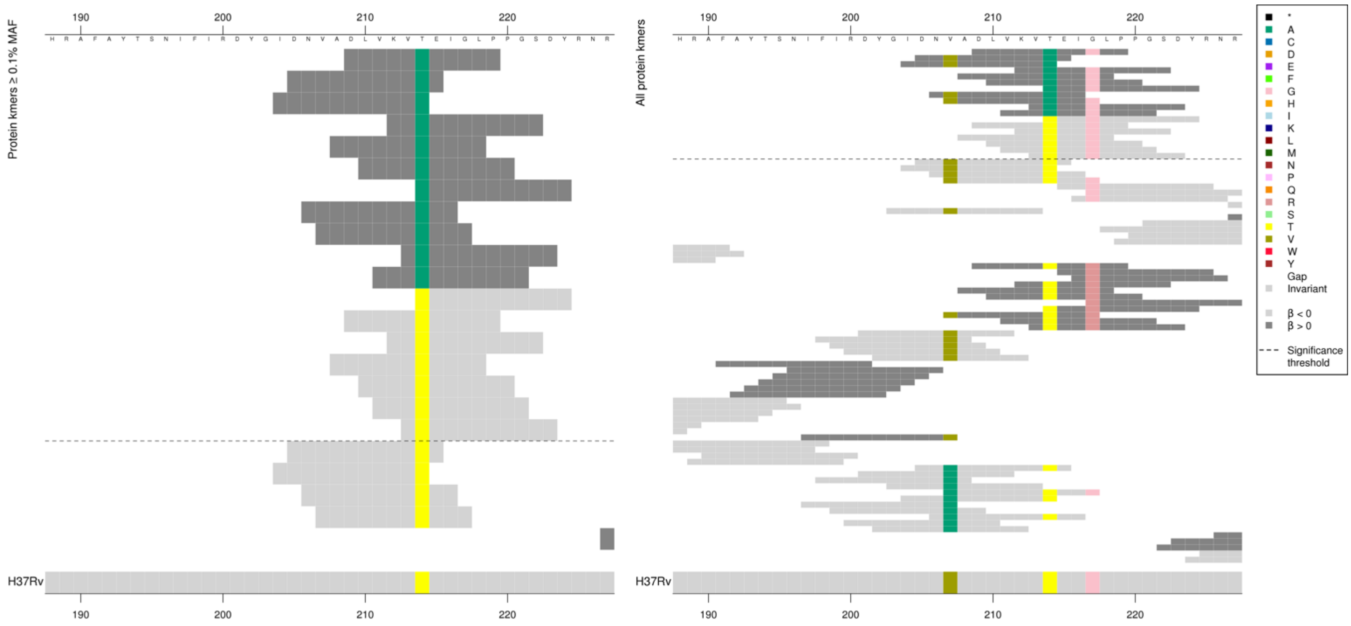

| Kmer | -log₁₀ p | β | MAC | MAF | Ps | Variants |
|---|---|---|---|---|---|---|
| DLVKVAEIGLP | 7.99 | 1.14 | 100 | 1.37 | 209 | 214A |
| TEIGLPPGSDY | 7.93 | -0.96 | 106 | 1.45 | 214 | 214T/217G |

#### *mez*

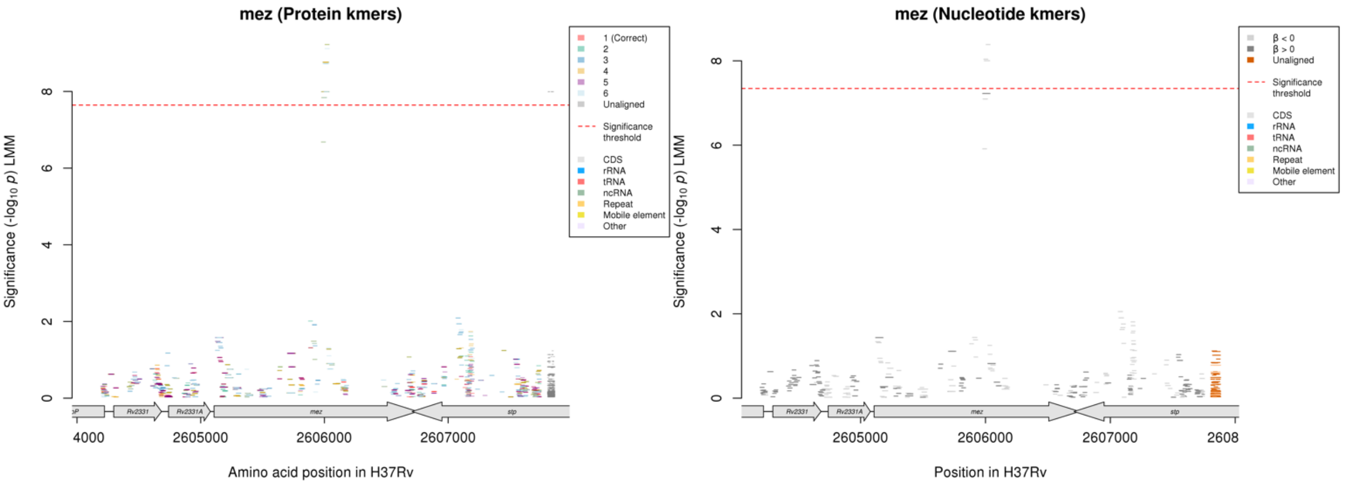

No significant kmers aligned to the correct reading frame for the protein kmers.
Nucleotide kmers:

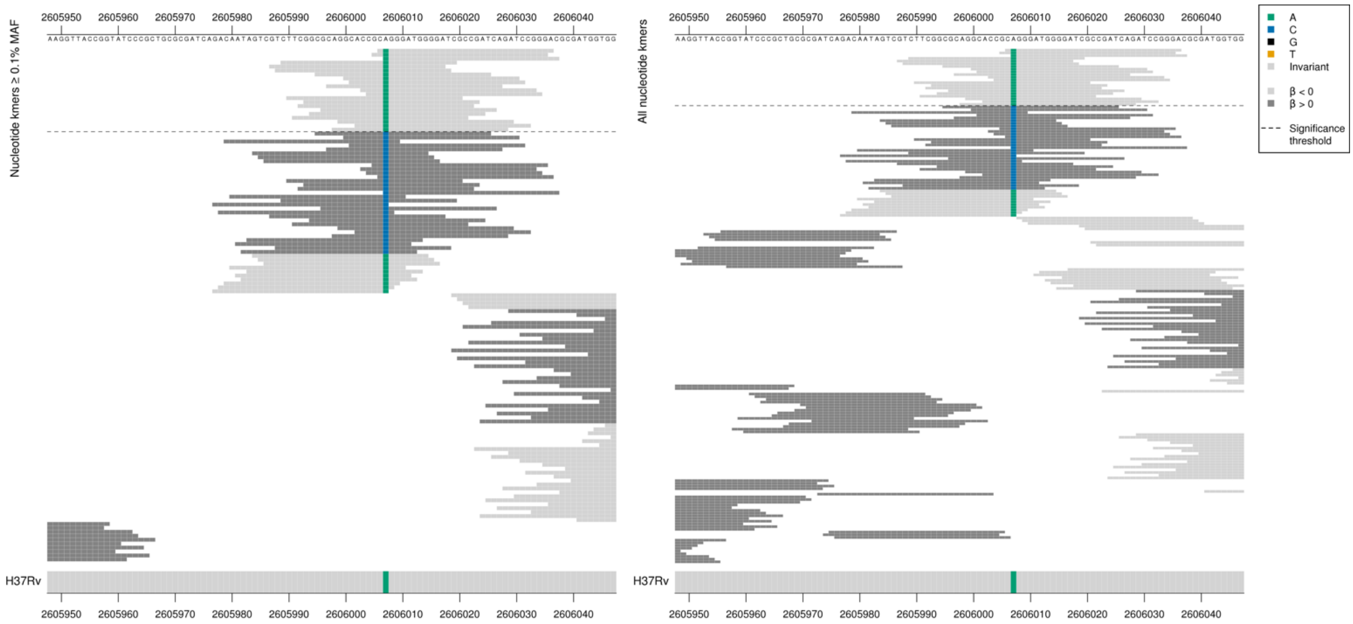

*mez* begins at position 2605108
Synonymous mutation at position 2606007 (900)
GCA = Ala / A
GCC = Ala / A

| Kmer | $-log_{10}\,p$ | β | MAC | MAF | Ps | Variants |
|---|---|---|---|---|---|---|
| CAGGGATGGGGATCGCCGATCAGATCCGGGA | 8.39 | -1.13 | 103 | 1.41 | 2606006 | 2606007/900A |

# *Rv2390c*

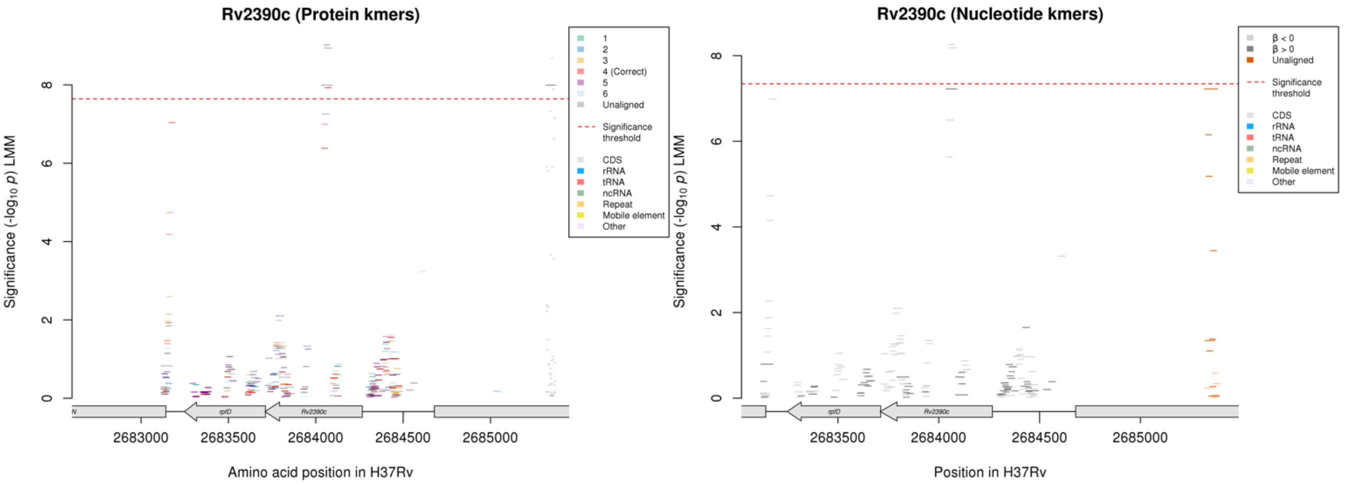

Protein kmers:

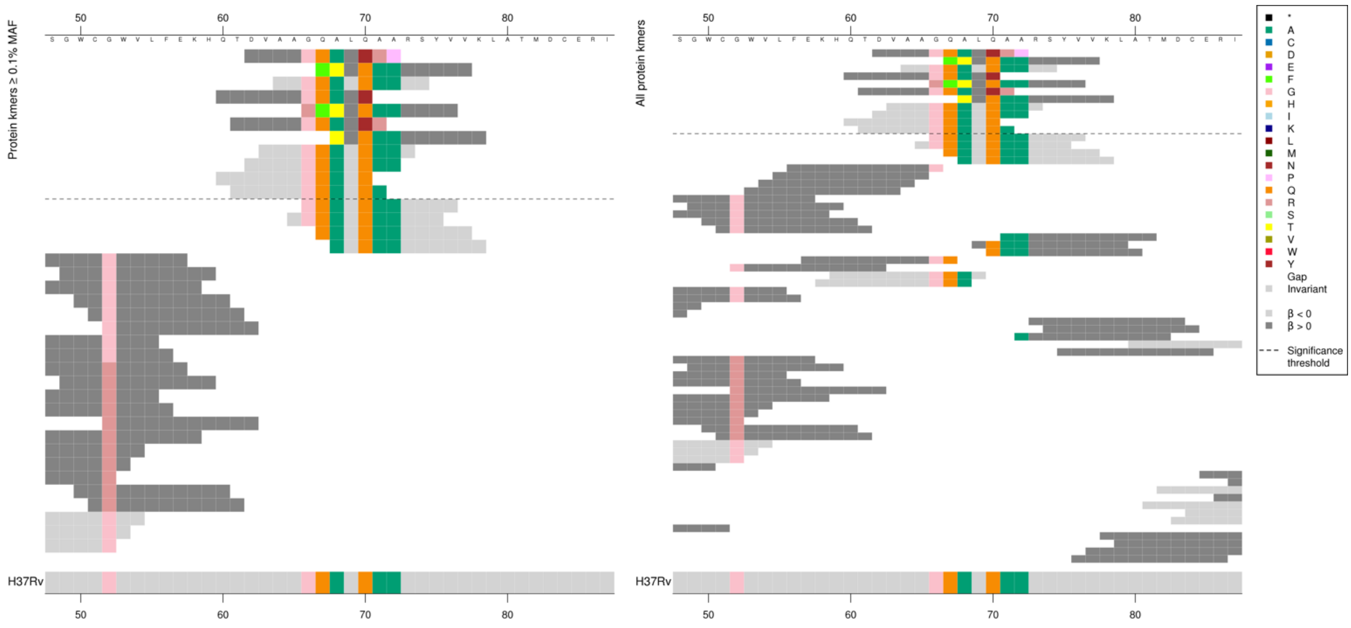

The same region for the nucleotide kmer analysis:

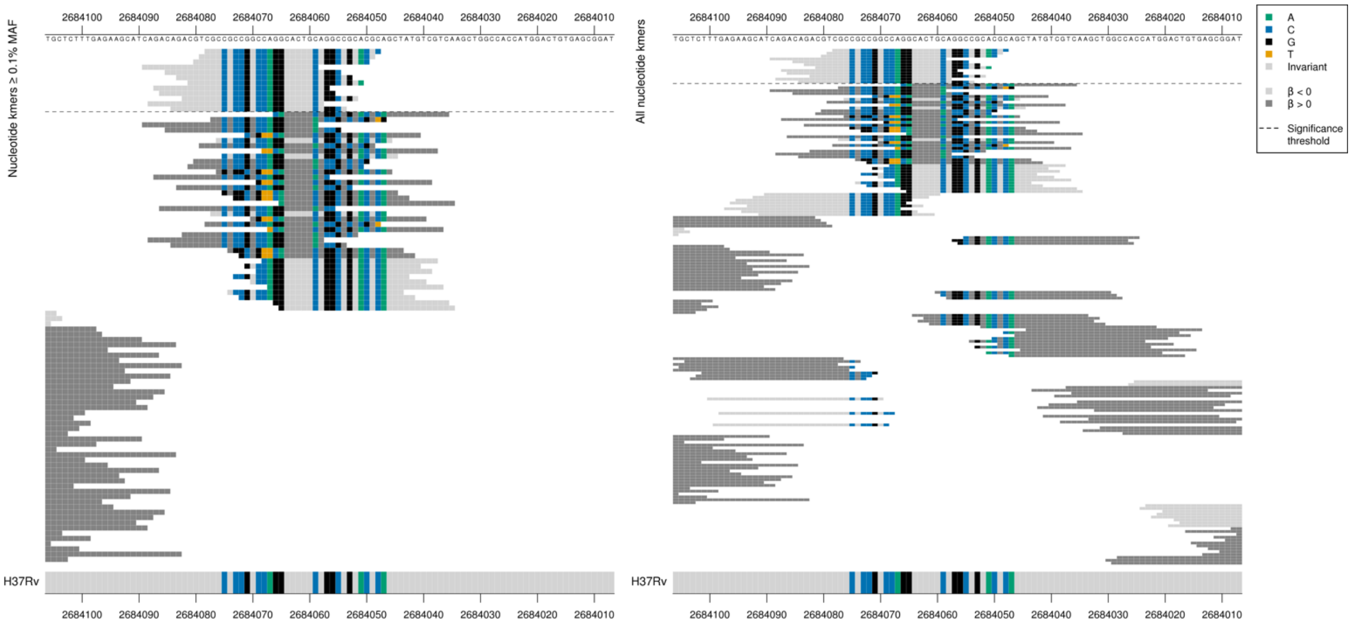

Gene begins at position 2684266.

Low and high-MIC associated kmers were significant in the protein kmer analysis, only low-MIC kmers were significantly associated in the nucleotide kmer analysis.

Run of mutations/possible indel between positions 2684075-2684065 (192-202) and between positions 2684059-2684047 (208-220) in the nucleotide kmers that were not significant, compared to low MIC-associated kmers that were the same as the reference.

| Kmer | -log$_{10}$ *p* | β | MAC | MAF | Ps | Variants |
|---|---|---|---|---|---|---|
| DVAAGQALNRP | 7.99 | 1.14 | 100 | 1.37 | 62 | Indels |
| AAGQALQAARS | 7.99 | -1.14 | 100 | 1.37 | 64 | WT |

#### *yrbE3B*

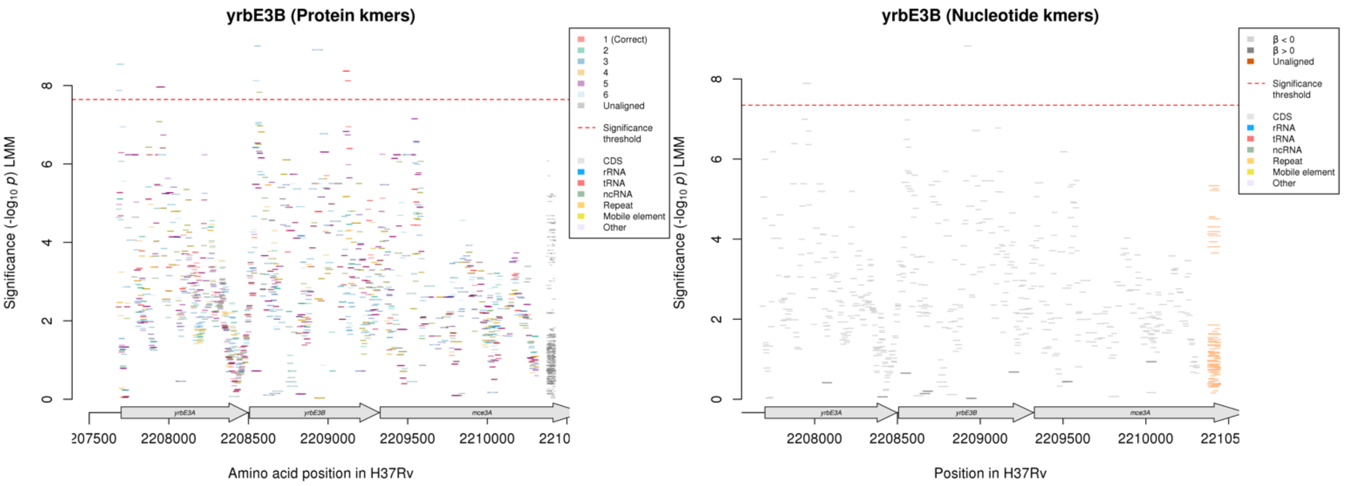

Protein kmers:

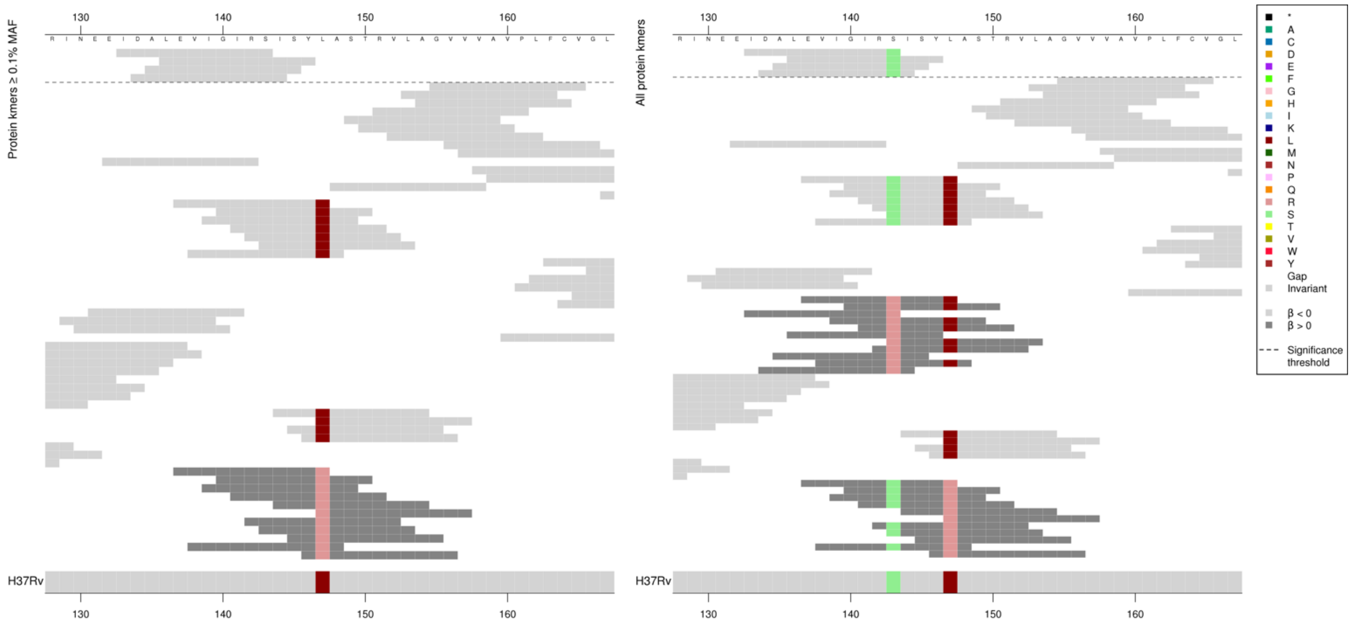

Kmers below the significance threshold = 143R

| Kmer | -log$_{10}$ *p* | β | MAC | MAF | Ps | Variants |
|---|---|---|---|---|---|---|
| IDALEVIGIRS | 8.91 | -1.44 | 13 | 0.18 | 133 | WT / 143S |

#### Protein kmers:

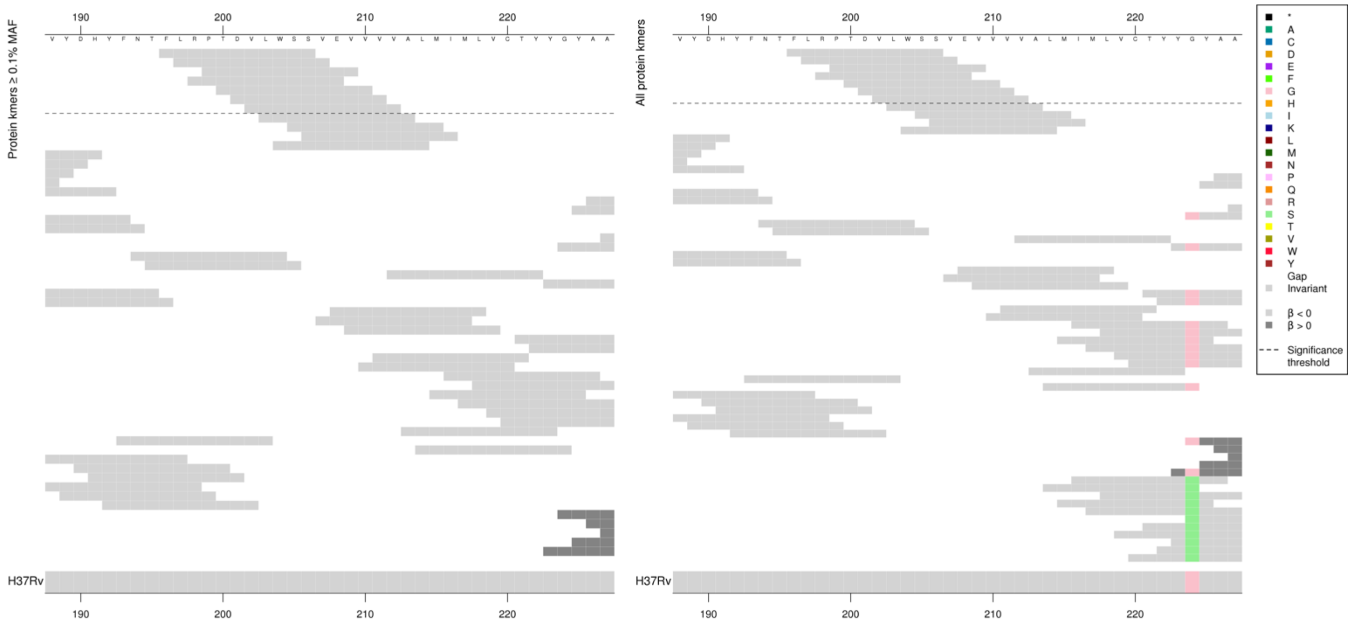

| Kmer | $-\log_{10} p$ | β | MAC | MAF | Ps | Variants |
|---|---|---|---|---|---|---|
| FLRPTDVLWSS | 8.37 | -1.39 | 13 | 0.18 | 196 | WT |

## *Rv0207c*

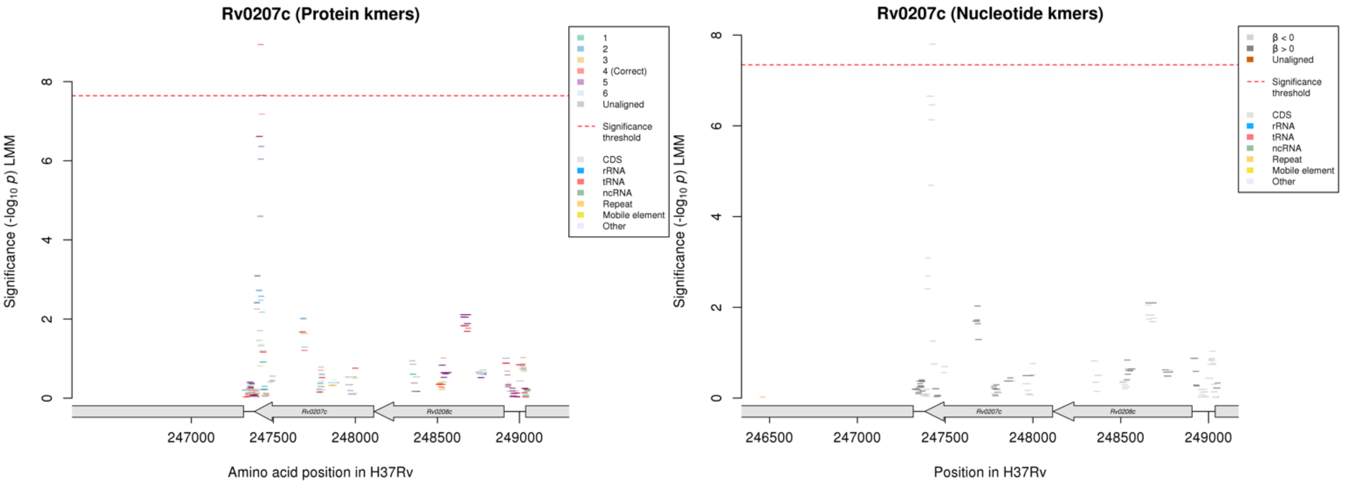

Protein kmers:

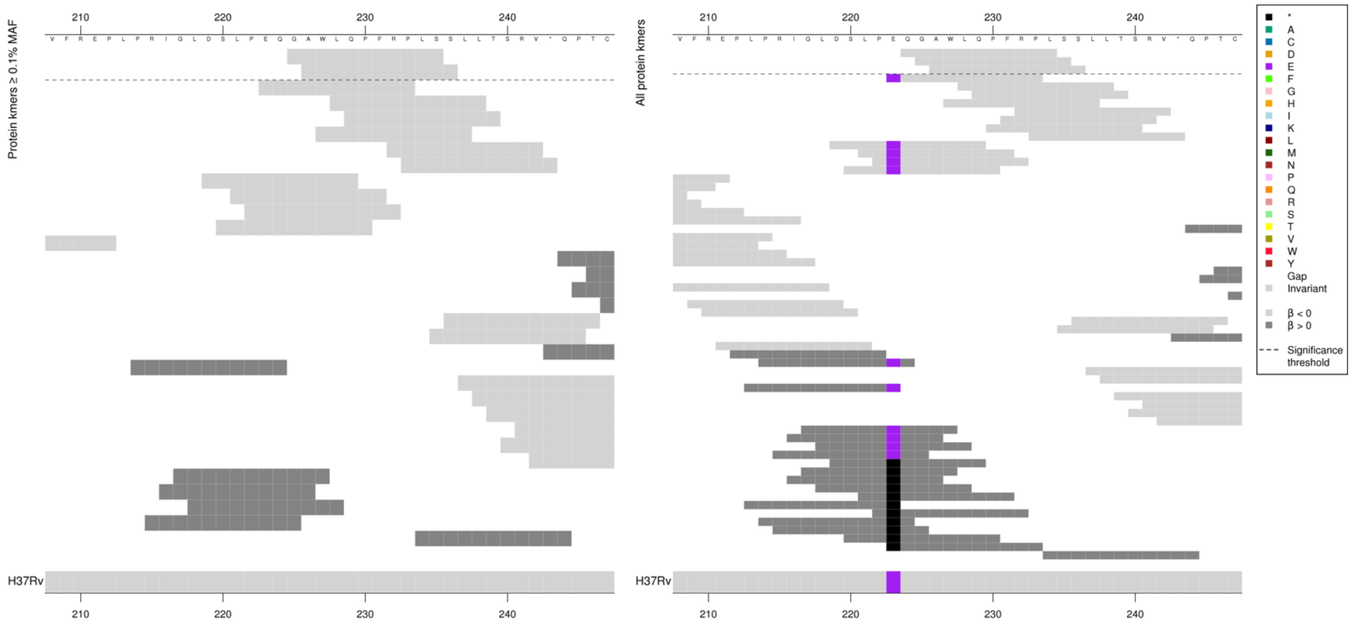

No variants identified, likewise for the nucleotide kmer analysis.

| Kmer | $-\log_{10} p$ | β | MAC | MAF | Ps | Variants |
|---|---|---|---|---|---|---|
| GAWLQPFRPLS | 8.94 | -1.77 | 8 | 0.11 | 225 | WT |

#### *argS*

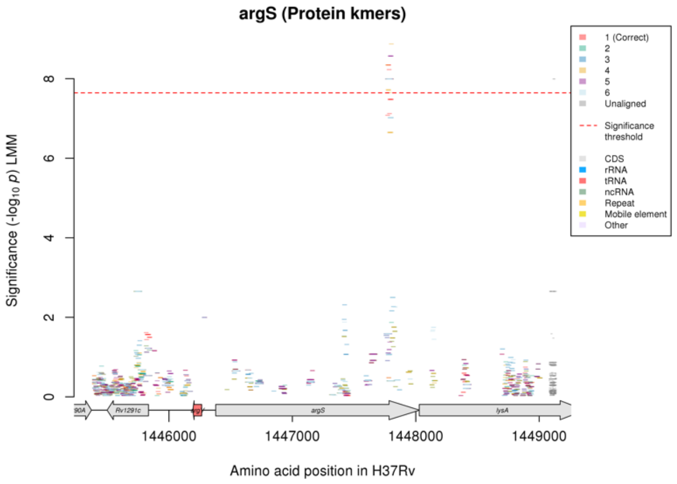

Not significant in nucleotide kmer analysis.

### Clofazimine (CFZ)

#### Protein kmers:

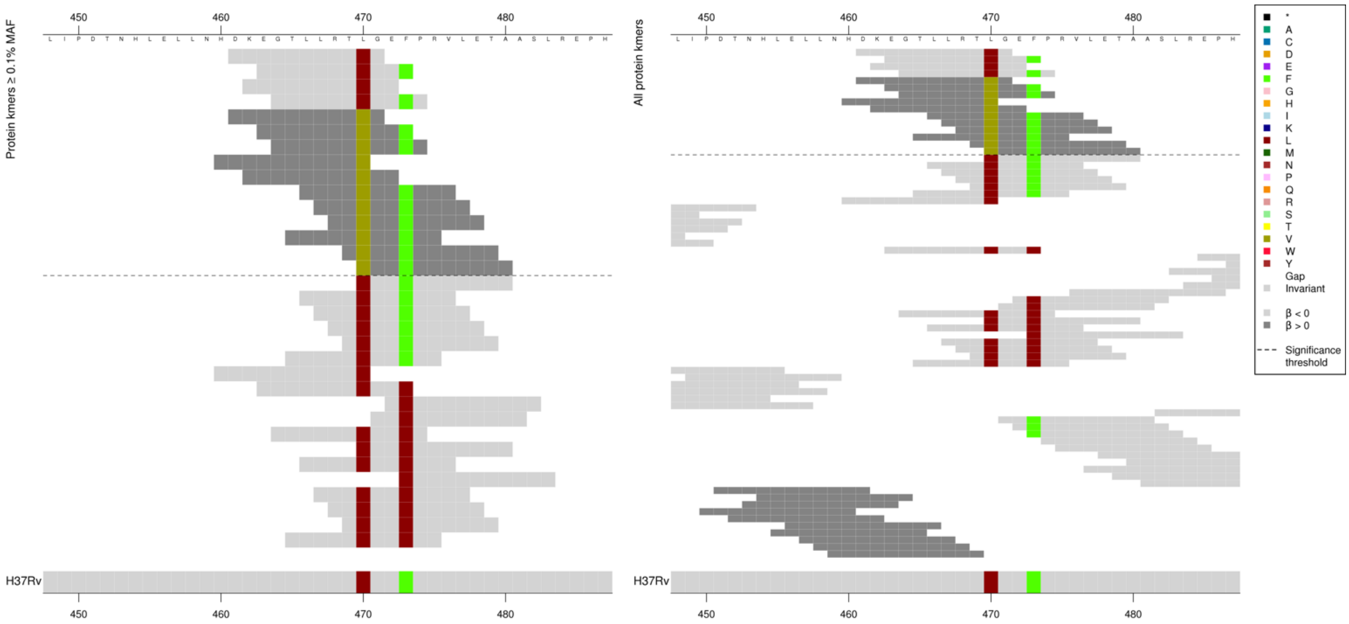

| Kmer | $-\log_{10} p$ | β | MAC | MAF | Ps | Variants |
|---|---|---|---|---|---|---|
| DKEGTLLRTLG | 8.34 | -1.13 | 101 | 1.38 | 461 | 470L |
| DKEGTLLRTVG | 7.99 | 1.14 | 100 | 1.37 | 461 | 470V |

### Delamanid (DLM)

#### *ddn*

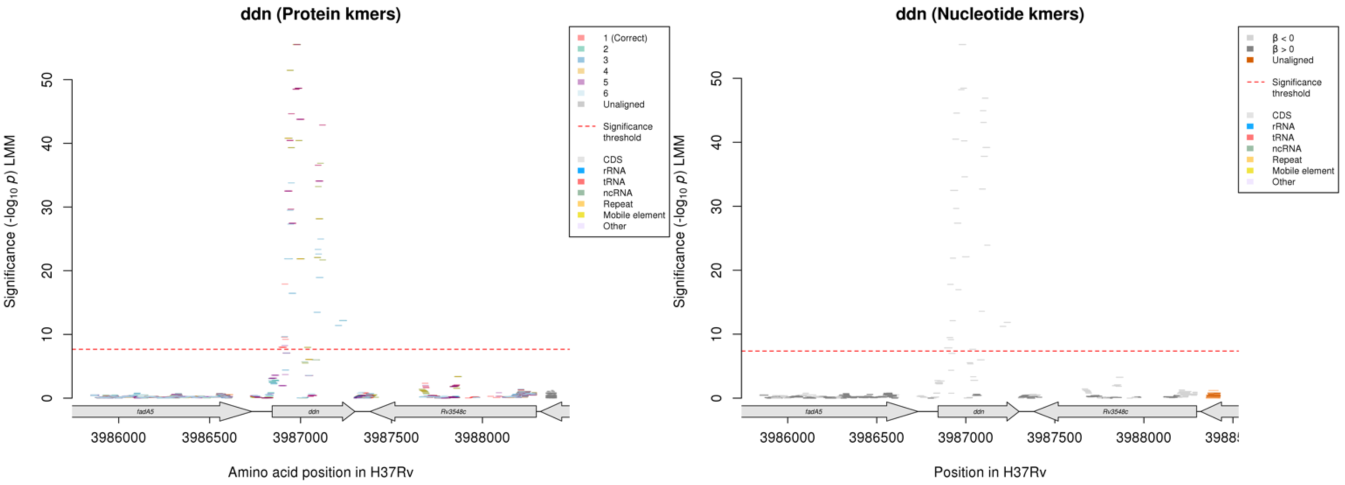

Protein kmers:

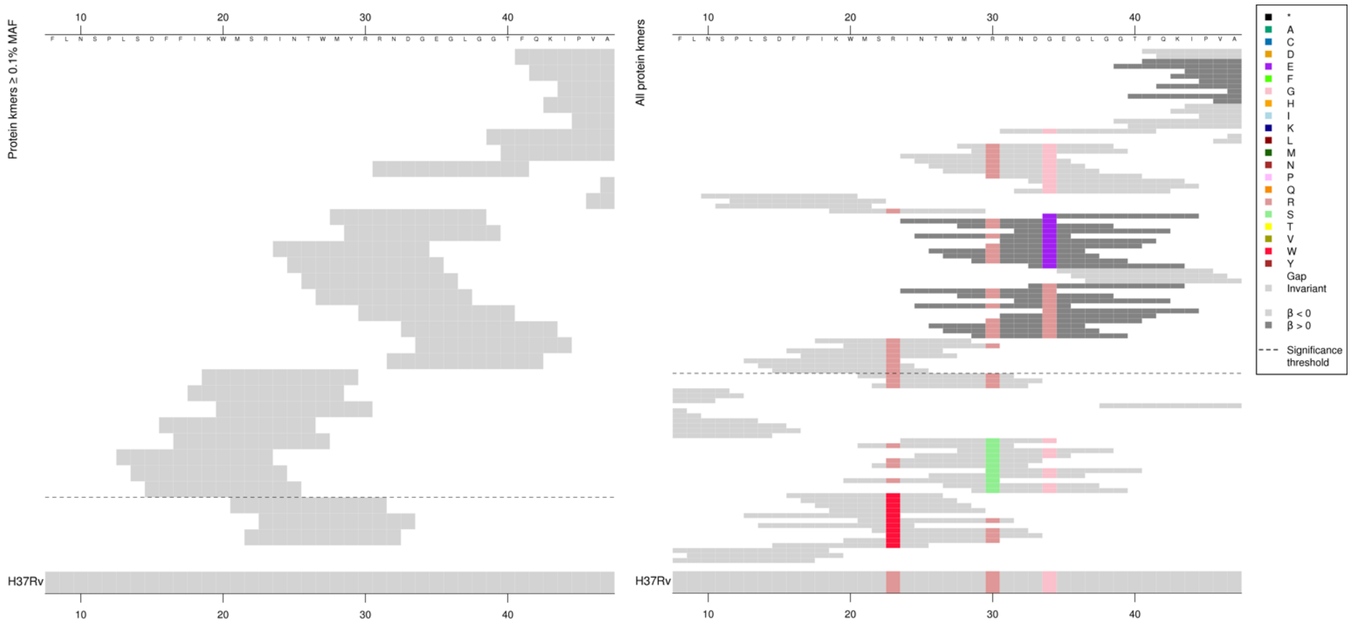

WT significant kmers were possibly linked to the substitutions, or capturing regions with rare variation not analysed here. The reference-sequence kmers capturing a region with no variation were also capturing no variation when looking at the nucleotide kmers, so they were not capturing synonymous substitutions. They could be capturing rare variants below the threshold of being seen in five genomes across the full dataset.

Low MAF kmers = WT (ps 10, 35), 34E, 34R

| Kmer | $-\log_{10} p$ | β | MAC | MAF | Ps | Variants |
|---|---|---|---|---|---|---|
| RNDGEGLGGTF | 44.65 | -3.33 | 14 | 0.19 | 31 | 34G |
| MYRRNDGEGLG | 40.45 | -2.70 | 19 | 0.25 | 28 | 30R, 34G |
| KWMSRINTWMY | 17.91 | -2.71 | 9 | 0.12 | 19 | 23R |
| WMSRINTWMYR | 9.24 | -1.49 | 14 | 0.19 | 20 | 23R, 30R |

Protein kmers:

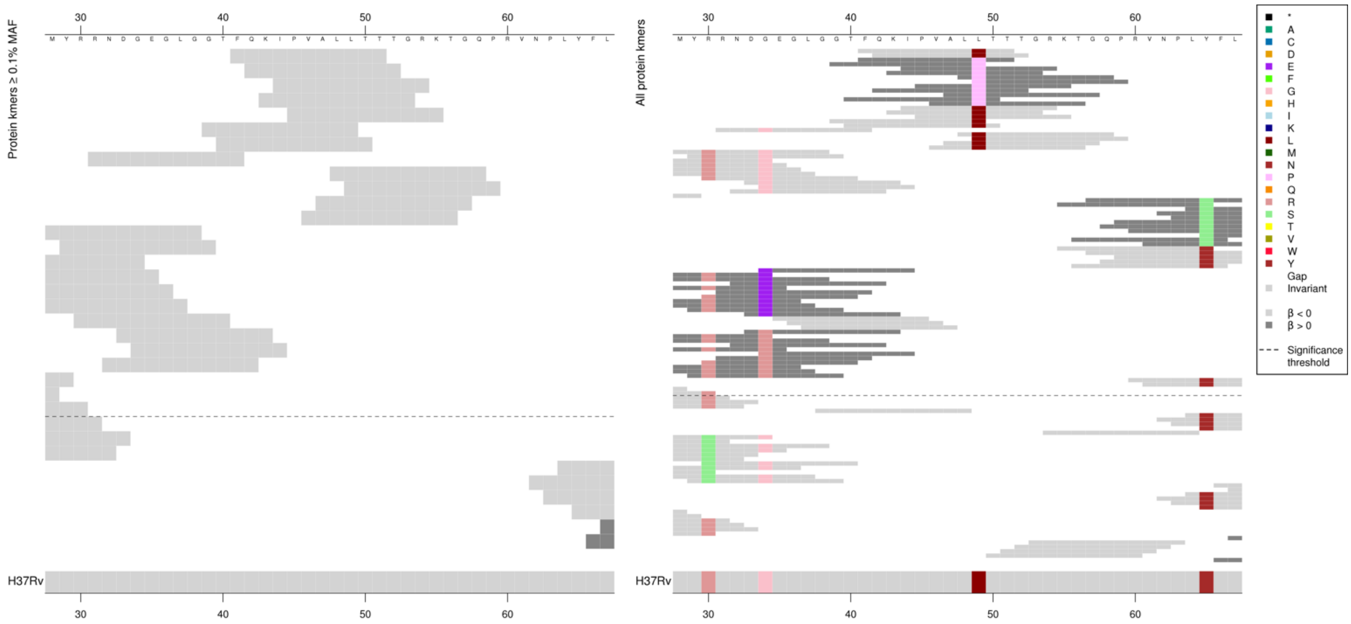

Low MAF kmers = 49P.

| Kmer | $-\log_{10} p$ | β | MAC | MAF | Ps | Variants |
|---|---|---|---|---|---|---|
| FQKIPVALLTT | 55.50 | -4.75 | 9 | 0.12 | 41 | 49L |

Protein kmers:

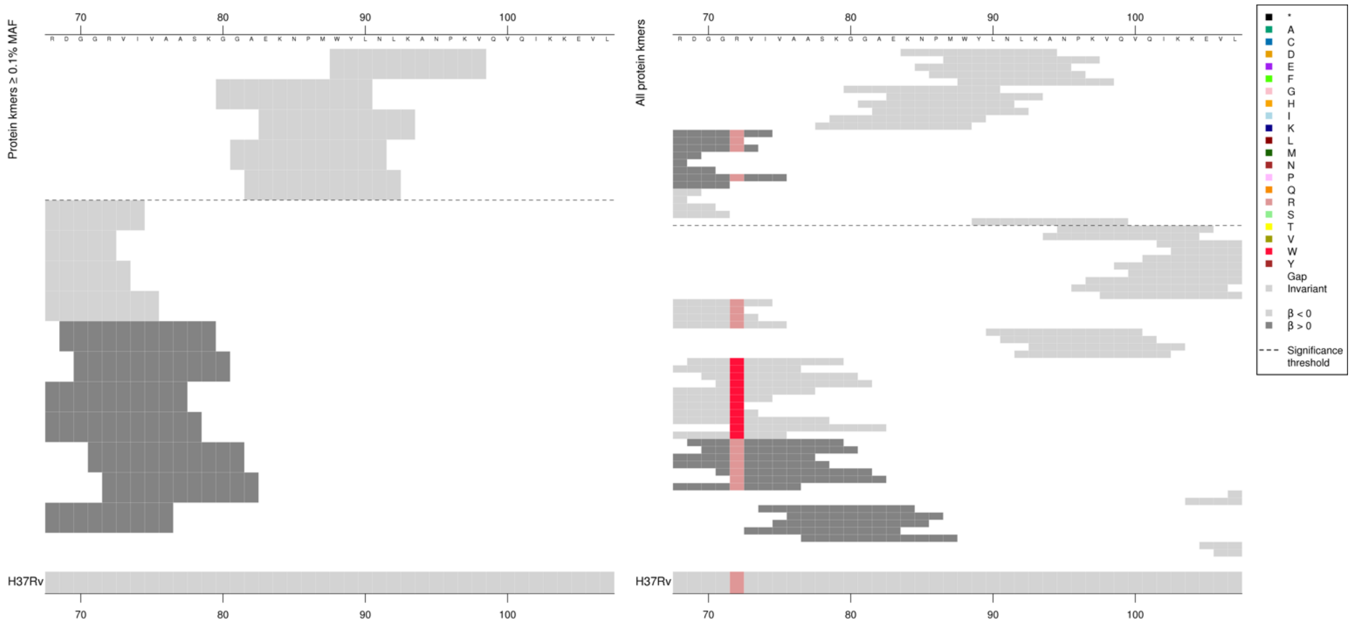

The same region as above covering codons ~80-100 in the nucleotide kmer analysis:

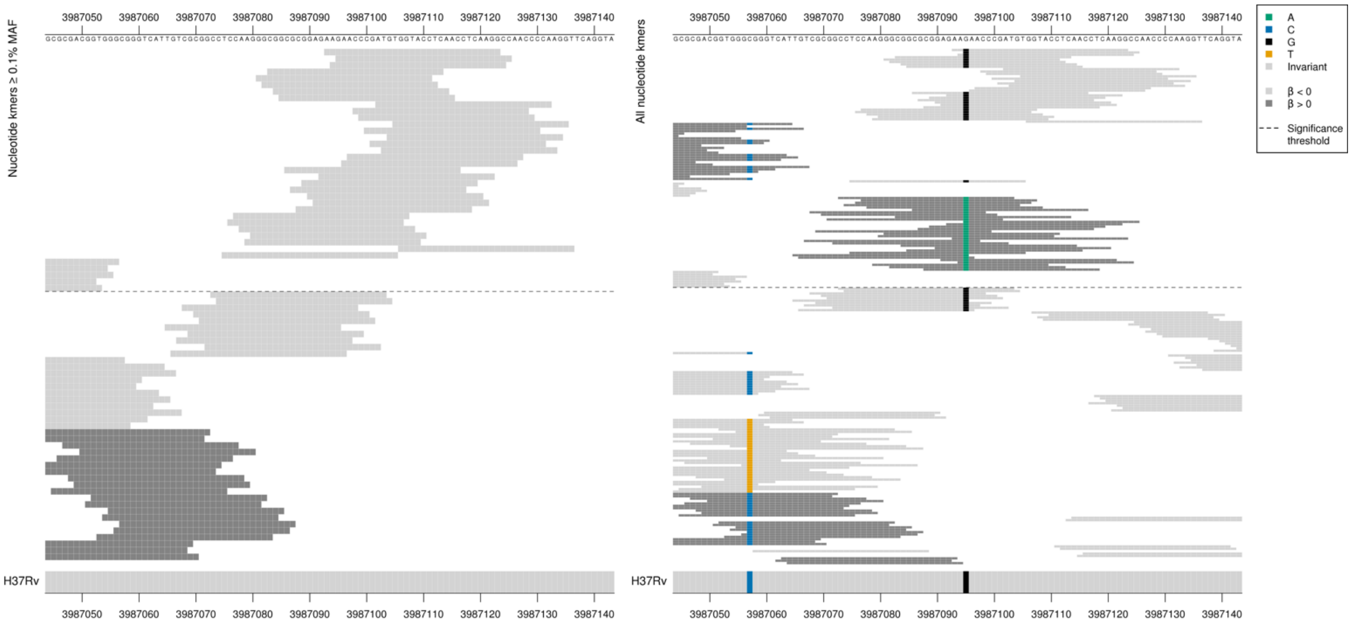

Capturing a synonymous variant at position 3987095/252, codon 84
Low MIC-associated kmers = AAG = Lys / K
High MIC-associated kmers = AAA = Lys / K

There were also low MIC-associated WT kmers not covering this variant which were possibly in LD, or capturing a region with rare variation not shown in these figures.

Low MAF kmers = 3987095/252 A (84K aa).

| Kmer | $-\log_{10} p$ | β | MAC | MAF | Ps | Variants |
|---|---|---|---|---|---|---|
| AAGAACCCGATGTGGTACCTCAACCTCAAGG | 46.89 | -3.70 | 12 | 0.16 | 3987093 | 3987095/252 G (84K aa) |
| ATGTGGTACCTCAACCTCAAGGCCAACCCCA | 39.19 | -4.23 | 8 | 0.11 | 3987102 | WT |

#### *fadE22*

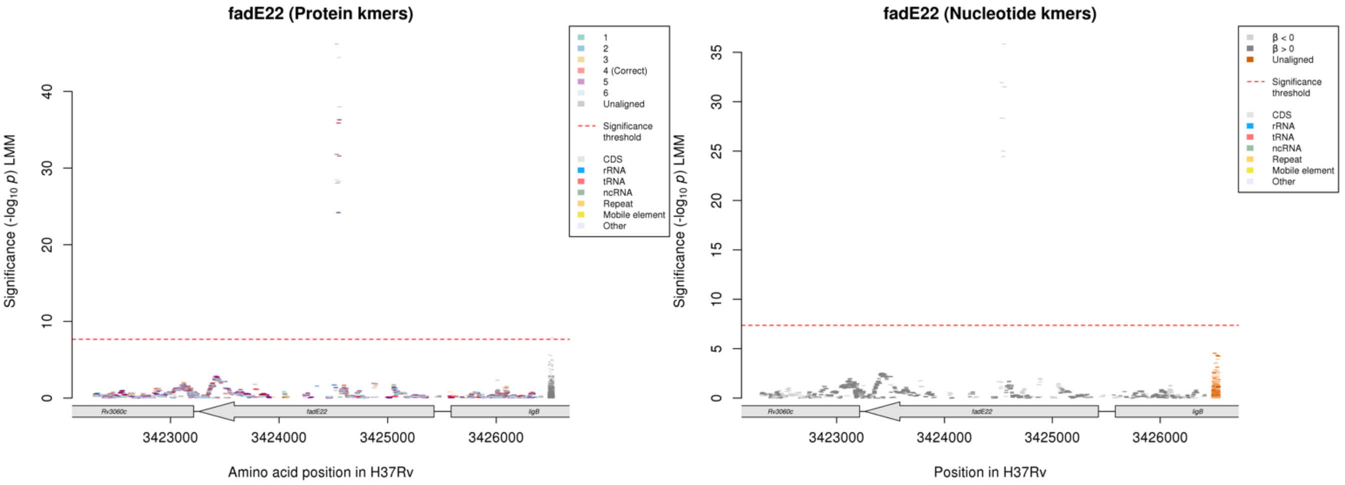

Protein kmers:

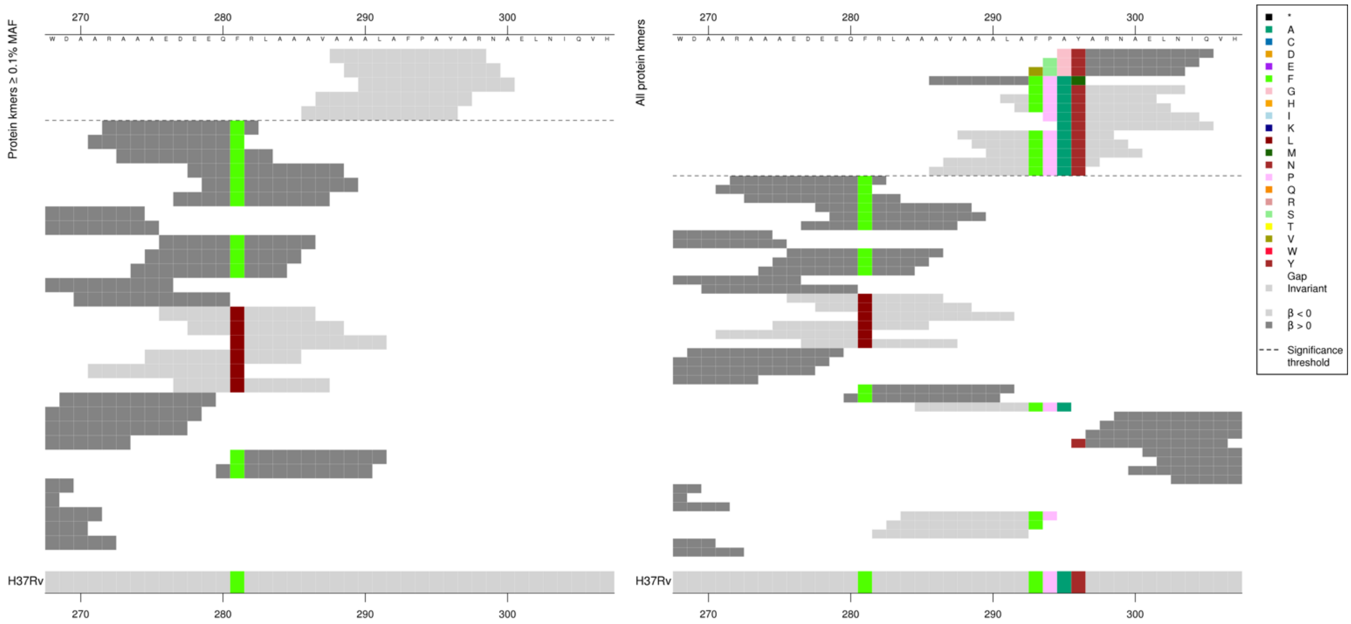

The same region for the nucleotide kmer analysis:

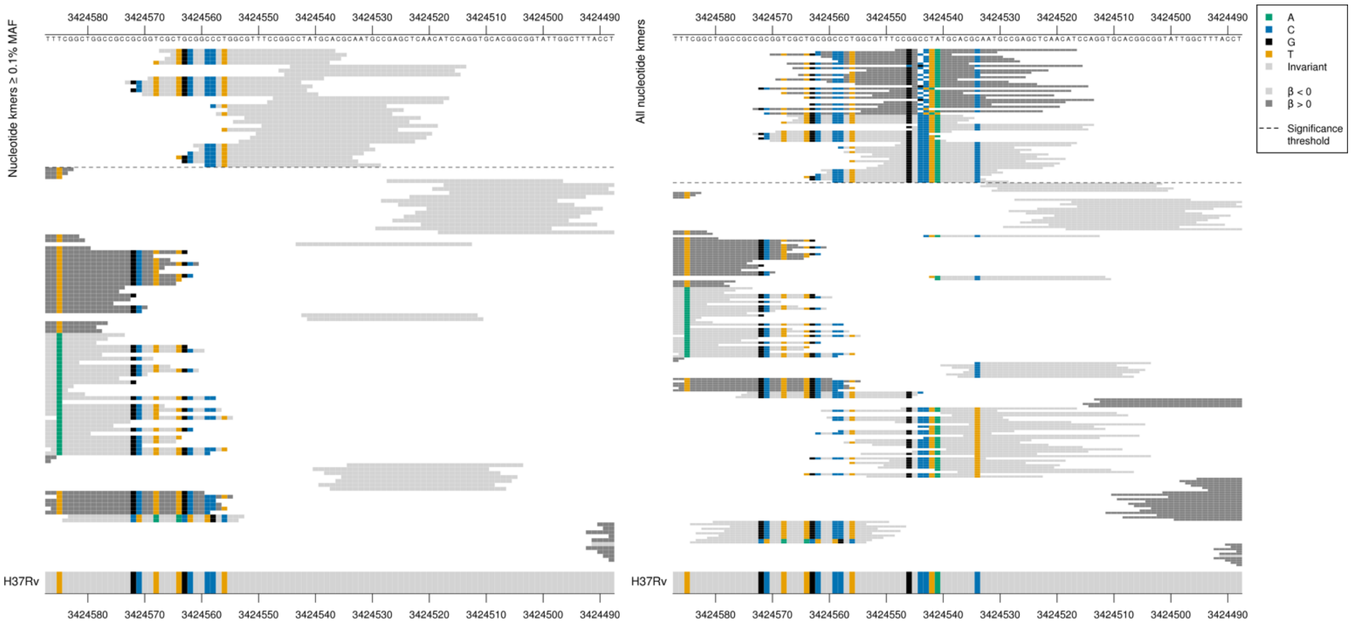

Low MAF kmers = C deletion between 3424544-3424543/884-885

| Kmer | $-\log_{10} p$ | β | MAC | MAF | Ps | Variants |
|---|---|---|---|---|---|---|
| CGCTGCGGCCCTGGCGTTTCCGGCCTATGCA | 35.83 | -3.49 | 10 | 0.13 | 3424567 | WT |

### *fba*

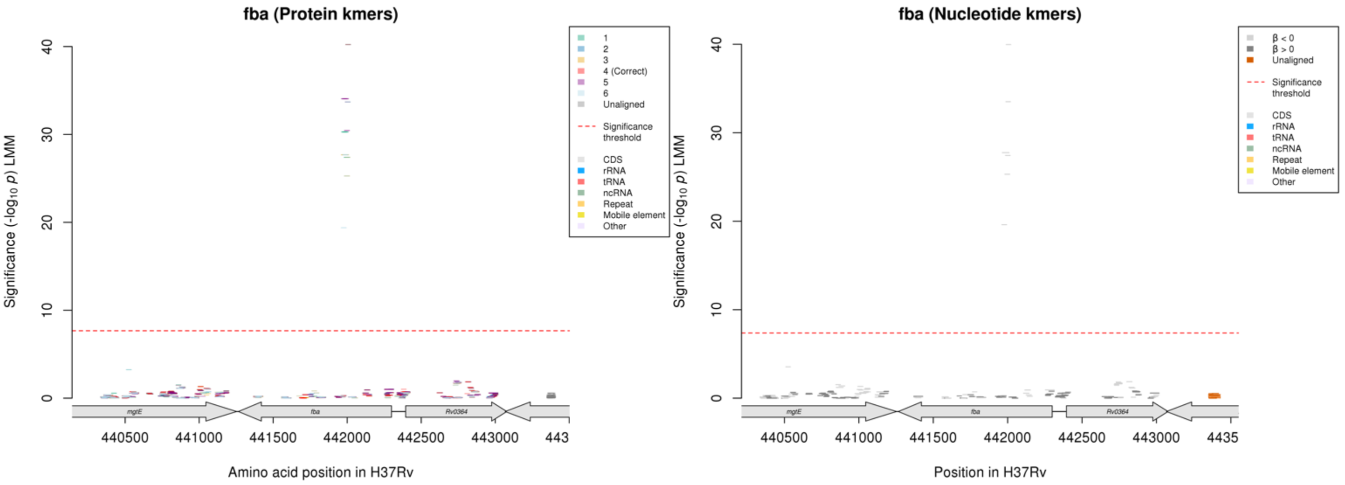

No significant kmers aligned to the correct reading frame in the protein kmer analysis.

Nucleotide kmers:

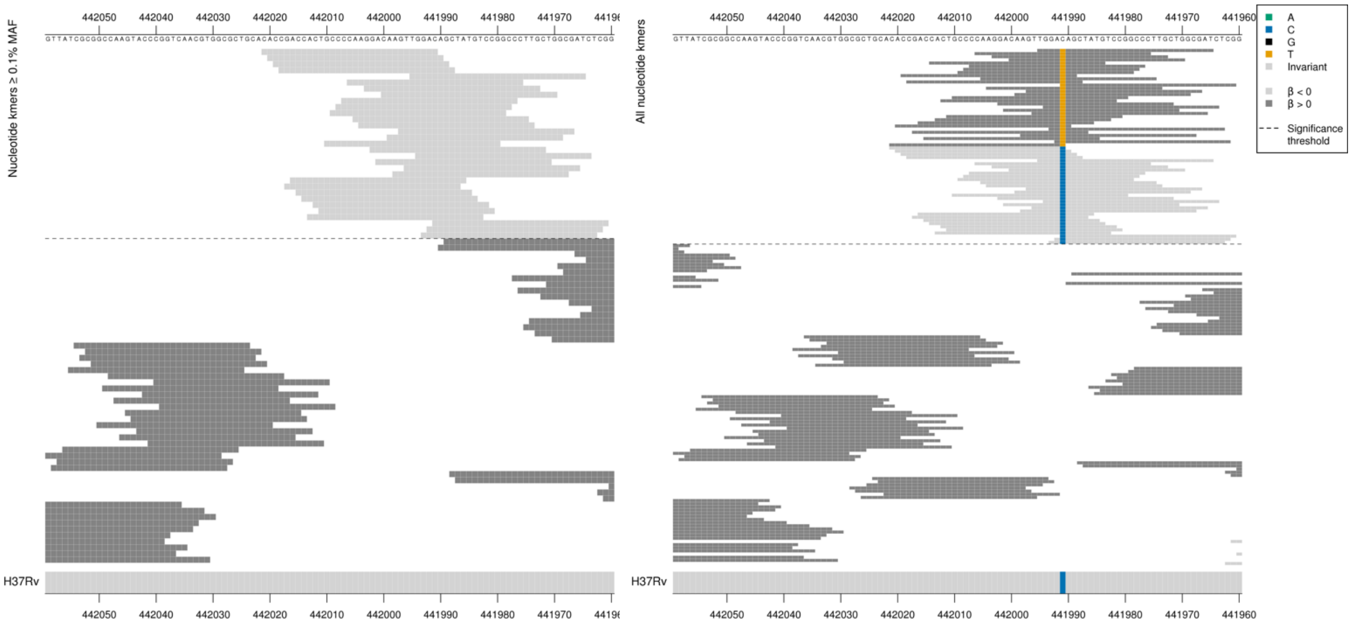

Capturing a synonymous mutation at position 441991/309, codon 103.

Low MIC-associated kmers = GAC = Asp / D

High MIC-associated low frequency kmers = GAT = Asp / D

Low MAF kmers = 441991/309T (103D aa)

| Kmer | -log₁₀ *p* | β | MAC | MAF | Ps | Variants |
|---|---|---|---|---|---|---|
| CACCGACCACTGCCCCAAGGACAAGTTGGAC | 39.97 | -4.21 | 8 | 0.11 | 442021 | 441991/309C (103D aa) |

# Rv2180c

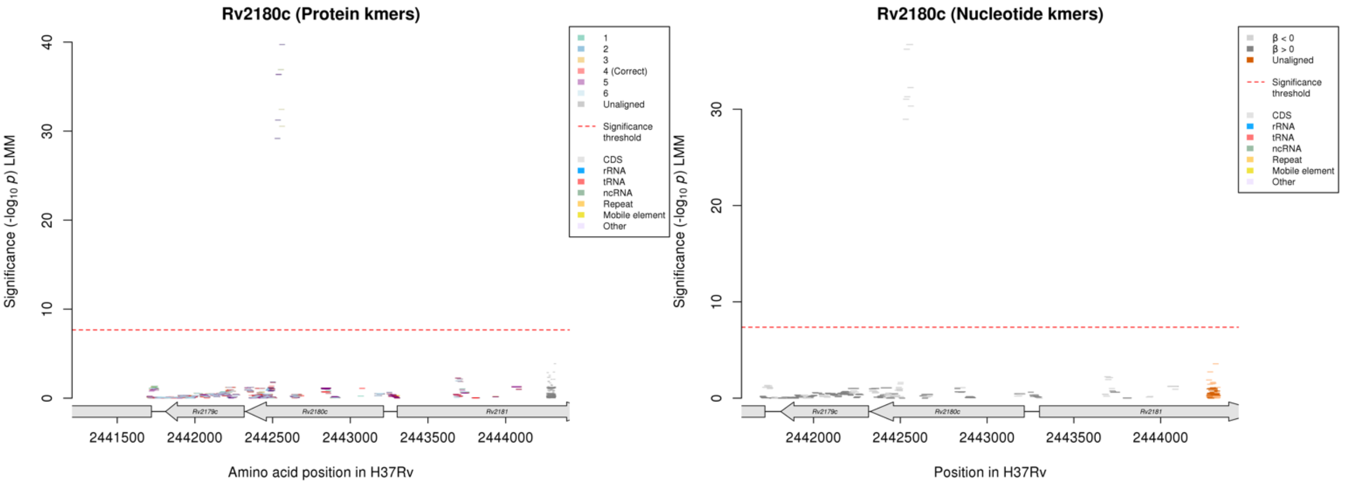

No significant kmers aligned to the correct reading frame for the protein kmer analysis.

Nucleotide kmers:

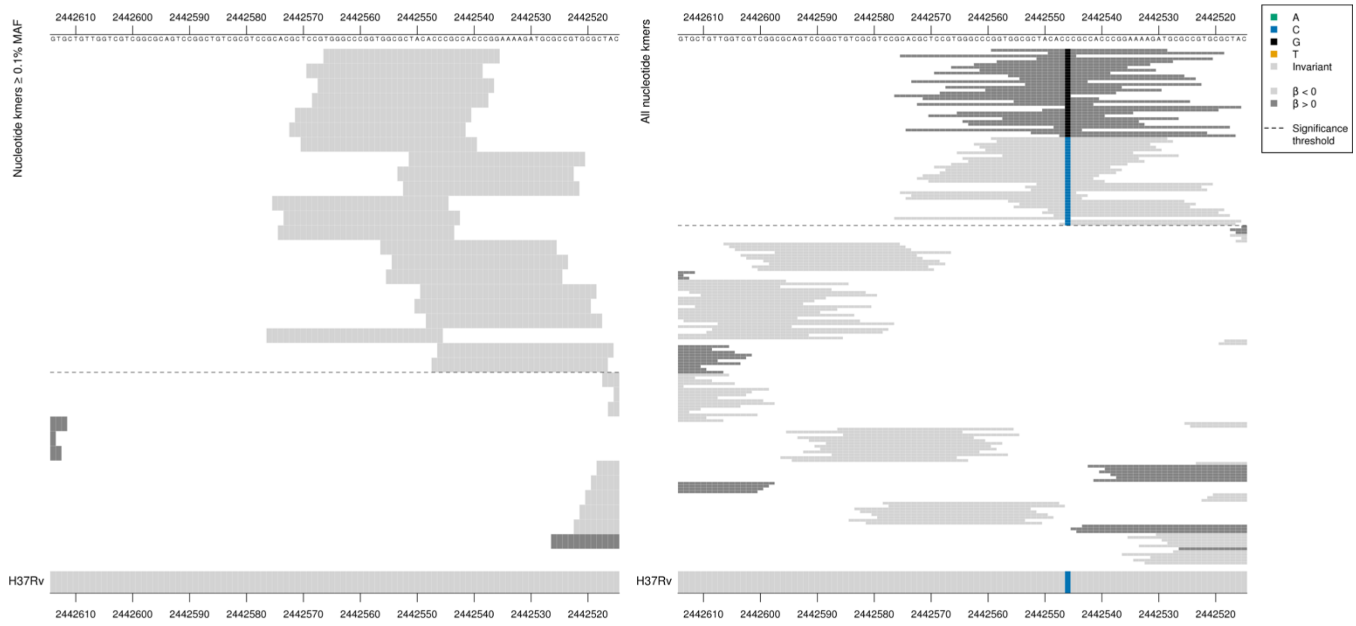

Capturing a synonymous mutation at position 2442546/669.
Low MIC-associated kmers = ACC = Thr / T
High MIC-associated low frequency kmers = ACG = Thr / T

Low MAF kmers = 2442546/669G (223T aa), 2442546/669C (223T aa)

| Kmer | $-\log_{10} p$ | β | MAC | MAF | Ps | Variants |
|---|---|---|---|---|---|---|
| TGGGCCCGGTGGCGCTACACCCGCCACCCGG | 36.71 | -3.91 | 8 | 0.11 | 2442566 | 2442546/669C (223T aa) |

#### *gap*

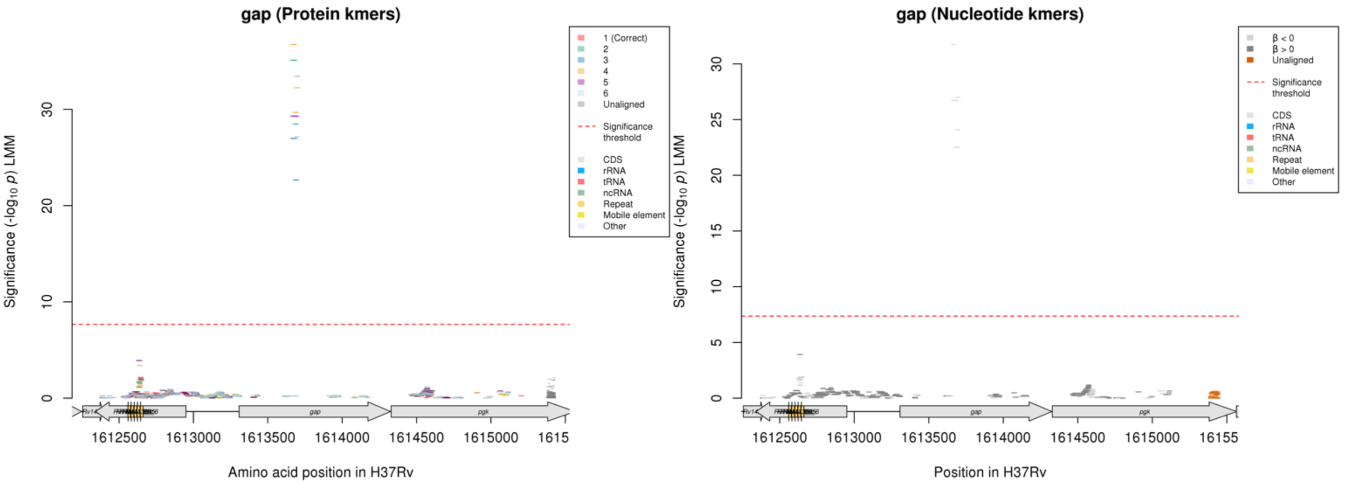

No significant kmers aligned to the correct reading frame in the protein kmer analysis.

Nucleotide kmers:

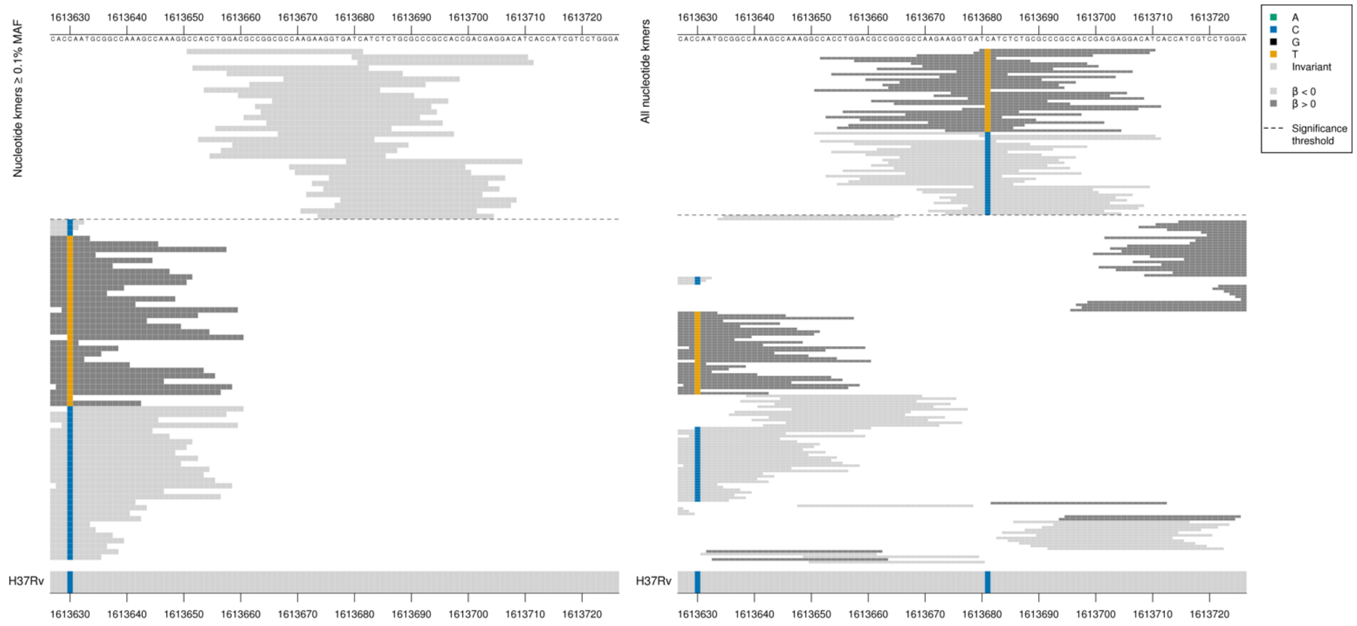

*gap* begins at position 1613307.
Capturing a synonymous mutation at position 1613681/375
High MIC-associated kmers = ATT = Ile / I
Low MIC-associated kmers = ATC = Ile / I

Low MAF kmers = 1613681/375T (125I aa)

| Kmer | -log$_{10}$ *p* | β | MAC | MAF | Ps | Variants |
|---|---|---|---|---|---|---|
| CCACCTGGACGCCGGCGCCAAGAAGGTGATC | 31.73 | -3.30 | 10 | 0.13 | 1613651 | 1613681/375C (125I aa) |

#### *lprF:Rv1371* [R]

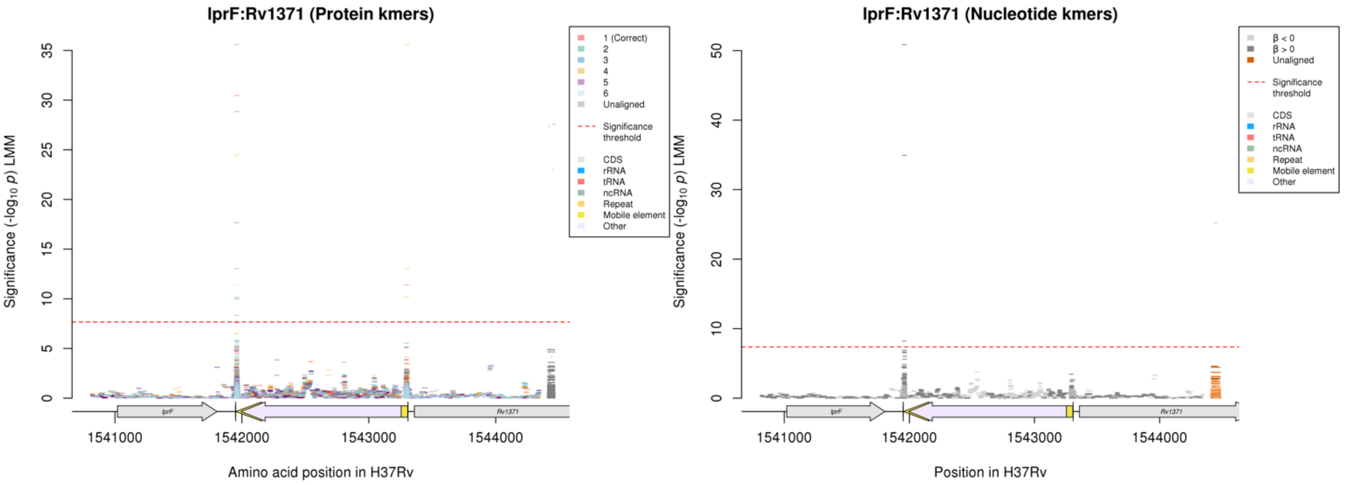

Repeat region at the end of a mobile element.
Nucleotide kmers:

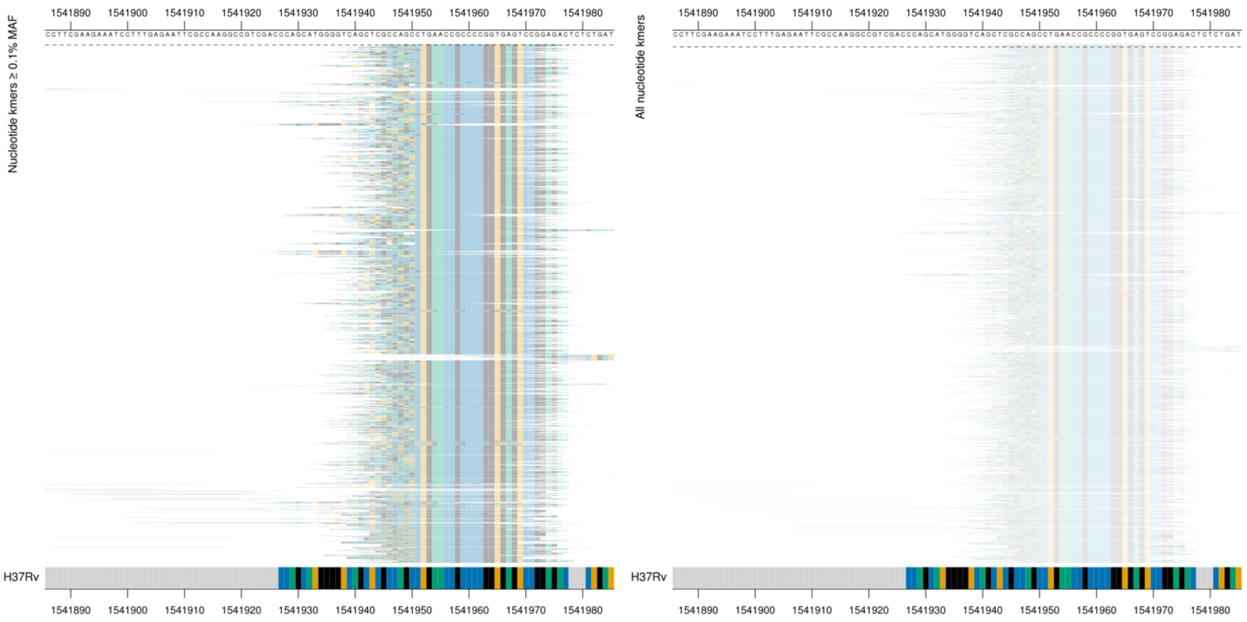

| Kmer | -log$_{10}$ *p* | β | MAC | MAF | Ps | Variants |
|---|---|---|---|---|---|---|
| AGCTGACTGAACCGCCCCGGTGAGTCCGGAG | 50.87 | 4.35 | 8 | 0.11 | 1541945 | Repeat region |

## *Rv0914c*

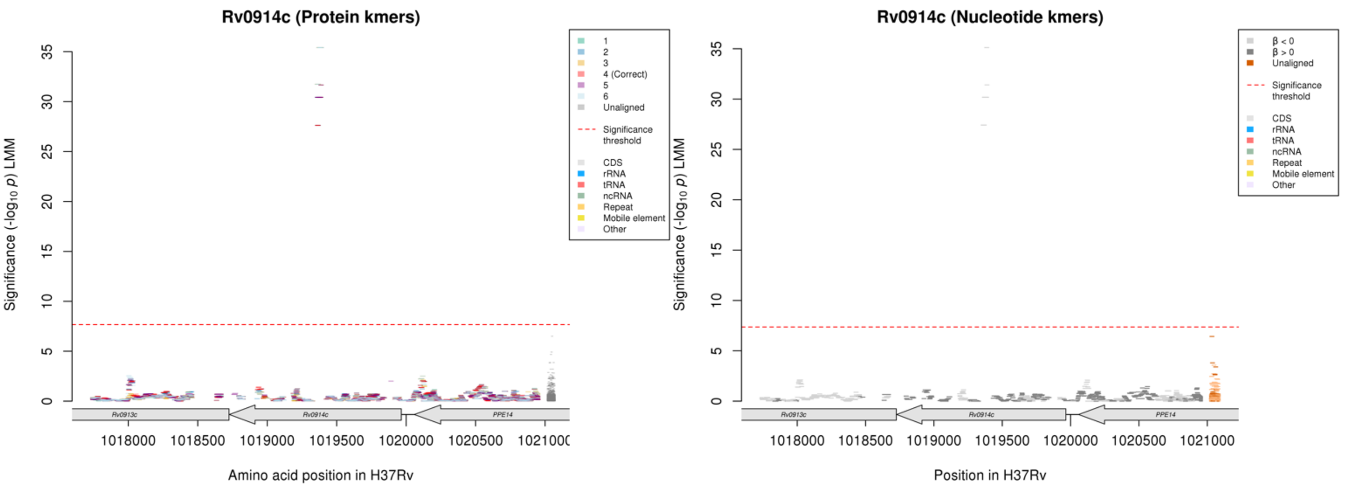

Protein kmers:

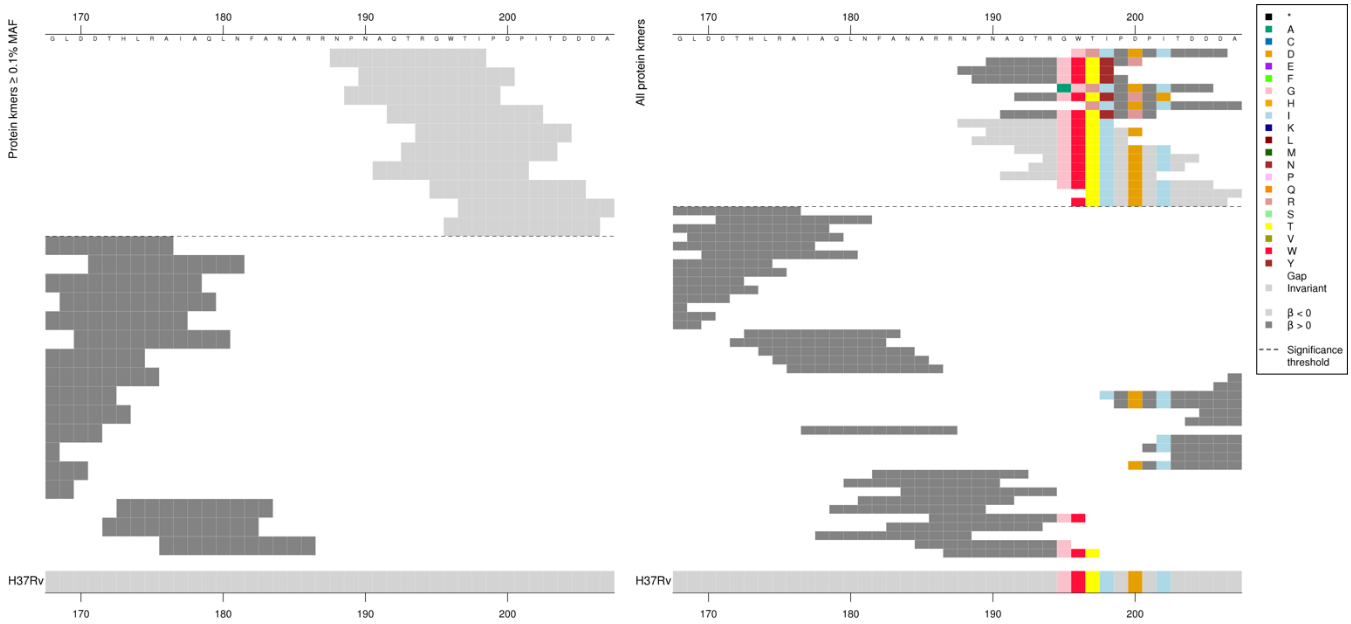

The same region for the nucleotide kmer analysis:

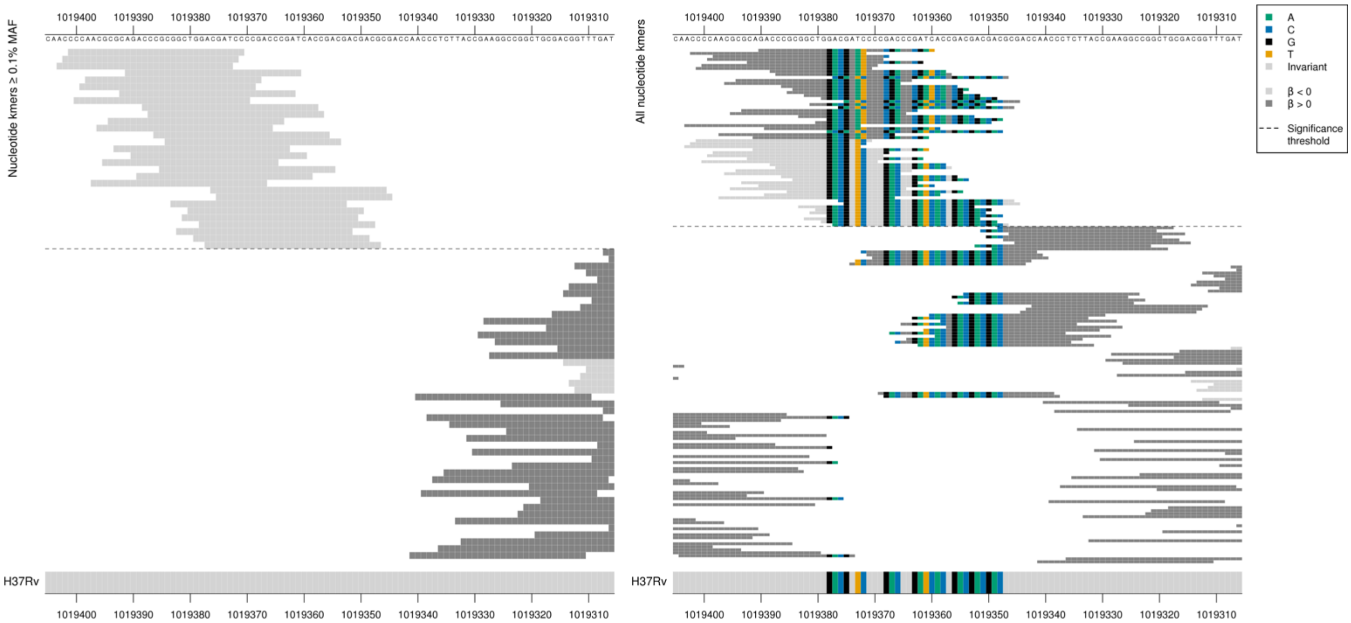

High MIC-associated kmers = A insertion between positions 1019375-1019373/591-593.

Low MAF kmers = A insertion between positions 1019375-1019373/591-593

| Kmer | $-\log_{10} p$ | β | MAC | MAF | Ps | Variants |
|---|---|---|---|---|---|---|
| CCCAACGCGCAGACCCGCGGCTGGACGATCC | 35.11 | -3.70 | 9 | 0.12 | 1019401 | WT |

Delamanid (DLM)

# *Rv1200*

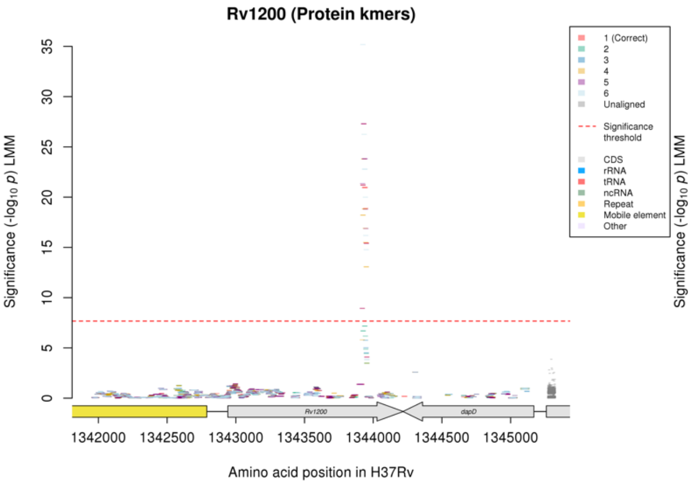

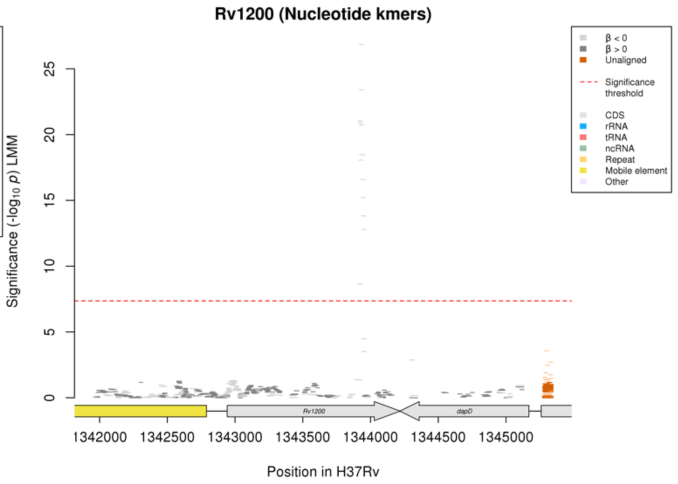

Protein kmers:

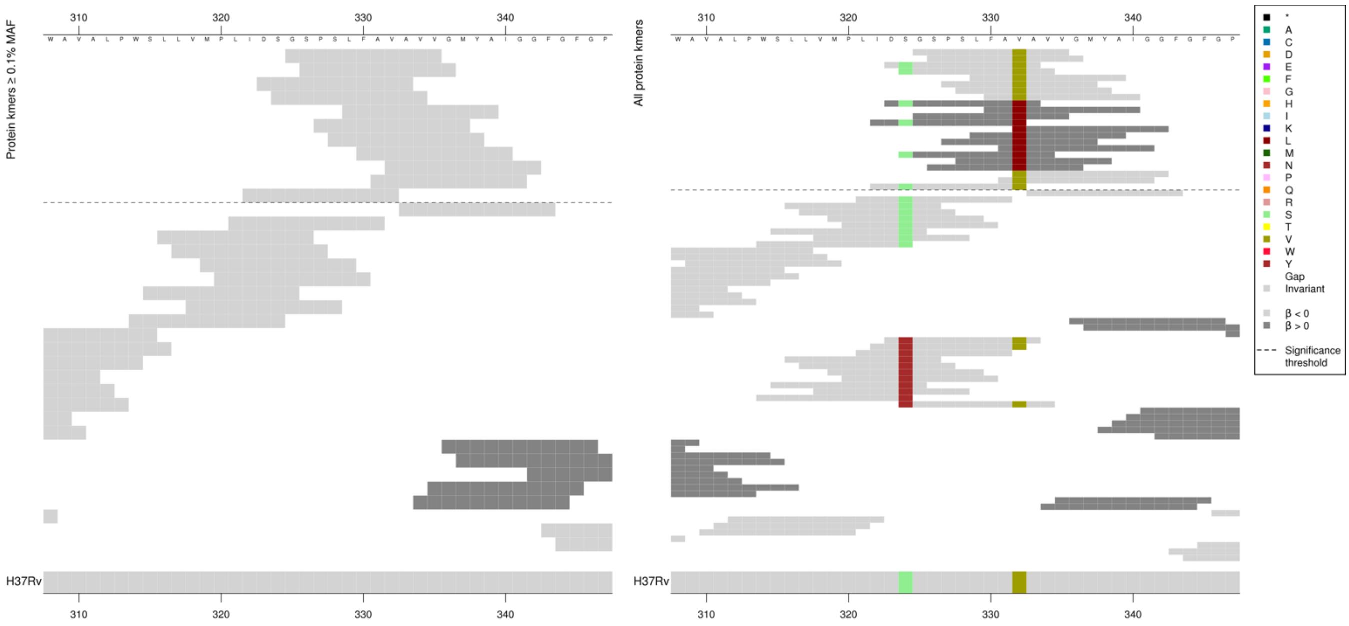

Low MAF kmers = 332L.

| Kmer | $-\log_{10} p$ | β | MAC | MAF | Ps | Variants |
|---|---|---|---|---|---|---|
| GSPSLFAVAVV | 27.30 | -3.55 | 8 | 0.11 | 325 | 332V |
| DSGSPSLFAVA | 21.20 | -2.30 | 14 | 0.19 | 323 | 324S, 332V |

### *fadE10*

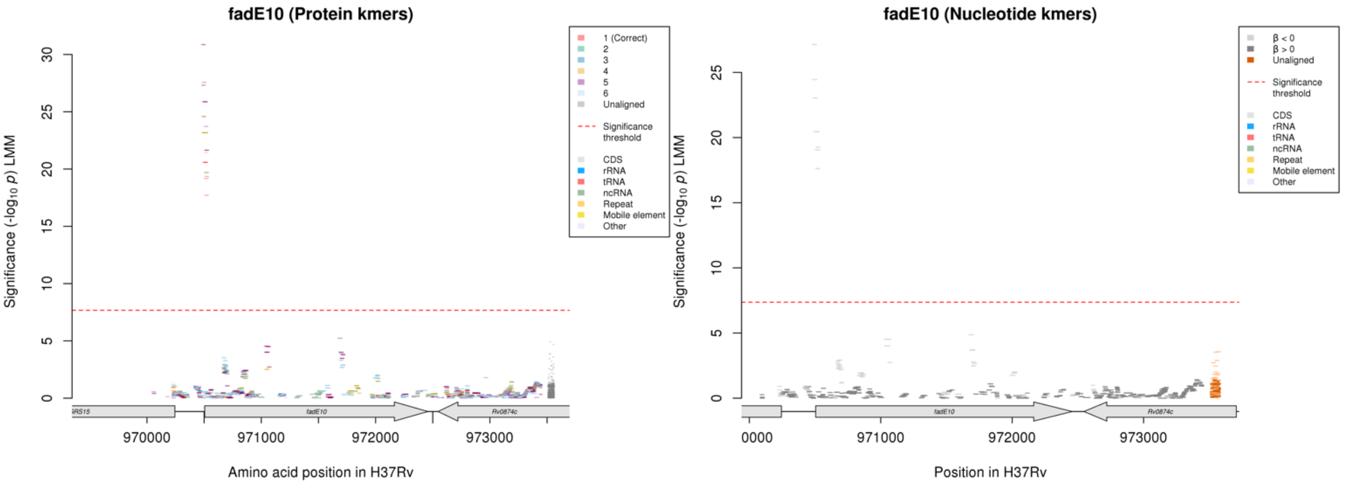

#### Protein kmers:

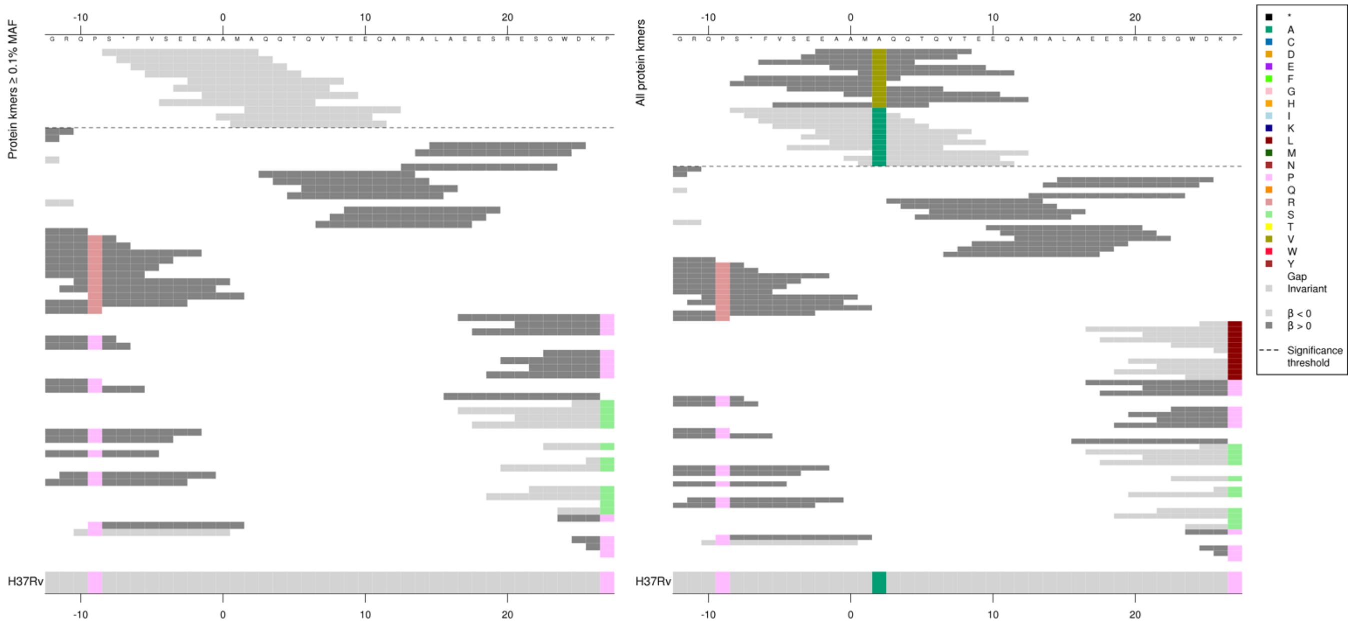

Low MAF kmers = 2V.

| Kmer | -log$_{10}$ *p* | β | MAC | MAF | Ps | Variants |
|---|---|---|---|---|---|---|
| S*FVSEEAAMA | 27.32 | -2.95 | 11 | 0.15 | -8 | 2A |

### *dinP*

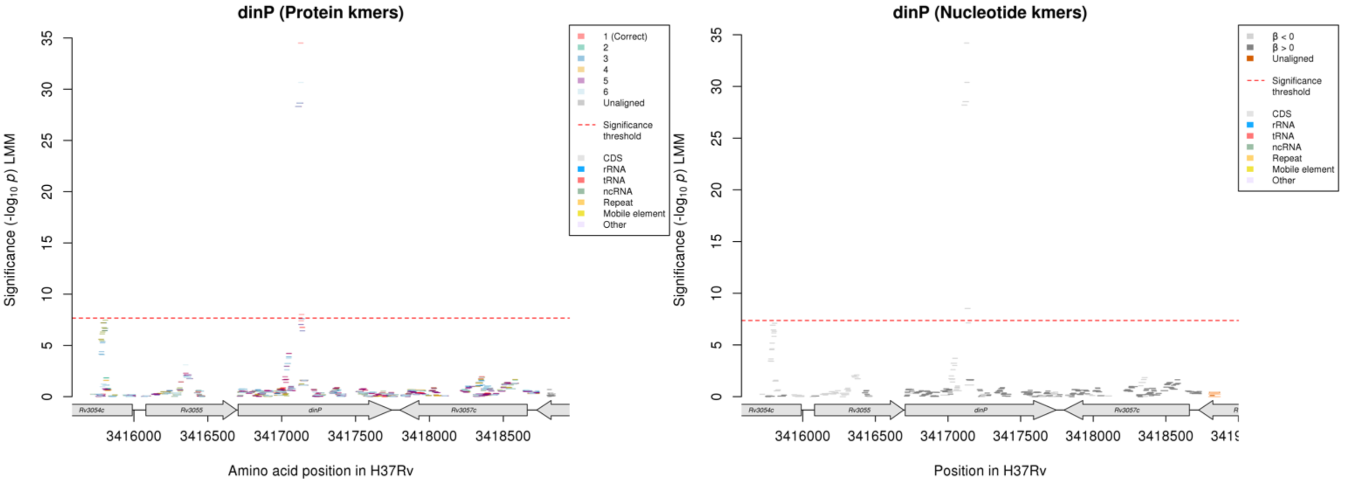

Protein kmers:

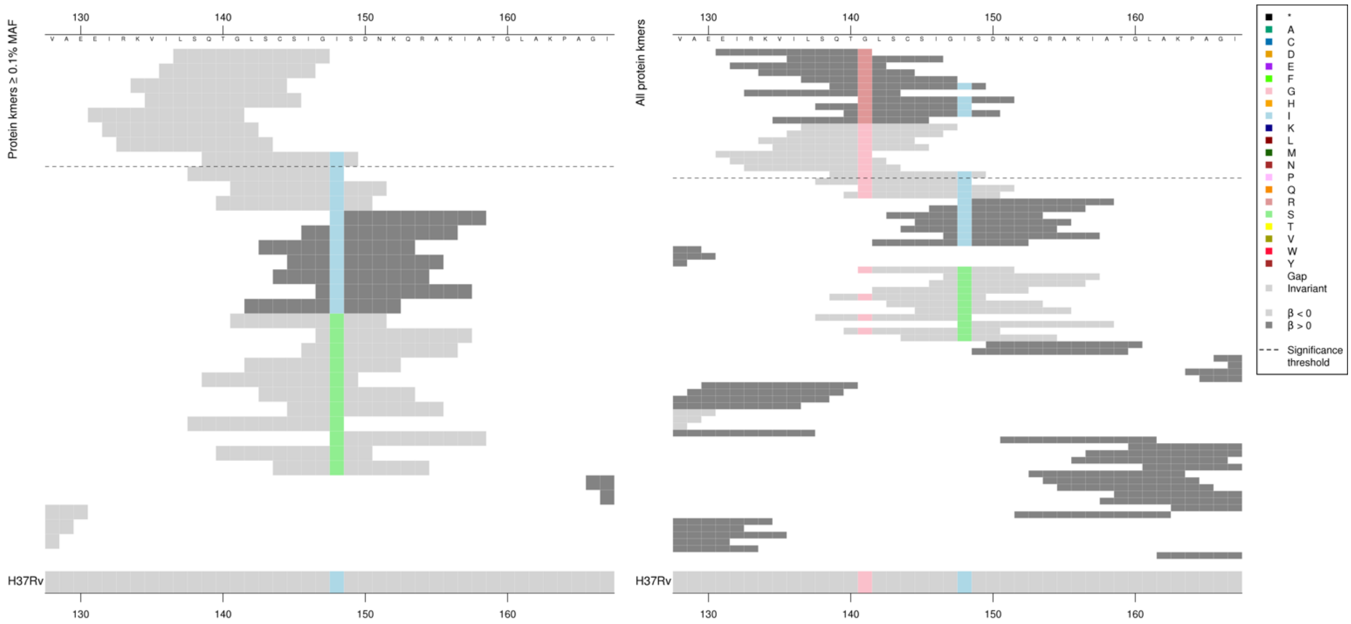

141G present in between 8-10 genomes.

Low MAF kmers = 141R.

| Kmer | $-\log_{10} p$ | β | MAC | MAF | Ps | Variants |
|---|---|---|---|---|---|---|
| LSQTGLSCSIG | 34.50 | -3.71 | 8 | 0.11 | 137 | 141G |
| QTGLSCSIGIS | 8.01 | -1.05 | 59 | 0.78 | 139 | 141G, 148I |

#### *mmpL8*

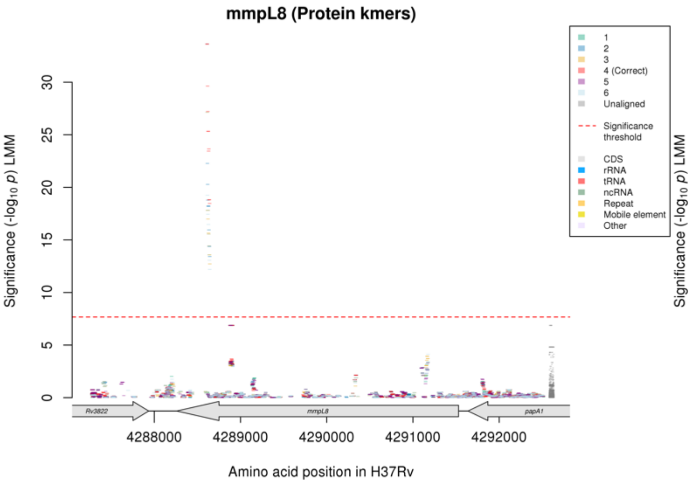

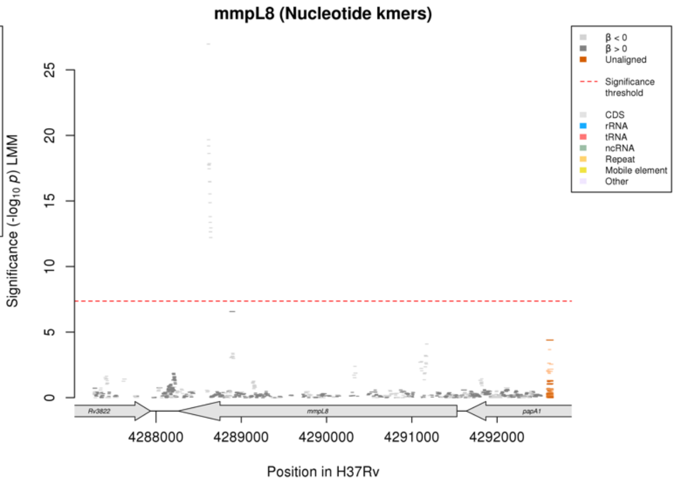

Protein kmers:

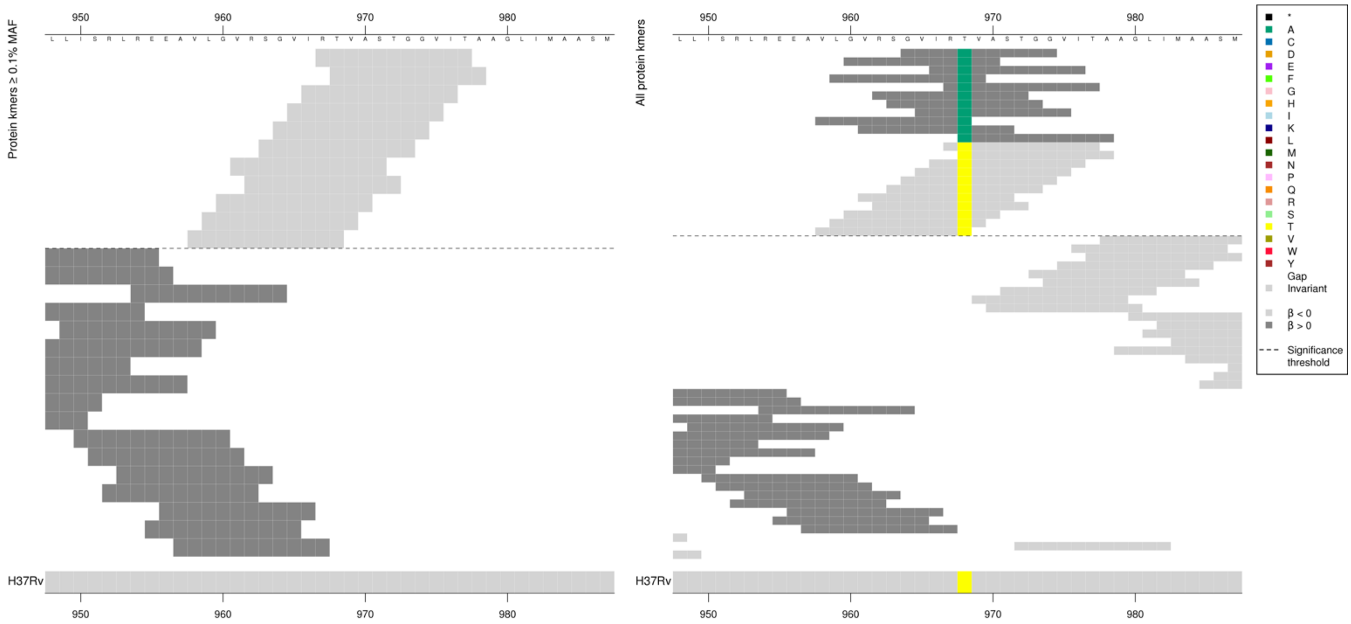

Low MAF kmers = 968A.

| Kmer | $-\log_{10} p$ | β | MAC | MAF | Ps | Variants |
|---|---|---|---|---|---|---|
| RTVASTGGVIT | 33.62 | -3.33 | 10 | 0.13 | 967 | 968T |

#### *cut1*

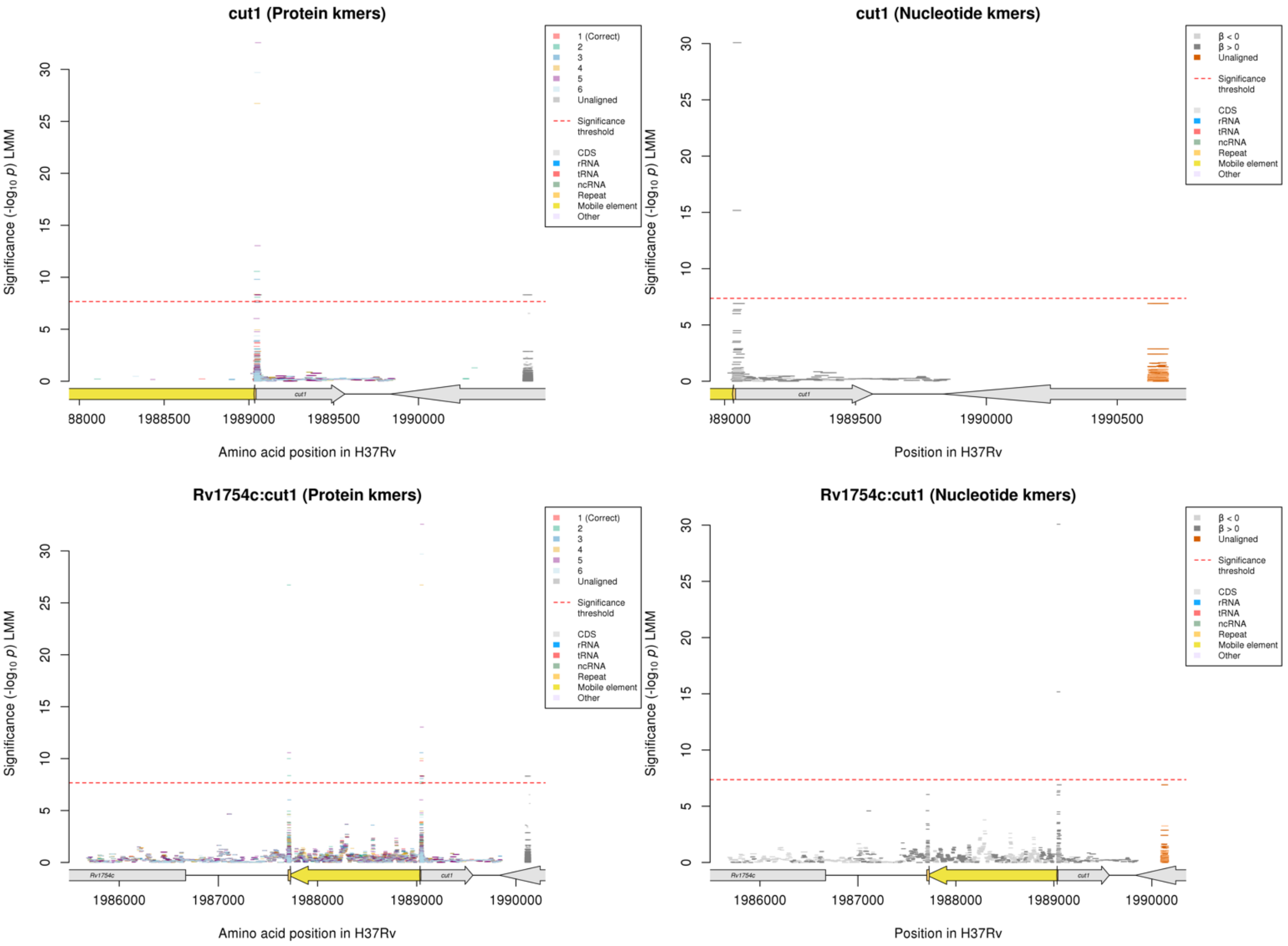

Nucleotide kmers:

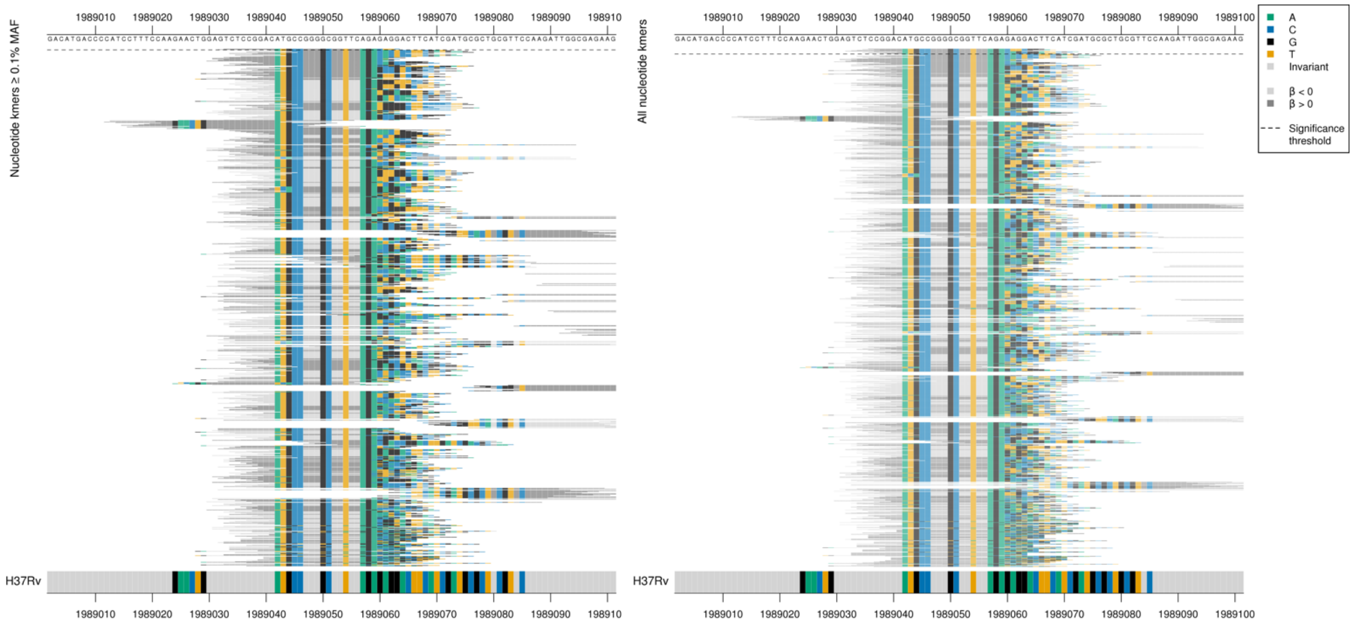

Repeat region

Range of MACs in region up to 213, showing the top kmer with no MAF threshold and the top kmer above the MAF threshold.

| Kmer | $-\log_{10} p$ | β | MAC | MAF | Ps | Variants |
|---|---|---|---|---|---|---|
| TCTCCGGACATGCCGGGGCGGTTCAGACTTC | 30.08 | 2.59 | 14 | 0.19 | 1989033 | Repeat region (same kmer for PPE39) |

#### *PPE39*

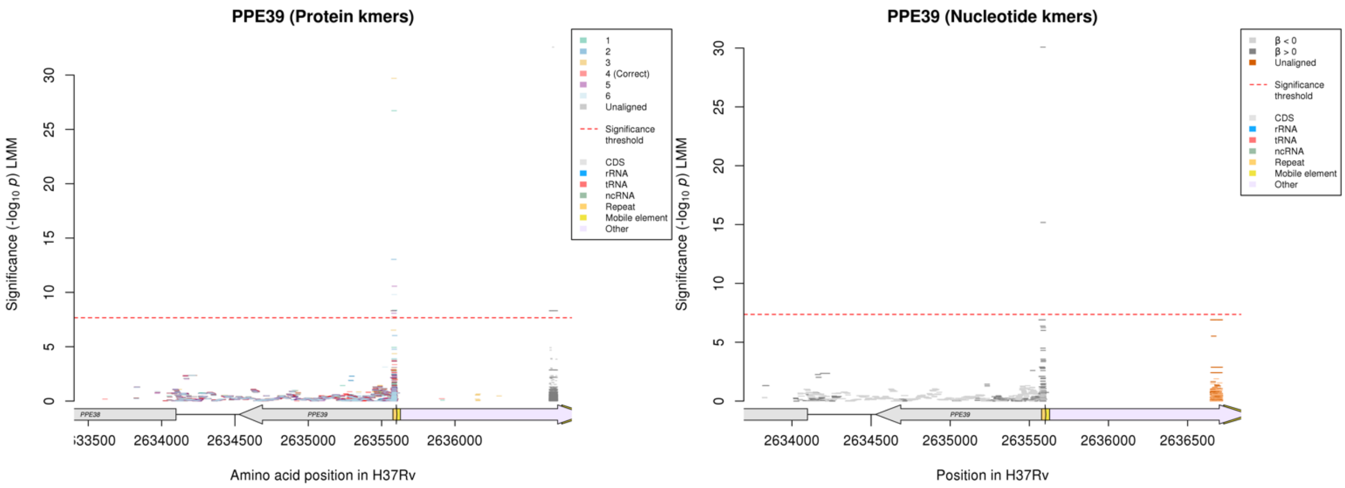

#### Nucleotide kmers:

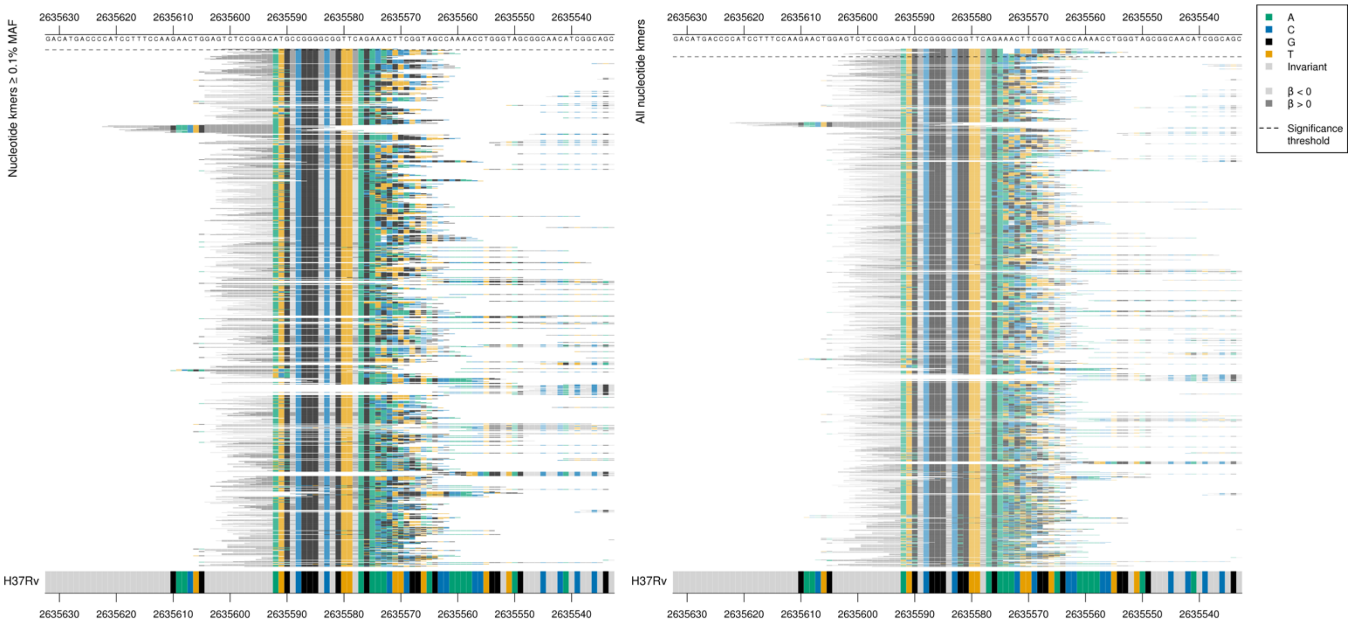

#### Repeat region

| Kmer | $-\log_{10} p$ | β | MAC | MAF | Ps | Variants |
|---|---|---|---|---|---|---|
| TCTCCGGACATGCCGGGGCGGTTCAGACTTC | 30.08 | 2.59 | 14 | 0.19 | 2635601 | Repeat region (same kmer for cut1) |

#### *Rv3430a:gadB*

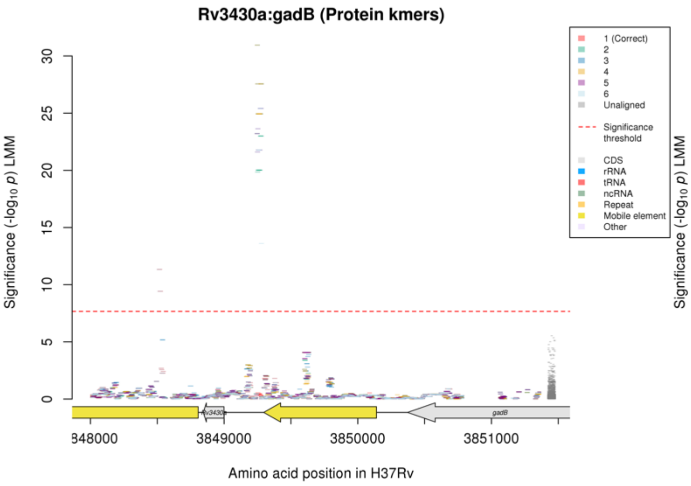

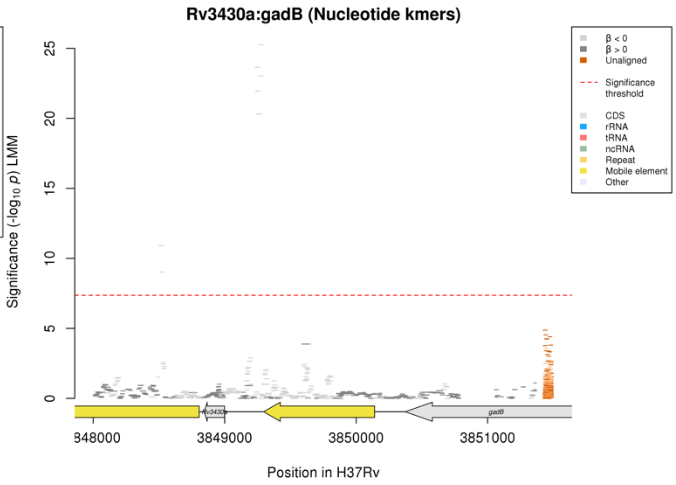

Nucleotide kmers:

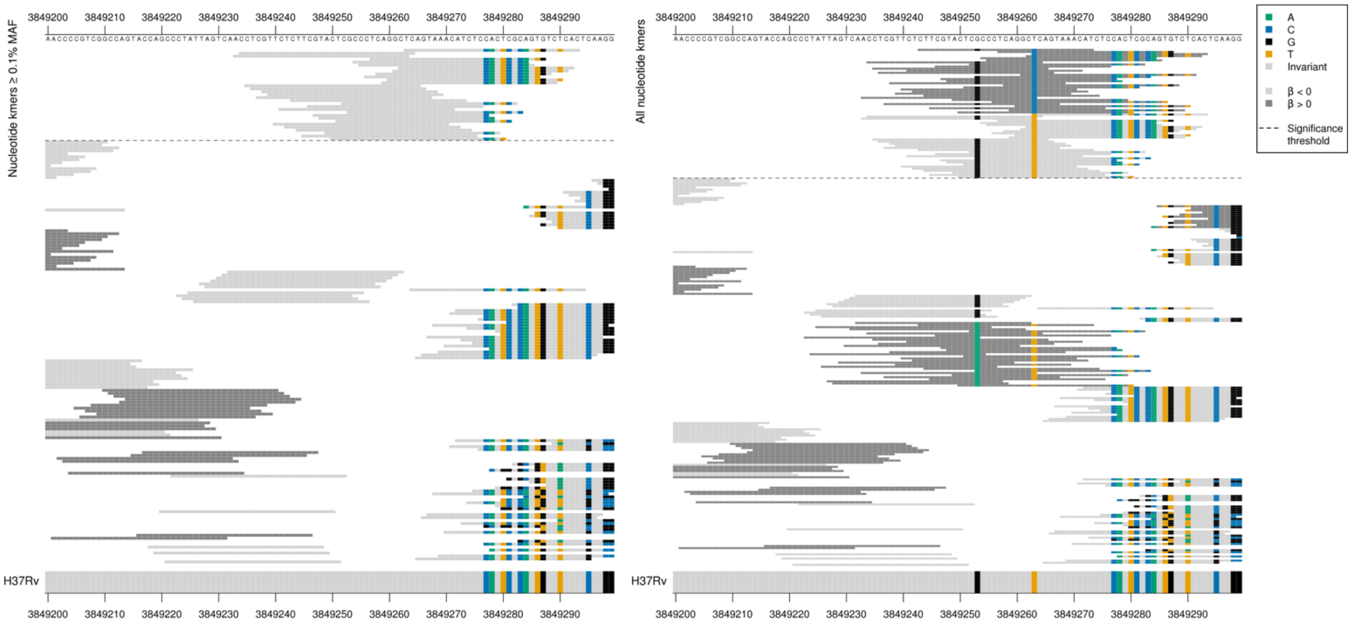

All high MIC-associated kmers were the same *p*-value, just capturing the variant at position 3849263, but the low MIC-associated kmers were a range of *p*-values, possibly influenced by the surrounding variants.

Low MAF kmers = 3849263C

| Kmer | -log$_{10}$ *p* | β | MAC | MAF | Ps | Variants |
|---|---|---|---|---|---|---|
| TCAGTAAACATCTCCACTCGCAGTGTCTCAC | 25.25 | -2.48 | 13 | 0.17 | 3849263 | 3849263T |
| CCTCGTTCTCTTCGTACTCGCCCTCAGGCTC | 23.65 | -2.28 | 17 | 0.22 | 3849234 | 3849253G, 3849263T |

## *Rv1429*

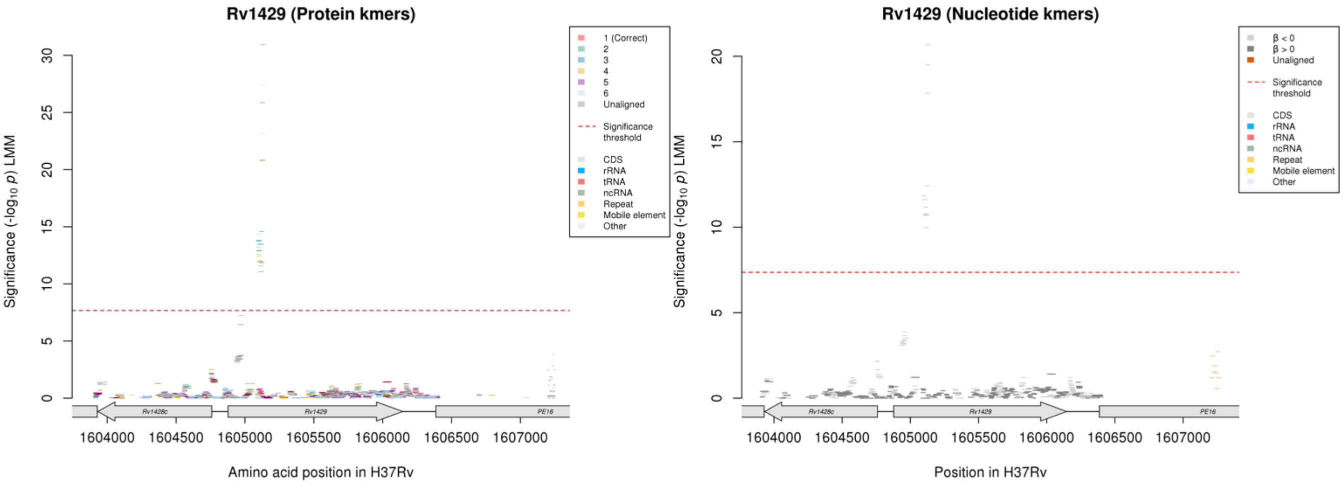

No significant kmers aligned to the correct reading frame in the protein kmer analysis.

Nucleotide kmers:

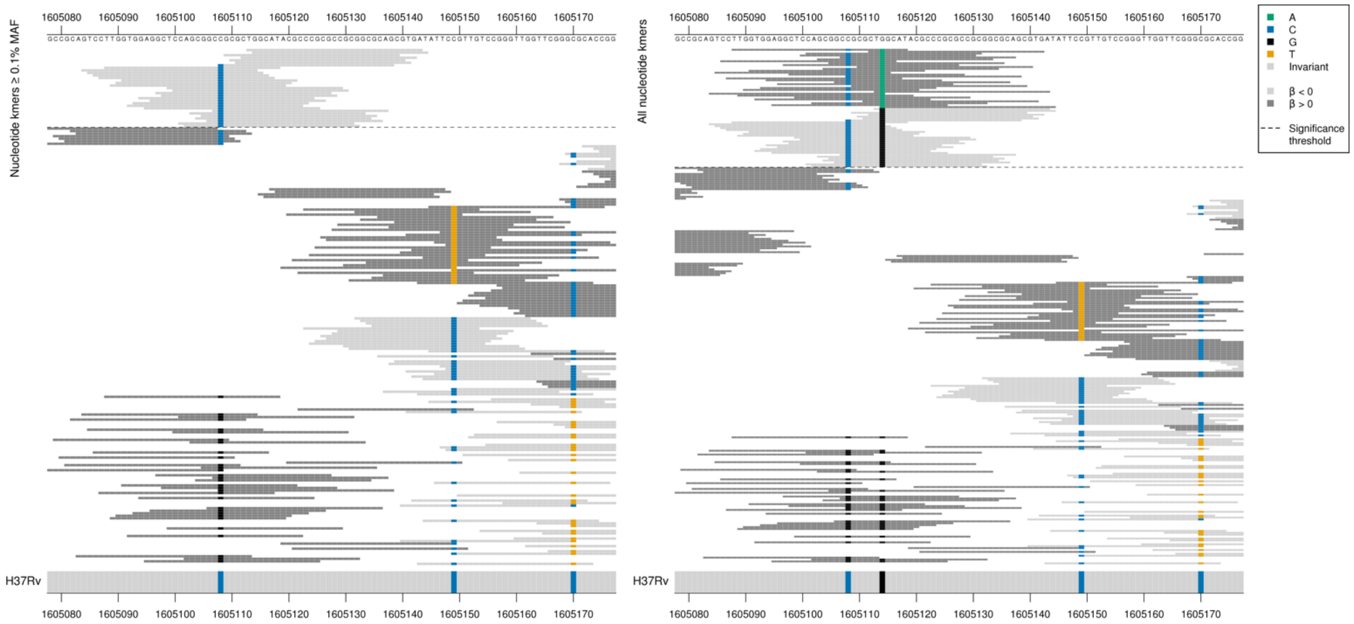

All high MIC-associated kmers were the same *p*-value, just capturing the mutation at position 1605114/237. The low MIC-associated kmers covered a range of *p*-values, showing the top kmer for the two allele combinations in the table.

Low MAF kmers = 1605114/237A.

| Kmer | $-\log_{10} p$ | β | MAC | MAF | Ps | Variants |
|---|---|---|---|---|---|---|
| TGGCATACGCCCGCGCCGCGGCGCAGCGTGA | 20.68 | -2.21 | 14 | 0.19 | 1605113 | 1605114/237G |
| CGCGCTGGCATACGCCCGCGCCGCGGCGCAG | 12.42 | -1.34 | 24 | 0.32 | 1605108 | 1605108/77C, 1605114/237G |

## *Rv3847*

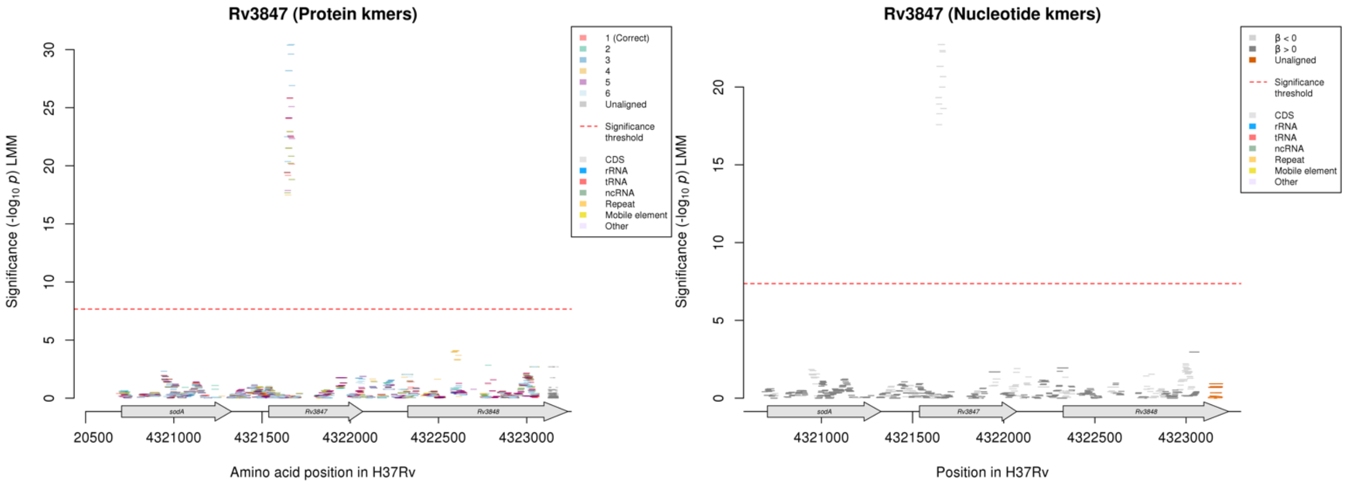

Nucleotide kmers:

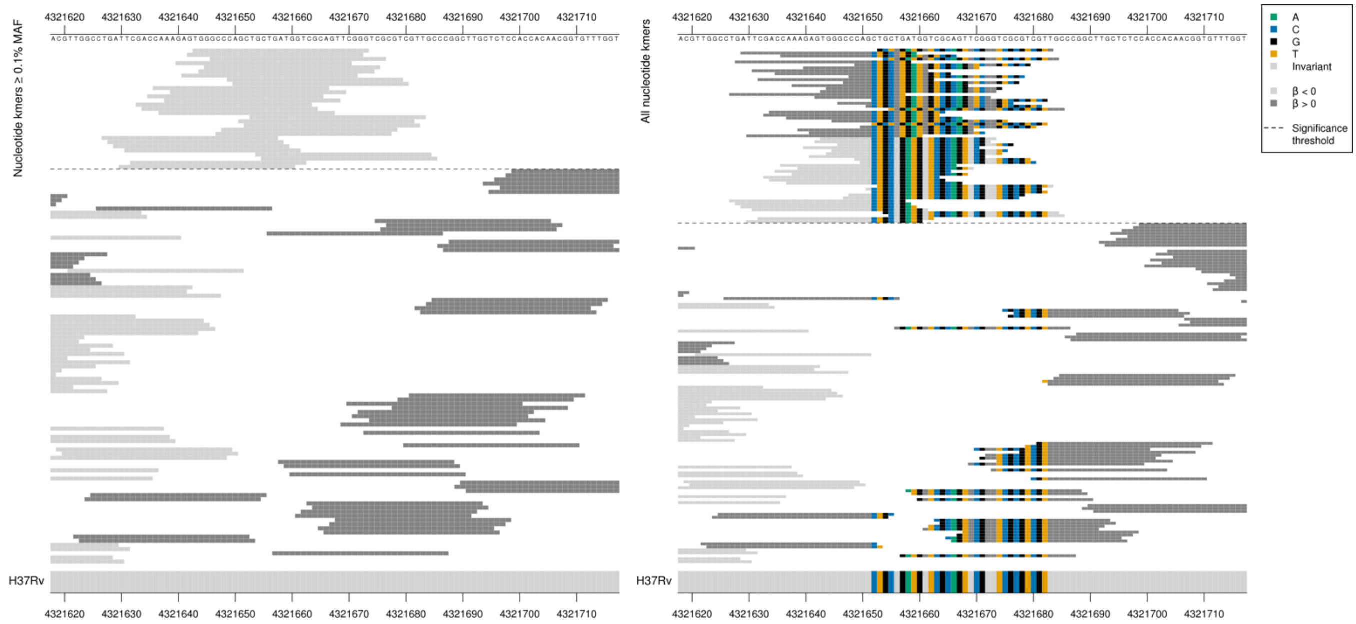

Low MAF kmers = T insertion between 4321655-4321657/118-120

| Kmer | $-\log_{10} p$ | β | MAC | MAF | Ps | Variants |
|---|---|---|---|---|---|---|
| TGGGCCCAGCTGCTGATGGTCGCAGTTCGGG | 22.73 | -2.48 | 12 | 0.16 | 4321643 | WT |

#### *pknH*

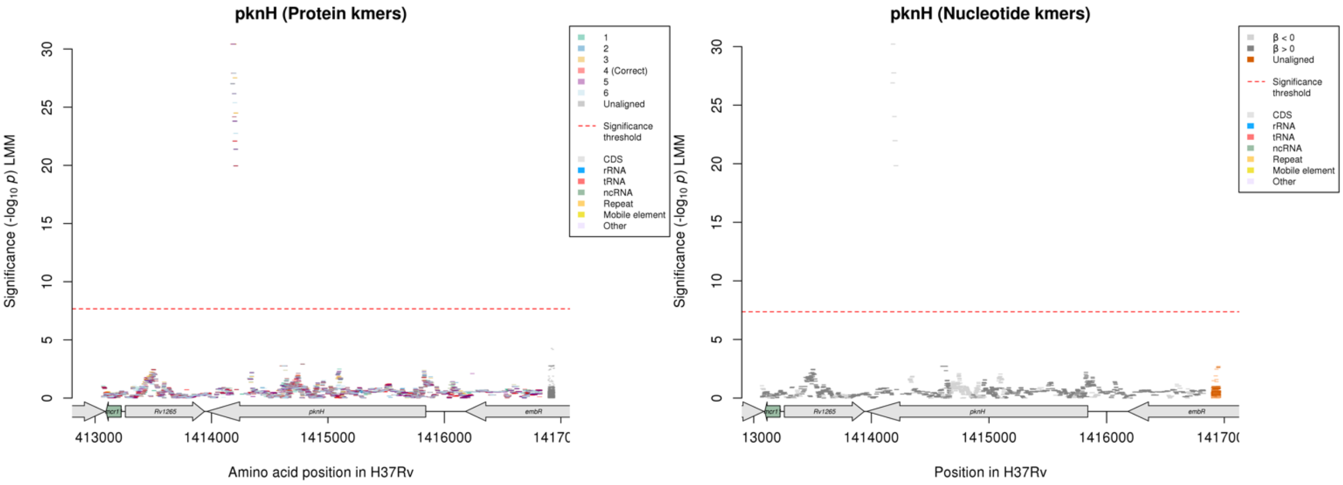

Protein kmers:

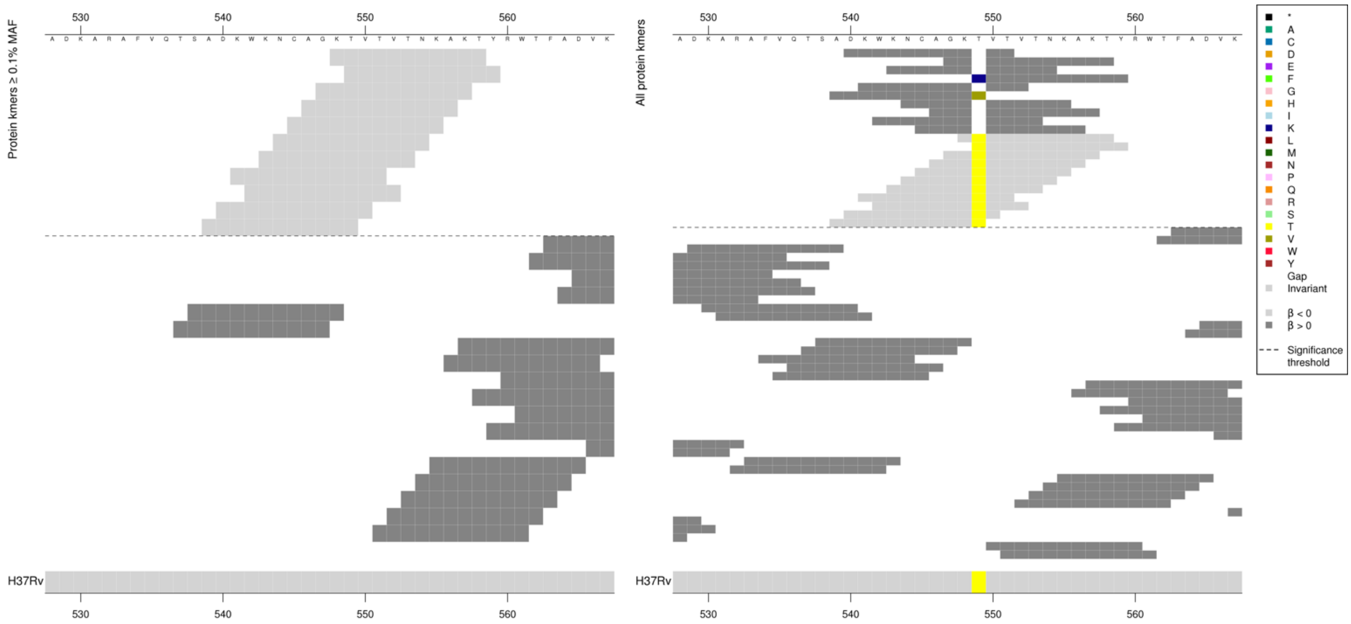

The same region for the nucleotide kmers:

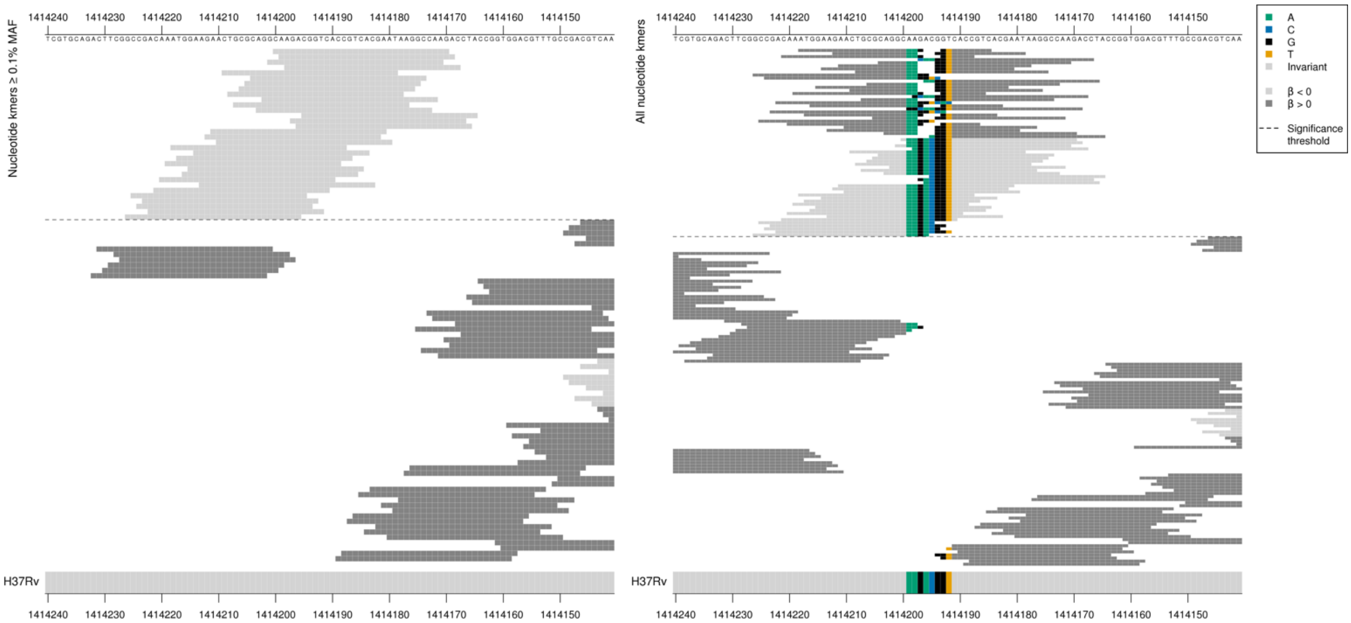

All high MIC-associated kmers were the same *p*-value. Low MIC-associated kmers covered a range of *p*-values.
Low MAF kmers were capturing either a deletion of ACG positions 1414196-1414194/1645-1647 or GAC positions 1414197-1414195/1644-1646.

| Kmer | $-\log_{10} p$ | β | MAC | MAF | Ps | Variants |
|---|---|---|---|---|---|---|
| CAAGACGGTCACCGTCACGAATAAGGCCAAG | 30.21 | -3.29 | 9 | 0.12 | 1414200 | WT |

### *plsC*

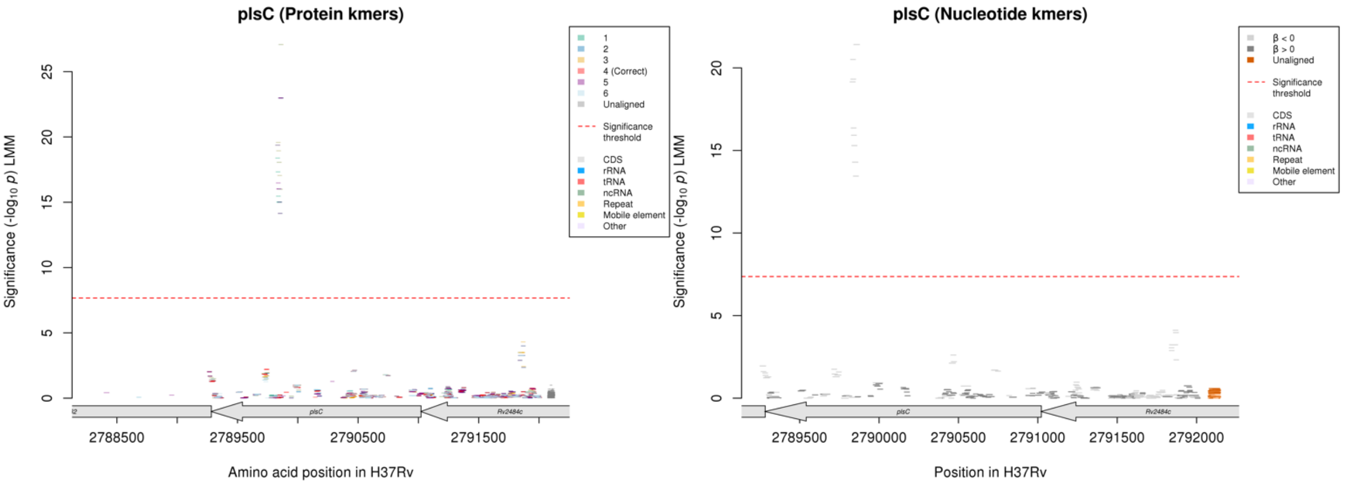

No significant kmers above the MAF threshold aligned to the correct reading frame for the protein kmer analysis.

Nucleotide kmers:

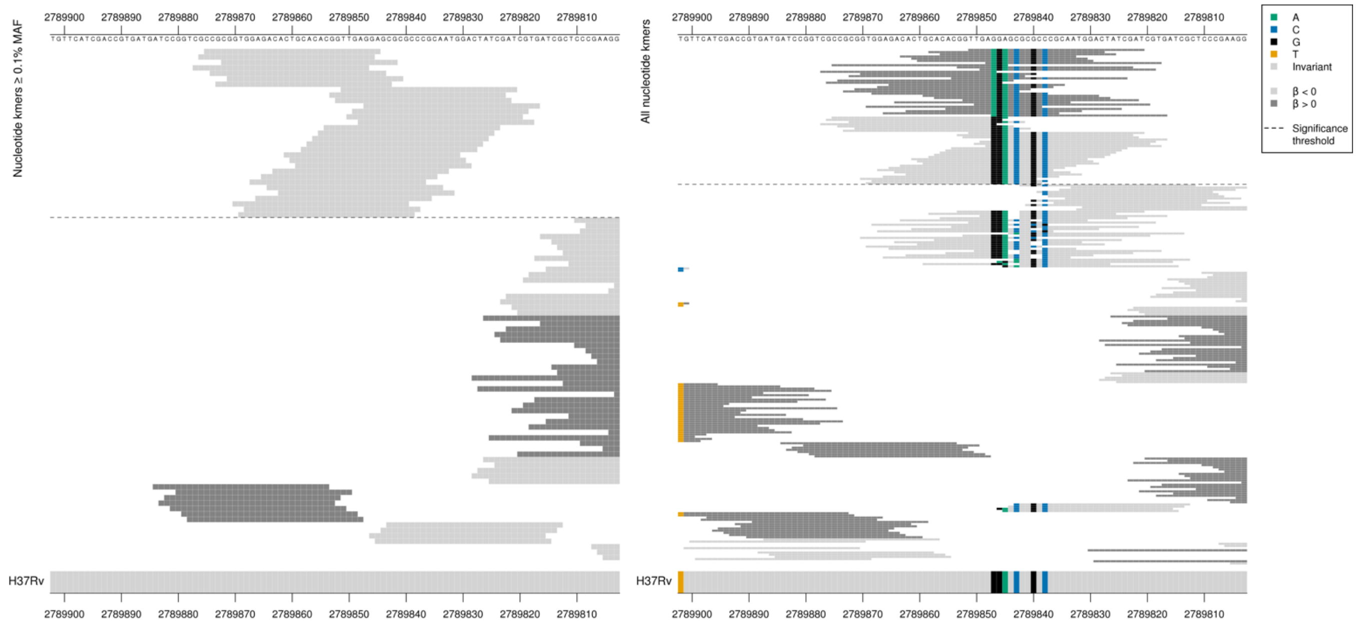

plsC begins on the reverse strand at position 2791022

Low MIC-associated kmer p-values varied depending on whether they covered the region containing deletions in non-significant kmers.

Low MAF kmers = 2789847A

| Kmer | -log10 *p* | β | MAC | MAF | Ps | Variants |
|---|---|---|---|---|---|---|
| CCGCGGTGGAGACACTGCACACGGTTGAGGA | 21.41 | -2.24 | 14 | 0.19 | 2789875 | 2789847G |

### *agpS*

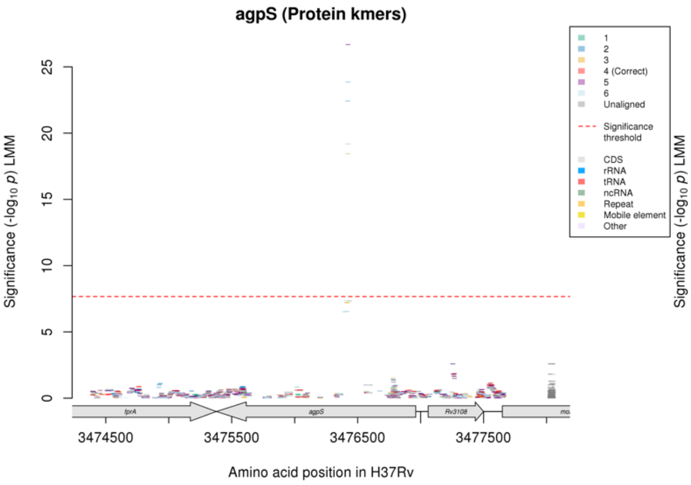

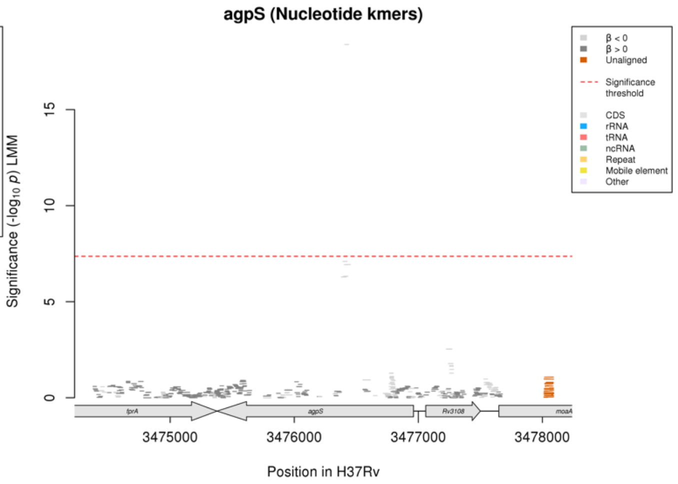

#### Nucleotide kmers:

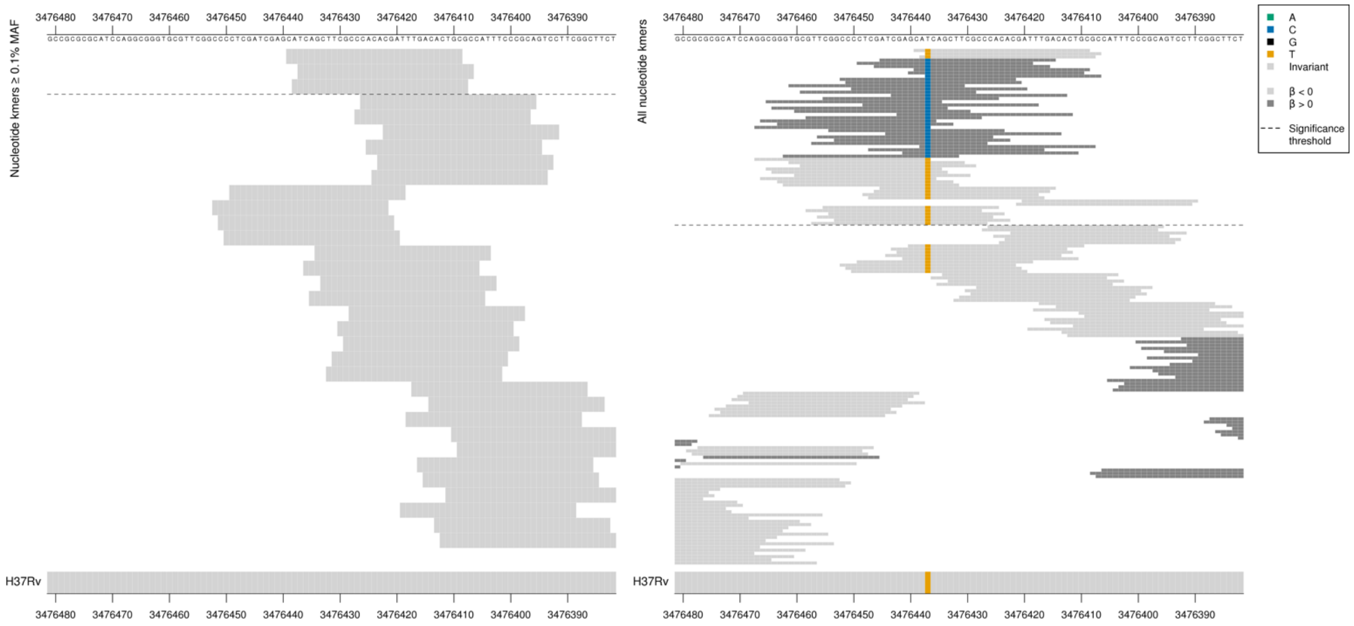

#### Low MAF kmers = 3476437C

| Kmer | $-\log_{10} p$ | β | MAC | MAF | Ps | Variants |
|---|---|---|---|---|---|---|
| CATCAGCTTCGCCCACACGATTTGACACTGC | 18.39 | -2.44 | 10 | 0.13 | 3476439 | 3476437T |

# *Rv3263*

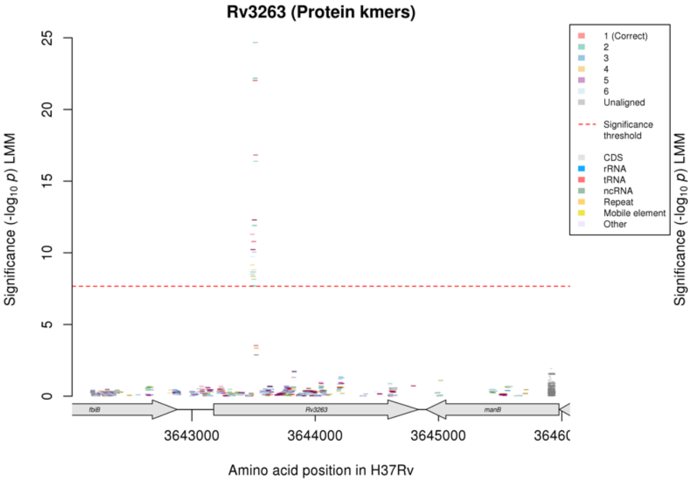

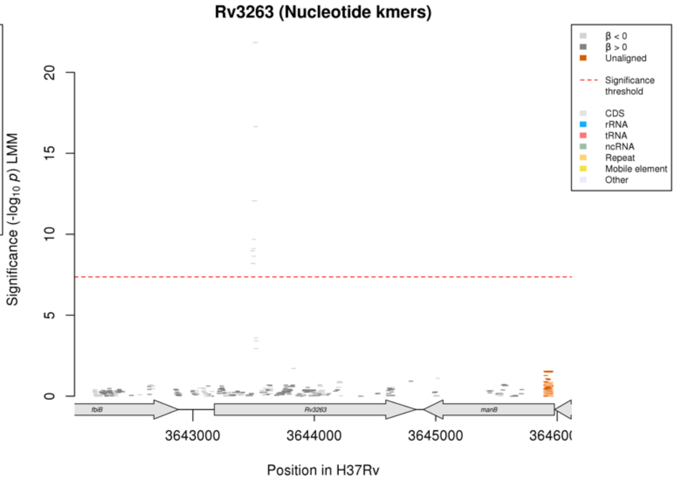

Protein kmers:

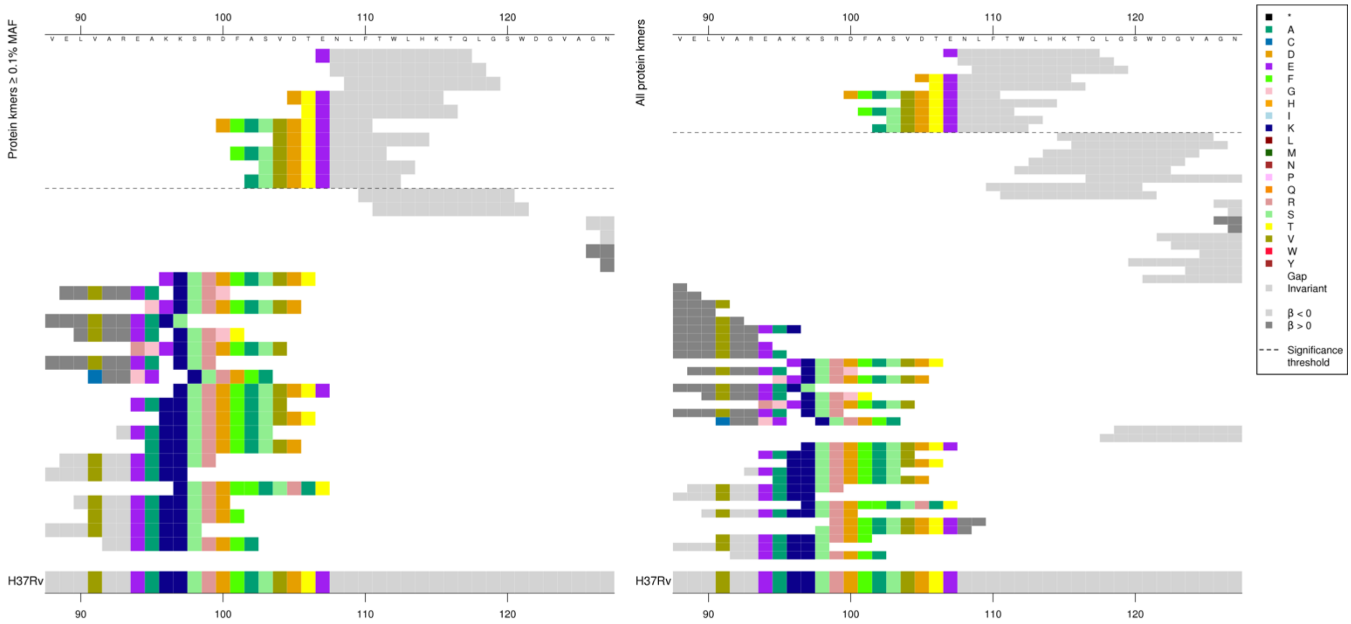

Nucleotide kmers:

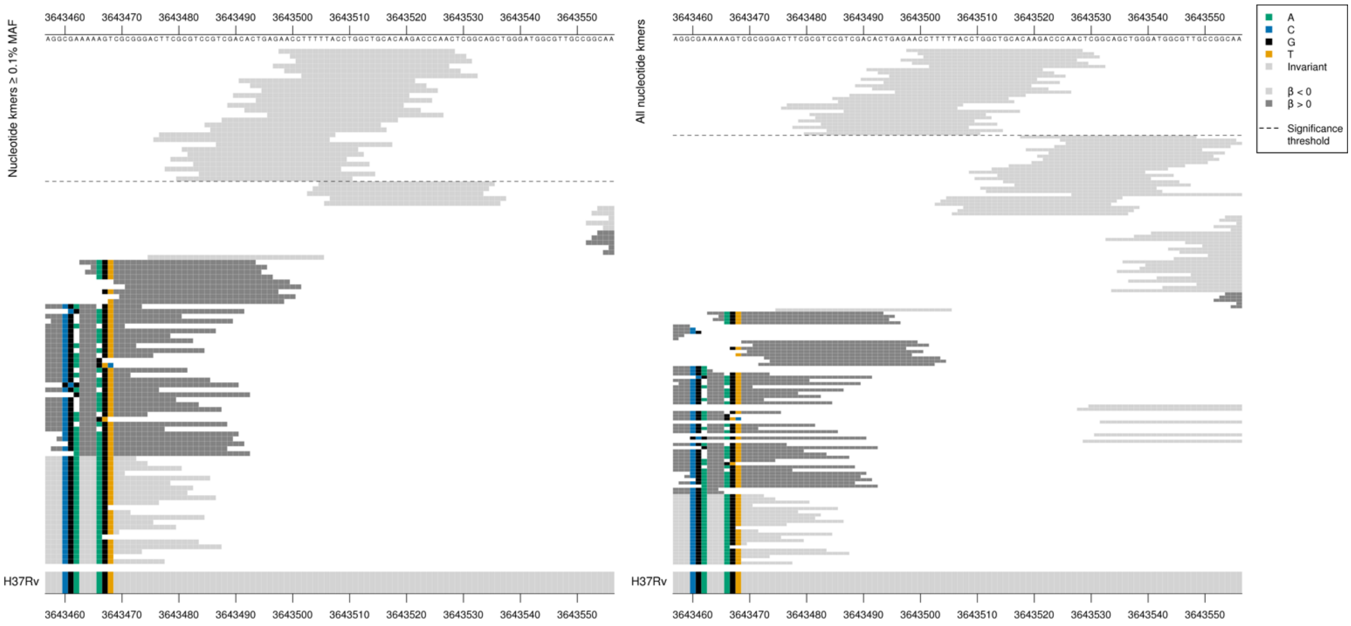

Rv3263 begins at position 3643177

Possibly in LD with other regions, or capturing very rare variation not analysed here.

| Kmer | $-\log_{10} p$ | β | MAC | MAF | Ps | Variants |
|---|---|---|---|---|---|---|
| ENLFTWLHKTQ | 22.02 | -3.00 | 10 | 0.13 | 107 | WT |

### Ethambutol (EMB)

#### *embB*

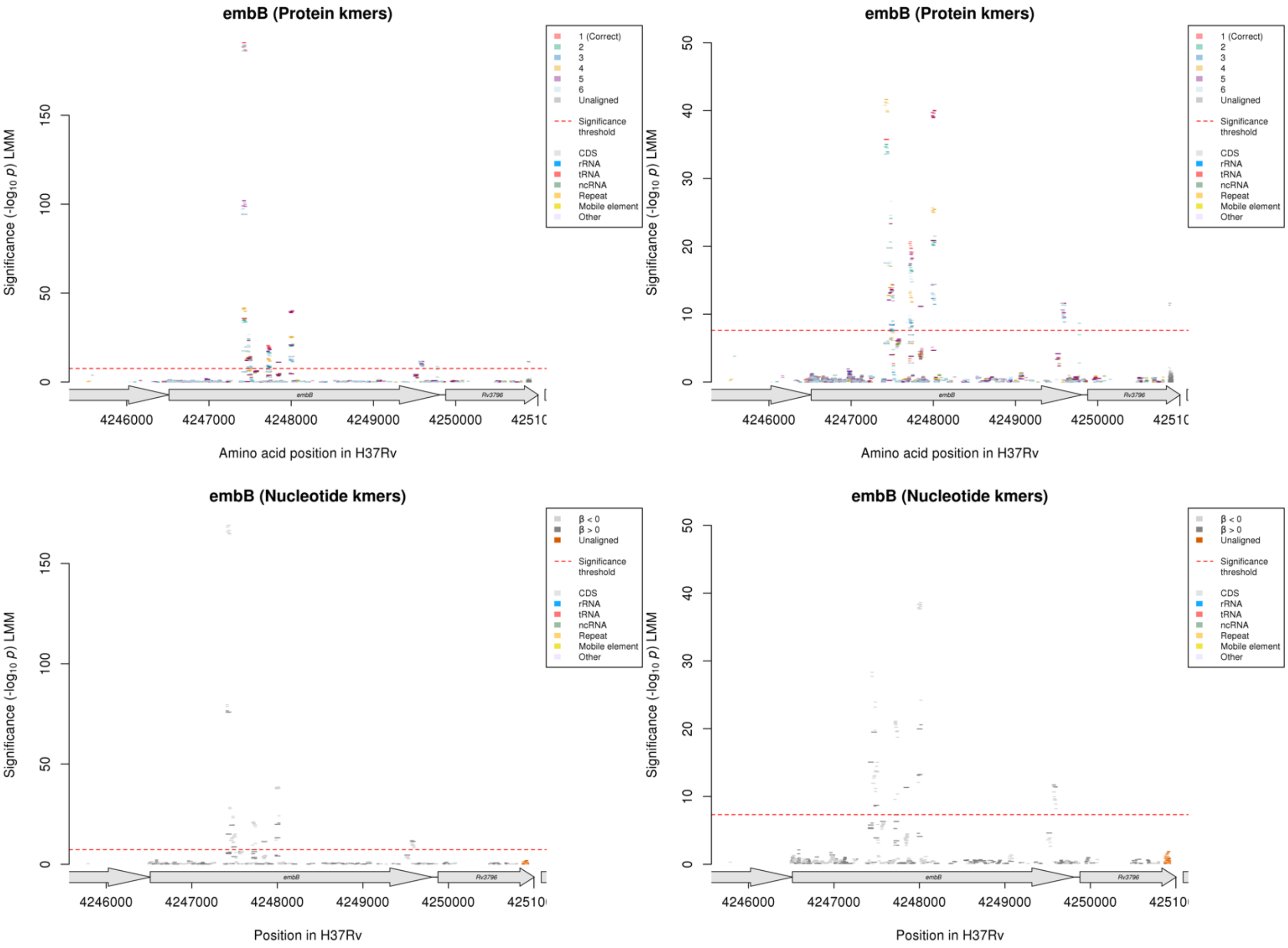

Protein kmers:

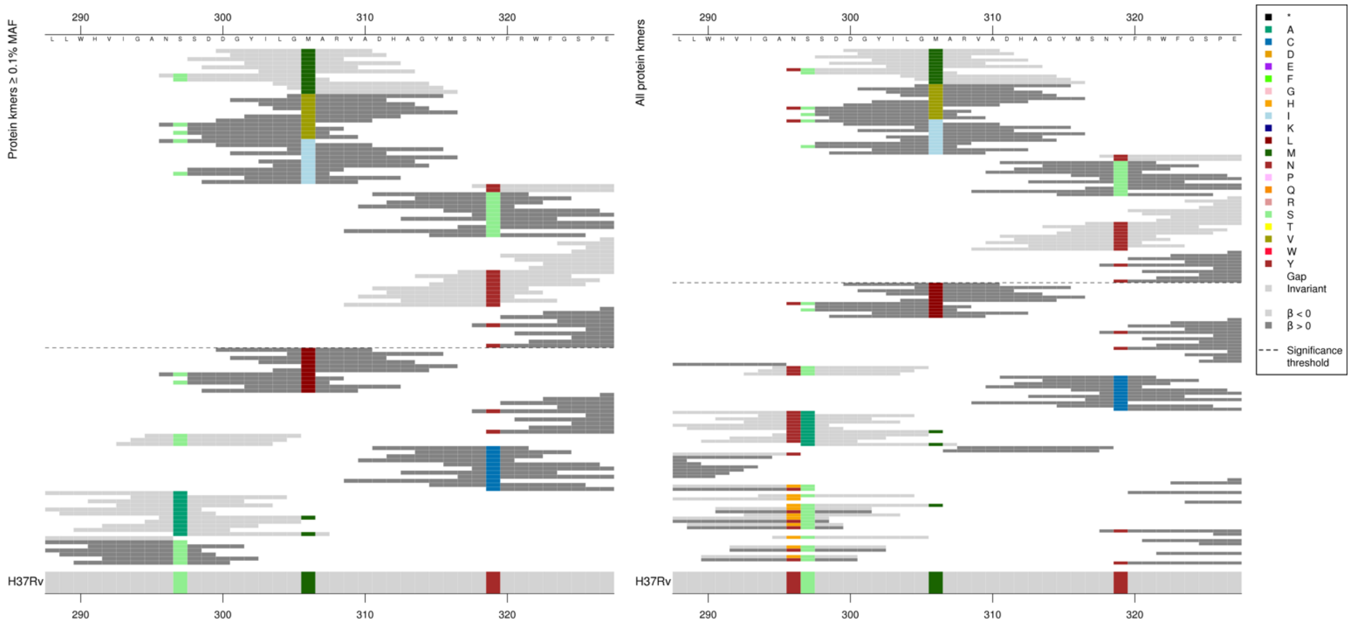

Codon 306

Low MIC-associated kmers = 306M

High MIC-associated kmers = 306V/I

Kmers below significance threshold = 306L

| Kmer | $-\log_{10} p$ | β | MAC | MAF | Ps | Variants |
|---|---|---|---|---|---|---|
| DGYILGMARVA | 190.76 | -1.54 | 1454 | 20.53 | 300 | 306M |
| GVARVADHAGY | 94.33 | 1.34 | 795 | 11.23 | 305 | 306V |
| NSSDDGYILGI | 35.78 | 0.83 | 630 | 8.90 | 296 | 306I |

Protein kmers:

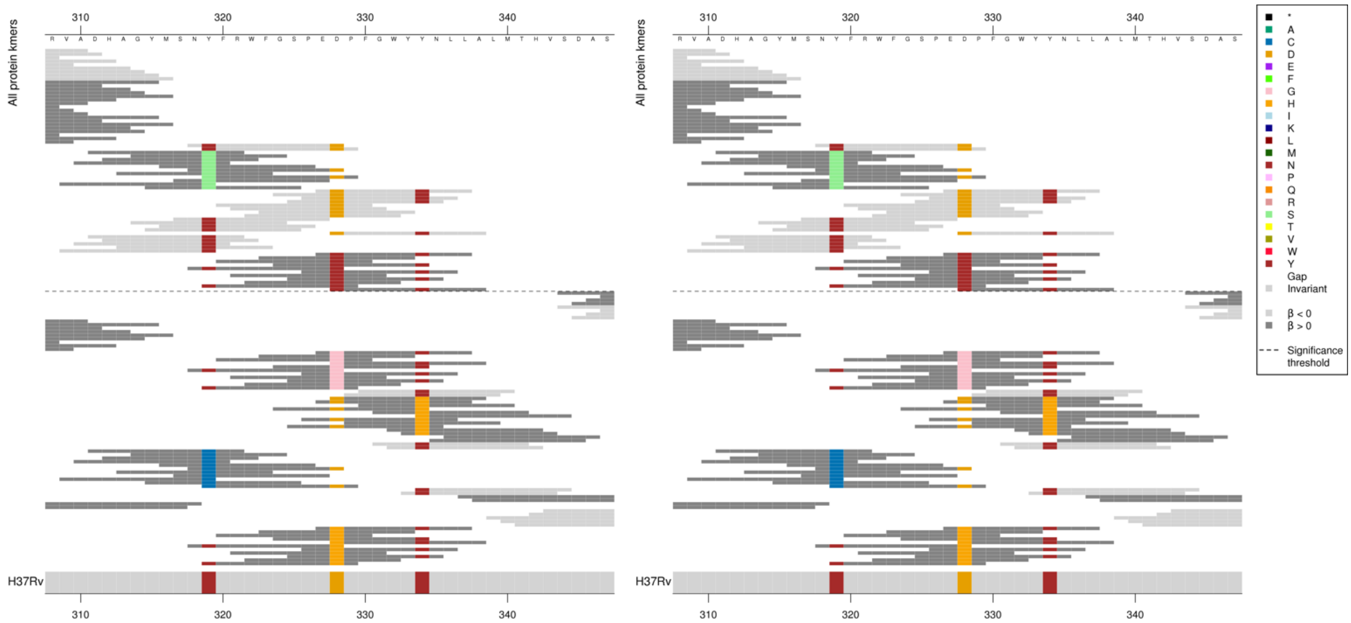

Codon 319
Low MIC-associated kmers = 319Y
High MIC-associated kmers = 319S
Kmers below significance threshold = 319C

Codon 328
Low MIC-associated kmers = 328D
High MIC-associated kmers = 328Y
Kmers below significance threshold = 328G/H

Codon 334
Low MIC-associated kmers = 334Y
Kmers below significance threshold = 334H

| Kmer | $-\log_{10} p$ | β | MAC | MAF | Ps | Variants |
|---|---|---|---|---|---|---|
| NYFRWFGSPED | 23.33 | -1.28 | 156 | 2.20 | 318 | 319Y, 328D |
| DHAGYMSNSFR | 19.77 | 1.88 | 117 | 1.65 | 311 | 319S |
| EDPFGWYYNLL | 14.34 | -1.43 | 33 | 0.47 | 327 | 328D, 334Y |
| FRWFGSPEDPF | 13.70 | -1.45 | 28 | 0.40 | 320 | 328D |
| SNYFRWFGSPE | 13.21 | -1.20 | 133 | 1.88 | 317 | 319Y |
| EYPFGWYYNLL | 8.51 | 1.96 | 9 | 0.13 | 327 | 328Y |

Protein kmers:

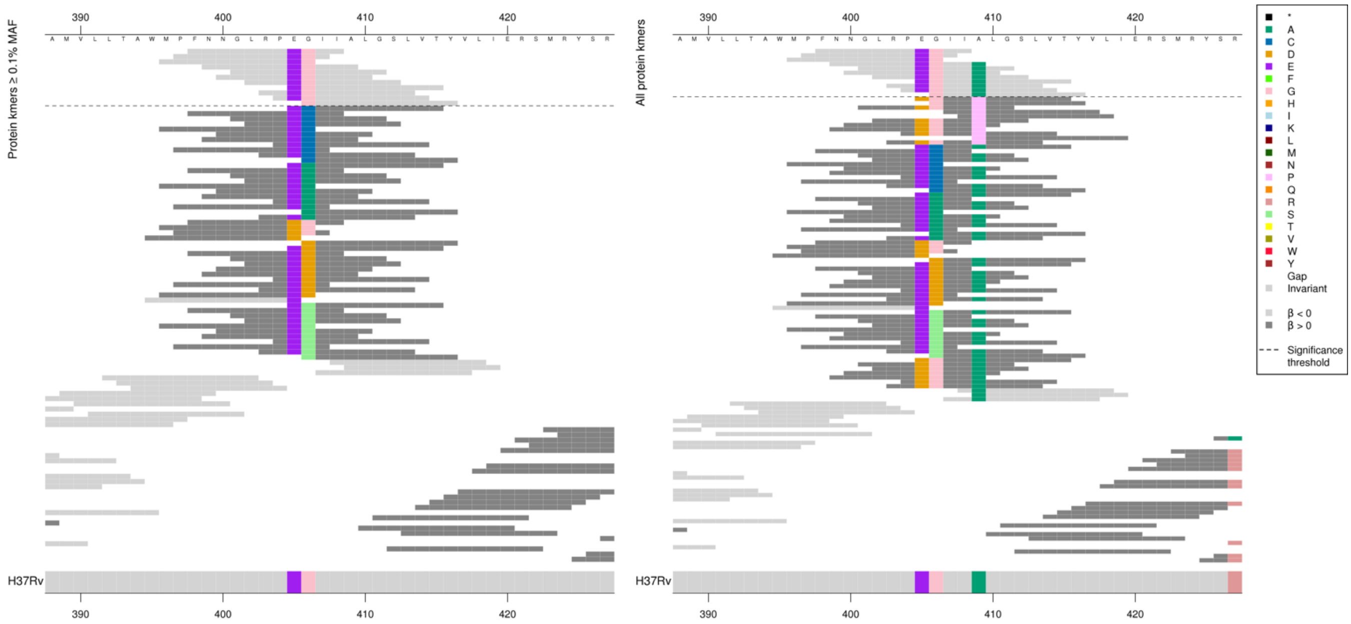

The only position covered by all significant kmers was codon 406, but the kmers have different *p*-values, so were likely effected by the other substitutions.

Codon 405: Kmers below significance threshold = E/D
Codon 406: Kmers below significance threshold = G/C/A/D/S
Codon 409: Kmers below significance threshold = P/A

| Kmer | $-\log_{10} p$ | β | MAC | MAF | Ps | Variants |
|---|---|---|---|---|---|---|
| FNNGLRPEGII | 20.68 | -0.72 | 402 | 5.68 | 398 | 405E, 406G |
| NNGLRPEGIIA | 20.10 | -0.71 | 403 | 5.69 | 399 | 405E, 406G, 409A |
| GIIALGSLVTY | 18.27 | -0.68 | 396 | 5.59 | 406 | 406G, 409A |

Protein kmers:

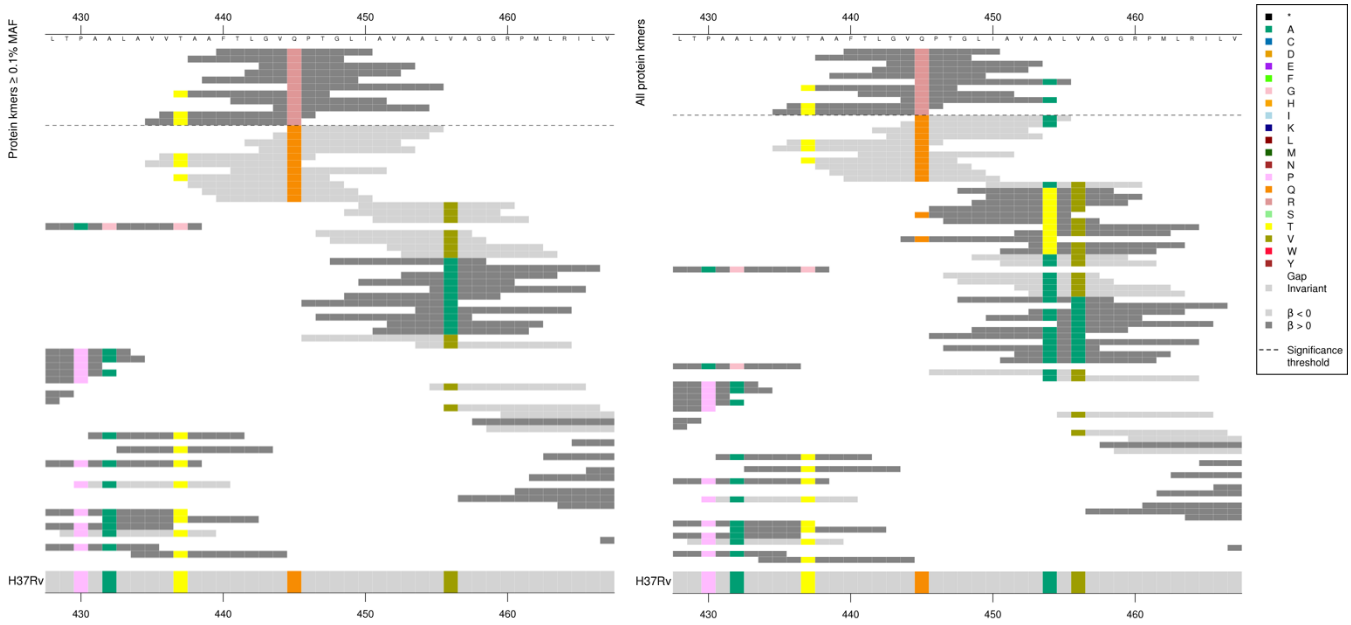

Codon 445: Kmers below significance threshold = Q

| Kmer | $-\log_{10} p$ | β | MAC | MAF | Ps | Variants |
|---|---|---|---|---|---|---|
| FTLGVRPTGLI | 11.14 | 1.68 | 37 | 0.52 | 440 | 445R |

#### Protein kmers:

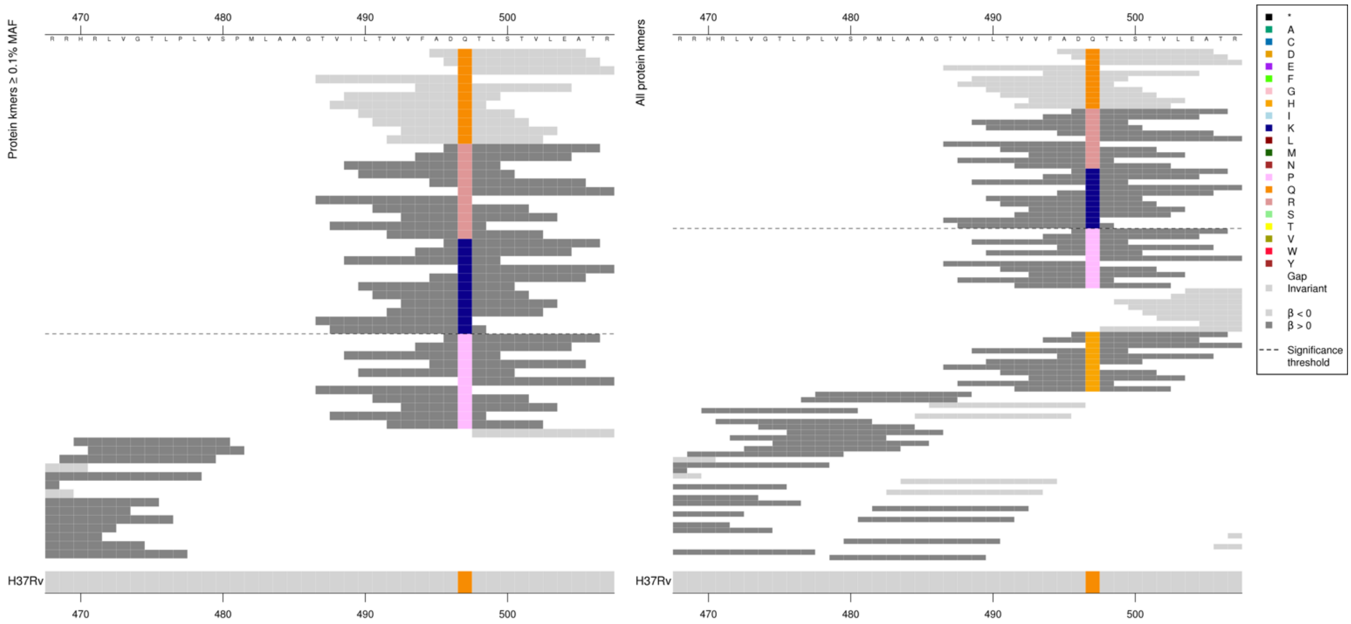

Codon 497: Kmers below significance threshold = P/H

| Kmer | -log$_{10}$ *p* | β | MAC | MAF | Ps | Variants |
|---|---|---|---|---|---|---|
| ADQTLSTVLEA | 39.98 | -1.11 | 300 | 4.24 | 495 | 497Q |
| DRTLSTVLEAT | 20.84 | 0.98 | 227 | 3.21 | 496 | 497R |
| DKTLSTVLEAT | 14.40 | 1.37 | 40 | 0.56 | 496 | 497K |

#### Protein kmers:

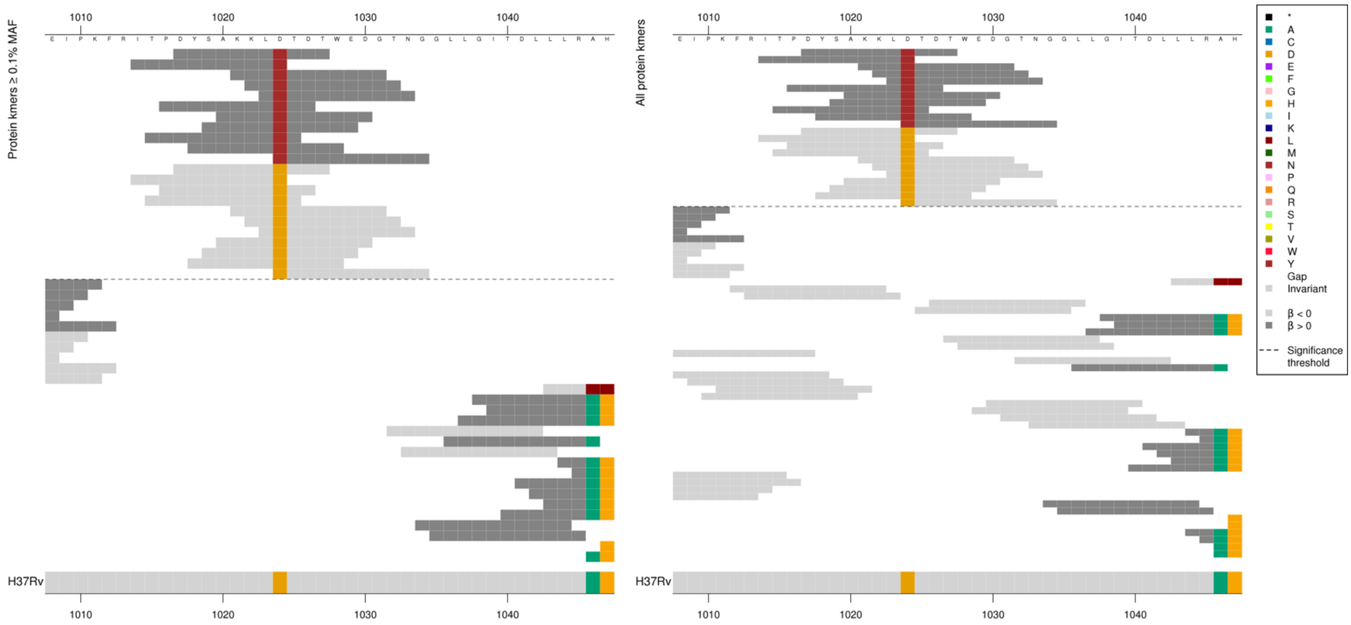

| Kmer | -log$_{10}$ *p* | β | MAC | MAF | Ps | Variants |
|---|---|---|---|---|---|---|
| DYSAKKLNTDT | 11.61 | 1.02 | 56 | 0.79 | 1017 | 1024N |
| DYSAKKLDTDT | 10.66 | -0.99 | 58 | 0.82 | 1017 | 1024D |

### *rpoB*

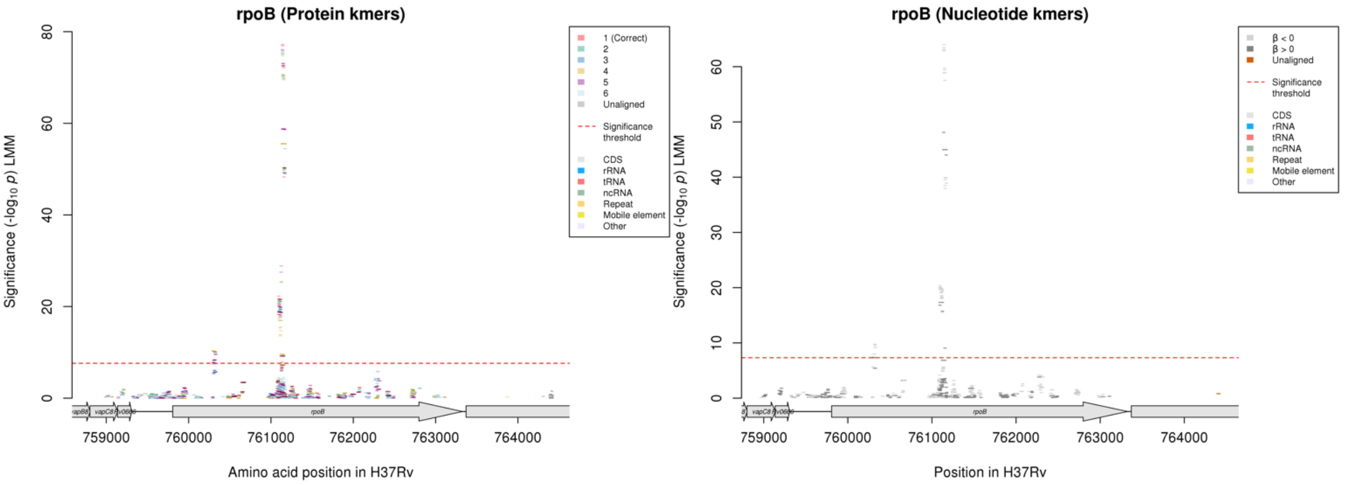

Protein kmers:

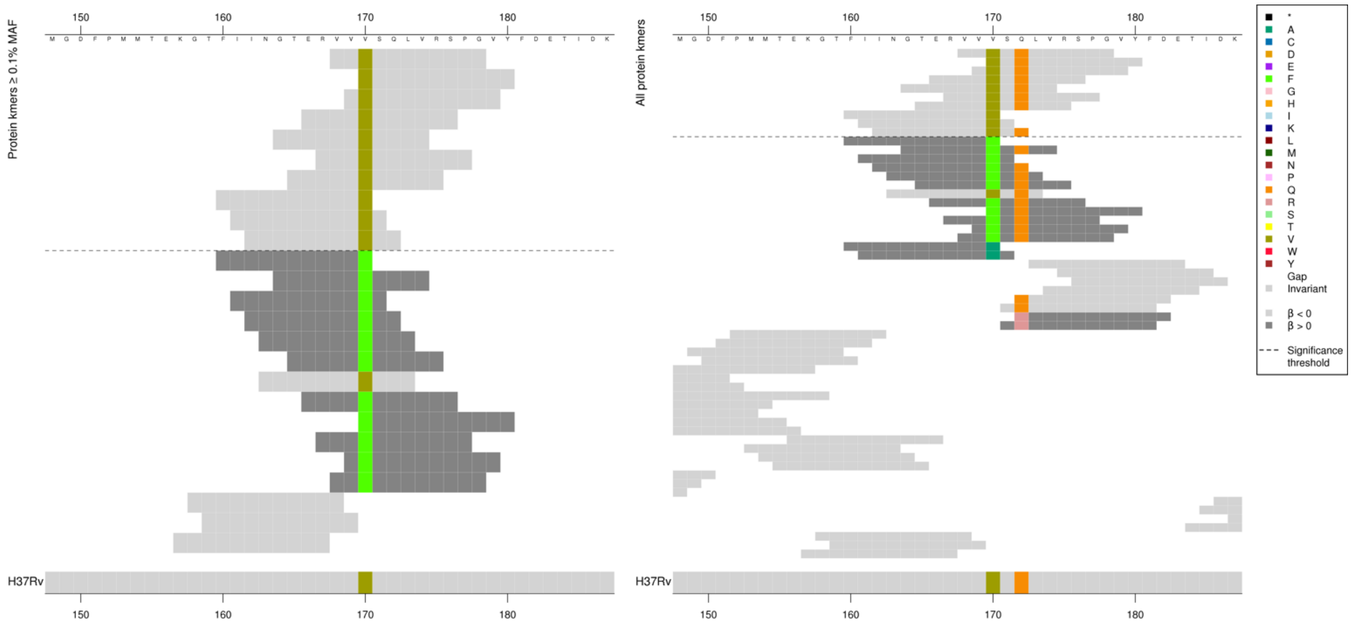

| Kmer | $-\log_{10} p$ | β | MAC | MAF | Ps | Variants |
|---|---|---|---|---|---|---|
| VVVSQLVRSPG | 10.00 | -1.36 | 24 | 0.34 | 168 | 170V, 172Q |
| FIINGTERVVV | 7.74 | -1.08 | 29 | 0.41 | 160 | 170V |

Protein kmers:

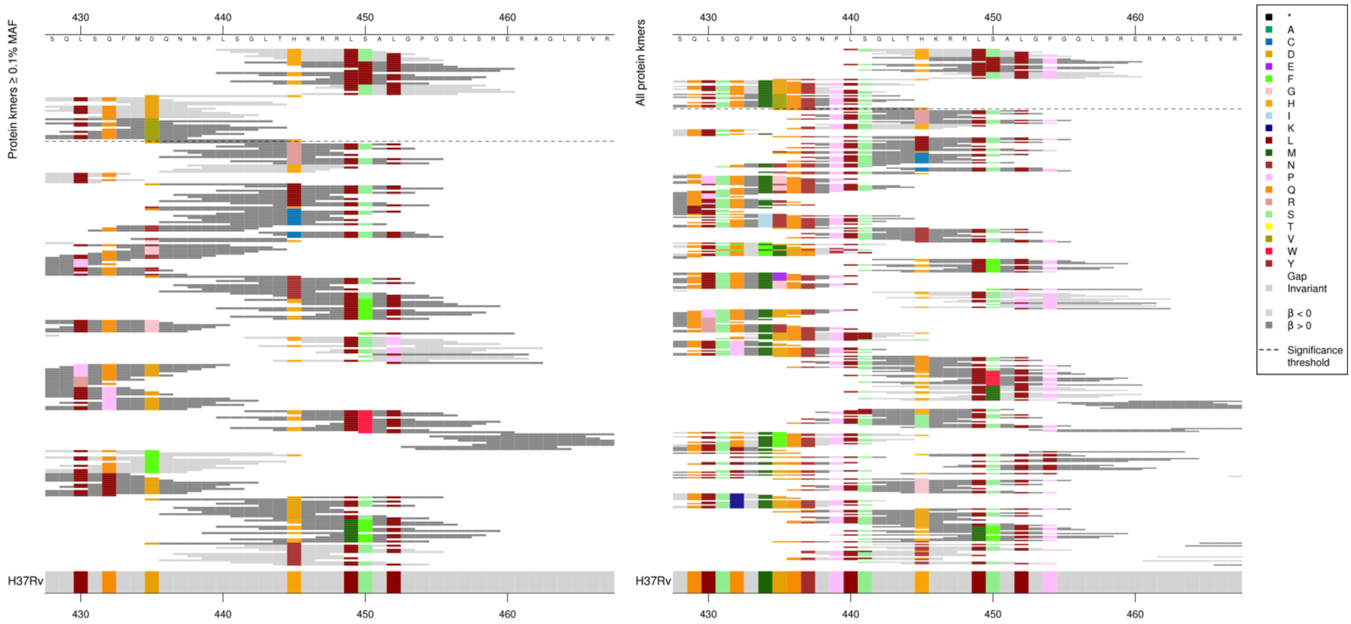

Main variants that were being captured:
Codon 450S/L and codon 435D/V
However presence of the other surrounding alleles affected the significance of the low-MIC associated kmers containing the above variants, just reporting the main variants in the table here.
450L kmers all the same p-value.
435V kmers all the same p-value apart from the last kmer, which is present in one fewer genome.

| Kmer | $-\log_{10} p$ | β | MAC | MAF | Ps | Variants |
|---|---|---|---|---|---|---|
| LSGLTHKRRLS | 77.19 | -0.93 | 2187 | 30.89 | 440 | 450S |
| GLTHKRRLLAL | 55.53 | 0.87 | 1848 | 26.10 | 442 | 450L |
| DQNNPLSGLTH | 28.85 | -0.62 | 804 | 11.35 | 435 | 435D |
| FGTSQLSQFMV | 18.76 | 0.82 | 382 | 5.39 | 425 | 435V |

#### *katG*

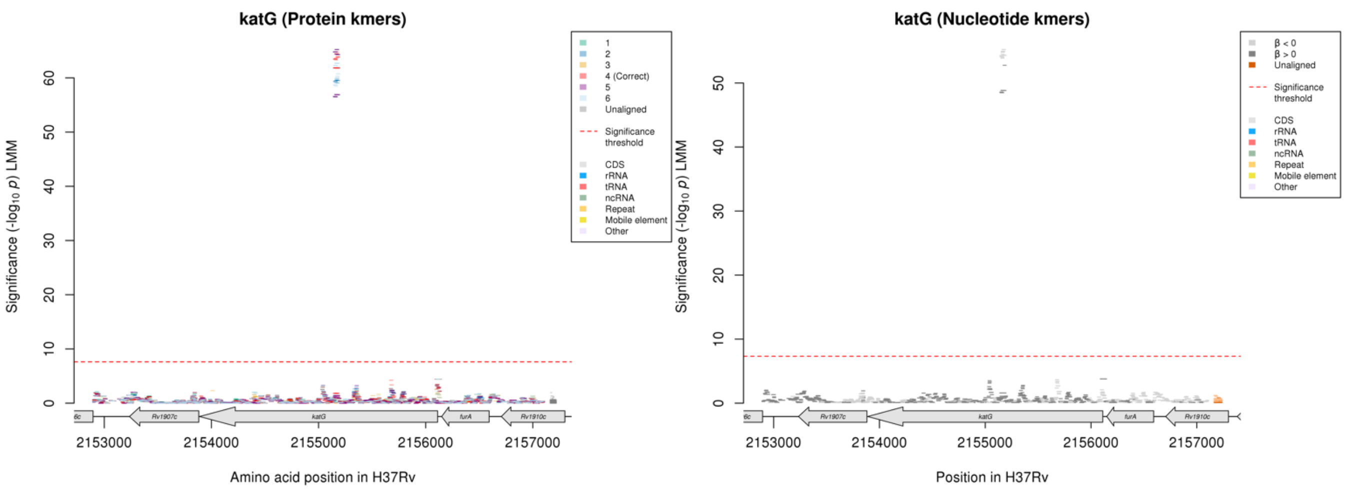

#### Protein kmers:

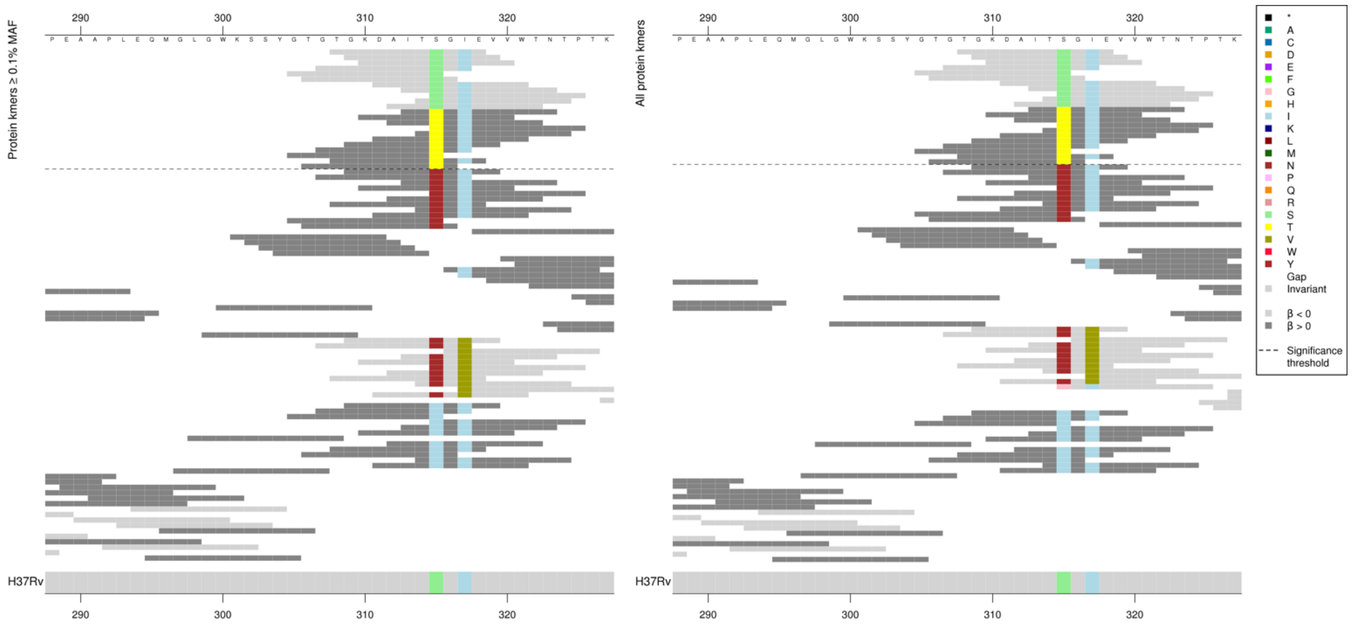

| Kmer | $-\log_{10} p$ | β | MAC | MAF | Ps | Variants |
|---|---|---|---|---|---|---|
| TGKDAITSGIE | 64.77 | -1.00 | 2697 | 38.09 | 308 | 315S |
| ITTGIEVVWTN | 61.83 | 1.00 | 2629 | 37.13 | 313 | 315T |

#### *embA*

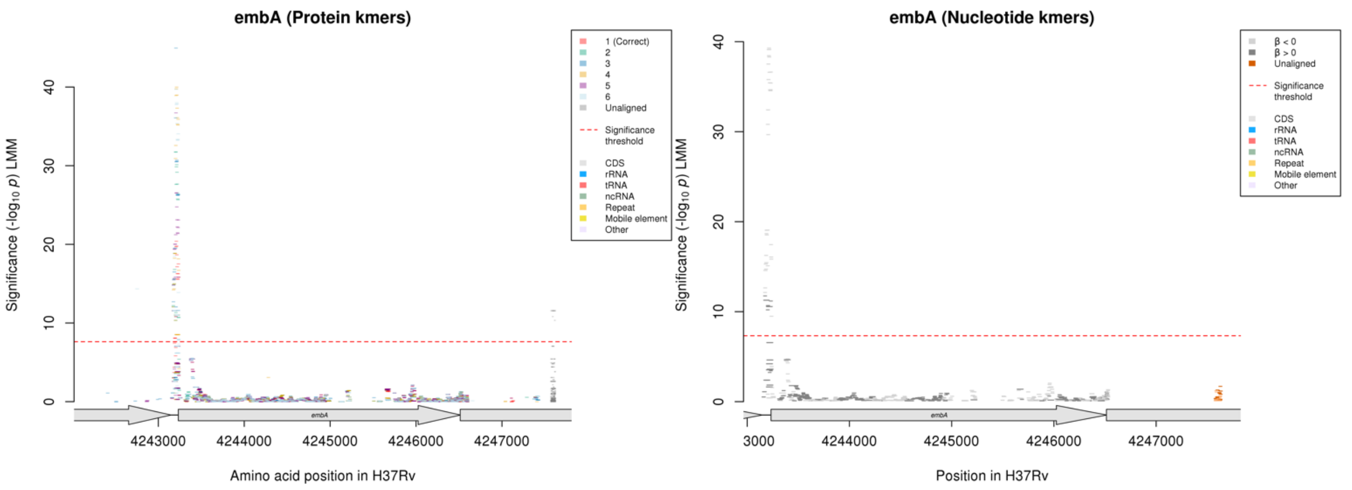

Nucleotide kmers:

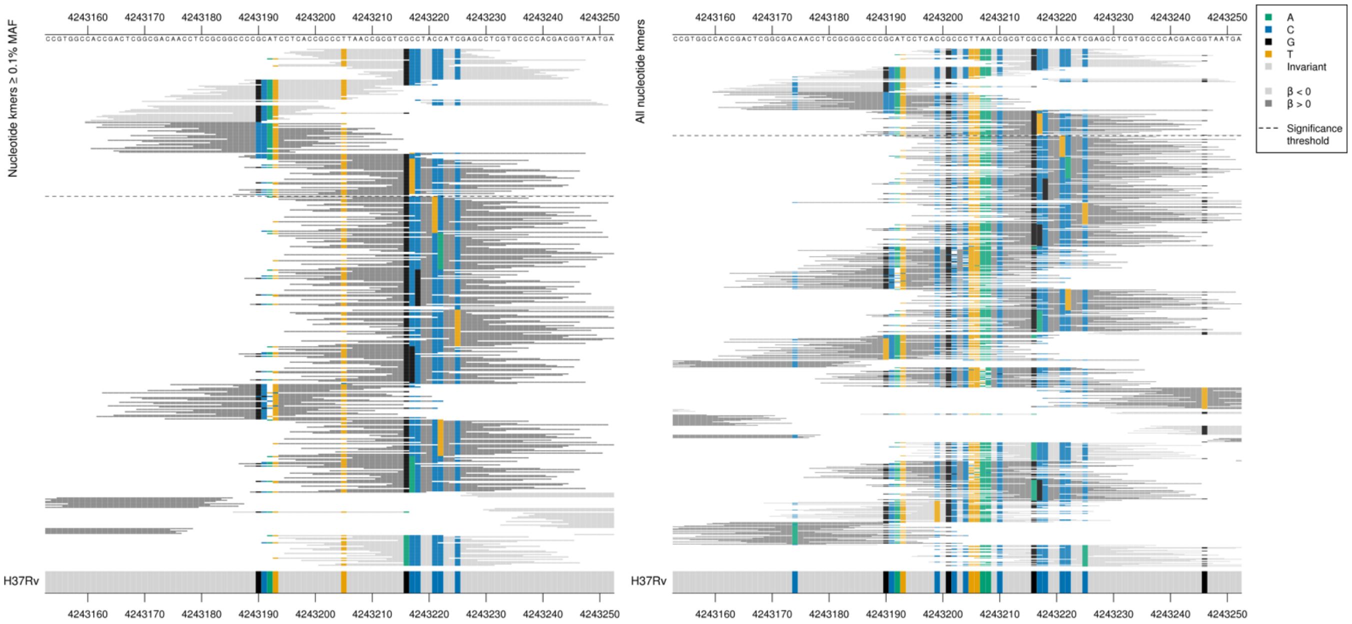

*embA* start position is 4243233.

Main mutations that were captured (where both alleles were significant) were at positions 4243217 and 4243190. Other positions were likely important and impacted the significance of the kmers. Reporting the top kmers that captured the mutations at positions 4243217 and 4243190.

| Kmer | $-\log_{10} p$ | β | MAC | MAF | Ps | Variants |
|---|---|---|---|---|---|---|
| CCGCCCTTAACCGCGTCGCCTACCATCGAGC | 39.29 | -0.95 | 355 | 5.01 | 4243199 | 4243217/-16C [+ others] |
| CCGCATCCTCACCGCCCTTAACCGCGTCGCC | 32.42 | -0.98 | 242 | 3.42 | 4243188 | 4243190/-43G [+ others] |
| CACCGACTCGGCGACAACCTCCGCGGCCCCC | 11.75 | 1.37 | 35 | 0.49 | 4243160 | 4243190/-43C [+ others] |
| TCTACCATCGAGCCTCGTGCCCCACGACGGT | 11.18 | 0.94 | 88 | 1.24 | 4243217 | 4243217/-16T [+ others] |

Two mutations captured at positions:
4243217:
Low MIC-associated kmers: C
High MIC-associated kmers: T
Mutation in possible promoter region, position -16 (4243217) plus others

4243190:
Low MIC-associated kmers: G
High MIC-associated kmers: C
Mutation in possible promoter region, position -43 (4243190) plus others

4243192:
Low MIC: A
High MIC: deletion
A deletion in possible promoter region, position -41 (4243192) in not significant kmers

#### *pncA*

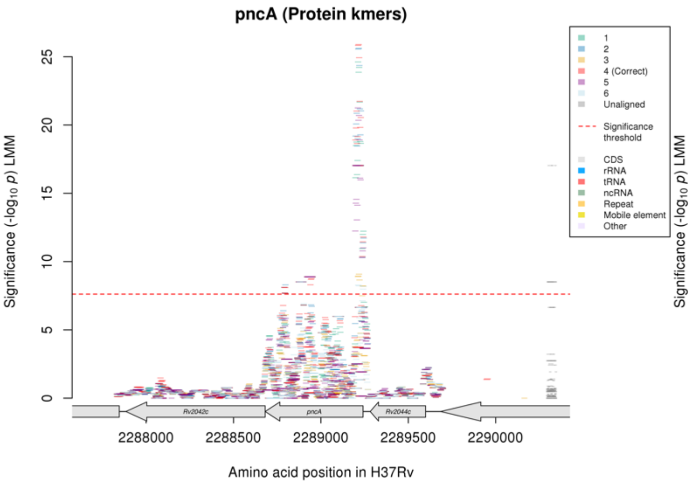

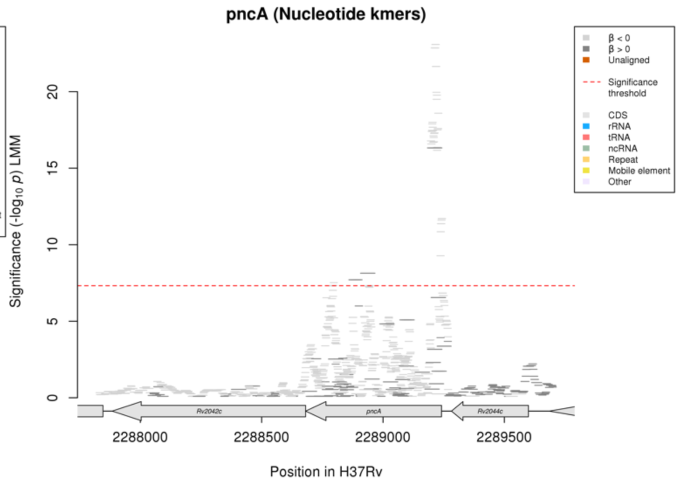

Protein kmers:

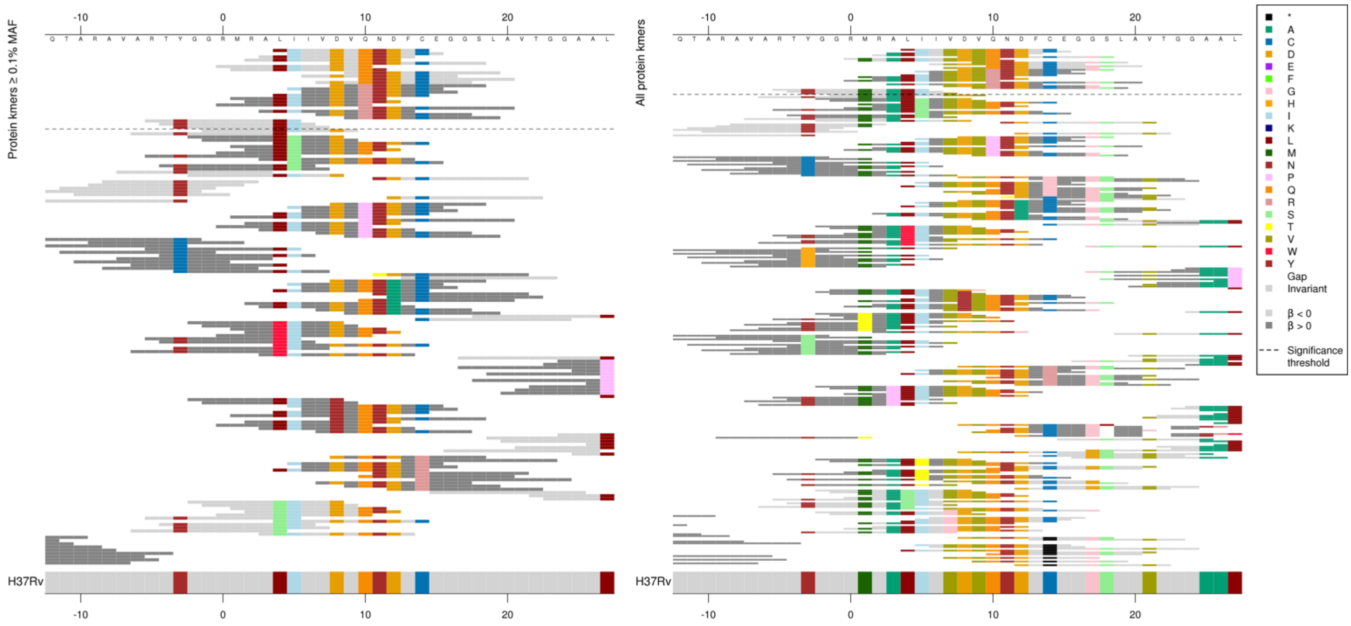

The same region for nucleotide kmers:

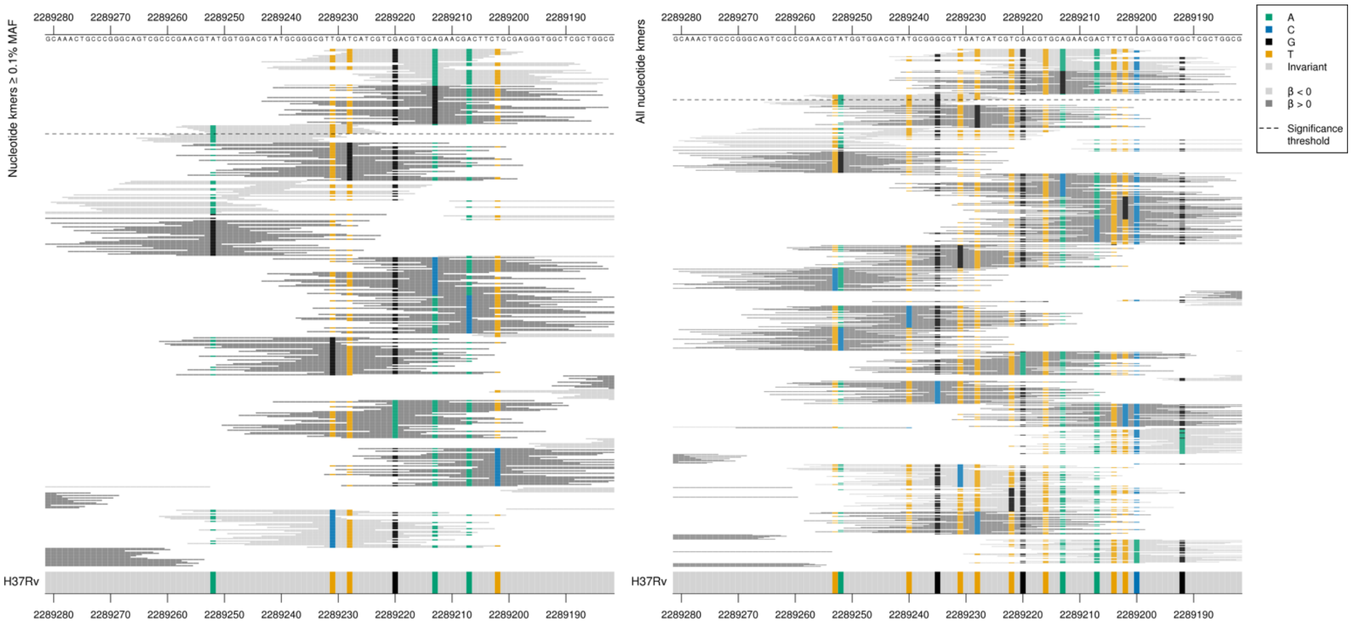

*pncA* begins at position 2289241 on the reverse strand.

| Kmer | -log$_{10}$ *p* | β | MAC | MAF | Ps | Variants |
|---|---|---|---|---|---|---|
| LIIVDVQNDFC | 25.89 | -0.80 | 370 | 5.23 | 4 | 10Q |
| DVRNDFCEGGS | 17.03 | 1.64 | 124 | 1.75 | 8 | 10R |
| RTYGGRMRALI | 11.80 | -0.65 | 196 | 2.77 | -5 | 2289252/-11 A (nucleotide) and 4L, 5I |

Protein kmers:

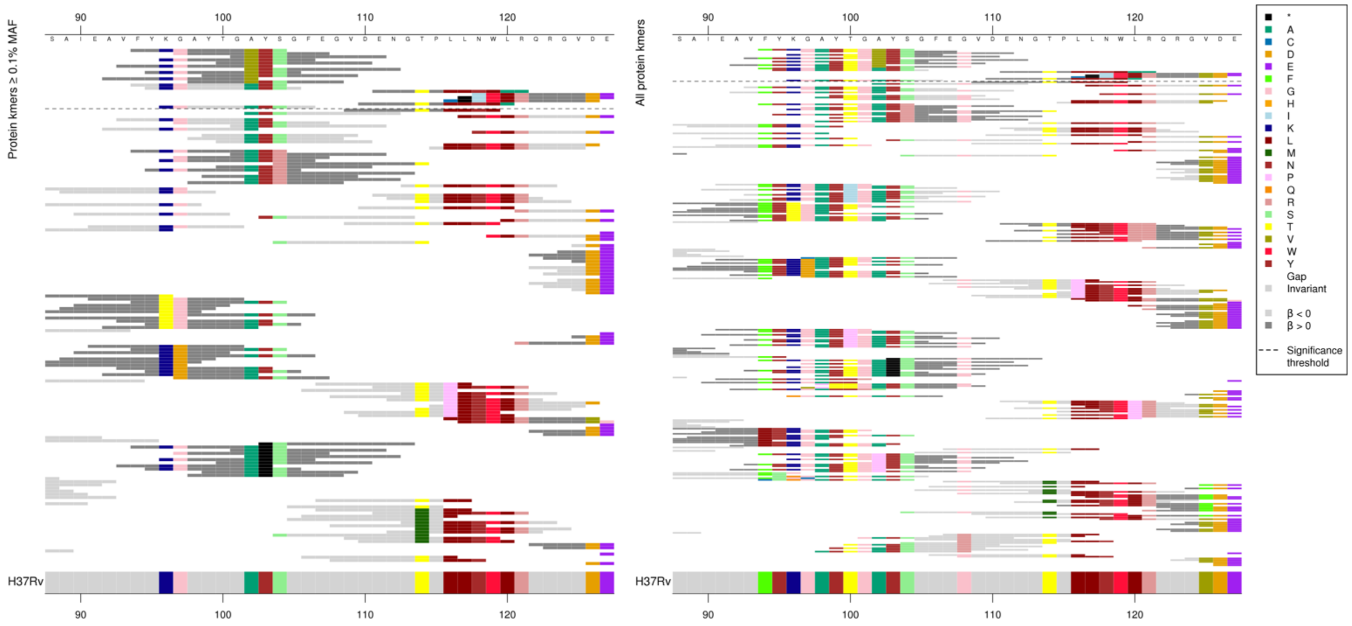

High MIC-associated kmers around codon 102 all only covered codon 102, and they were all the same *p*-value, so the other alleles don’t appear to be influencing the significance. This isn’t the case for the low MIC-associated kmers which had different *p*-values.

| Kmer | -log$_{10}$ *p* | β | MAC | MAF | Ps | Variants |
|---|---|---|---|---|---|---|
| FYKGAYTGVYS | 8.88 | 2.35 | 10 | 0.14 | 94 | 102V |
| FYKGAYTGAYS | 8.74 | -0.56 | 181 | 2.56 | 94 | 102A [+ others] |

Protein kmers:

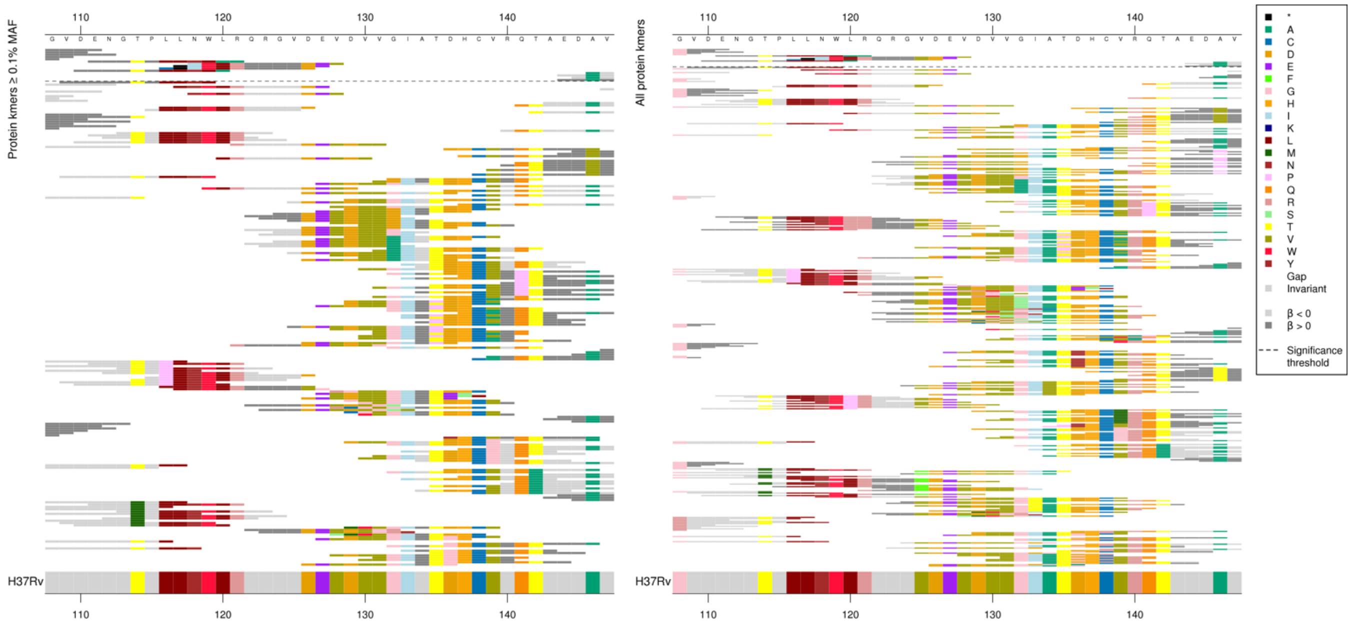

The same region for nucleotide kmers:

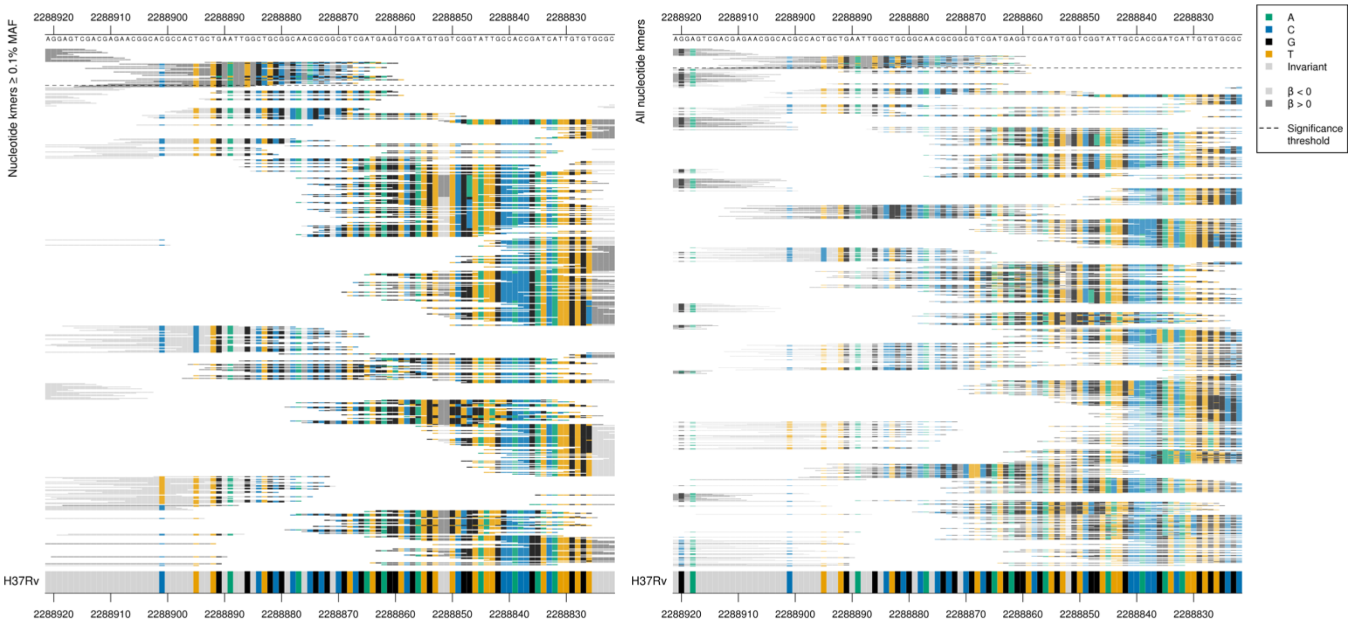

High MIC-associated kmers around codon 120 contained insertions compared to the reference, a T between reference nucleotide positions 2288889-2288886/353-356 (codons 118-119).

| Kmer | $-\log_{10} p$ | β | MAC | MAF | Ps | Variants |
|---|---|---|---|---|---|---|
| CTGAATTTGGCTGCGGCAACGCGGCGTCGAT | 7.71 | 2.05 | 10 | 0.14 | 2288893 | T insertion between 2288889-2288886/353-356 |

Protein kmers:

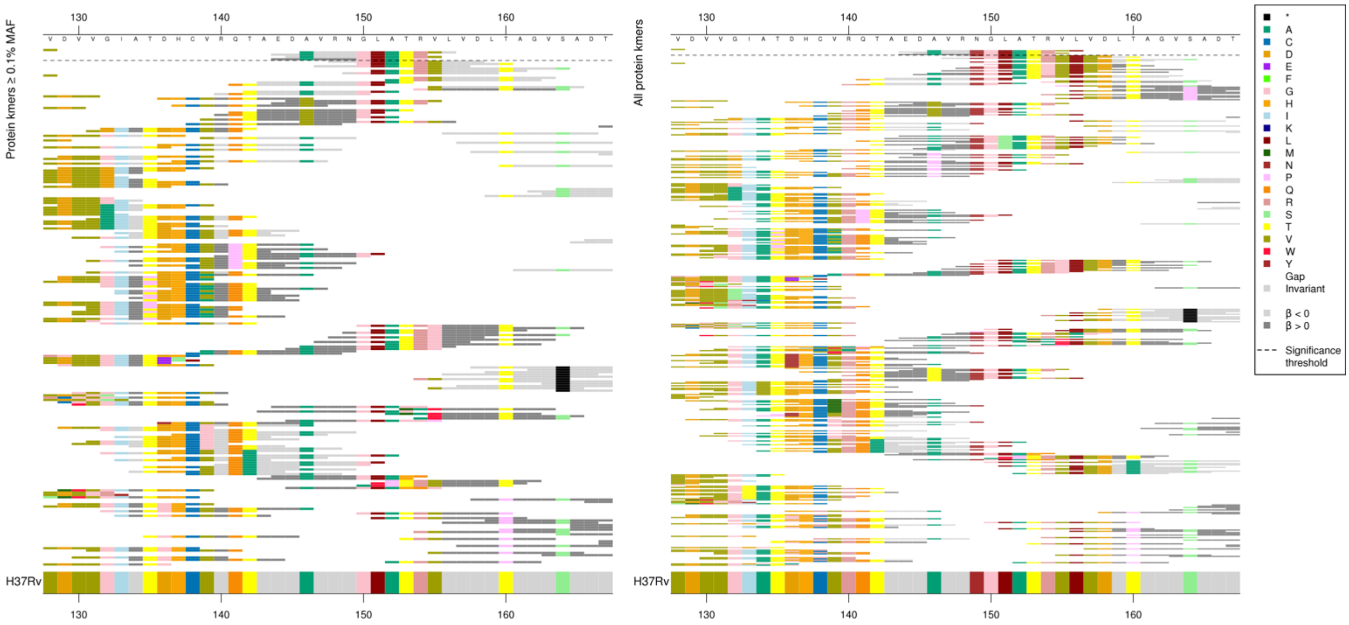

Lots of variants in this area, the three low MIC-associated kmers were all the same as the reference, and all covered positions 146-154 (146A, 154R). The single high MIC-associated kmer was identical to the reference except at position 154 where it was a G. Other positions were possibly also important.

| Kmer | $-\log_{10} p$ | β | MAC | MAF | Ps | Variants |
|---|---|---|---|---|---|---|
| AVRNGLATRVL | 8.11 | -0.62 | 136 | 1.92 | 146 | WT / 146A, 154R |
| EDAVRNGLATG | 7.66 | 1.74 | 11 | 0.16 | 144 | 154G |

#### *gyrA*

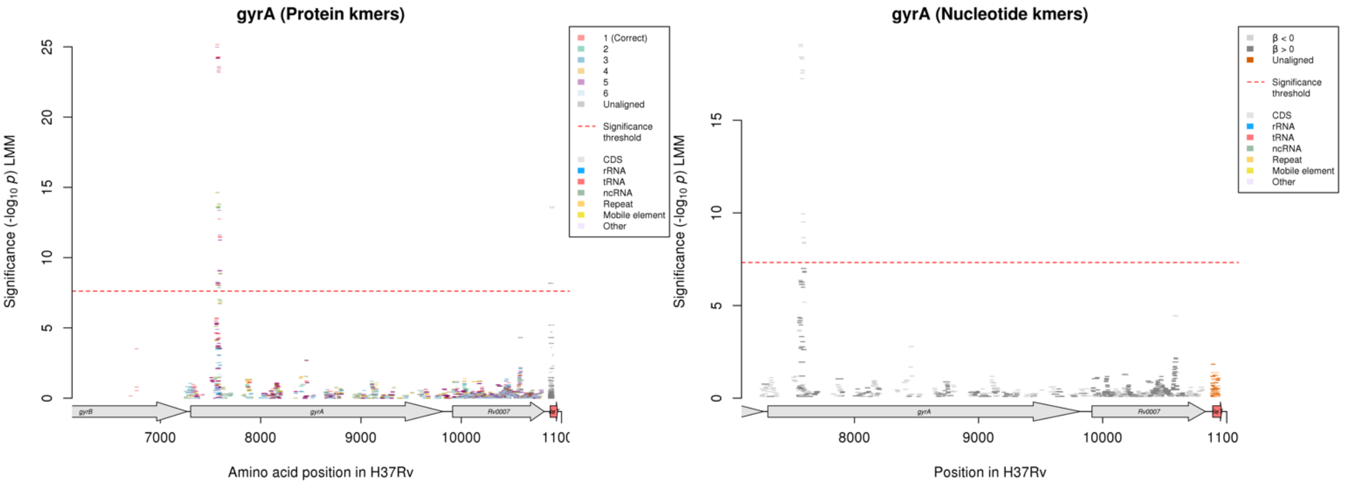

Protein kmers:

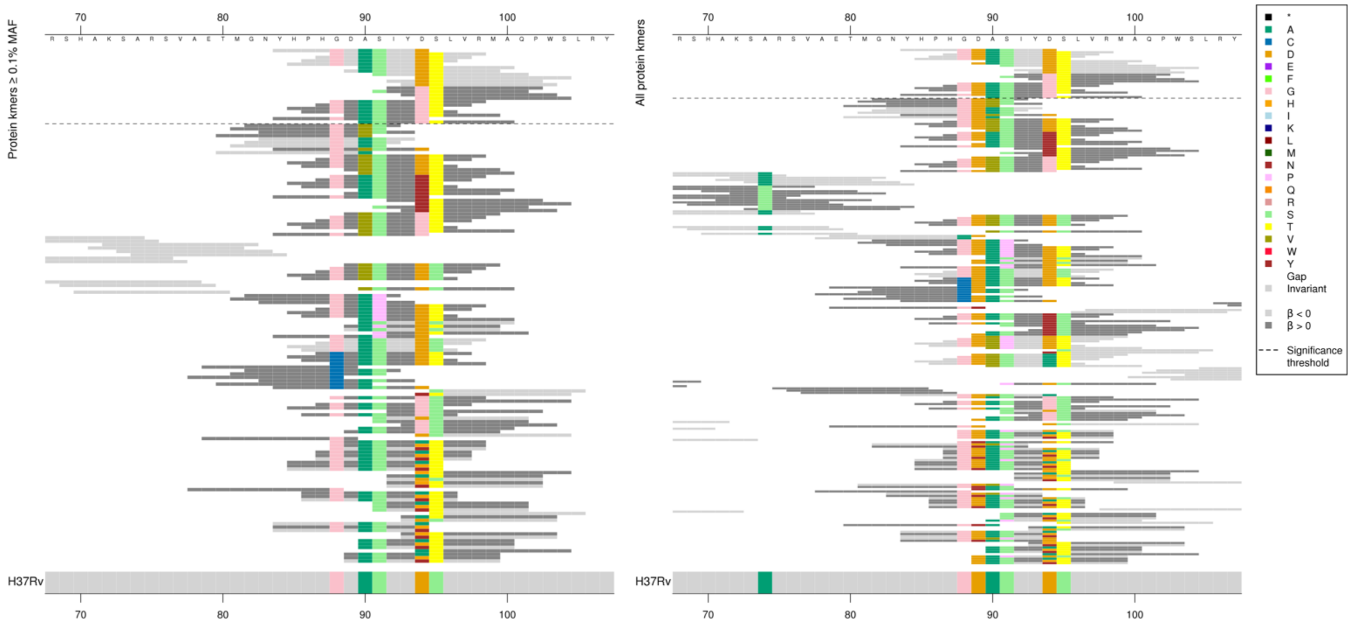

All significant kmers covered just codon 94, however the kmers were different *p*-values dependent on the alleles so showing the top result for each combination of alleles.

| Kmer | $-\log_{10} p$ | β | MAC | MAF | Ps | Variants |
|---|---|---|---|---|---|---|
| YHPHGDASIYD | 25.16 | -0.54 | 1206 | 17.03 | 84 | 88G, 89D, 90A, 91S, 94D |
| GDASIYDTLVR | 24.26 | -0.54 | 1586 | 22.40 | 88 | 88G, 89D, 90A, 91S, 94D, 95T |
| ASIYDTLVRMA | 23.46 | -0.53 | 1564 | 22.09 | 90 | 90A, 91S, 94D, 95T |
| DASIYDTLVRM | 23.19 | -0.53 | 1569 | 22.16 | 89 | 89D, 90A, 91S, 94D, 95T |
| SIYDTLVRMAQ | 12.76 | -0.39 | 1257 | 17.75 | 91 | 91S, 94D, 95T |
| DTLVRMAQPWS | 11.47 | -0.38 | 1198 | 16.92 | 94 | 94D, 95T |
| IYGTLVRMAQP | 9.07 | 0.38 | 540 | 7.63 | 92 | 94G, 95T |
| SIYGTLVRMAQ | 9.07 | 0.38 | 540 | 7.63 | 91 | 91S, 94G, 95T |
| GDASIYGTLVR | 8.17 | 0.36 | 529 | 7.47 | 88 | 88G, 89D, 90A, 91S, 94G, 95T |
| DASIYGTLVRM | 8.17 | 0.36 | 529 | 7.47 | 89 | 89D, 90A, 91S, 94G, 95T |
| YHPHGDASIYG | 8.06 | 0.35 | 541 | 7.64 | 84 | 88G, 89D, 90A, 91S, 94G |
| ASIYGTLVRMA | 8.03 | 0.36 | 531 | 7.50 | 90 | 90A, 91S, 94G, 95T |

Ethambutol (EMB)

#### *rpsL*

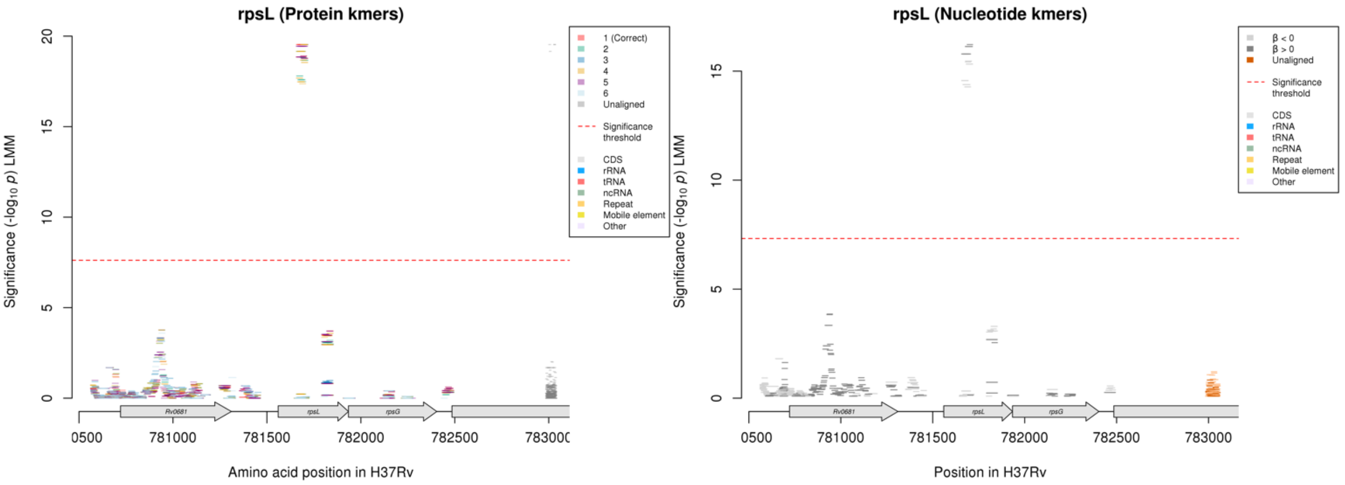

Protein kmers:

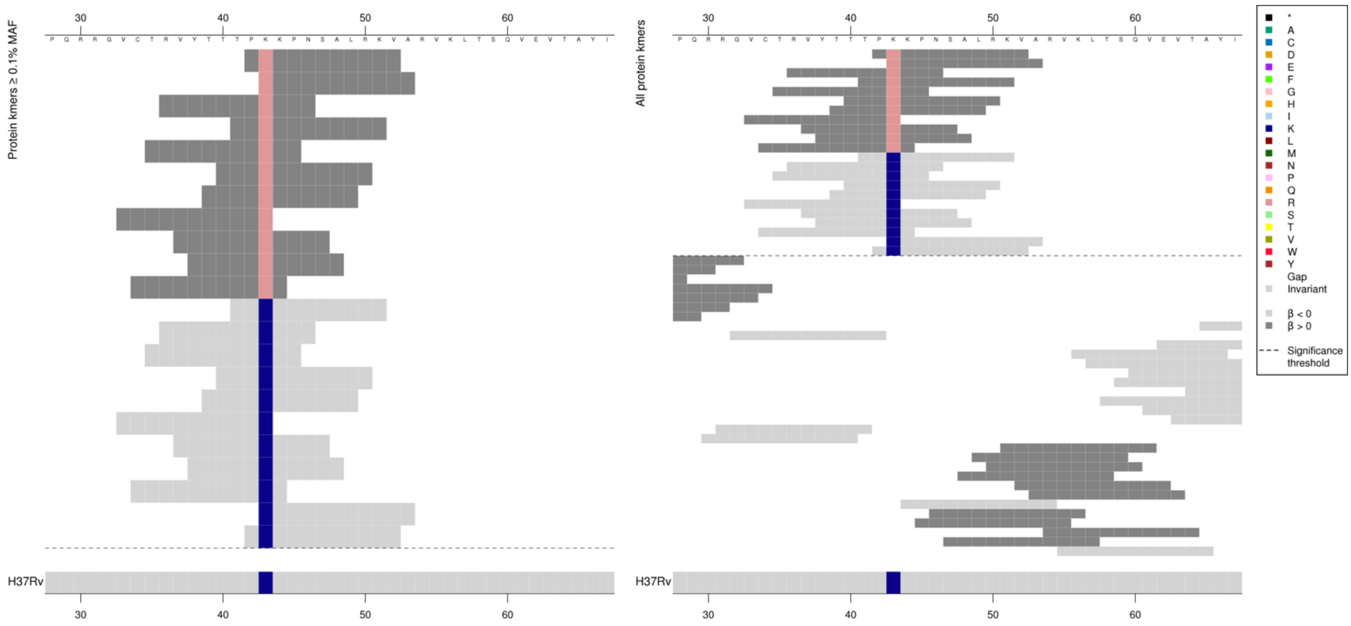

| Kmer | $-\log_{10} p$ | β | MAC | MAF | Ps | Variants |
|---|---|---|---|---|---|---|
| PRKPNSALRKV | 19.53 | 0.63 | 1281 | 18.09 | 42 | 43R |
| TPKKPNSALRK | 18.91 | -0.62 | 1280 | 18.08 | 41 | 43K |

## *Rv1565c*

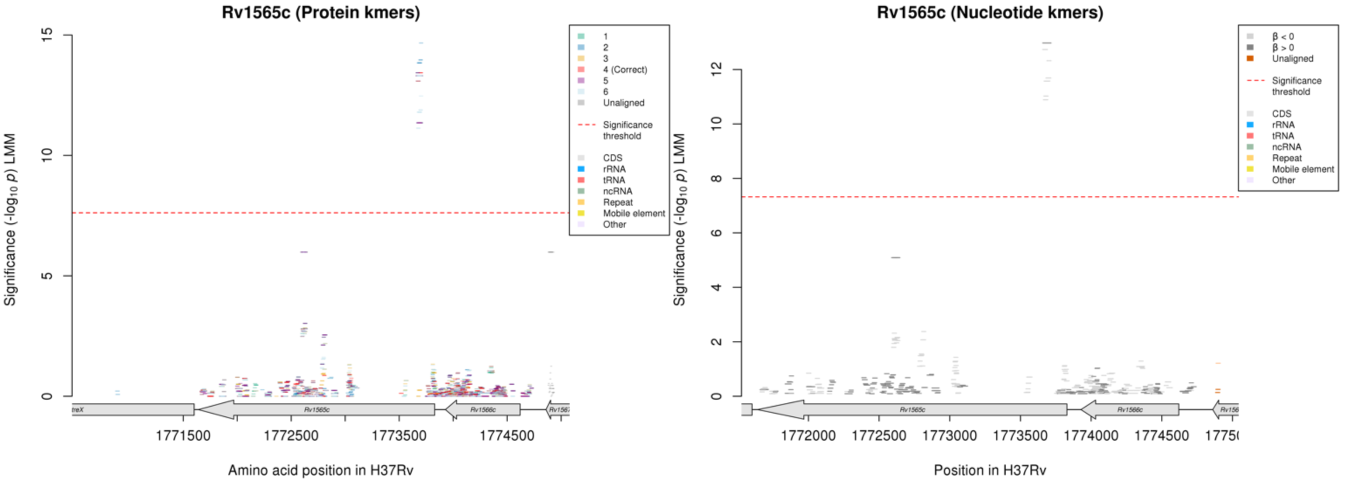

#### Protein kmers:

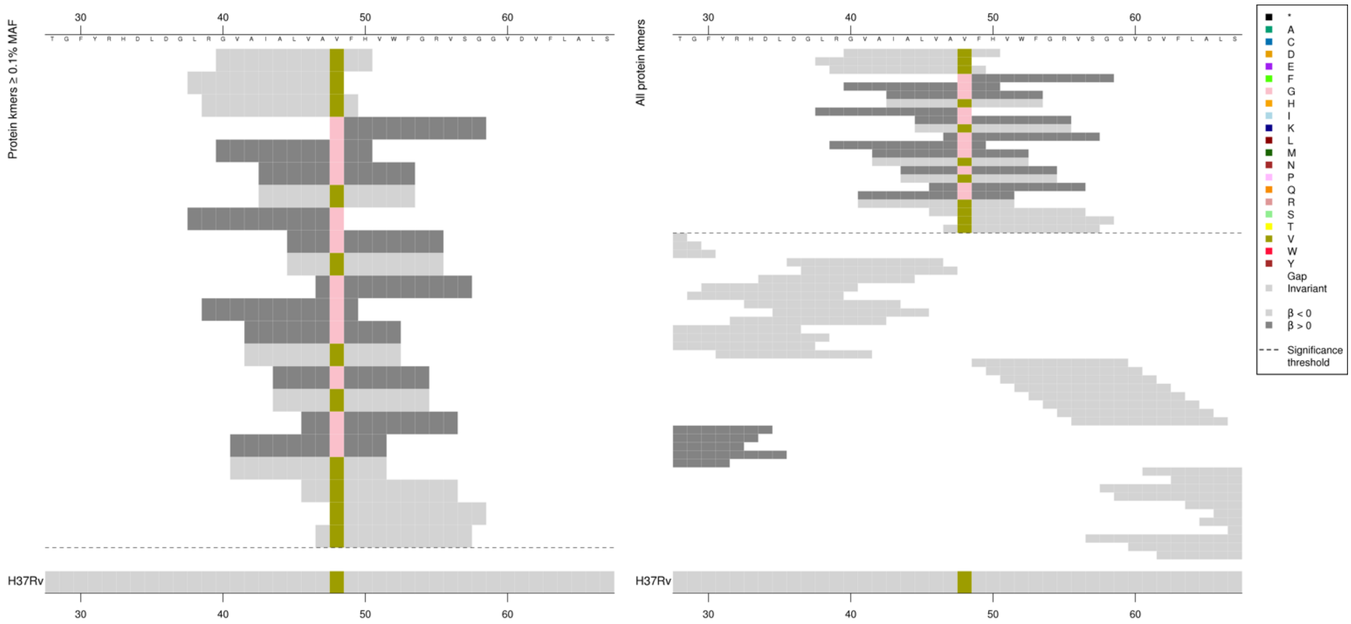

| Kmer | -log₁₀ p | β | MAC | MAF | Ps | Variants |
|---|---|---|---|---|---|---|
| GVAIALVAVFH | 13.43 | -1.63 | 122 | 1.72 | 40 | 48V |
| GFHVWFGRVSG | 13.31 | 1.63 | 120 | 1.69 | 48 | 48G |

## *Rv2478c:Rv2481c* [R]

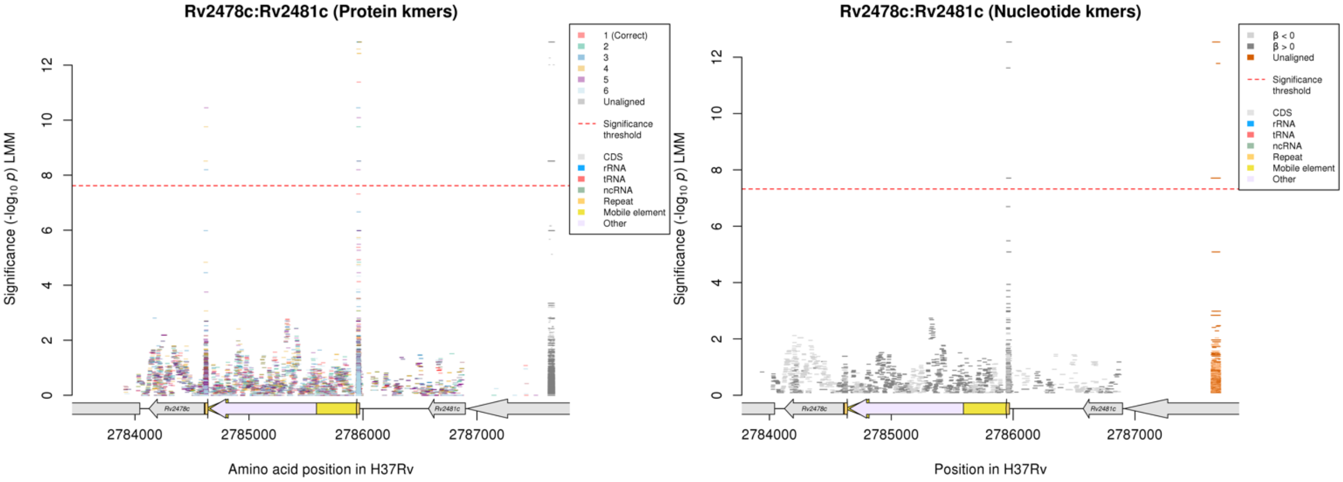

#### Nucleotide kmers:

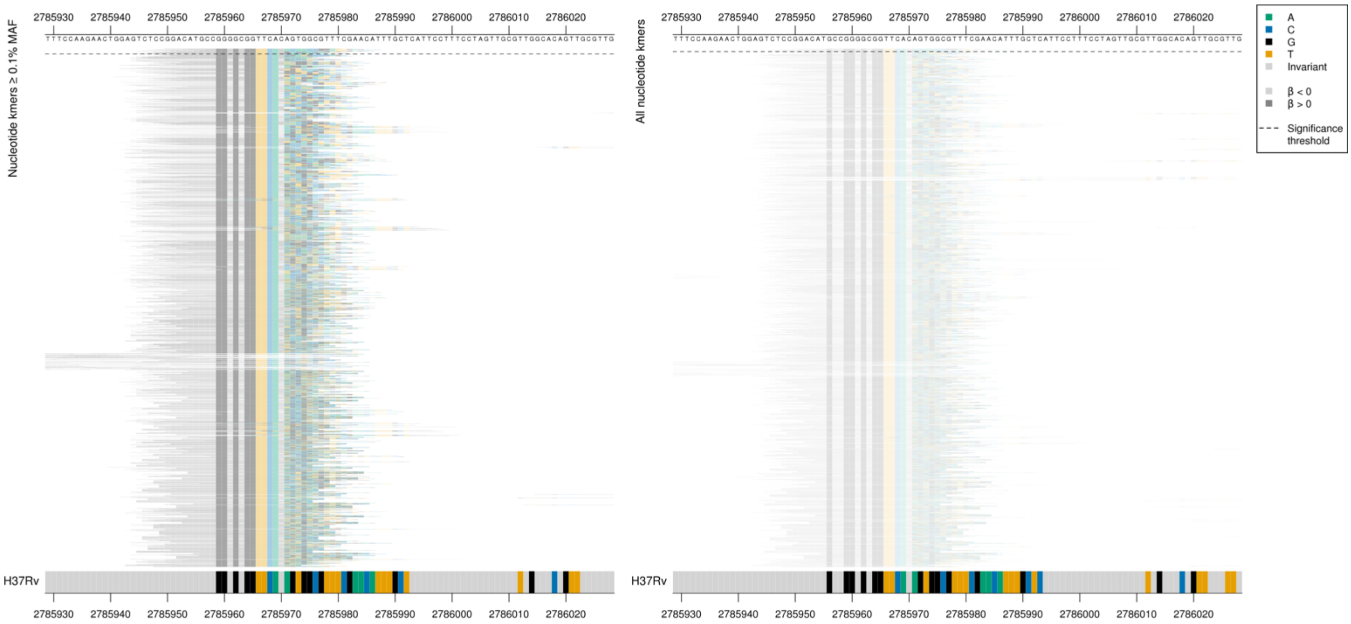

| Kmer | $-\log_{10} p$ | β | MAC | MAF | Ps | Variants |
|---|---|---|---|---|---|---|
| GGACATGCCGGGGCGGTTCACACAACCGGAT | 12.54 | 1.52 | 110 | 1.55 | 2785950 | Repeat region |

## *Rv1752*

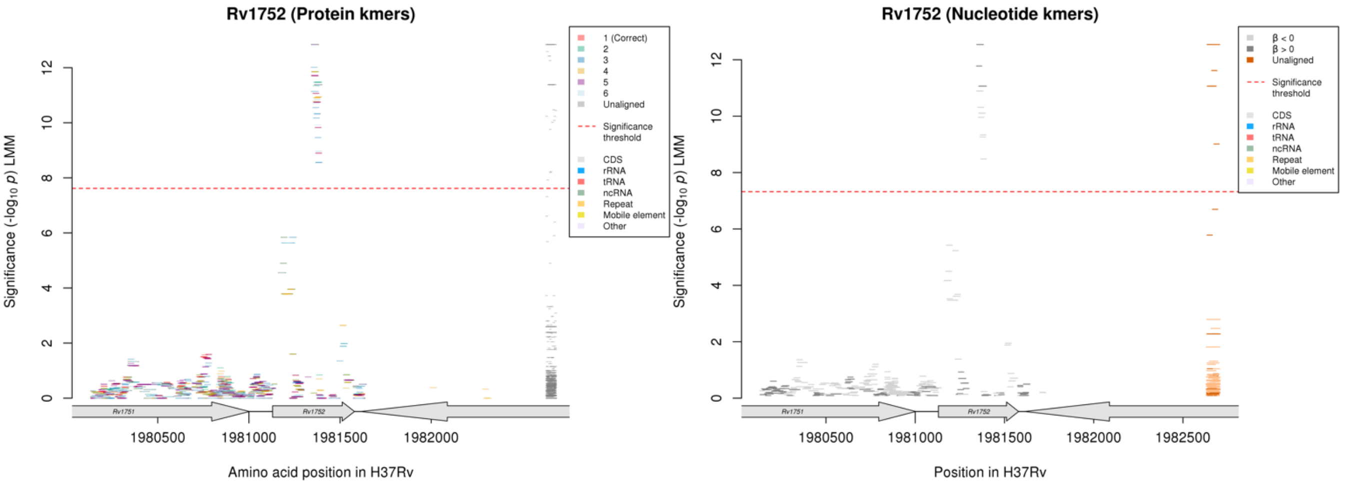

Protein kmers:

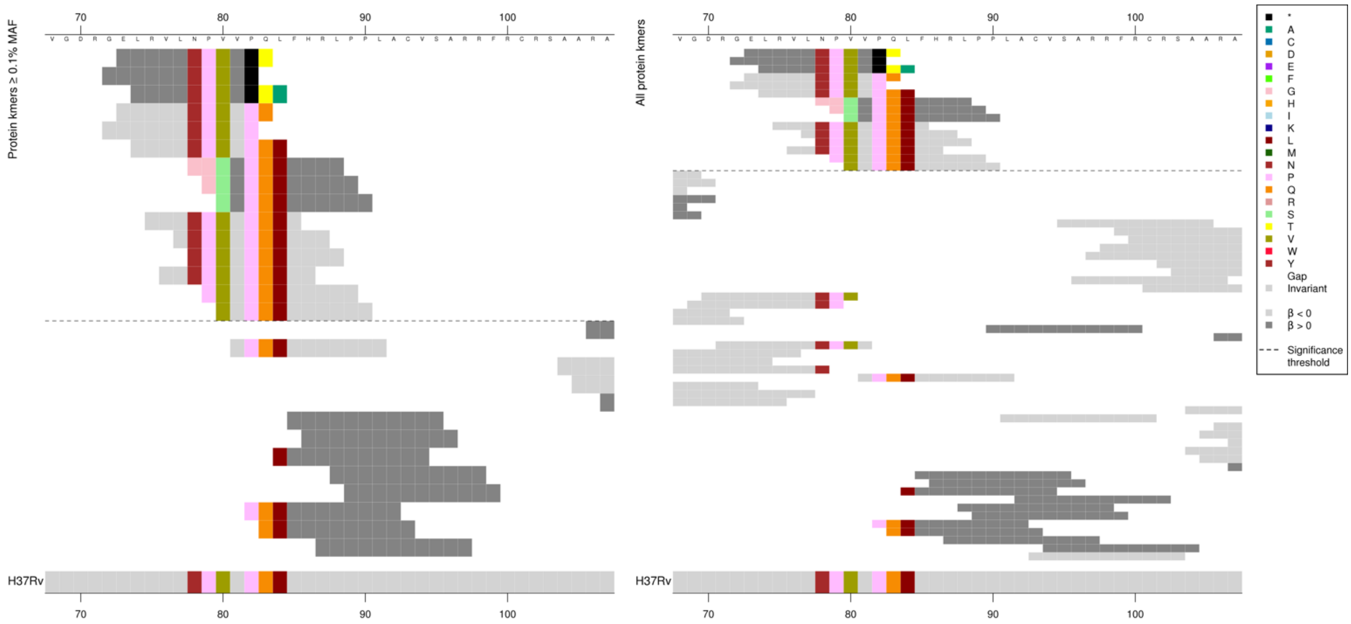

Same region for nucleotide kmers:

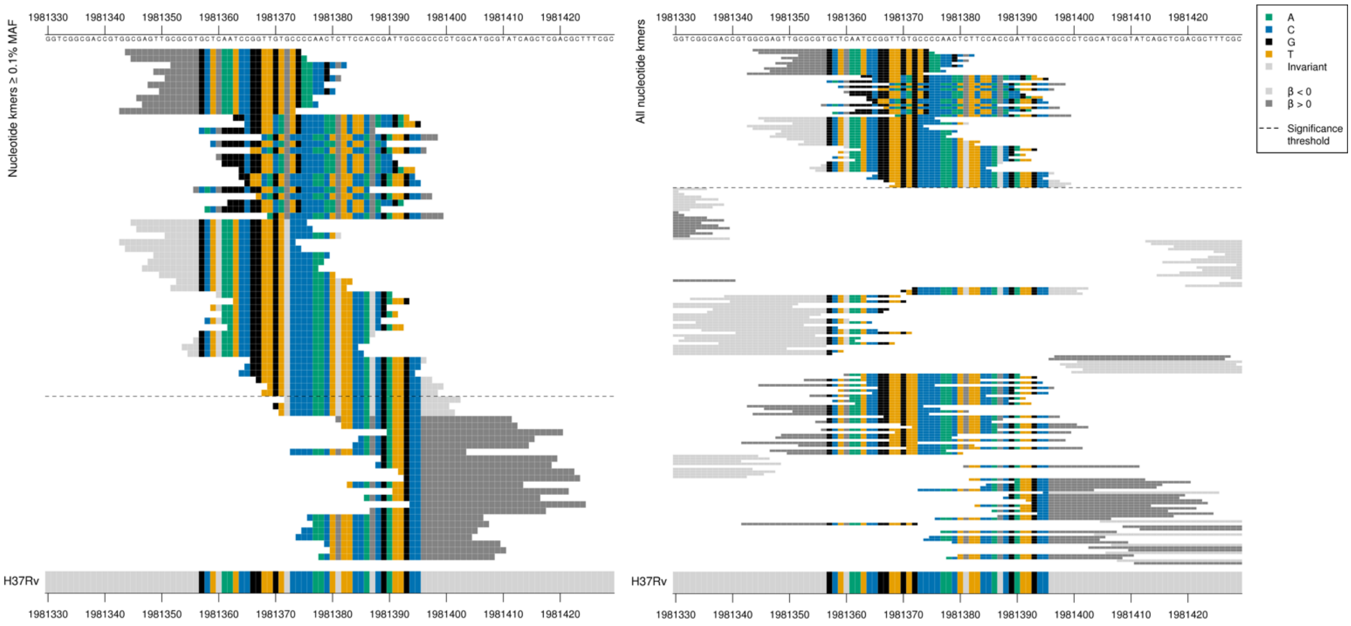

Gene begins position 1981130. Capturing multiple differences to the reference in the nine most significant high MIC-associated kmers. Most of the kmer aligned perfectly but then the rest of the kmers were very different to the reference, possibly capturing an indel. The second group of significant high MIC-associated kmers contained a CA insertion between the T of position 1981369 and the G of position 1981370. The top unaligned kmer had also been assigned to *Rv2478c:Rv2481c*. Low MIC-associated kmers all covered positions 1981369-1981373/240-244, the first 9 were more significant and appear to be covering the region of the most significant high MIC-associated kmers, the second set covered the region of the kmers with the CA insertion.
The top unaligned kmers were only assigned to *Rv1752* and *Rv2478c:Rv2481c*.

| Kmer | -log10 *p* | β | MAC | MAF | Ps | Variants |
|---|---|---|---|---|---|---|
| GCGAGTTGCGCGTGCTCAATCCGGTTGTGTG | 12.54 | 1.52 | 110 | 1.55 | 1981344 | Multiple differences to reference between positions 1981373-1981382/244-253 resulting in possible stop codon |
| GGCGGTTCAGTGCCCCAACTCTTCCACCGAT | 11.06 | 1.40 | 109 | 1.54 | 1981363 | CA insertion between positions 1981369-1981370/240-241 |
| CGAGTTGCGCGTGCTCAATCCGGTTGTGCCC | 10.89 | -1.32 | 117 | 1.65 | 1981345 | WT |

## Rv3183:Rv3188$^{R}$

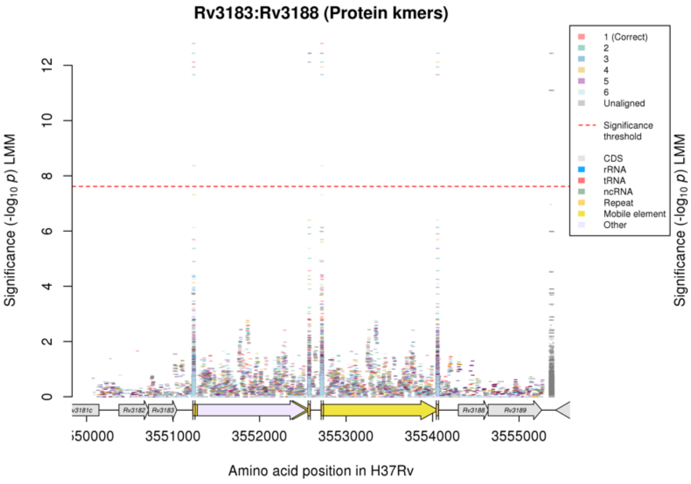

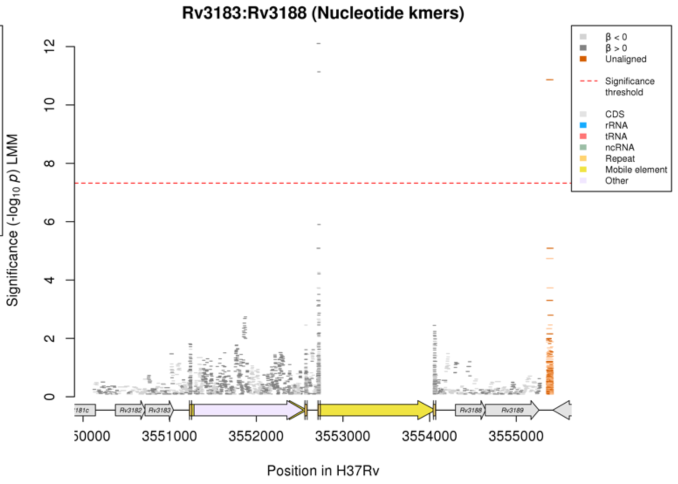

Repeat regions at the end of mobile elements

Nucleotide kmers:

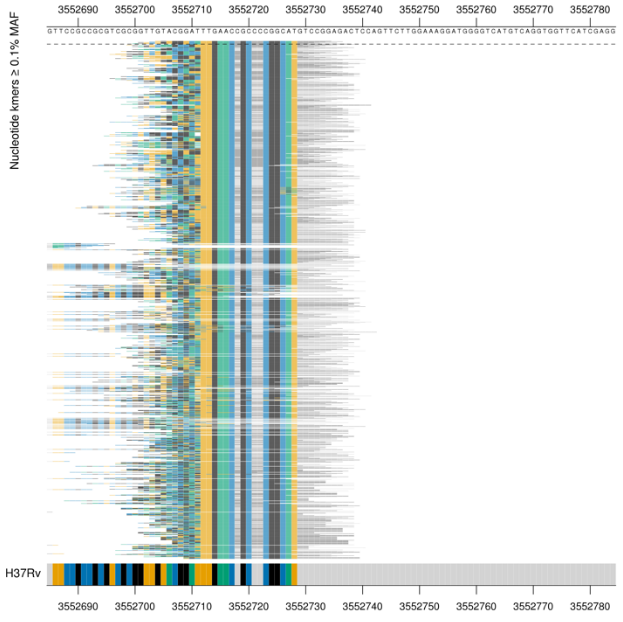

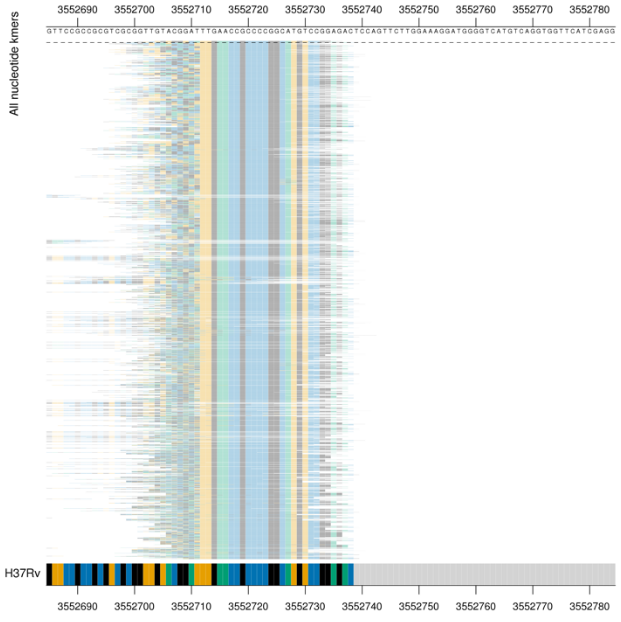

| Kmer | -log10 *p* | β | MAC | MAF | Ps | Variants |
|---|---|---|---|---|---|---|
| TGTGGTCATTGAACCGCCCCGGCATGTCCGG | 12.11 | 1.46 | 121 | 1.71 | 3552704 | Repeat region |

### *dxs2:Rv3382c* R

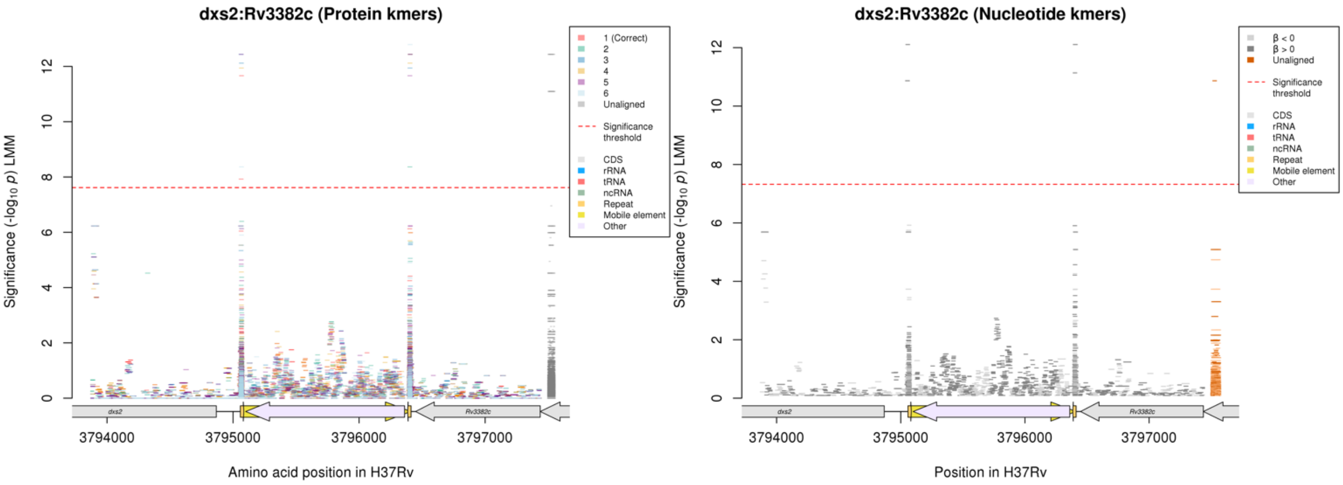

Repeat regions at the end of a mobile element

Nucleotide kmers:

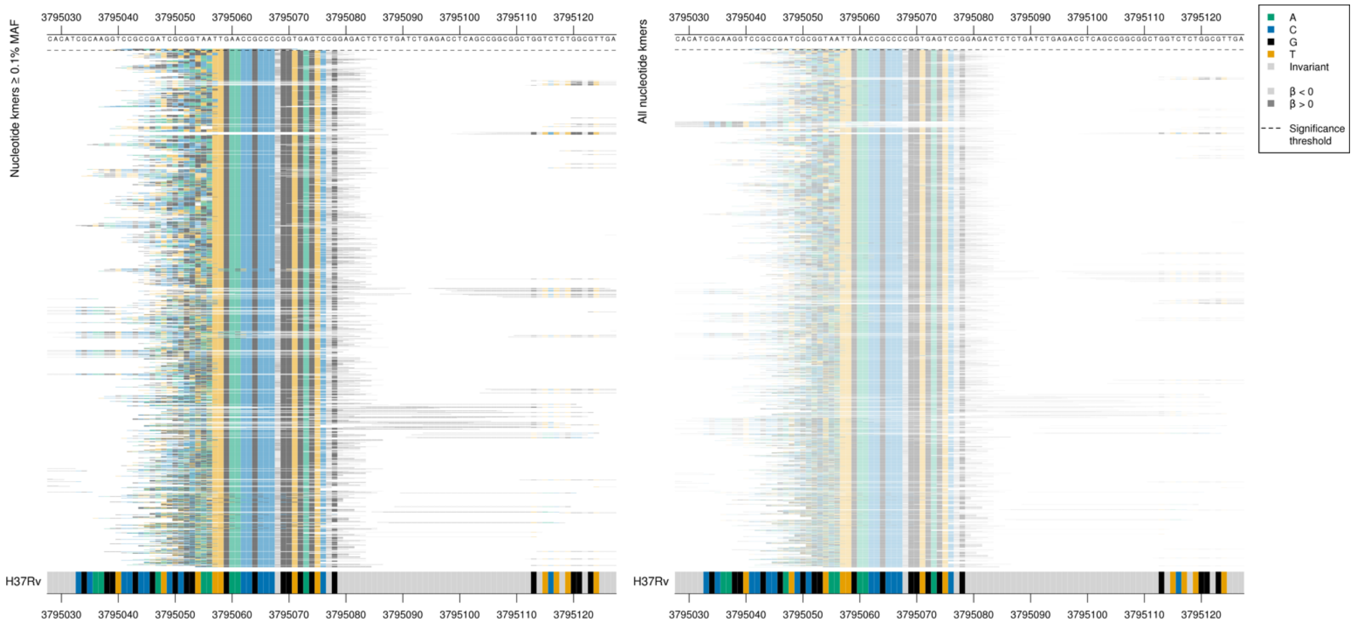

| Kmer | -log10 *p* | β | MAC | MAF | Ps | Variants |
|---|---|---|---|---|---|---|
| CCGGAGTCTGTGGTCATTGAACCGCCCCGGC | 12.11 | 1.46 | 121 | 1.71 | 3795041 | Repeat region |

Nucleotide kmers:

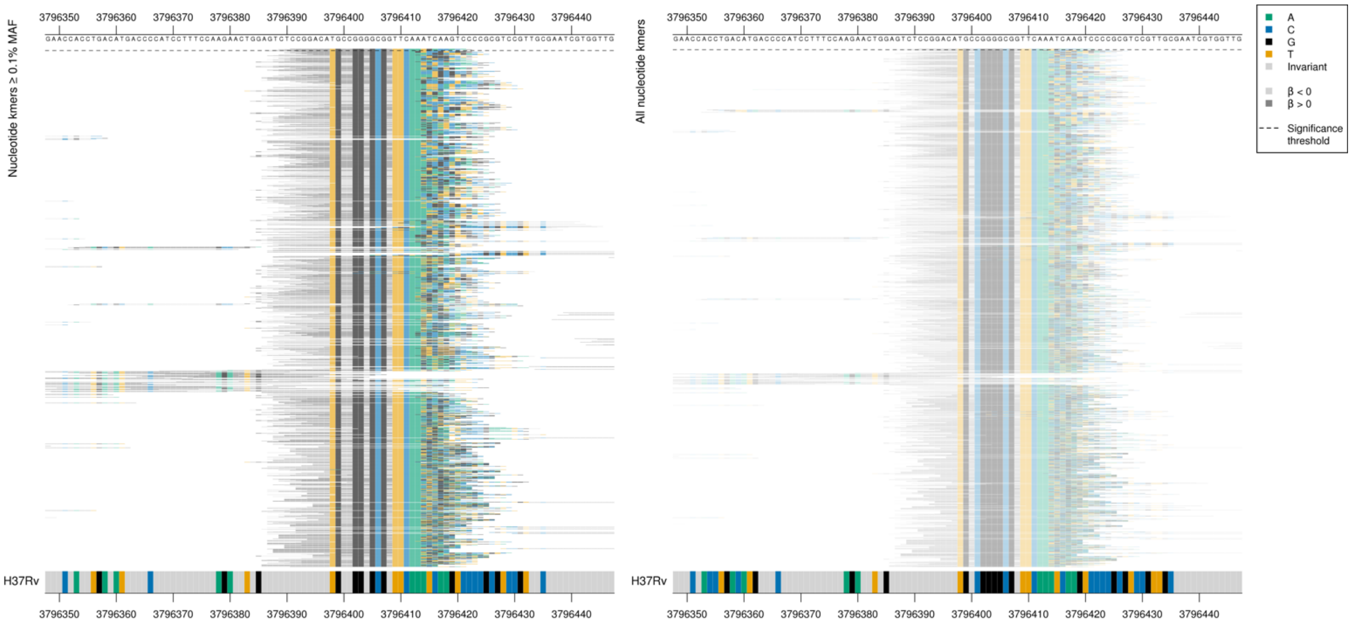

| Kmer | $-\log_{10} p$ | β | MAC | MAF | Ps | Variants |
|---|---|---|---|---|---|---|
| CCGGACATGCCGGGGCGGTTCAATGACCACA | 12.11 | 1.46 | 121 | 1.71 | 3796391 | Repeat region |

#### *rpsA/coaE*

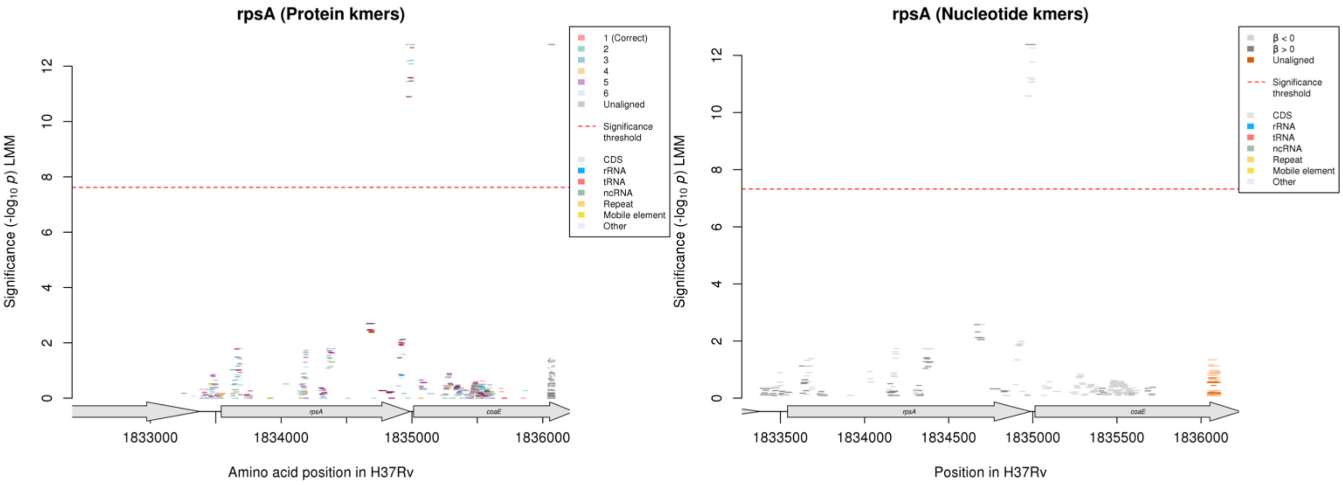

In the *rpsA* reading frame:

Protein kmers:

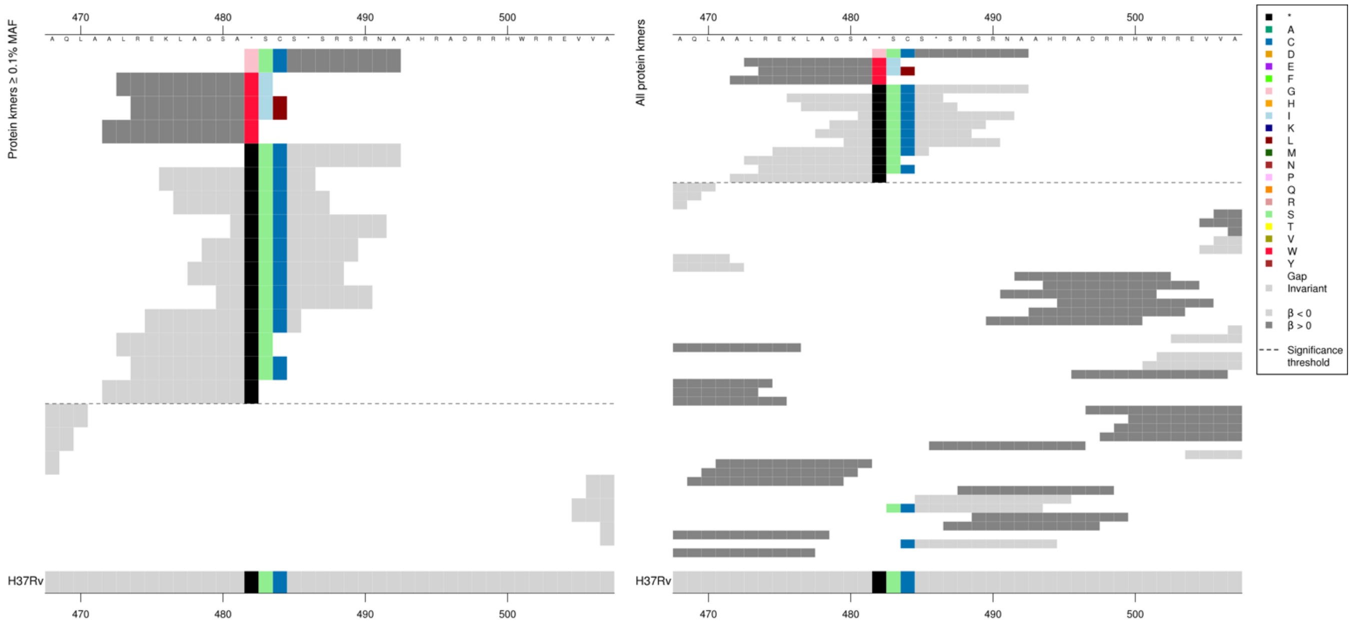

The same region for nucleotide kmers:

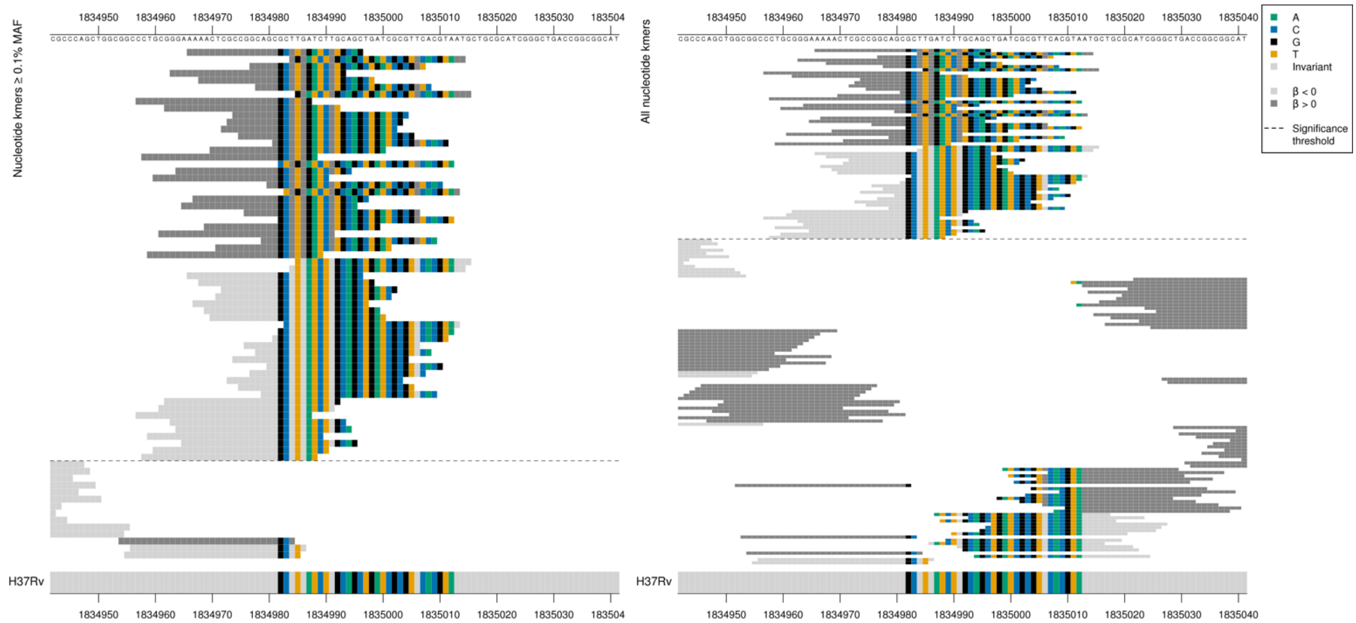

*rpsA* ends at position 1834987. G insertion in high MIC-associated kmers changing stop codon TGA to TGG (Trp / W), read through mutation, will read through to gene *coaE*

| Kmer | $-\log_{10} p$ | β | MAC | MAF | Ps | Variants |
|---|---|---|---|---|---|---|
| AAAACTCGCCGGCAGCGCTTGGATCTTGCAG | 12.38 | 1.41 | 114 | 1.61 | 1834966 | G insertion changing stop codon TGA to TGG (Trp / W), read through |
| TGATCTTGCAGCTGATCGCGTTCACGTAATG | 12.26 | -1.39 | 117 | 1.65 | 1834985 | WT |

#### *ctpI*

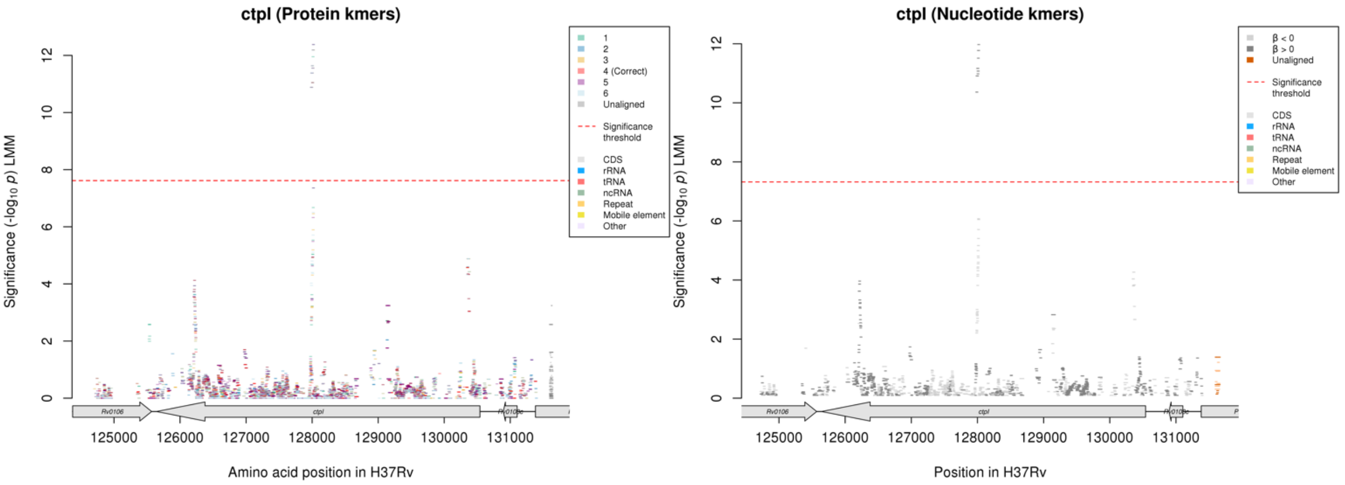

Nucleotide kmers:

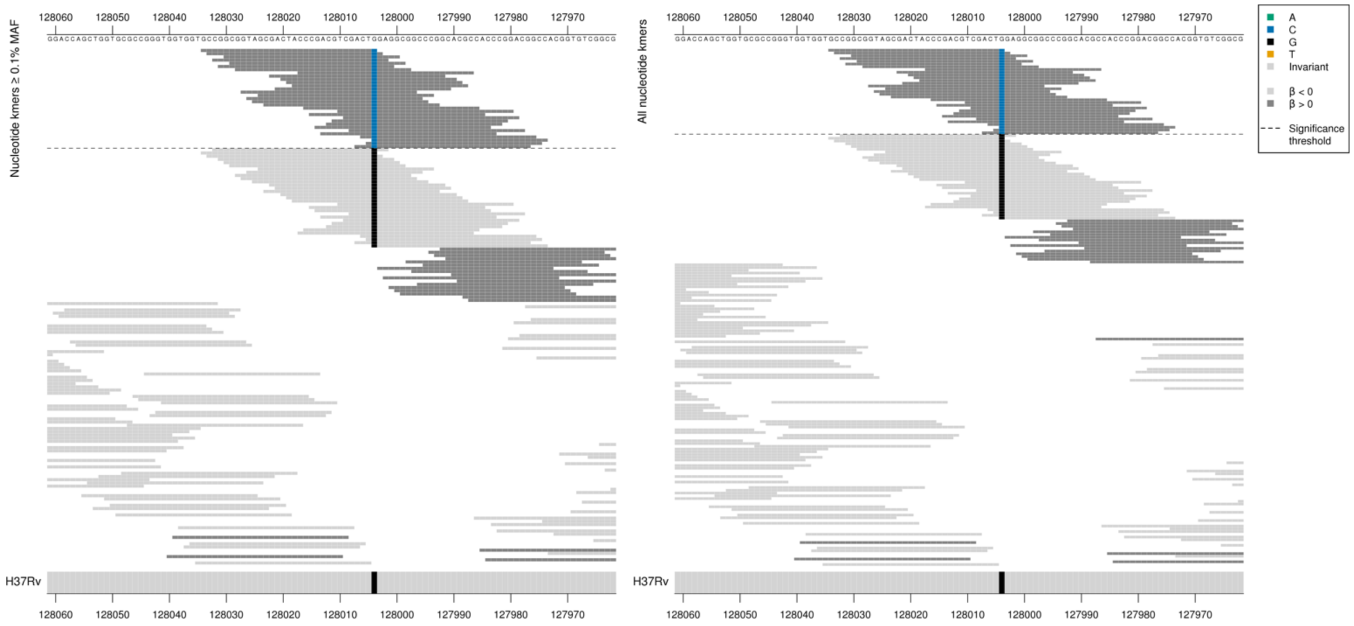

Capturing a synonymous mutation at position 128004/2538.
High MIC-associated significant kmers = CTC = Leu / L
Kmers below significance threshold = CTG = Leu / L

| Kmer | $-\log_{10} p$ | β | MAC | MAF | Ps | Variants |
|---|---|---|---|---|---|---|
| GCCGGCGGTAGCGACTACCCGACGTCGACTC | 11.97 | 1.50 | 113 | 1.60 | 128034 | 128004/2538G |

### *guaA*

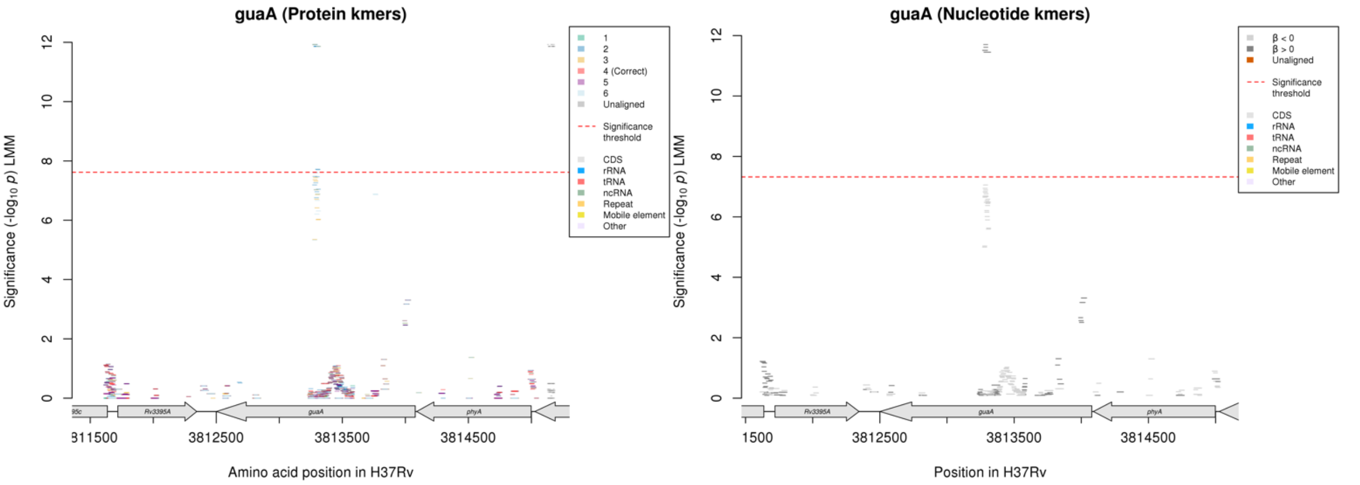

Nucleotide kmers:

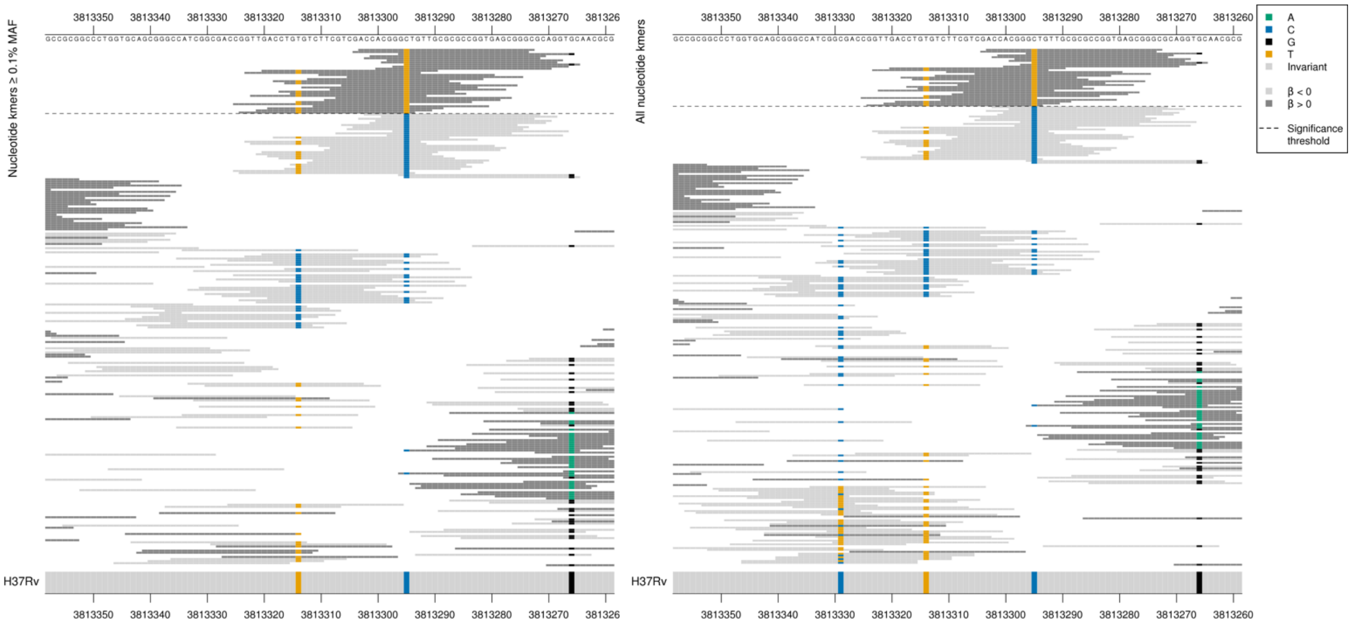

Capturing a synonymous mutation at position 3813295/784 codon 262.
High MIC-associated significant kmers = TTG = Leu / L
Kmers below significance threshold = CTG = Leu / L

| Kmer | -log$_{10}$ $p$ | β | MAC | MAF | Ps | Variants |
|---|---|---|---|---|---|---|
| ACCACGGGTTGTTGCGCGCCGGTGAGCGGGC | 11.70 | 1.49 | 111 | 1.57 | 3813303 | 3813295/784T |

#### *moaC3:Rv3327* [R]

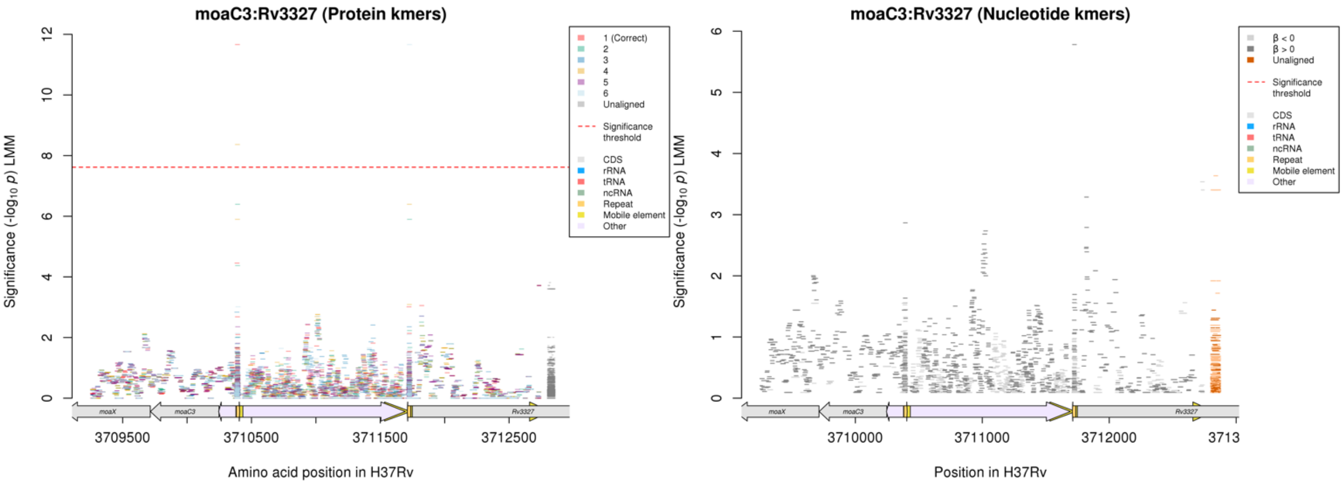

Not significant in nucleotide kmer analysis.
Repeat region

| Kmer | $-\log_{10} p$ | β | MAC | MAF | Ps | Variants |
|---|---|---|---|---|---|---|
| WSLNRPGMSGD | 11.66 | 1.23 | 132 | 1.86 | 43 | Protein kmer out of frame as falls in an intergenic region |

#### *lprF:Rv1371* [R]

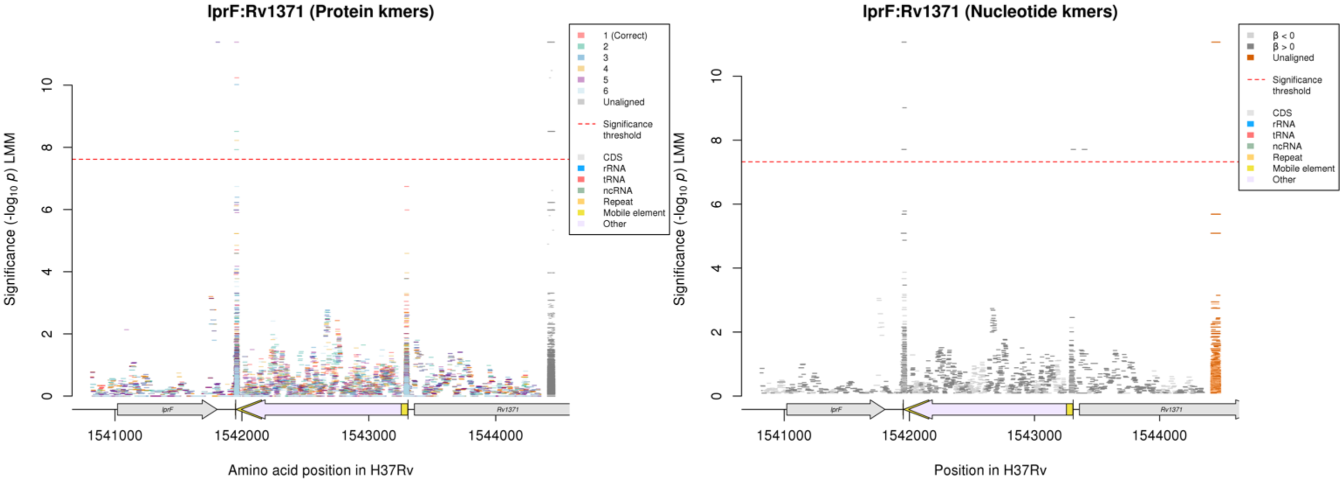

Repeat region

#### Nucleotide kmers:

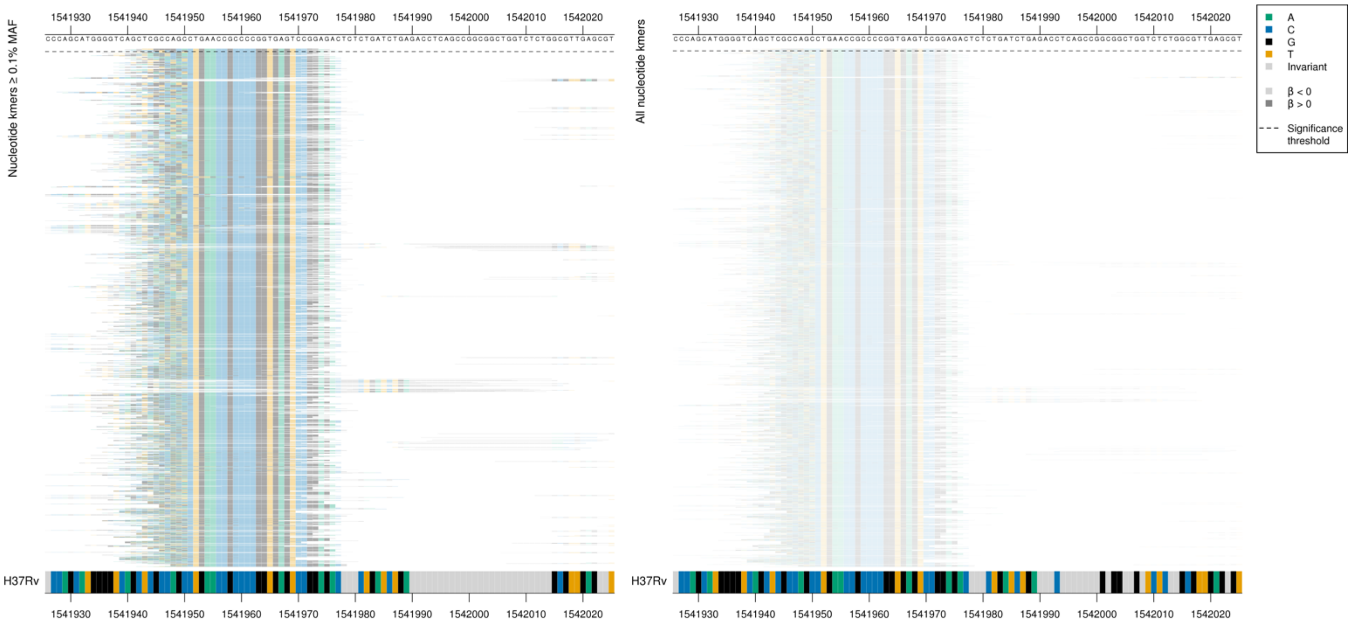

| Kmer | -$\log_{10} p$ | β | MAC | MAF | Ps | Variants |
|---|---|---|---|---|---|---|
| TTGGGGCACTGAACCGCCCCGGTGAGTCCGG | 11.06 | 1.40 | 109 | 1.54 | 1541943 | Repeat region |

#### Nucleotide kmers:

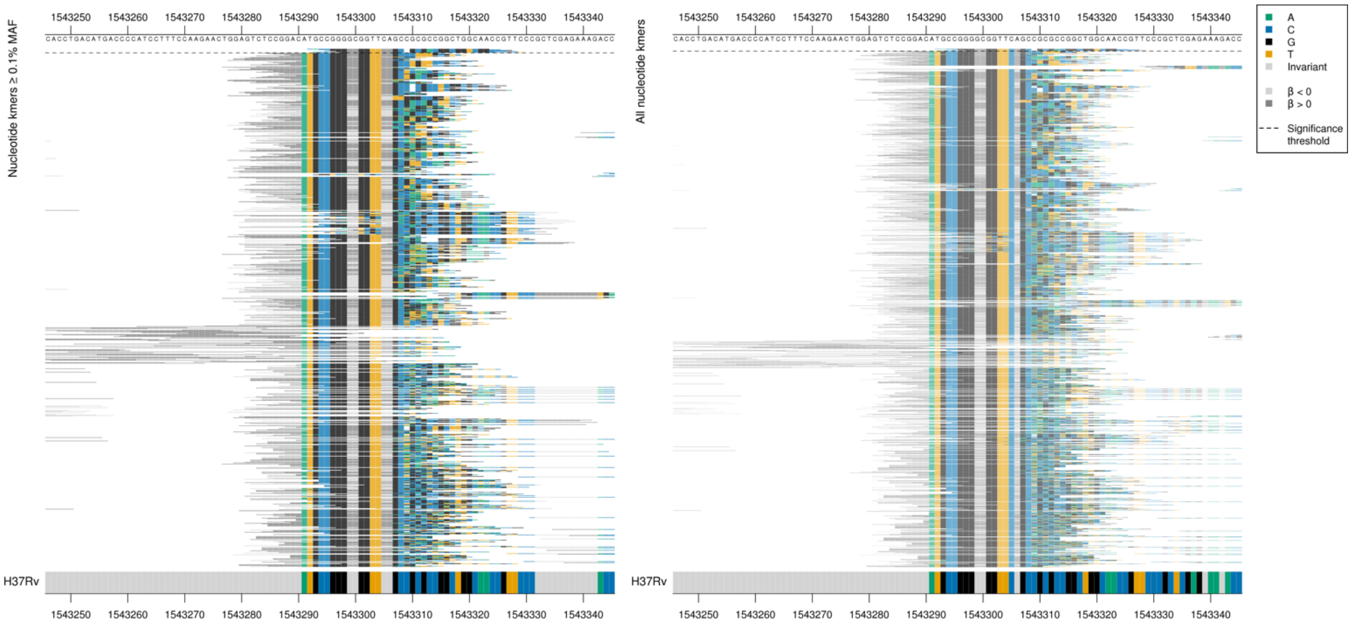

| Kmer | -$\log_{10} p$ | β | MAC | MAF | Ps | Variants |
|---|---|---|---|---|---|---|
| CACCGGGGCGGTTCAGCGCGACGGCGGTCGG | 7.71 | 2.05 | 10 | 0.14 | 1543292 | Repeat region |

### *fabG1*

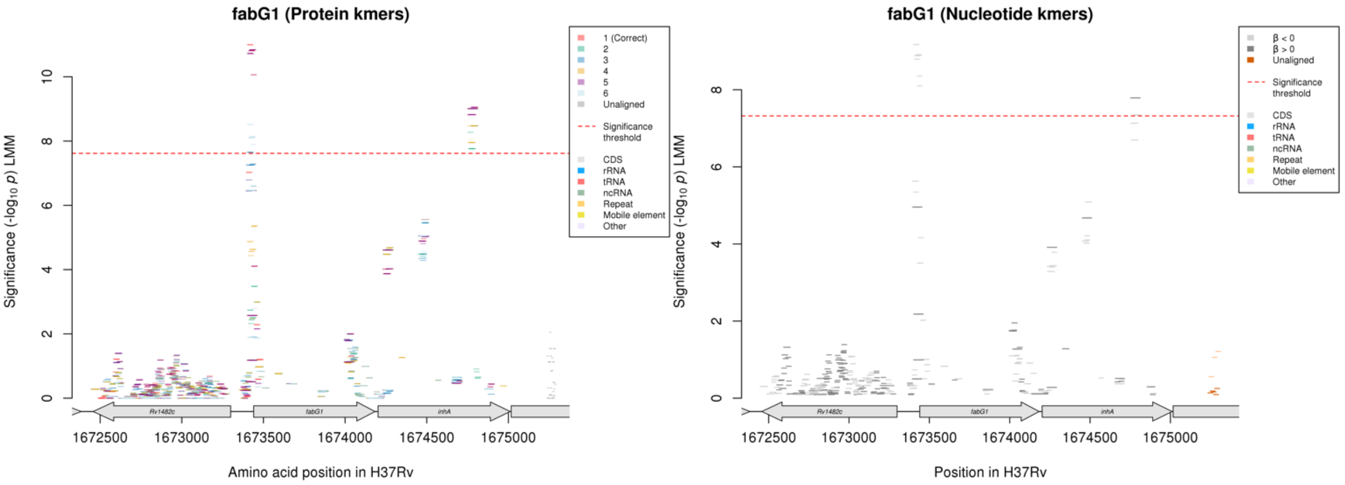

Nucleotide kmers:

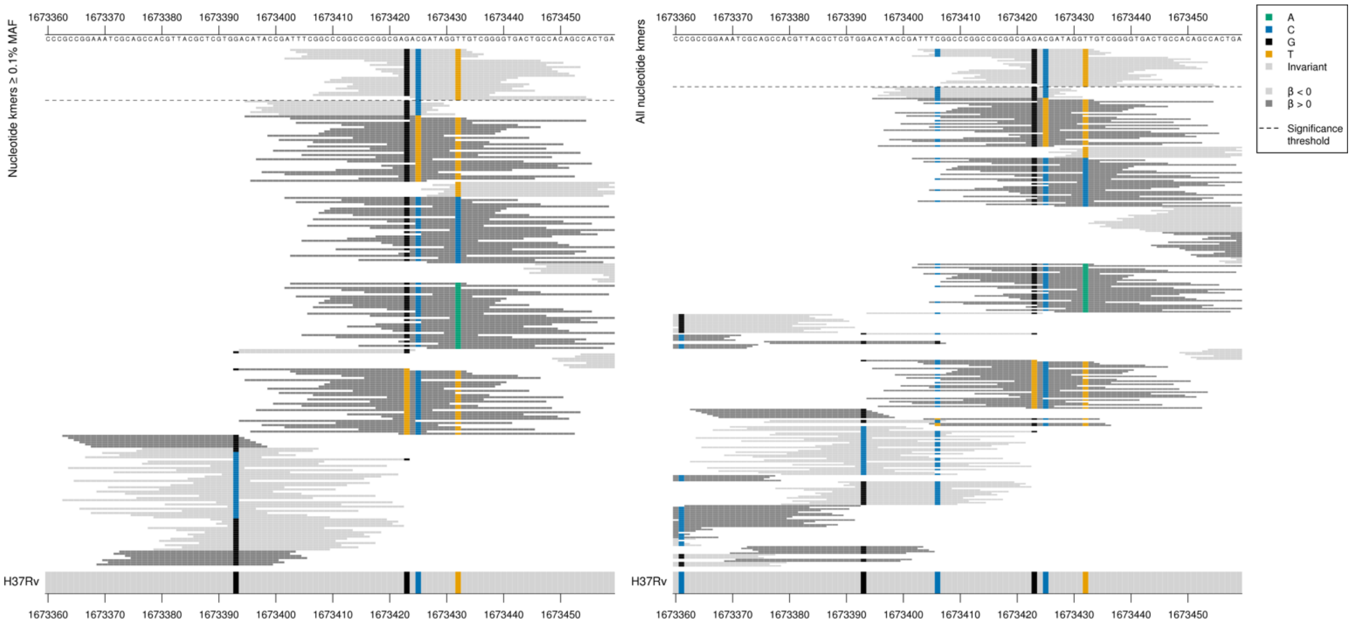

*fabG1* begins at position 1673440.

1673425:
Low MIC-associated kmers = C
Kmers below significance threshold = C/T

1673432:
Low MIC-associated kmers = T
Kmers below the significance threshold = T/C/A

1673423:
Low MIC-associated kmers = G
Kmers below significance threshold = G/T

1673406:
Low MIC-associated kmers = C
Kmers below significance threshold = C/G/T

| Kmer | $-\log_{10} p$ | β | MAC | MAF | Ps | Variants |
|---|---|---|---|---|---|---|
| TTTCGGCCCGGCCGCGGCGAGACGATAGGTT | 9.17 | -0.39 | 905 | 12.78 | 1673403 | 1673406/-34C, 1673423/-17G, 1673425/-15C, 1673432/-8T |
| GCGGCGAGACGATAGGTTGTCGGGGTGACTG | 8.91 | -0.38 | 895 | 12.64 | 1673416 | 1673423/-17G, 1673425/-15C, 1673432/-8T |
| ACGATAGGTTGTCGGGGTGACTGCCACAGCC | 8.35 | -0.38 | 764 | 10.79 | 1673424 | 1673425/-15C, 1673432/-8T |

### *spoU*

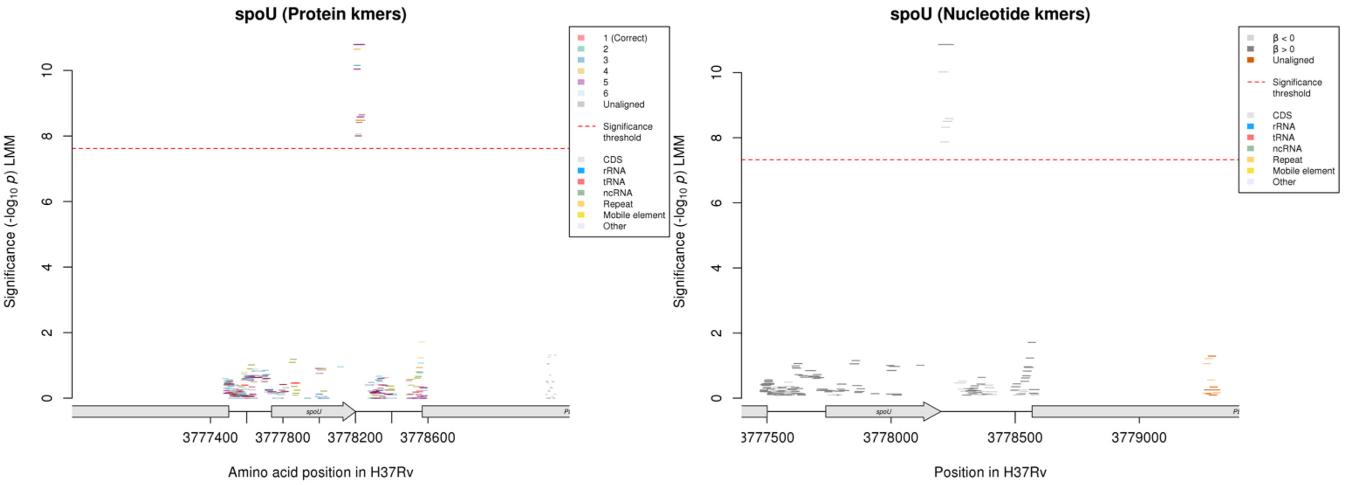

Nucleotide kmers:

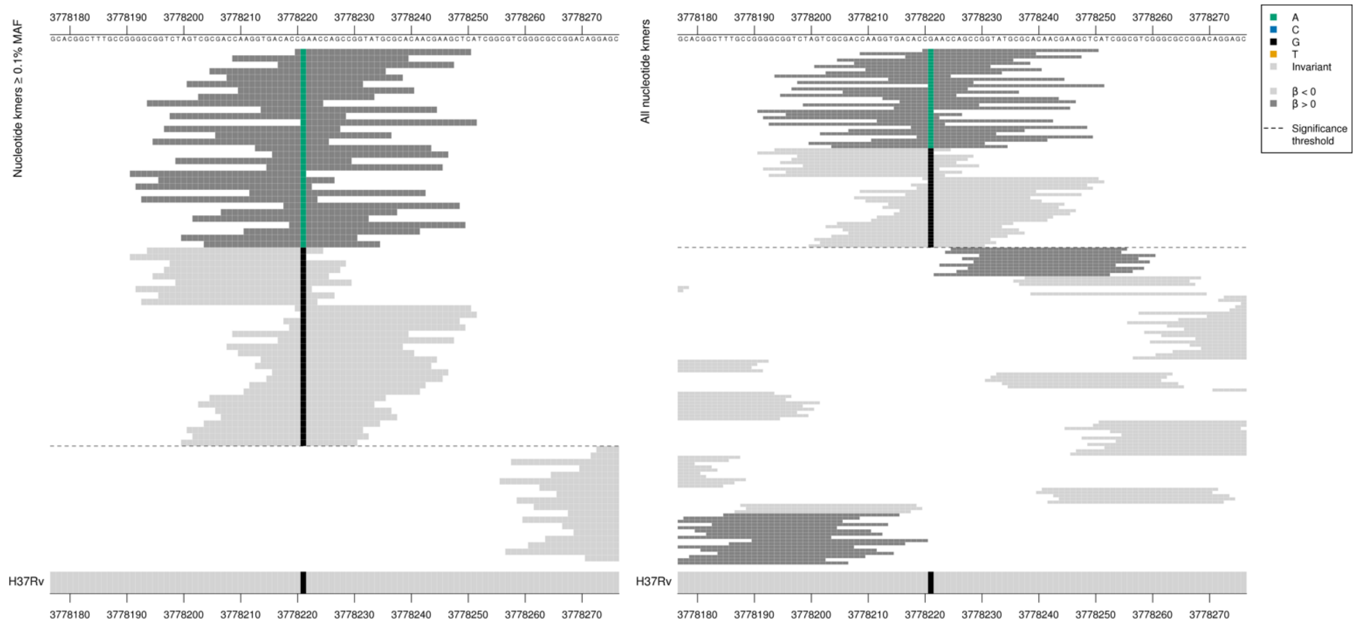

*spoU* ends at position 3778201, the kmers overlap this position. PE_*PGRS51* is the following gene but does not start until 3778568, this variant is 347 bases upstream.
Mutation 20bp downstream of the stop codon (3778221).

| Kmer | $-\log_{10} p$ | β | MAC | MAF | Ps | Variants |
|---|---|---|---|---|---|---|
| CAAACCAGCCGGTATGCGCACAACGAAGCTC | 10.86 | 1.36 | 163 | 2.30 | 3778220 | 3778221A (20bp after stopcodon) |
| CGGTCTAGTCGCGACCAAGGTGACACCGAAC | 10.02 | -1.25 | 166 | 2.34 | 3778194 | 3778221G (20bp after stopcodon) |

### *glpK*

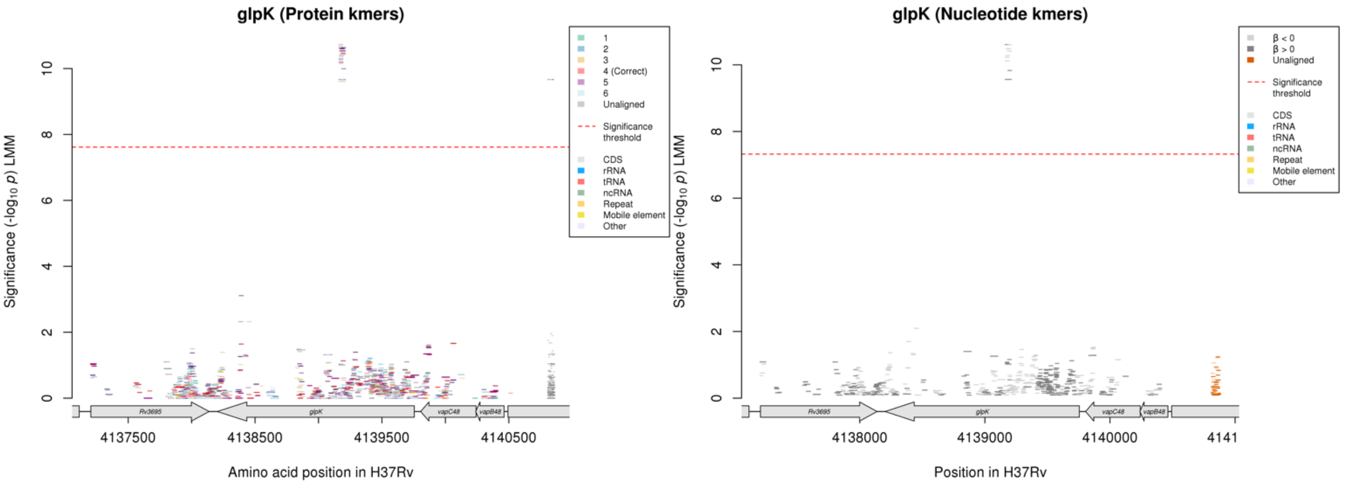

#### Protein kmers:

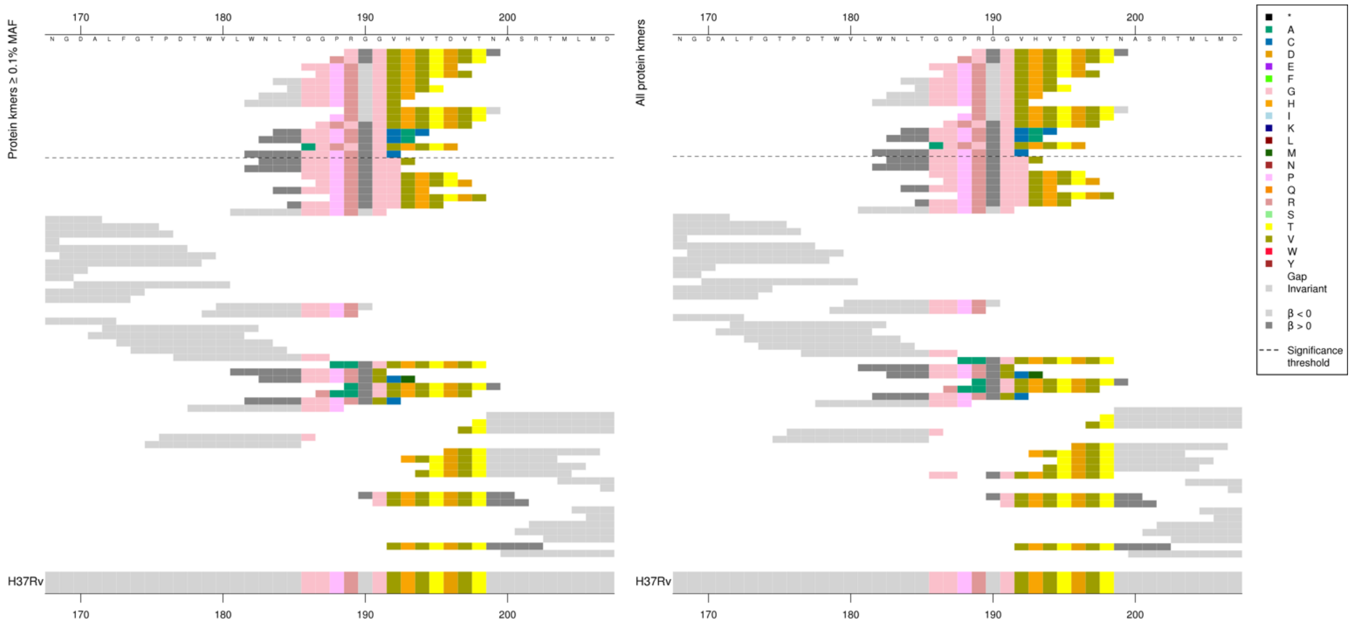

#### Nucleotide kmers:

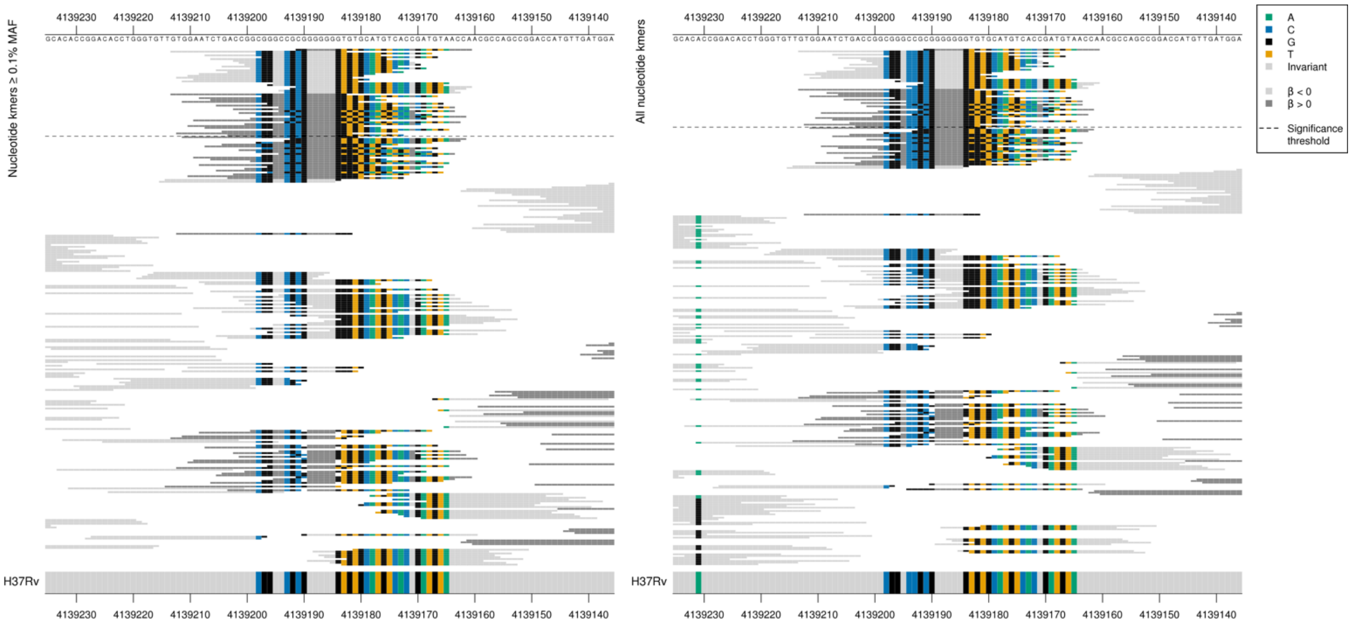

| Kmer | $-\log_{10} p$ | β | MAC | MAF | Ps | Variants |
|---|---|---|---|---|---|---|
| GGGGGGGGTGTGCATGTCACCGATGTAACCA | 10.61 | 0.57 | 181 | 2.56 | 4139191 | G insertion between 4139191-4139183/565-573 |
| GTGGAATCTGACCGGCGGGCCGCGGGGGGGT | 10.60 | -0.54 | 212 | 2.99 | 4139213 | WT |

### Ethionamide (ETH)

#### *fabG1*

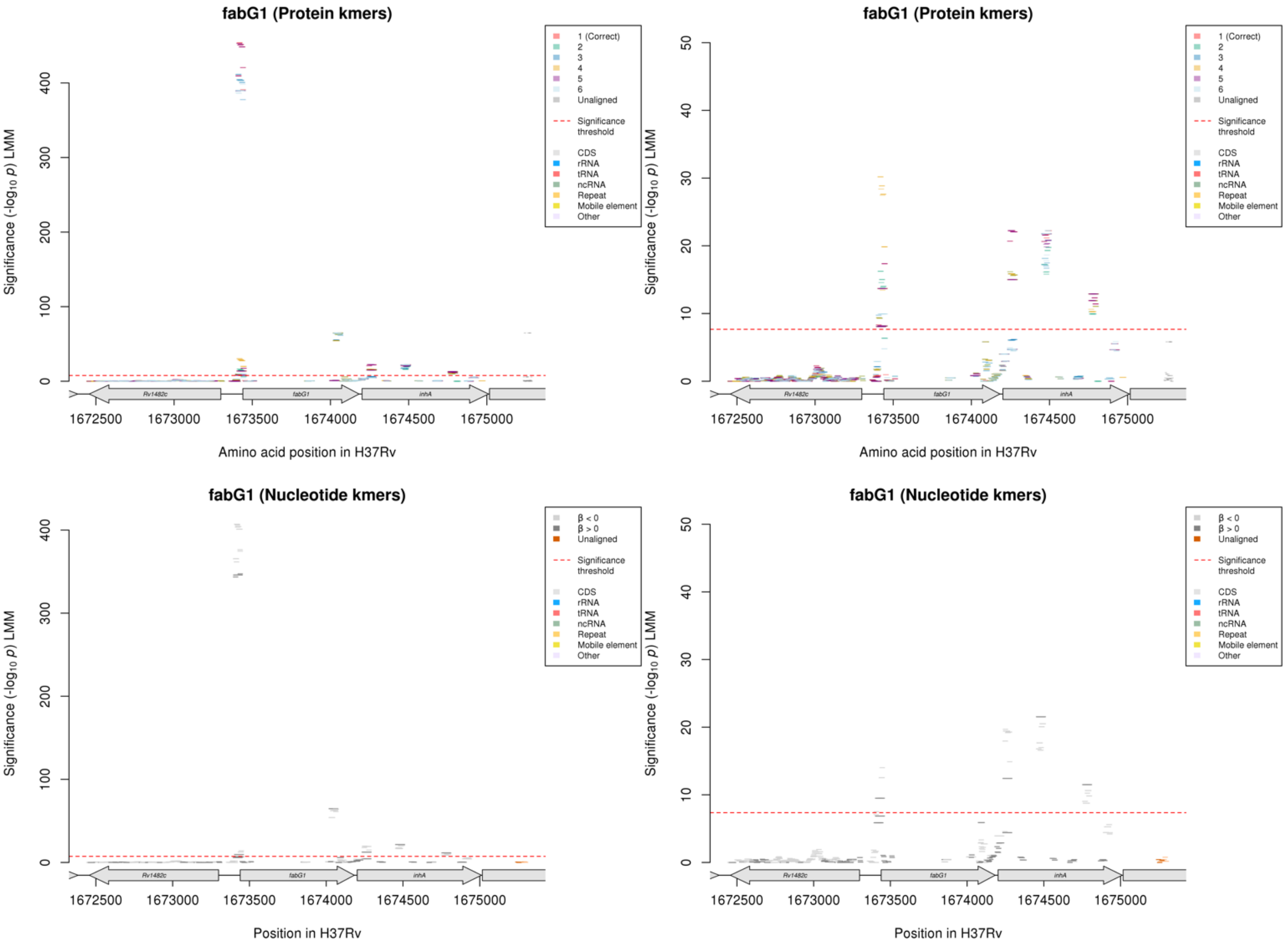

Nucleotide kmers:

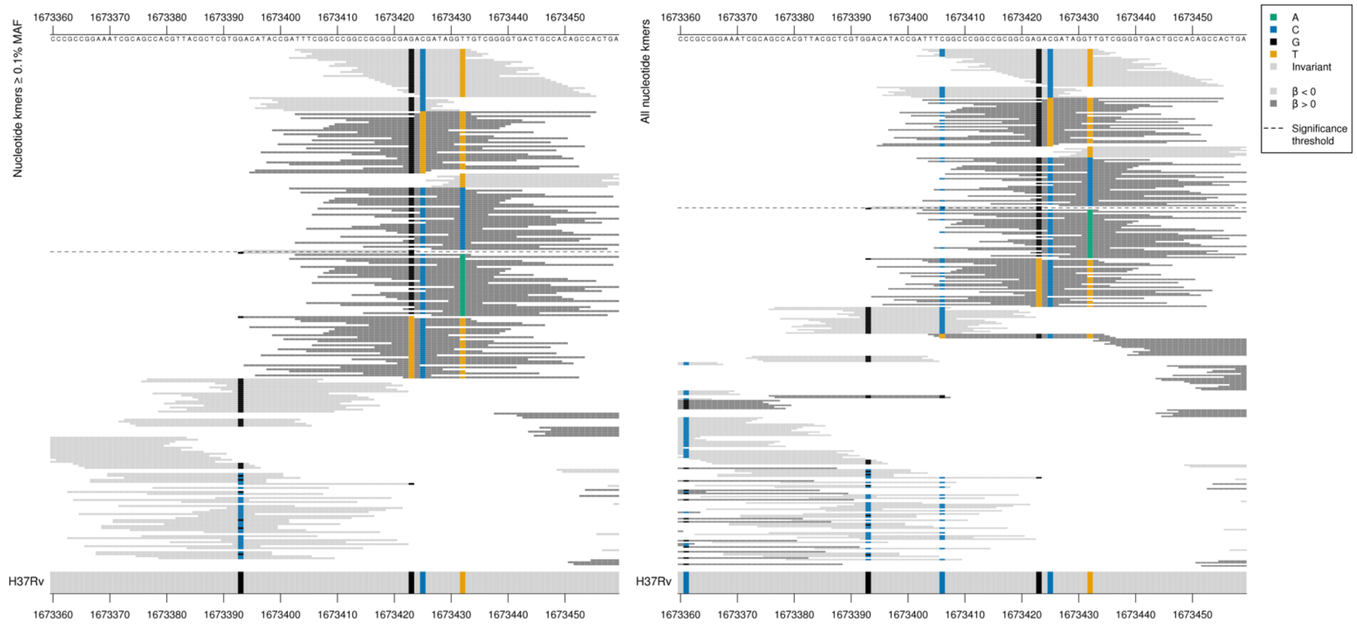

*fabG1* begins at position 1673440. Mainly seems to be capturing mutations in the promoter region at positions 1673425 and 1673432, but the *p*-values varied depending on the allele at position 1673423.

1673425:
Low MIC-associated kmers = C
Some high MIC-associated kmers = T

1673432:
Low MIC-associated kmers = T
Some high MIC-associated kmers = C (significant) or A (just below significance)

1673423:
Low MIC-associated kmers = G
Kmers below significance threshold = T

| Kmer | $-\log_{10} p$ | β | MAC | MAF | Ps | Variants |
|---|---|---|---|---|---|---|
| TTTCGGCCCGGCCGCGGCGAGACGATAGGTT | 406.99 | -2.57 | 1035 | 12.49 | 1673403 | 1673406/-34C, 1673423/-17G, 1673425/-15C, 1673432/-8T |
| GGCCCGGCCGCGGCGAGACGATAGGTTGTCG | 405.92 | -2.58 | 1028 | 12.40 | 1673407 | 1673423/-17G, 1673425/-15C, 1673432/-8T |
| ACGATAGGTTGTCGGGGTGACTGCCACAGCC | 376.34 | -2.56 | 881 | 10.63 | 1673424 | 1673425/-15C, 1673432/-8T |
| CATACCGATTTCGGCCCGGCCGCGGCGAGAC | 365.47 | -2.58 | 944 | 11.39 | 1673395 | 1673406/-34C, 1673423/-17G, 1673425/-15C |
| TGATAGGTTGTCGGGGTGACTGCCACAGCCA | 347.25 | 2.62 | 788 | 9.51 | 1673425 | 1673425/-15T, 1673432/-8T |
| TTTCGGCCCGGCCGCGGCGAGATGATAGGTT | 345.87 | 2.62 | 787 | 9.50 | 1673403 | 1673423/-17G, 1673425/-15T, 1673432/-8T |
| CCGATTTCGGCCCGGCCGCGGCGAGATGATA | 345.87 | 2.62 | 787 | 9.50 | 1673399 | 1673423/-17G, 1673425/-15T |
| TTGTCGGGGTGACTGCCACAGCCACTGAAGG | 14.01 | -1.14 | 95 | 1.15 | 1673432 | 1673432/-8T |
| ATTTCGGCCCGGCCGCGGCGAGACGATAGGC | 9.50 | 1.17 | 51 | 0.62 | 1673402 | 1673432/-8C |
| ACATACCGATTTCGGCCCGGCCGCGGCGAGA | 7.51 | -0.93 | 161 | 1.94 | 1673394 | 1673406/-34C, 1673423/-17G |

Nucleotide kmers:

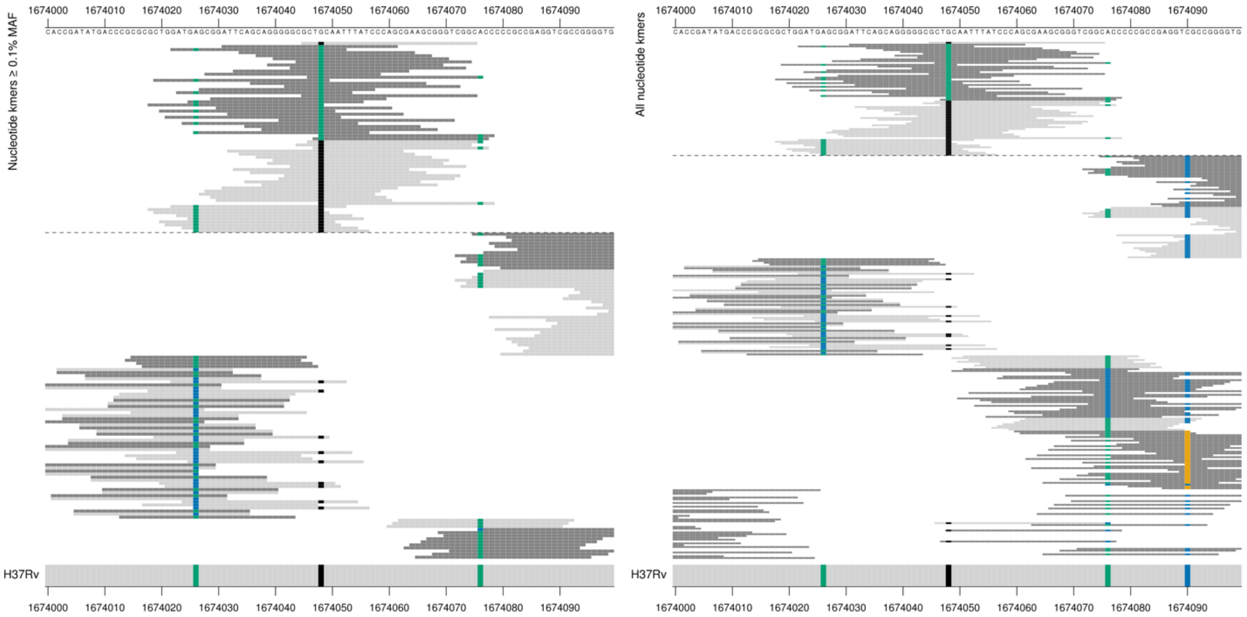

CTA/CTG
Synonymous L203L

| Kmer | $-\log_{10} p$ | β | MAC | MAF | Ps | Variants |
|---|---|---|---|---|---|---|
| GCTGCAATTTATCCCAGCGAAGCGGGTCGGC | 64.80 | -2.27 | 111 | 1.34 | 1674045 | 1674048/609G |
| ATTCAGCAGGGGGCGCTACAATTTATCCCAG | 64.68 | 2.30 | 107 | 1.29 | 1674031 | 1674048/609A |
| CTGCAATTTATCCCAGCGAAGCGGGTCGGCA | 64.18 | -2.25 | 112 | 1.35 | 1674046 | 1674048/609G, 1674076/637A |
| GATGAGCGGATTCAGCAGGGGGCGCTGCAAT | 54.00 | -1.99 | 121 | 1.46 | 1674022 | 1674026/587A, 1674048/609G |

### *ethA*

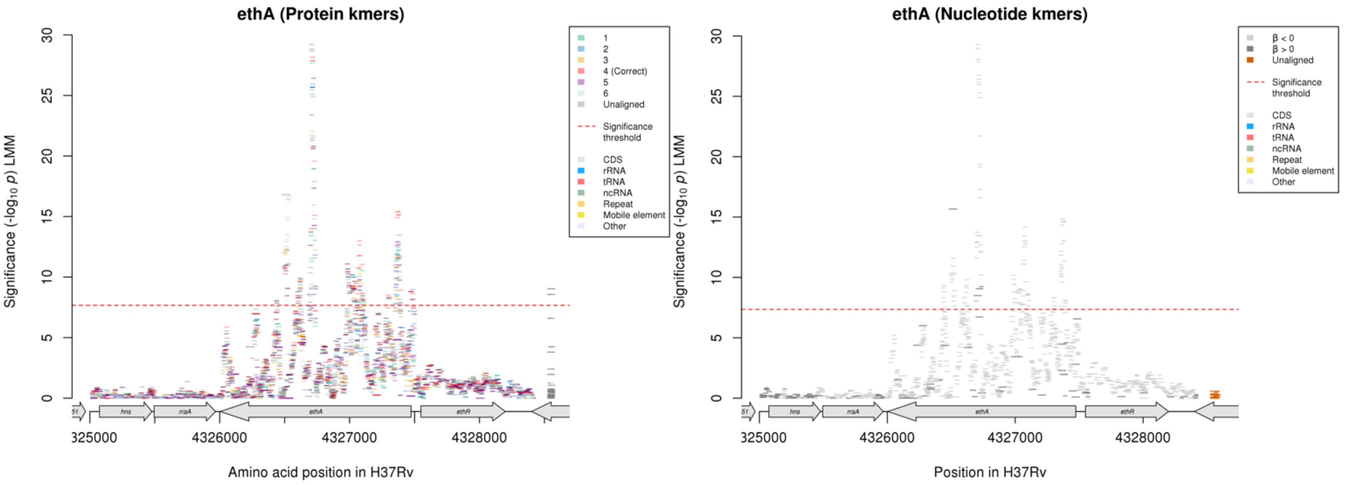

Protein kmers:

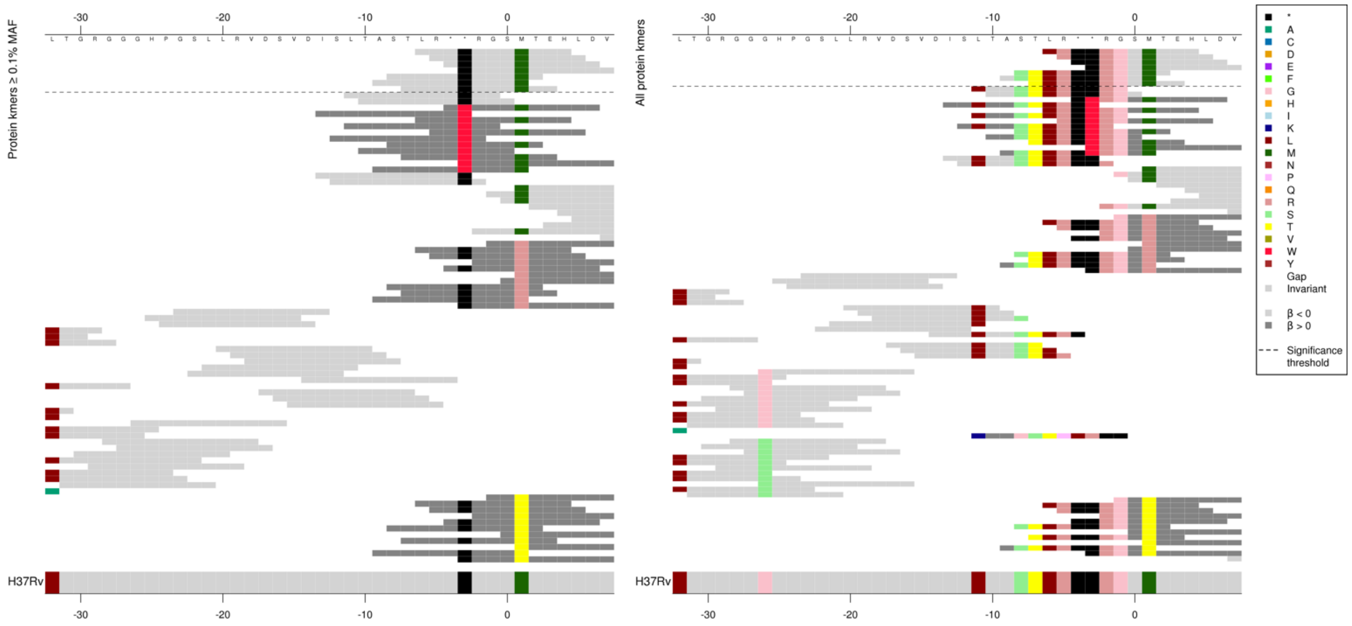

Nucleotide kmers were not significant in this region, but showing the alignments to interpret what the protein kmers were capturing here:

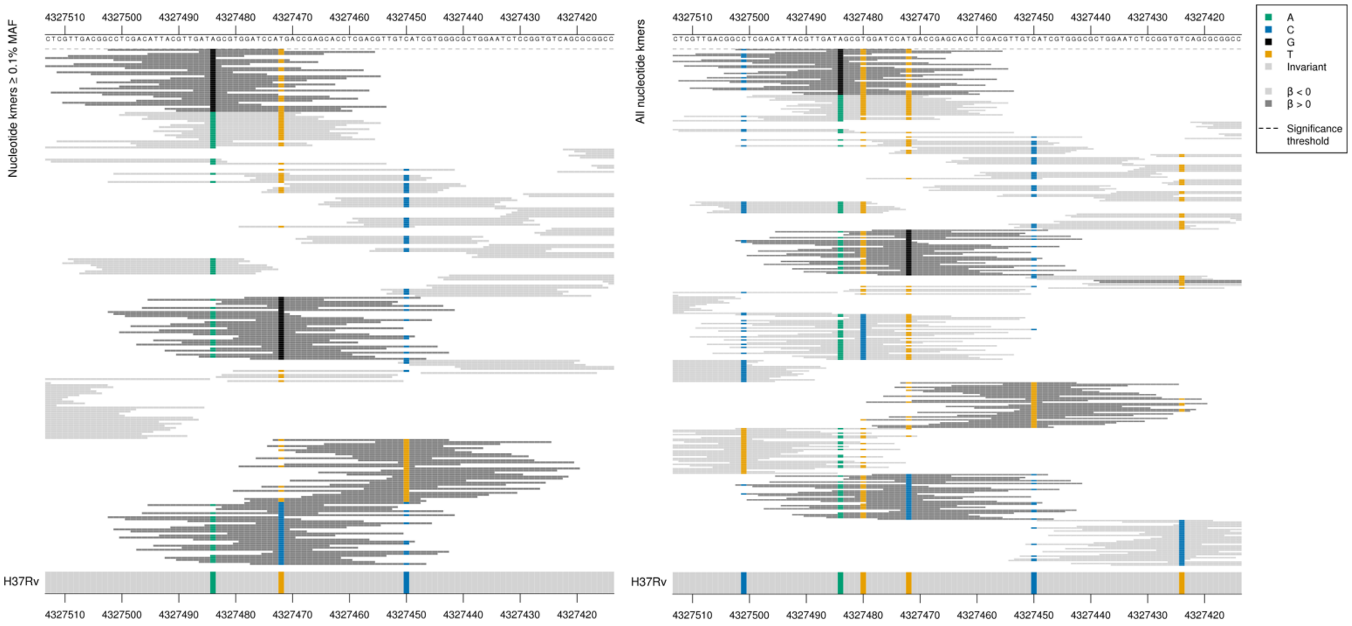

*ethA* begins at position 4327473.

Low MIC-associated kmers capturing the reference allele at a combination of mutations at positions 4327484/-11 and 4327472/2.

4327484/-11:
Kmers below significance threshold = G

4327472/2:
Kmers below significance threshold = G/C changing start codon ATG to AGG/ACG

| Kmer | $-\log_{10} p$ | β | MAC | MAF | Ps | Variants |
|---|---|---|---|---|---|---|
| LR**RGSMTEH | 8.80 | -0.99 | 177 | 2.14 | -6 | 4327484/-11A (nucleotide), 4327472/2T (nucleotide, 1M amino acid) (Protein kmer doesn't exist as spanning the UTR) |

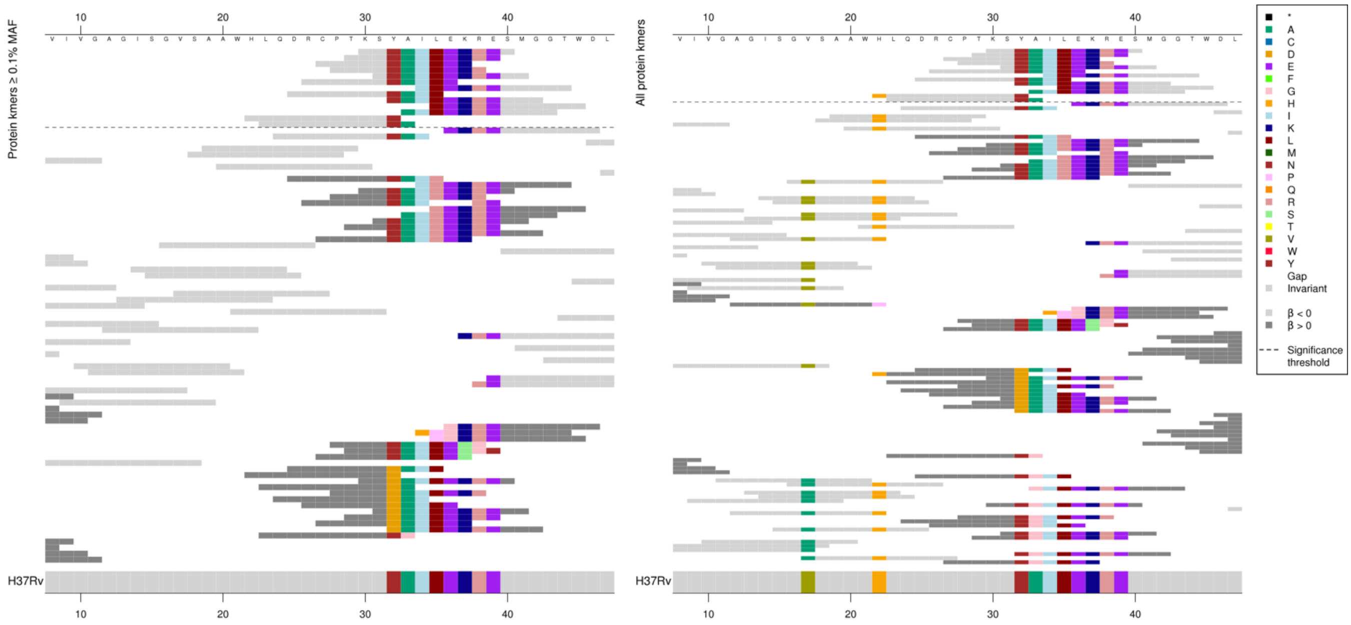

Nucleotide kmers in the same region:

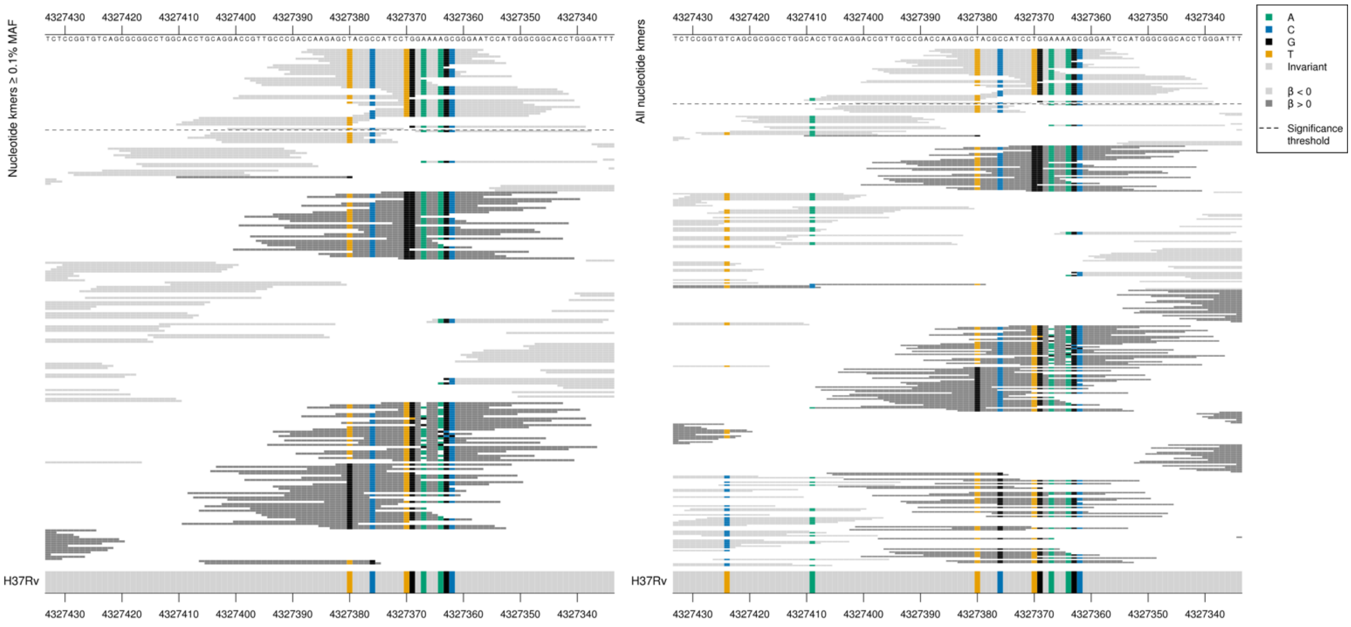

First set of kmers appear to be mainly capturing 35L, but other amino acid changes may be influencing the *p*-value. Second set of reference sequence low MIC-associated kmers were possibly capturing 32Y plus other positions.

| Kmer | $-log_{10}$ *p* | β | MAC | MAF | Ps | Variants |
|---|---|---|---|---|---|---|
| KSYAILEKRES | 15.40 | -1.23 | 177 | 2.14 | 30 | WT / 35L |
| HLQDRCPTKSY | 8.92 | -1.10 | 141 | 1.70 | 22 | WT / 22H, 32Y |
| LQDRCPTKSYA | 7.87 | -1.07 | 146 | 1.76 | 23 | WT / 32Y, 33A |

Protein kmers:

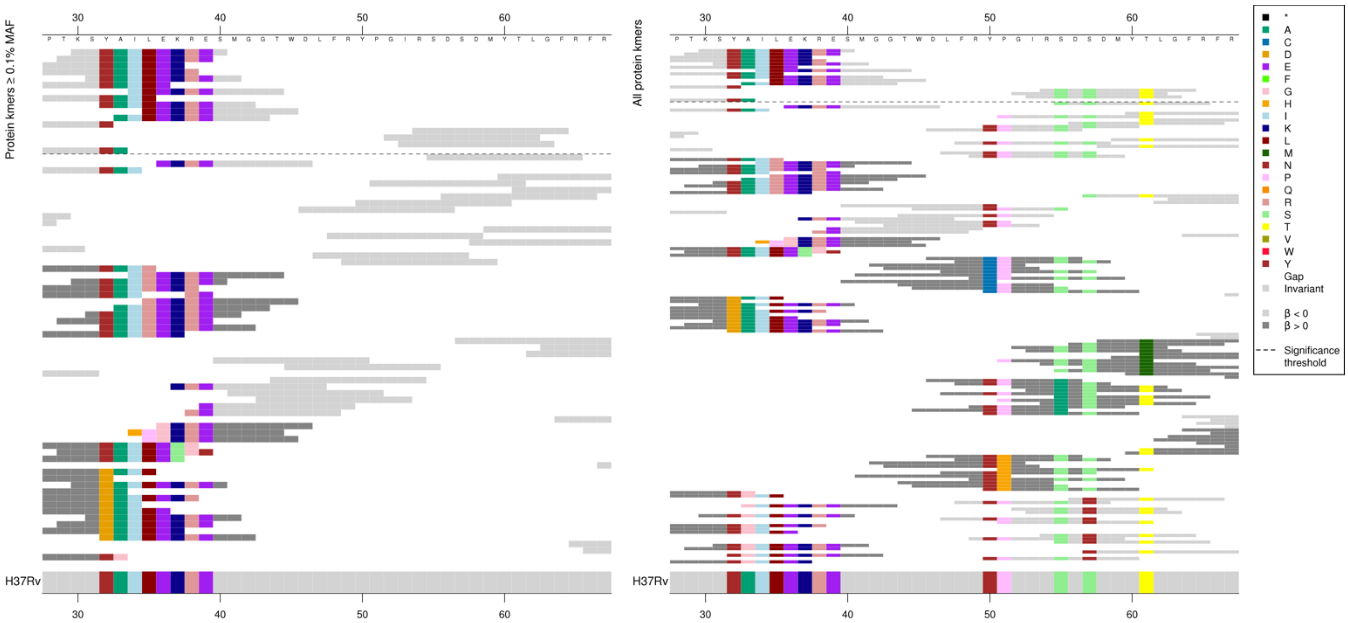

Kmers below significance threshold = 55A, 57Y, 61M

| Kmer | $-log_{10}$ *p* | β | MAC | MAF | Ps | Variants |
|---|---|---|---|---|---|---|
| RSDSDMYTLGF | 8.50 | -0.94 | 150 | 1.81 | 54 | WT / 55S, 57S, 61T |

Protein kmers:

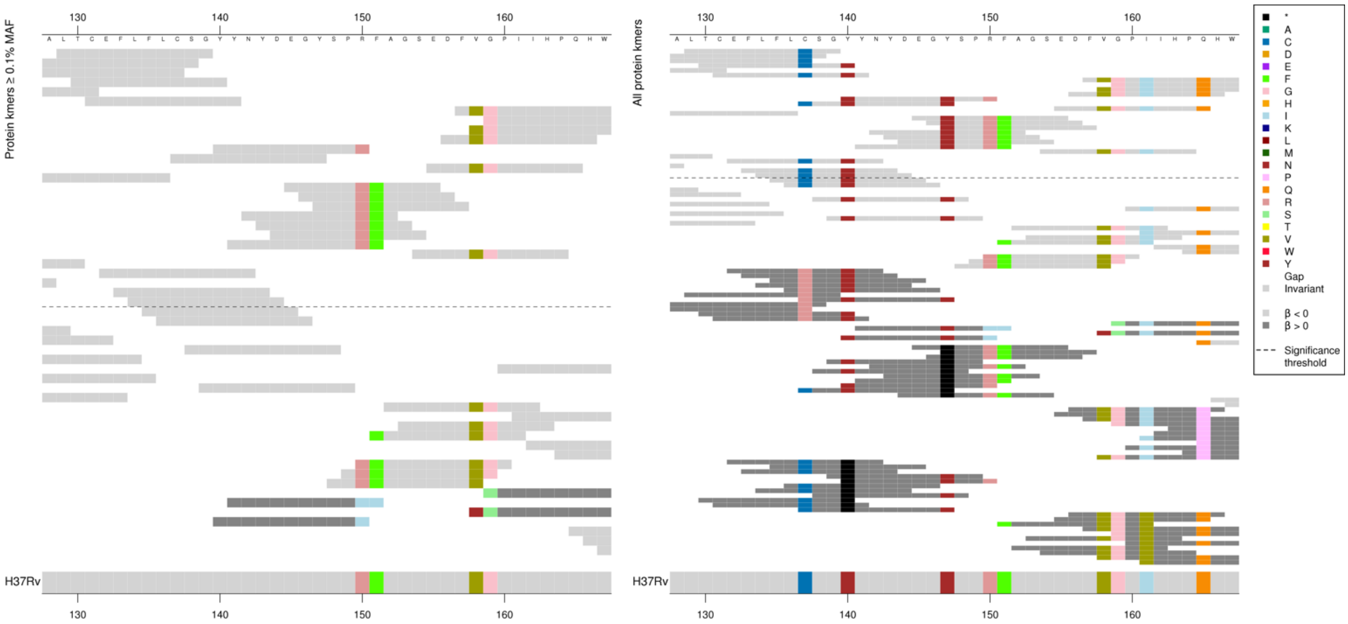

Same region for nucleotide kmers:

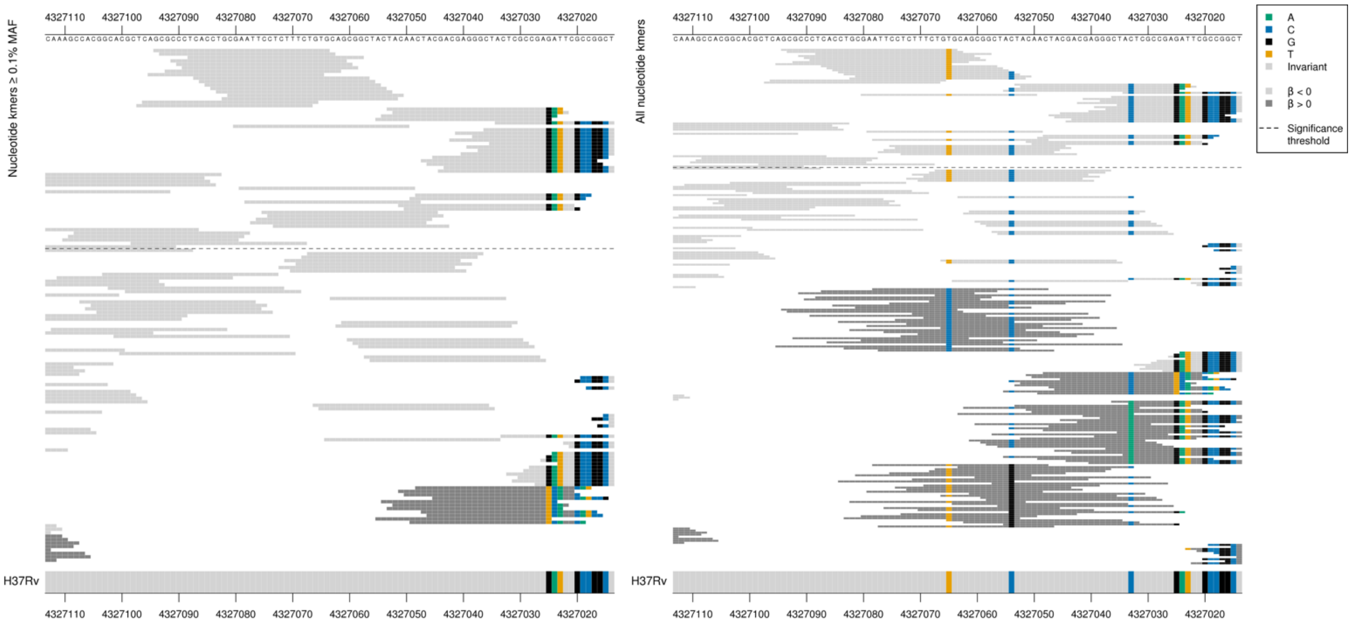

Low MIC-associated kmers all capturing the reference sequence in this region. Showing just those where there was variation at sites within the kmer for this region. The main positions (but not significant) substitutions that these were likely capturing: 137R, 140*, 147*,161V, 165P.

| Kmer | $-\log_{10} p$ | β | MAC | MAF | Ps | Variants |
|---|---|---|---|---|---|---|
| LTCEFLFLCSG | 13.00 | -1.37 | 123 | 1.48 | 129 | WT / 137C |
| TCEFLFLCSGY | 11.76 | -1.22 | 129 | 1.56 | 130 | WT / 137C, 140Y |
| FVGPIIHPQHW | 10.44 | -1.08 | 145 | 1.75 | 157 | WT / 158V, 159G, 161I, 165Q |
| GPIIHPQHWPE | 10.44 | -1.08 | 145 | 1.75 | 159 | WT / 159G, 161I, 165Q |
| YYNYDEGYSPR | 9.64 | -1.17 | 134 | 1.62 | 140 | WT / 140Y, 147Y, 150R |
| CSGYYNYDEGY | 9.49 | -1.12 | 129 | 1.56 | 137 | WT / 137C, 140Y, 147Y |
| EGYSPRFAGSE | 9.25 | -1.24 | 128 | 1.54 | 145 | WT / 147Y, 150R, 151F |
| SEDFVGPIIHP | 8.66 | -1.12 | 132 | 1.59 | 154 | WT / 158V, 159G, 161I |

Protein kmers:

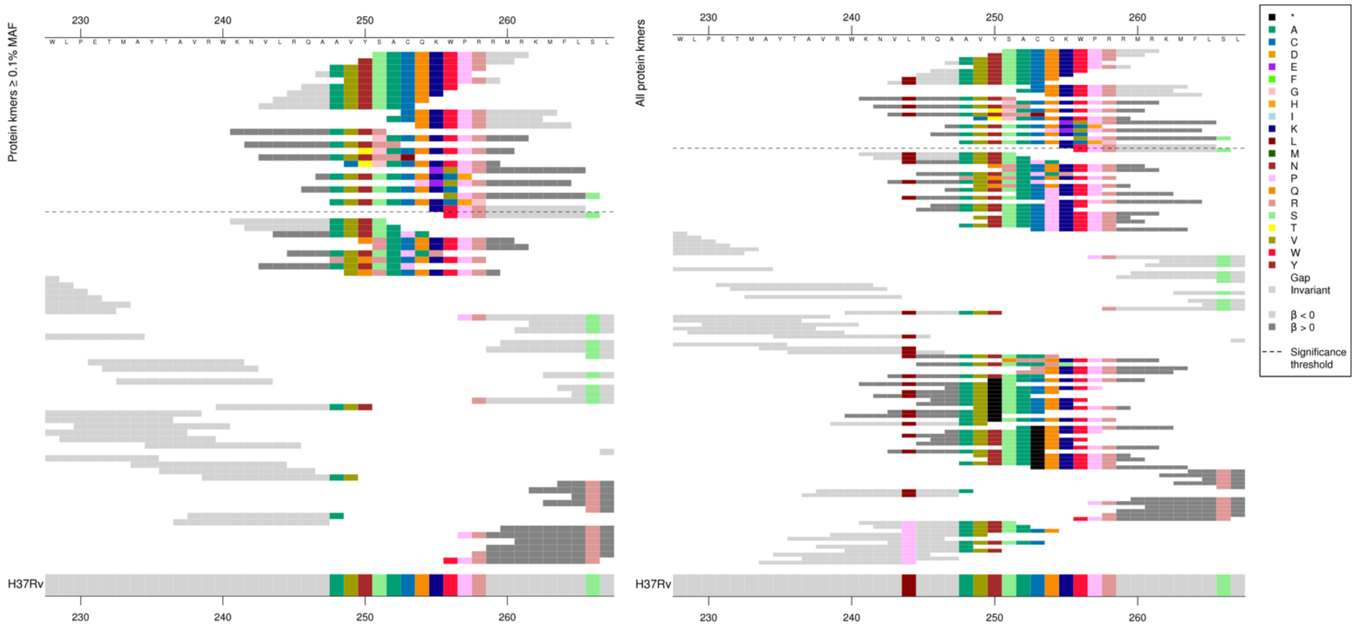

First alignment shows low MIC-associated kmers which were identical to the reference. The high MIC-associated kmers were capturing multiple amino acid changes.

Same region for nucleotide kmers:

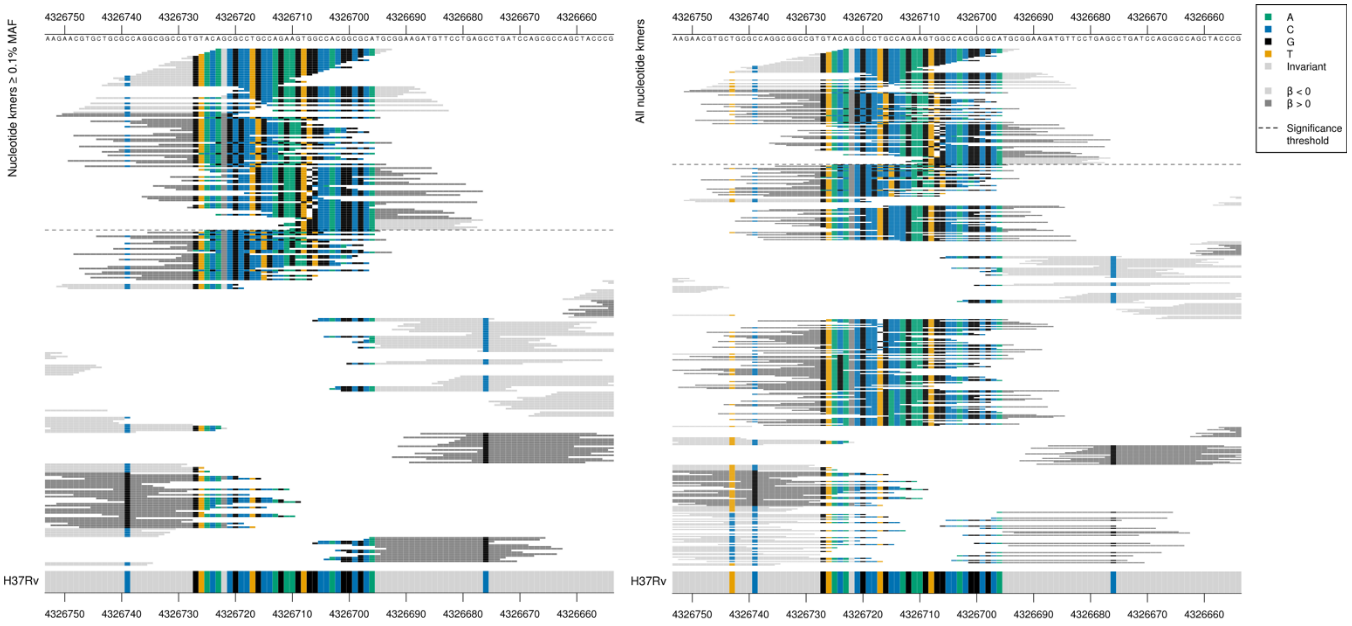

First set of high MIC-associated kmers deletion of a G between positions 4326707-4326706/767-768. Second set of high MIC-associated kmers, insertion of a G between positions 4326723-4326722/751-752.

| Kmer | $-\log_{10} p$ | β | MAC | MAF | Ps | Variants |
|---|---|---|---|---|---|---|
| SACQKWPRRMR | 29.20 | -1.43 | 92 | 1.11 | 251 | WT codons 243-264 |
| AACGTGCTGCGCCAGGCGGCCGTGTACAGGC | 9.23 | 1.53 | 18 | 0.22 | 4326750 | G insertion between positions 4326723-4326722/751-752 |
| TGCCAGAAGTGCCACGGCGCATGCGGAAGAT | 8.50 | 2.67 | 10 | 0.12 | 4326717 | G deletion between positions 4326707-4326706/767-768 |

Protein kmers:

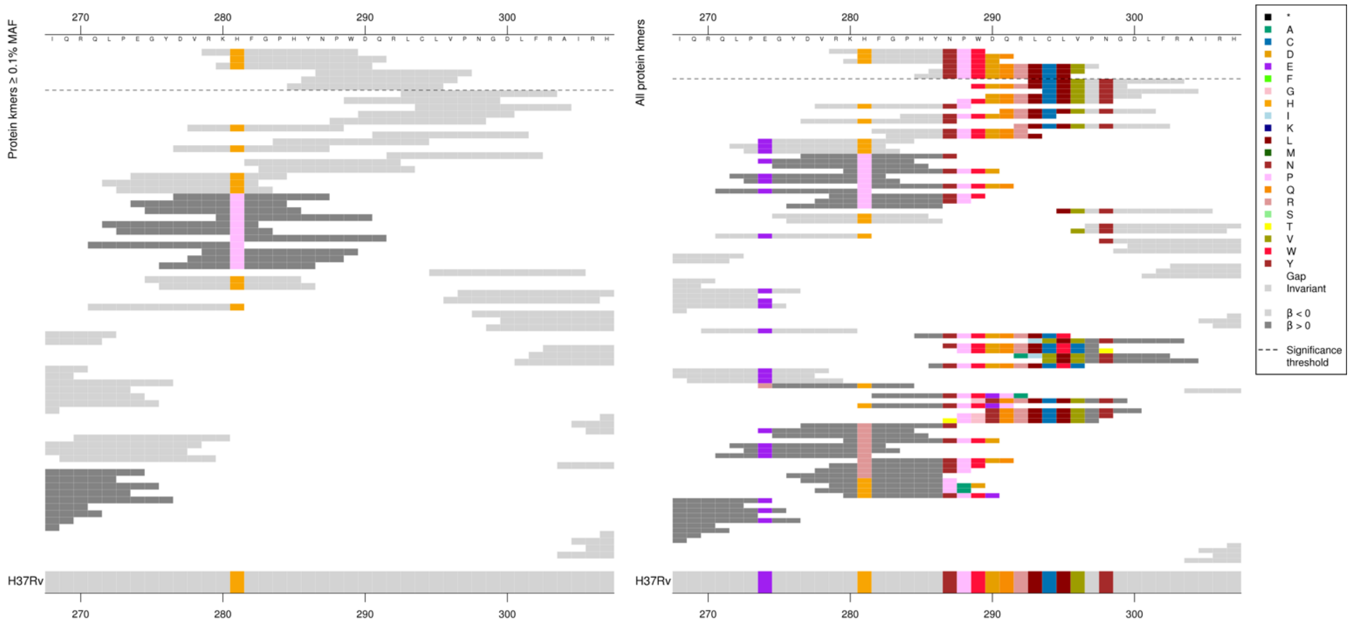

Same region for the nucleotide kmers:

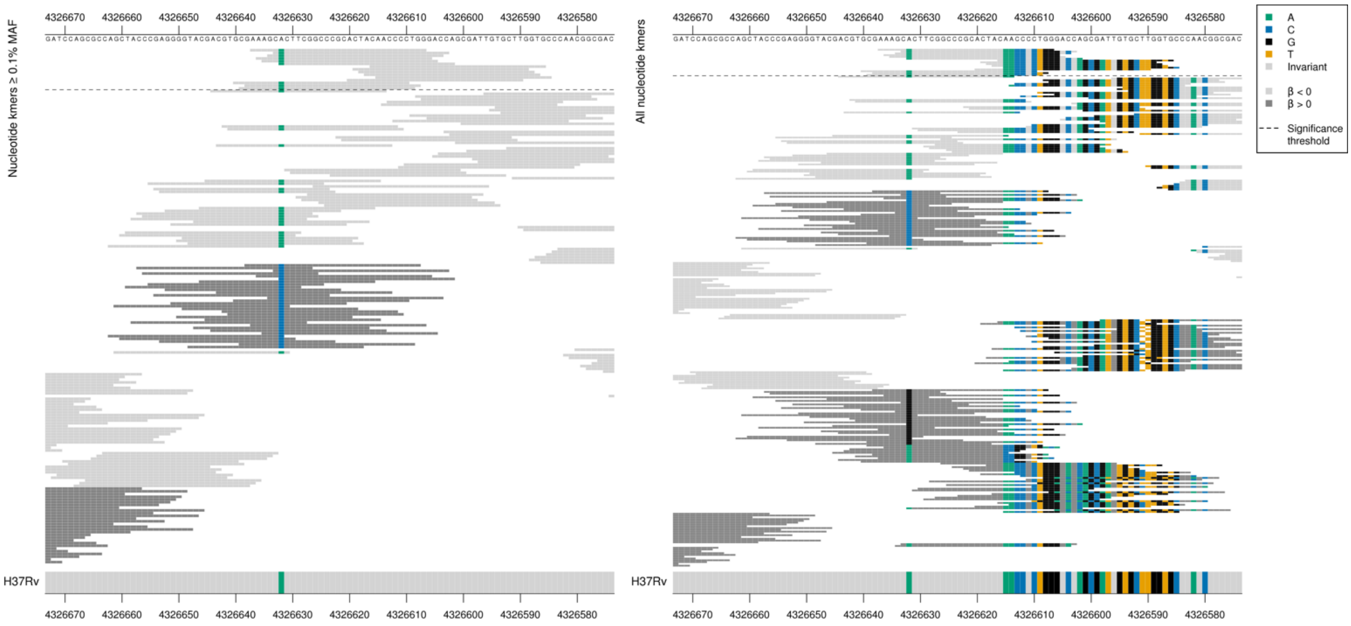

Low MIC-associated kmers were all wild-type covering the codons 279-297. The kmers that were not significant were likely to be capturing:

Amino acid 281P/R

T deletion between positions 4326591-4326590/883-884

A insertion between positions 4326606-4326604/868-870

| Kmer | $-\log_{10} p$ | β | MAC | MAF | Ps | Variants |
|---|---|---|---|---|---|---|
| RKHFGPHYNPW | 9.77 | -1.02 | 71 | 0.86 | 279 | WT codons 279-297 |

Protein kmers:

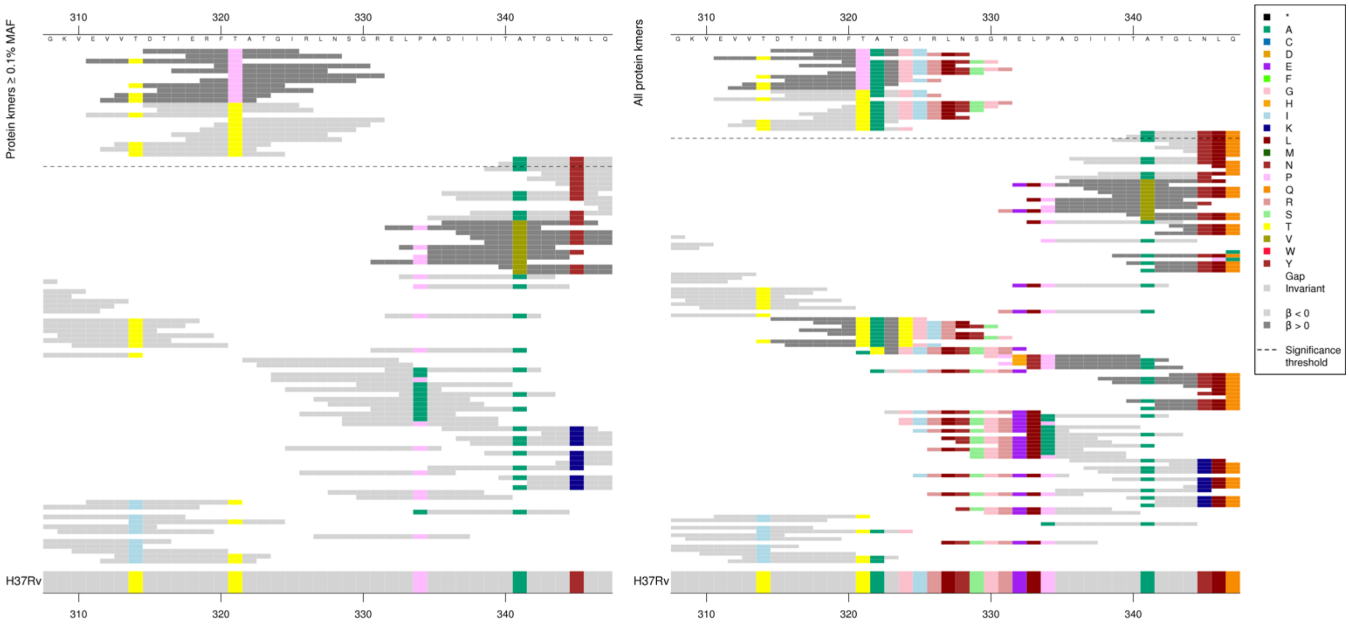

High MIC-associated kmers appear to only be capturing 321P as they were all the same *p*-value, the low MIC-associated kmers all had different *p*-values so could be affected by the other substitutions.

| Kmer | -log$_{10}$ *p* | β | MAC | MAF | Ps | Variants |
|---|---|---|---|---|---|---|
| DTIERFPATGI | 16.80 | 2.27 | 27 | 0.33 | 315 | 321P |
| DTIERFTATGI | 12.21 | -1.28 | 66 | 0.80 | 315 | 321T |

Protein kmers:

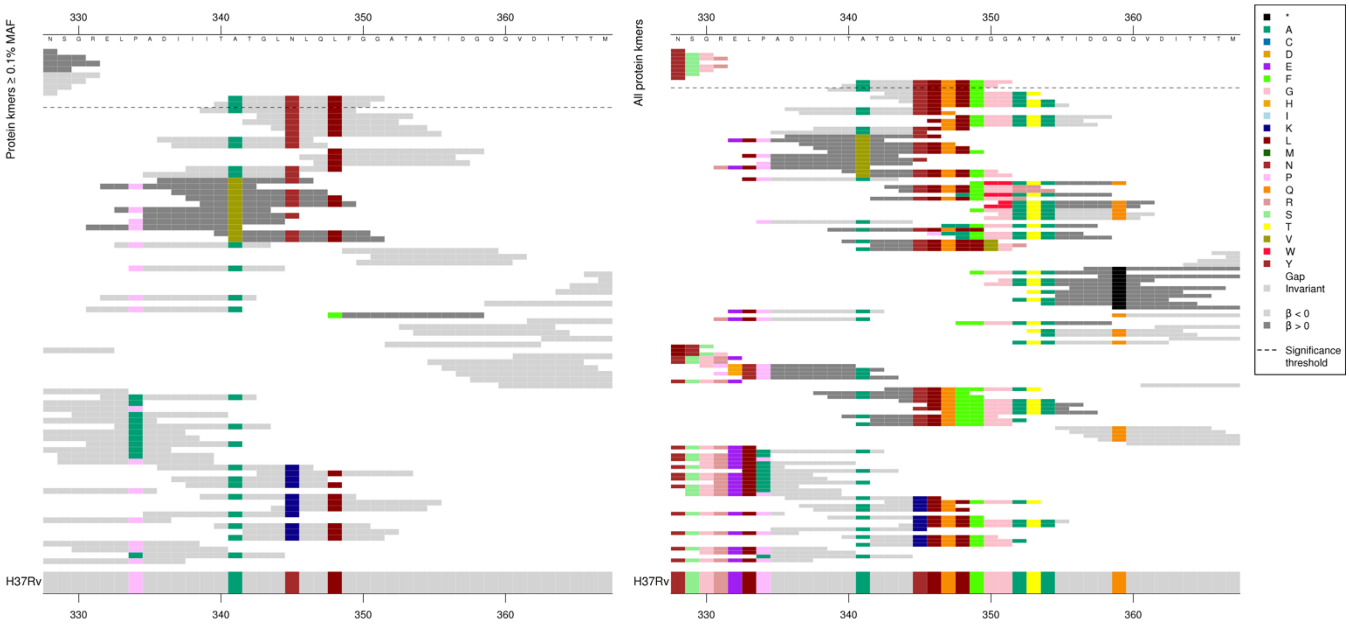

Kmers below significance threshold = 341V plus others.

| Kmer | -log$_{10}$ *p* | β | MAC | MAF | Ps | Variants |
|---|---|---|---|---|---|---|
| ATGLNLQLFGG | 8.01 | -1.03 | 178 | 2.15 | 341 | WT codons 340-351 |

#### *rpoB*

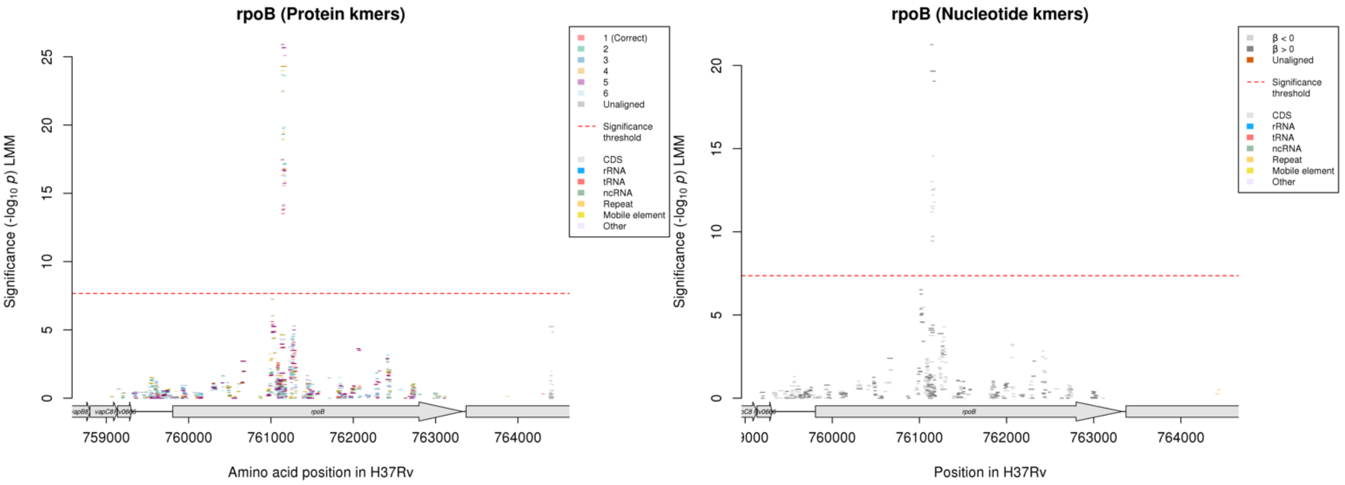

Protein kmers:

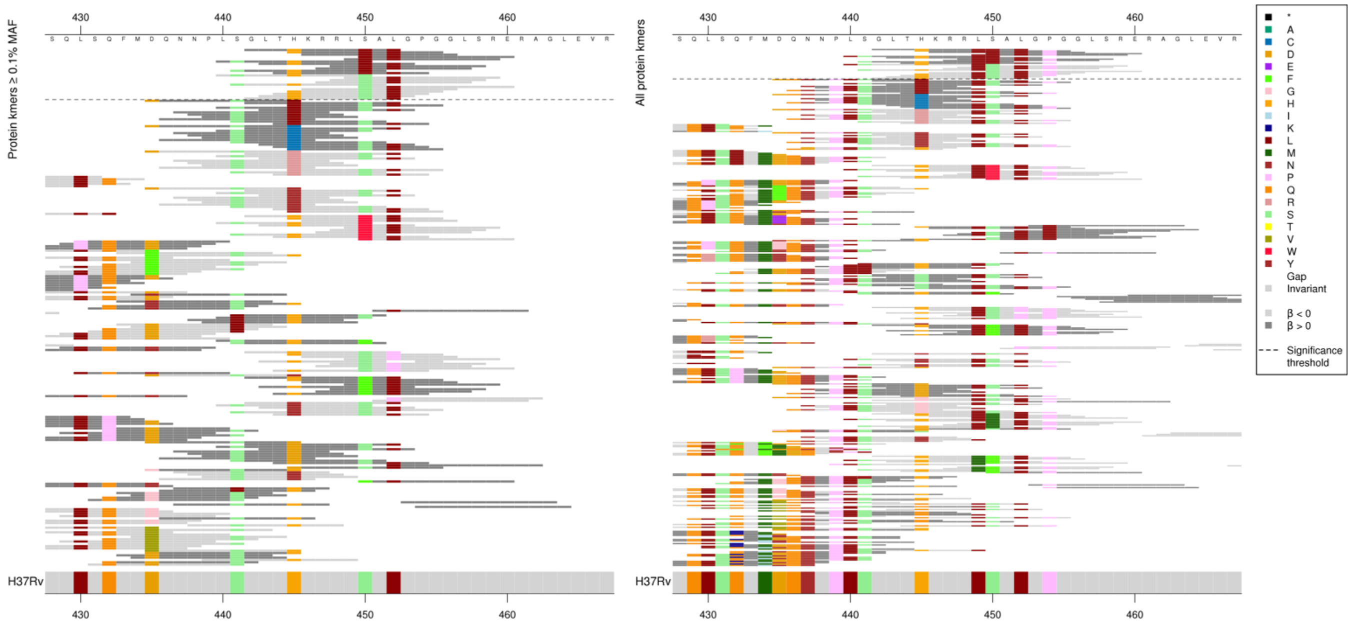

High MIC-associated kmers only capturing 450L as they were all the same *p*-value, the low MIC-associated had different *p*-values so the other substitutions could be affecting the significance.

| Kmer | -log10 *p* | β | MAC | MAF | Ps | Variants |
|---|---|---|---|---|---|---|
| GLTHKRRLLAL | 24.30 | 0.60 | 1986 | 23.97 | 442 | 450L |
| LSGLTHKRRLS | 17.43 | -0.45 | 2384 | 28.77 | 440 | 450S |

#### *gyrA*

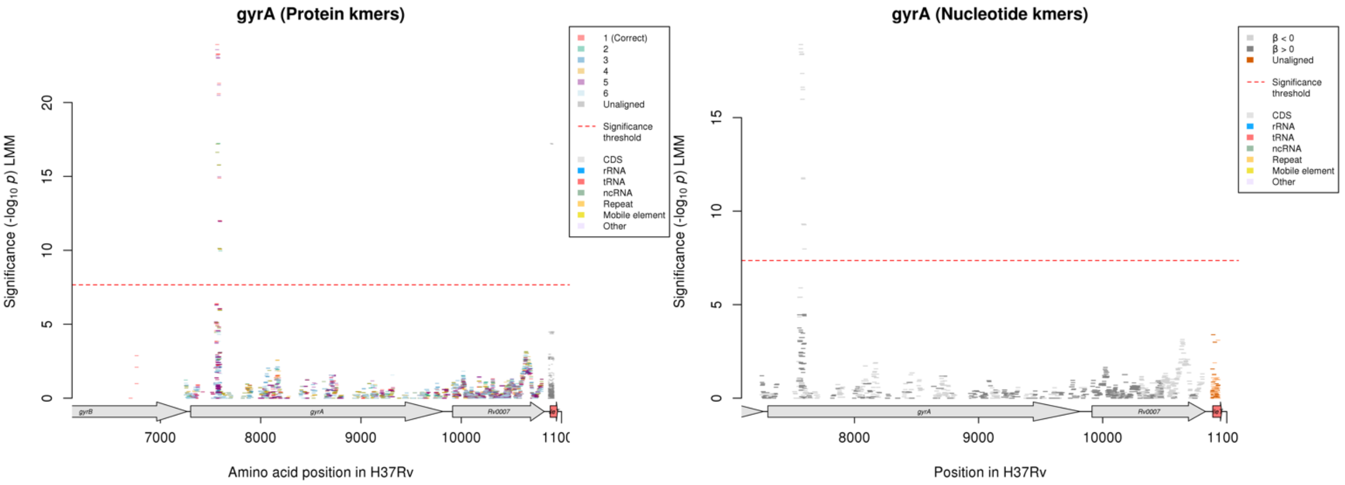

#### Protein kmers:

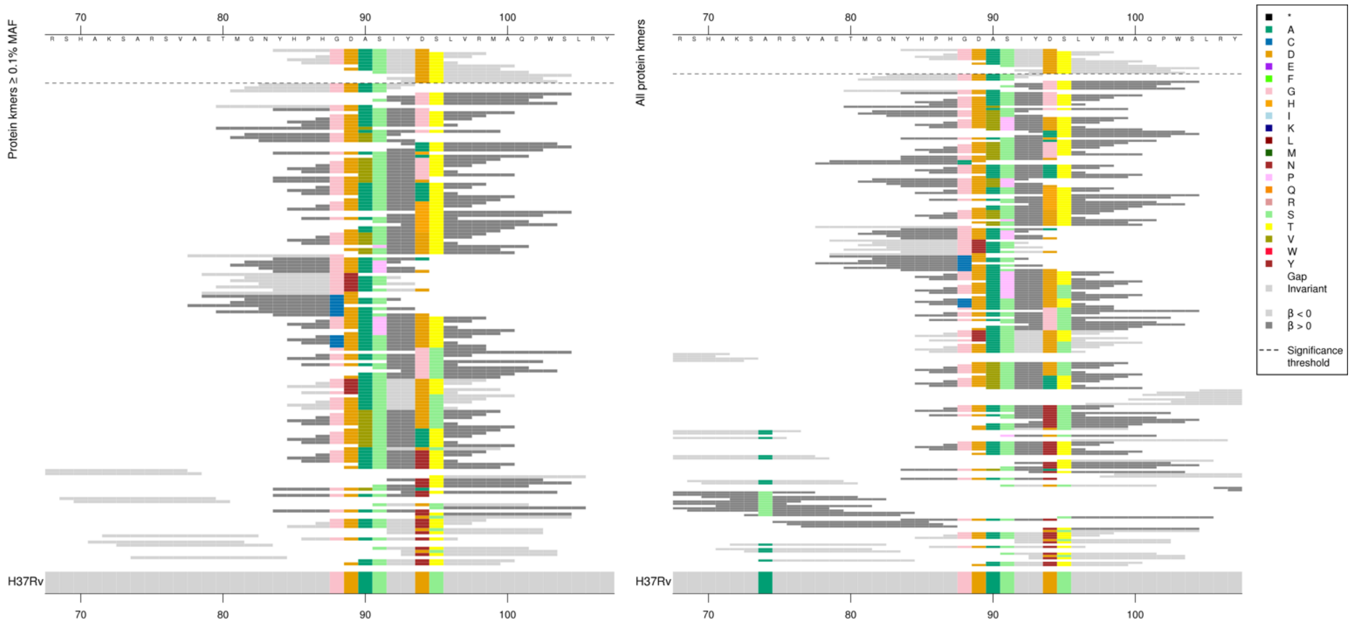

| Kmer | $-\log_{10} p$ | β | MAC | MAF | Ps | Variants |
|---|---:|---:|---:|---:|---:|---|
| YHPHGDASIYD | 23.92 | -0.54 | 1261 | 15.22 | 84 | 88G, 89D, 90A, 91S, 94D |
| GDASIYDTLVR | 23.26 | -0.55 | 1738 | 20.97 | 88 | 88G, 89D, 90A, 91S, 94D, 95T |
| ASIYDTLVRMA | 21.29 | -0.52 | 1716 | 20.71 | 90 | 90A, 91S, 94D, 95T |
| DASIYDTLVRM | 20.59 | -0.51 | 1724 | 20.80 | 89 | 89D, 90A, 91S, 94D, 95T |
| SIYDTLVRMAQ | 14.89 | -0.44 | 1402 | 16.92 | 91 | 91S, 94D, 95T |
| DTLVRMAQPWS | 11.99 | -0.41 | 1340 | 16.17 | 94 | 94D, 95T |

### *inhA*

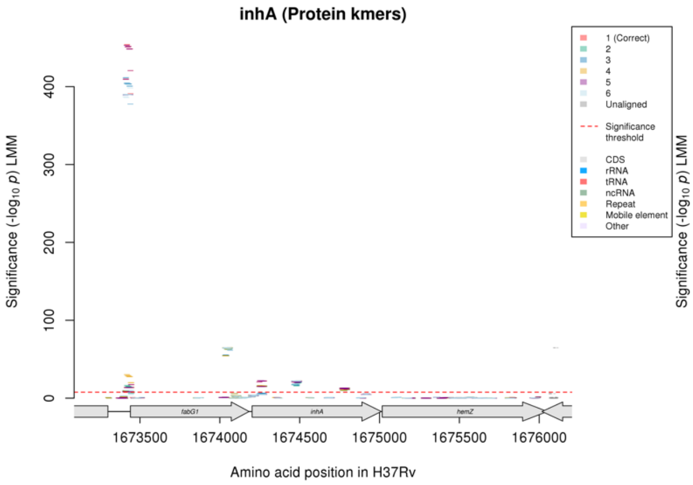

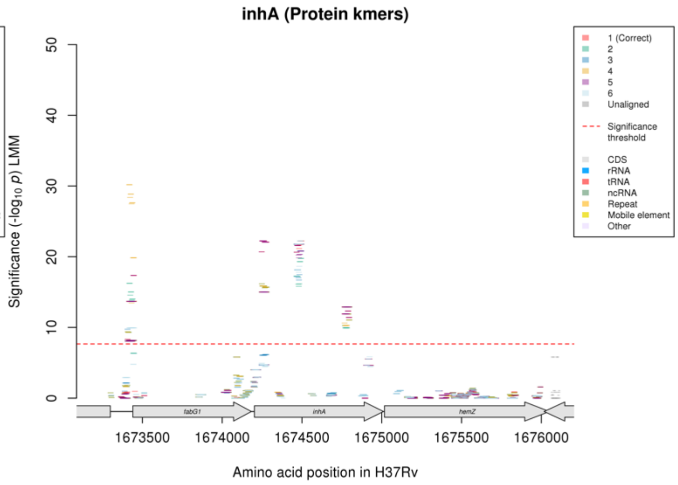

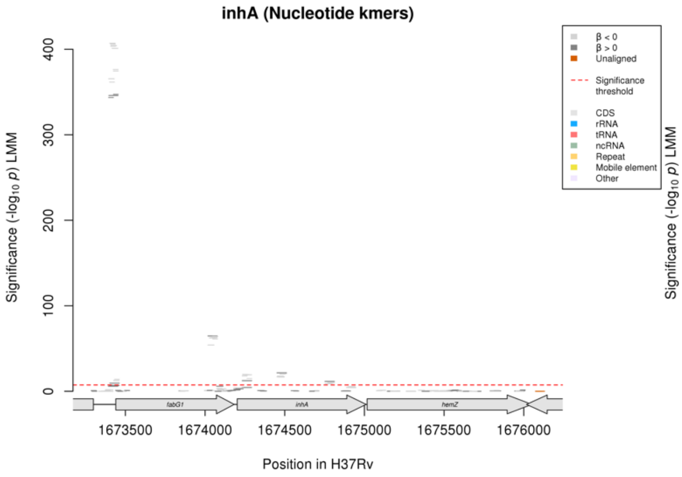

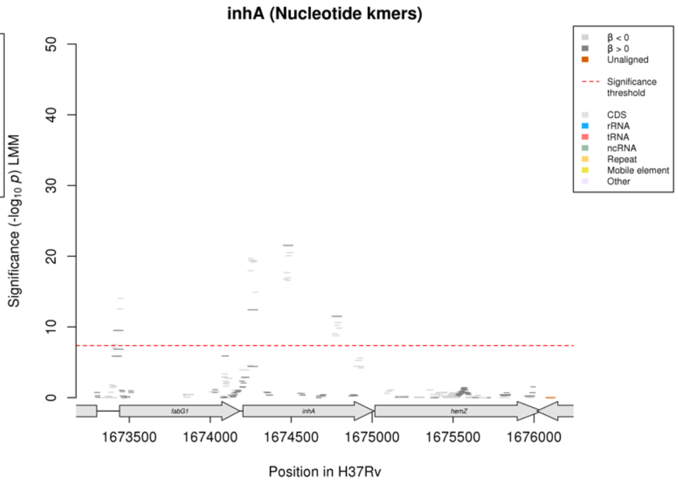

#### Protein kmers:

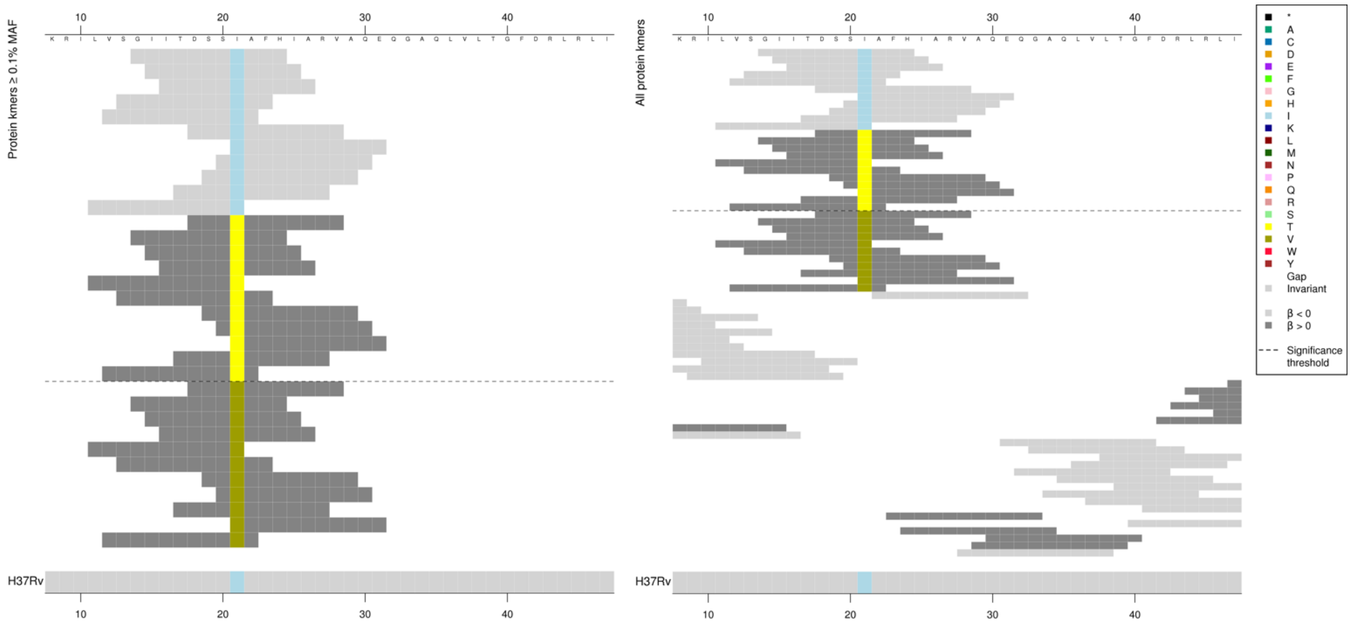

Kmers below significance threshold = 21V

| Kmer | $-\log_{10} p$ | β | MAC | MAF | Ps | Variants |
|---|---|---|---|---|---|---|
| GIITDSSIAFH | 22.24 | -1.55 | 68 | 0.82 | 14 | 21I |
| DSSTAFHIARV | 15.00 | 1.50 | 53 | 0.64 | 18 | 21T |

#### Protein kmers:

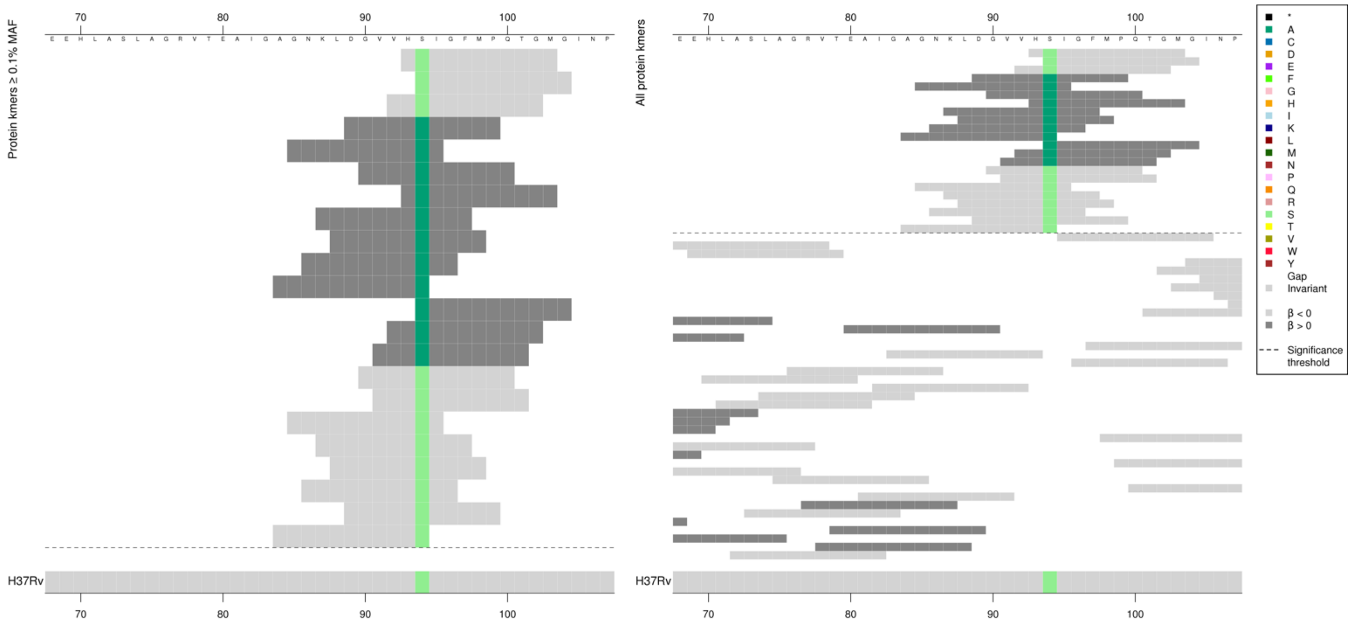

| Kmer | $-\log_{10} p$ | β | MAC | MAF | Ps | Variants |
|---|---|---|---|---|---|---|
| HSIGFMPQTGM | 22.23 | -2.14 | 34 | 0.41 | 93 | 94S |
| DGVVHAIGFMP | 21.76 | 2.19 | 30 | 0.36 | 89 | 94A |

#### Protein kmers:

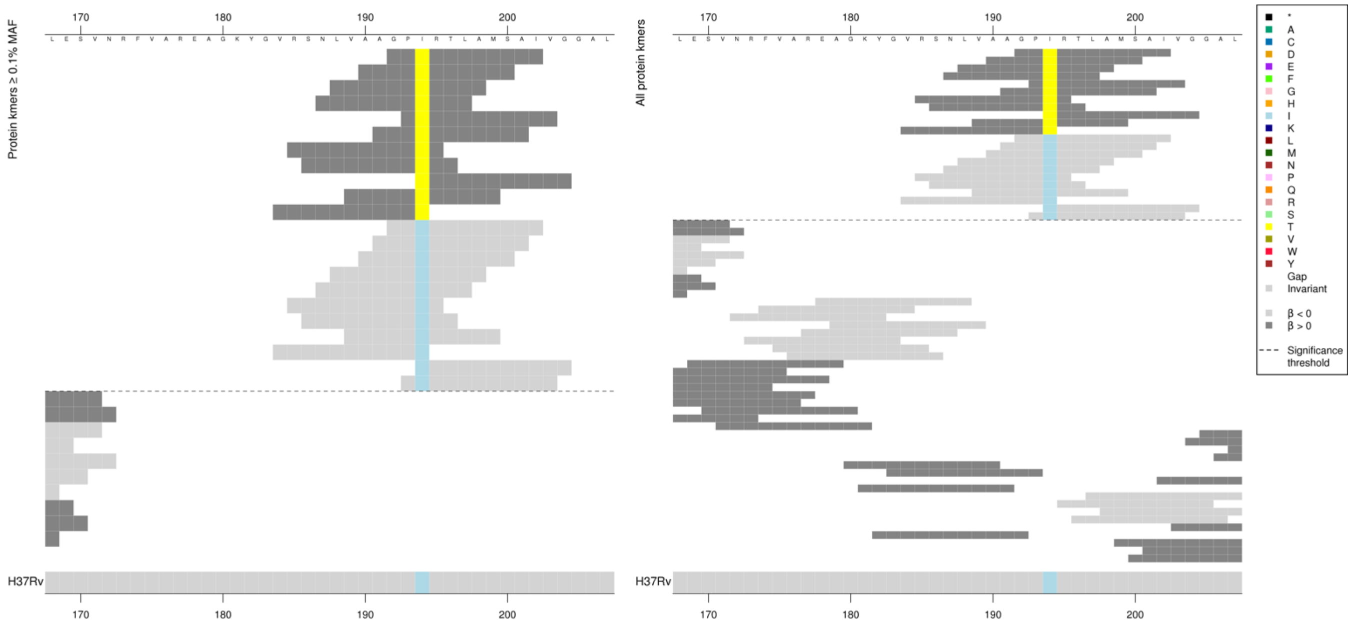

| Kmer | $-\log_{10} p$ | β | MAC | MAF | Ps | Variants |
|---|---|---|---|---|---|---|
| GPTRTLAMSAI | 12.89 | 1.53 | 53 | 0.64 | 192 | 194T |
| GPIRTLAMSAI | 12.32 | -1.51 | 53 | 0.64 | 192 | 194I |

#### *whiB7*

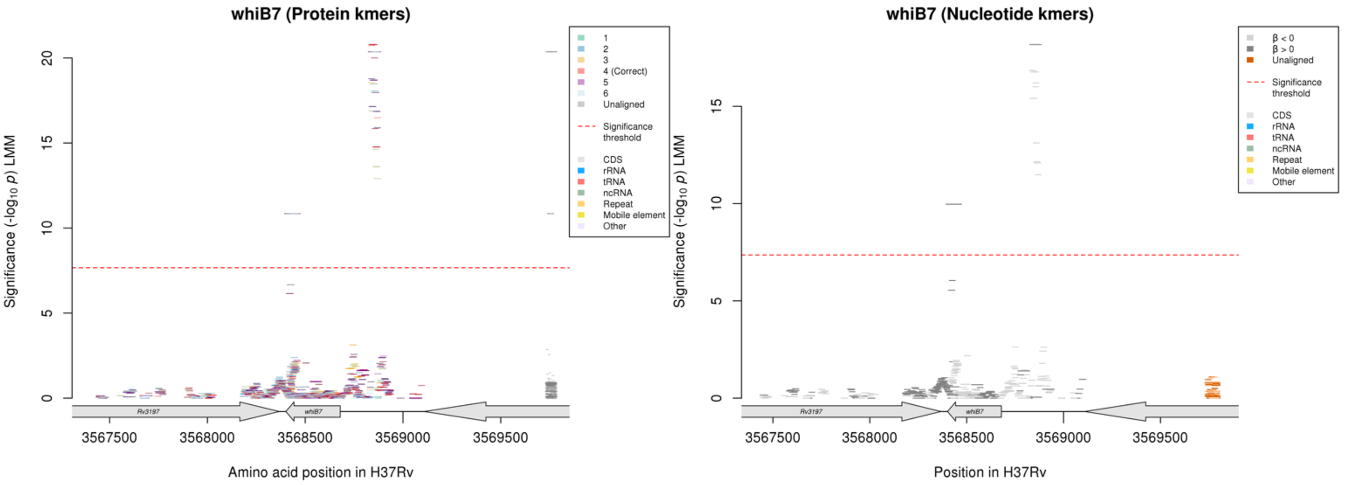

Nucleotide kmers:

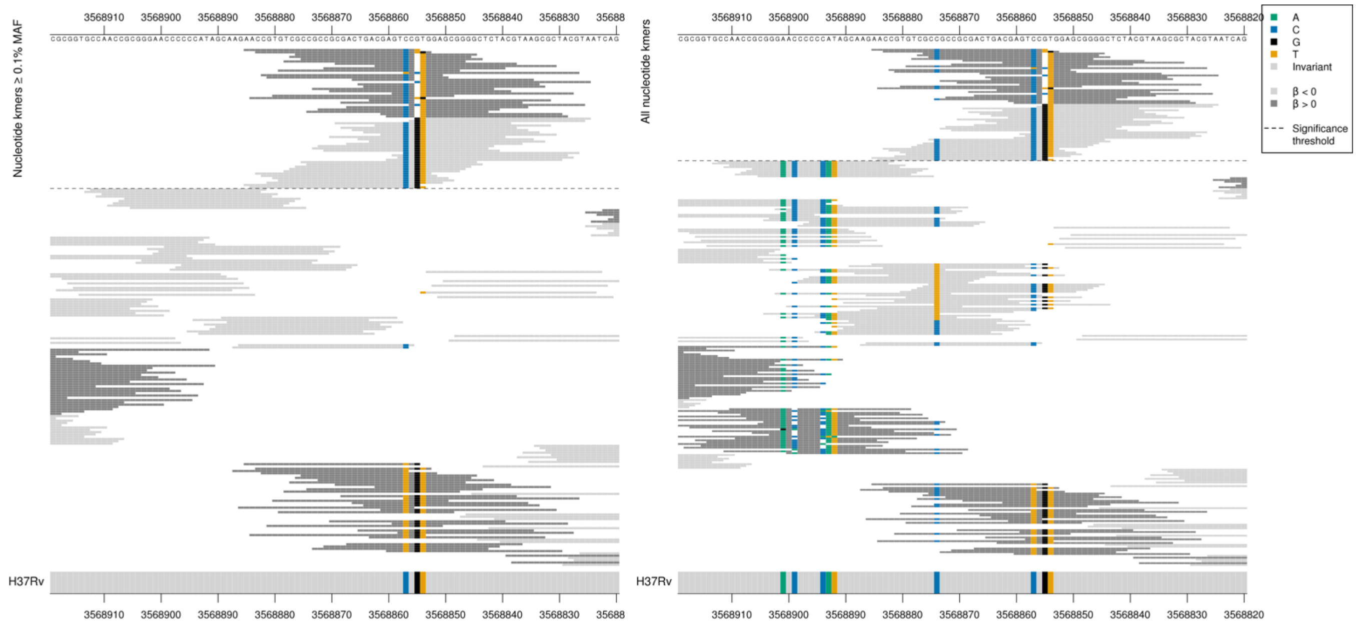

Upstream intergenic region – *whiB7* begins at position 3568679.
Capturing a high MIC-associated deletion of the G at position 3568855, 177bp upstream of the start codon for *whiB7*.
Low MIC-associated kmers were all the same as the reference.

| Kmer | $-\log_{10} p$ | β | MAC | MAF | Ps | Variants |
|---|---|---|---|---|---|---|
| AACCGTGTCGCCGCCGCGACTGACGAGTCCT | 18.18 | 2.16 | 46 | 0.56 | 3568885 | G deletion at position 3568855/-177 whiB7 |
| GTGGAGCGGGGCTCTACGTAAGCGCTACGTA | 16.84 | -1.83 | 54 | 0.65 | 3568855 | WT / 3568855G |

Protein kmers:

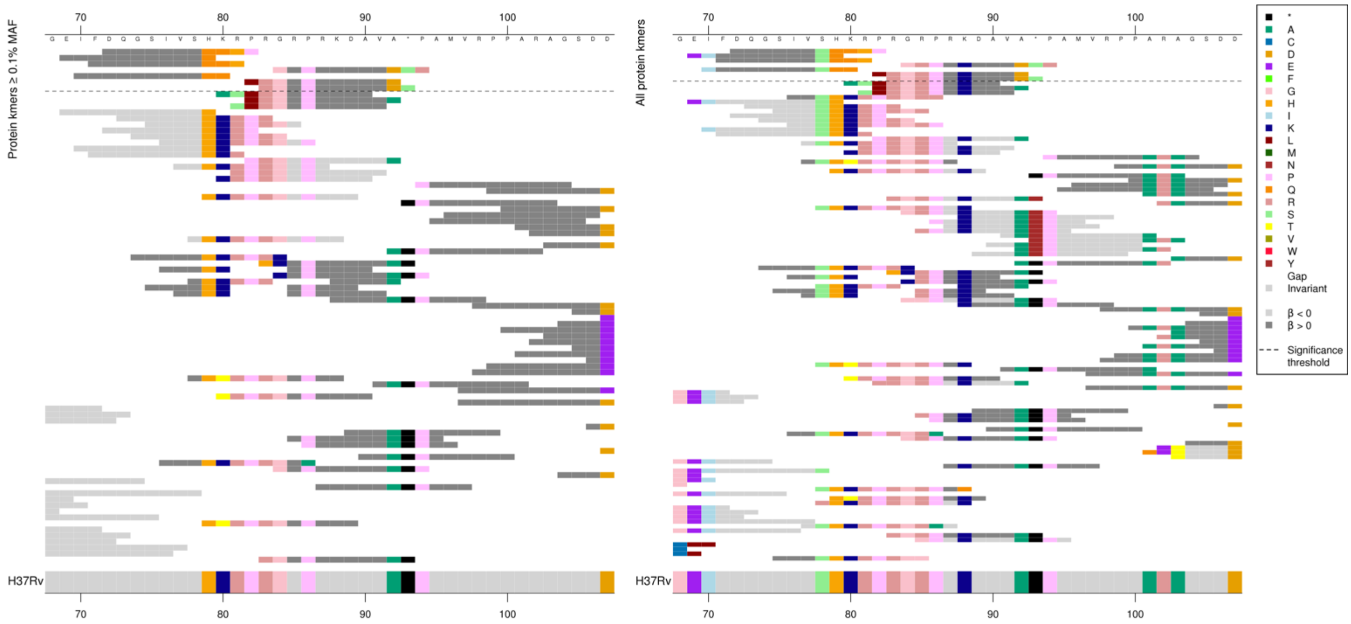

Same region for nucleotide kmers:

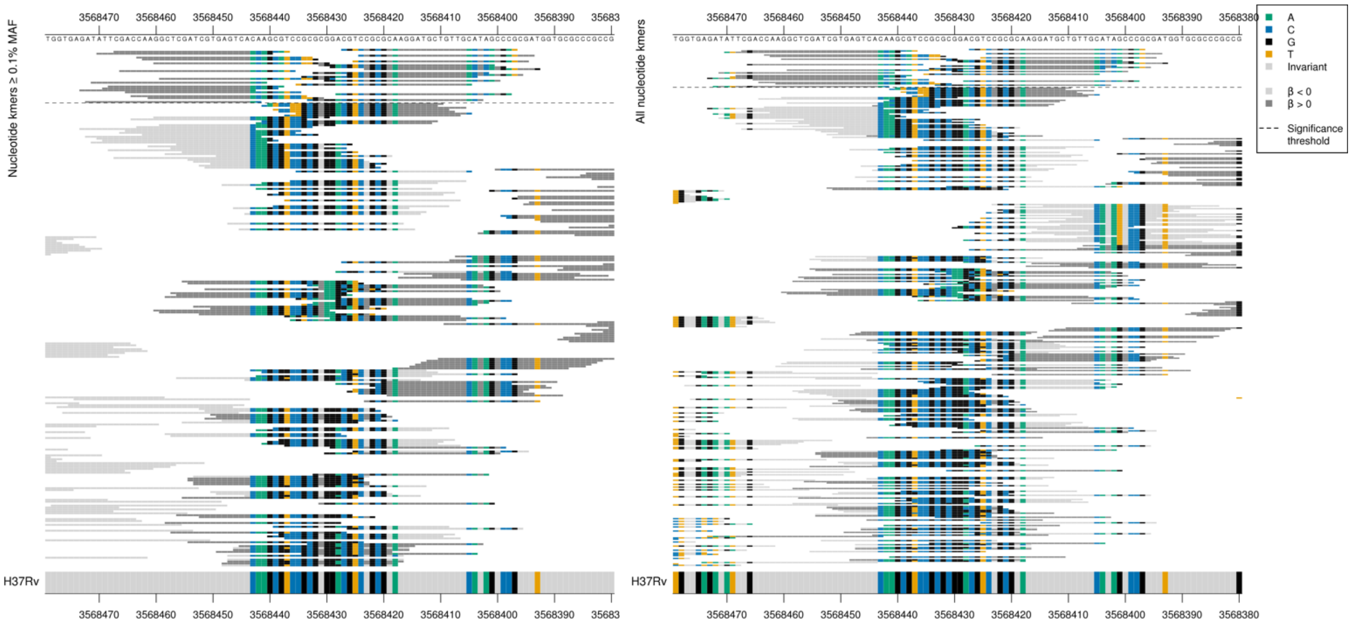

Gene ends at position 3568401. All 28 high MIC-associated had the same *p*-value, all have a long region of identity to the reference and then a region of many differences.

| Kmer | $-\log_{10} p$ | β | MAC | MAF | Ps | Variants |
|---|---|---|---|---|---|---|
| DQGSIVSQQHP | 10.85 | 1.96 | 22 | 0.27 | 72 | Multiple differences to reference at end of gene from codon 79 and overlapping stop codon |

### *PPE3*

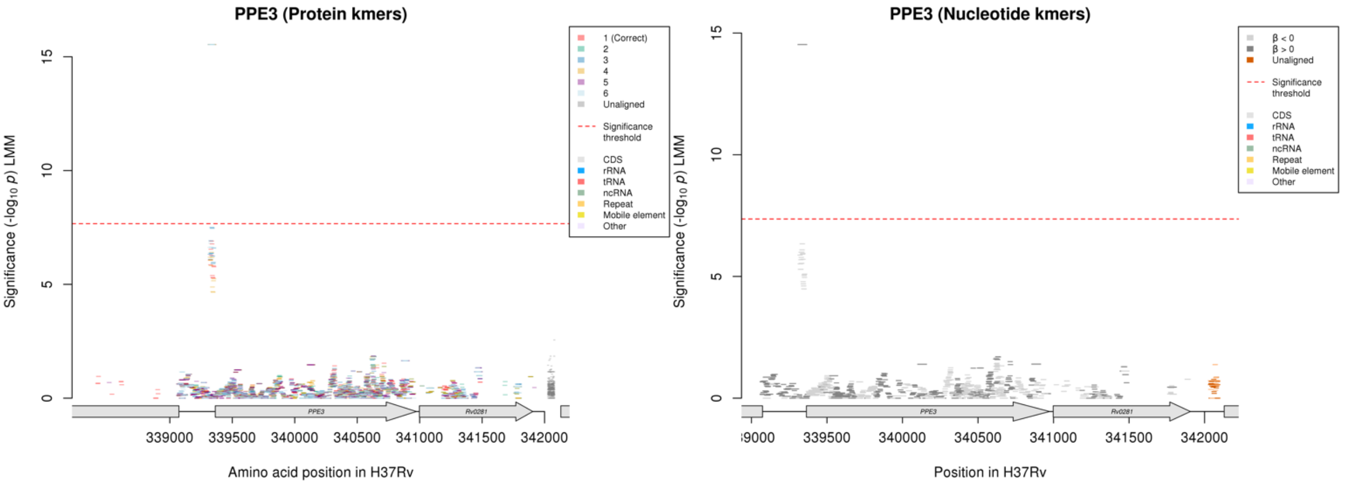

Nucleotide kmers:

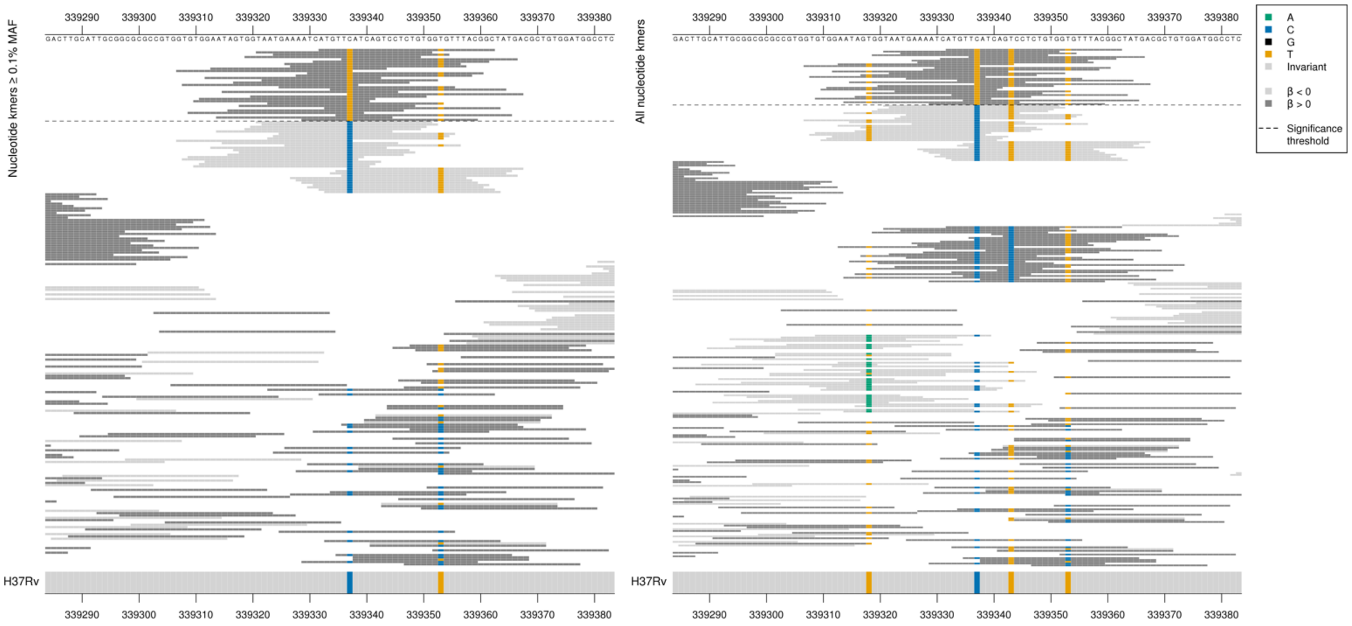

*PPE3* starts at position 339364.

Capturing a mutation at position 339337/-27.

Kmers below significance threshold = 339337/-27C.

| Kmer | $-\log_{10} p$ | β | MAC | MAF | Ps | Variants |
|---|---|---|---|---|---|---|
| ATGTTTATCAGTCCTCTGTGGTGTTTACGGC | 14.53 | 2.02 | 29 | 0.35 | 339332 | 339337/-27T |

#### *mpt53*

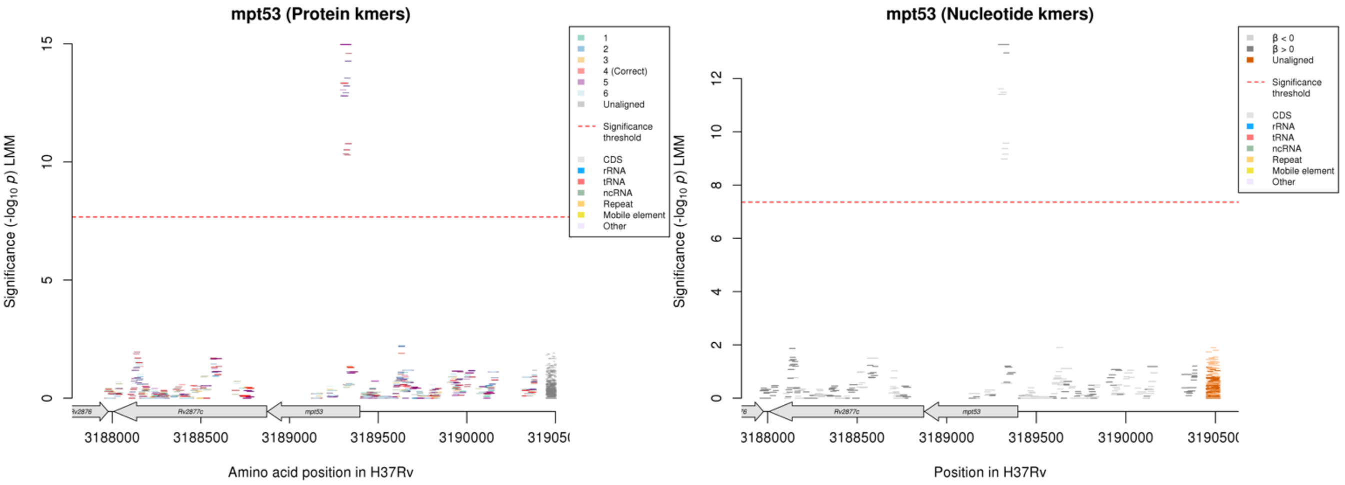

Protein kmers:

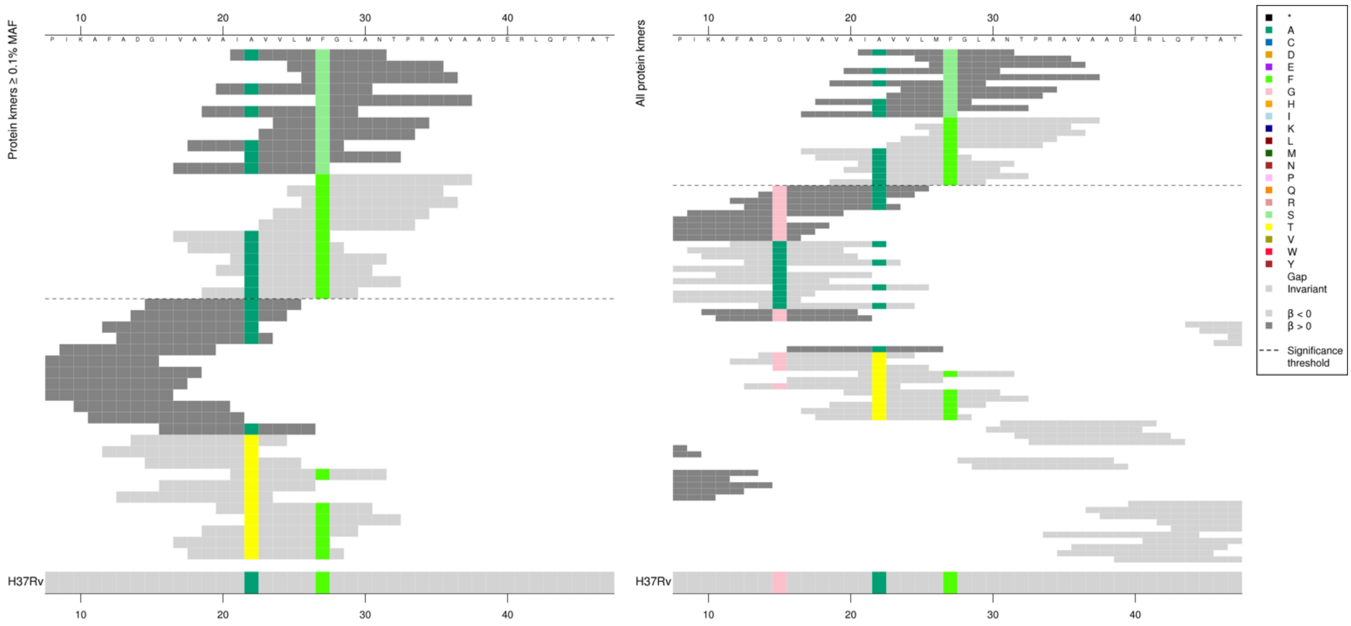

Main codon being captured is codon 27. An A at codon 22 was more significantly associated with low MICs than a T.

| Kmer | $-\log_{10} p$ | β | MAC | MAF | Ps | Variants |
|---|---|---|---|---|---|---|
| IAVVLMSGLAN | 14.97 | 1.96 | 50 | 0.60 | 21 | 27S |
| FGLANTPRAVA | 13.33 | -1.66 | 56 | 0.68 | 27 | 27F |
| VAVAIAVVLMF | 10.77 | -1.48 | 63 | 0.76 | 17 | 27F, 22A |

#### *embB*

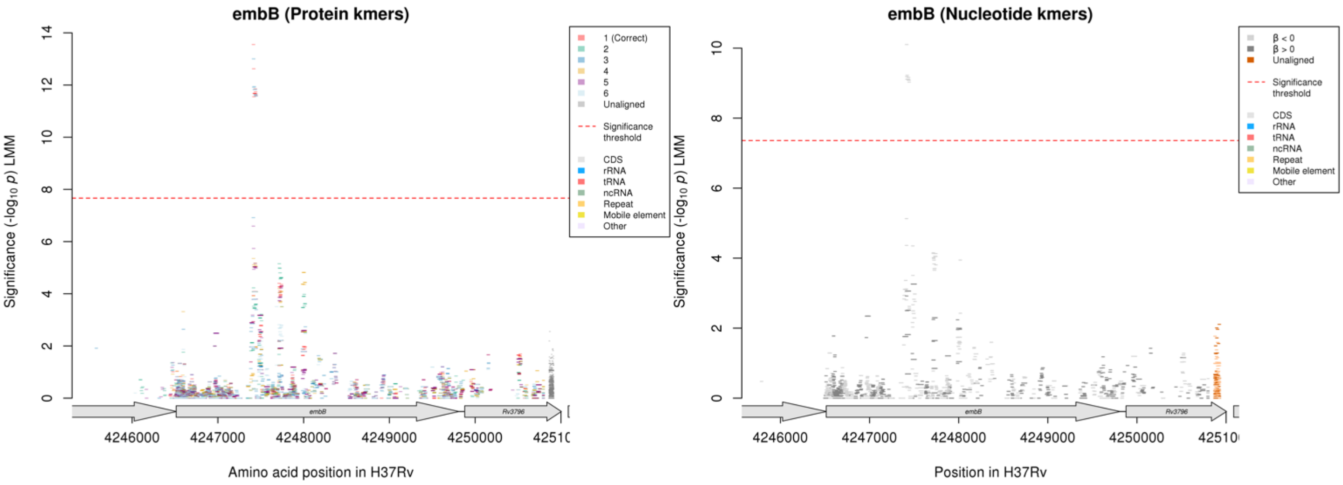

Protein kmers:

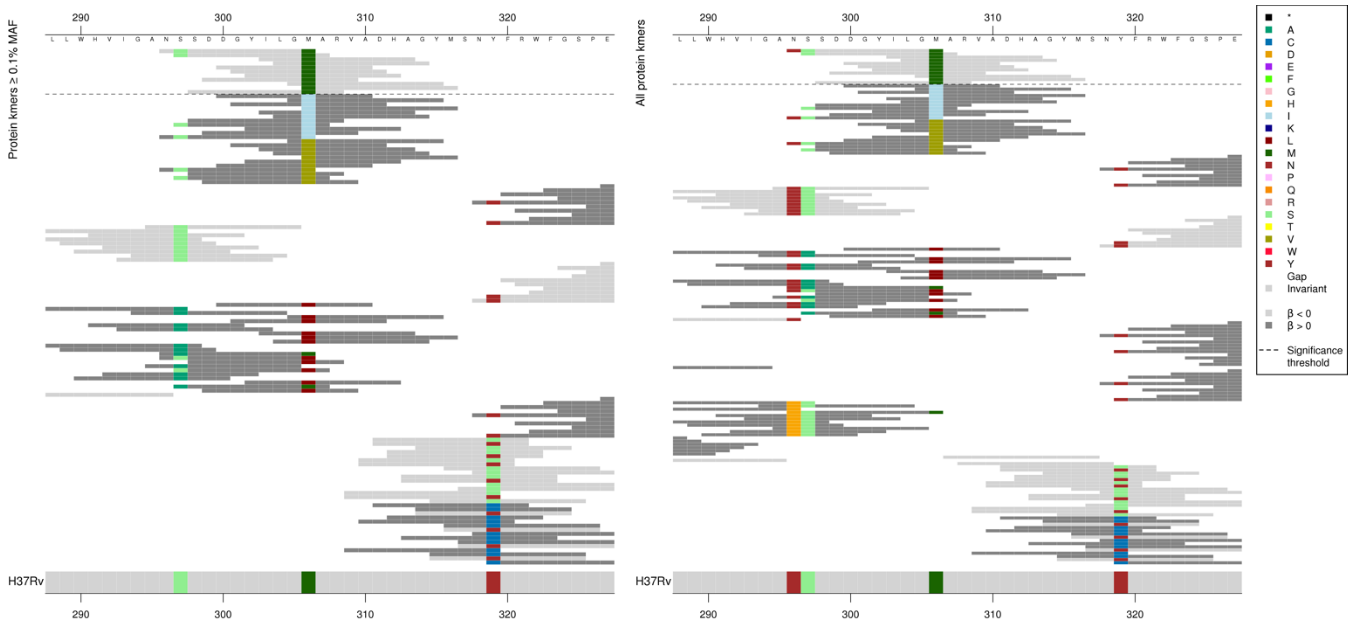

Low MIC-associated kmers mainly capturing 306M as that is the only position they all covered, but the kmers had different *p*-values depending on the other positions they covered.

Kmers below significance threshold = Codon 296H, 297A, 306I/V/L

| Kmer | $-\log_{10} p$ | β | MAC | MAF | Ps | Variants |
|---|---|---|---|---|---|---|
| NSSDDGYILGM | 13.55 | -0.43 | 1603 | 19.34 | 296 | 306M |

### *eccA1*

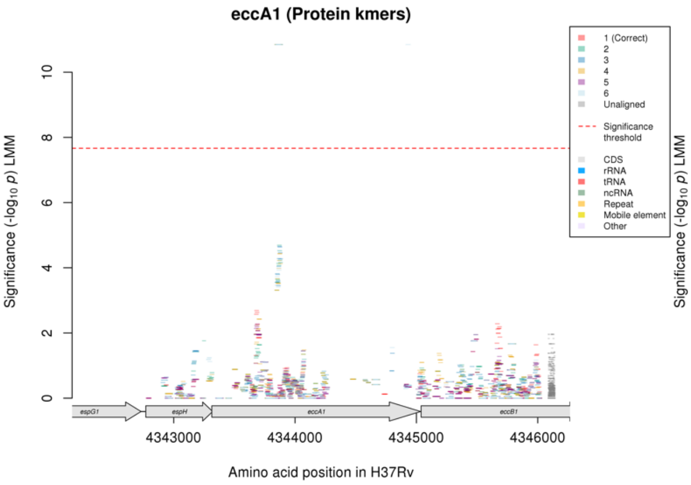

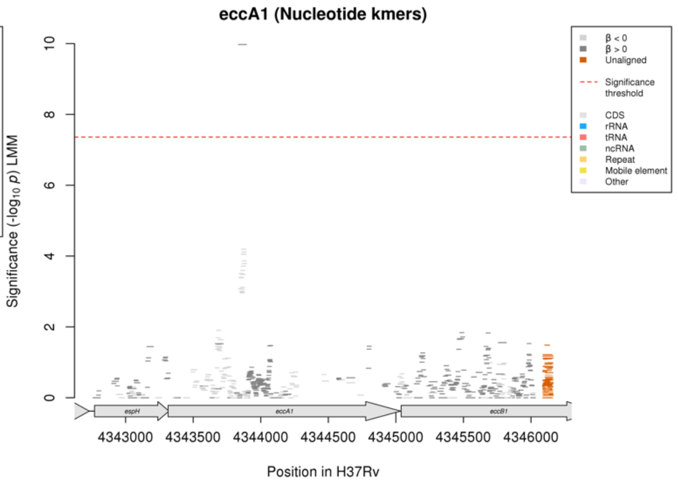

Nucleotide kmers:

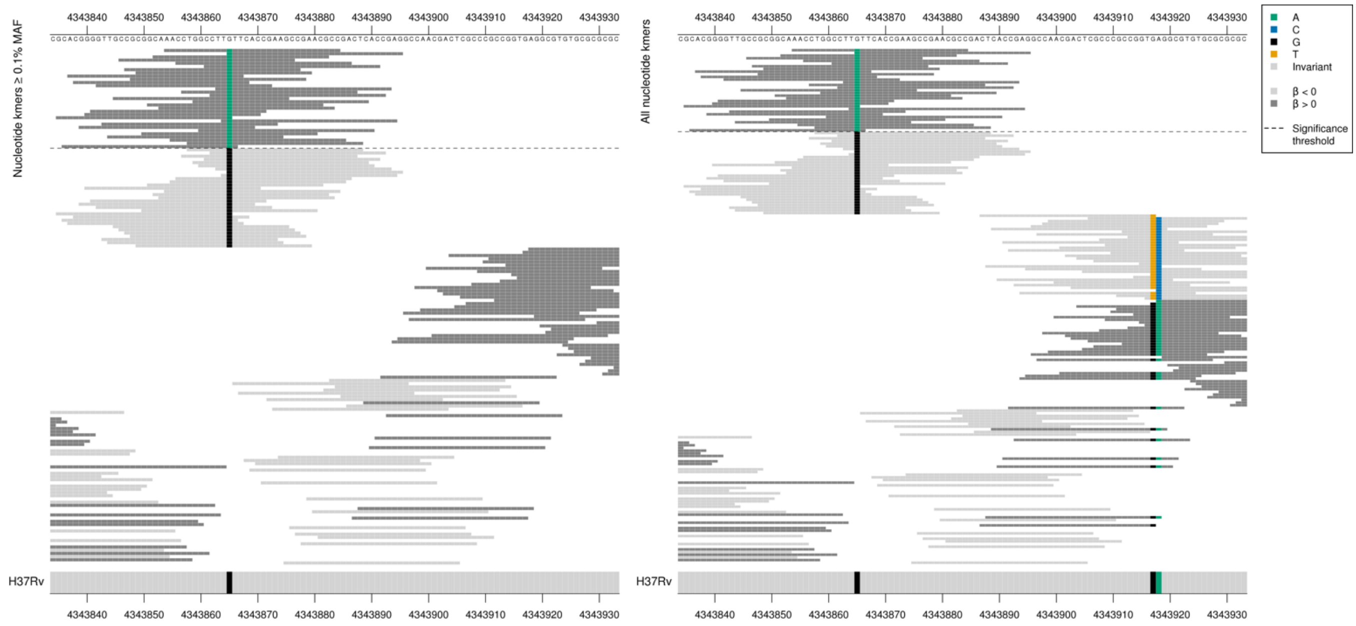

Capturing a synonymous mutation at position 4343865/552.
High MIC-associated significant kmers = TTA = Leu / L
Kmers below significance threshold = TTG = Leu / L

| Kmer | $-\log_{10} p$ | β | MAC | MAF | Ps | Variants |
|---|---|---|---|---|---|---|
| AACCTGGCCTTATTCACCGAAGCCGAACGCC | 9.97 | 1.85 | 22 | 0.27 | 4343854 | 4343865/552A |

#### embA

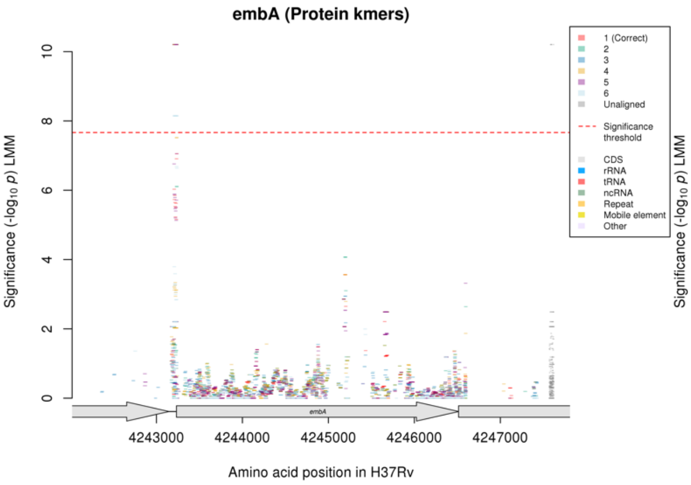

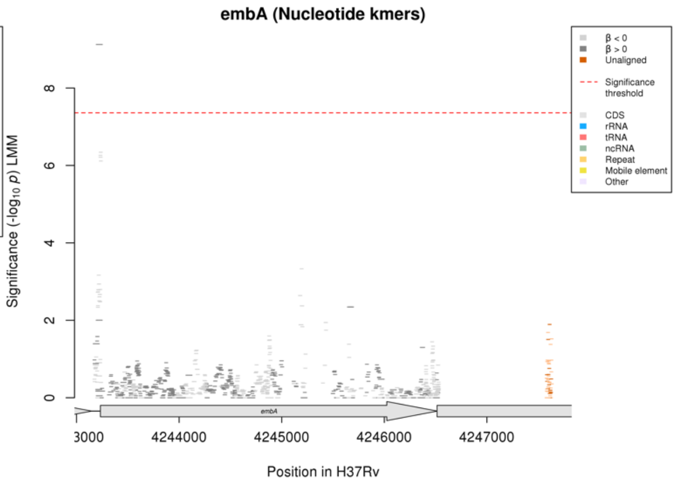

Nucleotide kmers:

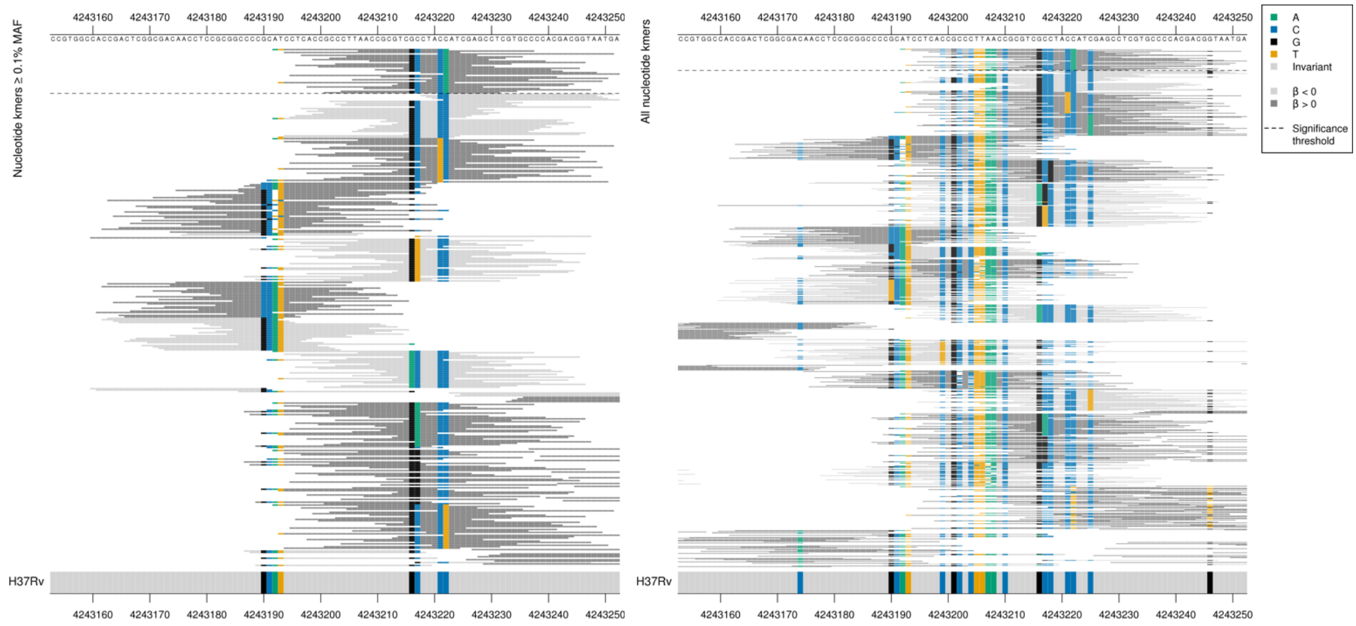

All high MIC-associated kmers were the same *p*-value, so seem to be capturing 4243222/-11A.

| Kmer | -log10 *p* | β | MAC | MAF | Ps | Variants |
|---|---|---|---|---|---|---|
| ATCCTCACCGCCCTTAACCGCGTCGCCTACA | 9.12 | 1.15 | 66 | 0.80 | 4243192 | 4243222/-11A |

## Rv0565c

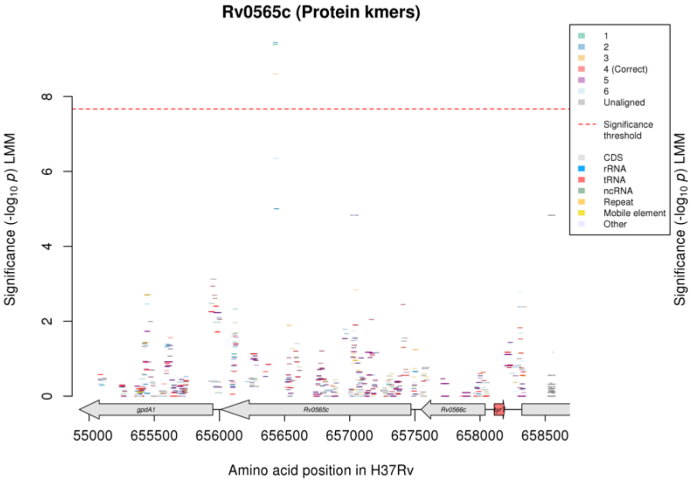

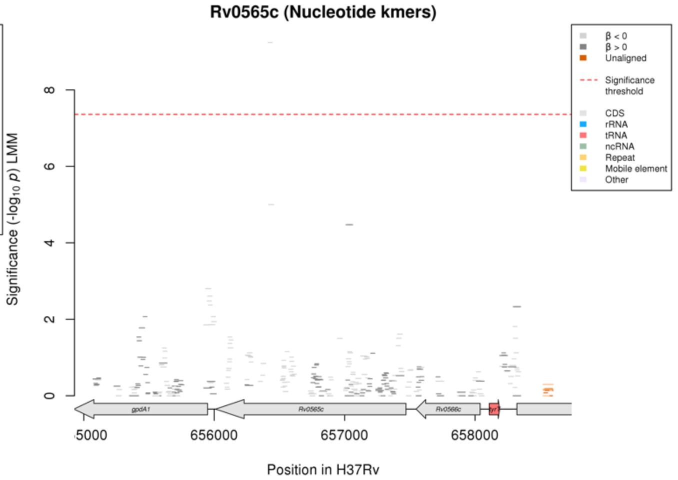

Nucleotide kmers:

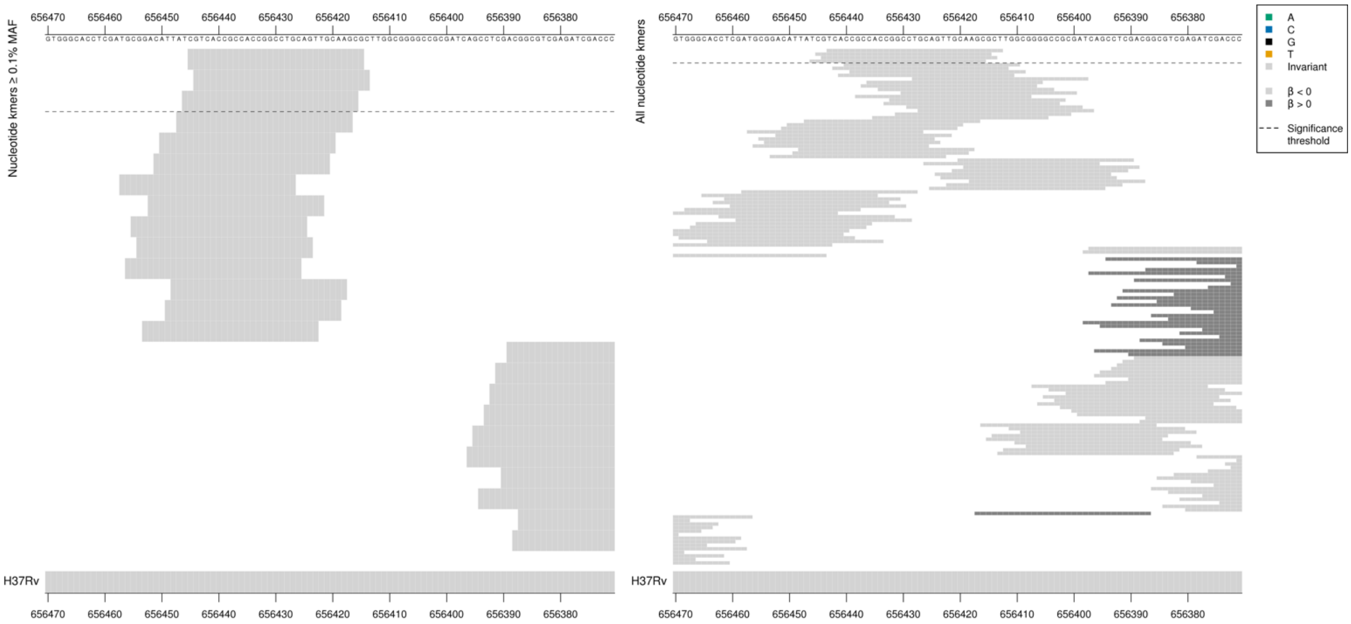

Removing the threshold requiring a kmer to be aligned to a gene/intergenic region in at least five genomes across the full dataset. Showing the nucleotide kmer alignment after lowering the threshold to just one genome:

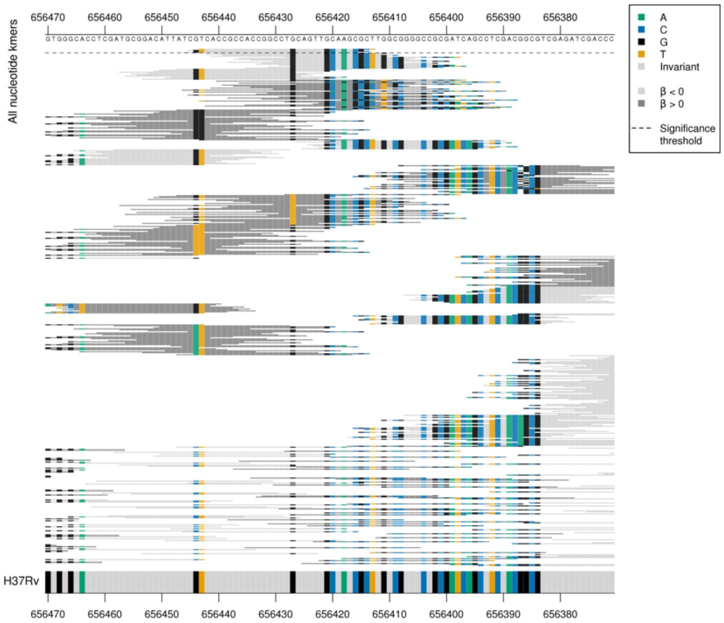

Low MIC-associated reference-sequence kmers covered a region with lots of rare alternative alleles.

| Kmer | $-\log_{10} p$ | β | MAC | MAF | Ps | Variants |
|---|---|---|---|---|---|---|
| CGTCACCGCCACCGGCCTGCAGTTGCAAGCG | 9.24 | -2.09 | 12 | 0.14 | 656445 | WT |

### *fadB4*

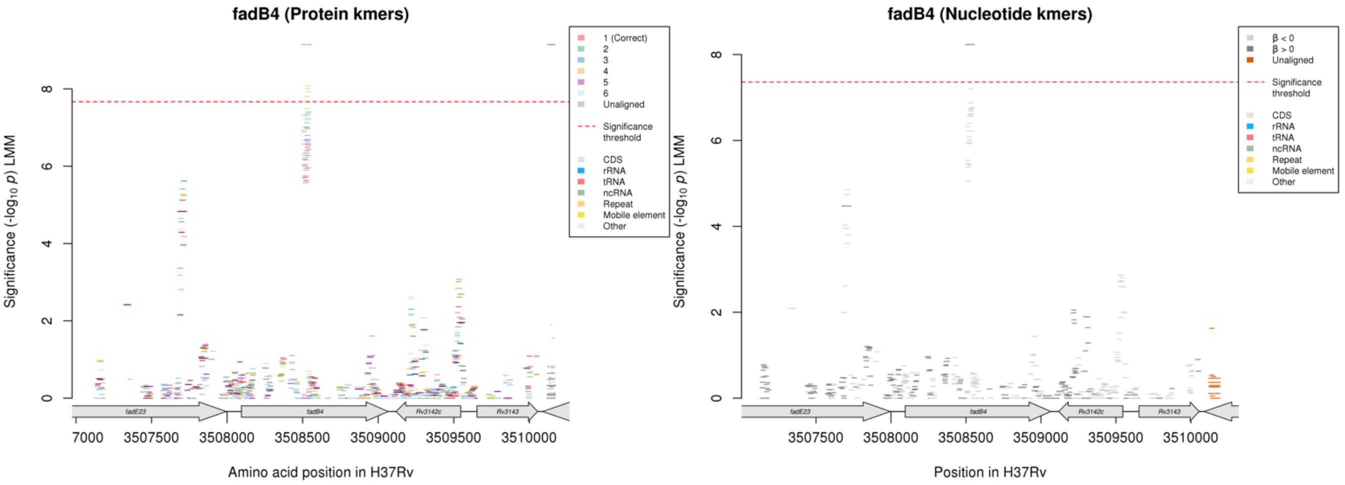

#### Protein kmers:

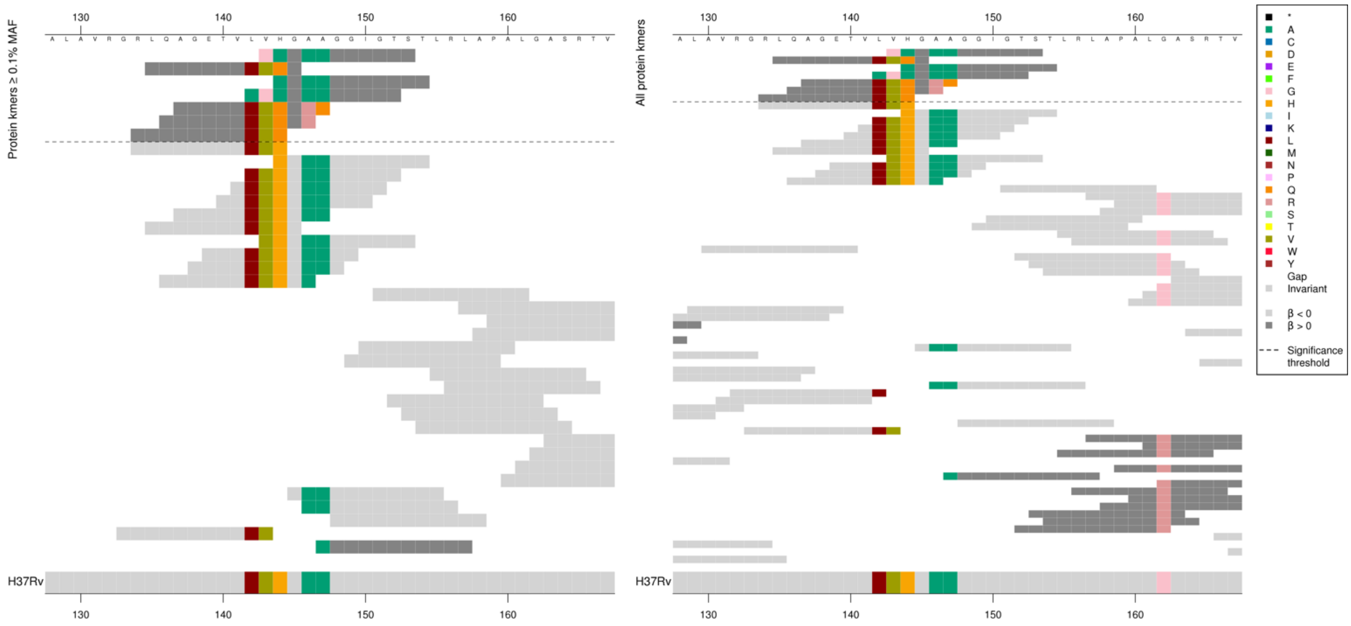

#### The same region for the nucleotide kmers:

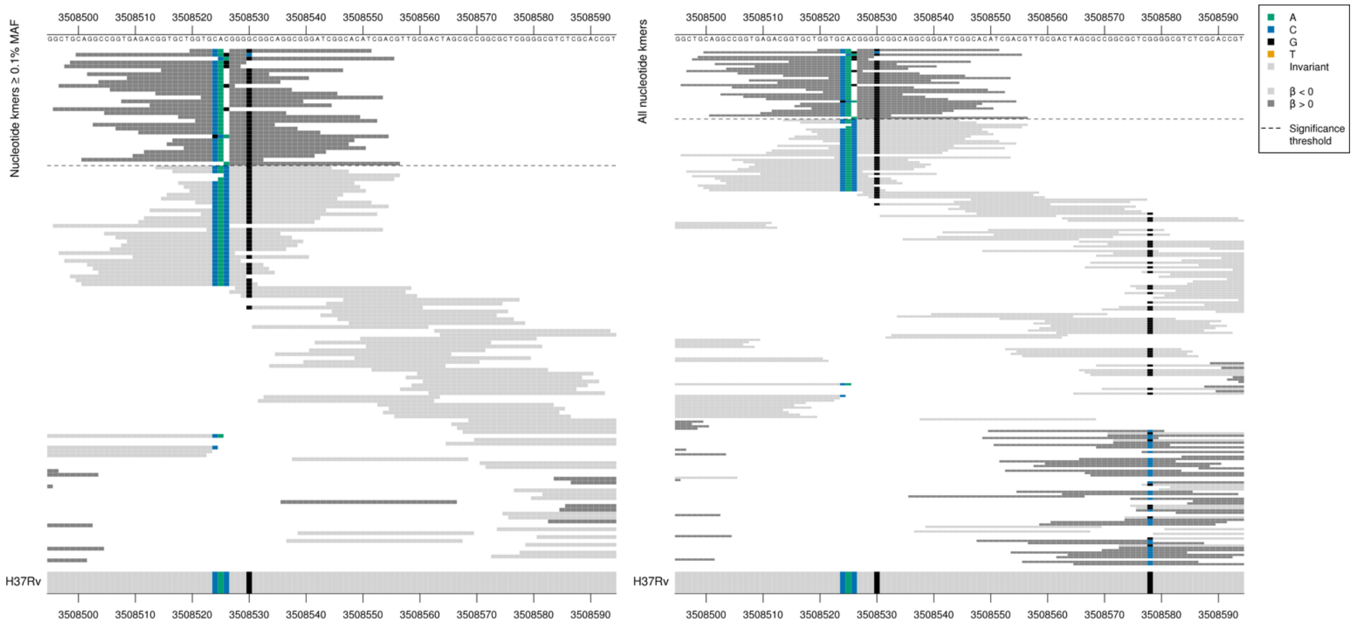

#### Deletion of a C at position 3508526/432.

| Kmer | -log$_{10}$ *p* | β | MAC | MAF | Ps | Variants |
|---|---|---|---|---|---|---|
| GGTGCAGGGGCGGCAGGCGGGATCGGCACAT | 8.23 | 2.02 | 10 | 0.12 | 3508520 | C deletion at position 3508526/432 |

### *plsC*

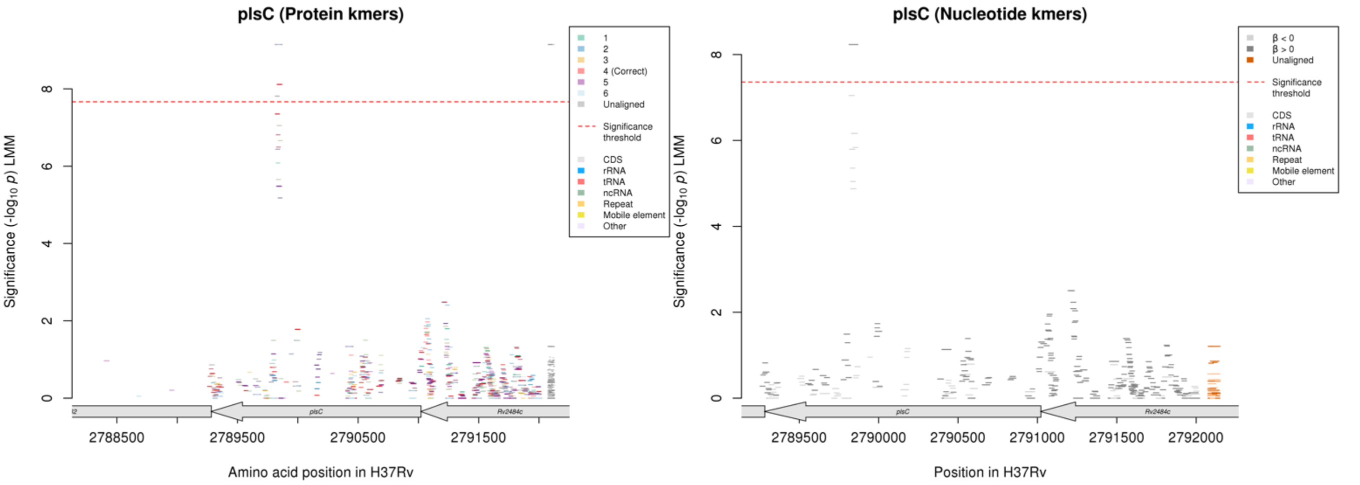

Protein kmers:

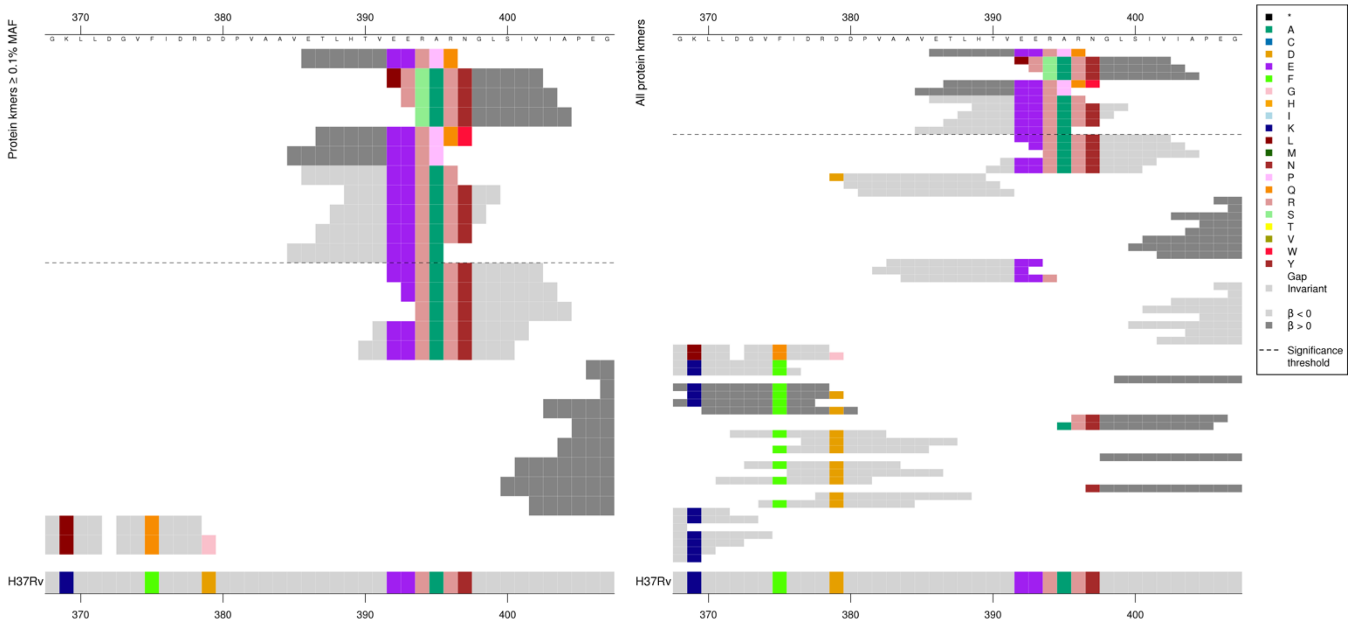

Same region for nucleotide kmers:

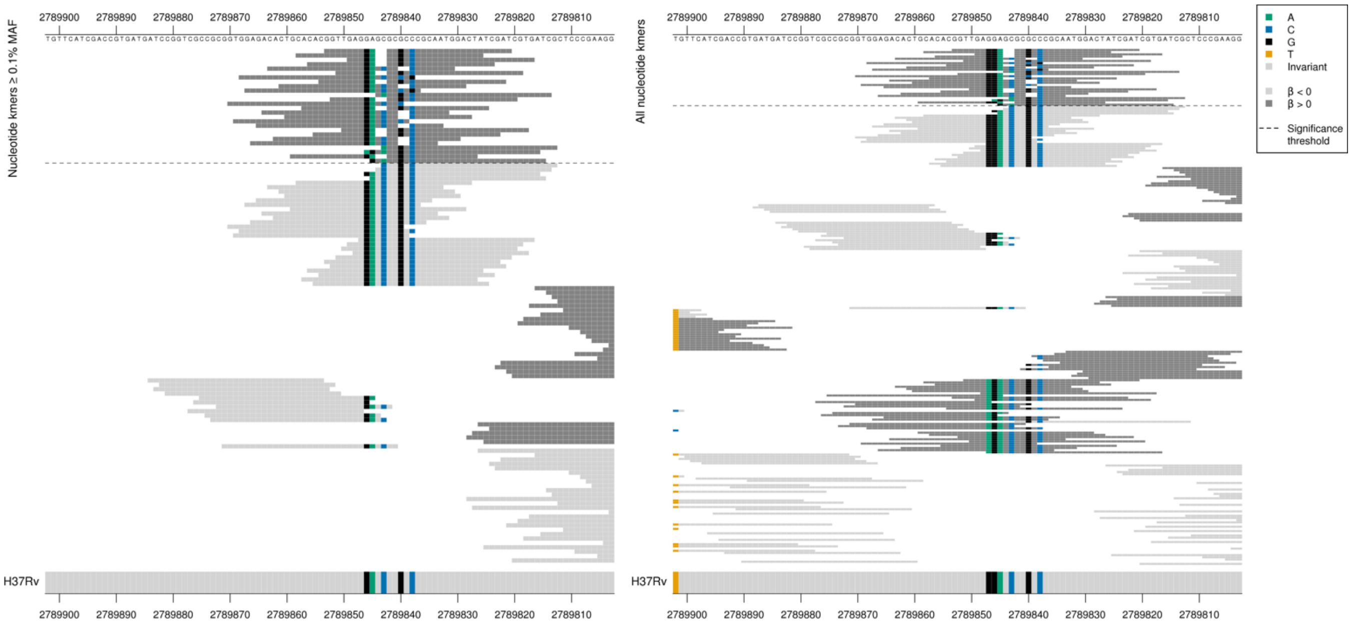

Deletion of the bases GC within the repetitive region “GCGCGC” positions 2789839-2789844 associated with high MIC.

| Kmer | -log$_{10}$ *p* | β | MAC | MAF | Ps | Variants |
|---|---|---|---|---|---|---|
| GGTTGAGGAGCGCCCGCAATGGACTATCGAT | 8.23 | 2.02 | 10 | 0.12 | 2789853 | GC deletion within repetitive region “GCGCGC” positions 2789844-2789839/1179-1184 |

# *Rv0920c*

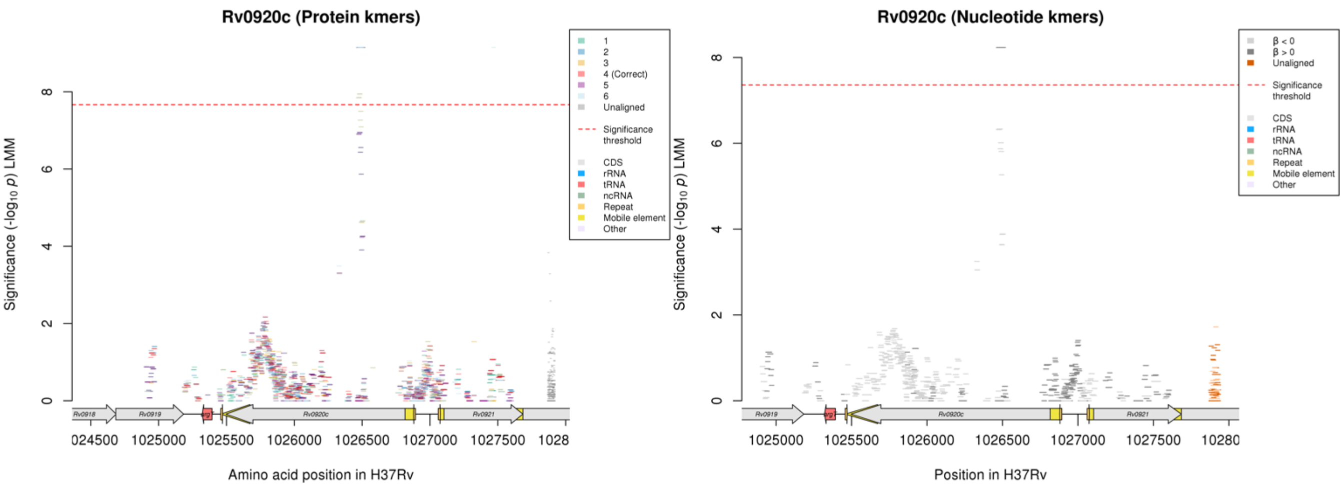

No significant protein kmers aligned to the correct reading frame.
Nucleotide kmers:

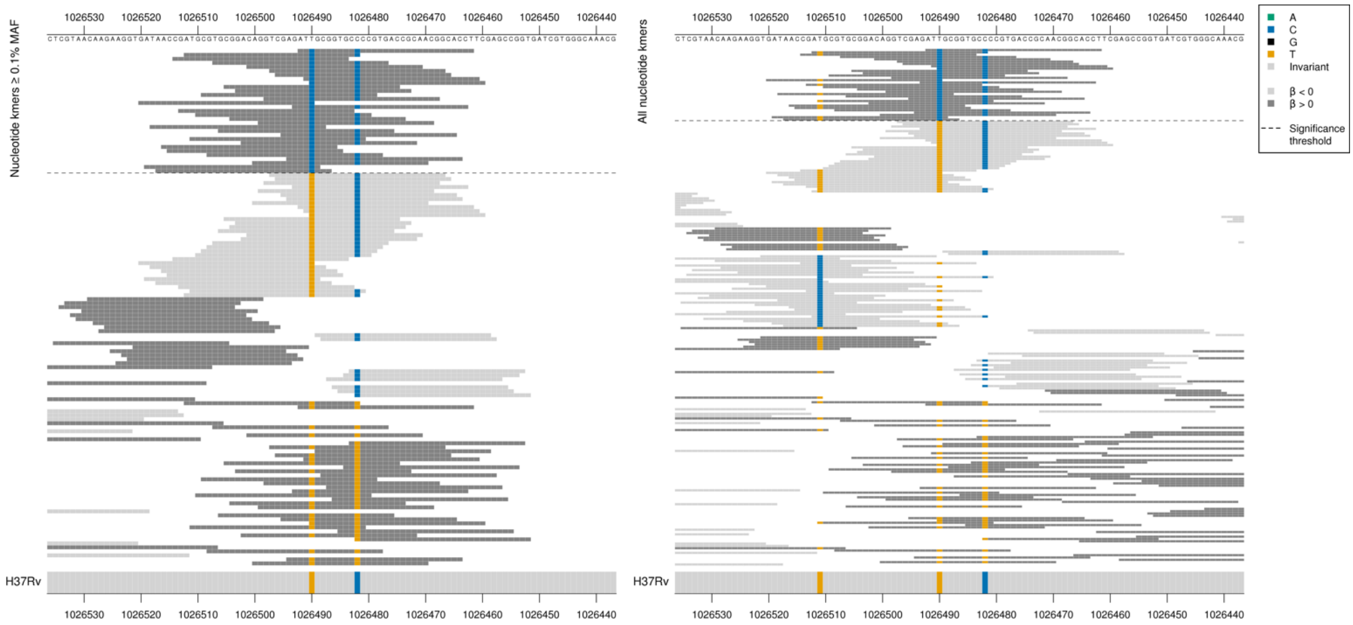

All high MIC-associated kmers were the same *p*-value, capturing the mutation at position 1026490/327.

High MIC-associated kmers = ATC = Ile / I
Kmers below significance threshold = ATT = Ile / I
Synonymous mutation.

| Kmer | -log$_{10}$ *p* | β | MAC | MAF | Ps | Variants |
|---|---|---|---|---|---|---|
| ATCGCGGTGCCCCGTGACCGCAACGGCACCT | 8.23 | 2.02 | 10 | 0.12 | 1026492 | 1026490/327C |

# *Rv3698*

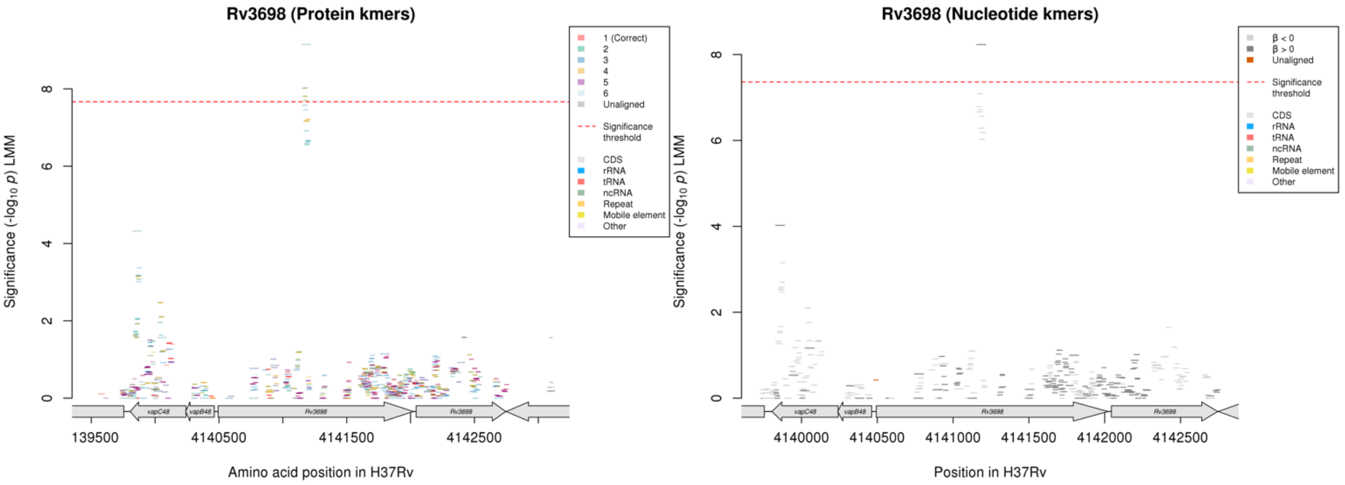

No significant protein kmers align to the correct reading frame.
Nucleotide kmers:

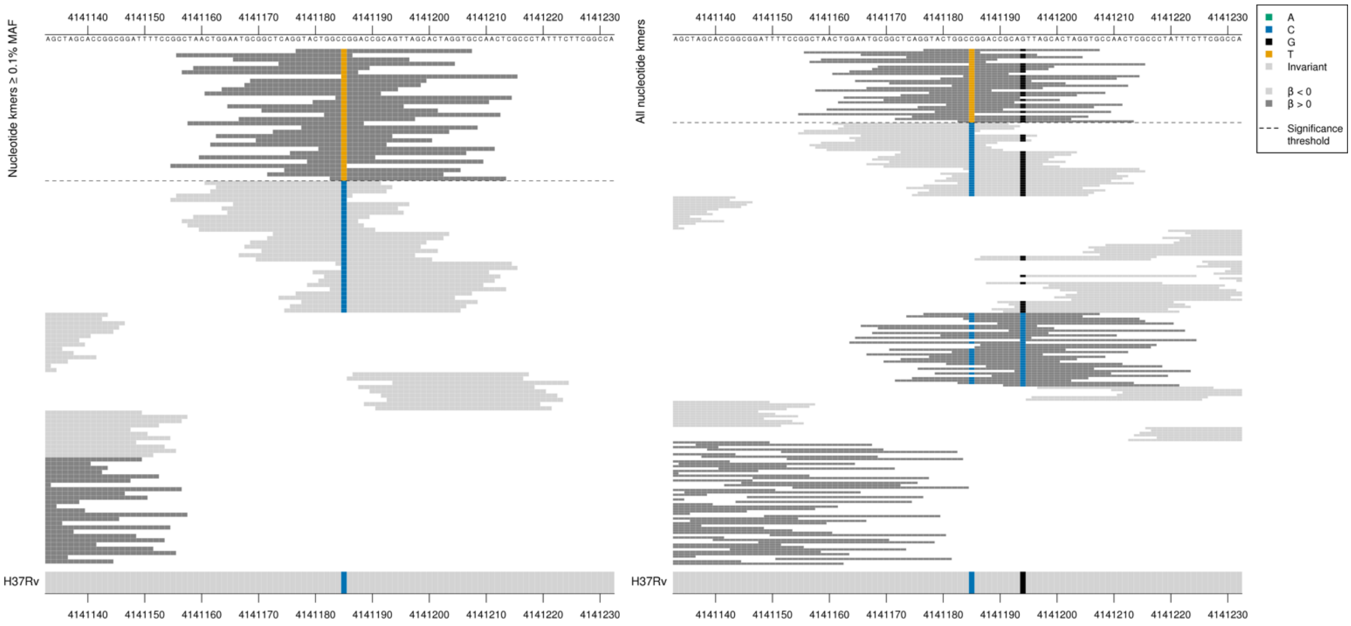

All high MIC-associated kmers were the same *p*-value, capturing the mutation at position 4141185/693 (codon 231).

High MIC-associated kmers = GCT = Ala / A
Kmers below significance threshold = GCC = Ala / A

| Kmer | $-\log_{10} p$ | β | MAC | MAF | Ps | Variants |
|---|---|---|---|---|---|---|
| GTACTGGCTGGACCGCAGTTAGCACTAGGTG | 8.23 | 2.02 | 10 | 0.12 | 4141177 | 4141185/693T |

#### *rrS*

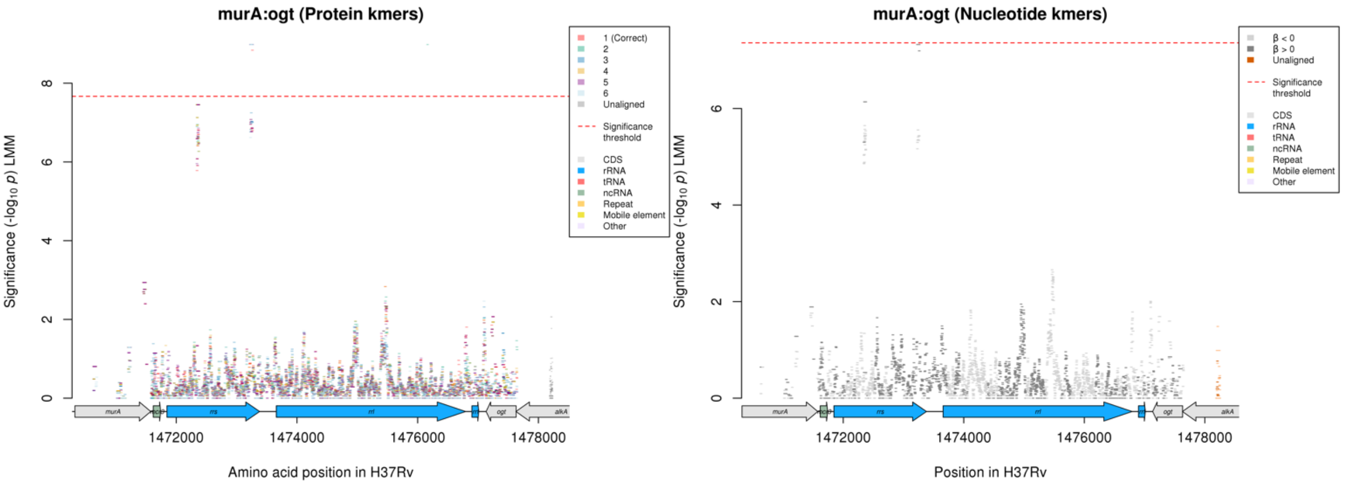

Not significant in the nucleotide kmer analysis. The region that is significant in the protein kmer analysis is just below the significance threshold for the nucleotide kmer analysis, showing the nucleotide kmers in the region below:

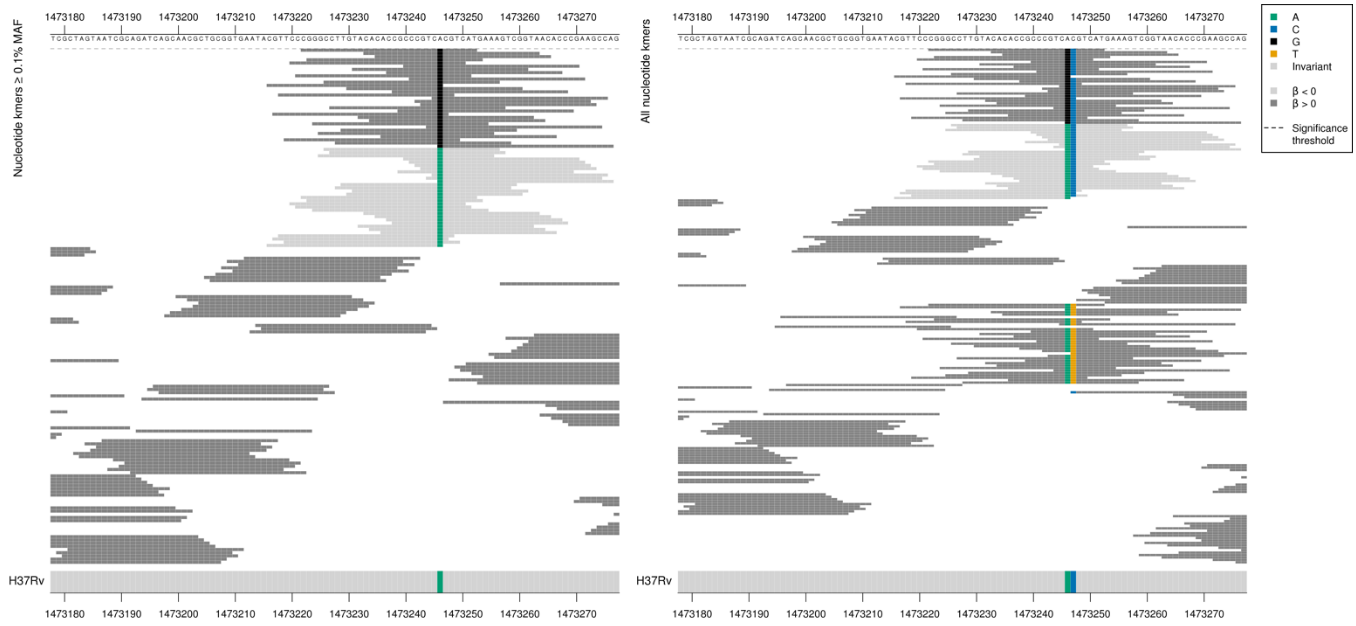

The top kmer from the protein kmer analysis for this region had a $-\log_{10} p$ = 8.98, beta = 0.48, MAC = 452, MAF = 5.45.

| Kmer | $-\log_{10} p$ | β | MAC | MAF | Ps | Variants |
|---|---|---|---|---|---|---|
| CGGGCCTTGTACACACCGCCCGTCGCGTCAT | 7.32 | 0.45 | 452 | 5.45 | 1473222 | 1473246/1401G |

### *pncA*

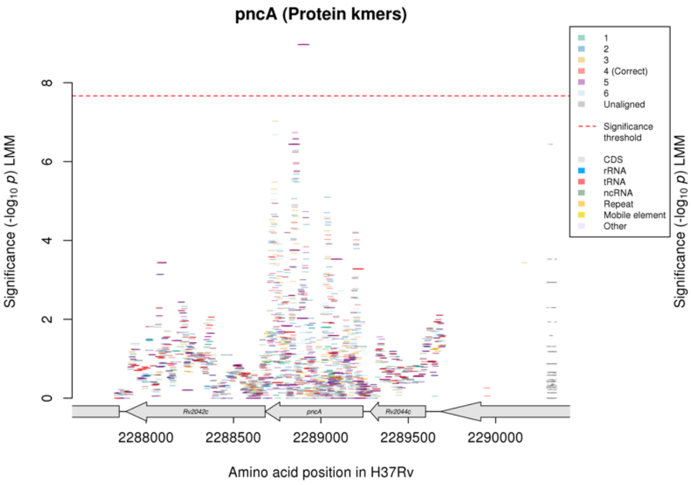

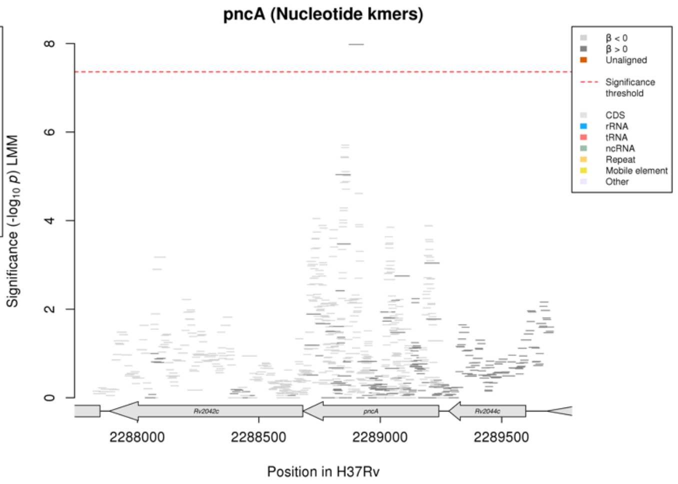

#### Protein kmers:

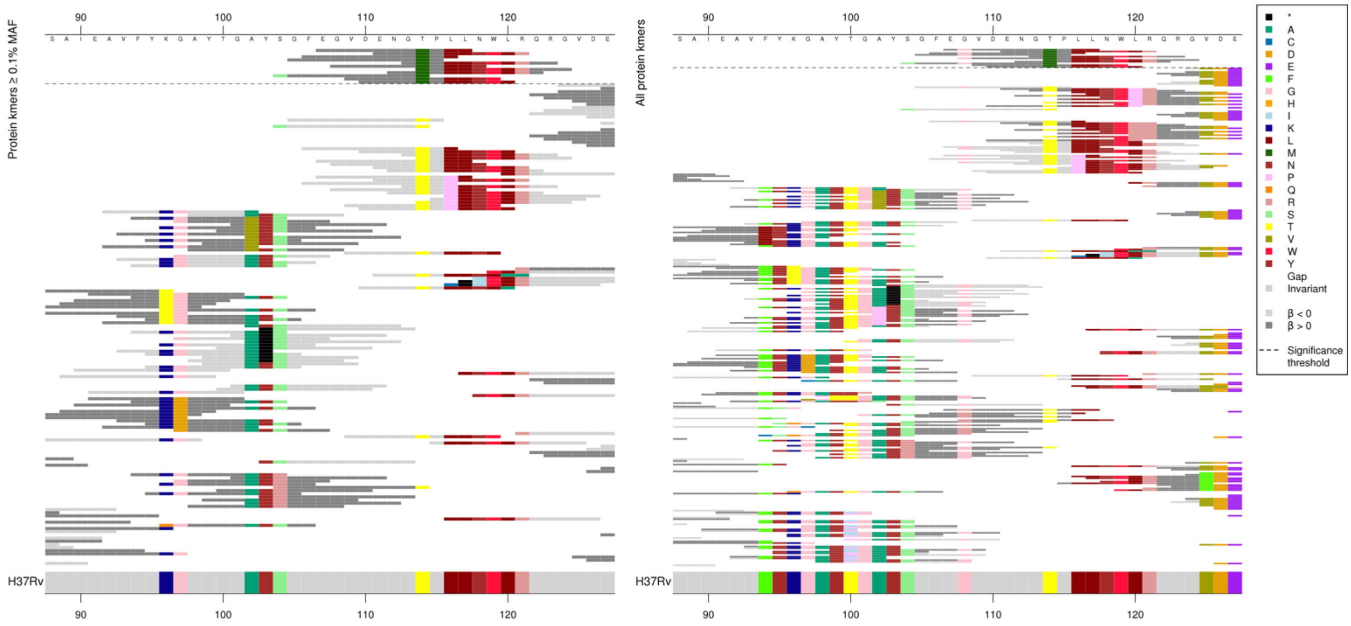

#### Same region for nucleotide kmers:

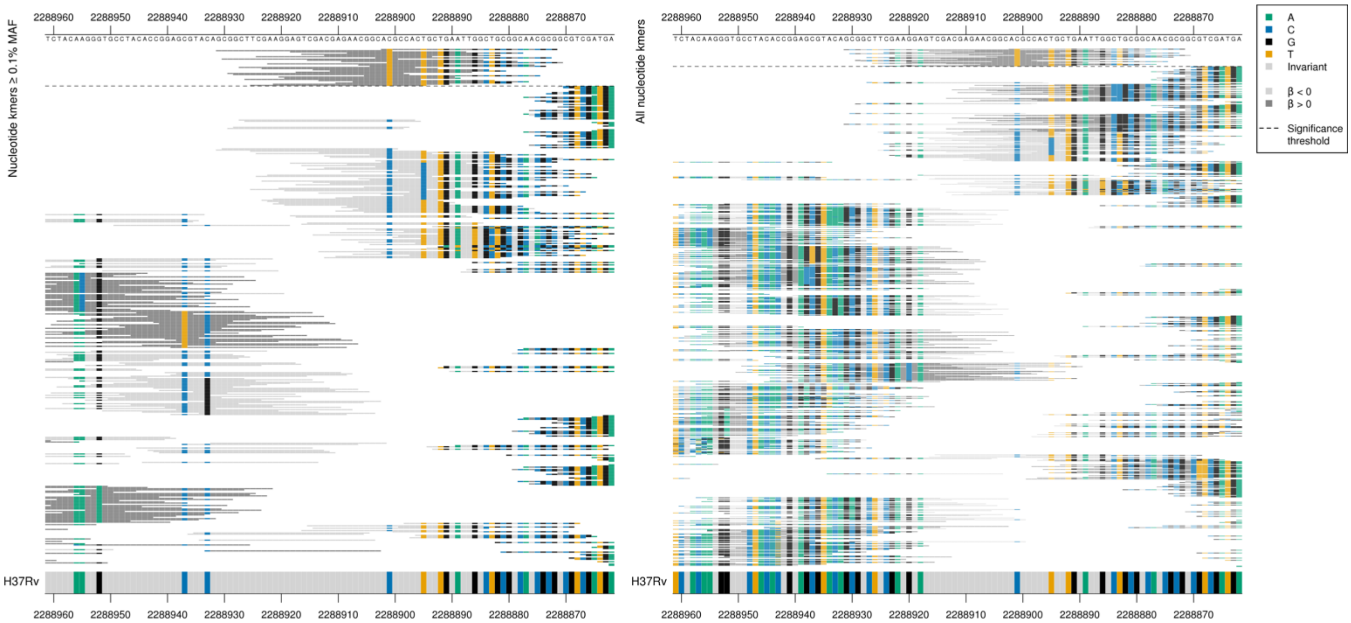

| Kmer | $-\log_{10} p$ | β | MAC | MAF | Ps | Variants |
|---|---|---|---|---|---|---|
| EGVDENGMPLL | 8.97 | 1.87 | 13 | 0.16 | 107 | 114M |

### *PPE56*

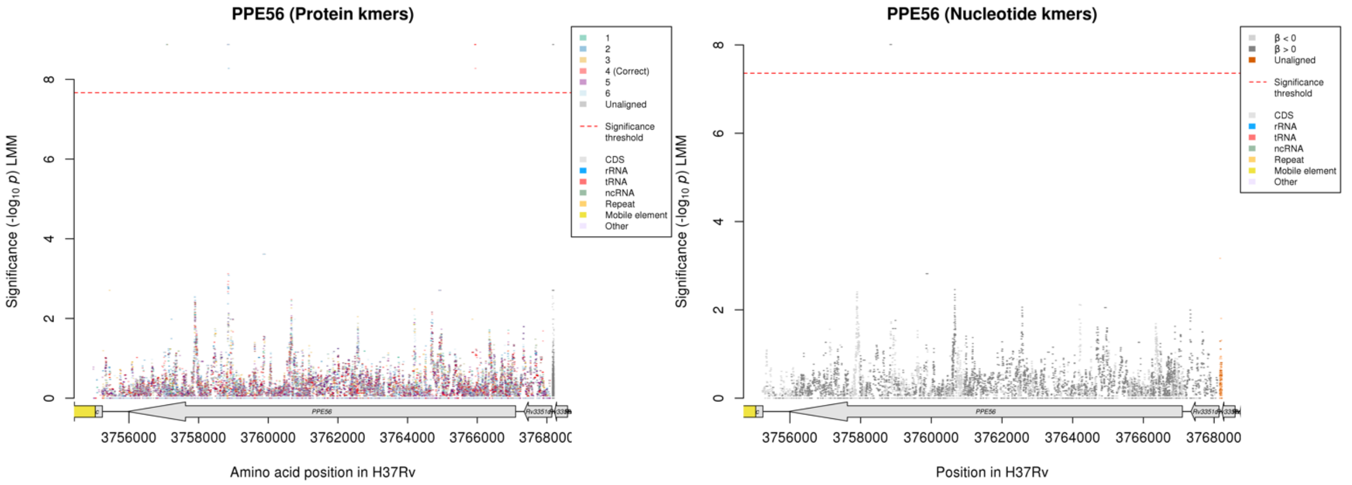

Nucleotide kmers:

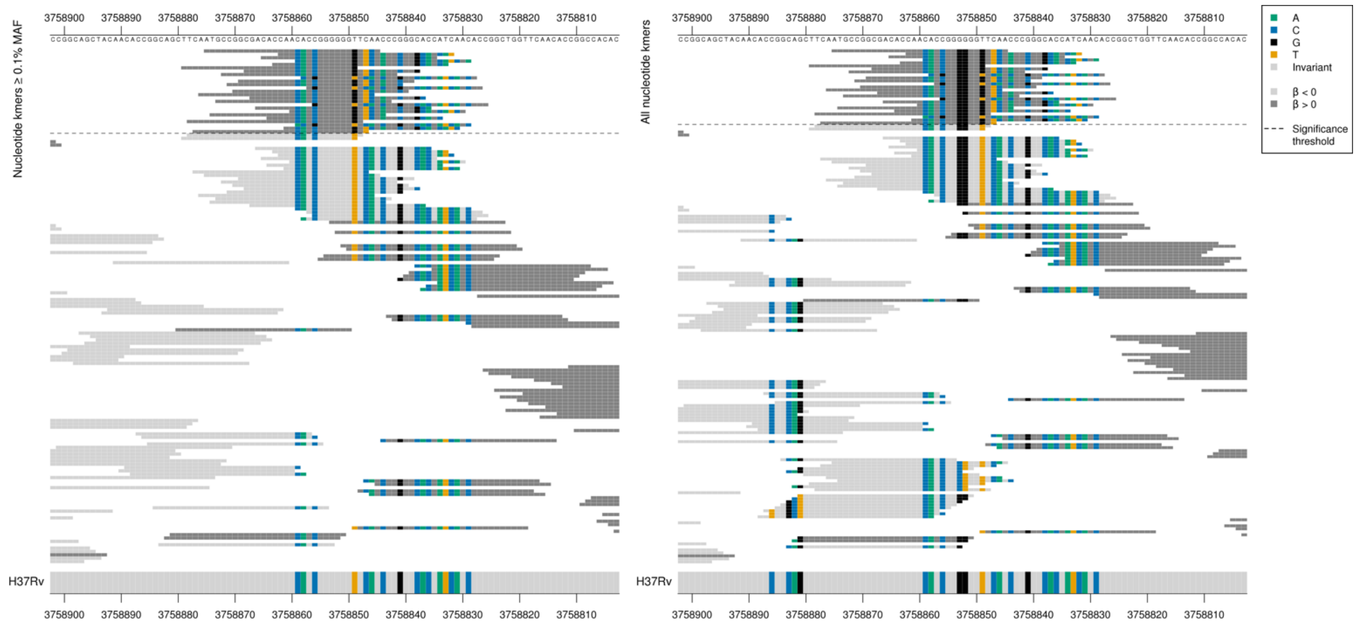

G insertion in increasing MIC-associated kmers adjacent to the seven bases long G region at positions 8248-8253 (3758855-3758850).

| Kmer | $-\log_{10} p$ | β | MAC | MAF | Ps | Variants |
|---|---|---|---|---|---|---|
| ATGCCGGCGACACCAACACCGGGGGGGTTCA | 8.01 | 1.93 | 11 | 0.13 | 3758875 | G insertion within "GGGGGGG" 3758855-3758850/8248-8253 |

# *Rv2019*

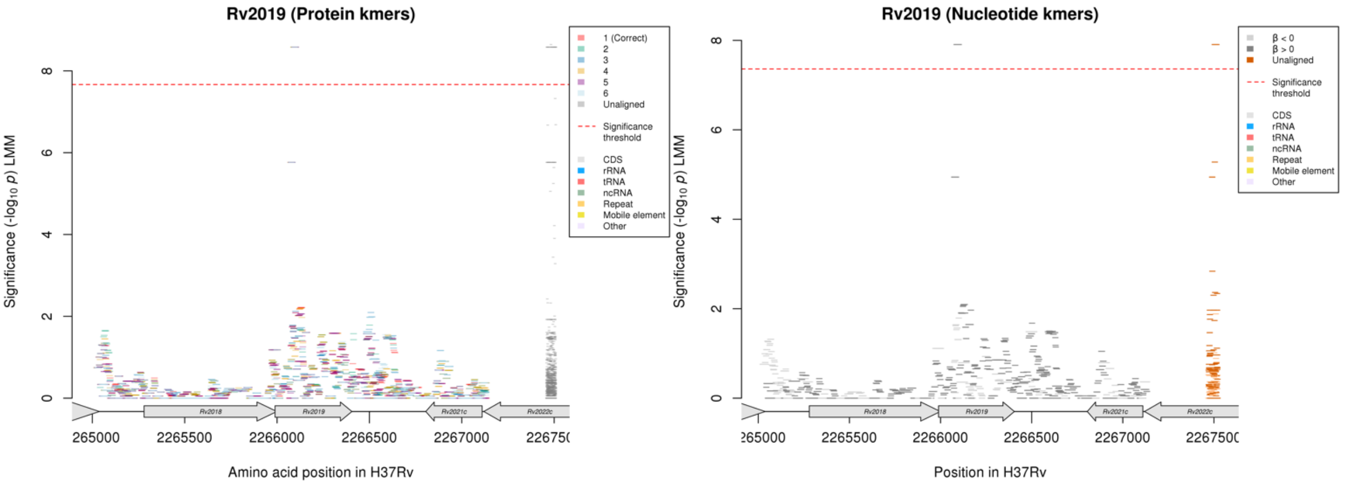

#### Protein kmers:

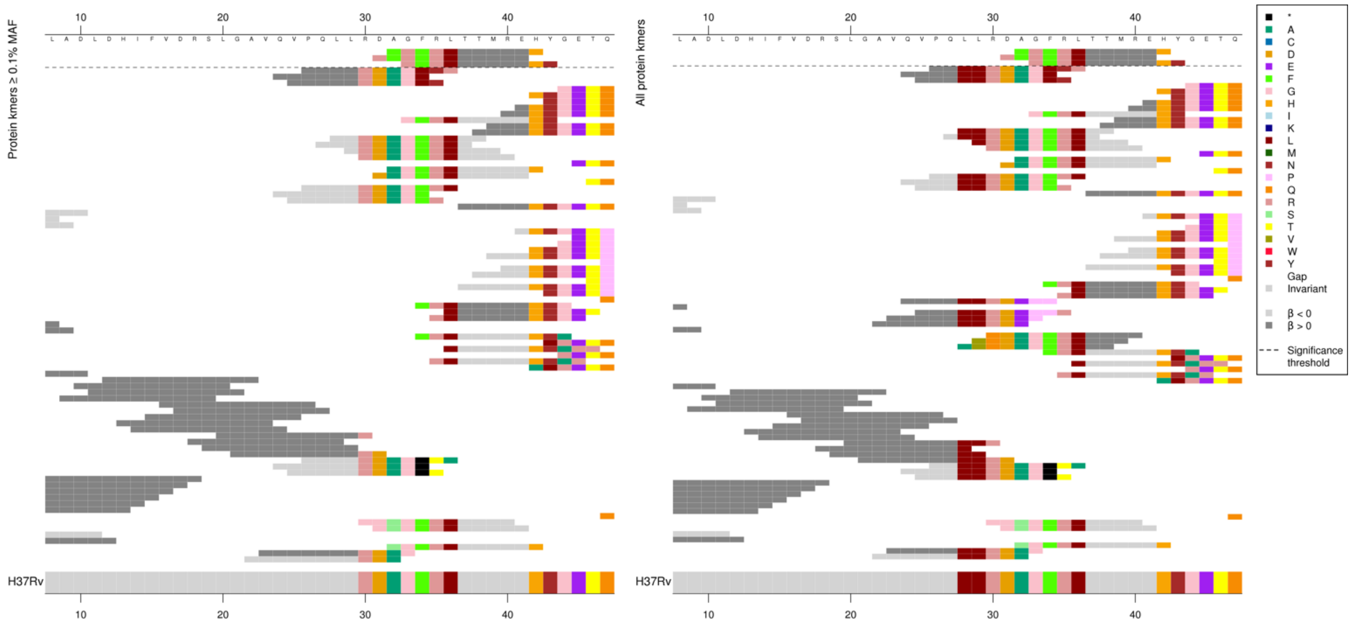

#### Same region for nucleotide kmers:

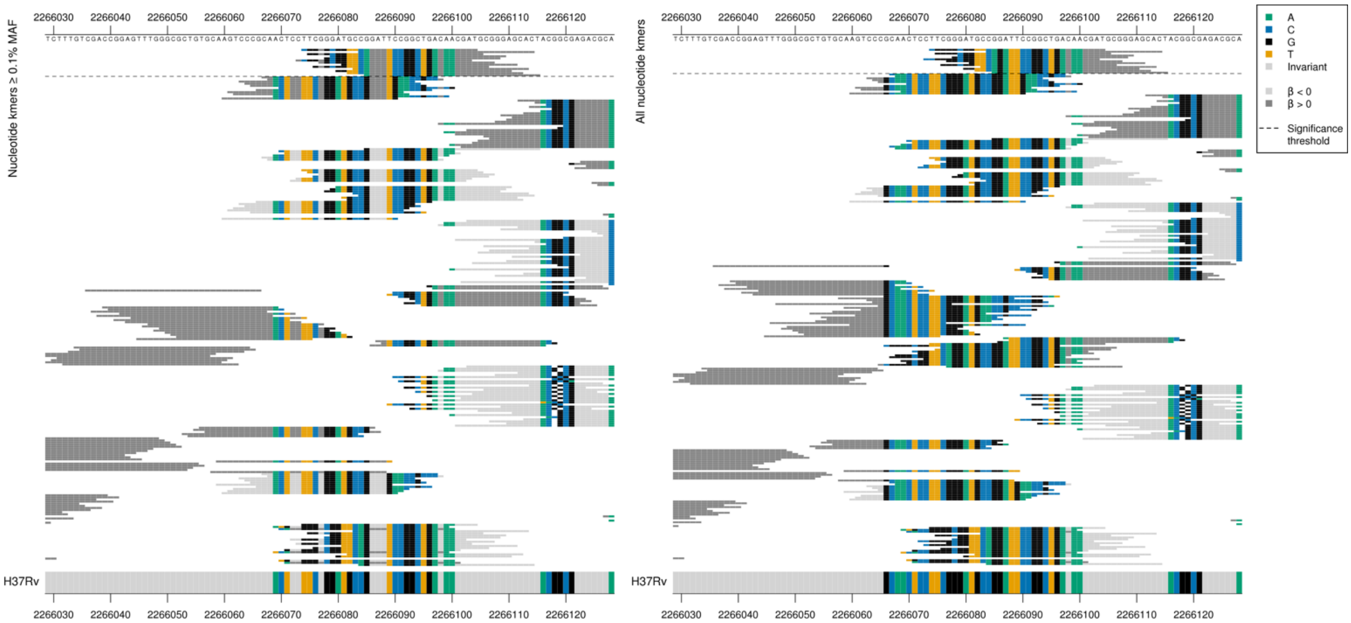

| Kmer | $-\log_{10} p$ | β | MAC | MAF | Ps | Variants |
|---|---|---|---|---|---|---|
| CGGGGCGGTTCAGATTCCGGCTGACAACGAT | 7.91 | 2.18 | 28 | 0.34 | 2266074 | Multiple differences to reference, possible indel |

### *lprF:Rv1371*$^{R}$

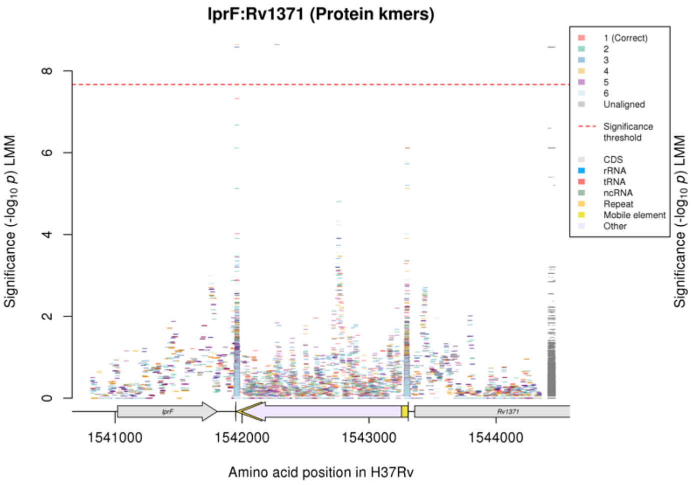

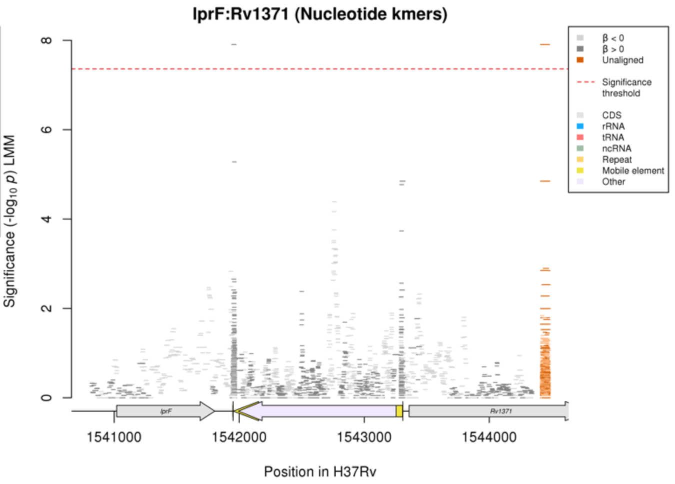

Repeat region.

Nucleotide kmers:

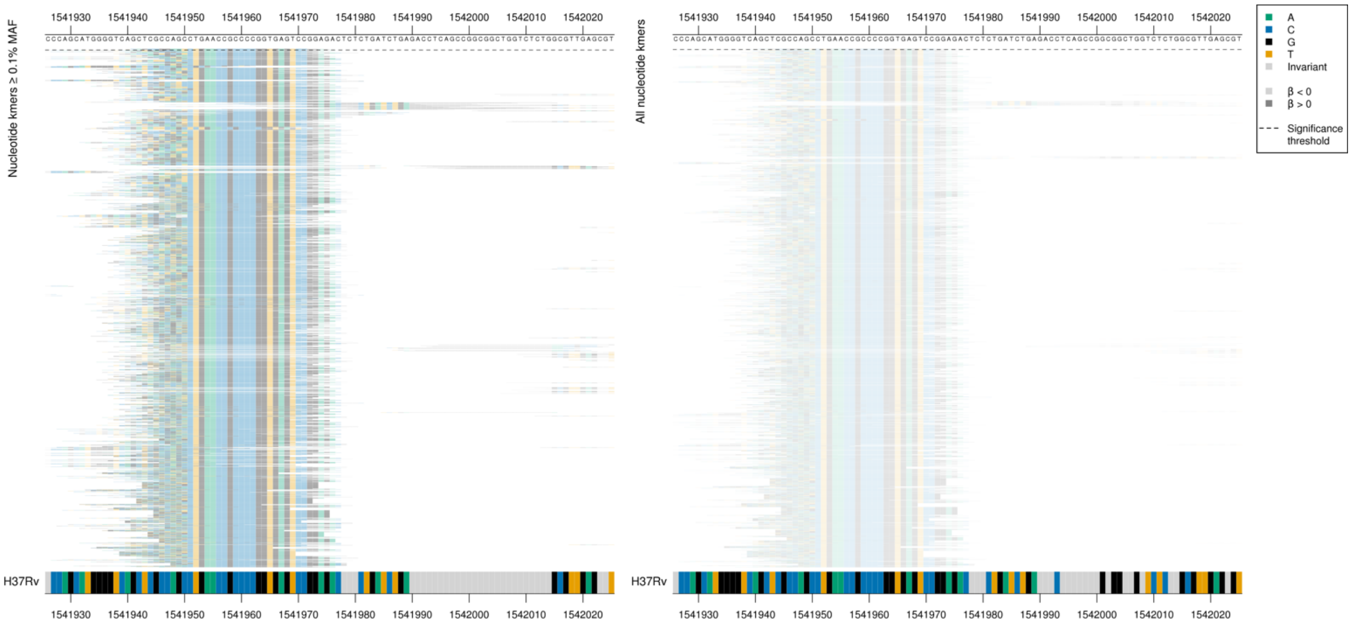

| Kmer | $-\log_{10} p$ | β | MAC | MAF | Ps | Variants |
|---|---|---|---|---|---|---|
| CAGCCGGAATCTGAACCGCCCCGGTGAGTCC | 7.91 | 2.18 | 28 | 0.34 | 1541941 | Repeat region |

### Isoniazid (INH)

#### *katG*

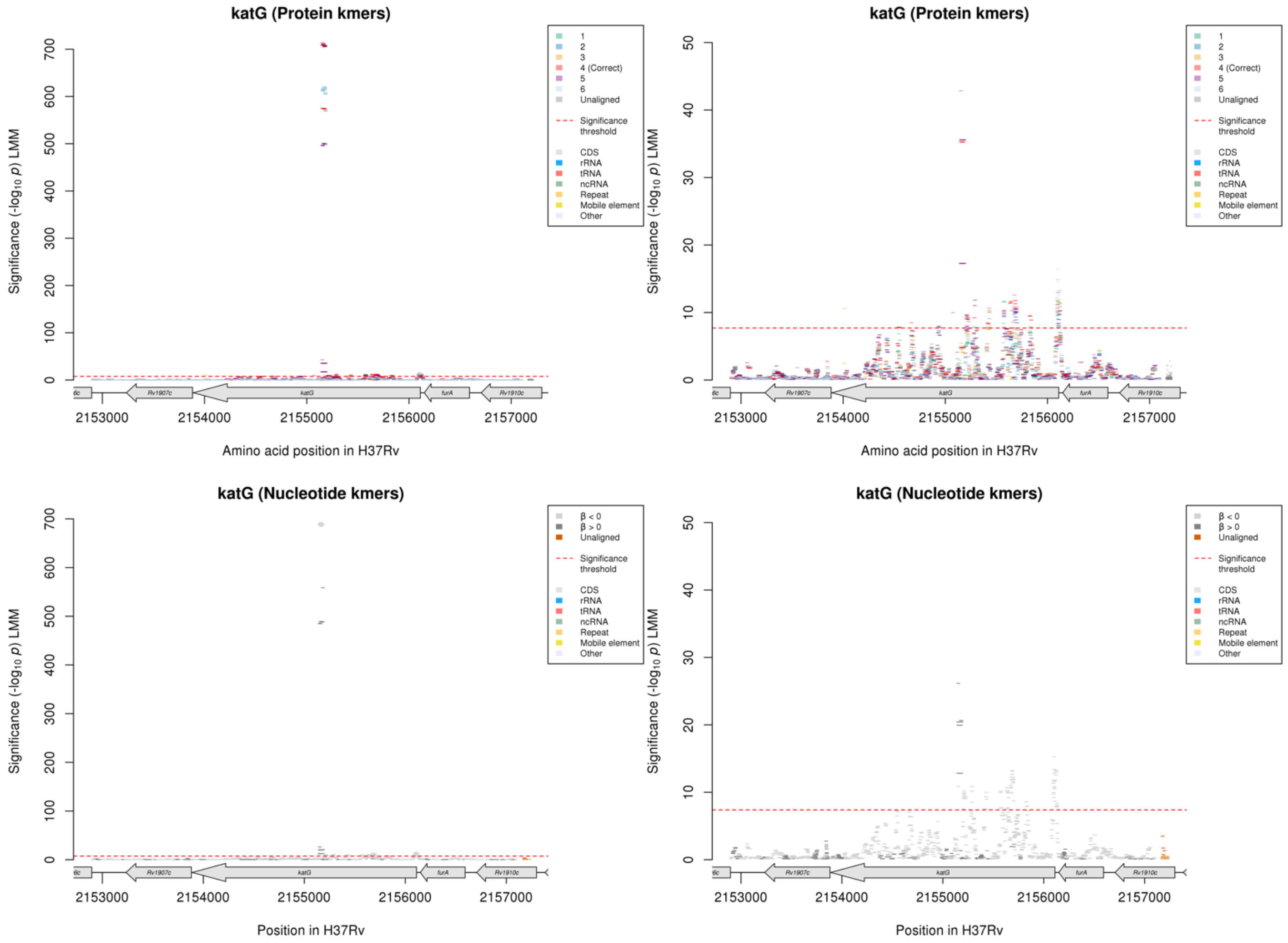

Protein kmers:

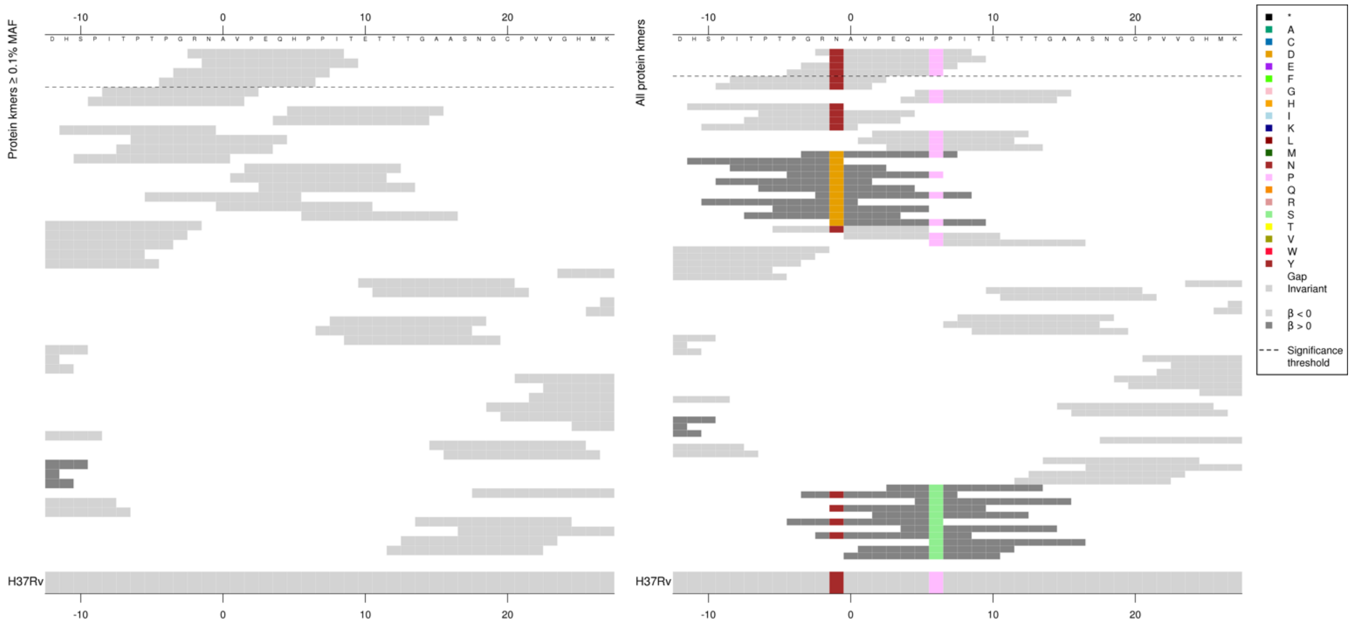

Same region for nucleotide kmers:

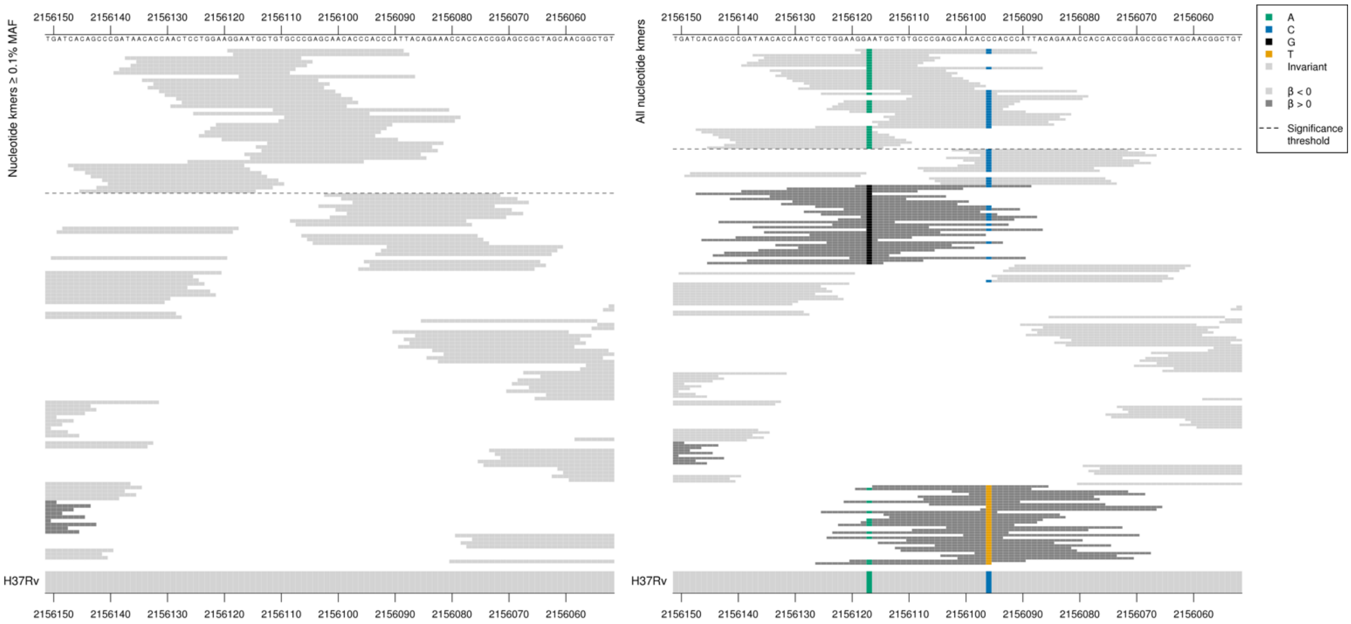

*katG* starts at position 2156111, *furA* ends at position 2156149.
Capturing a mutation 6bp upstream from the start codon and at position 16 in the gene.
2156117/-6:
Kmers below significance threshold = G
2156096/16 (codon 6):
Kmers below significance threshold = T (amino acid S)
For the codon 6 substitution, it is divergence from the reference that was associated with lower MIC.

| Kmer | $-\log_{10} p$ | β | MAC | MAF | Ps | Variants |
|---|---|---|---|---|---|---|
| GGAATGCTGTGCCCGAGCAACACCCACCCAT | 15.28 | -2.42 | 42 | 0.47 | 2156119 | 2156117/-6A, 2156096/16C (6P aa) |
| TAACACCAACTCCTGGAAGGAATGCTGTGCC | 13.24 | -2.18 | 43 | 0.48 | 2156137 | 2156117/-6A |
| GTGCCCGAGCAACACCCACCCATTACAGAAA | 11.72 | -2.44 | 29 | 0.32 | 2156111 | 2156096/16C (6P aa) |

Protein kmers:

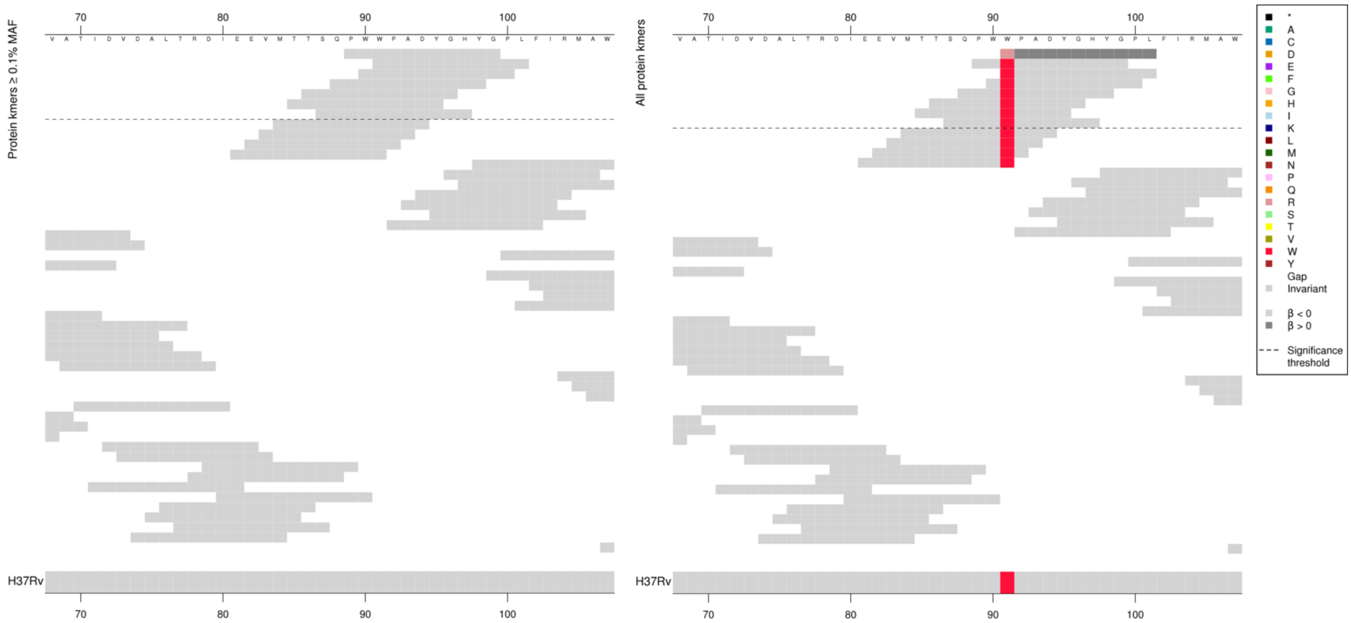

Low MAF kmers = 91R.

| Kmer | $-\log_{10} p$ | β | MAC | MAF | Ps | Variants |
|---|---|---|---|---|---|---|
| PWWPADYGHYG | 9.49 | -2.75 | 25 | 0.28 | 89 | 91W |

Protein kmers:

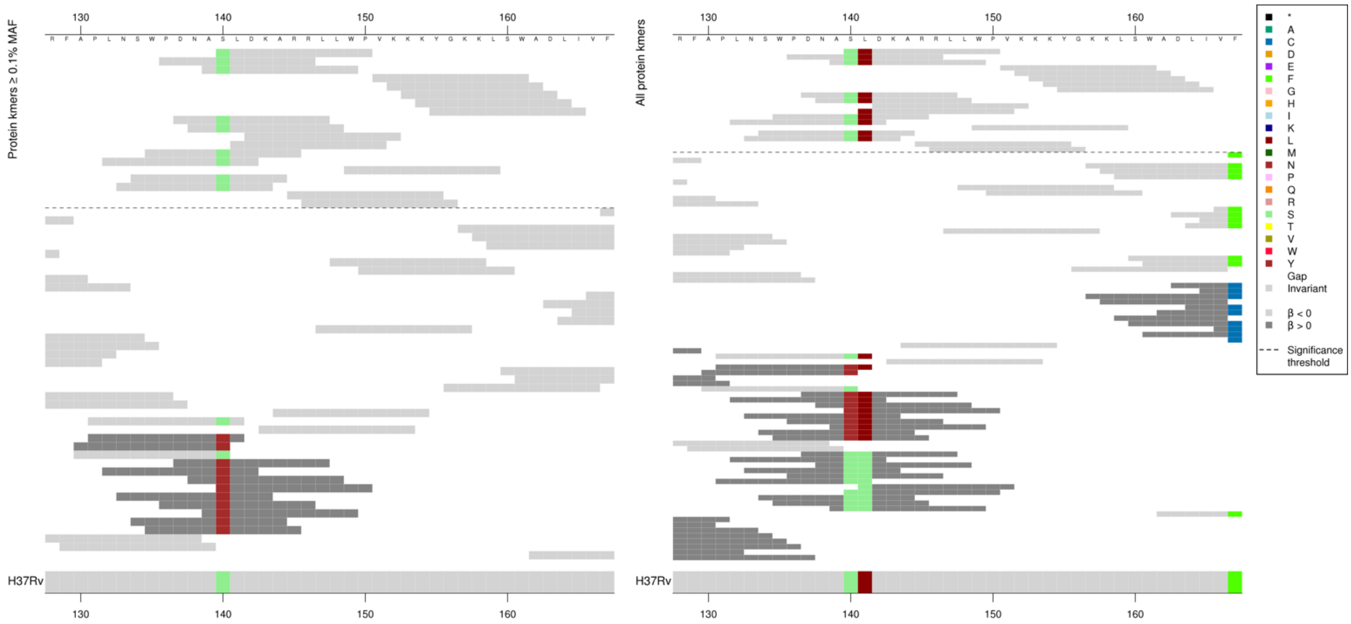

Low MIC-associated kmers covering codons 140-141 all have different *p*-values.
Kmers below significance threshold = 140N, 141S
WT kmers could be in LD with other variants or there could be rare variation not analysed here.
Keeping just the top WT kmer for the table.

| Kmer | -log₁₀ *p* | β | MAC | MAF | Ps | Variants |
|---|---|---|---|---|---|---|
| SLDKARRLLWP | 12.59 | -2.22 | 44 | 0.49 | 140 | 140S, 141L |
| VKKKYGKKLSW | 11.68 | -2.65 | 32 | 0.36 | 151 | WT |
| LDKARRLLWPV | 10.63 | -2.23 | 35 | 0.39 | 141 | 141L |

Protein kmers:

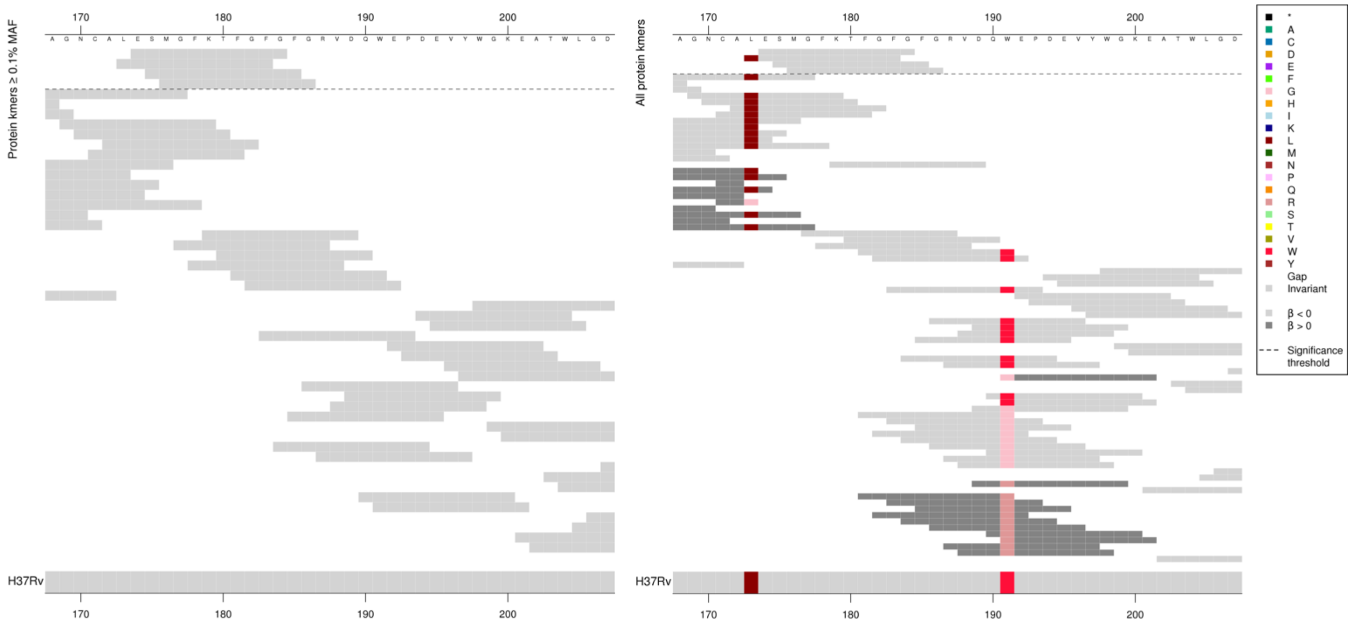

Possibly in LD with other regions or capturing rare variation.

| Kmer | -log₁₀ *p* | β | MAC | MAF | Ps | Variants |
|---|---|---|---|---|---|---|
| ESMGFKTFGFG | 10.28 | -2.92 | 21 | 0.24 | 174 | WT covering codons 173-186 |

Protein kmers:

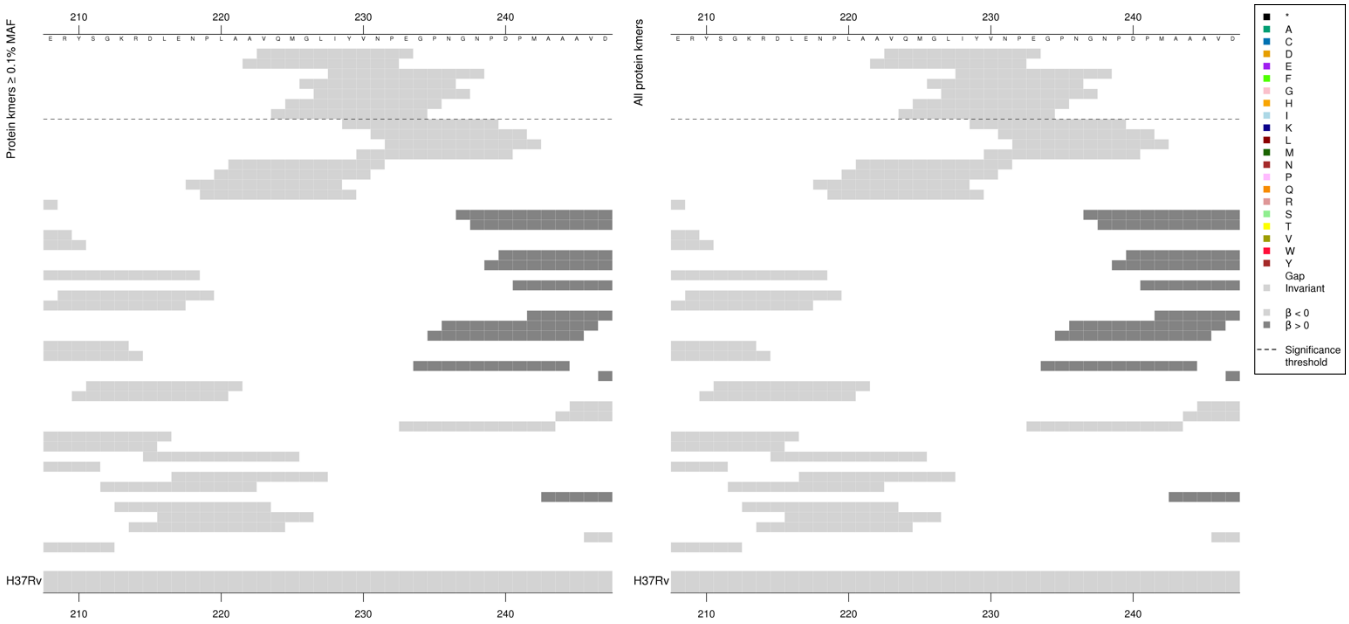

Possibly in LD with other regions or capturing rare variation.

| Kmer | -log$_{10}$ *p* | β | MAC | MAF | Ps | Variants |
|---|---|---|---|---|---|---|
| VQMGLIYVNPE | 10.64 | -2.82 | 23 | 0.26 | 223 | WT covering codons 222-238 |

Protein kmers:

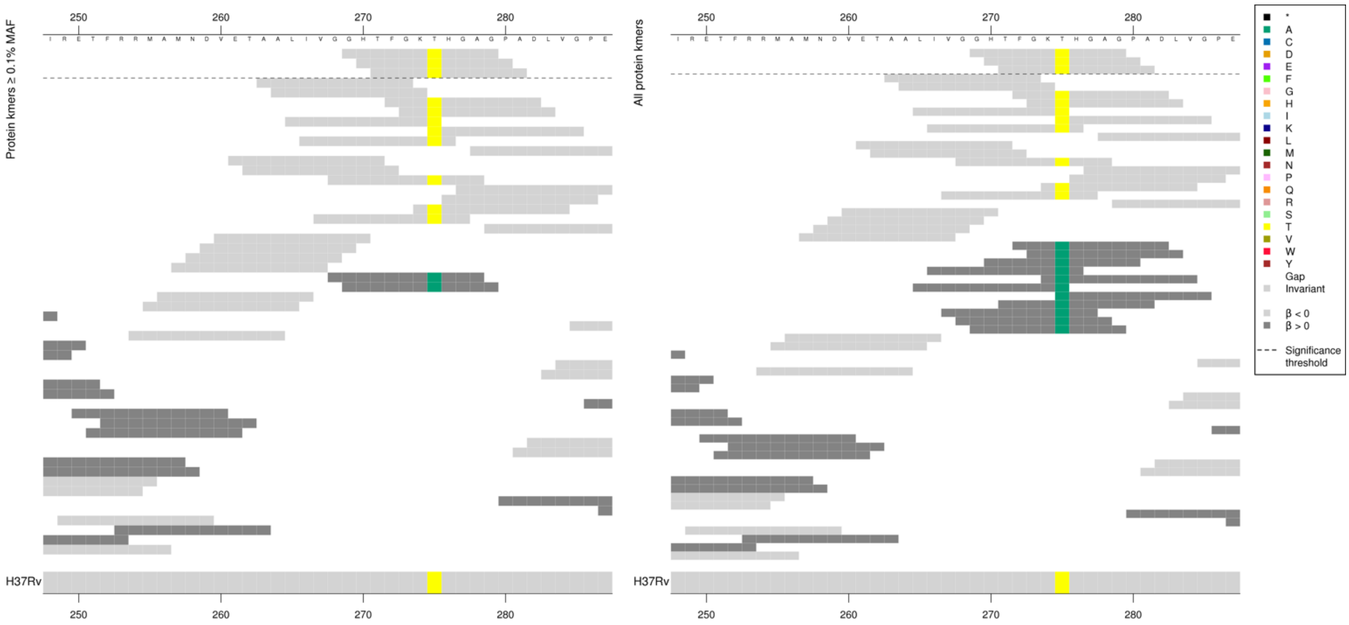

Kmers below significance threshold = 275A

| Kmer | -log$_{10}$ *p* | β | MAC | MAF | Ps | Variants |
|---|---|---|---|---|---|---|
| GHTFGKTHGAG | 11.84 | -2.36 | 32 | 0.36 | 269 | WT / 275T |

Protein kmers:

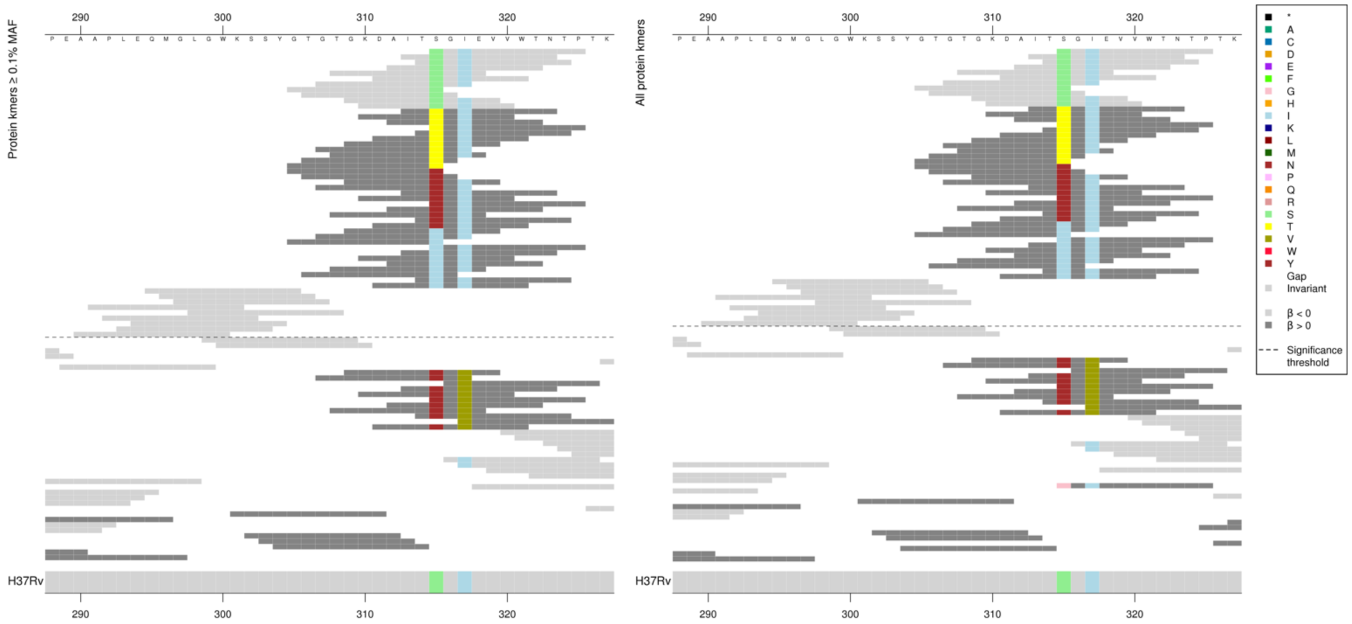

Kmer *p*-values were possibly influenced by the variant at codon 317, although the low MIC-associated kmers that did not cover codon 317 were the same *p*-value as another kmer that does. The high MIC-associated 315T kmers were a range of *p*-values but they did not change in line with covering 317. The 315N kmers that did not cover 317 were more significant that those that did. The 315I kmers were all the same *p*-value.

Kmers below significance threshold = 317V

WT kmers were possibly in LD with the kmers capturing codon 315 or were capturing a region with rare variation not analysed here.

| Kmer | $-\log_{10} p$ | β | MAC | MAF | Ps | Variants |
|---|---|---|---|---|---|---|
| SGIEVVWTNTP | 714.16 | -5.11 | 3372 | 37.76 | 315 | 315S |
| ITTGIEVVWTN | 574.79 | 4.90 | 3290 | 36.85 | 313 | 315T |
| GTGTGKDAITN | 35.57 | 2.86 | 56 | 0.63 | 305 | 315N |
| GKDAITIGIEV | 17.27 | 3.91 | 10 | 0.11 | 309 | 315I |
| QMGLGWKSSYG | 9.64 | -2.58 | 29 | 0.32 | 295 | WT covering codons 290-308 |

Protein kmers:

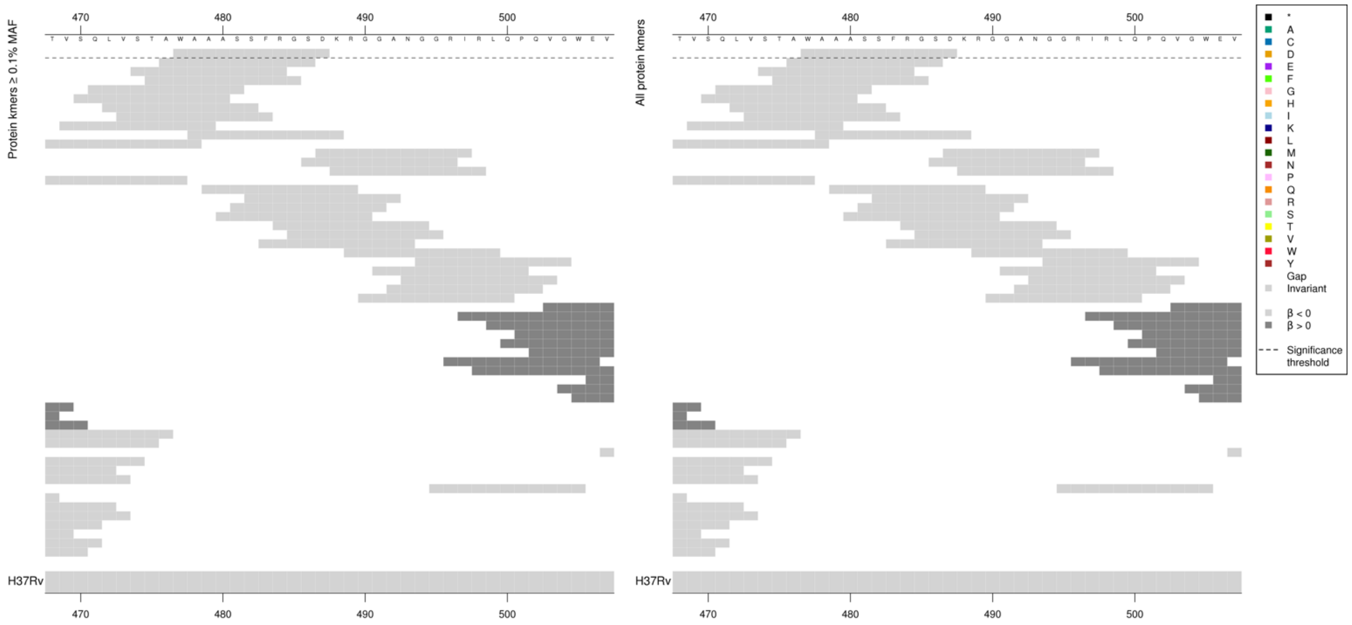

Possibly in LD with other regions or capturing rare variation not analysed here.

| Kmer | $-\log_{10} p$ | β | MAC | MAF | Ps | Variants |
|---|---|---|---|---|---|---|
| WAAASSFRGSD | 8.50 | -2.90 | 23 | 0.26 | 477 | WT covering codons 477-487 |

Protein kmers:

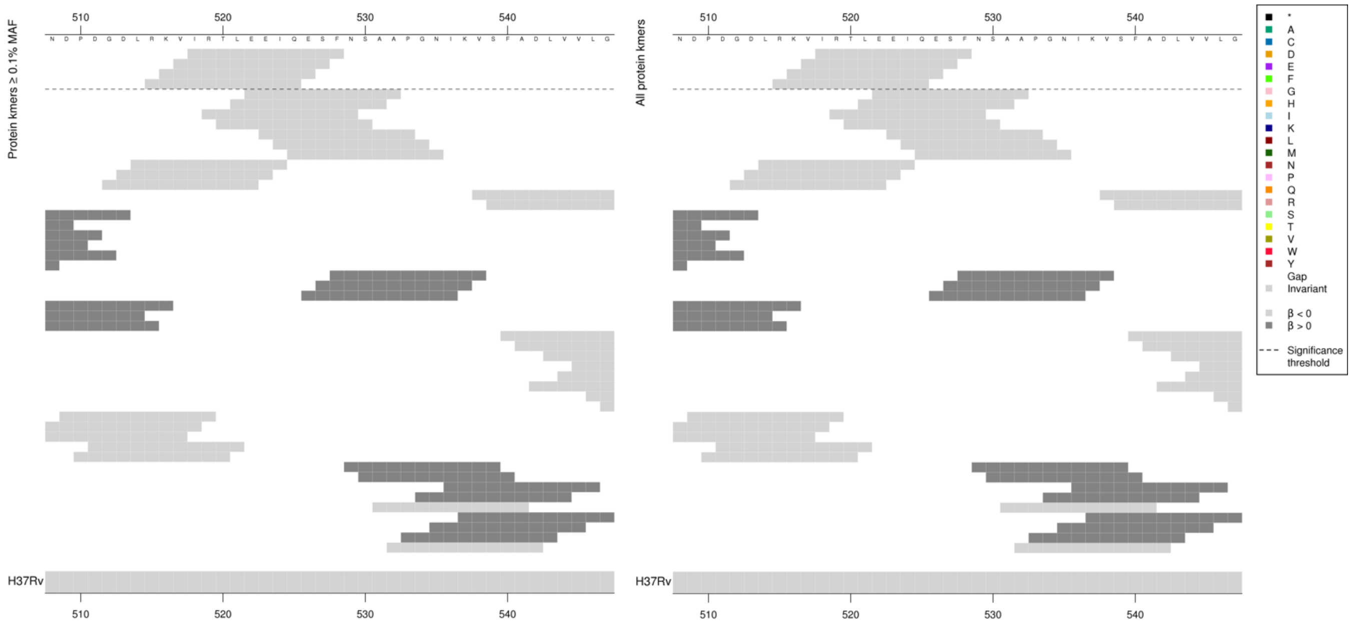

Possibly in LD with other regions or capturing rare variation not analysed here.

| Kmer | $-\log_{10} p$ | β | MAC | MAF | Ps | Variants |
|---|---|---|---|---|---|---|
| IRTLEEIQESF | 7.81 | -2.88 | 21 | 0.24 | 518 | WT covering codons 515-528 |

#### *proA:ahpC*

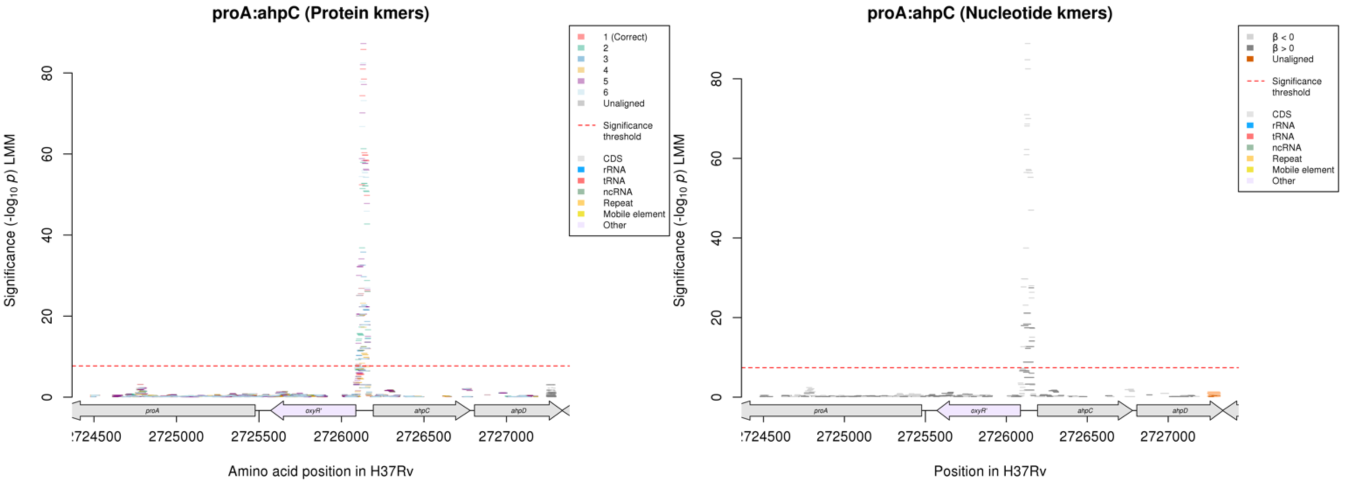

Nucleotide kmers:

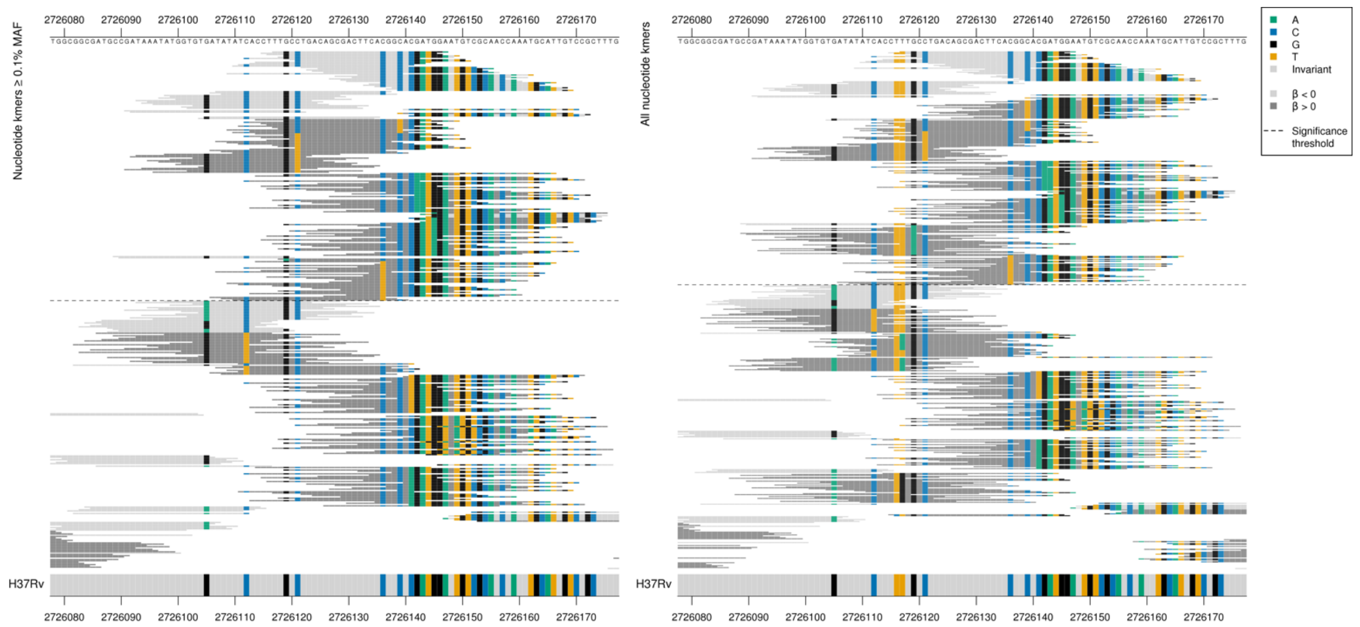

oxyR’ begins at position 2726087, *ahpC* begins at position 2726193, so this is in the intergenic region upstream of both.

Main variants = 2726105, 2726117, 2726119, 2726121, 2726136, 2726139, 2726142, 2726145

Low MAF kmers = 2726139T

| Kmer | $-\log_{10} p$ | β | MAC | MAF | Ps | Variants |
|---|---|---|---|---|---|---|
| TTGCCTGACAGCGACTTCACGGCACGATGGA | 88.85 | -3.03 | 188 | 2.11 | 2726117 | WT / 2726139C + others |
| TATATCACCTTTGCCTGACAGCGACTTCACG | 37.48 | -3.03 | 68 | 0.76 | 2726107 | WT / 2726112C, 2726116T, 2726117T, 2726119G, 2726121C, 2726136C |
| ATGGTGTGATATATCACCTTTGCCTGACAGC | 29.75 | -3.36 | 718 | 8.04 | 2726098 | WT / 2726105G, 2726116T, 2726117T, 2726119G, 2726121C |
| TACGATGGAATGTCGCAACCAAATGCATTGT | 27.49 | 5.34 | 10 | 0.11 | 2726139 | 2726139T |
| GATGGAATGTCGCAACCAAATGCATTGTCCG | 26.34 | -2.35 | 88 | 0.99 | 2726142 | WT / 2726142G + others |
| CGATGGAATGTCGCAACCAAATGCATTGTCC | 24.91 | -1.90 | 137 | 1.53 | 2726141 | WT / 2726141C, 2726142G + others |
| CCTTTGCCTGACAGCGACTTCACGGTACGAT | 21.11 | 5.07 | 9 | 0.10 | 2726114 | 2726119G, 2726139T |
| CCTTTGCTTGACAGCGACTTCACGGCACGAT | 18.36 | 4.55 | 14 | 0.16 | 2726114 | 2726121T, 2726136C |
| ATGGTGTGATATATCACCTTTGCTTGACAGC | 17.97 | 4.62 | 13 | 0.15 | 2726098 | 2726105G, 2726121T |
| CGGCACAATGGAATGTCGCAACCAAATGCAT | 17.42 | 3.15 | 21 | 0.24 | 2726136 | 2726142A |
| AGAATGTCGCAACCAAATGCATTGTCCGCTT | 17.34 | 2.47 | 38 | 0.43 | 2726145 | 2726145A |
| ATGGAATGTCGCAACCAAATGCATTGTCCGC | 15.07 | -2.05 | 69 | 0.77 | 2726143 | WT / 2726145G |
| CCGATAAATATGGTGTGATATATCACCTTTG | 12.66 | -2.41 | 709 | 7.94 | 2726089 | WT / 2726119G |
| TACCTGACAGCGACTTCACGGCACGATGGAA | 12.25 | 4.14 | 9 | 0.10 | 2726118 | 2726119A |
| CCTTTGCCTGACAGCGACTTCATGGCACGAT | 8.81 | 3.19 | 11 | 0.12 | 2726114 | 2726136T |

### *fabG1*

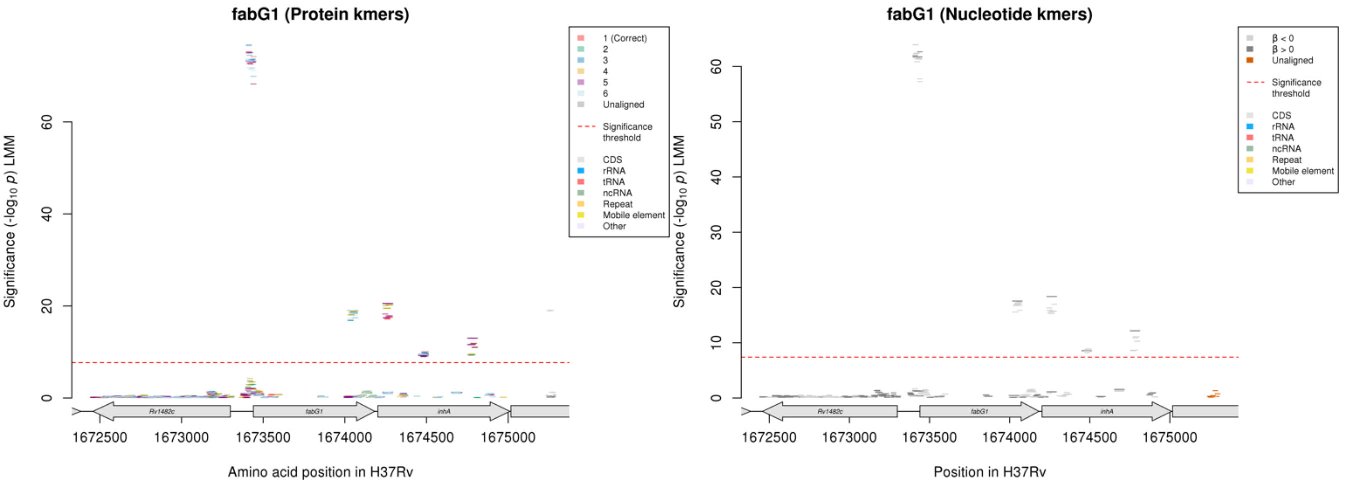

Nucleotide kmers:

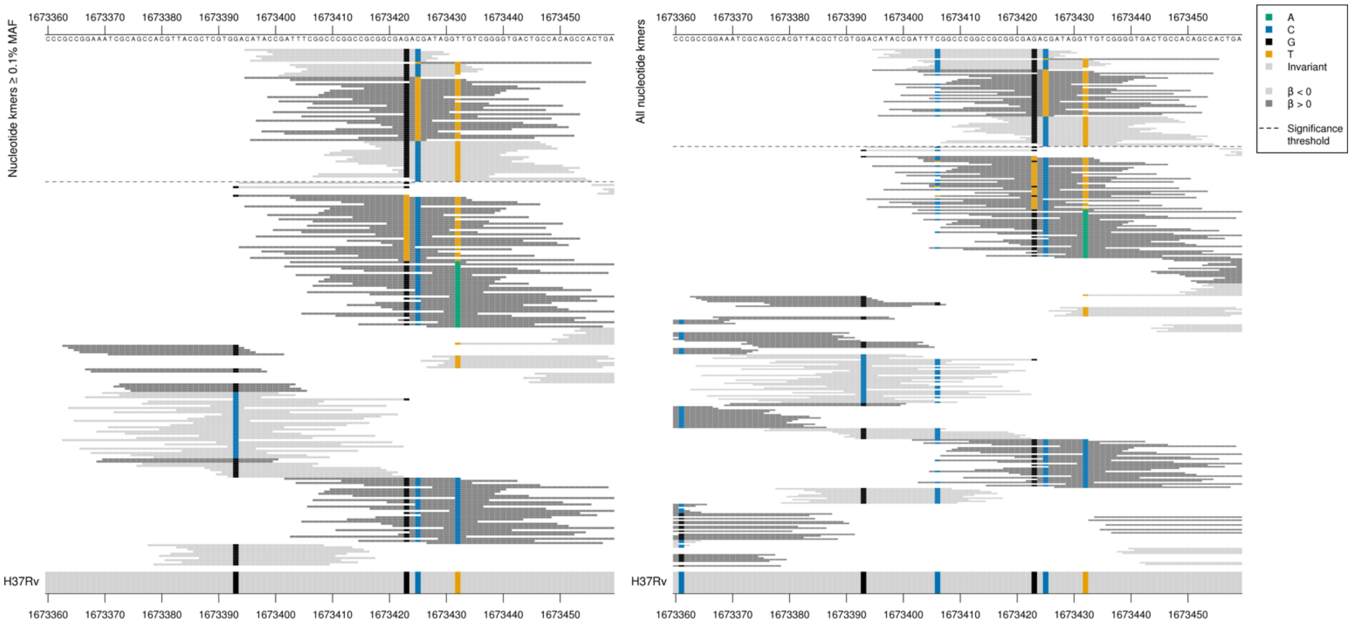

fabG1 begins at position 1673440.
Capturing a combination of variants. Divergence from the reference sequence was associated with increasing MICs.

1673406:
Low MIC-associated kmers = C
Some kmers below significance threshold = G

1673423:
Low MIC-associated kmers = G
Some kmers below significance threshold = T

1673425:
Low MIC-associated kmers = C
Some high MIC-associated kmers = T

1673432:
Low MIC-associated kmers = T
Some kmers below significance threshold = A/C

Main mutation being captured is 1673425/-15. The other mutations affect the significance. The low MIC-associated kmers not covering position 1673432/-8 were more significant than those which did.

| Kmer | $-\log_{10} p$ | β | MAC | MAF | Ps | Variants |
|---|---|---|---|---|---|---|
| CATACCGATTTCGGCCCGGCCGCGGCGAGAC | 63.92 | -1.94 | 998 | 11.18 | 1673395 | 1673406/-34C, 1673423/-17G, 1673425/-15C |
| TGATAGGTTGTCGGGGTGACTGCCACAGCCA | 62.65 | 2.01 | 801 | 8.97 | 1673425 | 1673425/-15T, 1673432/-8T |
| TTTCGGCCCGGCCGCGGCGAGACGATAGGTT | 62.39 | -1.81 | 1094 | 12.25 | 1673403 | 1673406/-34C, 1673423/-17G, 1673425/-15C, 1673432/-8T |
| CATACCGATTTCGGCCCGGCCGCGGCGAGAT | 61.95 | 2.00 | 801 | 8.97 | 1673395 | 1673406/-34C, 1673423/-17G, 1673425/-15T |
| TTTCGGCCCGGCCGCGGCGAGATGATAGGTT | 61.73 | 2.00 | 800 | 8.96 | 1673403 | 1673425/-15T |
| GCGGCGAGACGATAGGTTGTCGGGGTGACTG | 61.49 | -1.82 | 1082 | 12.12 | 1673416 | 1673423/-17G, 1673425/-15C, 1673432/-8T |
| ACGATAGGTTGTCGGGGTGACTGCCACAGCC | 57.76 | -1.79 | 898 | 10.06 | 1673424 | 1673425/-15C, 1673432/-8T |

Nucleotide kmers:

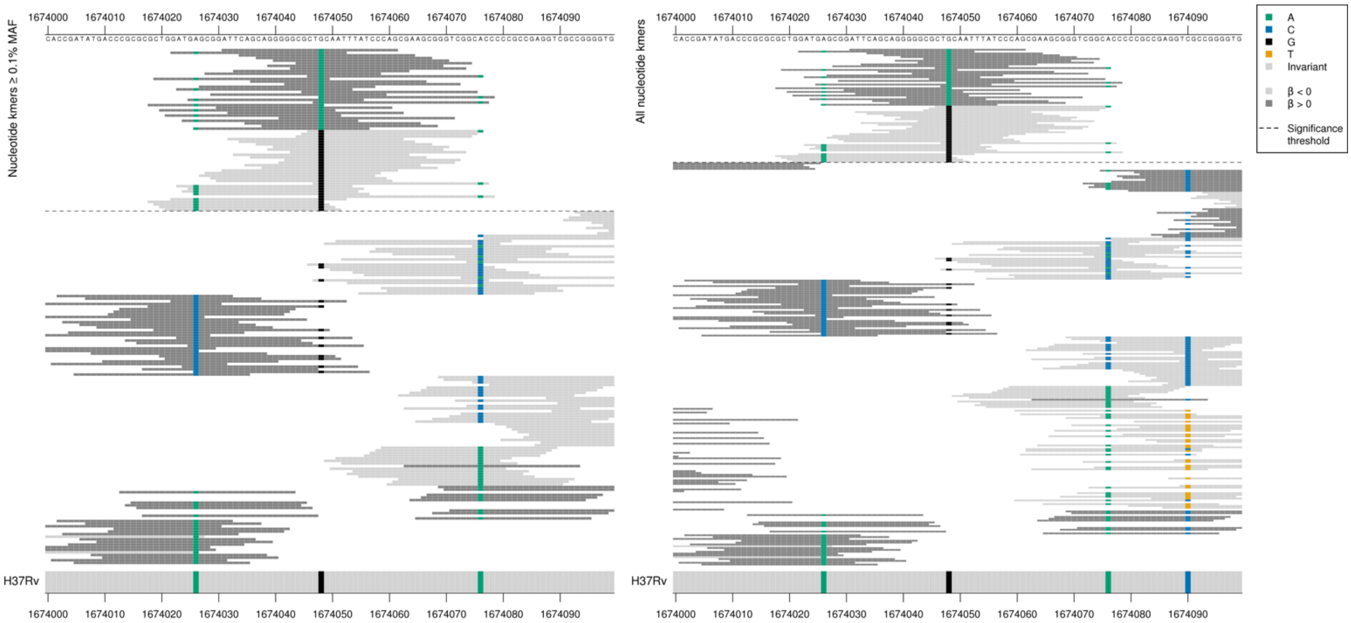

Synonymous mutation
High MIC-associated kmers = CTA = Leu / L
Low MIC-associated kmers = CTG = Leu / L

| Kmer | $-\log_{10} p$ | β | MAC | MAF | Ps | Variants |
|---|---|---|---|---|---|---|
| ATTCAGCAGGGGGCGCTACAATTTATCCCAG | 17.54 | 1.74 | 107 | 1.20 | 1674031 | 1674048/609A |
| CTGCAATTTATCCCAGCGAAGCGGGTCGGCA | 17.39 | -1.71 | 114 | 1.28 | 1674046 | 1674048/609G |

#### *rpoB*

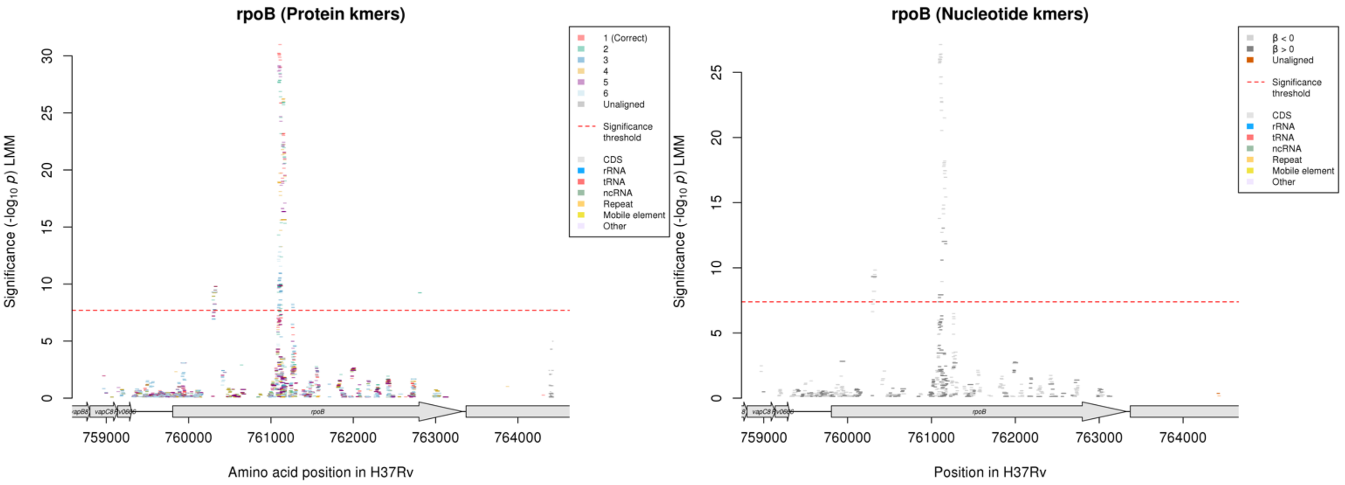

Protein kmers:

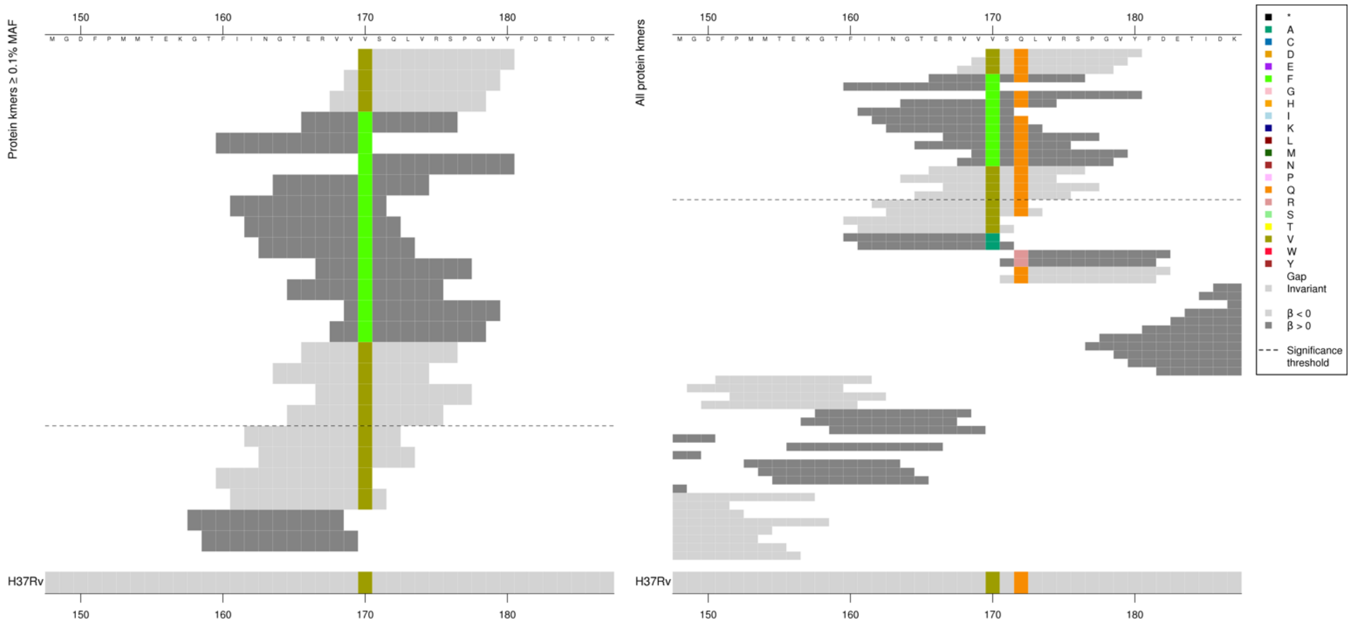

Codon at position 172 possibly also important. High MIC-associated kmers were all the same *p*-value. Low MIC-associated kmers were different *p*-values but they all covered codons 170-172.
Some kmers below significance threshold = 170A, 172R

| Kmer | $-\log_{10} p$ | β | MAC | MAF | Ps | Variants |
|---|---|---|---|---|---|---|
| VSQLVRSPGVY | 9.79 | -1.98 | 26 | 0.29 | 170 | 170V |
| ERVVFSQLVRS | 9.28 | 2.24 | 20 | 0.22 | 166 | 170F |

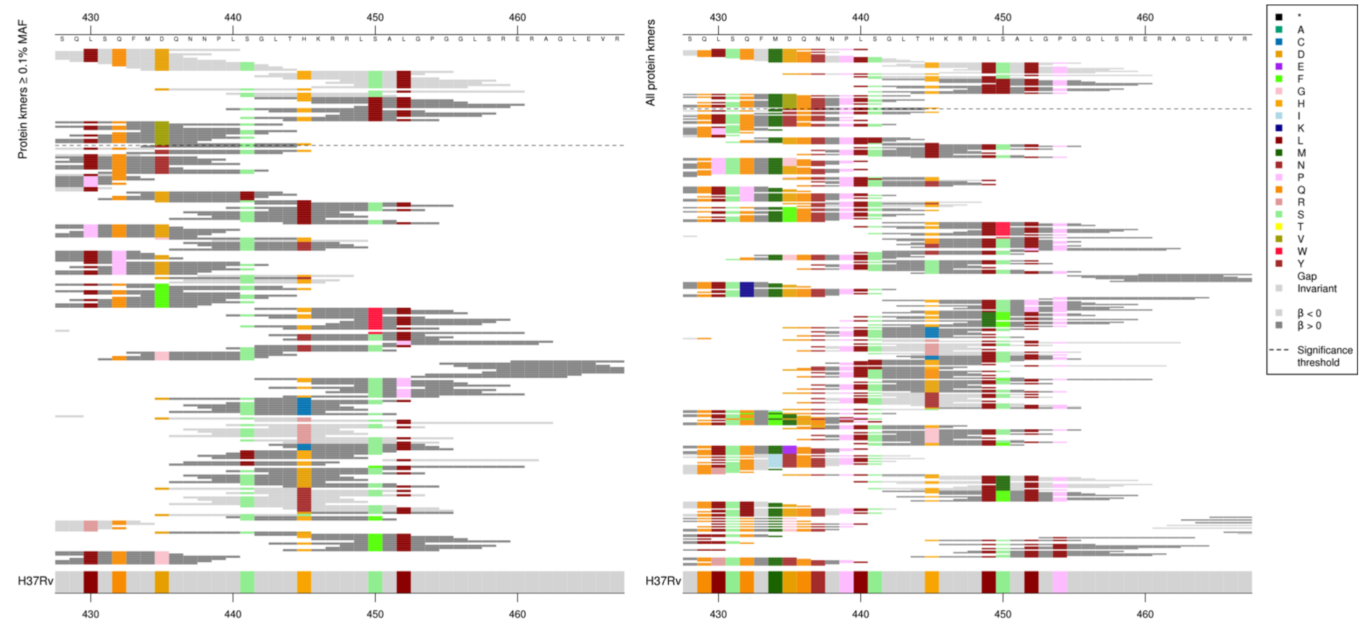

Main substitutions being captured were at codons 435 and 450. The low MIC-associated kmers capturing these had different *p*-values depending possibly on other substitutions in the region, likewise for the high MIC-associated kmers capturing 435V. The high MIC-associated kmers capturing 450L were all the same *p*-value and not affected by other substitutions.

| Kmer | $-\log_{10} p$ | β | MAC | MAF | Ps | Variants |
|---|---|---|---|---|---|---|
| LSQFMDQNNPL | 30.99 | -1.17 | 783 | 8.77 | 430 | 435D |
| HKRRLSALGPG | 23.20 | -0.83 | 2718 | 30.44 | 445 | 450S |
| GLTHKRRLLAL | 15.64 | 0.79 | 2182 | 24.44 | 442 | 450L |
| FGTSQLSQFMV | 9.88 | 0.94 | 461 | 5.16 | 425 | 435V |

#### *inhA*

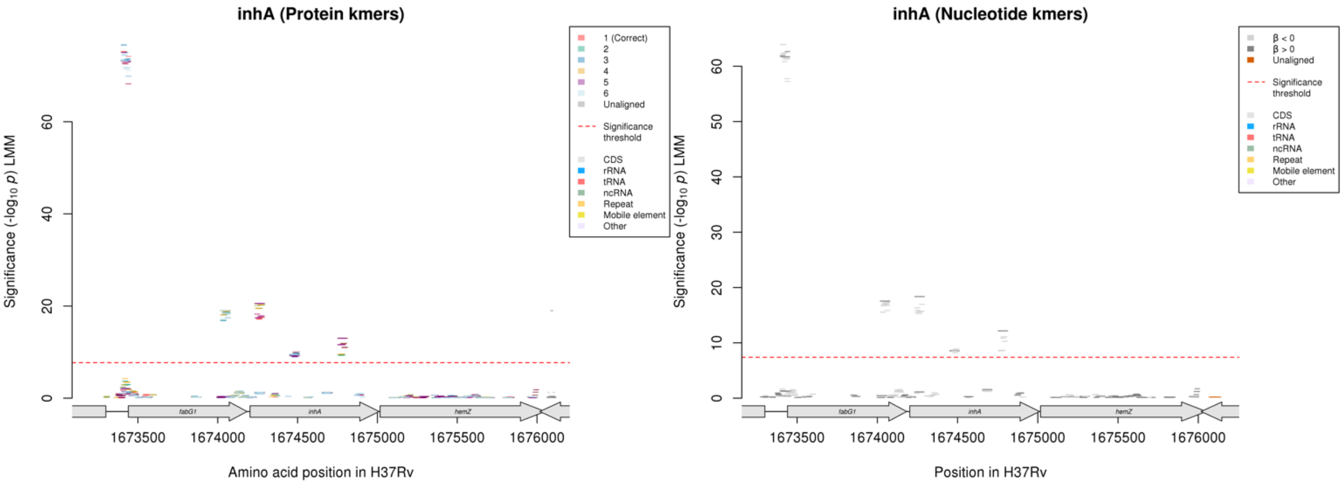

Protein kmers:

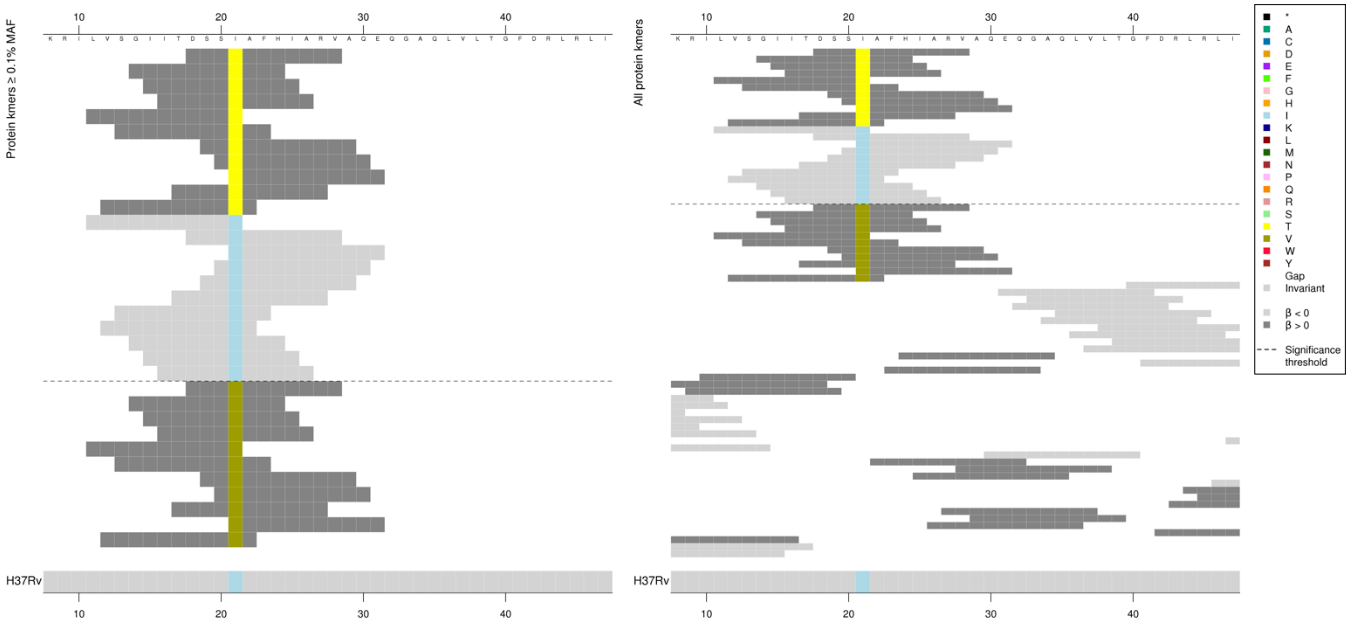

Kmers below significance threshold = 21V

| Kmer | -log$_{10}$ *p* | β | MAC | MAF | Ps | Variants |
|---|---|---|---|---|---|---|
| DSSTAFHIARV | 20.54 | 2.77 | 43 | 0.48 | 18 | 21T |
| LVSGIITDSSI | 18.26 | -2.13 | 58 | 0.65 | 11 | 21I |

Protein kmers:

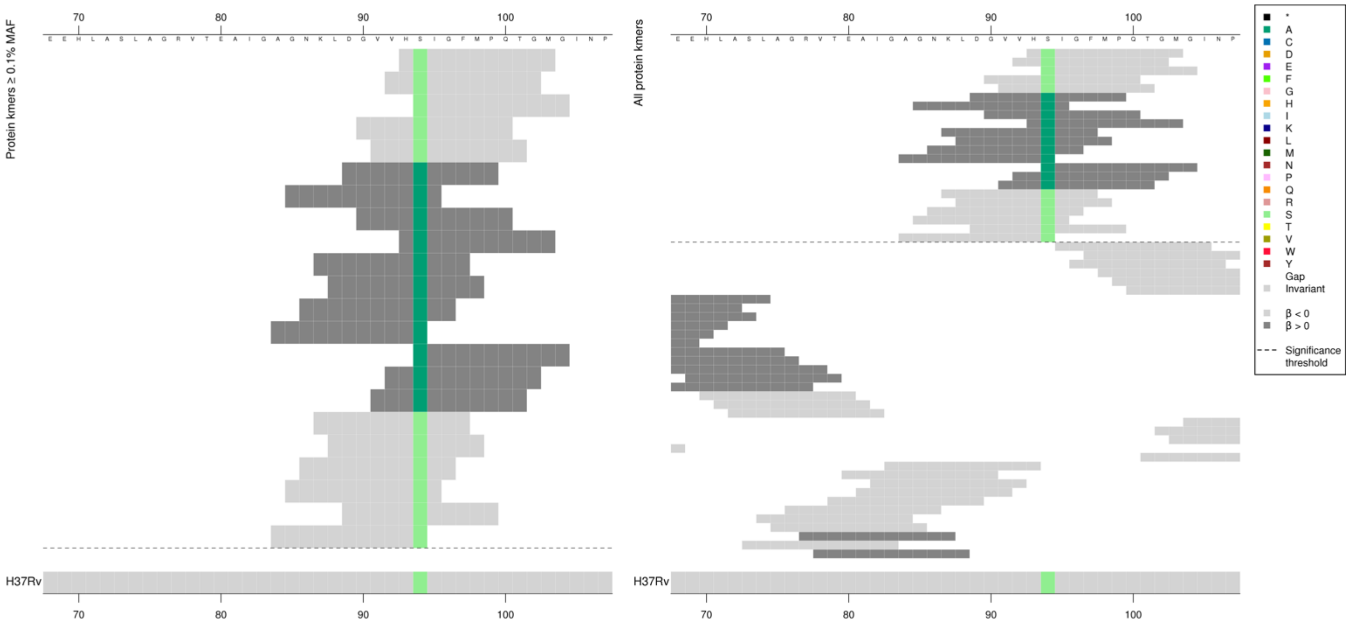

| Kmer | -log$_{10}$ *p* | β | MAC | MAF | Ps | Variants |
|---|---|---|---|---|---|---|
| HSIGFMPQTGM | 10.09 | -2.16 | 33 | 0.37 | 93 | 94S |
| DGVVHAIGFMP | 9.44 | 2.13 | 30 | 0.34 | 89 | 94A |

Protein kmers:

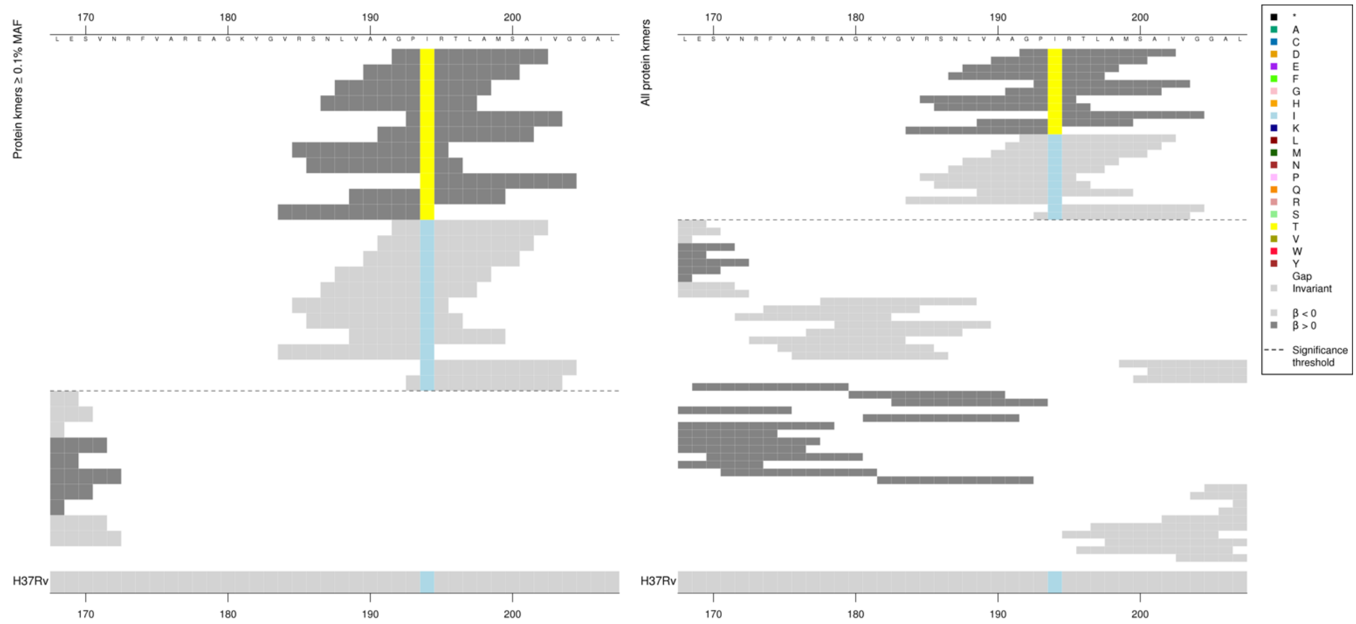

| Kmer | $-\log_{10} p$ | β | MAC | MAF | Ps | Variants |
|---|---|---|---|---|---|---|
| GPTRTLAMSAI | 12.99 | 2.32 | 46 | 0.52 | 192 | 194T |
| GPIRTLAMSAI | 11.85 | -2.23 | 46 | 0.52 | 192 | 194I |

#### *embB*

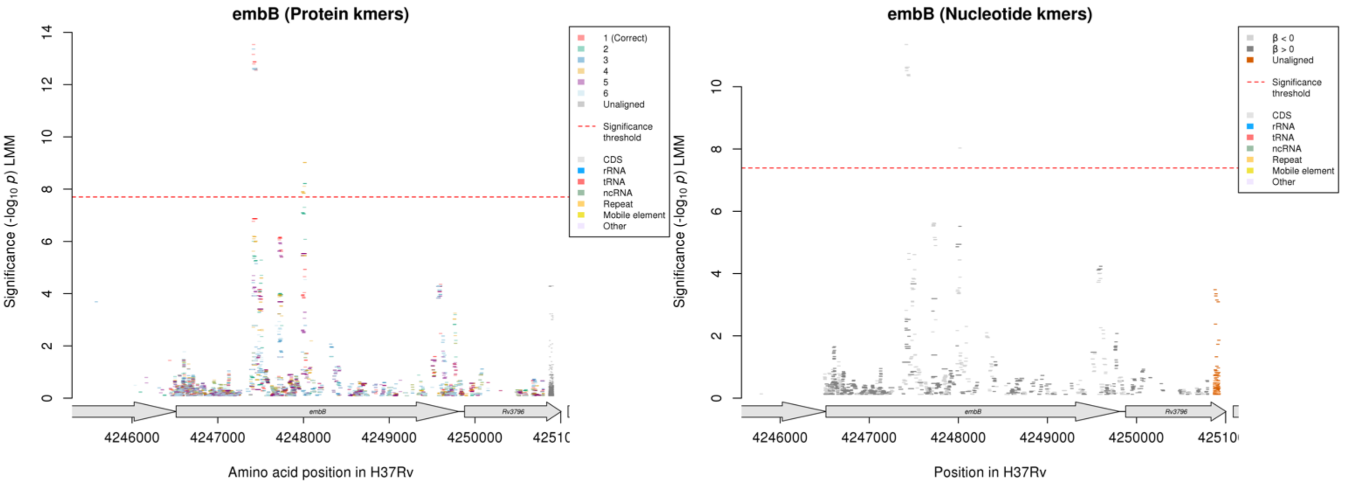

Only the first *embB* peak contains significant kmers aligning to the correct reading frame

Protein kmers:

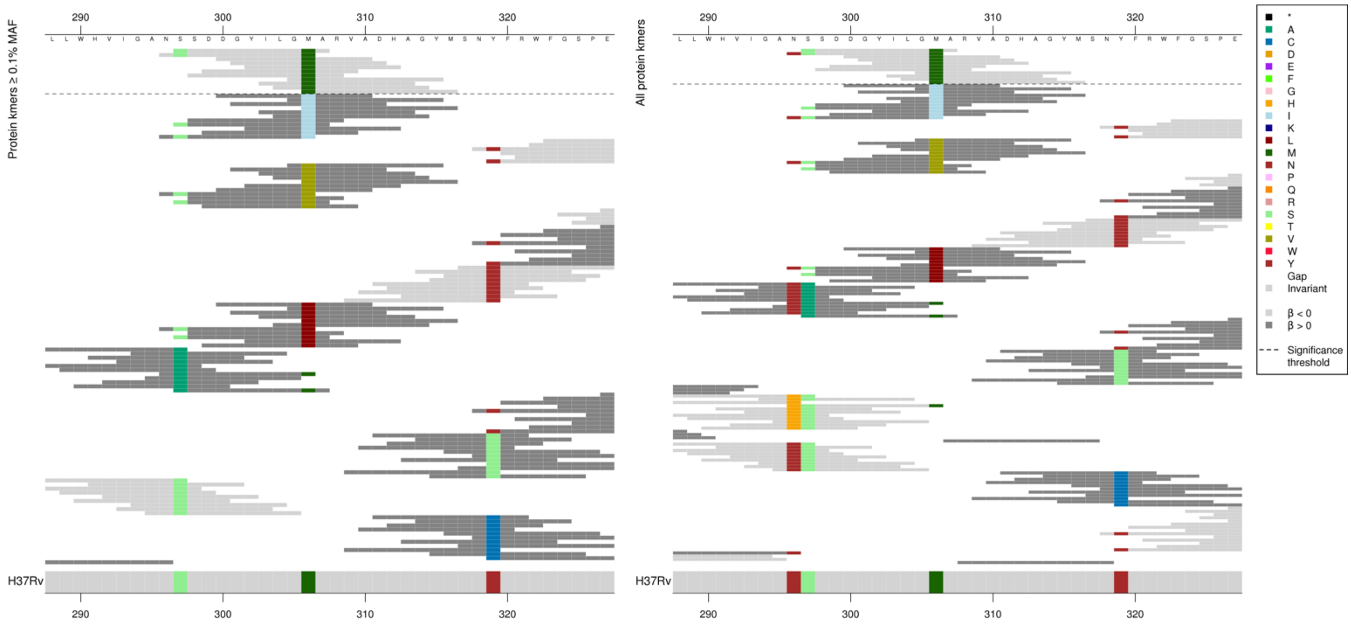

Mostly capturing the substitution at position 306, but the *p*-values appear to be affected by the other surrounding substitutions.
Kmers below significance threshold = Codon 296H, 297A, 306I/V/L

| Kmer | $-\log_{10} p$ | β | MAC | MAF | Ps | Variants |
|---|---|---|---|---|---|---|
| SSDDGYILGMA | 13.53 | -0.71 | 1752 | 19.62 | 297 | 306M |

Nucleotide kmers:

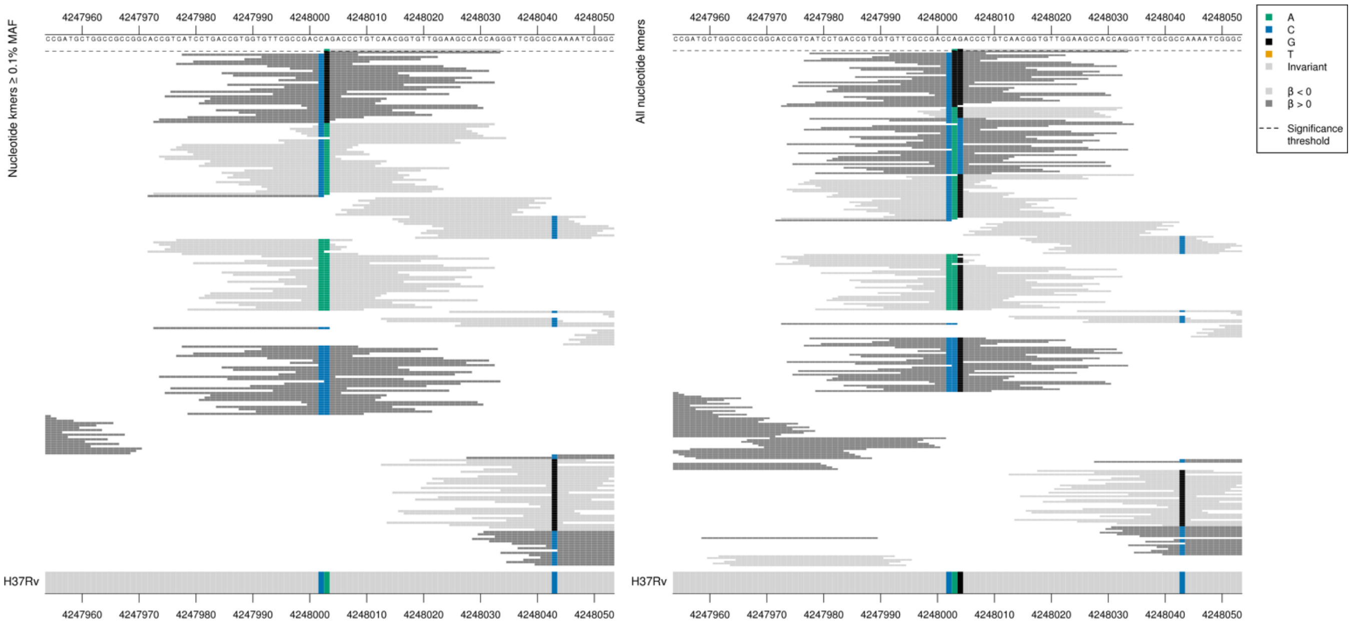

Codon 497
Low MIC-associated kmers = CAG = Gln / Q
Kmers below significance threshold = CGG = Arg / R
Kmers below significance threshold = CAC = His / H
Kmers below significance threshold = CCG = Pro / P
Kmers below significance threshold = AAG = Lys / K

| Kmer | $-\log_{10} p$ | β | MAC | MAF | Ps | Variants |
|---|---|---|---|---|---|---|
| AGACCCTGTCAACGGTGTTGGAAGCCACCAG | 8.03 | -0.87 | 299 | 3.35 | 4248003 | 4248003/1490A, 4248004/1491G both codon 497 |

## *Rv1139c:Rv1140*

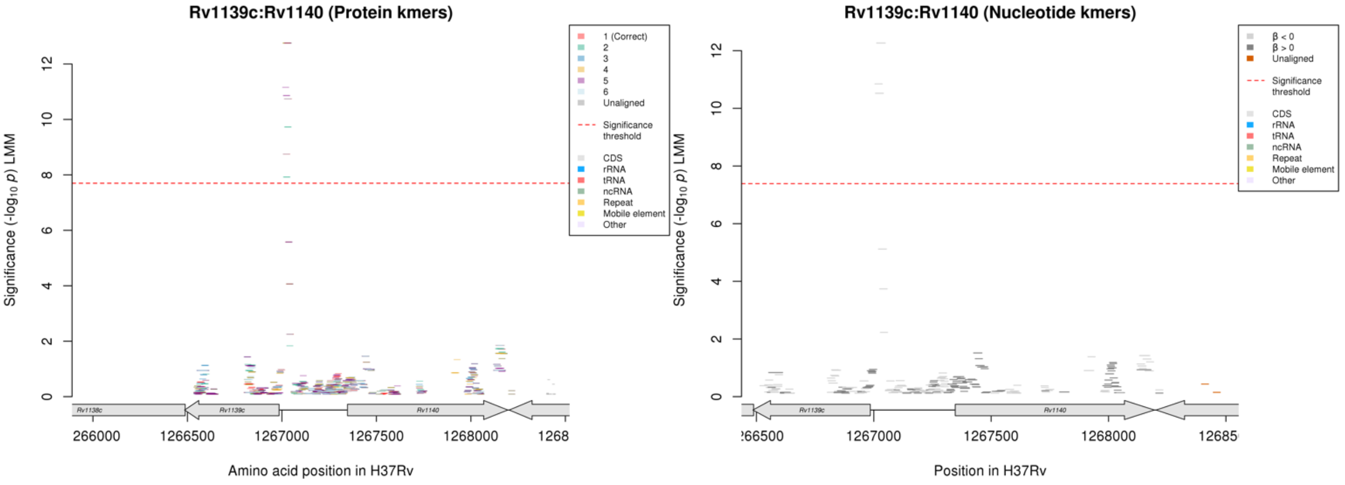

Nucleotide kmers:

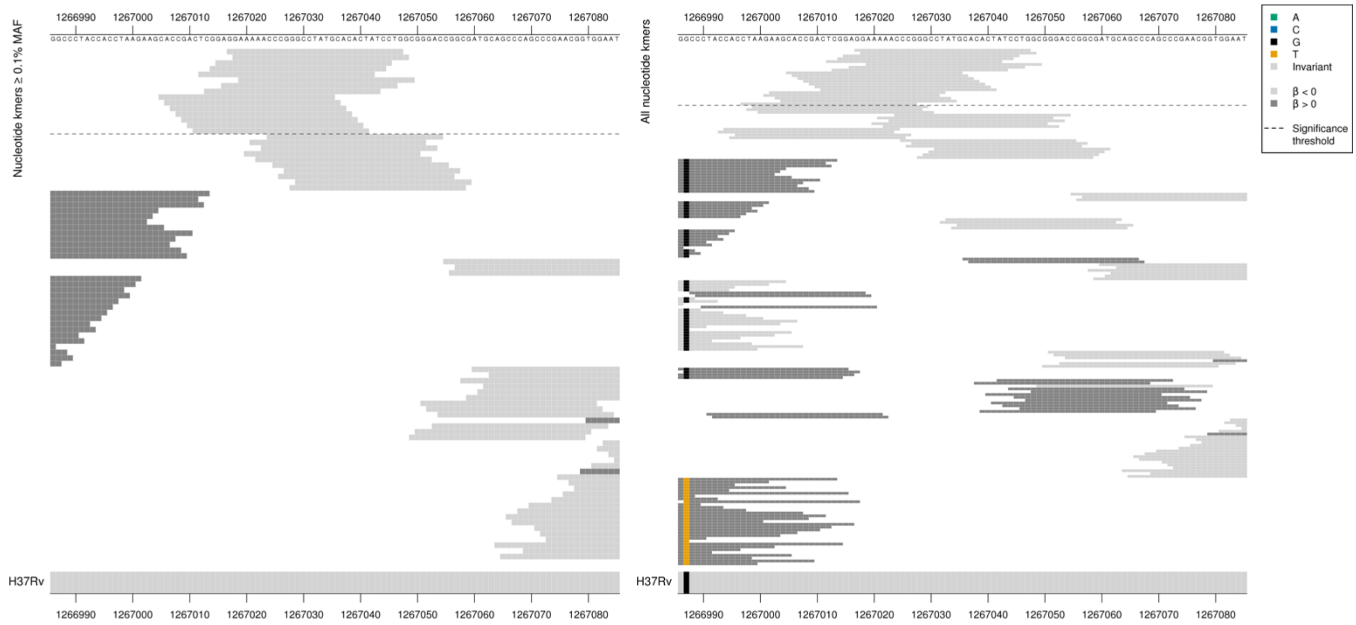

Removing the threshold requiring a kmer to be aligned to a gene/intergenic region in at least five genomes across the full dataset. Showing the nucleotide kmer alignment after lowering the threshold to just one genome:

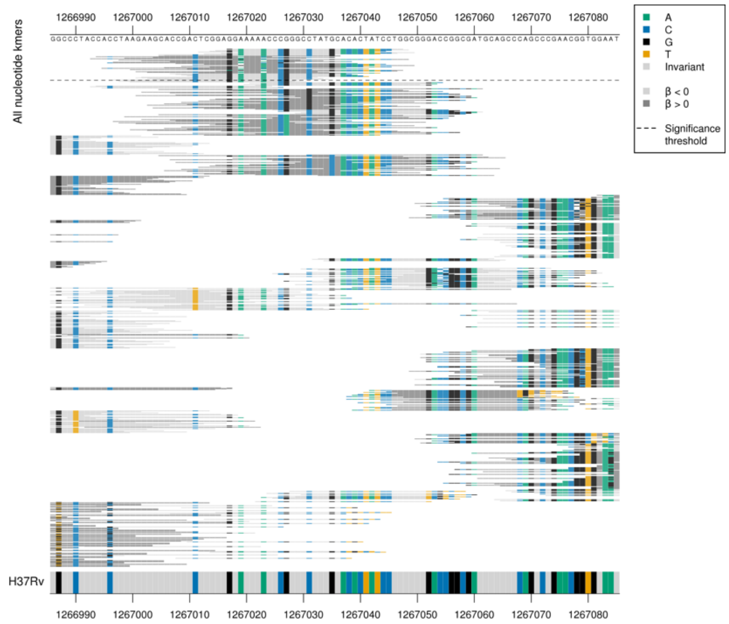

| Kmer | $-\log_{10} p$ | β | MAC | MAF | Ps | Variants |
|---|---|---|---|---|---|---|
| GGAAAAACCCGGGCCTATGCACACTATCCTG | 12.26 | -4.30 | 10 | 0.11 | 1267017 | WT covering positions 1267001-1267049 |

## *Rv1158c*

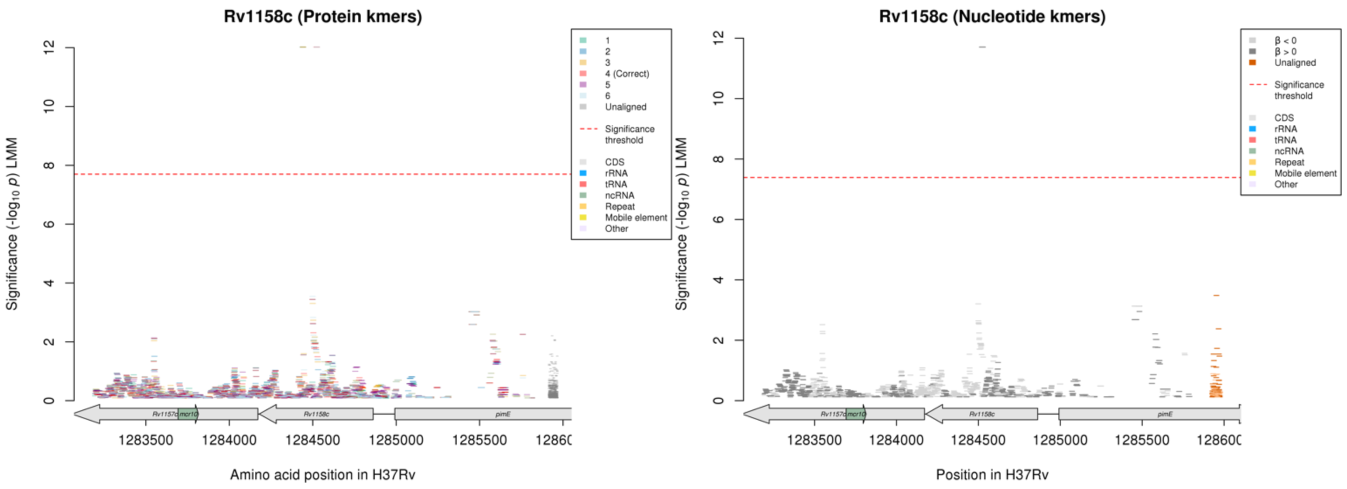

#### Protein kmers:

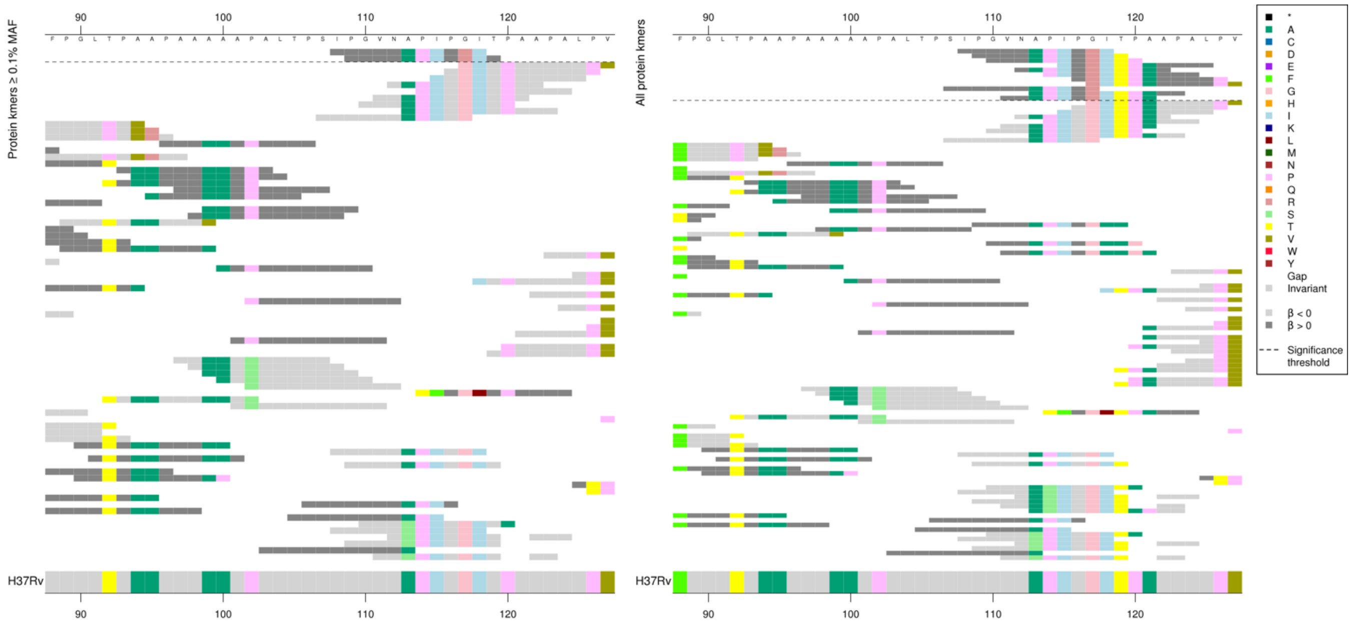

#### Nucleotide kmers in the same region:

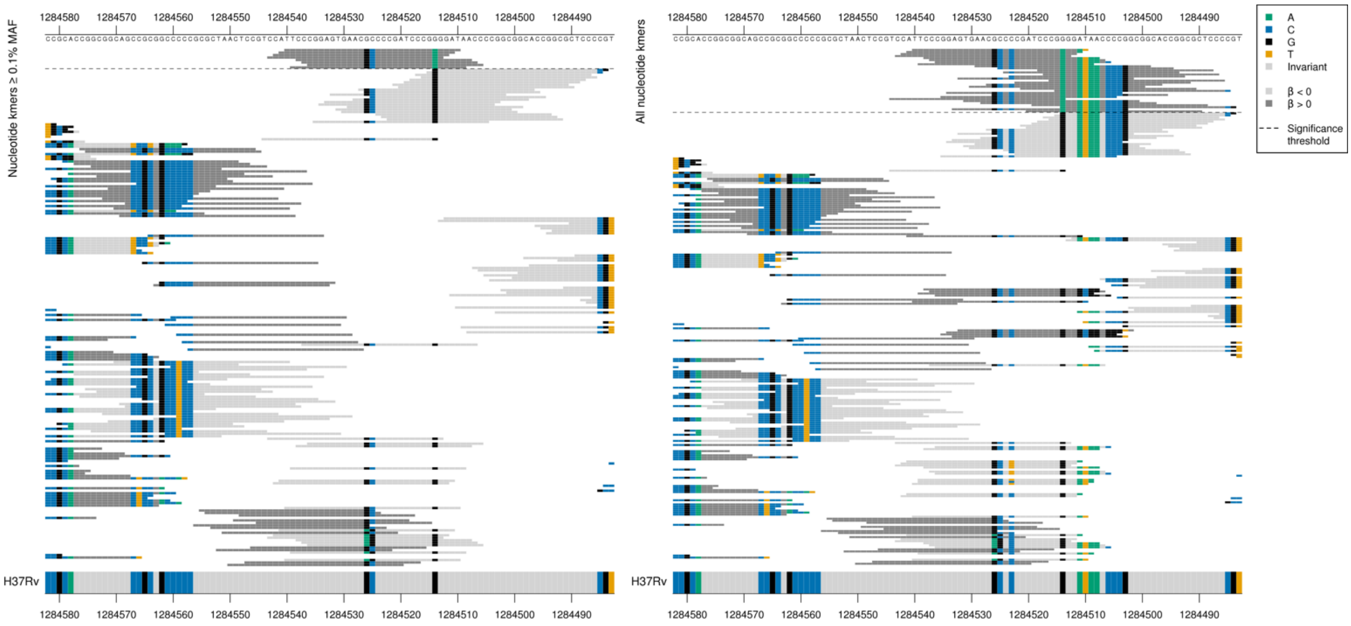

| Kmer | $-\log_{10} p$ | β | MAC | MAF | Ps | Variants |
|---|---|---|---|---|---|---|
| IPGVNAPIPRI | 12.02 | 3.52 | 9 | 0.10 | 108 | 117R |

#### *rpsL*

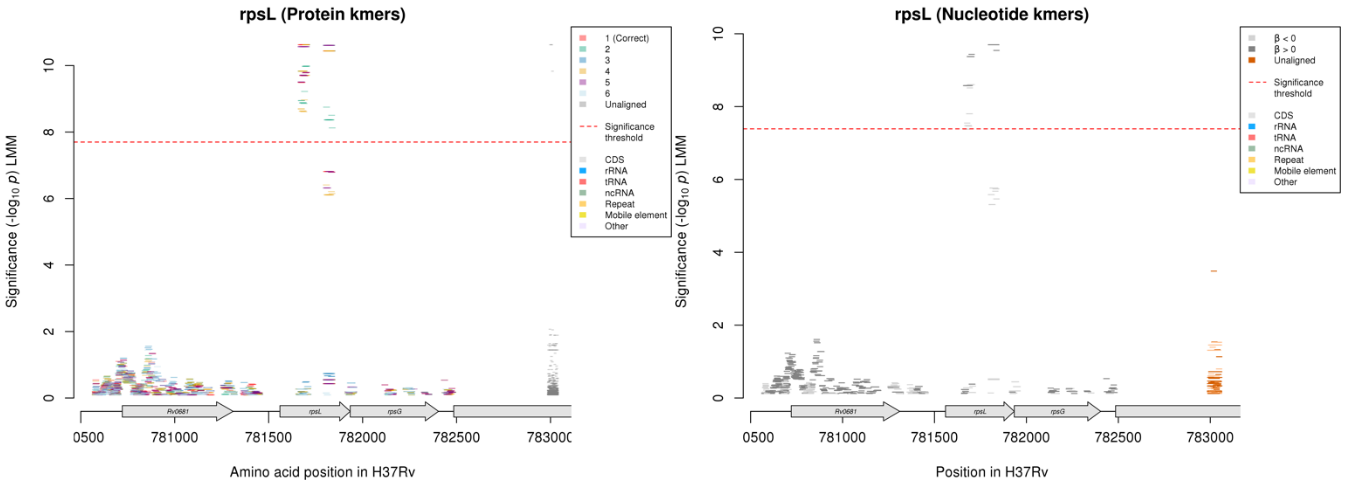

Protein kmers:

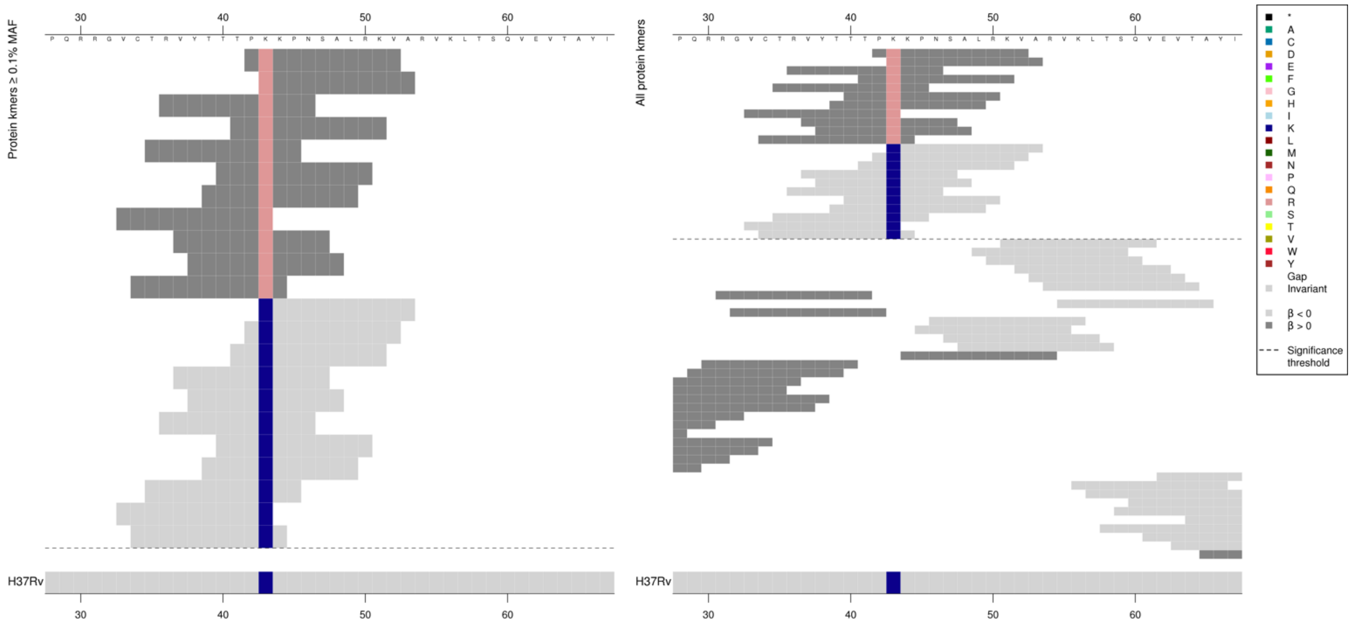

| Kmer | $-\log_{10} p$ | β | MAC | MAF | Ps | Variants |
|---|---|---|---|---|---|---|
| PRKPNSALRKV | 10.63 | 0.77 | 1553 | 17.39 | 42 | 42R |
| KKPNSALRKVA | 9.79 | -0.74 | 1553 | 17.39 | 43 | 42K |

Protein kmers:

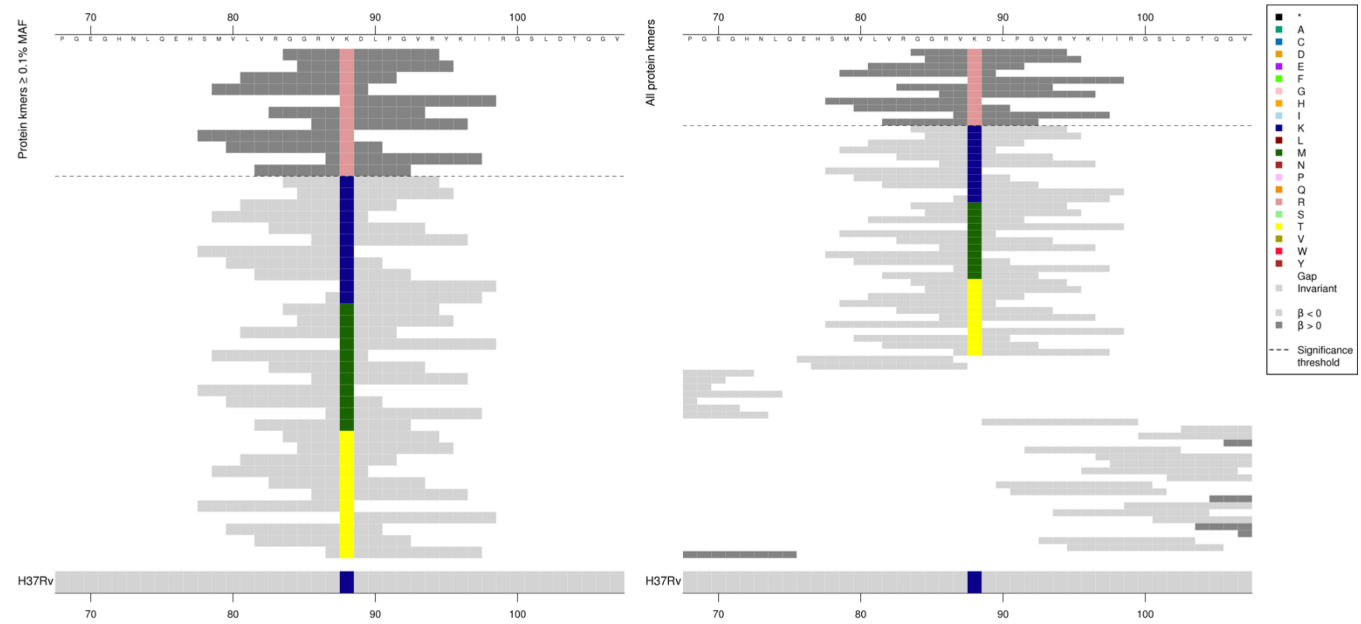

Kmers below significance threshold = 88K/M/T

| Kmer | -log$_{10}$ *p* | β | MAC | MAF | Ps | Variants |
|---|---|---|---|---|---|---|
| GGRVRDLPGVR | 10.44 | 1.10 | 223 | 2.50 | 84 | 88R |

# *Rv1219c*

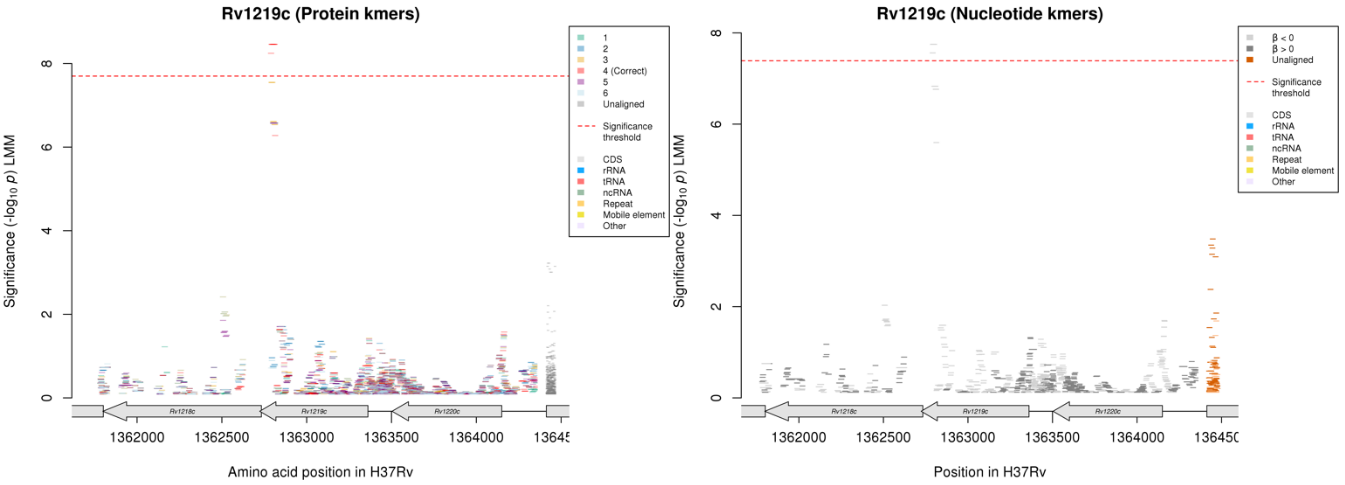

*Rv1219c* is on the reverse strand, begins at position 1363361 and ends at position 1362723, 213 amino acids long. *Rv1219c* is a transcriptional regulator.

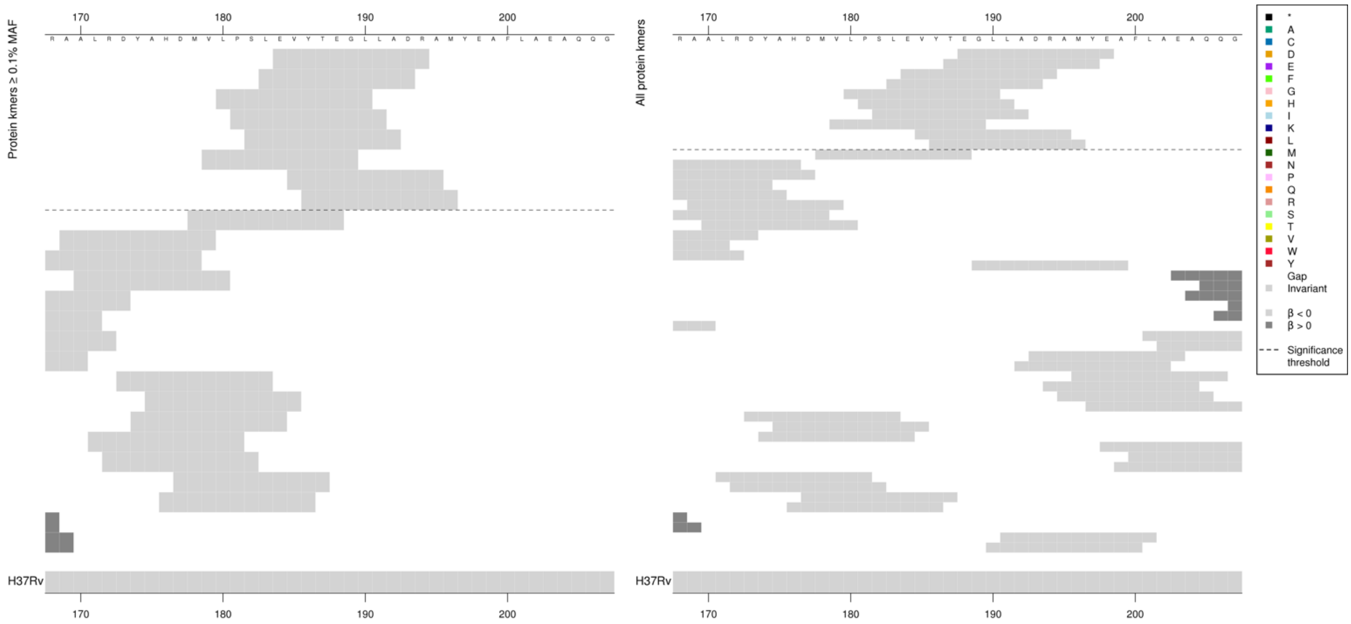

Removing the threshold requiring a kmer to be aligned to a gene/intergenic region in at least five genomes across the full dataset. Showing the protein kmer alignment after lowering the threshold to just one genome:

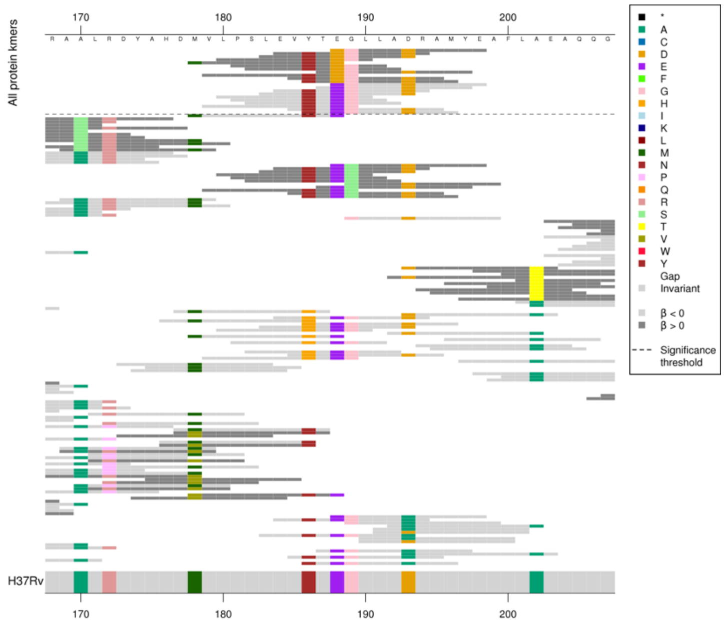

Significant reference-sequence low MIC-associated kmers, plus rare alternative alleles in the region. Just showing those above the threshold of five in the tables.
Most significant kmer in this region was very low MAF, so also showing the result for the top kmer above 0.1% MAF.

| Kmer | $-\log_{10} p$ | β | MAC | MAF | Ps | Variants |
|---|---|---|---|---|---|---|
| EVYTEGLLADR | 8.46 | -3.63 | 10 | 0.11 | 184 | WT codons 179-195 |

### *ftsK/Rv2749*

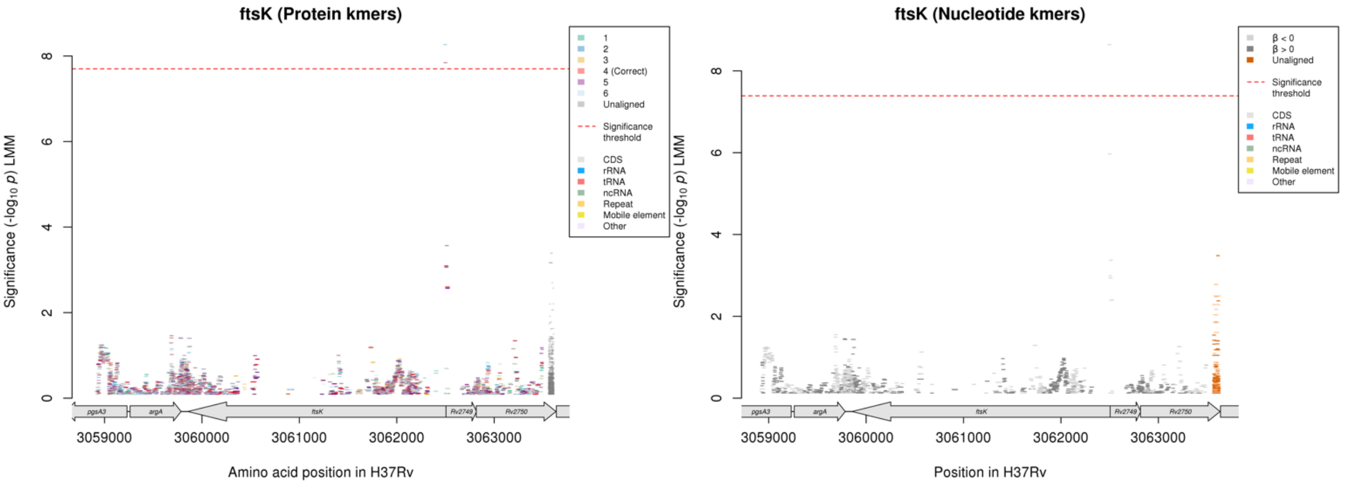

*ftsK*:

Protein kmers:

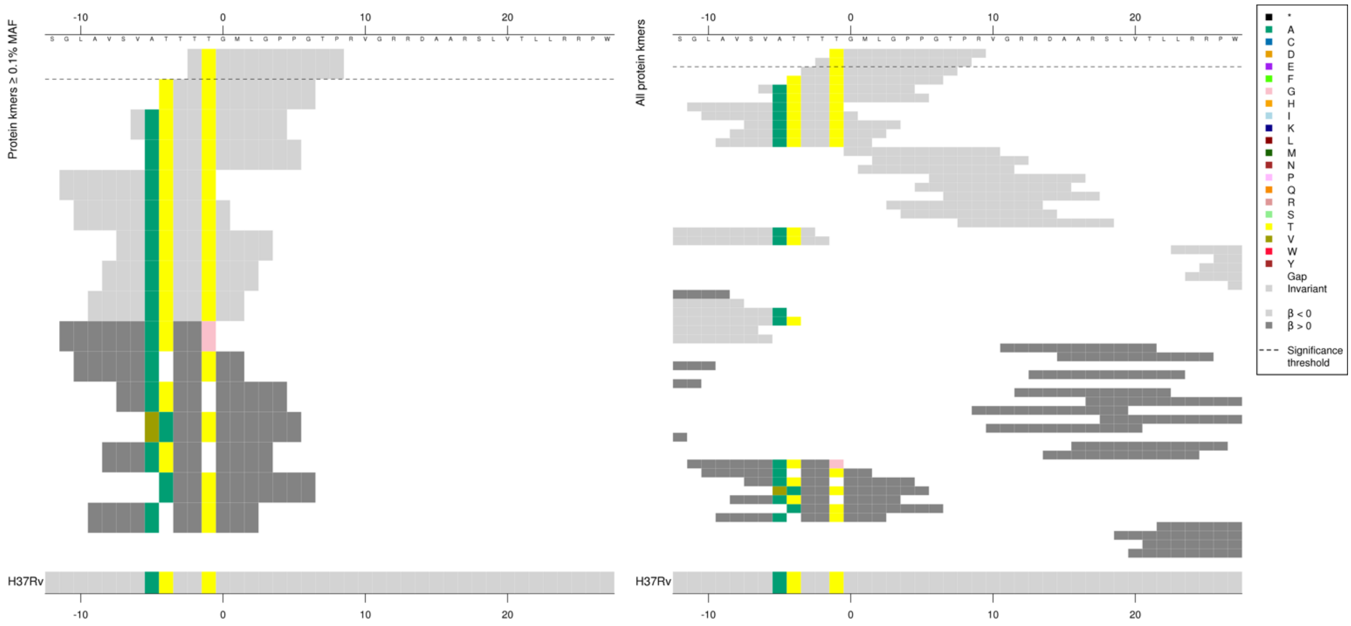

Same region for nucleotide kmers:

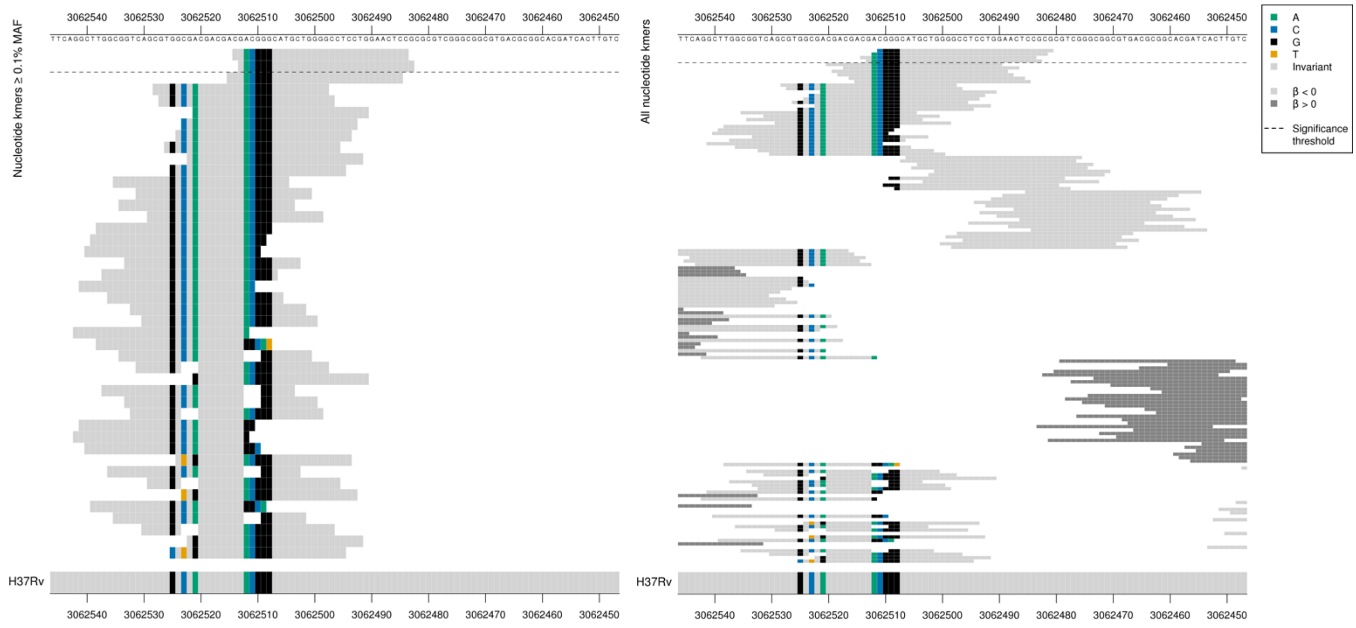

*Rv2749*:

Protein kmers:

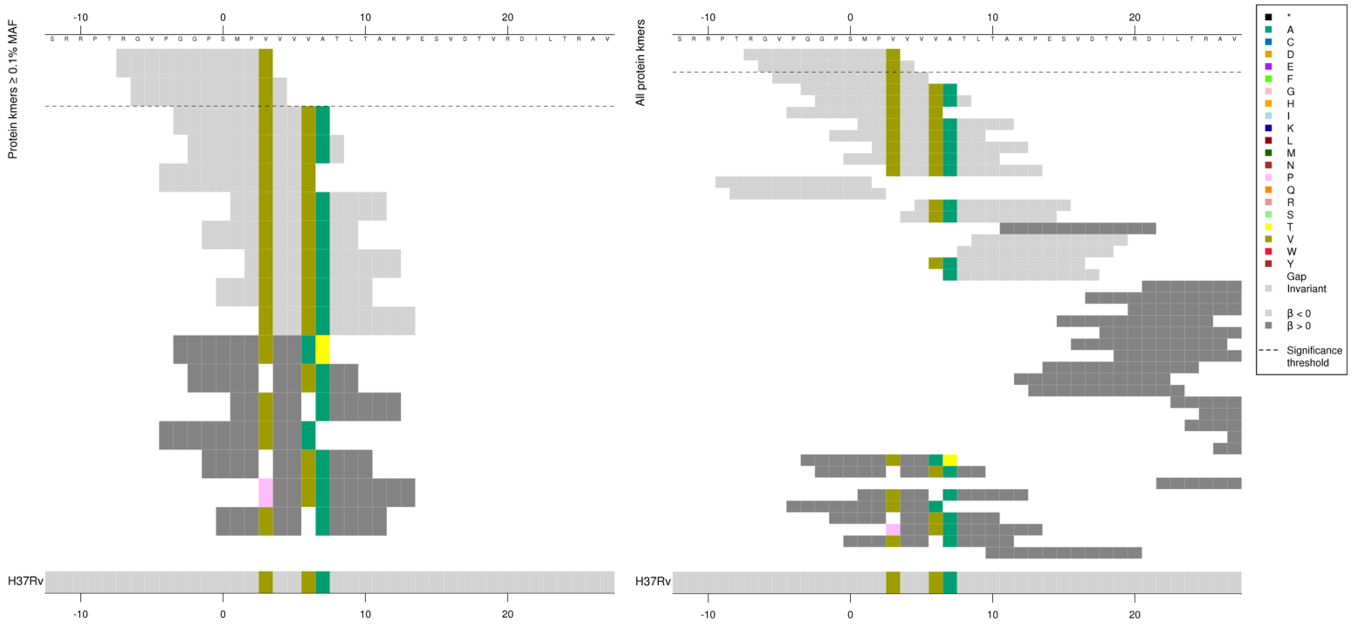

Same region for nucleotide kmers:

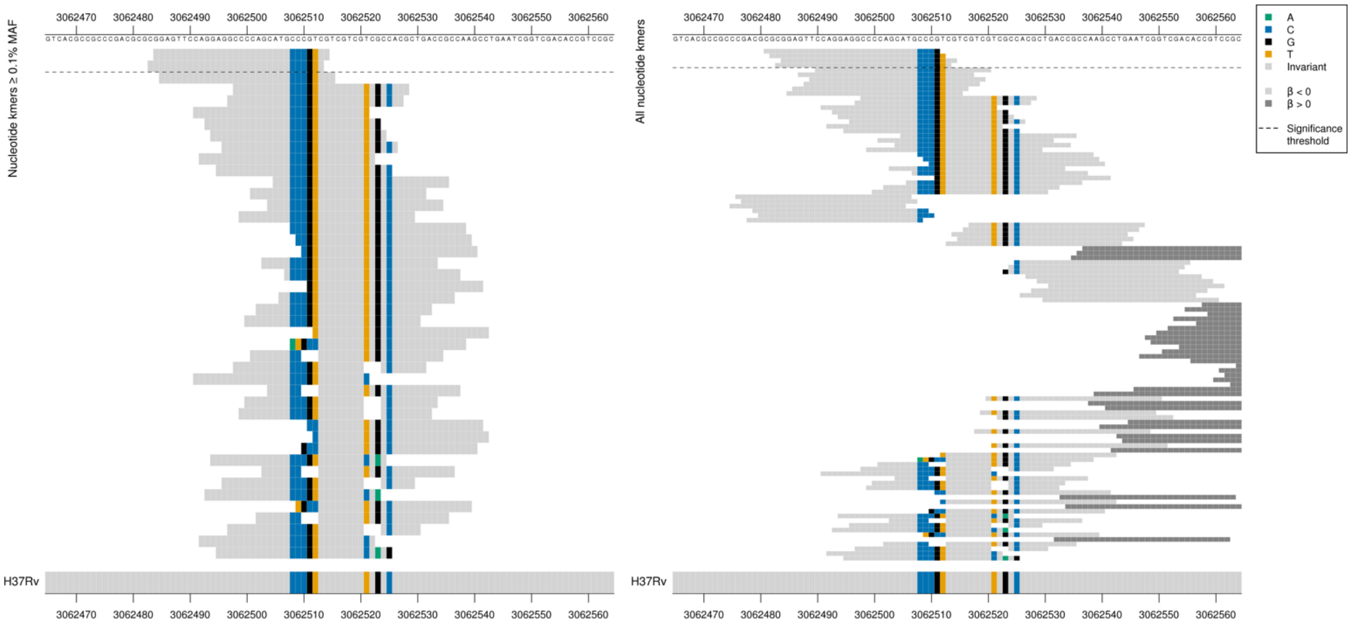

*ftsK* begins at position 3062506, *Rv2749* begins at position 3062505.
Kmers overlap both genes. In high MIC (not significant) kmers = deletion of 3bp, ‘CGT’ on the forward strand, in a region of four repeats of ‘CGT’ positions 3063511-3063518. This would result in a deletion of codon 3 for *Rv2749*. Some high MIC (not significant) kmers had a deletion of ‘TCG’ on the forward strand at positions 3063521-3063523, but the significant low MIC-associated kmers did not cover this region.

Kmers were capturing wild type at the positions 3063511-3063518, just upstream of the *ftsK* start codon, just after the start codon for *Rv2749*.

| Kmer | $-\log_{10} p$ | β | MAC | MAF | Ps | Variants |
|---|---|---|---|---|---|---|
| CGACGGGCATGCTGGGGCCTCCTGGAACTCC | 8.64 | -3.72 | 9 | 0.10 | 3062514 | WT around start codon of ftsK and Rv2749 |

#### *gid*

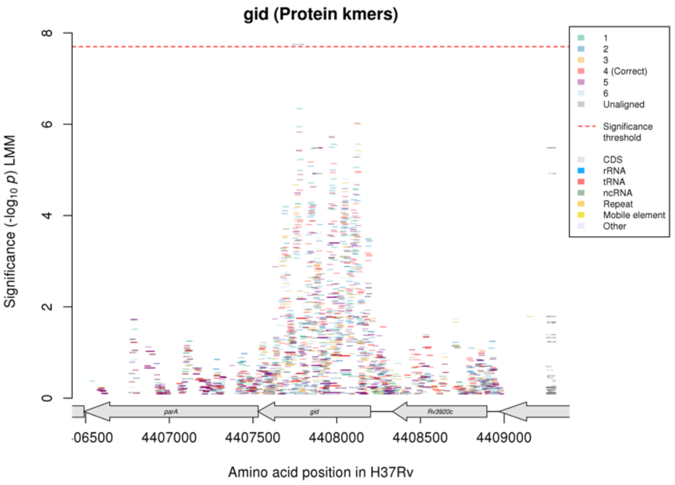

Not significant in nucleotide kmer gwas.

Protein kmers:

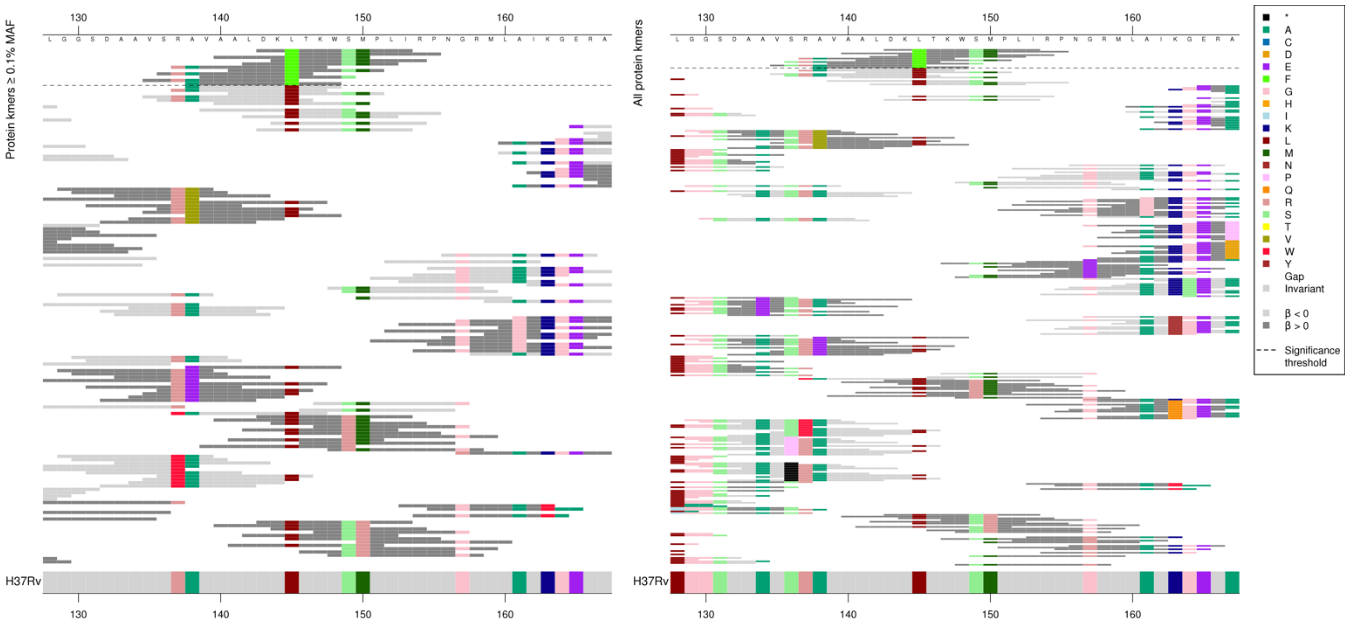

| Kmer | $-\log_{10} p$ | β | MAC | MAF | Ps | Variants |
|---|---|---|---|---|---|---|
| DKFTKWSMPLI | 7.74 | 2.33 | 18 | 0.20 | 143 | 145F |

### Kanamycin (KAN)

#### *rrs*

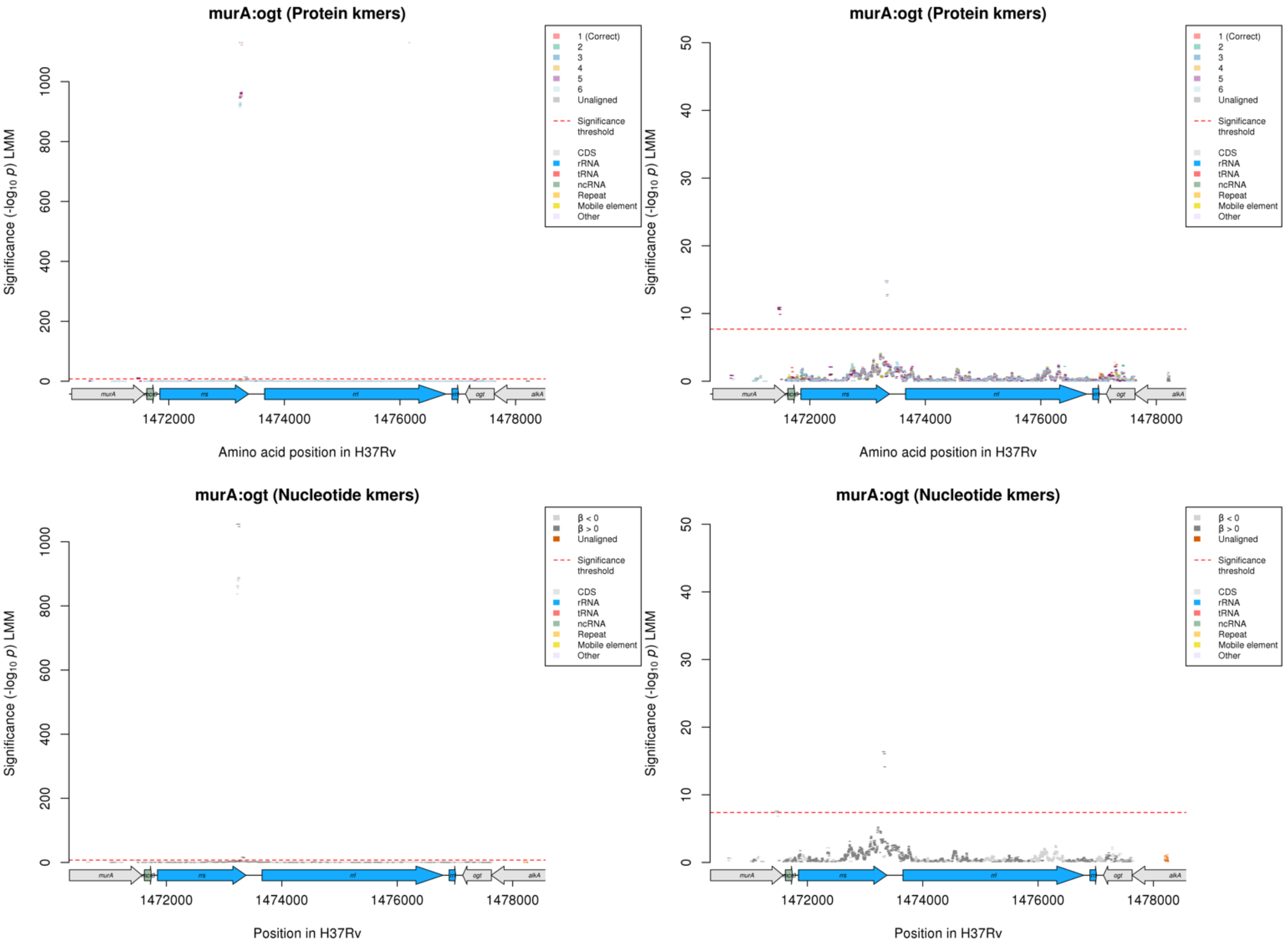

Nucleotide kmers:

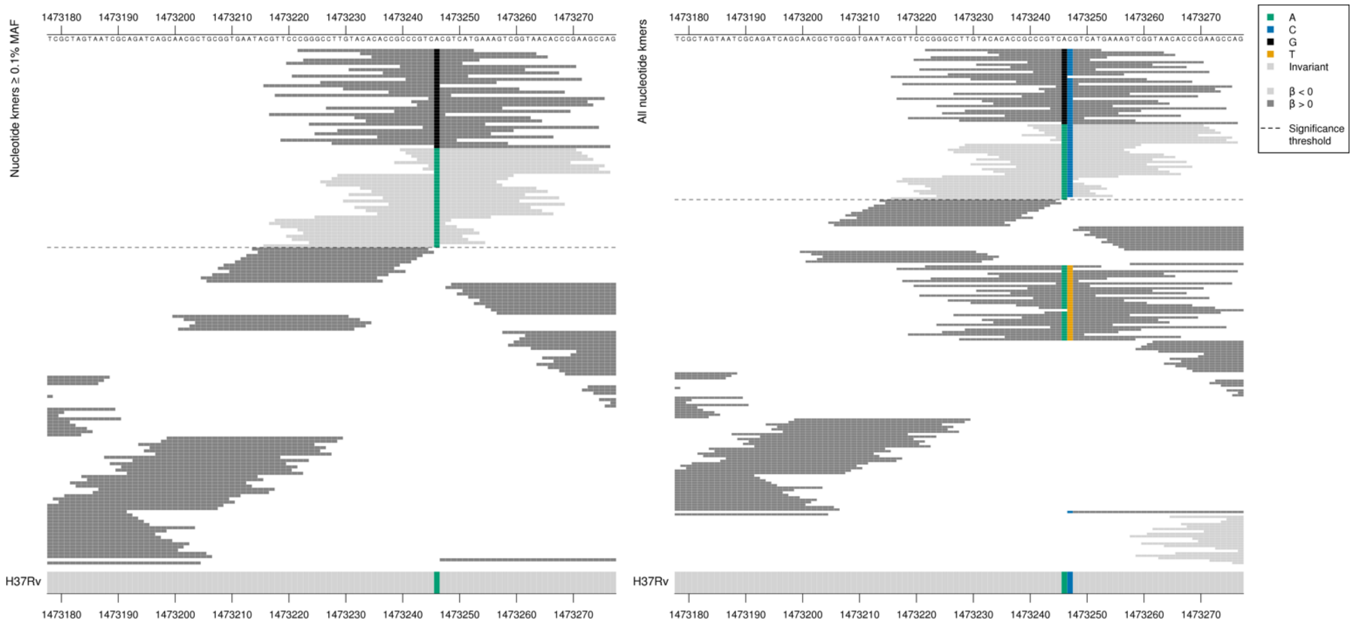

These kmers were capturing low MIC-associated kmers containing an A at position 1473246/1401 in *rrs* and high MIC-associated kmers containing a G at position 1473246.

There were low frequency kmers capturing an additional variant at position 1402 in *rrs* that were not significant. This variant did possibly impact the *p*-values of the significant kmers. All but the last high MIC-associated kmer were the same *p*-value, so significance was not affected by covering position 1402. The low MIC-associated kmers covered a range of *p*-values.

| Kmer | $-\log_{10} p$ | β | MAC | MAF | Ps | Variants |
|---|---|---|---|---|---|---|
| CGGGCCTTGTACACACCGCCCGTCGCGTCAT | 1054.65 | 3.83 | 536 | 6.13 | 1473222 | 1473246/1401G, 1473247/1402C |
| CCCGTCACGTCATGAAAGTCGGTAACACCCG | 889.02 | -3.50 | 589 | 6.73 | 1473240 | 1473246/1401A, 1473247/1402C |
| CGTTCCCGGGCCTTGTACACACCGCCCGTCA | 837.23 | -3.41 | 594 | 6.79 | 1473216 | 1473246/1401A |

Nucleotide kmers:

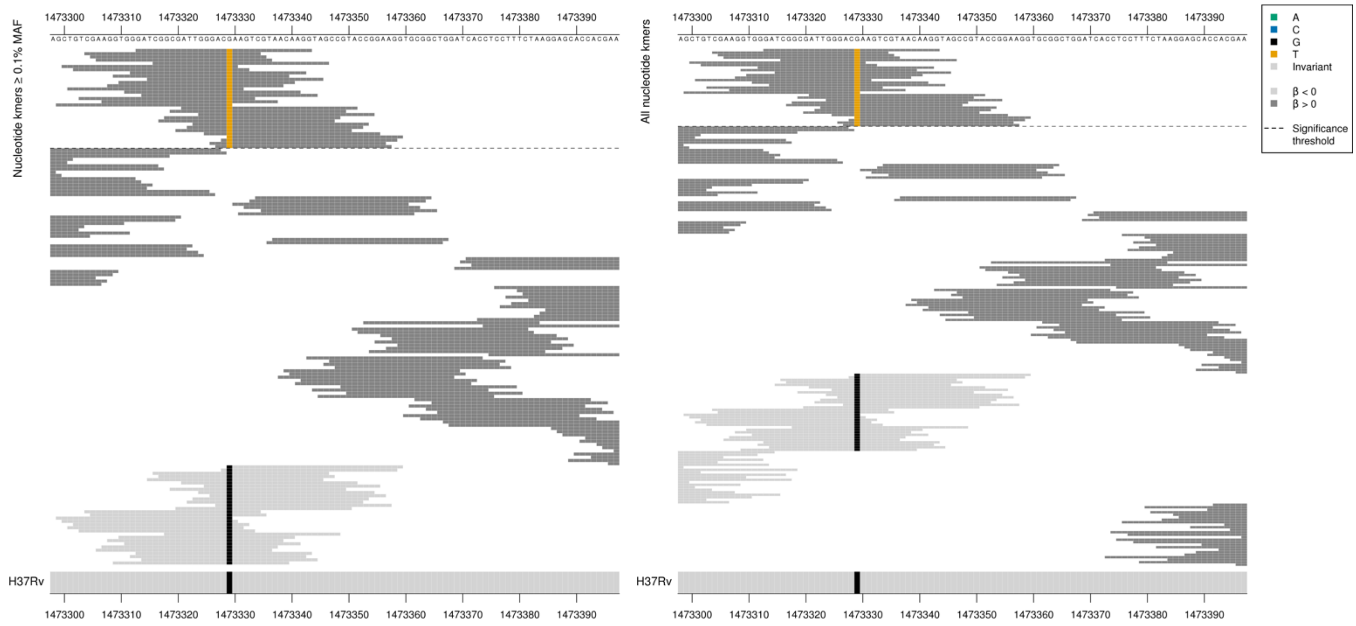

| Kmer | $-\log_{10} p$ | β | MAC | MAF | Ps | Variants |
|---|---|---|---|---|---|---|
| GATCGGCGATTGGGACTAAGTCGTAACAAGG | 16.38 | 2.26 | 11 | 0.13 | 1473313 | 1473329/1484T |

#### *eis*

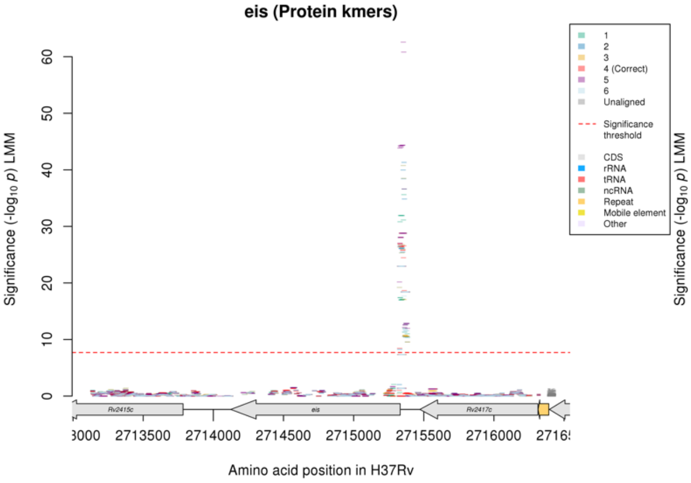

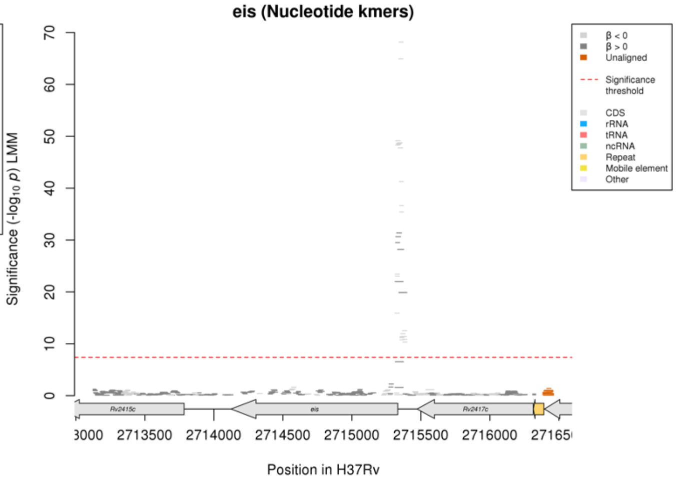

Nucleotide kmers:

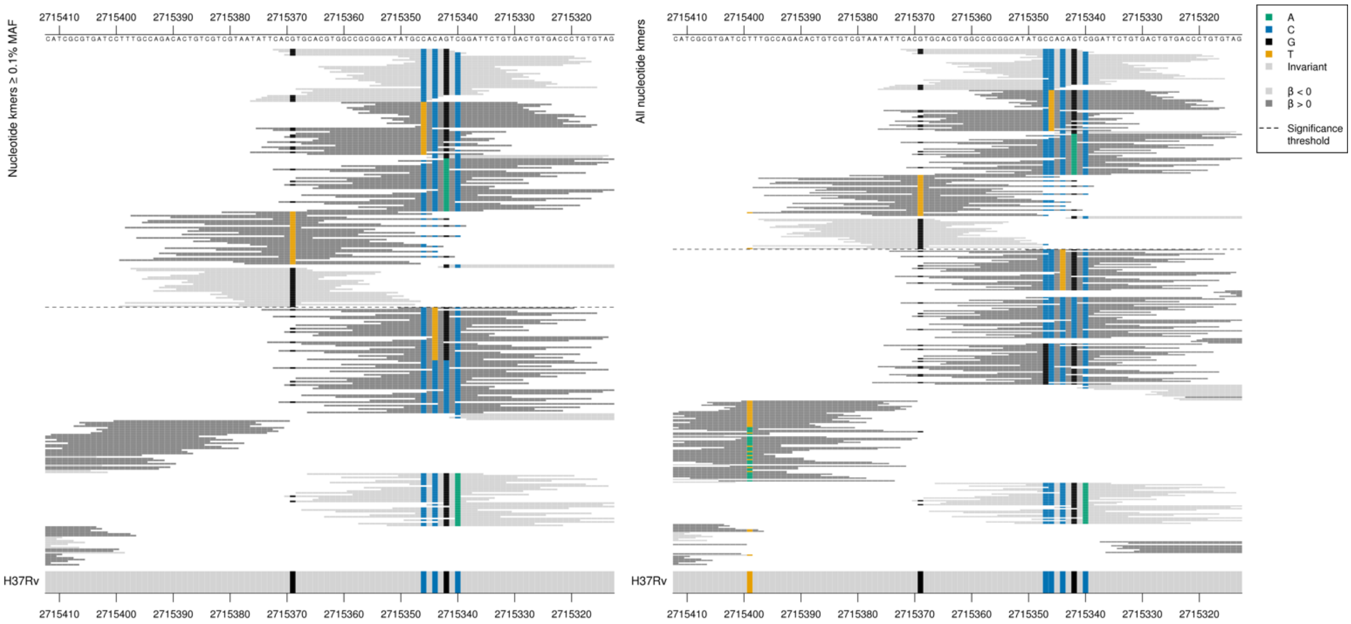

*eis* begins at position 2715332, capturing mutations just upstream of the start codon.

| Kmer | $-\log_{10} p$ | β | MAC | MAF | Ps | Variants |
|---|---|---|---|---|---|---|
| CACGTGCACGTGGCCGCGGCATATGCCACAG | 68.16 | -1.40 | 220 | 2.51 | 2715372 | WT / 2715369G, 2715347C, 2715346C, 2715344C, 2715342G |
| GTGCACGTGGCCGCGGCATATGCCACAGTCG | 64.93 | -1.32 | 242 | 2.77 | 2715369 | WT / 2715369G, 2715347C, 2715346C, 2715344C, 2715342G, 2715340C |
| ATGCCACAGTCGGATTCTGTGACTGTGACCC | 49.14 | -1.20 | 219 | 2.50 | 2715350 | WT / 2715347C, 2715346C, 2715344C, 2715342G, 2715340C |
| CACAGTCGGATTCTGTGACTGTGACCCTGTG | 48.31 | -1.21 | 214 | 2.45 | 2715346 | WT / 2715346C, 2715344C, 2715342G, 2715340C |
| TCACGTGCACGTGGCCGCGGCATATGCCACA | 41.28 | -1.28 | 153 | 1.75 | 2715373 | WT / 2715369G, 2715347C, 2715346C, 2715344C |
| ATTCACGTGCACGTGGCCGCGGCATATGCCA | 36.64 | -1.50 | 75 | 0.86 | 2715375 | WT / 2715369G, 2715347C, 2715346C |
| GCCGCGGCATATGCTACAGTCGGATTCTGTG | 31.36 | 1.79 | 40 | 0.46 | 2715360 | 2715346T [+ 2715347C, 2715344C, 2715342G, 2715340C] |
| TACAGTCGGATTCTGTGACTGTGACCCTGTG | 29.51 | 1.76 | 39 | 0.45 | 2715346 | 2715346T [+ 2715344C, 2715342G, 2715340C] |
| ATTCACGTGCACGTGGCCGCGGCATATGCTA | 28.20 | 1.66 | 42 | 0.48 | 2715375 | 2715346T [+2715347C] |
| CAGTCGGATTCTGTGACTGTGACCCTGTGTA | 23.42 | -0.91 | 176 | 2.01 | 2715344 | WT / 2715344C, 2715342G, 2715340C |
| ATCGGATTCTGTGACTGTGACCCTGTGTAGC | 22.00 | 1.40 | 49 | 0.56 | 2715342 | 2715342A |
| CGTAATATTCACTTGCACGTGGCCGCGGCAT | 19.87 | 1.84 | 23 | 0.26 | 2715381 | 2715369T |
| GTCGGATTCTGTGACTGTGACCCTGTGTAGC | 15.91 | -0.89 | 99 | 1.13 | 2715342 | WT / 2715342G, 2715340C |
| TGCCAGACACTGTCGTCGTAATATTCACGTG | 12.53 | -1.33 | 30 | 0.34 | 2715397 | WT / 2715369G |
| TTTGCCAGACACTGTCGTCGTAATATTCACG | 10.32 | -1.10 | 37 | 0.42 | 2715399 | WT / 2715399T, 2715369G |

#### *gyrA*

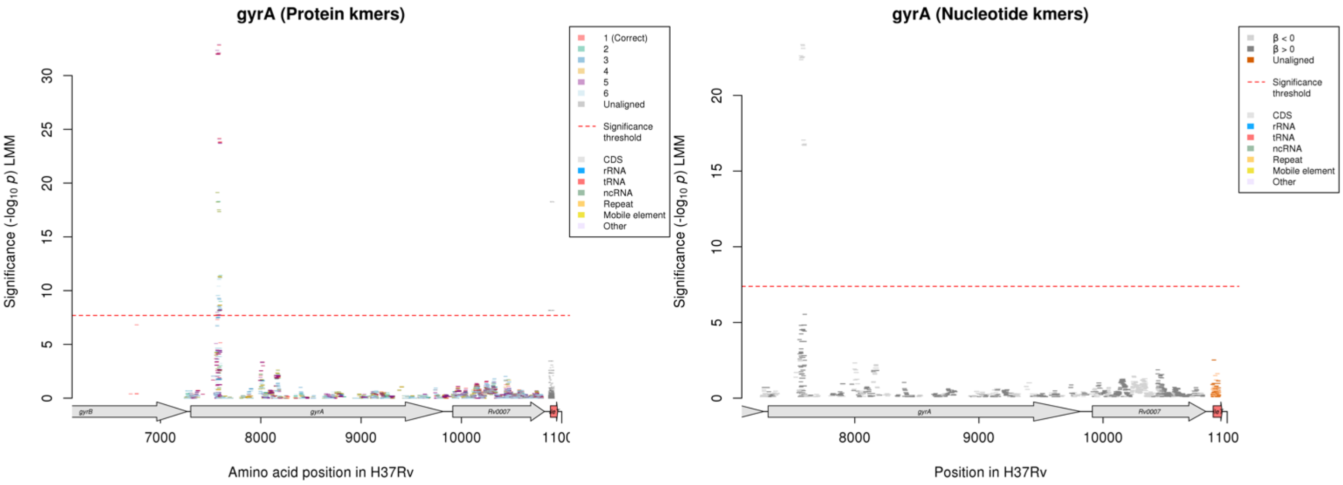

Protein kmers:

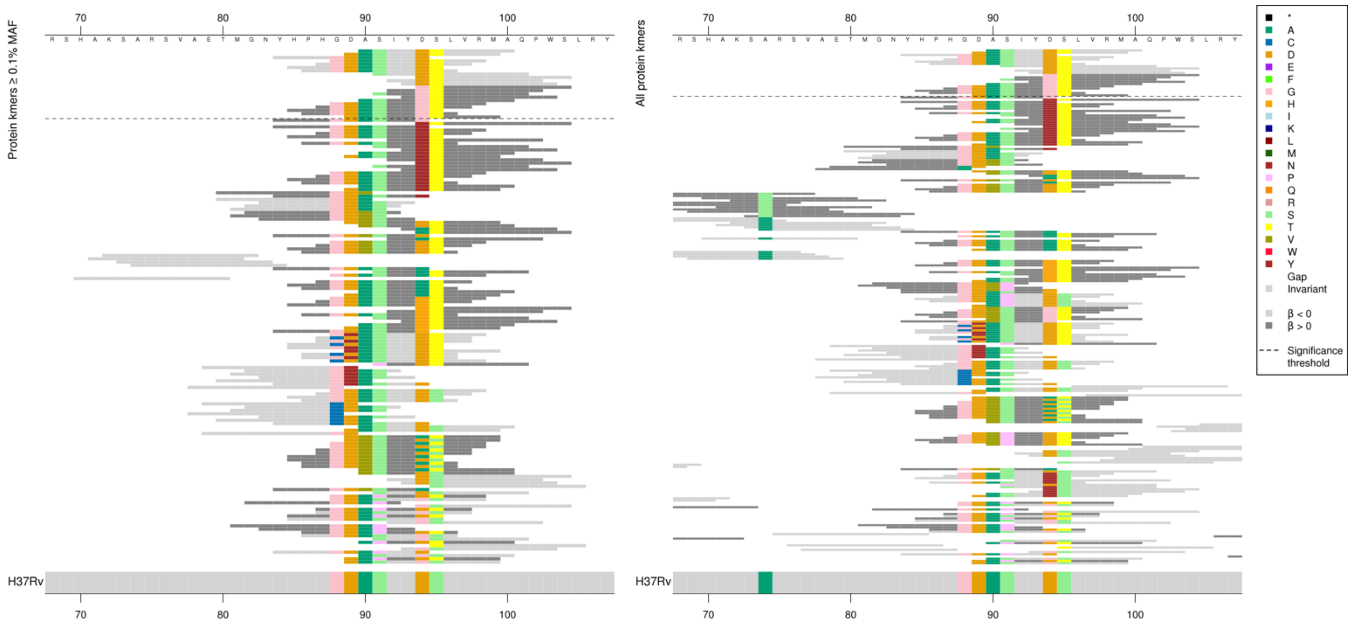

Codon captured by all kmers was 94, and 94/95 for all high MIC-associated kmers, p-values varied based on which other codons were covered, just showing the top for the alleles at 94/95 in the table.

| Kmer | $-\log_{10} p$ | β | MAC | MAF | Ps | Variants |
|---|---|---|---|---|---|---|
| ASIYDTLVRMA | 32.82 | -0.55 | 1813 | 20.72 | 90 | 94D |
| GTLVRMAQPWS | 8.17 | 0.33 | 602 | 6.88 | 94 | 94G, 95T |

### *rpoB*

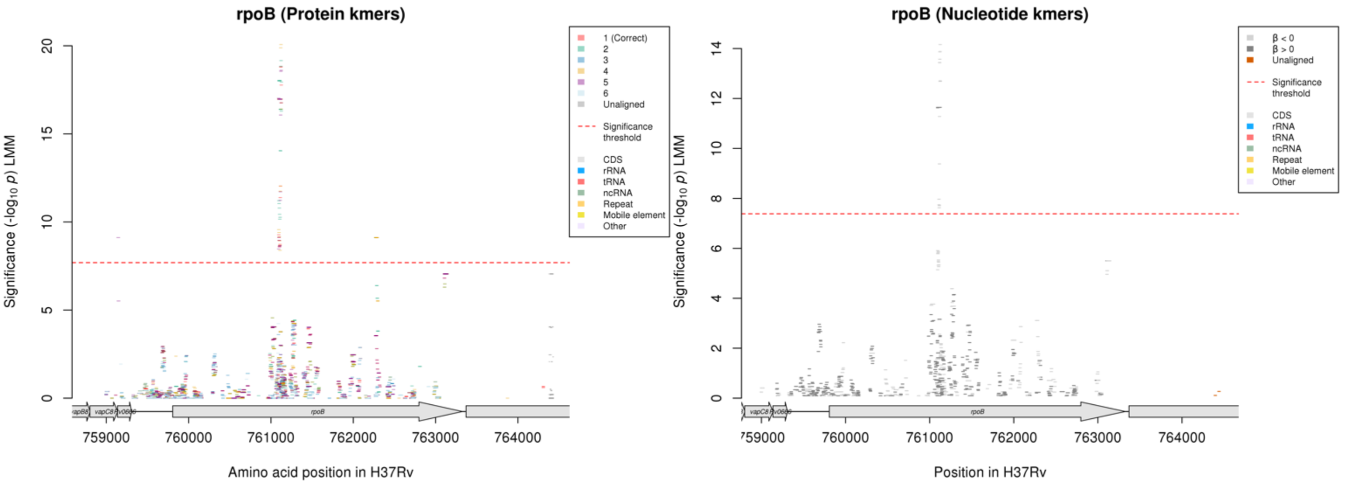

Protein kmers:

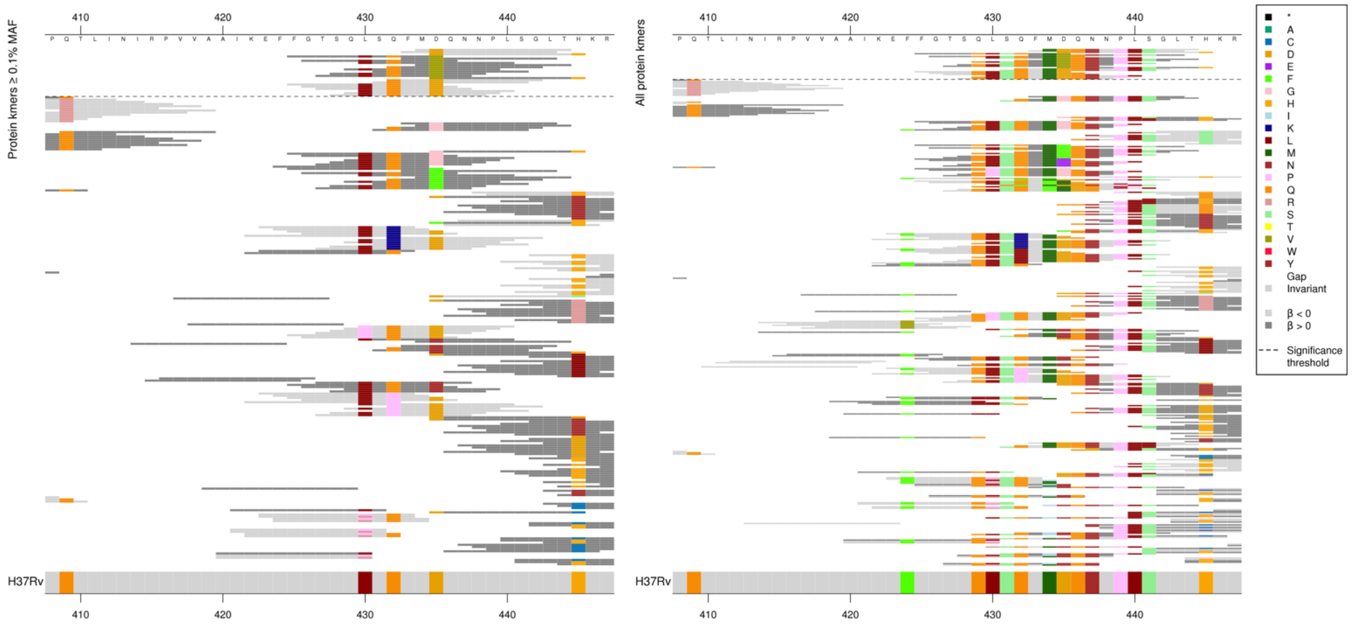

Main substitution being captured was at codon 435. The low MIC-associated kmers capturing these had different *p*-values depending possibly on other substitutions in the region, likewise for the high MIC-associated kmers capturing 435V.

| Kmer | $-\log_{10} p$ | β | MAC | MAF | Ps | Variants |
|---|---|---|---|---|---|---|
| MDQNNPLSGLT | 18.81 | -0.57 | 636 | 7.27 | 434 | 435D |
| FGTSQLSQFMV | 16.97 | 0.70 | 449 | 5.13 | 425 | 435V |

### *ethA*

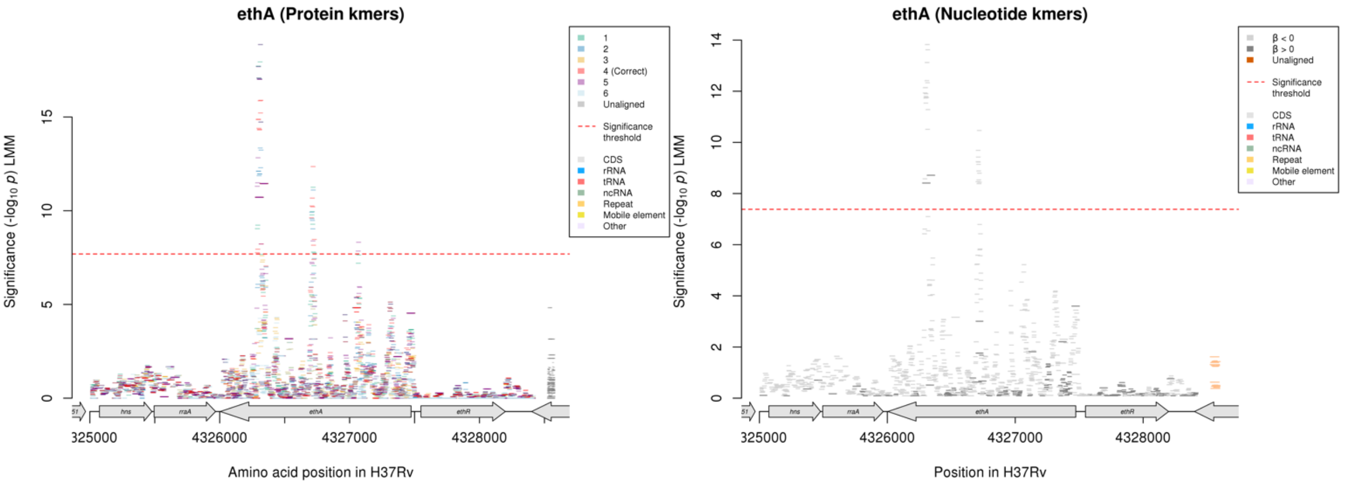

The region in *ethA* that is significant in the protein kmer analysis and not the nucleotide kmer analysis does not have any kmers aligned to the correct reading frame, so it is not analysed here.

Protein kmers:

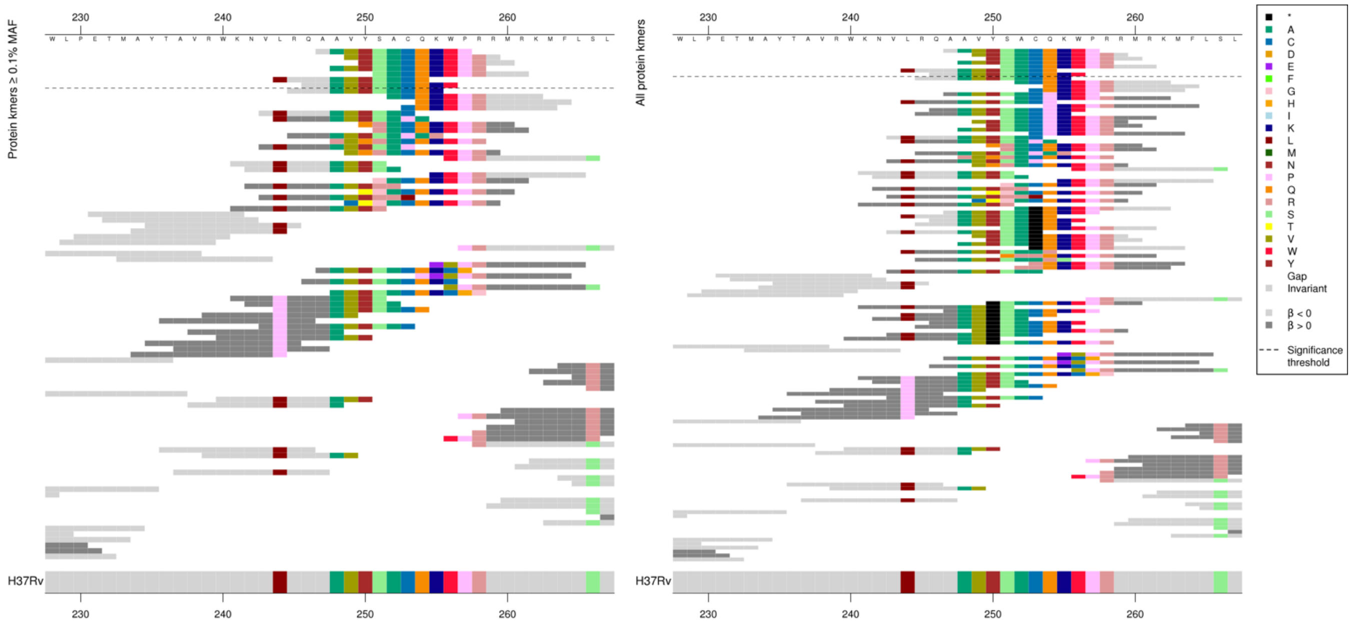

First alignment shows low MIC-associated kmers which were identical to the reference. The high MIC (not significant) kmers were capturing multiple amino acid changes.

Same region for nucleotide kmers:

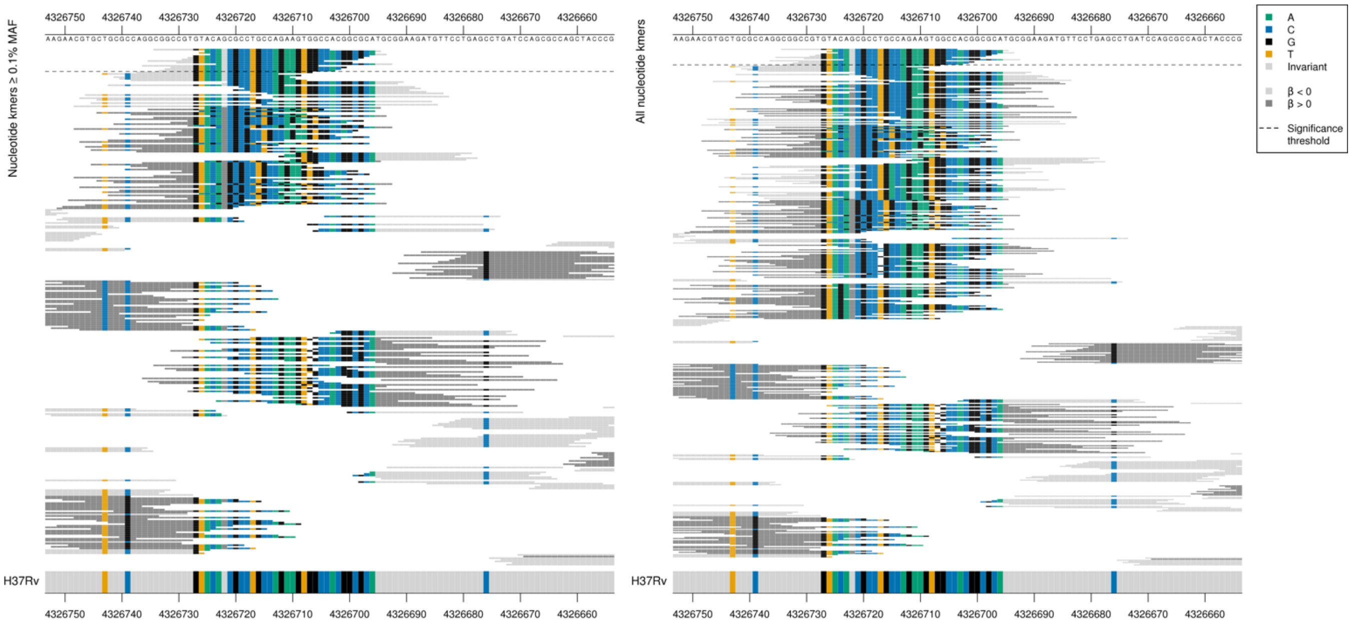

The kmers below the significance threshold were capturing indels.

| Kmer | $-\log_{10} p$ | β | MAC | MAF | Ps | Variants |
|---|---|---|---|---|---|---|
| AAVYSACQKWP | 12.36 | -0.74 | 114 | 1.30 | 247 | WT codons 244-261 |

Protein kmers:

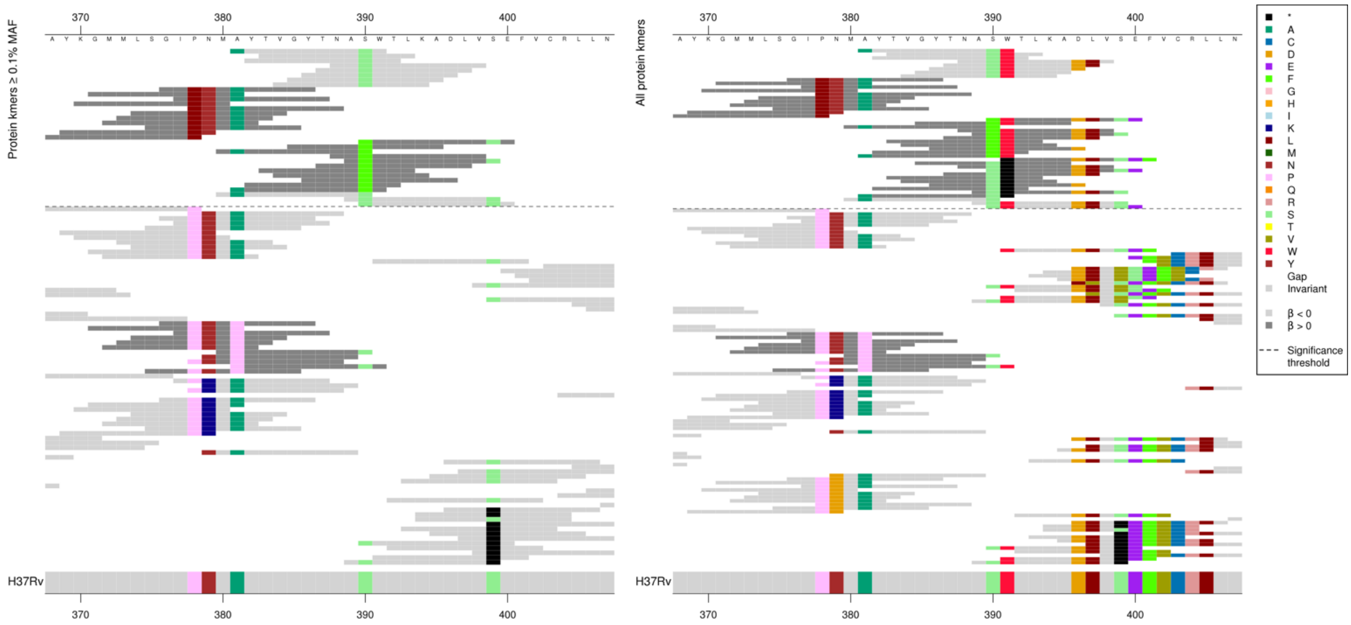

This region of *ethA* was significantly associated for AMI and KAN, but not for ETH.
The *p*-value of the low MIC-associated kmers varied depending on which codons were covered. Reporting the top two covering 381 and 390 below. The high MIC-associated kmers were all only capturing the single substitution as for each they were all the same *p*-value.
Kmers below significance threshold = 381P

Low MAF kmers = 391*.

| Kmer | $-\log_{10} p$ | β | MAC | MAF | Ps | Variants |
|---|---|---|---|---|---|---|
| AYTVGYTNASW | 15.89 | -1.32 | 234 | 2.67 | 381 | 381A, 390S |
| TVGYTNASWTL | 15.87 | -1.21 | 66 | 0.75 | 383 | 390S |
| GILNMAYTVGY | 11.45 | 1.87 | 45 | 0.51 | 376 | 378L |
| FWTLKADLVSE | 10.72 | 2.09 | 18 | 0.21 | 390 | 390F |

#### *fabG1*

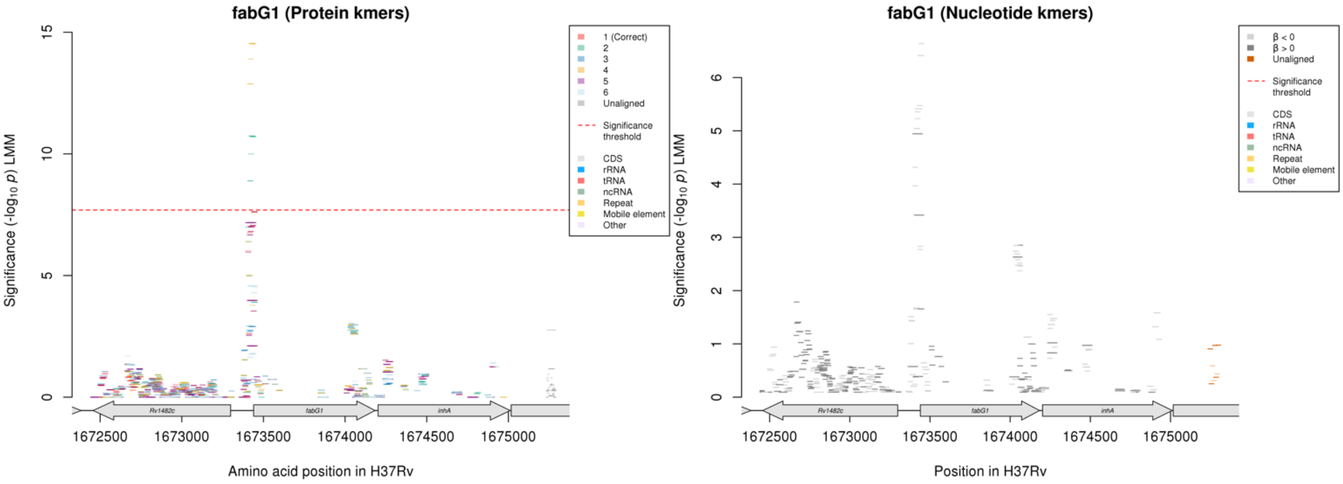

Not significant in the nucleotide kmer analysis, and the protein kmers that aligned to the correct reading frame were just below the significance threshold.

The significant protein kmers in this region were capturing a variant upstream of the start codon. This region was not significant by the nucleotide kmer analysis, but the alignment in the region is as follows:

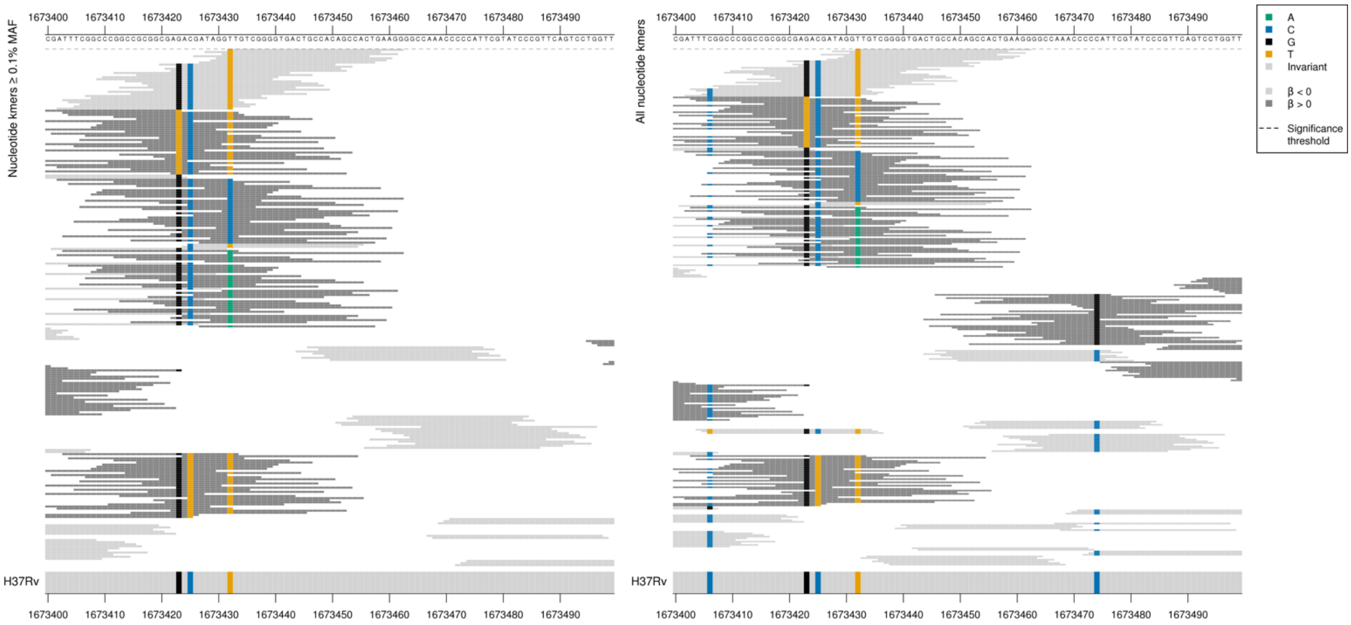

Therefore the protein kmer analysis was capturing the WT sequence where the high MIC (not significant) kmers were either 1673432/-8C or 1673432/-8A.

| Kmer | $-\log_{10} p$ | β | MAC | MAF | Ps | Variants |
|---|---|---|---|---|---|---|
| TTGTCGGGGTGACTGCCACAGCCACTGAAGG | 6.64 | -0.66 | 91 | 1.04 | 1673432 | WT |

## *Rv1830*

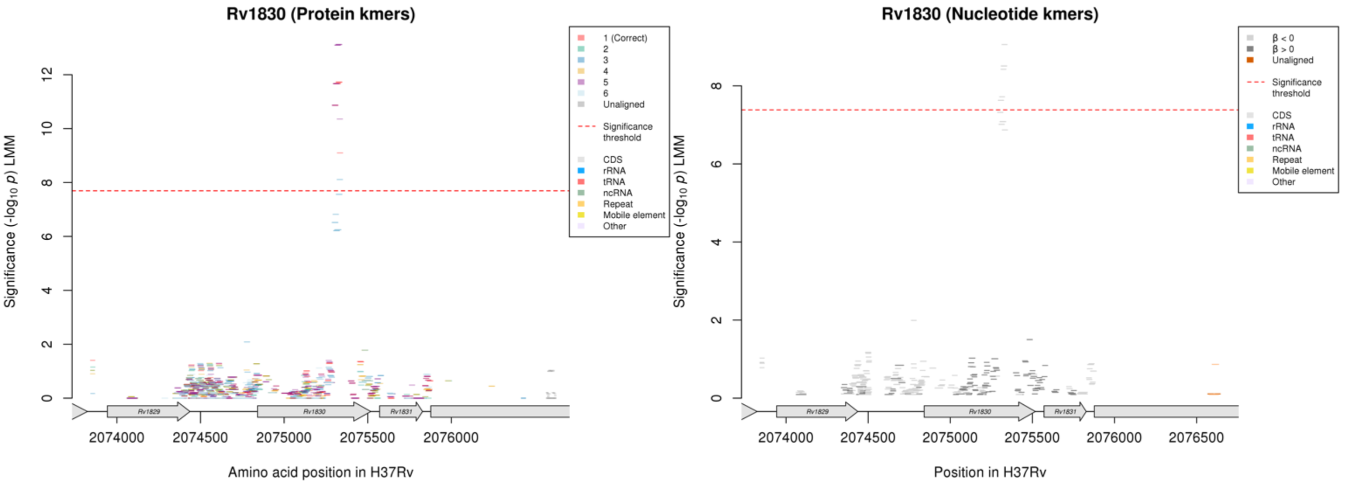

Protein kmers:

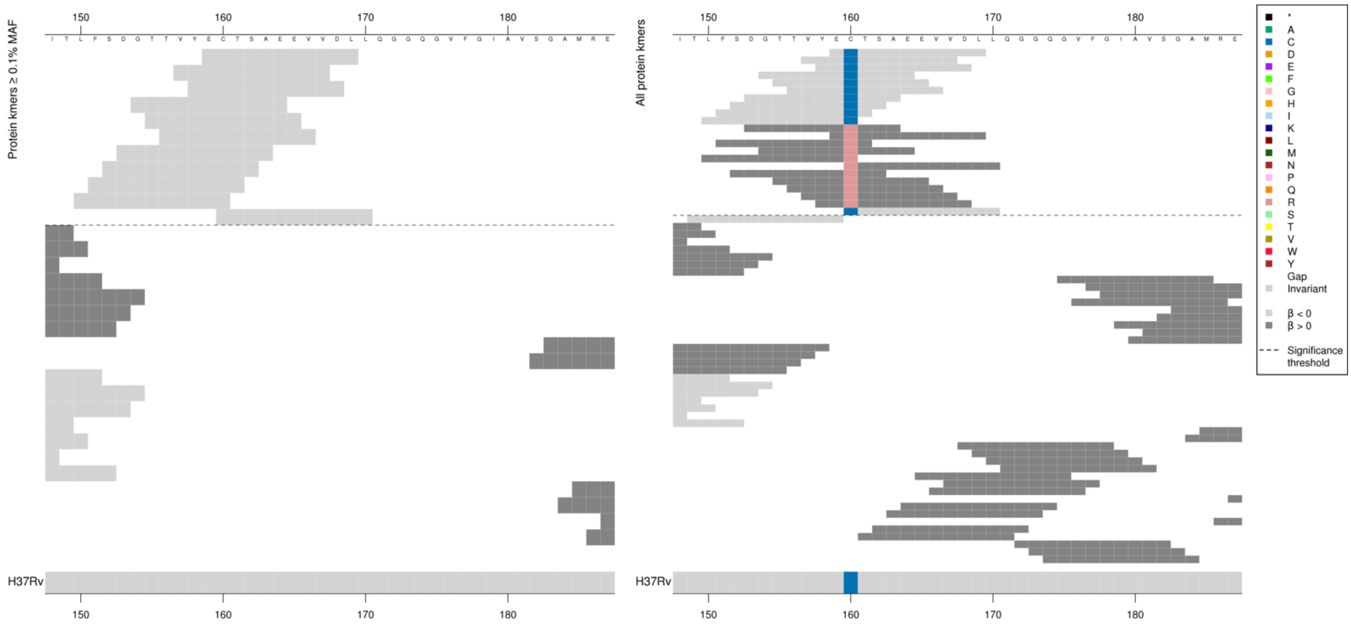

Low MAF kmers = 160R.

| Kmer | $-\log_{10} p$ | β | MAC | MAF | Ps | Variants |
|---|---|---|---|---|---|---|
| ECTSAEEVVDL | 11.72 | -2.31 | 11 | 0.13 | 159 | 160C |

#### *ptbB*

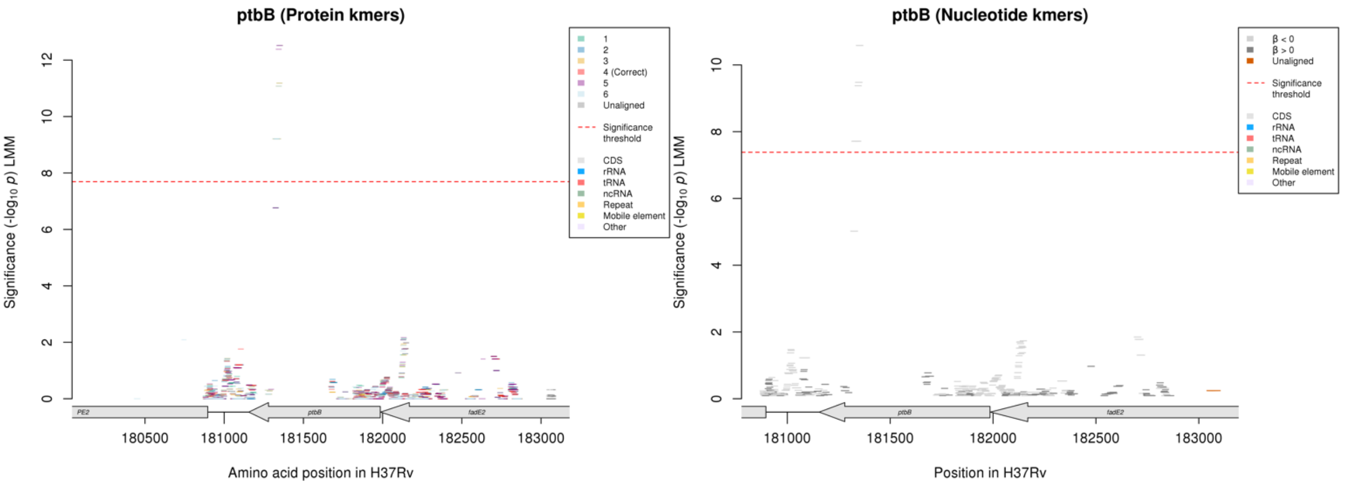

Nucleotide kmers:

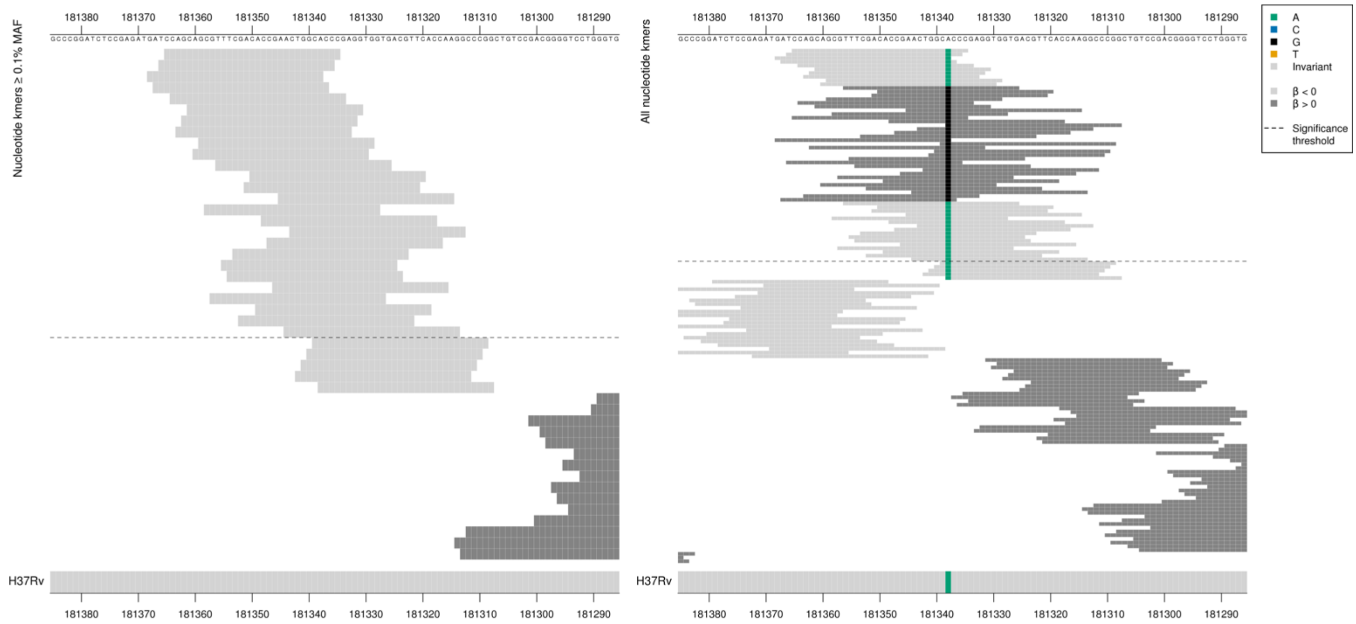

Synonymous mutation at position 181338/648
Low MIC associated kmers = GCA = Ala / A MAC between 9-11
High MIC associated kmers = GCG = Ala / A

Low MAF kmers = 181338/648G.

| Kmer | -log$_{10}$ *p* | β | MAC | MAF | Ps | Variants |
|---|---|---|---|---|---|---|
| CCAGCAGCGTTTCGACACCGAACTGGCACCC | 10.58 | -2.35 | 9 | 0.10 | 181365 | 181338/648A |

#### *PPE42*

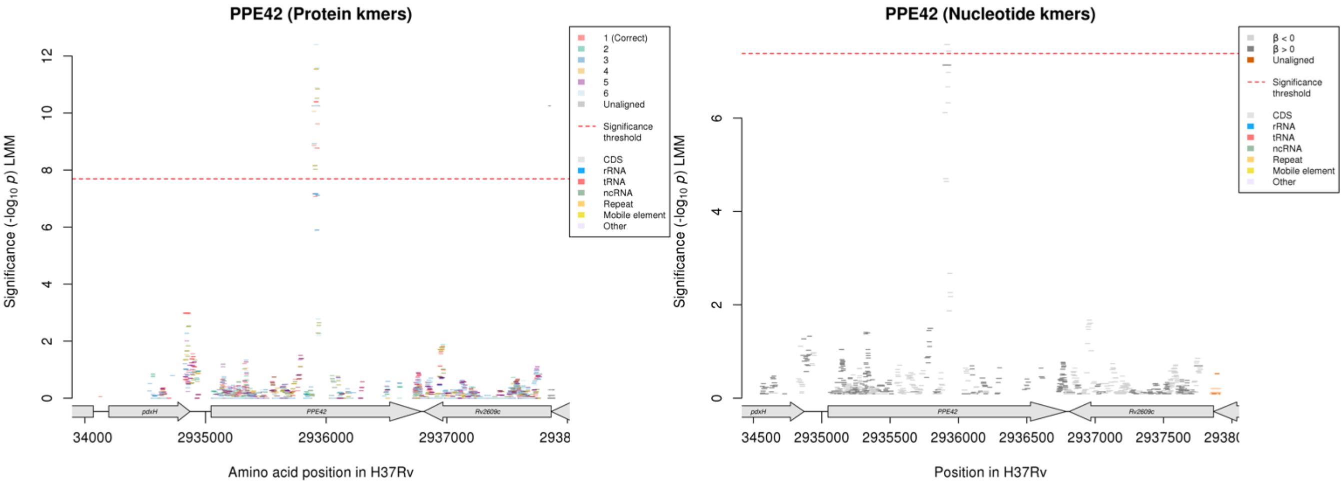

Protein kmers:

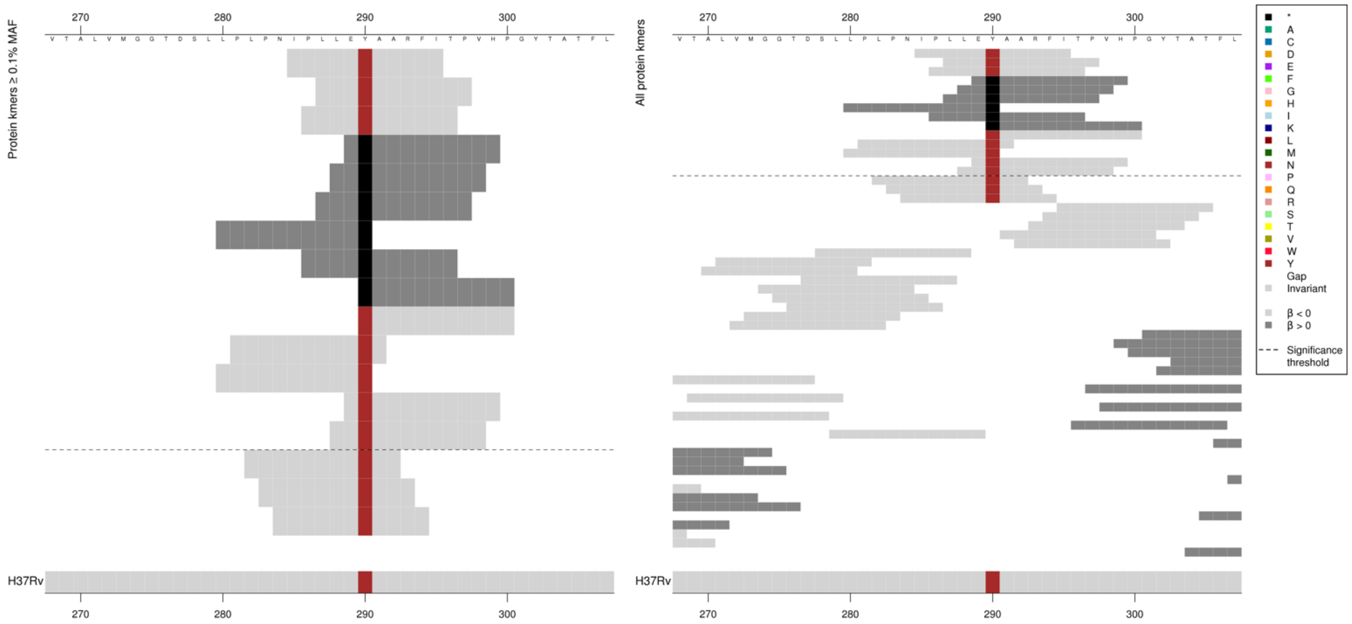

High MIC-associated kmers were all the same *p*-value. Low MIC-associated kmers were different *p*-values.

| Kmer | $-\log_{10} p$ | β | MAC | MAF | Ps | Variants |
|---|---|---|---|---|---|---|
| IPLLEYAARFI | 10.39 | -1.93 | 45 | 0.51 | 285 | 290Y |
| E*AARFITPVH | 10.25 | 2.33 | 40 | 0.46 | 289 | 290* |

#### *echA8*

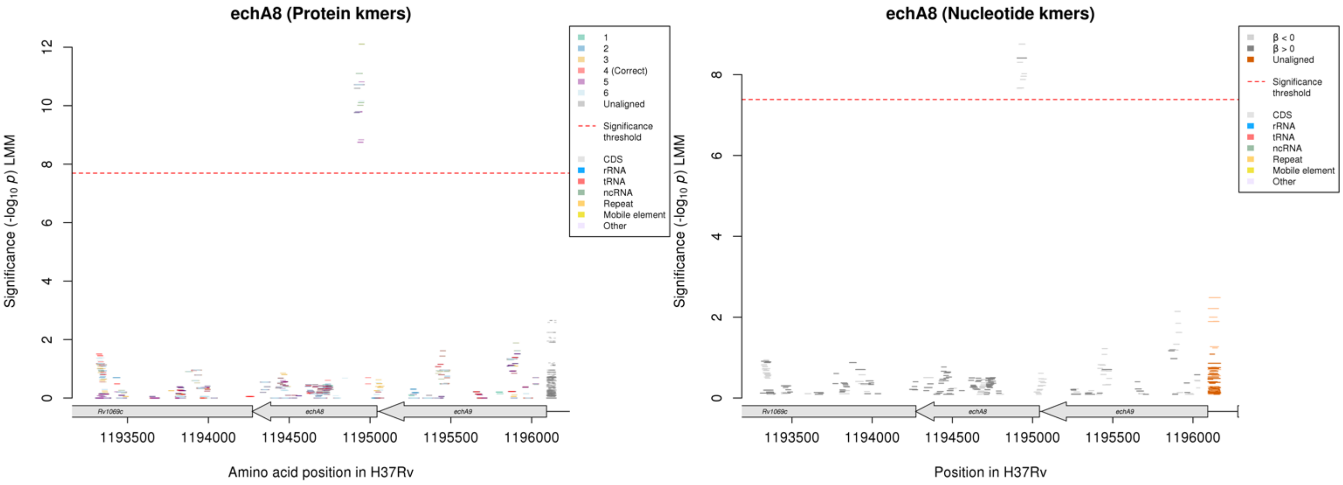

Nucleotide kmers:

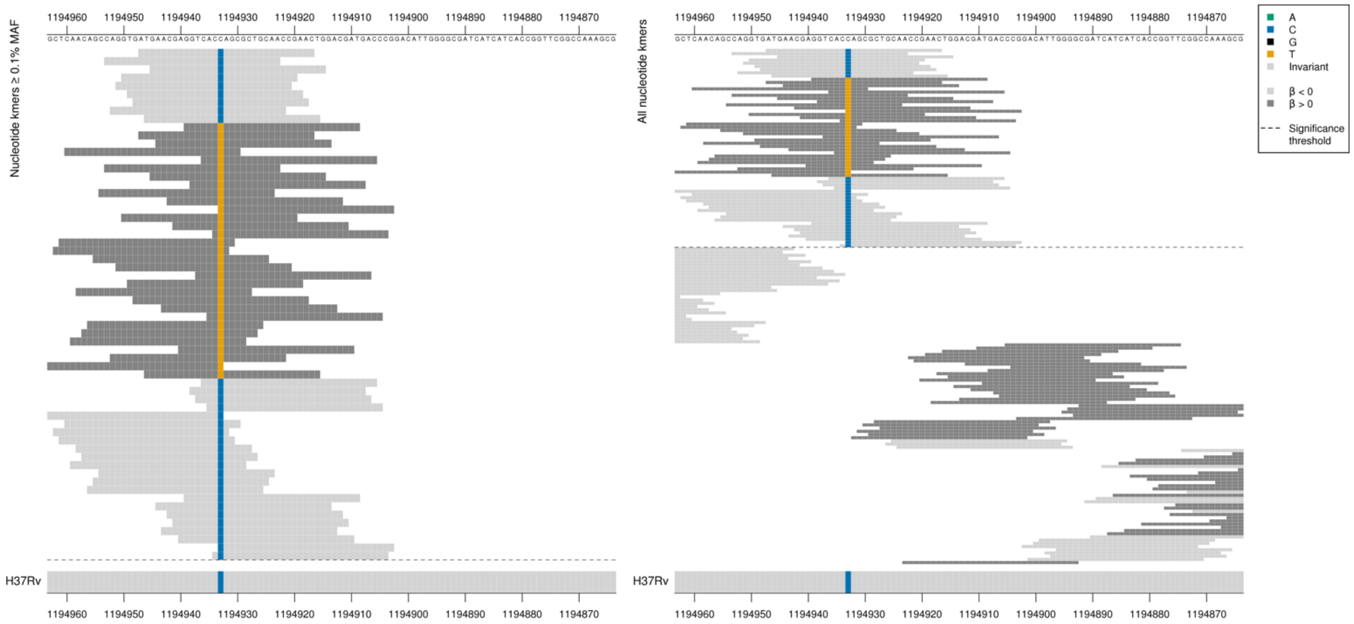

Low MIC-associated = ACC = Thr / T
High MIC-associated = ACT = Thr / T
Synonymous mutation at position 1194933/111

| Kmer | $-\log_{10} p$ | β | MAC | MAF | Ps | Variants |
|---|---|---|---|---|---|---|
| ATGAACGAGGTCACCAGCGCTGCAACCGAAC | 8.75 | -1.72 | 21 | 0.24 | 1194947 | 1194933/111C |
| GGTCACTAGCGCTGCAACCGAACTGGACGAT | 8.41 | 1.87 | 18 | 0.21 | 1194939 | 1194933/111T |

#### *lprF:Rv1371*$^{R}$

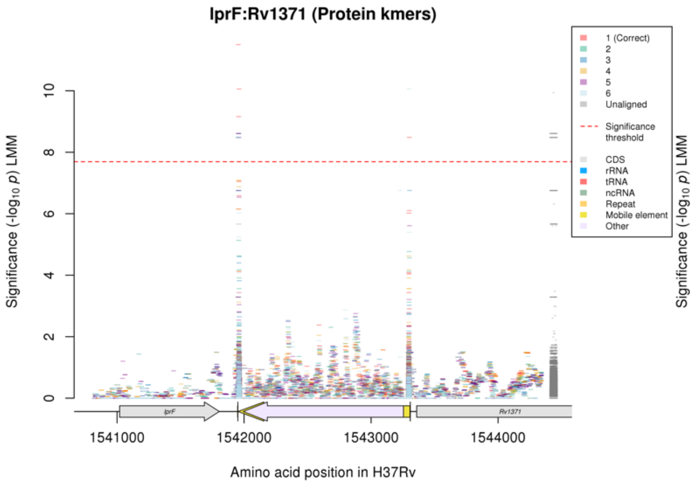

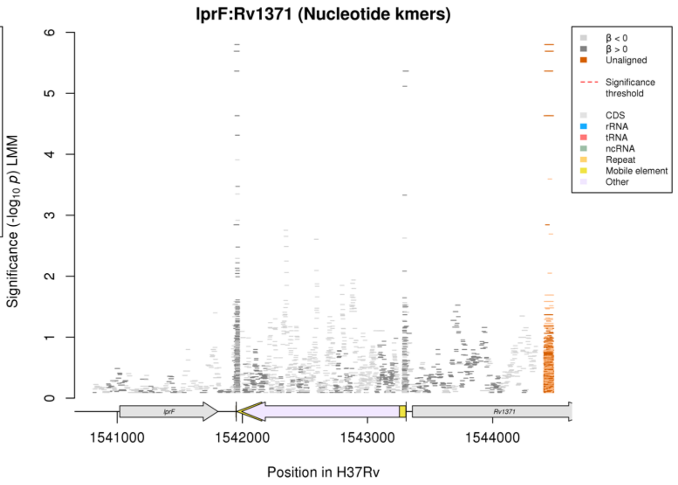

Repeat regions at the end of a mobile element

Not significant in the nucleotide kmer analysis, but showing the alignment of the most significant region (in both the protein and nucleotide kmer analyses) below:

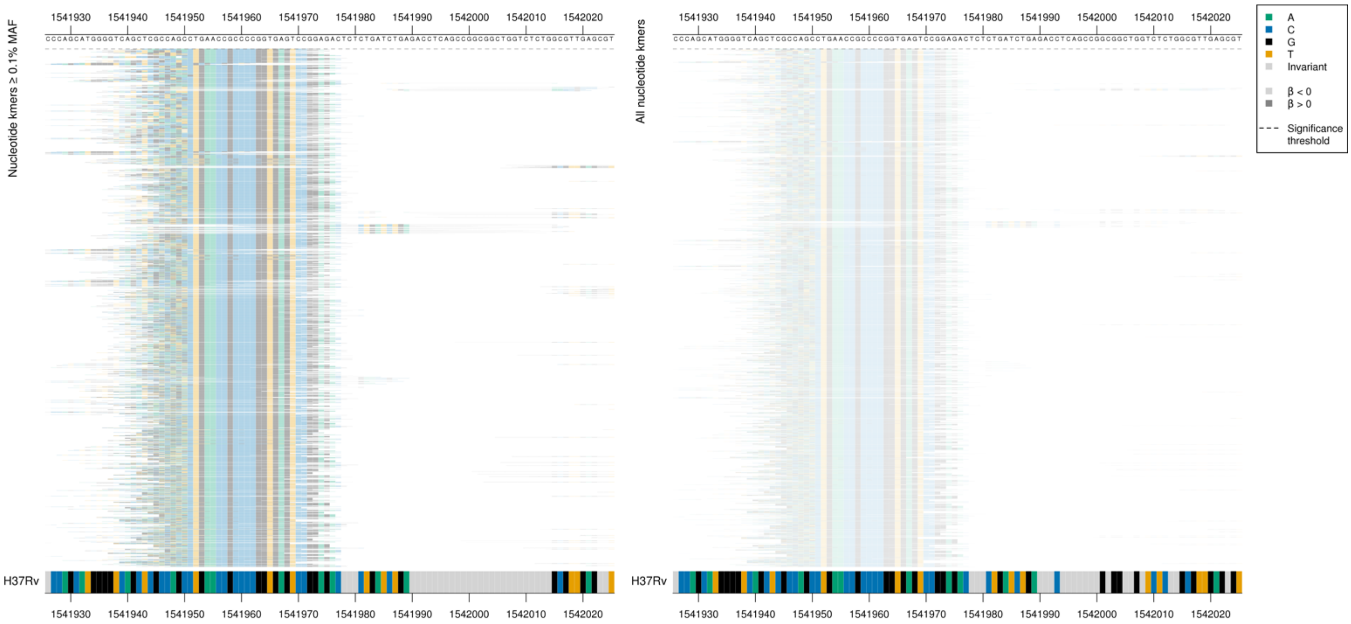

Most significant protein kmer in the region (LARSTEPPR*V): -log10p = 11.50, beta = 2.07, MAC = 41, MAF = 0.47, codon ps = 45.

| Kmer | $-\log_{10} p$ | β | MAC | MAF | Ps | Variants |
|---|---|---|---|---|---|---|
| CAACCCAACTGAACCGCCCCGGTGAGTCCGG | 5.80 | 1.83 | 41 | 0.47 | 1541943 | Repeat region |

#### *Rv2348c:plcC*

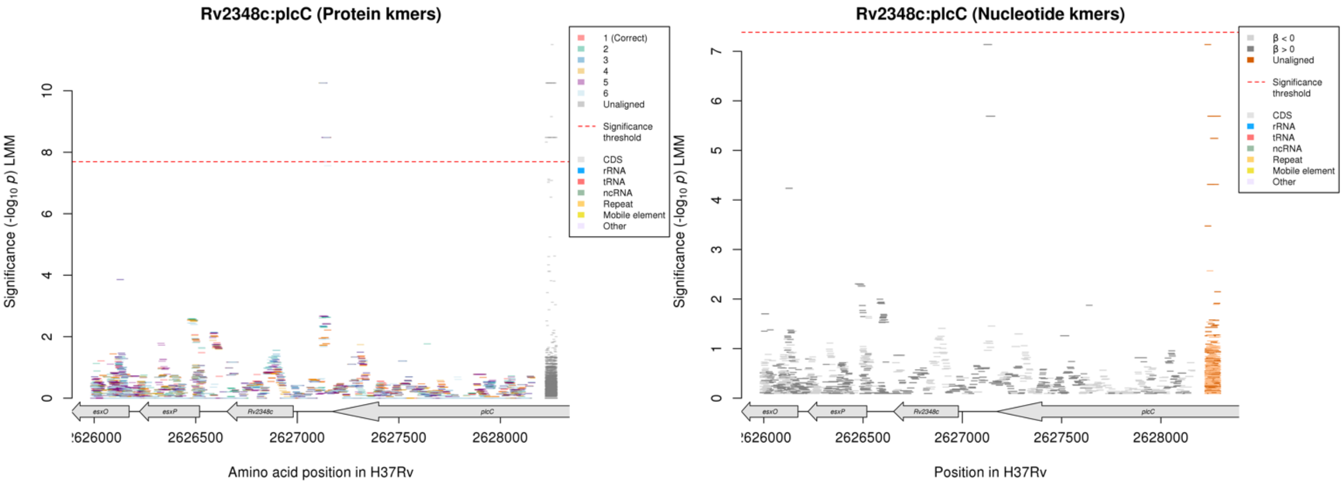

Not significant in the nucleotide kmer analysis.

Nucleotide kmers:

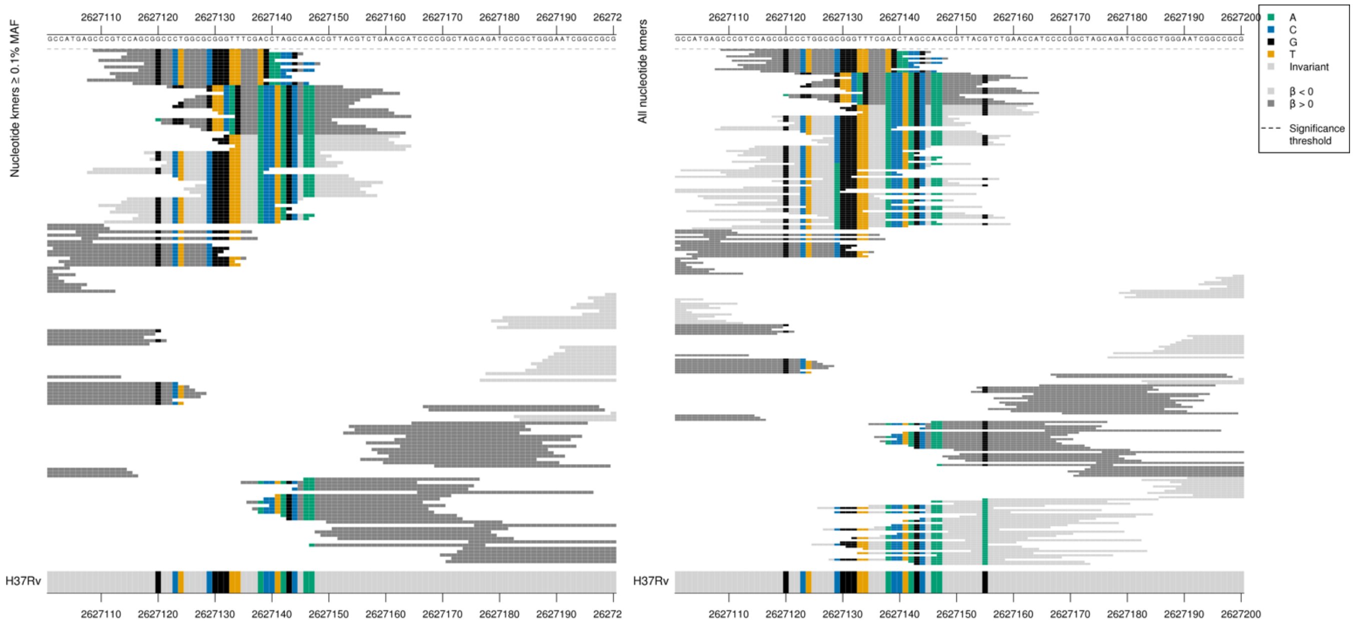

There were two groups of high MIC-associated kmers captured by the protein kmer analysis. For each group, the nucleotide kmers matched the reference for one half of the kmer, then diverge from the reference, possibly a deletion or insertion.

| Kmer | $-\log_{10} p$ | β | MAC | MAF | Ps | Variants |
|---|---|---|---|---|---|---|
| CCCGTCCAGCGGCCCTGGCGCGGGTTTCGTG | 7.14 | 2.03 | 40 | 0.46 | 2627209 | Multiple differences to reference |

#### *narU*

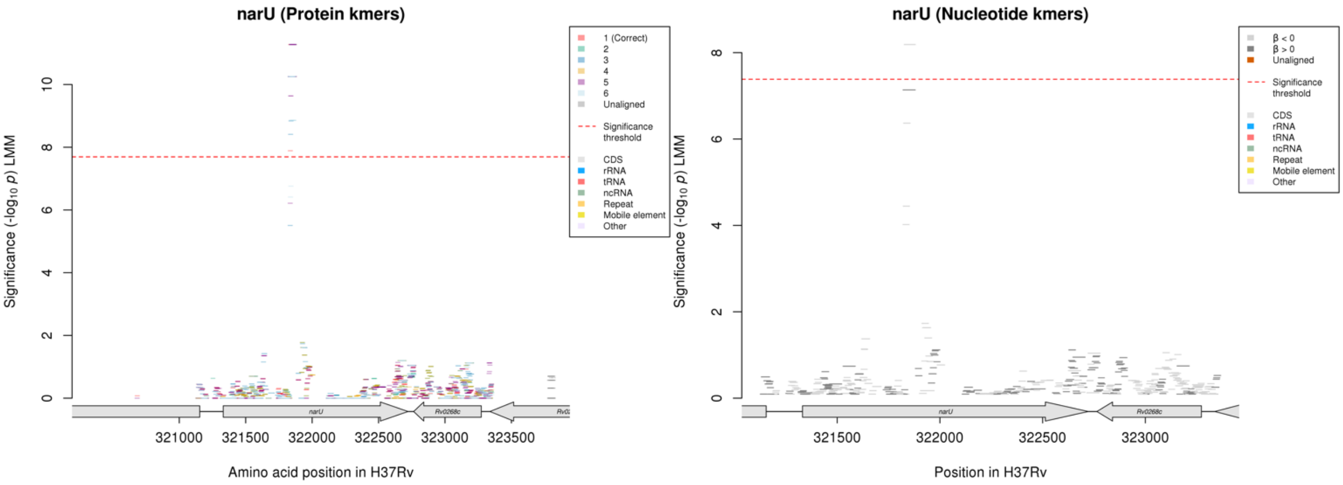

#### Protein kmers:

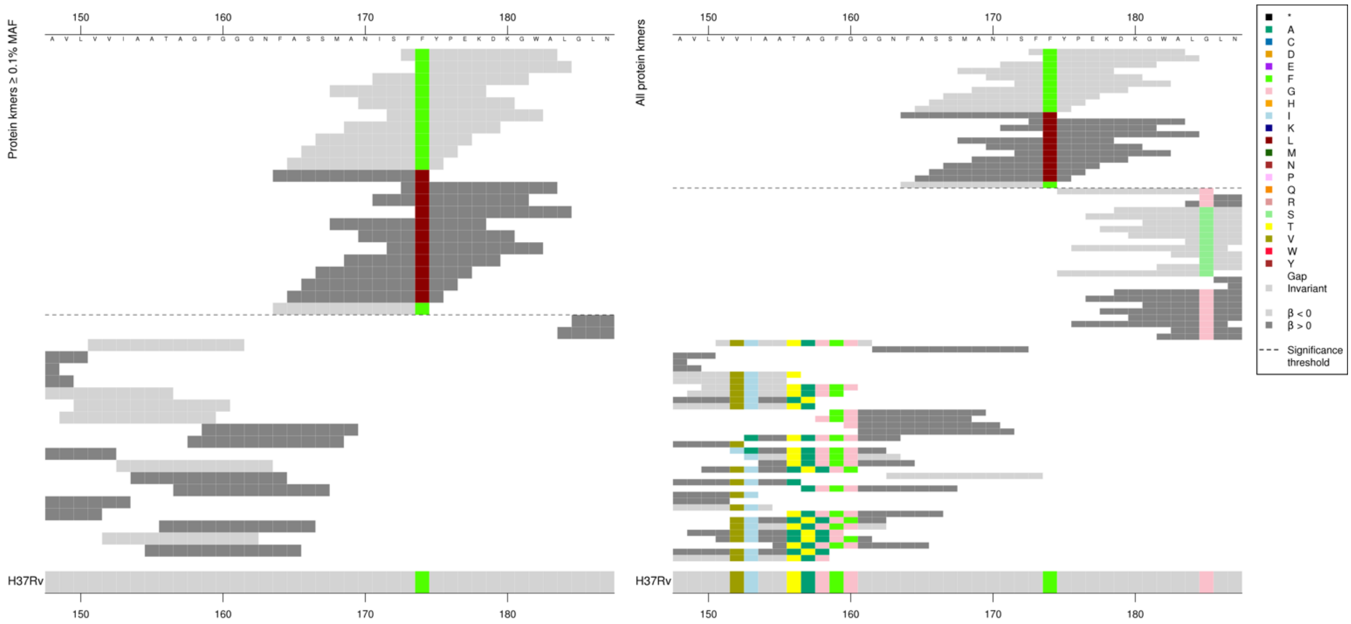

| Kmer | -log₁₀ *p* | β | MAC | MAF | Ps | Variants |
|---|---|---|---|---|---|---|
| FFYPEKDKGWA | 11.27 | -2.09 | 44 | 0.50 | 173 | 174F |
| FASSMANISFL | 10.25 | 2.33 | 40 | 0.46 | 164 | 174L |

#### *pgi*

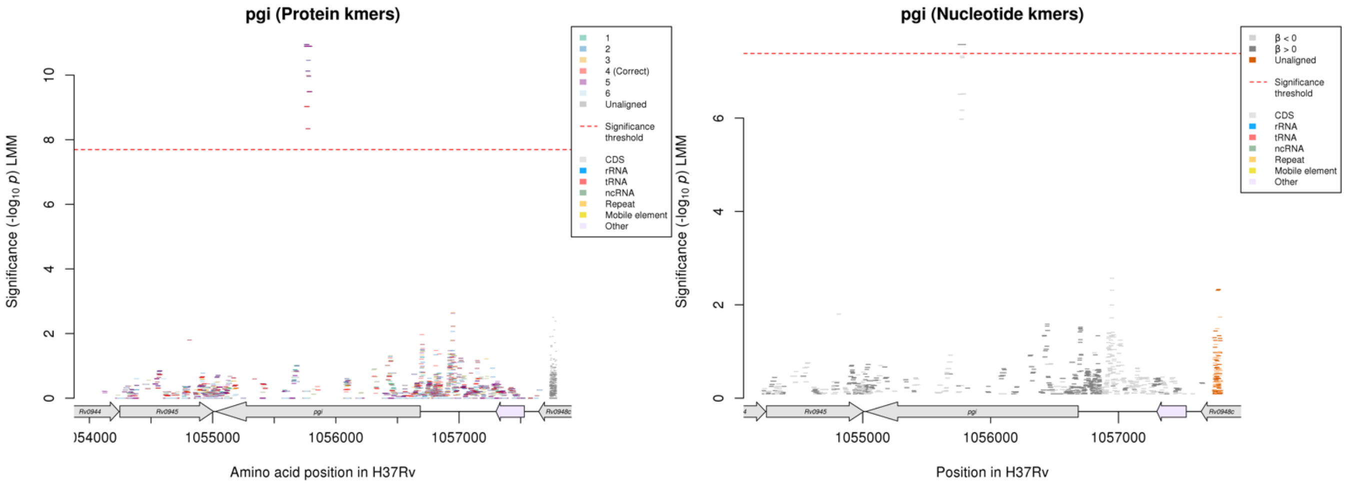

Protein kmers:

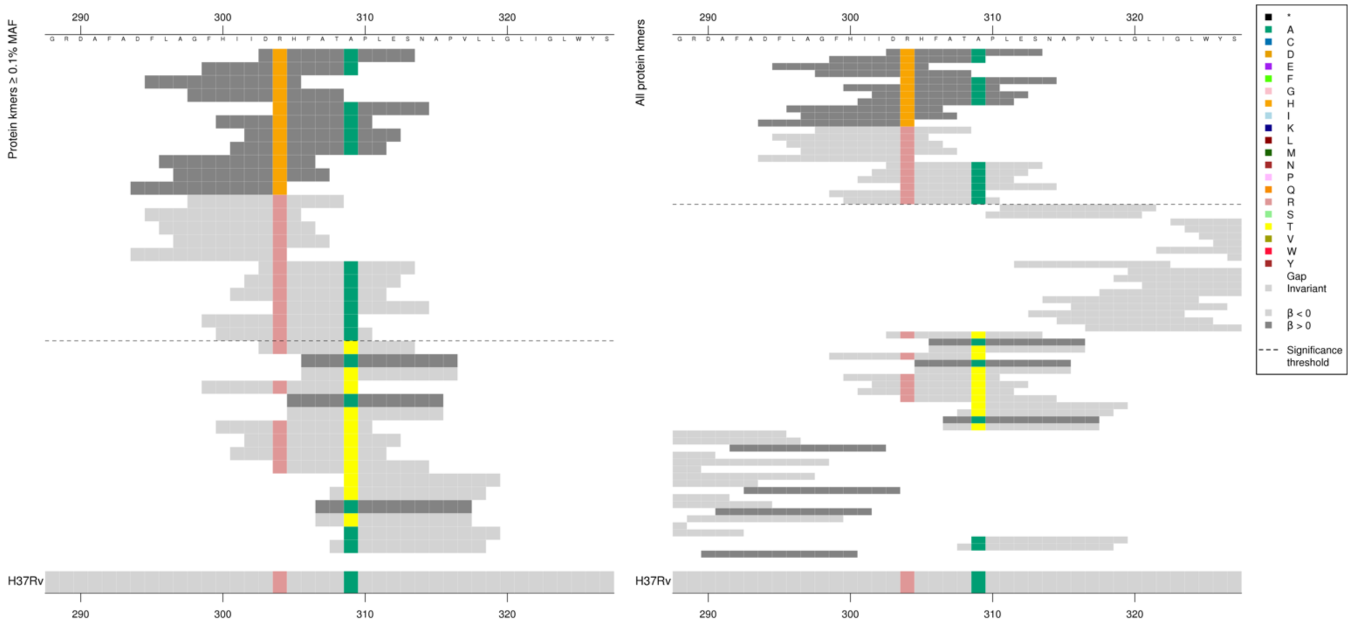

| Kmer | $-\log_{10} p$ | β | MAC | MAF | Ps | Variants |
|---|---|---|---|---|---|---|
| DHHFATAPLES | 10.89 | 1.35 | 128 | 1.46 | 303 | 304H |
| GFHIIDRHFAT | 9.98 | -1.26 | 130 | 1.49 | 298 | 304R |
| DRHFATAPLES | 9.03 | -1.14 | 143 | 1.63 | 303 | 304R, 309A |

#### *mmaA4*

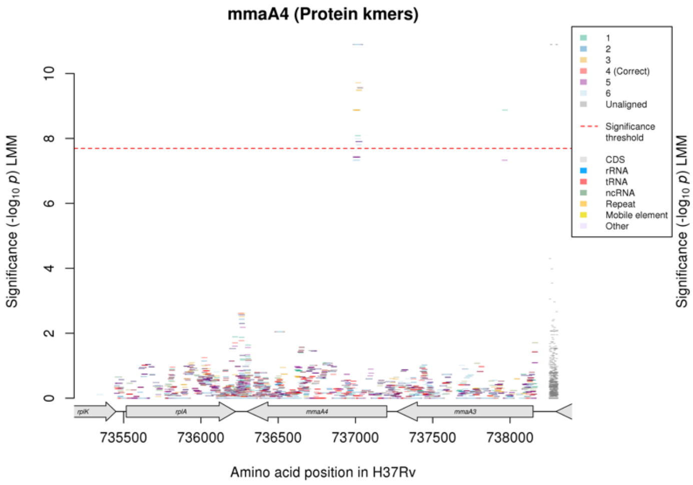

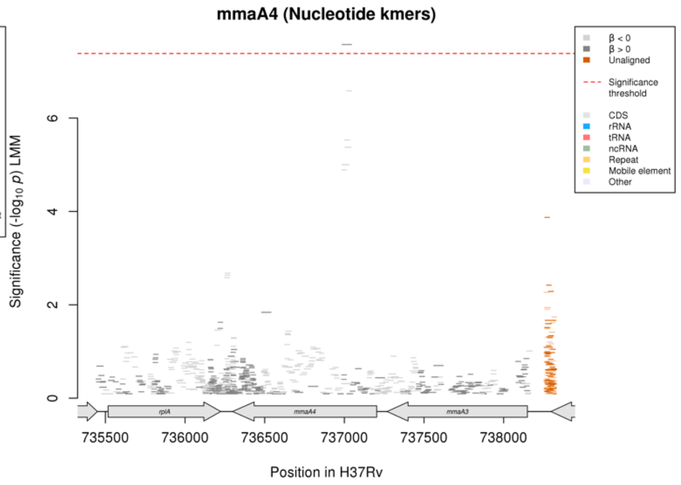

No significant kmers aligned to the correct reading frame for the protein kmer analysis.

Nucleotide kmers:

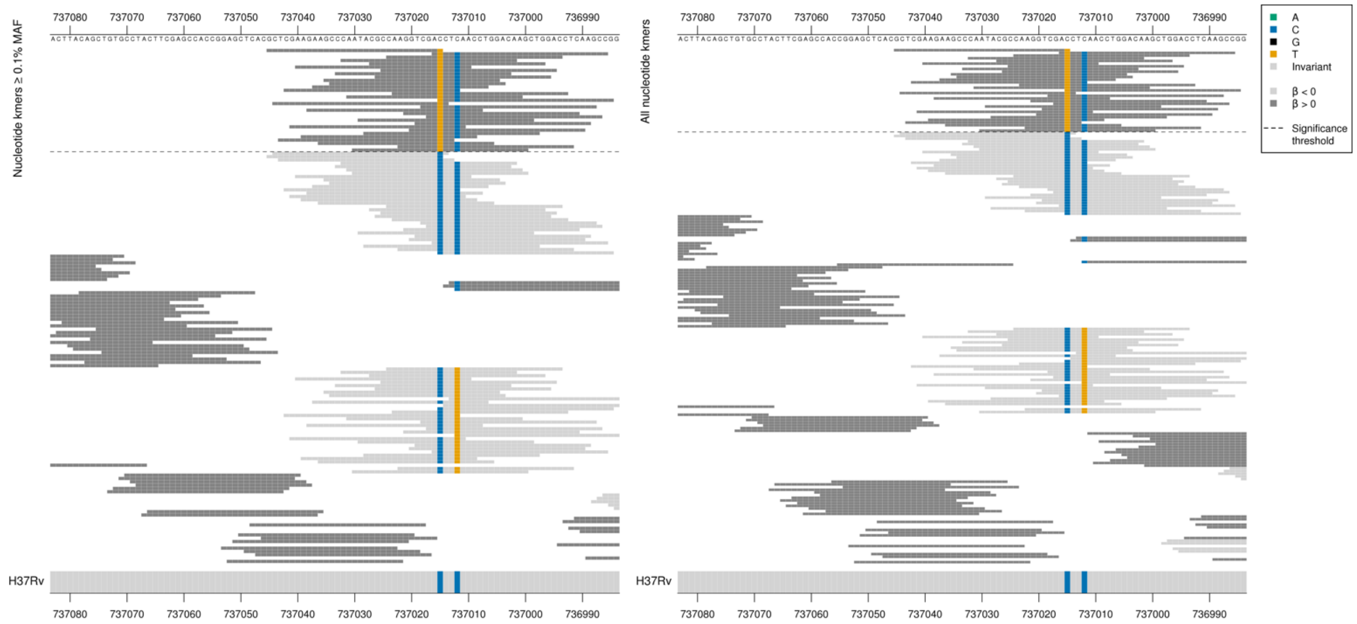

mmaA4 begins on the reverse strand at 737203
High MIC-associated kmeres = GAT = Asp
Kmers below significance threshold = GAC = Asp

| Kmer | -log$_{10}$ *p* | β | MAC | MAF | Ps | Variants |
|---|---|---|---|---|---|---|
| GCTCGAAGAAGCCCAATACGCCAAGGTCGAT | 7.57 | 1.15 | 128 | 1.46 | 737045 | 737015/189T |

#### *pncA*

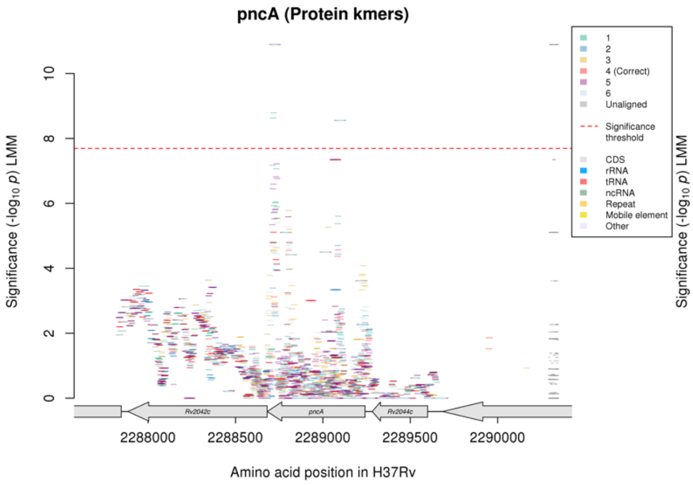

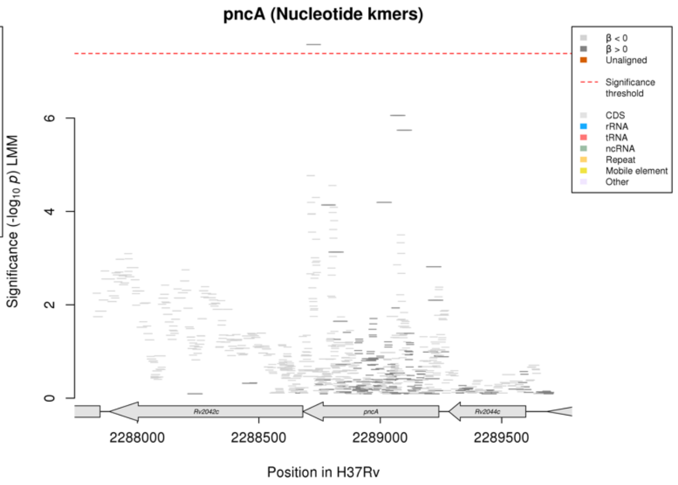

#### Protein kmers:

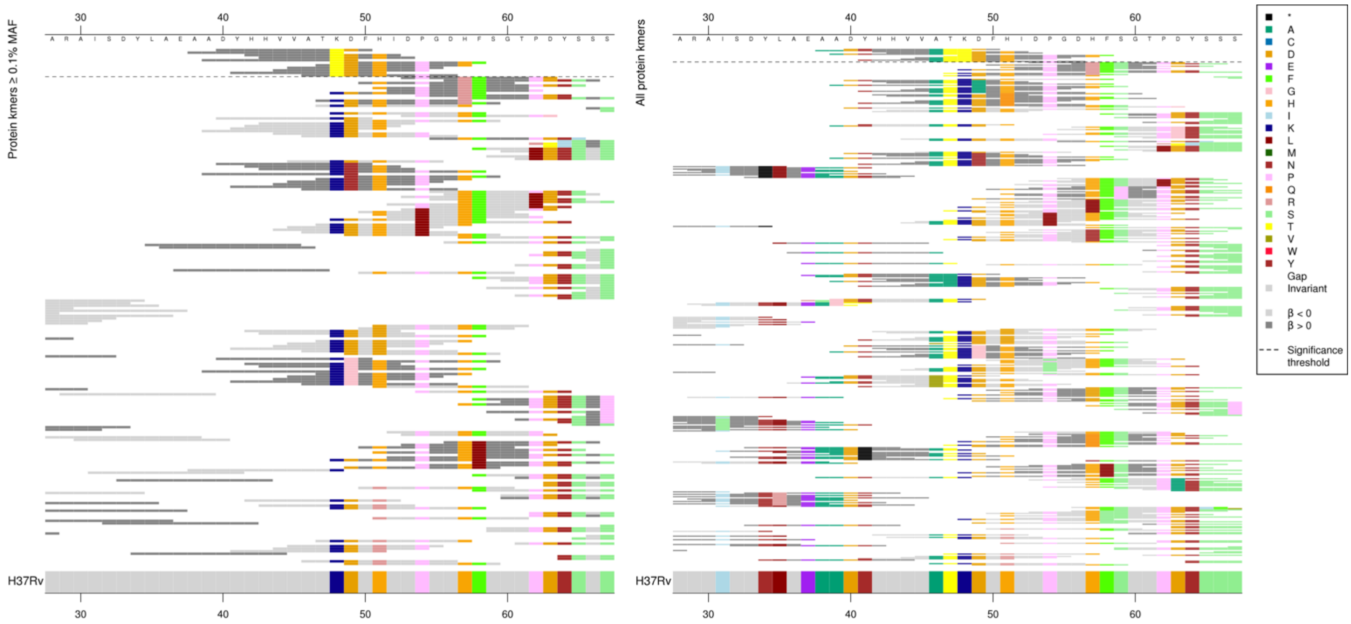

| Kmer | $-\log_{10} p$ | β | MAC | MAF | Ps | Variants |
|---|---|---|---|---|---|---|
| DYHHVVATTDF | 8.55 | 1.96 | 42 | 0.48 | 40 | 48T |

#### Protein kmers:

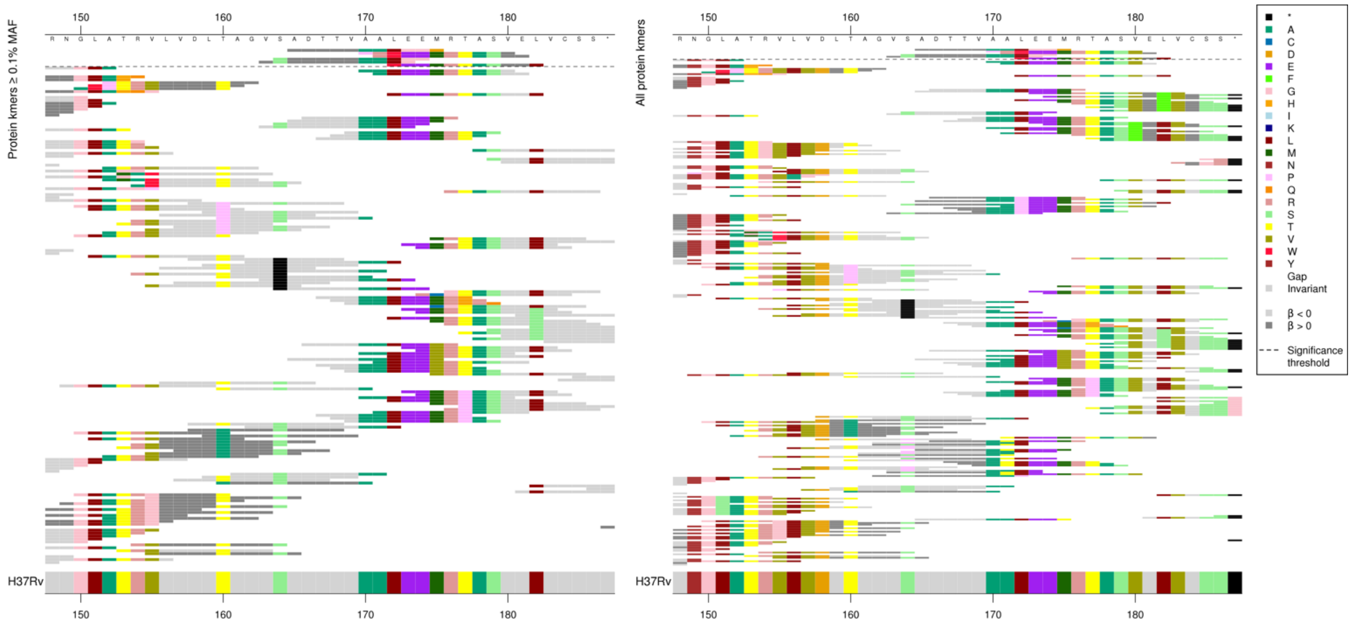

Same region for nucleotide kmers:

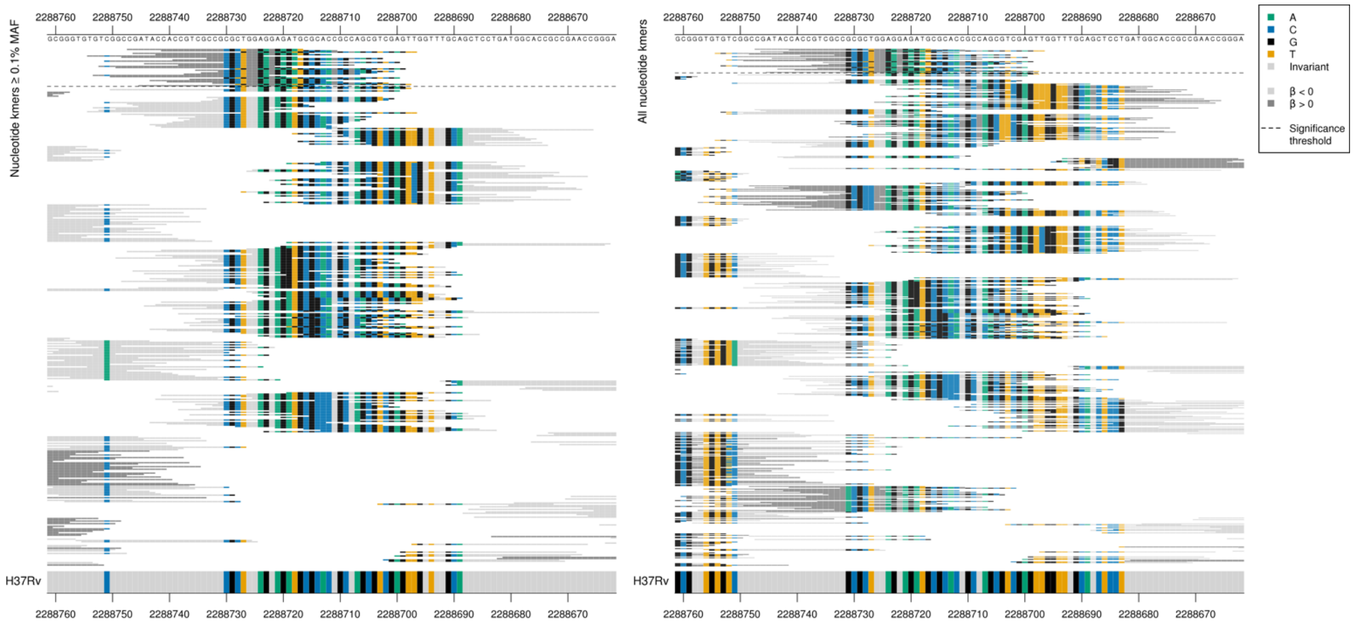

| Kmer | -log₁₀ *p* | β | MAC | MAF | Ps | Variants |
|---|---|---|---|---|---|---|
| CGATACCACCGTCGCCGCGCTGGGAGGAGAT | 7.57 | 1.15 | 128 | 1.46 | 2288747 | G insertion between 2288727-2288724/515-518 |

### *viuB*

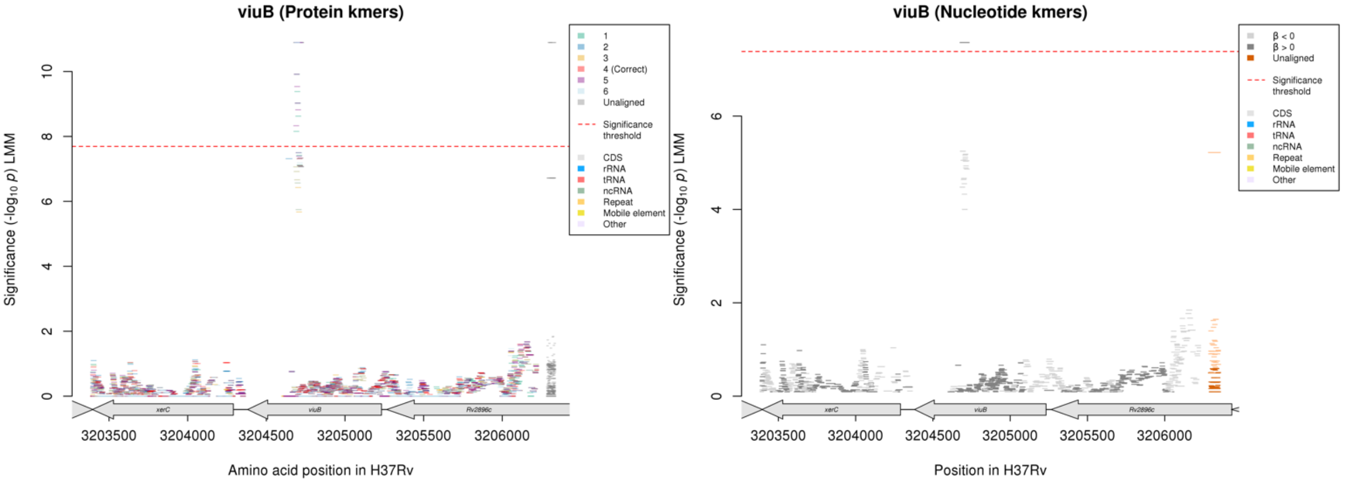

No significant kmers aligned to the correct reading frame in the protein kmer analysis.

Nucleotide kmers:

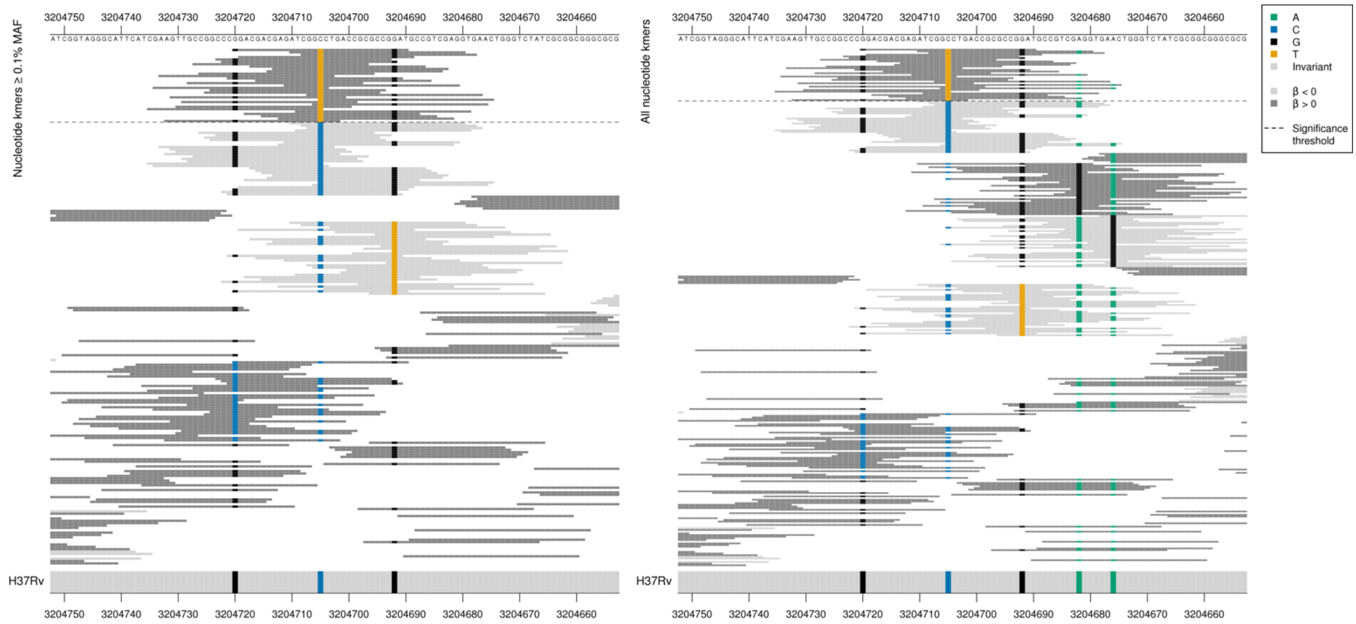

High MIC-associated kmers = GGC = G/Gly

Kmers below significance threshold = GGT = G/Gly

| Kmer | $-\log_{10} p$ | β | MAC | MAF | Ps | Variants |
|---|---|---|---|---|---|---|
| GGACGACGAGATCGGTCTGACCGCGCCGGAT | 7.57 | 1.15 | 128 | 1.46 | 3204720 | 3204705/528T |

#### *lprC*

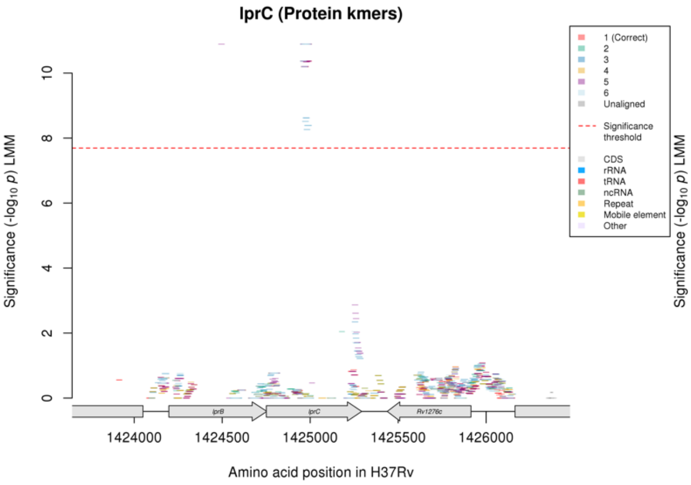

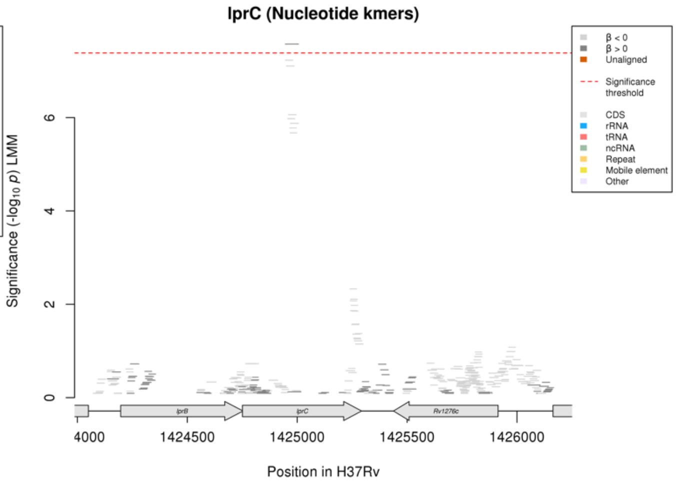

Protein kmers:

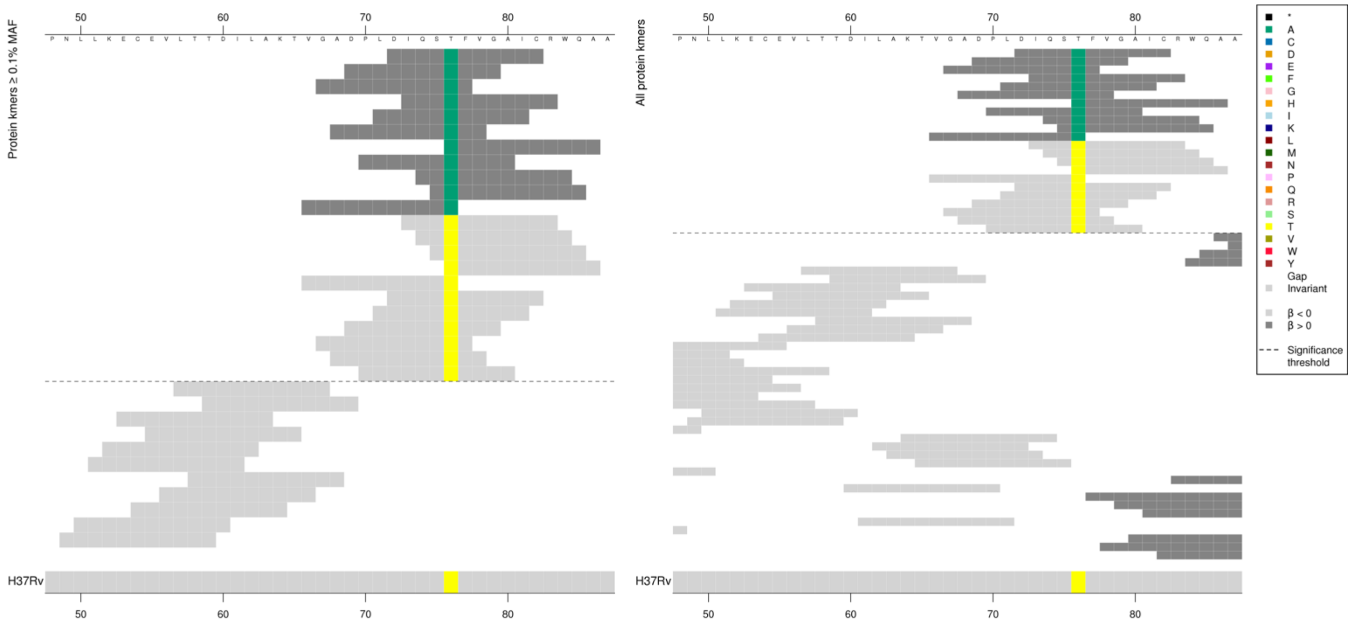

| Kmer | -log₁₀ $p$ | β | MAC | MAF | Ps | Variants |
|---|---|---|---|---|---|---|
| DIQSAFVGAIC | 10.89 | 1.35 | 128 | 1.46 | 72 | 76A |
| IQSTFVGAICR | 10.37 | -1.28 | 133 | 1.52 | 73 | 76T |

#### *murA*

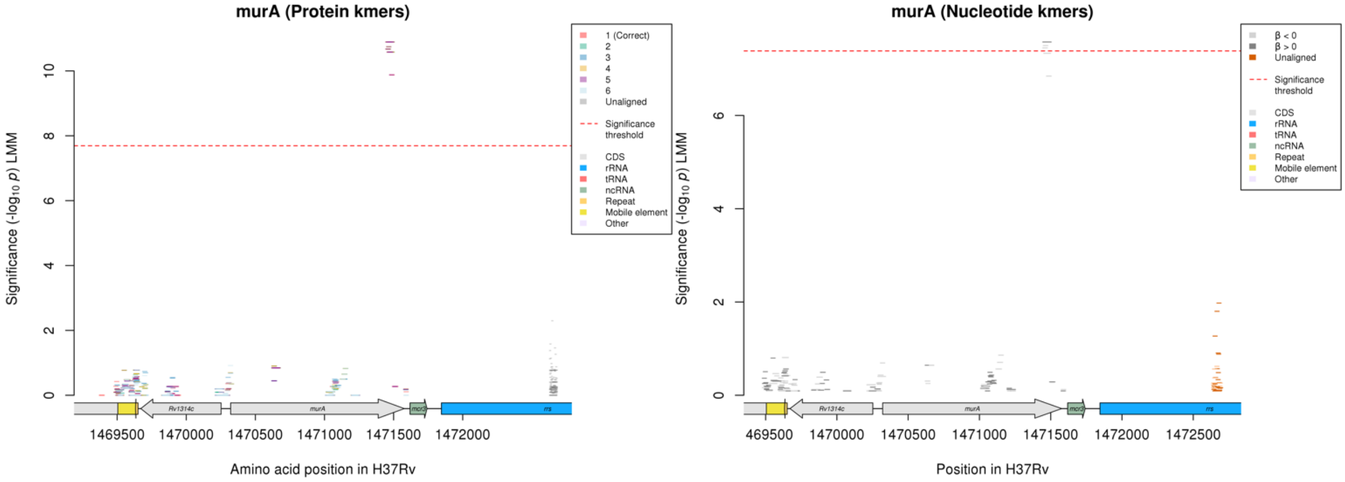

Protein kmers:

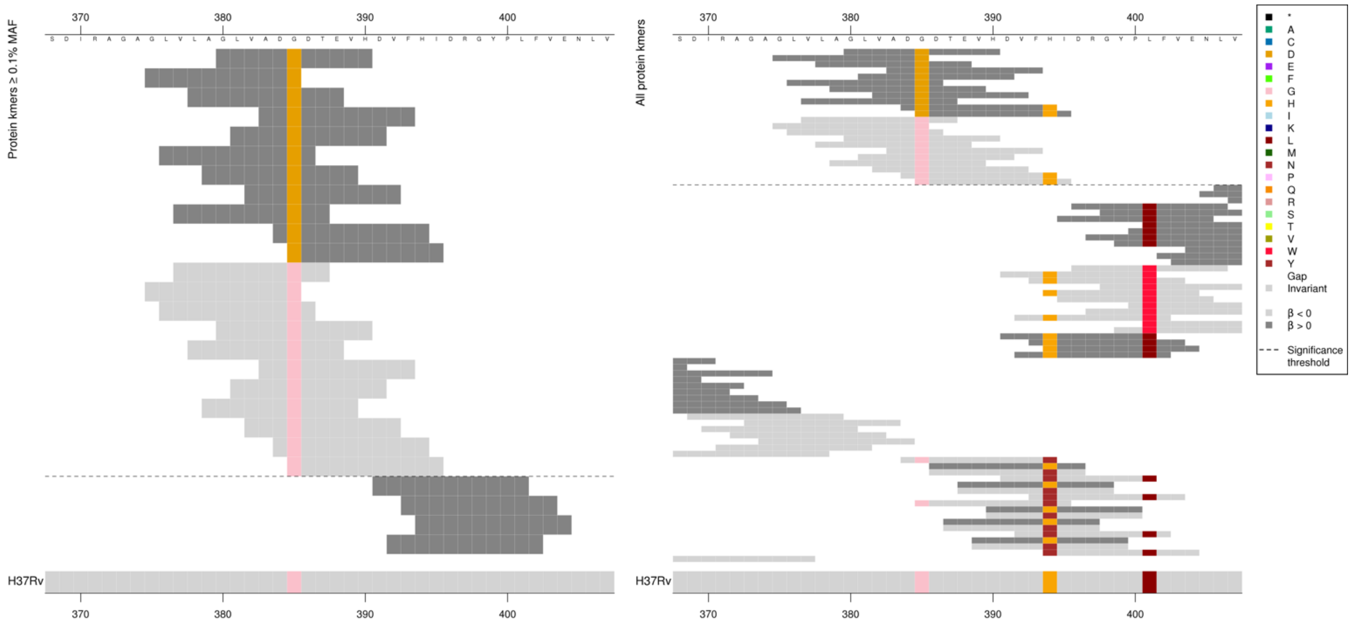

| Kmer | -log₁₀ *p* | β | MAC | MAF | Ps | Variants |
|---|---|---|---|---|---|---|
| GLVADDDTEVH | 10.89 | 1.35 | 128 | 1.46 | 380 | 385D |
| VLAGLVADGDT | 10.75 | -1.31 | 130 | 1.49 | 377 | 385G |

## *Rv1393c*

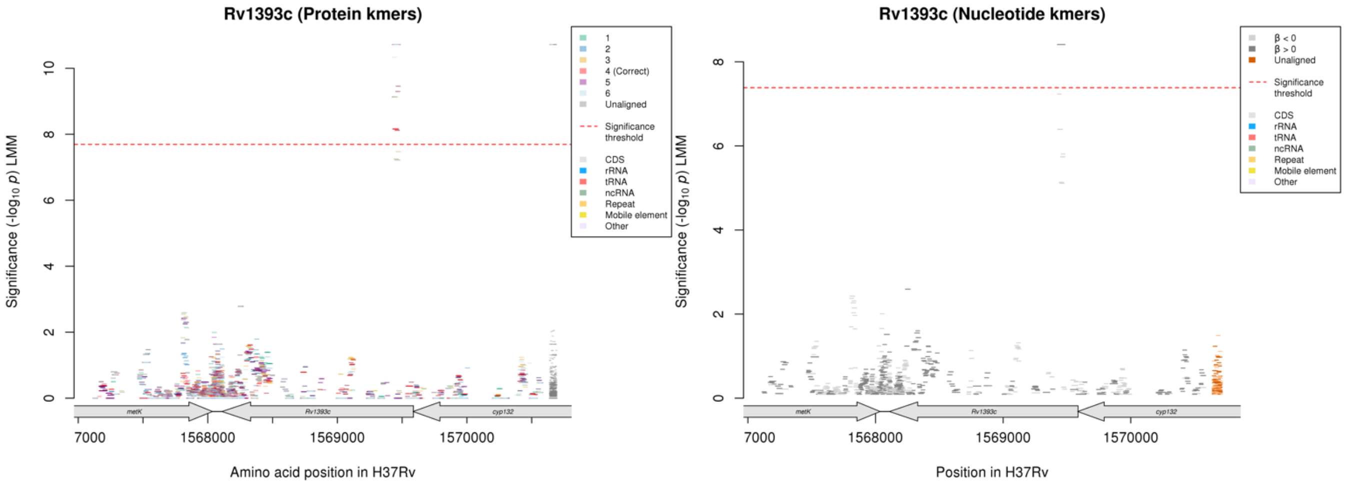

#### Protein kmers:

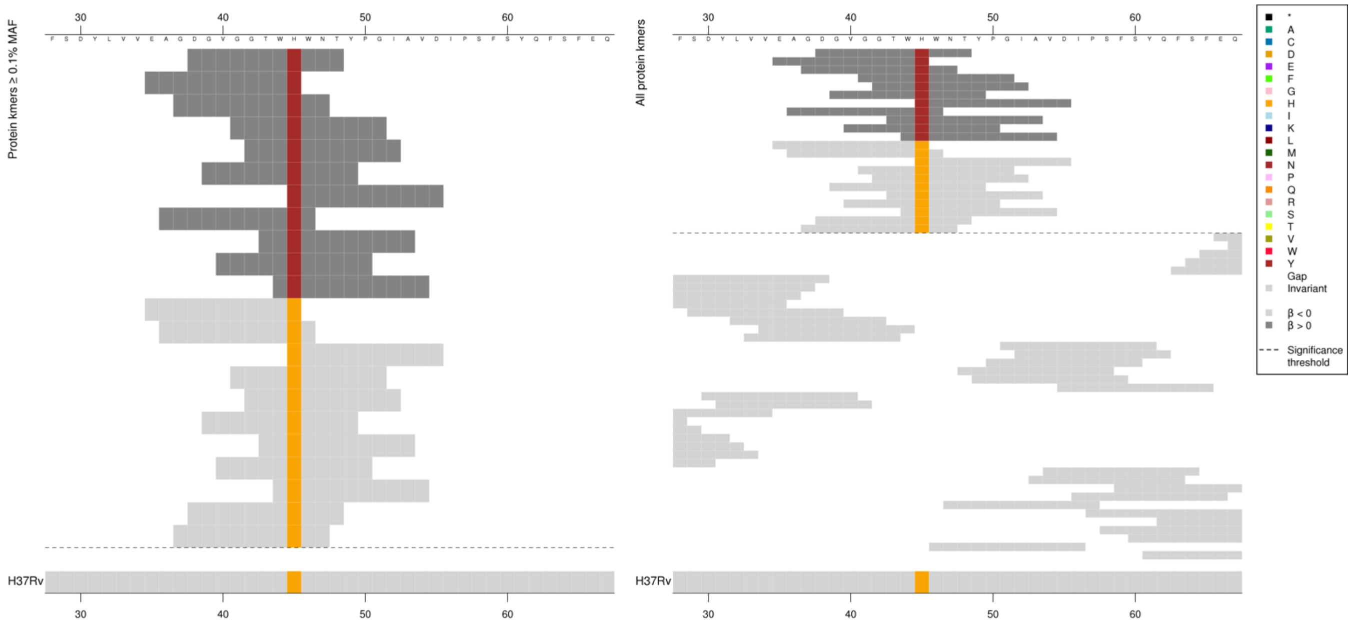

| Kmer | $-log_{10}\,p$ | β | MAC | MAF | Ps | Variants |
|---|---|---|---|---|---|---|
| DGVGGTWNWNT | 10.72 | 2.09 | 18 | 0.21 | 38 | 45N |
| EAGDGVGGTWH | 9.46 | -1.64 | 26 | 0.30 | 35 | 45H |

## *Rv0579*

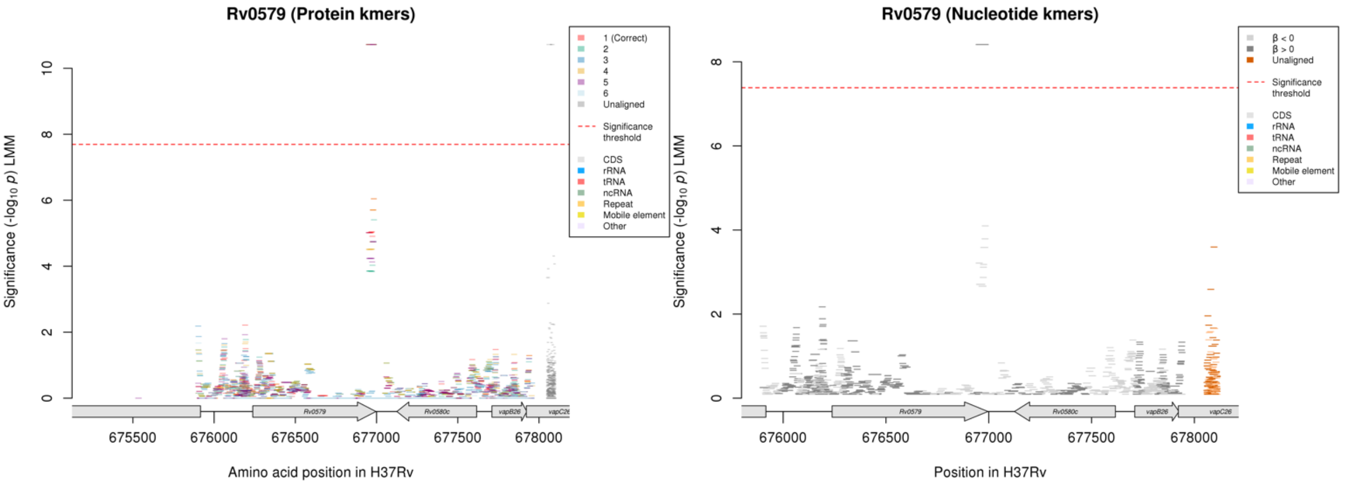

Protein kmers:

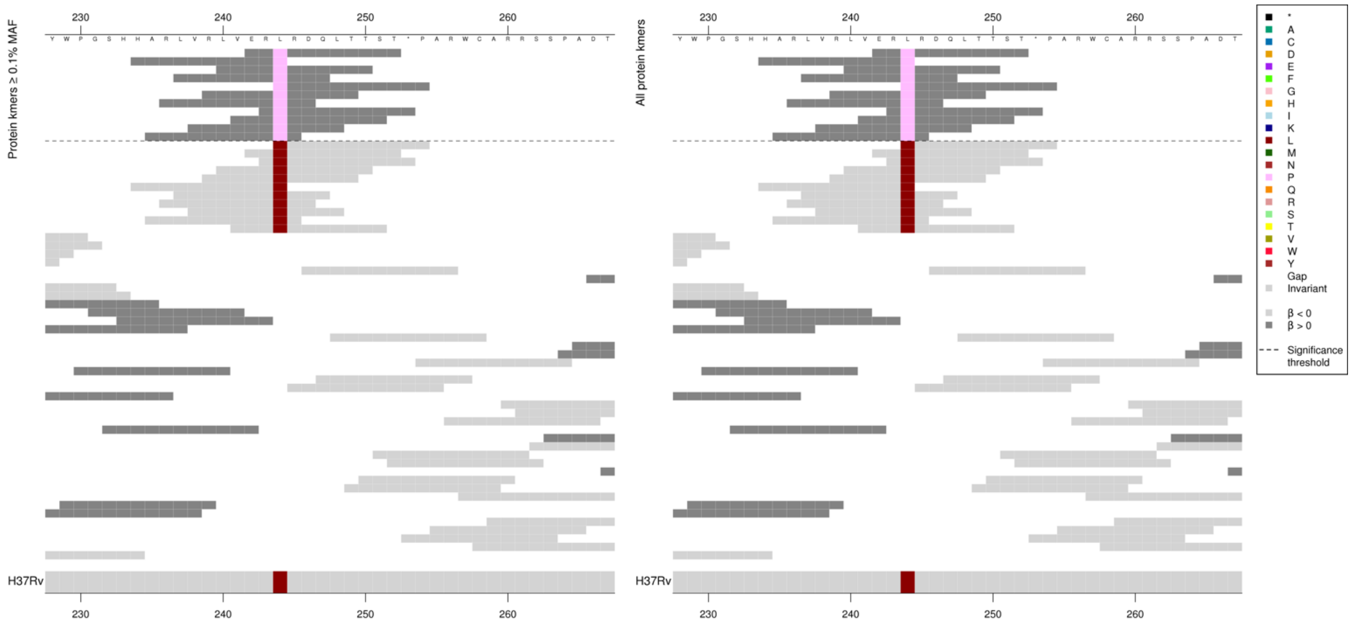

Stop codon is codon 253. The high MIC-associated P containing kmers were significantly associated, however the low MIC associated L kmers were not significant.

| Kmer | $-\log_{10} p$ | β | MAC | MAF | Ps | Variants |
|---|---|---|---|---|---|---|
| ERPRDQLTTST | 10.72 | 2.09 | 18 | 0.21 | 242 | 244P |

#### *glnE*

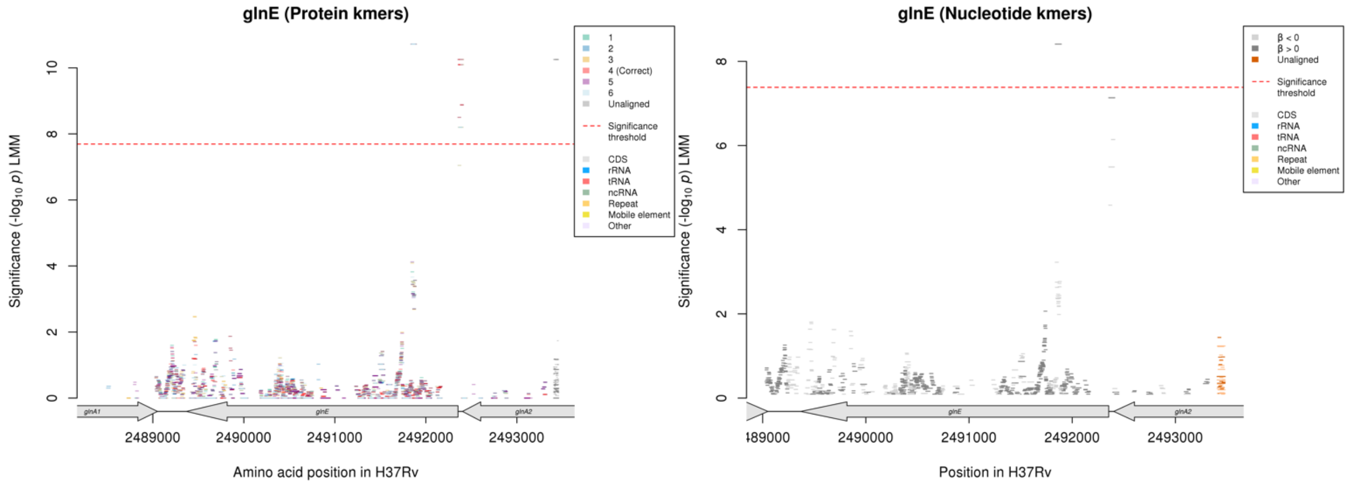

Protein kmers:

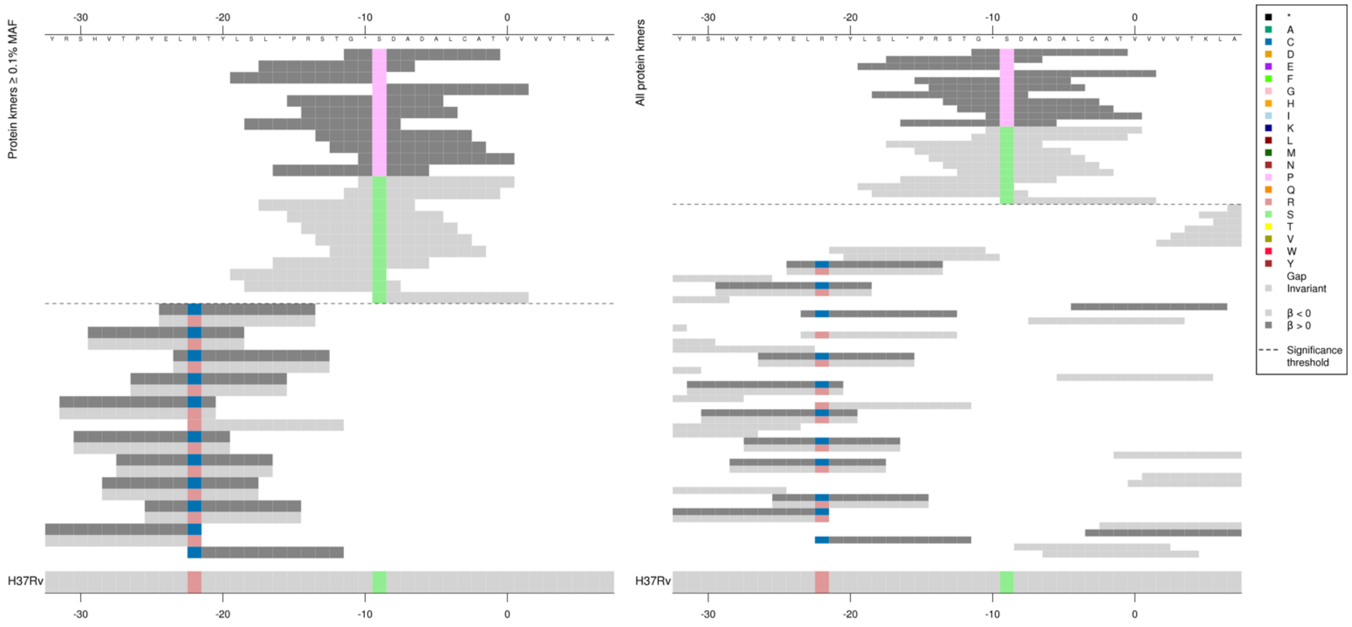

The variant captured in this region by the protein kmer analysis is upstream of the start codon. This region was not significant in the nucleotide kmer analysis, but the alignments are shown below:

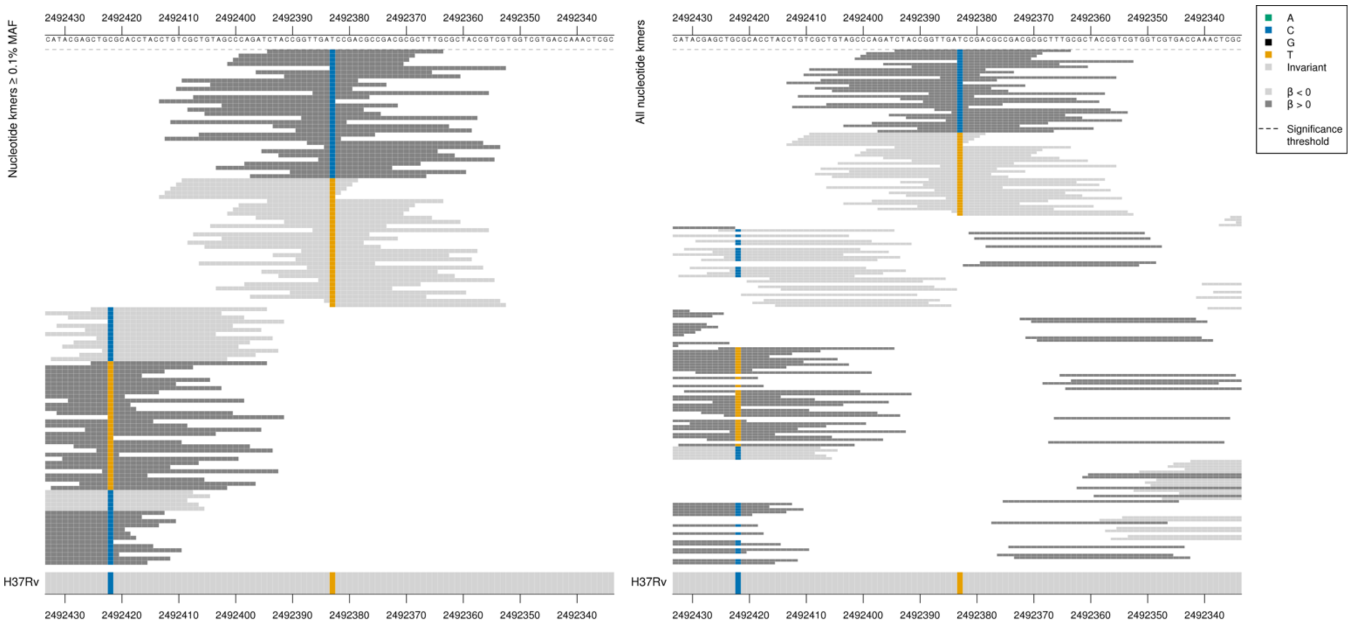

*glnE* begins at position 2492353.
Capturing a mutation at position 2492383/-30.

| Kmer | $-\log_{10} p$ | β | MAC | MAF | Ps | Variants |
|---|---|---|---|---|---|---|
| CTACCGGTTGACCCGACGCCGACGCGCTTTG | 7.14 | 2.03 | 40 | 0.46 | 2492394 | 2492383/-30C |
| CGCTGTAGCCCAGATCTACCGGTTGATCCGA | 6.14 | -1.66 | 42 | 0.48 | 2492409 | 2492383/-30T |

#### Kanamycin (KAN)

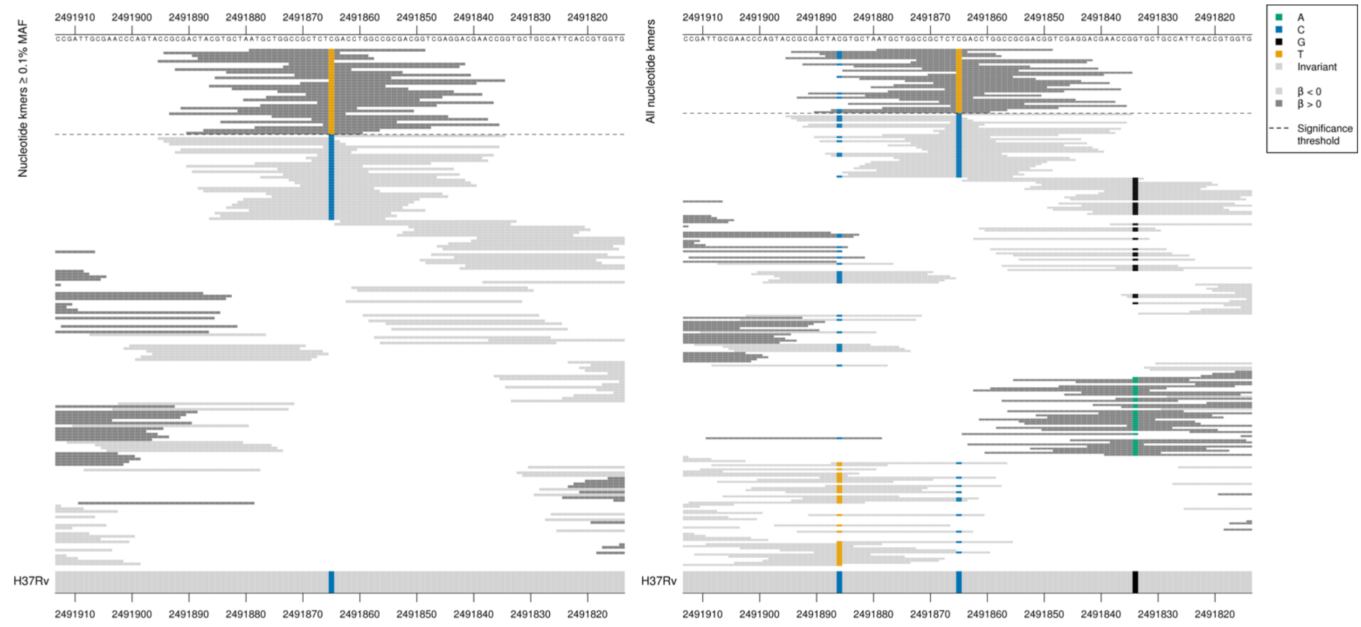

Synonymous mutation at position 2491865/489
High MIC-associated kmers = CTT = Leu / L
Kmers below significance threshold = CTC = Leu / L

| Kmer | -log10 *p* | β | MAC | MAF | Ps | Variants |
|---|---|---|---|---|---|---|
| ATGCTGGCCGCTCTTGACCTGGCCGCGACGG | 8.41 | 1.87 | 18 | 0.21 | 2491879 | 2491865/489T |

#### *rne*

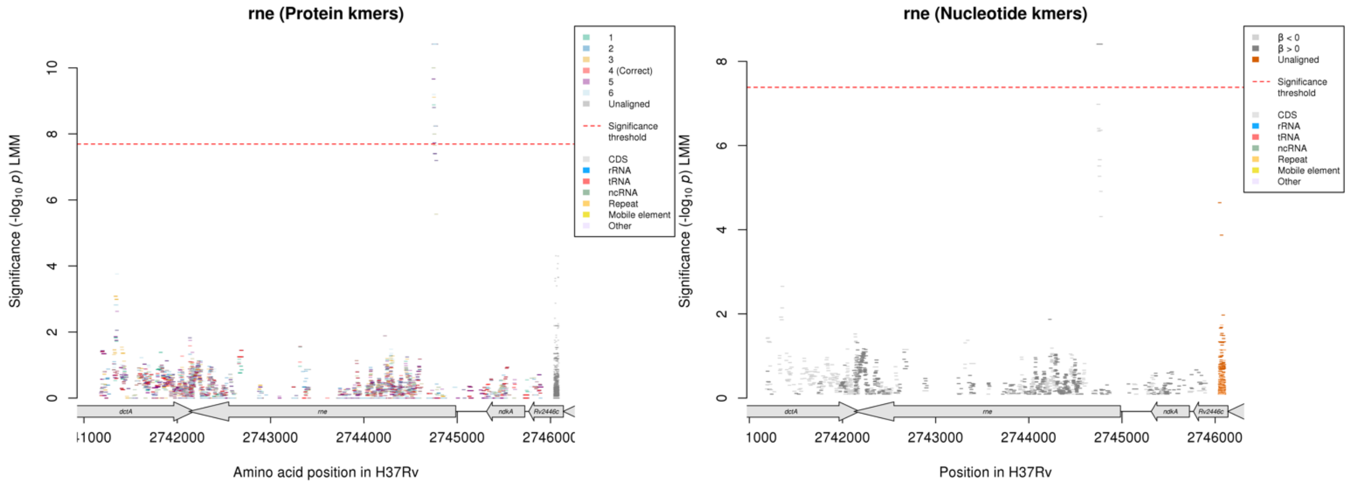

Nucleotide kmers:

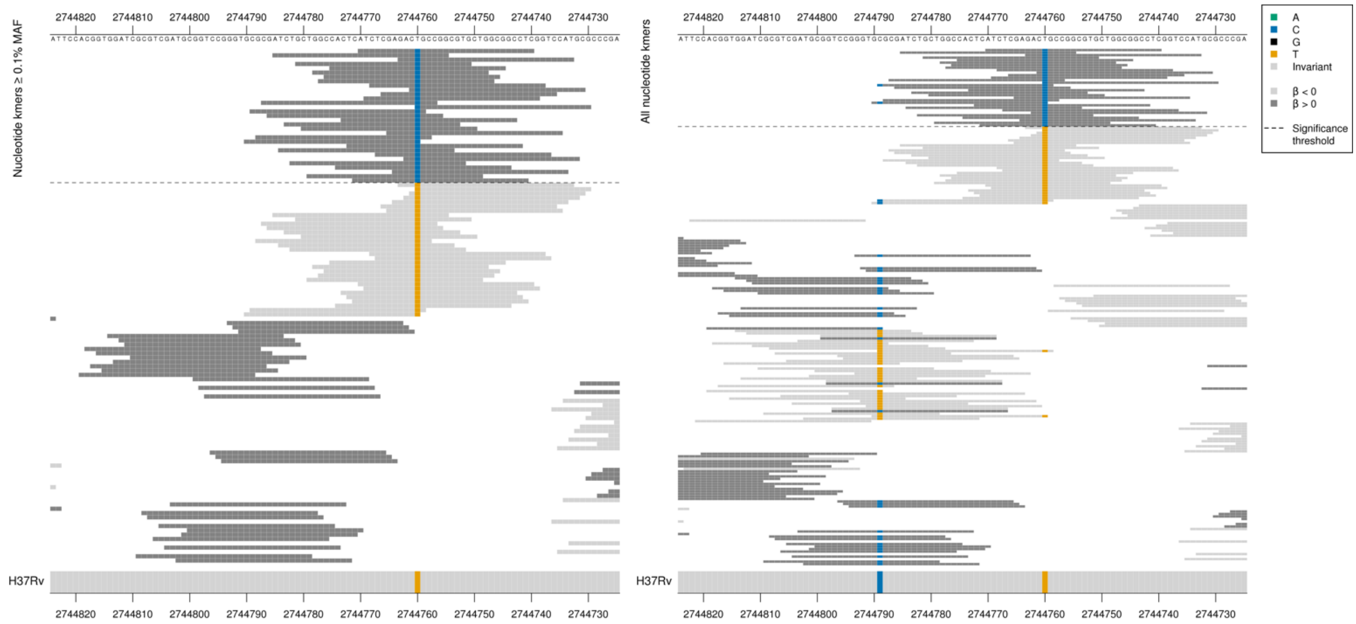

High MIC associated kmers = ACC = Thr / T
Low MIC associated kmers = ACT = Thr / T
Synonymous mutation at position 2744760/225

| Kmer | $-\log_{10} p$ | β | MAC | MAF | Ps | Variants |
|---|---|---|---|---|---|---|
| ATCTCGAGACCGCCGGCGTGCTGGCGGCCTC | 8.41 | 1.87 | 18 | 0.21 | 2744770 | 2744760/225C |

## *Rv1362c*

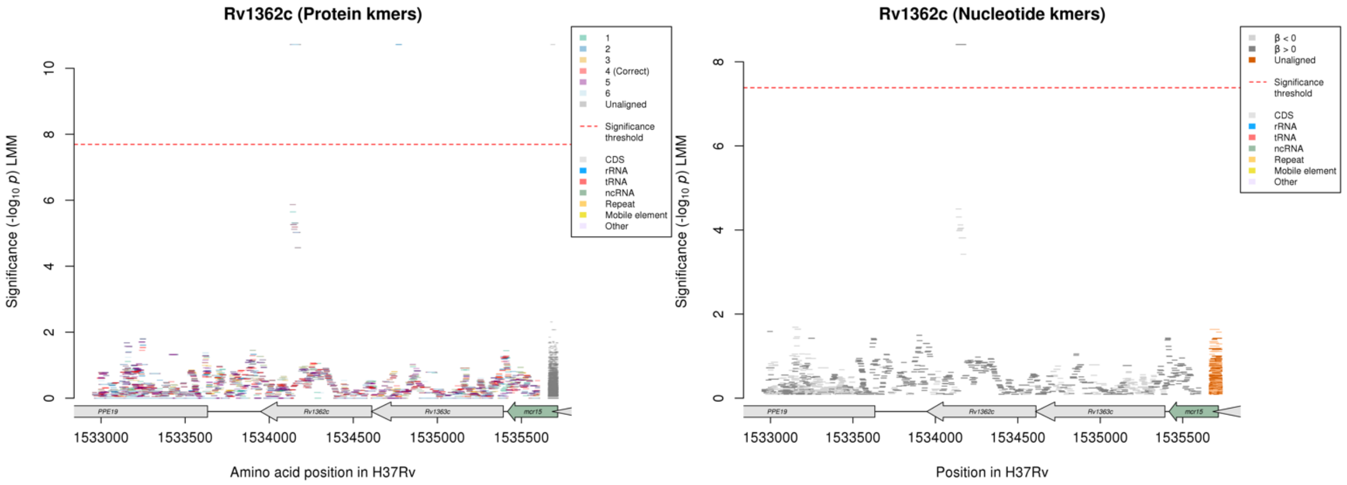

No significant kmers aligned to the correct reading frame in the protein kmer analysis.

#### Nucleotide kmers:

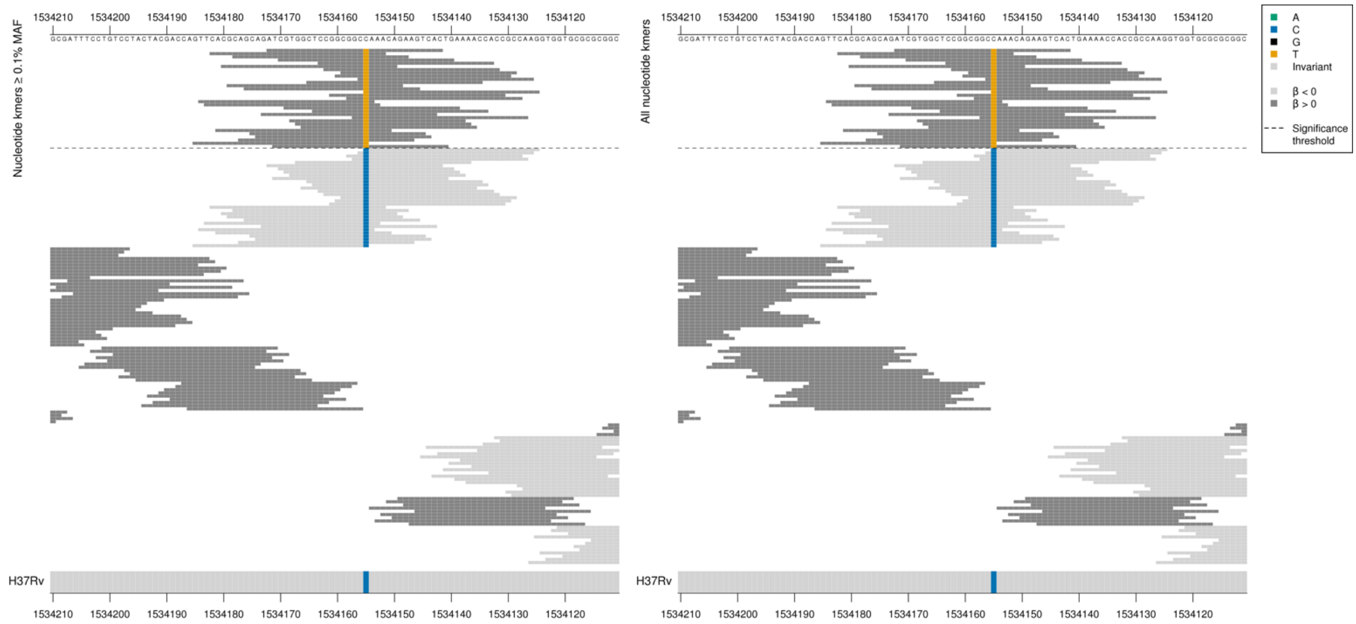

High MIC-associated kmers = GCT = Ala / A
Kmers below significance threshold = GCC = Ala / A

| Kmer | $-\log_{10} p$ | β | MAC | MAF | Ps | Variants |
|---|---|---|---|---|---|---|
| ATCGTGGCTCCGGCGGCTAAACAGAAGTCAC | 8.41 | 1.87 | 18 | 0.21 | 1534172 | 1534155/456T |

## *Rv0208c:Rv0209*

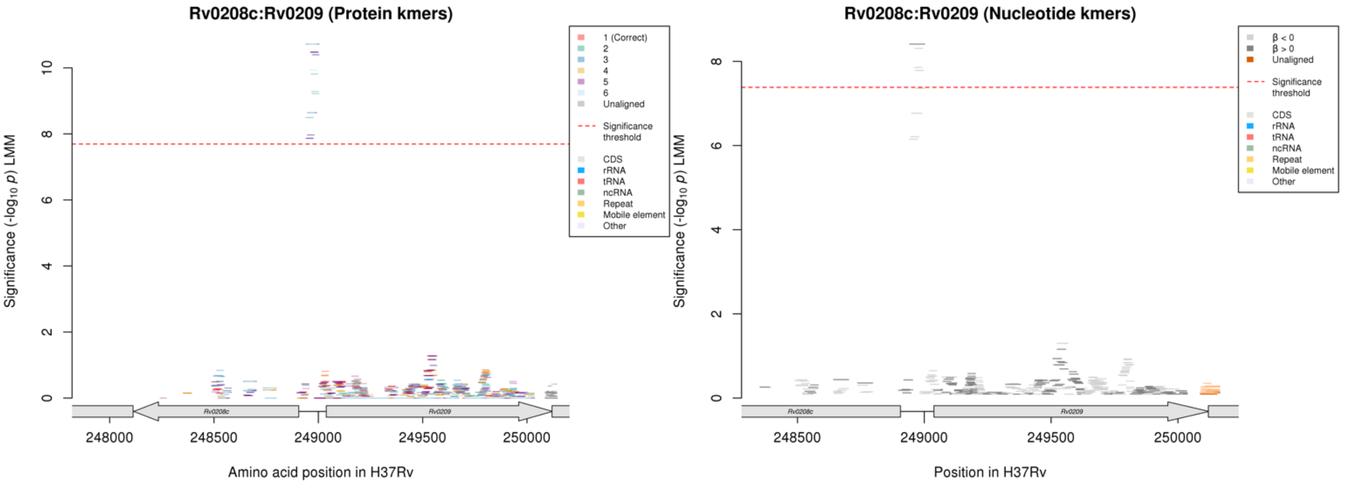

Nucleotide kmers:

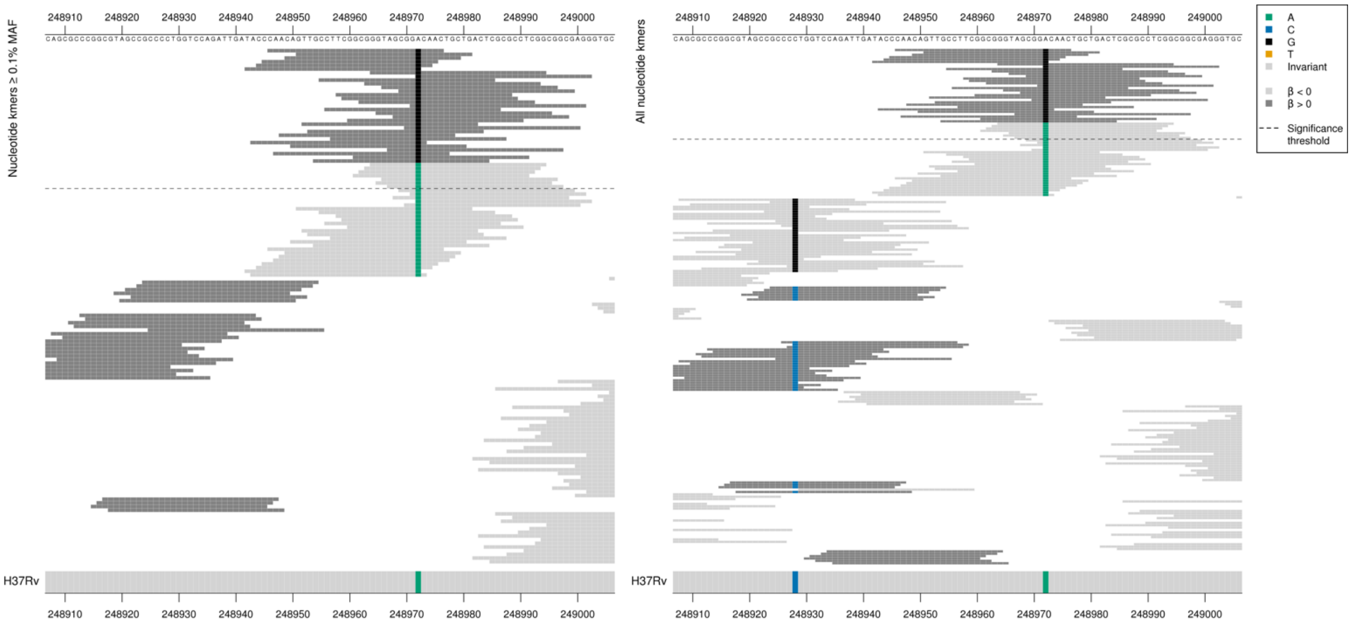

Gene *Rv0208c* starts at position 248906 on the reverse strand and *Rv0209* starts at position 249038 on the forward strand. The kmers were capturing a mutation in the intergenic region, position 248972, 67bp from both genes.

Significant kmers on the forward strand:

High MIC-associated kmers = G

Low MIC-associated kmers = A

| Kmer | $-\log_{10} p$ | β | MAC | MAF | Ps | Variants |
|---|---|---|---|---|---|---|
| CAACAGTTGCCTTCGGCGGGTAGCGGGCAAC | 8.41 | 1.87 | 18 | 0.21 | 248946 | 248972 (67bp from both genes) G (forward strand) |
| GGTAGCGGACAACTGCTGACTCGCGCCTCGG | 8.31 | -1.76 | 20 | 0.23 | 248964 | 248972 (67bp from both genes) A (forward strand) |

### Levofloxacin (LEV)

#### *gyrA*

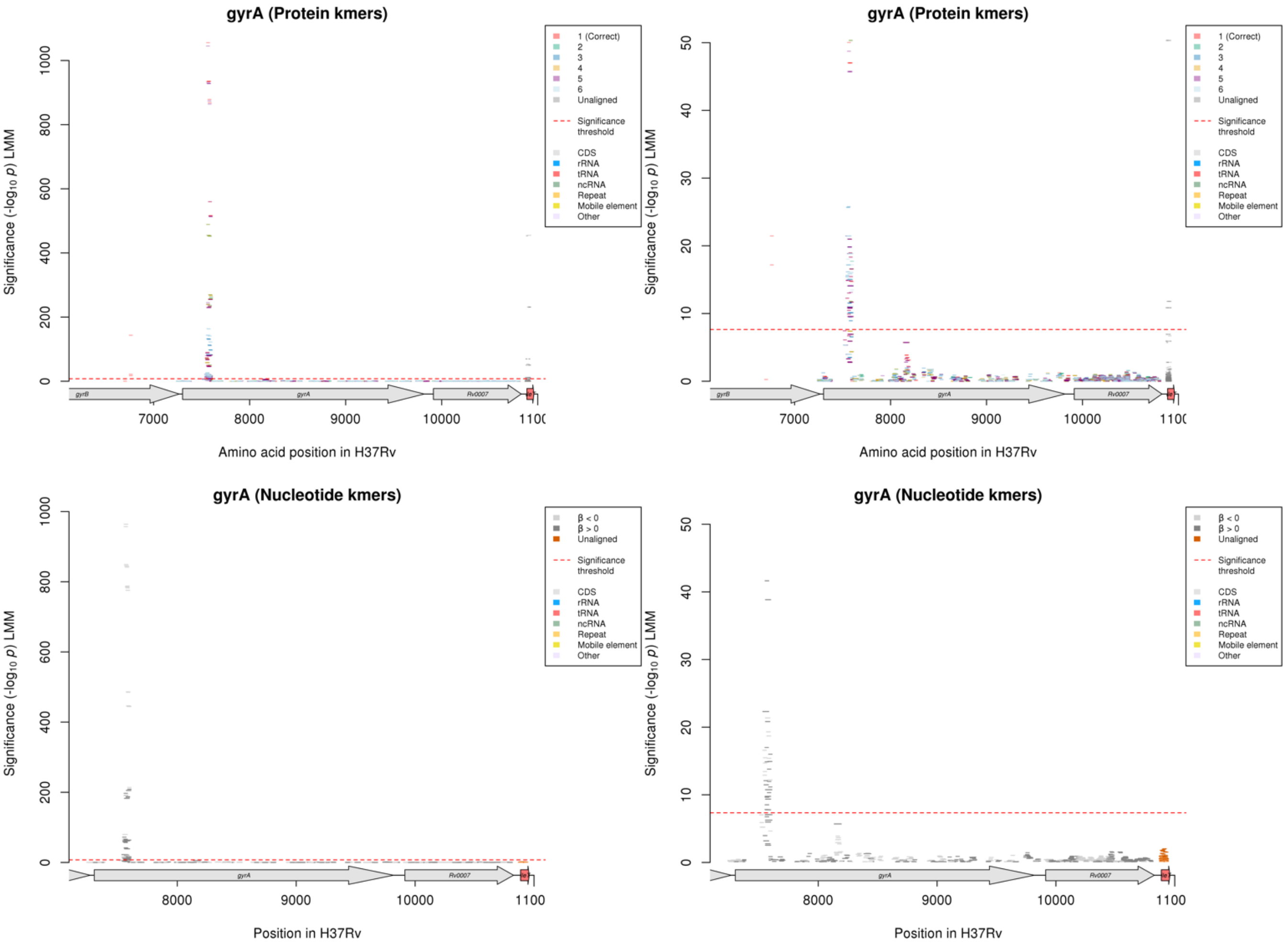

Protein kmers:

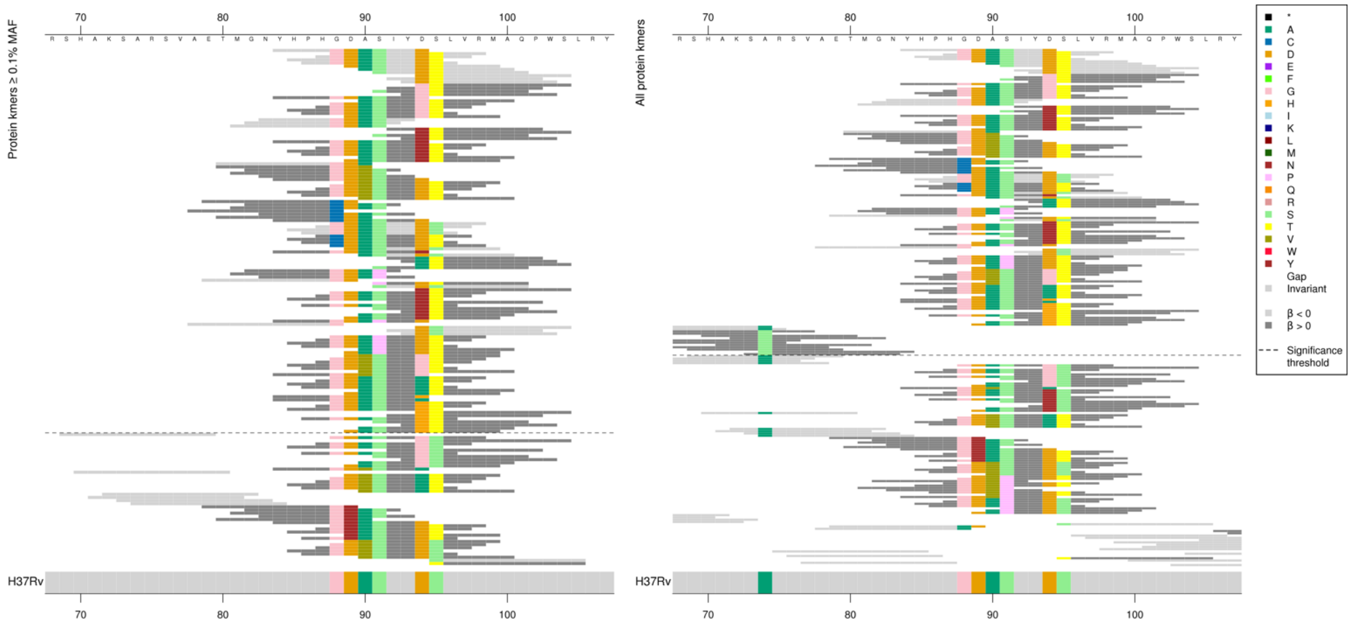

Main codons being captured: 74, 88, 89, 90, 91, 94, 95

Kmers below significance threshold: 89N, 74A

| Kmer | $-\log_{10} p$ | β | MAC | MAF | Ps | Variants |
|---|---|---|---|---|---|---|
| YHPHGDASIYD | 1055.64 | -3.48 | 1106 | 15.15 | 84 | 94D [+ 88G, 89D, 90A, 91S] |
| GDASIYDTLVR | 935.11 | -3.38 | 1489 | 20.40 | 88 | 94D [+ 88G, 89D, 90A, 91S, 95T] |
| DASIYDTLVRM | 878.69 | -3.32 | 1476 | 20.22 | 89 | 94D [+ 89D, 90A, 91S, 95T] |
| ASIYDTLVRMA | 868.75 | -3.31 | 1468 | 20.11 | 90 | 94D [+ 90A, 91S, 95T] |
| SIYDTLVRMAQ | 559.60 | -2.97 | 1207 | 16.53 | 91 | 94D [+ 91S, 95T] |
| DTLVRMAQPWS | 516.20 | -2.94 | 1159 | 15.88 | 94 | 94D [+ 95T] |
| GTLVRMAQPWS | 255.09 | 2.49 | 517 | 7.08 | 94 | 94G [+ 95T] |
| SIYGTLVRMAQ | 255.09 | 2.49 | 517 | 7.08 | 91 | 94G [+ 91S, 95T] |
| YHPHGDASIYG | 238.79 | 2.40 | 517 | 7.08 | 84 | 94G [+ 88G, 89D, 90A, 91S] |
| ASIYGTLVRMA | 233.54 | 2.41 | 506 | 6.93 | 90 | 94G [+ 90A, 91S] |
| GDASIYGTLVR | 229.82 | 2.39 | 504 | 6.90 | 88 | 94G [+ 88G, 89D, 90A, 91S, 95T] |
| DASIYGTLVRM | 229.82 | 2.39 | 504 | 6.90 | 89 | 94G [+ 89D, 90A, 91S, 95T] |
| NYHPHGDASIY | 89.13 | -1.54 | 354 | 4.85 | 83 | 88G, 89D, 90A, 91S |
| IYNTLVRMAQP | 81.53 | 2.36 | 96 | 1.32 | 92 | 94N [+ 95T] |
| SIYNTLVRMAQ | 81.53 | 2.36 | 96 | 1.32 | 91 | 94N [+ 91S, 95T] |
| YHPHGDASIYN | 81.38 | 2.36 | 94 | 1.29 | 84 | 94N [+ 88G, 89D, 90A, 91S] |
| GDASIYNTLVR | 79.03 | 2.34 | 93 | 1.27 | 88 | 94N [+ 88G, 89D, 90A, 91S, 95T] |
| ASIYNTLVRMA | 79.03 | 2.34 | 93 | 1.27 | 90 | 94N [+ 90A, 91S, 95T] |
| DASIYNTLVRM | 79.03 | 2.34 | 93 | 1.27 | 89 | 94N [+ 89D, 90A, 91S, 95T] |
| TMGNYHPHGDA | 73.56 | -1.49 | 310 | 4.25 | 80 | 88G, 89D, 90A |
| TMGNYHPHGDV | 71.85 | 1.48 | 311 | 4.26 | 80 | 90V [+ 88G, 89D] |
| GNYHPHGDVSI | 68.24 | 1.46 | 305 | 4.18 | 82 | 90V [+ 88G, 89D, 91S] |
| YHPHGDVSIYD | 50.03 | 1.28 | 281 | 3.85 | 84 | 90V [+ 88G, 89D, 91S, 94D] |
| DVSIYDTLVRM | 47.01 | 1.26 | 269 | 3.68 | 89 | 90V [+ 89D, 91S, 94D, 95T] |
| GDVSIYDTLVR | 47.01 | 1.26 | 269 | 3.68 | 88 | 90V [+ 88G, 89D, 91S, 94D, 95T] |
| VSIYDTLVRMA | 47.01 | 1.26 | 269 | 3.68 | 90 | 90V [+ 91S, 94D, 95T] |
| ETMGNYHPHCD | 21.45 | 3.23 | 12 | 0.16 | 79 | 88C [+ 89D] |
| GNYHPHCDASI | 21.45 | 3.23 | 12 | 0.16 | 82 | 88C [+ 89D, 90A, 91S] |
| AETMGNYHPHC | 21.45 | 3.23 | 12 | 0.16 | 78 | 88C |
| TMGNYHPHCDA | 21.45 | 3.23 | 12 | 0.16 | 80 | 88C [+ 89D, 90A] |
| YHPHCDASIYD | 21.45 | 3.23 | 12 | 0.16 | 84 | 88C [+ 89D, 90A, 91S, 94D] |
| GDASIYDSLVR | 20.98 | -2.12 | 383 | 5.25 | 88 | 88G, 89D, 90A, 91S, 94D, 95S |
| HCDASIYDTLV | 19.82 | 3.23 | 11 | 0.15 | 87 | 88C [+ 89D, 90A, 91S, 94D, 95T] |
| DASIYDSLVRM | 18.96 | -2.02 | 384 | 5.26 | 89 | 89D, 90A, 91S, 94D, 95S |
| YHPHGDASIYY | 18.84 | 1.89 | 36 | 0.49 | 84 | 94Y [+88G, 89D, 90A, 91S] |
| ASIYDSLVRMA | 18.33 | -2.00 | 385 | 5.27 | 90 | 90A, 91S, 94D, 95S |
| IYATLVRMAQP | 16.58 | 1.10 | 92 | 1.26 | 92 | 94A [+ 95T] |
| SIYATLVRMAQ | 15.46 | 1.07 | 91 | 1.25 | 91 | 94A [+ 91S, 95T] |
| GNYHPHGDAPI | 14.91 | 1.40 | 52 | 0.71 | 82 | 91P [+ 88G, 89D, 90A] |
| ETMGNYHPHGD | 14.88 | -1.88 | 28 | 0.38 | 79 | 88G, 89D |
| PIYDTLVRMAQ | 14.81 | 1.41 | 52 | 0.71 | 91 | 91P [+ 94D, 95T] |

| | | | | | | |
|---|---|---|---|---|---|---|
| SIYDSLVRMAQ | 14.79 | -1.97 | 397 | 5.44 | 91 | 91S, 94D, 95S |
| YTLVRMAQPWS | 14.72 | 1.73 | 31 | 0.42 | 94 | 94Y [+ 95T] |
| GDASIYYTLVR | 14.08 | 1.71 | 30 | 0.41 | 88 | 94Y [+ 88G, 89D, 90A, 91S, 95T] |
| SIYYTLVRMAQ | 14.08 | 1.71 | 30 | 0.41 | 91 | 94Y [+ 91S, 95T] |
| ASIYYTLVRMA | 14.08 | 1.71 | 30 | 0.41 | 90 | 94Y [+ 90A, 91S, 95T] |
| DASIYYTLVRM | 14.08 | 1.71 | 30 | 0.41 | 89 | 94Y [+89D, 90A, 91S, 95T] |
| YHPHGDAPIYD | 13.02 | 1.34 | 50 | 0.68 | 84 | 91P [+ 88G, 89D, 90A, 94D] |
| AETMGNYHPHG | 12.26 | -2.11 | 18 | 0.25 | 78 | 88G |
| DSLVRMAQPWS | 11.67 | -1.86 | 402 | 5.51 | 94 | 94D, 95S |
| GDAPIYDTLVR | 10.95 | 1.30 | 45 | 0.62 | 88 | 91P [+88G, 89D, 90A, 94D, 95T] |
| APIYDTLVRMA | 10.95 | 1.30 | 45 | 0.62 | 90 | 91P [+ 90A, 94D, 95T] |
| DAPIYDTLVRM | 10.95 | 1.30 | 45 | 0.62 | 89 | 91P [+ 89D, 90A, 94D, 95T] |
| DVSIYGTLVRM | 10.85 | 2.68 | 10 | 0.14 | 89 | 90V, 94G [+ 89D, 91S, 95T] |
| GDVSIYGTLVR | 10.85 | 2.68 | 10 | 0.14 | 88 | 90V, 94G [+ 88G, 89D, 91S, 95T] |
| VSIYGTLVRMA | 10.85 | 2.68 | 10 | 0.14 | 90 | 90V, 94G [+ 91S, 95T] |
| YHPHGDVSIYG | 10.85 | 2.68 | 10 | 0.14 | 84 | 90V, 94G [+ 88G, 89D, 91S] |
| GDASIYATLVR | 10.79 | 0.94 | 80 | 1.10 | 88 | 94A [+ 88G, 89D, 90A, 91S, 95T] |
| ASIYATLVRMA | 10.79 | 0.94 | 80 | 1.10 | 90 | 94A [+ 90A, 91S, 95T] |
| DASIYATLVRM | 10.79 | 0.94 | 80 | 1.10 | 89 | 94A [+ 89D, 90A, 91S, 95T] |
| YHPHGDASIYH | 10.47 | 1.36 | 33 | 0.45 | 84 | 94H [+ 88G, 89D, 90A, 91S] |
| YHPHGDASIYA | 9.91 | 0.89 | 81 | 1.11 | 84 | 94A [+ 88G, 89D, 90A, 91S] |
| GDASIYHTLVR | 9.55 | 1.32 | 32 | 0.44 | 88 | 94H [+ 88G, 89D, 90A, 91S, 95T] |
| HTLVRMAQPWS | 9.55 | 1.32 | 32 | 0.44 | 94 | 94H [+ 95T] |
| SIYHTLVRMAQ | 9.55 | 1.32 | 32 | 0.44 | 91 | 94H [+ 91S, 95T] |
| ASIYHTLVRMA | 9.55 | 1.32 | 32 | 0.44 | 90 | 94H [+ 90A, 91S, 95T] |
| DASIYHTLVRM | 9.55 | 1.32 | 32 | 0.44 | 89 | 94H [+ 89D, 90A, 91S, 95T] |

#### *rrs*

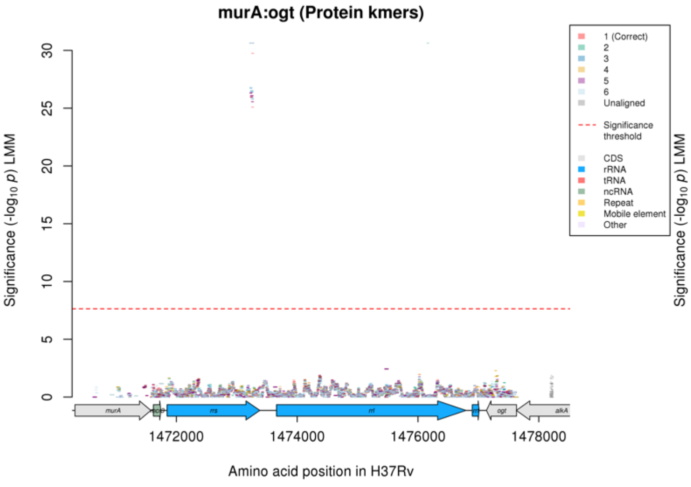

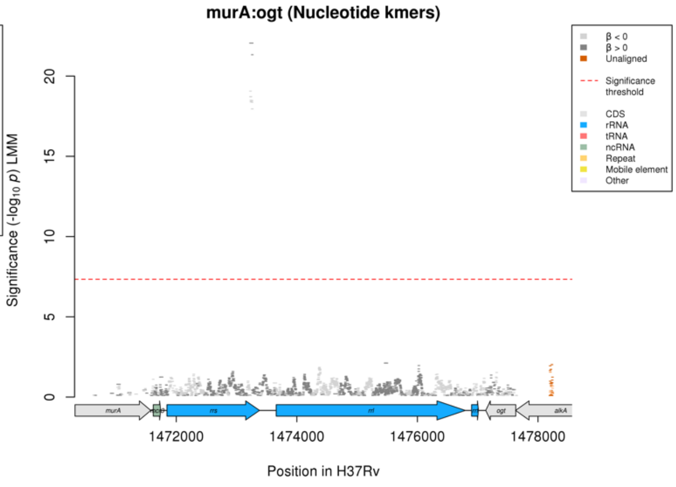

### Levofloxacin (LEV)

#### Nucleotide kmers:

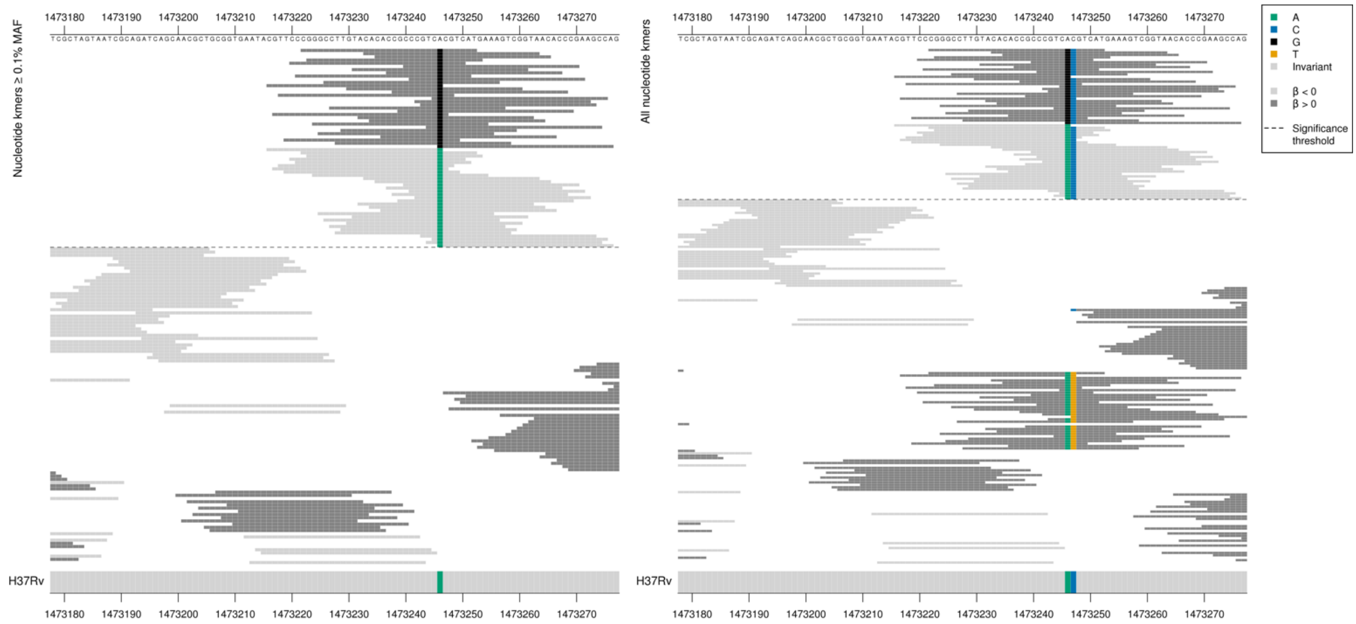

| Kmer | $-\log_{10} p$ | β | MAC | MAF | Ps | Variants |
|---|---|---|---|---|---|---|
| CGGGCCTTGTACACACCGCCCGTCGCGTCAT | 22.05 | 0.89 | 424 | 5.81 | 1473222 | 1473246/1401G |
| CGTTCCCGGGCCTTGTACACACCGCCCGTCA | 19.06 | -0.81 | 460 | 6.30 | 1473216 | 1473246/1401A |

#### *gyrB*

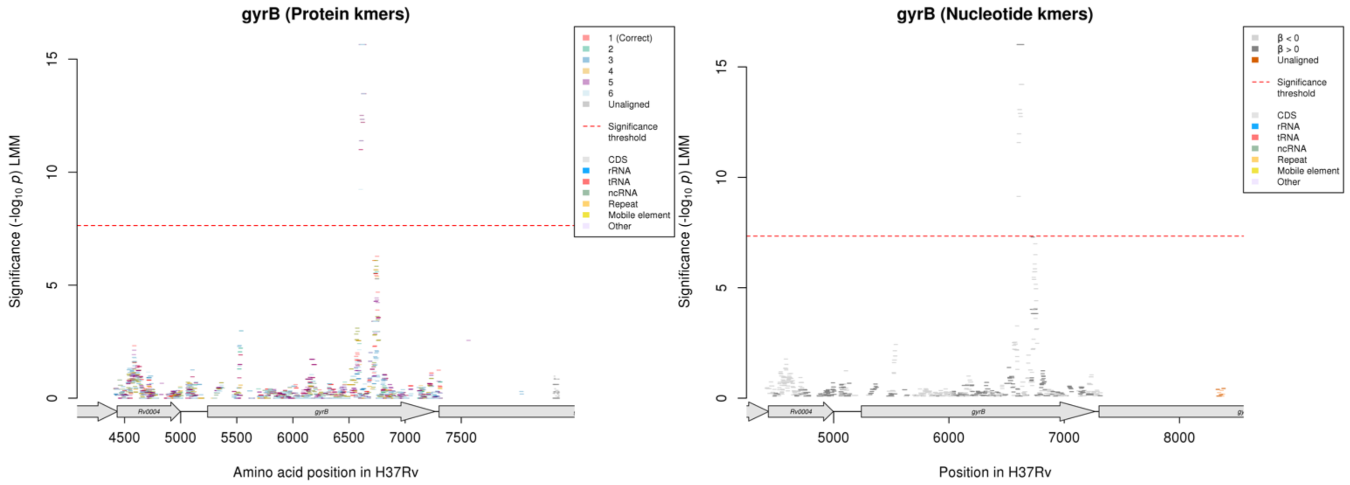

### Levofloxacin (LEV)

#### Protein kmers:

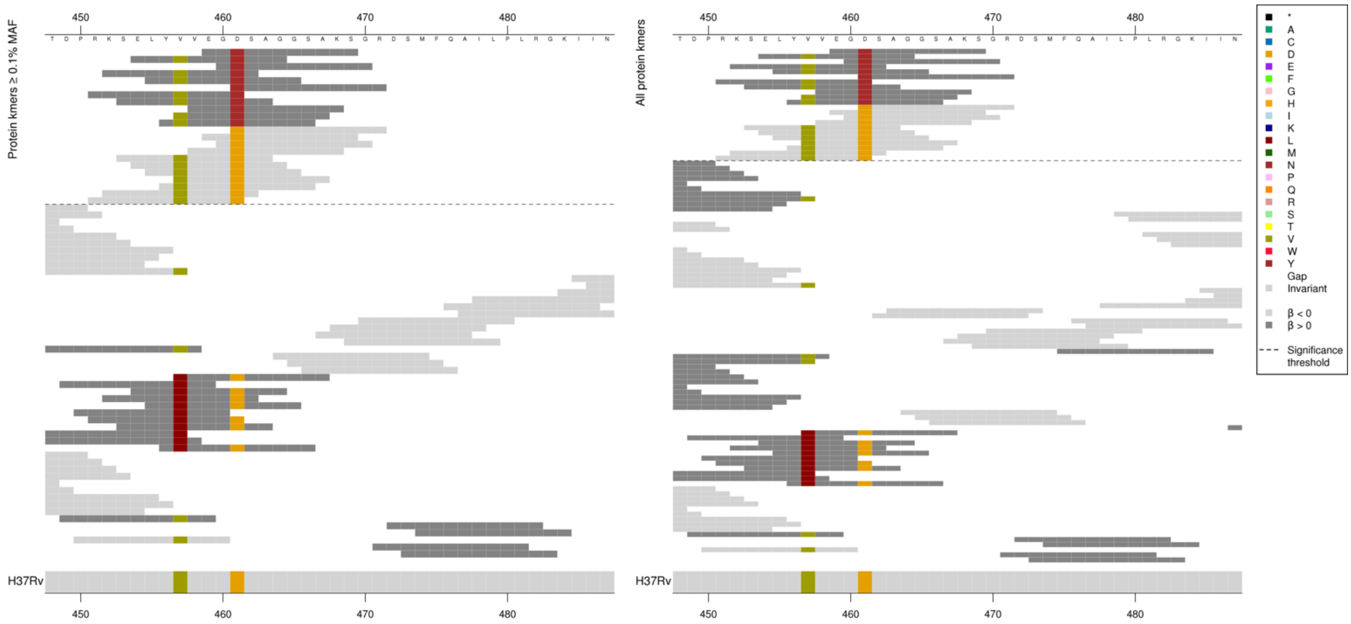

Kmers below significance threshold = 457L

| Kmer | $-\log_{10} p$ | β | MAC | MAF | Ps | Variants |
|---|---|---|---|---|---|---|
| EGNSAGGSAKS | 15.65 | 2.46 | 15 | 0.21 | 459 | 461N |
| DSAGGSAKSGR | 13.47 | -2.14 | 22 | 0.30 | 461 | 461D |
| SELYVVEGDSA | 12.51 | -1.96 | 28 | 0.38 | 453 | 457V, 461D |

#### *embB*

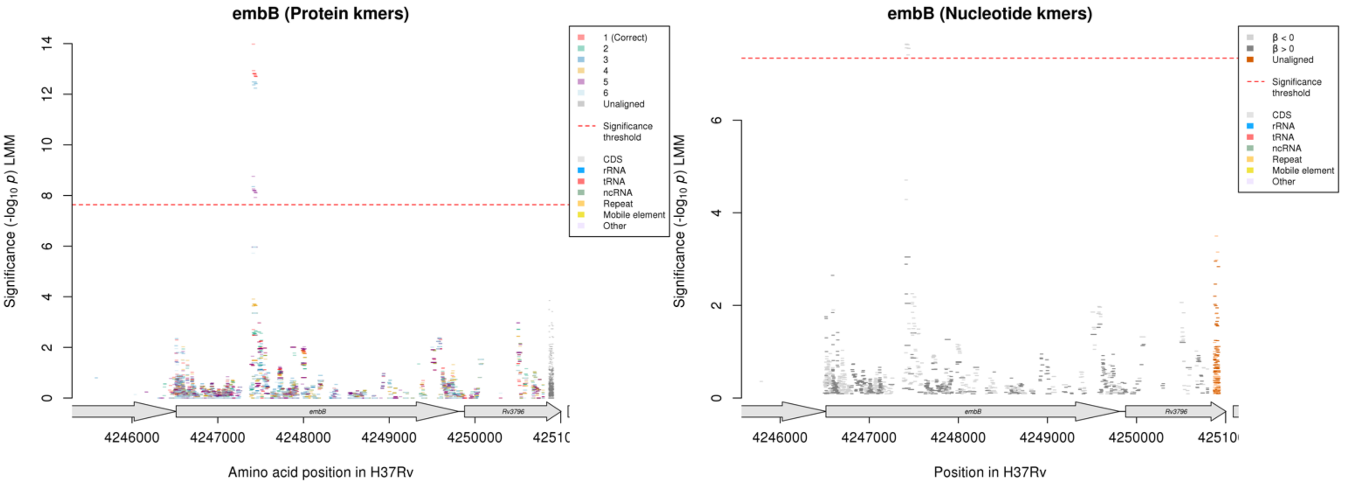

Protein kmers:

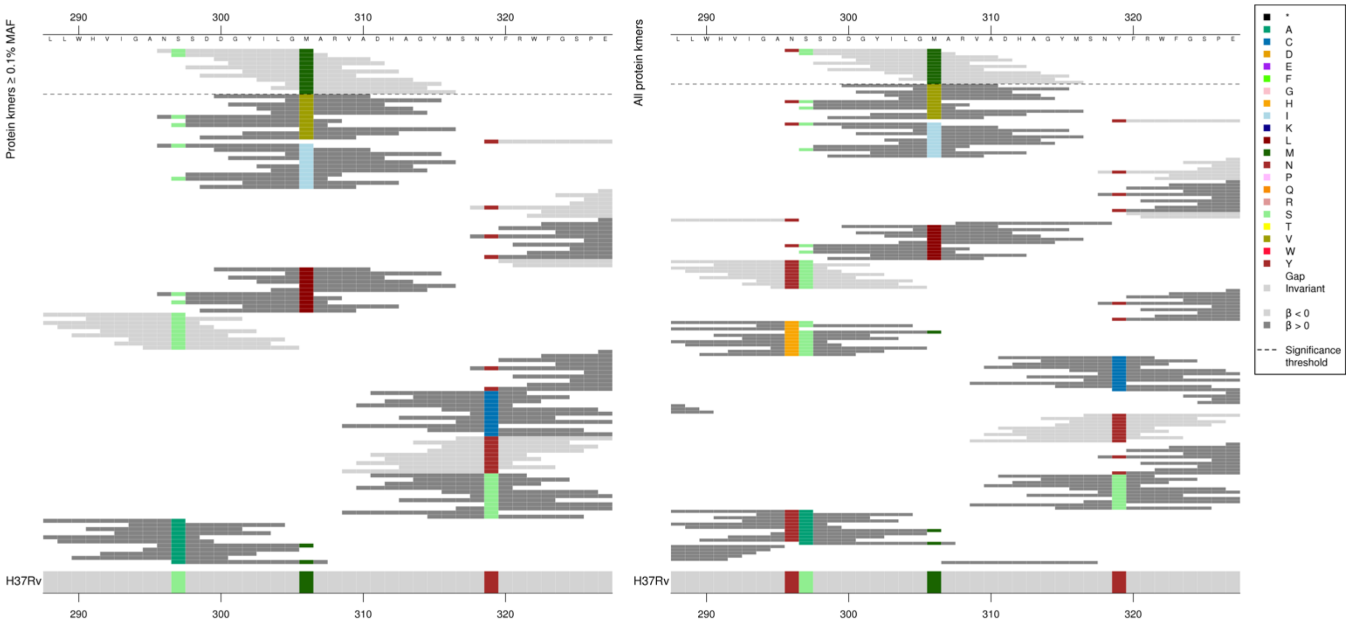

Kmers below significance threshold at codon 306 = V/I/L

| Kmer | -log$_{10}$ *p* | β | MAC | MAF | Ps | Variants |
|---|---|---|---|---|---|---|
| NSSDDGYILGM | 13.98 | -0.50 | 1405 | 19.25 | 296 | 306M |

*rpoB*

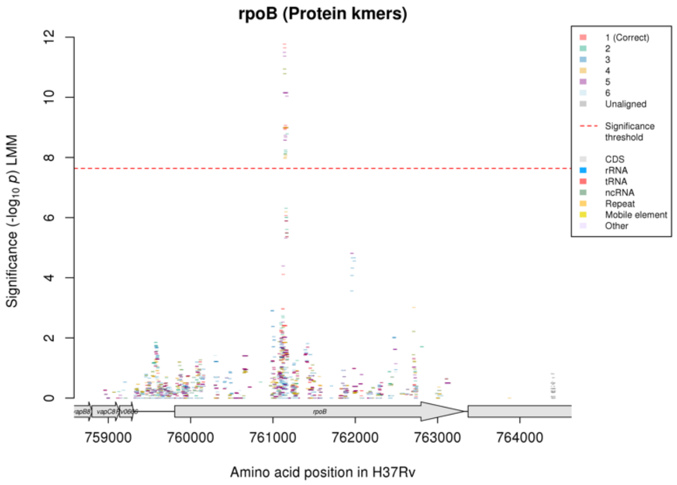

Not significant for nucleotide kmer analysis.

### Levofloxacin (LEV)

Protein kmers:

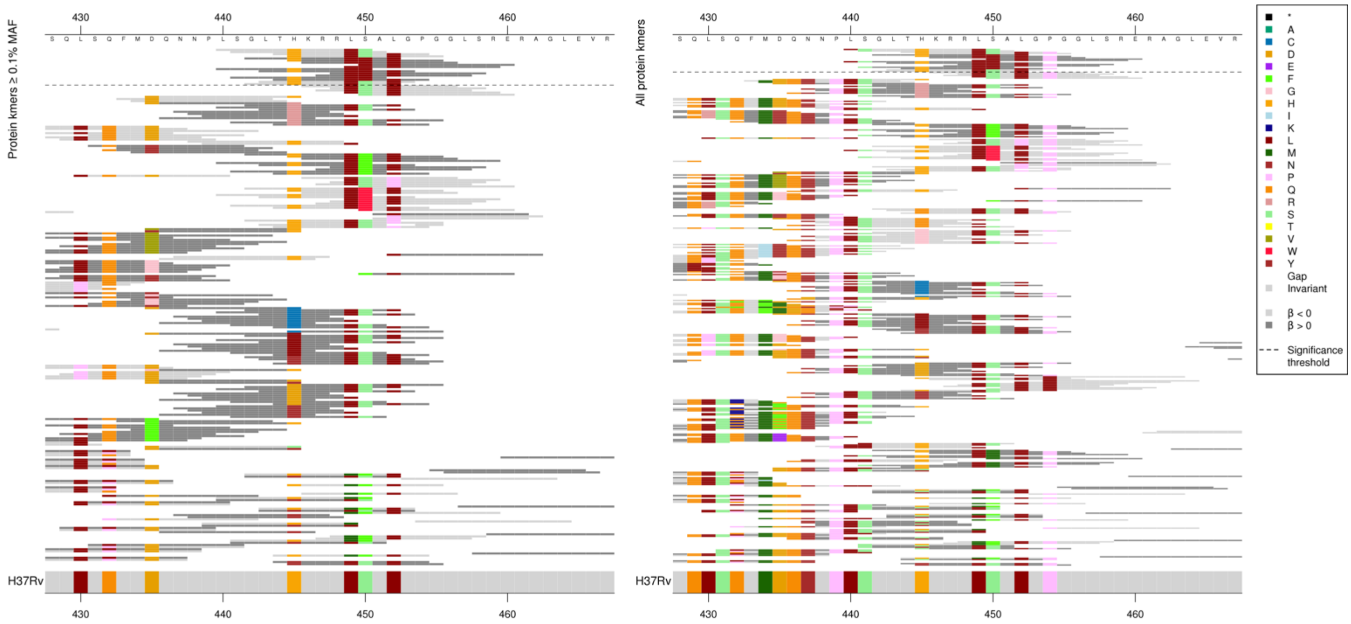

Main association is codon 450. Low MIC-associated kmer *p*-values varied depending on other positions, high MIC-associated kmers were all the same *p*-value and just capturing codon 450L.

| Kmer | $-\log_{10} p$ | β | MAC | MAF | Ps | Variants |
|---|---|---|---|---|---|---|
| LSGLTHKRRLS | 11.77 | -0.41 | 2108 | 28.88 | 440 | 440L, 441S, 445H, 449L, 450S |
| SGLTHKRRLSA | 11.64 | -0.41 | 2110 | 28.90 | 441 | 441S, 445H, 449L, 450S |
| THKRRLSALGP | 9.07 | -0.36 | 2189 | 29.99 | 444 | 445H, 449L, 450S, 452L, 454P |
| GLTHKRRLLAL | 8.99 | 0.40 | 1749 | 23.96 | 442 | 450L |
| LTHKRRLSALG | 8.96 | -0.35 | 2182 | 29.89 | 443 | 445H, 449L, 450S, 452L |

#### *vapC36*

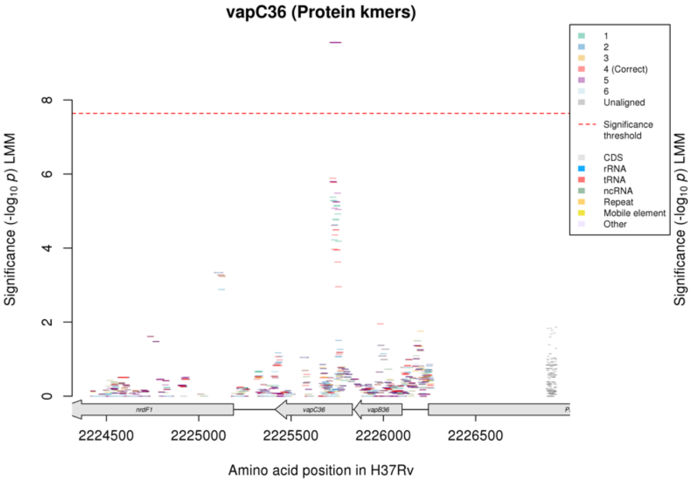

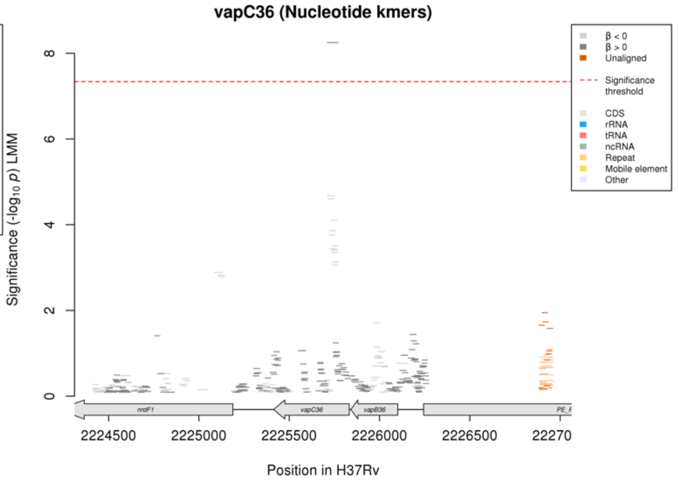

#### Protein kmers:

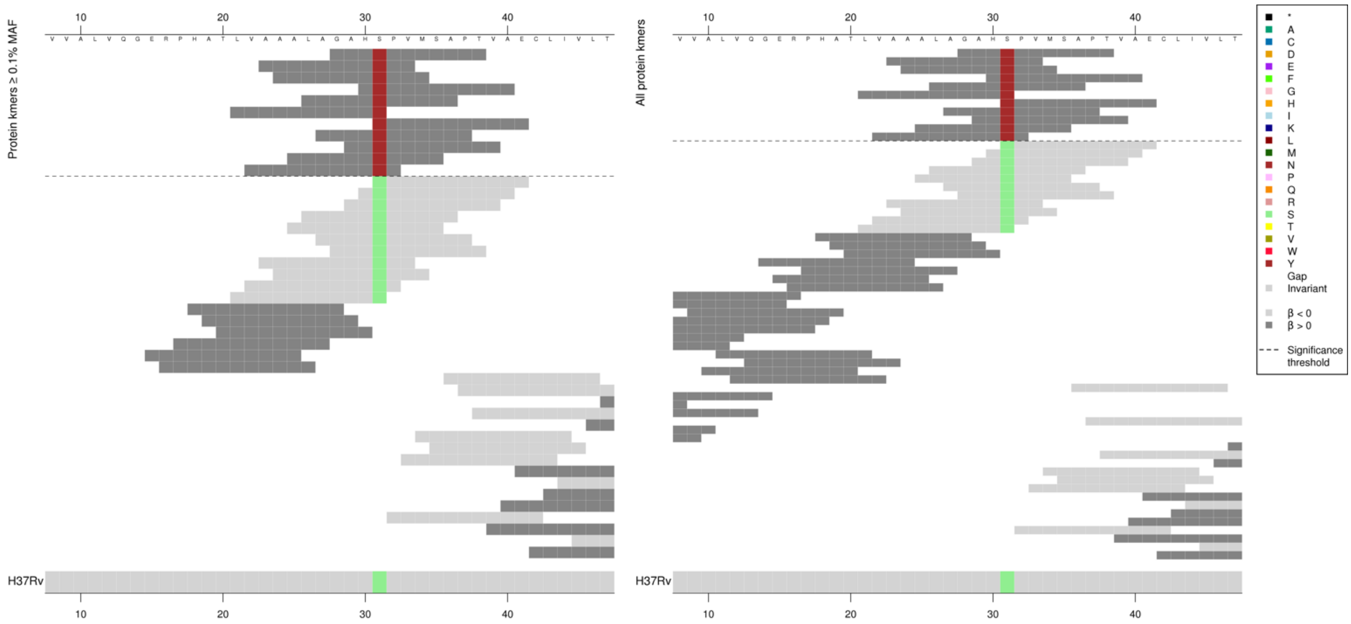

Kmers below significance threshold = 31S

| Kmer | -log$_{10}$ *p* | β | MAC | MAF | Ps | Variants |
|---|---|---|---|---|---|---|
| GAHNPVMSAPT | 9.55 | 2.84 | 8 | 0.11 | 28 | 31N |

#### *mce2F*

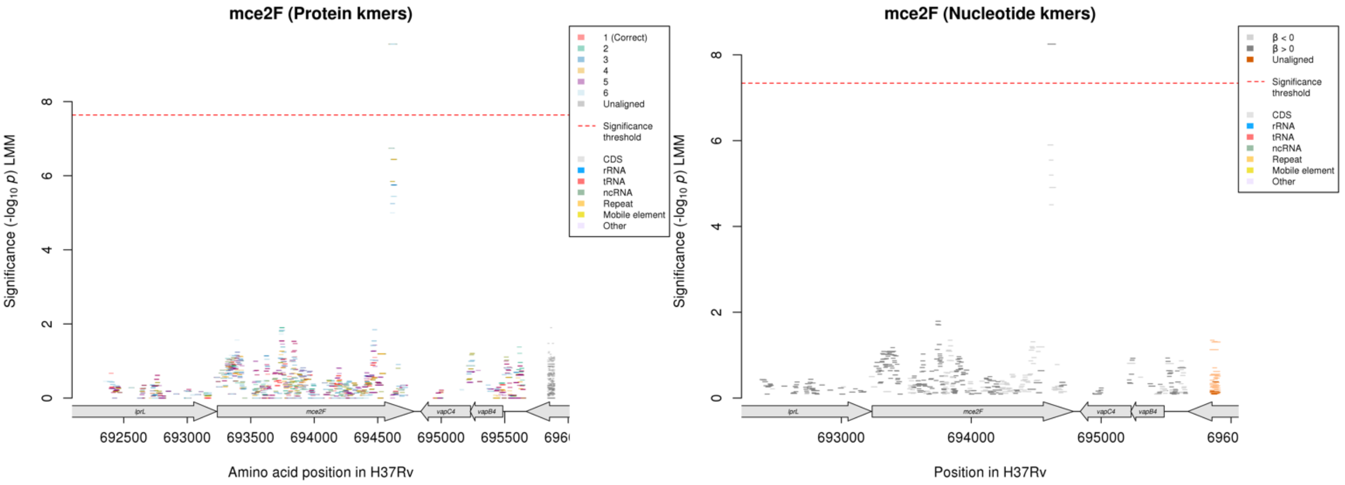

Protein kmers:

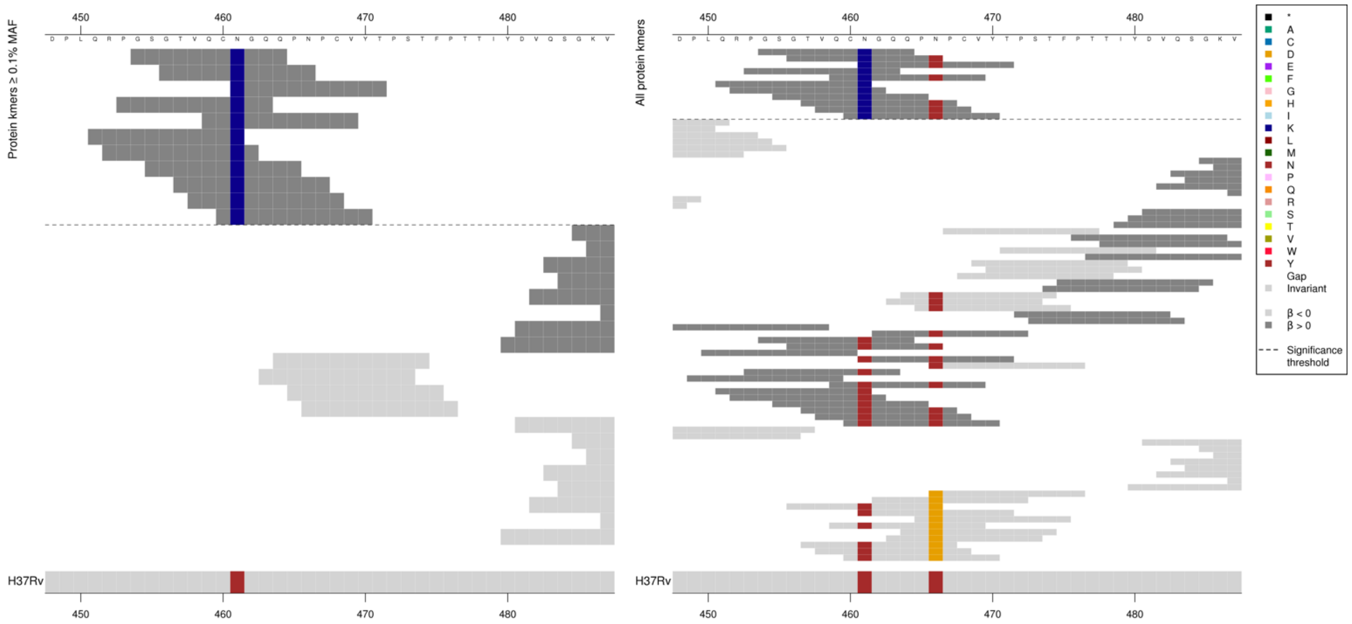

No kmers associated with low MIC in this region, although the estimate of the effect for the kmers below the significance threshold is not significant.
Kmers below significance threshold = 461N

| Kmer | $-\log_{10} p$ | β | MAC | MAF | Ps | Variants |
|---|---|---|---|---|---|---|
| GSGTVQCKGQQ | 9.550761 | 2.84 | 8 | 0.11 | 454 | 461K |

#### *fabG1*

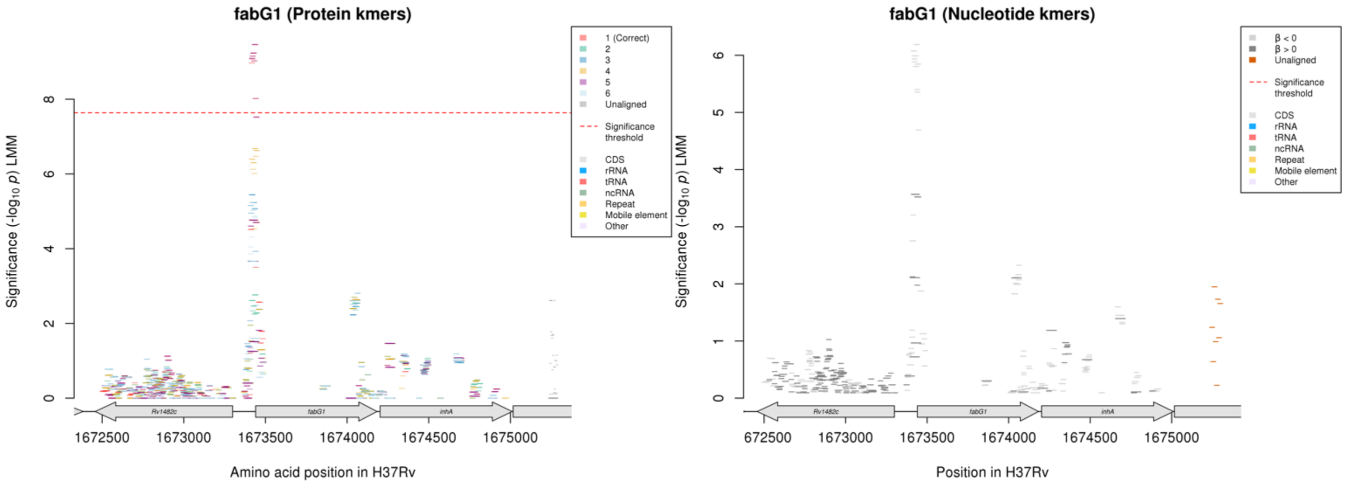

Not significant in nucleotide kmer analysis.
Protein kmer analysis is capturing a variant before the start codon.

Protein kmers:

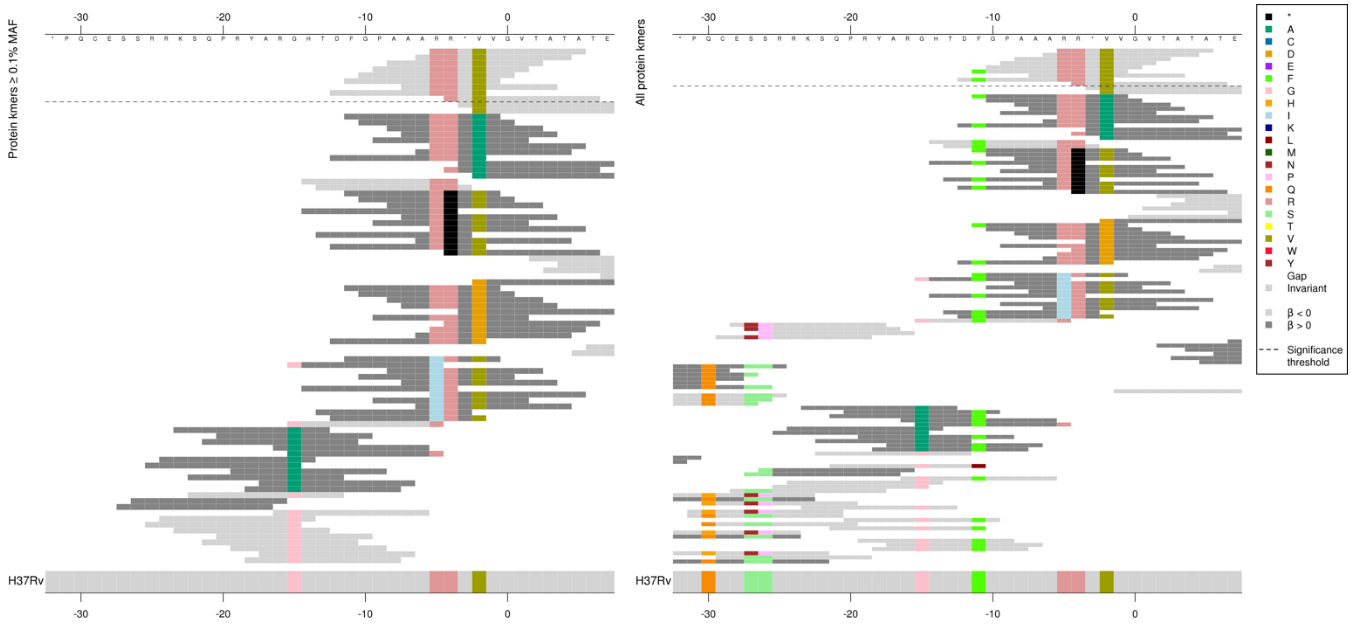

The kmers in the same region in the nucleotide kmer analysis:

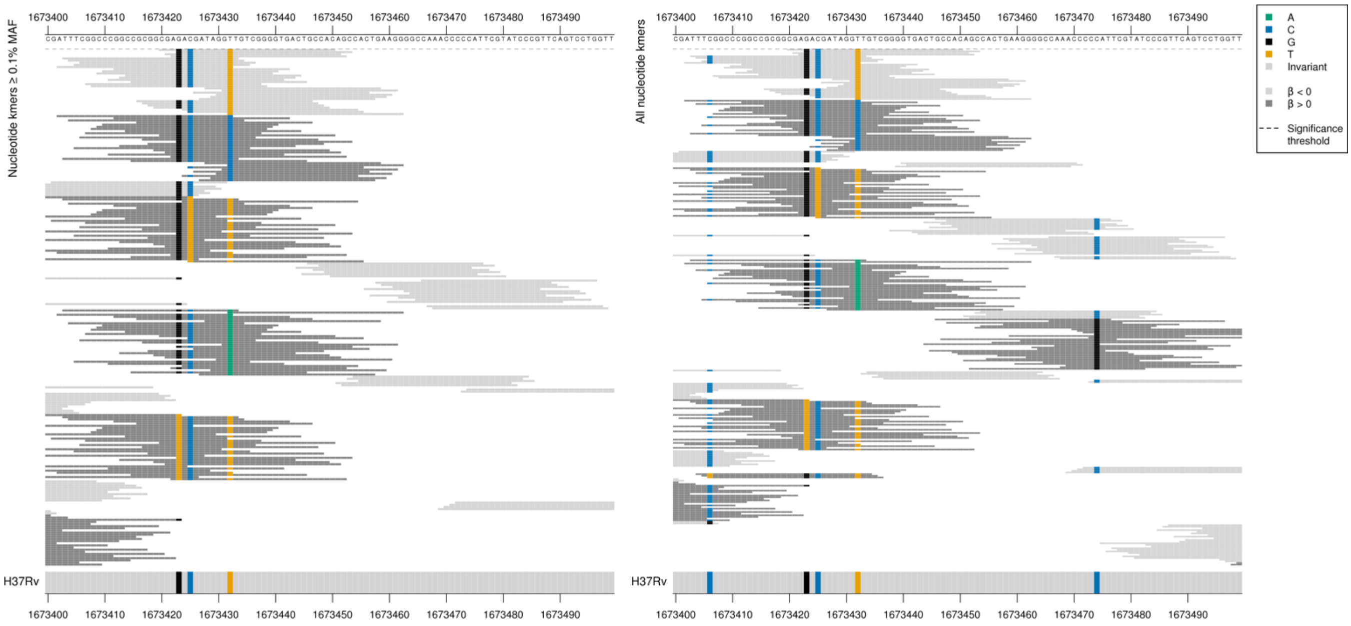

The top kmer in the protein kmer analysis: $-\log_{10} p$ = 9.46, beta = -0.45, MAC = 849, MAF = 11.63.

| Kmer | $-\log_{10} p$ | β | MAC | MAF | Ps | Variants |
|---|---|---|---|---|---|---|
| CGAGACGATAGGTTGTCGGGGTGACTGCCAC | 6.19 | -0.38 | 849 | 11.63 | 1673420 | WT |

### *katG*

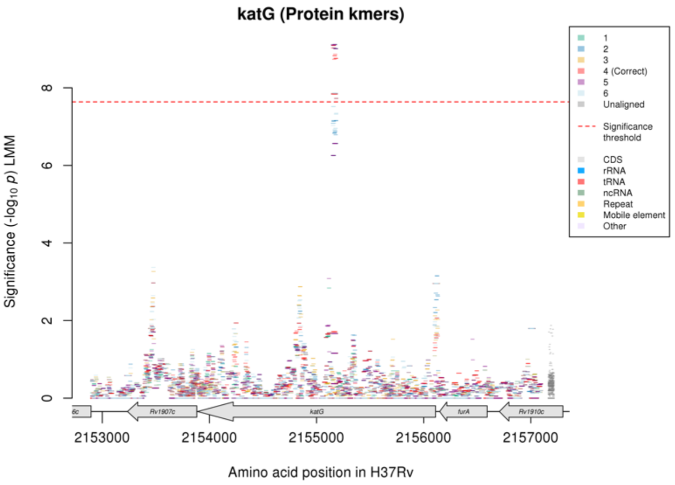

Not significant in the nucleotide kmer analysis.

Protein kmers:

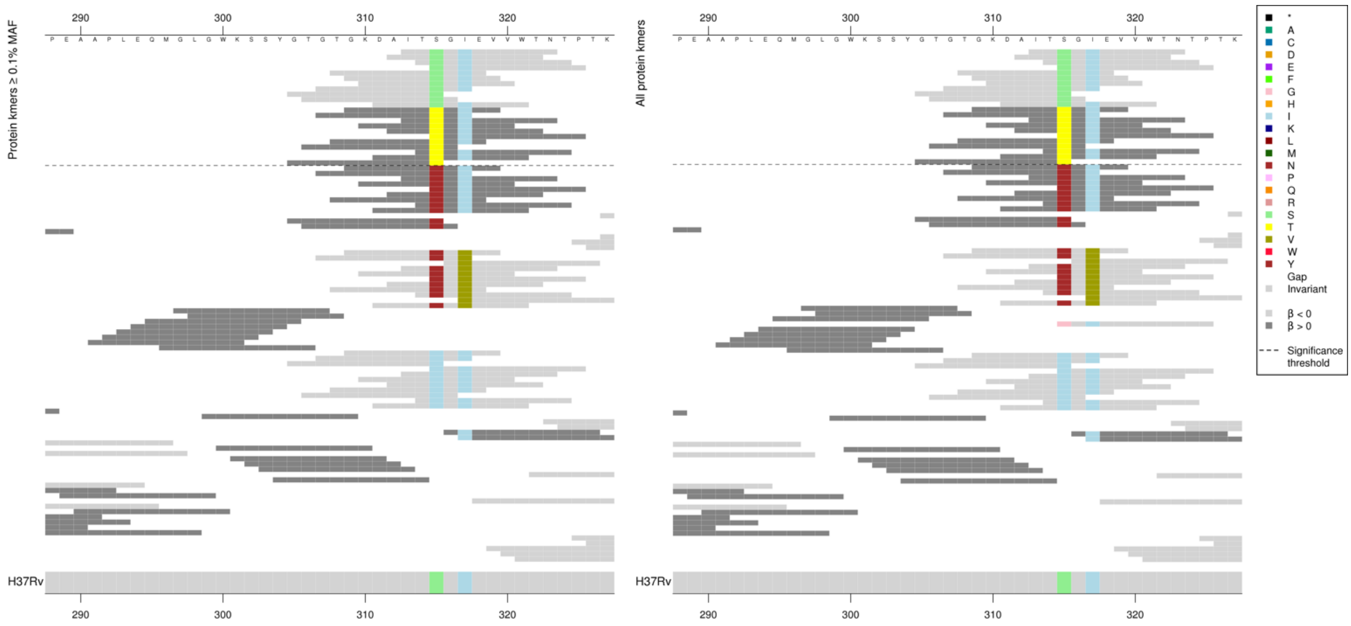

Significant low MIC-associated kmer *p*-values varied, but not in line with whether they covered codon 317. All high MIC-associated 315T kmers were the same *p*-value apart from the last kmer which was slightly less significant.

Kmers below significance threshold = 315N, 315I, 317V

| Kmer | $-\log_{10} p$ | β | MAC | MAF | Ps | Variants |
|---|---|---|---|---|---|---|
| ITSGIEVVWTN | 9.09 | -0.41 | 2715 | 37.19 | 313 | 315S |
| GKDAITTGIEV | 7.84 | 0.39 | 2638 | 36.14 | 309 | 315T |

### *folC*

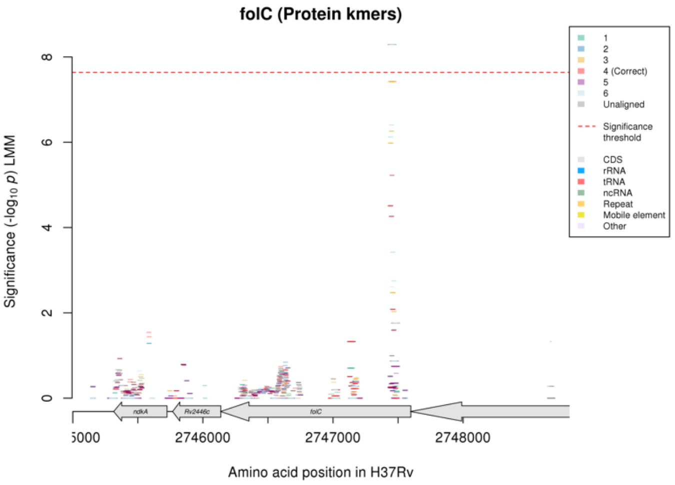

Not significant in nucleotide kmer analysis.
Protein kmers:

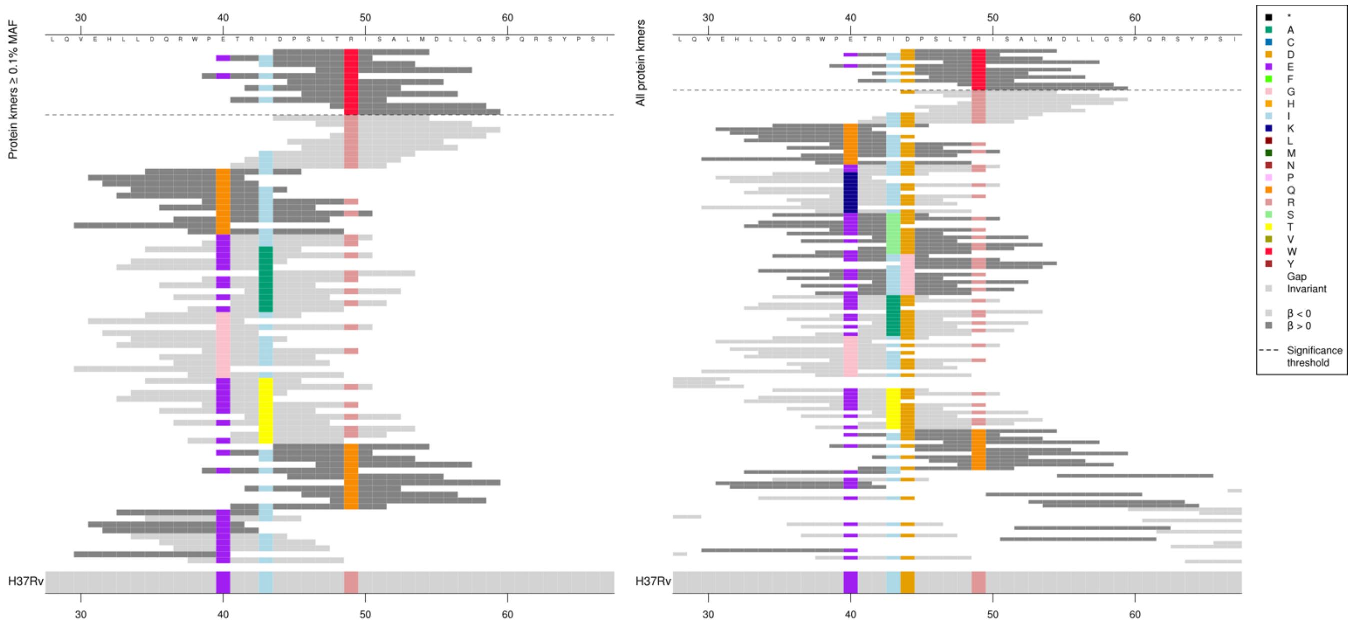

| Kmer | $-\log_{10} p$ | β | MAC | MAF | Ps | Variants |
|---|---|---|---|---|---|---|
| DPSLTWISALM | 8.29 | 2.02 | 14 | 0.19 | 44 | 49W |

### *tlyA*

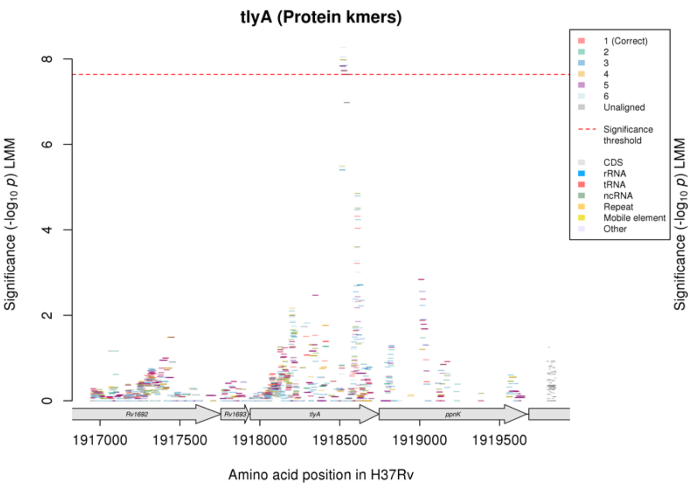

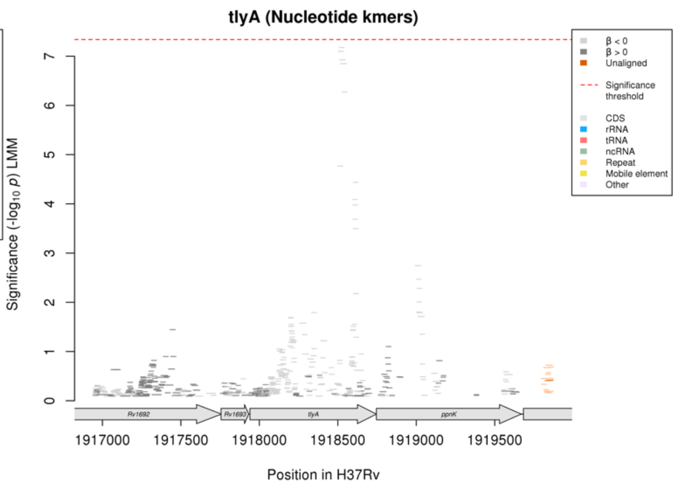

Protein kmers:

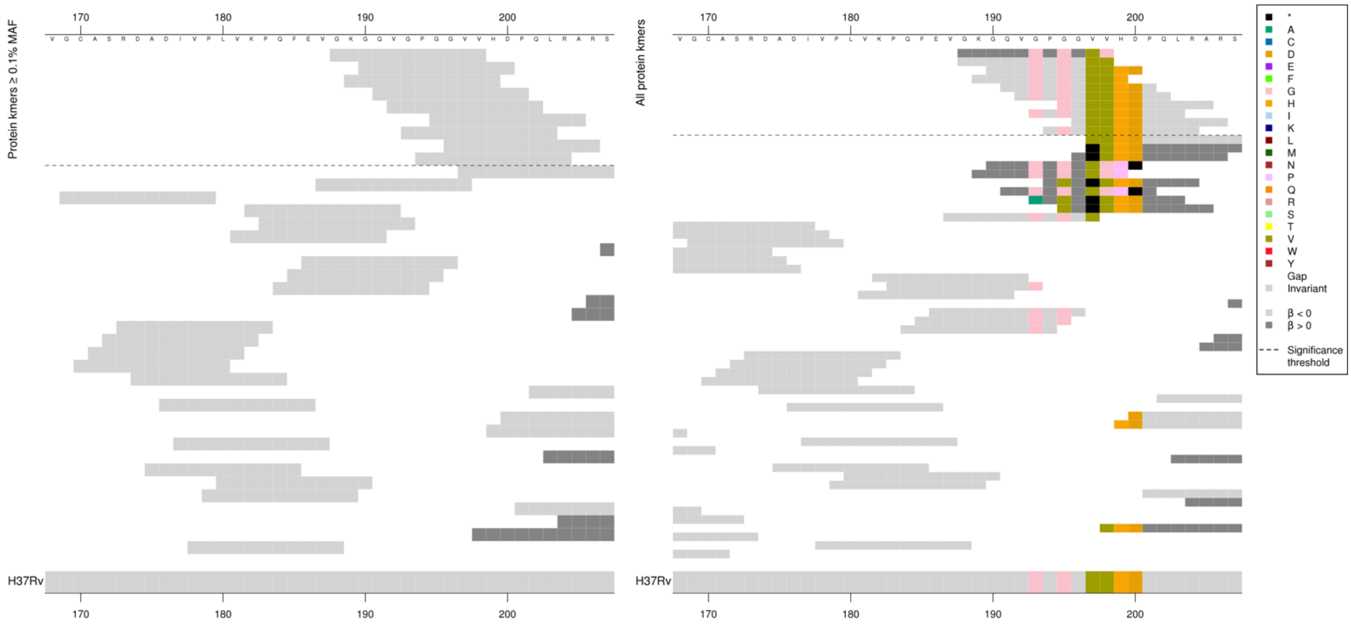

The same region for the nucleotide kmer analysis:

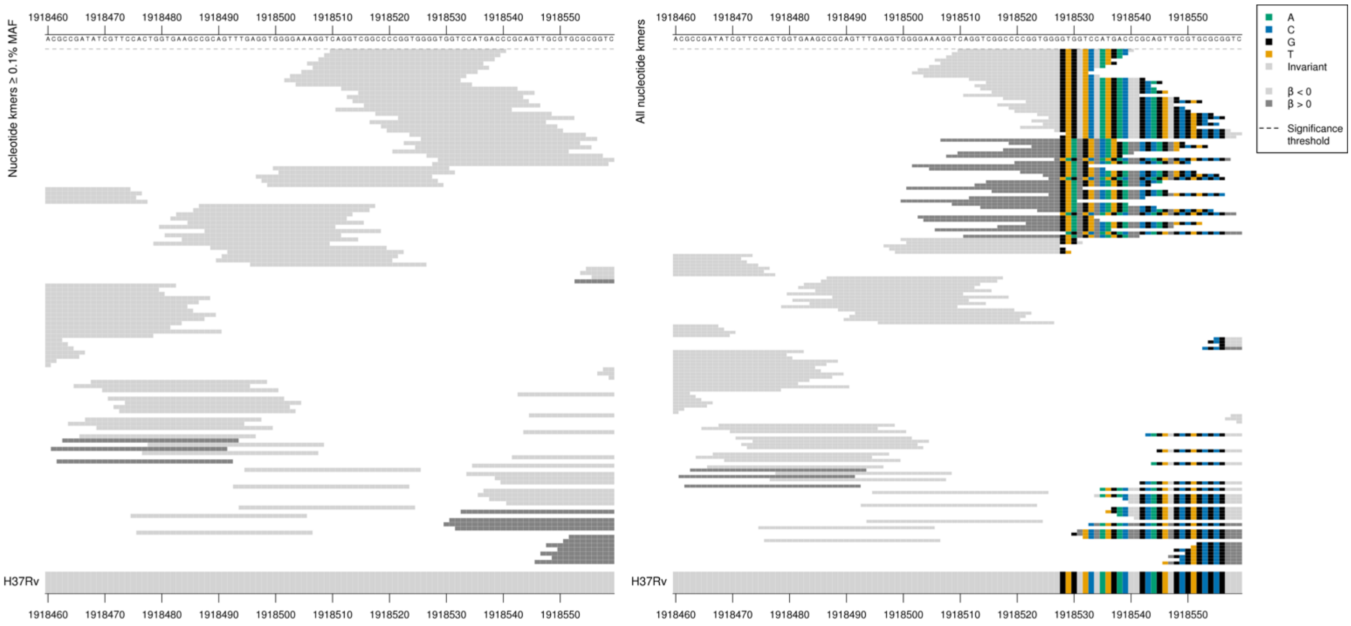

Low MAF kmers = Insertion of an A between positions 1918529-1918530/590-591 in codon 198.
Codon 198 stays as V, as WT = GTG = Val / V, with the insertion GTA = Val / V
The insertion changes from codons 199 onwards, so codon 199 changes from GTC = Val / V to GGT = Gly / G, as seen in the protein kmer alignment.

Significant kmers cover codons 188-202 (nucleotide bases 562-606).

| Kmer | $-\log_{10} p$ | β | MAC | MAF | Ps | Variants |
|---|---|---|---|---|---|---|
| GKGQVGPGGVV | 7.83 | -1.86 | 19 | 0.26 | 188 | WT codons 188-206 |

### *ethA*

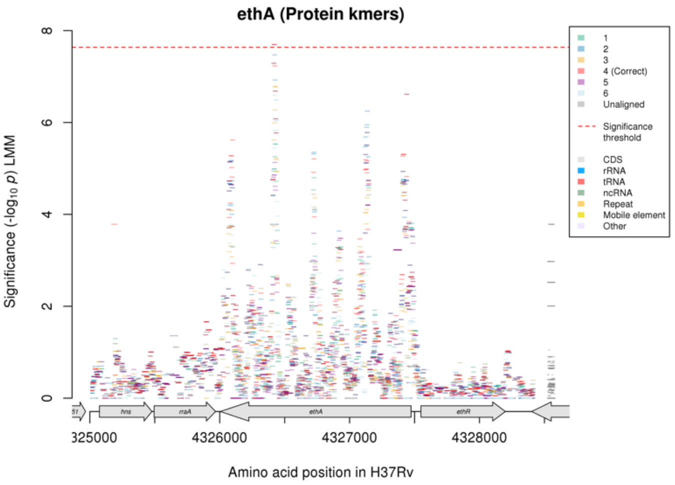

Not significant in nucleotide kmer analysis.
Protein kmers:

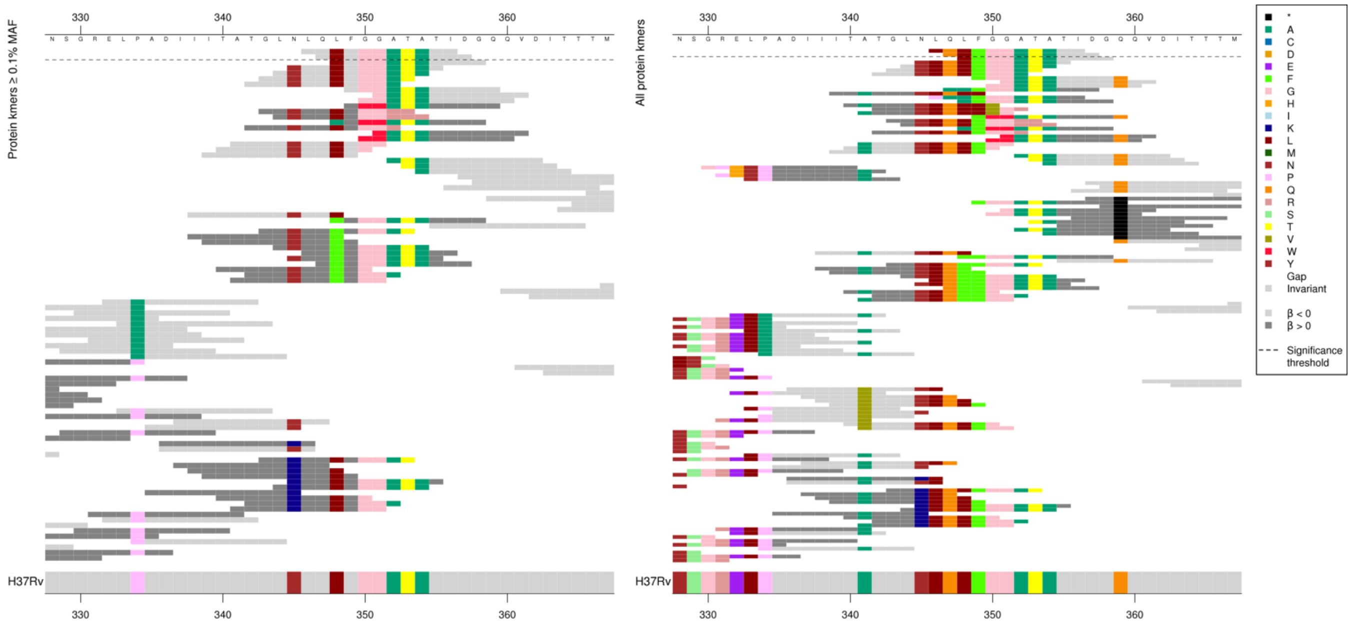

Kmers below significance threshold = 348F, 345K, plus others.

| Kmer | $-\log_{10} p$ | β | MAC | MAF | Ps | Variants |
|---|---|---|---|---|---|---|
| LQLFGGATATI | 7.70 | -1.25 | 50 | 0.68 | 346 | WT codons 346-357 |

# *Rv0228*

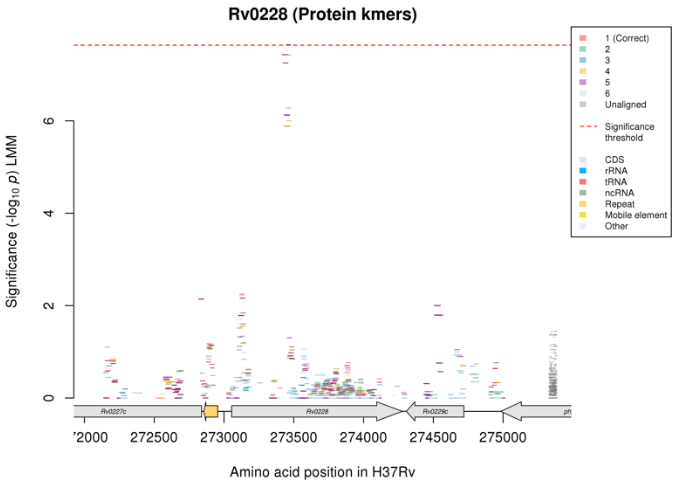

Not significant in nucleotide kmer analysis.
Protein kmers:

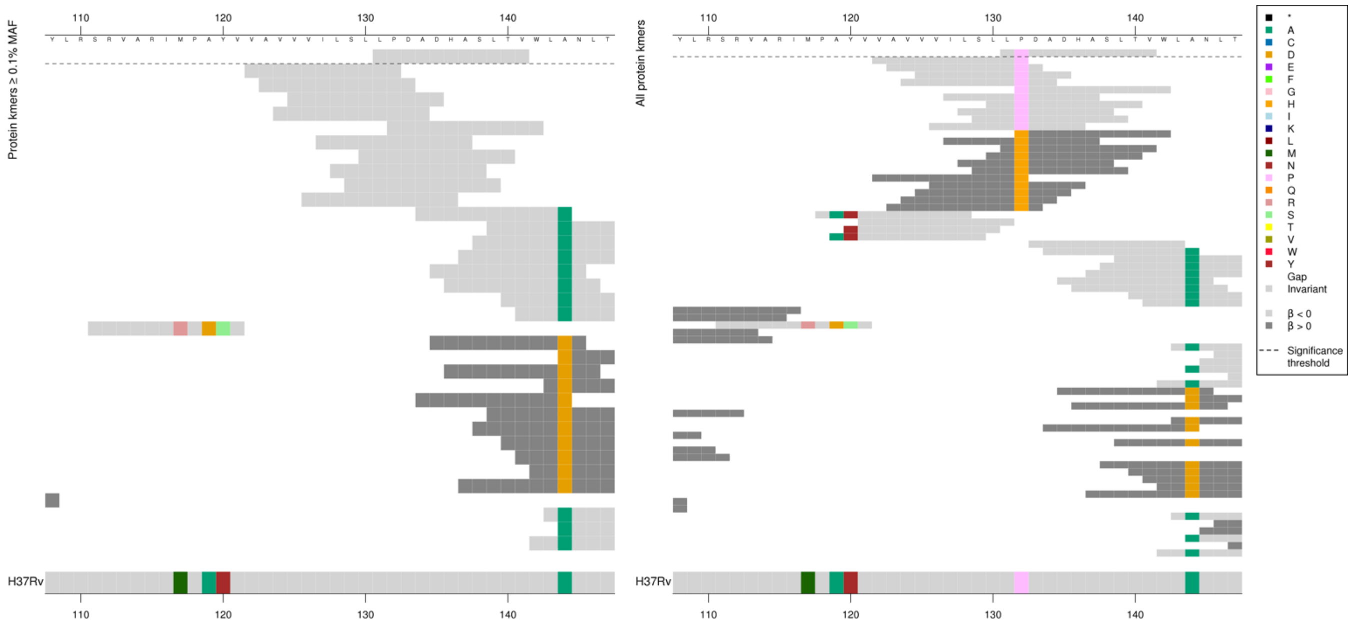

| Kmer | $-\log_{10} p$ | β | MAC | MAF | Ps | Variants |
|---|---|---|---|---|---|---|
| LPDADHASLTV | 7.65 | -2.52 | 12 | 0.16 | 131 | 132P |

### Linezolid (LZD)

#### *rplC*

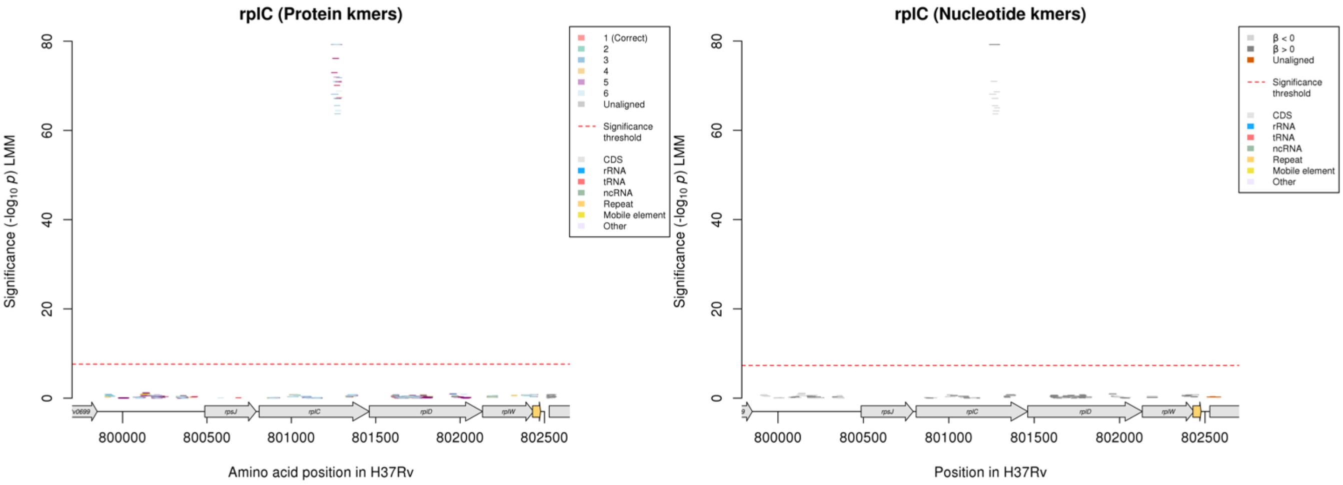

Protein kmers:

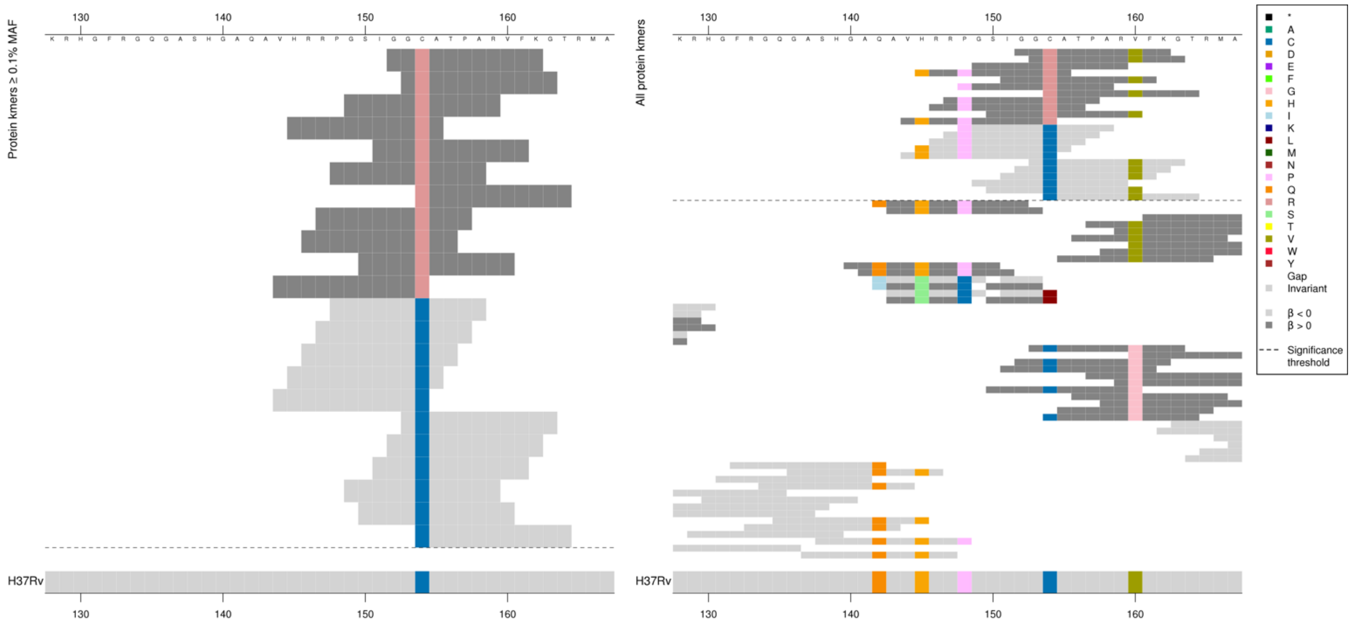

All high MIC-associated kmers were the same *p*-value and only captured 154R, the low MIC-associated kmers were different *p*-values depending on the other substitutions in the region.

| Kmer | $-\log_{10} p$ | β | MAC | MAF | Ps | Variants |
|---|---|---|---|---|---|---|
| GGRATPARVFK | 79.24 | 3.15 | 26 | 0.39 | 152 | 154R |
| PGSIGGCATPA | 76.14 | -3.14 | 25 | 0.37 | 148 | 148P, 154C |
| HRRPGSIGGCA | 72.95 | -3.04 | 26 | 0.39 | 145 | 145H, 148P, 154C |
| GCATPARVFKG | 70.92 | -2.96 | 27 | 0.40 | 153 | 154C, 160V |
| GSIGGCATPAR | 70.10 | -2.93 | 27 | 0.40 | 149 | 154C |

### *rpoB*

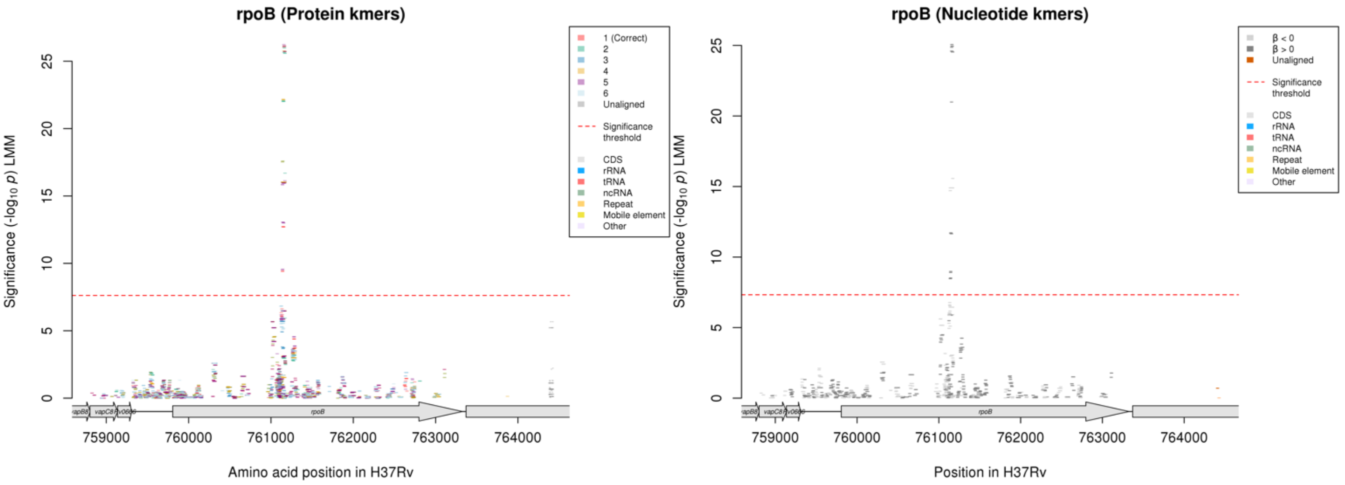

Protein kmers:

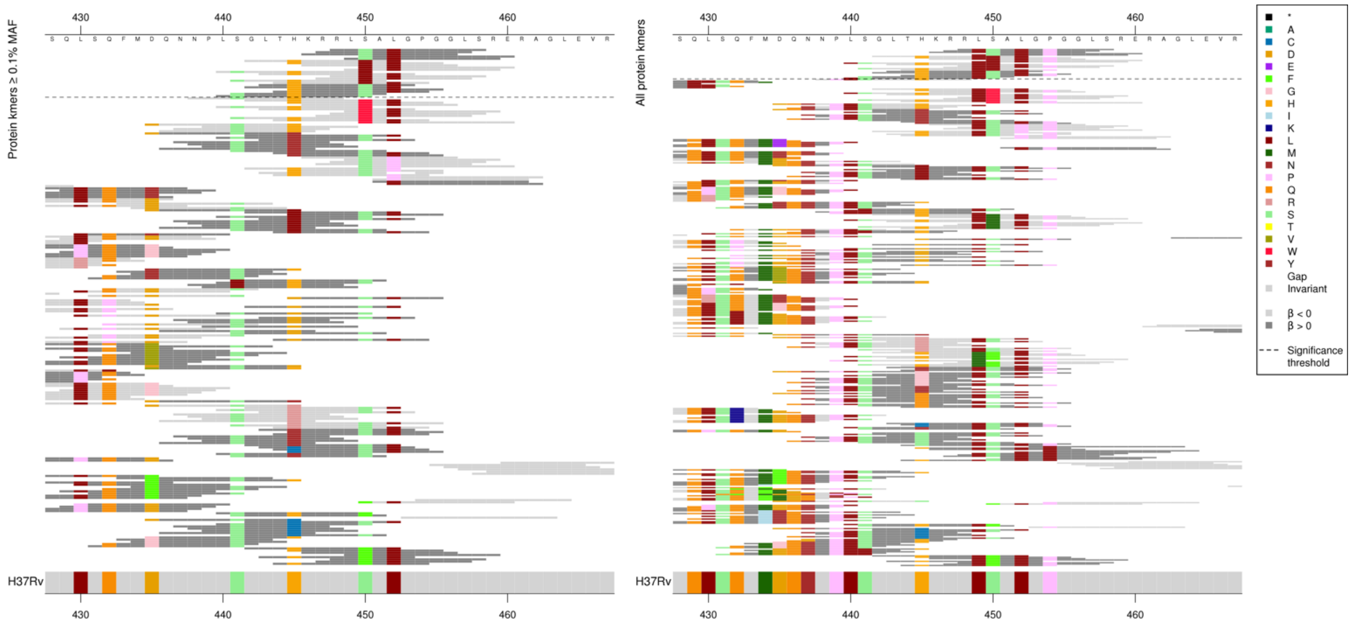

The low MIC-associated kmers were all the same *p*-value, capturing 450L. The high MIC-associated kmers had different *p*-values depending on the other substututions.
The associations were the opposite direction to their effect on RIF MIC.

| Kmer | $-\log_{10} p$ | β | MAC | MAF | Ps | Variants |
|---|---|---|---|---|---|---|
| RLSALGPGGLS | 26.13 | 0.35 | 1828 | 27.15 | 448 | 449L, 450S, 452L, 454P |
| GLTHKRRLLAL | 16.02 | -0.28 | 1686 | 25.04 | 442 | 450L |
| LTHKRRLSALG | 12.73 | 0.23 | 2072 | 30.78 | 443 | 445H, 449L, 450S, 452L |
| HKRRLSALGPG | 12.71 | 0.23 | 2076 | 30.84 | 445 | 445H, 449L, 450S, 452L, 454P |
| SGLTHKRRLSA | 9.44 | 0.20 | 2003 | 29.75 | 441 | 441S, 445H, 449L, 450S |
| LSGLTHKRRLS | 9.39 | 0.20 | 2001 | 29.72 | 440 | 440L, 441S, 445H, 449L, 450S |

#### *emrB*

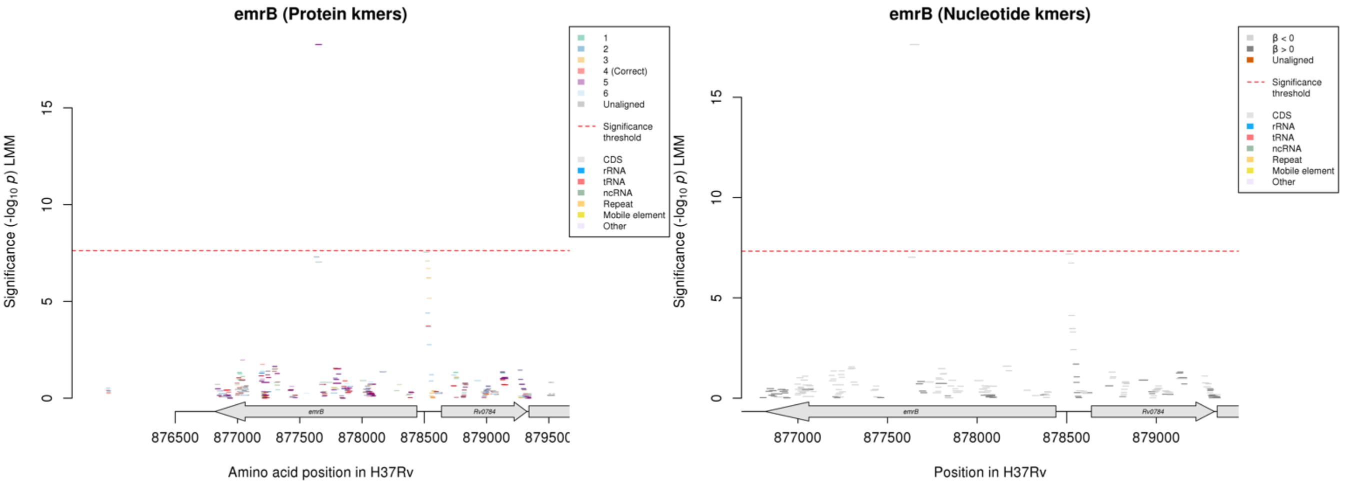

Nucleotide kmers:

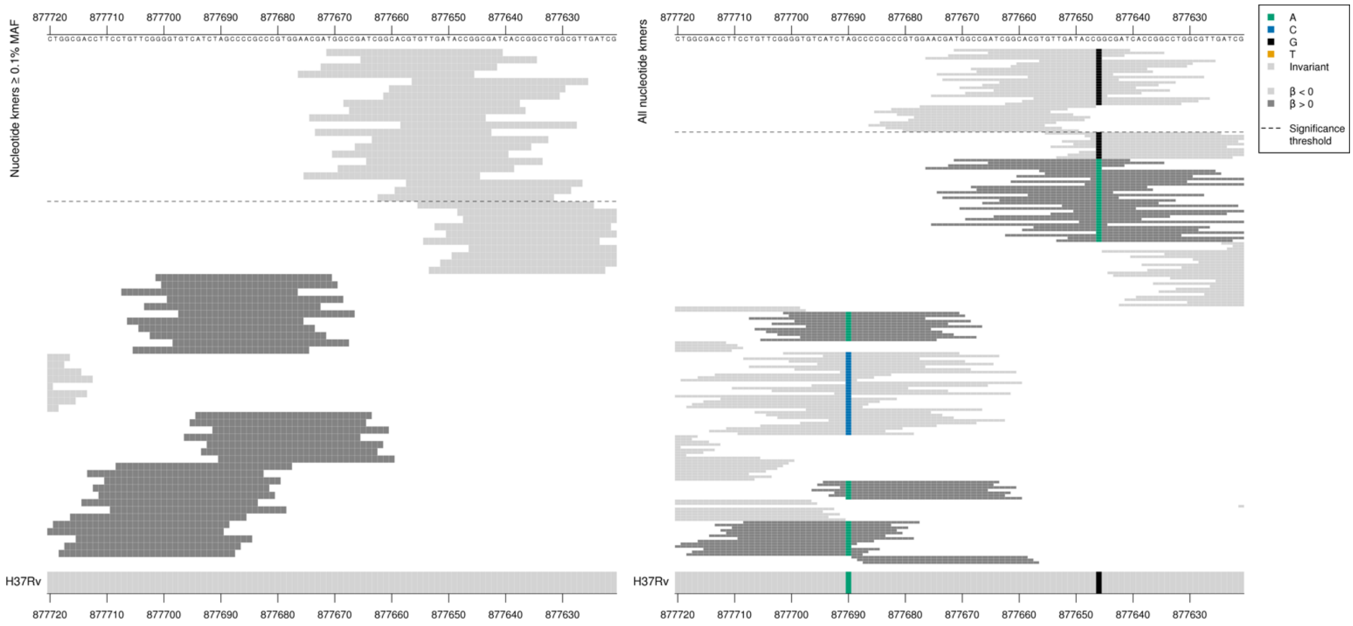

Synonymous mutation at position 877646/795

Kmers below significance threshold = 877646/795A

CCG = Pro / P

CCA = Pro / P

Also some low MAF WT kmers adjacent, they could be in LD.

| Kmer | $-\log_{10} p$ | β | MAC | MAF | Ps | Variants |
|---|---|---|---|---|---|---|
| TGGCCGATCGGCACGTGTTGATACCGGCGAT | 17.63 | -2.59 | 11 | 0.16 | 877671 | 877646/795G |

# *Rv3552*

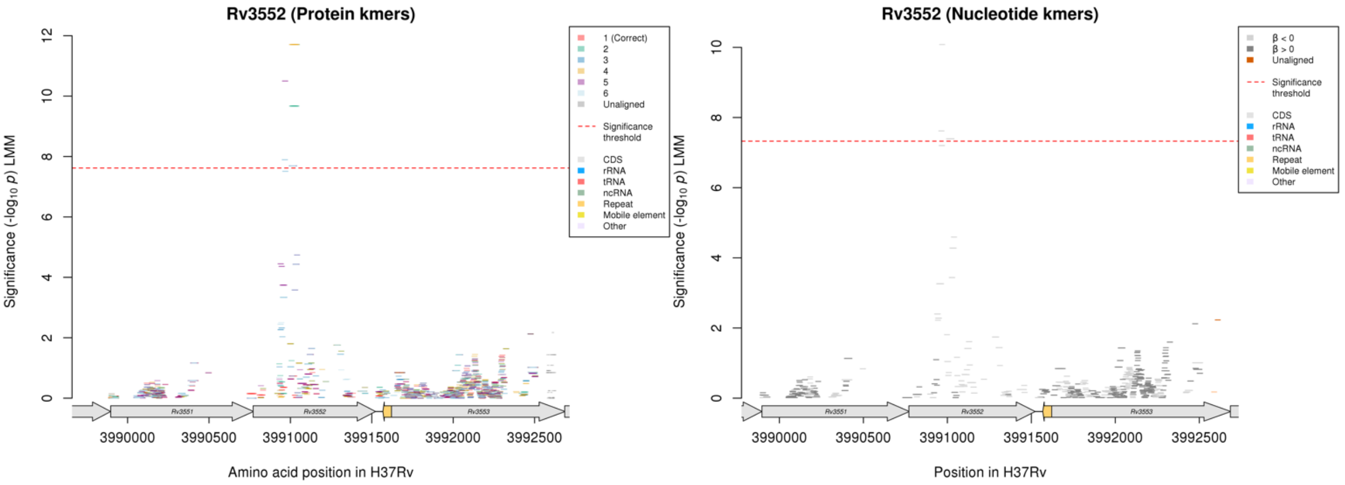

No significant kmers aligned to the correct reading frame in the protein kmer analysis.

Nucleotide kmers:

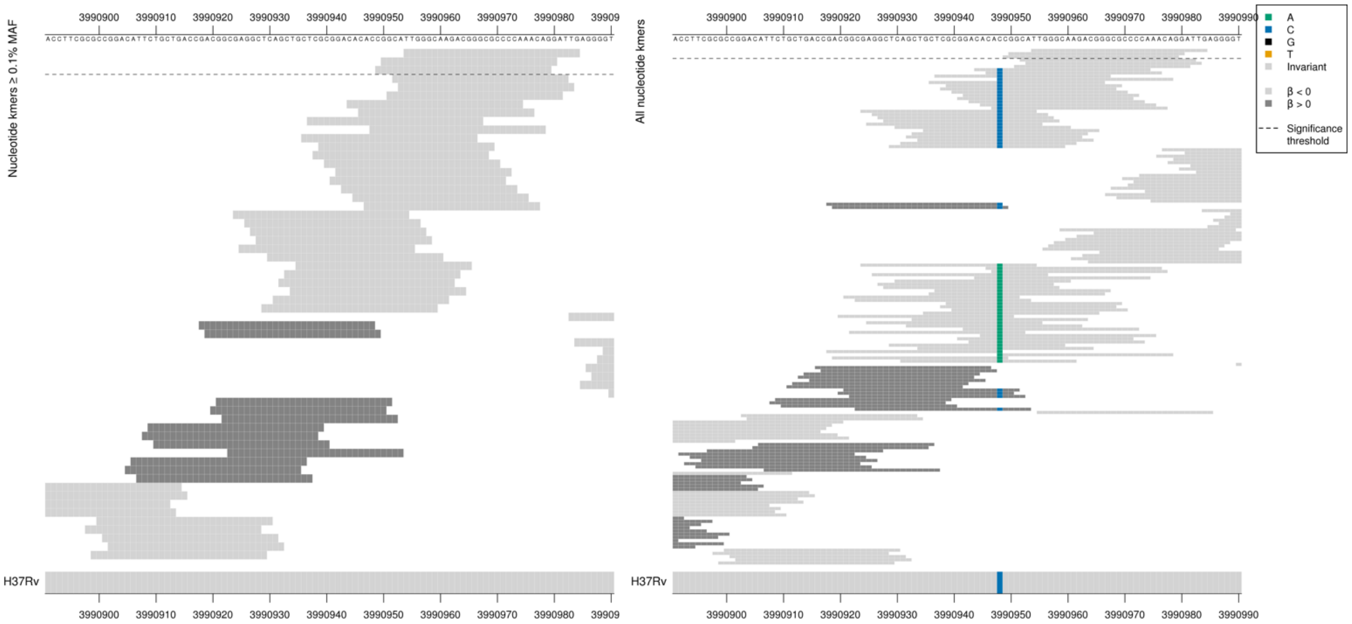

Removing the threshold requiring a kmer to be aligned to a gene/intergenic region in at least five genomes across the full dataset. Showing the nucleotide kmer alignment after lowering the threshold to just one genome:

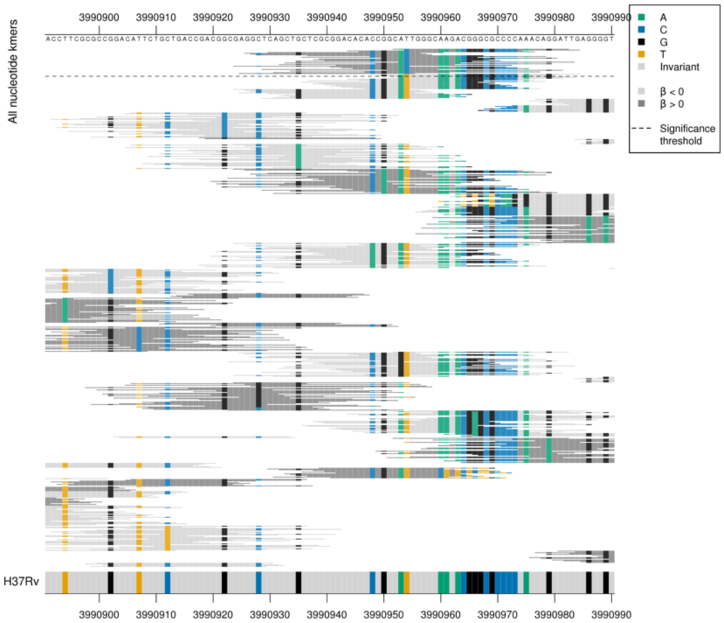

Alternative alleles were present in the region but were present in fewer than five genomes.
The top kmer had a MAC of only 7, so also showing the second kmer which had a slightly higher MAC of 9.

| Kmer | $-\log_{10} p$ | β | MAC | MAF | Ps | Variants |
|---|---|---|---|---|---|---|
| TTGGGCAAGACGGGCGCCCCAAACAGGATTG | 10.08 | -2.66 | 7 | 0.10 | 3990954 | WT |
| GGCATTGGGCAAGACGGGCGCCCCAAACAGG | 7.62 | -1.91 | 9 | 0.13 | 3990950 | WT |

Nucleotide kmers

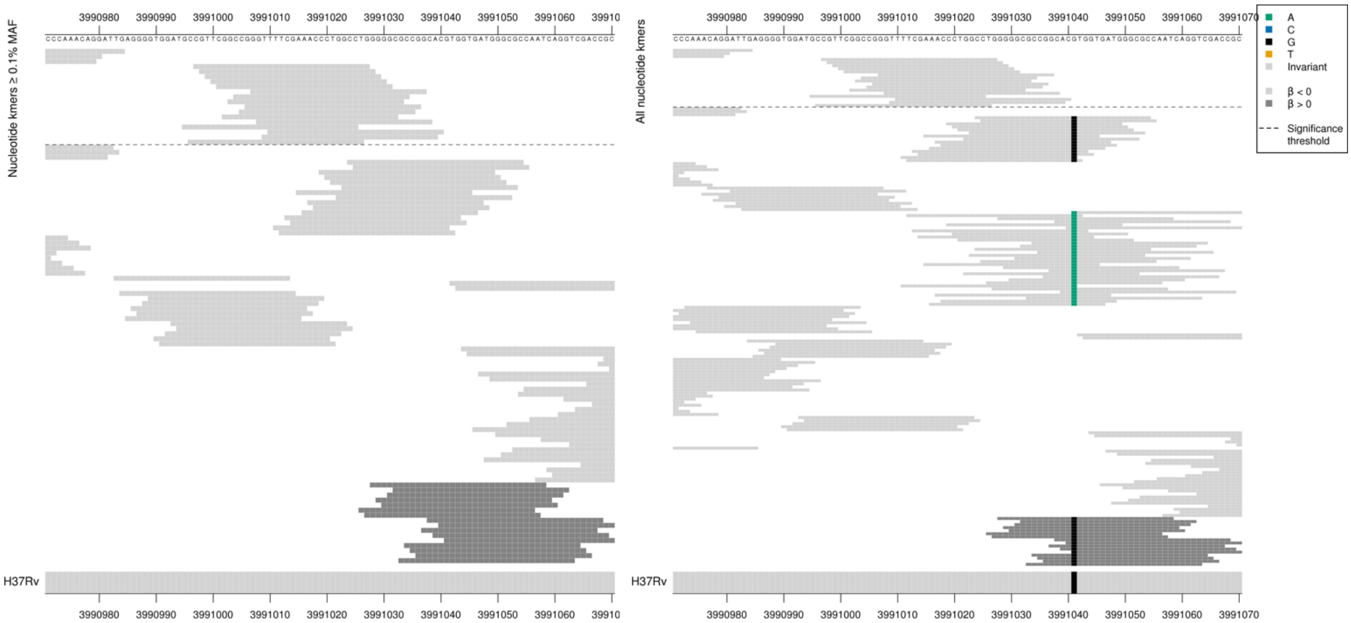

Removing the threshold requiring a kmer to be aligned to a gene/intergenic region in at least five genomes across the full dataset. Showing the nucleotide kmer alignment after lowering the threshold to just one genome:

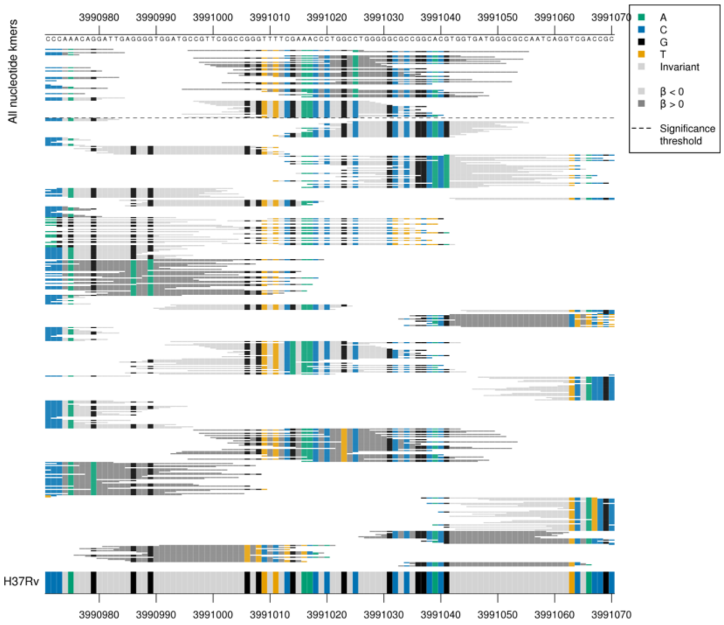

Again rare variation present in fewer than five genomes.

| Kmer | $-\log_{10} p$ | β | MAC | MAF | Ps | Variants |
|---|---|---|---|---|---|---|
| CGTTCGGCCGGGTTTTCGAAACCCTGGCCTG | 7.39 | -1.64 | 12 | 0.18 | 3990997 | WT |

#### *add*

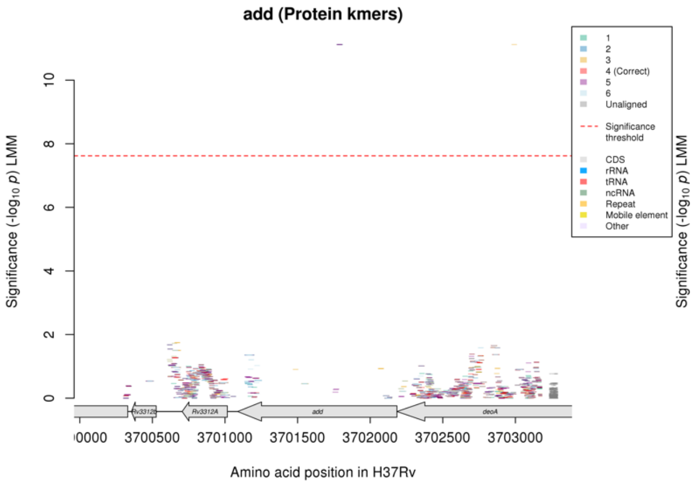

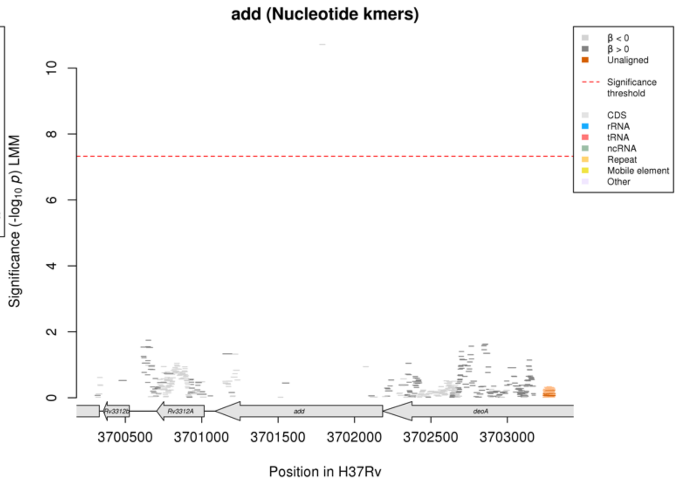

No significant kmers aligned to the correct reading frame in the protein kmer analysis. The only significant kmers in this region in the protein kmer analysis were below 0.1% MAF. Not many kmers aligned to the gene.

Protein kmers:

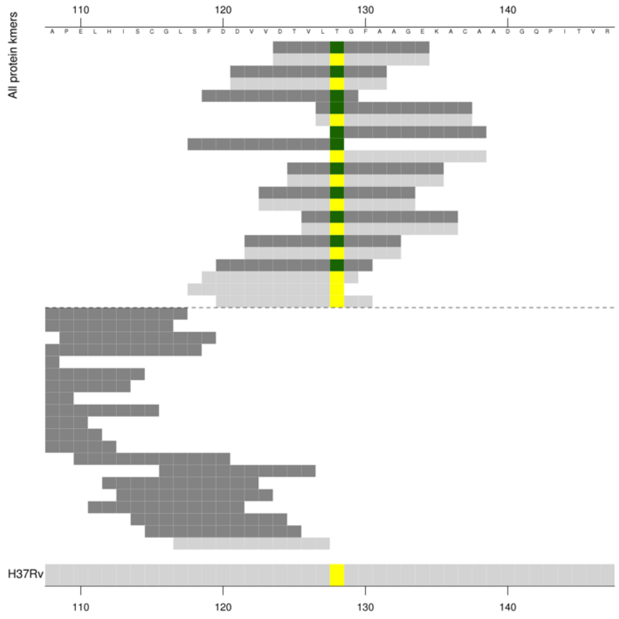

Nucleotide kmers in the same region:

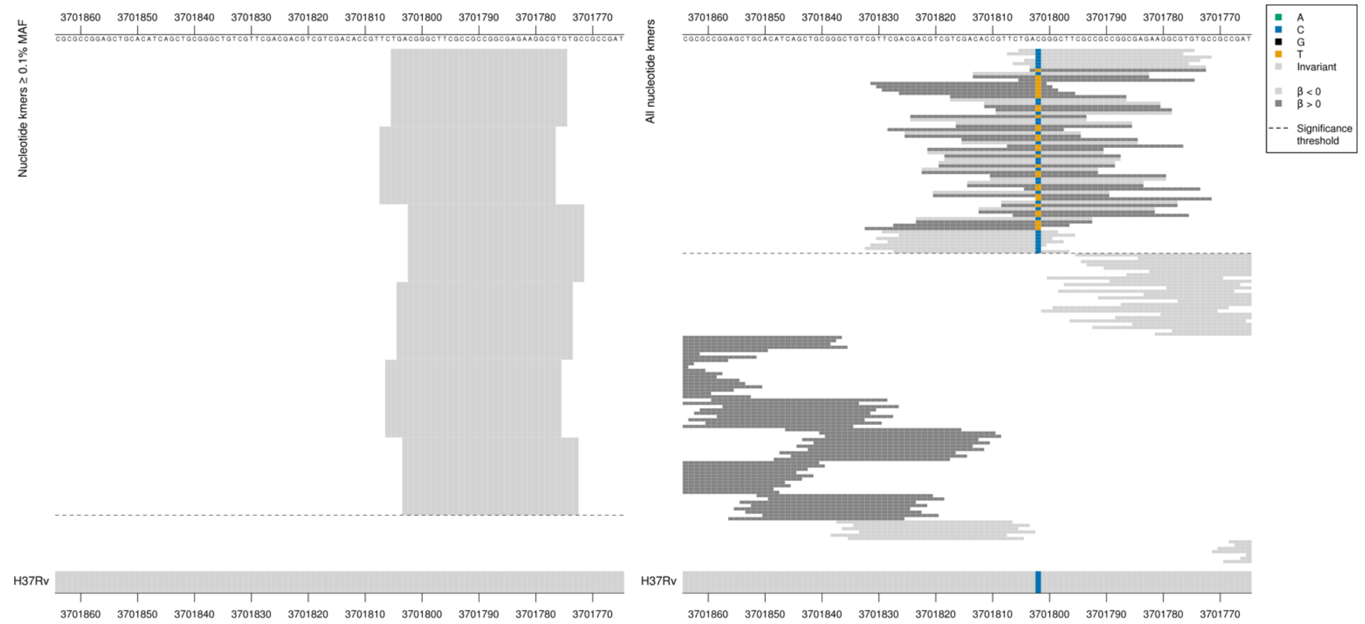

Non-synonymous mutation at position 3701802/128.
Low MIC-associated kmers = ACG = Thr / T
High MIC-associated kmers = ATG = Met / M

Showing the kmers for the nucleotide kmer analysis as some were above the 0.1% MAF threshold. However the top kmer for the protein kmer analysis: $-\log_{10} p$ = 10.85, beta = 2.86, MAC = 5, MAF = 0.07, ps = 124.

Low MAF kmers = 3701802/383T (128M aa)

| Kmer | $-\log_{10} p$ | β | MAC | MAF | Ps | Variants |
|---|---|---|---|---|---|---|
| TGACGGGCTTCGCCGCCGGCGAGAAGGCGTG | 10.71 | -2.40 | 7 | 0.10 | 3701805 | 3701802/383C (128T aa) |

#### *vapC33*

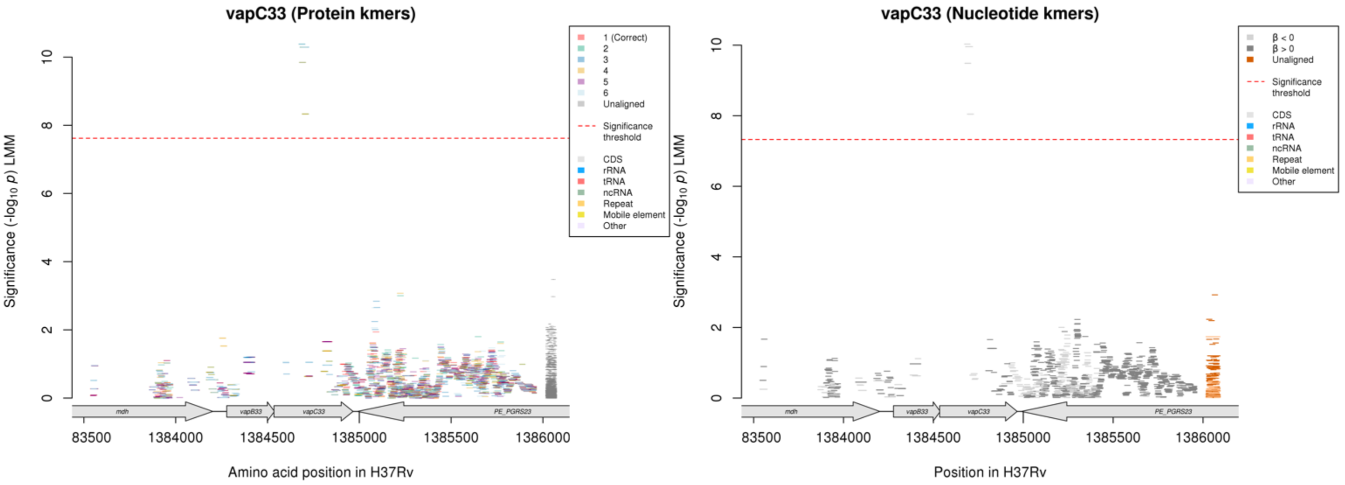

Nucleotide kmers:

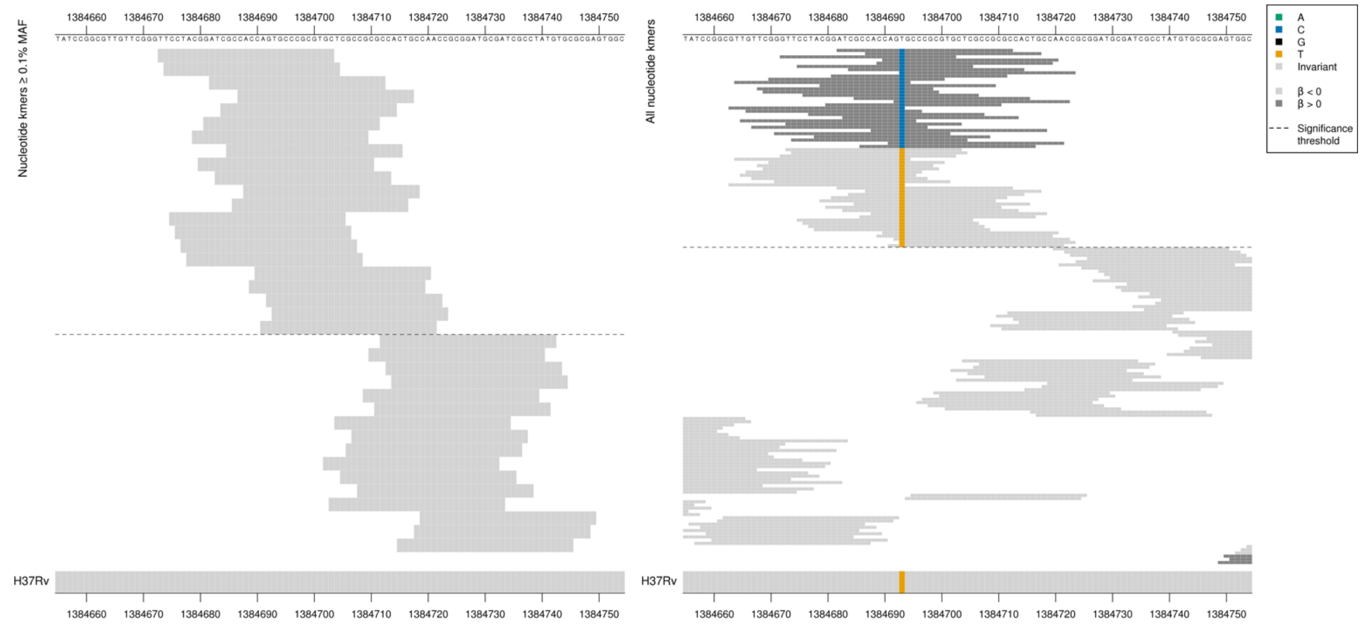

Synonymous mutation at position 1384693/159.
AGT = Ser / S
AGC = Ser / S

Low MAF kmers = 1384693/159C

| Kmer | $-\log_{10} p$ | β | MAC | MAF | Ps | Variants |
|---|---|---|---|---|---|---|
| TTCCTACGGATCGCCACCAGTGCCCGCGTGC | 10.03 | -2.34 | 7 | 0.10 | 1384673 | 1384693/159T |

#### *ppgK*

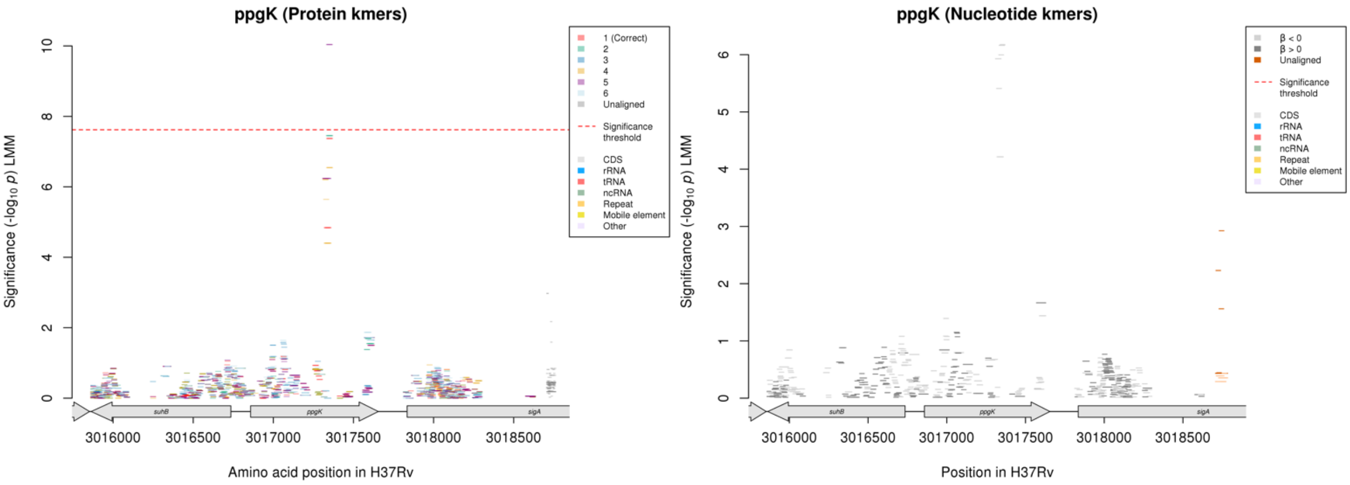

Not significant in the nucleotide analysis.

Protein kmers:

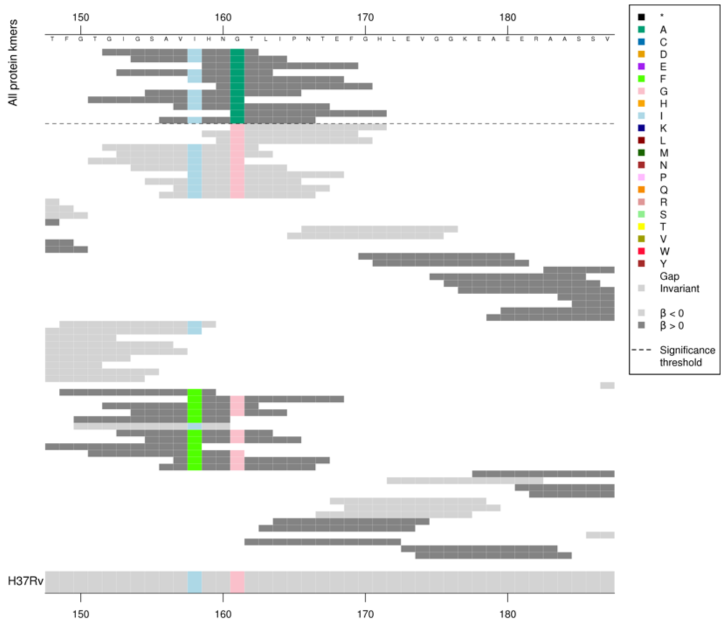

No significant protein kmers above the 0.1% MAF cutoff aligned to the correct reading frame. There were protein kmers above the MAF threshold that align to frame 5 (NSVLGINVPLW, $-\log_{10}p$ = 10.04, beta = -2.36, MAC = 7)

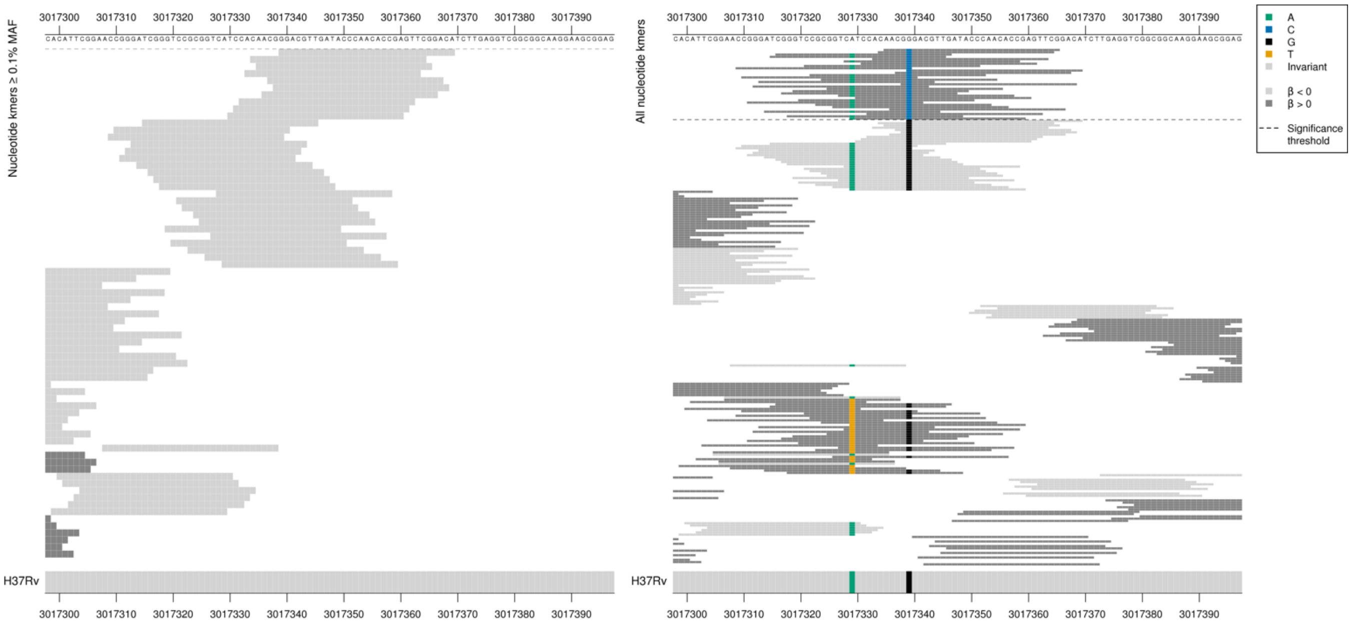

Kmers below significance threshold = 161G

| Kmer | $-\log_{10}p$ | β | MAC | MAF | Ps | Variants |
|---|---|---|---|---|---|---|
| GGACGTTGATACCCAACACCGAGTTCGGACA | 6.18 | -1.62 | 8 | 0.12 | 3017339 | WT |

### *pncB1:Rv1331*

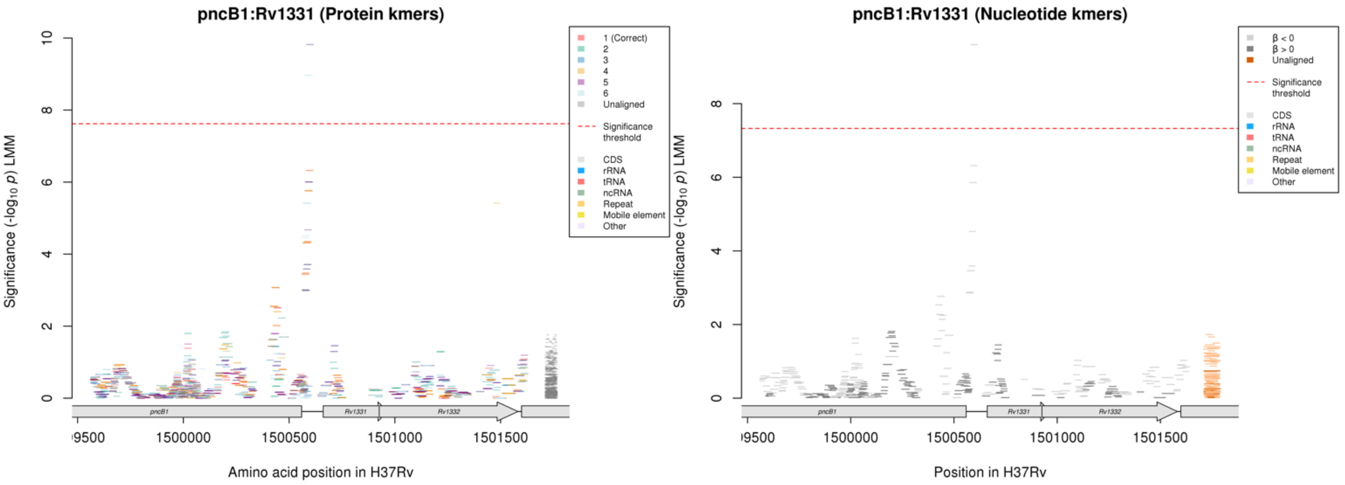

Nucleotide kmers:

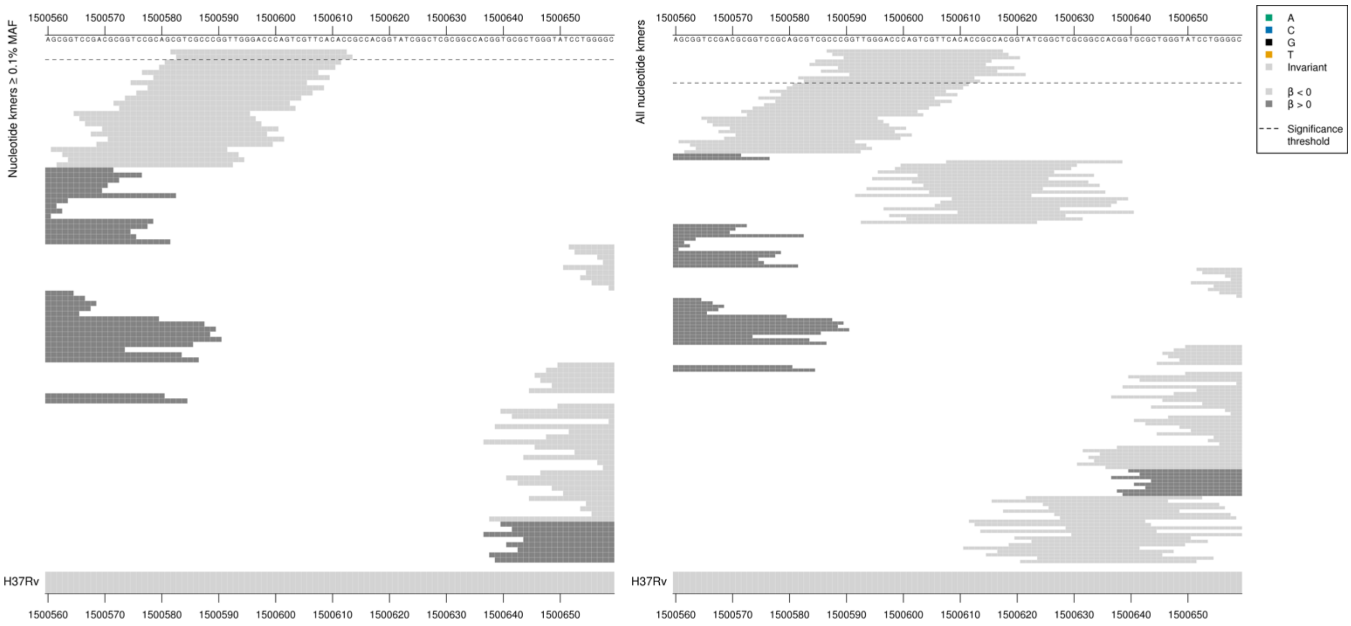

Removing the threshold requiring a kmer to be aligned to a gene/intergenic region in at least five genomes across the full dataset. Showing the nucleotide kmer alignment after lowering the threshold to just one genome:

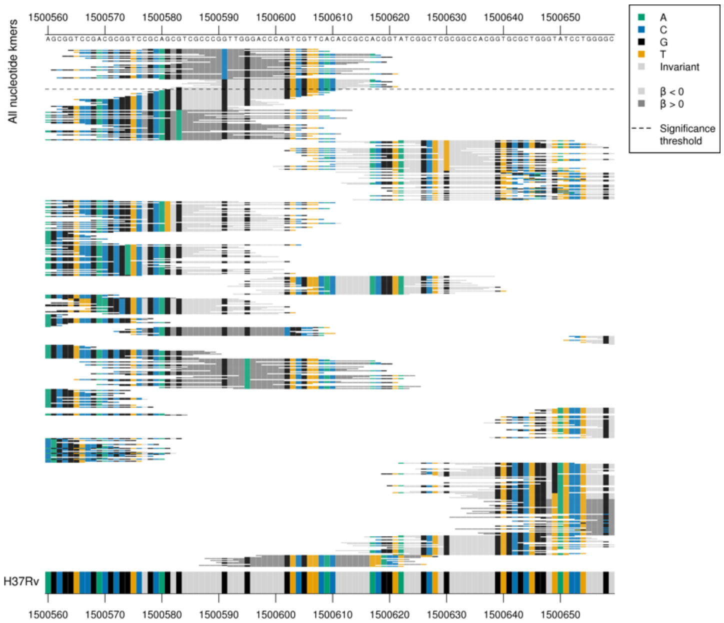

Rare variation present in the region, present in fewer than five genomes.

*pncB1* starts at position 1500559, *Rv1331* starts at position 1500661.

| Kmer | -log10 *p* | β | MAC | MAF | Ps | Variants |
|---|---|---|---|---|---|---|
| CGTCGCCCGGTTGGGACCCAGTCGTTCACAC | 9.60 | -2.11 | 7 | 0.10 | 1500582 | WT |

### *lprA*

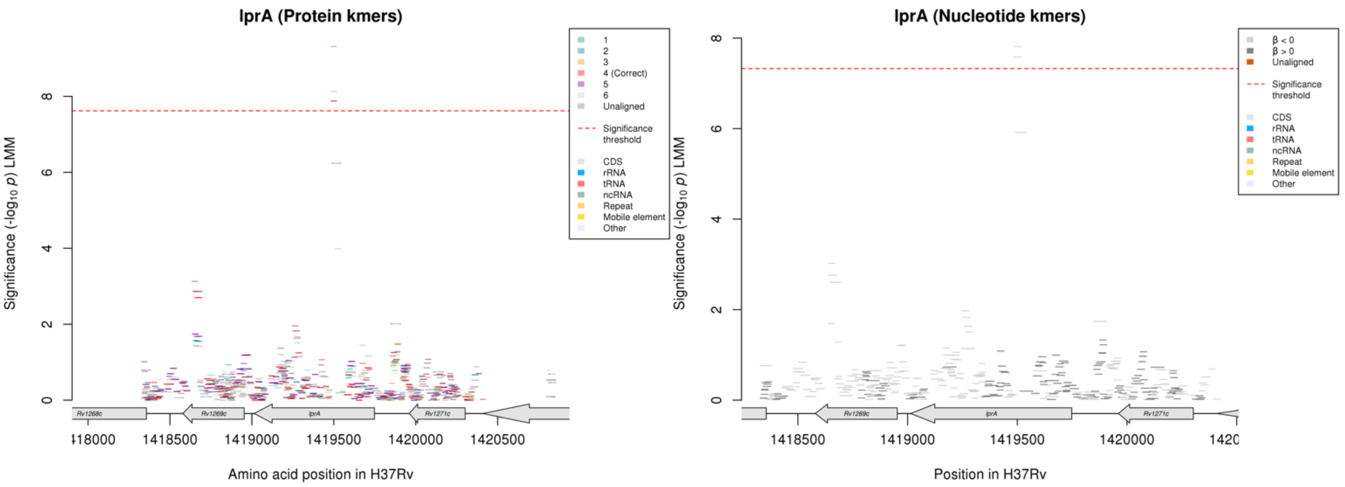

Protein kmers:

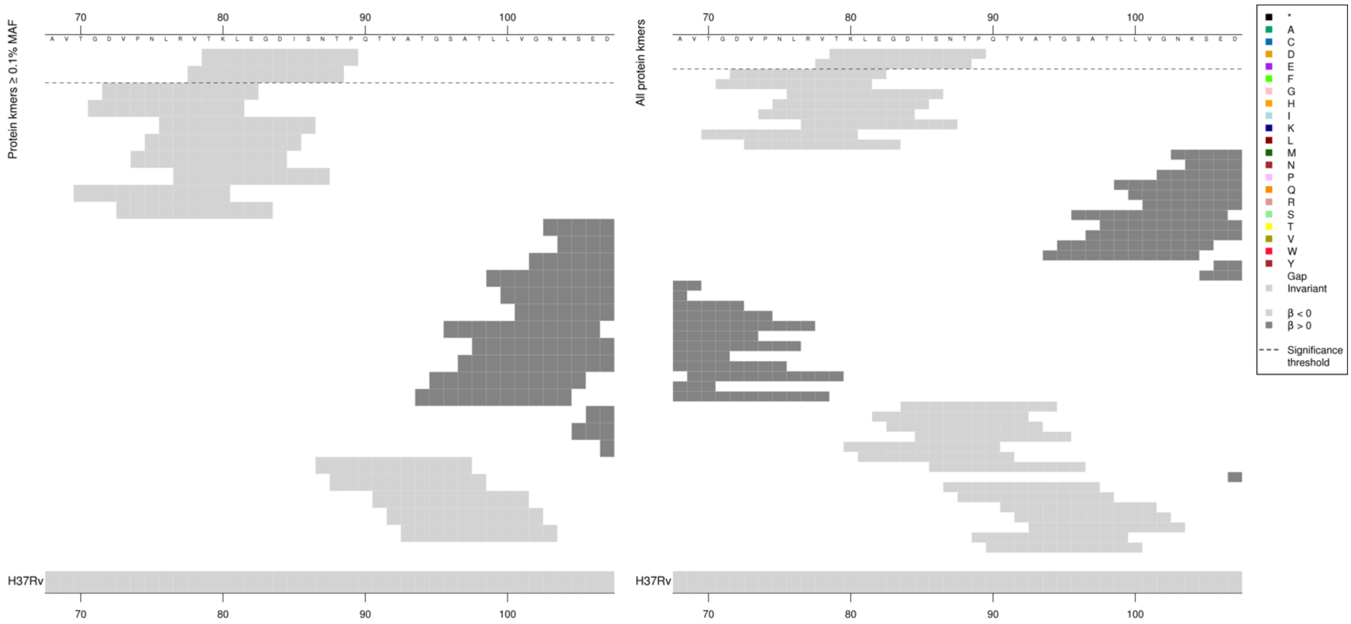

Removing the threshold requiring a kmer to be aligned to a gene/intergenic region in at least five genomes across the full dataset. Showing the protein kmer alignment after lowering the threshold to just one genome:

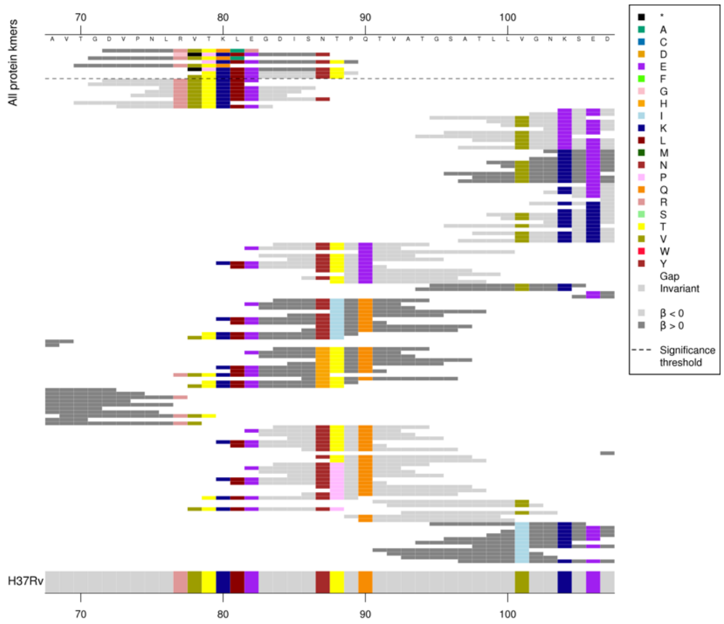

Rare variation is present in the region – variants in fewer than five genomes, possibly indels.

| Kmer | $-log_{10}$ *p* | β | MAC | MAF | Ps | Variants |
|---|---|---|---|---|---|---|
| TKLEGDISNTP | 7.88 | -1.93 | 8 | 0.12 | 79 | WT |

#### *pafA*

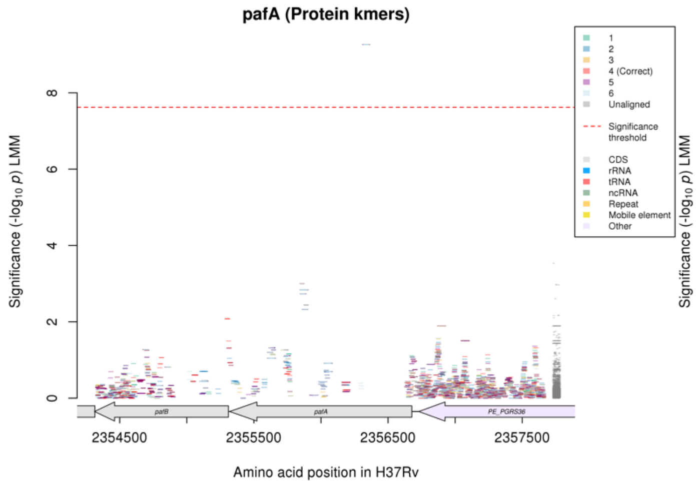

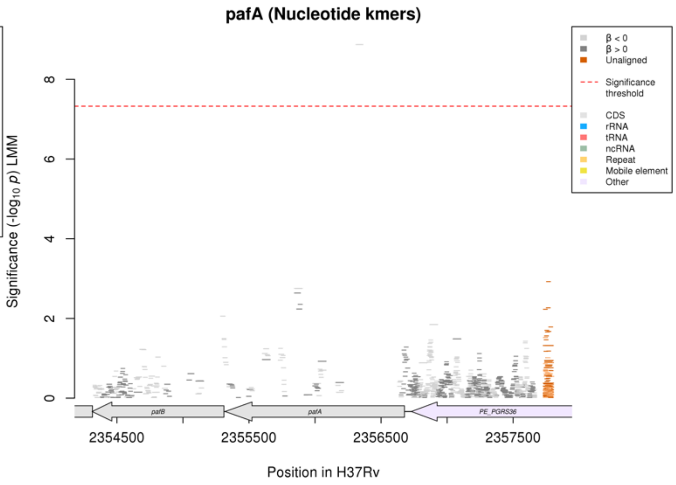

Nucleotide kmers:

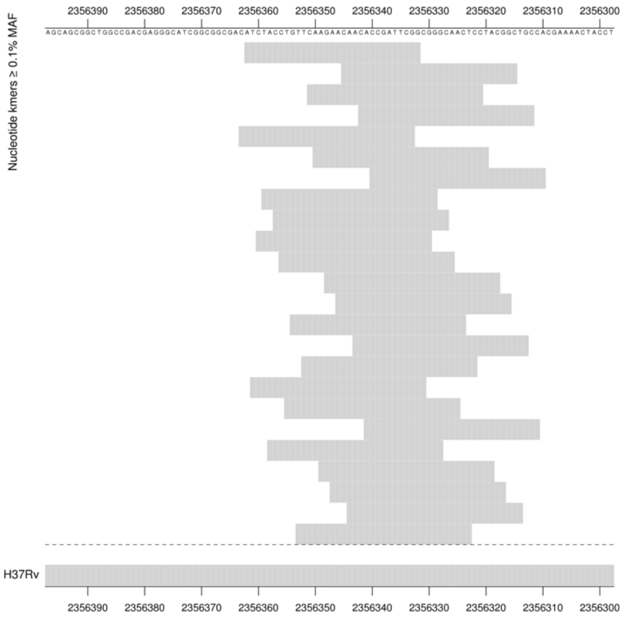

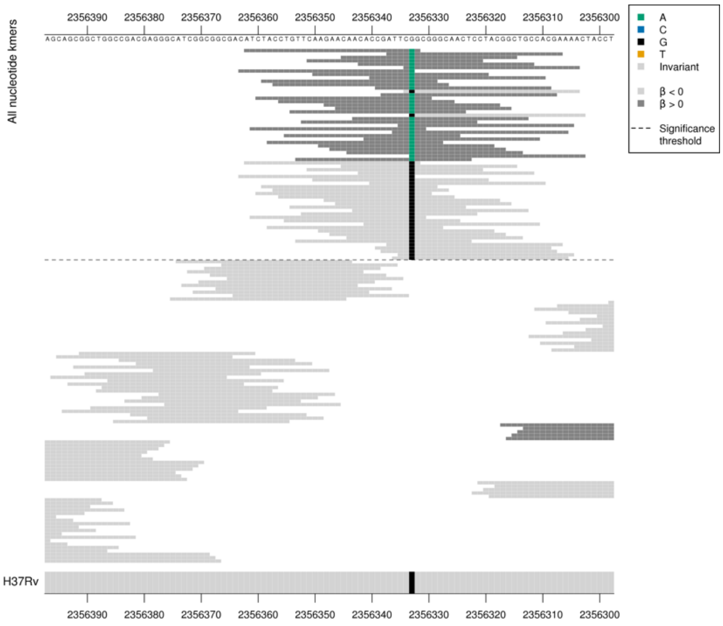

Synonymous mutation at position 2356333/345.
TCG = Ser / S
TCA = Ser / S

Low MAF kmers = 2356333/345A, 2356333/345G

| Kmer | -log₁₀ *p* | β | MAC | MAF | Ps | Variants |
|---|---|---|---|---|---|---|
| ATCTACCTGTTCAAGAACAACACCGATTCGG | 8.87 | -2.24 | 7 | 0.10 | 2356362 | 2356333/345G |

### *PE_PGRS6*

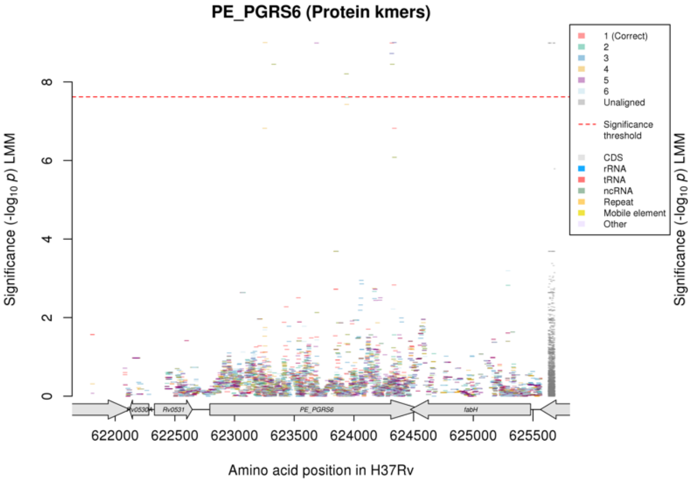

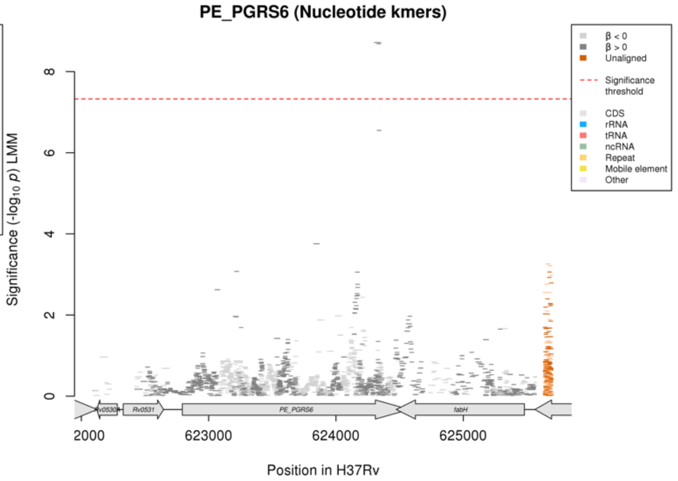

Protein kmers:

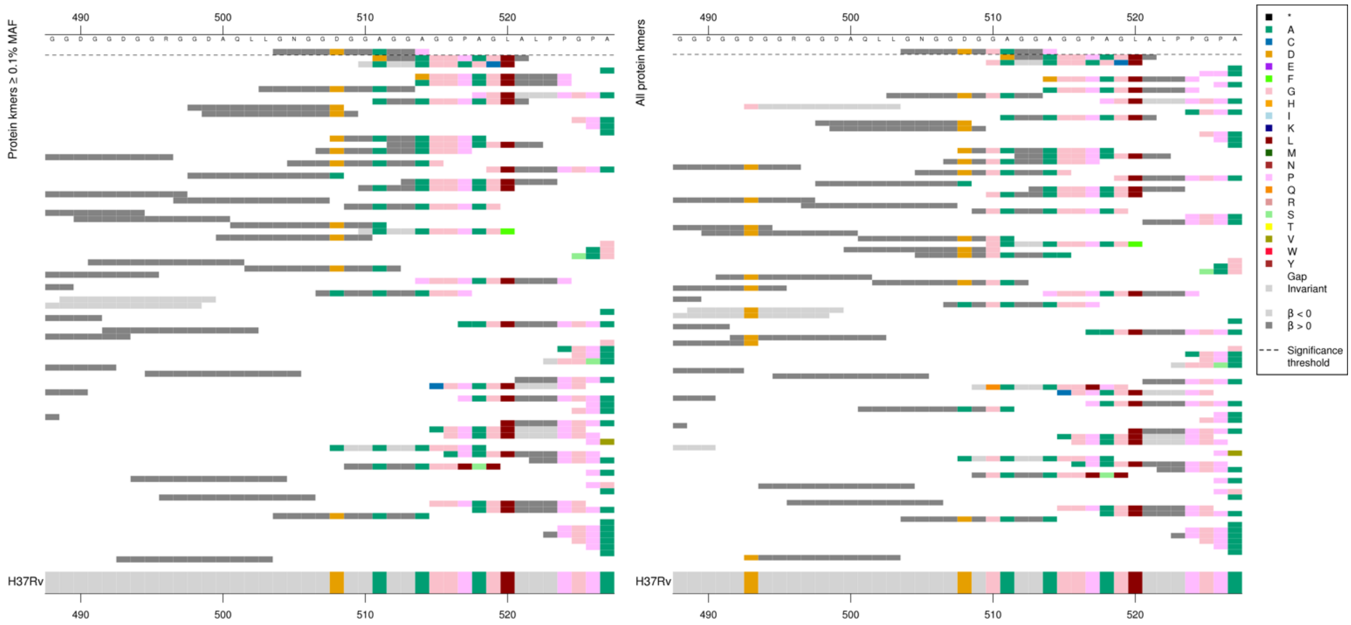

Same region for nucleotide kmers:

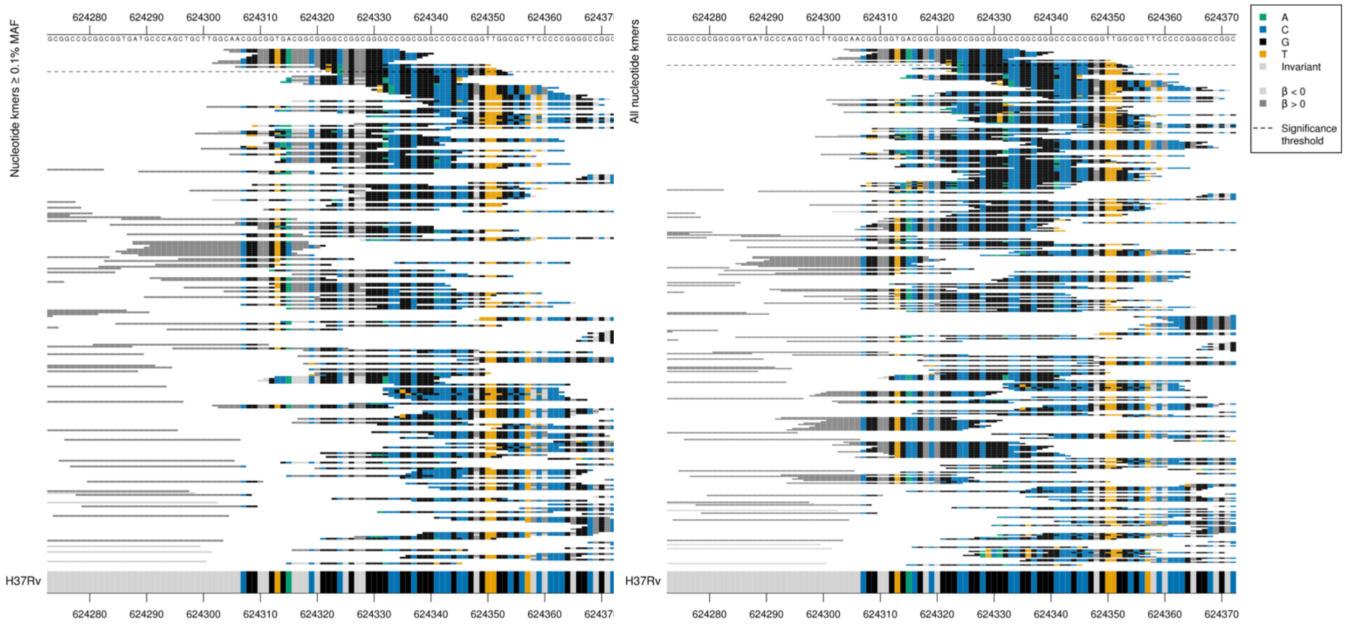

Differences compared to the reference but it’s unclear whether it is due to a point mutation or an indel.

| Kmer | -log₁₀ p | β | MAC | MAF | Ps | Variants |
|---|---|---|---|---|---|---|
| GNGGDGGAGGP | 8.99 | 1.66 | 11 | 0.16 | 504 | Multiple differences compared to reference |

#### *vapB20*

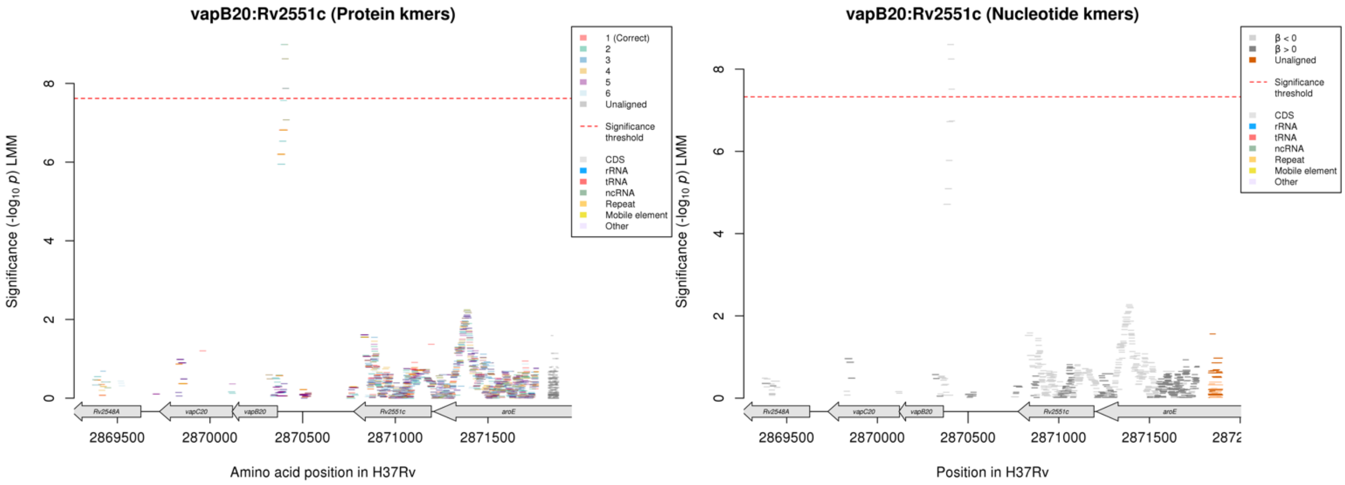

### Linezolid (LZD)

Nucleotide kmers:

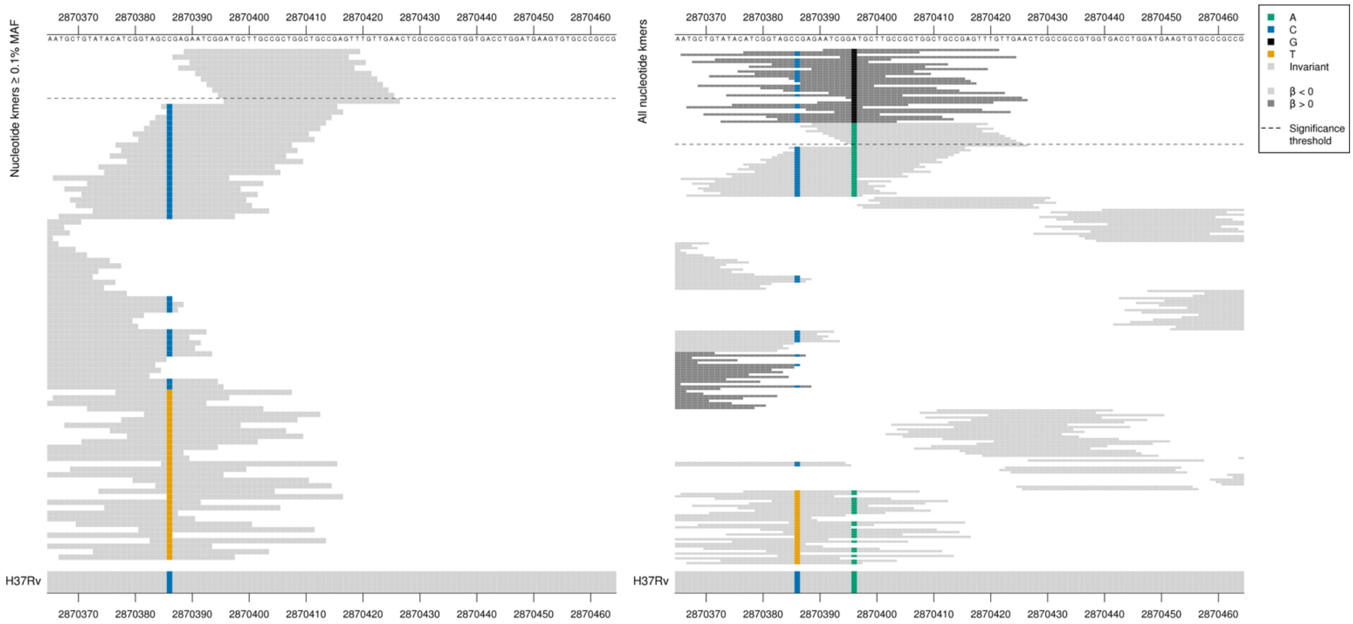

*vapB20* begins at position 2870364 on reverse strand. Possibly capturing a promoter region mutation.
On the correct strand for *vapB20*:
Low MIC-associated kmers = 2870396/-33 T

Low MAF kmers = 2870396/-33 C

| Kmer | $-\log_{10} p$ | β | MAC | MAF | Ps | Variants |
|---|---|---|---|---|---|---|
| GAATCGGATGCTTGCCGCTGGCTGCCGAGTT | 8.60 | -2.02 | 8 | 0.12 | 2870389 | 2870396/-33T (vapB20) |

# *Rv0061c*

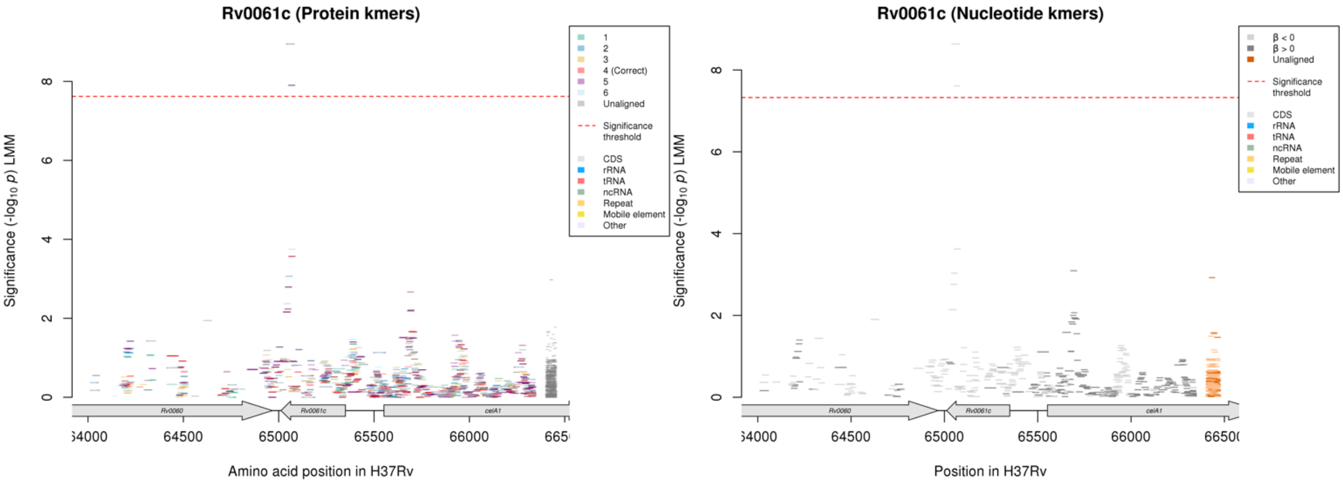

Protein kmers:

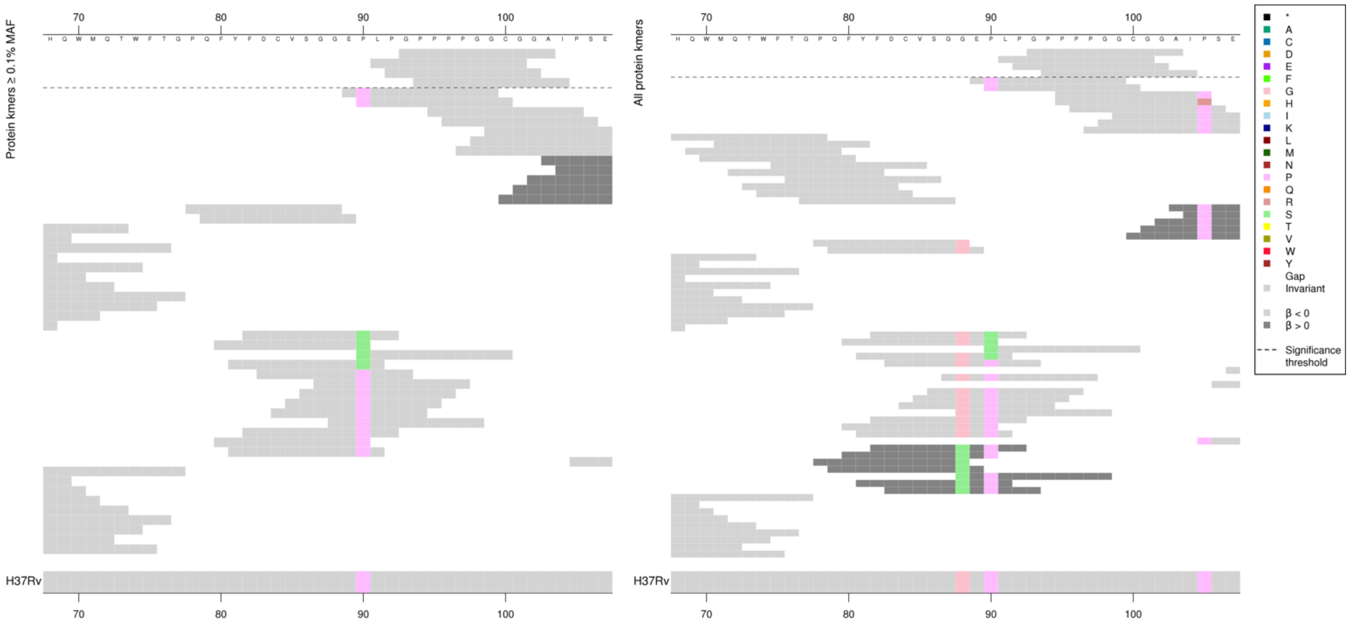

Removing the threshold requiring a kmer to be aligned to a gene/intergenic region in at least five genomes across the full dataset. Showing the protein kmer alignment after lowering the threshold to just one genome:

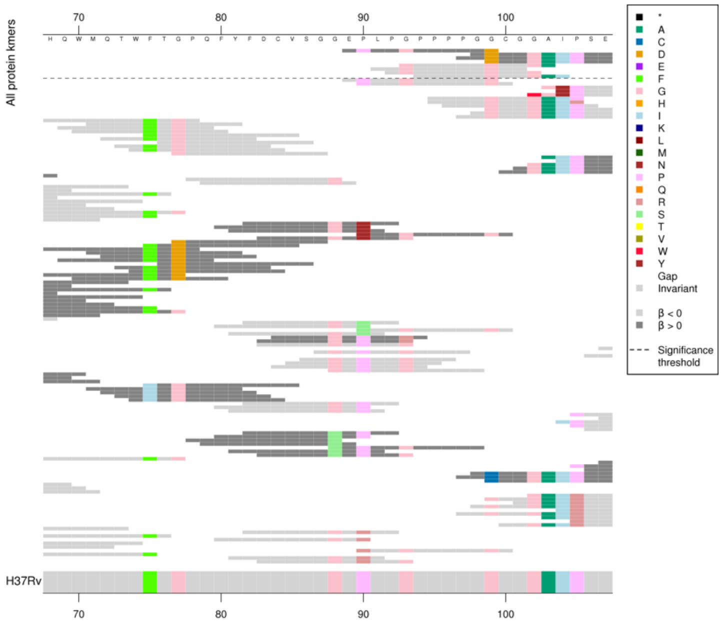

Rare variation present that is in fewer than five genomes.

| Kmer | $-\log_{10} p$ | β | MAC | MAF | Ps | Variants |
|---|---|---|---|---|---|---|
| GPPPPGGCGGA | 8.95 | -1.99 | 8 | 0.12 | 93 | WT codons 91-104 |

### *PE_PGRS4*

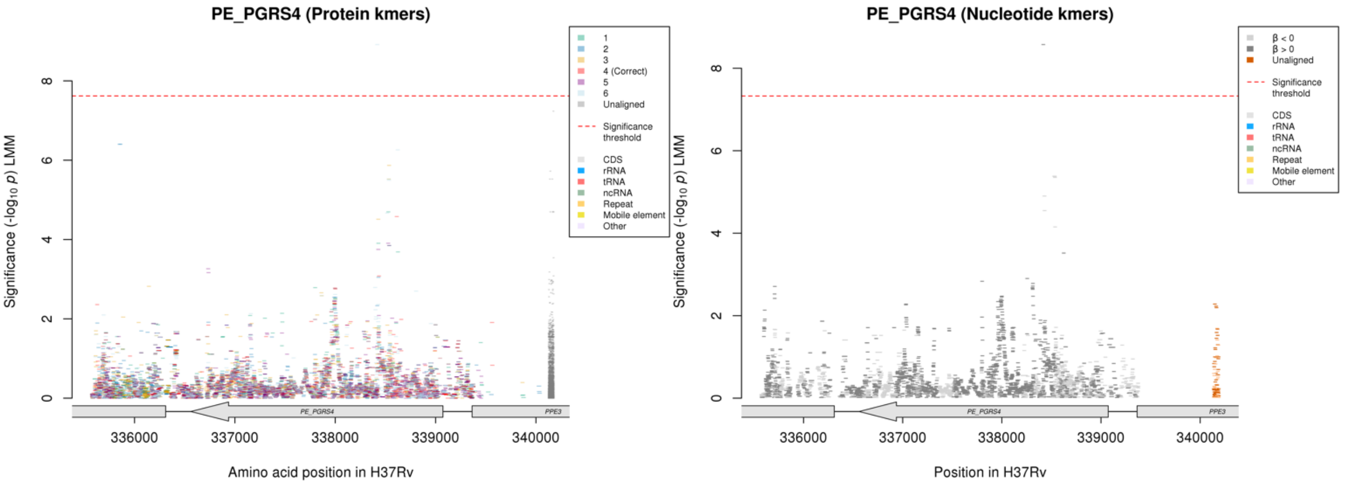

No significant protein kmers aligned to the correct reading frame for the gene *PE_PGRS4*.
Nucleotide kmers:

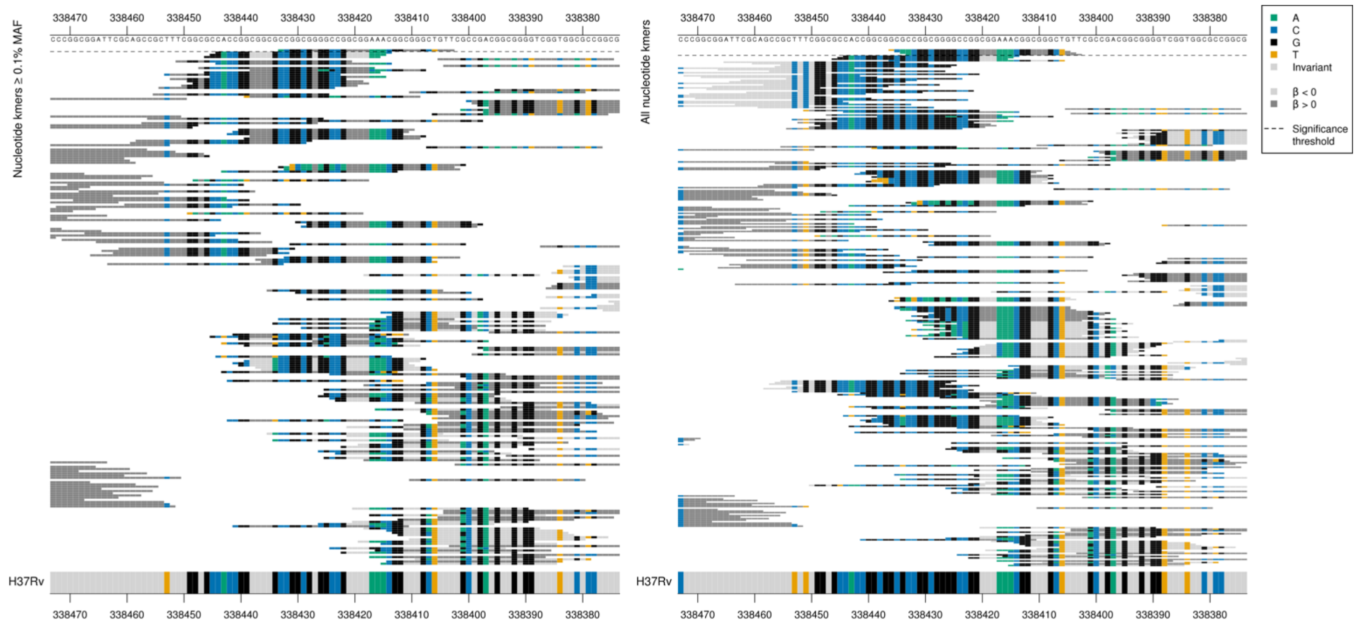

Multiple differences to the reference at one end of the kmers, possibly an insertion or deletion. Almost all of the kmer is identical to the reference.

| Kmer | -log₁₀ *p* | β | MAC | MAF | Ps | Variants |
|---|---|---|---|---|---|---|
| ACGGCGGGGCCGGCGGAAACGGCGGGCTGTT | 8.57 | 2.08 | 7 | 0.10 | 338433 | Multiple differences to reference - possible indel |

# *Rv1049*

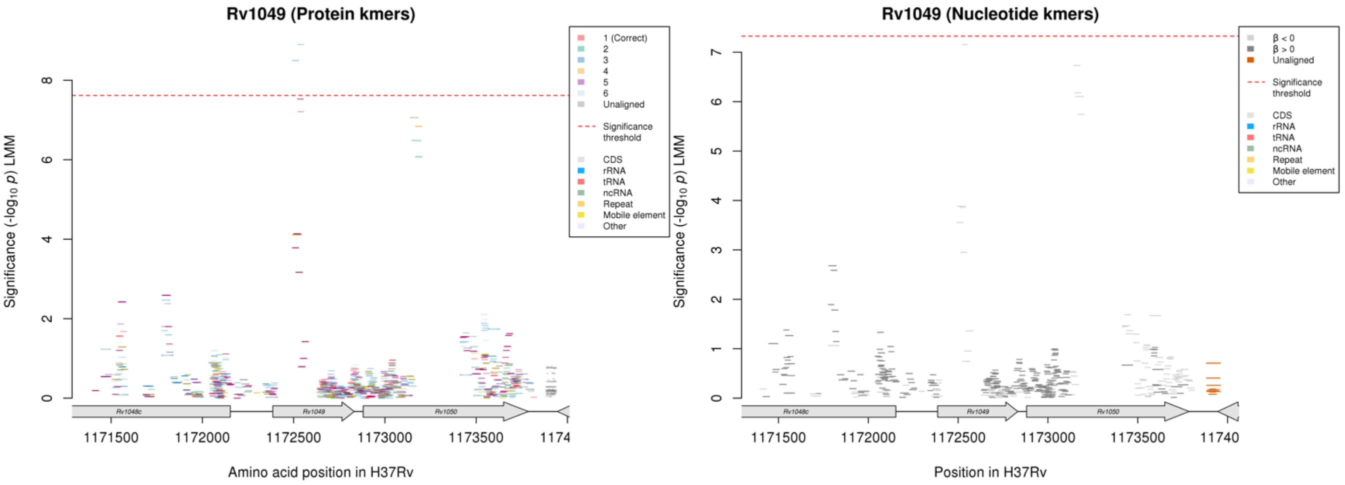

No significant kmers aligned to the correct reading frame for the protein kmer analysis.
Region just below the significance threshold for nucleotide kmers for those above the MAF threshold.

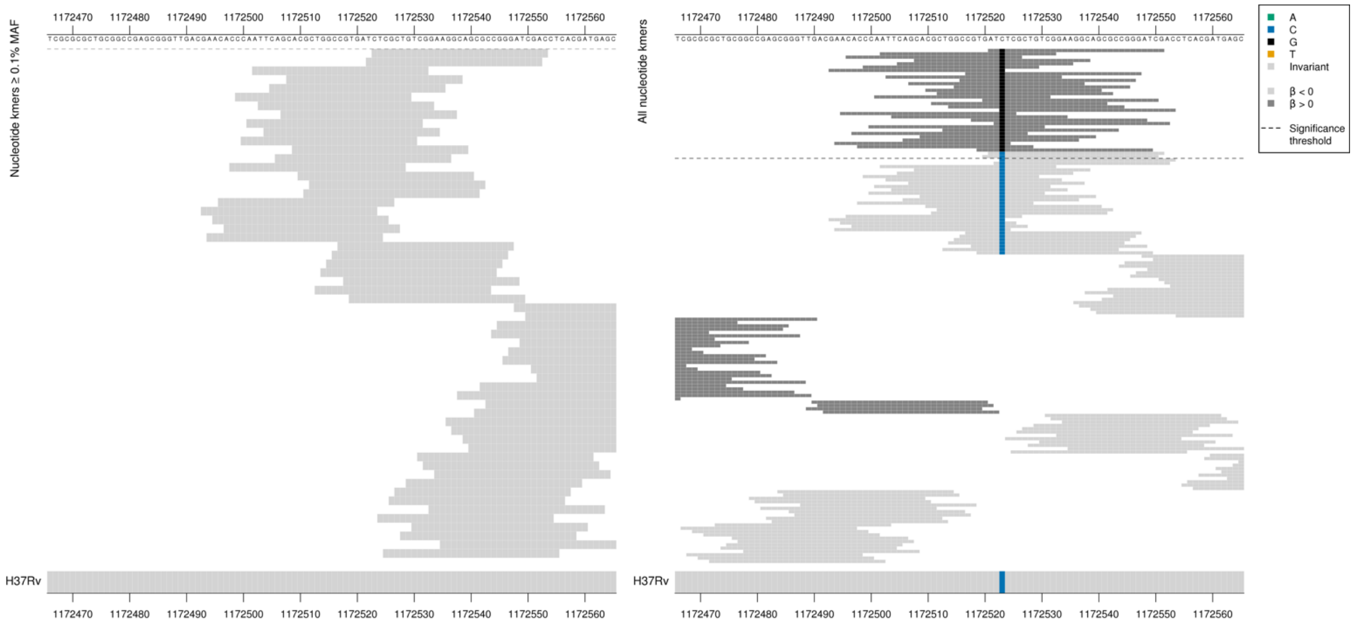

ATG = Met / M
ATC = Ile / I

Low MAF kmers = 1172523/138G, 1172523/138C

| Kmer | -log$_{10}$ *p* | β | MAC | MAF | Ps | Variants |
|---|---|---|---|---|---|---|
| CTCGCTGTCGGAAGGCAGCGCCGGGATCGAC | 7.16 | -1.78 | 8 | 0.12 | 1172523 | 1172523/138C |

### *lprF:Rv1371* R

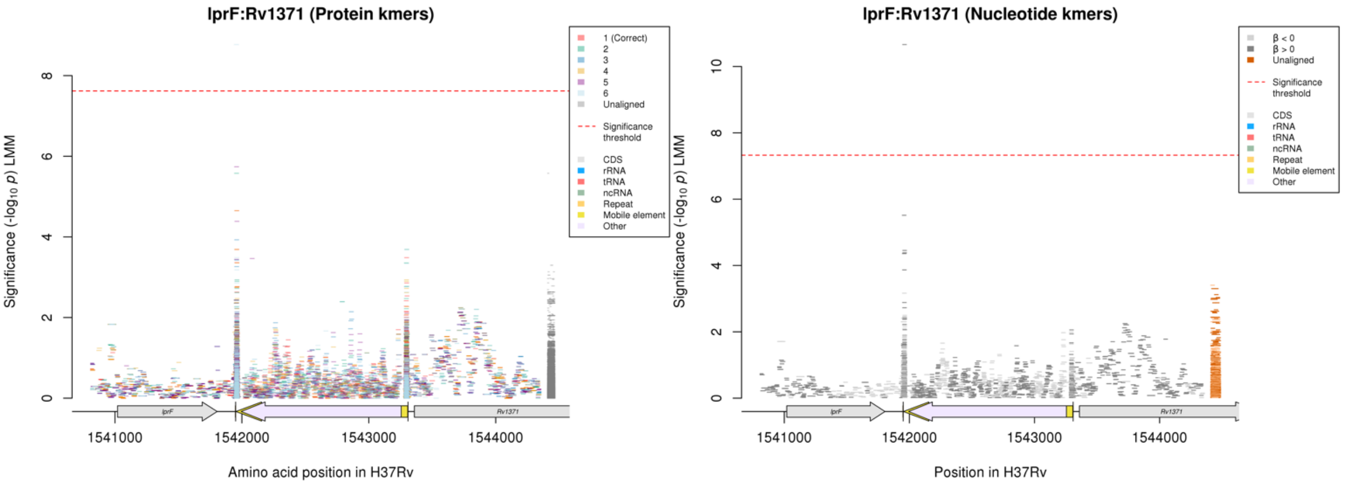

Nucleotide kmers:

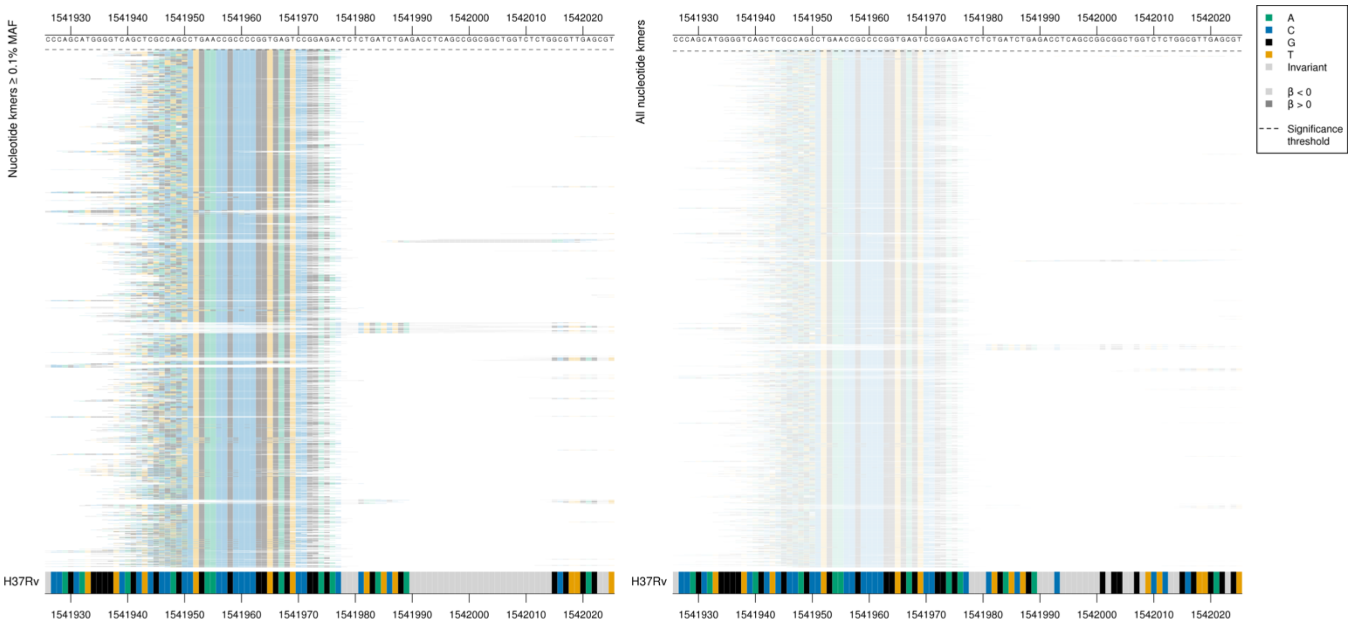

| Kmer | $-\log_{10} p$ | β | MAC | MAF | Ps | Variants |
|---|---|---|---|---|---|---|
| CACCACTGAACCGCCCCGGTGAGTCCGGAGA | 10.66 | 2.15 | 8 | 0.12 | 1541946 | Repeat region |

# *Rv3183:Rv3188* R

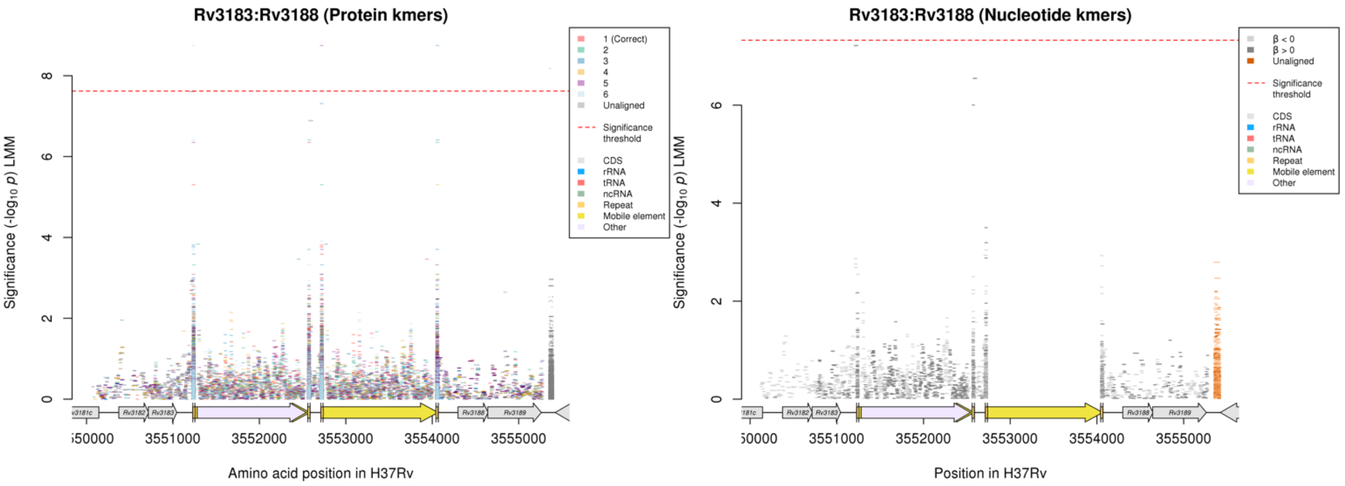

Showing the region for the top nucleotide kmers, just below the significance threshold:

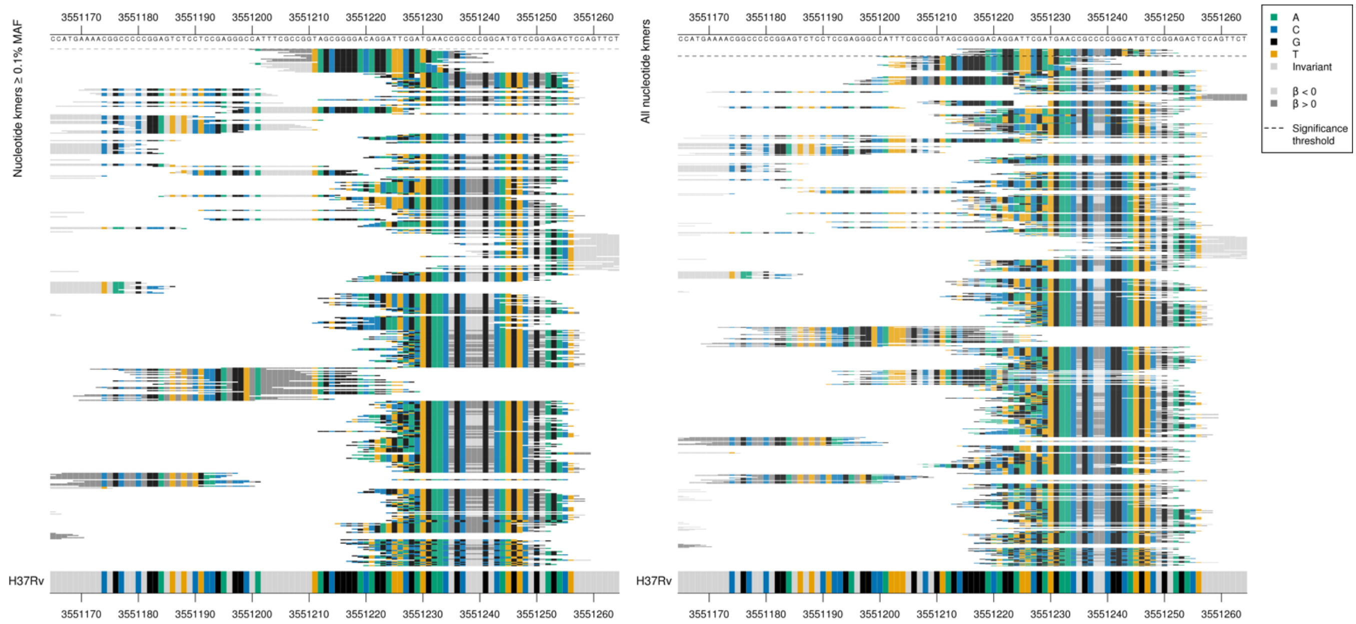

| Kmer | $-\log_{10} p$ | β | MAC | MAF | Ps | Variants |
|---|---|---|---|---|---|---|
| CATTTCGCCGGTAGCGGGGACAGGATTCGAT | 7.22 | 1.90 | 7 | 0.10 | 3551200 | Repeat region |

#### *dxs2:Rv3382c* [R]

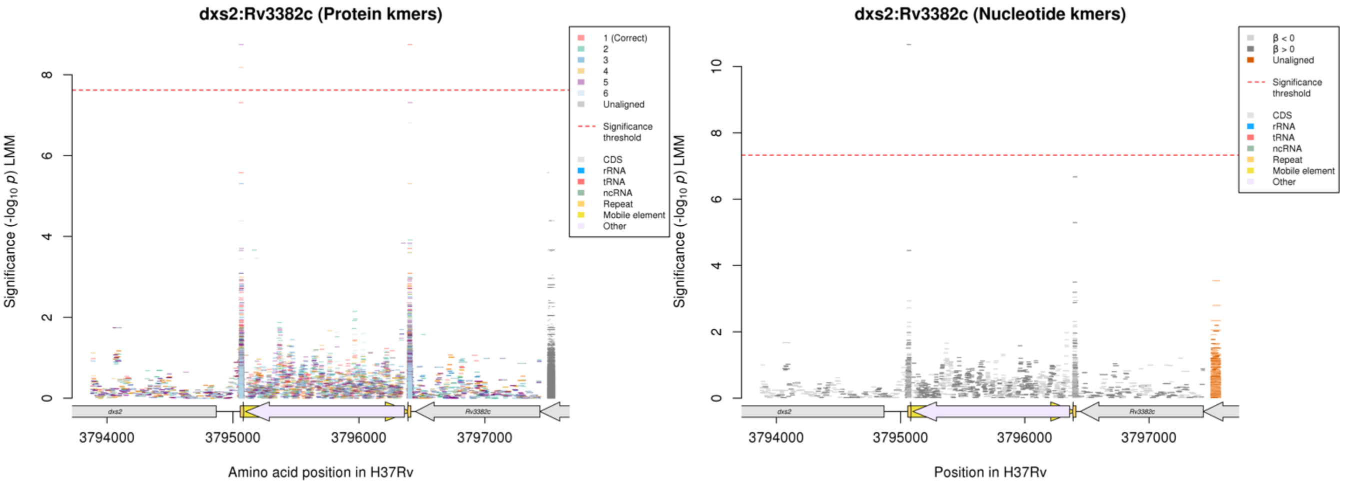

#### Repeat region

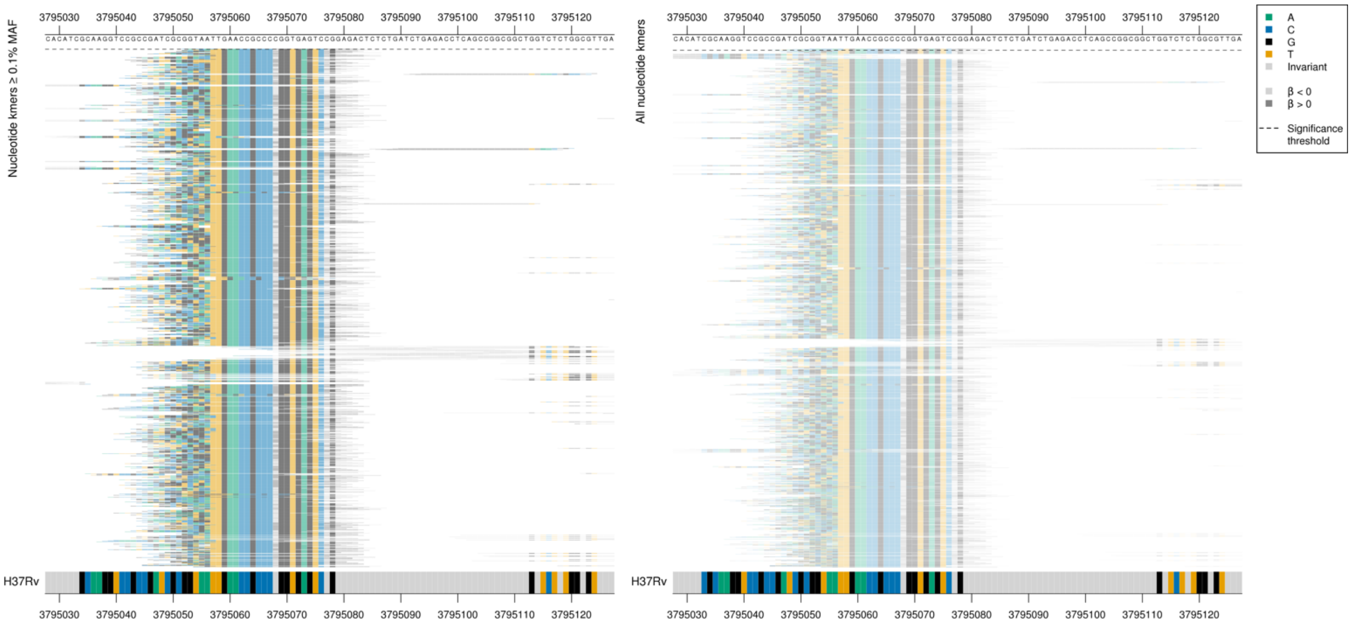

| Kmer | -log$_{10}$ *p* | β | MAC | MAF | Ps | Variants |
|---|---|---|---|---|---|---|
| CACCACTGAACCGCCCCGGTGAGTCCGGAGA | 10.66 | 2.15 | 8 | 0.12 | 3795052 | Repeat region |

# *Rv0556*

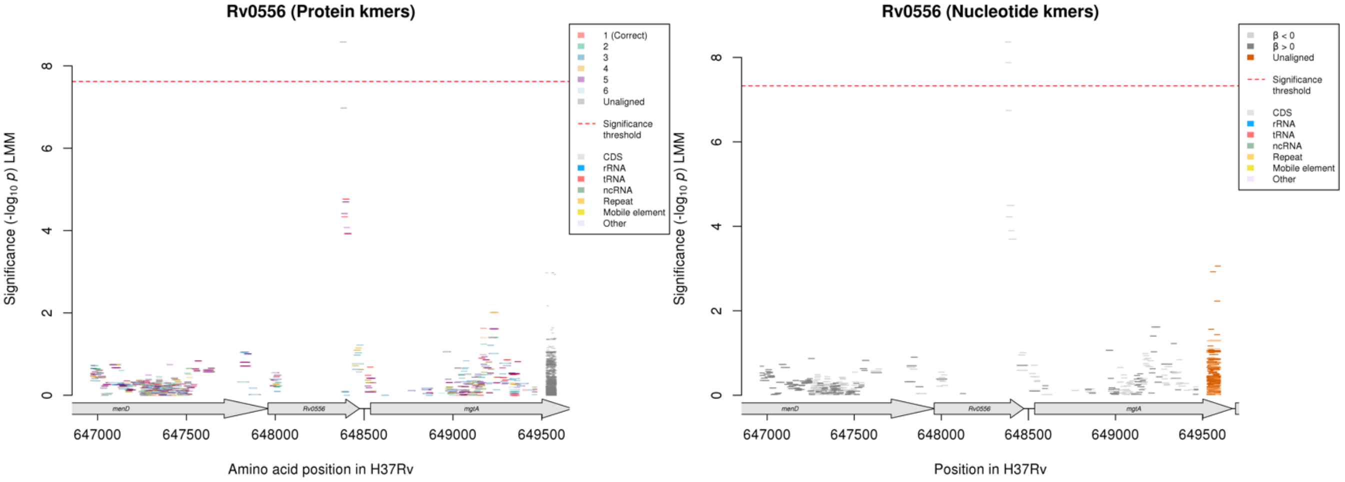

No significant protein kmers above the MAF cutoff aligned to the correct reading frame

Nucleotide kmers:

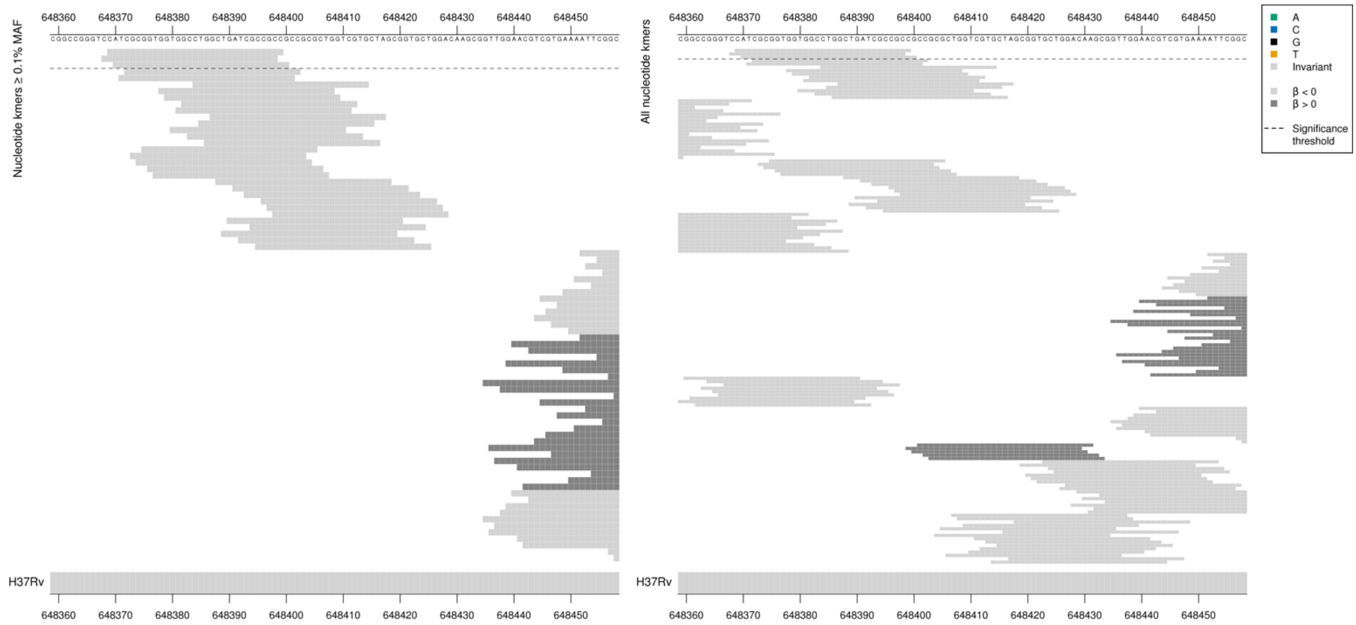

Removing the threshold requiring a kmer to be aligned to a gene/intergenic region in at least five genomes across the full dataset. Showing the nucleotide kmer alignment after lowering the threshold to just one genome:

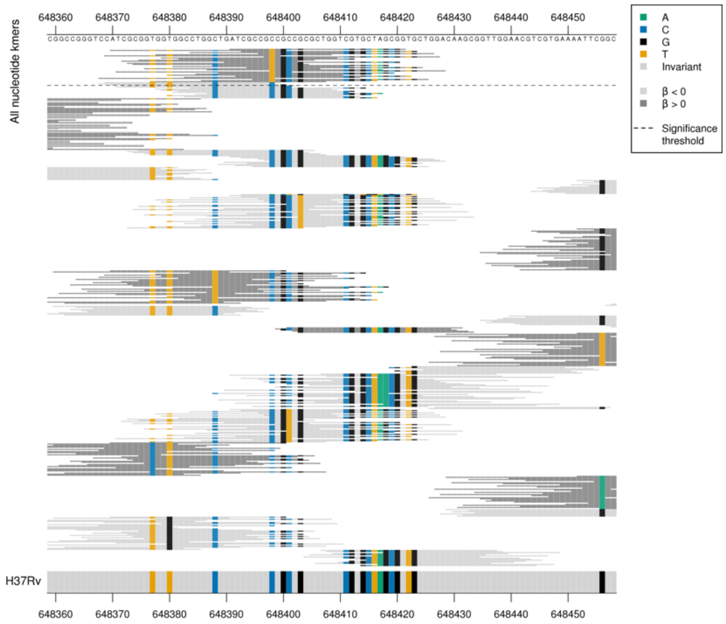

Rare variation in the region present in fewer than five genomes.
Kmer MAC between 8-9.

| Kmer | $-\log_{10} p$ | β | MAC | MAF | Ps | Variants |
|---|---|---|---|---|---|---|
| CATCGCGGTGGTGGCCTGGCTGATCGCCGCC | 8.36 | -1.98 | 8 | 0.12 | 648369 | WT |

## *Rv0514*

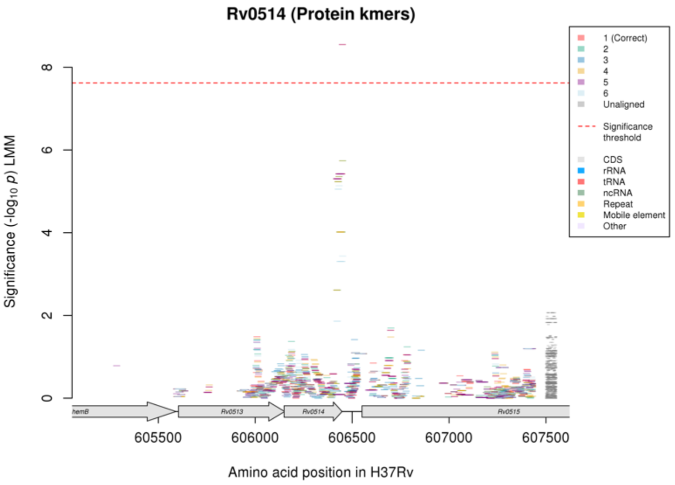

Not significant in the nucleotide kmer analysis.

Protein kmers:

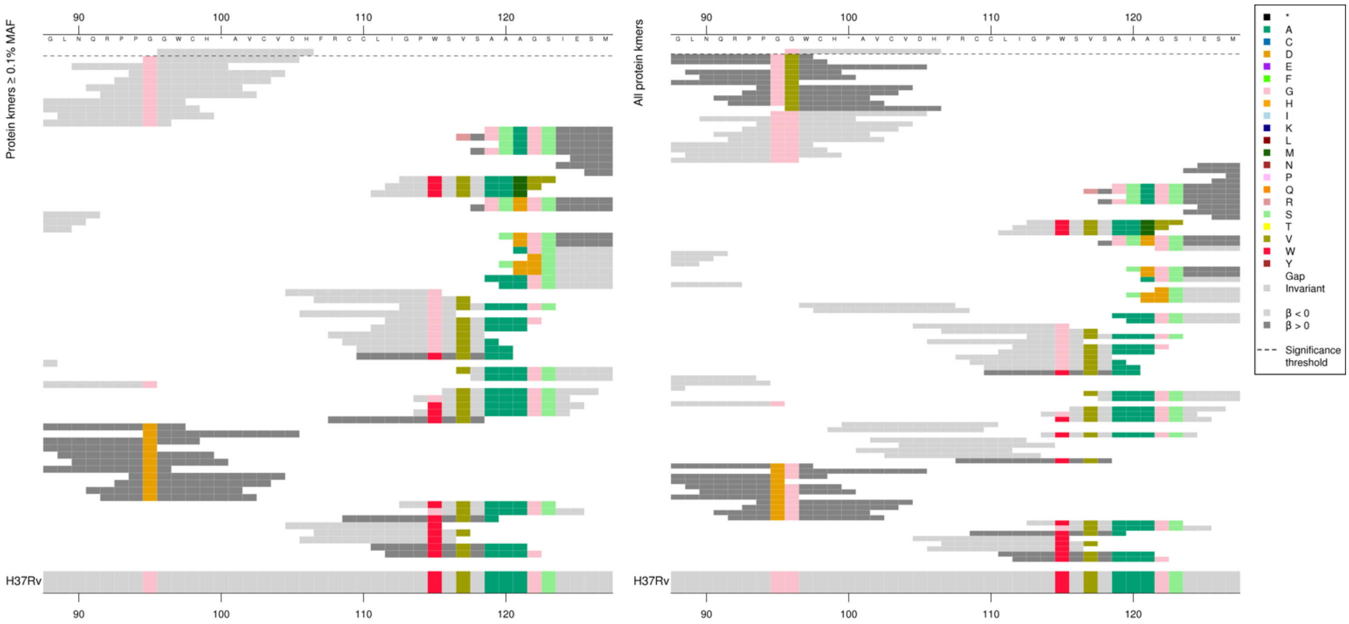

Rv0514 is 99 amino acids long

| Kmer | $-\log_{10} p$ | β | MAC | MAF | Ps | Variants |
|---|---|---|---|---|---|---|
| GWCH*AVCVDH | 8.55 | -1.87 | 10 | 0.15 | 96 | WT / 96G |

### Moxifloxacin (MXF)

#### *gyrA*

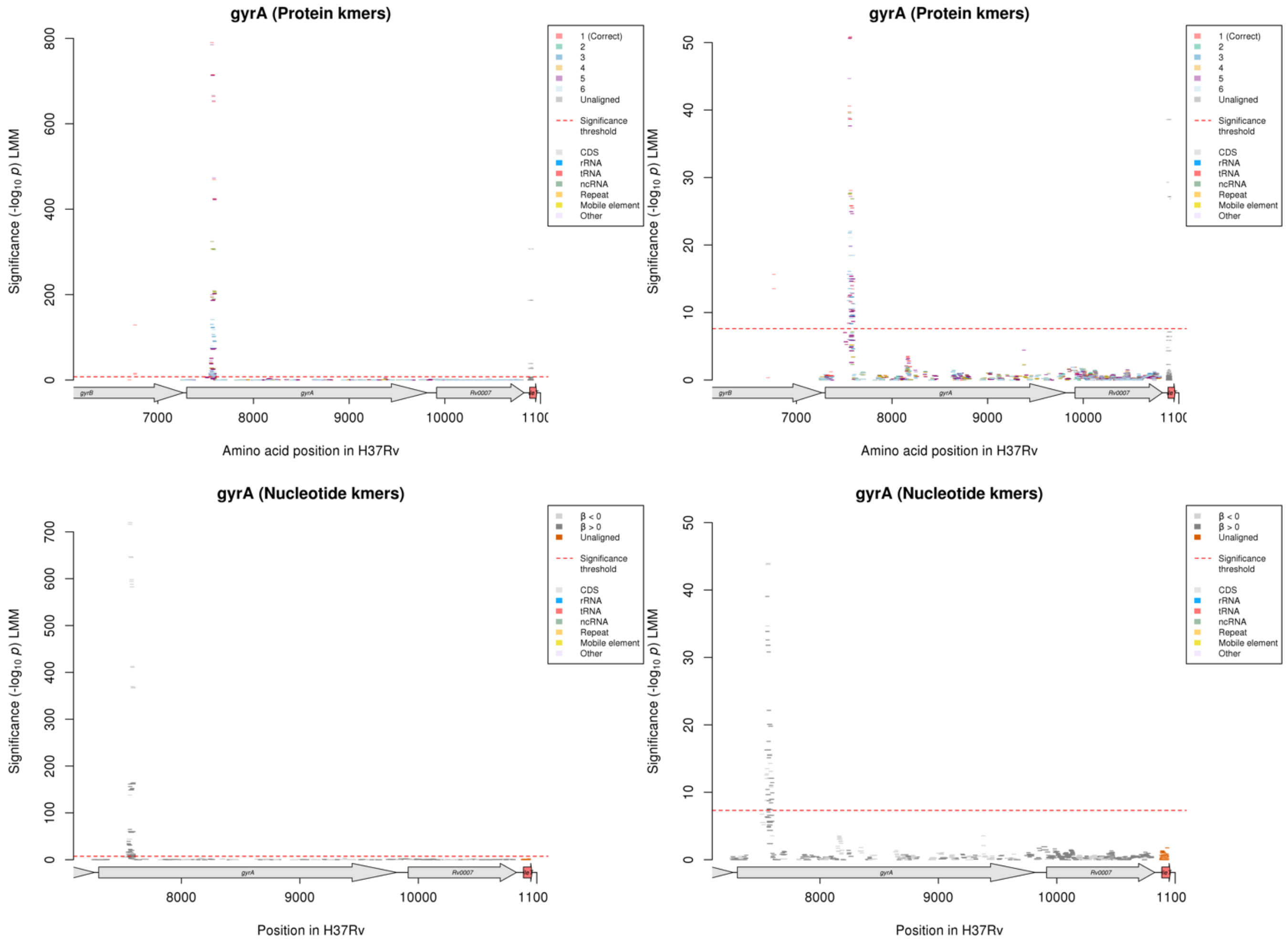

Protein kmers:

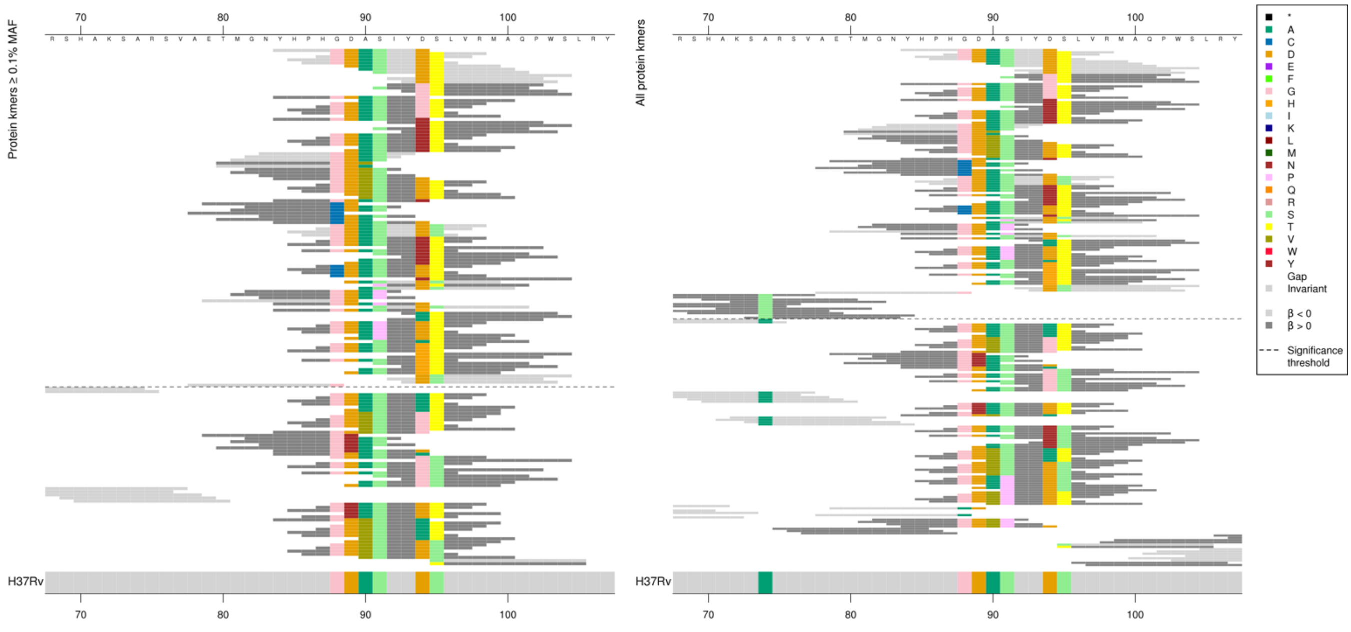

| Kmer | -log₁₀ *p* | β | MAC | MAF | Ps | Variants |
|---|---|---|---|---|---|---|
| YHPHGDASIYD | 790.20 | -3.53 | 1102 | 17.25 | 84 | 88G, 89D, 90A, 91S, 94D |
| GDASIYDTLVR | 714.41 | -3.44 | 1391 | 21.78 | 88 | 88G, 89D, 90A, 91S, 94D, 95T |
| DASIYDTLVRM | 663.70 | -3.36 | 1375 | 21.52 | 89 | 89D, 90A, 91S, 94D, 95T |
| ASIYDTLVRMA | 651.41 | -3.34 | 1368 | 21.42 | 90 | 90A, 91S, 94D, 95T |
| SIYDTLVRMAQ | 469.35 | -3.09 | 1104 | 17.28 | 91 | 91S, 94D, 95T |
| DTLVRMAQPWS | 423.62 | -3.01 | 1051 | 16.45 | 94 | 94D, 95T |
| IYGTLVRMAQP | 202.31 | 2.49 | 501 | 7.84 | 92 | 94G [+ 95T] |
| SIYGTLVRMAQ | 202.31 | 2.49 | 501 | 7.84 | 91 | 94G [+ 91S, 95T] |
| YHPHGDASIYG | 193.84 | 2.42 | 501 | 7.84 | 84 | 94G [+ 88G, 89D, 90A, 91S] |
| ASIYGTLVRMA | 188.48 | 2.43 | 490 | 7.67 | 90 | 94G [+ 90A, 91S, 95T] |
| DASIYGTLVRM | 186.40 | 2.42 | 488 | 7.64 | 89 | 94G [+ 89D, 90A, 91S, 95T] |
| YHPHGDASIYN | 73.79 | 2.48 | 99 | 1.55 | 84 | 94N [+ 88G, 89D, 90A, 91S] |
| IYNTLVRMAQP | 73.73 | 2.48 | 100 | 1.57 | 92 | 94N [+ 95T] |
| SIYNTLVRMAQ | 73.73 | 2.48 | 100 | 1.57 | 91 | 94N [+ 91S, 95T] |
| GDASIYNTLVR | 72.29 | 2.47 | 98 | 1.53 | 88 | 94N [+ 88G, 89D, 90A, 91S, 95T] |
| ASIYNTLVRMA | 72.29 | 2.47 | 98 | 1.53 | 90 | 94N [+ 90A, 91S, 95T] |
| DASIYNTLVRM | 72.29 | 2.47 | 98 | 1.53 | 89 | 94N [+ 89D, 90A, 91S, 95T] |
| NYHPHGDASIY | 50.87 | -1.29 | 360 | 5.64 | 83 | 88G, 89D, 90A, 91S |
| TMGNYHPHGDV | 40.58 | 1.24 | 312 | 4.88 | 80 | 90V [+ 88G, 89D] |
| TMGNYHPHGDA | 39.73 | -1.22 | 312 | 4.88 | 80 | 88G, 89D, 90A |
| GNYHPHGDVSI | 38.64 | 1.22 | 305 | 4.77 | 82 | 90V [+ 88G, 89D, 91S] |
| YHPHGDVSIYD | 28.08 | 1.06 | 283 | 4.43 | 84 | 90V [+ 88G, 89D, 91S, 94D] |
| GDVSIYDTLVR | 25.80 | 1.03 | 272 | 4.26 | 88 | 90V [+ 88G, 89D, 91S, 94D, 95T] |
| DVSIYDTLVRM | 25.49 | 1.03 | 273 | 4.27 | 89 | 90V [+ 89D, 91S, 94D, 95T] |
| VSIYDTLVRMA | 25.49 | 1.03 | 273 | 4.27 | 90 | 90V [+ 91S, 94D, 95T] |
| YHPHGDASIYY | 18.45 | 2.02 | 39 | 0.61 | 84 | 94Y [+ 88G, 89D, 90A, 91S] |
| ETMGNYHPHCD | 15.65 | 2.70 | 16 | 0.25 | 79 | 88C [+ 89D] |
| GNYHPHCDASI | 15.65 | 2.70 | 16 | 0.25 | 82 | 88C [+ 89D, 90A, 91S] |
| AETMGNYHPHC | 15.65 | 2.70 | 16 | 0.25 | 78 | 88C |
| TMGNYHPHCDA | 15.65 | 2.70 | 16 | 0.25 | 80 | 88C [+ 89D, 90A] |
| GDASIYDSLVR | 15.37 | -2.01 | 289 | 4.52 | 88 | 88G, 89D, 90A, 91S, 94D, 95S |
| GDASIYYTLVR | 14.98 | 1.90 | 33 | 0.52 | 88 | 94Y [+ 88G, 89D, 90A, 91S, 95T] |
| IYYTLVRMAQP | 14.98 | 1.90 | 33 | 0.52 | 92 | 94Y [+ 95T] |
| SIYYTLVRMAQ | 14.98 | 1.90 | 33 | 0.52 | 91 | 94Y [+ 91S, 95T] |
| ASIYYTLVRMA | 14.98 | 1.90 | 33 | 0.52 | 90 | 94Y [+ 90A, 91S, 95T] |
| DASIYYTLVRM | 14.98 | 1.90 | 33 | 0.52 | 89 | 94Y [+ 89D, 90A, 91S, 95T] |
| HCDASIYDTLV | 14.65 | 2.69 | 15 | 0.23 | 87 | 88C [+ 89D, 90A, 91S, 94D, 95T] |
| DASIYDSLVRM | 13.98 | -1.91 | 290 | 4.54 | 89 | 89D, 90A, 91S, 94D, 95S |
| PIYDTLVRMAQ | 13.49 | 1.41 | 56 | 0.88 | 91 | 91P [+ 94D, 95T] |
| ASIYDSLVRMA | 12.83 | -1.84 | 291 | 4.56 | 90 | 90A, 91S, 94D, 95S |
| GNYHPHGDAPI | 12.57 | 1.36 | 55 | 0.86 | 82 | 91P [+ 88G, 89D, 90A] |
| ETMGNYHPHGD | 12.35 | -1.87 | 28 | 0.44 | 79 | 88G, 89D |
| YHPHGDAPIYD | 11.61 | 1.34 | 53 | 0.83 | 84 | 91P [+ 88G, 89D, 90A, 94D] |

| | | | | | | |
|---|---|---|---|---|---|---|
| SIYDSLVRMAQ | 10.43 | -1.79 | 301 | 4.71 | 91 | 91S, 94D, 95S |
| YHPHGDASIYH | 10.43 | 1.55 | 32 | 0.50 | 84 | 94H [+ 88G, 89D, 90A, 91S] |
| YATLVRMAQPW | 10.25 | 0.99 | 86 | 1.35 | 93 | 94A [+95T] |
| GDAPIYDTLVR | 10.14 | 1.29 | 49 | 0.77 | 88 | 91P [+ 88G, 89D, 90A, 94D, 95T] |
| APIYDTLVRMA | 10.14 | 1.29 | 49 | 0.77 | 90 | 91P [+ 90A, 94D, 95T] |
| DAPIYDTLVRM | 10.14 | 1.29 | 49 | 0.77 | 89 | 91P [+ 89D, 90A, 94D, 95T] |
| SIYATLVRMAQ | 9.71 | 0.96 | 86 | 1.35 | 91 | 91A [+ 91S, 95T] |
| GDASIYHTLVR | 9.40 | 1.52 | 30 | 0.47 | 88 | 94H [+ 88G, 89D, 90A, 91S, 95T] |
| HTLVRMAQPWS | 9.40 | 1.52 | 30 | 0.47 | 94 | 94H [+ 95T] |
| SIYHTLVRMAQ | 9.40 | 1.52 | 30 | 0.47 | 91 | 94H [+ 91S, 95T] |
| ASIYHTLVRMA | 9.40 | 1.52 | 30 | 0.47 | 90 | 94H [+ 90A, 91S, 95T] |
| DASIYHTLVRM | 9.40 | 1.52 | 30 | 0.47 | 89 | 94H [+ 89D, 90A, 91S, 95T] |
| DSLVRMAQPWS | 8.67 | -1.71 | 306 | 4.79 | 94 | 94D, 95S |
| AETMGNYHPHG | 8.41 | -1.82 | 20 | 0.31 | 78 | 88G |

*rrs*

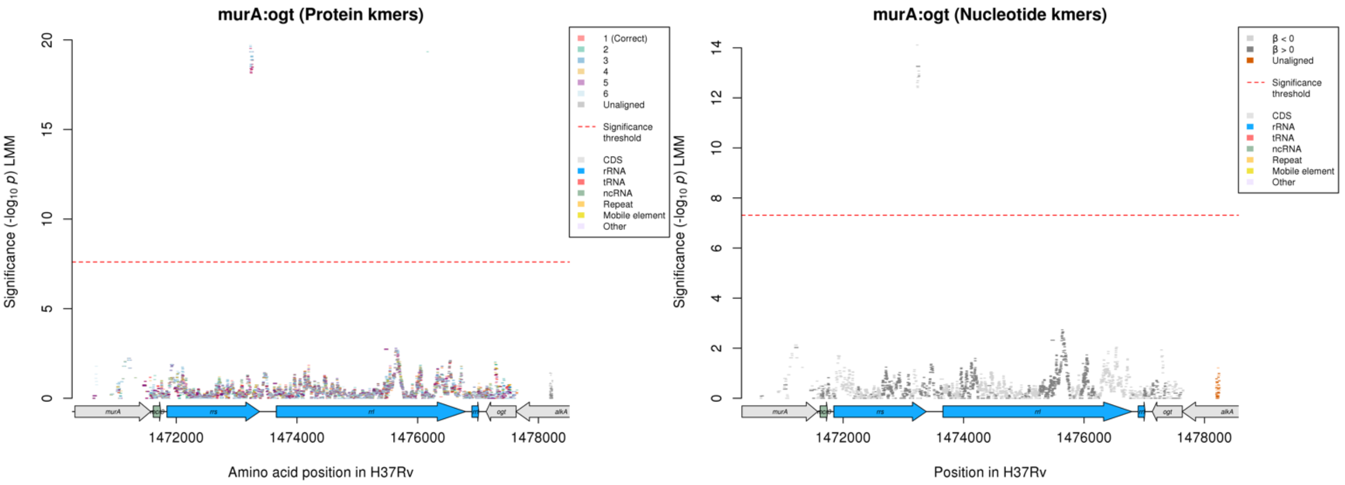

Nucleotide kmers:

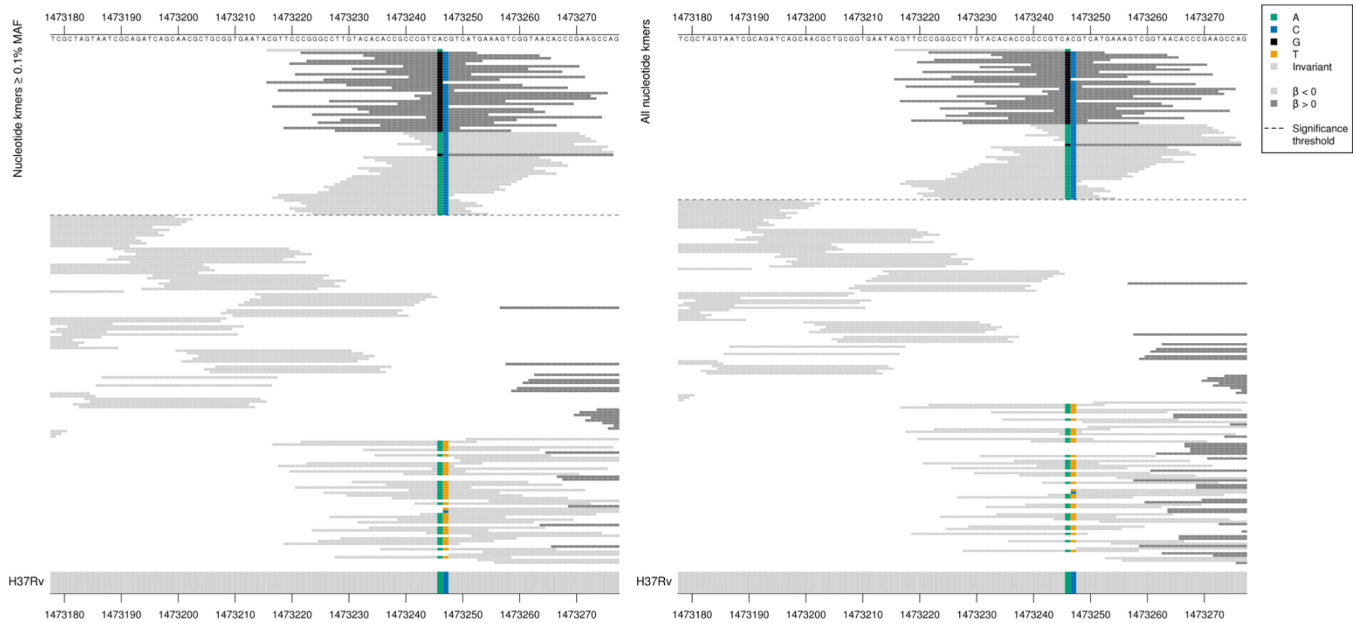

The low MIC-associated kmers were a range of *p*-values, not directly corresponding with whether they covered position 1473247/1402 or not. All but the last high MIC-associated kmer was the same *p*-value, so seem to just be capturing position 1473246/1401.

| Kmer | -log₁₀ *p* | β | MAC | MAF | Ps | Variants |
|---|---|---|---|---|---|---|
| CGTTCCCGGGCCTTGTACACACCGCCCGTCA | 14.11 | -0.78 | 459 | 7.19 | 1473216 | 1473246/1401A |
| CGGGCCTTGTACACACCGCCCGTCGCGTCAT | 13.26 | 0.77 | 428 | 6.70 | 1473222 | 1473246/1401G |

#### *rpoB*

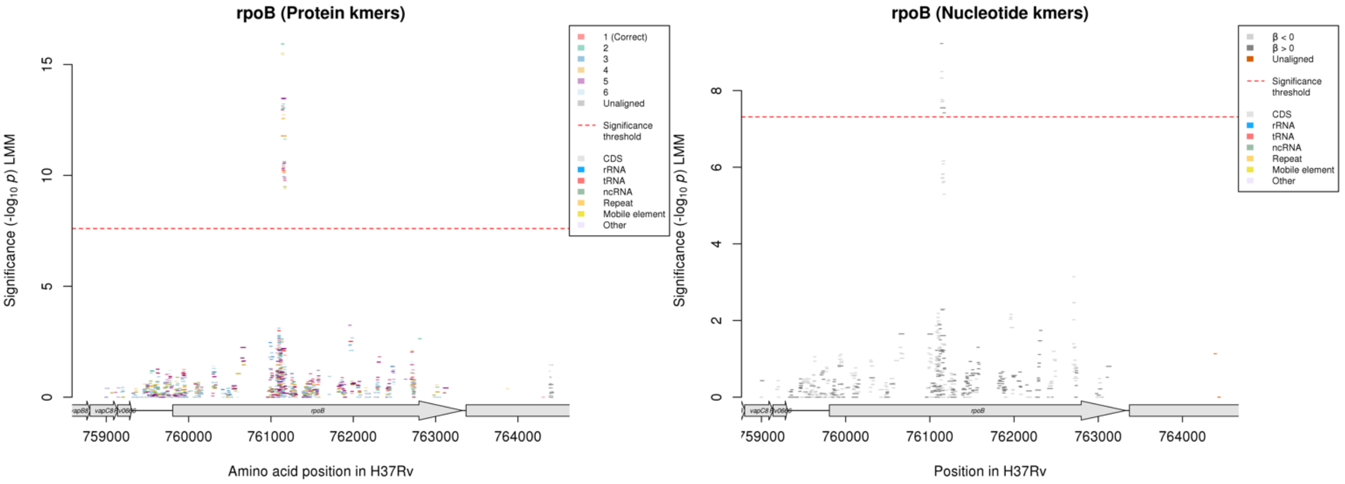

Protein kmers:

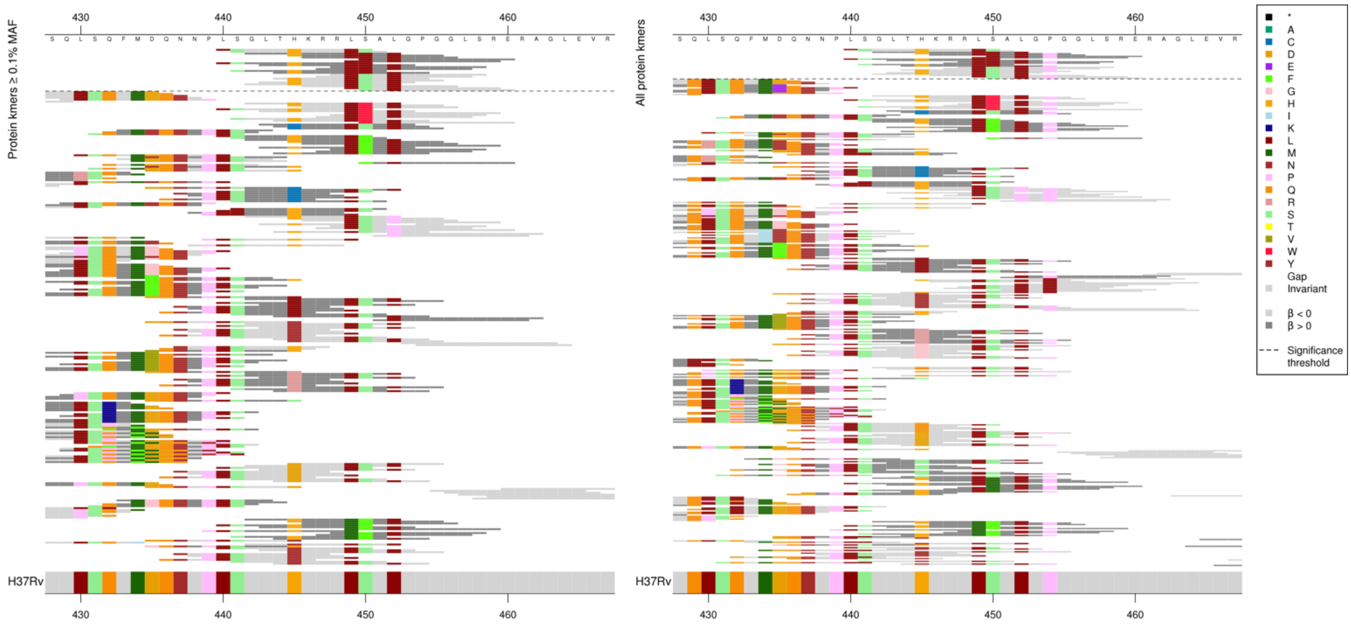

High MIC-associated kmers were all the same *p*-value and were just capturing 450L, the low MIC-associated kmers varied in *p*-value depending on the other substitutions covered.

| Kmer | -log₁₀ *p* | β | MAC | MAF | Ps | Variants |
|---|---|---|---|---|---|---|
| LSGLTHKRRLS | 12.97 | -0.50 | 1955 | 30.60 | 440 | 440L, 441S, 445H, 449L, 450S |
| SGLTHKRRLSA | 12.94 | -0.50 | 1958 | 30.65 | 441 | 441S, 445H, 449L, 450S |
| GLTHKRRLLAL | 11.78 | 0.52 | 1632 | 25.55 | 442 | 450L |
| RLSALGPGGLS | 10.56 | -0.48 | 1767 | 27.66 | 448 | 449L, 450S, 452L, 454P |
| THKRRLSALGP | 10.39 | -0.44 | 2017 | 31.57 | 444 | 445H, 449L, 450S, 452L, 454P |
| LTHKRRLSALG | 10.23 | -0.44 | 2014 | 31.53 | 443 | 445H, 449L, 450S, 452L |
| SALGPGGLSRE | 9.77 | -0.46 | 1765 | 27.63 | 450 | 450S, 452L, 454P |

#### *gyrB*

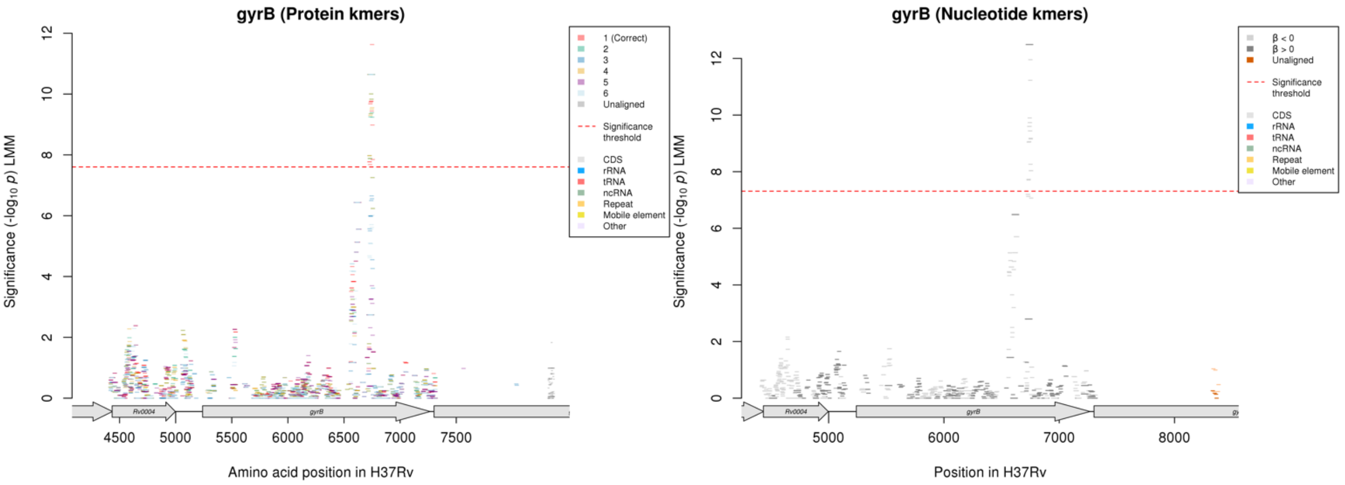

Peak is in a different location to the peak for LEV. The upstream peak just below significance is where the significant kmers fall for LEV, capturing codon 461.

Protein kmers:

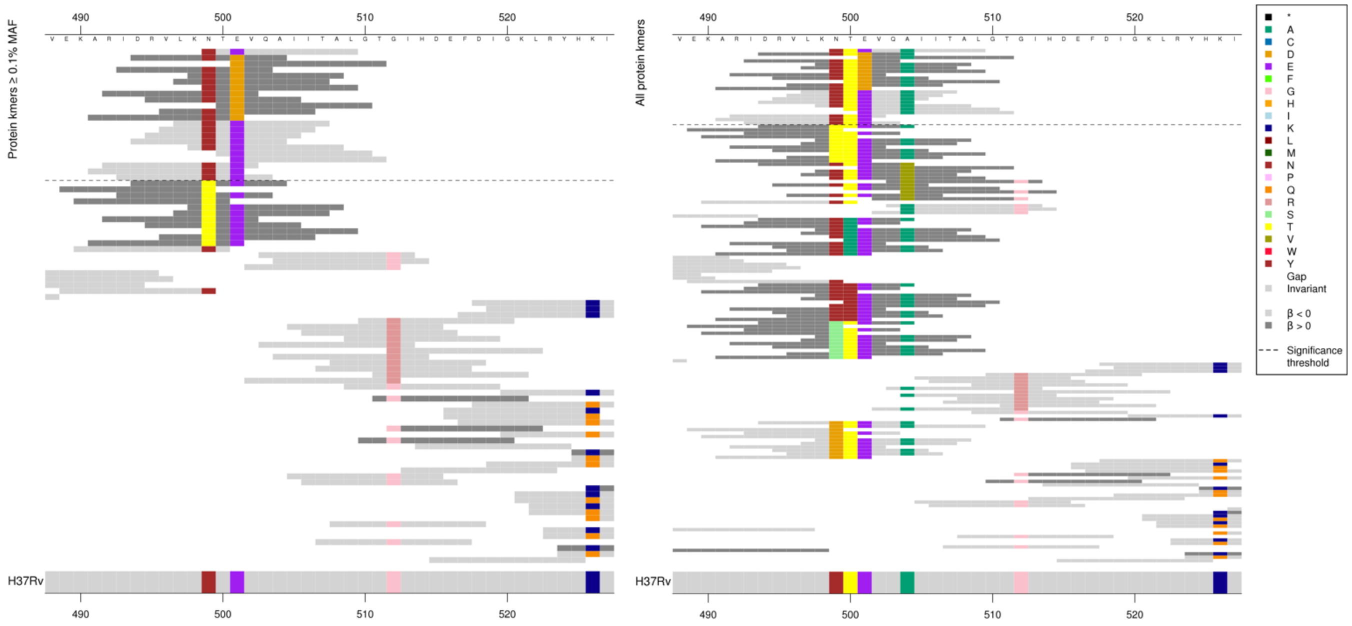

High MIC-associated kmers were all the same *p*-value and just captured 501D. Low MIC-associated kmers had different p-values, sometimes for the same combination of substitutions.

Kmers below significance threshold = 499T/S, 500A/N, 504V

| Kmer | -log$_{10}$ *p* | β | MAC | MAF | Ps | Variants |
|---|---|---|---|---|---|---|
| NTEVQAIITAL | 11.63 | -1.33 | 56 | 0.88 | 499 | 499N, 500T, 501E, 504A |
| DRVLKNTDVQA | 10.64 | 1.86 | 23 | 0.36 | 494 | 501D |
| TEVQAIITALG | 8.98 | -1.40 | 39 | 0.61 | 500 | 500T, 501E, 504A |
| EVQAIITALGT | 7.85 | -1.36 | 35 | 0.55 | 501 | 501E, 504A |
| RIDRVLKNTEV | 7.77 | -1.14 | 48 | 0.75 | 492 | 499N, 500T, 501E |

### *embB*

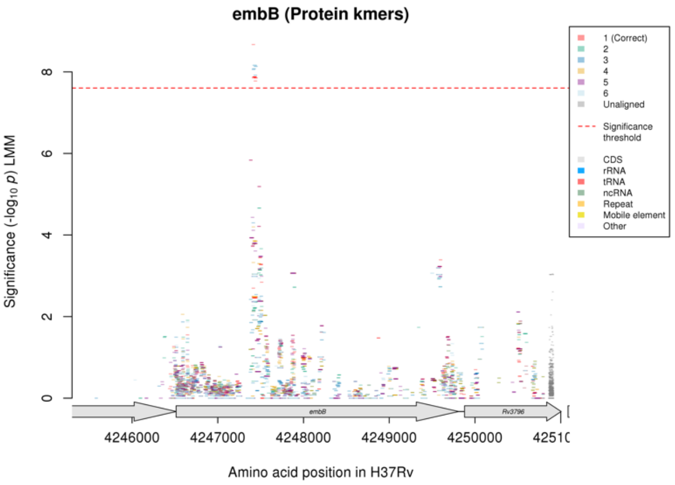

Not significant in the nucleotide kmer analysis.
Protein kmers:

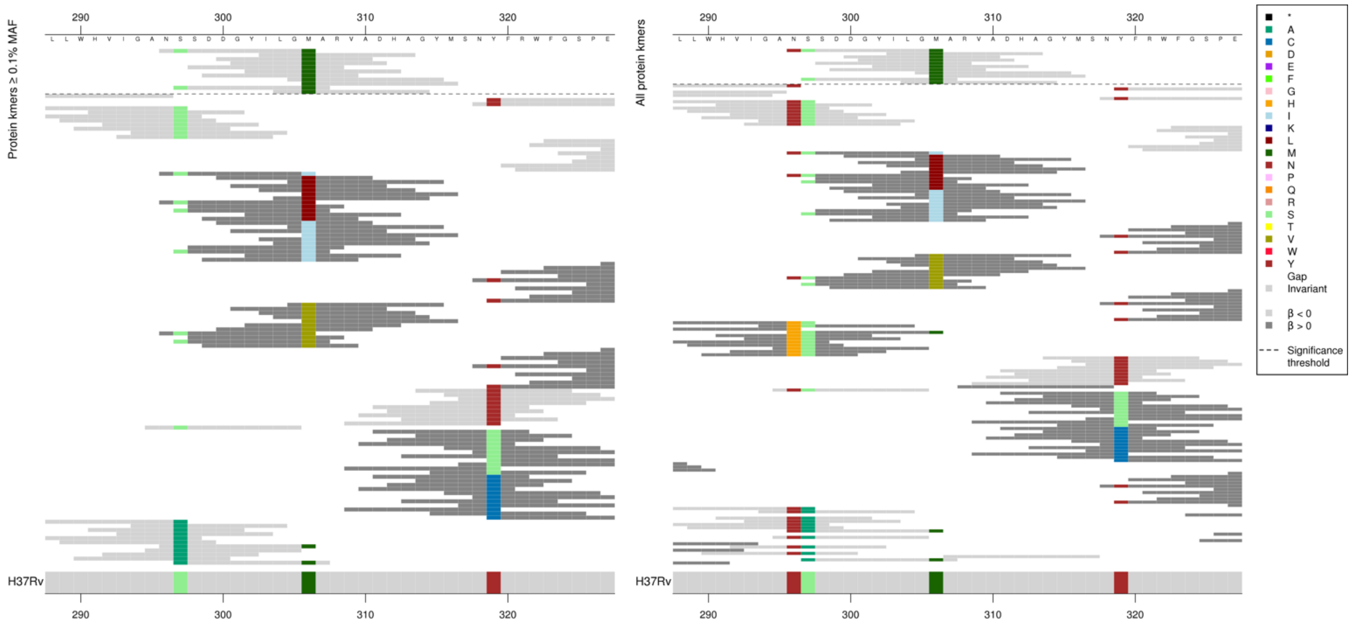

*P*-values varied for the low MIC-associated kmers, but mainly captured 306M so just showing the top p-value for 306M.

Kmers below significance threshold = 296H, 297A, 306M/V/I

| Kmer | -log$_{10}$ *p* | β | MAC | MAF | Ps | Variants |
|---|---|---|---|---|---|---|
| NSSDDGYILGM | 8.67 | -0.44 | 1339 | 20.96 | 296 | 306M |

### *katG*

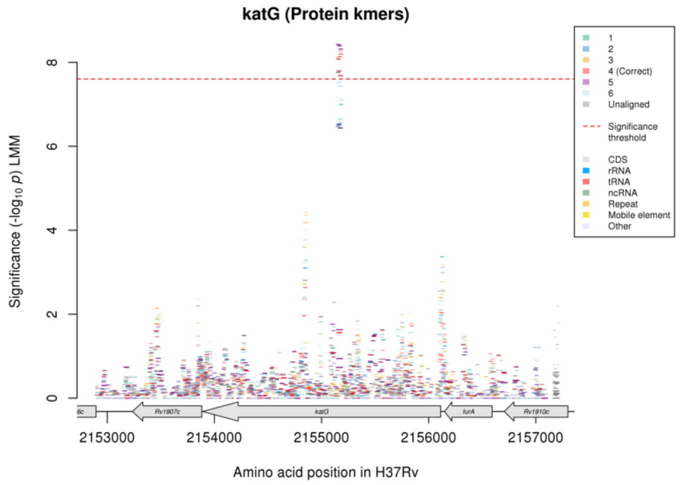

Not significant in the nucleotide kmer analysis.
Protein kmers:

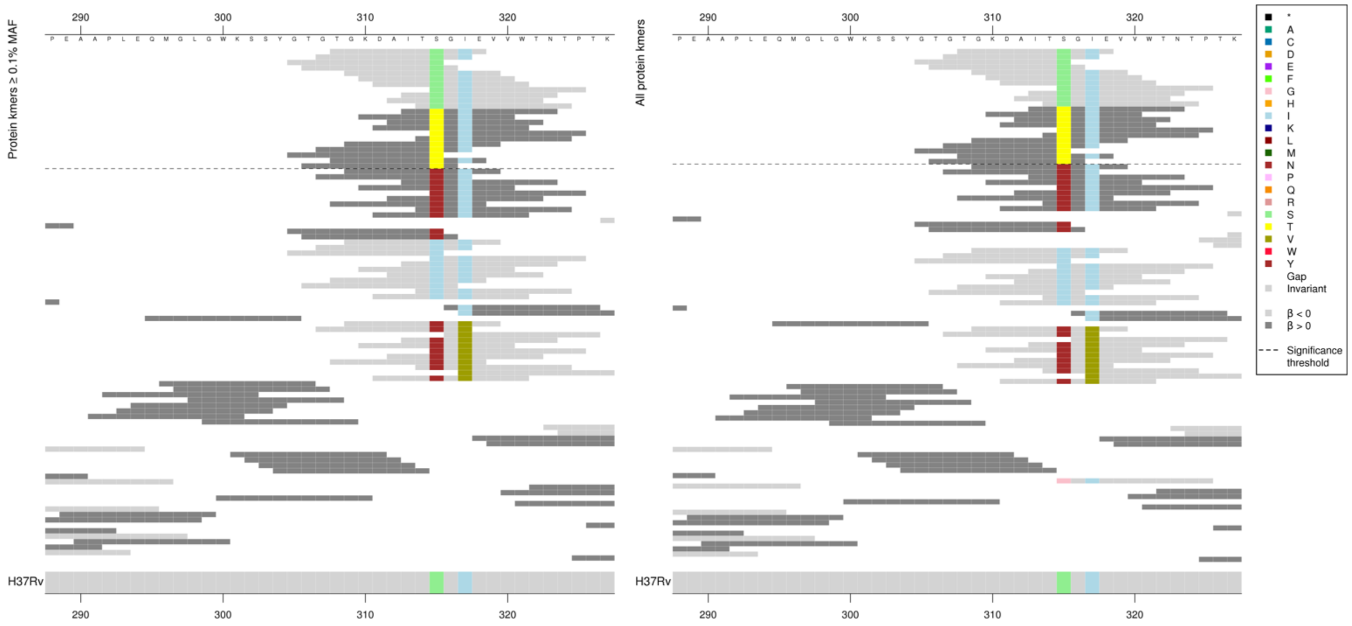

*P*-values varied for both low MIC-associated and high MIC-associated kmers.

| Kmer | $-\log_{10} p$ | β | MAC | MAF | Ps | Variants |
|---|---|---|---|---|---|---|
| TGKDAITSGIE | 8.27 | -0.45 | 2477 | 38.78 | 308 | 315S |
| ITTGIEVVWTN | 7.79 | 0.45 | 2409 | 37.71 | 313 | 315T |

### Rifabutin (RFB)

#### *rpoB*

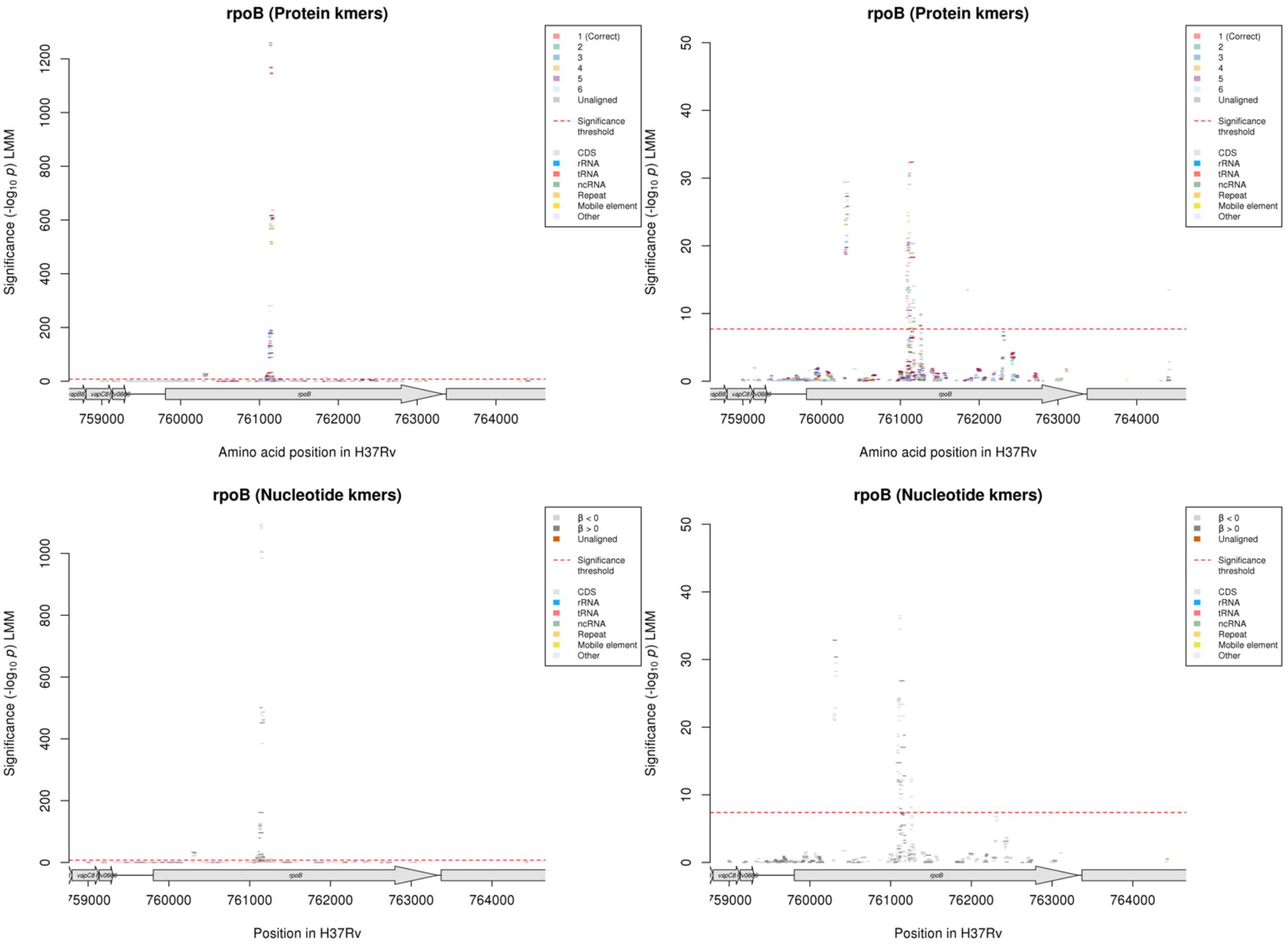

Protein kmers:

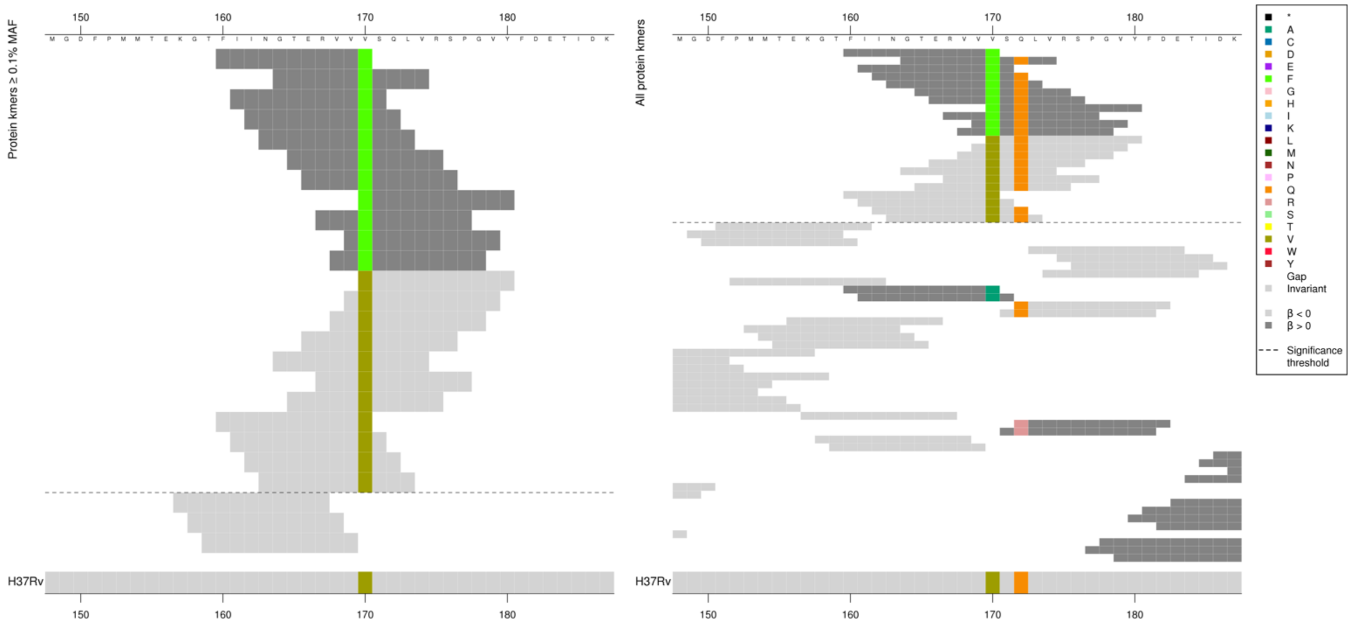

The last five high MIC-associated kmers were slightly less significant than the first six, but this does not correspond with whether they covered codon 172. Low MIC-associated kmers also covered a range of *p*-values but this doesn’t directly correspond to whether they covered codon 172 either.

Kmers below significance threshold = 170A, 172R

| Kmer | -log₁₀ *p* | β | MAC | MAF | Ps | Variants |
|---|---|---|---|---|---|---|
| FIINGTERVVF | 29.42 | 3.95 | 16 | 0.17 | 160 | 170F |
| VSQLVRSPGVY | 25.82 | -3.23 | 21 | 0.22 | 170 | 170V |

Protein kmers:

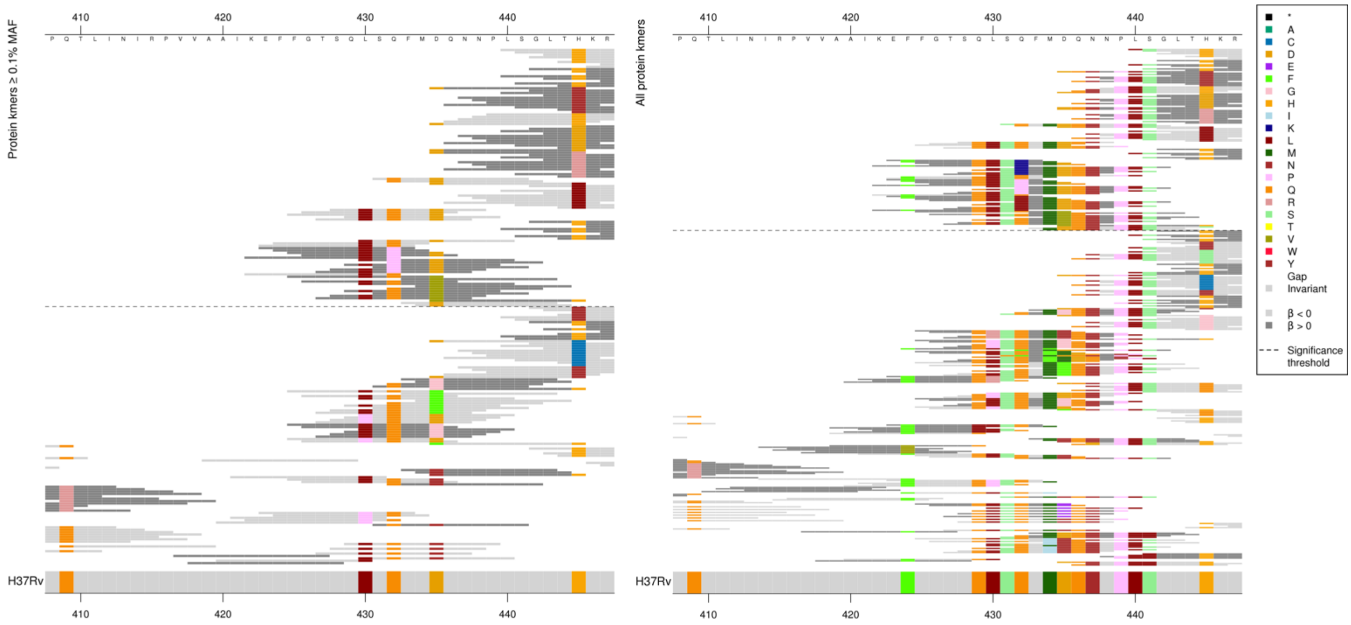

Main substitutions captured between kmers that were significant was at codons 432 and 435, but for the low MIC-associated kmers the *p*-values varied depending on other substitutions. For the high MIC-associated kmers, all 432K, 432P and 435V kmers were the same *p*-value, and all 432L kmers were the same *p*-value apart from the last kmer, so these do not appear to be as affected by the other substitutions.

Main significant substitutions:
Low MIC-associated kmers: 432Q, 435D
High MIC-associated kmers: 432K/P/L, 435V

Low MAF kmers = 432K, 432L, (435D, 436Q, 437N, 439P, 440L, 441S, 445S)

| Kmer | -log₁₀ *p* | β | MAC | MAF | Ps | Variants |
|---|---|---|---|---|---|---|
| SQFMDQNNPLS | 30.32 | -1.17 | 577 | 6.13 | 431 | 431S, 432Q, 434M, 435D, 436Q, 437N, 439P, 440L, 441S |
| QFMDQNNPLSG | 29.08 | -1.15 | 576 | 6.12 | 432 | 432Q, 434M, 435D, 436Q, 437N, 439P, 440L, 441S |
| SQLSQFMDQNN | 20.27 | -0.86 | 665 | 7.06 | 428 | 429Q, 430L, 431S, 432Q, 434M, 435D, 436Q, 437N |
| GTSQLSQFMDQ | 20.13 | -0.86 | 663 | 7.04 | 426 | 429Q, 430L, 431S, 432Q, 434M, 435D, 436Q |
| FGTSQLSQFMD | 20.11 | -0.86 | 661 | 7.02 | 425 | 429Q, 430L, 431S, 432Q, 434M, 435D |
| QLSQFMDQNNP | 19.35 | -0.84 | 662 | 7.03 | 429 | 429Q, 430L, 431S, 432Q, 434M, 435D, 436Q, 437N, 439P |
| TSQLSQFMDQN | 19.35 | -0.84 | 665 | 7.06 | 427 | 429Q, 430L, 431S, 432Q, 434M, 435D, 436Q, 437N |
| LSQFMDQNNPL | 17.49 | -0.80 | 656 | 6.97 | 430 | 430L, 431S, 432Q, 434M, 435D, 436Q, 437N, 439P, 440L |
| FFGTSQLSQFM | 16.20 | -1.01 | 177 | 1.88 | 424 | 424F, 429Q, 430L, 431S, 432Q, 434M |
| EFFGTSQLSQF | 14.74 | -0.98 | 164 | 1.74 | 423 | 424F, 429Q, 430L, 431S, 432Q |
| EFFGTSQLSPF | 13.47 | 3.79 | 12 | 0.13 | 423 | 432P |
| FGTSQLSQFMV | 10.47 | 0.92 | 368 | 3.91 | 425 | 435V |
| FMDQNNPLSGL | 9.65 | -0.66 | 542 | 5.75 | 433 | 434M, 435D, 436Q, 437N, 439P, 440L, 441S |

Protein kmers:

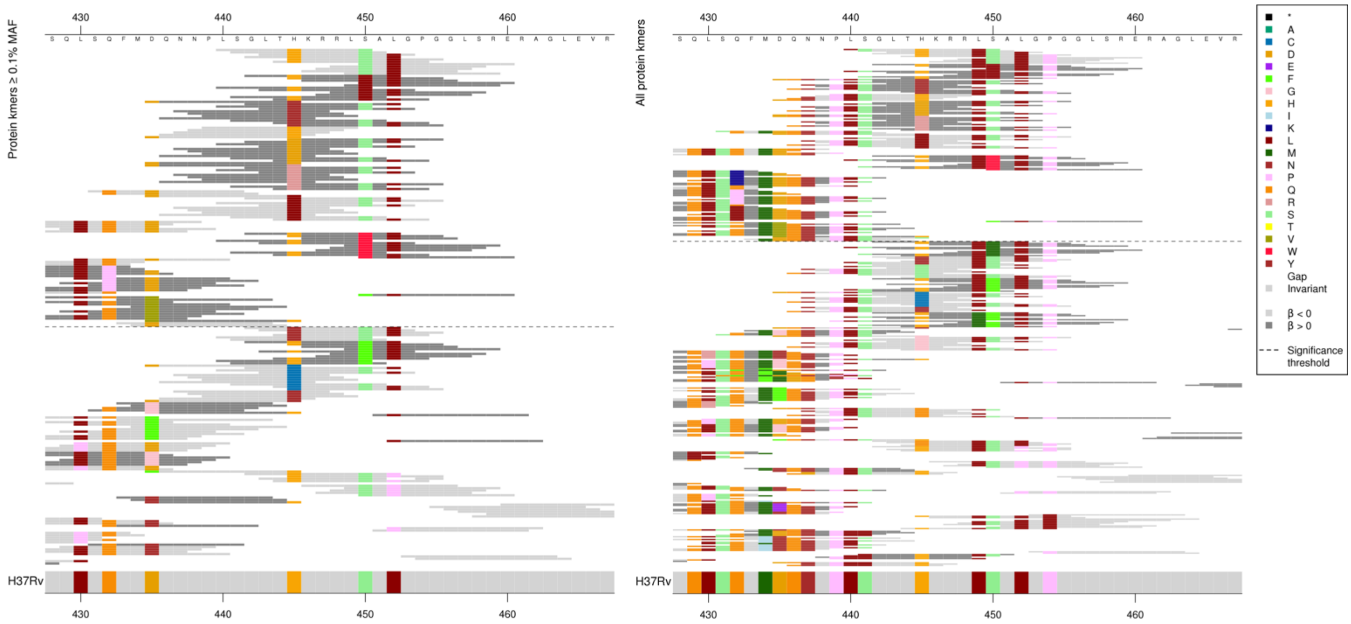

The main codons being captured were 445 and 450, but the *p*-values of the low MIC-associated kmer varied depending on the codons covered (apart from those capturing 445L), but the high MIC-associated kmers for each substitution captured didn't vary depending on other codons in the same way, so just showing the top kmer for each of them. Kmers capturing 450L, 445Y, 445R, 445L, 450W were all the same p-value. First and last kmer for 445D were slightly different p-values, MAC range from 93-95.

| Kmer | $-\log_{10} p$ | β | MAC | MAF | Ps | Variants |
|---|---|---|---|---|---|---|
| LSGLTHKRRLS | 1257.54 | -4.73 | 2683 | 28.49 | 440 | 440L, 441S, 445H, 449L, 450S |
| SGLTHKRRLSA | 1249.39 | -4.73 | 2686 | 28.52 | 441 | 441S, 445H, 449L, 450S |
| LTHKRRLSALG | 1167.73 | -4.62 | 2763 | 29.34 | 443 | 445H, 449L, 450S, 452L |
| HKRRLSALGPG | 1144.70 | -4.57 | 2771 | 29.42 | 445 | 445H, 449L, 450S, 452L, 454P |
| SALGPGGLSRE | 608.97 | -4.04 | 2411 | 25.60 | 450 | 450S, 452L, 454P |
| RLSALGPGGLS | 606.78 | -4.05 | 2412 | 25.61 | 448 | 449L, 450S, 452L, 454P |
| GLTHKRRLLAL | 567.53 | 4.07 | 2235 | 23.73 | 442 | 450L |
| DQNNPLSGLTY | 177.97 | 3.97 | 130 | 1.38 | 435 | 445Y |
| PLSGLTHKRRL | 148.71 | -2.44 | 383 | 4.07 | 439 | 439P, 440L, 441S, 445H, 449L |
| NPLSGLTHKRR | 141.53 | -2.41 | 376 | 3.99 | 438 | 439P, 440L, 441S, 445H |
| NNPLSGLTHKR | 131.13 | -2.29 | 384 | 4.08 | 437 | 437N, 439P, 440L, 441S, 445H |
| QNNPLSGLTHK | 131.00 | -2.29 | 384 | 4.08 | 436 | 436Q, 437N, 439P, 440L, 441S, 445H |
| DQNNPLSGLTH | 127.36 | -1.81 | 890 | 9.45 | 435 | 435D, 436Q, 437N, 439P, 440L, 441S, 445H |
| DKRRLSALGPG | 105.49 | 3.61 | 95 | 1.01 | 445 | 445D |
| DQNNPLSGLTR | 32.37 | 3.26 | 32 | 0.34 | 435 | 445R |
| DQNNPLSGLTL | 20.30 | -2.14 | 47 | 0.50 | 435 | 445L |
| GLTHKRRLWAL | 18.30 | 1.95 | 44 | 0.47 | 442 | 450W |
| FALGPGGLSRE | 10.88 | 1.68 | 37 | 0.39 | 450 | 450F, 452L, 454P |

Protein kmers:

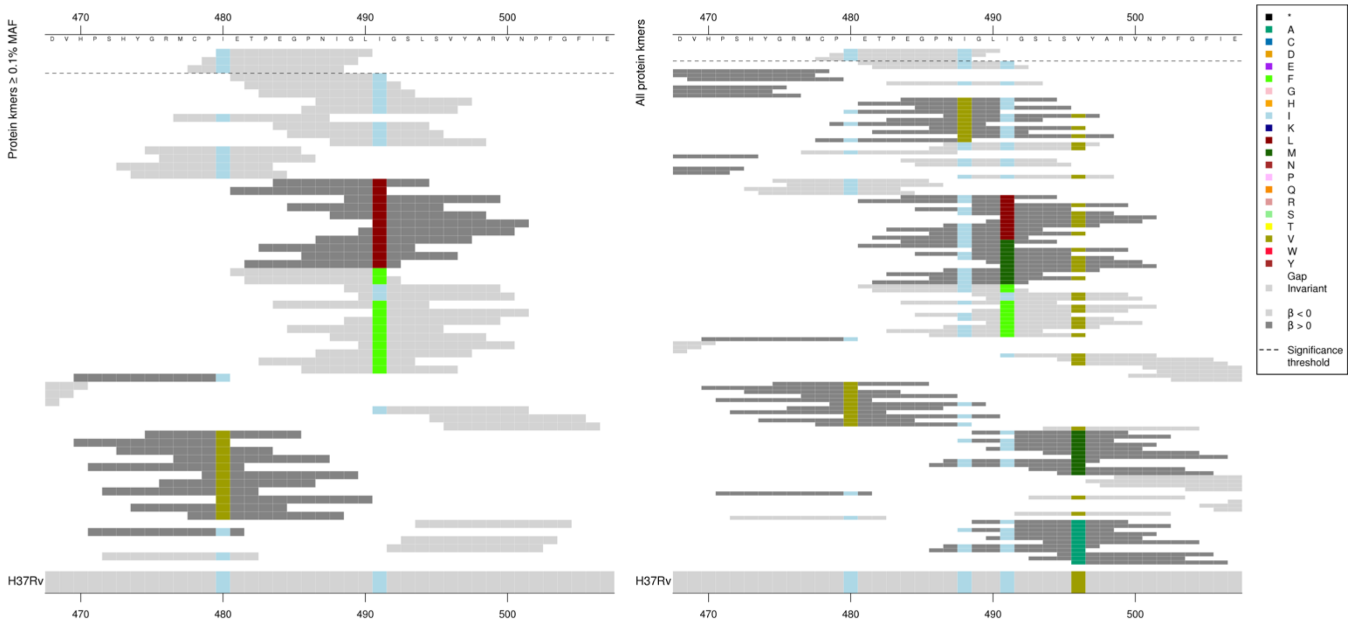

For RIF, the significant low MIC-associated kmers covered and included codon 480I, 488I and 491I, however the significant kmers here for RFB did not include codon 491I.

Kmers below significance threshold = 480V, 488V, 491L/M/F

| Kmer | $-\log_{10} p$ | β | MAC | MAF | Ps | Variants |
|---|---|---|---|---|---|---|
| IETPEGPNIGL | 9.71 | -1.73 | 29 | 0.31 | 480 | 480I, 488I |

#### *embB*

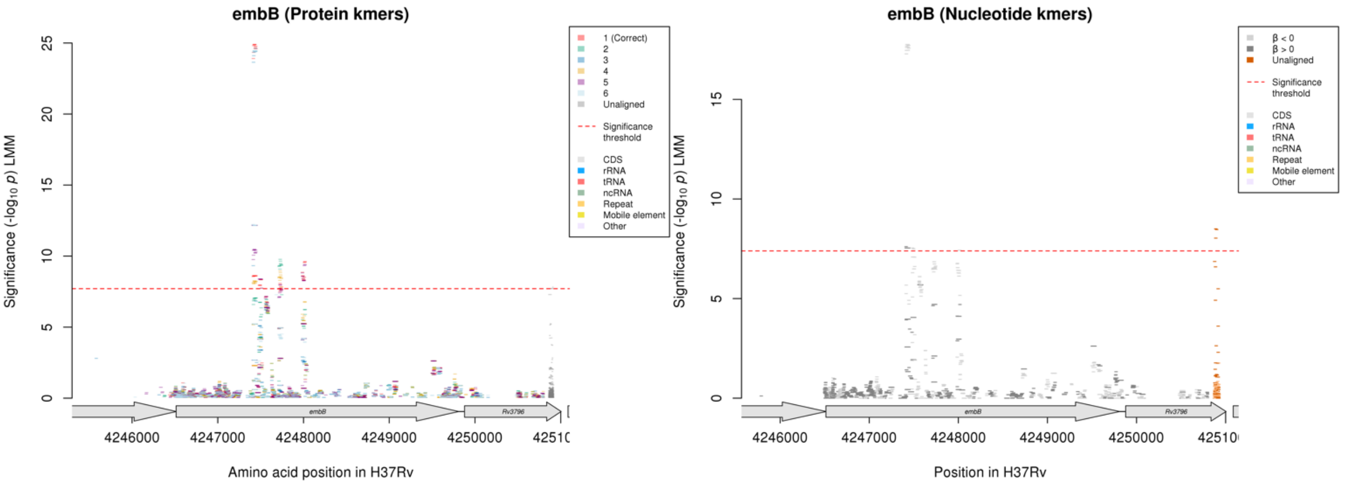

Protein kmers:

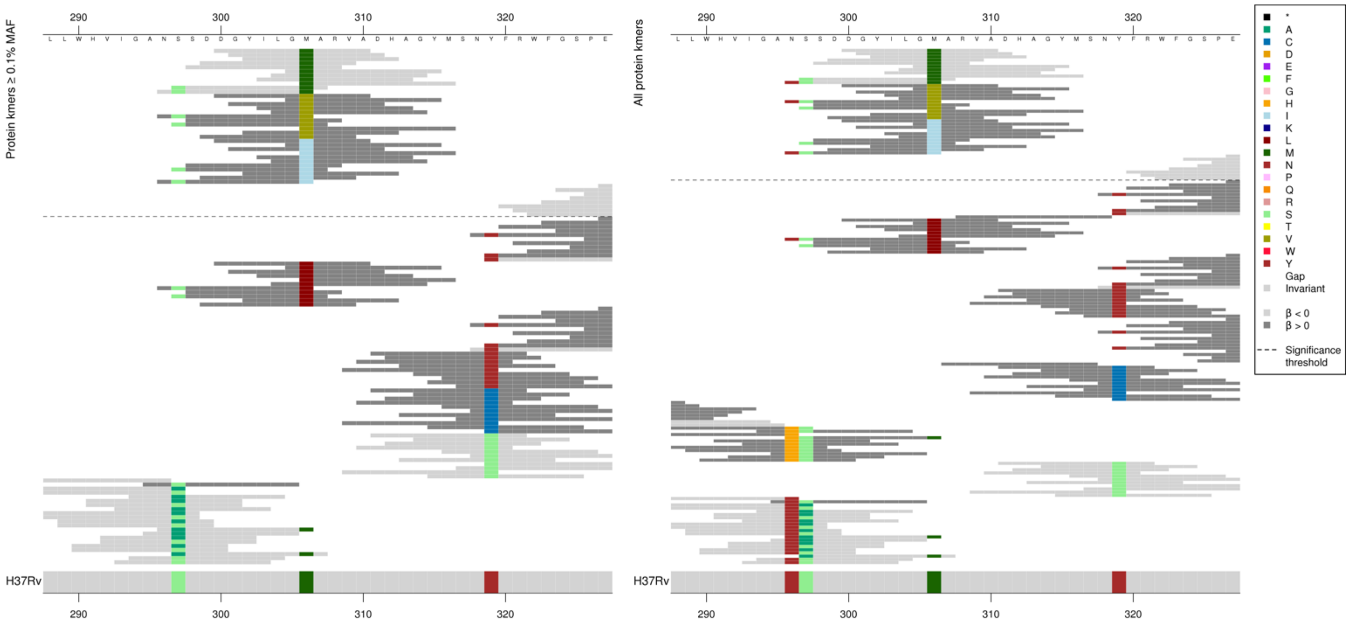

P-values varied slightly for all variants.

| Kmer | -log₁₀ p | β | MAC | MAF | Ps | Variants |
|---|---|---|---|---|---|---|
| DGYILGMARVA | 24.89 | -0.90 | 1679 | 17.83 | 300 | 306M |
| DGYILGVARVA | 12.17 | 0.73 | 911 | 9.67 | 300 | 306V |
| DGYILGIARVA | 8.61 | 0.61 | 737 | 7.83 | 300 | 306I |

Protein kmers:

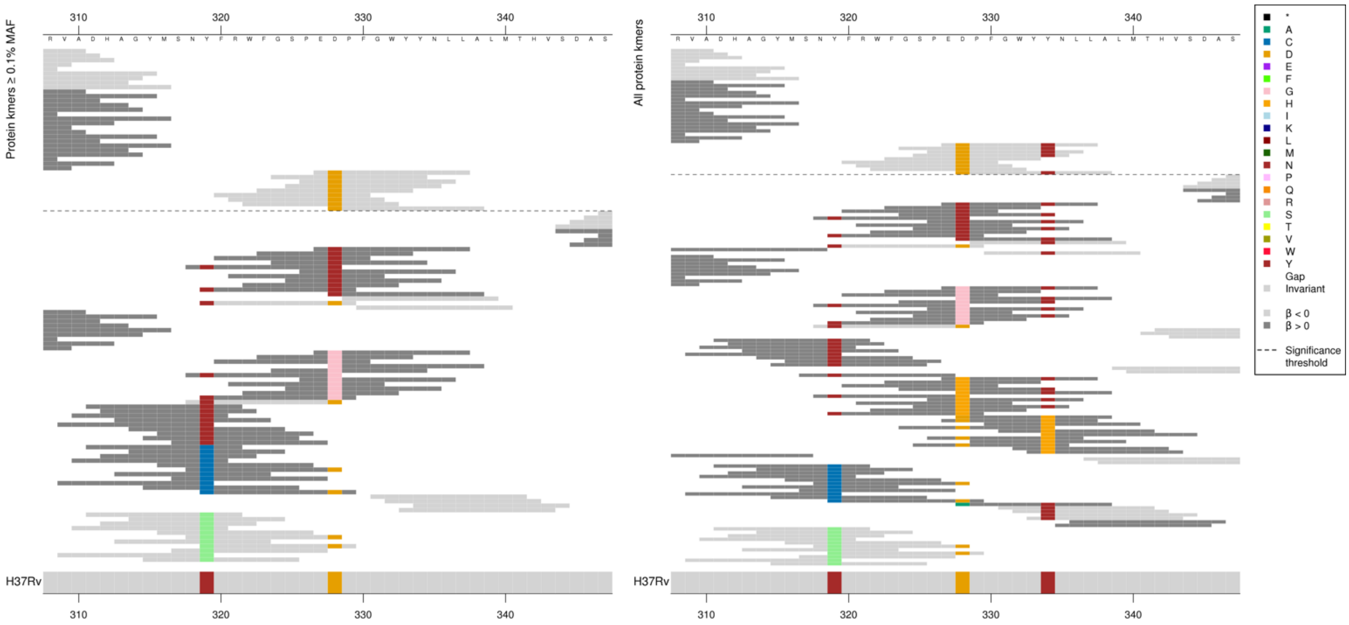

| Kmer | -log₁₀ p | β | MAC | MAF | Ps | Variants |
|---|---|---|---|---|---|---|
| EDPFGWYYNLL | 8.34 | -1.27 | 48 | 0.51 | 327 | 328D, 334Y |
| FGSPEDPFGWY | 7.95 | -1.26 | 42 | 0.45 | 323 | 328D |

#### Protein kmers:

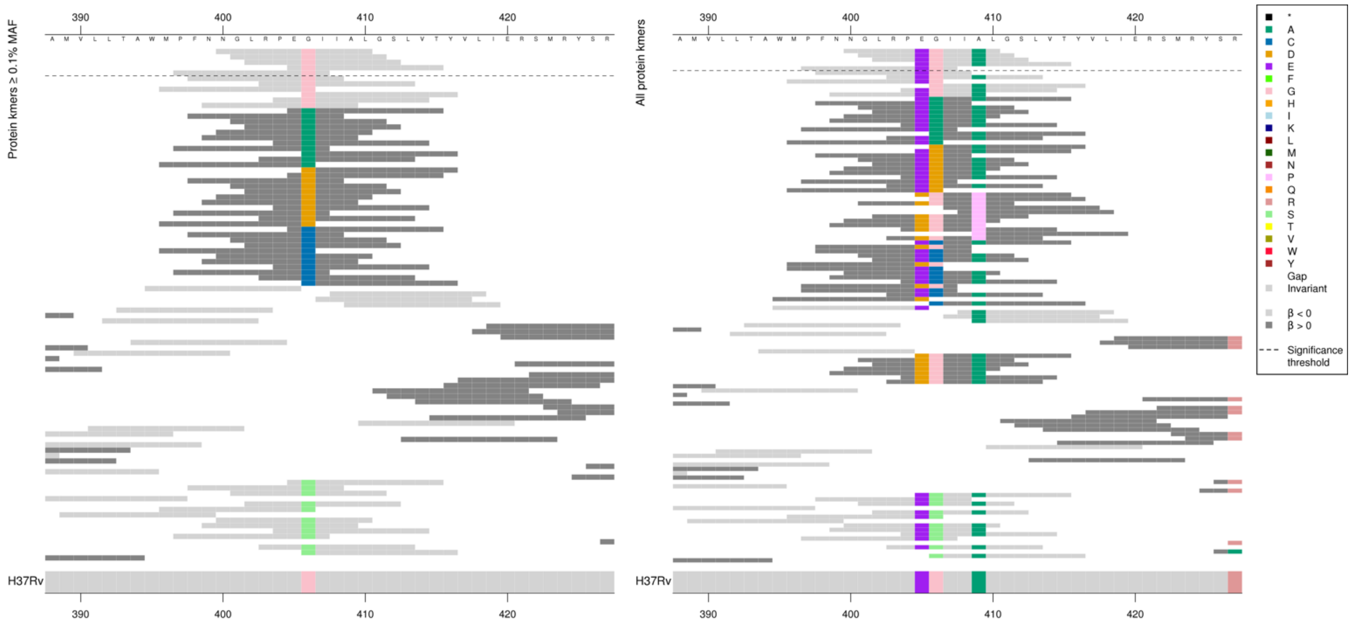

| Kmer | -log$_{10}$ *p* | β | MAC | MAF | Ps | Variants |
|---|---|---|---|---|---|---|
| NGLRPEGIIAL | 8.08 | -0.62 | 502 | 5.33 | 400 | 405E, 406G, 409A |
| PFNNGLRPEGI | 7.72 | -0.61 | 499 | 5.30 | 397 | 405E, 406G |

#### Protein kmers:

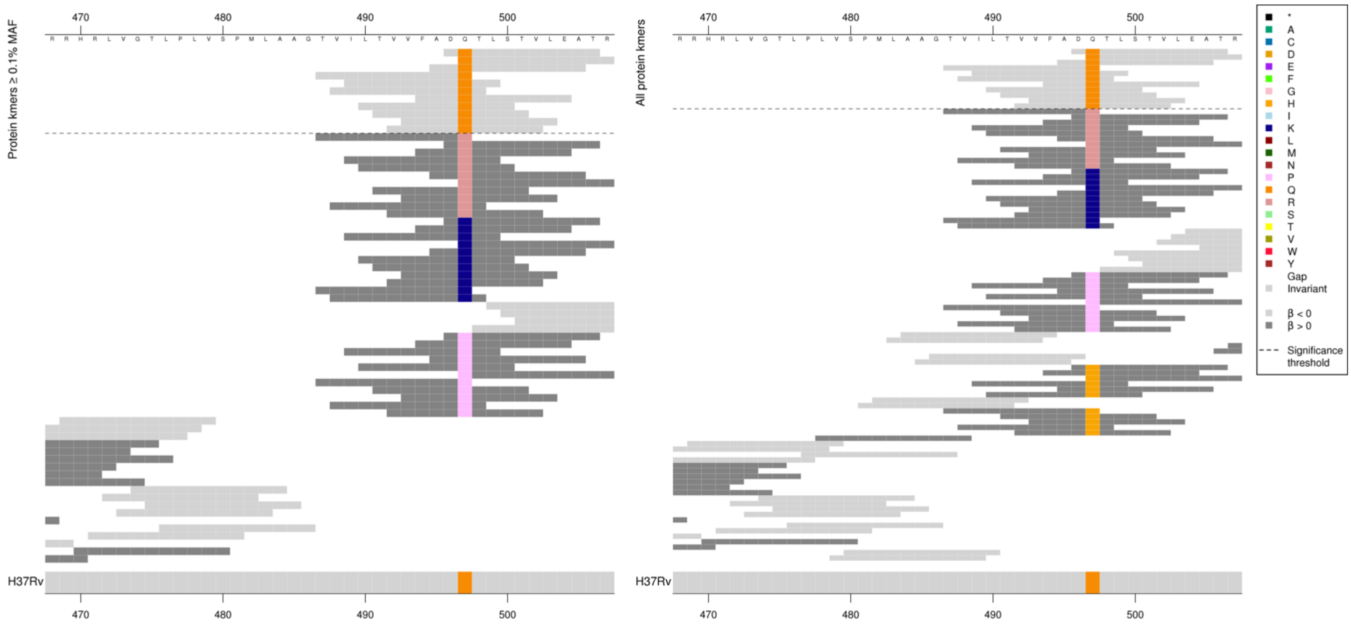

| Kmer | -log$_{10}$ *p* | β | MAC | MAF | Ps | Variants |
|---|---|---|---|---|---|---|
| DQTLSTVLEAT | 9.60 | -0.75 | 358 | 3.80 | 496 | 497Q |

### *katG*

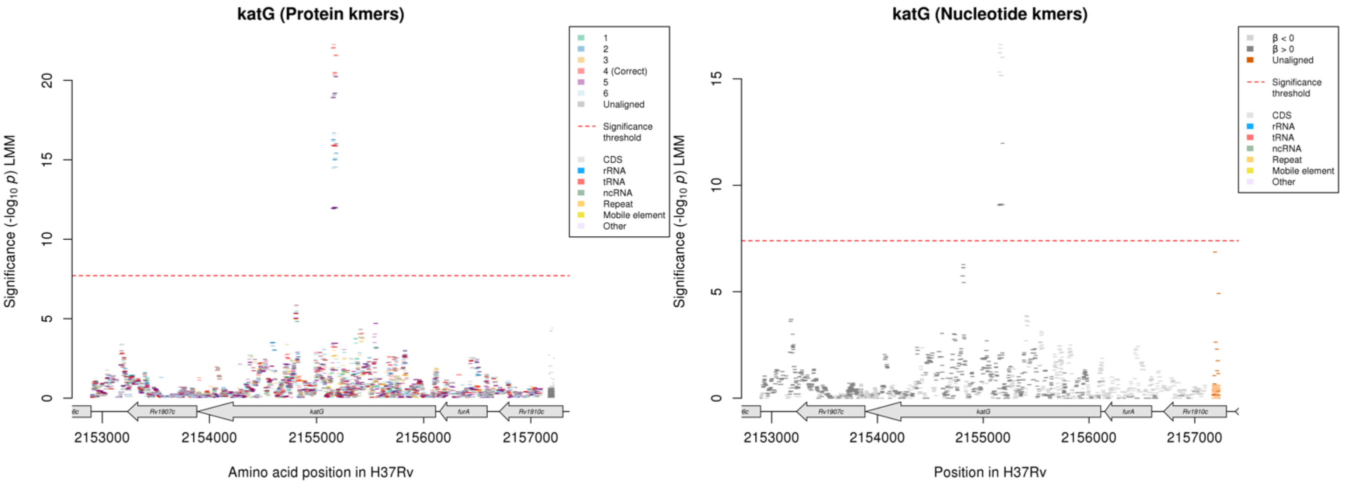

Protein kmers:

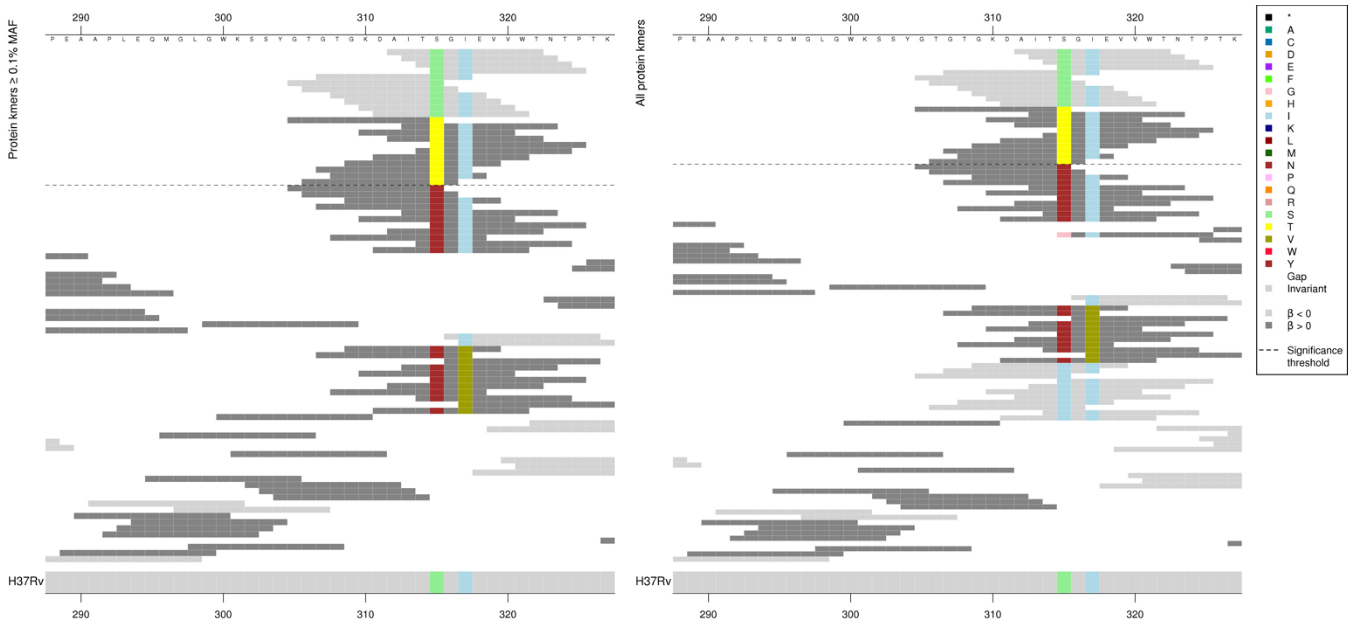

*P*-values varied for both low MIC-associated and high MIC-associated kmers, but not in line with whether they covered codon 317.

| Kmer | $-\log_{10} p$ | β | MAC | MAF | Ps | Variants |
|---|---|---|---|---|---|---|
| AITSGIEVVWT | 22.25 | -0.94 | 3324 | 35.29 | 312 | 315S |
| GTGTGKDAITT | 16.01 | 0.84 | 3233 | 34.33 | 305 | 315T |

#### *rpoC*

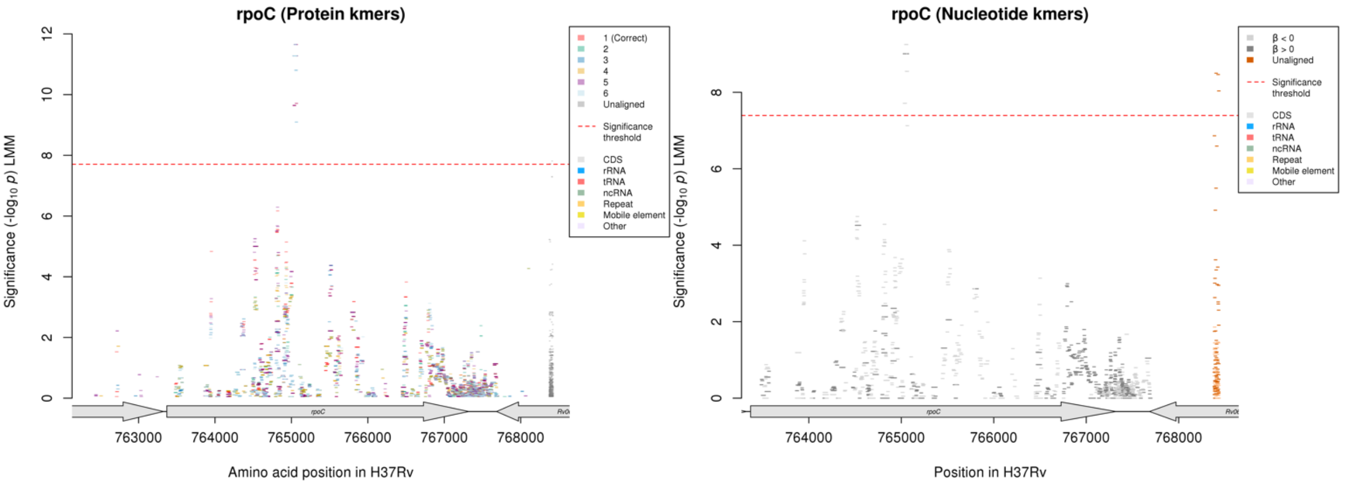

Protein kmers:

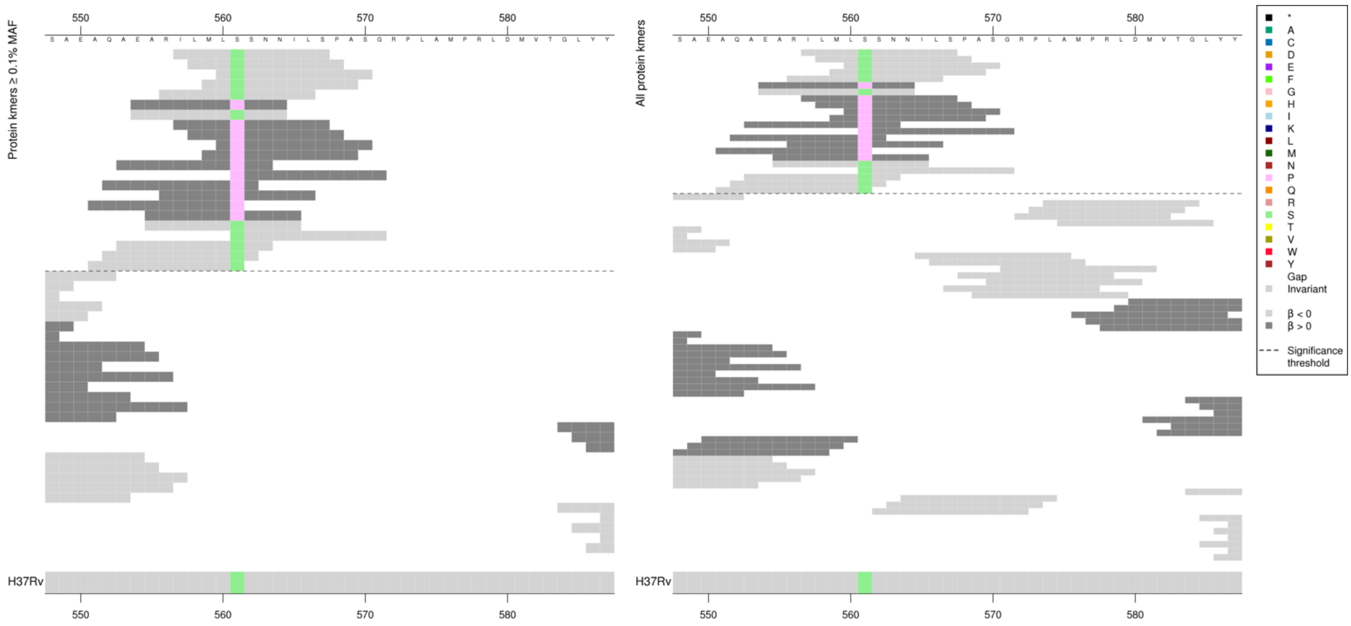

| Kmer | $-\log_{10} p$ | β | MAC | MAF | Ps | Variants |
|---|---|---|---|---|---|---|
| ILMLSSNNILS | 11.65 | -3.10 | 13 | 0.14 | 557 | 561S |
| EARILMLPSNN | 11.28 | 3.13 | 12 | 0.13 | 554 | 561P |

## *Rv0810c*

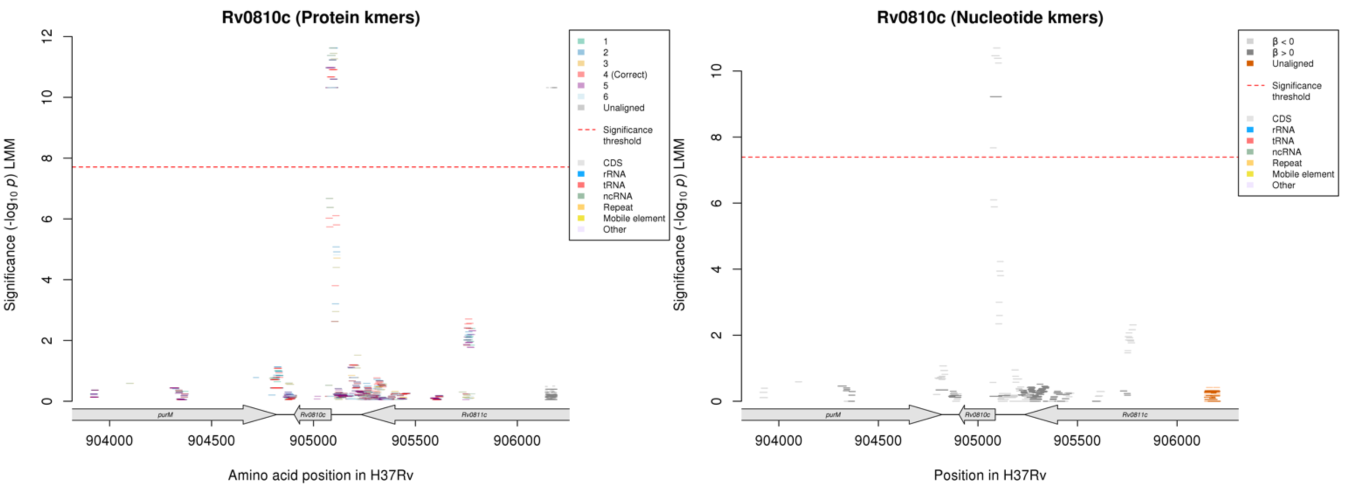

Nucleotide kmers:

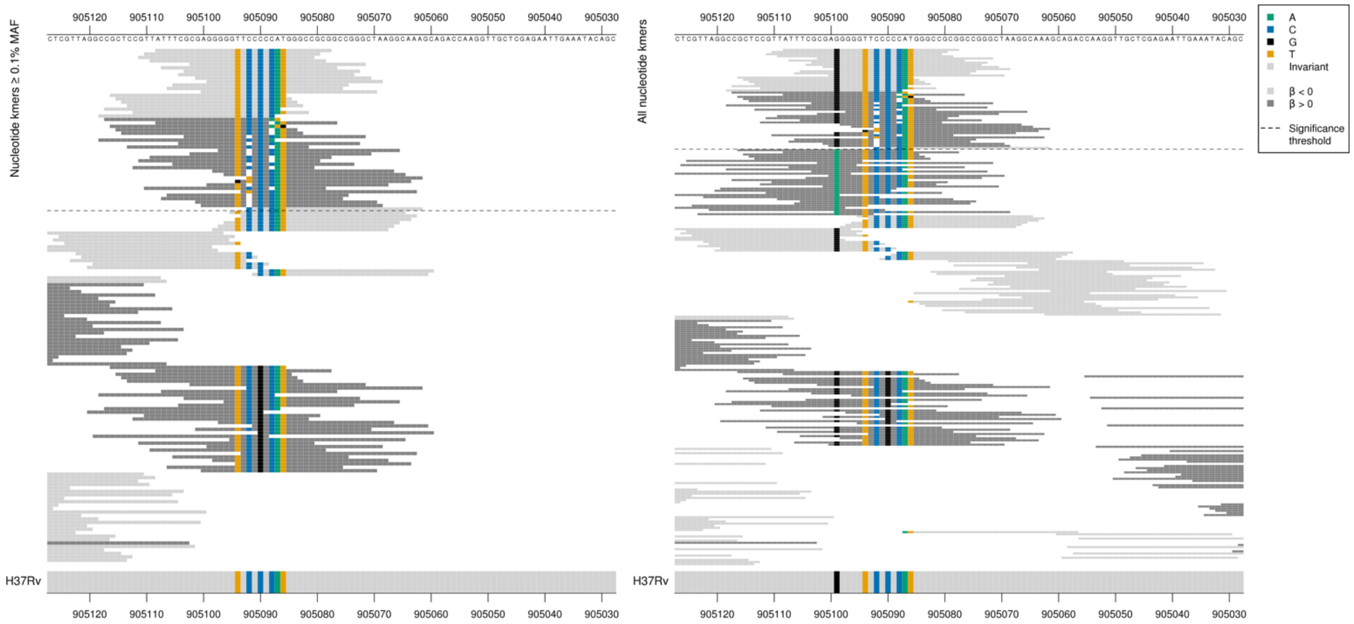

*Rv0810c* begins at position 905087.
High MIC-associated kmers = C deletion in the ‘CCCCC’ region positions -1 to -5 (905088-905092) directly upstream of the start codon
Kmers below significance threshold = 905099/-12 A

All high MIC-associated kmers had the same *p*-values, all capturing the C deletion. Low MIC-associated kmers varied depending on the positions covered, showing the top kmer.

| Kmer | -log₁₀ *p* | β | MAC | MAF | Ps | Variants |
|---|---|---|---|---|---|---|
| ATTTCGCGAGGGGGTTCCCCCATGGCCGCG | 10.70 | -1.54 | 54 | 0.57 | 905108 | WT |
| CGCTCCGTTATTTCGCGAGGGGGTTCCCCAT | 9.22 | 2.11 | 31 | 0.33 | 905117 | C deletion between 905088-905092/-1 to -5 |

## *Rv2478c:Rv2481c* R

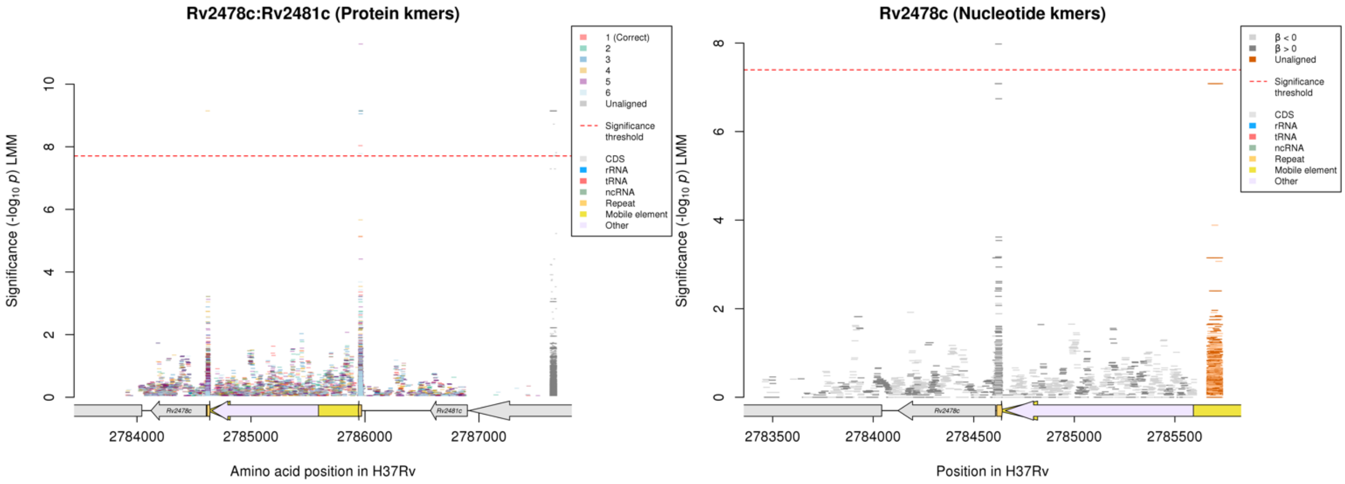

Nucleotide kmers:

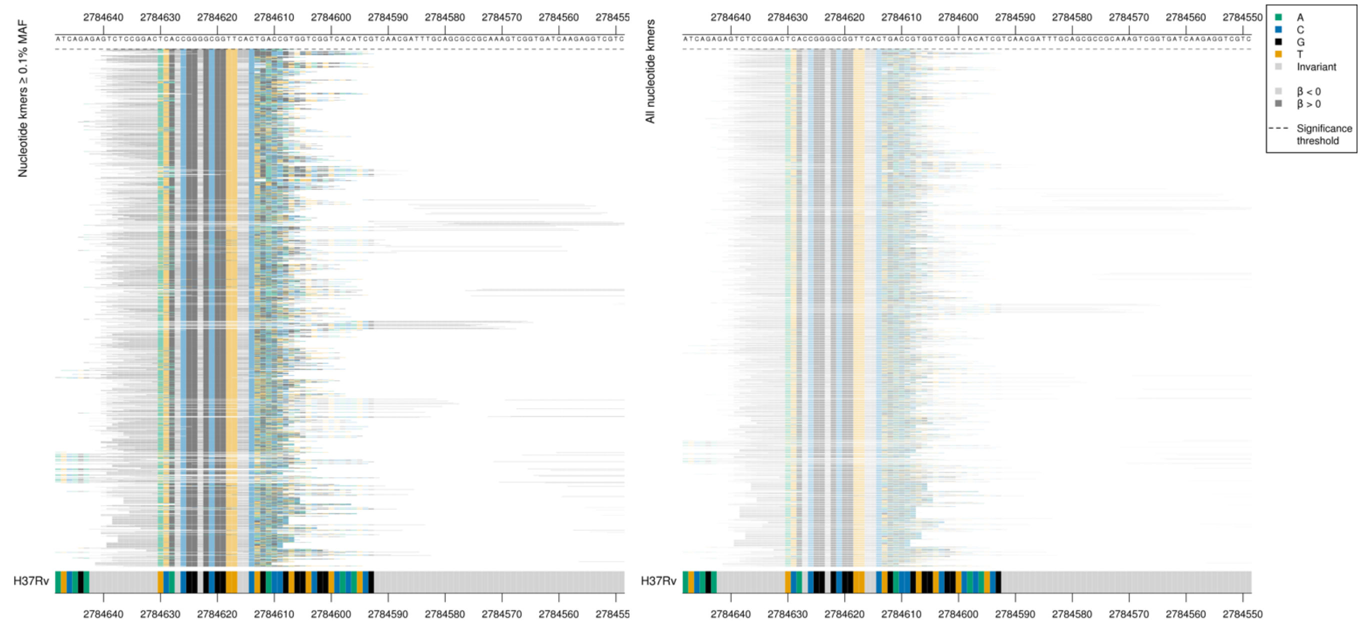

Repeat region at the end of a mobile element

| Kmer | $-\log_{10} p$ | β | MAC | MAF | Ps | Variants |
|---|---|---|---|---|---|---|
| CTCCGGACATGCCGGGGCGGTTCACGACGAC | 7.98 | 2.33 | 16 | 0.17 | 2784638 | Repeat region |

## *Rv2647:Rv2650c* [R]

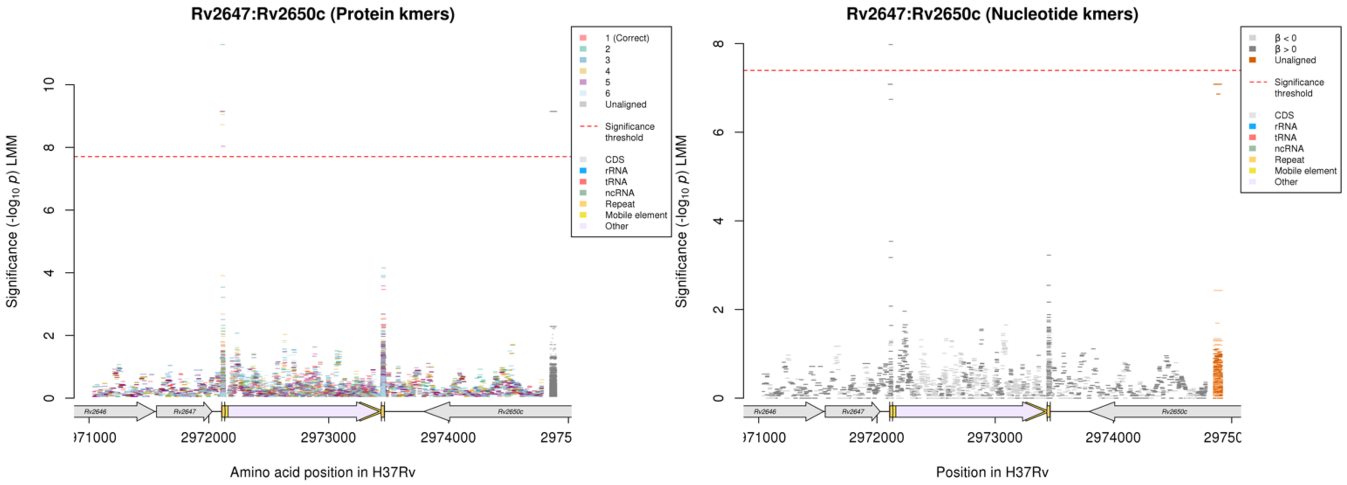

Nucleotide kmers:

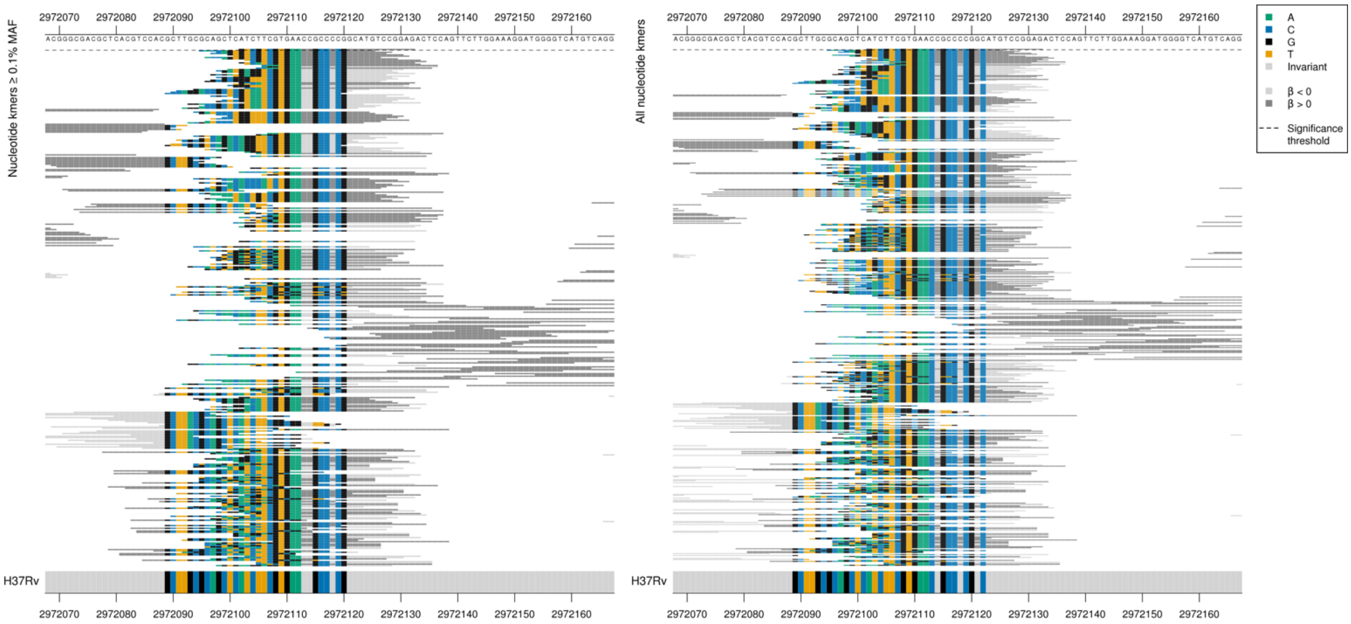

Repeat region at the end of a mobile element.
Just one significant nucleotide kmer, with two differences to the reference.

| Kmer | $-\log_{10} p$ | β | MAC | MAF | Ps | Variants |
|---|---|---|---|---|---|---|
| CTCCGGACATGCCGGGGCGGTTCACGACGAC | 7.98 | 2.33 | 16 | 0.17 | 2972102 | Repeat region |

#### *rplP*

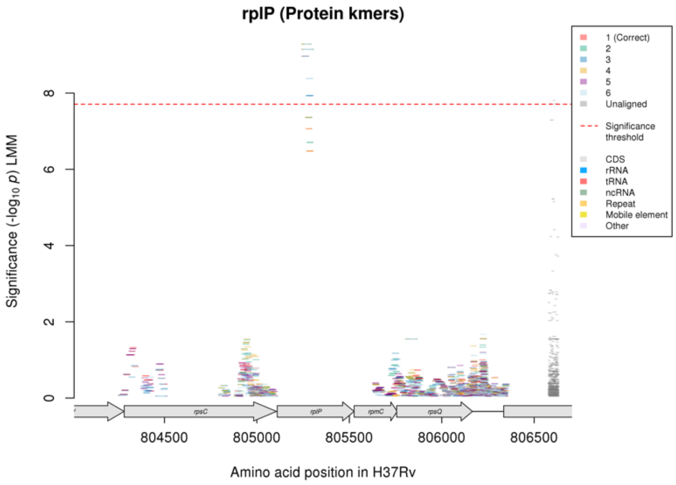

Not significant in nucleotide kmer analysis.

Protein kmers:

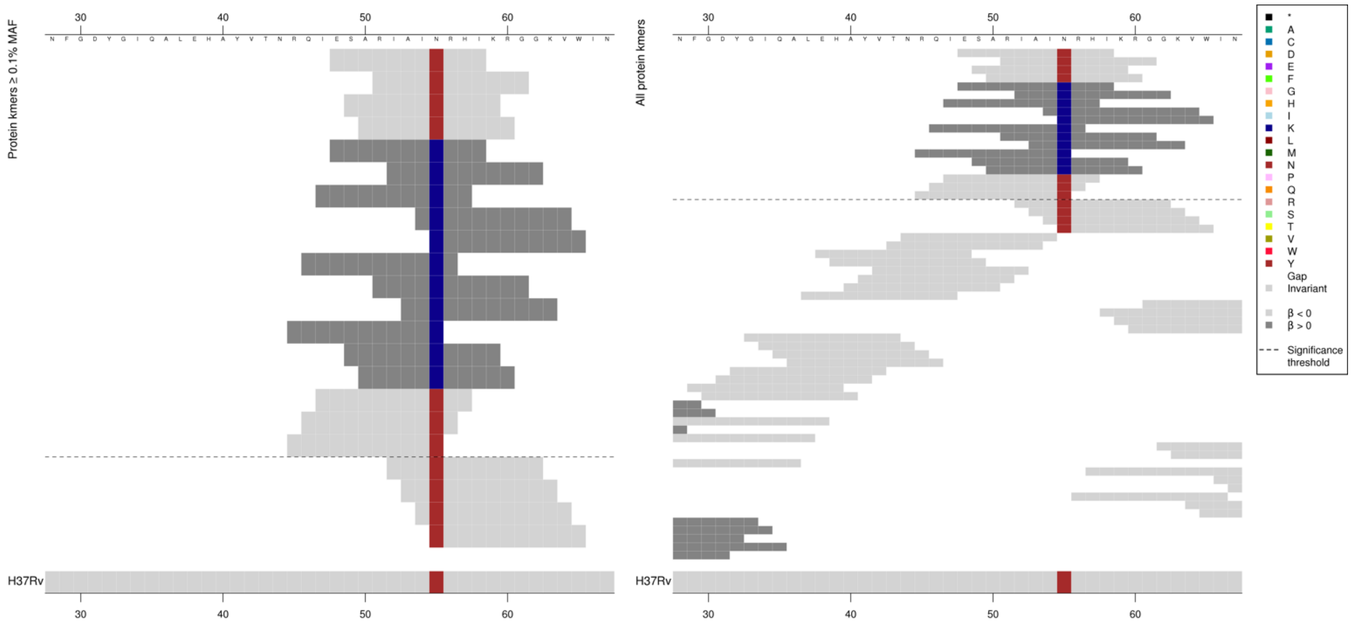

| Kmer | $-\log_{10} p$ | β | MAC | MAF | Ps | Variants |
|---|---|---|---|---|---|---|
| ESARIAINRHI | 9.28 | -3.30 | 11 | 0.12 | 48 | 55N |
| ESARIAIKRHI | 9.15 | 3.29 | 10 | 0.11 | 48 | 55K |

# *Rv2797c*

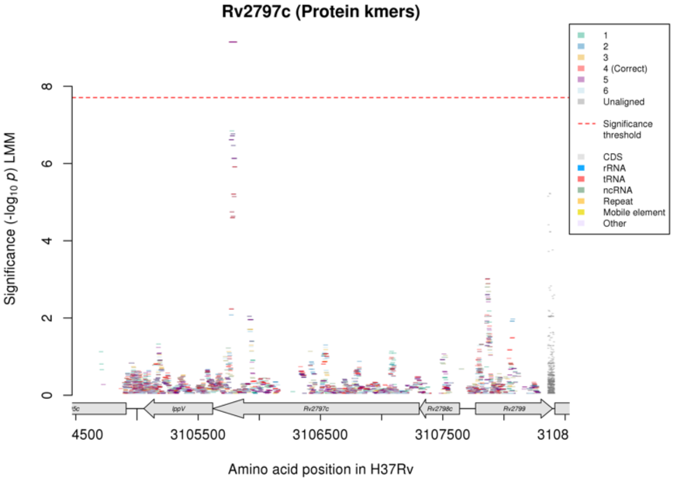

Not significant in the nucleotide kmer analysis.

Protein kmers:

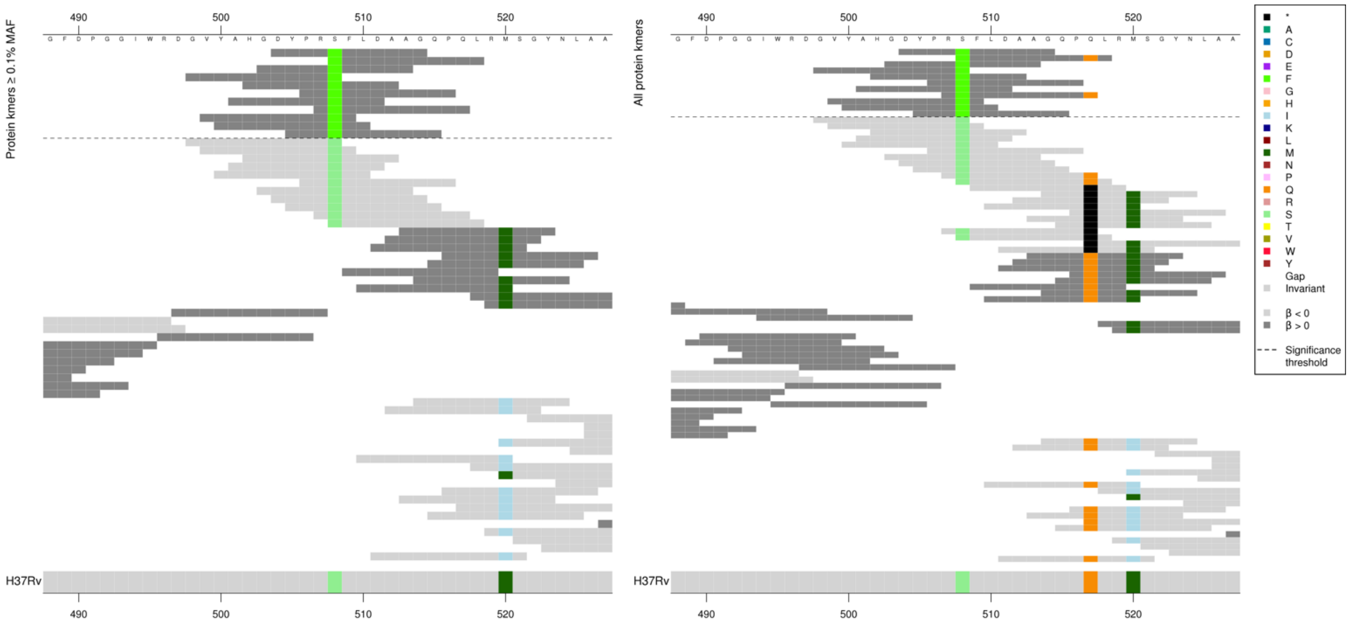

Kmers below significance threshold = 508S

| Kmer | $-\log_{10} p$ | β | MAC | MAF | Ps | Variants |
|---|---|---|---|---|---|---|
| DYPRFFLDAAG | 9.15 | 3.29 | 10 | 0.11 | 504 | 508F |

#### *cpsY*

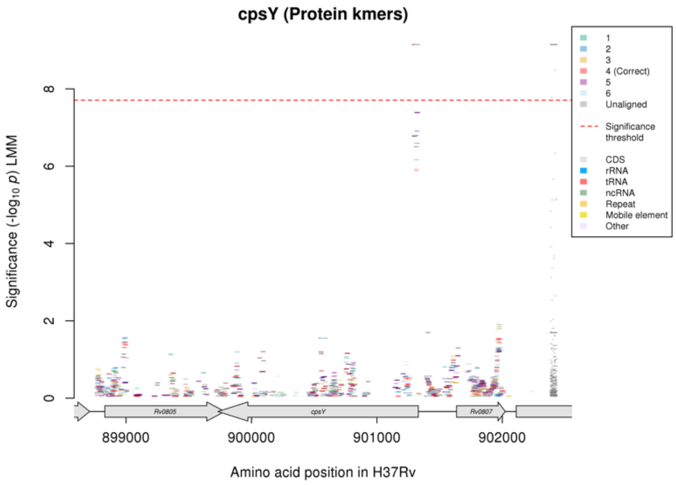

Not significant in the nucleotide kmer analysis.

Protein kmers:

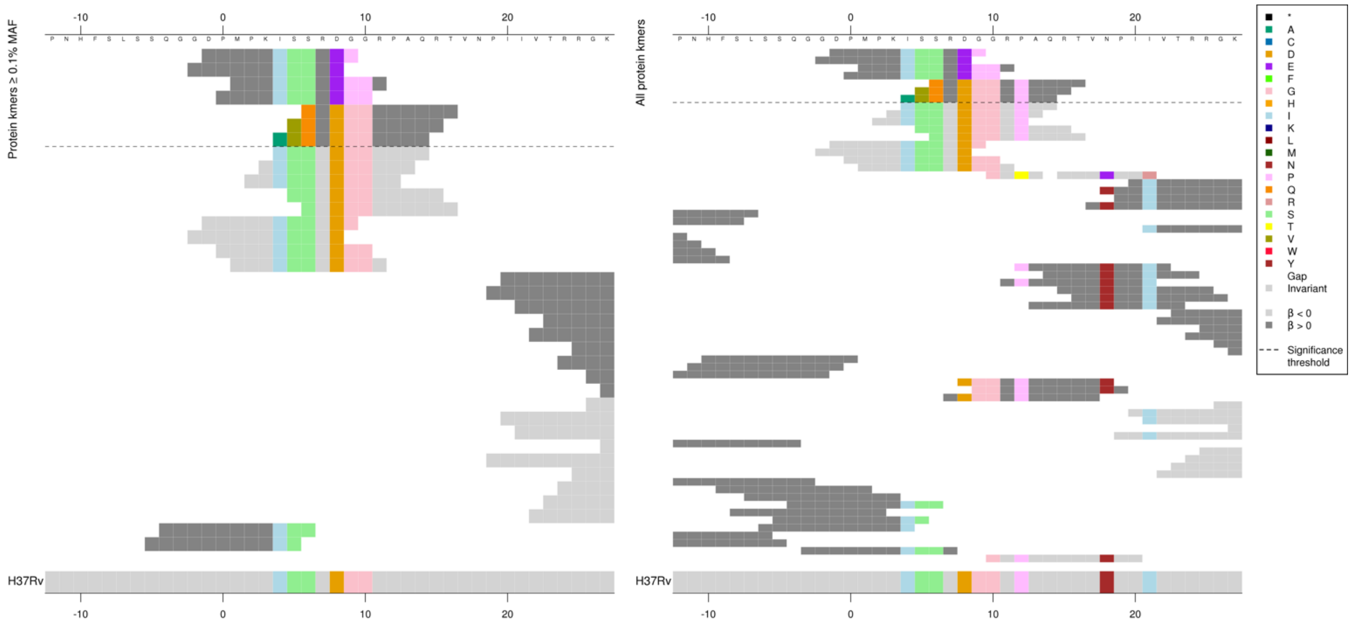

The kmers were identical to the reference for most of the kmer, then there were differences which begin within the CGCGCG repeat region. It is not clear if the differences were due to deletions or insertions.

| Kmer | $-\log_{10} p$ | β | MAC | MAF | Ps | Variants |
|---|---|---|---|---|---|---|
| DPMPKISSREP | 9.15 | 3.29 | 10 | 0.11 | -1 | Multiple differences to reference just after the start codon |

#### *lysA*

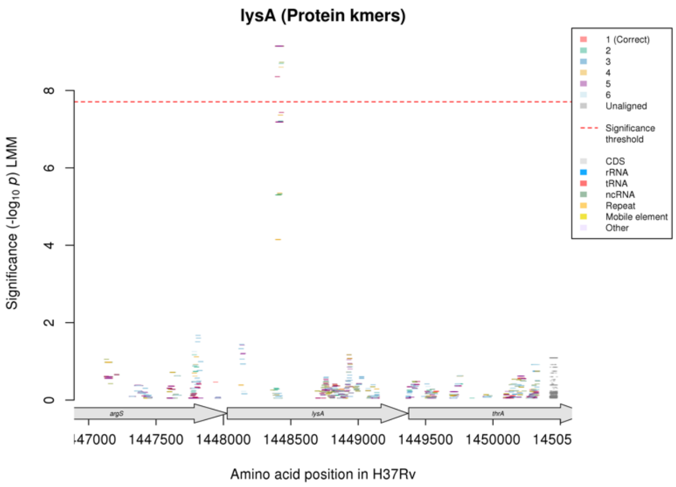

Not significant in nucleotide kmer analysis.

Protein kmers:

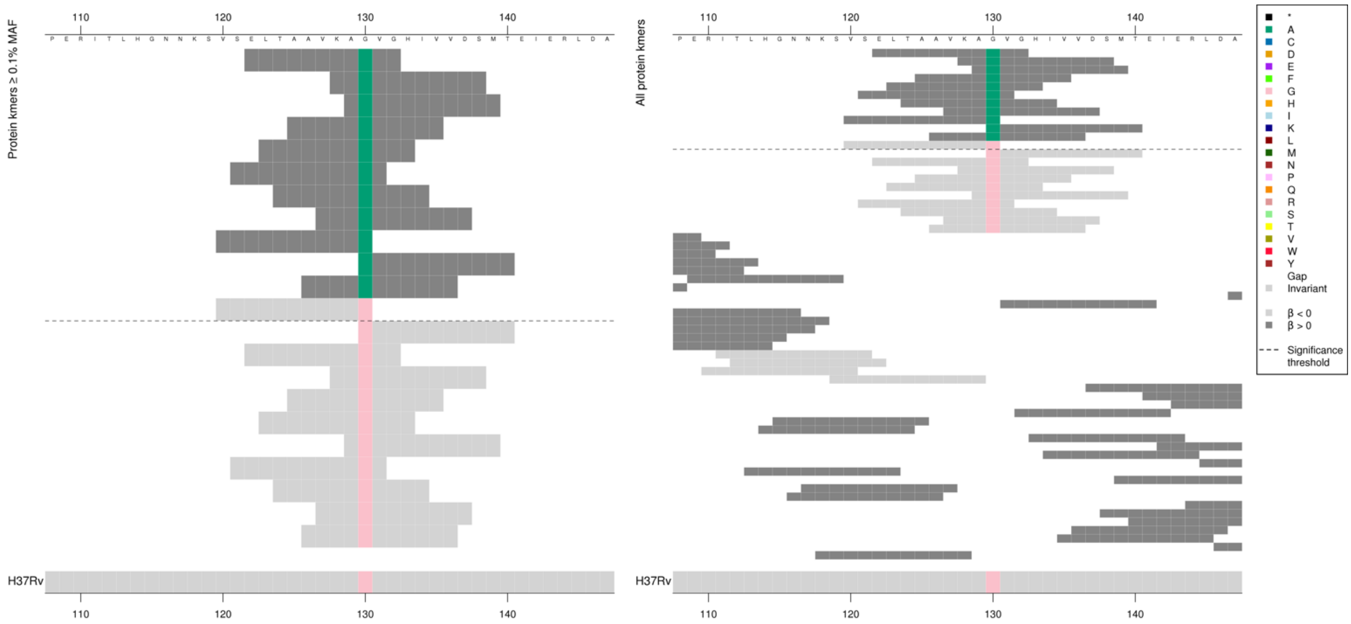

| Kmer | -log$_{10}$ *p* | β | MAC | MAF | Ps | Variants |
|---|---|---|---|---|---|---|
| ELTAAVKAAVG | 9.15 | 3.29 | 10 | 0.11 | 122 | 130A |
| VSELTAAVKAG | 8.36 | -2.94 | 14 | 0.15 | 120 | 130G |

#### *mprB*

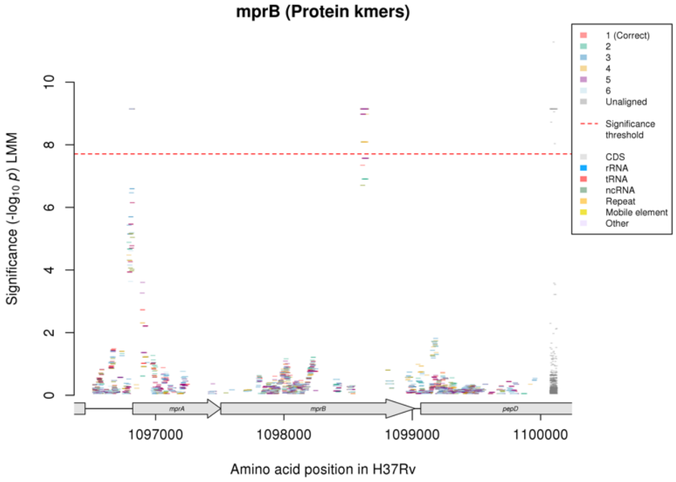

Not significant in the nucleotide kmer analysis.

Protein kmers:

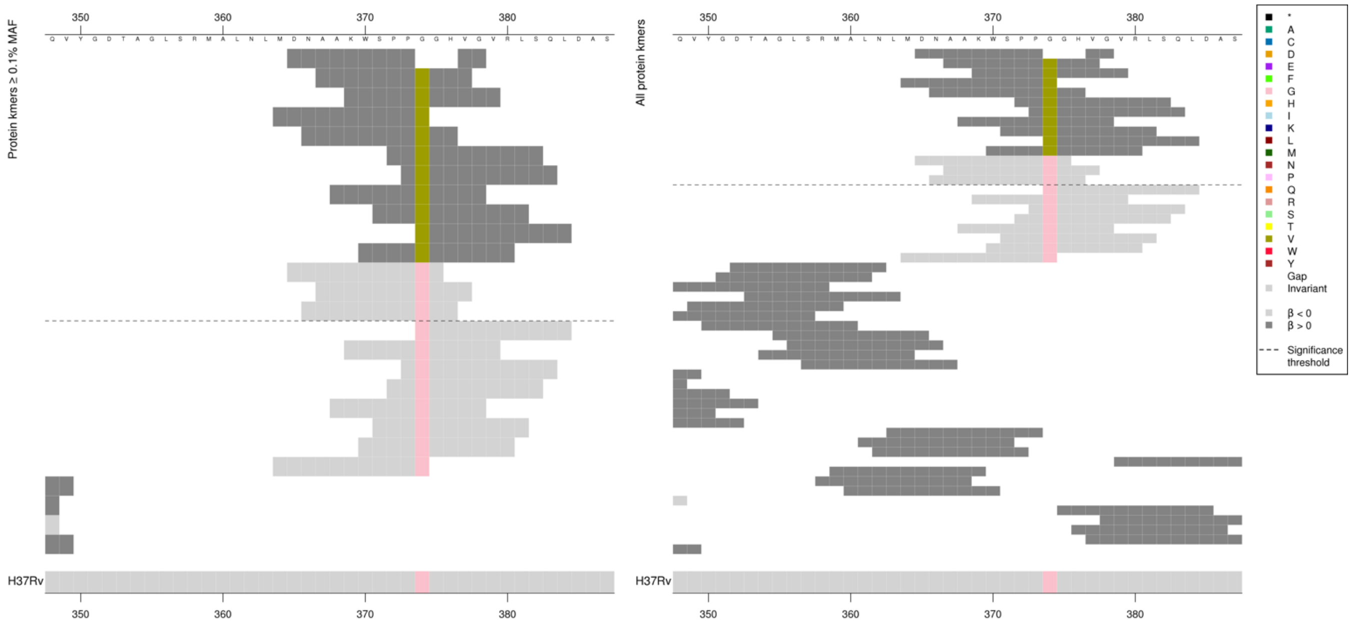

There is no deletion/gap in the top high MIC-associated kmer, BLAST has just inserted a gap rather than create a mismatch at codon 374.

| Kmer | -log$_{10}$ *p* | β | MAC | MAF | Ps | Variants |
|---|---|---|---|---|---|---|
| DNAAKWSPPVG | 9.15 | 3.29 | 10 | 0.11 | 365 | 374V |
| DNAAKWSPPGG | 8.98 | -3.24 | 11 | 0.12 | 365 | 374G |

#### *mprA*

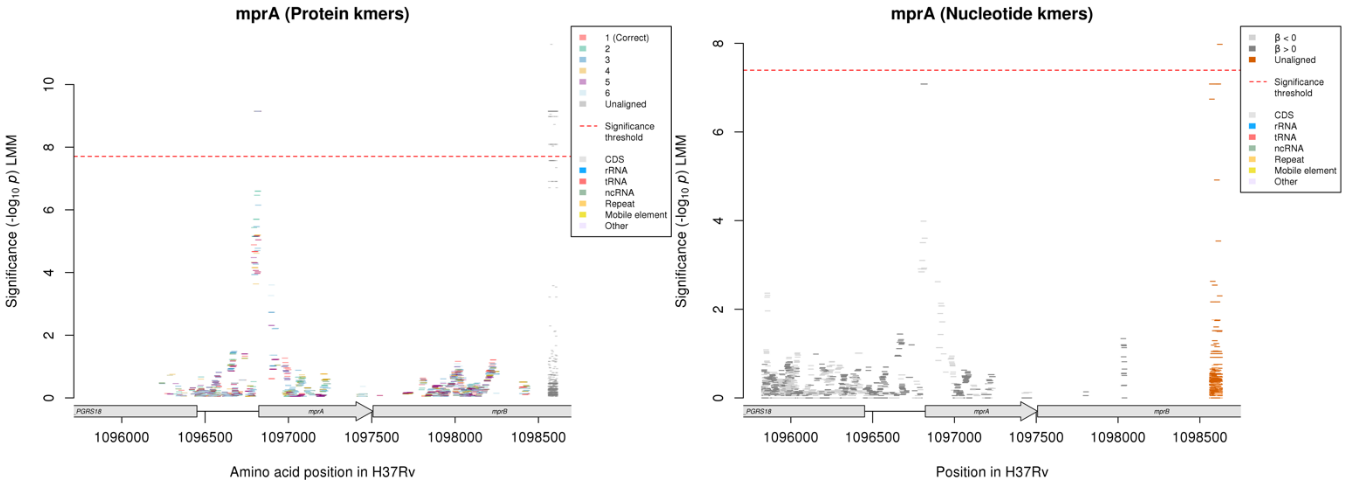

The intergenic region upstream of mprA is just below the significance threshold in the nucleotide kmer analysis.

Protein kmers:

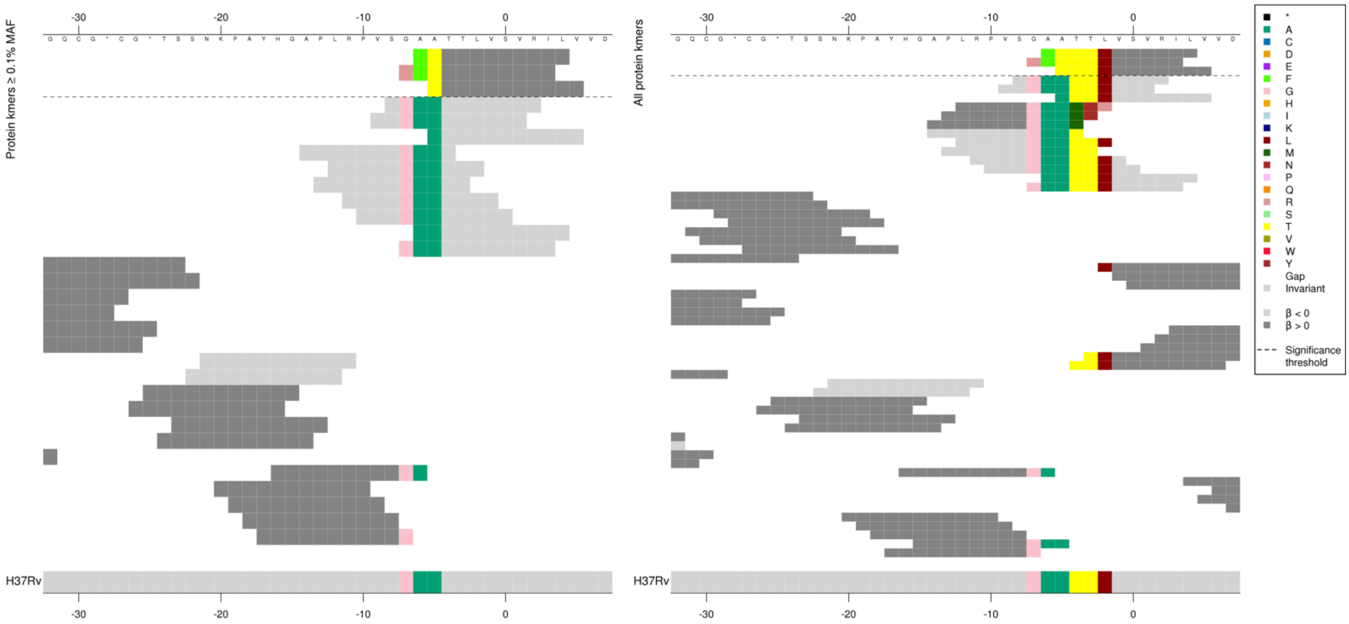

Capturing variants upstream of the start codon. This region is not significant in the nucleotide kmer analysis, but the alignments are below:

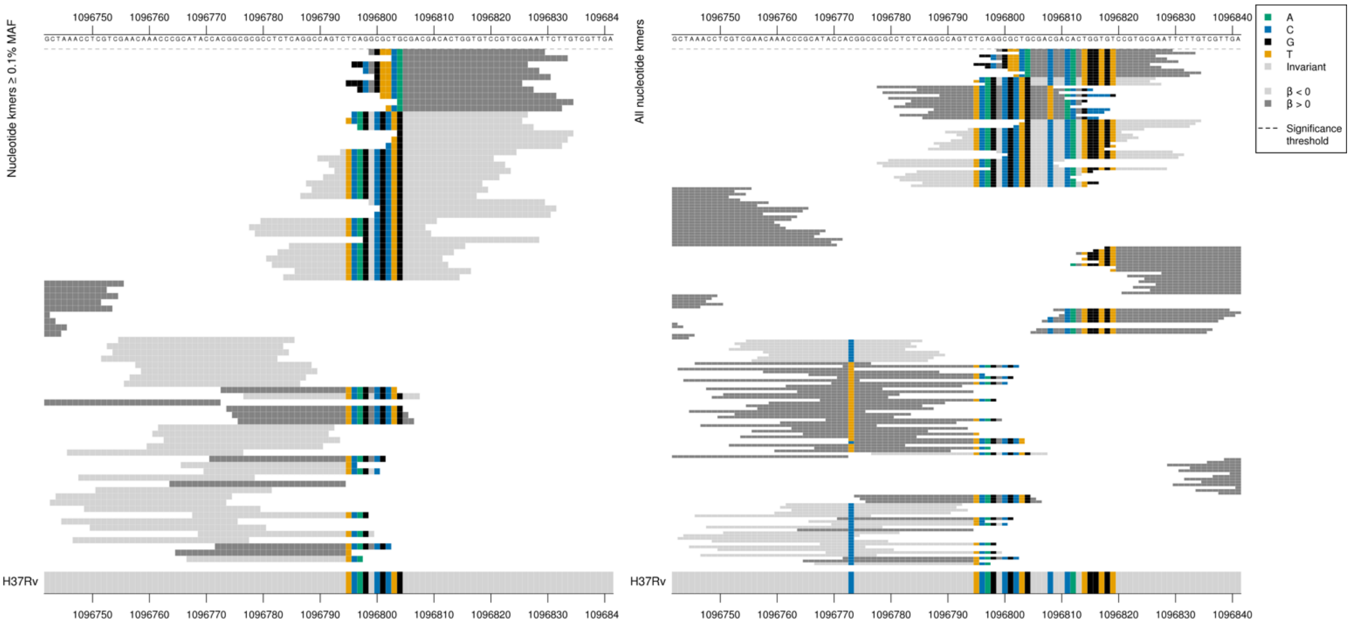

*mprA* begins at position 1096822.
Multiple differences compared to the reference, possibly a deletion or an insertion, most of the kmer is identical to the reference.

| Kmer | $-\log_{10} p$ | β | MAC | MAF | Ps | Variants |
|---|---|---|---|---|---|---|
| FTTTLVSVRIL | 9.15 | 3.29 | 10 | 0.11 | -6 | Multiple differences to reference upstream of start codon |

# *Rv3228*

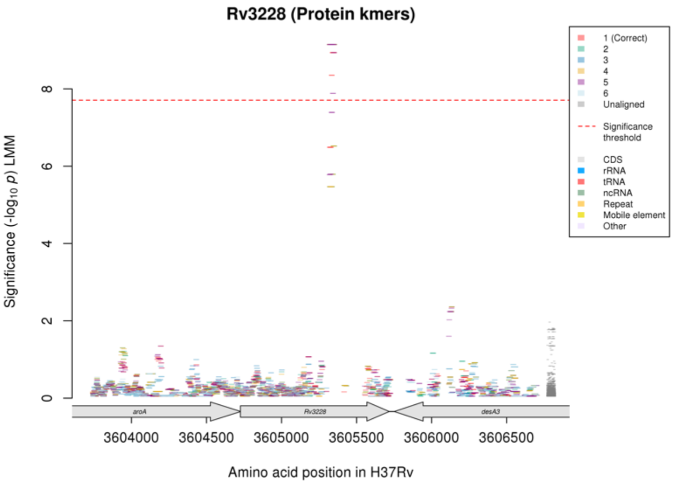

Not significant in the nucleotide kmer analysis.
Protein kmers:

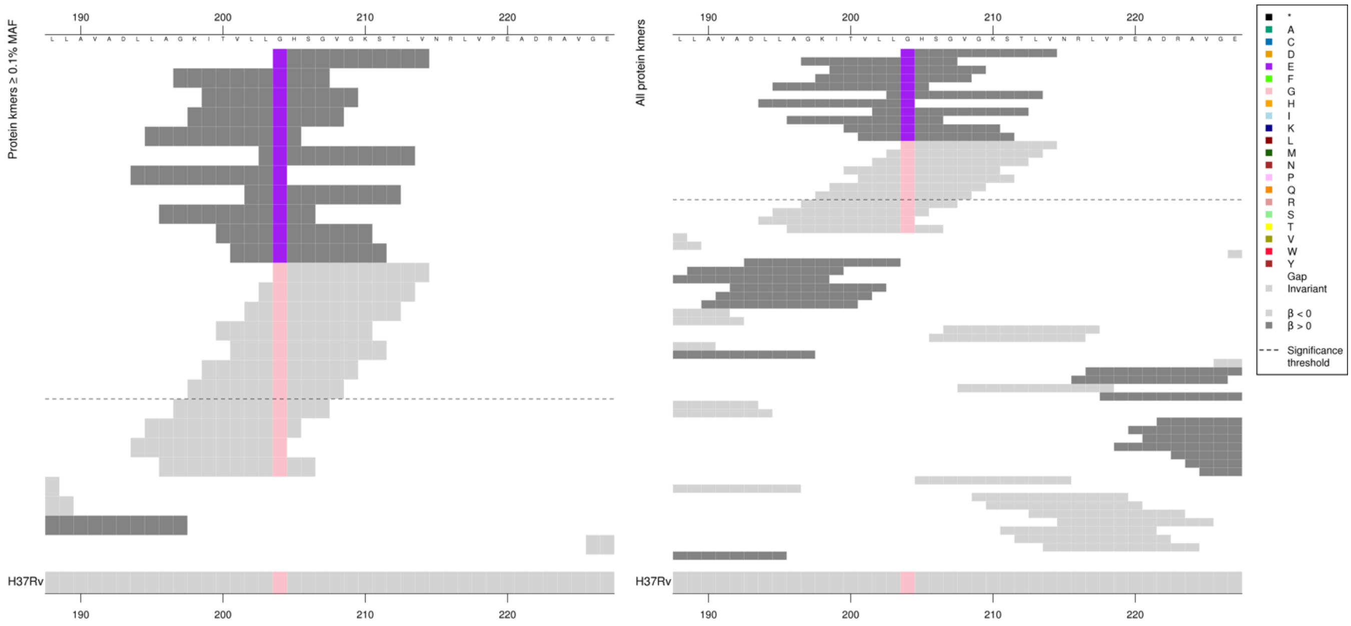

| Kmer | $-\log_{10} p$ | β | MAC | MAF | Ps | Variants |
|---|---|---|---|---|---|---|
| EHSGVGKSTLV | 9.15 | 3.29 | 10 | 0.11 | 204 | 204E |
| GHSGVGKSTLV | 8.94 | -3.19 | 12 | 0.13 | 204 | 204G |

# Rv1290c

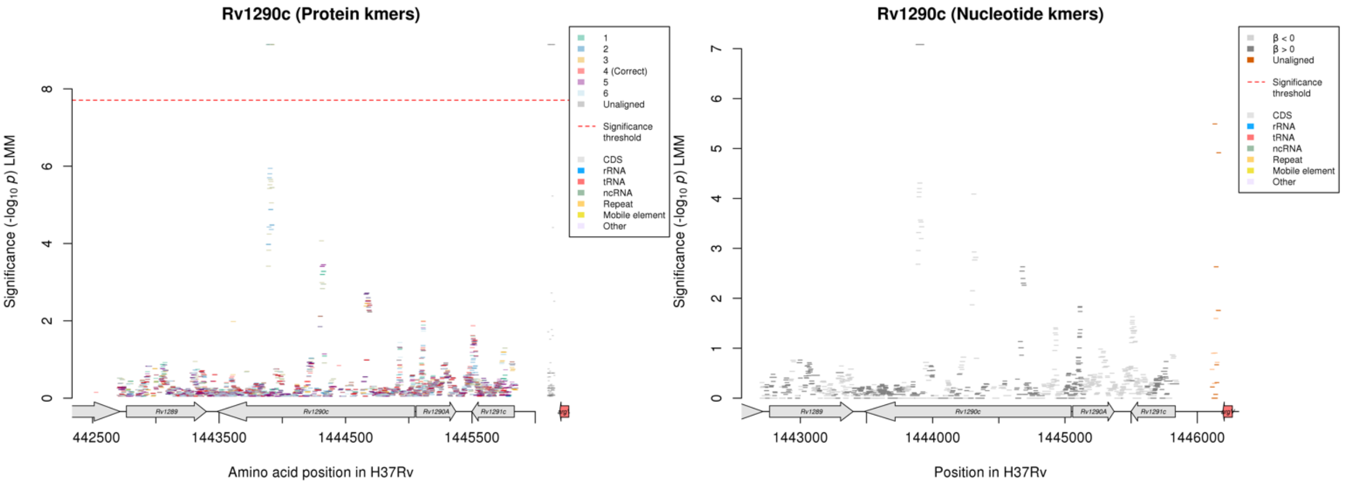

No significant kmers aligned to the correct reading frame for the protein kmer analysis.
Region is just below the significance threshold for nucleotide analysis.
Begins on reverse strand at 1445047
Nucleotide kmers:

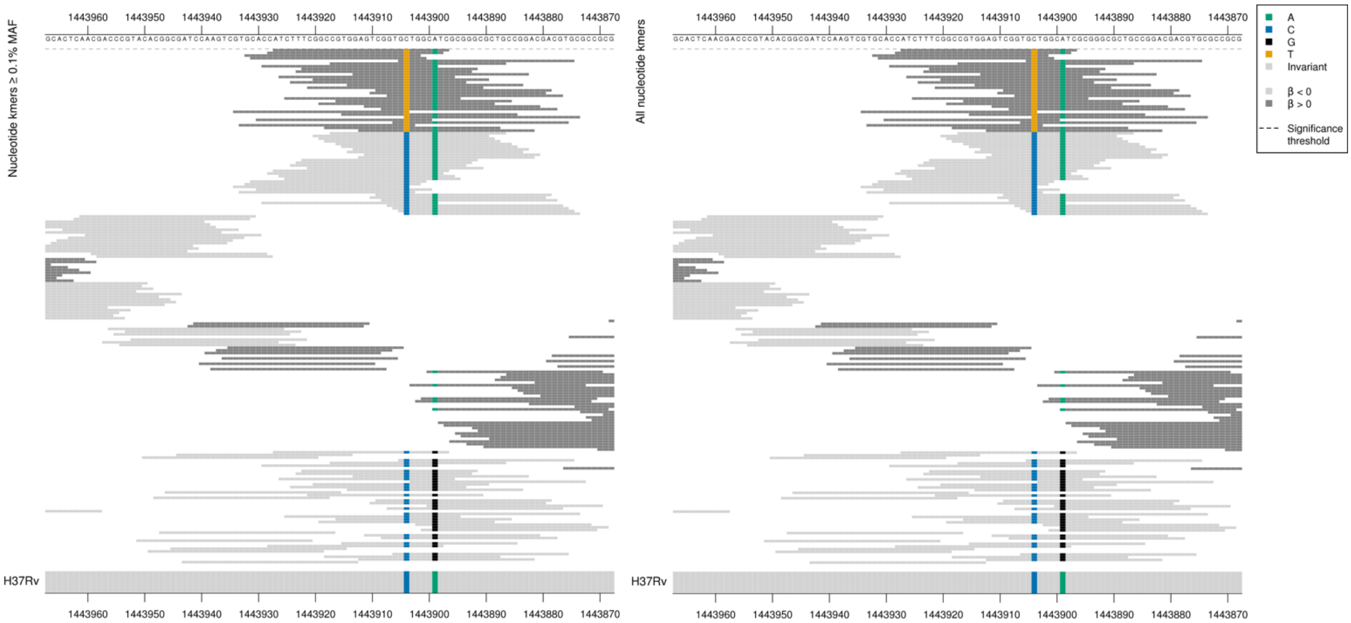

CTG = Leu / L
TTG = Leu / L

| Kmer | $-\log_{10} p$ | β | MAC | MAF | Ps | Variants |
|---|---|---|---|---|---|---|
| ATCTTTCGGCCGTGGAGTCGGTGTTGGCATC | 7.08 | 2.82 | 10 | 0.11 | 1443927 | 1443904T |

#### *pncA*

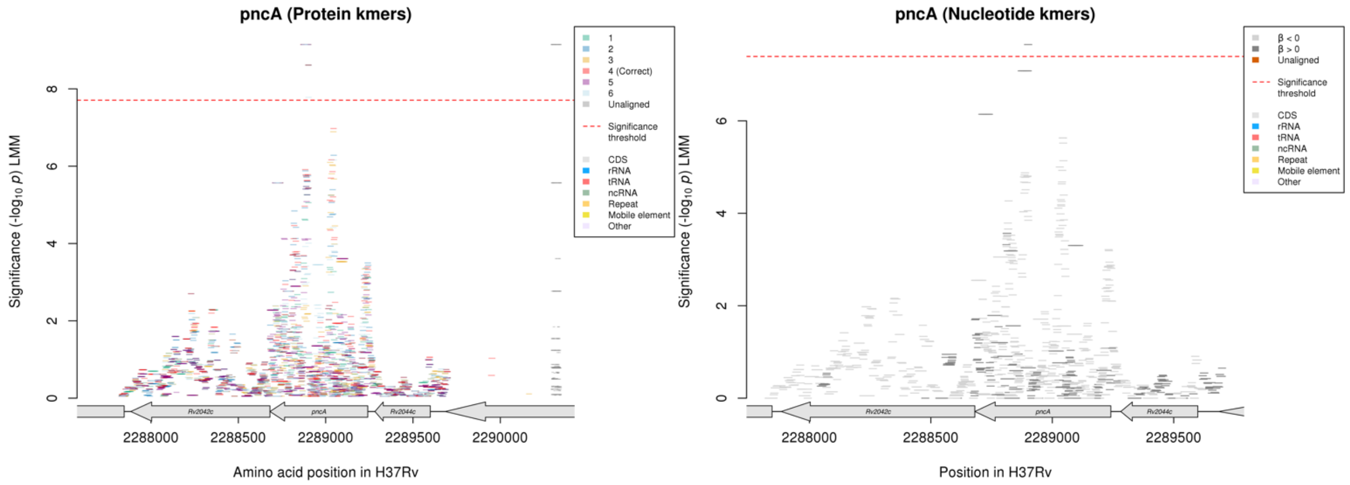

Protein kmers:

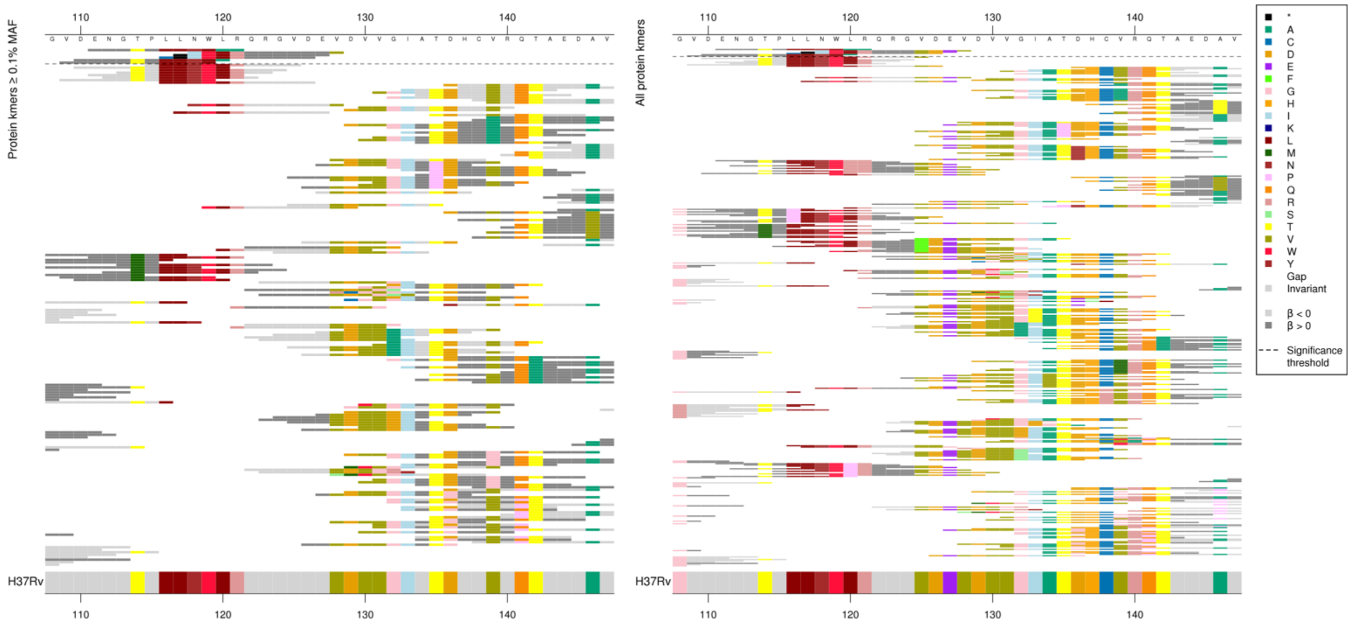

Same region for nucleotide kmers:

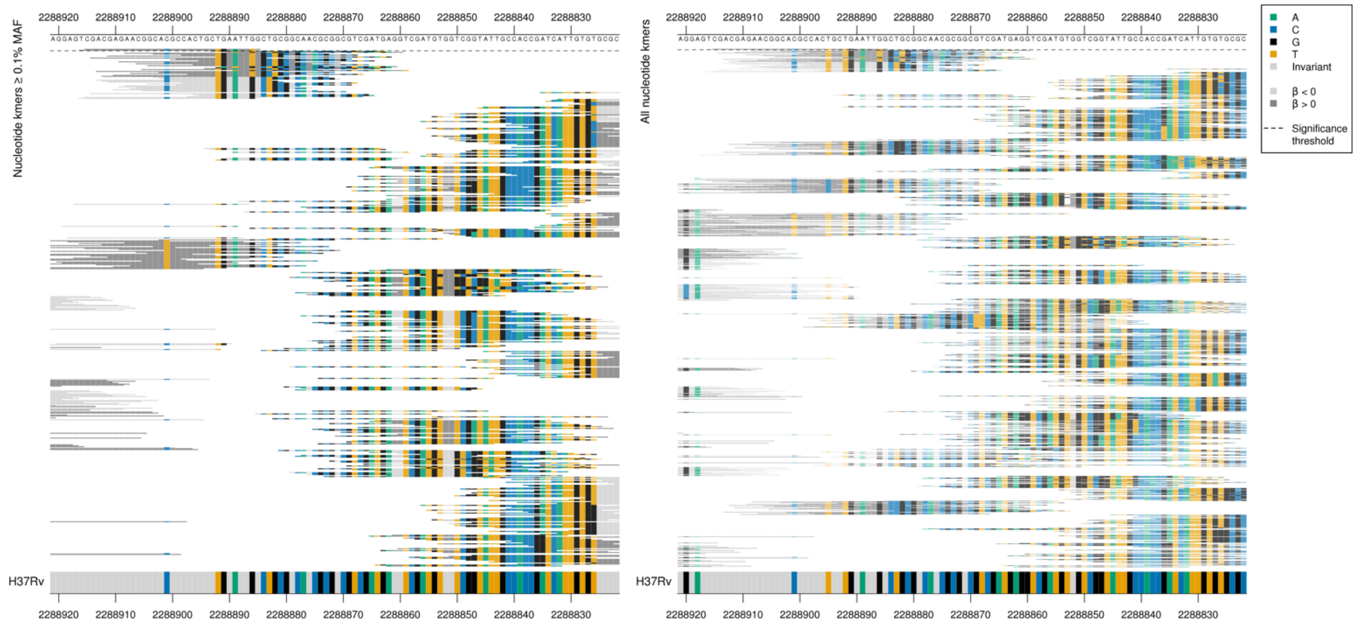

High MIC-associated kmers had a T insertion between position 2288888-2288886/354-356.

Highest protein kmer in this region: $-\log_{10} p$ = 9.14, beta = 3.29, MAC = 10, MAF = 0.11, pos = 111.

| Kmer | -log$_{10}$ *p* | β | MAC | MAF | Ps | Variants |
|---|---|---|---|---|---|---|
| CGACGAGAACGGCACGCCACTGCTGAATTTG | 7.65 | 2.11 | 14 | 0.15 | 2288915 | T insertion between position 2288888-2288886/354-356 |

#### *Rv2277c:pitB* $^{R}$

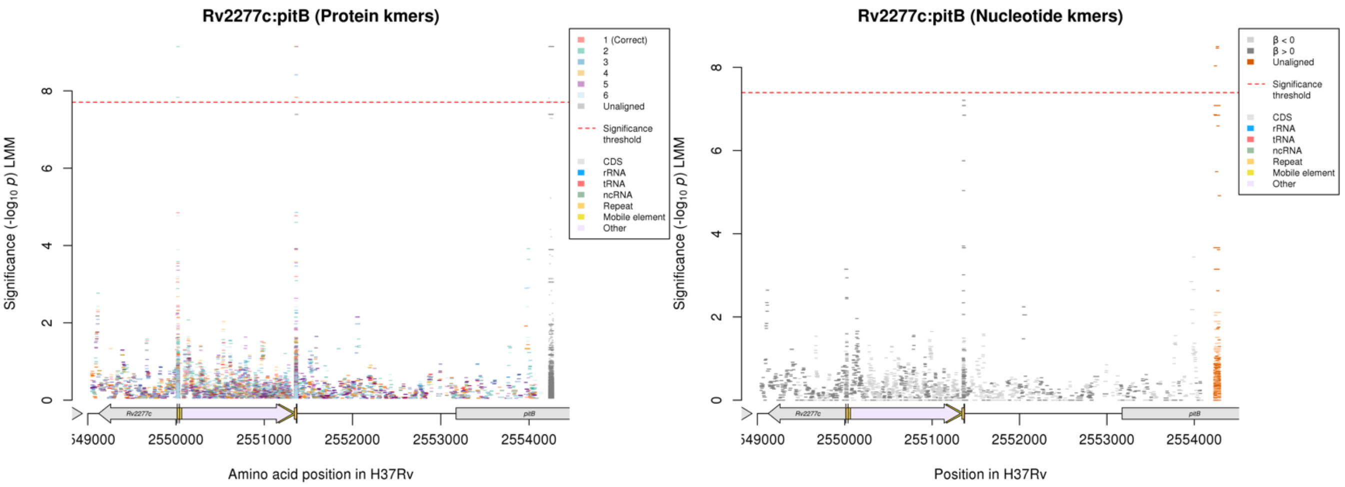

No significant kmers aligned to the region in the nucleotide kmer analysis, there were significant kmers that were assigned to the region that didn't align.

Nucleotide kmers in one of the regions significant in the protein kmer analysis, that is just below significance in the nucleotide kmer analysis:

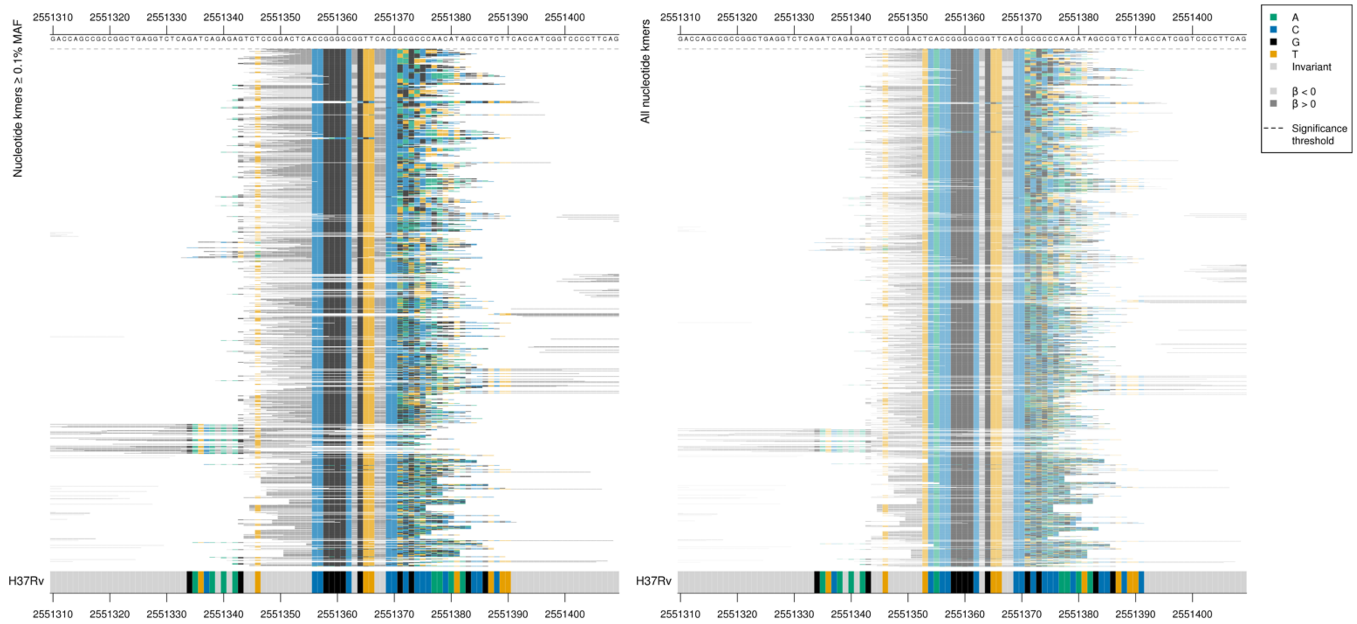

Repeat region at a mobile element.

| Kmer | -log$_{10}$ *p* | β | MAC | MAF | Ps | Variants |
|---|---|---|---|---|---|---|
| CCGGACTCACCGGGGCGGTTCACCTAGGCGG | 7.21 | 2.99 | 16 | 0.17 | 2551347 | Repeat region |

# *Rv0726c*

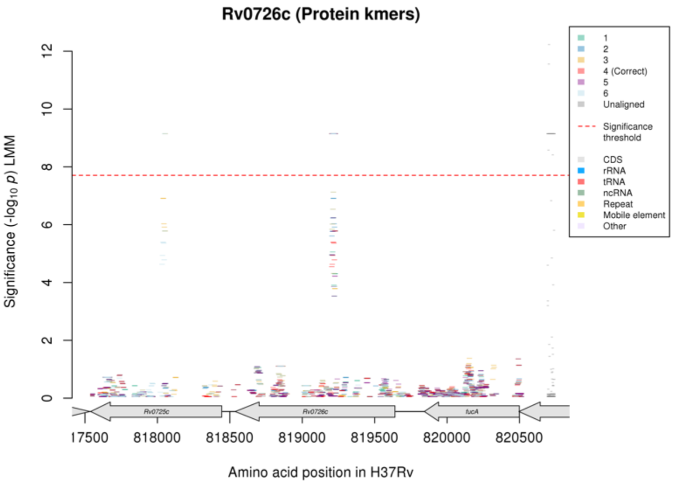

Not significant in the nucleotide kmer analysis.
No matches for top kmers on BLAST online. Top two kmers (-log$_{10}$ $p$ = 12.1 and 11.4) were not assigned to any other genes (four other assignments but each only seen in one genome, assignment to *Rv0726c* seen in 10 genomes).

Protein kmers:

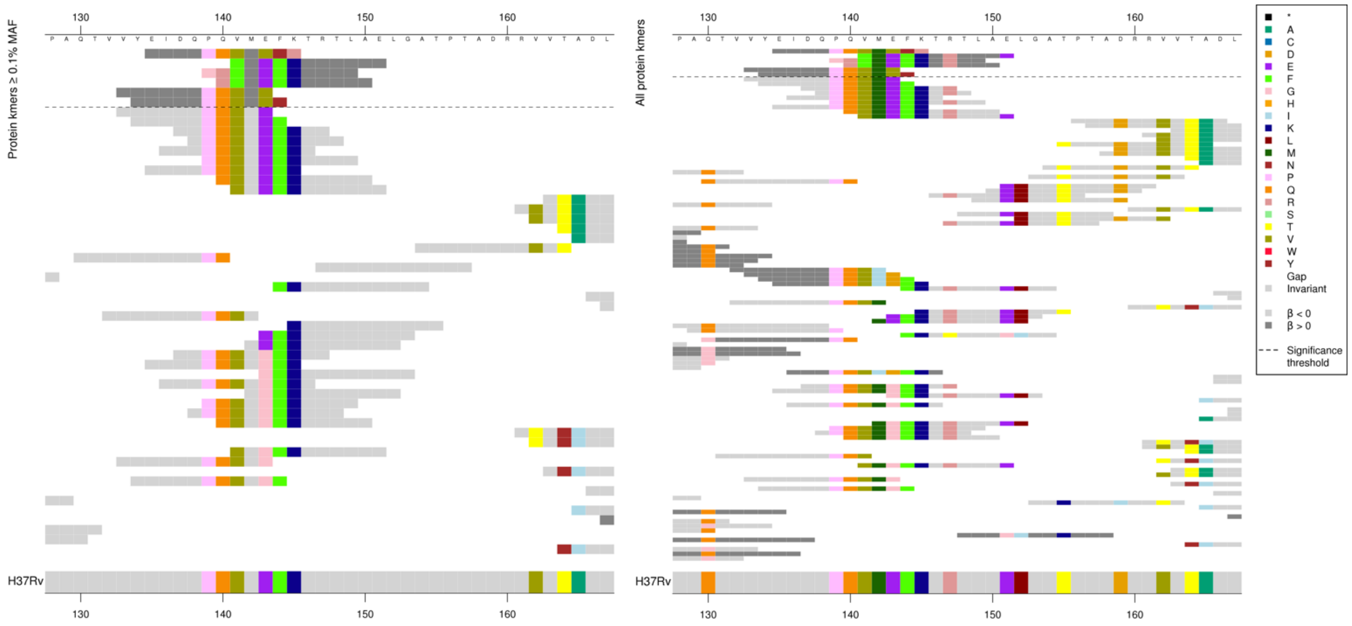

The significant high MIC-associated kmers were all the same as the reference for most of the kmer then there were multiple differences, but it is unclear if this is due to a potential insertion or deletion.

| Kmer | -log$_{10}$ $p$ | β | MAC | MAF | Ps | Variants |
|---|---|---|---|---|---|---|
| EIDQPQVMVNR | 9.15 | 3.29 | 10 | 0.11 | 135 | Multiple differences to the reference |

### cysA3/cysA2

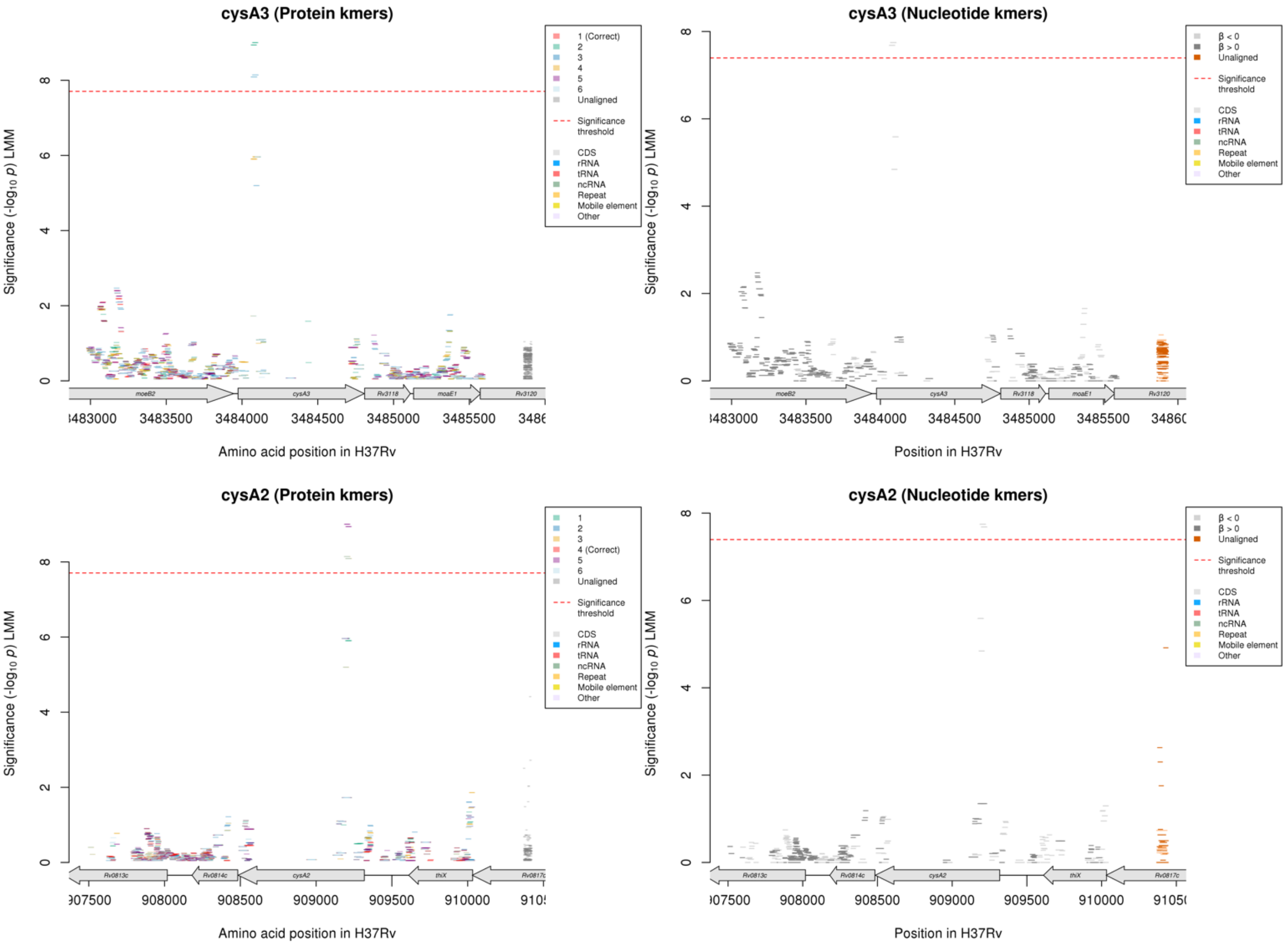

#### Nucleotide kmers:

*cysA3*:

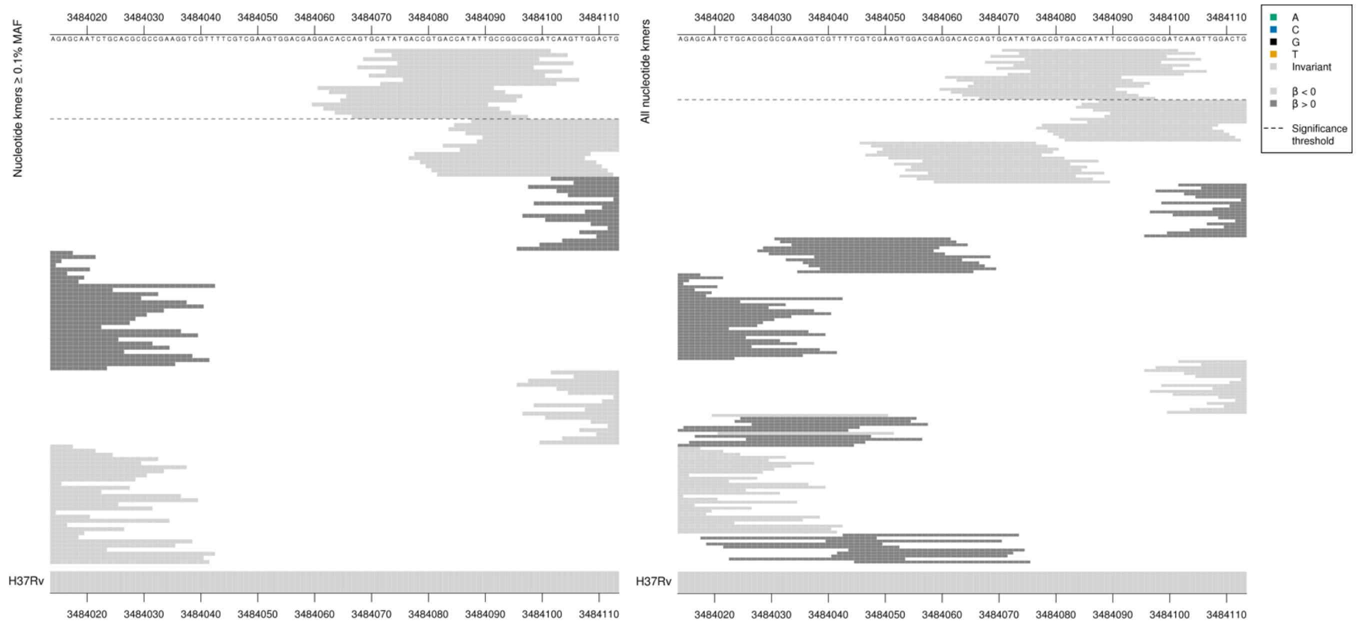

*cysA2*:

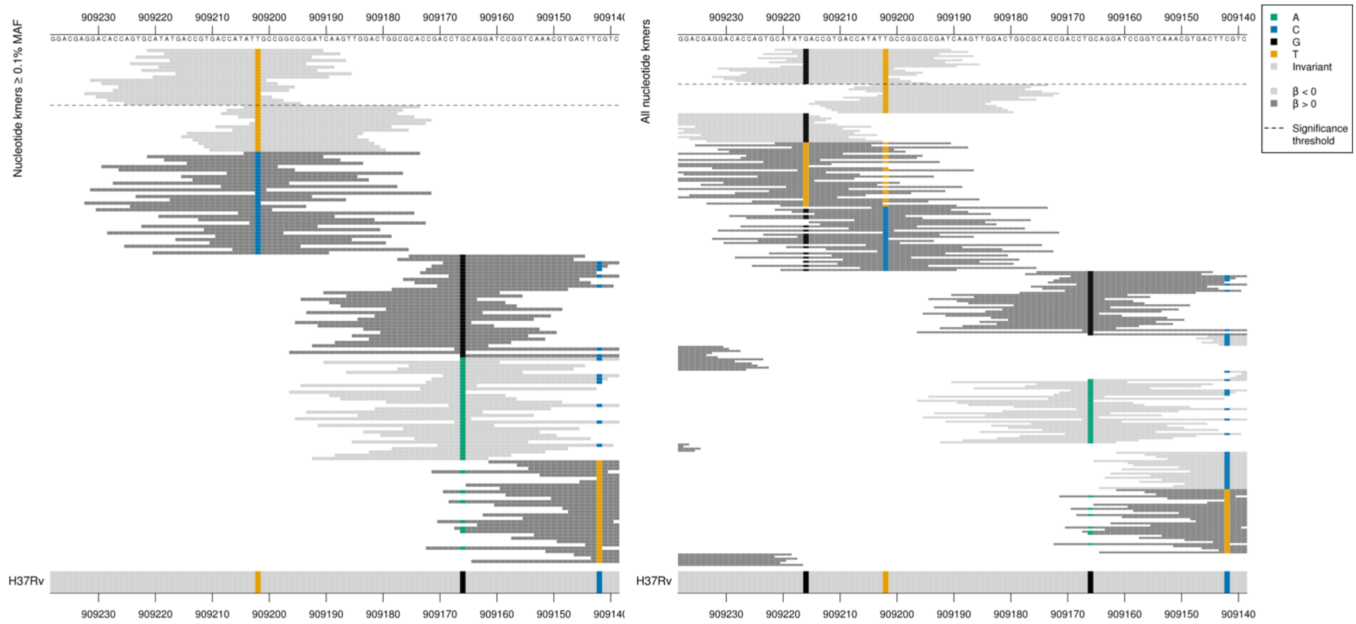

Non-synonymous mutation at position 909216/103 (codon 35)
GAC = Asp / D
TAC = Tyr / Y
Synonymous mutation at position 909202/117 (codon 39)
ATT = Ile / I
ATC = Ile / I

| Kmer | $-\log_{10} p$ | β | MAC | MAF | Ps | Variants |
|---|---|---|---|---|---|---|
| CATATGACCGTGACCATATTGCCGGCGCGAT | 7.74 | -2.65 | 22 | 0.23 | 909221 | 909216/103G (35D aa), 909202/117T (cysA2 positions, same kmer as cysA3) |

## Rv0914c

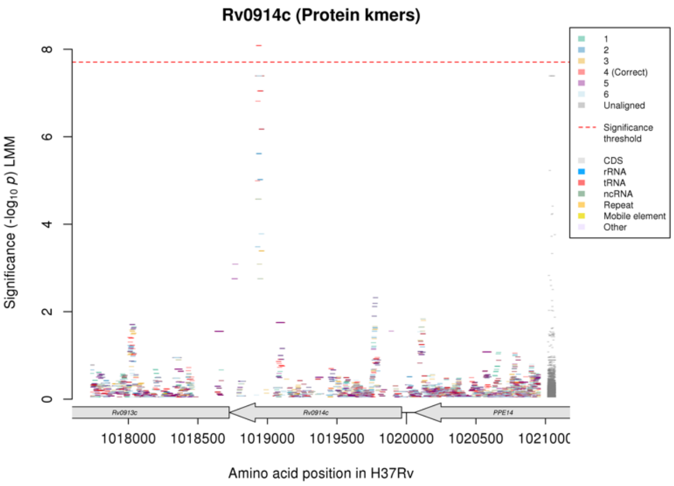

Not significant in the nucleotide kmer analysis.

Protein kmers:

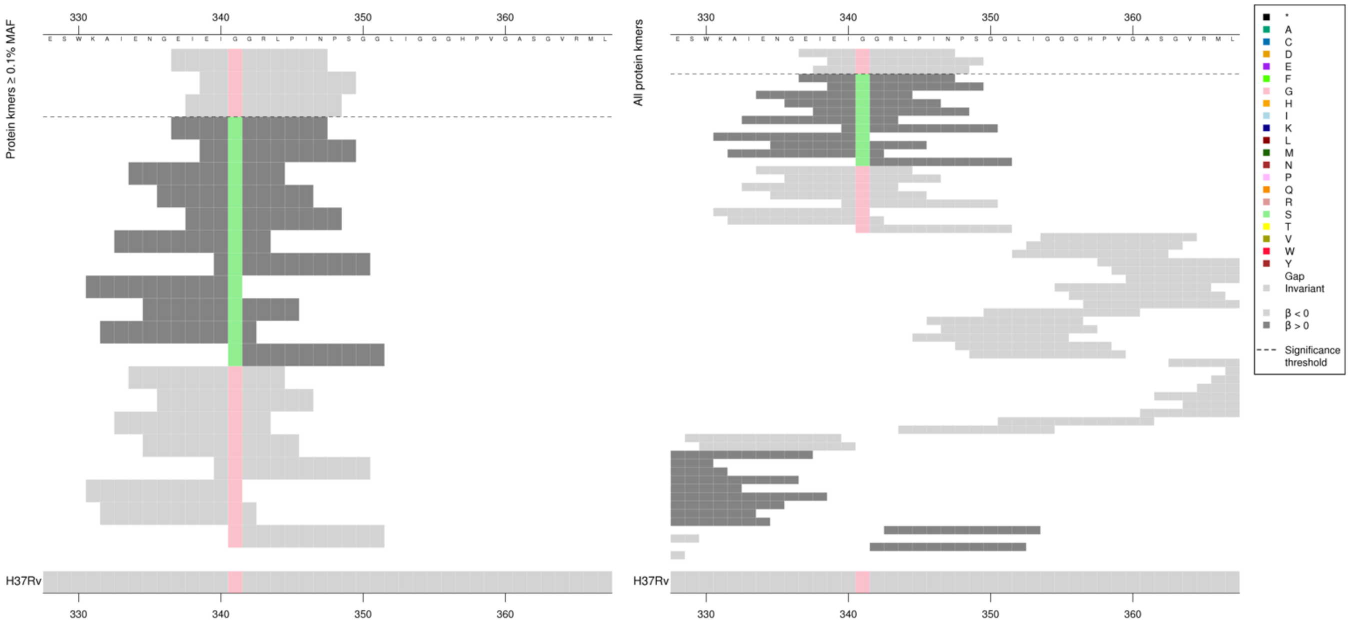

Kmers below significance threshold = 341S

| Kmer | $-\log_{10} p$ | β | MAC | MAF | Ps | Variants |
|---|---|---|---|---|---|---|
| EIEIGGRLPIN | 8.08 | -2.91 | 19 | 0.20 | 337 | 341G |

### Rifampicin (RIF)

#### *rpoB*

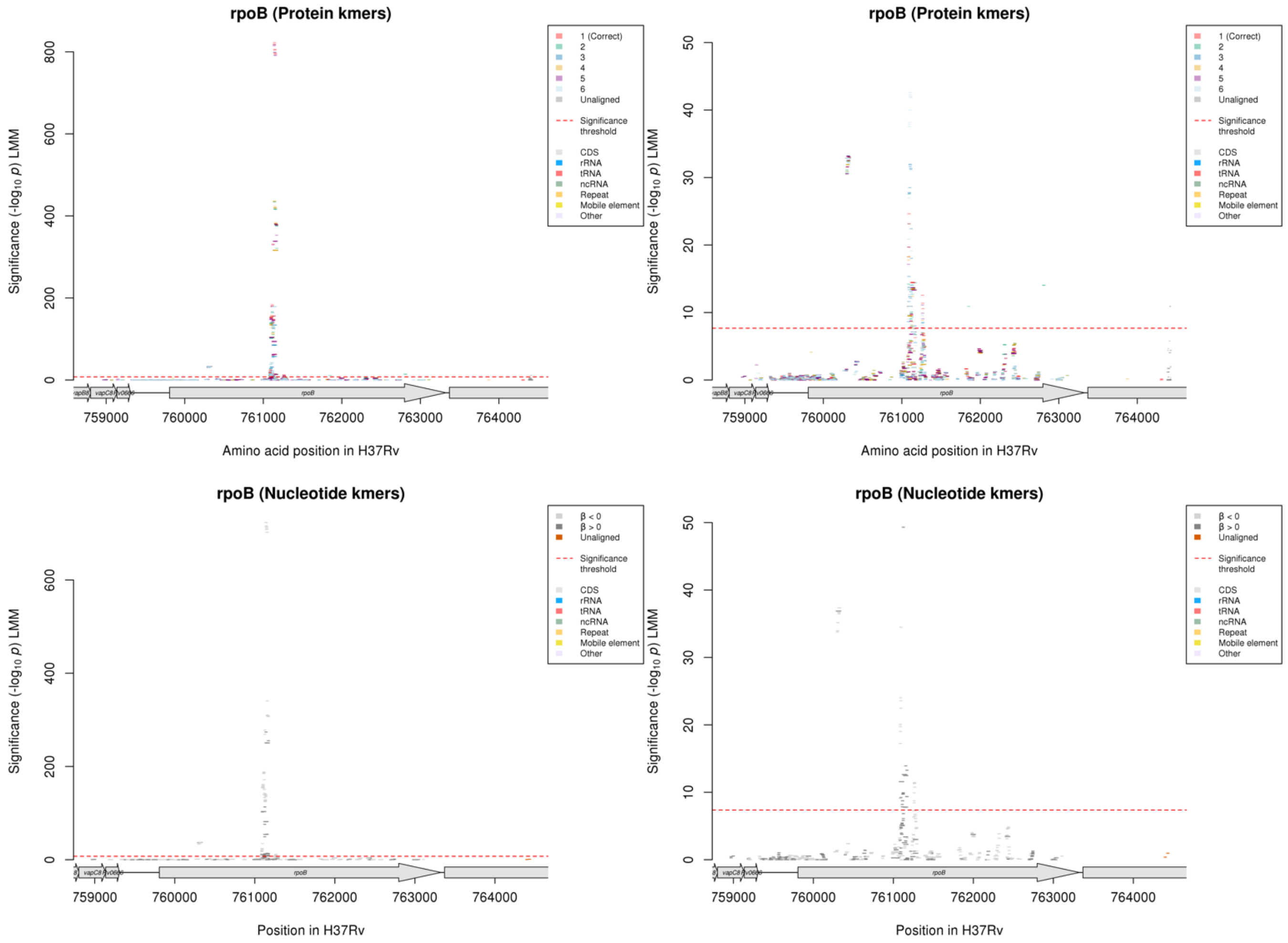

Protein kmers:

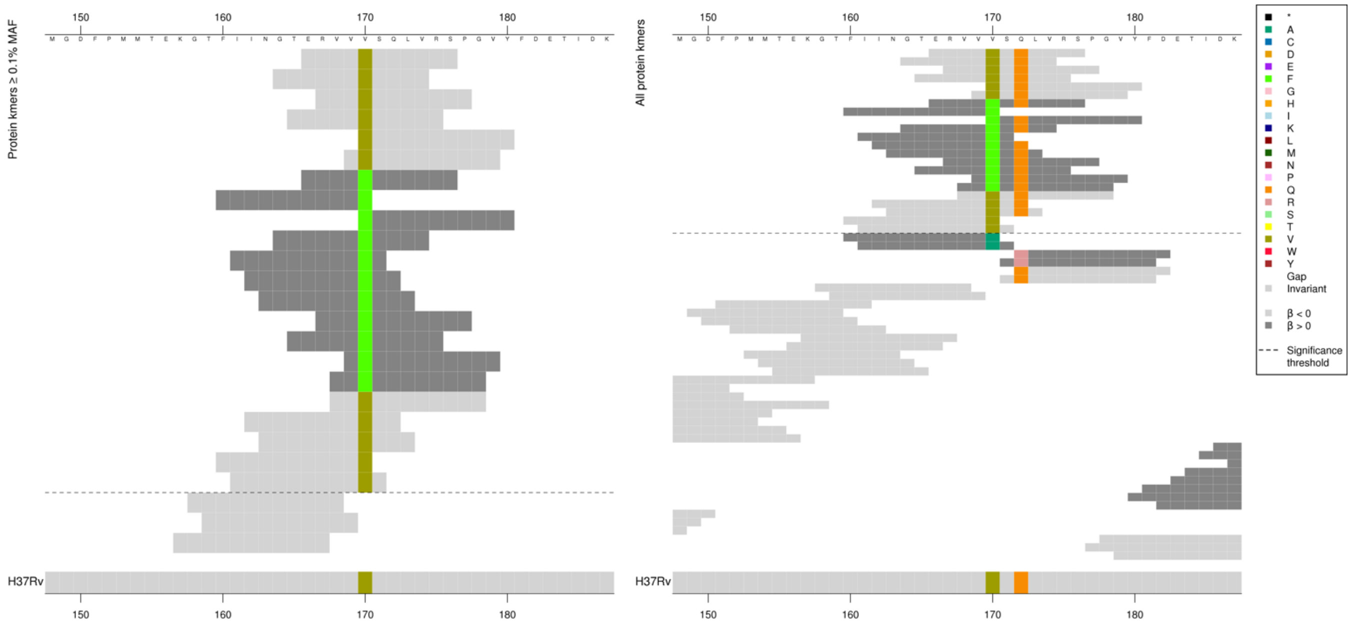

Range of *p*-values for the low MIC-associated kmers, all high MIC-associated kmers were the same *p*-value. Kmers below significance threshold = 170A, 172R

| Kmer | $-\log_{10} p$ | β | MAC | MAF | Ps | Variants |
|---|---|---|---|---|---|---|
| ERVVVSQLVRS | 33.14 | -4.56 | 30 | 0.36 | 166 | 170V, 172Q |
| ERVVFSQLVRS | 32.87 | 5.36 | 20 | 0.24 | 166 | 170F |
| FIINGTERVVV | 30.56 | -4.46 | 29 | 0.35 | 160 | 170V |

Protein kmers:

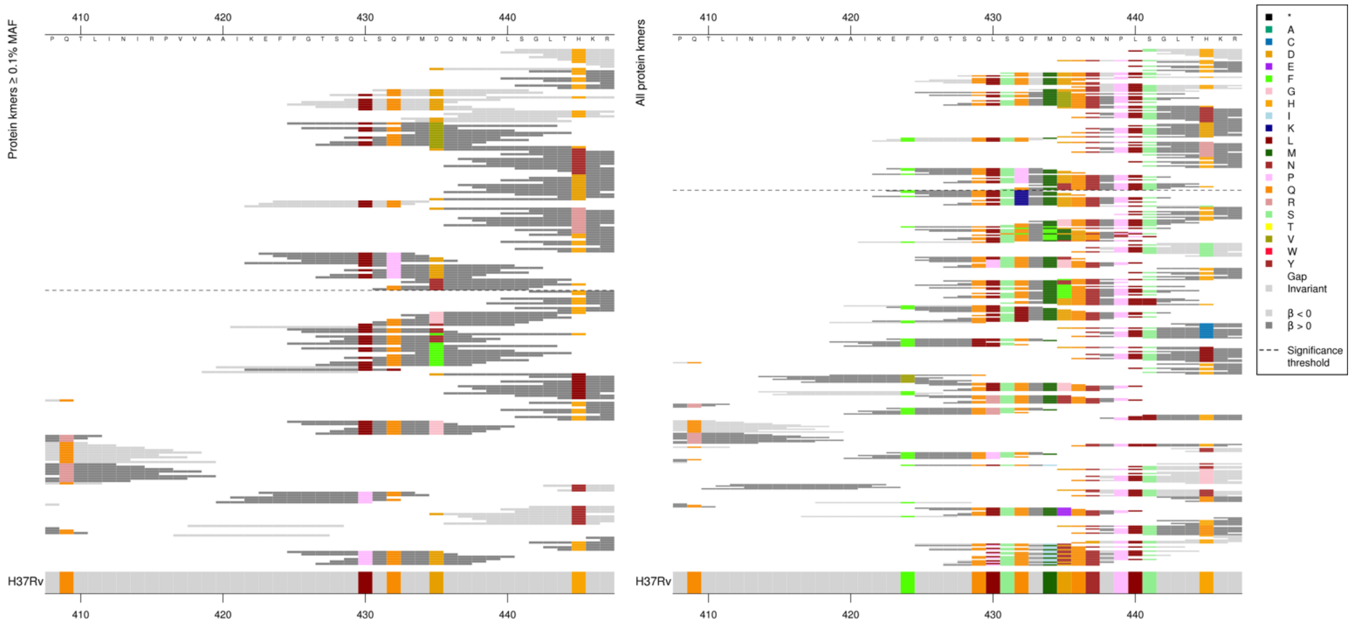

Focusing on the kmer centered on 435 for this alignment.
Kmers capturing 435V, 432P were all the same p-value as each other. 435Y kmers were slightly different p-values, just showing the top one here. Kmers capturing 435Y were not significant for RFB.

| Kmer | $-\log_{10} p$ | β | MAC | MAF | Ps | Variants |
|---|---|---|---|---|---|---|
| SQFMDQNNPLS | 183.36 | -3.94 | 605 | 7.21 | 431 | 431S, 432Q 434M, 435D, 436Q, 437N, 439P, 440L, 441S |
| QFMDQNNPLSG | 180.56 | -3.91 | 603 | 7.18 | 432 | 432Q, 434M, 435D, 436Q, 437N, 439P, 440L, 441S |
| SQLSQFMDQNN | 156.89 | -3.36 | 678 | 8.08 | 428 | 429Q, 430L, 431S, 432Q 434M, 435D, 436Q, 437N |
| QLSQFMDQNNP | 153.22 | -3.32 | 675 | 8.04 | 429 | 429Q, 430L, 431S, 432Q 434M, 435D, 436Q, 437N, 439P |
| GTSQLSQFMDQ | 152.84 | -3.32 | 675 | 8.04 | 426 | 429Q, 430L, 431S, 432Q 434M, 435D, 436Q |
| FGTSQLSQFMD | 152.34 | -3.32 | 673 | 8.02 | 425 | 429Q, 430L, 431S, 432Q 434M, 435D |
| TSQLSQFMDQN | 151.14 | -3.29 | 678 | 8.08 | 427 | 429Q, 430L, 431S, 432Q 434M, 435D, 436Q, 437N |
| LSQFMDQNNPL | 149.12 | -3.29 | 670 | 7.98 | 430 | 430L, 431S, 432Q 434M, 435D, 436Q, 437N, 439P, 440L |
| FMDQNNPLSGL | 139.20 | -3.59 | 564 | 6.72 | 433 | 434M, 435D, 436Q, 437N, 439P, 440L |
| FGTSQLSQFMV | 104.03 | 3.97 | 410 | 4.88 | 425 | 435V |
| FFGTSQLSQFM | 24.62 | -1.78 | 166 | 1.98 | 424 | 424F, 429Q, 430L, 431S, 432Q, 434M |
| EFFGTSQLSQF | 19.68 | -1.63 | 152 | 1.81 | 423 | 424F, 429Q, 430L, 431S, 432Q |
| EFFGTSQLSPF | 10.91 | 4.34 | 14 | 0.17 | 423 | 432P |
| MYQNNPLSGLT | 9.77 | 1.68 | 62 | 0.74 | 434 | 435Y |

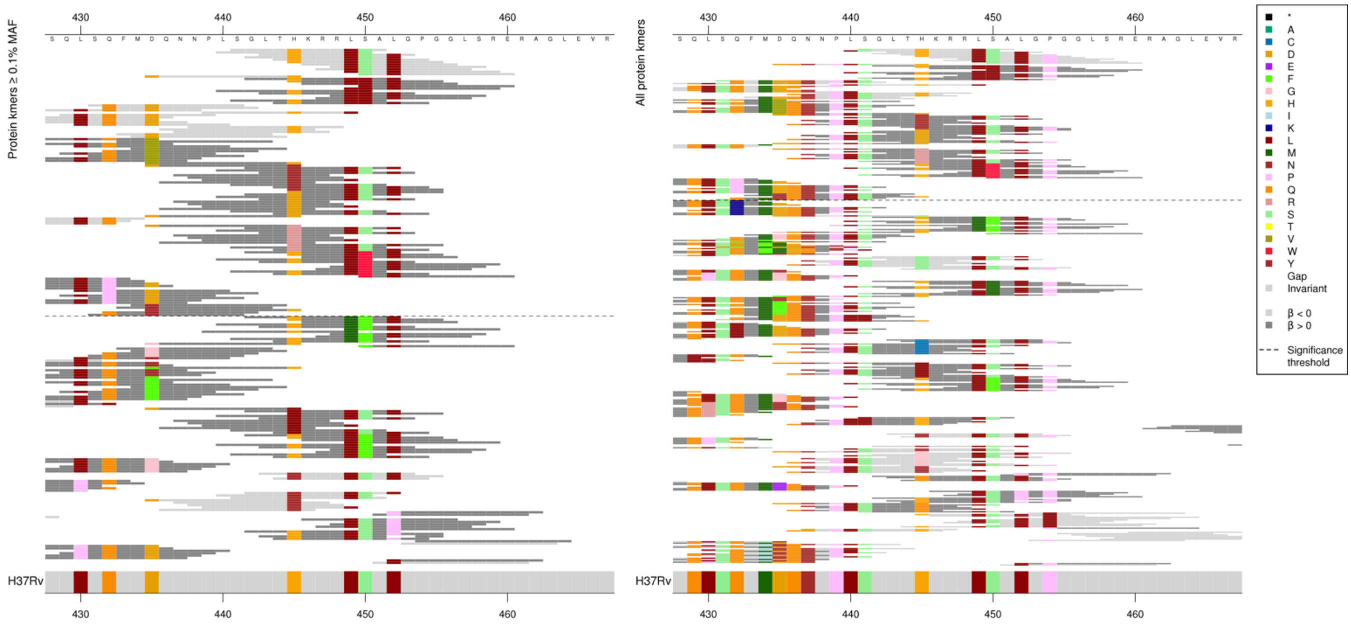

Focusing on the kmers capturing variants centered on 445-450 from this alignment.
Kmers capturing 450L, 445Y, 445R, 450W were all the same p-value. 445D kmers were slightly different p-values.

| Kmer | $-\log_{10} p$ | β | MAC | MAF | Ps | Variants |
|---|---|---|---|---|---|---|
| SGLTHKRRLSA | 822.66 | -5.43 | 2664 | 31.74 | 441 | 441S, 445H, 449L, 450S |
| LSGLTHKRRLS | 818.61 | -5.42 | 2661 | 31.70 | 440 | 440L, 441S, 445H, 449L, 450S |
| LTHKRRLSALG | 805.37 | -5.37 | 2717 | 32.37 | 443 | 445H, 449L, 450S, 452L |
| THKRRLSALGP | 795.64 | -5.34 | 2725 | 32.46 | 444 | 445H, 449L, 450S, 452L, 454P |
| KRRLSALGPGG | 380.96 | -4.47 | 2400 | 28.59 | 446 | 449L, 450S, 452L, 454P |
| SALGPGGLSRE | 377.40 | -4.45 | 2395 | 28.53 | 450 | 450S, 452L, 454P |
| DQNNPLSGLTH | 331.19 | -3.97 | 887 | 10.57 | 435 | 435D, 436Q, 437N, 439P, 440L, 441S, 445H |
| GLTHKRRLLAL | 316.29 | 4.27 | 2237 | 26.65 | 442 | 450L |
| PLSGLTHKRRL | 156.46 | -3.50 | 358 | 4.26 | 439 | 439P, 440L, 441S, 445H, 449L |
| NPLSGLTHKRR | 147.97 | -3.45 | 350 | 4.17 | 438 | 439P, 440L, 441S, 445H |
| QNNPLSGLTHK | 146.33 | -3.38 | 358 | 4.26 | 436 | 436Q, 437N, 439P, 440L, 441S, 445H |
| NNPLSGLTHKR | 146.05 | -3.38 | 358 | 4.26 | 437 | 437N, 439P, 440L, 441S, 445H |
| DQNNPLSGLTY | 93.87 | 4.02 | 130 | 1.55 | 435 | 445Y |
| DKRRLSALGPG | 62.80 | 3.88 | 96 | 1.14 | 445 | 445D |
| DQNNPLSGLTR | 14.44 | 3.04 | 30 | 0.36 | 435 | 445R |
| GLTHKRRLWAL | 13.35 | 2.28 | 45 | 0.54 | 442 | 450W |

Protein kmers:

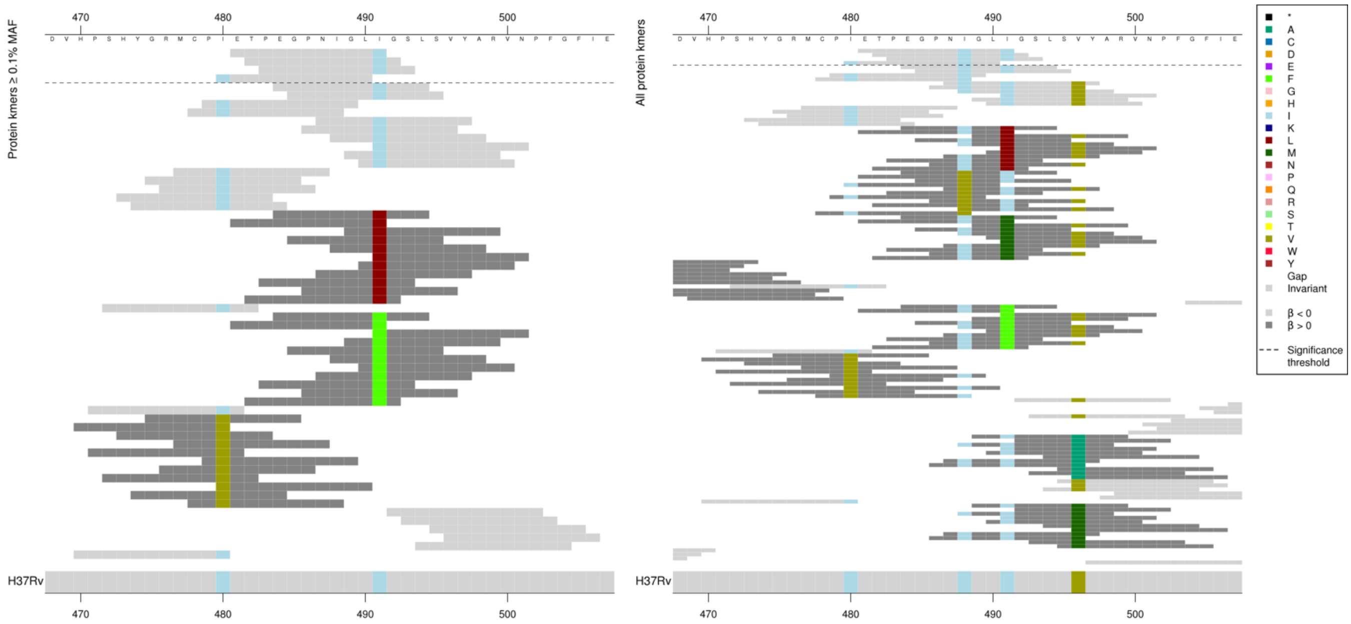

For RIF, the significant low MIC-associated kmers did not cover position 491I.

Kmers below significance threshold = 488V, 491L/M/F

| Kmer | $-\log_{10} p$ | β | MAC | MAF | Ps | Variants |
|---|---|---|---|---|---|---|
| ETPEGPNIGLI | 12.56 | -2.16 | 64 | 0.76 | 481 | 488I, 491I |
| IETPEGPNIGL | 7.71 | -2.24 | 27 | 0.32 | 480 | 480I, 488I |

#### *katG*

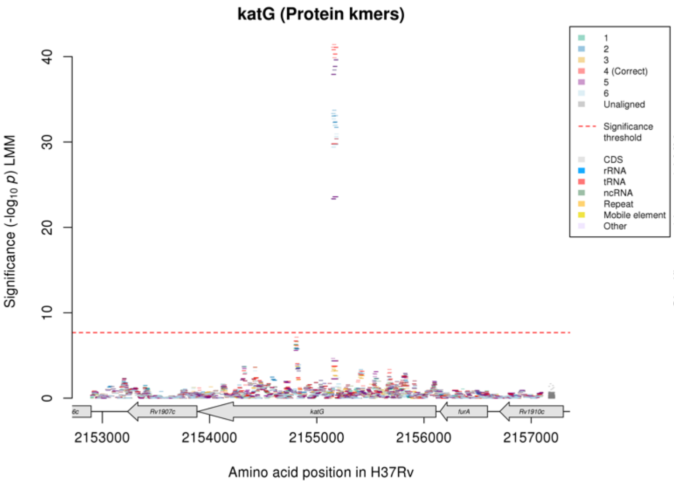

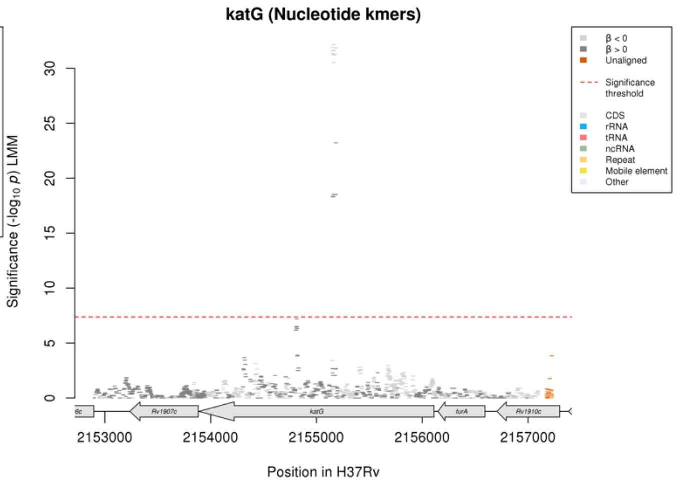

Protein kmers:

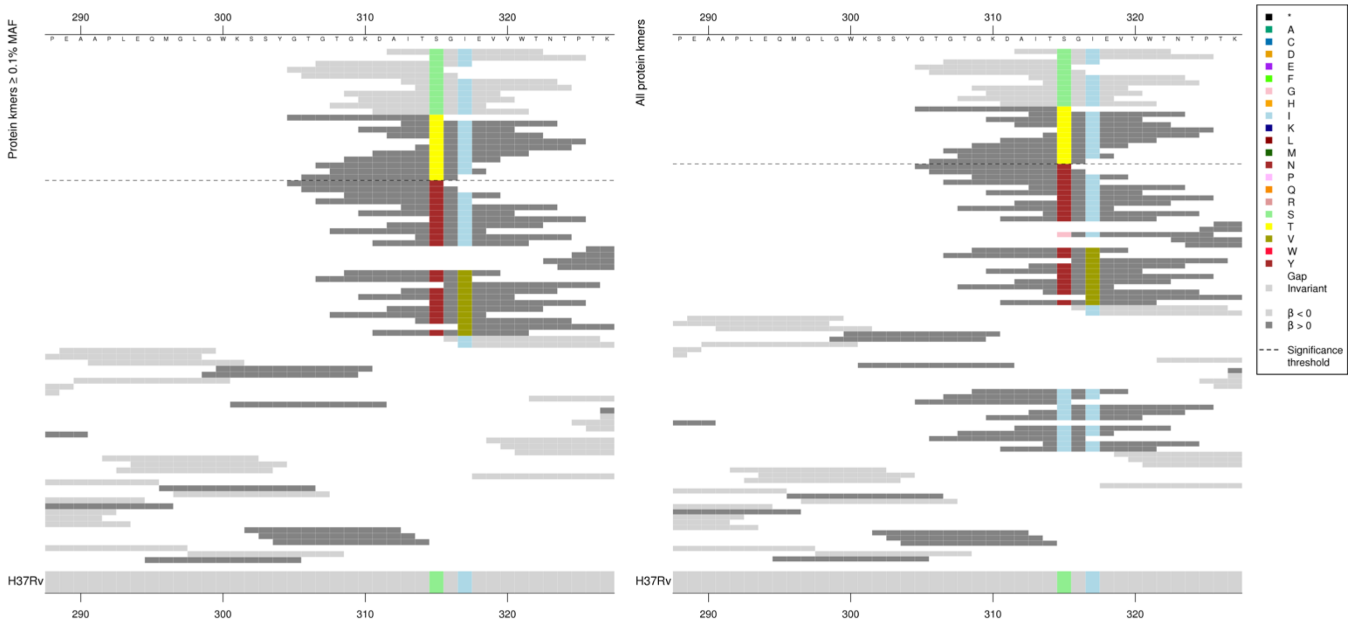

*P*-values varied for both low MIC-associated and high MIC-associated kmers, but not in line with whether they covered codon 317.

| Kmer | $-\log_{10} p$ | β | MAC | MAF | Ps | Variants |
|---|---|---|---|---|---|---|
| AITSGIEVVWT | 41.41 | -1.78 | 3201 | 38.13 | 312 | 315S |
| GTGTGKDAITT | 30.37 | 1.58 | 3116 | 37.12 | 305 | 315T |

#### *embB*

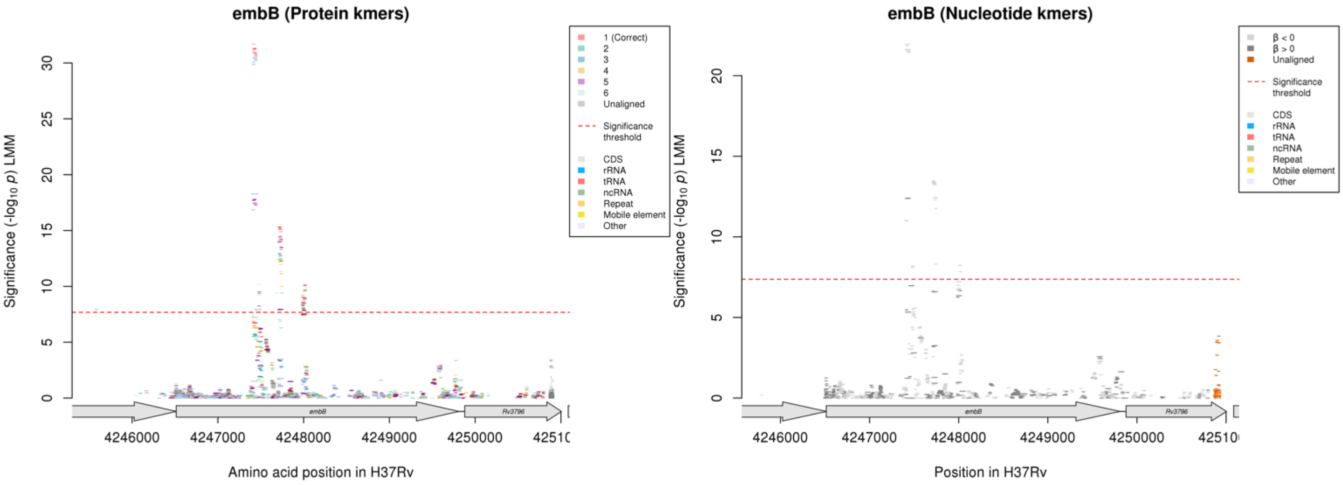

Protein kmers:

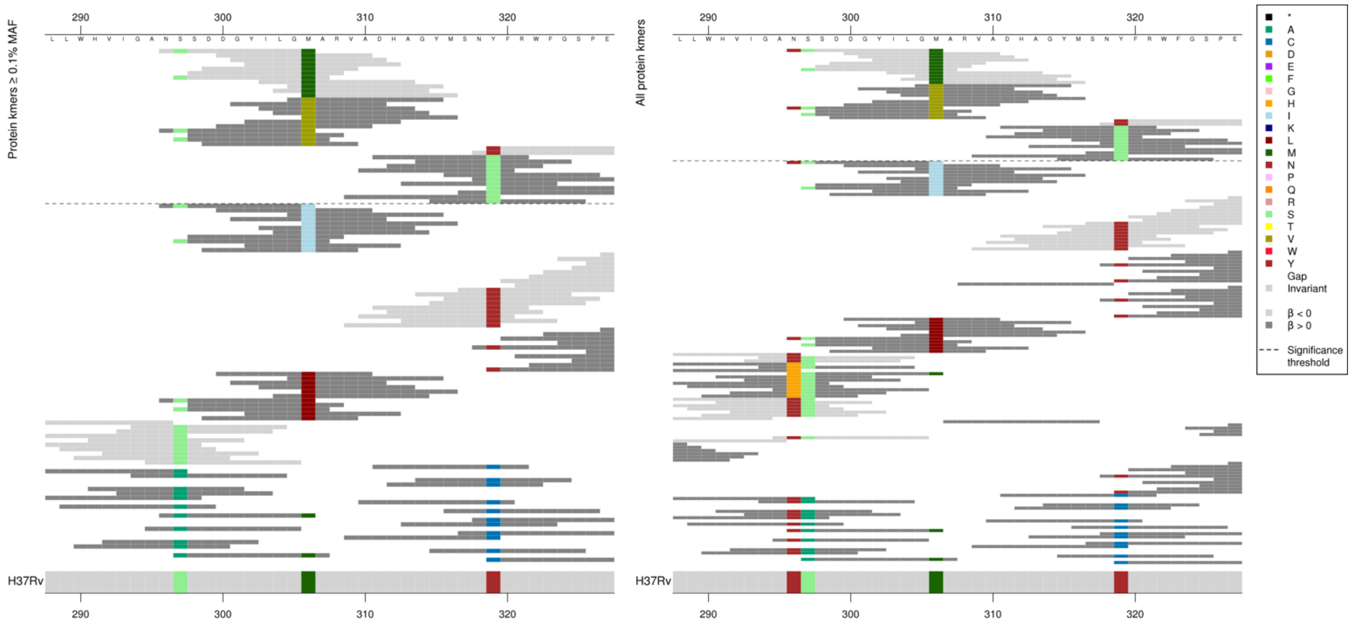

Focusing on the kmers centred on codon 306 from this alignment, and those around 319 in the next alignment.

Range of *p*-values for the low MIC-associated 306M kmers, but changes in *p*-values did not directly correspond with whether they covered codons 296-297. High MIC-associated 306V kmers also varied slightly.

| Kmer | $-\log_{10} p$ | β | MAC | MAF | Ps | Variants |
|---|---|---|---|---|---|---|
| NSSDDGYILGM | 31.69 | -1.38 | 1675 | 19.95 | 296 | 306M |
| GVARVADHAGY | 18.28 | 1.24 | 891 | 10.61 | 305 | 306V |

Protein kmers:

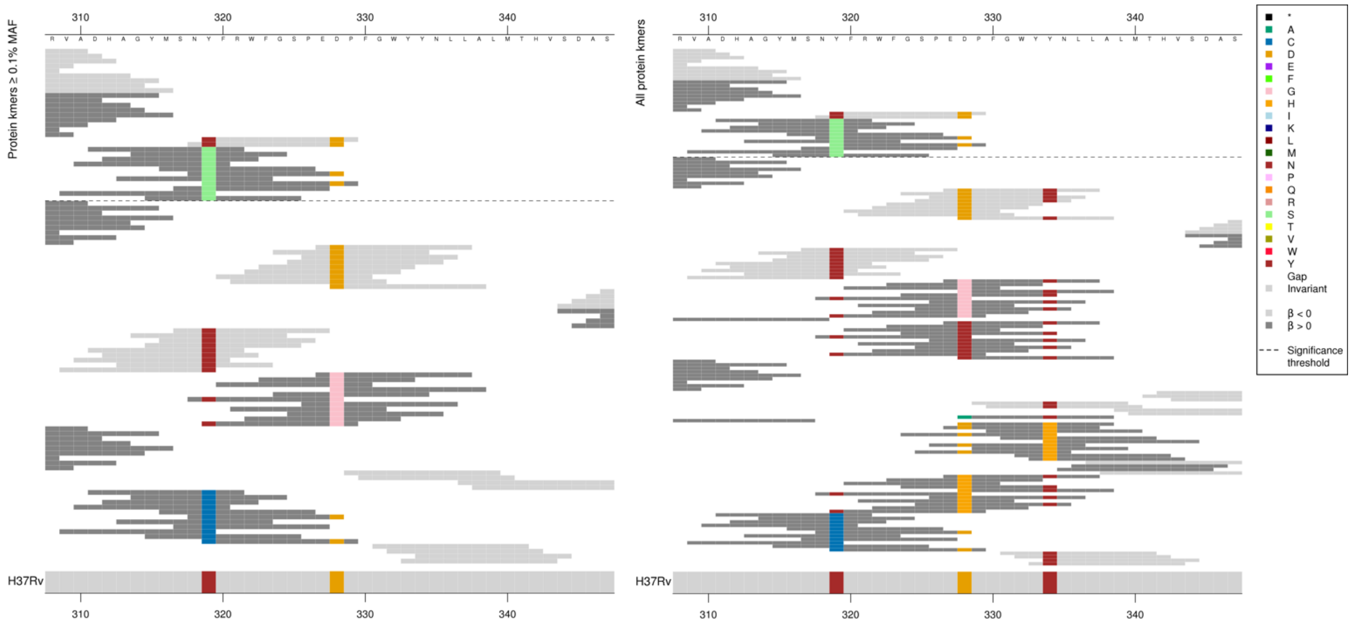

This variant is not significant for RFB.

Kmers below significance threshold = 319C, 328G/Y/H

| Kmer | $-\log_{10} p$ | β | MAC | MAF | Ps | Variants |
|---|---|---|---|---|---|---|
| YFRWFGSPEDP | 9.51 | -1.54 | 159 | 1.89 | 319 | 319Y, 328D |
| DHAGYMSNSFR | 7.92 | 2.46 | 109 | 1.30 | 311 | 319S |

Protein kmers:

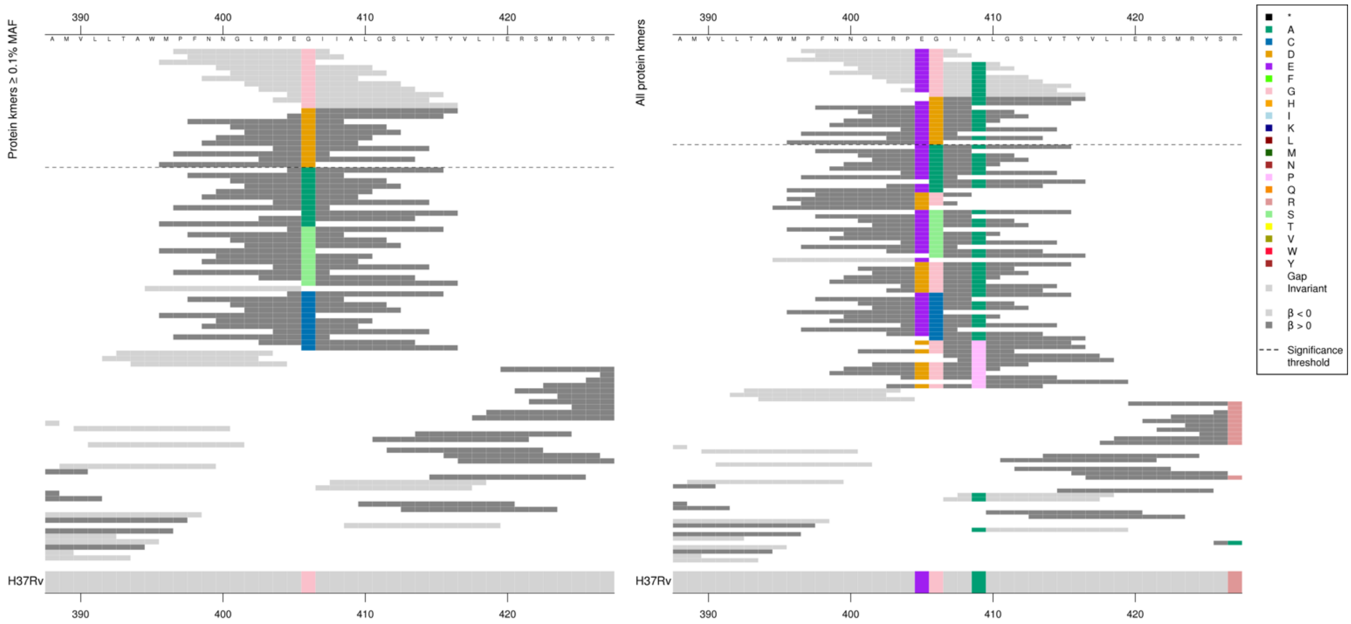

There were no significant high MIC-associated kmers for EMB for this variant.
All of the high MIC-associated kmers here were the same *p*-value except for the last which was slightly lower, with a MAC one lower at 73.

| Kmer | $-\log_{10} p$ | β | MAC | MAF | Ps | Variants |
|---|---|---|---|---|---|---|
| PFNNGLRPEGI | 15.31 | -1.21 | 471 | 5.61 | 397 | 405E, 406G |
| NGLRPEGIIAL | 15.21 | -1.20 | 474 | 5.65 | 400 | 405E, 406G, 409A |
| GIIALGSLVTY | 13.49 | -1.14 | 469 | 5.59 | 406 | 406G, 409A |
| DIIALGSLVTY | 7.93 | 1.46 | 74 | 0.88 | 406 | 406D |

Protein kmers:

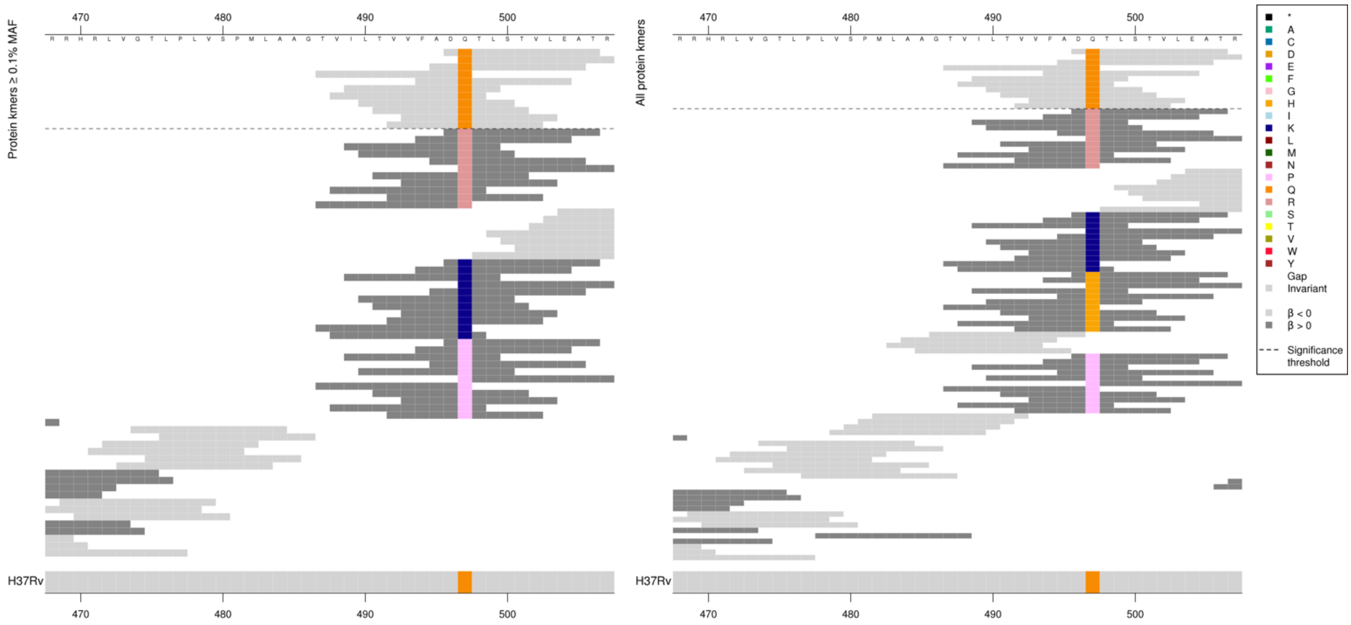

| Kmer | $-\log_{10} p$ | β | MAC | MAF | Ps | Variants |
|---|---|---|---|---|---|---|
| DQTLSTVLEAT | 10.12 | -1.07 | 353 | 4.21 | 496 | 497Q |

# *Rv1565c*

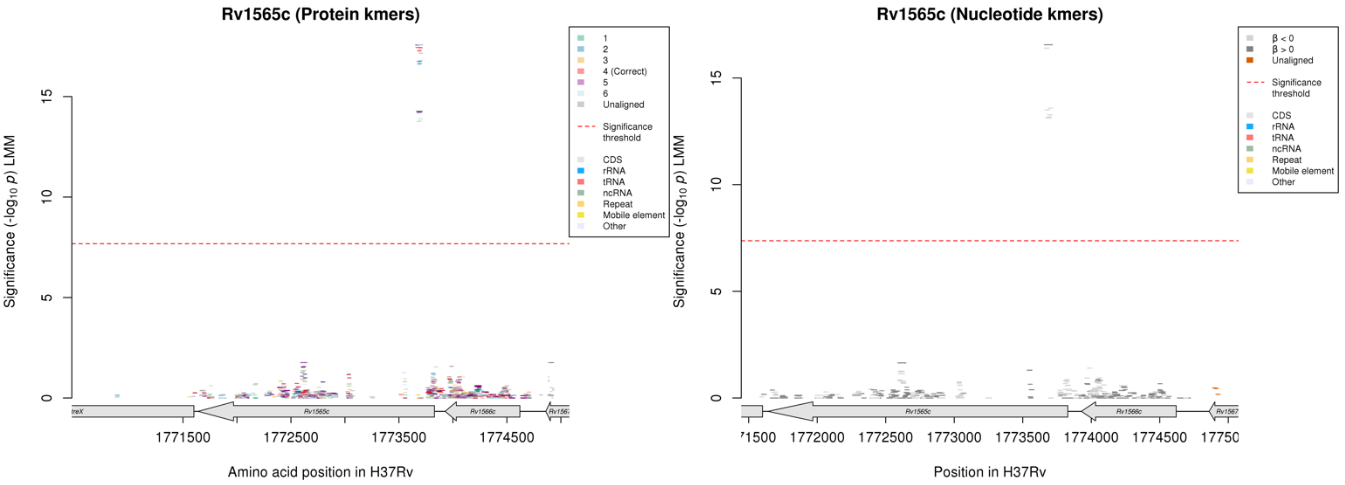

#### Protein kmers:

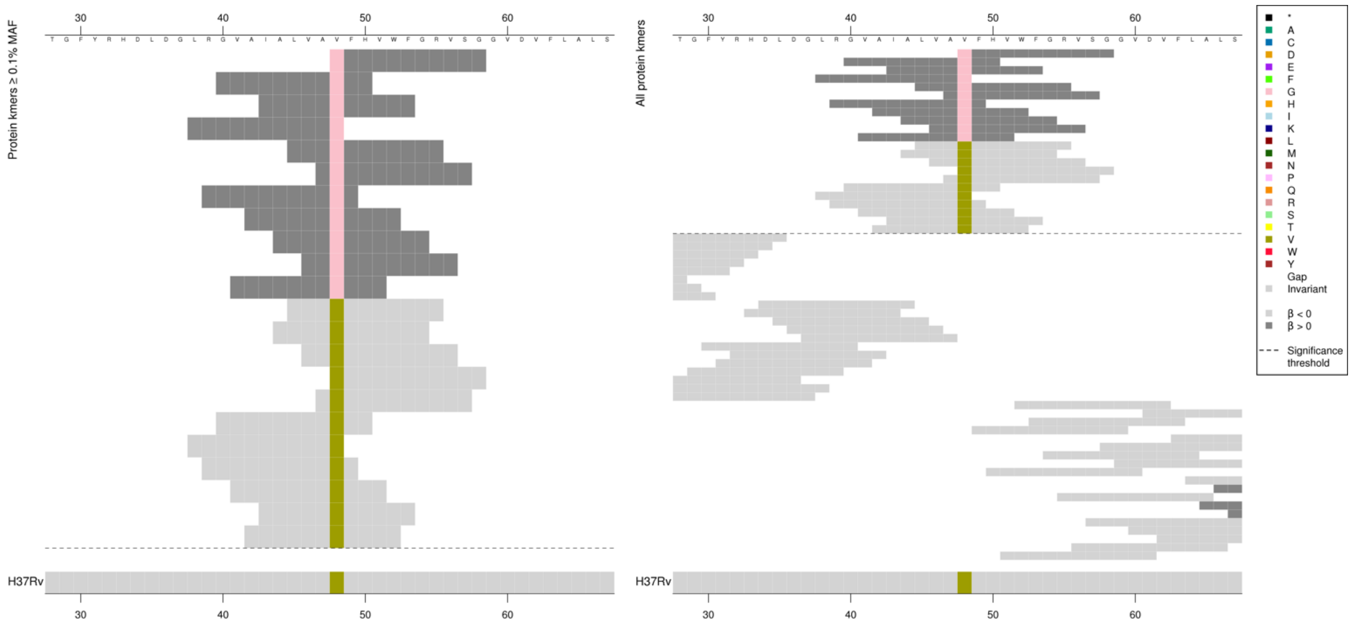

| Kmer | $-\log_{10} p$ | β | MAC | MAF | Ps | Variants |
|---|---|---|---|---|---|---|
| GFHVWFGRVSG | 17.58 | 4.05 | 114 | 1.36 | 48 | 48G |
| LVAVFHVWFGR | 17.44 | -4.01 | 117 | 1.39 | 45 | 48V |

### guaA

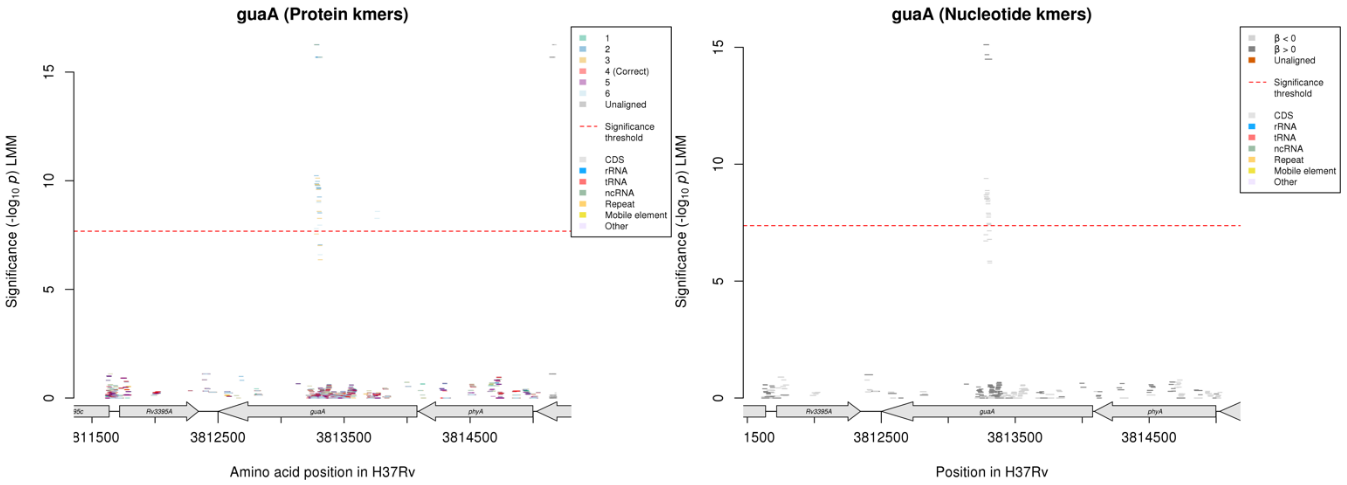

Nucleotide kmers:

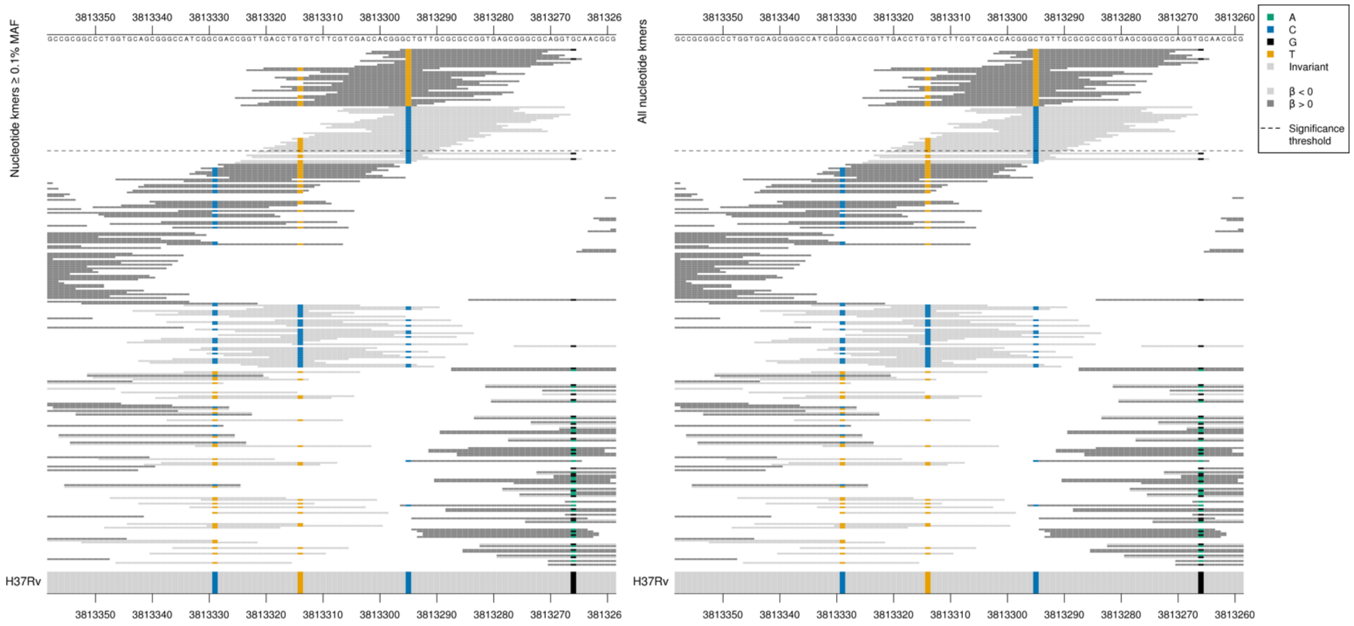

First 9 high MIC-associated kmers had the same *p*-value, the 10th was slightly less significant, then the remaining 21 were all the same *p*-value regardless of whether they covered the variant at position 3813314. The low MIC-associated kmers were a range of *p*-values, showing the top for the two allele combinations.

Synonymous mutation at position 3813295/784.
High MIC-associated significant kmers = TTG = Leu / L
Low MIC-associated kmers = CTG = Leu / L

Synonymous mutation at position 3813314/765.
High MIC-associated significant kmers = TGT = Cys / C
Kmers below significance threshold = TGC = Cys / C

| Kmer | $-\log_{10} p$ | β | MAC | MAF | Ps | Variants |
|---|---|---|---|---|---|---|
| GTTGTTGCGCGCCGGTGAGCGGGCGCAGGTG | 15.11 | 3.66 | 107 | 1.27 | 3813296 | 3813295/784T |
| GGGCTGTTGCGCGCCGGTGAGCGGGCGCAGG | 9.39 | -2.12 | 292 | 3.48 | 3813298 | 3813295/784C |
| GTGTCTTCGTCGACCACGGGCTGTTGCGCGC | 7.91 | -1.82 | 349 | 4.16 | 3813315 | 3813314/765T, 3813295/784C |

#### *ctpI*

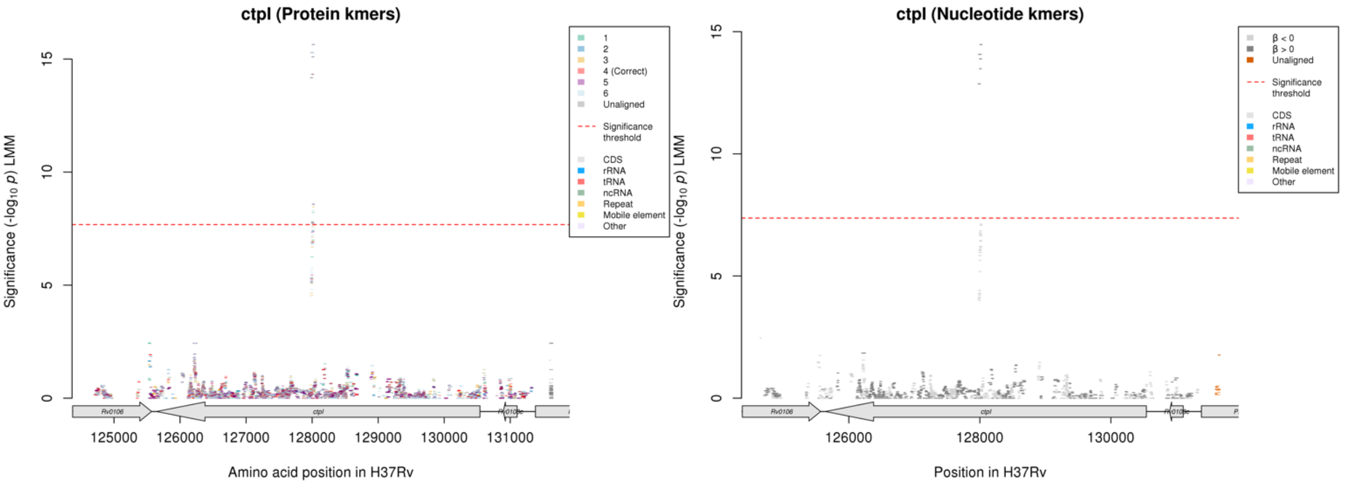

Nucleotide kmers:

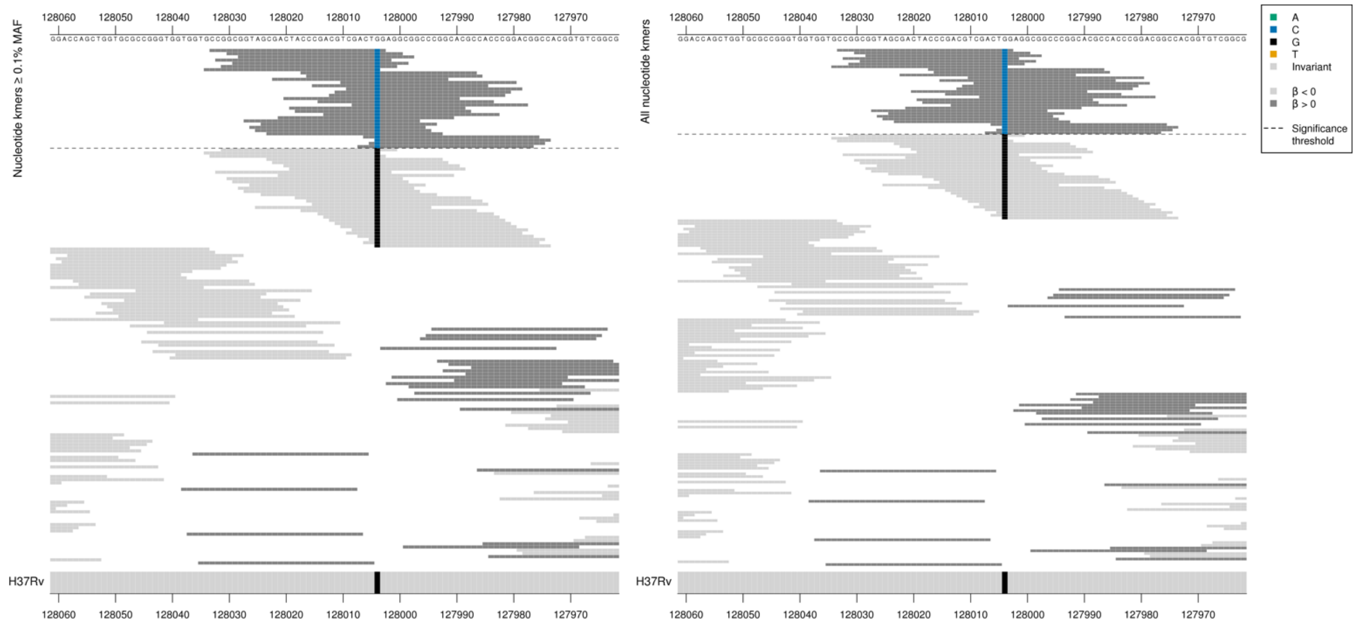

Gene starts at position 130541.
Synonymous mutation at position 128004/2538
High MIC-associated kmers = CTC = Leu / L
Kmers below significance threshold = CTG = Leu / L

| Kmer | $-\log_{10} p$ | β | MAC | MAF | Ps | Variants |
|---|---|---|---|---|---|---|
| CCGGCGGTAGCGACTACCCGACGTCGACTCG | 14.47 | 3.54 | 107 | 1.27 | 128033 | 128004/2538C |

### *spoU*

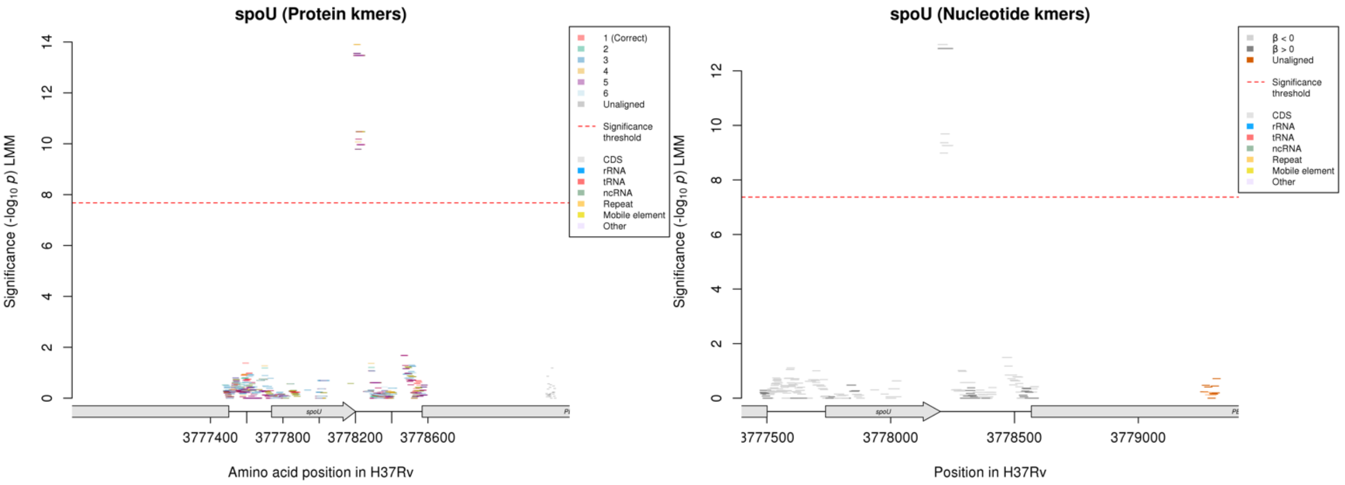

Nucleotide kmers:

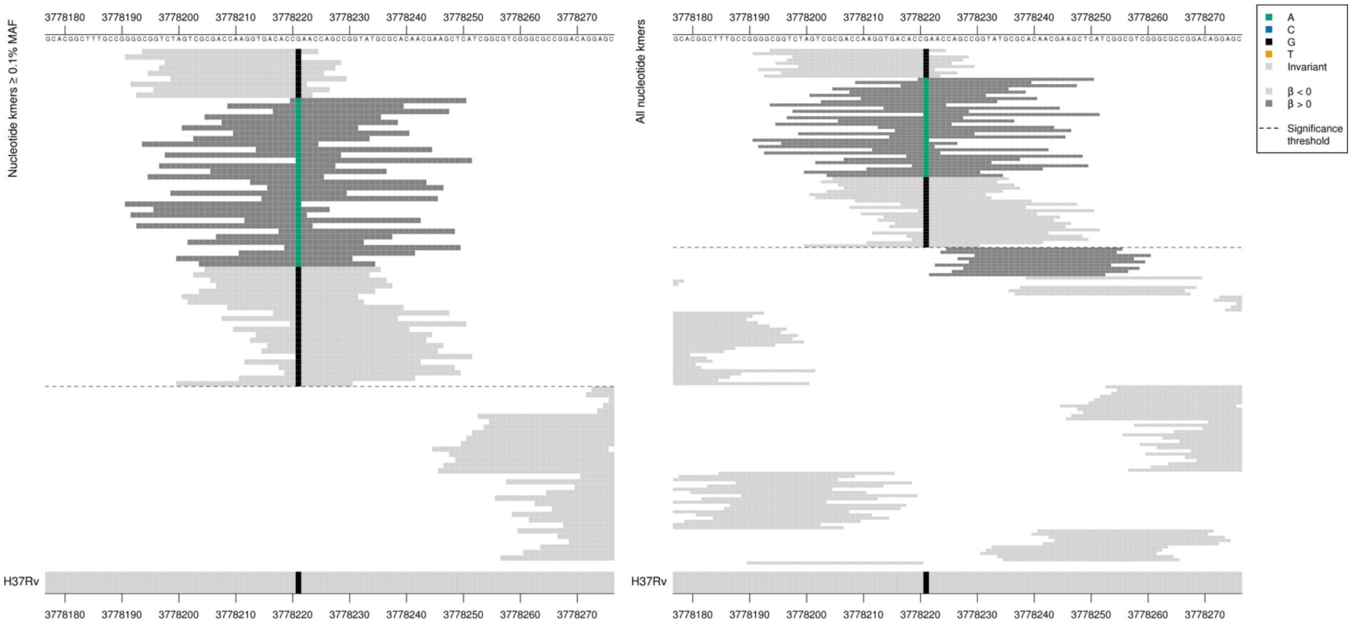

*spoU* ends at position 3778201, the kmers overlap this position. PE_*PGRS51* is the following gene but does not start until 3778568, this variant is 347 bases upstream.
Mutation 20bp downstream of the stop codon (3778221).

| Kmer | -log$_{10}$ *p* | β | MAC | MAF | Ps | Variants |
|---|---|---|---|---|---|---|
| CGGTCTAGTCGCGACCAAGGTGACACCGAAC | 12.96 | -3.07 | 163 | 1.94 | 3778194 | 3778221G (20bp after stopcodon) |
| CAAACCAGCCGGTATGCGCACAACGAAGCTC | 12.82 | 3.19 | 159 | 1.89 | 3778220 | 3778221A (20bp after stopcodon) |

### *dxs2:Rv3382c* [R]

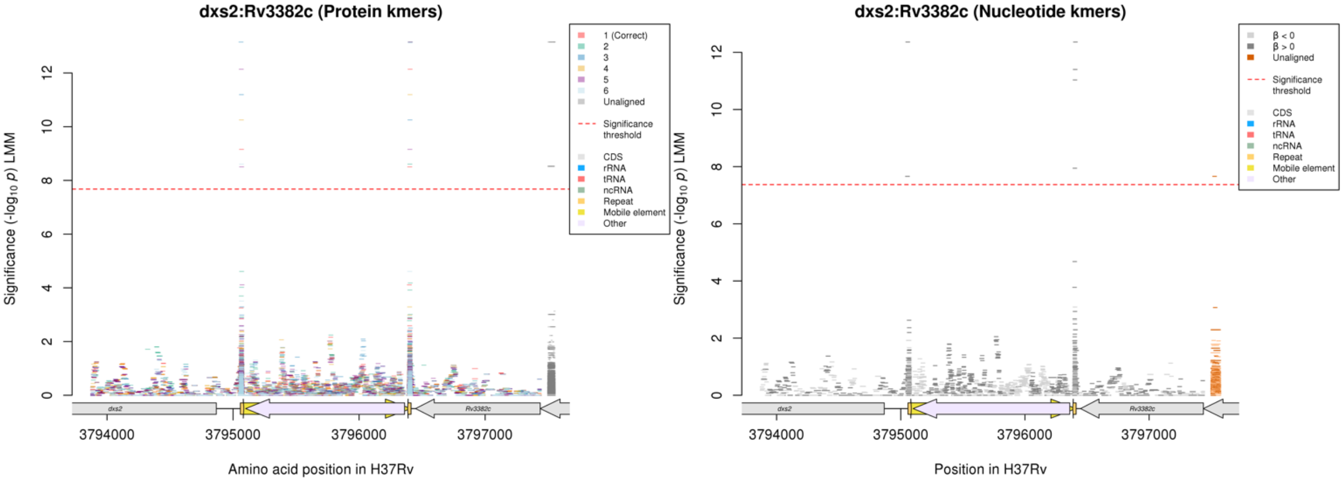

#### Repeat regions at the ends of mobile elements

Nucleotide kmers:

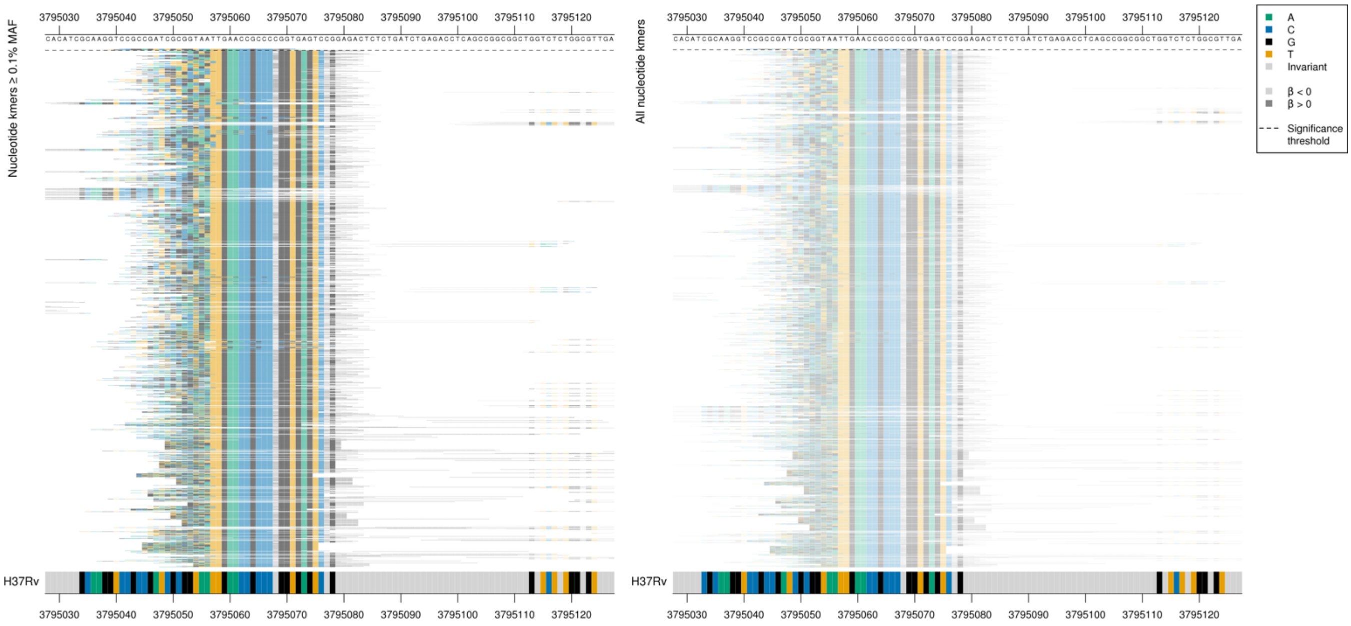

| Kmer | $-\log_{10} p$ | β | MAC | MAF | Ps | Variants |
|---|---|---|---|---|---|---|
| CCGGAGTCTGTGGTCATTGAACCGCCCCGGC | 12.36 | 3.01 | 118 | 1.41 | 3795041 | Repeat region |

#### Nucleotide kmers:

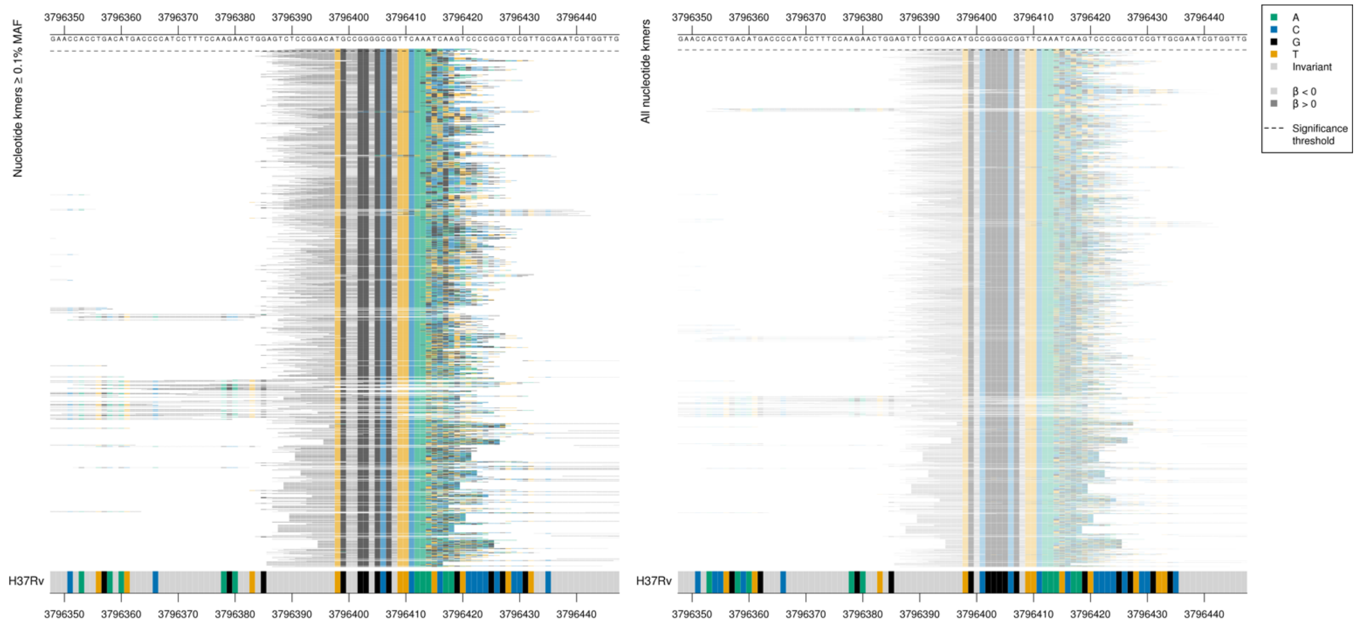

| Kmer | $-\log_{10} p$ | β | MAC | MAF | Ps | Variants |
|---|---|---|---|---|---|---|
| CCGGACATGCCGGGGCGGTTCAATGACCACA | 12.36 | 3.01 | 118 | 1.41 | 3796391 | Repeat region |

## *Rv3183:Rv3188* [R]

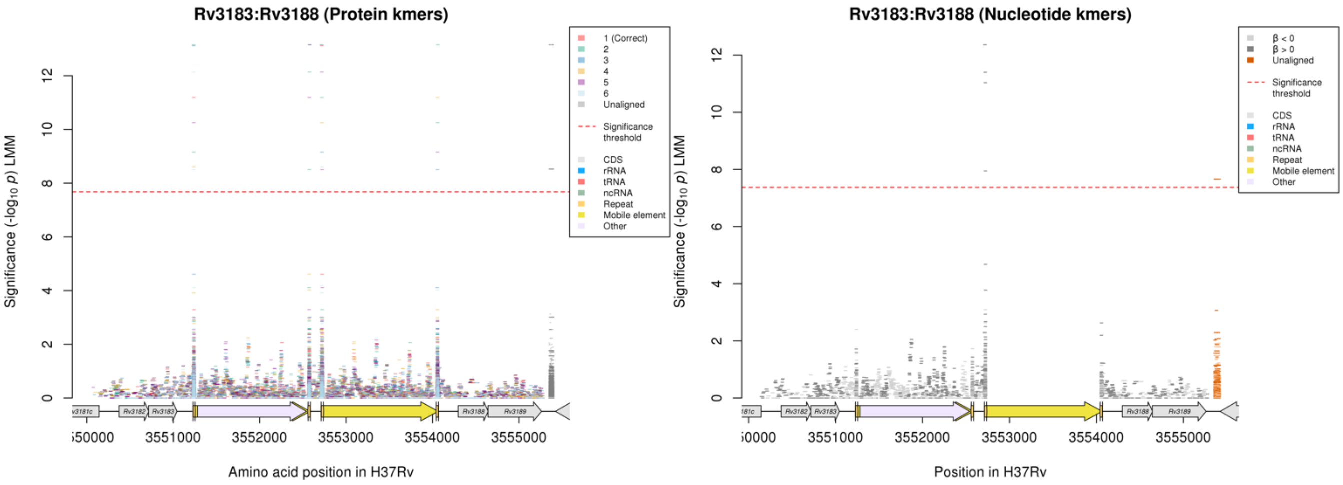

Repeat regions at the ends of mobile elements

#### Nucleotide kmers:

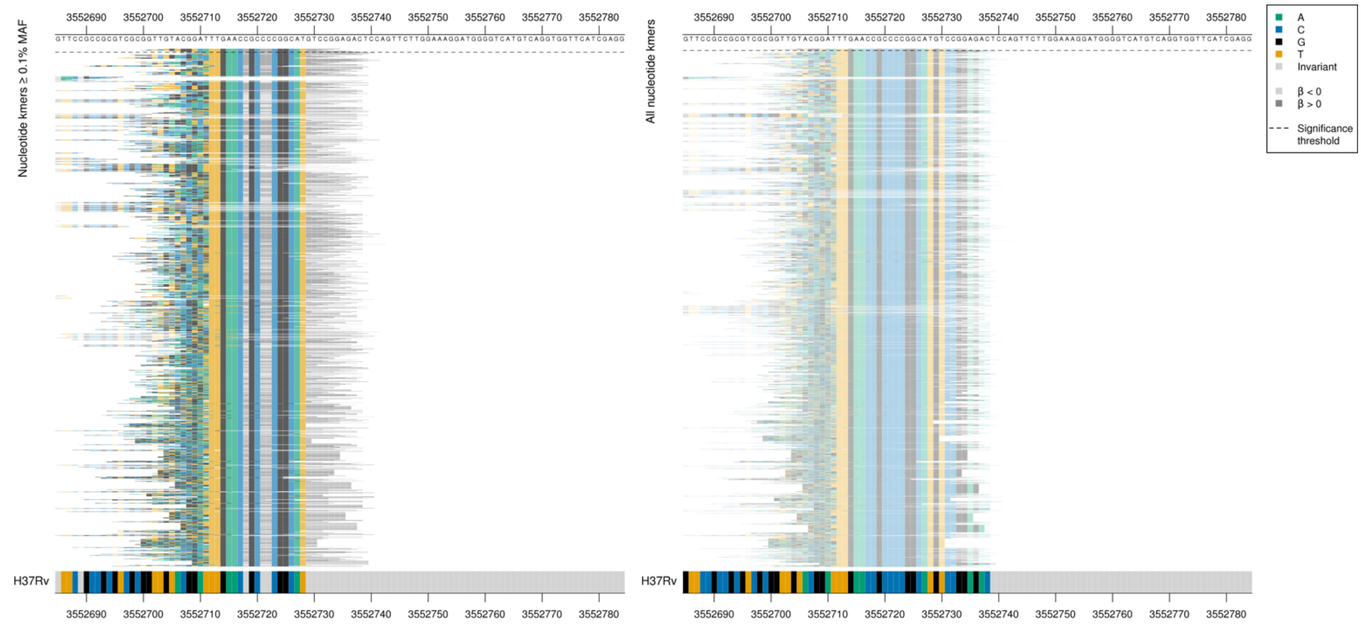

| Kmer | -log$_{10}$ *p* | β | MAC | MAF | Ps | Variants |
|---|---|---|---|---|---|---|
| TGTGGTCATTGAACCGCCCCGGCATGTCCGG | 12.36 | 3.01 | 118 | 1.41 | 3552704 | Repeat region |

#### *relA*

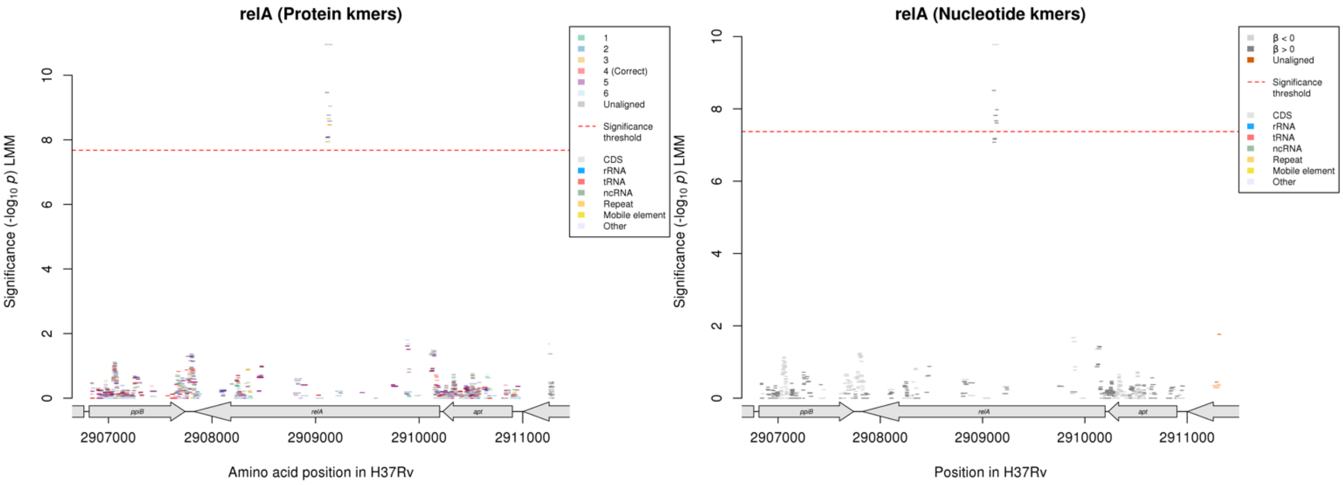

#### Nucleotide kmers:

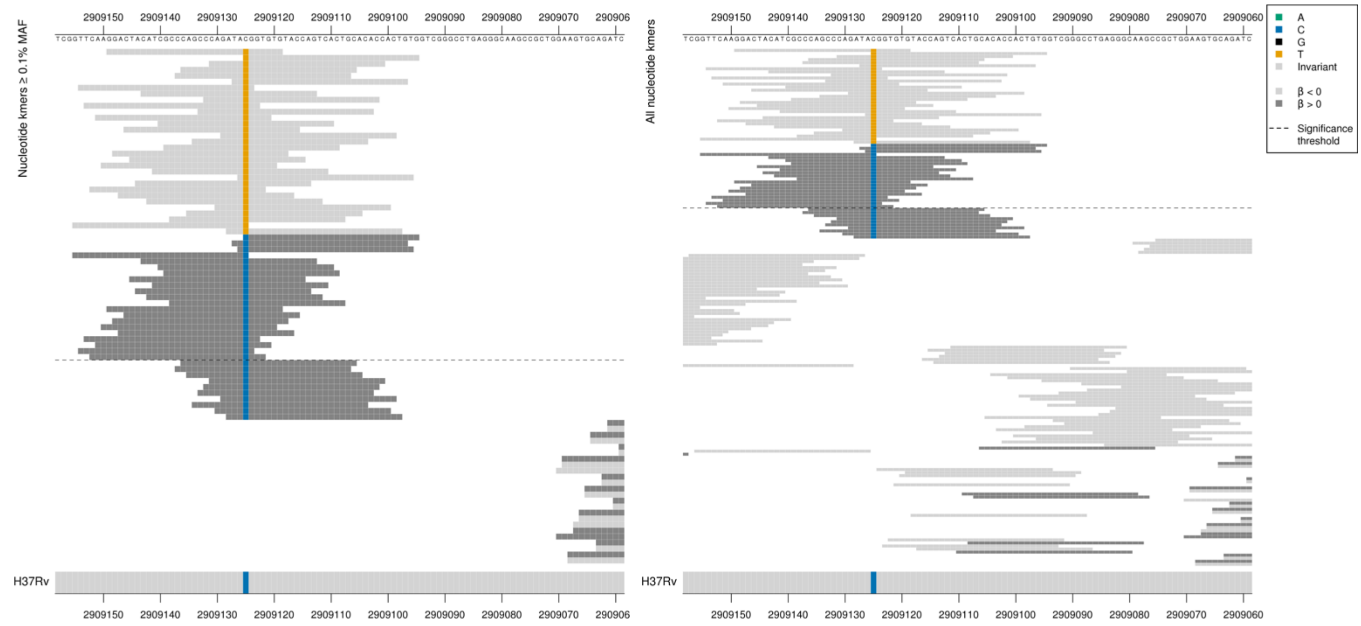

Synonymous mutation at position 2909125/1074. Low MIC-associated kmers were different to reference.
High MIC-associated kmers = TAC = Tyr / Y
Low MIC-associated kmers = TAT = Tyr / Y

| Kmer | $-\log_{10} p$ | β | MAC | MAF | Ps | Variants |
|---|---|---|---|---|---|---|
| GGACTACATCGCCCAGCCCAGATATGGTGTG | 9.78 | -3.40 | 16 | 0.19 | 2909149 | 2909125/1074T |
| CGGTGTGTACCAGTCACTGCACACCACTGTG | 8.51 | 2.95 | 19 | 0.23 | 2909125 | 2909125/1074C |

### *proA:ahpC*

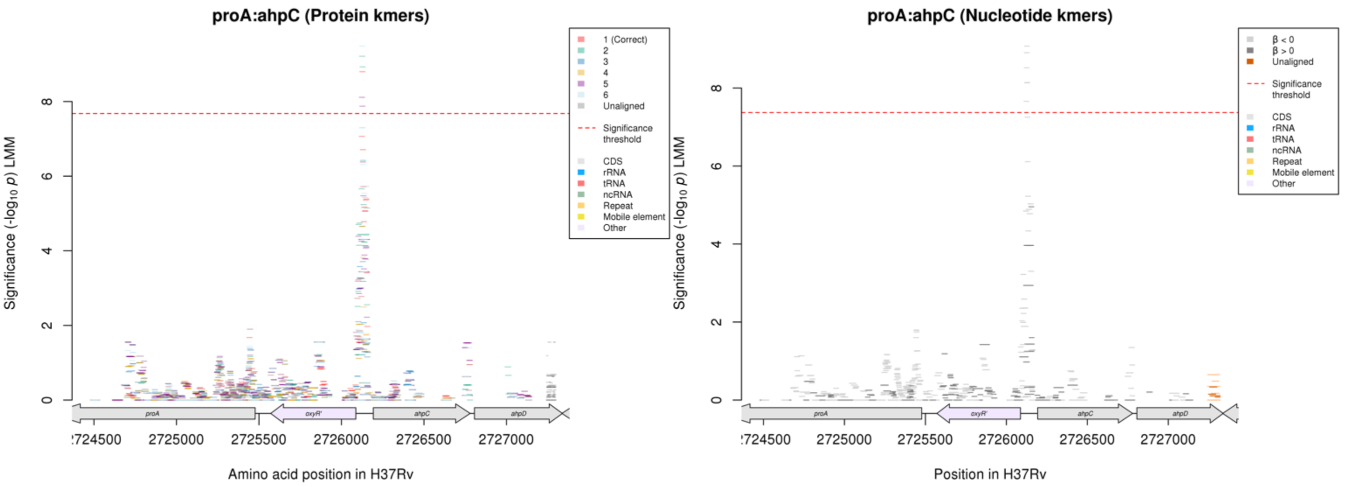

Nucleotide kmers:

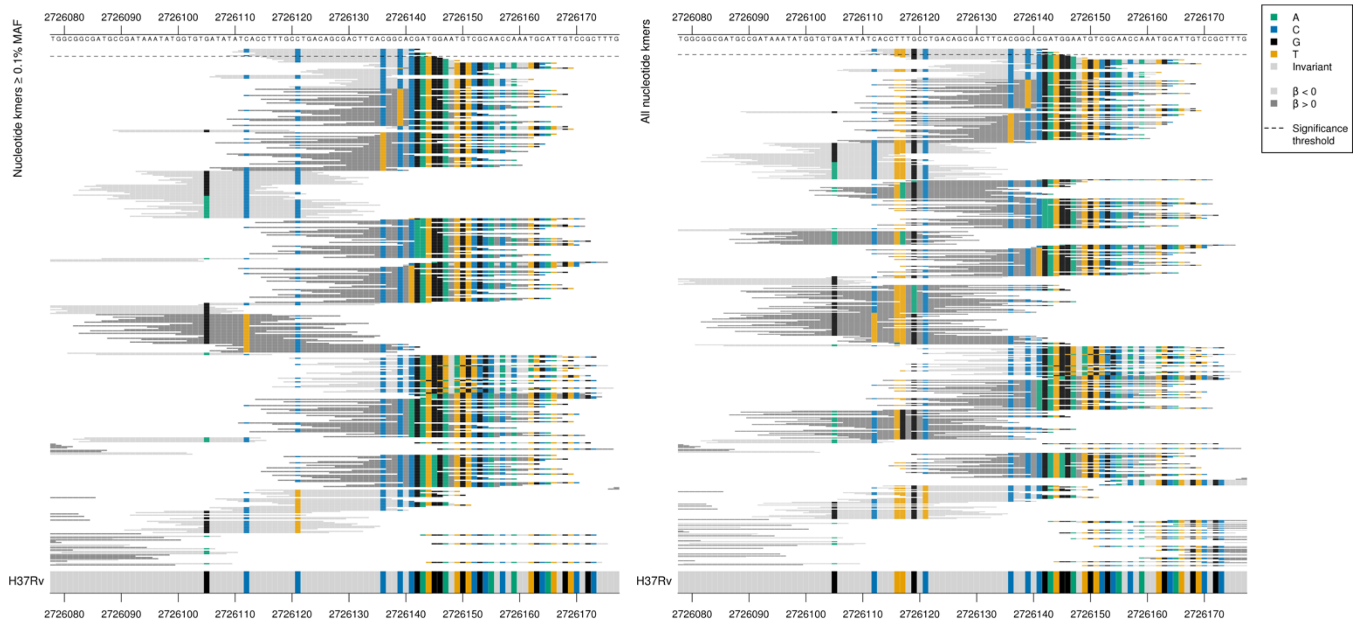

oxyR' begins at position 2726087, *ahpC* begins at position 2726193, so this is in the intergenic region upstream of both.

Significant low MIC-associated kmers were all the same as the reference. See result for INH for further interpretation of the variation captured by the kmers which in this analysis were high MIC and not significant.

| Kmer | $-\log_{10} p$ | β | MAC | MAF | Ps | Variants |
|---|---|---|---|---|---|---|
| CACCTTTGCCTGACAGCGACTTCACGGCACG | 9.07 | -1.22 | 144 | 1.72 | 2726112 | WT / 2726112C, 2726121C. 2726136C, 2726139C, 2726141C, 2726142G |
| TCACCTTTGCCTGACAGCGACTTCACGGCAC | 8.91 | -1.29 | 128 | 1.52 | 2726111 | WT / 2726112C, 2726121C. 2726136C, 2726139C, 2726141C |
| ATCACCTTTGCCTGACAGCGACTTCACGGCA | 8.52 | -1.54 | 77 | 0.92 | 2726110 | WT / 2726112C, 2726121C. 2726136C, 2726139C |
| CCTTTGCCTGACAGCGACTTCACGGCACGAT | 8.14 | -1.19 | 134 | 1.60 | 2726114 | WT / 2726121C. 2726136C, 2726139C, 2726141C, 2726142G, 2726143A, 2726144T |
| ACCTTTGCCTGACAGCGACTTCACGGCACGA | 8.13 | -1.18 | 135 | 1.61 | 2726113 | WT / 2726121C. 2726136C, 2726139C, 2726141C, 2726142G, 2726143A |
| TATCACCTTTGCCTGACAGCGACTTCACGGC | 7.66 | -1.48 | 75 | 0.89 | 2726109 | WT / 2726112C, 2726121C. 2726136C, 2726139C |

#### *fabG1*

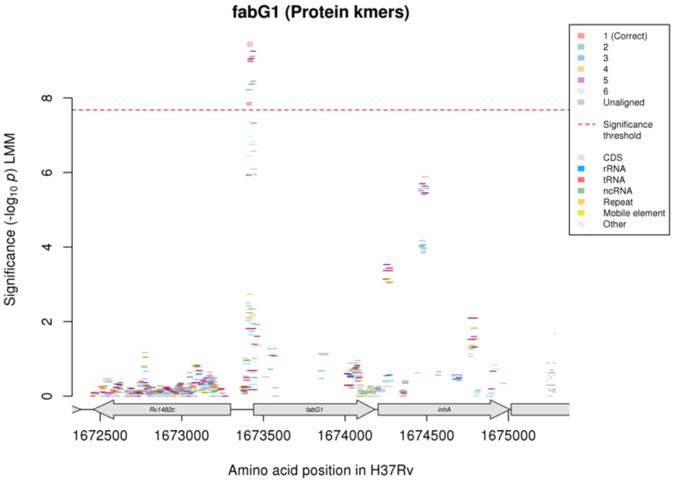

Not significant in the nucleotide kmer analysis.

Protein kmers:

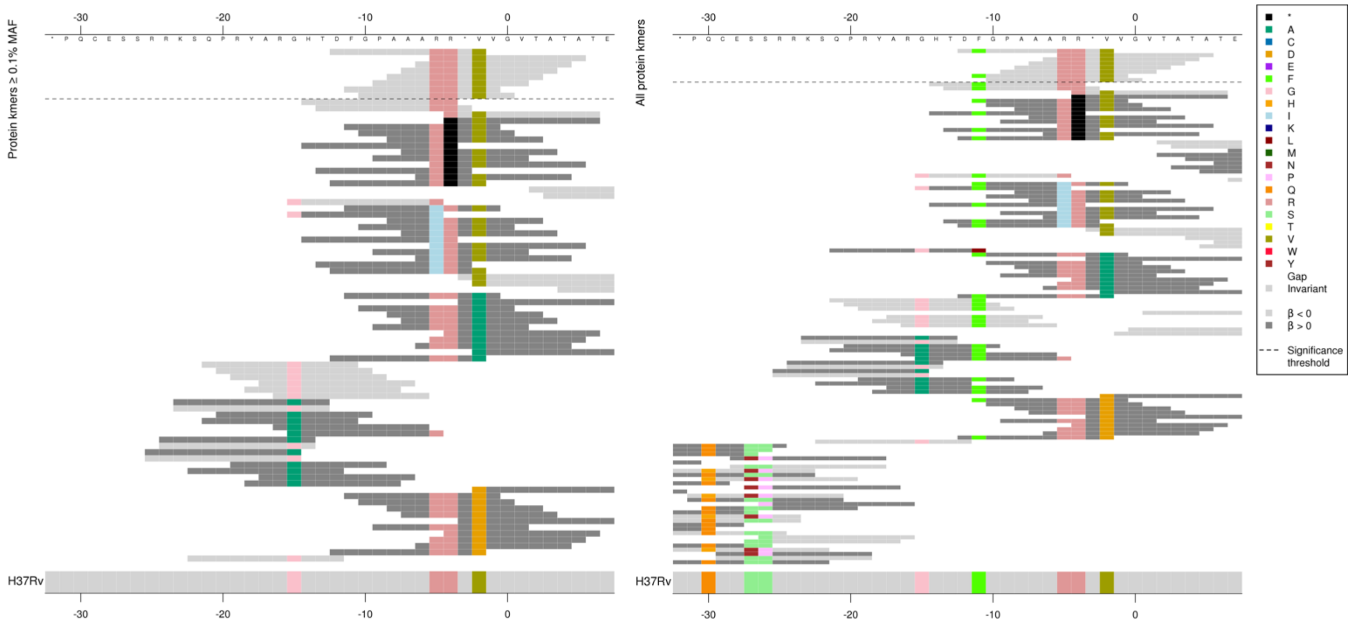

The kmers were capturing upstream of the start codon. The significant kmers were capturing the reference sequence.

| Kmer | -log₁₀ p | β | MAC | MAF | Ps | Variants |
|---|---|---|---|---|---|---|
| DFGPAAARR*V | 9.39 | -0.80 | 1074 | 12.79 | -12 | WT – out of frame, upstream of gene |

#### *moaC3:Rv3327*[R]

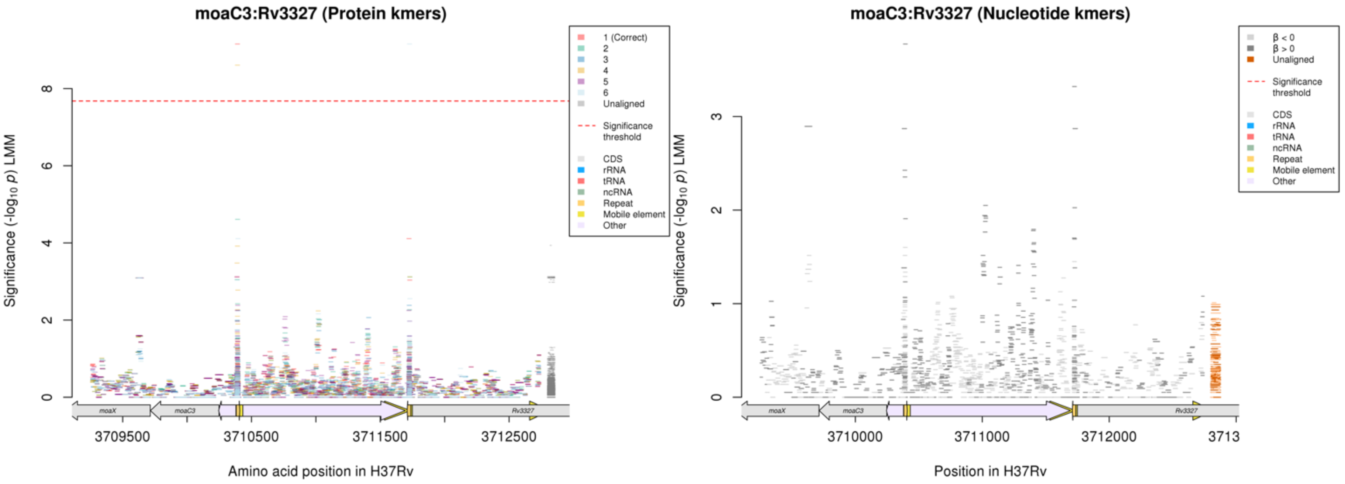

Not significant in the nucleotide kmer analysis.
Repeat region at the end of a mobile element.

| Kmer | -log₁₀ p | β | MAC | MAF | Ps | Variants |
|---|---|---|---|---|---|---|
| WSLNRPGMSGD | 9.15 | 2.06 | 134 | 1.60 | - | Repeat region, out of frame as in intergenic region |

## *Rv0810c*

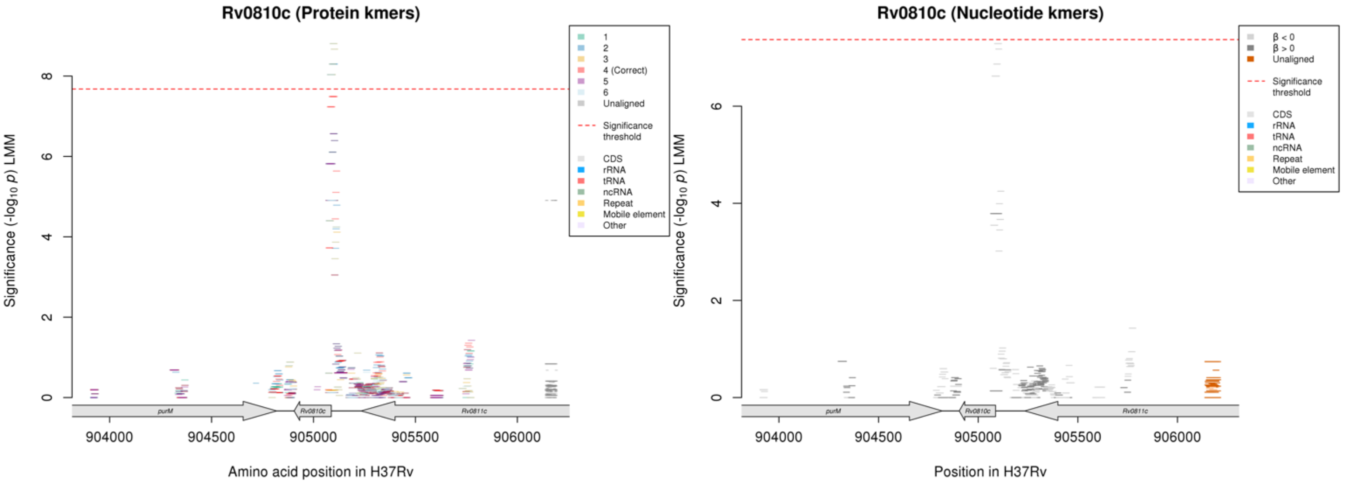

Not significant in nucleotide kmer analysis, just below the significance threshold.
Capturing variants upstream of the start codon. Although not significant in the nucleotide kmer analysis, alignments are below:

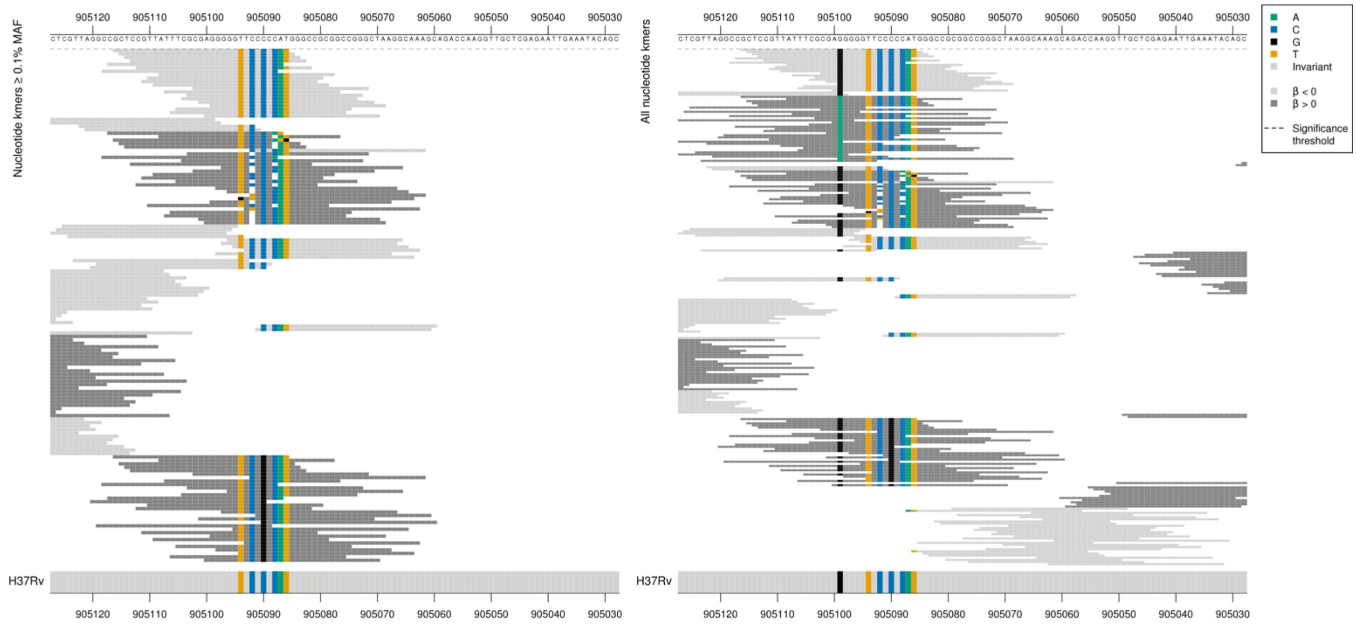

*Rv0810c* begins at position 905087.

Kmers below significance threshold = C deletion in the ‘CCCCC’ region positions -1 to -5 (905088-905092) directly upstream of the start codon

Kmers below significance threshold = 905099/-12 A

| Kmer | $-\log_{10} p$ | β | MAC | MAF | Ps | Variants |
|---|---|---|---|---|---|---|
| GCTCCGTTATTTCGCGAGGGGGTTCCCCCAT | 7.29 | -1.62 | 61 | 0.73 | 905116 | WT |

#### *fadD9*

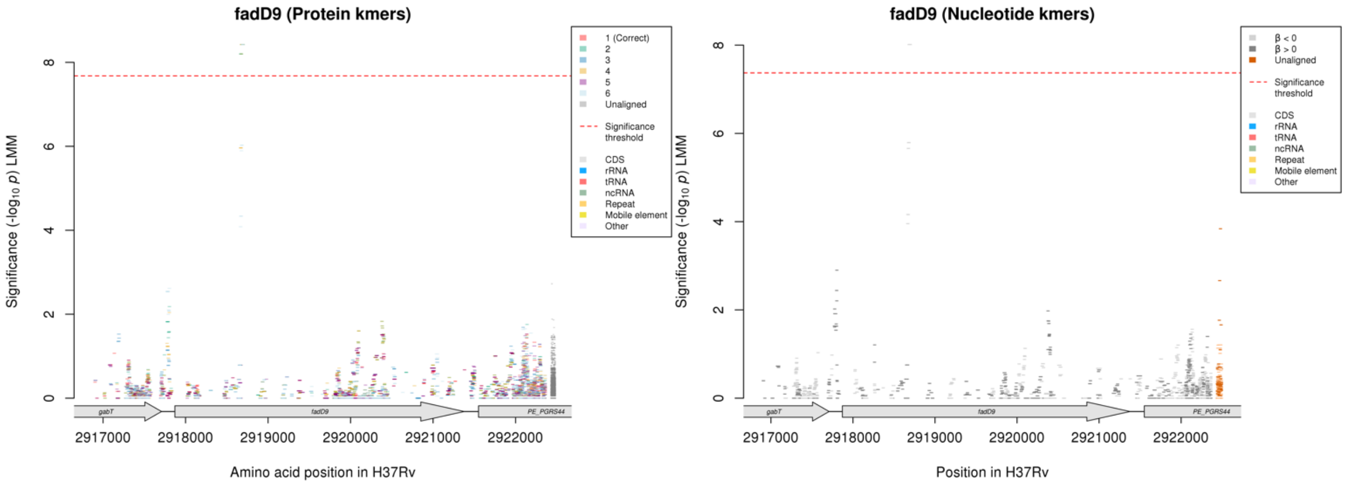

No significant kmers aligned to the correct reading frame in the protein kmer analysis.

Nucleotide kmers:

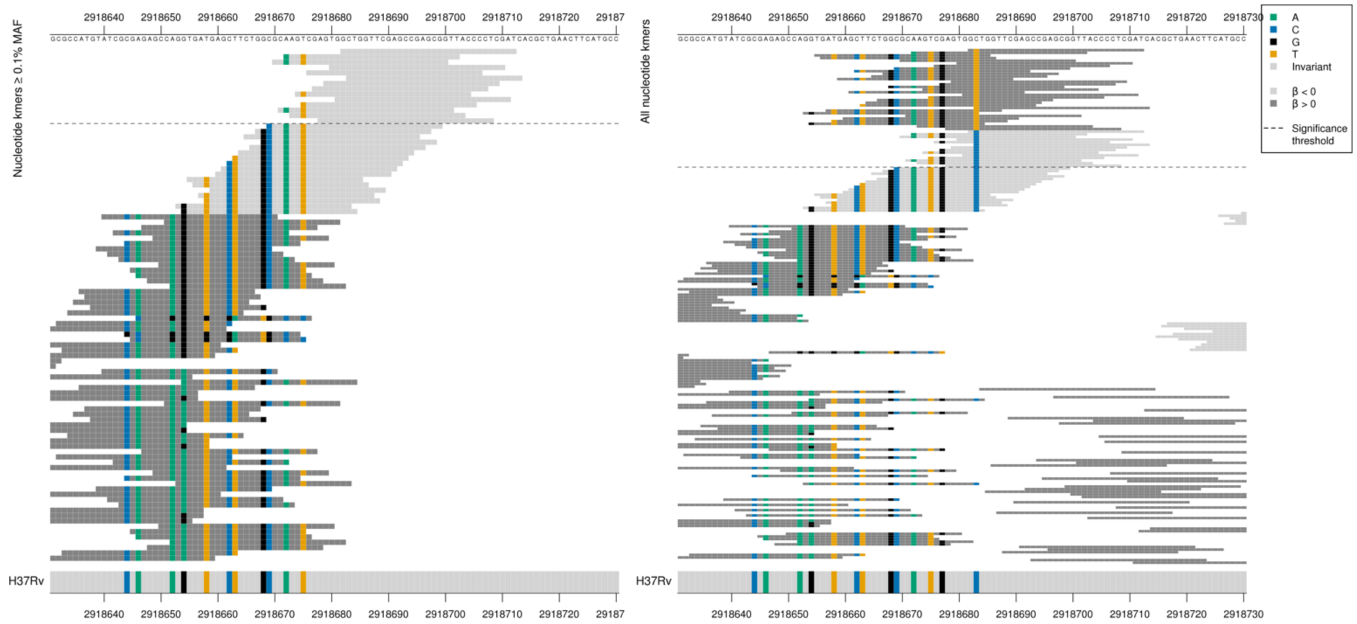

High MIC-associated kmers = GGT = Gly / G
Low MIC-associated kmers = GGC = Gly / G

Low MAF kmers = 2918683T

| Kmer | $-\log_{10} p$ | β | MAC | MAF | Ps | Variants |
|---|---|---|---|---|---|---|
| GCTGGTTCGAGCCGAGCGGTTACCCCTCGAT | 8.01 | -3.71 | 10 | 0.12 | 2918682 | WT / 2918683C |

## *Rv3779*

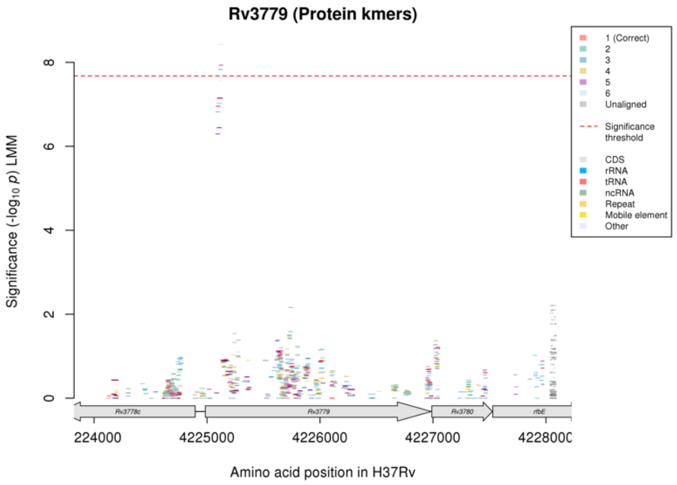

Not significant in nucleotide kmer analysis.

Protein kmers:

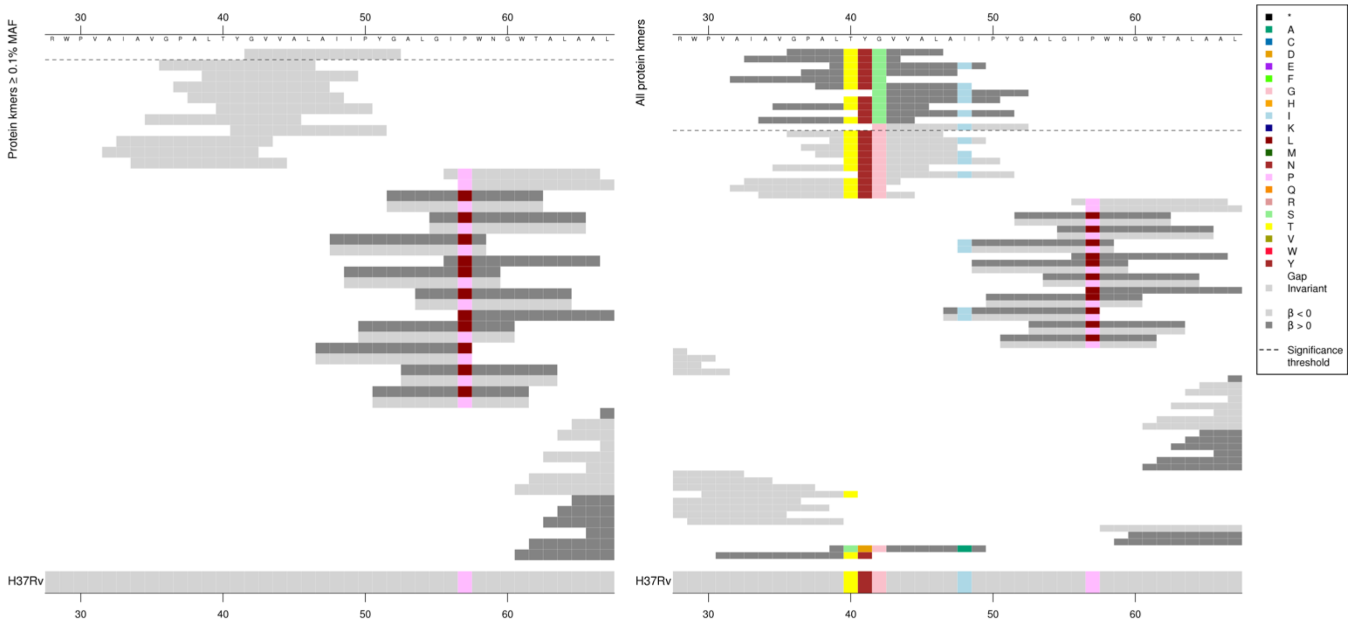

Low MAF kmers = 42S.

| Kmer | $-\log_{10} p$ | β | MAC | MAF | Ps | Variants |
|---|---|---|---|---|---|---|
| GVVALAIIPYG | 7.93 | -3.63 | 9 | 0.11 | 42 | 42G, 48I |

#### *rpsL*

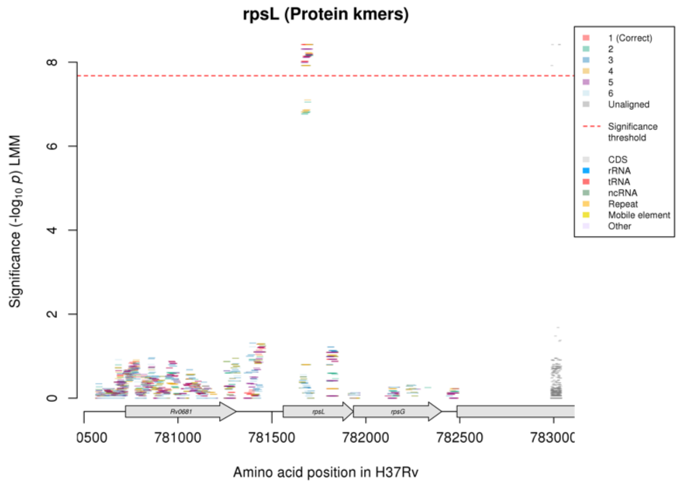

Not significant in the nucleotide kmer analysis.

Protein kmers:

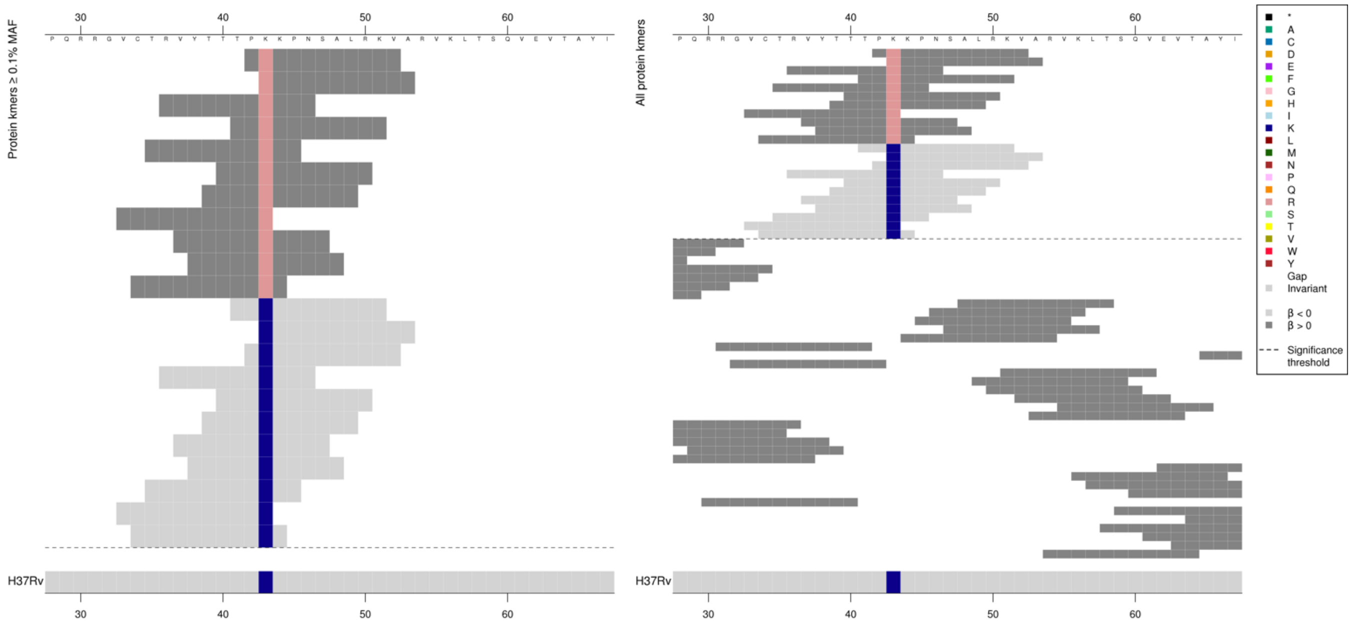

| Kmer | $-\log_{10} p$ | β | MAC | MAF | Ps | Variants |
|---|---|---|---|---|---|---|
| PRKPNSALRKV | 8.42 | 0.85 | 1518 | 18.08 | 42 | 43R |
| TPKKPNSALRK | 8.19 | -0.84 | 1519 | 18.10 | 41 | 43K |

#### *rpoC*

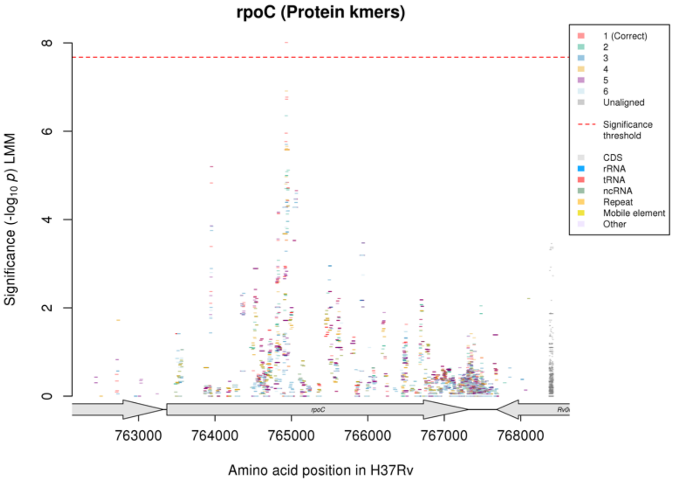

Not significant in the nucleotide kmer analysis.

Protein kmers:

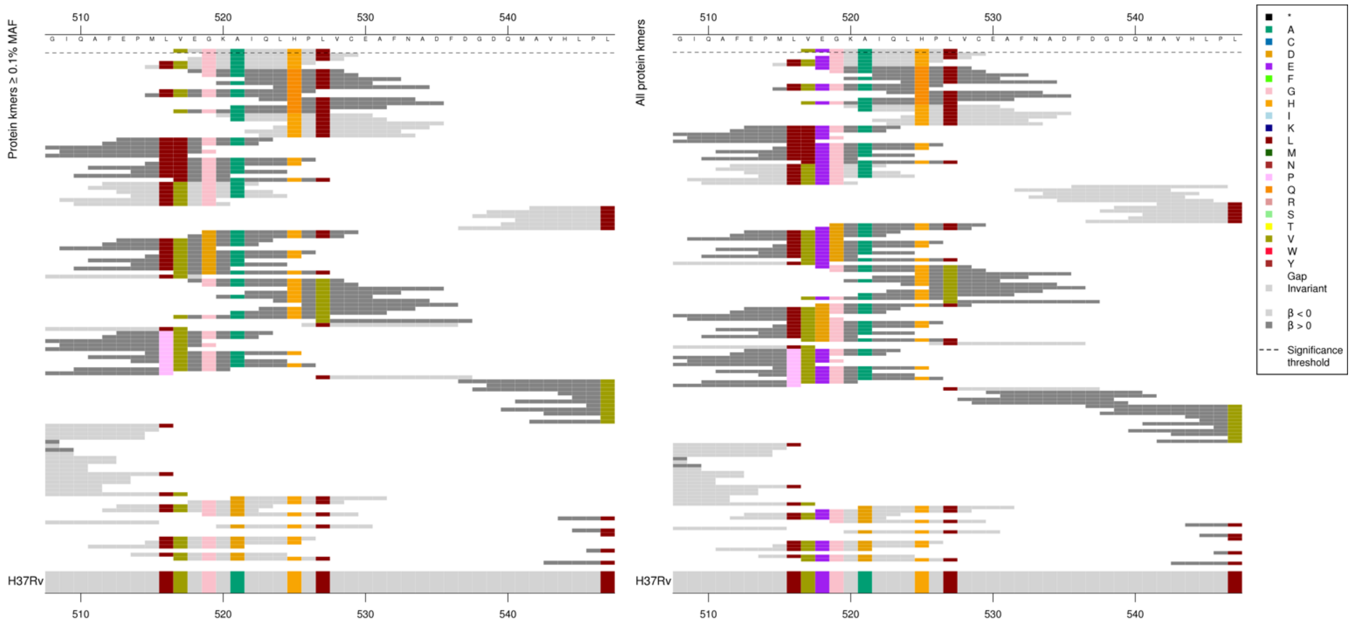

Different region in *rpoC* to the region that is significant for RFB.

| Kmer | -log$_{10}$ *p* | β | MAC | MAF | Ps | Variants |
|---|---|---|---|---|---|---|
| VEGKAIQLHPL | 8.01 | -1.42 | 103 | 1.23 | 517 | WT |

## *Rv2190c:Rv2191*

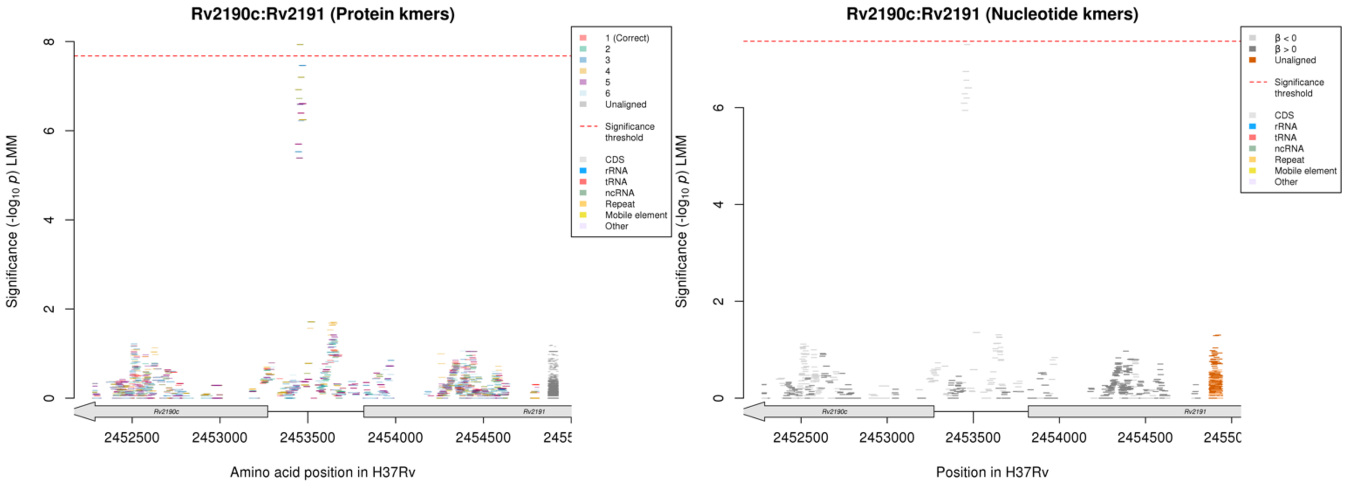

Just below the significance threshold for nucleotide kmers.

Nucleotide kmers:

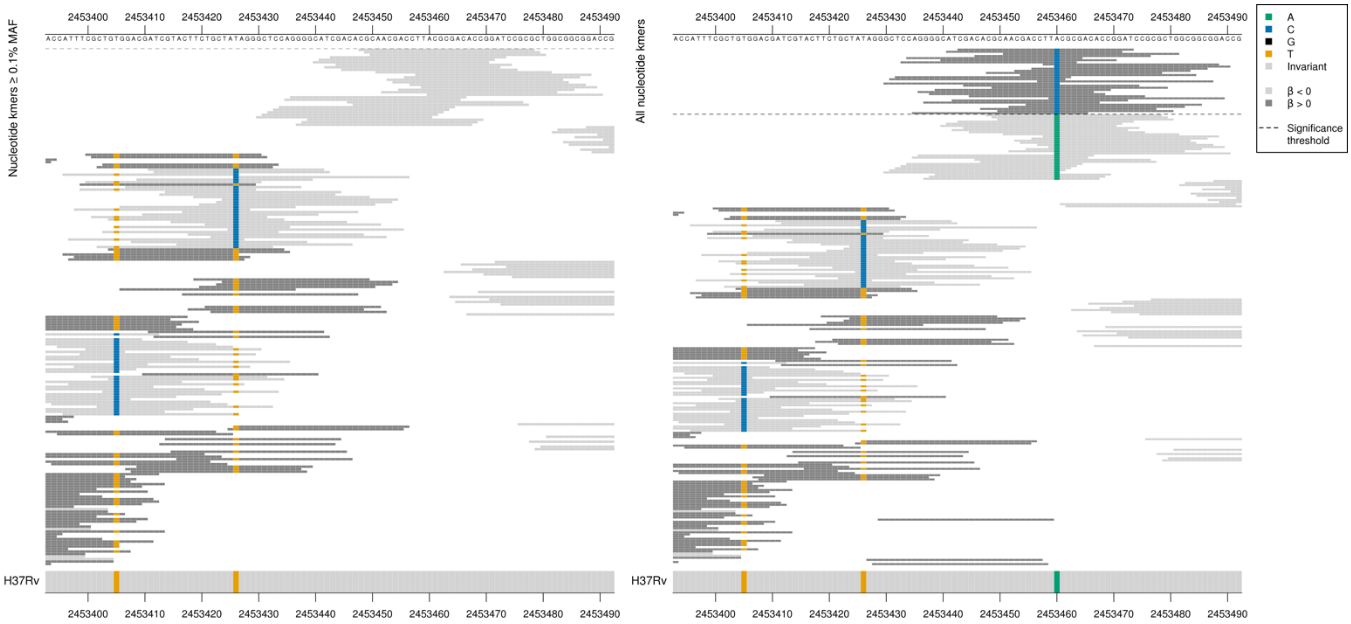

Low MAF kmers = 2453460C.

| Kmer | -log₁₀ *p* | β | MAC | MAF | Ps | Variants |
|---|---|---|---|---|---|---|
| CGCAACGACCTTACGCGACACCGGATCCGCG | 7.30 | -3.42 | 13 | 0.15 | 2453448 | 2453460A |
